# The paralogues MAGOH and MAGOHB are oncogenic factors in high-grade gliomas and safeguard the splicing of cell division and cell cycle genes

**DOI:** 10.1101/2022.12.20.521107

**Authors:** Rodrigo A. S. Barreiro, Gabriela D. A. Guardia, Fabiana M. Meliso, Xiufen Lei, Wei-Qing Li, Andre Savio, Martin Fellermeyer, Helena B. Conceição, Rafael L. V. Mercuri, Tesha Landry, Mei Qiao, Lorea Blazquez, Jernej Ule, Luiz O. F. Penalva, Pedro A. F. Galante

## Abstract

The exon junction complex (EJC) plays key roles throughout the lifespan of RNA and is particularly relevant in the nervous system. We investigated the roles of two EJC members, the paralogs MAGOH and MAGOHB, with respect to brain tumor development. High MAGOH/MAGOHB expression was observed in 14 tumor types; glioblastoma (GBM) showed the greatest difference compared to normal tissue. Increased MAGOH/MAGOHB expression was associated with poor prognosis in glioma patients, while knockdown of MAGOH/MAGOHB affected different cancer phenotypes. Reduced MAGOH/MAGOHB expression in GBM cells caused alterations in the splicing profile, including re-splicing and skipping of multiple exons. The binding profiles of EJC proteins indicated that exons affected by MAGOH/MAGOHB knockdown accumulated fewer complexes on average, providing a possible explanation for their sensitivity to MAGOH/MAGOHB knockdown. Transcripts (genes) showing alterations in the splicing profile are mainly implicated in cell division, cell cycle, splicing, and translation. We propose that high MAGOH/MAGOHB levels are required to safeguard the splicing of genes in high demand in scenarios requiring increased cell proliferation (brain development and GBM growth), ensuring efficient cell division, cell cycle regulation, and gene expression (splicing and translation). Since differentiated neuronal cells do not require increased MAGOH/MAGOHB expression, targeting these paralogs is a potential option for treating GBM.

## 1 INTRODUCTION

RNA binding proteins (RBPs) regulate multiple steps of gene expression, from RNA processing to translation (Glisovic et al., 2008). Aberrant RBP expression occurs in multiple cancer types and is linked to the acquisition of different cancer phenotypes (Marcelino Meliso et al., 2017; Gebauer et al., 2021). Identification and characterization of oncogenic RBPs are especially relevant in malignancies with no treatment options such as glioblastoma (GBM), the most common and lethal type of brain tumor (Ostrom et al., 2022). Musashi1 (Uren et al., 2015; Velasco et al., 2019; Baroni et al., 2021b), SERBP1 (Kosti et al., 2020), SNRPB(Correa et al., 2016), PTB (McCutcheon et al., 2004), IGF2BP3 (Suvasini et al., 2011; Bhargava et al., 2018; Sun et al., 2021) and HuR (Filippova et al., 2011; Guha et al., 2022b) are examples of RBPs established as oncogenic factors in GBM(Marcelino Meliso et al., 2017).

The exon junction complex (EJC) is a dynamic multi-protein complex deposited onto 20 nucleotides upstream from exon junctions of recently spliced mRNAs. The EJC, formed by core proteins eIF4A3, RBM8A, and MAGOH or MAGOHB, serve as platform for binding of other nuclear and cytoplasmic RBPs to safeguard mRNA processing, transport, decay, and translation(Boehm and Gehring, 2016; Deka and Singh, 2017). Mutations (or copy number variations) in EJC genes have been linked to intellectual disability, autism, microcephaly, and other pathologies(McMahon et al., 2016; Bartkowska et al., 2018).

In the context of cancer, the EJC components EIF4A3 and RBM8A, have been well-characterized. EIF4A3 is implicated in GBM, hepatocellular carcinoma, and pancreatic cancer, among other cancer types. Its impact on tumorigenesis involves interactions with lncRNA and circRNAs(Ye et al., 2021). RBM8A also affects GBM development via the Notch/STAT3 pathway(Lin et al., 2021), modulates TP53 expression(Lu et al., 2017), and affects sensitivity to DNA damage agents(Lu et al., 2017). However, MAGOH and MAGOHB are still poorly understood in the context of cancer.

MAGOH and MAGOHB are paralog genes(Singh et al., 2013) that, in addition to their function in the EJC, are involved in multiple biological processes including neurogenesis, brain development (Silver et al., 2013; Mao et al., 2015), cell cycle regulation (Silver et al., 2010; Inaki et al., 2011), and apoptosis (Michelle et al., 2012; Silver et al., 2013; Sheehan et al., 2020). Magoh-/-mice are embryonically lethal and Magoh-haplo-insufficient mice have smaller brains due to defects in neuronal stem cell division (Silver et al., 2010; Mao et al., 2015). Furthermore, hemizygous MAGOH deletion affects cell viability by compromising splicing and RNA surveillance (Viswanathan et al., 2018). MAGOH and MAGOHB are aberrantly expressed in different tumor types (Correa et al., 2016; Stricker et al., 2017; Zhou et al., 2020) while their knockdown affected the development of gastric cancer (Zhou et al., 2020).

In the present study, we determined that MAGOH and MAGOHB are highly expressed in multiple tumor types. In gliomas, high levels of MAGOH/MAGOHB were associated with poor overall survival and worse response to treatments. Their simultaneous knockdown affected cancer-related phenotypes in GBM cells but not astrocytes. High MAGOH/MAGOHB expression in GBM cells prevented re-splicing and aberrant splicing of genes (transcripts) implicated in cell cycle regulation, cell division, translation, and splicing. Since fully differentiated neuronal cells and GBM cells have very different requirements for MAGOH/MAGOHB function, targeting these paralogs is suggested as a potential therapeutic option for GBM.

## 2 RESULTS

### 2.1 MAGOH and MAGOHB expression are increased in different tumor types

We investigated MAGOH and MAGOHB expression using datasets from GTex (Genotype-Tissue Expression (GTEx Consortium, 2013)) and TCGA (The Cancer Genome Atlas). In 4,894 samples from 13 normal tissues, expression of MAGOH (blue boxplots) was higher than MAGOHB (yellow boxplots) and more variable across tissue types (Figure 1A and Supplementary Table 1). Despite these differences, MAGOH and MAGOHB expression levels were highly correlated in all tissues (rho > 0.8; Figure 1A; blue circles).

**Figure 1.**
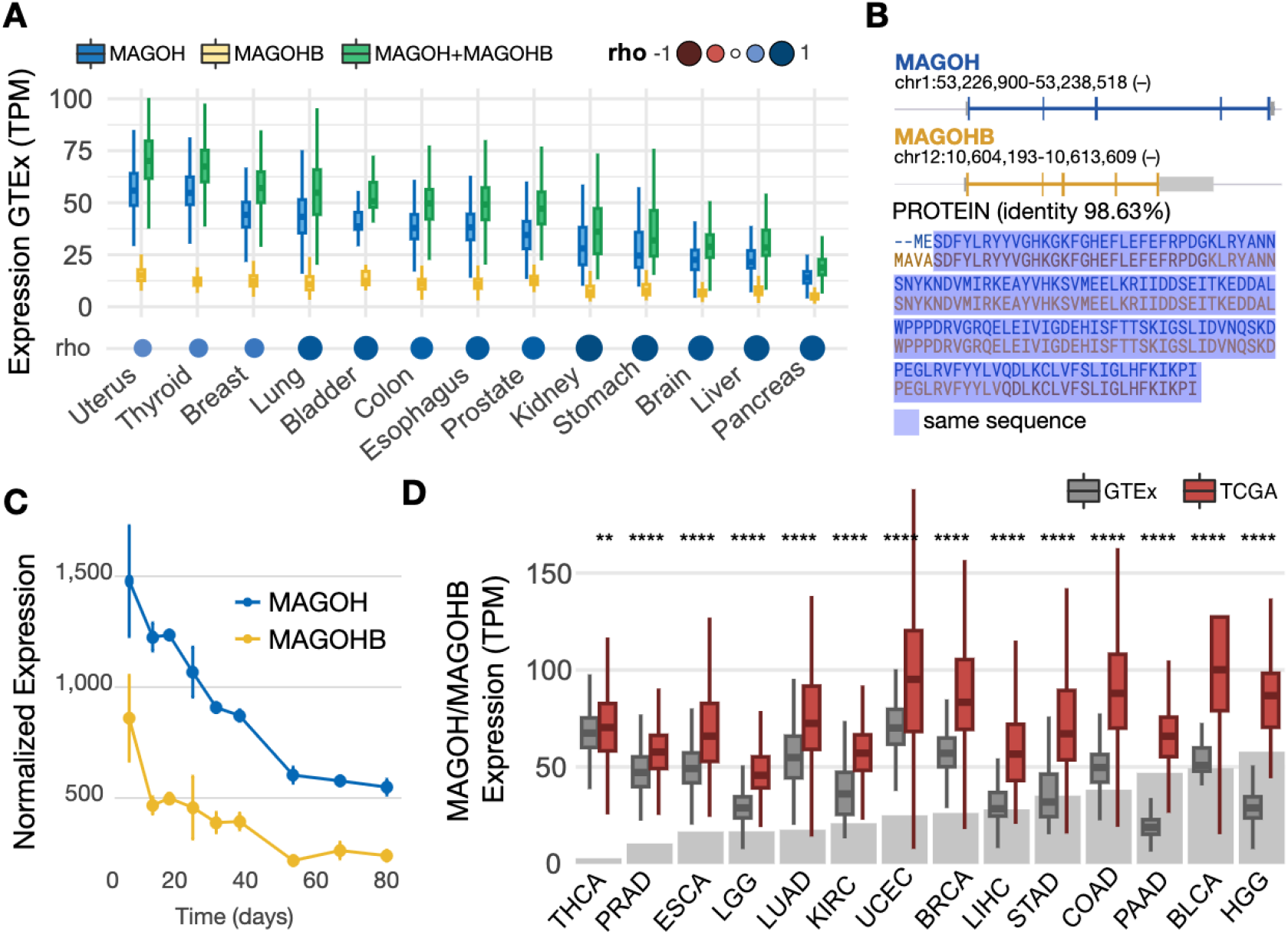
MAGOH and MAGOHB are overexpressed in multiple cancer types. A) MAGOH, MAGOHB, and MAGOH plus MAGOHB expression in multiple normal tissues from the GTEx database; Spearman correlation coefficients (rho) are shown between MAGOH and MAGOHB. B) MAGOH and MAGOHB genomic locations (top) and protein sequence alignment (bottom). Untranslated regions are shown in gray. C) MAGOH and MAGOHB expression (DESeq2 normalized) during cortex development according to Cortecon(van de Leemput et al., 2014). D) Combined MAGOH/MAGOHB expression levels in TCGA tumor samples vs. corresponding normal tissues from the GTEx database. Gray bars indicate the fold change between tumor versus normal expression (median). THCA, thyroid carcinoma; PRAD, prostate adenocarcinoma; ESCA, esophageal carcinoma; LGG, brain lower grade glioma; LUAD, lung adenocarcinoma; KIRC, kidney renal clear cell carcinoma; UCEC, uterine corpus endometrial carcinoma; BRCA, breast invasive carcinoma; LIHC, liver hepatocellular carcinoma; STAD, stomach adenocarcinoma; COAD, colon adenocarcinoma; PAAD, pancreatic adenocarcinoma; BLCA, bladder urothelial carcinoma; HGG, high grade (grade 4) glioma. Distributions of MAGOH/MAGOHB expression in tumor versus normal tissues were compared with Wilcoxon rank-sum tests. *P-value ≤ 0*.*01* (**) and *p-value ≤ 0*.*0001* (****).

Interestingly, the shared identity between these genes is high (98.6%; Figure 1B) at the protein level, but low at the nucleotide level (86%, Supplementary Figure 1A). Nonsynonymous (dN) to synonymous substitution (dS) rate ratio (**ω**) is 0.0056 (Supplementary Figure 1B), suggesting strong purifying selection on these genes and functional redundancy(Viswanathan et al., 2018).

MAGOH and MAGOHB expression are relatively low in different regions from the brain compared to other tissues (Figure 1A). In line with these results, MAGOH and MAGOHB are highly expressed in the early stages of corticogenesis (van de Leemput et al., 2014) (usually enriched with neuronal precursor cells) followed by a sharp decrease as cortex morphogenesis progresses (Figure 1C).

Next, we investigated MAGOH and MAGOHB expression (here, considered MAGOH plus MAGOHB expression) in 5,715 tumor samples from 14 cancer types. MAGOH/MAGOHB were overexpressed (p-value < 0.01, Wilcoxon rank sum test) in all analyzed cancers compared to their normal counterparts. Notably, the most pronounced expression difference in cancer vs. normal tissue was observed in high-grade glioma (HGG) (gray bar plot, Figure 1D and Supplementary Table 2).

### 2.2 MAGOH and MAGOHB levels are associated with survival in glioma patients and affect cancer-relevant phenotypes

Next, we investigated MAGOH and MAGOHB expression in 674 glioma samples (grade 4 samples, n=161; grade 3 samples, n=267; and grade 2 samples, n=246). Expression of both was lower in grade 2 gliomas and higher in grade 4 gliomas (p-value < 0.0001; Wilcoxon test) (Figure 2A and Supplementary Table 3). MAGOH/MAGOHB expression was also evaluated by immunostaining in an independent glioma cohort from the Shanghai ChangZheng Hospital. Levels of MAGOH/MAGOHB expression were positively correlated with malignancy with 93.2% of grade 4 gliomas, 83.3% of grade 3 gliomas, but only 44.4% of grade 2 gliomas (p-value < 0.001; displaying MAGOH/MAGOHB expression (Supplementary Table 4). Moreover, 61% (72/118) of grade 4 gliomas showed highly positive staining for MAGOH/MAGOHB expression, significantly more than in grade 2 (29.6%) and grade 3 (41.17%) gliomas (p-value = 0.005, Figure 2B).

**Figure 2.**
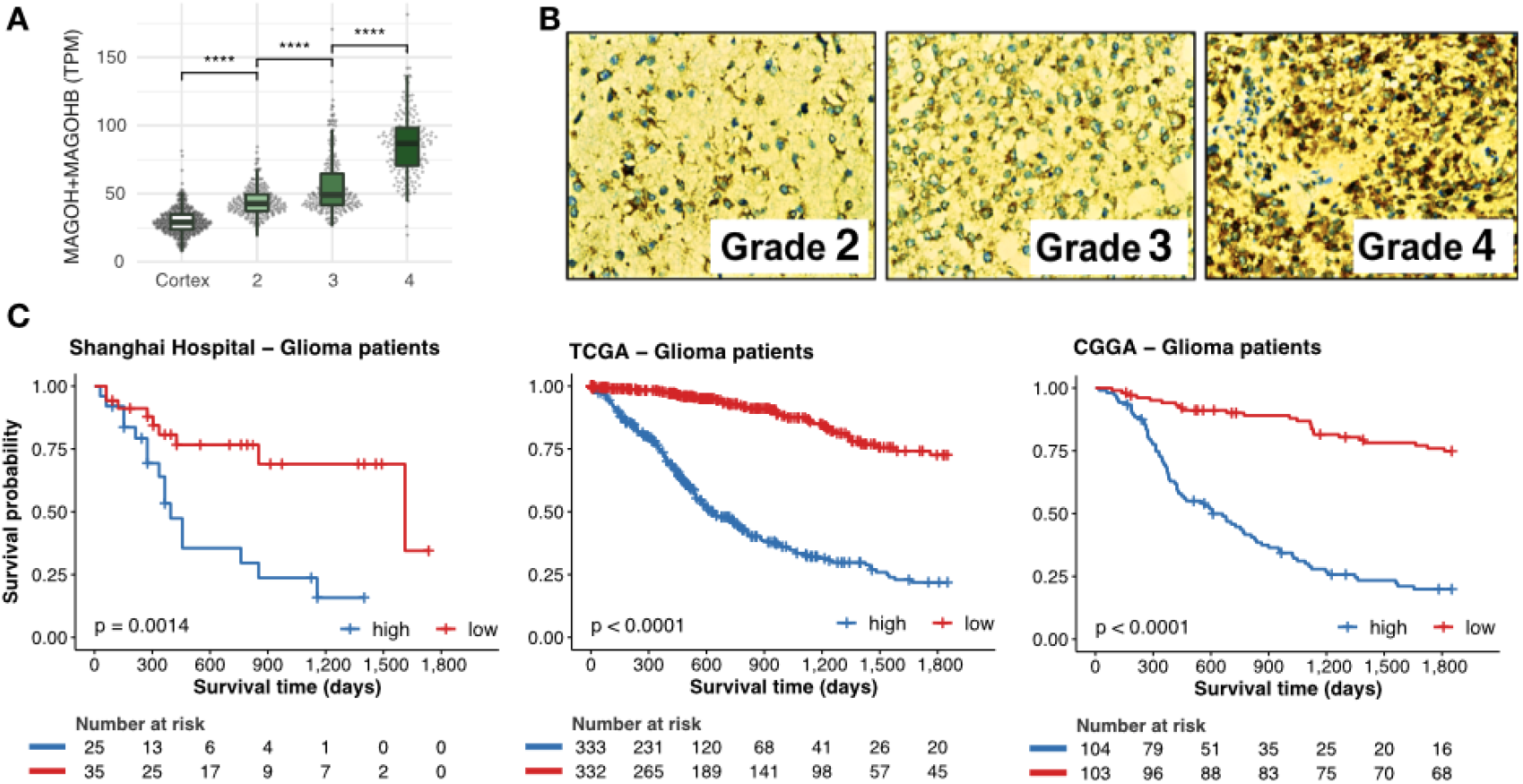
Levels of MAGOH and MAGOHB expression correlate with degree of malignancy and predict outcomes in glioma patients. **A)** Expression of MAGOH/MAGOHB in the brain cortex for grades 2, 3, and 4 gliomas (****: p-value < 0.0001; Wilcoxon rank-sum test). **B)** Immunohistochemical staining for MAGOH and MAGOHB in glioma samples from the Shanghai ChangZheng Hospital cohort. **C)** Survival curves of glioma patients with high (blue) and low (red) combined MAGOH and MAGOHB expression in three independent cohorts.

High expression of MAGOH/MAGOHB was correlated with worse overall survival in over 900 glioma patients from three independent cohorts (Shanghai ChangZheng Hospital, 60 patients; TCGA, 665 patients; CGGA, 207 patients) even after a co-factor stratification (Figure 2C and Supplementary Figure 2).

### 2.3 High expression of MAGOH and MAGOHB affect cancer-relevant phenotypes

We next investigated whether high expression of MAGOH/MAGOHB is necessary to maintain cancer-relevant phenotypes. Since both genes are highly conserved paralogs (Figure 1B) and potentially redundant in function, we knocked down both genes simultaneously in U251 and U343 GBM cells (Supplementary Figure 3).

MAGOH/MAGOHB silencing significantly reduced cell viability (Figure 3A) and proliferation (Figure 3B), increased levels of apoptosis (Figure 3C), and affected cell cycle distribution (G1 arrest) (Figure 3D). MAGOH/MAGOHB knockdown also affected viability of glioma stem cells (1919 and 3565). But in astrocytes, decreased levels of MAGOH/MAGOHB did not affect cell viability (Figure 1A) or levels of apoptosis (Figure 1C), suggesting that high MAGOH/MAGOHB expression is not required in normal neuronal cells.

**Figure 3.**
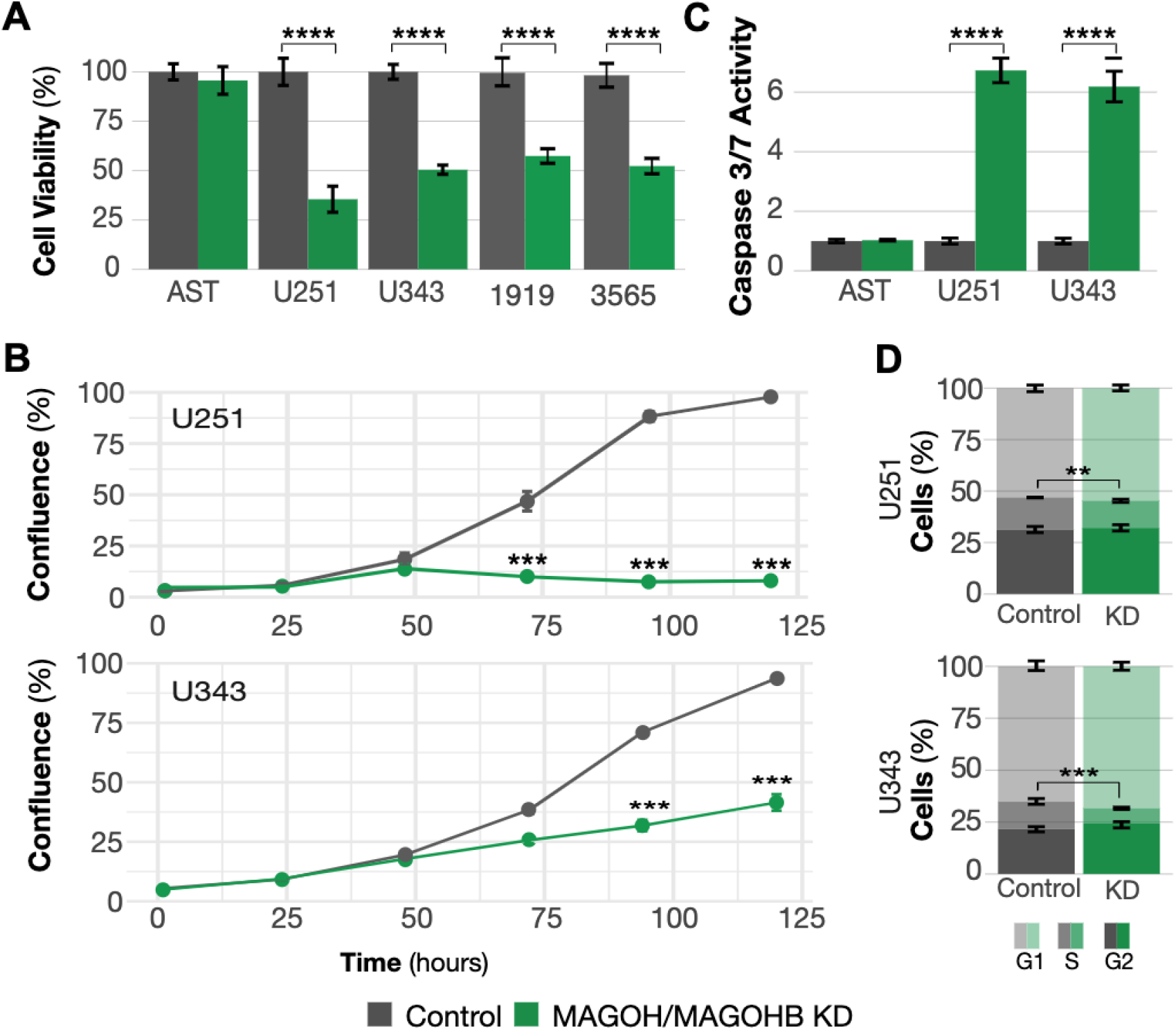
MAGOH and MAGOHB knockdown impacts cancer-relevant phenotypes. A) Cell viability was determined by MTS assays. MAGOH/MAGOHB knockdown significantly reduced viability of U251 and U343 glioblastoma cells and 1919 and 3565 glioma stem cells (**p-value <0.01, *p-value < 0.05; both, Student’s T-test). No changes were observed in astrocytes (AST). B) Proliferation was reduced in U251 and U343 MAGOH/MAGOHB knockdown cells compared to siControl cells; measured on an IncuCyte^®^ system (***p-value <0.001, ANOVA). C) Caspase-3/7 activity indicated increased apoptosis in U251 and U343 MAGOH/MAGOHB knockdown cells versus siControl cells. Astrocytes (AST) showed no changes in caspase activity (**p-value <0.01, T-test). D) Reduced MAGOH/MAGOHB expression caused changes in cell cycle distribution with G1 arrest in U251 and U343 cells (***p-value <0.001, t-test).

### 2.4 MAGOH and MAGOHB knockdown affects preferentially splicing events in genes implicated in cell division, cell cycle, translation, and RNA processing

As core components of the EJC, MAGOH and MAGOHB influence RNA processing. We investigated changes in the splicing profile in U251 and U343 cells after MAGOH/MAGOHB knockdown. A high number of splicing events were affected: 692 in U343 cells, 3,467 in U251 cells, and 190 events in both cell lines (Figure 4A). Among the different types of splicing alterations, exon skipping was prevalent (Figure 4B).

**Figure 4.**
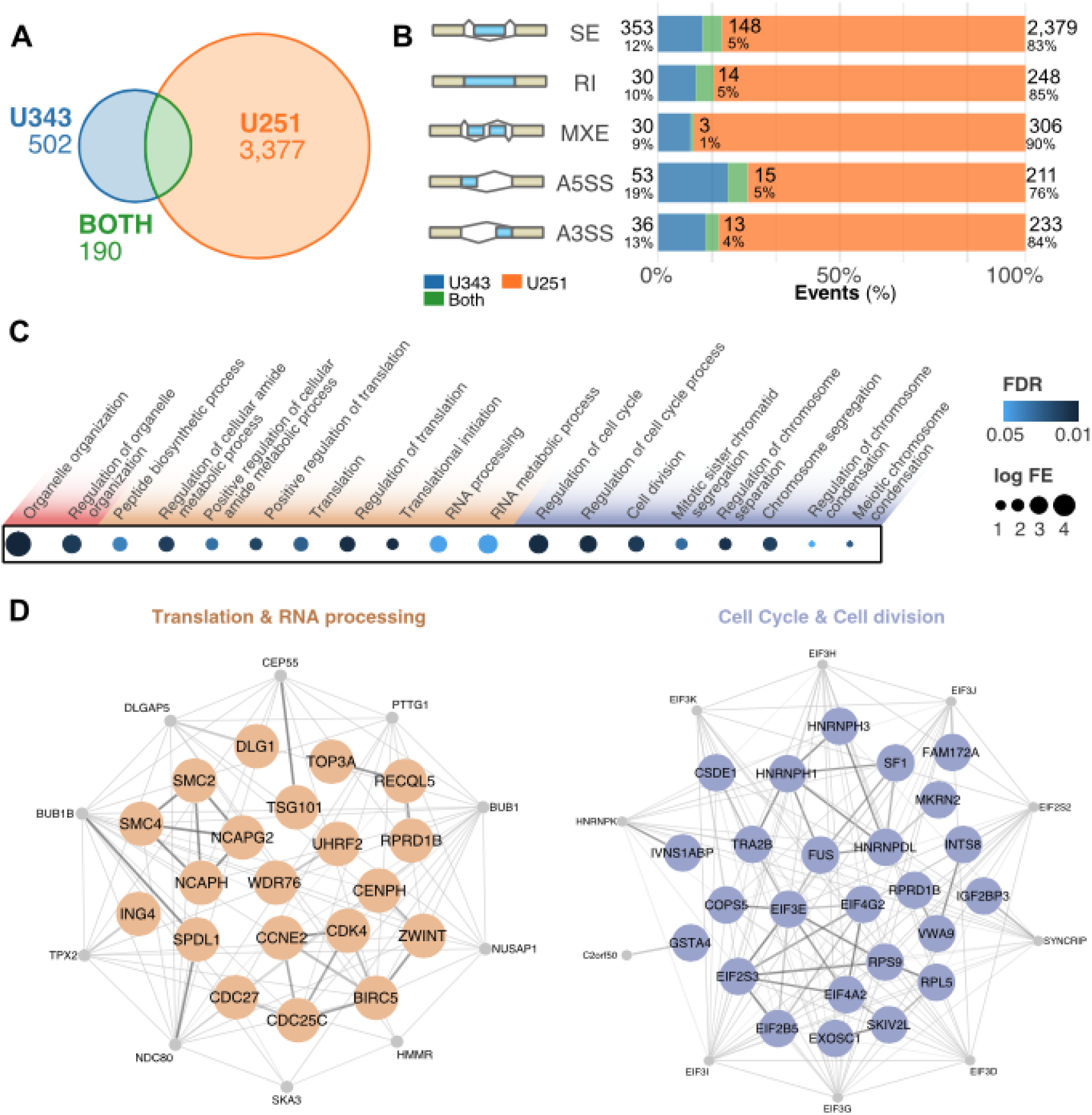
MAGOH/MAGOHB knockdown alters the splicing profile of GBM cells. (A) Splicing events showing changes after MAGOH/MAGOHB knockdown in GBM cell lines (|ΔPSI| > 0.2 and false discovery rate [FDR] < 0.01). (B) Altered splicing events in U251 and U343 cells stratified by event type: skipped exons (SE), retained introns (RI), mutually exclusive exons (MXE), and alternative 5′ and 3′ splice sites (A5SS and A3SS). (C) Enriched GO terms (biological processes) for genes displaying splicing alterations upon MAGOH/MAGOHB KD in both U251 and U343 cells (hypergeometric test; FDR < 0.05). (D) Protein-protein interaction networks of genes displaying the same splicing alteration in MAGOH/MAGOHB KD cells in both GBM cell lines. Node genes are shown in gray.

Gene Ontology (GO) enrichment analyses of genes displaying splicing alterations in MAGOH/MAGOHB KD cells identified cell cycle/division, regulation of RNA splicing, RNA processing, translation, and organelle organization as the top terms (Figure 4C). Similar GO terms were observed in all three sets (U251, U343, and U251-U343 overlap) (Figure 4C and Supplementary Table 5). Network analyses of RNA splicing/translation (Figure 4D) and cell cycle/division genes (Figure 4E) suggested that MAGOH/MAGOHB safeguard the splicing of highly connected genes in these processes.

To build on the specificity and relevance of MAGOH/MAGOHB impact on splicing, we analyzed another dataset(Viswanathan et al., 2018) in which MAGOH/MAGOHB expression levels were modulated in ChagoK1, a cell line derived from a non-small cell lung cancer that contains an hemizygous *MAGOH*-deletion. Using the same methodology and filters employed in analyses of U251 and U343 cells, we identified 3,801 alterations in splicing events in MAGOH/MAGOHB^high^ vs. MAGOH/MAGOHB^low^ ChagoK1 cells. A third of these altered splicing events were also observed in our analyses in U251 and U343 cells (Supplementary Figure 4). Moreover, GO analyses of genes displaying splicing alterations in ChagoK1 cells showed enrichment for the same biological processes as in our GBM analyses: cell division/cycle, RNA processing, and translation (Supplementary Figure 4). These results further indicate that MAGOH/MAGOHB preferentially modulates splicing of genes in these biological processes.

### 2.5 High expression of MAGOH/MAGOHB in GBM cells prevents aberrant splicing isoforms

EJC association with pre-mRNA is essential to prevent aberrant splicing(Joseph and Lai, 2021) by repressing cryptic splice sites that mediate recursive splicing (re-splicing) events(Blazquez et al., 2018; Boehm et al., 2018) and coordinating the correct order of intron excision(Le Hir et al., 2000; Lykke-Andersen et al., 2001). We examined our datasets for the presence of aberrant splicing events in MAGOH/MAGOHB KD cells. First, we checked whether splice junctions particularly affected by MAGOH/MAGOHB KD displayed any differences in EJC association (Figure 5A). Based on genome-wide evaluation of EJC binding sites(Patton et al., 2020), exons located upstream from those that were skipped in MAGOH/MAGOHB KD cells had a higher rate of EJC occupancy (Figure 5B; p-value = 0.0059), suggesting that a stronger EJC presence is required to prevent these aberrant splicing (potentially, re-splicing) events.

**Figure 5.**
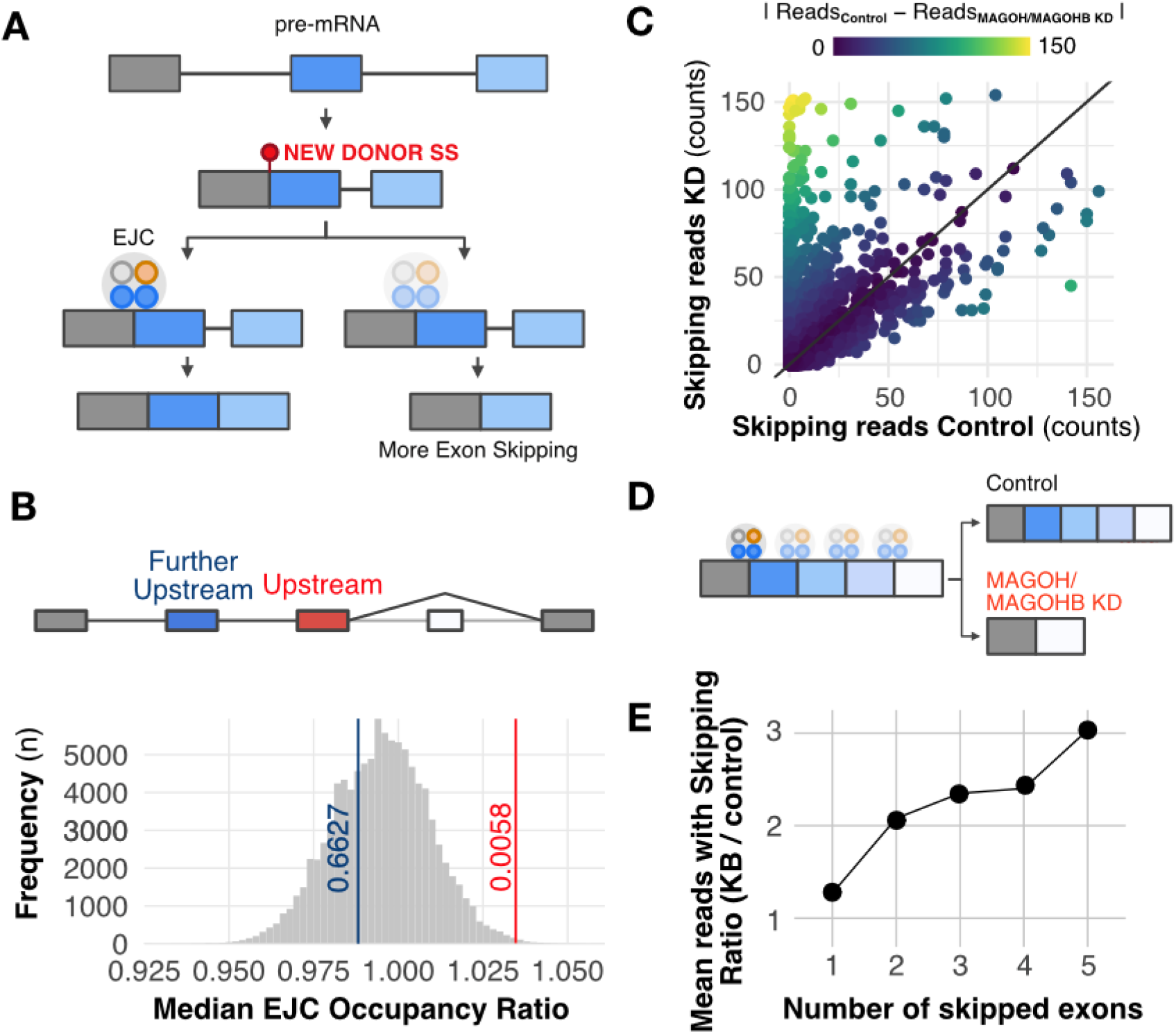
MAGOH/MAGOHB knockdown promotes multiple exon-skipping and aberrant splicing events. A) Model displaying how the EJC can prevent recursive splicing. After splicing, a new splice site can originate in the newly formed exon junction. EJC occupancy near this new junction can prevent a new splicing event (recursive splicing). B) EJC occupancy is elevated in the exon upstream to the skipped exon(s) regulated by MAGOH/MAGOHB compared to other exons in the same gene (excluding the last exon). Bootstrap curves (100,000 re-samplings) of random selected exons indicate that exons immediately upstream to skipped exons (red line) have a higher ratio of EJC occupancy than the remaining exons. Values obtained for the further upstream exons (red line) are similar to those of other exons. C) MAGOH/MAGOHB knockdown cells had more RNA-seq reads (reflecting skipping of one or more exons) compared to controls. D) Model proposing that exons with high EJC occupancy are more sensitive to MAGOH/MAGOHB KD, resulting in multiple exon-skipping events. E) MAGOH/MAGOHB knockdown cells display a higher ratio (median) of reads, supporting skipping of two or more exons (e.g., 2x and 3x for skipping 2 and 5 exons, respectively).

Additionally, MAGOH/MAGOHB KD sequencing data showed a higher number of reads supporting exon-skipping events, some exclusively found in MAGOH/MAGOHB KD cells (Figure 5C). Finally, we observed that MAGOH/MAGOHB KD cells have more exon-skipping events (two or more exons) than control cells (Figure 5D and E, Supplementary Table 6), suggesting that they also have a higher number of aberrant splicing events.

Altogether, these results indicate that reduced expression of MAGOH/MAGOHB can create aberrant splicing events in glioblastoma cells, and suggest that high MAGOH/MAGOHB expression in tumors has an important role in preventing such events that are potentially harmful to tumor cells.

### 2.6 Genes presenting with aberrant splicing events in MAGOH/MAGOHB knockdown cells are mainly associated with regulation of cell cycle and cell division

We then analyzed the impact of aberrant splicing on gene function. Genes related to main GO enriched terms in Figure 4C with multiple exon-skipping events in U251 and U343 MAGOH/MAGOHB KD cells are depicted in Figure 6A. For genes related to cell cycle/division, we determined that in 85% of cases, aberrant splicing events alter protein domains with complete or partial loss through the inclusion of premature stop codons or frameshift events (Figure 6B). These alterations affect genes regulating different cell cycle phases (Figure 6C). Thus, we propose that in GBM cells, high MAGOH/MAGOHB expression would prevent aberrant splicing events that could ultimately compromise cell cycle and division, critical steps for tumor growth.

**Figure 6.**
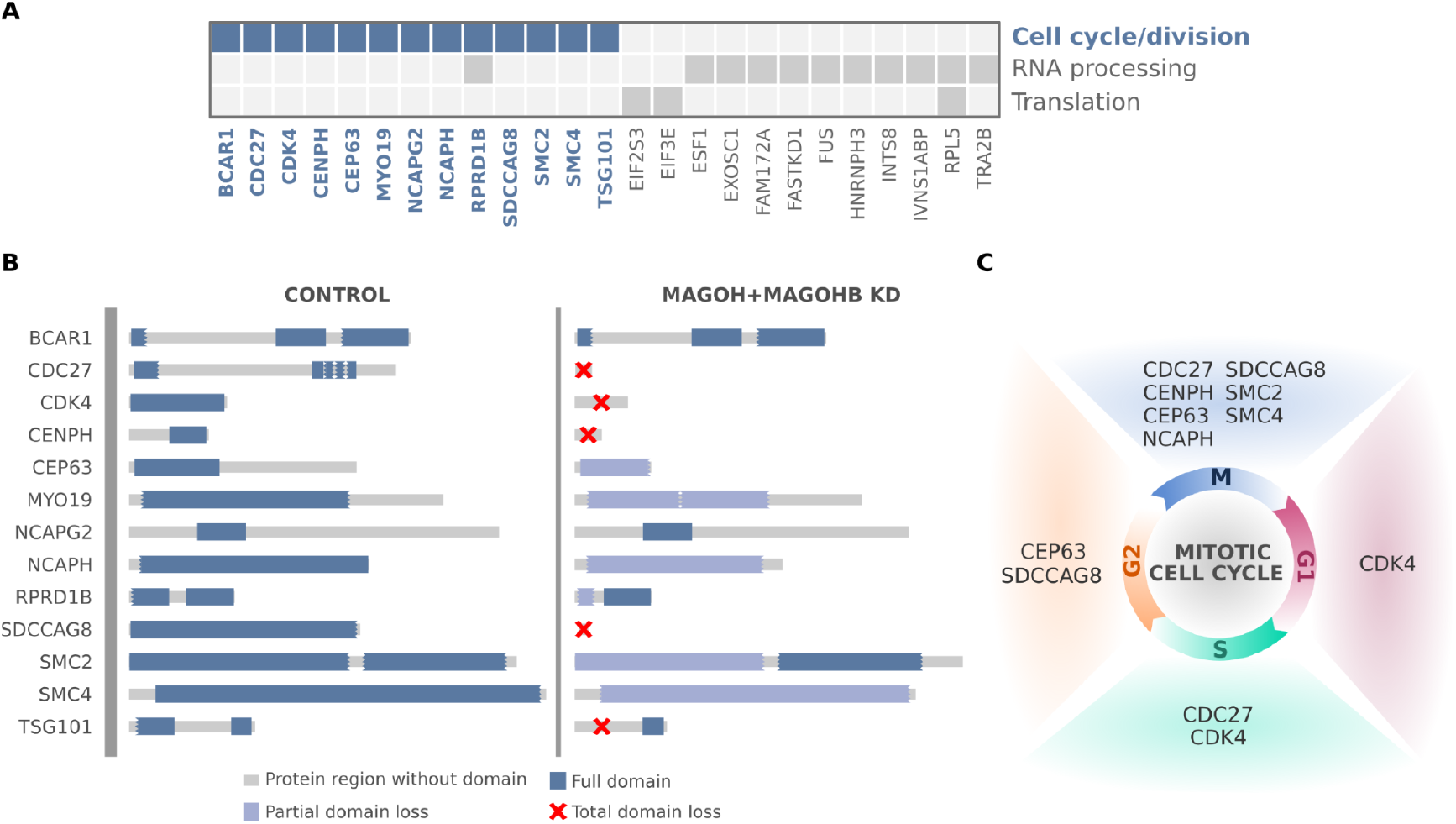
Genes affected by MAGOH/MAGOHB knockdown presenting with multiple exon-skipping events are mainly associated with regulation of cell cycle/division. (A) Genes with multiple exon-skipping events in both U251 and U343 MAGOH/MAGOHB KD cells and their functions. (B) Cell cycle and cell division genes whose protein domains were compromised (partial or total domain losses) due to multiple exon-skipping events in MAGOH/MAGOHB KD cells. (C) Genes presenting partial or total domain losses due to multiple exon-skipping events caused by MAGOH/MAGOHB KD are implicated in different phases of the cell cycle.

## DISCUSSION

RNA binding proteins (RBPs) modulate numerous steps in gene expression and are particularly relevant in the nervous system (neurogenesis and brain development (Popper et al., 2021; Prashad and Gopal, 2021)), where they orchestrate changes in splicing, translation, and mRNA decay. Alterations in RBP expression levels and function have been associated with neurological disorders (e. g., HNRNPA1 (Jahanbazi Jahan-Abad et al., 2022), MATR3 (Johnson et al., 2014), TIA1 (Mackenzie et al., 2017), and TARDBP (Valdmanis et al., 2009)) and brain tumor development (e.g., Musashi1 (Toda et al., 2001; Galante et al., 2009; Velasco et al., 2019; Baroni et al., 2021a), SERBP1(Correa et al., 2016), PTB (McCutcheon et al., 2004) and HuR (Filippova et al., 2011; Guha et al., 2022a)). Modulation of their expression or target genes with specific inhibitors has started to be explored therapeutically (Hong, 2017). For example, the EJC proteins MAGOH and MAGOHB were among top hits identified in a functional genomic screening to identify RNA binding proteins with oncogenic potential in glioblastoma cells (Correa et al., 2016; Marcelino Meliso et al., 2017).

Our analyses showed that MAGOH/MAGOHB display increased expression in the 14 cancer types analyzed in relation to normal counterparts, suggesting a broad involvement in tumor development. Among them, grade 4 glioma showed the greatest difference in expression versus normal tissues. High MAGOH/MAGOHB expression in grade 4 glioma (Correa et al., 2016) seems to be particularly important, as loss of Chromosome 1p – where the MAGOH gene is located – is observed in some tumor types but rarely in grade 4 glioma (Viswanathan et al., 2018).

As members of the exon-exon junction complex (EJC), MAGOH and MAGOHB participate in different stages of RNA processing, transport, and translation. Knockdown or depletion of EJC members lead to alterations in splicing profiles, including the occurrence of recursive splicing and generation of aberrant splicing isoforms. These effects were observed both in humans and Drosophila cells, indicating that the EJC has a conserved function in “safe-guarding” the splicing process (Blazquez et al., 2018; Joseph and Lai, 2021; Otani et al., 2021). Genomic studies have shown differences in EJC loading across junctions within a given transcript (Saulière et al., 2010; Singh et al., 2012). Furthermore, only a fraction of splicing events is affected by depletion of EJC members (Wang et al., 2014; Fukumura et al., 2016; Viswanathan et al., 2018). According to binding site profiles for EJC members (Patton et al., 2020), exon junctions associated with splicing events affected by MAGOH/ MAGOHB knockdown display on average fewer associations than other junctions in the same transcript, providing one possible explanation for their increased sensitivity.

Certain cell types and tissues show high sensitivity to changes in expression of EJC members (Asthana et al., 2022). MAGOH, MAGOHB, and EIF4A3 display high expression levels during early stages of cortex formation (van de Leemput et al., 2014) and reduced levels in differentiated neuronal cells. Members of the EJC are critical during brain development and their mutation in humans or knockout in mouse models result in microcephaly (Mao et al., 2015, 2016). In GBM cells, we observed that MAGOH and MAGOHB knockdown preferentially affected splicing of transcripts (genes) implicated in splicing, cell cycle, and especially cell division. All common mutations causing microcephaly are present in genes implicated in cell division (MCPH1, ASPM, CDK5RAP2, CENPJ, STIL, WDR62, and CEP152) (Vue et al., 2020; Phan and Holland, 2021; Zaqout and Kaindl, 2021). We suggest that in these two scenarios (brain and glioblastoma development), highly proliferating cells demand efficient expression of cell cycle and division genes, and increased levels of MAGOH and MAGOHB are required to prevent defects in the splicing of these transcripts.

Targeting of microcephaly-associated genes has been proposed as an alternative to microtubule-targeting agents to treat brain tumors (Iegiani et al., 2021). In line with this idea, our results showed that MAGOH and MAGOHB knockdown is well-tolerated in astrocytes but not in GBM cells. Therefore, considering the impact of MAGOH and MAGOHB on cancer phenotypes and the dissimilar sensitivity to their knockdown between GBM and differentiated neuronal cells, targeting EJC members such as MAGOH and MAGOHB may be an alternative therapeutic strategy to treat GBM.

## METHODS

### MAGOH and MAGOHB expression profiles in healthy tissues

MAGOH and MAGOHB expression levels were evaluated using RNA sequencing (RNAseq) data from 4,894 samples of healthy tissues from 13 different sites, including bladder (21 samples), breast mammary tissue (459 samples), sigmoid and transverse colon (373 and 406 samples), esophageal mucosa (555 samples), kidney cortex (85 samples), brain cortex and frontal cortex (255 and 209 samples), liver (226 samples), lung (578 samples), pancreas (328 samples), prostate (245 samples), stomach (359 samples), thyroid (653 samples) and uterus (142 samples). Gene expression (in transcripts per million [TPM]) from each sample was extracted from the GTEx portal (GTEx Consortium, 2013) (version 8).

### MAGOH and MAGOHB expression profiles in tumor versus healthy tissues

We downloaded TCGA RNA sequencing data from 14 cancer types with matching normal tissues available from GTEx (GTEx Consortium, 2013). In total, 5,715 raw RNA sequencing (FASTq files) were downloaded from the GDC data portal (https://portal.gdc.cancer.gov/) and locally processed using Kallisto (Kallenberg et al., 2017) (default parameters; version 0.43.1) and txImport (Soneson et al., 2015) (default parameters; version 1.24.0) to obtain gene-level expression in TPM. For these analyses, the expression of MAGOH and MAGOHB (MAGOH/MAGOHB) were considered combined (summed).

### MAGOH and MAGOHB expression and impact on patient survival

To evaluate whether MAGOH and MAGOHB expression levels could predict outcomes of glioma patients, we stratified patients in the TCGA and CGGA cohorts, based on MAGOH/MAGOHB median expression levels, in high vs. low expression groups. In each cohort, we performed survival analyses between groups using log-rank tests. Kaplan-Meier survival curves were built using R (https://www.r-project.org/) packages survival (version 3.4.0) and survminer (version 0.4.9).

Another survival study was conducted using data from a cohort at the Changzheng Hospital (Naval Medical University, Shanghai, China). All cases were obtained from the Department of Pathology, and were graded by the two pathologists separately using the World Health Organization grading system. A cohort of 177 patients underwent resection in the Department of Neurosurgery from January 2011 to August 2016. All aspects of the study were reviewed and approved by the Specialty Committee on Ethics of Biomedicine Research, Second Military Medical University. The clinicopathologic characteristics of the patients are summarized in Supplementary Table 1.

### Immunohistochemical assays

Immunohistochemistry was performed using 5 mm thick tissue sections from the Changzheng Hospital cohort; sections were paraffin-embedded and mounted on pre-coated glass slides. The anti-MAGOH/MAGOHB antibody (ab170944) was purchased from Abcam. The staining intensity of tumor cells had a predominantly nuclear focal pattern, but in some cases, we also observed a weaker cytoplasmic focal staining pattern. Samples were scored 0 to 4 based on staining. The percentages of tumor cells with staining were graded as 0 (<5%), 1 (5-25%), 2 (25-50%), 3 (50-75%), or 4 (75-100%). We then classified staining intensity and percentages of cells stained as strong (+++, final score 7-9), moderate (++, final score =4-6) and weak (+, final score =0-3). To conduct survival analyses, we compared “high” (+++) vs. “low” (++ and below) groups and used log-rank tests for comparisons. Kaplan-Meier survival curves were built in R.

### Cell growth and transfection

U251 and U343 glioblastoma cells were obtained from the University of Uppsala (Sweden) and cultured in Dulbecco’s Modified Eagle medium with 10% fetal bovine serum, 1% penicillin and streptomycin (Life Technologies, Carlsbad, CA). Astrocytes (ScienCell #1800) were cultured in DMEM:F12 with 10% FBS and 1% penicillin and streptomycin. Glioma stem cell lines (MES3565 (3565)(Jin et al., 2017) and PM1919 (1919)(Jin et al., 2017)) were cultured in serum-free media consisting of Neurobasal-A media supplemented with B-27, sodium pyruvate, Glutamax, 1% penicillin and streptomycin, 20 ng/ml EGF (ThermoFisher), and 20 ng/ml hFGF (PeproTech). Glioma stem cells were pulsed every 72 h with EGF/FGF and then disassociated with Accutase (ThermoFisher) at room temperature for 10 min. Cells were transfected with control siRNA or MAGOH/MAGOHB small interfering RNA (siRNA) using Lipofectamine RNAiMax reagent (Invitrogen). All experiments were performed in triplicate. siRNAs were obtained from Dharmacon, MAGOH#L-011327-00-0005, MAGOHB#L-016706-02-0005, Control # D-001810-10-05. Quantitative PCR was performed using TaqMan Universal PCR Master Mix and probes, and reactions performed on a ViiA− 7 Real-Time PCR System (all, Applied Biosystems, Waltham, MA). Data were acquired using the system’s RUO software and analyzed using the 2−ΔΔCT method, with GAPDH as an endogenous control.

### Cell viability and caspase 3/7 activity assays

Transfected cells were seeded in a 96-well tissue culture plate. 48 hours later, cell viability and caspase 3/7 activity were evaluated using CellTiter-Glo and Caspases-Glo 3/7 assays (both from Promega, Madison, WI), according to the manufacturer’s instructions. Absorbance and luminescence were measured with a Molecular Devices SpectraMax M5 microplate reader. Data were evaluated using T-tests and presented as mean ± standard errors. All experiments were performed in triplicate.

### Cell proliferation assays

U251 and U343 transfected cells were grown in 96-well tissue culture plates (8×10^2^ cell/well). The percentage of confluent cells was monitored for 125 hours using a high-definition automated imaging system (IncuCyte^®^; Sartorius, Goettingen, Germany). Data were evaluated using ANOVA and presented as mean ± standard errors. All experiments were performed in triplicate.

### Cell cycle assays

Flow cytometry was used to perform cell cycle analyses. U251 and U343 cells were plated using a 24-well plate (4 x 10^4^ cells/well). Cells were transfected with either siControl or siMAGOH + siMAGOHB and incubated for 48 hours. Cells were harvested using trypsin, washed with phosphate-buffered saline, and then fixed with 75% ethanol. Cells were spun down and resuspended in 500 µL of propidium iodide and incubated for a minimum of 20 minutes. A FACS BD Caliber instrument was used for flow cytometry. All experiments were performed in triplicate.

### RNA extraction and RNA sequencing assays

Total RNA from transfected U251 and U343 cells (siRNA control vs. siRNA MAGOH/MAGOHB) was extracted using the TRIzol reagent (Invitrogen, Carlsbad, CA) according to the manufacturer’s instructions. Reverse transcription was performed using a High-Capacity cDNA reverse transcription kit (Applied Biosystems, Warrington, WA) with random priming. Libraries used in RNA sequencing were prepared using TruSeq RNA Library Preparation Kit (Illumina, San Diego, CA), following manufacturer’s instructions and sequenced in a HiSeq-2000 machine at UTHSCSA’s Genome Sequencing Facility. All experiments were performed in triplicate.

### Splicing analyses

To identify splicing alterations induced by MAGOH/MAGOHB knockdown, raw RNA-Seq reads of control and knockdown samples were aligned against the human reference genome (version GRCh38) and matching GENCODE transcriptome (v29) using STAR (Dobin et al., 2013) (default parameters; version 2.7.7.a). The mapped reads from control and knockdown samples of U251 and U343 cell lines were processed using rMATS (Shen et al., 2014) (default parameters; version v4.1.2) to characterize splicing events. Events were divided into exon skipping, mutually exclusive exons, intron retention, and alternative donor or acceptor sites (A3SS). Events were classified as statistically significant if p-values (adjusted for false discovery rate) were below 0.01 and absolute delta for percent splice in (deltaPSI) was above 0.2. For further quantification of exons spliced, since there are many possible transcript configurations, we defined MANE transcripts as our reference and only evaluated genes with three or more exons.

To corroborate the observed splicing alterations driven by MAGOH/MAGOHB knockdown in U251 and U343 cells, we obtained FASTq files from Viswanathan et al. (Viswanathan et al., 2018) and compared the splicing profiles of ChagoK1 cells (MAGOH + MAGOHB high vs. MAGOH + MAGOHB low). Data were processed using the same pipelines and following the same parameters described above.

### Gene Ontology analyses

Gene Ontology (GO) analyses were performed with coding genes showing differences in splicing patterns in control vs. MAGOH and MAGOHB knockdown cells in both cell lines analyzed. We used Panther (Mi et al., 2019) (default parameters; version 17.0), and considered all human coding genes as background. Only GO terms with FDR < 0.05 and fold enrichment greater than two were considered. Protein-protein interaction analyses were performed for enriched GO gene sets using the STRING database (Szklarczyk et al., 2019) coupled to Cytoscape (Shannon et al., 2003).

### Analyses of EJC crosslink immunoprecipitation data

Exon-level EJC binding quantifications were obtained from Patton et al. (Patton et al., 2020). We aggregated the count from all chemical crosslink experiments. For these analyses, we kept only genes with at least one read on the available RNA-Seq count columns. For the bootstrap analyses of exon-skipping events, we calculated the mean of reads from all exons except the first and last exons (since the skipping of these exons is often caused by other events) and then we calculated the ratio reads mapped to the upstream exon of a exon-skipped event to the calculated mean reads. We performed this calculation for each gene and exon with exon-skipped events in any of the three splicing analyses (U251, U343, and ChagoK1). For the 100,000 resampling, we randomly selected exons of any filtered gene list and calculated the described ratio. The next step was to compare the mean of the ratio obtained from exons upstream the skipping event with the ratios achieved by the resampling, and calculated p-values using two-tailed tests.

### Ka/Ks Analysis

To calculate the nonsynonymous (dN) to synonymous substitution (dS) rate ratio (ω), we extracted the nucleotide and amino acid sequences from MAGOH and MAGOHB and aligned them using ClustalW (McWilliam et al., 2013). PAL2NAL (Yang, 2007) (codeml) was used to calculate the synonymous (dS) and non-synonymous (dN) substitution rates.

## Code availability

Details on the Python, R, and shell script codes used in this manuscript will be available for non-commercial academic purposes at GitHub (https://github.com/galantelab/magoh).

## Data availability

RNA sequencing data have been deposited at the European Nucleotide Archive (ENA) under accession number PRJEB55365.

## Acknowledgements

We would like to thank Karen Klein for the editorial services. This work was supported by NIH grant R21CA205475-01 to L.O.F.P. P.A.F.G was supported by grants from Serrapilheira Foundation and Fundação de Amparo à Pesquisa do Estado de São Paulo (FAPESP) [2018/15579-8]. G.D.A.G. was supported by a fellowship from FAPESP [2017/19541-2] and Young Scientist program, Hospital Sírio-Libanês. H.B.C. was supported by a fellowship from FAPESP [2018/13613-4] and CAPES program. R.L.V.M. was supported by a fellowship from FAPESP [2020/02413-4].

## Author Contributions

Conceptualization, L.O.F.P and P.A.F.G.; Funding acquisition, L.O.F.P. and P.A.F.G.; Major bioinformatics analyzes, R.A.S.B. and G.D.A.G.; Further bioinformatics analyzes F.M.M., H.B.C., R.L.V.M.; RNASeq experiments, M.Q.; Pathological and survival analyses, W.Q.L; Biological assays and expression analysis, X.L., T.L., M.F., A.S.; Concept and input in splicing analyzes, JL and LB; Writing - original draft, L.O.F.P., P.A.F.G., G.D.A.G., R.A.S.B.; Writing - review and editing, all authors.

