## Supplemental Tables for "The paralogues MAGOH and MAGOHB are oncogenic factors in high-grade gliomas and safeguard the splicing of cell division and cell cycle genes"

| Table S1. MAGOH and MAGOHB expression in normal (GTEX) data. | | | | |
|---|---|---|---|---|
| sample_id | tissue | tcga_correspondence | MAGOH_tpm | MAGOHB_tpm |
| GTEX-OIZF-1926-SM-7PBZS | Bladder | TCGA-BLCA | 55.71 | 16.99 |
| GTEX-P44H-2226-SM-E9U4P | Bladder | TCGA-BLCA | 32.63 | 7.83 |
| GTEX-QEL4-1826-SM-EZ6KU | Bladder | TCGA-BLCA | 45.37 | 18.36 |
| GTEX-RN64-2426-SM-EZ6L2 | Bladder | TCGA-BLCA | 41.14 | 14.94 |
| GTEX-RU72-2526-SM-EWRML | Bladder | TCGA-BLCA | 46.03 | 17.46 |
| GTEX-S32W-1126-SM-4AD5V | Bladder | TCGA-BLCA | 47.02 | 20.19 |
| GTEX-S33H-1926-SM-EYYVH | Bladder | TCGA-BLCA | 40.53 | 17.42 |
| GTEX-S3XE-1226-SM-4AD4L | Bladder | TCGA-BLCA | 38.57 | 8.93 |
| GTEX-S4Q7-0926-SM-4AD5D | Bladder | TCGA-BLCA | 32.99 | 9.69 |
| GTEX-S4UY-0926-SM-4AD6O | Bladder | TCGA-BLCA | 36.4 | 11.43 |
| GTEX-S95S-0626-SM-EZ6L8 | Bladder | TCGA-BLCA | 38.63 | 14.11 |
| GTEX-SE5C-1026-SM-4BRUG | Bladder | TCGA-BLCA | 35.39 | 9.02 |
| GTEX-SNMC-0826-SM-4DM66 | Bladder | TCGA-BLCA | 37.25 | 11.48 |
| GTEX-SNOS-0526-SM-4DM54 | Bladder | TCGA-BLCA | 45.97 | 13.84 |
| GTEX-T2YK-2326-SM-EZ6LA | Bladder | TCGA-BLCA | 101.93 | 31 |
| GTEX-T5JW-1026-SM-EZ6LR | Bladder | TCGA-BLCA | 44.92 | 11.03 |
| GTEX-T6MN-2226-SM-EVYAM | Bladder | TCGA-BLCA | 29.08 | 14.35 |
| GTEX-TMMY-1526-SM-4DXST | Bladder | TCGA-BLCA | 39.18 | 9.62 |
| GTEX-U3ZM-0826-SM-4DXU6 | Bladder | TCGA-BLCA | 37.62 | 10.56 |
| GTEX-U3ZN-1226-SM-4DXUD | Bladder | TCGA-BLCA | 38.35 | 12.41 |
| GTEX-U4B1-1226-SM-4DXT7 | Bladder | TCGA-BLCA | 37.75 | 12.19 |
| GTEX-1117F-3226-SM-5N9CT | Brain - Cortex | TCGA-LGG | 15.76 | 4.25 |
| GTEX-111FC-3126-SM-5GZZ2 | Brain - Cortex | TCGA-LGG | 21.71 | 6.9 |
| GTEX-1128S-2726-SM-5H12C | Brain - Cortex | TCGA-LGG | 21.08 | 5.25 |
| GTEX-117XS-3026-SM-5N9CA | Brain - Cortex | TCGA-LGG | 25.59 | 5.51 |
| GTEX-1192X-3126-SM-5N9BY | Brain - Cortex | TCGA-LGG | 30.48 | 8.67 |
| GTEX-11DXW-1126-SM-5H12Q | Brain - Cortex | TCGA-LGG | 18.96 | 4.94 |
| GTEX-11DXY-3226-SM-5GIDE | Brain - Cortex | TCGA-LGG | 27.54 | 4.97 |
| GTEX-11EI6-3026-SM-5GZZO | Brain - Cortex | TCGA-LGG | 20.1 | 4.44 |
| GTEX-11EMC-3226-SM-5EGKW | Brain - Cortex | TCGA-LGG | 14.14 | 5.06 |

| | | | | |
|---|---|---|---|---|
| GTEX-11GS4-3126-SM-5A5LH | Brain - Cortex | TCGA-LGG | 15.96 | 6.1 |
| GTEX-11GSO-2926-SM-5HL73 | Brain - Cortex | TCGA-LGG | 24.7 | 6.84 |
| GTEX-11GSP-3226-SM-5986O | Brain - Cortex | TCGA-LGG | 14.82 | 5.64 |
| GTEX-11NUK-2926-SM-5A5MD | Brain - Cortex | TCGA-LGG | 23.28 | 7 |
| GTEX-11NV4-2126-SM-5N9DS | Brain - Cortex | TCGA-LGG | 9.67 | 4.26 |
| GTEX-11O72-2926-SM-5BC4V | Brain - Cortex | TCGA-LGG | 21.28 | 5.02 |
| GTEX-11OC5-0726-SM-5P9JK | Brain - Cortex | TCGA-LGG | 32.86 | 7.96 |
| GTEX-11OF3-2926-SM-5HL78 | Brain - Cortex | TCGA-LGG | 23.01 | 7.48 |
| GTEX-11ONC-2926-SM-5P9JM | Brain - Cortex | TCGA-LGG | 23.73 | 6.04 |
| GTEX-11PRG-2926-SM-5987A | Brain - Cortex | TCGA-LGG | 18.79 | 6.62 |
| GTEX-11TTK-2926-SM-5PNYP | Brain - Cortex | TCGA-LGG | 15.14 | 4.1 |
| GTEX-11WQK-3026-SM-5EQL6 | Brain - Cortex | TCGA-LGG | 19.68 | 6.48 |
| GTEX-11ZUS-2926-SM-5FQSL | Brain - Cortex | TCGA-LGG | 20.48 | 4.5 |
| GTEX-12126-1026-SM-5P9JJ | Brain - Cortex | TCGA-LGG | 18.74 | 5.63 |
| GTEX-12WSA-2926-SM-5EQ4D | Brain - Cortex | TCGA-LGG | 9.83 | 3.3 |
| GTEX-12WSC-3026-SM-5GCNF | Brain - Cortex | TCGA-LGG | 23.89 | 5.99 |
| GTEX-12WSD-3126-SM-5HL7P | Brain - Cortex | TCGA-LGG | 15.67 | 6.32 |
| GTEX-12WSF-3126-SM-6M478 | Brain - Cortex | TCGA-LGG | 19.37 | 5.05 |
| GTEX-12WSH-3026-SM-5CVNI | Brain - Cortex | TCGA-LGG | 23.93 | 6.27 |
| GTEX-12ZZW-2926-SM-5LZUP | Brain - Cortex | TCGA-LGG | 19.84 | 3.75 |
| GTEX-12ZZX-2926-SM-5GCOQ | Brain - Cortex | TCGA-LGG | 26.9 | 7.59 |
| GTEX-12ZZY-3026-SM-5GCOU | Brain - Cortex | TCGA-LGG | 25.15 | 8.06 |
| GTEX-12ZZZ-3026-SM-5BC67 | Brain - Cortex | TCGA-LGG | 22.58 | 6.4 |
| GTEX-1313W-3126-SM-5LZUI | Brain - Cortex | TCGA-LGG | 20.34 | 7.62 |
| GTEX-131XW-3126-SM-5LZUC | Brain - Cortex | TCGA-LGG | 14 | 3.81 |
| GTEX-131YS-3126-SM-5KLYT | Brain - Cortex | TCGA-LGG | 25.1 | 7.71 |
| GTEX-132Q8-3026-SM-5PNVG | Brain - Cortex | TCGA-LGG | 26.42 | 6.31 |
| GTEX-1399T-3026-SM-5KLZC | Brain - Cortex | TCGA-LGG | 22.1 | 6.86 |
| GTEX-139T8-1026-SM-5J2MC | Brain - Cortex | TCGA-LGG | 18.63 | 3.29 |
| GTEX-13FHO-3026-SM-5J1O9 | Brain - Cortex | TCGA-LGG | 26.15 | 7.9 |
| GTEX-13FHP-3026-SM-5IJBS | Brain - Cortex | TCGA-LGG | 17.72 | 6.75 |

| | | | | |
|---|---|---|---|---|
| GTEX-13FLW-1426-SM-5K7YE | Brain - Cortex | TCGA-LGG | 25.71 | 5.84 |
| GTEX-13FXS-3126-SM-5J2NN | Brain - Cortex | TCGA-LGG | 24.6 | 7.55 |
| GTEX-13G51-3026-SM-5IJB8 | Brain - Cortex | TCGA-LGG | 18.11 | 8.53 |
| GTEX-13IVO-2926-SM-5L3CZ | Brain - Cortex | TCGA-LGG | 19.55 | 5.45 |
| GTEX-13JUV-2926-SM-5LZX7 | Brain - Cortex | TCGA-LGG | 14.94 | 4.63 |
| GTEX-13JVG-3126-SM-5L3FH | Brain - Cortex | TCGA-LGG | 19.05 | 7.78 |
| GTEX-13NYB-3026-SM-5IJD7 | Brain - Cortex | TCGA-LGG | 25.19 | 7.3 |
| GTEX-13NYC-2826-SM-5K7WR | Brain - Cortex | TCGA-LGG | 34.28 | 10.16 |
| GTEX-13NYS-3126-SM-5KLYV | Brain - Cortex | TCGA-LGG | 30.07 | 8.29 |
| GTEX-13O3O-3126-SM-5KM3H | Brain - Cortex | TCGA-LGG | 23.19 | 4.9 |
| GTEX-13O3Q-2926-SM-5KM45 | Brain - Cortex | TCGA-LGG | 19.16 | 7.07 |
| GTEX-13OVH-3026-SM-5MR4N | Brain - Cortex | TCGA-LGG | 25.56 | 5.14 |
| GTEX-13OVJ-2826-SM-5L3GW | Brain - Cortex | TCGA-LGG | 27.45 | 8.67 |
| GTEX-13OW6-3026-SM-5J2MI | Brain - Cortex | TCGA-LGG | 28 | 8.62 |
| GTEX-13OW7-3026-SM-5L3GY | Brain - Cortex | TCGA-LGG | 15.23 | 5.41 |
| GTEX-13OW8-2826-SM-5L3GC | Brain - Cortex | TCGA-LGG | 20.06 | 5.8 |
| GTEX-13PL6-3126-SM-5LUAR | Brain - Cortex | TCGA-LGG | 22.3 | 8.09 |
| GTEX-13PVQ-3026-SM-5SI93 | Brain - Cortex | TCGA-LGG | 20.61 | 7.25 |
| GTEX-13QIC-2926-SM-5J2NF | Brain - Cortex | TCGA-LGG | 22.18 | 6.28 |
| GTEX-13S7M-3126-SM-5RQJQ | Brain - Cortex | TCGA-LGG | 20.78 | 3.93 |
| GTEX-13SLX-3126-SM-5S2Q5 | Brain - Cortex | TCGA-LGG | 27.75 | 6.53 |
| GTEX-13VXU-2926-SM-5LU5C | Brain - Cortex | TCGA-LGG | 19.82 | 6.63 |
| GTEX-13X6J-3026-SM-5Q5CU | Brain - Cortex | TCGA-LGG | 16.01 | 4.31 |
| GTEX-13X6K-2926-SM-5Q5D5 | Brain - Cortex | TCGA-LGG | 16.92 | 5.31 |
| GTEX-1445S-3026-SM-5O9BR | Brain - Cortex | TCGA-LGG | 13.51 | 4.74 |
| GTEX-144FL-3026-SM-5O99C | Brain - Cortex | TCGA-LGG | 24.07 | 7.68 |
| GTEX-144GL-3026-SM-5Q5CW | Brain - Cortex | TCGA-LGG | 22.66 | 5.08 |
| GTEX-145LS-3126-SM-5Q5BY | Brain - Cortex | TCGA-LGG | 22.17 | 7.14 |
| GTEX-145MF-2726-SM-5O995 | Brain - Cortex | TCGA-LGG | 17.92 | 5.71 |
| GTEX-145MG-3026-SM-5RQJA | Brain - Cortex | TCGA-LGG | 25.16 | 7 |
| GTEX-145MH-3026-SM-5Q5DZ | Brain - Cortex | TCGA-LGG | 25.1 | 7.67 |

| | | | | |
|---|---|---|---|---|
| GTEX-147GR-3026-SM-5S2ML | Brain - Cortex | TCGA-LGG | 10.72 | 6.78 |
| GTEX-14A5I-2926-SM-5Q5CQ | Brain - Cortex | TCGA-LGG | 23.81 | 5.75 |
| GTEX-14ASI-3026-SM-5S2PN | Brain - Cortex | TCGA-LGG | 23.64 | 6.45 |
| GTEX-14BIL-3026-SM-7EWD4 | Brain - Cortex | TCGA-LGG | 15.85 | 5.92 |
| GTEX-14BIM-3026-SM-7EWCY | Brain - Cortex | TCGA-LGG | 23.45 | 5.55 |
| GTEX-14BIN-3226-SM-62LDR | Brain - Cortex | TCGA-LGG | 21.25 | 6.35 |
| GTEX-14BMV-3026-SM-5S2PQ | Brain - Cortex | TCGA-LGG | 29.03 | 8.48 |
| GTEX-14C39-3126-SM-5ZZW6 | Brain - Cortex | TCGA-LGG | 21.54 | 5.99 |
| GTEX-14C5O-3026-SM-5YYB2 | Brain - Cortex | TCGA-LGG | 21.96 | 3.76 |
| GTEX-14DAQ-3126-SM-62LDS | Brain - Cortex | TCGA-LGG | 27.54 | 6.01 |
| GTEX-14JG1-3026-SM-5YYAA | Brain - Cortex | TCGA-LGG | 21.85 | 5.87 |
| GTEX-14JIY-2926-SM-69LPT | Brain - Cortex | TCGA-LGG | 23.77 | 7.7 |
| GTEX-14LZ3-3026-SM-5ZZVU | Brain - Cortex | TCGA-LGG | 21.95 | 5.5 |
| GTEX-14PJM-3126-SM-6EU2R | Brain - Cortex | TCGA-LGG | 22.68 | 6.69 |
| GTEX-15DCD-2926-SM-7KULT | Brain - Cortex | TCGA-LGG | 23.38 | 7.31 |
| GTEX-15EO6-3026-SM-7KUMA | Brain - Cortex | TCGA-LGG | 30.05 | 6.28 |
| GTEX-15ER7-3126-SM-7KUGH | Brain - Cortex | TCGA-LGG | 23.23 | 6.29 |
| GTEX-15G19-2926-SM-7KUFL | Brain - Cortex | TCGA-LGG | 29.63 | 10.26 |
| GTEX-15RJE-3026-SM-7KUM1 | Brain - Cortex | TCGA-LGG | 17.37 | 4.72 |
| GTEX-15SHU-3126-SM-6M46H | Brain - Cortex | TCGA-LGG | 17.89 | 7.37 |
| GTEX-15UF6-3026-SM-7KUKL | Brain - Cortex | TCGA-LGG | 13.54 | 4.34 |
| GTEX-16GPK-3026-SM-79ON9 | Brain - Cortex | TCGA-LGG | 12.38 | 4.57 |
| GTEX-16XZZ-3026-SM-79OMJ | Brain - Cortex | TCGA-LGG | 17.72 | 5.25 |
| GTEX-16YQH-3026-SM-79OMM | Brain - Cortex | TCGA-LGG | 16.14 | 4.36 |
| GTEX-16Z82-3026-SM-79ON3 | Brain - Cortex | TCGA-LGG | 17.1 | 6.18 |
| GTEX-17EVP-3026-SM-7IGQL | Brain - Cortex | TCGA-LGG | 13.35 | 4.54 |
| GTEX-17EVQ-2826-SM-7IGQK | Brain - Cortex | TCGA-LGG | 20.67 | 5.9 |
| GTEX-17F97-3026-SM-79OKC | Brain - Cortex | TCGA-LGG | 23.99 | 7.11 |
| GTEX-17HG3-2926-SM-79ON8 | Brain - Cortex | TCGA-LGG | 15.63 | 5.26 |
| GTEX-17HHY-3026-SM-79OKP | Brain - Cortex | TCGA-LGG | 23.06 | 6.62 |
| GTEX-17HII-3026-SM-7EWFM | Brain - Cortex | TCGA-LGG | 14.9 | 5.24 |

| | | | | |
|---|---|---|---|---|
| GTEX-17JCI-3126-SM-7LTA2 | Brain - Cortex | TCGA-LGG | 20.46 | 7.24 |
| GTEX-17MF6-2926-SM-7LTAW | Brain - Cortex | TCGA-LGG | 9.46 | 2.94 |
| GTEX-183WM-2826-SM-731C2 | Brain - Cortex | TCGA-LGG | 15.7 | 6.91 |
| GTEX-18465-3026-SM-7LG6Y | Brain - Cortex | TCGA-LGG | 24.07 | 5.22 |
| GTEX-18A6Q-2926-SM-718AR | Brain - Cortex | TCGA-LGG | 19.13 | 6.21 |
| GTEX-1A3MX-3026-SM-731BF | Brain - Cortex | TCGA-LGG | 20.68 | 5.29 |
| GTEX-1A8G6-2926-SM-731CK | Brain - Cortex | TCGA-LGG | 14.6 | 5.07 |
| GTEX-1A8G7-3126-SM-7IGP7 | Brain - Cortex | TCGA-LGG | 13.74 | 5.87 |
| GTEX-1AX9I-3026-SM-73KUH | Brain - Cortex | TCGA-LGG | 16.19 | 4.27 |
| GTEX-1B8L1-3026-SM-7EPHK | Brain - Cortex | TCGA-LGG | 17.58 | 4.84 |
| GTEX-1B8SF-3126-SM-73KU1 | Brain - Cortex | TCGA-LGG | 19.25 | 5.98 |
| GTEX-1B8SG-1726-SM-73KXK | Brain - Cortex | TCGA-LGG | 8.66 | 4.4 |
| GTEX-1B933-3126-SM-7P8RL | Brain - Cortex | TCGA-LGG | 21.43 | 5.25 |
| GTEX-1C64N-3026-SM-7MGVU | Brain - Cortex | TCGA-LGG | 8.28 | 2.87 |
| GTEX-1C6VQ-3026-SM-7IGPU | Brain - Cortex | TCGA-LGG | 14.67 | 6.66 |
| GTEX-1C6VS-3026-SM-79OLA | Brain - Cortex | TCGA-LGG | 22.83 | 7.98 |
| GTEX-1C6WA-3026-SM-7P8RQ | Brain - Cortex | TCGA-LGG | 15.81 | 4.23 |
| GTEX-1CAV2-2726-SM-7IGNN | Brain - Cortex | TCGA-LGG | 5.81 | 2.24 |
| GTEX-1CB4H-3126-SM-7IGN4 | Brain - Cortex | TCGA-LGG | 21.67 | 6.69 |
| GTEX-1E1VI-3026-SM-7RHHU | Brain - Cortex | TCGA-LGG | 21.41 | 6.47 |
| GTEX-1E2YA-3026-SM-7IGO7 | Brain - Cortex | TCGA-LGG | 10.11 | 3.21 |
| GTEX-1EKGG-2926-SM-7EPIH | Brain - Cortex | TCGA-LGG | 19.2 | 5.03 |
| GTEX-1EMGI-3026-SM-7P8P6 | Brain - Cortex | TCGA-LGG | 24.18 | 7.11 |
| GTEX-1EN7A-3026-SM-7PBYT | Brain - Cortex | TCGA-LGG | 9.48 | 3.89 |
| GTEX-1EWIQ-3126-SM-7MXTI | Brain - Cortex | TCGA-LGG | 19.63 | 6.3 |
| GTEX-1EX96-3026-SM-7RHH5 | Brain - Cortex | TCGA-LGG | 21.38 | 6.57 |
| GTEX-1F48J-2826-SM-7P8R4 | Brain - Cortex | TCGA-LGG | 24.67 | 6.89 |
| GTEX-1F52S-3126-SM-7PC3M | Brain - Cortex | TCGA-LGG | 12.65 | 4.03 |
| GTEX-1F6I4-3126-SM-7P8SD | Brain - Cortex | TCGA-LGG | 10.82 | 3.54 |
| GTEX-1F6RS-3126-SM-9MQK7 | Brain - Cortex | TCGA-LGG | 24.41 | 6.91 |
| GTEX-1F75A-3026-SM-9KNU2 | Brain - Cortex | TCGA-LGG | 22.91 | 5.78 |

| | | | | |
|---|---|---|---|---|
| GTEX-1F75B-3026-SM-7PBZ7 | Brain - Cortex | TCGA-LGG | 21.15 | 5.93 |
| GTEX-1F7RK-1826-SM-7RHI4 | Brain - Cortex | TCGA-LGG | 15.71 | 7.77 |
| GTEX-1F88E-2826-SM-7RHHE | Brain - Cortex | TCGA-LGG | 12.39 | 4.75 |
| GTEX-1GF9V-2926-SM-7PBXO | Brain - Cortex | TCGA-LGG | 17.21 | 5.47 |
| GTEX-1GF9W-2626-SM-7P8TA | Brain - Cortex | TCGA-LGG | 22.43 | 4.41 |
| GTEX-1GMR8-3026-SM-9KNVF | Brain - Cortex | TCGA-LGG | 22.61 | 4.42 |
| GTEX-1GMRU-3026-SM-7P8TC | Brain - Cortex | TCGA-LGG | 27.13 | 7.91 |
| GTEX-1GN1V-3126-SM-7PBZG | Brain - Cortex | TCGA-LGG | 7.69 | 3.23 |
| GTEX-1GN2E-3026-SM-9WG6I | Brain - Cortex | TCGA-LGG | 13.86 | 3.31 |
| GTEX-1GN73-3126-SM-9KNTY | Brain - Cortex | TCGA-LGG | 22.75 | 5.76 |
| GTEX-1GZ2Q-3226-SM-9WG6L | Brain - Cortex | TCGA-LGG | 22.52 | 5.52 |
| GTEX-1GZ4I-3026-SM-9KNTX | Brain - Cortex | TCGA-LGG | 27.2 | 6.97 |
| GTEX-1GZHY-3126-SM-9QEHH | Brain - Cortex | TCGA-LGG | 7.85 | 3.5 |
| GTEX-1H1DG-2926-SM-9WYU3 | Brain - Cortex | TCGA-LGG | 19.83 | 9.44 |
| GTEX-1H1ZS-3026-SM-9KNUJ | Brain - Cortex | TCGA-LGG | 24.78 | 6.56 |
| GTEX-1H23P-2926-SM-9WYU8 | Brain - Cortex | TCGA-LGG | 16.63 | 4.25 |
| GTEX-1H2FU-3126-SM-9WYUJ | Brain - Cortex | TCGA-LGG | 24.13 | 6.24 |
| GTEX-1H3NZ-3026-SM-9JGHW | Brain - Cortex | TCGA-LGG | 25.09 | 7.87 |
| GTEX-1H3O1-1726-SM-9WYSR | Brain - Cortex | TCGA-LGG | 13.16 | 5.96 |
| GTEX-1H3VE-2726-SM-9WPP3 | Brain - Cortex | TCGA-LGG | 11.53 | 4.07 |
| GTEX-1HB9E-2926-SM-D4P3S | Brain - Cortex | TCGA-LGG | 17.39 | 7.77 |
| GTEX-1HBPH-3126-SM-9WYUV | Brain - Cortex | TCGA-LGG | 18.65 | 5.22 |
| GTEX-1HBPI-2826-SM-ARZNO | Brain - Cortex | TCGA-LGG | 15.62 | 5.87 |
| GTEX-1HBPM-2926-SM-CL54E | Brain - Cortex | TCGA-LGG | 6.53 | 2.75 |
| GTEX-1HCU6-3126-SM-CL54L | Brain - Cortex | TCGA-LGG | 30.36 | 7.38 |
| GTEX-1HCVE-3026-SM-CL54G | Brain - Cortex | TCGA-LGG | 21.07 | 7.55 |
| GTEX-1HFI6-3026-SM-CNPPJ | Brain - Cortex | TCGA-LGG | 15.25 | 6.87 |
| GTEX-1HGF4-3126-SM-CL54M | Brain - Cortex | TCGA-LGG | 15.95 | 6.31 |
| GTEX-1HR98-3026-SM-CNPPK | Brain - Cortex | TCGA-LGG | 21.75 | 6.81 |
| GTEX-1HSKV-3026-SM-A96SL | Brain - Cortex | TCGA-LGG | 16.22 | 3.68 |
| GTEX-1HSMO-3026-SM-A9G3F | Brain - Cortex | TCGA-LGG | 23.81 | 6.92 |

| | | | | |
|---|---|---|---|---|
| GTEX-1HT8W-3026-SM-CGQEO | Brain - Cortex | TCGA-LGG | 24.97 | 6.21 |
| GTEX-1HUB1-2926-SM-ACKXA | Brain - Cortex | TCGA-LGG | 15.38 | 4.2 |
| GTEX-1I19N-3026-SM-CL55E | Brain - Cortex | TCGA-LGG | 18.7 | 4.88 |
| GTEX-1I1CD-3026-SM-B2LY2 | Brain - Cortex | TCGA-LGG | 17.68 | 6.7 |
| GTEX-1I1GQ-3026-SM-CL55K | Brain - Cortex | TCGA-LGG | 15.67 | 5.32 |
| GTEX-1I1GR-2926-SM-CNNQG | Brain - Cortex | TCGA-LGG | 22.82 | 6.06 |
| GTEX-1I1GV-3126-SM-A96SF | Brain - Cortex | TCGA-LGG | 15.12 | 3.77 |
| GTEX-1I1HK-2826-SM-CKZPL | Brain - Cortex | TCGA-LGG | 18.79 | 5.94 |
| GTEX-1ICLZ-2826-SM-CL53U | Brain - Cortex | TCGA-LGG | 16.34 | 6.06 |
| GTEX-1IDFM-2826-SM-ACKWJ | Brain - Cortex | TCGA-LGG | 8.2 | 3.77 |
| GTEX-1IDJH-3026-SM-CL53B | Brain - Cortex | TCGA-LGG | 23.22 | 7.6 |
| GTEX-1IKJJ-2926-SM-A9G2N | Brain - Cortex | TCGA-LGG | 10.79 | 3.32 |
| GTEX-1IKK5-2926-SM-ARU7Q | Brain - Cortex | TCGA-LGG | 23.26 | 7.41 |
| GTEX-1IL2U-3026-SM-C1YSH | Brain - Cortex | TCGA-LGG | 20.16 | 6.28 |
| GTEX-1IOXB-3126-SM-CKZO1 | Brain - Cortex | TCGA-LGG | 8.22 | 2.47 |
| GTEX-1IY9M-3026-SM-A9G37 | Brain - Cortex | TCGA-LGG | 17.92 | 5.61 |
| GTEX-1J1OQ-3226-SM-A9G2Q | Brain - Cortex | TCGA-LGG | 20.13 | 5.24 |
| GTEX-1J8Q2-3026-SM-CKZN8 | Brain - Cortex | TCGA-LGG | 22.91 | 5.77 |
| GTEX-1JJ6O-3126-SM-CNNP8 | Brain - Cortex | TCGA-LGG | 22.5 | 5.3 |
| GTEX-1JJE9-2926-SM-CY8HV | Brain - Cortex | TCGA-LGG | 6.12 | 2.45 |
| GTEX-1JJEA-2826-SM-ARU8R | Brain - Cortex | TCGA-LGG | 15.66 | 3.84 |
| GTEX-1JMLX-2926-SM-CL54C | Brain - Cortex | TCGA-LGG | 16.56 | 5.02 |
| GTEX-1JMOU-2726-SM-CY8IO | Brain - Cortex | TCGA-LGG | 21.98 | 7.98 |
| GTEX-1JMPZ-2726-SM-ARZNF | Brain - Cortex | TCGA-LGG | 21.98 | 5.67 |
| GTEX-1JMQK-3026-SM-ARL95 | Brain - Cortex | TCGA-LGG | 19.72 | 5 |
| GTEX-1JMQL-2826-SM-ARU91 | Brain - Cortex | TCGA-LGG | 24.8 | 7.32 |
| GTEX-1KD4Q-2926-SM-D3L9N | Brain - Cortex | TCGA-LGG | 23.85 | 7.16 |
| GTEX-1KWVE-2326-SM-D4P3N | Brain - Cortex | TCGA-LGG | 22.73 | 5.35 |
| GTEX-1LC47-2926-SM-CXZKC | Brain - Cortex | TCGA-LGG | 20.14 | 5.54 |
| GTEX-1LG7Y-3026-SM-D5OVI | Brain - Cortex | TCGA-LGG | 23.77 | 5.94 |
| GTEX-1LKK1-3126-SM-EWRNY | Brain - Cortex | TCGA-LGG | 20.33 | 5.44 |

| | | | | |
|---|---|---|---|---|
| GTEX-1M4P7-2726-SM-E9TJQ | Brain - Cortex | TCGA-LGG | 11.91 | 4.14 |
| GTEX-1N2DW-2826-SM-E9TK5 | Brain - Cortex | TCGA-LGG | 13.11 | 4.67 |
| GTEX-1N5O9-2926-SM-EXUSN | Brain - Cortex | TCGA-LGG | 16.67 | 4.27 |
| GTEX-1NV5F-3226-SM-EXUSL | Brain - Cortex | TCGA-LGG | 14.57 | 3.53 |
| GTEX-1NV8Z-3126-SM-E76R3 | Brain - Cortex | TCGA-LGG | 19.48 | 5.66 |
| GTEX-1O9I2-3026-SM-E76PX | Brain - Cortex | TCGA-LGG | 17.31 | 5.89 |
| GTEX-1OJC4-2826-SM-E6CQX | Brain - Cortex | TCGA-LGG | 23.69 | 4.97 |
| GTEX-1PIIG-2726-SM-E76PQ | Brain - Cortex | TCGA-LGG | 26.66 | 6.95 |
| GTEX-1PPGY-3126-SM-EXUSR | Brain - Cortex | TCGA-LGG | 22.75 | 5.31 |
| GTEX-1PWST-3126-SM-EXUSE | Brain - Cortex | TCGA-LGG | 21.64 | 6.35 |
| GTEX-1QMI2-3126-SM-EXUT5 | Brain - Cortex | TCGA-LGG | 25.04 | 7.21 |
| GTEX-1QP6S-2926-SM-DTXF8 | Brain - Cortex | TCGA-LGG | 15.77 | 8.64 |
| GTEX-1R9K4-3126-SM-DTX9Y | Brain - Cortex | TCGA-LGG | 20.88 | 5.22 |
| GTEX-1R9PO-2926-SM-EVYBR | Brain - Cortex | TCGA-LGG | 29.88 | 8.75 |
| GTEX-1RQEC-2926-SM-EVR5A | Brain - Cortex | TCGA-LGG | 18.1 | 4.81 |
| GTEX-N7MT-1126-SM-CYPQY | Brain - Cortex | TCGA-LGG | 13.84 | 7.01 |
| GTEX-NPJ7-2726-SM-2I3FT | Brain - Cortex | TCGA-LGG | 16.46 | 5.27 |
| GTEX-NPJ8-1526-SM-2D7VU | Brain - Cortex | TCGA-LGG | 27.45 | 8.62 |
| GTEX-OHPN-2526-SM-2I5H3 | Brain - Cortex | TCGA-LGG | 30.61 | 8.54 |
| GTEX-OXRN-2426-SM-2I5EQ | Brain - Cortex | TCGA-LGG | 17.93 | 3.88 |
| GTEX-P44G-2426-SM-2I5ER | Brain - Cortex | TCGA-LGG | 20.86 | 7.12 |
| GTEX-PVOW-2526-SM-2XCF7 | Brain - Cortex | TCGA-LGG | 25.06 | 8.18 |
| GTEX-PWO3-0926-SM-2I5EY | Brain - Cortex | TCGA-LGG | 12.21 | 3.98 |
| GTEX-Q2AG-2926-SM-2HMJ3 | Brain - Cortex | TCGA-LGG | 13.96 | 4.8 |
| GTEX-QDT8-2926-SM-32PKC | Brain - Cortex | TCGA-LGG | 21.73 | 7.94 |
| GTEX-QMR6-1426-SM-32PLA | Brain - Cortex | TCGA-LGG | 20.04 | 5.51 |
| GTEX-QVJO-1426-SM-2S1QY | Brain - Cortex | TCGA-LGG | 21.01 | 6.08 |
| GTEX-QVUS-2826-SM-3GADB | Brain - Cortex | TCGA-LGG | 24.87 | 6 |
| GTEX-RNOR-2326-SM-2TF4I | Brain - Cortex | TCGA-LGG | 23.5 | 6.82 |
| GTEX-RU72-3026-SM-CYPRV | Brain - Cortex | TCGA-LGG | 22.22 | 5.62 |
| GTEX-T2IS-3026-SM-32QPM | Brain - Cortex | TCGA-LGG | 19.66 | 6.41 |

| | | | | |
|---|---|---|---|---|
| GTEX-T5JC-2426-SM-3NMDB | Brain - Cortex | TCGA-LGG | 22.66 | 7.09 |
| GTEX-T6MN-2626-SM-32PMQ | Brain - Cortex | TCGA-LGG | 24.96 | 5.29 |
| GTEX-TSE9-3026-SM-3DB76 | Brain - Cortex | TCGA-LGG | 19.9 | 5.29 |
| GTEX-UTHO-3026-SM-3GAFB | Brain - Cortex | TCGA-LGG | 26.71 | 8.68 |
| GTEX-WHSE-3026-SM-3P5ZH | Brain - Cortex | TCGA-LGG | 22.28 | 6.51 |
| GTEX-WL46-2926-SM-3LK82 | Brain - Cortex | TCGA-LGG | 22.27 | 5.45 |
| GTEX-WVLH-3026-SM-3MJG9 | Brain - Cortex | TCGA-LGG | 23.1 | 5.78 |
| GTEX-WWYW-3126-SM-3NB39 | Brain - Cortex | TCGA-LGG | 24.09 | 7.88 |
| GTEX-WZTO-2926-SM-3NM9I | Brain - Cortex | TCGA-LGG | 26.32 | 7.94 |
| GTEX-X4XX-3026-SM-3NMB2 | Brain - Cortex | TCGA-LGG | 28.92 | 6.48 |
| GTEX-X4XY-3026-SM-EWRM9 | Brain - Cortex | TCGA-LGG | 18.82 | 5.92 |
| GTEX-X585-3026-SM-46MWF | Brain - Cortex | TCGA-LGG | 34.45 | 10.35 |
| GTEX-XLM4-3026-SM-4AT6L | Brain - Cortex | TCGA-LGG | 24.05 | 6.72 |
| GTEX-Y111-2726-SM-4TT3N | Brain - Cortex | TCGA-LGG | 14.2 | 5.81 |
| GTEX-Y8DK-0826-SM-4TT3T | Brain - Cortex | TCGA-LGG | 21.65 | 6.38 |
| GTEX-YFC4-3126-SM-5PNV6 | Brain - Cortex | TCGA-LGG | 24.01 | 7.44 |
| GTEX-YJ89-3026-SM-5IFJI | Brain - Cortex | TCGA-LGG | 19.4 | 5.45 |
| GTEX-Z93S-2926-SM-57WB9 | Brain - Cortex | TCGA-LGG | 15.57 | 4.46 |
| GTEX-ZAJG-3126-SM-5HL9J | Brain - Cortex | TCGA-LGG | 28.02 | 5.51 |
| GTEX-ZAK1-3026-SM-5S2MJ | Brain - Cortex | TCGA-LGG | 24.39 | 7.11 |
| GTEX-ZDXO-2126-SM-4WKFI | Brain - Cortex | TCGA-LGG | 16.03 | 5.14 |
| GTEX-ZE7O-3126-SM-5HL5X | Brain - Cortex | TCGA-LGG | 21.09 | 6.25 |
| GTEX-ZE9C-3026-SM-4WKHB | Brain - Cortex | TCGA-LGG | 24.8 | 6.03 |
| GTEX-ZF28-3026-SM-4WKHP | Brain - Cortex | TCGA-LGG | 20.95 | 6.68 |
| GTEX-ZUA1-3026-SM-59HJC | Brain - Cortex | TCGA-LGG | 21.44 | 4.65 |
| GTEX-ZVT3-3026-SM-5E43N | Brain - Cortex | TCGA-LGG | 26.35 | 8.9 |
| GTEX-ZYFD-3026-SM-5E44C | Brain - Cortex | TCGA-LGG | 23.04 | 6.24 |
| GTEX-ZYY3-3126-SM-5SI9L | Brain - Cortex | TCGA-LGG | 27.85 | 8.34 |
| GTEX-ZZPT-3026-SM-5GZXH | Brain - Cortex | TCGA-LGG | 18.54 | 4.48 |
| GTEX-1192X-0011-R10a-SM-DO941 | Brain - Frontal Cortex (BA | TCGA-LGG | 41.07 | 9.75 |
| GTEX-11DXY-0011-R10b-SM-DO12 | Brain - Frontal Cortex (BA | TCGA-LGG | 30.44 | 6.95 |

| | | | | |
|---|---|---|---|---|
| GTEX-11DYG-0011-R10b-SM-DNZZ | Brain - Frontal Cortex (B | TCGA-LGG | 27.08 | 7.24 |
| GTEX-11DZ1-0011-R10b-SM-DO94 | Brain - Frontal Cortex (B | TCGA-LGG | 32.88 | 7.99 |
| GTEX-11EI6-0011-R10a-SM-DO93R | Brain - Frontal Cortex (B | TCGA-LGG | 16.89 | 6.02 |
| GTEX-11EMC-0011-R10a-SM-DNZZ | Brain - Frontal Cortex (B | TCGA-LGG | 29.38 | 7.94 |
| GTEX-11GSP-0011-R10a-SM-5NQ7 | Brain - Frontal Cortex (B | TCGA-LGG | 18.28 | 5.45 |
| GTEX-11H98-0011-R10b-SM-5NQ9 | Brain - Frontal Cortex (B | TCGA-LGG | 33.18 | 9.07 |
| GTEX-11PRG-0011-R10b-SM-DO93 | Brain - Frontal Cortex (B | TCGA-LGG | 27.01 | 7.05 |
| GTEX-11TTK-0011-R10b-SM-DO92 | Brain - Frontal Cortex (B | TCGA-LGG | 15.89 | 8.53 |
| GTEX-11TUW-0011-R10a-SM-DO92 | Brain - Frontal Cortex (B | TCGA-LGG | 27.07 | 13.28 |
| GTEX-11WQC-0011-R10a-SM-57W | Brain - Frontal Cortex (B | TCGA-LGG | 17.41 | 6.69 |
| GTEX-11WQK-0011-R10a-SM-5BC6 | Brain - Frontal Cortex (B | TCGA-LGG | 19.31 | 5.64 |
| GTEX-11ZTS-0011-R10a-SM-DO93 | Brain - Frontal Cortex (B | TCGA-LGG | 22.2 | 6.4 |
| GTEX-12126-0011-R10b-SM-5BC6T | Brain - Frontal Cortex (B | TCGA-LGG | 20.55 | 7.5 |
| GTEX-12WSA-0011-R10b-SM-5P9E | Brain - Frontal Cortex (B | TCGA-LGG | 19.09 | 3.72 |
| GTEX-12WSC-0011-R10a-SM-5GU | Brain - Frontal Cortex (B | TCGA-LGG | 32.07 | 7.21 |
| GTEX-12WSE-0011-R10b-SM-5P9J | Brain - Frontal Cortex (B | TCGA-LGG | 27.97 | 8.18 |
| GTEX-12WSF-0011-R10a-SM-5LZV | Brain - Frontal Cortex (B | TCGA-LGG | 30.56 | 9.57 |
| GTEX-12WSH-0011-R10a-SM-5LZU | Brain - Frontal Cortex (B | TCGA-LGG | 27.94 | 7.86 |
| GTEX-12WSM-0011-R10a-SM-5DU | Brain - Frontal Cortex (B | TCGA-LGG | 27.11 | 8.18 |
| GTEX-12ZZW-0011-R10b-SM-5HL9 | Brain - Frontal Cortex (B | TCGA-LGG | 16.02 | 4.7 |
| GTEX-12ZZX-0011-R10b-SM-5DUW | Brain - Frontal Cortex (B | TCGA-LGG | 35.8 | 9.66 |
| GTEX-12ZZY-0011-R10b-SM-5HL7 | Brain - Frontal Cortex (B | TCGA-LGG | 38.45 | 11.28 |
| GTEX-12ZZZ-0011-R10a-SM-5P9H | Brain - Frontal Cortex (B | TCGA-LGG | 24.38 | 6.31 |
| GTEX-1313W-0011-R10b-SM-5DUX | Brain - Frontal Cortex (B | TCGA-LGG | 25.79 | 7.96 |
| GTEX-131XH-0011-R10a-SM-5EGL | Brain - Frontal Cortex (B | TCGA-LGG | 26.94 | 7.59 |
| GTEX-131XW-0011-R10a-SM-5DU | Brain - Frontal Cortex (B | TCGA-LGG | 8.9 | 3.28 |
| GTEX-131YS-0011-R10b-SM-5EQ5 | Brain - Frontal Cortex (B | TCGA-LGG | 27 | 6.05 |
| GTEX-132Q8-0011-R10b-SM-5DUW | Brain - Frontal Cortex (B | TCGA-LGG | 27.94 | 5.04 |
| GTEX-139TS-0011-R10a-SM-5K7TU | Brain - Frontal Cortex (B | TCGA-LGG | 17.44 | 7.09 |
| GTEX-13CZV-0011-R10b-SM-5LZY | Brain - Frontal Cortex (B | TCGA-LGG | 30.03 | 7.25 |
| GTEX-13FHO-0011-R10b-SM-5J2M | Brain - Frontal Cortex (B | TCGA-LGG | 30 | 6.36 |

| | | | | |
|---|---|---|---|---|
| GTEX-13FLV-0011-R10b-SM-5LZZ2 | Brain - Frontal Cortex (B. | TCGA-LGG | 23.61 | 7.85 |
| GTEX-13FLW-0011-R10b-SM-5LZZI | Brain - Frontal Cortex (B. | TCGA-LGG | 29.28 | 6.42 |
| GTEX-13FTZ-0011-R10b-SM-5KLZS | Brain - Frontal Cortex (B. | TCGA-LGG | 20.07 | 5.19 |
| GTEX-13FXS-0011-R10a-SM-5J2M. | Brain - Frontal Cortex (B. | TCGA-LGG | 23.88 | 5.2 |
| GTEX-13G51-0011-R10b-SM-5LZYI | Brain - Frontal Cortex (B. | TCGA-LGG | 16.64 | 4.63 |
| GTEX-13IVO-0011-R10a-SM-5LZY3 | Brain - Frontal Cortex (B. | TCGA-LGG | 13.78 | 2.66 |
| GTEX-13JUV-0011-R10b-SM-5LZXI | Brain - Frontal Cortex (B. | TCGA-LGG | 19.86 | 7.02 |
| GTEX-13JVG-0011-R10b-SM-5KM2 | Brain - Frontal Cortex (B. | TCGA-LGG | 13.2 | 8.1 |
| GTEX-13N1W-0011-R10b-SM-5MR4 | Brain - Frontal Cortex (B. | TCGA-LGG | 30.78 | 8.03 |
| GTEX-13N2G-0011-R10a-SM-5MR3 | Brain - Frontal Cortex (B. | TCGA-LGG | 61.29 | 20.02 |
| GTEX-13NYB-0011-R10a-SM-5KM4 | Brain - Frontal Cortex (B. | TCGA-LGG | 26.84 | 6.56 |
| GTEX-13NYS-0011-R10a-SM-5MR4 | Brain - Frontal Cortex (B. | TCGA-LGG | 43.03 | 7.7 |
| GTEX-13NZA-0011-R10b-SM-5KM5 | Brain - Frontal Cortex (B. | TCGA-LGG | 12.43 | 4.3 |
| GTEX-13O3O-0011-R10a-SM-5LUA | Brain - Frontal Cortex (B. | TCGA-LGG | 33.7 | 7.11 |
| GTEX-13O3Q-0011-R10b-SM-5KM3 | Brain - Frontal Cortex (B. | TCGA-LGG | 33.13 | 9.74 |
| GTEX-13OVJ-0011-R10b-SM-5L3H | Brain - Frontal Cortex (B. | TCGA-LGG | 24.82 | 9.61 |
| GTEX-13OVL-0011-R10a-SM-5L3G | Brain - Frontal Cortex (B. | TCGA-LGG | 22.66 | 8.26 |
| GTEX-13OW6-0011-R10a-SM-5L3H | Brain - Frontal Cortex (B. | TCGA-LGG | 31.31 | 10.97 |
| GTEX-13OW7-0011-R10a-SM-5L3H | Brain - Frontal Cortex (B. | TCGA-LGG | 27.76 | 8.22 |
| GTEX-13OW8-0011-R10a-SM-5L3H | Brain - Frontal Cortex (B. | TCGA-LGG | 19.36 | 5.08 |
| GTEX-13QIC-0011-R10a-SM-5O9C | Brain - Frontal Cortex (B. | TCGA-LGG | 30.26 | 10.57 |
| GTEX-13QJC-0011-R10a-SM-5SI63 | Brain - Frontal Cortex (B. | TCGA-LGG | 39.69 | 6.56 |
| GTEX-13RTJ-0011-R10b-SM-5O9C | Brain - Frontal Cortex (B. | TCGA-LGG | 35.17 | 6.63 |
| GTEX-13S7M-0011-R10b-SM-5PNZ | Brain - Frontal Cortex (B. | TCGA-LGG | 32.32 | 8.51 |
| GTEX-13SLW-0011-R10a-SM-5S2U | Brain - Frontal Cortex (B. | TCGA-LGG | 23.95 | 5.23 |
| GTEX-13SLX-0011-R10a-SM-5P9H | Brain - Frontal Cortex (B. | TCGA-LGG | 39.06 | 8.91 |
| GTEX-13X6I-0011-R10a-SM-5PNWI | Brain - Frontal Cortex (B. | TCGA-LGG | 29.43 | 8.27 |
| GTEX-13X6J-0011-R10b-SM-5PNW | Brain - Frontal Cortex (B. | TCGA-LGG | 24.65 | 6.19 |
| GTEX-13X6K-0011-R10a-SM-5P9JI | Brain - Frontal Cortex (B. | TCGA-LGG | 31.54 | 8.77 |
| GTEX-1445S-0011-R10a-SM-5PNW | Brain - Frontal Cortex (B. | TCGA-LGG | 17.61 | 5.57 |
| GTEX-145LS-0011-R10a-SM-5PNU | Brain - Frontal Cortex (B. | TCGA-LGG | 26.93 | 8.81 |

| | | | | |
|---|---|---|---|---|
| GTEX-145LU-0011-R10a-SM-5PNW | Brain - Frontal Cortex (B/ | TCGA-LGG | 23.17 | 7.72 |
| GTEX-145MG-0011-R10a-SM-5PNV | Brain - Frontal Cortex (B/ | TCGA-LGG | 36.24 | 9.15 |
| GTEX-145MH-0011-R10b-SM-5PNU | Brain - Frontal Cortex (B/ | TCGA-LGG | 30.51 | 9.38 |
| GTEX-145MI-0011-R10a-SM-5PNZI | Brain - Frontal Cortex (B/ | TCGA-LGG | 27.82 | 8.01 |
| GTEX-14753-0011-R10b-SM-5S2WI | Brain - Frontal Cortex (B/ | TCGA-LGG | 8.98 | 2.68 |
| GTEX-1477Z-0011-R10b-SM-5S2RI | Brain - Frontal Cortex (B/ | TCGA-LGG | 11.48 | 4.06 |
| GTEX-147F4-0011-R10b-SM-5S2W | Brain - Frontal Cortex (B/ | TCGA-LGG | 17.77 | 6.3 |
| GTEX-147GR-0011-R10b-SM-5S2U | Brain - Frontal Cortex (B/ | TCGA-LGG | 25.68 | 8.6 |
| GTEX-14ASI-0011-R10b-SM-5S2VZ | Brain - Frontal Cortex (B/ | TCGA-LGG | 38.24 | 11.08 |
| GTEX-14BIL-0011-R10a-SM-5SI75 | Brain - Frontal Cortex (B/ | TCGA-LGG | 35.82 | 9.86 |
| GTEX-14BIN-0011-R10a-SM-5S2U/ | Brain - Frontal Cortex (B/ | TCGA-LGG | 29.87 | 7.23 |
| GTEX-14C5O-0011-R10a-SM-6872I | Brain - Frontal Cortex (B/ | TCGA-LGG | 16.96 | 3.56 |
| GTEX-14E6D-0011-R10a-SM-5YY9 | Brain - Frontal Cortex (B/ | TCGA-LGG | 33.23 | 11.16 |
| GTEX-14E7W-0011-R10a-SM-5YYA | Brain - Frontal Cortex (B/ | TCGA-LGG | 38.05 | 10.76 |
| GTEX-14JIY-0011-R10a-SM-6AJAO | Brain - Frontal Cortex (B/ | TCGA-LGG | 29.57 | 9.2 |
| GTEX-14LZ3-0011-R10b-SM-6AJA9 | Brain - Frontal Cortex (B/ | TCGA-LGG | 19.93 | 8.32 |
| GTEX-14PJM-0011-R10a-SM-686ZI | Brain - Frontal Cortex (B/ | TCGA-LGG | 27.97 | 8.62 |
| GTEX-14PJO-0011-R10a-SM-686ZS | Brain - Frontal Cortex (B/ | TCGA-LGG | 21.49 | 6.67 |
| GTEX-14PN4-0011-R10b-SM-69LP2 | Brain - Frontal Cortex (B/ | TCGA-LGG | 31.22 | 8.07 |
| GTEX-14PQA-0011-R10a-SM-6AJA | Brain - Frontal Cortex (B/ | TCGA-LGG | 28.39 | 6.91 |
| GTEX-15CHQ-0011-R10b-SM-6AJB | Brain - Frontal Cortex (B/ | TCGA-LGG | 30.28 | 9.01 |
| GTEX-15DCD-0011-R10b-SM-6LPII | Brain - Frontal Cortex (B/ | TCGA-LGG | 29.38 | 8.97 |
| GTEX-15DDE-0011-R10a-SM-7KUK | Brain - Frontal Cortex (B/ | TCGA-LGG | 19.32 | 3.87 |
| GTEX-15G19-0011-R10b-SM-7KUG | Brain - Frontal Cortex (B/ | TCGA-LGG | 39.44 | 13.15 |
| GTEX-15UF6-0011-R10b-SM-9OSV | Brain - Frontal Cortex (B/ | TCGA-LGG | 9.65 | 3.76 |
| GTEX-16GPK-0011-R10b-SM-7MKF | Brain - Frontal Cortex (B/ | TCGA-LGG | 16.37 | 3.47 |
| GTEX-16NPV-0011-R10a-SM-79OK | Brain - Frontal Cortex (B/ | TCGA-LGG | 16.64 | 5.44 |
| GTEX-16XZZ-0011-R10b-SM-7LT91 | Brain - Frontal Cortex (B/ | TCGA-LGG | 26.85 | 8.46 |
| GTEX-16YQH-0011-R10b-SM-7LT8I | Brain - Frontal Cortex (B/ | TCGA-LGG | 22.91 | 6.01 |
| GTEX-16Z82-0011-R10a-SM-79OJX | Brain - Frontal Cortex (B/ | TCGA-LGG | 19.18 | 6.17 |
| GTEX-17EVP-0011-R10a-SM-7IGLF | Brain - Frontal Cortex (B/ | TCGA-LGG | 34.51 | 12.92 |

| | | | | |
|---|---|---|---|---|
| GTEX-17F97-0011-R10b-SM-793BE | Brain - Frontal Cortex (B. | TCGA-LGG | 27.09 | 7.52 |
| GTEX-17HG3-0011-R10b-SM-7LT97 | Brain - Frontal Cortex (B. | TCGA-LGG | 36.77 | 9.34 |
| GTEX-17HII-0011-R10a-SM-79OMY | Brain - Frontal Cortex (B. | TCGA-LGG | 21.04 | 4.02 |
| GTEX-17JCI-0011-R10b-SM-718A2 | Brain - Frontal Cortex (B. | TCGA-LGG | 24.71 | 8.59 |
| GTEX-18464-0011-R10a-SM-72D6X | Brain - Frontal Cortex (B. | TCGA-LGG | 29.14 | 8.03 |
| GTEX-18A6Q-0011-R10b-SM-72D7. | Brain - Frontal Cortex (B. | TCGA-LGG | 31.14 | 9.4 |
| GTEX-1A3MX-0011-R10b-SM-7P8P | Brain - Frontal Cortex (B. | TCGA-LGG | 27.25 | 7.37 |
| GTEX-1A8G6-0011-R10b-SM-7P8PI | Brain - Frontal Cortex (B. | TCGA-LGG | 17.87 | 6.07 |
| GTEX-1B8L1-0011-R10b-SM-7P8Q4 | Brain - Frontal Cortex (B. | TCGA-LGG | 31.4 | 7.27 |
| GTEX-1B8SF-0011-R10b-SM-7P8S' | Brain - Frontal Cortex (B. | TCGA-LGG | 31.19 | 7.86 |
| GTEX-1B933-0011-R10b-SM-9OSVI | Brain - Frontal Cortex (B. | TCGA-LGG | 26.6 | 10.2 |
| GTEX-1B996-0011-R10b-SM-CJI3T | Brain - Frontal Cortex (B. | TCGA-LGG | 45 | 12.35 |
| GTEX-1BAJH-0011-R10a-SM-7MXT | Brain - Frontal Cortex (B. | TCGA-LGG | 19.99 | 4.28 |
| GTEX-1C6VQ-0011-R10b-SM-7P8C | Brain - Frontal Cortex (B. | TCGA-LGG | 13.91 | 6.83 |
| GTEX-1C6VR-0011-R10b-SM-7P8Q | Brain - Frontal Cortex (B. | TCGA-LGG | 20.29 | 6.1 |
| GTEX-1CB4G-0011-R10a-SM-9QEH | Brain - Frontal Cortex (B. | TCGA-LGG | 34.66 | 7.78 |
| GTEX-1CB4H-0011-R10b-SM-9QEH | Brain - Frontal Cortex (B. | TCGA-LGG | 29.89 | 8.51 |
| GTEX-1CB4J-0011-R10a-SM-9WG6 | Brain - Frontal Cortex (B. | TCGA-LGG | 59.3 | 18.39 |
| GTEX-1E1VI-0011-R10b-SM-ARU8I | Brain - Frontal Cortex (B. | TCGA-LGG | 22.47 | 7.73 |
| GTEX-1EH9U-0011-R10a-SM-CJI2A | Brain - Frontal Cortex (B. | TCGA-LGG | 17.61 | 4.78 |
| GTEX-1EMGI-0011-R10a-SM-9OSW | Brain - Frontal Cortex (B. | TCGA-LGG | 23.63 | 6.6 |
| GTEX-1EX96-0011-R10b-SM-CE6Q | Brain - Frontal Cortex (B. | TCGA-LGG | 35.26 | 10.18 |
| GTEX-1F48J-0011-R10a-SM-9QEIB | Brain - Frontal Cortex (B. | TCGA-LGG | 31.8 | 9.89 |
| GTEX-1F6I4-0011-R10b-SM-9OSWT | Brain - Frontal Cortex (B. | TCGA-LGG | 27.85 | 8.28 |
| GTEX-1F6IF-0011-R10a-SM-9QEIN | Brain - Frontal Cortex (B. | TCGA-LGG | 24.64 | 8.04 |
| GTEX-1F75W-0011-R10b-SM-ARU7 | Brain - Frontal Cortex (B. | TCGA-LGG | 19.23 | 5.81 |
| GTEX-1F7RK-0011-R10b-SM-ARU7 | Brain - Frontal Cortex (B. | TCGA-LGG | 22.33 | 6.59 |
| GTEX-1GF9V-0011-R10b-SM-CKZN | Brain - Frontal Cortex (B. | TCGA-LGG | 39.89 | 9.08 |
| GTEX-1GF9W-0011-R10a-SM-9OSX | Brain - Frontal Cortex (B. | TCGA-LGG | 33.63 | 8.61 |
| GTEX-1GMR8-0011-R10a-SM-CJI2 | Brain - Frontal Cortex (B. | TCGA-LGG | 28.14 | 7.1 |
| GTEX-1GMRU-0011-R10b-SM-CJI2 | Brain - Frontal Cortex (B. | TCGA-LGG | 31.17 | 7.78 |

| | | | | |
|---|---|---|---|---|
| GTEX-1GN1U-0011-R10b-SM-CKZN | Brain - Frontal Cortex (B. | TCGA-LGG | 39.09 | 11.99 |
| GTEX-1GN1V-0011-R10a-SM-CKZN | Brain - Frontal Cortex (B. | TCGA-LGG | 10.45 | 2.37 |
| GTEX-1GN2E-0011-R10a-SM-CE6F | Brain - Frontal Cortex (B. | TCGA-LGG | 18.94 | 5.14 |
| GTEX-1GN73-0011-R10b-SM-CKZN | Brain - Frontal Cortex (B. | TCGA-LGG | 35.04 | 10.75 |
| GTEX-1GTWX-0011-R10b-SM-CJI2 | Brain - Frontal Cortex (B. | TCGA-LGG | 22.52 | 5.77 |
| GTEX-1GZ2Q-0011-R10b-SM-CE6F | Brain - Frontal Cortex (B. | TCGA-LGG | 27.32 | 6.44 |
| GTEX-1GZ4I-0011-R10a-SM-9OSX | Brain - Frontal Cortex (B. | TCGA-LGG | 27.81 | 8.73 |
| GTEX-1H1CY-0011-R10a-SM-CE6S | Brain - Frontal Cortex (B. | TCGA-LGG | 16.12 | 5.37 |
| GTEX-1H1DG-0011-R10b-SM-CE6S | Brain - Frontal Cortex (B. | TCGA-LGG | 28.06 | 7.99 |
| GTEX-1H1ZS-0011-R10b-SM-A9SK | Brain - Frontal Cortex (B. | TCGA-LGG | 28.08 | 7.97 |
| GTEX-1H23P-0011-R10a-SM-A96R | Brain - Frontal Cortex (B. | TCGA-LGG | 21.39 | 4.65 |
| GTEX-1H3O1-0011-R10a-SM-CJI3 | Brain - Frontal Cortex (B. | TCGA-LGG | 20.98 | 5.56 |
| GTEX-1H3VE-0011-R10b-SM-A9SK | Brain - Frontal Cortex (B. | TCGA-LGG | 19.57 | 7.76 |
| GTEX-1H4P4-0011-R10b-SM-CE6S | Brain - Frontal Cortex (B. | TCGA-LGG | 17.03 | 5.81 |
| GTEX-1HB9E-0011-R10a-SM-CJI3H | Brain - Frontal Cortex (B. | TCGA-LGG | 31.47 | 9.08 |
| GTEX-1HBPH-0011-R10a-SM-CE6S | Brain - Frontal Cortex (B. | TCGA-LGG | 16.63 | 6.04 |
| GTEX-1HBPI-0011-R10a-SM-A96R | Brain - Frontal Cortex (B. | TCGA-LGG | 28 | 7.54 |
| GTEX-1HBPM-0011-R10a-SM-CJI3 | Brain - Frontal Cortex (B. | TCGA-LGG | 7.86 | 2.5 |
| GTEX-1HCU6-0011-R10a-SM-CKZF | Brain - Frontal Cortex (B. | TCGA-LGG | 24.75 | 9.52 |
| GTEX-1HCVE-0011-R10a-SM-ARZL | Brain - Frontal Cortex (B. | TCGA-LGG | 27.78 | 7.2 |
| GTEX-1HFI6-0011-R10b-SM-ARZL | Brain - Frontal Cortex (B. | TCGA-LGG | 30.45 | 11.15 |
| GTEX-1HGF4-0011-R10b-SM-CE6S | Brain - Frontal Cortex (B. | TCGA-LGG | 23.17 | 8.73 |
| GTEX-1HR9M-0011-R10a-SM-CM2 | Brain - Frontal Cortex (B. | TCGA-LGG | 14.22 | 4.99 |
| GTEX-1HSKV-0011-R10b-SM-CMK | Brain - Frontal Cortex (B. | TCGA-LGG | 22.03 | 6.12 |
| GTEX-1HSMO-0011-R10a-SM-CL5 | Brain - Frontal Cortex (B. | TCGA-LGG | 19.02 | 6.34 |
| GTEX-1I1CD-0011-R10b-SM-CJI3W | Brain - Frontal Cortex (B. | TCGA-LGG | 15.23 | 4.47 |
| GTEX-1I1GV-0011-R10a-SM-CE6S | Brain - Frontal Cortex (B. | TCGA-LGG | 25.7 | 5.09 |
| GTEX-1I1HK-0011-R10b-SM-CJI3M | Brain - Frontal Cortex (B. | TCGA-LGG | 23.56 | 6.09 |
| GTEX-1ICLZ-0011-R10a-SM-A9SM | Brain - Frontal Cortex (B. | TCGA-LGG | 26.2 | 7.48 |
| GTEX-1IDJH-0011-R10a-SM-ARZL | Brain - Frontal Cortex (B. | TCGA-LGG | 27.76 | 6.59 |
| GTEX-1IKK5-0011-R10b-SM-CE6S | Brain - Frontal Cortex (B. | TCGA-LGG | 16.24 | 6.6 |

| | | | | |
|---|---|---|---|---|
| GTEX-1IL2V-0011-R10a-SM-CL55A | Brain - Frontal Cortex (B. | TCGA-LGG | 6.02 | 4.47 |
| GTEX-1IOXB-0011-R10a-SM-D3L8I | Brain - Frontal Cortex (B. | TCGA-LGG | 31.55 | 8.6 |
| GTEX-1J1OQ-0011-R10a-SM-A9G1 | Brain - Frontal Cortex (B. | TCGA-LGG | 26.92 | 7.25 |
| GTEX-1J8Q2-0011-R10b-SM-D4P2( | Brain - Frontal Cortex (B. | TCGA-LGG | 27.12 | 7.14 |
| GTEX-1LC47-0011-R10b-SM-EYYW | Brain - Frontal Cortex (B. | TCGA-LGG | 21.89 | 6.28 |
| GTEX-N7MS-0011-R10A-SM-2HMJI | Brain - Frontal Cortex (B. | TCGA-LGG | 23.58 | 7.96 |
| GTEX-N7MT-0011-R10A-SM-2I3E1 | Brain - Frontal Cortex (B. | TCGA-LGG | 27.41 | 8.07 |
| GTEX-NL3H-0011-R10A-SM-2I3E9 | Brain - Frontal Cortex (B. | TCGA-LGG | 31.11 | 7.56 |
| GTEX-NL4W-0011-R10A-SM-2I3DY | Brain - Frontal Cortex (B. | TCGA-LGG | 23.13 | 8.32 |
| GTEX-NPJ7-0011-R10A-SM-2I3E5 | Brain - Frontal Cortex (B. | TCGA-LGG | 32.09 | 7.42 |
| GTEX-NPJ8-0011-R10A-SM-2YUM( | Brain - Frontal Cortex (B. | TCGA-LGG | 39.79 | 11.36 |
| GTEX-OHPN-0011-R10A-SM-33HBI | Brain - Frontal Cortex (B. | TCGA-LGG | 13.34 | 4.67 |
| GTEX-OXRN-0011-R10A-SM-2I5GC | Brain - Frontal Cortex (B. | TCGA-LGG | 20.05 | 5.93 |
| GTEX-OXRO-0011-R10A-SM-2I5EH | Brain - Frontal Cortex (B. | TCGA-LGG | 15.98 | 5.69 |
| GTEX-P44G-0011-R10A-SM-2I3FF | Brain - Frontal Cortex (B. | TCGA-LGG | 24.96 | 8.55 |
| GTEX-P44H-0011-R10A-SM-2XCEK | Brain - Frontal Cortex (B. | TCGA-LGG | 32.58 | 7.32 |
| GTEX-PVOW-0011-R10A-SM-EXOH | Brain - Frontal Cortex (B. | TCGA-LGG | 37.65 | 11.46 |
| GTEX-Q2AG-0011-R10A-SM-2HML. | Brain - Frontal Cortex (B. | TCGA-LGG | 26.02 | 8.27 |
| GTEX-QDT8-0011-R10A-SM-32PKC | Brain - Frontal Cortex (B. | TCGA-LGG | 35.49 | 13.24 |
| GTEX-QMR6-0011-R10A-SM-32PK( | Brain - Frontal Cortex (B. | TCGA-LGG | 33.15 | 9.9 |
| GTEX-QVJO-0011-R10A-SM-2S1QJ | Brain - Frontal Cortex (B. | TCGA-LGG | 28.17 | 8.72 |
| GTEX-QVUS-0011-R10A-SM-3GIK3 | Brain - Frontal Cortex (B. | TCGA-LGG | 35.78 | 8.37 |
| GTEX-S7SE-0011-R10A-SM-2XCDF | Brain - Frontal Cortex (B. | TCGA-LGG | 23.29 | 7.13 |
| GTEX-T5JC-0011-R10A-SM-32PM2 | Brain - Frontal Cortex (B. | TCGA-LGG | 25.68 | 8.54 |
| GTEX-T6MN-0011-R10A-SM-32QP7 | Brain - Frontal Cortex (B. | TCGA-LGG | 36.52 | 9.42 |
| GTEX-TSE9-0011-R10A-SM-3DB7C | Brain - Frontal Cortex (B. | TCGA-LGG | 17.92 | 5.99 |
| GTEX-UTHO-0011-R10A-SM-3GIJC | Brain - Frontal Cortex (B. | TCGA-LGG | 34.34 | 15.1 |
| GTEX-WHSE-0011-R10A-SM-EYYV | Brain - Frontal Cortex (B. | TCGA-LGG | 19.04 | 8.39 |
| GTEX-WL46-0011-R10A-SM-3MJFC | Brain - Frontal Cortex (B. | TCGA-LGG | 33.85 | 7.55 |
| GTEX-WVLH-0011-R10A-SM-3MJFI | Brain - Frontal Cortex (B. | TCGA-LGG | 37.03 | 11.45 |
| GTEX-WWYW-0011-R10A-SM-3NB3 | Brain - Frontal Cortex (B. | TCGA-LGG | 41.59 | 11.41 |

| | | | | |
|---|---|---|---|---|
| GTEX-WZTO-0011-R10B-SM-4E3KI | Brain - Frontal Cortex (B. | TCGA-LGG | 26.55 | 8.45 |
| GTEX-X261-0011-R10B-SM-4E3JT | Brain - Frontal Cortex (B. | TCGA-LGG | 17.78 | 8.32 |
| GTEX-X4EP-0011-R10B-SM-4QASJ | Brain - Frontal Cortex (B. | TCGA-LGG | 48.96 | 18.12 |
| GTEX-X4XX-0011-R10B-SM-46MW( | Brain - Frontal Cortex (B. | TCGA-LGG | 37.88 | 9.58 |
| GTEX-X4XY-0011-R10B-SM-46MWS | Brain - Frontal Cortex (B. | TCGA-LGG | 4.24 | 3.4 |
| GTEX-X585-0011-R10A-SM-46MUY | Brain - Frontal Cortex (B. | TCGA-LGG | 34.85 | 12.18 |
| GTEX-XLM4-0011-R10A-SM-4AT5P | Brain - Frontal Cortex (B. | TCGA-LGG | 27.94 | 6.48 |
| GTEX-XMD1-0011-R10A-SM-4AT4A | Brain - Frontal Cortex (B. | TCGA-LGG | 18.36 | 6.78 |
| GTEX-Y8DK-0011-R10A-SM-4SOK1 | Brain - Frontal Cortex (B. | TCGA-LGG | 27.95 | 6.47 |
| GTEX-YFC4-0011-R10a-SM-4SOK5 | Brain - Frontal Cortex (B. | TCGA-LGG | 27.09 | 8.36 |
| GTEX-YJ89-0011-R10a-SM-4SOK9 | Brain - Frontal Cortex (B. | TCGA-LGG | 24.56 | 7.68 |
| GTEX-Z93S-0011-R10a-SM-4RGNM | Brain - Frontal Cortex (B. | TCGA-LGG | 23.76 | 6.45 |
| GTEX-ZAB4-0011-R10a-SM-4SOKH | Brain - Frontal Cortex (B. | TCGA-LGG | 22.26 | 7.07 |
| GTEX-ZDXO-0011-R10a-SM-4WWD | Brain - Frontal Cortex (B. | TCGA-LGG | 24.61 | 9.93 |
| GTEX-ZE7O-0011-R10a-SM-57WAZ | Brain - Frontal Cortex (B. | TCGA-LGG | 17.1 | 11.92 |
| GTEX-ZF28-0011-R10a-SM-4WWEI | Brain - Frontal Cortex (B. | TCGA-LGG | 27.45 | 7.15 |
| GTEX-ZUA1-0011-R10a-SM-51MT6 | Brain - Frontal Cortex (B. | TCGA-LGG | 21.51 | 5.17 |
| GTEX-ZV68-0011-R10a-SM-51MT7 | Brain - Frontal Cortex (B. | TCGA-LGG | 11.82 | 3.92 |
| GTEX-ZVT3-0011-R10b-SM-57WB6 | Brain - Frontal Cortex (B. | TCGA-LGG | 20.35 | 6.07 |
| GTEX-ZVZQ-0011-R10b-SM-51MRT | Brain - Frontal Cortex (B. | TCGA-LGG | 28.14 | 10.49 |
| GTEX-ZXG5-0011-R10a-SM-57WDI | Brain - Frontal Cortex (B. | TCGA-LGG | 11.7 | 4.97 |
| GTEX-1117F-2826-SM-5GZXL | Breast - Mammary Tissu | TCGA-BRCA | 43.37 | 14.14 |
| GTEX-111YS-1926-SM-5GICC | Breast - Mammary Tissu | TCGA-BRCA | 58.15 | 10.65 |
| GTEX-1122O-1226-SM-5H113 | Breast - Mammary Tissu | TCGA-BRCA | 42.58 | 9.31 |
| GTEX-117XS-1926-SM-5GICO | Breast - Mammary Tissu | TCGA-BRCA | 34.82 | 5.97 |
| GTEX-117YX-1426-SM-5H12H | Breast - Mammary Tissu | TCGA-BRCA | 35.87 | 9.49 |
| GTEX-1192X-2326-SM-5987X | Breast - Mammary Tissu | TCGA-BRCA | 54.03 | 14.16 |
| GTEX-11DXW-0626-SM-5N9ER | Breast - Mammary Tissu | TCGA-BRCA | 37.05 | 14.85 |
| GTEX-11DXY-2326-SM-5GICW | Breast - Mammary Tissu | TCGA-BRCA | 45.5 | 10.34 |
| GTEX-11DXZ-1926-SM-5GZZL | Breast - Mammary Tissu | TCGA-BRCA | 33.04 | 7.89 |
| GTEX-11DZ1-0326-SM-5N9BN | Breast - Mammary Tissu | TCGA-BRCA | 63.22 | 15.74 |

| | | | | |
|---|---|---|---|---|
| GTEX-11EI6-0626-SM-5985T | Breast - Mammary Tissu | TCGA-BRCA | 54.39 | 15.81 |
| GTEX-11EM3-1326-SM-5N9C6 | Breast - Mammary Tissu | TCGA-BRCA | 44.87 | 11.23 |
| GTEX-11EMC-2026-SM-5A5JV | Breast - Mammary Tissu | TCGA-BRCA | 51.04 | 20.22 |
| GTEX-11EQ9-1826-SM-5Q5AJ | Breast - Mammary Tissu | TCGA-BRCA | 47.2 | 11.25 |
| GTEX-11GS4-2126-SM-5A5KR | Breast - Mammary Tissu | TCGA-BRCA | 54.76 | 14.54 |
| GTEX-11GSO-1926-SM-5A5K3 | Breast - Mammary Tissu | TCGA-BRCA | 32.49 | 6.33 |
| GTEX-11GSP-0926-SM-9WYSG | Breast - Mammary Tissu | TCGA-BRCA | 42.07 | 15.05 |
| GTEX-11I78-2226-SM-5PNYA | Breast - Mammary Tissu | TCGA-BRCA | 34.4 | 6.54 |
| GTEX-11LCK-2426-SM-5HL5F | Breast - Mammary Tissu | TCGA-BRCA | 35.85 | 10.37 |
| GTEX-11NSD-0926-SM-5N9DR | Breast - Mammary Tissu | TCGA-BRCA | 43.87 | 13.46 |
| GTEX-11NV4-2026-SM-5N9DG | Breast - Mammary Tissu | TCGA-BRCA | 41.9 | 12.15 |
| GTEX-11O72-2126-SM-5N9FO | Breast - Mammary Tissu | TCGA-BRCA | 39.35 | 13.98 |
| GTEX-11OF3-1926-SM-59889 | Breast - Mammary Tissu | TCGA-BRCA | 45.13 | 11.16 |
| GTEX-11ONC-2126-SM-5HL6E | Breast - Mammary Tissu | TCGA-BRCA | 36.51 | 9.82 |
| GTEX-11P7K-0726-SM-5EGKX | Breast - Mammary Tissu | TCGA-BRCA | 52.84 | 10.58 |
| GTEX-11P81-1926-SM-5BC53 | Breast - Mammary Tissu | TCGA-BRCA | 38.48 | 14.2 |
| GTEX-11P82-1326-SM-5HL62 | Breast - Mammary Tissu | TCGA-BRCA | 48.21 | 10.78 |
| GTEX-11PRG-0826-SM-5EQ6A | Breast - Mammary Tissu | TCGA-BRCA | 54.42 | 16.17 |
| GTEX-11TT1-2126-SM-5GU5Y | Breast - Mammary Tissu | TCGA-BRCA | 62.61 | 19.57 |
| GTEX-11TUW-1826-SM-5BC5D | Breast - Mammary Tissu | TCGA-BRCA | 63.13 | 10.47 |
| GTEX-11WQC-1726-SM-5GU4W | Breast - Mammary Tissu | TCGA-BRCA | 58.14 | 16.05 |
| GTEX-11WQK-2426-SM-5GU5C | Breast - Mammary Tissu | TCGA-BRCA | 71.58 | 13.02 |
| GTEX-11ZTT-2326-SM-5EQLG | Breast - Mammary Tissu | TCGA-BRCA | 27.56 | 12.92 |
| GTEX-11ZUS-0826-SM-5FQUY | Breast - Mammary Tissu | TCGA-BRCA | 31.9 | 10.31 |
| GTEX-1211K-1926-SM-5EQLB | Breast - Mammary Tissu | TCGA-BRCA | 45.37 | 12.78 |
| GTEX-1269C-2426-SM-5FQSN | Breast - Mammary Tissu | TCGA-BRCA | 44.13 | 14.45 |
| GTEX-12BJ1-1826-SM-5HL9N | Breast - Mammary Tissu | TCGA-BRCA | 50.74 | 6.37 |
| GTEX-12KS4-0126-SM-5Q5A5 | Breast - Mammary Tissu | TCGA-BRCA | 40.08 | 8.68 |
| GTEX-12WSK-2226-SM-5GCO5 | Breast - Mammary Tissu | TCGA-BRCA | 56.34 | 9.03 |
| GTEX-12WSM-1726-SM-5BC6J | Breast - Mammary Tissu | TCGA-BRCA | 48.02 | 9.04 |
| GTEX-12WSN-1326-SM-5GCNT | Breast - Mammary Tissu | TCGA-BRCA | 37.9 | 12.13 |

| | | | | |
|---|---|---|---|---|
| GTEX-12ZZX-1126-SM-5EGKB | Breast - Mammary Tissu | TCGA-BRCA | 52.41 | 14.91 |
| GTEX-13113-5013-SM-7SB7G | Breast - Mammary Tissu | TCGA-BRCA | 44.92 | 9.89 |
| GTEX-1313W-0826-SM-5EQ4T | Breast - Mammary Tissu | TCGA-BRCA | 56.65 | 14.72 |
| GTEX-1314G-1226-SM-5BC6D | Breast - Mammary Tissu | TCGA-BRCA | 53.11 | 16.02 |
| GTEX-131XW-0726-SM-5EGK3 | Breast - Mammary Tissu | TCGA-BRCA | 38.32 | 18.93 |
| GTEX-131YS-0626-SM-5EGKL | Breast - Mammary Tissu | TCGA-BRCA | 41.67 | 14.31 |
| GTEX-132AR-0826-SM-5EGK6 | Breast - Mammary Tissu | TCGA-BRCA | 44.69 | 13.35 |
| GTEX-132NY-0826-SM-5K7Y7 | Breast - Mammary Tissu | TCGA-BRCA | 39.83 | 14.01 |
| GTEX-132Q8-1426-SM-5EGK7 | Breast - Mammary Tissu | TCGA-BRCA | 29.39 | 8.87 |
| GTEX-133LE-1726-SM-5K7VQ | Breast - Mammary Tissu | TCGA-BRCA | 48.14 | 13.1 |
| GTEX-1399U-1826-SM-5PNZ1 | Breast - Mammary Tissu | TCGA-BRCA | 50.61 | 11.25 |
| GTEX-139T6-1626-SM-5PNYZ | Breast - Mammary Tissu | TCGA-BRCA | 36.02 | 9.35 |
| GTEX-139T8-0826-SM-5L3DE | Breast - Mammary Tissu | TCGA-BRCA | 34.92 | 11.48 |
| GTEX-139TU-0626-SM-5KM3X | Breast - Mammary Tissu | TCGA-BRCA | 52.62 | 11.63 |
| GTEX-13CF2-2026-SM-5K7VI | Breast - Mammary Tissu | TCGA-BRCA | 46.06 | 13.5 |
| GTEX-13CF3-2126-SM-5IFJP | Breast - Mammary Tissu | TCGA-BRCA | 42.39 | 11.89 |
| GTEX-13D11-1026-SM-5IJFB | Breast - Mammary Tissu | TCGA-BRCA | 48.34 | 9.56 |
| GTEX-13FHO-0826-SM-5L3E8 | Breast - Mammary Tissu | TCGA-BRCA | 44.82 | 13.05 |
| GTEX-13FTW-1426-SM-5LZWZ | Breast - Mammary Tissu | TCGA-BRCA | 32.22 | 9.97 |
| GTEX-13FTX-1126-SM-5N9EN | Breast - Mammary Tissu | TCGA-BRCA | 39.21 | 9.26 |
| GTEX-13FTY-2226-SM-5J1ND | Breast - Mammary Tissu | TCGA-BRCA | 50.39 | 22.06 |
| GTEX-13N11-1726-SM-5J1OJ | Breast - Mammary Tissu | TCGA-BRCA | 49.06 | 14.48 |
| GTEX-13N1W-0626-SM-5MR4U | Breast - Mammary Tissu | TCGA-BRCA | 57.61 | 7.57 |
| GTEX-13NZ8-0126-SM-5IJCT | Breast - Mammary Tissu | TCGA-BRCA | 58.23 | 19.63 |
| GTEX-13NZ9-1026-SM-5MR5K | Breast - Mammary Tissu | TCGA-BRCA | 59.18 | 17.05 |
| GTEX-13NZB-2126-SM-5MR4Y | Breast - Mammary Tissu | TCGA-BRCA | 61.15 | 14.93 |
| GTEX-13O3O-0826-SM-5K7WE | Breast - Mammary Tissu | TCGA-BRCA | 66.92 | 14.68 |
| GTEX-13O3P-0826-SM-5L3DH | Breast - Mammary Tissu | TCGA-BRCA | 44.96 | 13.97 |
| GTEX-13O3Q-2226-SM-5KM4O | Breast - Mammary Tissu | TCGA-BRCA | 50.71 | 15.95 |
| GTEX-13O61-1826-SM-5KM4I | Breast - Mammary Tissu | TCGA-BRCA | 53.39 | 10.64 |
| GTEX-13OW5-2226-SM-5L3HC | Breast - Mammary Tissu | TCGA-BRCA | 51.9 | 17.79 |

| | | | | |
|---|---|---|---|---|
| GTEX-13OW8-0226-SM-5K7UP | Breast - Mammary Tissu | TCGA-BRCA | 46.04 | 10.08 |
| GTEX-13PL6-2926-SM-5L3I2 | Breast - Mammary Tissu | TCGA-BRCA | 47.65 | 28.19 |
| GTEX-13PVQ-1026-SM-5KM3M | Breast - Mammary Tissu | TCGA-BRCA | 62.08 | 7.33 |
| GTEX-13PVR-2226-SM-7DHKP | Breast - Mammary Tissu | TCGA-BRCA | 36.74 | 8.87 |
| GTEX-13QIC-2326-SM-5LU5N | Breast - Mammary Tissu | TCGA-BRCA | 43.97 | 16.12 |
| GTEX-13QJ3-0826-SM-7DHKK | Breast - Mammary Tissu | TCGA-BRCA | 37.95 | 11.46 |
| GTEX-13S86-1226-SM-5S2OA | Breast - Mammary Tissu | TCGA-BRCA | 61.58 | 17.29 |
| GTEX-13SLW-2526-SM-62LDQ | Breast - Mammary Tissu | TCGA-BRCA | 57.39 | 13.75 |
| GTEX-13SLX-2326-SM-5ZZWE | Breast - Mammary Tissu | TCGA-BRCA | 54.53 | 17.55 |
| GTEX-13VXU-2826-SM-664MA | Breast - Mammary Tissu | TCGA-BRCA | 61.79 | 15.27 |
| GTEX-13W3W-1226-SM-5LU4H | Breast - Mammary Tissu | TCGA-BRCA | 43.89 | 13.9 |
| GTEX-13W46-0826-SM-5LU3H | Breast - Mammary Tissu | TCGA-BRCA | 51.59 | 9.39 |
| GTEX-144GL-2026-SM-5LU3O | Breast - Mammary Tissu | TCGA-BRCA | 40.12 | 9.99 |
| GTEX-144GM-0926-SM-5O994 | Breast - Mammary Tissu | TCGA-BRCA | 42.86 | 7.33 |
| GTEX-145LT-0726-SM-5S2VM | Breast - Mammary Tissu | TCGA-BRCA | 55.61 | 13.81 |
| GTEX-145ME-1526-SM-5Q5F2 | Breast - Mammary Tissu | TCGA-BRCA | 30.89 | 6.72 |
| GTEX-145MF-2226-SM-7EPIR | Breast - Mammary Tissu | TCGA-BRCA | 36.57 | 9.93 |
| GTEX-145MN-1926-SM-5SIAI | Breast - Mammary Tissu | TCGA-BRCA | 38.32 | 8.92 |
| GTEX-145MO-0826-SM-5NQBL | Breast - Mammary Tissu | TCGA-BRCA | 39.1 | 9.33 |
| GTEX-146FH-0826-SM-5SI8T | Breast - Mammary Tissu | TCGA-BRCA | 52.34 | 16.85 |
| GTEX-14753-2426-SM-5LU8U | Breast - Mammary Tissu | TCGA-BRCA | 50.96 | 9.28 |
| GTEX-147F4-2826-SM-5NQBN | Breast - Mammary Tissu | TCGA-BRCA | 46.3 | 14.29 |
| GTEX-14A5I-0726-SM-5TDEB | Breast - Mammary Tissu | TCGA-BRCA | 54.48 | 19.32 |
| GTEX-14ABY-2126-SM-5QGQZ | Breast - Mammary Tissu | TCGA-BRCA | 64.05 | 14.89 |
| GTEX-14AS3-1626-SM-5S2OY | Breast - Mammary Tissu | TCGA-BRCA | 60.49 | 12.64 |
| GTEX-14B4R-1226-SM-5TDDT | Breast - Mammary Tissu | TCGA-BRCA | 31.57 | 8.95 |
| GTEX-14BMU-1626-SM-5TDE7 | Breast - Mammary Tissu | TCGA-BRCA | 51.45 | 14.2 |
| GTEX-14BMV-0626-SM-793AU | Breast - Mammary Tissu | TCGA-BRCA | 38.17 | 10.07 |
| GTEX-14DAR-1326-SM-7DUEG | Breast - Mammary Tissu | TCGA-BRCA | 30.18 | 7.48 |
| GTEX-14E6C-1326-SM-62LEQ | Breast - Mammary Tissu | TCGA-BRCA | 71 | 7.43 |
| GTEX-14E6E-1326-SM-5S2NR | Breast - Mammary Tissu | TCGA-BRCA | 38.47 | 13.81 |

| | | | | |
|---|---|---|---|---|
| GTEX-14E7W-0826-SM-62LEJ | Breast - Mammary Tissu | TCGA-BRCA | 41.76 | 11.22 |
| GTEX-14H4A-2526-SM-5YYAY | Breast - Mammary Tissu | TCGA-BRCA | 42.56 | 8.67 |
| GTEX-14ICK-2426-SM-6EU27 | Breast - Mammary Tissu | TCGA-BRCA | 64.14 | 12.88 |
| GTEX-14LLW-0626-SM-62LFC | Breast - Mammary Tissu | TCGA-BRCA | 48.17 | 13.33 |
| GTEX-14PHY-1926-SM-5YY95 | Breast - Mammary Tissu | TCGA-BRCA | 47.86 | 12.3 |
| GTEX-14PJ4-2126-SM-6ETZJ | Breast - Mammary Tissu | TCGA-BRCA | 44.96 | 13.1 |
| GTEX-14PJO-0726-SM-69LO8 | Breast - Mammary Tissu | TCGA-BRCA | 48.17 | 13.84 |
| GTEX-14PKU-0426-SM-6EU1P | Breast - Mammary Tissu | TCGA-BRCA | 57.05 | 13.19 |
| GTEX-14PN4-0626-SM-62LFP | Breast - Mammary Tissu | TCGA-BRCA | 64.4 | 9.22 |
| GTEX-15EOM-5019-SM-793DK | Breast - Mammary Tissu | TCGA-BRCA | 48.72 | 20.41 |
| GTEX-15ER7-1626-SM-6PAMZ | Breast - Mammary Tissu | TCGA-BRCA | 55.28 | 13.75 |
| GTEX-15ETS-0626-SM-7KUMX | Breast - Mammary Tissu | TCGA-BRCA | 43.81 | 11.84 |
| GTEX-15FZZ-0726-SM-7KUFZ | Breast - Mammary Tissu | TCGA-BRCA | 35.24 | 13.12 |
| GTEX-15G19-2126-SM-6M48J | Breast - Mammary Tissu | TCGA-BRCA | 82.22 | 30.33 |
| GTEX-15RJE-2626-SM-7KFT1 | Breast - Mammary Tissu | TCGA-BRCA | 35.76 | 6.76 |
| GTEX-15SHW-1326-SM-6PAL8 | Breast - Mammary Tissu | TCGA-BRCA | 43.16 | 12.68 |
| GTEX-15UF6-0126-SM-6PAMB | Breast - Mammary Tissu | TCGA-BRCA | 57.34 | 14.3 |
| GTEX-15UF7-0726-SM-6M46D | Breast - Mammary Tissu | TCGA-BRCA | 40.13 | 14.2 |
| GTEX-169BO-1126-SM-7MGWN | Breast - Mammary Tissu | TCGA-BRCA | 35.63 | 6.6 |
| GTEX-16BQI-1026-SM-7KUEA | Breast - Mammary Tissu | TCGA-BRCA | 41.2 | 10.39 |
| GTEX-16NGA-0826-SM-718AF | Breast - Mammary Tissu | TCGA-BRCA | 48.58 | 11.19 |
| GTEX-16YQH-2826-SM-6PAMY | Breast - Mammary Tissu | TCGA-BRCA | 40.43 | 11.55 |
| GTEX-17EUY-1926-SM-7DUF6 | Breast - Mammary Tissu | TCGA-BRCA | 52.84 | 9.64 |
| GTEX-17EVP-0226-SM-79OND | Breast - Mammary Tissu | TCGA-BRCA | 49.41 | 13.02 |
| GTEX-17EVQ-0426-SM-7LG57 | Breast - Mammary Tissu | TCGA-BRCA | 50.76 | 10.09 |
| GTEX-17F96-2426-SM-7IGLN | Breast - Mammary Tissu | TCGA-BRCA | 36.67 | 8.92 |
| GTEX-17F97-2526-SM-7EWDV | Breast - Mammary Tissu | TCGA-BRCA | 50.91 | 10.09 |
| GTEX-17F98-0526-SM-79OK5 | Breast - Mammary Tissu | TCGA-BRCA | 38.35 | 17.07 |
| GTEX-17F9E-2526-SM-7MGW2 | Breast - Mammary Tissu | TCGA-BRCA | 24.03 | 4.78 |
| GTEX-17GQL-0326-SM-7LG5U | Breast - Mammary Tissu | TCGA-BRCA | 39.19 | 9.14 |
| GTEX-17HG3-0126-SM-7IGNH | Breast - Mammary Tissu | TCGA-BRCA | 54.65 | 18.37 |

| | | | | |
|---|---|---|---|---|
| GTEX-17HGU-1326-SM-79OKB | Breast - Mammary Tissu | TCGA-BRCA | 42.84 | 11.15 |
| GTEX-17HHE-1426-SM-7EPH4 | Breast - Mammary Tissu | TCGA-BRCA | 38.85 | 10.46 |
| GTEX-17HHY-0926-SM-793C1 | Breast - Mammary Tissu | TCGA-BRCA | 49.27 | 19.31 |
| GTEX-17JCI-0726-SM-7EPH1 | Breast - Mammary Tissu | TCGA-BRCA | 47.57 | 15.26 |
| GTEX-17KNJ-2026-SM-7LG53 | Breast - Mammary Tissu | TCGA-BRCA | 44.12 | 10.33 |
| GTEX-17MF6-0326-SM-7EPH5 | Breast - Mammary Tissu | TCGA-BRCA | 43.06 | 6.86 |
| GTEX-17MFQ-0926-SM-7LG4S | Breast - Mammary Tissu | TCGA-BRCA | 42.16 | 8.92 |
| GTEX-183FY-1126-SM-7DHLJ | Breast - Mammary Tissu | TCGA-BRCA | 37.3 | 11.55 |
| GTEX-183WM-0726-SM-7LTAA | Breast - Mammary Tissu | TCGA-BRCA | 38.64 | 7.39 |
| GTEX-18465-2026-SM-718AP | Breast - Mammary Tissu | TCGA-BRCA | 36.46 | 12.1 |
| GTEX-18A66-1726-SM-7LG5N | Breast - Mammary Tissu | TCGA-BRCA | 55.66 | 14.47 |
| GTEX-18A6Q-0926-SM-7LG4N | Breast - Mammary Tissu | TCGA-BRCA | 36.29 | 11.67 |
| GTEX-18A7A-0726-SM-7LTAI | Breast - Mammary Tissu | TCGA-BRCA | 47.46 | 17.53 |
| GTEX-18A7B-2626-SM-7LG55 | Breast - Mammary Tissu | TCGA-BRCA | 41.48 | 9.1 |
| GTEX-18D9A-1526-SM-7LG4J | Breast - Mammary Tissu | TCGA-BRCA | 31.04 | 8.13 |
| GTEX-18QFQ-0826-SM-718AX | Breast - Mammary Tissu | TCGA-BRCA | 47.66 | 12.68 |
| GTEX-1A3MV-1626-SM-731C1 | Breast - Mammary Tissu | TCGA-BRCA | 38.4 | 13.86 |
| GTEX-1A3MX-2726-SM-718B6 | Breast - Mammary Tissu | TCGA-BRCA | 48.03 | 11.7 |
| GTEX-1A8G7-2426-SM-731AK | Breast - Mammary Tissu | TCGA-BRCA | 50.42 | 14.39 |
| GTEX-1AMEY-1026-SM-718AA | Breast - Mammary Tissu | TCGA-BRCA | 48.05 | 8.43 |
| GTEX-1AX8Z-0926-SM-731AW | Breast - Mammary Tissu | TCGA-BRCA | 52.41 | 18.11 |
| GTEX-1AX9I-0726-SM-73KWV | Breast - Mammary Tissu | TCGA-BRCA | 56.01 | 12.72 |
| GTEX-1AX9J-1126-SM-731B7 | Breast - Mammary Tissu | TCGA-BRCA | 43.95 | 8.46 |
| GTEX-1B8KE-1226-SM-73KWK | Breast - Mammary Tissu | TCGA-BRCA | 52.9 | 8.04 |
| GTEX-1B8KZ-1526-SM-7DUG7 | Breast - Mammary Tissu | TCGA-BRCA | 58.99 | 19.55 |
| GTEX-1B8L1-2426-SM-9WYTE | Breast - Mammary Tissu | TCGA-BRCA | 40.56 | 13.55 |
| GTEX-1B8SG-1226-SM-7IGPF | Breast - Mammary Tissu | TCGA-BRCA | 53.17 | 10.79 |
| GTEX-1B932-0826-SM-73KXG | Breast - Mammary Tissu | TCGA-BRCA | 50.06 | 23.01 |
| GTEX-1B933-2526-SM-7IGO5 | Breast - Mammary Tissu | TCGA-BRCA | 43.63 | 12.13 |
| GTEX-1B97I-0426-SM-79OL7 | Breast - Mammary Tissu | TCGA-BRCA | 56.77 | 14.32 |
| GTEX-1B97J-0426-SM-79OLQ | Breast - Mammary Tissu | TCGA-BRCA | 32.22 | 14.74 |

| | | | | |
|---|---|---|---|---|
| GTEX-1BAJH-0826-SM-7EWEF | Breast - Mammary Tissu | TCGA-BRCA | 46.1 | 15.87 |
| GTEX-1C475-1226-SM-7SB72 | Breast - Mammary Tissu | TCGA-BRCA | 46.87 | 11.42 |
| GTEX-1C64O-0726-SM-7DUFU | Breast - Mammary Tissu | TCGA-BRCA | 48.26 | 17.31 |
| GTEX-1C6VQ-0426-SM-79OOX | Breast - Mammary Tissu | TCGA-BRCA | 47.6 | 15.69 |
| GTEX-1C6VS-0726-SM-7EPHF | Breast - Mammary Tissu | TCGA-BRCA | 63.53 | 13.55 |
| GTEX-1C6WA-2426-SM-7SB86 | Breast - Mammary Tissu | TCGA-BRCA | 35.95 | 7.64 |
| GTEX-1CAMR-1426-SM-793BO | Breast - Mammary Tissu | TCGA-BRCA | 30.94 | 8.24 |
| GTEX-1CAMS-1426-SM-7IGPM | Breast - Mammary Tissu | TCGA-BRCA | 48.92 | 13.83 |
| GTEX-1CB4E-0226-SM-79OLW | Breast - Mammary Tissu | TCGA-BRCA | 58.41 | 14.57 |
| GTEX-1CB4G-2326-SM-79OOI | Breast - Mammary Tissu | TCGA-BRCA | 42.91 | 12.54 |
| GTEX-1CB4J-1826-SM-7EWF9 | Breast - Mammary Tissu | TCGA-BRCA | 60.41 | 9.32 |
| GTEX-1E2YA-2726-SM-7IGPW | Breast - Mammary Tissu | TCGA-BRCA | 46.32 | 8.36 |
| GTEX-1EKGG-2626-SM-7IGPY | Breast - Mammary Tissu | TCGA-BRCA | 38.31 | 12.58 |
| GTEX-1EN7A-1426-SM-7PBY4 | Breast - Mammary Tissu | TCGA-BRCA | 47.11 | 10.41 |
| GTEX-1EU9M-2826-SM-7EWFH | Breast - Mammary Tissu | TCGA-BRCA | 31.77 | 12.88 |
| GTEX-1EWIQ-1726-SM-7SB7T | Breast - Mammary Tissu | TCGA-BRCA | 37.9 | 14.88 |
| GTEX-1F5PK-2126-SM-7PC28 | Breast - Mammary Tissu | TCGA-BRCA | 30.89 | 8.14 |
| GTEX-1F5PL-1026-SM-7RHI1 | Breast - Mammary Tissu | TCGA-BRCA | 53.07 | 13.03 |
| GTEX-1F6I4-2526-SM-7SB8F | Breast - Mammary Tissu | TCGA-BRCA | 48.2 | 19.95 |
| GTEX-1F75A-1826-SM-7SB8G | Breast - Mammary Tissu | TCGA-BRCA | 40.03 | 13.73 |
| GTEX-1F75B-0626-SM-7SB8A | Breast - Mammary Tissu | TCGA-BRCA | 40.92 | 8.21 |
| GTEX-1F75I-2026-SM-7SB7N | Breast - Mammary Tissu | TCGA-BRCA | 47.13 | 15.04 |
| GTEX-1GF9U-1726-SM-7MGY5 | Breast - Mammary Tissu | TCGA-BRCA | 45.5 | 14.31 |
| GTEX-1GF9V-2526-SM-7SB8K | Breast - Mammary Tissu | TCGA-BRCA | 48.88 | 17.72 |
| GTEX-1GF9X-1026-SM-ACKX2 | Breast - Mammary Tissu | TCGA-BRCA | 54.25 | 15.02 |
| GTEX-1GMR2-1926-SM-9MQLZ | Breast - Mammary Tissu | TCGA-BRCA | 30.49 | 7.82 |
| GTEX-1GMR3-2626-SM-7RHHG | Breast - Mammary Tissu | TCGA-BRCA | 48.84 | 17.95 |
| GTEX-1GMR8-0726-SM-9KNVI | Breast - Mammary Tissu | TCGA-BRCA | 32.04 | 14.99 |
| GTEX-1GMRU-0826-SM-9WYT7 | Breast - Mammary Tissu | TCGA-BRCA | 46.77 | 14.38 |
| GTEX-1GN1U-0526-SM-9KNTT | Breast - Mammary Tissu | TCGA-BRCA | 28.92 | 17.48 |
| GTEX-1GN1W-1926-SM-7P8TH | Breast - Mammary Tissu | TCGA-BRCA | 48.58 | 14.15 |

| | | | | |
|---|---|---|---|---|
| GTEX-1GN2E-2226-SM-9WG6J | Breast - Mammary Tissu | TCGA-BRCA | 59.43 | 11.22 |
| GTEX-1GPI6-0626-SM-7RHHJ | Breast - Mammary Tissu | TCGA-BRCA | 50.41 | 17.23 |
| GTEX-1GPI7-0926-SM-7PC3J | Breast - Mammary Tissu | TCGA-BRCA | 35.48 | 11.13 |
| GTEX-1GZ2Q-2626-SM-7MXV9 | Breast - Mammary Tissu | TCGA-BRCA | 50.04 | 16.73 |
| GTEX-1GZ4H-0326-SM-7P8QX | Breast - Mammary Tissu | TCGA-BRCA | 45.67 | 9.3 |
| GTEX-1GZ4I-2826-SM-9MQK5 | Breast - Mammary Tissu | TCGA-BRCA | 39.95 | 12.69 |
| GTEX-1GZHY-0726-SM-9WG6B | Breast - Mammary Tissu | TCGA-BRCA | 36.93 | 13.15 |
| GTEX-1H11D-1526-SM-9MQJY | Breast - Mammary Tissu | TCGA-BRCA | 42.87 | 14.34 |
| GTEX-1H1CY-2226-SM-9WG7E | Breast - Mammary Tissu | TCGA-BRCA | 37.99 | 16.98 |
| GTEX-1H1DG-0526-SM-9YFLN | Breast - Mammary Tissu | TCGA-BRCA | 69.34 | 19.4 |
| GTEX-1H1E6-0926-SM-9YFLT | Breast - Mammary Tissu | TCGA-BRCA | 59.97 | 16.5 |
| GTEX-1H1ZS-0126-SM-9YFKQ | Breast - Mammary Tissu | TCGA-BRCA | 37.12 | 9.66 |
| GTEX-1H23P-0726-SM-9WG7K | Breast - Mammary Tissu | TCGA-BRCA | 46.31 | 9.26 |
| GTEX-1H3VY-0926-SM-ACKXE | Breast - Mammary Tissu | TCGA-BRCA | 35.67 | 10.39 |
| GTEX-1H4P4-2226-SM-9MQKQ | Breast - Mammary Tissu | TCGA-BRCA | 39.78 | 12.88 |
| GTEX-1HBPM-0126-SM-ARL9E | Breast - Mammary Tissu | TCGA-BRCA | 36.15 | 11.93 |
| GTEX-1HBPN-2326-SM-9WPPU | Breast - Mammary Tissu | TCGA-BRCA | 26.55 | 6.47 |
| GTEX-1HC8U-0826-SM-ADEIL | Breast - Mammary Tissu | TCGA-BRCA | 30.45 | 9.87 |
| GTEX-1HCU7-1026-SM-ADEIP | Breast - Mammary Tissu | TCGA-BRCA | 55.87 | 13.83 |
| GTEX-1HCU8-0126-SM-ADEHH | Breast - Mammary Tissu | TCGA-BRCA | 43.33 | 9.45 |
| GTEX-1HCU9-0326-SM-9WYSC | Breast - Mammary Tissu | TCGA-BRCA | 40.3 | 9.78 |
| GTEX-1HCUA-0726-SM-ACKVU | Breast - Mammary Tissu | TCGA-BRCA | 44.85 | 10.34 |
| GTEX-1HCVE-2626-SM-ADEJ8 | Breast - Mammary Tissu | TCGA-BRCA | 42.9 | 16.2 |
| GTEX-1HFI7-0126-SM-CGQFQ | Breast - Mammary Tissu | TCGA-BRCA | 29.36 | 9.43 |
| GTEX-1HKZK-1826-SM-ADEJ2 | Breast - Mammary Tissu | TCGA-BRCA | 44.24 | 14.29 |
| GTEX-1HR98-0426-SM-COH36 | Breast - Mammary Tissu | TCGA-BRCA | 44.19 | 9.62 |
| GTEX-1HR9M-2426-SM-ACKXI | Breast - Mammary Tissu | TCGA-BRCA | 48.54 | 15.36 |
| GTEX-1HSEH-2026-SM-ADEHN | Breast - Mammary Tissu | TCGA-BRCA | 34.26 | 11.52 |
| GTEX-1HSGN-1526-SM-A9SMO | Breast - Mammary Tissu | TCGA-BRCA | 48.67 | 17.5 |
| GTEX-1HSKV-2026-SM-CMKGX | Breast - Mammary Tissu | TCGA-BRCA | 49.14 | 18.87 |
| GTEX-1HSMO-0826-SM-ADEIV | Breast - Mammary Tissu | TCGA-BRCA | 46.44 | 11.1 |

| | | | | |
|---|---|---|---|---|
| GTEX-1HT8W-2226-SM-CGQEC | Breast - Mammary Tissu | TCGA-BRCA | 40.36 | 11.89 |
| GTEX-1HUB1-1626-SM-CM2SL | Breast - Mammary Tissu | TCGA-BRCA | 51.47 | 11.64 |
| GTEX-1I19N-2826-SM-CL55D | Breast - Mammary Tissu | TCGA-BRCA | 33 | 11.2 |
| GTEX-1I1GP-0426-SM-COH3N | Breast - Mammary Tissu | TCGA-BRCA | 41.51 | 9.49 |
| GTEX-1I1GT-1126-SM-ARL7J | Breast - Mammary Tissu | TCGA-BRCA | 39.82 | 9.64 |
| GTEX-1I1GU-1026-SM-ADEHJ | Breast - Mammary Tissu | TCGA-BRCA | 52.53 | 15.28 |
| GTEX-1I1GV-2426-SM-CGQFM | Breast - Mammary Tissu | TCGA-BRCA | 31.43 | 8.72 |
| GTEX-1I1HK-2426-SM-ADEHX | Breast - Mammary Tissu | TCGA-BRCA | 46.82 | 14.42 |
| GTEX-1I4MK-1626-SM-CM2TJ | Breast - Mammary Tissu | TCGA-BRCA | 39.68 | 11.94 |
| GTEX-1I6K6-0826-SM-ARL7Z | Breast - Mammary Tissu | TCGA-BRCA | 45.27 | 10.04 |
| GTEX-1I6K7-1026-SM-A9SLK | Breast - Mammary Tissu | TCGA-BRCA | 51.46 | 12.29 |
| GTEX-1ICG6-1326-SM-ACKWR | Breast - Mammary Tissu | TCGA-BRCA | 30.91 | 15.55 |
| GTEX-1ICLY-1726-SM-B2LVT | Breast - Mammary Tissu | TCGA-BRCA | 52.3 | 12.22 |
| GTEX-1ICLZ-0426-SM-B2LW2 | Breast - Mammary Tissu | TCGA-BRCA | 45.55 | 9.53 |
| GTEX-1IDFM-0426-SM-B2LVV | Breast - Mammary Tissu | TCGA-BRCA | 48.81 | 10.88 |
| GTEX-1IDJE-0826-SM-CKZNZ | Breast - Mammary Tissu | TCGA-BRCA | 46.92 | 14.06 |
| GTEX-1IDJF-2326-SM-ARL86 | Breast - Mammary Tissu | TCGA-BRCA | 31.65 | 11.57 |
| GTEX-1IDJH-2226-SM-E6CJV | Breast - Mammary Tissu | TCGA-BRCA | 35.41 | 6.87 |
| GTEX-1IDJU-1826-SM-CMKHV | Breast - Mammary Tissu | TCGA-BRCA | 26.97 | 7.44 |
| GTEX-1IE54-0626-SM-ARL8D | Breast - Mammary Tissu | TCGA-BRCA | 47.96 | 11.26 |
| GTEX-1IGQW-0926-SM-ADEH6 | Breast - Mammary Tissu | TCGA-BRCA | 53.51 | 16.21 |
| GTEX-1IKJJ-2326-SM-COH3U | Breast - Mammary Tissu | TCGA-BRCA | 50.44 | 16.16 |
| GTEX-1IKOE-1326-SM-CE6T6 | Breast - Mammary Tissu | TCGA-BRCA | 42.54 | 12.11 |
| GTEX-1IKOH-2326-SM-CM2TW | Breast - Mammary Tissu | TCGA-BRCA | 41.79 | 15.5 |
| GTEX-1IOXB-1926-SM-ARL8E | Breast - Mammary Tissu | TCGA-BRCA | 38.58 | 12.08 |
| GTEX-1IY9M-2726-SM-CMKG5 | Breast - Mammary Tissu | TCGA-BRCA | 45.57 | 17.61 |
| GTEX-1J8EW-2626-SM-CL53E | Breast - Mammary Tissu | TCGA-BRCA | 42.15 | 15.38 |
| GTEX-1J8Q2-2726-SM-A9G2X | Breast - Mammary Tissu | TCGA-BRCA | 36.85 | 12.05 |
| GTEX-1J8Q3-1326-SM-CMKI1 | Breast - Mammary Tissu | TCGA-BRCA | 42.12 | 10.96 |
| GTEX-1J8QM-2026-SM-ARZMW | Breast - Mammary Tissu | TCGA-BRCA | 47.24 | 12.93 |
| GTEX-1JJE9-2326-SM-ARL8Q | Breast - Mammary Tissu | TCGA-BRCA | 48.39 | 12.99 |

| | | | | |
|---|---|---|---|---|
| GTEX-1JJEA-2726-SM-ARL8V | Breast - Mammary Tissu | TCGA-BRCA | 59.08 | 14.06 |
| GTEX-1JK1U-1826-SM-ARL8W | Breast - Mammary Tissu | TCGA-BRCA | 48.47 | 14.08 |
| GTEX-1JKYN-0726-SM-ARZMY | Breast - Mammary Tissu | TCGA-BRCA | 29.35 | 7.04 |
| GTEX-1JKYR-0926-SM-CMKGJ | Breast - Mammary Tissu | TCGA-BRCA | 45.23 | 7.17 |
| GTEX-1JMLX-2026-SM-CMKGN | Breast - Mammary Tissu | TCGA-BRCA | 49.85 | 10.92 |
| GTEX-1JMOU-2026-SM-CMKGU | Breast - Mammary Tissu | TCGA-BRCA | 35.34 | 9.2 |
| GTEX-1JMPY-0526-SM-CL54O | Breast - Mammary Tissu | TCGA-BRCA | 47.55 | 11.06 |
| GTEX-1JMPZ-1626-SM-COH31 | Breast - Mammary Tissu | TCGA-BRCA | 53.06 | 20.53 |
| GTEX-1JMQK-2626-SM-COH2W | Breast - Mammary Tissu | TCGA-BRCA | 44.21 | 13.71 |
| GTEX-1JN1M-3026-SM-CMKGK | Breast - Mammary Tissu | TCGA-BRCA | 83.15 | 23.65 |
| GTEX-1JN6P-2426-SM-ARL99 | Breast - Mammary Tissu | TCGA-BRCA | 34.5 | 14.25 |
| GTEX-1K2DA-1726-SM-CGQGS | Breast - Mammary Tissu | TCGA-BRCA | 48.23 | 14.69 |
| GTEX-1K9T9-1426-SM-DIPFQ | Breast - Mammary Tissu | TCGA-BRCA | 49.67 | 15.46 |
| GTEX-1KAFJ-0226-SM-DIPEL | Breast - Mammary Tissu | TCGA-BRCA | 52.61 | 12.7 |
| GTEX-1KANA-2026-SM-DIPFB | Breast - Mammary Tissu | TCGA-BRCA | 57.86 | 12.31 |
| GTEX-1KANC-1426-SM-CYPT9 | Breast - Mammary Tissu | TCGA-BRCA | 42.93 | 13.39 |
| GTEX-1KD4Q-2326-SM-D4P3M | Breast - Mammary Tissu | TCGA-BRCA | 47.78 | 16.19 |
| GTEX-1KD5A-1926-SM-DKPQA | Breast - Mammary Tissu | TCGA-BRCA | 41.43 | 12.21 |
| GTEX-1KXAM-1026-SM-CY8IA | Breast - Mammary Tissu | TCGA-BRCA | 40.52 | 11.79 |
| GTEX-1L5NE-2126-SM-CY8IH | Breast - Mammary Tissu | TCGA-BRCA | 34.29 | 9.26 |
| GTEX-1LBAC-0926-SM-CXKY9 | Breast - Mammary Tissu | TCGA-BRCA | 41.38 | 10.65 |
| GTEX-1LC47-1226-SM-D4P44 | Breast - Mammary Tissu | TCGA-BRCA | 49.94 | 13.61 |
| GTEX-1LG7Z-0726-SM-CXKYD | Breast - Mammary Tissu | TCGA-BRCA | 43.42 | 10.01 |
| GTEX-1LGRB-1826-SM-CNPQL | Breast - Mammary Tissu | TCGA-BRCA | 49.88 | 17.23 |
| GTEX-1LNCM-0226-SM-DIPF4 | Breast - Mammary Tissu | TCGA-BRCA | 46.23 | 13.81 |
| GTEX-1LSNL-2826-SM-E9U5L | Breast - Mammary Tissu | TCGA-BRCA | 42.11 | 11.35 |
| GTEX-1LSNM-1326-SM-E9TJ1 | Breast - Mammary Tissu | TCGA-BRCA | 48.7 | 11.44 |
| GTEX-1LSVX-2426-SM-E6CJX | Breast - Mammary Tissu | TCGA-BRCA | 30 | 10.68 |
| GTEX-1LVAM-0926-SM-E9TJ2 | Breast - Mammary Tissu | TCGA-BRCA | 56.42 | 13.86 |
| GTEX-1LVAN-1326-SM-CNPQE | Breast - Mammary Tissu | TCGA-BRCA | 39.51 | 16.65 |
| GTEX-1M4P7-2626-SM-E9U5F | Breast - Mammary Tissu | TCGA-BRCA | 44.28 | 11.35 |

| | | | | |
|---|---|---|---|---|
| GTEX-1M5QR-2526-SM-E9U5P | Breast - Mammary Tissu | TCGA-BRCA | 47.67 | 16.23 |
| GTEX-1MA7W-1026-SM-DKPQK | Breast - Mammary Tissu | TCGA-BRCA | 53.88 | 10.94 |
| GTEX-1MA7X-1626-SM-E9U5B | Breast - Mammary Tissu | TCGA-BRCA | 42.62 | 9.62 |
| GTEX-1MCC2-2126-SM-EWRON | Breast - Mammary Tissu | TCGA-BRCA | 66.17 | 16.6 |
| GTEX-1MCYP-0126-SM-E6CIS | Breast - Mammary Tissu | TCGA-BRCA | 40.51 | 11.07 |
| GTEX-1MJK2-2226-SM-EVYCR | Breast - Mammary Tissu | TCGA-BRCA | 55.56 | 18.39 |
| GTEX-1MJK3-1126-SM-E9J3F | Breast - Mammary Tissu | TCGA-BRCA | 28.47 | 8.36 |
| GTEX-1N2EE-2126-SM-E8VNN | Breast - Mammary Tissu | TCGA-BRCA | 45.21 | 15.49 |
| GTEX-1N2EF-2226-SM-E9J33 | Breast - Mammary Tissu | TCGA-BRCA | 75.53 | 8.57 |
| GTEX-1N5O9-1126-SM-DTXEQ | Breast - Mammary Tissu | TCGA-BRCA | 58.77 | 15.18 |
| GTEX-1NHNU-0926-SM-E9J2T | Breast - Mammary Tissu | TCGA-BRCA | 38.69 | 12.51 |
| GTEX-1NUQO-2226-SM-E9U68 | Breast - Mammary Tissu | TCGA-BRCA | 31.84 | 8.95 |
| GTEX-1NV8Z-2826-SM-E8VME | Breast - Mammary Tissu | TCGA-BRCA | 53.2 | 11.77 |
| GTEX-1OKEX-0826-SM-E9J2B | Breast - Mammary Tissu | TCGA-BRCA | 34.59 | 8.61 |
| GTEX-1P4AB-1826-SM-E76PU | Breast - Mammary Tissu | TCGA-BRCA | 42.88 | 14.02 |
| GTEX-1PDJ9-1126-SM-EXUSW | Breast - Mammary Tissu | TCGA-BRCA | 40.16 | 13.87 |
| GTEX-1POEN-0926-SM-DTX88 | Breast - Mammary Tissu | TCGA-BRCA | 40.06 | 15.95 |
| GTEX-1PPGY-1526-SM-EVYBK | Breast - Mammary Tissu | TCGA-BRCA | 54.51 | 11.15 |
| GTEX-1PPH7-0826-SM-E8VNX | Breast - Mammary Tissu | TCGA-BRCA | 36.59 | 8.43 |
| GTEX-1QCLY-1626-SM-DTXDW | Breast - Mammary Tissu | TCGA-BRCA | 45.99 | 14.38 |
| GTEX-1QCLZ-2326-SM-DTXDO | Breast - Mammary Tissu | TCGA-BRCA | 39.17 | 12.93 |
| GTEX-1QEPI-0926-SM-DTXF3 | Breast - Mammary Tissu | TCGA-BRCA | 46.2 | 14.46 |
| GTEX-1QMI2-0726-SM-E9J4F | Breast - Mammary Tissu | TCGA-BRCA | 42.9 | 10.79 |
| GTEX-1QP28-1226-SM-E9J2M | Breast - Mammary Tissu | TCGA-BRCA | 37.99 | 5.85 |
| GTEX-1QP29-1726-SM-E8VMQ | Breast - Mammary Tissu | TCGA-BRCA | 73.01 | 18.38 |
| GTEX-1QP9N-1826-SM-E8VMP | Breast - Mammary Tissu | TCGA-BRCA | 23.55 | 6.71 |
| GTEX-1QPFJ-2026-SM-EVYBE | Breast - Mammary Tissu | TCGA-BRCA | 37.99 | 7.69 |
| GTEX-1QW4Y-1026-SM-DTXE2 | Breast - Mammary Tissu | TCGA-BRCA | 35.96 | 9.93 |
| GTEX-1R46S-2526-SM-E8VN2 | Breast - Mammary Tissu | TCGA-BRCA | 48.07 | 10.78 |
| GTEX-1R7EU-1226-SM-E8VMR | Breast - Mammary Tissu | TCGA-BRCA | 45.57 | 11.37 |
| GTEX-1R7EV-1526-SM-E8VOD | Breast - Mammary Tissu | TCGA-BRCA | 19.01 | 5.64 |

| | | | | |
|---|---|---|---|---|
| GTEX-1R9JW-0326-SM-EAZ45 | Breast - Mammary Tissu | TCGA-BRCA | 33.84 | 8.08 |
| GTEX-1R9PM-1826-SM-EVYBH | Breast - Mammary Tissu | TCGA-BRCA | 30.49 | 6.07 |
| GTEX-1R9PN-0326-SM-DTX85 | Breast - Mammary Tissu | TCGA-BRCA | 47.37 | 14.73 |
| GTEX-1RAZQ-1226-SM-EWRNL | Breast - Mammary Tissu | TCGA-BRCA | 49.87 | 14.7 |
| GTEX-1RAZS-1226-SM-E8VMT | Breast - Mammary Tissu | TCGA-BRCA | 40.12 | 12.46 |
| GTEX-1RLM8-2226-SM-EVYD1 | Breast - Mammary Tissu | TCGA-BRCA | 29.07 | 8.26 |
| GTEX-1RQEC-2026-SM-DTXDM | Breast - Mammary Tissu | TCGA-BRCA | 51.37 | 15.56 |
| GTEX-1S82U-1126-SM-E9U6I | Breast - Mammary Tissu | TCGA-BRCA | 41.09 | 10.54 |
| GTEX-1S82Z-0726-SM-E6CHX | Breast - Mammary Tissu | TCGA-BRCA | 27.26 | 6.6 |
| GTEX-PSDG-1626-SM-48TCQ | Breast - Mammary Tissu | TCGA-BRCA | 26.68 | 8.42 |
| GTEX-Q2AG-0326-SM-48U1O | Breast - Mammary Tissu | TCGA-BRCA | 44.06 | 13.77 |
| GTEX-QDT8-0626-SM-48TYW | Breast - Mammary Tissu | TCGA-BRCA | 43.78 | 17.34 |
| GTEX-QEG5-0726-SM-D4P1T | Breast - Mammary Tissu | TCGA-BRCA | 62.1 | 22.75 |
| GTEX-QEL4-2126-SM-447AE | Breast - Mammary Tissu | TCGA-BRCA | 71.73 | 19.84 |
| GTEX-QMRM-1626-SM-D4P1U | Breast - Mammary Tissu | TCGA-BRCA | 27.98 | 8.92 |
| GTEX-QVJO-1826-SM-447C9 | Breast - Mammary Tissu | TCGA-BRCA | 32.85 | 7.89 |
| GTEX-R3RS-0626-SM-48FE1 | Breast - Mammary Tissu | TCGA-BRCA | 49.79 | 23.87 |
| GTEX-R53T-1526-SM-48FEK | Breast - Mammary Tissu | TCGA-BRCA | 40.41 | 10.29 |
| GTEX-R55D-0826-SM-48FEA | Breast - Mammary Tissu | TCGA-BRCA | 37.8 | 7.83 |
| GTEX-REY6-2426-SM-48FF5 | Breast - Mammary Tissu | TCGA-BRCA | 46.19 | 10.51 |
| GTEX-RU1J-0626-SM-4WAWY | Breast - Mammary Tissu | TCGA-BRCA | 39.63 | 11.98 |
| GTEX-RU72-0626-SM-46MUI | Breast - Mammary Tissu | TCGA-BRCA | 47.06 | 18.3 |
| GTEX-RUSQ-2026-SM-4GIAK | Breast - Mammary Tissu | TCGA-BRCA | 41.03 | 17.23 |
| GTEX-RWS6-1926-SM-47JXY | Breast - Mammary Tissu | TCGA-BRCA | 37.19 | 13.58 |
| GTEX-S32W-2026-SM-4AD6E | Breast - Mammary Tissu | TCGA-BRCA | 39.56 | 10.8 |
| GTEX-S33H-0326-SM-4AD6N | Breast - Mammary Tissu | TCGA-BRCA | 37.64 | 16.4 |
| GTEX-S341-1526-SM-4AD6K | Breast - Mammary Tissu | TCGA-BRCA | 40.32 | 15 |
| GTEX-S4P3-1326-SM-4AD6V | Breast - Mammary Tissu | TCGA-BRCA | 45.11 | 16.26 |
| GTEX-S4Q7-1126-SM-4AD6R | Breast - Mammary Tissu | TCGA-BRCA | 36.2 | 8.67 |
| GTEX-S4UY-0726-SM-4AD6X | Breast - Mammary Tissu | TCGA-BRCA | 36.03 | 11.66 |
| GTEX-S7SE-0826-SM-4AT4D | Breast - Mammary Tissu | TCGA-BRCA | 54.99 | 13.17 |

| | | | | |
|---|---|---|---|---|
| GTEX-SE5C-2126-SM-4BRUJ | Breast - Mammary Tissu | TCGA-BRCA | 44.67 | 10.44 |
| GTEX-T2IS-1526-SM-32QPR | Breast - Mammary Tissu | TCGA-BRCA | 66.09 | 15.41 |
| GTEX-T2YK-2226-SM-32QPT | Breast - Mammary Tissu | TCGA-BRCA | 42.95 | 21.8 |
| GTEX-T5JC-2126-SM-32PMO | Breast - Mammary Tissu | TCGA-BRCA | 38.75 | 16.29 |
| GTEX-T5JW-2026-SM-4DM63 | Breast - Mammary Tissu | TCGA-BRCA | 41.63 | 18.85 |
| GTEX-T6MN-0726-SM-32PML | Breast - Mammary Tissu | TCGA-BRCA | 57.75 | 16.79 |
| GTEX-T6MO-0326-SM-32QOK | Breast - Mammary Tissu | TCGA-BRCA | 34.67 | 11.63 |
| GTEX-TKQ1-0226-SM-33HB5 | Breast - Mammary Tissu | TCGA-BRCA | 27.98 | 9.33 |
| GTEX-TKQ2-1826-SM-33HB2 | Breast - Mammary Tissu | TCGA-BRCA | 39.13 | 14.77 |
| GTEX-TML8-1226-SM-32QON | Breast - Mammary Tissu | TCGA-BRCA | 39.02 | 18.33 |
| GTEX-TMMY-0726-SM-33HBE | Breast - Mammary Tissu | TCGA-BRCA | 45.11 | 12.55 |
| GTEX-U3ZH-1426-SM-4DXSR | Breast - Mammary Tissu | TCGA-BRCA | 51.12 | 18.78 |
| GTEX-U3ZN-1926-SM-4DXSG | Breast - Mammary Tissu | TCGA-BRCA | 40.75 | 13.26 |
| GTEX-U412-1826-SM-4DXTJ | Breast - Mammary Tissu | TCGA-BRCA | 42.13 | 8.4 |
| GTEX-U8XE-0826-SM-4E3J1 | Breast - Mammary Tissu | TCGA-BRCA | 36.19 | 14.99 |
| GTEX-UJHI-1426-SM-3DB9C | Breast - Mammary Tissu | TCGA-BRCA | 47.32 | 10.03 |
| GTEX-UPK5-2326-SM-3P5Z8 | Breast - Mammary Tissu | TCGA-BRCA | 49.79 | 14.58 |
| GTEX-UTHO-1026-SM-3GAF7 | Breast - Mammary Tissu | TCGA-BRCA | 39.76 | 11.91 |
| GTEX-V955-2026-SM-3GAFA | Breast - Mammary Tissu | TCGA-BRCA | 30.23 | 18.44 |
| GTEX-VJWN-0726-SM-3GIJ8 | Breast - Mammary Tissu | TCGA-BRCA | 54.84 | 17.54 |
| GTEX-VUSG-2226-SM-4KKZO | Breast - Mammary Tissu | TCGA-BRCA | 53.67 | 13.84 |
| GTEX-W5X1-2326-SM-3GIL6 | Breast - Mammary Tissu | TCGA-BRCA | 43.67 | 23 |
| GTEX-WFG8-2026-SM-EXOHO | Breast - Mammary Tissu | TCGA-BRCA | 51.09 | 16.07 |
| GTEX-WFON-1826-SM-3GILG | Breast - Mammary Tissu | TCGA-BRCA | 34.08 | 10.17 |
| GTEX-WHSB-1926-SM-EXOHH | Breast - Mammary Tissu | TCGA-BRCA | 23.86 | 7.38 |
| GTEX-WI4N-1426-SM-3LK7H | Breast - Mammary Tissu | TCGA-BRCA | 35.56 | 15.9 |
| GTEX-WOFL-0826-SM-3MJG1 | Breast - Mammary Tissu | TCGA-BRCA | 49.99 | 13.75 |
| GTEX-WRHU-0326-SM-3MJFY | Breast - Mammary Tissu | TCGA-BRCA | 41.04 | 14.38 |
| GTEX-WXYG-2226-SM-4E3IM | Breast - Mammary Tissu | TCGA-BRCA | 40.54 | 14.93 |
| GTEX-WY7C-2726-SM-3NB3P | Breast - Mammary Tissu | TCGA-BRCA | 30.65 | 11.86 |
| GTEX-WYBS-0926-SM-3NM94 | Breast - Mammary Tissu | TCGA-BRCA | 35.69 | 17.05 |

| | | | | |
|---|---|---|---|---|
| GTEX-WYJK-1326-SM-3NB2T | Breast - Mammary Tissu | TCGA-BRCA | 50.23 | 10.72 |
| GTEX-WYVS-1726-SM-3NMAY | Breast - Mammary Tissu | TCGA-BRCA | 48.09 | 18.65 |
| GTEX-X15G-1626-SM-3NMB3 | Breast - Mammary Tissu | TCGA-BRCA | 30.71 | 15.37 |
| GTEX-X261-0626-SM-3NMD9 | Breast - Mammary Tissu | TCGA-BRCA | 54.89 | 11.17 |
| GTEX-X4EP-2926-SM-3P5YQ | Breast - Mammary Tissu | TCGA-BRCA | 45.12 | 17.5 |
| GTEX-X4XY-0926-SM-4E3JD | Breast - Mammary Tissu | TCGA-BRCA | 43.66 | 18.71 |
| GTEX-X62O-0526-SM-EWRMA | Breast - Mammary Tissu | TCGA-BRCA | 50.18 | 12.27 |
| GTEX-XBED-1626-SM-47JYN | Breast - Mammary Tissu | TCGA-BRCA | 46.32 | 9.98 |
| GTEX-XGQ4-0926-SM-4AT4U | Breast - Mammary Tissu | TCGA-BRCA | 31.89 | 6.88 |
| GTEX-XMD1-0826-SM-4AT52 | Breast - Mammary Tissu | TCGA-BRCA | 36.43 | 12.43 |
| GTEX-XMD2-0926-SM-4WWEF | Breast - Mammary Tissu | TCGA-BRCA | 48.39 | 14.05 |
| GTEX-XMK1-1126-SM-4IHJ8 | Breast - Mammary Tissu | TCGA-BRCA | 21.52 | 5.92 |
| GTEX-XOT4-0726-SM-4GIAW | Breast - Mammary Tissu | TCGA-BRCA | 45.47 | 17.87 |
| GTEX-XQ3S-1326-SM-4BOPQ | Breast - Mammary Tissu | TCGA-BRCA | 33.04 | 11.16 |
| GTEX-XQ8I-2426-SM-4WAXY | Breast - Mammary Tissu | TCGA-BRCA | 35.09 | 10.51 |
| GTEX-XUW1-2326-SM-4BOO5 | Breast - Mammary Tissu | TCGA-BRCA | 41.09 | 9.54 |
| GTEX-XUZC-1626-SM-4BRVP | Breast - Mammary Tissu | TCGA-BRCA | 37.4 | 14.08 |
| GTEX-XV7Q-2326-SM-4BRVZ | Breast - Mammary Tissu | TCGA-BRCA | 38.6 | 11.47 |
| GTEX-XYKS-1326-SM-4BRUN | Breast - Mammary Tissu | TCGA-BRCA | 55.79 | 18.43 |
| GTEX-Y111-2026-SM-4SOJA | Breast - Mammary Tissu | TCGA-BRCA | 31.75 | 10.99 |
| GTEX-Y114-2026-SM-4TT7L | Breast - Mammary Tissu | TCGA-BRCA | 46.17 | 12.28 |
| GTEX-Y3I4-1526-SM-4TT7K | Breast - Mammary Tissu | TCGA-BRCA | 42.15 | 8.71 |
| GTEX-Y3IK-2326-SM-4WWDT | Breast - Mammary Tissu | TCGA-BRCA | 42.17 | 15.28 |
| GTEX-Y5LM-1726-SM-4VDSX | Breast - Mammary Tissu | TCGA-BRCA | 45.28 | 12.94 |
| GTEX-Y5V5-2126-SM-4WWFO | Breast - Mammary Tissu | TCGA-BRCA | 46.57 | 14.27 |
| GTEX-Y5V6-2126-SM-4WWFX | Breast - Mammary Tissu | TCGA-BRCA | 36.48 | 15.37 |
| GTEX-Y8E4-1626-SM-5S2MW | Breast - Mammary Tissu | TCGA-BRCA | 47.67 | 10.53 |
| GTEX-Y8LW-1626-SM-5IFHX | Breast - Mammary Tissu | TCGA-BRCA | 46.34 | 11.38 |
| GTEX-Y9LG-1426-SM-5IFJZ | Breast - Mammary Tissu | TCGA-BRCA | 36.66 | 7.33 |
| GTEX-YB5E-1726-SM-5IFJ3 | Breast - Mammary Tissu | TCGA-BRCA | 25.48 | 5.45 |
| GTEX-YB5K-1626-SM-5IFIN | Breast - Mammary Tissu | TCGA-BRCA | 32.95 | 9.15 |

| | | | | |
|---|---|---|---|---|
| GTEX-YEC3-1026-SM-5IFI5 | Breast - Mammary Tissu | TCGA-BRCA | 42.82 | 9.32 |
| GTEX-YECK-0726-SM-EZ6LV | Breast - Mammary Tissu | TCGA-BRCA | 41.83 | 13.46 |
| GTEX-YFC4-1426-SM-5IFJG | Breast - Mammary Tissu | TCGA-BRCA | 50.21 | 17.34 |
| GTEX-YFCO-1826-SM-4W1YH | Breast - Mammary Tissu | TCGA-BRCA | 44.02 | 10.06 |
| GTEX-YJ8O-2226-SM-5IFHW | Breast - Mammary Tissu | TCGA-BRCA | 46.99 | 14.47 |
| GTEX-Z9EW-1526-SM-EZ6LU | Breast - Mammary Tissu | TCGA-BRCA | 32.46 | 13.25 |
| GTEX-ZA64-1526-SM-5CVMD | Breast - Mammary Tissu | TCGA-BRCA | 43.36 | 8.91 |
| GTEX-ZAB4-2526-SM-5HL8M | Breast - Mammary Tissu | TCGA-BRCA | 43.08 | 8.92 |
| GTEX-ZAJG-0626-SM-5HL8X | Breast - Mammary Tissu | TCGA-BRCA | 47.86 | 14.28 |
| GTEX-ZAK1-1326-SM-EVYAS | Breast - Mammary Tissu | TCGA-BRCA | 41.15 | 15.81 |
| GTEX-ZC5H-2626-SM-5J2MG | Breast - Mammary Tissu | TCGA-BRCA | 43.8 | 18.2 |
| GTEX-ZDTT-2126-SM-DKPOM | Breast - Mammary Tissu | TCGA-BRCA | 41.61 | 12.27 |
| GTEX-ZDXO-0126-SM-5S2ND | Breast - Mammary Tissu | TCGA-BRCA | 36.24 | 14.74 |
| GTEX-ZDYS-1126-SM-5K7UB | Breast - Mammary Tissu | TCGA-BRCA | 34.95 | 12.02 |
| GTEX-ZEX8-2226-SM-57WC6 | Breast - Mammary Tissu | TCGA-BRCA | 39.35 | 12.96 |
| GTEX-ZF29-1926-SM-5S2P1 | Breast - Mammary Tissu | TCGA-BRCA | 55.55 | 14.95 |
| GTEX-ZF2S-2026-SM-5E461 | Breast - Mammary Tissu | TCGA-BRCA | 51.49 | 18.19 |
| GTEX-ZF3C-2326-SM-5S2NC | Breast - Mammary Tissu | TCGA-BRCA | 52.22 | 18.79 |
| GTEX-ZGAY-1726-SM-DO91P | Breast - Mammary Tissu | TCGA-BRCA | 49.1 | 12 |
| GTEX-ZLFU-2126-SM-DO931 | Breast - Mammary Tissu | TCGA-BRCA | 52.69 | 10.33 |
| GTEX-ZLV1-1426-SM-DO92Z | Breast - Mammary Tissu | TCGA-BRCA | 40.45 | 10.92 |
| GTEX-ZLWG-1926-SM-DO92J | Breast - Mammary Tissu | TCGA-BRCA | 38.5 | 11.63 |
| GTEX-ZPIC-1126-SM-5BC7F | Breast - Mammary Tissu | TCGA-BRCA | 33.94 | 10.34 |
| GTEX-ZQG8-0726-SM-5P9H9 | Breast - Mammary Tissu | TCGA-BRCA | 28.88 | 8.48 |
| GTEX-ZQUD-1926-SM-51MSA | Breast - Mammary Tissu | TCGA-BRCA | 27.36 | 7.39 |
| GTEX-ZT9W-2026-SM-51MRA | Breast - Mammary Tissu | TCGA-BRCA | 34.33 | 6.76 |
| GTEX-ZTPG-2826-SM-57WGA | Breast - Mammary Tissu | TCGA-BRCA | 32.61 | 7.32 |
| GTEX-ZTTD-1026-SM-51MRD | Breast - Mammary Tissu | TCGA-BRCA | 68.99 | 18.69 |
| GTEX-ZTX8-1226-SM-4YCE9 | Breast - Mammary Tissu | TCGA-BRCA | 68.81 | 10.49 |
| GTEX-ZU9S-1926-SM-5NQBP | Breast - Mammary Tissu | TCGA-BRCA | 60.41 | 16.06 |
| GTEX-ZUA1-1526-SM-59HLS | Breast - Mammary Tissu | TCGA-BRCA | 45.21 | 12.02 |

| | | | | |
|---|---|---|---|---|
| GTEX-ZV6S-1826-SM-5NQ8D | Breast - Mammary Tissu | TCGA-BRCA | 42.33 | 10.93 |
| GTEX-ZV7C-1826-SM-5NQ83 | Breast - Mammary Tissu | TCGA-BRCA | 48.62 | 10.59 |
| GTEX-ZVE2-1226-SM-5NQ8R | Breast - Mammary Tissu | TCGA-BRCA | 44.38 | 16.73 |
| GTEX-ZVT2-1826-SM-5NQ8W | Breast - Mammary Tissu | TCGA-BRCA | 54.35 | 11.96 |
| GTEX-ZVT4-1026-SM-57WC4 | Breast - Mammary Tissu | TCGA-BRCA | 60.6 | 7.62 |
| GTEX-ZVTK-0326-SM-51MRR | Breast - Mammary Tissu | TCGA-BRCA | 50.43 | 14.83 |
| GTEX-ZVZQ-0826-SM-51MRF | Breast - Mammary Tissu | TCGA-BRCA | 41.51 | 12.07 |
| GTEX-ZWKS-2826-SM-5NQ74 | Breast - Mammary Tissu | TCGA-BRCA | 44.71 | 10.95 |
| GTEX-ZXES-0826-SM-5E43C | Breast - Mammary Tissu | TCGA-BRCA | 46.11 | 9.63 |
| GTEX-ZY6K-1626-SM-5GZWV | Breast - Mammary Tissu | TCGA-BRCA | 55.23 | 13.05 |
| GTEX-ZYFC-0826-SM-5E44K | Breast - Mammary Tissu | TCGA-BRCA | 42.8 | 10.27 |
| GTEX-ZYT6-0126-SM-5E45J | Breast - Mammary Tissu | TCGA-BRCA | 39.74 | 8.46 |
| GTEX-ZYW4-0826-SM-5GIDG | Breast - Mammary Tissu | TCGA-BRCA | 46.24 | 7.24 |
| GTEX-ZZ64-1226-SM-5E43R | Breast - Mammary Tissu | TCGA-BRCA | 40.28 | 6.27 |
| GTEX-ZZPU-0626-SM-5E43T | Breast - Mammary Tissu | TCGA-BRCA | 52.39 | 13.03 |
| GTEX-111CU-1226-SM-5EGIN | Colon - Sigmoid | TCGA-COAD | 45.6 | 13.69 |
| GTEX-111YS-1826-SM-5GIEP | Colon - Sigmoid | TCGA-COAD | 44.35 | 14.81 |
| GTEX-1122O-1426-SM-5H11R | Colon - Sigmoid | TCGA-COAD | 44 | 11.71 |
| GTEX-117XS-2226-SM-5GIE1 | Colon - Sigmoid | TCGA-COAD | 34.06 | 12.5 |
| GTEX-117YX-0526-SM-5EGJH | Colon - Sigmoid | TCGA-COAD | 43.45 | 12.86 |
| GTEX-11DXX-1926-SM-5EGJK | Colon - Sigmoid | TCGA-COAD | 52.59 | 10.69 |
| GTEX-11DXY-2026-SM-5N9CZ | Colon - Sigmoid | TCGA-COAD | 33.06 | 11.99 |
| GTEX-11DXZ-1626-SM-5GIDI | Colon - Sigmoid | TCGA-COAD | 36.44 | 13.39 |
| GTEX-11EI6-2626-SM-5PNVT | Colon - Sigmoid | TCGA-COAD | 40.37 | 13.5 |
| GTEX-11EM3-1626-SM-5N9CO | Colon - Sigmoid | TCGA-COAD | 61.05 | 20.91 |
| GTEX-11EMC-1426-SM-5N9E4 | Colon - Sigmoid | TCGA-COAD | 38.52 | 18.02 |
| GTEX-11EQ8-2626-SM-5N9BO | Colon - Sigmoid | TCGA-COAD | 89.73 | 16.64 |
| GTEX-11EQ9-1526-SM-5PNY7 | Colon - Sigmoid | TCGA-COAD | 37.2 | 11.1 |
| GTEX-11GS4-1826-SM-5HL4T | Colon - Sigmoid | TCGA-COAD | 37.47 | 14.21 |
| GTEX-11I78-1326-SM-5PNY9 | Colon - Sigmoid | TCGA-COAD | 55.68 | 11.24 |
| GTEX-11ONC-1926-SM-5GU75 | Colon - Sigmoid | TCGA-COAD | 32.37 | 14.45 |

| | | | | |
|---|---|---|---|---|
| GTEX-11P7K-1826-SM-5HL5A | Colon - Sigmoid | TCGA-COAD | 39.44 | 13.11 |
| GTEX-11P81-1426-SM-5P9JN | Colon - Sigmoid | TCGA-COAD | 50.39 | 10.7 |
| GTEX-11PRG-2026-SM-5GU5F | Colon - Sigmoid | TCGA-COAD | 27.48 | 11.14 |
| GTEX-11TT1-0926-SM-5GU5M | Colon - Sigmoid | TCGA-COAD | 56.62 | 24.05 |
| GTEX-11TUW-2326-SM-5EQMO | Colon - Sigmoid | TCGA-COAD | 44.37 | 8.62 |
| GTEX-11VI4-1426-SM-5GU6O | Colon - Sigmoid | TCGA-COAD | 36.61 | 11.49 |
| GTEX-11ZTS-1526-SM-5N9G7 | Colon - Sigmoid | TCGA-COAD | 34.03 | 10.98 |
| GTEX-11ZTT-1626-SM-5EQKR | Colon - Sigmoid | TCGA-COAD | 42.85 | 16.09 |
| GTEX-11ZU8-1126-SM-5EQ5K | Colon - Sigmoid | TCGA-COAD | 55.08 | 16.19 |
| GTEX-1211K-1826-SM-5EGJ2 | Colon - Sigmoid | TCGA-COAD | 42.16 | 9.01 |
| GTEX-1212Z-1826-SM-5FQSH | Colon - Sigmoid | TCGA-COAD | 38.6 | 13.12 |
| GTEX-12696-1926-SM-5EGJT | Colon - Sigmoid | TCGA-COAD | 41.74 | 7.5 |
| GTEX-12BJ1-2026-SM-5FQUH | Colon - Sigmoid | TCGA-COAD | 47.49 | 8.39 |
| GTEX-12KS4-2326-SM-5EGJB | Colon - Sigmoid | TCGA-COAD | 36.94 | 14.03 |
| GTEX-12WSD-2626-SM-5EGJE | Colon - Sigmoid | TCGA-COAD | 42.71 | 12.94 |
| GTEX-12WSG-2426-SM-5EQLZ | Colon - Sigmoid | TCGA-COAD | 32.27 | 15.39 |
| GTEX-12WSJ-1426-SM-5GCON | Colon - Sigmoid | TCGA-COAD | 42.94 | 11.89 |
| GTEX-12WSL-1826-SM-5LZW9 | Colon - Sigmoid | TCGA-COAD | 46.06 | 11.74 |
| GTEX-12WSM-1426-SM-5GCPA | Colon - Sigmoid | TCGA-COAD | 55.69 | 19.77 |
| GTEX-12ZZW-2426-SM-5DUWE | Colon - Sigmoid | TCGA-COAD | 40.62 | 11.76 |
| GTEX-12ZZX-2726-SM-5N9FT | Colon - Sigmoid | TCGA-COAD | 38.63 | 11.48 |
| GTEX-12ZZY-2726-SM-5EQ45 | Colon - Sigmoid | TCGA-COAD | 36.66 | 16.47 |
| GTEX-13111-1926-SM-5GCOL | Colon - Sigmoid | TCGA-COAD | 37.98 | 12.07 |
| GTEX-13113-1426-SM-5EGHU | Colon - Sigmoid | TCGA-COAD | 39.19 | 6.51 |
| GTEX-131XE-1026-SM-5EGKE | Colon - Sigmoid | TCGA-COAD | 57.21 | 13.05 |
| GTEX-131XG-1526-SM-5GCNQ | Colon - Sigmoid | TCGA-COAD | 48.1 | 16.33 |
| GTEX-131YS-2926-SM-5EGKI | Colon - Sigmoid | TCGA-COAD | 44.06 | 11.51 |
| GTEX-132AR-2826-SM-5IJFV | Colon - Sigmoid | TCGA-COAD | 72.46 | 14.69 |
| GTEX-132NY-2726-SM-5PNY2 | Colon - Sigmoid | TCGA-COAD | 38.65 | 13.67 |
| GTEX-1339X-1726-SM-5P9J9 | Colon - Sigmoid | TCGA-COAD | 45.05 | 10.18 |
| GTEX-1399R-2126-SM-5K7WZ | Colon - Sigmoid | TCGA-COAD | 43.5 | 14.2 |

| | | | | |
|---|---|---|---|---|
| GTEX-1399U-2226-SM-5KLZI | Colon - Sigmoid | TCGA-COAD | 41.39 | 10.26 |
| GTEX-139D8-2926-SM-5IJBM | Colon - Sigmoid | TCGA-COAD | 45.85 | 12.89 |
| GTEX-139T6-1426-SM-5P9J7 | Colon - Sigmoid | TCGA-COAD | 42.07 | 9.03 |
| GTEX-139TS-2226-SM-5J2N1 | Colon - Sigmoid | TCGA-COAD | 28.81 | 10.49 |
| GTEX-139UW-2326-SM-5KM1Y | Colon - Sigmoid | TCGA-COAD | 30.54 | 14.37 |
| GTEX-13CF2-1826-SM-5N9GK | Colon - Sigmoid | TCGA-COAD | 28.72 | 8.61 |
| GTEX-13CF3-1326-SM-5LZXZ | Colon - Sigmoid | TCGA-COAD | 40.59 | 13.89 |
| GTEX-13D11-0626-SM-5LZYY | Colon - Sigmoid | TCGA-COAD | 41.77 | 9.98 |
| GTEX-13FH7-1626-SM-5J2O9 | Colon - Sigmoid | TCGA-COAD | 49.64 | 11.81 |
| GTEX-13FHO-2726-SM-5K7VG | Colon - Sigmoid | TCGA-COAD | 29.15 | 11.41 |
| GTEX-13FTW-2126-SM-5K7YG | Colon - Sigmoid | TCGA-COAD | 39.52 | 10.75 |
| GTEX-13G51-2126-SM-5IJD9 | Colon - Sigmoid | TCGA-COAD | 38.89 | 14.26 |
| GTEX-13N2G-2226-SM-5IJCU | Colon - Sigmoid | TCGA-COAD | 34.72 | 16.03 |
| GTEX-13O1R-2426-SM-5KLZZ | Colon - Sigmoid | TCGA-COAD | 50.43 | 11.31 |
| GTEX-13O61-1526-SM-5KM3P | Colon - Sigmoid | TCGA-COAD | 44.36 | 10.69 |
| GTEX-13OVI-2126-SM-5KM3U | Colon - Sigmoid | TCGA-COAD | 44.74 | 7.3 |
| GTEX-13OVJ-1826-SM-5KM2D | Colon - Sigmoid | TCGA-COAD | 34.91 | 14.18 |
| GTEX-13OVK-1026-SM-7KFTC | Colon - Sigmoid | TCGA-COAD | 52.76 | 24.83 |
| GTEX-13OW5-2826-SM-5KM1G | Colon - Sigmoid | TCGA-COAD | 35.25 | 9.72 |
| GTEX-13OW6-2226-SM-5N9FL | Colon - Sigmoid | TCGA-COAD | 40.18 | 15.73 |
| GTEX-13OW8-2726-SM-5L3FZ | Colon - Sigmoid | TCGA-COAD | 35.1 | 11.24 |
| GTEX-13PL7-2426-SM-5J2N3 | Colon - Sigmoid | TCGA-COAD | 28.55 | 13.7 |
| GTEX-13PVQ-2726-SM-5L3H7 | Colon - Sigmoid | TCGA-COAD | 36 | 12.14 |
| GTEX-13PVR-1726-SM-5Q5EC | Colon - Sigmoid | TCGA-COAD | 50.14 | 9.85 |
| GTEX-13QBU-2026-SM-5KLZB | Colon - Sigmoid | TCGA-COAD | 37.97 | 14.22 |
| GTEX-13RTJ-1926-SM-5YYA2 | Colon - Sigmoid | TCGA-COAD | 35.53 | 11.74 |
| GTEX-13SLW-2826-SM-5Q5ES | Colon - Sigmoid | TCGA-COAD | 43.33 | 13.82 |
| GTEX-13U4I-2626-SM-5SI9C | Colon - Sigmoid | TCGA-COAD | 55.36 | 13.47 |
| GTEX-13VXT-2026-SM-5L3EW | Colon - Sigmoid | TCGA-COAD | 45.55 | 8.77 |
| GTEX-13W3W-2226-SM-5LU4M | Colon - Sigmoid | TCGA-COAD | 44.77 | 12.4 |
| GTEX-13X6I-2726-SM-5Q5ET | Colon - Sigmoid | TCGA-COAD | 33.93 | 7.65 |

| | | | | |
|---|---|---|---|---|
| GTEX-13X6J-2826-SM-7EPGA | Colon - Sigmoid | TCGA-COAD | 30 | 12.22 |
| GTEX-13X6K-2726-SM-5O9DG | Colon - Sigmoid | TCGA-COAD | 41.03 | 12.05 |
| GTEX-144GM-1126-SM-79OJK | Colon - Sigmoid | TCGA-COAD | 44.95 | 10.52 |
| GTEX-145LS-2926-SM-5O99G | Colon - Sigmoid | TCGA-COAD | 39.85 | 14.77 |
| GTEX-145LT-0826-SM-5S2QL | Colon - Sigmoid | TCGA-COAD | 34.36 | 12.8 |
| GTEX-145ME-1126-SM-5SIAT | Colon - Sigmoid | TCGA-COAD | 56.6 | 13.96 |
| GTEX-145MH-2726-SM-5QGQ6 | Colon - Sigmoid | TCGA-COAD | 32.63 | 15.64 |
| GTEX-145MO-2626-SM-5QGPD | Colon - Sigmoid | TCGA-COAD | 38.82 | 11.9 |
| GTEX-146FH-2426-SM-5Q5EW | Colon - Sigmoid | TCGA-COAD | 54.68 | 17.62 |
| GTEX-1477Z-2126-SM-5Q5CM | Colon - Sigmoid | TCGA-COAD | 38.47 | 11.46 |
| GTEX-147GR-2826-SM-5RQK8 | Colon - Sigmoid | TCGA-COAD | 34.38 | 15.27 |
| GTEX-147JS-2626-SM-5S2UX | Colon - Sigmoid | TCGA-COAD | 35.47 | 12.33 |
| GTEX-148VJ-2426-SM-5NQ9Y | Colon - Sigmoid | TCGA-COAD | 37.61 | 10.7 |
| GTEX-14A5H-1626-SM-5SIAN | Colon - Sigmoid | TCGA-COAD | 30.92 | 11.11 |
| GTEX-14A5I-2426-SM-5Q5CP | Colon - Sigmoid | TCGA-COAD | 31.68 | 12.07 |
| GTEX-14AS3-1026-SM-5TDD7 | Colon - Sigmoid | TCGA-COAD | 60.74 | 10.74 |
| GTEX-14BIN-2426-SM-5TDCF | Colon - Sigmoid | TCGA-COAD | 31.82 | 10.75 |
| GTEX-14BMU-1326-SM-5RQIH | Colon - Sigmoid | TCGA-COAD | 58.76 | 18.32 |
| GTEX-14C39-1626-SM-5S2P4 | Colon - Sigmoid | TCGA-COAD | 52.53 | 16.62 |
| GTEX-14C5O-2626-SM-5RQI5 | Colon - Sigmoid | TCGA-COAD | 32.73 | 11.23 |
| GTEX-14DAQ-2926-SM-664MV | Colon - Sigmoid | TCGA-COAD | 42.41 | 11.78 |
| GTEX-14E1K-1626-SM-664MO | Colon - Sigmoid | TCGA-COAD | 30.82 | 15.31 |
| GTEX-14E6C-2226-SM-5QGQG | Colon - Sigmoid | TCGA-COAD | 30.6 | 8.99 |
| GTEX-14E6E-1726-SM-5S2R6 | Colon - Sigmoid | TCGA-COAD | 48.3 | 10.86 |
| GTEX-14JG6-1626-SM-5YYBC | Colon - Sigmoid | TCGA-COAD | 39.31 | 13.29 |
| GTEX-14PHX-1126-SM-5YYA5 | Colon - Sigmoid | TCGA-COAD | 37.77 | 12.64 |
| GTEX-14PJ3-1226-SM-5ZZVY | Colon - Sigmoid | TCGA-COAD | 37.13 | 12.35 |
| GTEX-14PJ4-1826-SM-5YY97 | Colon - Sigmoid | TCGA-COAD | 39.96 | 7.83 |
| GTEX-14PK6-1826-SM-69LOM | Colon - Sigmoid | TCGA-COAD | 48.58 | 8.08 |
| GTEX-14PKV-1926-SM-686YU | Colon - Sigmoid | TCGA-COAD | 38.42 | 9.64 |
| GTEX-14PN3-1926-SM-6EU1Y | Colon - Sigmoid | TCGA-COAD | 54.08 | 10.5 |

| | | | | |
|---|---|---|---|---|
| GTEX-15CHQ-2626-SM-686YZ | Colon - Sigmoid | TCGA-COAD | 40.47 | 13.63 |
| GTEX-15DYW-2426-SM-7KUG8 | Colon - Sigmoid | TCGA-COAD | 45.59 | 12.84 |
| GTEX-15DZA-1026-SM-6PAND | Colon - Sigmoid | TCGA-COAD | 45.77 | 10.9 |
| GTEX-15EO6-2326-SM-6LPKK | Colon - Sigmoid | TCGA-COAD | 40.77 | 15.64 |
| GTEX-15ER7-2826-SM-6PAN2 | Colon - Sigmoid | TCGA-COAD | 67.83 | 21.14 |
| GTEX-15ETS-2326-SM-7KUGC | Colon - Sigmoid | TCGA-COAD | 43.24 | 12.54 |
| GTEX-15F5U-1426-SM-7KUN2 | Colon - Sigmoid | TCGA-COAD | 45.38 | 11.13 |
| GTEX-15FZZ-1826-SM-6LLJP | Colon - Sigmoid | TCGA-COAD | 39.62 | 11.73 |
| GTEX-15G19-2726-SM-6LPKG | Colon - Sigmoid | TCGA-COAD | 62.76 | 22.79 |
| GTEX-15G1A-1926-SM-7KUFF | Colon - Sigmoid | TCGA-COAD | 54.71 | 17.3 |
| GTEX-15RIE-2026-SM-7KFRW | Colon - Sigmoid | TCGA-COAD | 51.15 | 14.34 |
| GTEX-15RIF-1426-SM-7KUKQ | Colon - Sigmoid | TCGA-COAD | 41.77 | 13.26 |
| GTEX-15RJ7-1726-SM-6PAL3 | Colon - Sigmoid | TCGA-COAD | 46.75 | 14.01 |
| GTEX-15RJE-2226-SM-6M46A | Colon - Sigmoid | TCGA-COAD | 27.64 | 11.23 |
| GTEX-15UF6-2726-SM-7KULY | Colon - Sigmoid | TCGA-COAD | 32.7 | 12.1 |
| GTEX-15UF7-1326-SM-7KUKV | Colon - Sigmoid | TCGA-COAD | 45.95 | 11.82 |
| GTEX-16BQI-2026-SM-7KULQ | Colon - Sigmoid | TCGA-COAD | 29.72 | 10.42 |
| GTEX-16MTA-1626-SM-6M47N | Colon - Sigmoid | TCGA-COAD | 50.64 | 7.95 |
| GTEX-16NGA-0926-SM-718AG | Colon - Sigmoid | TCGA-COAD | 44 | 17.97 |
| GTEX-16XZZ-2726-SM-7EWDO | Colon - Sigmoid | TCGA-COAD | 31.11 | 9.23 |
| GTEX-16Z82-2426-SM-7KUMS | Colon - Sigmoid | TCGA-COAD | 29.02 | 12.19 |
| GTEX-17EUY-1726-SM-79OK4 | Colon - Sigmoid | TCGA-COAD | 42.19 | 11.93 |
| GTEX-17F96-2126-SM-7KFSM | Colon - Sigmoid | TCGA-COAD | 38.25 | 14.05 |
| GTEX-17F9Y-2626-SM-7IGN9 | Colon - Sigmoid | TCGA-COAD | 46.11 | 10.48 |
| GTEX-17GQL-1626-SM-7LG69 | Colon - Sigmoid | TCGA-COAD | 38.58 | 10.19 |
| GTEX-17HG3-1126-SM-7DUFC | Colon - Sigmoid | TCGA-COAD | 23.21 | 7.06 |
| GTEX-17HGU-2626-SM-79OKN | Colon - Sigmoid | TCGA-COAD | 42.58 | 11.82 |
| GTEX-17HHE-1626-SM-7DHLN | Colon - Sigmoid | TCGA-COAD | 30.43 | 12.56 |
| GTEX-17JCI-2826-SM-7IGNA | Colon - Sigmoid | TCGA-COAD | 30.66 | 11.51 |
| GTEX-17KNJ-1826-SM-7LT9N | Colon - Sigmoid | TCGA-COAD | 53.6 | 17.18 |
| GTEX-17MFQ-1226-SM-7KFSA | Colon - Sigmoid | TCGA-COAD | 45.9 | 14.39 |

| | | | | |
|---|---|---|---|---|
| GTEX-183FY-1426-SM-7DHLM | Colon - Sigmoid | TCGA-COAD | 47.8 | 10.28 |
| GTEX-183WM-2526-SM-731BP | Colon - Sigmoid | TCGA-COAD | 38.92 | 12.57 |
| GTEX-18465-2726-SM-7LT9I | Colon - Sigmoid | TCGA-COAD | 30.4 | 12.38 |
| GTEX-18A66-2626-SM-72D67 | Colon - Sigmoid | TCGA-COAD | 40.62 | 11.88 |
| GTEX-18A67-2226-SM-7LT9Z | Colon - Sigmoid | TCGA-COAD | 47.83 | 14.46 |
| GTEX-18A6Q-2226-SM-7LG61 | Colon - Sigmoid | TCGA-COAD | 30.86 | 11.32 |
| GTEX-18D9A-1326-SM-7LT8P | Colon - Sigmoid | TCGA-COAD | 51.71 | 11.2 |
| GTEX-18QFQ-1726-SM-731C9 | Colon - Sigmoid | TCGA-COAD | 34.23 | 8.29 |
| GTEX-1A32A-2626-SM-73KX9 | Colon - Sigmoid | TCGA-COAD | 46.56 | 11.39 |
| GTEX-1A3MV-1026-SM-72D5U | Colon - Sigmoid | TCGA-COAD | 40.51 | 17.16 |
| GTEX-1A3MX-2326-SM-718B4 | Colon - Sigmoid | TCGA-COAD | 36.6 | 6.47 |
| GTEX-1AMEY-1826-SM-72D5L | Colon - Sigmoid | TCGA-COAD | 39.34 | 11.82 |
| GTEX-1AMFI-1826-SM-73KWB | Colon - Sigmoid | TCGA-COAD | 35.72 | 12.38 |
| GTEX-1AX9J-2526-SM-73KUY | Colon - Sigmoid | TCGA-COAD | 35.91 | 10.48 |
| GTEX-1AX9K-1626-SM-73KUS | Colon - Sigmoid | TCGA-COAD | 41.18 | 13.16 |
| GTEX-1AYCT-1926-SM-793AE | Colon - Sigmoid | TCGA-COAD | 38.95 | 12.08 |
| GTEX-1B8KE-1526-SM-7DUG2 | Colon - Sigmoid | TCGA-COAD | 38.51 | 7.64 |
| GTEX-1B8L1-1926-SM-7IGND | Colon - Sigmoid | TCGA-COAD | 45.95 | 13.9 |
| GTEX-1B932-2126-SM-7IGMU | Colon - Sigmoid | TCGA-COAD | 44.35 | 14.09 |
| GTEX-1B97I-1526-SM-73KUK | Colon - Sigmoid | TCGA-COAD | 44.35 | 10.56 |
| GTEX-1B996-2726-SM-7DUGG | Colon - Sigmoid | TCGA-COAD | 44.32 | 10.28 |
| GTEX-1BAJH-2726-SM-7IGNJ | Colon - Sigmoid | TCGA-COAD | 33.65 | 11.22 |
| GTEX-1C2JI-1726-SM-73KUW | Colon - Sigmoid | TCGA-COAD | 31.93 | 10.45 |
| GTEX-1C475-1526-SM-7DHM3 | Colon - Sigmoid | TCGA-COAD | 40.6 | 7.78 |
| GTEX-1C4CL-1626-SM-7DUF3 | Colon - Sigmoid | TCGA-COAD | 40.6 | 12.51 |
| GTEX-1C64N-2226-SM-7SB7C | Colon - Sigmoid | TCGA-COAD | 40.52 | 14.87 |
| GTEX-1C64O-2426-SM-7P8R3 | Colon - Sigmoid | TCGA-COAD | 28.62 | 8.64 |
| GTEX-1C6VQ-2626-SM-7IGMQ | Colon - Sigmoid | TCGA-COAD | 40.12 | 14.71 |
| GTEX-1CAMR-1026-SM-7P8QI | Colon - Sigmoid | TCGA-COAD | 36.04 | 9.93 |
| GTEX-1CAMS-2626-SM-7PBY3 | Colon - Sigmoid | TCGA-COAD | 46.32 | 14.97 |
| GTEX-1CB4F-1926-SM-7MKFN | Colon - Sigmoid | TCGA-COAD | 32.5 | 17.18 |

| | | | | |
|---|---|---|---|---|
| GTEX-1CB4G-2726-SM-79OO4 | Colon - Sigmoid | TCGA-COAD | 37.67 | 14.17 |
| GTEX-1E2YA-2526-SM-7IGOU | Colon - Sigmoid | TCGA-COAD | 20.94 | 8.96 |
| GTEX-1EH9U-1926-SM-7P8QM | Colon - Sigmoid | TCGA-COAD | 33.61 | 10.55 |
| GTEX-1EMGI-2626-SM-7IGNR | Colon - Sigmoid | TCGA-COAD | 36.07 | 11.28 |
| GTEX-1EU9M-2726-SM-7P8QD | Colon - Sigmoid | TCGA-COAD | 29.34 | 10.4 |
| GTEX-1F5PK-1726-SM-7RHHW | Colon - Sigmoid | TCGA-COAD | 42.47 | 14.34 |
| GTEX-1F5PL-1926-SM-7MXTP | Colon - Sigmoid | TCGA-COAD | 70.97 | 16.76 |
| GTEX-1F75W-2126-SM-7RHIA | Colon - Sigmoid | TCGA-COAD | 38.2 | 12.42 |
| GTEX-1GF9X-1426-SM-7RHGX | Colon - Sigmoid | TCGA-COAD | 48.72 | 12.49 |
| GTEX-1GL5R-1726-SM-9KNUX | Colon - Sigmoid | TCGA-COAD | 48.32 | 10.14 |
| GTEX-1GMR2-1826-SM-9KNUP | Colon - Sigmoid | TCGA-COAD | 33 | 9.82 |
| GTEX-1GMR3-2526-SM-7P8QS | Colon - Sigmoid | TCGA-COAD | 32.27 | 9.55 |
| GTEX-1GMR8-2926-SM-9KNW5 | Colon - Sigmoid | TCGA-COAD | 42.82 | 12.81 |
| GTEX-1GMRU-2626-SM-9WPPM | Colon - Sigmoid | TCGA-COAD | 33.31 | 11.67 |
| GTEX-1GN73-2726-SM-9WYUB | Colon - Sigmoid | TCGA-COAD | 33.68 | 11.18 |
| GTEX-1GPI7-1926-SM-9KNUH | Colon - Sigmoid | TCGA-COAD | 34.84 | 11.83 |
| GTEX-1GTWX-2626-SM-9WPOT | Colon - Sigmoid | TCGA-COAD | 34.45 | 12.88 |
| GTEX-1GZ4H-1326-SM-7P8TE | Colon - Sigmoid | TCGA-COAD | 46.59 | 9.87 |
| GTEX-1H1DE-1626-SM-9KNU5 | Colon - Sigmoid | TCGA-COAD | 37.85 | 9.7 |
| GTEX-1H1E6-1826-SM-A9G21 | Colon - Sigmoid | TCGA-COAD | 50.39 | 12.31 |
| GTEX-1H1ZS-2626-SM-9QEIX | Colon - Sigmoid | TCGA-COAD | 33.54 | 7.39 |
| GTEX-1H23P-2626-SM-9KNVN | Colon - Sigmoid | TCGA-COAD | 26.53 | 10.27 |
| GTEX-1H3NZ-2526-SM-9KNVO | Colon - Sigmoid | TCGA-COAD | 37.27 | 13.66 |
| GTEX-1H3VE-2326-SM-9KNVU | Colon - Sigmoid | TCGA-COAD | 36.69 | 12.07 |
| GTEX-1H3VY-2526-SM-9KNW1 | Colon - Sigmoid | TCGA-COAD | 43.44 | 14.32 |
| GTEX-1HB9E-2526-SM-D4P35 | Colon - Sigmoid | TCGA-COAD | 30.48 | 8.97 |
| GTEX-1HBPH-2526-SM-ACKXK | Colon - Sigmoid | TCGA-COAD | 42.18 | 13.39 |
| GTEX-1HBPI-2226-SM-9WYTY | Colon - Sigmoid | TCGA-COAD | 35.41 | 13.81 |
| GTEX-1HBPM-2626-SM-9WYU5 | Colon - Sigmoid | TCGA-COAD | 30.76 | 10.79 |
| GTEX-1HBPN-2026-SM-B2LWA | Colon - Sigmoid | TCGA-COAD | 33.27 | 9.41 |
| GTEX-1HCU6-2926-SM-B2LWV | Colon - Sigmoid | TCGA-COAD | 35.4 | 12.62 |

| | | | | |
|---|---|---|---|---|
| GTEX-1HCU7-2726-SM-A9G3D | Colon - Sigmoid | TCGA-COAD | 45.04 | 14.45 |
| GTEX-1HCUA-1926-SM-B2LWE | Colon - Sigmoid | TCGA-COAD | 42.05 | 10.87 |
| GTEX-1HCVE-2326-SM-9WYUD | Colon - Sigmoid | TCGA-COAD | 24.97 | 11.05 |
| GTEX-1HFI7-1626-SM-9WYTR | Colon - Sigmoid | TCGA-COAD | 42.9 | 15.74 |
| GTEX-1HKZK-2026-SM-9WYUF | Colon - Sigmoid | TCGA-COAD | 30.7 | 9.77 |
| GTEX-1HSEH-1926-SM-A96SB | Colon - Sigmoid | TCGA-COAD | 30.07 | 7.09 |
| GTEX-1HSGN-2226-SM-A9SL6 | Colon - Sigmoid | TCGA-COAD | 37.48 | 12.7 |
| GTEX-1HSKV-2326-SM-A9SL9 | Colon - Sigmoid | TCGA-COAD | 31.42 | 11.95 |
| GTEX-1HSMO-2526-SM-A9SKP | Colon - Sigmoid | TCGA-COAD | 47.12 | 13.01 |
| GTEX-1HSMQ-2526-SM-A9SMK | Colon - Sigmoid | TCGA-COAD | 48.3 | 13.43 |
| GTEX-1I1CD-2826-SM-A9SL4 | Colon - Sigmoid | TCGA-COAD | 29.52 | 13.79 |
| GTEX-1I1GP-2826-SM-A9SKZ | Colon - Sigmoid | TCGA-COAD | 36.8 | 15.95 |
| GTEX-1I1GR-2226-SM-ARL7F | Colon - Sigmoid | TCGA-COAD | 30.01 | 8.01 |
| GTEX-1I1GT-2026-SM-CKZPK | Colon - Sigmoid | TCGA-COAD | 37.57 | 10.08 |
| GTEX-1I6K6-1926-SM-A96SN | Colon - Sigmoid | TCGA-COAD | 41.06 | 9.33 |
| GTEX-1ICG6-2226-SM-B2LY7 | Colon - Sigmoid | TCGA-COAD | 40.35 | 16.27 |
| GTEX-1IDJH-2826-SM-CJI49 | Colon - Sigmoid | TCGA-COAD | 26.32 | 9.52 |
| GTEX-1IDJI-2226-SM-CKZPM | Colon - Sigmoid | TCGA-COAD | 34.58 | 9.76 |
| GTEX-1IDJU-1926-SM-CL53W | Colon - Sigmoid | TCGA-COAD | 45.93 | 11.42 |
| GTEX-1IKOE-2126-SM-A96T5 | Colon - Sigmoid | TCGA-COAD | 42.52 | 9.43 |
| GTEX-1IL2U-2526-SM-C1YQV | Colon - Sigmoid | TCGA-COAD | 37.77 | 12.69 |
| GTEX-1J1OQ-2826-SM-D3L88 | Colon - Sigmoid | TCGA-COAD | 44.41 | 11.48 |
| GTEX-1J1R8-1426-SM-CL55O | Colon - Sigmoid | TCGA-COAD | 43.5 | 9.23 |
| GTEX-1J8EW-2426-SM-CE6T5 | Colon - Sigmoid | TCGA-COAD | 29.34 | 12.1 |
| GTEX-1J8JJ-2426-SM-D4P4A | Colon - Sigmoid | TCGA-COAD | 28.56 | 10.19 |
| GTEX-1J8Q3-2426-SM-CL53S | Colon - Sigmoid | TCGA-COAD | 38.81 | 13.75 |
| GTEX-1J8QM-1226-SM-CY8HI | Colon - Sigmoid | TCGA-COAD | 39.74 | 12.81 |
| GTEX-1JJE9-2526-SM-ARU8K | Colon - Sigmoid | TCGA-COAD | 29.68 | 8.15 |
| GTEX-1JJEA-2526-SM-CY8HJ | Colon - Sigmoid | TCGA-COAD | 37.75 | 16.16 |
| GTEX-1JK1U-1426-SM-CNPOW | Colon - Sigmoid | TCGA-COAD | 50.66 | 14.72 |
| GTEX-1JKYN-2226-SM-ARZNE | Colon - Sigmoid | TCGA-COAD | 39.22 | 10.12 |

| | | | | |
|---|---|---|---|---|
| GTEX-1JKYR-2026-SM-CNPON | Colon - Sigmoid | TCGA-COAD | 61.05 | 15.03 |
| GTEX-1JMLX-2526-SM-CY8HX | Colon - Sigmoid | TCGA-COAD | 35.34 | 11.23 |
| GTEX-1JMPY-1026-SM-ARZN1 | Colon - Sigmoid | TCGA-COAD | 41.28 | 8.76 |
| GTEX-1JMPZ-2626-SM-ARU8Z | Colon - Sigmoid | TCGA-COAD | 27.73 | 13.32 |
| GTEX-1JMQJ-2626-SM-CY8HQ | Colon - Sigmoid | TCGA-COAD | 37.48 | 18.38 |
| GTEX-1JMQK-2326-SM-ARZN4 | Colon - Sigmoid | TCGA-COAD | 34.99 | 11.41 |
| GTEX-1JN6P-2126-SM-ARL98 | Colon - Sigmoid | TCGA-COAD | 12.84 | 5.67 |
| GTEX-1JN76-1326-SM-ARU94 | Colon - Sigmoid | TCGA-COAD | 34.85 | 12.19 |
| GTEX-1K2DA-1626-SM-CGQGB | Colon - Sigmoid | TCGA-COAD | 37.38 | 13.48 |
| GTEX-1K9T9-2926-SM-E9TKD | Colon - Sigmoid | TCGA-COAD | 69.48 | 21.86 |
| GTEX-1KANA-1726-SM-CY8IG | Colon - Sigmoid | TCGA-COAD | 40.62 | 10.73 |
| GTEX-1KXAM-1926-SM-D3LAG | Colon - Sigmoid | TCGA-COAD | 34.85 | 13.06 |
| GTEX-1L5NE-1926-SM-D3L9R | Colon - Sigmoid | TCGA-COAD | 60.51 | 13.64 |
| GTEX-1LB8K-2526-SM-DIPEQ | Colon - Sigmoid | TCGA-COAD | 29.82 | 13.71 |
| GTEX-1LBAC-1726-SM-D3L9W | Colon - Sigmoid | TCGA-COAD | 46.48 | 14.48 |
| GTEX-1LG7Y-2226-SM-E9U5K | Colon - Sigmoid | TCGA-COAD | 43.14 | 13.26 |
| GTEX-1LG7Z-1326-SM-DIPF3 | Colon - Sigmoid | TCGA-COAD | 42.65 | 9.74 |
| GTEX-1LGOU-1726-SM-D3L8P | Colon - Sigmoid | TCGA-COAD | 41.22 | 14.65 |
| GTEX-1LGRB-2026-SM-COH47 | Colon - Sigmoid | TCGA-COAD | 49.62 | 10.16 |
| GTEX-1LKK1-2426-SM-DIPES | Colon - Sigmoid | TCGA-COAD | 26.23 | 9.77 |
| GTEX-1LSNL-1626-SM-E9TJY | Colon - Sigmoid | TCGA-COAD | 30.04 | 7.91 |
| GTEX-1LSNM-1426-SM-E6CIH | Colon - Sigmoid | TCGA-COAD | 29.04 | 7.67 |
| GTEX-1LSVX-2026-SM-E9TJD | Colon - Sigmoid | TCGA-COAD | 29.61 | 10.66 |
| GTEX-1LVAM-1726-SM-DHXKQ | Colon - Sigmoid | TCGA-COAD | 52.96 | 11.88 |
| GTEX-1LVAN-2626-SM-CNPQF | Colon - Sigmoid | TCGA-COAD | 29.7 | 10.57 |
| GTEX-1LVAO-2826-SM-E9TJH | Colon - Sigmoid | TCGA-COAD | 32.29 | 9.09 |
| GTEX-1MGNQ-2126-SM-EV7AC | Colon - Sigmoid | TCGA-COAD | 44.74 | 13.14 |
| GTEX-1N2DV-1826-SM-E6CJ5 | Colon - Sigmoid | TCGA-COAD | 28.6 | 8.3 |
| GTEX-1N2EF-1726-SM-EXUSI | Colon - Sigmoid | TCGA-COAD | 41.11 | 13.81 |
| GTEX-1OFPY-1226-SM-E76OM | Colon - Sigmoid | TCGA-COAD | 34.12 | 8.99 |
| GTEX-1OJC3-1826-SM-DTX93 | Colon - Sigmoid | TCGA-COAD | 33.31 | 5.11 |

| | | | | |
|---|---|---|---|---|
| GTEX-1P4AB-2626-SM-DTXEU | Colon - Sigmoid | TCGA-COAD | 29.16 | 7.04 |
| GTEX-1PBJI-2526-SM-E6CP4 | Colon - Sigmoid | TCGA-COAD | 40.09 | 13.5 |
| GTEX-1PBJJ-2026-SM-DTX8U | Colon - Sigmoid | TCGA-COAD | 30.47 | 11.68 |
| GTEX-1PDJ9-2026-SM-E76PT | Colon - Sigmoid | TCGA-COAD | 48.72 | 11.52 |
| GTEX-1PFEY-1726-SM-DPRZ4 | Colon - Sigmoid | TCGA-COAD | 41.44 | 10.25 |
| GTEX-1PIEJ-1826-SM-EWRMN | Colon - Sigmoid | TCGA-COAD | 59.88 | 9.88 |
| GTEX-1PIGE-1526-SM-DTXFC | Colon - Sigmoid | TCGA-COAD | 39.49 | 14.03 |
| GTEX-1PIIG-2126-SM-EXUSD | Colon - Sigmoid | TCGA-COAD | 33.84 | 12.27 |
| GTEX-1PPH8-2226-SM-EWRN7 | Colon - Sigmoid | TCGA-COAD | 40.82 | 7.7 |
| GTEX-1QCLY-2026-SM-EVR4C | Colon - Sigmoid | TCGA-COAD | 38.04 | 10.99 |
| GTEX-1QCLZ-2026-SM-EAZ4A | Colon - Sigmoid | TCGA-COAD | 41.75 | 16.93 |
| GTEX-1QP28-2226-SM-DTX8C | Colon - Sigmoid | TCGA-COAD | 37.12 | 9.58 |
| GTEX-1QP29-2526-SM-E6CQ3 | Colon - Sigmoid | TCGA-COAD | 36.32 | 14.05 |
| GTEX-1QP2A-2226-SM-DTXF6 | Colon - Sigmoid | TCGA-COAD | 49.35 | 8.67 |
| GTEX-1QP67-1426-SM-E8VOO | Colon - Sigmoid | TCGA-COAD | 42.4 | 11.83 |
| GTEX-1QP9N-0826-SM-DTX9O | Colon - Sigmoid | TCGA-COAD | 33.35 | 11.66 |
| GTEX-1QPFJ-2426-SM-EVR3L | Colon - Sigmoid | TCGA-COAD | 41.8 | 10.66 |
| GTEX-1QW4Y-1426-SM-E76P3 | Colon - Sigmoid | TCGA-COAD | 47.41 | 14.03 |
| GTEX-1R7EU-1826-SM-DTX9L | Colon - Sigmoid | TCGA-COAD | 35.23 | 11.66 |
| GTEX-1R9JW-2126-SM-E76QT | Colon - Sigmoid | TCGA-COAD | 46.94 | 13.2 |
| GTEX-1R9K5-2226-SM-DTX9G | Colon - Sigmoid | TCGA-COAD | 36.38 | 8.44 |
| GTEX-1R9PO-2026-SM-E9J4N | Colon - Sigmoid | TCGA-COAD | 37.25 | 14.55 |
| GTEX-1RAZQ-2026-SM-E76P7 | Colon - Sigmoid | TCGA-COAD | 28.3 | 8.46 |
| GTEX-1RAZR-2026-SM-E9TKE | Colon - Sigmoid | TCGA-COAD | 45.06 | 13.58 |
| GTEX-1RDX4-1926-SM-E6CJJ | Colon - Sigmoid | TCGA-COAD | 20 | 7.68 |
| GTEX-1RLM8-1726-SM-E9TKF | Colon - Sigmoid | TCGA-COAD | 45.94 | 14.8 |
| GTEX-1RMOY-1726-SM-EVYC9 | Colon - Sigmoid | TCGA-COAD | 44.44 | 11.55 |
| GTEX-1RQEC-2626-SM-EVR3Z | Colon - Sigmoid | TCGA-COAD | 41.53 | 16.7 |
| GTEX-1S5ZA-1426-SM-E6CJQ | Colon - Sigmoid | TCGA-COAD | 30.24 | 8.34 |
| GTEX-1S82P-2026-SM-E9TIQ | Colon - Sigmoid | TCGA-COAD | 50.31 | 11.77 |
| GTEX-1S831-1726-SM-EWROJ | Colon - Sigmoid | TCGA-COAD | 51.2 | 12.9 |

| | | | | |
|---|---|---|---|---|
| GTEX-V955-1726-SM-4JBHF | Colon - Sigmoid | TCGA-COAD | 29.43 | 7.75 |
| GTEX-VJYA-2126-SM-4KL1O | Colon - Sigmoid | TCGA-COAD | 31.44 | 7.32 |
| GTEX-W5WG-2026-SM-4LMIB | Colon - Sigmoid | TCGA-COAD | 35.44 | 9.54 |
| GTEX-WFG7-1626-SM-4LVMF | Colon - Sigmoid | TCGA-COAD | 51.64 | 13.28 |
| GTEX-WFG8-1726-SM-4LVM6 | Colon - Sigmoid | TCGA-COAD | 40.34 | 11.34 |
| GTEX-WFON-1326-SM-4LVMN | Colon - Sigmoid | TCGA-COAD | 31.15 | 8.13 |
| GTEX-WH7G-1426-SM-4LVMU | Colon - Sigmoid | TCGA-COAD | 26.38 | 4.78 |
| GTEX-WHPG-1626-SM-4M1ZL | Colon - Sigmoid | TCGA-COAD | 35.43 | 13.6 |
| GTEX-WHSE-2726-SM-EZ6ML | Colon - Sigmoid | TCGA-COAD | 29.57 | 6.76 |
| GTEX-WHWD-1526-SM-4OORV | Colon - Sigmoid | TCGA-COAD | 52.08 | 13.33 |
| GTEX-WI4N-2226-SM-4OOS9 | Colon - Sigmoid | TCGA-COAD | 27.19 | 7.06 |
| GTEX-WOFM-1926-SM-4OOSP | Colon - Sigmoid | TCGA-COAD | 30.42 | 5.91 |
| GTEX-WQUQ-2626-SM-4MVNP | Colon - Sigmoid | TCGA-COAD | 47.33 | 13.47 |
| GTEX-WRHK-1126-SM-4MVOJ | Colon - Sigmoid | TCGA-COAD | 36.25 | 9.35 |
| GTEX-WXYG-1726-SM-4ONCU | Colon - Sigmoid | TCGA-COAD | 32.91 | 10.22 |
| GTEX-WY7C-1926-SM-4ONCI | Colon - Sigmoid | TCGA-COAD | 43.37 | 14.98 |
| GTEX-WYJK-2326-SM-4ONDN | Colon - Sigmoid | TCGA-COAD | 33.06 | 6.04 |
| GTEX-WYVS-1126-SM-4SOJX | Colon - Sigmoid | TCGA-COAD | 42.36 | 10.57 |
| GTEX-X15G-1526-SM-4PQZM | Colon - Sigmoid | TCGA-COAD | 31.21 | 12.02 |
| GTEX-X4EO-2726-SM-4E3HS | Colon - Sigmoid | TCGA-COAD | 33.68 | 11.63 |
| GTEX-X5EB-1126-SM-46MVV | Colon - Sigmoid | TCGA-COAD | 36 | 10.29 |
| GTEX-XAJ8-0726-SM-47JY5 | Colon - Sigmoid | TCGA-COAD | 40.68 | 12.49 |
| GTEX-XBED-1726-SM-47JYO | Colon - Sigmoid | TCGA-COAD | 48.15 | 13.89 |
| GTEX-XQ8I-2326-SM-4BOQC | Colon - Sigmoid | TCGA-COAD | 48.82 | 15.28 |
| GTEX-XUW1-1826-SM-4BOQD | Colon - Sigmoid | TCGA-COAD | 37.6 | 11.09 |
| GTEX-XUZC-1526-SM-4BRV4 | Colon - Sigmoid | TCGA-COAD | 30.6 | 8.98 |
| GTEX-XV7Q-2226-SM-4BRVY | Colon - Sigmoid | TCGA-COAD | 40.16 | 11.63 |
| GTEX-XXEK-1826-SM-4BRVC | Colon - Sigmoid | TCGA-COAD | 33.07 | 11.01 |
| GTEX-Y114-1626-SM-4TT7I | Colon - Sigmoid | TCGA-COAD | 41.43 | 11.63 |
| GTEX-Y3I4-1126-SM-4TT7Y | Colon - Sigmoid | TCGA-COAD | 41.22 | 10.5 |
| GTEX-Y5LM-1426-SM-5RQJL | Colon - Sigmoid | TCGA-COAD | 36.96 | 9.72 |

| | | | | |
|---|---|---|---|---|
| GTEX-Y5V6-1626-SM-4VDT2 | Colon - Sigmoid | TCGA-COAD | 25.34 | 7.81 |
| GTEX-Y8E4-2326-SM-EAZAZ | Colon - Sigmoid | TCGA-COAD | 24.03 | 9.1 |
| GTEX-Y8LW-1126-SM-4VDS4 | Colon - Sigmoid | TCGA-COAD | 28.6 | 7.49 |
| GTEX-Y9LG-1326-SM-4VBQB | Colon - Sigmoid | TCGA-COAD | 37.12 | 13.29 |
| GTEX-YEC3-2026-SM-DKPOL | Colon - Sigmoid | TCGA-COAD | 42.97 | 10.05 |
| GTEX-YECK-2826-SM-5IFHF | Colon - Sigmoid | TCGA-COAD | 37.72 | 12.1 |
| GTEX-YF7O-1626-SM-5IFII | Colon - Sigmoid | TCGA-COAD | 37.31 | 12.67 |
| GTEX-YFCO-1526-SM-5YYB8 | Colon - Sigmoid | TCGA-COAD | 41.58 | 11.62 |
| GTEX-YJ8A-1726-SM-5P9IQ | Colon - Sigmoid | TCGA-COAD | 50.55 | 16.17 |
| GTEX-YJ8O-1426-SM-5PNV9 | Colon - Sigmoid | TCGA-COAD | 43.9 | 15.29 |
| GTEX-Z93S-2626-SM-57WBX | Colon - Sigmoid | TCGA-COAD | 25.64 | 10.39 |
| GTEX-ZA64-1426-SM-5CVMC | Colon - Sigmoid | TCGA-COAD | 48.33 | 13.8 |
| GTEX-ZAB4-2426-SM-5HL8A | Colon - Sigmoid | TCGA-COAD | 31.98 | 11.17 |
| GTEX-ZAB5-1426-SM-5HL9D | Colon - Sigmoid | TCGA-COAD | 47.52 | 15.07 |
| GTEX-ZAJG-2626-SM-5S2NU | Colon - Sigmoid | TCGA-COAD | 62.82 | 17.99 |
| GTEX-ZC5H-2426-SM-4WAZ6 | Colon - Sigmoid | TCGA-COAD | 44.8 | 14.85 |
| GTEX-ZDTS-2726-SM-5L3E2 | Colon - Sigmoid | TCGA-COAD | 44.05 | 19.35 |
| GTEX-ZE7O-2626-SM-51MSS | Colon - Sigmoid | TCGA-COAD | 34.96 | 15.12 |
| GTEX-ZE9C-2126-SM-4WKGY | Colon - Sigmoid | TCGA-COAD | 49.72 | 15.15 |
| GTEX-ZF28-2526-SM-57WFH | Colon - Sigmoid | TCGA-COAD | 53.84 | 12.67 |
| GTEX-ZF29-1826-SM-4WKG8 | Colon - Sigmoid | TCGA-COAD | 32.71 | 6.85 |
| GTEX-ZF2S-1826-SM-4WKFF | Colon - Sigmoid | TCGA-COAD | 40.16 | 15.34 |
| GTEX-ZG7Y-1826-SM-4WWD3 | Colon - Sigmoid | TCGA-COAD | 32.91 | 12.34 |
| GTEX-ZLFU-1426-SM-4WWEU | Colon - Sigmoid | TCGA-COAD | 54.4 | 10.43 |
| GTEX-ZLV1-1326-SM-DO11Q | Colon - Sigmoid | TCGA-COAD | 55.83 | 14.05 |
| GTEX-ZLWG-1826-SM-DO11I | Colon - Sigmoid | TCGA-COAD | 56.71 | 15.71 |
| GTEX-ZP4G-1526-SM-57WE5 | Colon - Sigmoid | TCGA-COAD | 36.04 | 10.71 |
| GTEX-ZPCL-1626-SM-57WGC | Colon - Sigmoid | TCGA-COAD | 45.23 | 10.89 |
| GTEX-ZPIC-2626-SM-57WE2 | Colon - Sigmoid | TCGA-COAD | 50.7 | 17.78 |
| GTEX-ZPU1-1726-SM-4WWFV | Colon - Sigmoid | TCGA-COAD | 33.41 | 9.23 |
| GTEX-ZQG8-2426-SM-57WEE | Colon - Sigmoid | TCGA-COAD | 33.51 | 10.26 |

| | | | | |
|---|---|---|---|---|
| GTEX-ZQUD-1026-SM-4YCEZ | Colon - Sigmoid | TCGA-COAD | 24.11 | 4.37 |
| GTEX-ZT9W-1726-SM-51MS3 | Colon - Sigmoid | TCGA-COAD | 41.37 | 6.72 |
| GTEX-ZTSS-1226-SM-51MSZ | Colon - Sigmoid | TCGA-COAD | 59.85 | 11.23 |
| GTEX-ZTTD-2626-SM-57WEX | Colon - Sigmoid | TCGA-COAD | 55.2 | 20.75 |
| GTEX-ZV6S-1126-SM-57WE9 | Colon - Sigmoid | TCGA-COAD | 36.29 | 10.55 |
| GTEX-ZVT2-1726-SM-51MRO | Colon - Sigmoid | TCGA-COAD | 43.28 | 9.38 |
| GTEX-ZVT3-2826-SM-5GU6A | Colon - Sigmoid | TCGA-COAD | 38.59 | 13.86 |
| GTEX-ZVT4-2626-SM-5N9GJ | Colon - Sigmoid | TCGA-COAD | 42.35 | 12.18 |
| GTEX-ZVZQ-2626-SM-59HLB | Colon - Sigmoid | TCGA-COAD | 13.19 | 6.5 |
| GTEX-ZXES-1726-SM-5E43G | Colon - Sigmoid | TCGA-COAD | 37.23 | 8.43 |
| GTEX-ZY6K-1326-SM-5GZWP | Colon - Sigmoid | TCGA-COAD | 45.98 | 11.47 |
| GTEX-ZYFD-2226-SM-5E43P | Colon - Sigmoid | TCGA-COAD | 55.01 | 12.14 |
| GTEX-ZYFG-1826-SM-5GZWX | Colon - Sigmoid | TCGA-COAD | 35.28 | 9.94 |
| GTEX-ZYT6-2826-SM-5GICX | Colon - Sigmoid | TCGA-COAD | 38.48 | 14.01 |
| GTEX-ZYY3-2226-SM-5E45A | Colon - Sigmoid | TCGA-COAD | 40.87 | 11.5 |
| GTEX-ZZ64-0826-SM-5E449 | Colon - Sigmoid | TCGA-COAD | 48.1 | 9.75 |
| GTEX-111CU-1426-SM-5GZYP | Colon - Transverse | TCGA-COAD | 44.01 | 9.6 |
| GTEX-111VG-2226-SM-5N9DU | Colon - Transverse | TCGA-COAD | 36.23 | 9.37 |
| GTEX-111YS-1626-SM-5GZZ9 | Colon - Transverse | TCGA-COAD | 48.26 | 10.4 |
| GTEX-1122O-1526-SM-5N9CL | Colon - Transverse | TCGA-COAD | 29.64 | 3.5 |
| GTEX-1128S-1626-SM-5H12O | Colon - Transverse | TCGA-COAD | 42.13 | 12.72 |
| GTEX-117YW-1826-SM-5PNY5 | Colon - Transverse | TCGA-COAD | 31.46 | 6.96 |
| GTEX-117YX-0826-SM-5H11J | Colon - Transverse | TCGA-COAD | 43.74 | 12.19 |
| GTEX-11DXX-1826-SM-5H126 | Colon - Transverse | TCGA-COAD | 34.74 | 5.85 |
| GTEX-11DXZ-1726-SM-5H12M | Colon - Transverse | TCGA-COAD | 24.97 | 5.22 |
| GTEX-11EM3-1526-SM-5A5KN | Colon - Transverse | TCGA-COAD | 40.18 | 8.92 |
| GTEX-11EQ9-1426-SM-5987G | Colon - Transverse | TCGA-COAD | 50.99 | 11.24 |
| GTEX-11GSP-1626-SM-5986N | Colon - Transverse | TCGA-COAD | 23.9 | 9.18 |
| GTEX-11I78-1226-SM-5A5K6 | Colon - Transverse | TCGA-COAD | 31.03 | 6.05 |
| GTEX-11LCK-2026-SM-5A5MA | Colon - Transverse | TCGA-COAD | 23.64 | 6.2 |
| GTEX-11NSD-1626-SM-5986U | Colon - Transverse | TCGA-COAD | 42.3 | 8.23 |

| | | | | |
|---|---|---|---|---|
| GTEX-11O72-2226-SM-5PNW2 | Colon - Transverse | TCGA-COAD | 43.72 | 13.27 |
| GTEX-11ONC-2026-SM-5HL61 | Colon - Transverse | TCGA-COAD | 37.19 | 10.16 |
| GTEX-11P7K-1526-SM-5EGI7 | Colon - Transverse | TCGA-COAD | 30.4 | 7.14 |
| GTEX-11P81-1126-SM-5GU5G | Colon - Transverse | TCGA-COAD | 45.74 | 13.09 |
| GTEX-11P82-1026-SM-5BC5J | Colon - Transverse | TCGA-COAD | 25.73 | 5.85 |
| GTEX-11TT1-1026-SM-5PNW7 | Colon - Transverse | TCGA-COAD | 79.93 | 36.38 |
| GTEX-11TTK-2426-SM-5PNYO | Colon - Transverse | TCGA-COAD | 36.03 | 12.14 |
| GTEX-11TUW-2026-SM-5EQL8 | Colon - Transverse | TCGA-COAD | 41.71 | 12.3 |
| GTEX-11VI4-1526-SM-5EQKC | Colon - Transverse | TCGA-COAD | 33.56 | 8.34 |
| GTEX-11ZVC-2126-SM-5EGJ1 | Colon - Transverse | TCGA-COAD | 50.51 | 20.34 |
| GTEX-1211K-1726-SM-5FQUJ | Colon - Transverse | TCGA-COAD | 21.5 | 4.32 |
| GTEX-12BJ1-2126-SM-5BC5O | Colon - Transverse | TCGA-COAD | 29.38 | 4.06 |
| GTEX-12C56-0626-SM-5EGGC | Colon - Transverse | TCGA-COAD | 29.82 | 7.06 |
| GTEX-12KS4-2426-SM-5LU8P | Colon - Transverse | TCGA-COAD | 36.83 | 13.58 |
| GTEX-12WSJ-1526-SM-5GCNI | Colon - Transverse | TCGA-COAD | 39.29 | 12.24 |
| GTEX-12WSL-1626-SM-5GCMU | Colon - Transverse | TCGA-COAD | 51.27 | 15.16 |
| GTEX-12ZZZ-2426-SM-5GCNN | Colon - Transverse | TCGA-COAD | 38.36 | 11.14 |
| GTEX-13111-0726-SM-5DUVE | Colon - Transverse | TCGA-COAD | 29.71 | 8.85 |
| GTEX-131XE-0826-SM-5HL9W | Colon - Transverse | TCGA-COAD | 40.99 | 11.61 |
| GTEX-131XG-1826-SM-5LZV4 | Colon - Transverse | TCGA-COAD | 38.45 | 8.73 |
| GTEX-132AR-2726-SM-5PNVN | Colon - Transverse | TCGA-COAD | 49.91 | 14.33 |
| GTEX-1339X-1626-SM-5IJDN | Colon - Transverse | TCGA-COAD | 44.9 | 11.91 |
| GTEX-133LE-1526-SM-5IFEN | Colon - Transverse | TCGA-COAD | 35.31 | 6.55 |
| GTEX-1399R-2226-SM-5P9JB | Colon - Transverse | TCGA-COAD | 46.56 | 12.1 |
| GTEX-1399S-1926-SM-5J2M5 | Colon - Transverse | TCGA-COAD | 22.75 | 4.56 |
| GTEX-1399U-1926-SM-5IFH7 | Colon - Transverse | TCGA-COAD | 29.36 | 4.47 |
| GTEX-139T6-1326-SM-5IFFO | Colon - Transverse | TCGA-COAD | 27.55 | 5.02 |
| GTEX-139TS-1926-SM-5IJFO | Colon - Transverse | TCGA-COAD | 38.54 | 9.05 |
| GTEX-139YR-2126-SM-5KM11 | Colon - Transverse | TCGA-COAD | 47.01 | 8.75 |
| GTEX-13CF3-1426-SM-5K7YY | Colon - Transverse | TCGA-COAD | 60.45 | 11.77 |
| GTEX-13D11-0826-SM-5KLZJ | Colon - Transverse | TCGA-COAD | 30.21 | 4.89 |

| | | | | |
|---|---|---|---|---|
| GTEX-13FH7-1526-SM-5J2N6 | Colon - Transverse | TCGA-COAD | 43.61 | 13.95 |
| GTEX-13FHP-2426-SM-5KLZ5 | Colon - Transverse | TCGA-COAD | 48.06 | 15.05 |
| GTEX-13FTW-1826-SM-5IJEN | Colon - Transverse | TCGA-COAD | 35.29 | 7.23 |
| GTEX-13O21-2426-SM-5MR4A | Colon - Transverse | TCGA-COAD | 26.97 | 6.18 |
| GTEX-13O3O-2526-SM-5L3DR | Colon - Transverse | TCGA-COAD | 30.97 | 9.64 |
| GTEX-13O3P-2326-SM-5KM55 | Colon - Transverse | TCGA-COAD | 38.98 | 10.15 |
| GTEX-13O61-1626-SM-5KM3T | Colon - Transverse | TCGA-COAD | 42.27 | 9.56 |
| GTEX-13OVG-1426-SM-5K7VP | Colon - Transverse | TCGA-COAD | 45.8 | 7.18 |
| GTEX-13OVI-2026-SM-5J1MU | Colon - Transverse | TCGA-COAD | 39.01 | 4.55 |
| GTEX-13OVJ-1726-SM-5IJDZ | Colon - Transverse | TCGA-COAD | 37.38 | 7.89 |
| GTEX-13OVK-0826-SM-7KULZ | Colon - Transverse | TCGA-COAD | 39.85 | 12.31 |
| GTEX-13OW5-2626-SM-5MR3M | Colon - Transverse | TCGA-COAD | 34.24 | 8.06 |
| GTEX-13OW6-2126-SM-5N9GL | Colon - Transverse | TCGA-COAD | 45.27 | 21.03 |
| GTEX-13PL7-2526-SM-5IFGR | Colon - Transverse | TCGA-COAD | 29.44 | 6.52 |
| GTEX-13PVR-1626-SM-73KVF | Colon - Transverse | TCGA-COAD | 29.85 | 5.41 |
| GTEX-13QBU-1826-SM-5IJG3 | Colon - Transverse | TCGA-COAD | 27.3 | 3.95 |
| GTEX-13QJ3-2626-SM-9OSXQ | Colon - Transverse | TCGA-COAD | 28.72 | 9.07 |
| GTEX-13S86-1626-SM-5Q5C5 | Colon - Transverse | TCGA-COAD | 53.44 | 10.83 |
| GTEX-13U4I-2426-SM-5LU34 | Colon - Transverse | TCGA-COAD | 42.25 | 13.61 |
| GTEX-13VXT-2426-SM-5LU46 | Colon - Transverse | TCGA-COAD | 47.26 | 12.87 |
| GTEX-13W3W-2126-SM-5K7UX | Colon - Transverse | TCGA-COAD | 46.48 | 15.14 |
| GTEX-13X6H-1926-SM-7EWD3 | Colon - Transverse | TCGA-COAD | 49.9 | 14.71 |
| GTEX-144GL-2526-SM-5TDDL | Colon - Transverse | TCGA-COAD | 43.93 | 9.47 |
| GTEX-144GM-1226-SM-5LU52 | Colon - Transverse | TCGA-COAD | 35.16 | 7.19 |
| GTEX-144GN-1126-SM-5LU33 | Colon - Transverse | TCGA-COAD | 47.17 | 15.27 |
| GTEX-145LS-2726-SM-5Q5D8 | Colon - Transverse | TCGA-COAD | 56.21 | 18.28 |
| GTEX-145LT-0926-SM-5LU9C | Colon - Transverse | TCGA-COAD | 33.98 | 10.19 |
| GTEX-145ME-0826-SM-5O9AF | Colon - Transverse | TCGA-COAD | 46.47 | 8.36 |
| GTEX-145MN-1726-SM-5SIA6 | Colon - Transverse | TCGA-COAD | 33.54 | 5.17 |
| GTEX-145MO-2426-SM-5NQ9W | Colon - Transverse | TCGA-COAD | 34.77 | 8.26 |
| GTEX-146FQ-1926-SM-5NQBY | Colon - Transverse | TCGA-COAD | 60.69 | 13.59 |

| | | | | |
|---|---|---|---|---|
| GTEX-147F3-1926-SM-5SIB9 | Colon - Transverse | TCGA-COAD | 43.7 | 9.88 |
| GTEX-147GR-2726-SM-664OO | Colon - Transverse | TCGA-COAD | 33.88 | 11.08 |
| GTEX-147JS-2526-SM-5S2Q8 | Colon - Transverse | TCGA-COAD | 33.35 | 9.85 |
| GTEX-148VJ-2626-SM-5QGPI | Colon - Transverse | TCGA-COAD | 33.65 | 12 |
| GTEX-14AS3-1126-SM-5Q5CR | Colon - Transverse | TCGA-COAD | 54.98 | 12.63 |
| GTEX-14BIN-3026-SM-5ZZUS | Colon - Transverse | TCGA-COAD | 37.76 | 12.79 |
| GTEX-14BMU-1526-SM-5TDE6 | Colon - Transverse | TCGA-COAD | 41.17 | 8.52 |
| GTEX-14BMV-2426-SM-5S2PP | Colon - Transverse | TCGA-COAD | 41.83 | 12.42 |
| GTEX-14C39-1926-SM-5ZZW5 | Colon - Transverse | TCGA-COAD | 39.03 | 9.5 |
| GTEX-14C5O-2526-SM-73KU2 | Colon - Transverse | TCGA-COAD | 22.28 | 6.25 |
| GTEX-14DAQ-2726-SM-5S2QY | Colon - Transverse | TCGA-COAD | 40.45 | 13.02 |
| GTEX-14DAR-1526-SM-5RQIX | Colon - Transverse | TCGA-COAD | 22 | 5.5 |
| GTEX-14E1K-1826-SM-793DT | Colon - Transverse | TCGA-COAD | 42.72 | 13.33 |
| GTEX-14E6E-1526-SM-5RQIG | Colon - Transverse | TCGA-COAD | 18.92 | 3.73 |
| GTEX-14H4A-1926-SM-5ZZWN | Colon - Transverse | TCGA-COAD | 45.48 | 12.78 |
| GTEX-14ICK-1026-SM-6AJA4 | Colon - Transverse | TCGA-COAD | 38.16 | 6.33 |
| GTEX-14ICL-1226-SM-5S2RD | Colon - Transverse | TCGA-COAD | 47.19 | 8.16 |
| GTEX-14JG6-1726-SM-69LPL | Colon - Transverse | TCGA-COAD | 34.55 | 9.43 |
| GTEX-14JIY-3326-SM-6871E | Colon - Transverse | TCGA-COAD | 32.04 | 4.11 |
| GTEX-14PHX-1526-SM-68717 | Colon - Transverse | TCGA-COAD | 26.29 | 4.2 |
| GTEX-14PJ3-0926-SM-69LQG | Colon - Transverse | TCGA-COAD | 36.29 | 11.4 |
| GTEX-14PJ4-1926-SM-6LLHG | Colon - Transverse | TCGA-COAD | 32.72 | 5.9 |
| GTEX-14PJ5-0926-SM-6LLIK | Colon - Transverse | TCGA-COAD | 52.82 | 12.41 |
| GTEX-14PJM-2626-SM-686ZD | Colon - Transverse | TCGA-COAD | 47.4 | 15.39 |
| GTEX-14PK6-1726-SM-6871J | Colon - Transverse | TCGA-COAD | 23.3 | 4.6 |
| GTEX-14PKV-2126-SM-664O3 | Colon - Transverse | TCGA-COAD | 37.23 | 6.79 |
| GTEX-14XAO-2026-SM-69LOI | Colon - Transverse | TCGA-COAD | 31.12 | 8.53 |
| GTEX-15CHC-1626-SM-5ZZWS | Colon - Transverse | TCGA-COAD | 50.78 | 14.1 |
| GTEX-15CHQ-2526-SM-6EU24 | Colon - Transverse | TCGA-COAD | 43.79 | 15.9 |
| GTEX-15CHR-1426-SM-7DUGV | Colon - Transverse | TCGA-COAD | 34.56 | 7.75 |
| GTEX-15DCE-2026-SM-7KUFE | Colon - Transverse | TCGA-COAD | 46.03 | 10.18 |

| | | | | |
|---|---|---|---|---|
| GTEX-15DYW-2326-SM-6PAMU | Colon - Transverse | TCGA-COAD | 37.89 | 10.91 |
| GTEX-15DZA-0926-SM-69LQ3 | Colon - Transverse | TCGA-COAD | 46.01 | 10.27 |
| GTEX-15ER7-2926-SM-7KUFK | Colon - Transverse | TCGA-COAD | 45.94 | 14.42 |
| GTEX-15G1A-1826-SM-7KUKO | Colon - Transverse | TCGA-COAD | 44.89 | 11.41 |
| GTEX-15RIE-1826-SM-6M48P | Colon - Transverse | TCGA-COAD | 50.76 | 10.98 |
| GTEX-15RIF-1326-SM-6LPKO | Colon - Transverse | TCGA-COAD | 32.33 | 6.24 |
| GTEX-15SB6-1226-SM-7KUMP | Colon - Transverse | TCGA-COAD | 40.22 | 10.45 |
| GTEX-15SHU-2526-SM-6M47D | Colon - Transverse | TCGA-COAD | 33.52 | 11.46 |
| GTEX-15UF6-2626-SM-6M466 | Colon - Transverse | TCGA-COAD | 46.53 | 13.97 |
| GTEX-16AAH-1326-SM-7KFRG | Colon - Transverse | TCGA-COAD | 35.51 | 9.17 |
| GTEX-16MT8-2726-SM-7IGLJ | Colon - Transverse | TCGA-COAD | 24.53 | 7.67 |
| GTEX-16NGA-1026-SM-7KFS3 | Colon - Transverse | TCGA-COAD | 47.32 | 17.45 |
| GTEX-16NPV-2626-SM-6M484 | Colon - Transverse | TCGA-COAD | 38.88 | 13.01 |
| GTEX-16XZZ-2626-SM-6M47S | Colon - Transverse | TCGA-COAD | 29.77 | 8.51 |
| GTEX-16Z82-2226-SM-6LPJ7 | Colon - Transverse | TCGA-COAD | 31.8 | 10.12 |
| GTEX-178AV-1326-SM-6LPJX | Colon - Transverse | TCGA-COAD | 35.71 | 8.96 |
| GTEX-17EUY-1826-SM-7DUEZ | Colon - Transverse | TCGA-COAD | 33.04 | 8.31 |
| GTEX-17EVQ-1826-SM-7IGPH | Colon - Transverse | TCGA-COAD | 45.31 | 15.21 |
| GTEX-17F96-2026-SM-79ON4 | Colon - Transverse | TCGA-COAD | 35.53 | 11.5 |
| GTEX-17F98-1026-SM-7IGO8 | Colon - Transverse | TCGA-COAD | 42.48 | 13.54 |
| GTEX-17F9E-1826-SM-7IGMC | Colon - Transverse | TCGA-COAD | 35.7 | 10.95 |
| GTEX-17GQL-1526-SM-7LTAC | Colon - Transverse | TCGA-COAD | 27.92 | 6.35 |
| GTEX-17HGU-2426-SM-7EWDP | Colon - Transverse | TCGA-COAD | 35.08 | 9.28 |
| GTEX-17HHE-1826-SM-79OKE | Colon - Transverse | TCGA-COAD | 32.32 | 11.73 |
| GTEX-17HII-2726-SM-7MGVM | Colon - Transverse | TCGA-COAD | 27.4 | 7.96 |
| GTEX-17JCI-2626-SM-7EWE6 | Colon - Transverse | TCGA-COAD | 36.12 | 9.84 |
| GTEX-17KNJ-1726-SM-7LG6A | Colon - Transverse | TCGA-COAD | 38.06 | 8.2 |
| GTEX-17MF6-2426-SM-7LTAM | Colon - Transverse | TCGA-COAD | 40.28 | 10.25 |
| GTEX-18A66-2426-SM-7KFSQ | Colon - Transverse | TCGA-COAD | 55.11 | 16.54 |
| GTEX-18A7A-2226-SM-7LT8K | Colon - Transverse | TCGA-COAD | 42.03 | 12.74 |
| GTEX-18A7B-2026-SM-72D71 | Colon - Transverse | TCGA-COAD | 37.1 | 11.29 |

| | | | | |
|---|---|---|---|---|
| GTEX-18D9A-1126-SM-7KFRJ | Colon - Transverse | TCGA-COAD | 42.83 | 8.17 |
| GTEX-18D9B-2526-SM-718BN | Colon - Transverse | TCGA-COAD | 34.59 | 14.35 |
| GTEX-18D9U-1926-SM-7KFTM | Colon - Transverse | TCGA-COAD | 44.54 | 14.45 |
| GTEX-18QFQ-1526-SM-72D57 | Colon - Transverse | TCGA-COAD | 44.42 | 8.92 |
| GTEX-1A3MV-1126-SM-718BS | Colon - Transverse | TCGA-COAD | 40.51 | 12.22 |
| GTEX-1AMEY-1926-SM-73KVT | Colon - Transverse | TCGA-COAD | 29.77 | 6.15 |
| GTEX-1AMFI-1726-SM-7189L | Colon - Transverse | TCGA-COAD | 28.3 | 4.76 |
| GTEX-1AX8Z-2726-SM-73KUC | Colon - Transverse | TCGA-COAD | 51.94 | 19.43 |
| GTEX-1AX9I-2626-SM-73KU5 | Colon - Transverse | TCGA-COAD | 41.52 | 13.54 |
| GTEX-1AYCT-1826-SM-731EG | Colon - Transverse | TCGA-COAD | 38.39 | 9.99 |
| GTEX-1AYD5-1726-SM-7RHFW | Colon - Transverse | TCGA-COAD | 29.51 | 11.42 |
| GTEX-1B8KE-1926-SM-731EE | Colon - Transverse | TCGA-COAD | 29.44 | 5.72 |
| GTEX-1B8KZ-1326-SM-7DUG6 | Colon - Transverse | TCGA-COAD | 40.03 | 10.41 |
| GTEX-1B932-2826-SM-73KUD | Colon - Transverse | TCGA-COAD | 33.2 | 13.19 |
| GTEX-1B996-2626-SM-73KZ1 | Colon - Transverse | TCGA-COAD | 34.84 | 12.77 |
| GTEX-1BAJH-2326-SM-7IGMO | Colon - Transverse | TCGA-COAD | 37.45 | 11.44 |
| GTEX-1C2JI-1626-SM-7EPHC | Colon - Transverse | TCGA-COAD | 32.53 | 8.2 |
| GTEX-1C475-1626-SM-731FP | Colon - Transverse | TCGA-COAD | 35.73 | 7.98 |
| GTEX-1C4CL-2126-SM-7IGQC | Colon - Transverse | TCGA-COAD | 48.07 | 14.45 |
| GTEX-1C64O-2326-SM-73KU8 | Colon - Transverse | TCGA-COAD | 42.24 | 14.12 |
| GTEX-1C6VQ-2726-SM-73KUL | Colon - Transverse | TCGA-COAD | 37.96 | 17.07 |
| GTEX-1CAMR-0926-SM-7MXTD | Colon - Transverse | TCGA-COAD | 35.64 | 4.63 |
| GTEX-1CAMS-2526-SM-7EPH7 | Colon - Transverse | TCGA-COAD | 34.96 | 10.95 |
| GTEX-1CB4F-2026-SM-79OLU | Colon - Transverse | TCGA-COAD | 39.2 | 17.07 |
| GTEX-1CB4I-2226-SM-7P8SL | Colon - Transverse | TCGA-COAD | 45.77 | 12.19 |
| GTEX-1E2YA-2626-SM-7MGWW | Colon - Transverse | TCGA-COAD | 27.29 | 9.13 |
| GTEX-1EH9U-2026-SM-7PBYS | Colon - Transverse | TCGA-COAD | 42.04 | 13.12 |
| GTEX-1EMGI-2726-SM-7P8S4 | Colon - Transverse | TCGA-COAD | 34.92 | 12.41 |
| GTEX-1F5PK-1926-SM-7MXT7 | Colon - Transverse | TCGA-COAD | 31.58 | 5.9 |
| GTEX-1F5PL-2126-SM-7PC2O | Colon - Transverse | TCGA-COAD | 37.71 | 7.44 |
| GTEX-1F75B-2726-SM-7P8QO | Colon - Transverse | TCGA-COAD | 51.74 | 10.88 |

| | | | | |
|---|---|---|---|---|
| GTEX-1F75I-2426-SM-9WPOY | Colon - Transverse | TCGA-COAD | 49.99 | 12.82 |
| GTEX-1GF9U-1526-SM-7MXUW | Colon - Transverse | TCGA-COAD | 30.32 | 7.49 |
| GTEX-1GF9X-1526-SM-7P8QP | Colon - Transverse | TCGA-COAD | 49.69 | 9.99 |
| GTEX-1GL5R-2426-SM-7MXV1 | Colon - Transverse | TCGA-COAD | 45.61 | 9.18 |
| GTEX-1GMR2-1726-SM-9JGI2 | Colon - Transverse | TCGA-COAD | 31.71 | 11.07 |
| GTEX-1GMR3-2826-SM-9WPOG | Colon - Transverse | TCGA-COAD | 26.07 | 8.07 |
| GTEX-1GN73-2626-SM-9KNUQ | Colon - Transverse | TCGA-COAD | 46.34 | 15.19 |
| GTEX-1GPI7-1726-SM-7P8R1 | Colon - Transverse | TCGA-COAD | 36.57 | 9.23 |
| GTEX-1GZ4H-1226-SM-9KNUA | Colon - Transverse | TCGA-COAD | 51.13 | 10.51 |
| GTEX-1GZ4I-2426-SM-9WPPP | Colon - Transverse | TCGA-COAD | 35.68 | 16.01 |
| GTEX-1H11D-2626-SM-9JGH4 | Colon - Transverse | TCGA-COAD | 35.6 | 7.58 |
| GTEX-1H1DE-1526-SM-ARL9B | Colon - Transverse | TCGA-COAD | 37.35 | 9.5 |
| GTEX-1H1DG-2226-SM-9WPPC | Colon - Transverse | TCGA-COAD | 39.8 | 16.66 |
| GTEX-1H1E6-1926-SM-9OSWD | Colon - Transverse | TCGA-COAD | 31.05 | 5.26 |
| GTEX-1H3VY-2426-SM-9WPPD | Colon - Transverse | TCGA-COAD | 42.25 | 12.98 |
| GTEX-1HCU6-2726-SM-A9SKV | Colon - Transverse | TCGA-COAD | 39.48 | 14.68 |
| GTEX-1HCU7-2626-SM-C1YQO | Colon - Transverse | TCGA-COAD | 45.11 | 15.77 |
| GTEX-1HCU9-1326-SM-B2LXT | Colon - Transverse | TCGA-COAD | 27.39 | 5.2 |
| GTEX-1HCVE-2526-SM-A9G2G | Colon - Transverse | TCGA-COAD | 29.85 | 11.89 |
| GTEX-1HFI7-1726-SM-A9G22 | Colon - Transverse | TCGA-COAD | 30.5 | 11.63 |
| GTEX-1HKZK-2126-SM-A9SL3 | Colon - Transverse | TCGA-COAD | 38.73 | 11.16 |
| GTEX-1HSEH-1726-SM-B2LXH | Colon - Transverse | TCGA-COAD | 27.42 | 6.4 |
| GTEX-1HSGN-2626-SM-CE6S1 | Colon - Transverse | TCGA-COAD | 30.51 | 9.04 |
| GTEX-1HSMP-2326-SM-D3L8R | Colon - Transverse | TCGA-COAD | 51.98 | 19.41 |
| GTEX-1HSMQ-2726-SM-C1YRE | Colon - Transverse | TCGA-COAD | 34.79 | 7.47 |
| GTEX-1I1GR-2526-SM-CNNQF | Colon - Transverse | TCGA-COAD | 34.16 | 10.34 |
| GTEX-1I1GT-1926-SM-ARU7D | Colon - Transverse | TCGA-COAD | 36.63 | 6.87 |
| GTEX-1I4MK-1126-SM-B2LWS | Colon - Transverse | TCGA-COAD | 36.99 | 8.23 |
| GTEX-1I6K6-1826-SM-B2LX5 | Colon - Transverse | TCGA-COAD | 28.48 | 8.49 |
| GTEX-1ICLZ-2426-SM-CKZOD | Colon - Transverse | TCGA-COAD | 54.16 | 22.53 |
| GTEX-1IDJC-1326-SM-CL53H | Colon - Transverse | TCGA-COAD | 48.86 | 12.46 |

| | | | | |
|---|---|---|---|---|
| GTEX-1IDJD-2226-SM-A96TD | Colon - Transverse | TCGA-COAD | 40.98 | 11.98 |
| GTEX-1IDJF-1826-SM-CKZOE | Colon - Transverse | TCGA-COAD | 43.19 | 14.66 |
| GTEX-1IDJU-2026-SM-A9SLM | Colon - Transverse | TCGA-COAD | 29.9 | 6.61 |
| GTEX-1IGQW-2326-SM-CE6RV | Colon - Transverse | TCGA-COAD | 37.51 | 17.56 |
| GTEX-1IKK5-2726-SM-CNPQ6 | Colon - Transverse | TCGA-COAD | 33.19 | 9.77 |
| GTEX-1IKOE-2026-SM-AHZ31 | Colon - Transverse | TCGA-COAD | 54.32 | 18.62 |
| GTEX-1IOXB-2626-SM-CJI2N | Colon - Transverse | TCGA-COAD | 21.68 | 3.47 |
| GTEX-1J1OQ-2726-SM-AHZ3R | Colon - Transverse | TCGA-COAD | 39.77 | 14.28 |
| GTEX-1J1R8-1126-SM-CNPQ3 | Colon - Transverse | TCGA-COAD | 26.93 | 5.32 |
| GTEX-1JJ6O-2426-SM-ARU8J | Colon - Transverse | TCGA-COAD | 30.63 | 7.58 |
| GTEX-1JJE9-2426-SM-D3L8T | Colon - Transverse | TCGA-COAD | 40.58 | 12.65 |
| GTEX-1JK1U-1526-SM-AHZ46 | Colon - Transverse | TCGA-COAD | 28.86 | 8.33 |
| GTEX-1JKYN-2326-SM-CGQG7 | Colon - Transverse | TCGA-COAD | 24.72 | 4.11 |
| GTEX-1JKYR-1926-SM-C1YR7 | Colon - Transverse | TCGA-COAD | 44.72 | 10.34 |
| GTEX-1JMLX-2726-SM-CM2SB | Colon - Transverse | TCGA-COAD | 35.94 | 11.64 |
| GTEX-1JMPY-0426-SM-C1YQ3 | Colon - Transverse | TCGA-COAD | 31.66 | 6.94 |
| GTEX-1JMPZ-2426-SM-C1YQP | Colon - Transverse | TCGA-COAD | 39.39 | 13.15 |
| GTEX-1JN1M-2626-SM-CY8I3 | Colon - Transverse | TCGA-COAD | 38.65 | 12.95 |
| GTEX-1JN76-1226-SM-C1YRM | Colon - Transverse | TCGA-COAD | 41.43 | 10.15 |
| GTEX-1K2DA-2026-SM-CGQGK | Colon - Transverse | TCGA-COAD | 26.62 | 5.26 |
| GTEX-1K9T9-2826-SM-D4P3F | Colon - Transverse | TCGA-COAD | 45.98 | 16.37 |
| GTEX-1KANA-1926-SM-DHXKF | Colon - Transverse | TCGA-COAD | 29.33 | 4.18 |
| GTEX-1KANB-1526-SM-CXZK4 | Colon - Transverse | TCGA-COAD | 36.9 | 16.33 |
| GTEX-1KANC-0926-SM-D3LA9 | Colon - Transverse | TCGA-COAD | 24.71 | 5.35 |
| GTEX-1KXAM-2126-SM-CY8I9 | Colon - Transverse | TCGA-COAD | 33.75 | 10.78 |
| GTEX-1L5NE-2026-SM-CXZKR | Colon - Transverse | TCGA-COAD | 16.91 | 5.57 |
| GTEX-1LBAC-1626-SM-CXZKV | Colon - Transverse | TCGA-COAD | 45.59 | 12.61 |
| GTEX-1LG7Y-2126-SM-DIPER | Colon - Transverse | TCGA-COAD | 45.01 | 10.45 |
| GTEX-1LG7Z-1426-SM-CXKYV | Colon - Transverse | TCGA-COAD | 30.75 | 4.96 |
| GTEX-1LGOU-1626-SM-D3LAK | Colon - Transverse | TCGA-COAD | 38.54 | 10.51 |
| GTEX-1LGRB-1426-SM-CNNPT | Colon - Transverse | TCGA-COAD | 30.34 | 4.77 |

| | | | | |
|---|---|---|---|---|
| GTEX-1LH75-2126-SM-EVR5F | Colon - Transverse | TCGA-COAD | 42.14 | 10.46 |
| GTEX-1LSVX-1726-SM-DHXJP | Colon - Transverse | TCGA-COAD | 45.58 | 13.77 |
| GTEX-1LVAM-1626-SM-EV79Y | Colon - Transverse | TCGA-COAD | 28.4 | 4.69 |
| GTEX-1LVAN-2526-SM-CNPPC | Colon - Transverse | TCGA-COAD | 27.96 | 12.67 |
| GTEX-1MA7W-0626-SM-EV79Z | Colon - Transverse | TCGA-COAD | 38.89 | 7.02 |
| GTEX-1MCC2-1626-SM-DIPEW | Colon - Transverse | TCGA-COAD | 44.58 | 11.51 |
| GTEX-1MGNQ-2026-SM-EAZ51 | Colon - Transverse | TCGA-COAD | 32.42 | 7.76 |
| GTEX-1MJK3-0926-SM-EV7AY | Colon - Transverse | TCGA-COAD | 44.56 | 9.72 |
| GTEX-1MUQO-2626-SM-EWRO3 | Colon - Transverse | TCGA-COAD | 49.74 | 12.51 |
| GTEX-1N2DV-1426-SM-DPRYQ | Colon - Transverse | TCGA-COAD | 29.05 | 3.93 |
| GTEX-1N2EF-1926-SM-DTX8M | Colon - Transverse | TCGA-COAD | 73.07 | 13.57 |
| GTEX-1N5O9-2726-SM-E76QH | Colon - Transverse | TCGA-COAD | 41.36 | 15.01 |
| GTEX-1NHNU-2326-SM-E6CQE | Colon - Transverse | TCGA-COAD | 40.82 | 15.04 |
| GTEX-1NSGN-2326-SM-E9U5X | Colon - Transverse | TCGA-COAD | 31.66 | 10.79 |
| GTEX-1NV5F-2226-SM-DTX8O | Colon - Transverse | TCGA-COAD | 36.89 | 13.88 |
| GTEX-1OKEX-1226-SM-DTXEC | Colon - Transverse | TCGA-COAD | 51.18 | 16.66 |
| GTEX-1PFEY-1426-SM-E9U6U | Colon - Transverse | TCGA-COAD | 34.29 | 11.35 |
| GTEX-1PIEJ-2026-SM-E6CPA | Colon - Transverse | TCGA-COAD | 49.98 | 12.39 |
| GTEX-1PIGE-1426-SM-E9U61 | Colon - Transverse | TCGA-COAD | 47.17 | 11.82 |
| GTEX-1POEN-2026-SM-E6CRL | Colon - Transverse | TCGA-COAD | 35.63 | 13.86 |
| GTEX-1PPGY-2826-SM-DTX9H | Colon - Transverse | TCGA-COAD | 33.98 | 14.2 |
| GTEX-1PPH8-2026-SM-DTX7P | Colon - Transverse | TCGA-COAD | 27.43 | 6.5 |
| GTEX-1PWST-2426-SM-EWRMP | Colon - Transverse | TCGA-COAD | 45.74 | 12.47 |
| GTEX-1QCLY-1726-SM-DPRXY | Colon - Transverse | TCGA-COAD | 33.48 | 9.31 |
| GTEX-1QCLZ-1926-SM-E6CR3 | Colon - Transverse | TCGA-COAD | 34.71 | 9.31 |
| GTEX-1QMI2-2626-SM-E6CR4 | Colon - Transverse | TCGA-COAD | 38.86 | 11.07 |
| GTEX-1QP67-1326-SM-EAZ4D | Colon - Transverse | TCGA-COAD | 29.54 | 7.11 |
| GTEX-1QP9N-1026-SM-E6CPX | Colon - Transverse | TCGA-COAD | 27.46 | 7.98 |
| GTEX-1QPFJ-2326-SM-E6CR7 | Colon - Transverse | TCGA-COAD | 46.76 | 9.3 |
| GTEX-1QW4Y-1326-SM-EWRMZ | Colon - Transverse | TCGA-COAD | 47.8 | 15.86 |
| GTEX-1R7EU-1926-SM-E9U62 | Colon - Transverse | TCGA-COAD | 32.76 | 6.93 |

| | | | | |
|---|---|---|---|---|
| GTEX-1R9JW-2026-SM-EAZ4O | Colon - Transverse | TCGA-COAD | 36.33 | 10.75 |
| GTEX-1R9K5-2126-SM-E8VN1 | Colon - Transverse | TCGA-COAD | 48.74 | 10.55 |
| GTEX-1R9PM-1526-SM-DPRY8 | Colon - Transverse | TCGA-COAD | 42.28 | 10.71 |
| GTEX-1RAZA-1726-SM-DPS13 | Colon - Transverse | TCGA-COAD | 31.84 | 4.9 |
| GTEX-1RAZQ-2126-SM-DTX9V | Colon - Transverse | TCGA-COAD | 43.59 | 9.25 |
| GTEX-1RAZR-1926-SM-E6CRF | Colon - Transverse | TCGA-COAD | 41.21 | 10.38 |
| GTEX-1RDX4-1826-SM-EVR54 | Colon - Transverse | TCGA-COAD | 32.83 | 9.64 |
| GTEX-1RMOY-1626-SM-EVYC8 | Colon - Transverse | TCGA-COAD | 37.44 | 9.78 |
| GTEX-1S3DN-1826-SM-EV79K | Colon - Transverse | TCGA-COAD | 34.8 | 7.55 |
| GTEX-1S5VW-2926-SM-E9U6H | Colon - Transverse | TCGA-COAD | 47.66 | 10.9 |
| GTEX-1S82U-0726-SM-E9TKI | Colon - Transverse | TCGA-COAD | 48.23 | 8.31 |
| GTEX-1S83E-1626-SM-EV7B9 | Colon - Transverse | TCGA-COAD | 36.17 | 7.78 |
| GTEX-NFK9-2026-SM-3LK5K | Colon - Transverse | TCGA-COAD | 19.09 | 5.62 |
| GTEX-O5YT-1426-SM-3MJHC | Colon - Transverse | TCGA-COAD | 47.62 | 25.1 |
| GTEX-O5YW-1426-SM-3MJHF | Colon - Transverse | TCGA-COAD | 37.99 | 8.78 |
| GTEX-OHPM-1426-SM-3TW8Y | Colon - Transverse | TCGA-COAD | 32.74 | 6.36 |
| GTEX-OHPN-2026-SM-E9TIA | Colon - Transverse | TCGA-COAD | 34.41 | 10.53 |
| GTEX-OIZF-1426-SM-2HML2 | Colon - Transverse | TCGA-COAD | 38.44 | 9.5 |
| GTEX-OIZH-1426-SM-3NB1O | Colon - Transverse | TCGA-COAD | 34.3 | 9.08 |
| GTEX-OOBJ-1426-SM-E6CHH | Colon - Transverse | TCGA-COAD | 31.1 | 11.79 |
| GTEX-OXRK-1726-SM-3NB16 | Colon - Transverse | TCGA-COAD | 39.51 | 11.78 |
| GTEX-OXRL-1426-SM-3NM9E | Colon - Transverse | TCGA-COAD | 43.07 | 13.25 |
| GTEX-OXRN-2026-SM-E6CHK | Colon - Transverse | TCGA-COAD | 27.2 | 10.06 |
| GTEX-OXRO-0726-SM-E9U4C | Colon - Transverse | TCGA-COAD | 50.91 | 11.08 |
| GTEX-P4PP-1426-SM-3NM9L | Colon - Transverse | TCGA-COAD | 32.74 | 7.6 |
| GTEX-P4QT-1426-SM-3NMCX | Colon - Transverse | TCGA-COAD | 30.73 | 6.17 |
| GTEX-P78B-1726-SM-3P5ZV | Colon - Transverse | TCGA-COAD | 42.56 | 15.39 |
| GTEX-PLZ5-1126-SM-3P613 | Colon - Transverse | TCGA-COAD | 30.75 | 7.41 |
| GTEX-PLZ6-0926-SM-3P5ZQ | Colon - Transverse | TCGA-COAD | 34.57 | 8.27 |
| GTEX-PSDG-1826-SM-EAZ2S | Colon - Transverse | TCGA-COAD | 31.37 | 9.27 |
| GTEX-PWCY-1026-SM-48TD4 | Colon - Transverse | TCGA-COAD | 28.16 | 6.85 |

| | | | | |
|---|---|---|---|---|
| GTEX-PWN1-1426-SM-48TDF | Colon - Transverse | TCGA-COAD | 26.96 | 6.22 |
| GTEX-PX3G-1426-SM-48U1J | Colon - Transverse | TCGA-COAD | 35.85 | 8.45 |
| GTEX-Q2AH-1226-SM-48TZL | Colon - Transverse | TCGA-COAD | 38.15 | 11.31 |
| GTEX-Q2AI-0926-SM-48U1F | Colon - Transverse | TCGA-COAD | 39.83 | 11.67 |
| GTEX-Q734-1126-SM-48TZY | Colon - Transverse | TCGA-COAD | 40.99 | 10.03 |
| GTEX-QCQG-1626-SM-48U26 | Colon - Transverse | TCGA-COAD | 32.13 | 8.5 |
| GTEX-QDVJ-1326-SM-48U1X | Colon - Transverse | TCGA-COAD | 37.22 | 13.01 |
| GTEX-QDVN-1326-SM-48TZ3 | Colon - Transverse | TCGA-COAD | 49.34 | 13.1 |
| GTEX-QEL4-1926-SM-D3L8D | Colon - Transverse | TCGA-COAD | 45.87 | 11.55 |
| GTEX-QLQ7-1126-SM-CYPRB | Colon - Transverse | TCGA-COAD | 41.12 | 15.81 |
| GTEX-QLQW-0426-SM-447A7 | Colon - Transverse | TCGA-COAD | 33.15 | 7.27 |
| GTEX-QMRM-1226-SM-447C6 | Colon - Transverse | TCGA-COAD | 48.93 | 13.9 |
| GTEX-QV31-0826-SM-CKZNC | Colon - Transverse | TCGA-COAD | 37.33 | 11.12 |
| GTEX-R45C-2126-SM-EZ6KZ | Colon - Transverse | TCGA-COAD | 52.44 | 18.85 |
| GTEX-R53T-1326-SM-48FCQ | Colon - Transverse | TCGA-COAD | 48 | 11.22 |
| GTEX-R55C-1126-SM-48FCJ | Colon - Transverse | TCGA-COAD | 21.98 | 5.02 |
| GTEX-R55D-1826-SM-48FEF | Colon - Transverse | TCGA-COAD | 32.64 | 7.95 |
| GTEX-R55G-1226-SM-48FDC | Colon - Transverse | TCGA-COAD | 35.04 | 6.92 |
| GTEX-RM2N-0926-SM-48FD1 | Colon - Transverse | TCGA-COAD | 29.1 | 4.98 |
| GTEX-RU1J-1326-SM-46MUL | Colon - Transverse | TCGA-COAD | 30.65 | 7.47 |
| GTEX-RWS6-1826-SM-47JXX | Colon - Transverse | TCGA-COAD | 39.39 | 13.85 |
| GTEX-S341-1426-SM-4AD6U | Colon - Transverse | TCGA-COAD | 37.83 | 10.88 |
| GTEX-S3XE-1126-SM-4AD4N | Colon - Transverse | TCGA-COAD | 40.7 | 9.36 |
| GTEX-S4P3-1226-SM-4AD4Y | Colon - Transverse | TCGA-COAD | 25.11 | 8.31 |
| GTEX-S4Q7-0826-SM-4AD5E | Colon - Transverse | TCGA-COAD | 30.65 | 7.46 |
| GTEX-S4UY-0826-SM-4AD4Z | Colon - Transverse | TCGA-COAD | 25.31 | 5.4 |
| GTEX-S7SF-1926-SM-4AT5B | Colon - Transverse | TCGA-COAD | 33.69 | 7.36 |
| GTEX-SE5C-1526-SM-4BRWU | Colon - Transverse | TCGA-COAD | 28.16 | 8.66 |
| GTEX-SJXC-2226-SM-EVYAK | Colon - Transverse | TCGA-COAD | 45.49 | 16.56 |
| GTEX-SNMC-1126-SM-4DM5M | Colon - Transverse | TCGA-COAD | 35.57 | 11.16 |
| GTEX-SNOS-1226-SM-4DM5H | Colon - Transverse | TCGA-COAD | 35.37 | 7.29 |

| | | | | |
|---|---|---|---|---|
| GTEX-T5JW-1126-SM-4DM5V | Colon - Transverse | TCGA-COAD | 36.95 | 10.75 |
| GTEX-T6MO-0826-SM-4DM51 | Colon - Transverse | TCGA-COAD | 40.01 | 9.56 |
| GTEX-TKQ1-0626-SM-4DXTS | Colon - Transverse | TCGA-COAD | 39.49 | 13.85 |
| GTEX-TKQ2-1326-SM-4DXT9 | Colon - Transverse | TCGA-COAD | 26.2 | 9.7 |
| GTEX-TML8-1326-SM-4DXTO | Colon - Transverse | TCGA-COAD | 34.98 | 11.95 |
| GTEX-U3ZH-1326-SM-4DXSF | Colon - Transverse | TCGA-COAD | 36.45 | 11.42 |
| GTEX-U3ZM-1126-SM-4DXUB | Colon - Transverse | TCGA-COAD | 24.49 | 6.16 |
| GTEX-U3ZN-2126-SM-4DXU1 | Colon - Transverse | TCGA-COAD | 34.8 | 9.52 |
| GTEX-U412-1726-SM-EZ6KX | Colon - Transverse | TCGA-COAD | 41.69 | 16.41 |
| GTEX-U4B1-1126-SM-4DXV3 | Colon - Transverse | TCGA-COAD | 41.07 | 10.59 |
| GTEX-UJHI-1126-SM-4IHLN | Colon - Transverse | TCGA-COAD | 31.87 | 8.2 |
| GTEX-UJMC-1326-SM-4IHLS | Colon - Transverse | TCGA-COAD | 21.91 | 6.56 |
| GTEX-V955-1626-SM-4JBHJ | Colon - Transverse | TCGA-COAD | 47.51 | 12.29 |
| GTEX-VJYA-2026-SM-4KL1K | Colon - Transverse | TCGA-COAD | 28.73 | 6.9 |
| GTEX-W5WG-2426-SM-4LMI6 | Colon - Transverse | TCGA-COAD | 27.48 | 7.24 |
| GTEX-WEY5-1426-SM-4LMJ3 | Colon - Transverse | TCGA-COAD | 48.39 | 13.05 |
| GTEX-WFG7-1526-SM-4LVMG | Colon - Transverse | TCGA-COAD | 36.12 | 8.29 |
| GTEX-WFON-1426-SM-4LVMT | Colon - Transverse | TCGA-COAD | 29.5 | 3.83 |
| GTEX-WH7G-1326-SM-4LVMS | Colon - Transverse | TCGA-COAD | 26.67 | 4.86 |
| GTEX-WHSB-0926-SM-4M1XJ | Colon - Transverse | TCGA-COAD | 43.88 | 15.16 |
| GTEX-WHWD-1226-SM-4OOS1 | Colon - Transverse | TCGA-COAD | 33.45 | 7.22 |
| GTEX-WOFM-2026-SM-4OOSQ | Colon - Transverse | TCGA-COAD | 27.07 | 5.62 |
| GTEX-WQUQ-2526-SM-4MVNO | Colon - Transverse | TCGA-COAD | 46.92 | 12.66 |
| GTEX-WVJS-2426-SM-EZ6LI | Colon - Transverse | TCGA-COAD | 37.35 | 13.07 |
| GTEX-WY7C-1626-SM-4ONC8 | Colon - Transverse | TCGA-COAD | 54.73 | 15.79 |
| GTEX-WYJK-2226-SM-4ONDJ | Colon - Transverse | TCGA-COAD | 32.43 | 7.25 |
| GTEX-WYVS-1826-SM-4OND6 | Colon - Transverse | TCGA-COAD | 40.13 | 10.08 |
| GTEX-X15G-1326-SM-4PQZJ | Colon - Transverse | TCGA-COAD | 32.71 | 8.74 |
| GTEX-X3Y1-1726-SM-4PQZL | Colon - Transverse | TCGA-COAD | 36.93 | 9.51 |
| GTEX-X4EO-2526-SM-EWRM8 | Colon - Transverse | TCGA-COAD | 37.47 | 15.49 |
| GTEX-X5EB-0926-SM-46MVT | Colon - Transverse | TCGA-COAD | 36.48 | 13.51 |

| | | | | |
|---|---|---|---|---|
| GTEX-XAJ8-0426-SM-47JYJ | Colon - Transverse | TCGA-COAD | 41.83 | 20.63 |
| GTEX-XBED-1526-SM-4AT5W | Colon - Transverse | TCGA-COAD | 31.9 | 9.85 |
| GTEX-XBEW-1926-SM-4SOJO | Colon - Transverse | TCGA-COAD | 19.21 | 4.11 |
| GTEX-XGQ4-1226-SM-4AT67 | Colon - Transverse | TCGA-COAD | 33.56 | 9.45 |
| GTEX-XMK1-1726-SM-4B64Z | Colon - Transverse | TCGA-COAD | 38.63 | 11.99 |
| GTEX-XPVG-1826-SM-4B64X | Colon - Transverse | TCGA-COAD | 25.29 | 5.69 |
| GTEX-XQ8I-2126-SM-4BOOM | Colon - Transverse | TCGA-COAD | 27.76 | 5.7 |
| GTEX-XUW1-1926-SM-4BOP1 | Colon - Transverse | TCGA-COAD | 35.46 | 8.87 |
| GTEX-XUZC-1326-SM-4BRV2 | Colon - Transverse | TCGA-COAD | 32.12 | 7.86 |
| GTEX-XV7Q-2126-SM-4BRVX | Colon - Transverse | TCGA-COAD | 41.2 | 11.75 |
| GTEX-XXEK-1926-SM-4BRVD | Colon - Transverse | TCGA-COAD | 53.96 | 14.32 |
| GTEX-XYKS-2226-SM-4E3IU | Colon - Transverse | TCGA-COAD | 29.29 | 6.95 |
| GTEX-Y114-1526-SM-4TT79 | Colon - Transverse | TCGA-COAD | 41.24 | 10.62 |
| GTEX-Y3I4-1426-SM-4TT9C | Colon - Transverse | TCGA-COAD | 34.7 | 9.06 |
| GTEX-Y3IK-1826-SM-4WWDS | Colon - Transverse | TCGA-COAD | 33.73 | 8.28 |
| GTEX-Y5LM-1526-SM-4VDTA | Colon - Transverse | TCGA-COAD | 34.14 | 6.72 |
| GTEX-Y8E4-2426-SM-4WWDN | Colon - Transverse | TCGA-COAD | 36.44 | 9.62 |
| GTEX-Y8LW-1226-SM-4WWDP | Colon - Transverse | TCGA-COAD | 45.67 | 12.18 |
| GTEX-Y9LG-1526-SM-4WWDR | Colon - Transverse | TCGA-COAD | 24.23 | 3.38 |
| GTEX-YB5E-1626-SM-5IFIQ | Colon - Transverse | TCGA-COAD | 30.76 | 5.1 |
| GTEX-YEC3-2426-SM-DO11A | Colon - Transverse | TCGA-COAD | 24.58 | 5.86 |
| GTEX-YEC4-1926-SM-5IFHJ | Colon - Transverse | TCGA-COAD | 40.28 | 12.02 |
| GTEX-YF7O-1726-SM-5IFJ9 | Colon - Transverse | TCGA-COAD | 36.45 | 9.15 |
| GTEX-Z93S-2426-SM-5HL8N | Colon - Transverse | TCGA-COAD | 39.89 | 11.51 |
| GTEX-ZAB4-2226-SM-5HL97 | Colon - Transverse | TCGA-COAD | 31.5 | 10.46 |
| GTEX-ZDTT-1926-SM-5HL5W | Colon - Transverse | TCGA-COAD | 32.53 | 7.84 |
| GTEX-ZDYS-2326-SM-4WKG6 | Colon - Transverse | TCGA-COAD | 32.63 | 6.2 |
| GTEX-ZEX8-1926-SM-4WKFR | Colon - Transverse | TCGA-COAD | 37.37 | 9.73 |
| GTEX-ZF29-1726-SM-4WKFV | Colon - Transverse | TCGA-COAD | 41.5 | 8.74 |
| GTEX-ZF2S-1926-SM-4WWAN | Colon - Transverse | TCGA-COAD | 37.56 | 8.92 |
| GTEX-ZG7Y-2126-SM-DO117 | Colon - Transverse | TCGA-COAD | 35.67 | 8.21 |

| | | | | |
|---|---|---|---|---|
| GTEX-ZGAY-2126-SM-4YCFB | Colon - Transverse | TCGA-COAD | 32.95 | 7.42 |
| GTEX-ZLV1-1226-SM-4WWBX | Colon - Transverse | TCGA-COAD | 28.05 | 5.72 |
| GTEX-ZLWG-1726-SM-DNZZ2 | Colon - Transverse | TCGA-COAD | 31.43 | 6.69 |
| GTEX-ZP4G-1726-SM-4WWCM | Colon - Transverse | TCGA-COAD | 30.71 | 7.09 |
| GTEX-ZPCL-1826-SM-57WF1 | Colon - Transverse | TCGA-COAD | 32.45 | 6.28 |
| GTEX-ZPIC-2226-SM-57WDX | Colon - Transverse | TCGA-COAD | 31.54 | 6.4 |
| GTEX-ZPU1-1926-SM-57WDT | Colon - Transverse | TCGA-COAD | 33.99 | 8.94 |
| GTEX-ZT9W-1826-SM-51MQX | Colon - Transverse | TCGA-COAD | 26.26 | 4.14 |
| GTEX-ZTSS-1126-SM-5985O | Colon - Transverse | TCGA-COAD | 38.37 | 6.68 |
| GTEX-ZV6S-1226-SM-59HLH | Colon - Transverse | TCGA-COAD | 47.33 | 10.26 |
| GTEX-ZV7C-1526-SM-5NQ72 | Colon - Transverse | TCGA-COAD | 37.29 | 7.27 |
| GTEX-ZVP2-1526-SM-5NQ82 | Colon - Transverse | TCGA-COAD | 34.83 | 5.81 |
| GTEX-ZVT2-1926-SM-5GU5T | Colon - Transverse | TCGA-COAD | 24.35 | 3.58 |
| GTEX-ZXES-1626-SM-5NQ7L | Colon - Transverse | TCGA-COAD | 46.15 | 7.89 |
| GTEX-ZY6K-1226-SM-5GZYL | Colon - Transverse | TCGA-COAD | 45.1 | 9.18 |
| GTEX-ZYFC-2626-SM-5NQ6S | Colon - Transverse | TCGA-COAD | 36.76 | 11.55 |
| GTEX-ZYFG-1926-SM-5GID5 | Colon - Transverse | TCGA-COAD | 39.15 | 9.44 |
| GTEX-ZYVF-2826-SM-5GIDK | Colon - Transverse | TCGA-COAD | 52.56 | 14.53 |
| GTEX-ZZ64-0726-SM-5GZX4 | Colon - Transverse | TCGA-COAD | 48.6 | 7.74 |
| GTEX-111CU-0726-SM-5GZYD | Esophagus - Mucosa | TCGA-ESCA | 38.9 | 8.84 |
| GTEX-111YS-0926-SM-5EGGI | Esophagus - Mucosa | TCGA-ESCA | 50.31 | 18.44 |
| GTEX-1122O-1726-SM-5GICM | Esophagus - Mucosa | TCGA-ESCA | 32.89 | 6.91 |
| GTEX-117YW-1926-SM-5H121 | Esophagus - Mucosa | TCGA-ESCA | 19.48 | 6.04 |
| GTEX-11DXX-1126-SM-5GZZK | Esophagus - Mucosa | TCGA-ESCA | 56.32 | 18.63 |
| GTEX-11DXY-1426-SM-5N9CN | Esophagus - Mucosa | TCGA-ESCA | 29.45 | 6.92 |
| GTEX-11DXZ-1226-SM-5H11L | Esophagus - Mucosa | TCGA-ESCA | 29.6 | 9.95 |
| GTEX-11DYG-1326-SM-5N9DO | Esophagus - Mucosa | TCGA-ESCA | 75.74 | 21.03 |
| GTEX-11DZ1-2026-SM-5A5KG | Esophagus - Mucosa | TCGA-ESCA | 58.59 | 16.03 |
| GTEX-11EI6-1526-SM-5P9GL | Esophagus - Mucosa | TCGA-ESCA | 16.28 | 6.27 |
| GTEX-11EM3-0826-SM-5N9CC | Esophagus - Mucosa | TCGA-ESCA | 52.56 | 13.53 |
| GTEX-11EMC-2226-SM-5A5JW | Esophagus - Mucosa | TCGA-ESCA | 60 | 22.03 |

| | | | | |
|---|---|---|---|---|
| GTEX-11EQ9-0726-SM-5A5K2 | Esophagus - Mucosa | TCGA-ESCA | 34.12 | 9.66 |
| GTEX-11GSP-1826-SM-5N9DQ | Esophagus - Mucosa | TCGA-ESCA | 43.53 | 9.7 |
| GTEX-11I78-1526-SM-5A5K7 | Esophagus - Mucosa | TCGA-ESCA | 41.5 | 11.69 |
| GTEX-11LCK-1726-SM-5S2OQ | Esophagus - Mucosa | TCGA-ESCA | 46.79 | 18.25 |
| GTEX-11NSD-1126-SM-5N9BQ | Esophagus - Mucosa | TCGA-ESCA | 38.77 | 16.2 |
| GTEX-11NUK-1326-SM-5P9GN | Esophagus - Mucosa | TCGA-ESCA | 30.96 | 10.83 |
| GTEX-11ONC-1526-SM-5GU6S | Esophagus - Mucosa | TCGA-ESCA | 28.31 | 8.79 |
| GTEX-11P7K-1326-SM-5BC5G | Esophagus - Mucosa | TCGA-ESCA | 27.96 | 7.3 |
| GTEX-11P81-0826-SM-5P9GR | Esophagus - Mucosa | TCGA-ESCA | 46.1 | 14.44 |
| GTEX-11P82-0526-SM-5986P | Esophagus - Mucosa | TCGA-ESCA | 35.34 | 10.32 |
| GTEX-11TT1-0526-SM-5P9JO | Esophagus - Mucosa | TCGA-ESCA | 57.22 | 22.57 |
| GTEX-11TUW-1326-SM-5EQKV | Esophagus - Mucosa | TCGA-ESCA | 34.7 | 10.94 |
| GTEX-11VI4-0526-SM-5EQ48 | Esophagus - Mucosa | TCGA-ESCA | 36.72 | 8.95 |
| GTEX-11XUK-0826-SM-5GU6P | Esophagus - Mucosa | TCGA-ESCA | 43.03 | 9.59 |
| GTEX-11ZTS-2626-SM-5EGIC | Esophagus - Mucosa | TCGA-ESCA | 30.04 | 8.53 |
| GTEX-11ZU8-1526-SM-5EGHX | Esophagus - Mucosa | TCGA-ESCA | 36.23 | 9.02 |
| GTEX-1211K-0926-SM-5FQTL | Esophagus - Mucosa | TCGA-ESCA | 39.25 | 12.25 |
| GTEX-1212Z-2126-SM-5FQTM | Esophagus - Mucosa | TCGA-ESCA | 46.11 | 11.31 |
| GTEX-12696-1326-SM-5FQTJ | Esophagus - Mucosa | TCGA-ESCA | 40.93 | 12 |
| GTEX-1269C-1026-SM-5N9EY | Esophagus - Mucosa | TCGA-ESCA | 16.36 | 4.29 |
| GTEX-12BJ1-1626-SM-5HL8Y | Esophagus - Mucosa | TCGA-ESCA | 29.45 | 5.99 |
| GTEX-12C56-1226-SM-5FQSU | Esophagus - Mucosa | TCGA-ESCA | 35.91 | 8.66 |
| GTEX-12KS4-0926-SM-5BC5S | Esophagus - Mucosa | TCGA-ESCA | 40.23 | 16.31 |
| GTEX-12WSA-1426-SM-5EGJU | Esophagus - Mucosa | TCGA-ESCA | 46.57 | 12.2 |
| GTEX-12WSD-1826-SM-5GCOA | Esophagus - Mucosa | TCGA-ESCA | 36.12 | 14.13 |
| GTEX-12WSG-5007-SM-7DUGZ | Esophagus - Mucosa | TCGA-ESCA | 50.2 | 13.48 |
| GTEX-12WSK-1126-SM-5LZV8 | Esophagus - Mucosa | TCGA-ESCA | 35.47 | 6.43 |
| GTEX-12WSL-1526-SM-5GCP8 | Esophagus - Mucosa | TCGA-ESCA | 56.16 | 22.48 |
| GTEX-12WSN-5013-SM-7P8P9 | Esophagus - Mucosa | TCGA-ESCA | 62.37 | 24.12 |
| GTEX-12ZZW-1726-SM-5DUVD | Esophagus - Mucosa | TCGA-ESCA | 44.36 | 14.06 |
| GTEX-12ZZX-1326-SM-5DUWI | Esophagus - Mucosa | TCGA-ESCA | 42 | 12.14 |

| | | | | |
|---|---|---|---|---|
| GTEX-12ZZY-1426-SM-5N9EI | Esophagus - Mucosa | TCGA-ESCA | 28.58 | 9.16 |
| GTEX-13111-0826-SM-5LZV3 | Esophagus - Mucosa | TCGA-ESCA | 45.77 | 12.97 |
| GTEX-131XE-1426-SM-5HL7V | Esophagus - Mucosa | TCGA-ESCA | 37.53 | 10.71 |
| GTEX-131XF-0826-SM-5GICG | Esophagus - Mucosa | TCGA-ESCA | 48.14 | 12.9 |
| GTEX-131XG-1026-SM-5GCMX | Esophagus - Mucosa | TCGA-ESCA | 35.11 | 8.73 |
| GTEX-131YS-1426-SM-5IJF7 | Esophagus - Mucosa | TCGA-ESCA | 34.78 | 10.51 |
| GTEX-132AR-2226-SM-5J2NT | Esophagus - Mucosa | TCGA-ESCA | 44.8 | 10.62 |
| GTEX-132Q8-1826-SM-5EGIY | Esophagus - Mucosa | TCGA-ESCA | 39.89 | 10.23 |
| GTEX-132QS-1426-SM-5P9GE | Esophagus - Mucosa | TCGA-ESCA | 47.24 | 12.8 |
| GTEX-133LE-0626-SM-5IFFJ | Esophagus - Mucosa | TCGA-ESCA | 60.95 | 20.78 |
| GTEX-1399Q-1126-SM-5P9JA | Esophagus - Mucosa | TCGA-ESCA | 35.65 | 8.12 |
| GTEX-1399R-0826-SM-5IJEL | Esophagus - Mucosa | TCGA-ESCA | 34.6 | 9.26 |
| GTEX-1399S-1326-SM-5P9GH | Esophagus - Mucosa | TCGA-ESCA | 29.07 | 7.33 |
| GTEX-1399U-1026-SM-5J1OC | Esophagus - Mucosa | TCGA-ESCA | 58.61 | 13.56 |
| GTEX-139T6-0626-SM-5IFF9 | Esophagus - Mucosa | TCGA-ESCA | 43.65 | 11.59 |
| GTEX-139TU-1226-SM-5J2NL | Esophagus - Mucosa | TCGA-ESCA | 37.06 | 13.37 |
| GTEX-139UW-1126-SM-5IJEJ | Esophagus - Mucosa | TCGA-ESCA | 54.56 | 14.56 |
| GTEX-139YR-1726-SM-5IFFK | Esophagus - Mucosa | TCGA-ESCA | 38.39 | 11.77 |
| GTEX-13CF3-1626-SM-5IFG9 | Esophagus - Mucosa | TCGA-ESCA | 64.79 | 19.28 |
| GTEX-13D11-1926-SM-5IFGB | Esophagus - Mucosa | TCGA-ESCA | 42.48 | 10.4 |
| GTEX-13FH7-0526-SM-5IJF8 | Esophagus - Mucosa | TCGA-ESCA | 50.26 | 15.81 |
| GTEX-13FHO-1726-SM-DPRZX | Esophagus - Mucosa | TCGA-ESCA | 36.69 | 10.43 |
| GTEX-13FHP-1526-SM-5IJGE | Esophagus - Mucosa | TCGA-ESCA | 12.72 | 4.42 |
| GTEX-13FLV-1126-SM-5J2NO | Esophagus - Mucosa | TCGA-ESCA | 41.52 | 8.22 |
| GTEX-13FTW-1626-SM-5KM4N | Esophagus - Mucosa | TCGA-ESCA | 39.32 | 10.66 |
| GTEX-13FTX-0526-SM-5IFGS | Esophagus - Mucosa | TCGA-ESCA | 33.97 | 10.45 |
| GTEX-13G51-0926-SM-5IFFU | Esophagus - Mucosa | TCGA-ESCA | 35.93 | 9.13 |
| GTEX-13JVG-2026-SM-5KM1N | Esophagus - Mucosa | TCGA-ESCA | 37.4 | 11.12 |
| GTEX-13NYB-0926-SM-5IFFA | Esophagus - Mucosa | TCGA-ESCA | 33.07 | 8.49 |
| GTEX-13NZB-1226-SM-5J2MW | Esophagus - Mucosa | TCGA-ESCA | 29.71 | 8.27 |
| GTEX-13O1R-1326-SM-5IJF5 | Esophagus - Mucosa | TCGA-ESCA | 72.12 | 18.68 |

| | | | | |
|---|---|---|---|---|
| GTEX-13O21-0626-SM-5IJDI | Esophagus - Mucosa | TCGA-ESCA | 42.34 | 13.95 |
| GTEX-13O3Q-0826-SM-5IFGF | Esophagus - Mucosa | TCGA-ESCA | 32.7 | 8.41 |
| GTEX-13O61-0926-SM-5IFFE | Esophagus - Mucosa | TCGA-ESCA | 39.25 | 9.34 |
| GTEX-13OVG-0626-SM-5IFEP | Esophagus - Mucosa | TCGA-ESCA | 36.33 | 9.45 |
| GTEX-13OVI-2626-SM-5KM4T | Esophagus - Mucosa | TCGA-ESCA | 27.04 | 6.99 |
| GTEX-13OVJ-1226-SM-5L3HA | Esophagus - Mucosa | TCGA-ESCA | 55.91 | 17.79 |
| GTEX-13OVK-1126-SM-6PAMR | Esophagus - Mucosa | TCGA-ESCA | 37.42 | 14.68 |
| GTEX-13OW8-2326-SM-5L3I9 | Esophagus - Mucosa | TCGA-ESCA | 29.73 | 7.84 |
| GTEX-13PL7-0226-SM-731DT | Esophagus - Mucosa | TCGA-ESCA | 29.52 | 9.8 |
| GTEX-13PVQ-1726-SM-5IFF3 | Esophagus - Mucosa | TCGA-ESCA | 64.93 | 16.14 |
| GTEX-13PVR-1826-SM-5Q5CC | Esophagus - Mucosa | TCGA-ESCA | 40.47 | 8.94 |
| GTEX-13QBU-1026-SM-5LU3R | Esophagus - Mucosa | TCGA-ESCA | 33.24 | 10.46 |
| GTEX-13QJ3-1726-SM-5RQK9 | Esophagus - Mucosa | TCGA-ESCA | 23.36 | 6.4 |
| GTEX-13RTK-0526-SM-5S2QF | Esophagus - Mucosa | TCGA-ESCA | 41.77 | 10.52 |
| GTEX-13S86-0826-SM-5SI6J | Esophagus - Mucosa | TCGA-ESCA | 50.27 | 11.65 |
| GTEX-13SLX-1726-SM-5Q5E5 | Esophagus - Mucosa | TCGA-ESCA | 51.43 | 14.47 |
| GTEX-13U4I-1126-SM-5LU3S | Esophagus - Mucosa | TCGA-ESCA | 39.92 | 14.4 |
| GTEX-13W3W-1626-SM-5LU35 | Esophagus - Mucosa | TCGA-ESCA | 46.27 | 14.09 |
| GTEX-13W46-1926-SM-5J2MF | Esophagus - Mucosa | TCGA-ESCA | 26.33 | 7.79 |
| GTEX-13X6H-1326-SM-5LU42 | Esophagus - Mucosa | TCGA-ESCA | 59.98 | 20.19 |
| GTEX-13X6I-1026-SM-5LU4C | Esophagus - Mucosa | TCGA-ESCA | 64.92 | 21.28 |
| GTEX-13X6K-2126-SM-5O9D4 | Esophagus - Mucosa | TCGA-ESCA | 48.83 | 20.24 |
| GTEX-13YAN-1926-SM-7EPH9 | Esophagus - Mucosa | TCGA-ESCA | 30.43 | 7.2 |
| GTEX-144FL-2726-SM-5O98Z | Esophagus - Mucosa | TCGA-ESCA | 52.05 | 12.52 |
| GTEX-144GM-1426-SM-5LU31 | Esophagus - Mucosa | TCGA-ESCA | 40.52 | 8.96 |
| GTEX-144GN-0626-SM-5O9B2 | Esophagus - Mucosa | TCGA-ESCA | 48.7 | 16.45 |
| GTEX-144GO-2726-SM-5O9A2 | Esophagus - Mucosa | TCGA-ESCA | 68.38 | 15.86 |
| GTEX-145LS-1826-SM-5LUAZ | Esophagus - Mucosa | TCGA-ESCA | 39.94 | 14.76 |
| GTEX-145LT-1126-SM-5LUA1 | Esophagus - Mucosa | TCGA-ESCA | 36.13 | 8.96 |
| GTEX-145LU-0926-SM-79OJL | Esophagus - Mucosa | TCGA-ESCA | 18.84 | 7.6 |
| GTEX-145LV-1326-SM-5S2QI | Esophagus - Mucosa | TCGA-ESCA | 23.46 | 7.23 |

| | | | | |
|---|---|---|---|---|
| GTEX-145ME-5004-SM-7MGXP | Esophagus - Mucosa | TCGA-ESCA | 46.02 | 11.7 |
| GTEX-145MF-1426-SM-5O9B6 | Esophagus - Mucosa | TCGA-ESCA | 50.76 | 14.99 |
| GTEX-145MN-1026-SM-5NQ9G | Esophagus - Mucosa | TCGA-ESCA | 38.21 | 10.7 |
| GTEX-145MO-1426-SM-DTX9W | Esophagus - Mucosa | TCGA-ESCA | 42.45 | 12.04 |
| GTEX-146FH-1726-SM-5QGQ2 | Esophagus - Mucosa | TCGA-ESCA | 52.26 | 19.22 |
| GTEX-146FQ-1026-SM-5LU8I | Esophagus - Mucosa | TCGA-ESCA | 54.27 | 21.01 |
| GTEX-146FR-0426-SM-5SI97 | Esophagus - Mucosa | TCGA-ESCA | 45.77 | 13.11 |
| GTEX-147F3-0826-SM-5NQA7 | Esophagus - Mucosa | TCGA-ESCA | 39.14 | 8.63 |
| GTEX-148VI-1026-SM-5TDDJ | Esophagus - Mucosa | TCGA-ESCA | 57.02 | 17.09 |
| GTEX-148VJ-1026-SM-5LU98 | Esophagus - Mucosa | TCGA-ESCA | 26.09 | 5.74 |
| GTEX-1497J-2726-SM-5SI9W | Esophagus - Mucosa | TCGA-ESCA | 50.58 | 17.37 |
| GTEX-14A6H-1326-SM-5NQBC | Esophagus - Mucosa | TCGA-ESCA | 18.82 | 5.99 |
| GTEX-14AS3-0626-SM-5QGQD | Esophagus - Mucosa | TCGA-ESCA | 36.83 | 11.41 |
| GTEX-14B4R-0326-SM-5TDDP | Esophagus - Mucosa | TCGA-ESCA | 49.35 | 15.91 |
| GTEX-14BIM-1326-SM-7EWD1 | Esophagus - Mucosa | TCGA-ESCA | 47.68 | 14.91 |
| GTEX-14BIN-2726-SM-5RQIE | Esophagus - Mucosa | TCGA-ESCA | 32.31 | 13.07 |
| GTEX-14BMU-0926-SM-5S2QB | Esophagus - Mucosa | TCGA-ESCA | 67.69 | 17.43 |
| GTEX-14C38-2126-SM-5RQJO | Esophagus - Mucosa | TCGA-ESCA | 48.23 | 17.48 |
| GTEX-14C5O-1626-SM-73KYH | Esophagus - Mucosa | TCGA-ESCA | 22.58 | 8.68 |
| GTEX-14DAQ-1426-SM-5RQJD | Esophagus - Mucosa | TCGA-ESCA | 33.01 | 9.47 |
| GTEX-14DAR-0826-SM-73KYJ | Esophagus - Mucosa | TCGA-ESCA | 29.16 | 7.04 |
| GTEX-14E1K-1226-SM-5S2PD | Esophagus - Mucosa | TCGA-ESCA | 35.79 | 10.5 |
| GTEX-14E6E-0726-SM-5S2R3 | Esophagus - Mucosa | TCGA-ESCA | 55.14 | 15.72 |
| GTEX-14H4A-1626-SM-5SI69 | Esophagus - Mucosa | TCGA-ESCA | 34.35 | 8.83 |
| GTEX-14ICK-2026-SM-6AJB2 | Esophagus - Mucosa | TCGA-ESCA | 38.58 | 9.27 |
| GTEX-14ICL-0526-SM-5RQJ6 | Esophagus - Mucosa | TCGA-ESCA | 40.81 | 12.02 |
| GTEX-14JG6-0926-SM-6871Y | Esophagus - Mucosa | TCGA-ESCA | 48.65 | 10.02 |
| GTEX-14JIY-0926-SM-6871D | Esophagus - Mucosa | TCGA-ESCA | 28.34 | 8.27 |
| GTEX-14PHY-0626-SM-664NN | Esophagus - Mucosa | TCGA-ESCA | 35.74 | 11.31 |
| GTEX-14PJ3-0626-SM-6UGC5 | Esophagus - Mucosa | TCGA-ESCA | 39.24 | 10.17 |
| GTEX-14PJ4-1526-SM-664OA | Esophagus - Mucosa | TCGA-ESCA | 59.8 | 18.93 |

| | | | | |
|---|---|---|---|---|
| GTEX-14PJ5-0526-SM-664OE | Esophagus - Mucosa | TCGA-ESCA | 39.65 | 10.62 |
| GTEX-14PJ6-1026-SM-686ZB | Esophagus - Mucosa | TCGA-ESCA | 73.83 | 25.5 |
| GTEX-14PJM-1526-SM-664NY | Esophagus - Mucosa | TCGA-ESCA | 38.17 | 11.21 |
| GTEX-14PJN-0526-SM-68724 | Esophagus - Mucosa | TCGA-ESCA | 41.92 | 13.11 |
| GTEX-14PK6-1326-SM-686ZE | Esophagus - Mucosa | TCGA-ESCA | 63.65 | 23.01 |
| GTEX-14PKV-1726-SM-6871U | Esophagus - Mucosa | TCGA-ESCA | 41.08 | 10.59 |
| GTEX-15DCD-1626-SM-6AJBH | Esophagus - Mucosa | TCGA-ESCA | 49.24 | 17.06 |
| GTEX-15DCE-1326-SM-6LPJL | Esophagus - Mucosa | TCGA-ESCA | 76.41 | 24.64 |
| GTEX-15DYW-1826-SM-6LPIY | Esophagus - Mucosa | TCGA-ESCA | 30.97 | 7.74 |
| GTEX-15EO6-1026-SM-7KUG9 | Esophagus - Mucosa | TCGA-ESCA | 35.73 | 11.75 |
| GTEX-15EOM-5013-SM-7MKF9 | Esophagus - Mucosa | TCGA-ESCA | 34.49 | 15.2 |
| GTEX-15ER7-2126-SM-7939E | Esophagus - Mucosa | TCGA-ESCA | 44.39 | 10.05 |
| GTEX-15ETS-1926-SM-7KUN5 | Esophagus - Mucosa | TCGA-ESCA | 35.95 | 10.95 |
| GTEX-15EU6-1626-SM-6LPKF | Esophagus - Mucosa | TCGA-ESCA | 48.61 | 12.52 |
| GTEX-15F5U-5010-SM-AHZ2D | Esophagus - Mucosa | TCGA-ESCA | 57.49 | 20.45 |
| GTEX-15RIE-0826-SM-6M47X | Esophagus - Mucosa | TCGA-ESCA | 34.86 | 12.1 |
| GTEX-15RIF-0826-SM-6M47C | Esophagus - Mucosa | TCGA-ESCA | 37.91 | 7.94 |
| GTEX-15RJ7-0926-SM-6LPJO | Esophagus - Mucosa | TCGA-ESCA | 49.83 | 18.92 |
| GTEX-15RJE-0926-SM-6LLHZ | Esophagus - Mucosa | TCGA-ESCA | 40.29 | 12.28 |
| GTEX-15SB6-0726-SM-6M48F | Esophagus - Mucosa | TCGA-ESCA | 45.47 | 10.09 |
| GTEX-15SHU-1426-SM-6LPIN | Esophagus - Mucosa | TCGA-ESCA | 38.36 | 13.47 |
| GTEX-15UF6-2326-SM-6M465 | Esophagus - Mucosa | TCGA-ESCA | 57.48 | 16.06 |
| GTEX-15UF7-0926-SM-6M46F | Esophagus - Mucosa | TCGA-ESCA | 34.81 | 9.71 |
| GTEX-169BO-0526-SM-793AK | Esophagus - Mucosa | TCGA-ESCA | 41.1 | 15.82 |
| GTEX-16A39-1026-SM-6LPJU | Esophagus - Mucosa | TCGA-ESCA | 53.3 | 12.72 |
| GTEX-16AAH-0726-SM-793A9 | Esophagus - Mucosa | TCGA-ESCA | 35.87 | 10.16 |
| GTEX-16BQI-1526-SM-6LLIE | Esophagus - Mucosa | TCGA-ESCA | 51.65 | 13.68 |
| GTEX-16MT8-1326-SM-6M47R | Esophagus - Mucosa | TCGA-ESCA | 43.14 | 10.6 |
| GTEX-16MTA-0826-SM-6LLIC | Esophagus - Mucosa | TCGA-ESCA | 54.09 | 14.57 |
| GTEX-16NGA-1426-SM-718AC | Esophagus - Mucosa | TCGA-ESCA | 36.22 | 11.16 |
| GTEX-16NPX-1626-SM-6LLI5 | Esophagus - Mucosa | TCGA-ESCA | 40.19 | 12.7 |

| | | | | |
|---|---|---|---|---|
| GTEX-16XZY-0926-SM-7938M | Esophagus - Mucosa | TCGA-ESCA | 43.31 | 12.07 |
| GTEX-16Z82-1926-SM-6M47U | Esophagus - Mucosa | TCGA-ESCA | 18.52 | 4.56 |
| GTEX-178AV-0526-SM-6LPJH | Esophagus - Mucosa | TCGA-ESCA | 53.26 | 14.09 |
| GTEX-17EUY-1426-SM-7938X | Esophagus - Mucosa | TCGA-ESCA | 35.35 | 11.44 |
| GTEX-17EVQ-1926-SM-7IGOF | Esophagus - Mucosa | TCGA-ESCA | 26.84 | 8.66 |
| GTEX-17F96-2526-SM-7DHL3 | Esophagus - Mucosa | TCGA-ESCA | 31.47 | 7.35 |
| GTEX-17F97-1326-SM-7MGVO | Esophagus - Mucosa | TCGA-ESCA | 46.49 | 13.32 |
| GTEX-17F98-1726-SM-7938O | Esophagus - Mucosa | TCGA-ESCA | 39.67 | 16.8 |
| GTEX-17F9E-1926-SM-7IGQF | Esophagus - Mucosa | TCGA-ESCA | 62.25 | 19.84 |
| GTEX-17HGU-1426-SM-7938G | Esophagus - Mucosa | TCGA-ESCA | 43.03 | 14.27 |
| GTEX-17HHE-0926-SM-7938K | Esophagus - Mucosa | TCGA-ESCA | 23.86 | 10.41 |
| GTEX-17HII-1526-SM-9OSVD | Esophagus - Mucosa | TCGA-ESCA | 16.62 | 4.85 |
| GTEX-17JCI-1226-SM-793D9 | Esophagus - Mucosa | TCGA-ESCA | 26.17 | 8.02 |
| GTEX-17KNJ-1326-SM-7KFTG | Esophagus - Mucosa | TCGA-ESCA | 56.88 | 16.41 |
| GTEX-17MFQ-0526-SM-7KFTX | Esophagus - Mucosa | TCGA-ESCA | 51.88 | 15.87 |
| GTEX-183FY-1526-SM-7IGOW | Esophagus - Mucosa | TCGA-ESCA | 39.14 | 10.27 |
| GTEX-183WM-1726-SM-7KFT5 | Esophagus - Mucosa | TCGA-ESCA | 63.97 | 15.64 |
| GTEX-18465-1126-SM-7LG6E | Esophagus - Mucosa | TCGA-ESCA | 25.24 | 6.75 |
| GTEX-18A66-1426-SM-7KFRT | Esophagus - Mucosa | TCGA-ESCA | 43.33 | 15.27 |
| GTEX-18A67-2426-SM-7LT95 | Esophagus - Mucosa | TCGA-ESCA | 45.95 | 14.01 |
| GTEX-18A6Q-1226-SM-7KFS9 | Esophagus - Mucosa | TCGA-ESCA | 41.84 | 11.73 |
| GTEX-18A7A-2326-SM-7KFRH | Esophagus - Mucosa | TCGA-ESCA | 50.13 | 16.71 |
| GTEX-18D9A-0526-SM-7KFSW | Esophagus - Mucosa | TCGA-ESCA | 35.23 | 8.63 |
| GTEX-18D9B-2026-SM-7LT96 | Esophagus - Mucosa | TCGA-ESCA | 37.59 | 14.81 |
| GTEX-18D9U-2126-SM-7LG4Y | Esophagus - Mucosa | TCGA-ESCA | 34.05 | 10.52 |
| GTEX-18QFQ-1026-SM-7LG6B | Esophagus - Mucosa | TCGA-ESCA | 49.19 | 13.74 |
| GTEX-1A32A-1826-SM-72D5G | Esophagus - Mucosa | TCGA-ESCA | 42.38 | 12 |
| GTEX-1A3MV-1326-SM-7LG6C | Esophagus - Mucosa | TCGA-ESCA | 42.79 | 8.62 |
| GTEX-1A8FM-1526-SM-7MXTA | Esophagus - Mucosa | TCGA-ESCA | 31.64 | 10.48 |
| GTEX-1AMEY-0226-SM-7189Q | Esophagus - Mucosa | TCGA-ESCA | 38.66 | 12.21 |
| GTEX-1AMFI-0226-SM-731D5 | Esophagus - Mucosa | TCGA-ESCA | 28.57 | 7.45 |

| | | | | |
|---|---|---|---|---|
| GTEX-1AX8Z-5016-SM-AHZ2B | Esophagus - Mucosa | TCGA-ESCA | 70.04 | 28.31 |
| GTEX-1AX9I-1426-SM-7DHM8 | Esophagus - Mucosa | TCGA-ESCA | 24.8 | 8.35 |
| GTEX-1AX9J-1826-SM-731FM | Esophagus - Mucosa | TCGA-ESCA | 44.07 | 9.72 |
| GTEX-1AX9K-1126-SM-731DQ | Esophagus - Mucosa | TCGA-ESCA | 52.78 | 18.51 |
| GTEX-1AYD5-0926-SM-793AF | Esophagus - Mucosa | TCGA-ESCA | 29.1 | 14.11 |
| GTEX-1B8KE-1826-SM-731FD | Esophagus - Mucosa | TCGA-ESCA | 34.96 | 6.26 |
| GTEX-1B8KZ-0726-SM-7938W | Esophagus - Mucosa | TCGA-ESCA | 33.91 | 15.78 |
| GTEX-1B933-2026-SM-7EWFC | Esophagus - Mucosa | TCGA-ESCA | 26.13 | 10.97 |
| GTEX-1B97I-0926-SM-731DZ | Esophagus - Mucosa | TCGA-ESCA | 46.28 | 12.82 |
| GTEX-1B97J-1026-SM-7MXTE | Esophagus - Mucosa | TCGA-ESCA | 42.07 | 11.4 |
| GTEX-1B996-1826-SM-7MGY6 | Esophagus - Mucosa | TCGA-ESCA | 54.14 | 14.11 |
| GTEX-1BAJH-2026-SM-7P8OZ | Esophagus - Mucosa | TCGA-ESCA | 45.42 | 10.31 |
| GTEX-1C2JI-1126-SM-731FH | Esophagus - Mucosa | TCGA-ESCA | 45.17 | 12.86 |
| GTEX-1C475-0926-SM-7MKFI | Esophagus - Mucosa | TCGA-ESCA | 64.2 | 16.6 |
| GTEX-1C64O-1826-SM-7DHM4 | Esophagus - Mucosa | TCGA-ESCA | 34.06 | 11.34 |
| GTEX-1C6VQ-1826-SM-7EWEH | Esophagus - Mucosa | TCGA-ESCA | 76.29 | 22.32 |
| GTEX-1CAMQ-2726-SM-7MKGD | Esophagus - Mucosa | TCGA-ESCA | 18.82 | 4.73 |
| GTEX-1CB4E-1026-SM-793AB | Esophagus - Mucosa | TCGA-ESCA | 35.59 | 11.48 |
| GTEX-1CB4F-1526-SM-7MKFM | Esophagus - Mucosa | TCGA-ESCA | 38.17 | 11.68 |
| GTEX-1CB4G-1326-SM-7MKFP | Esophagus - Mucosa | TCGA-ESCA | 27.62 | 8.18 |
| GTEX-1CB4H-1526-SM-7MXU4 | Esophagus - Mucosa | TCGA-ESCA | 23.52 | 6.89 |
| GTEX-1CB4I-1026-SM-7MKFZ | Esophagus - Mucosa | TCGA-ESCA | 48.06 | 13.01 |
| GTEX-1CB4J-2426-SM-7MXVC | Esophagus - Mucosa | TCGA-ESCA | 74.42 | 21.21 |
| GTEX-1E1VI-1526-SM-7IGQ2 | Esophagus - Mucosa | TCGA-ESCA | 27.76 | 7.18 |
| GTEX-1EH9U-2426-SM-7EWFG | Esophagus - Mucosa | TCGA-ESCA | 17.9 | 5.01 |
| GTEX-1EKGG-1726-SM-7MXUP | Esophagus - Mucosa | TCGA-ESCA | 24.89 | 6.98 |
| GTEX-1EMGI-1926-SM-7938Y | Esophagus - Mucosa | TCGA-ESCA | 35.11 | 13.75 |
| GTEX-1EN7A-1226-SM-7MGX9 | Esophagus - Mucosa | TCGA-ESCA | 21.24 | 6.98 |
| GTEX-1EU9M-2326-SM-7MKGS | Esophagus - Mucosa | TCGA-ESCA | 34.07 | 10.49 |
| GTEX-1EWIQ-1526-SM-79OO7 | Esophagus - Mucosa | TCGA-ESCA | 15.16 | 5.08 |
| GTEX-1EX96-2226-SM-7MKGT | Esophagus - Mucosa | TCGA-ESCA | 25.99 | 6.67 |

| | | | | |
|---|---|---|---|---|
| GTEX-1F52S-1426-SM-7P8PU | Esophagus - Mucosa | TCGA-ESCA | 29.54 | 12.58 |
| GTEX-1F5PK-1526-SM-7MKGV | Esophagus - Mucosa | TCGA-ESCA | 44.74 | 10.59 |
| GTEX-1F5PL-1726-SM-9QEGL | Esophagus - Mucosa | TCGA-ESCA | 84.94 | 25.24 |
| GTEX-1F75I-1726-SM-7MKGW | Esophagus - Mucosa | TCGA-ESCA | 40.19 | 10.78 |
| GTEX-1F7RK-1526-SM-7P8T6 | Esophagus - Mucosa | TCGA-ESCA | 29.33 | 8.98 |
| GTEX-1F88F-1126-SM-7MKHL | Esophagus - Mucosa | TCGA-ESCA | 35.84 | 8.51 |
| GTEX-1GF9U-0826-SM-7MXTZ | Esophagus - Mucosa | TCGA-ESCA | 59.2 | 27.38 |
| GTEX-1GL5R-1526-SM-7MKHO | Esophagus - Mucosa | TCGA-ESCA | 45.32 | 11.13 |
| GTEX-1GMR2-1526-SM-7MXV6 | Esophagus - Mucosa | TCGA-ESCA | 33.18 | 9.19 |
| GTEX-1GMR3-1526-SM-9JGGL | Esophagus - Mucosa | TCGA-ESCA | 25.98 | 10.36 |
| GTEX-1GMR8-1826-SM-9WYUE | Esophagus - Mucosa | TCGA-ESCA | 59.69 | 18.99 |
| GTEX-1GN1U-1826-SM-9JGH2 | Esophagus - Mucosa | TCGA-ESCA | 27.03 | 9.26 |
| GTEX-1GN1W-2226-SM-9JGI5 | Esophagus - Mucosa | TCGA-ESCA | 39.01 | 9.62 |
| GTEX-1GN73-1926-SM-9JGGS | Esophagus - Mucosa | TCGA-ESCA | 61.76 | 17.18 |
| GTEX-1GPI7-1026-SM-7MGY2 | Esophagus - Mucosa | TCGA-ESCA | 49.91 | 14.42 |
| GTEX-1GTWX-1326-SM-9JGHU | Esophagus - Mucosa | TCGA-ESCA | 38.24 | 13.32 |
| GTEX-1H11D-2126-SM-9OSXM | Esophagus - Mucosa | TCGA-ESCA | 56.31 | 22.91 |
| GTEX-1H1DE-1026-SM-9JGH6 | Esophagus - Mucosa | TCGA-ESCA | 61.58 | 22.15 |
| GTEX-1H1DG-1226-SM-9JGH3 | Esophagus - Mucosa | TCGA-ESCA | 27.6 | 10.87 |
| GTEX-1H1ZS-1526-SM-9OSXR | Esophagus - Mucosa | TCGA-ESCA | 38.43 | 12.01 |
| GTEX-1H23P-1426-SM-9JGH5 | Esophagus - Mucosa | TCGA-ESCA | 34.92 | 10.19 |
| GTEX-1H3NZ-1326-SM-9JGHI | Esophagus - Mucosa | TCGA-ESCA | 46.64 | 17.25 |
| GTEX-1H3VY-2026-SM-9JGHK | Esophagus - Mucosa | TCGA-ESCA | 65.54 | 28.36 |
| GTEX-1HB9E-1526-SM-D3LA6 | Esophagus - Mucosa | TCGA-ESCA | 32.89 | 12.29 |
| GTEX-1HBPH-1126-SM-A96TU | Esophagus - Mucosa | TCGA-ESCA | 25.86 | 6.44 |
| GTEX-1HBPI-0626-SM-9WPP4 | Esophagus - Mucosa | TCGA-ESCA | 25.06 | 6.54 |
| GTEX-1HBPM-1426-SM-9WPPE | Esophagus - Mucosa | TCGA-ESCA | 19.5 | 5.7 |
| GTEX-1HC8U-1526-SM-9WPO4 | Esophagus - Mucosa | TCGA-ESCA | 50.86 | 16.73 |
| GTEX-1HCU6-1326-SM-A96TZ | Esophagus - Mucosa | TCGA-ESCA | 55.66 | 11.99 |
| GTEX-1HCU7-2126-SM-A96SG | Esophagus - Mucosa | TCGA-ESCA | 37.56 | 14.49 |
| GTEX-1HCU8-2226-SM-9WPOV | Esophagus - Mucosa | TCGA-ESCA | 34.48 | 8.01 |

| | | | | |
|---|---|---|---|---|
| GTEX-1HCUA-1426-SM-9WPP7 | Esophagus - Mucosa | TCGA-ESCA | 41.93 | 11.52 |
| GTEX-1HCVE-1226-SM-9WPPH | Esophagus - Mucosa | TCGA-ESCA | 25.94 | 9 |
| GTEX-1HFI6-1226-SM-A96SK | Esophagus - Mucosa | TCGA-ESCA | 29.95 | 9.77 |
| GTEX-1HGF4-0926-SM-9WPP8 | Esophagus - Mucosa | TCGA-ESCA | 39.95 | 13.37 |
| GTEX-1HKZK-1426-SM-A96U1 | Esophagus - Mucosa | TCGA-ESCA | 44.43 | 14.13 |
| GTEX-1HSGN-1626-SM-A96U2 | Esophagus - Mucosa | TCGA-ESCA | 39.09 | 10.54 |
| GTEX-1HSKV-2426-SM-A8N8K | Esophagus - Mucosa | TCGA-ESCA | 26.18 | 7.12 |
| GTEX-1HSMQ-2026-SM-B2LY1 | Esophagus - Mucosa | TCGA-ESCA | 46.27 | 14.7 |
| GTEX-1I1CD-1726-SM-A9G3P | Esophagus - Mucosa | TCGA-ESCA | 28.31 | 8.09 |
| GTEX-1I1GP-1026-SM-C1YRZ | Esophagus - Mucosa | TCGA-ESCA | 87.27 | 21.34 |
| GTEX-1I1GQ-1726-SM-B2LWZ | Esophagus - Mucosa | TCGA-ESCA | 28.61 | 13.26 |
| GTEX-1I1GR-1326-SM-ARU7B | Esophagus - Mucosa | TCGA-ESCA | 20.94 | 8.35 |
| GTEX-1I1GS-2126-SM-ARU7G | Esophagus - Mucosa | TCGA-ESCA | 57.18 | 13.77 |
| GTEX-1I1GU-1426-SM-B2LX3 | Esophagus - Mucosa | TCGA-ESCA | 33.11 | 9.4 |
| GTEX-1I1GV-2226-SM-ARU7E | Esophagus - Mucosa | TCGA-ESCA | 31.37 | 7.03 |
| GTEX-1I4MK-0426-SM-B2LWQ | Esophagus - Mucosa | TCGA-ESCA | 50.42 | 16.3 |
| GTEX-1I6K7-1426-SM-B2LXP | Esophagus - Mucosa | TCGA-ESCA | 60.35 | 23.56 |
| GTEX-1ICG6-1926-SM-ARU7L | Esophagus - Mucosa | TCGA-ESCA | 38.71 | 11.83 |
| GTEX-1IDFM-1126-SM-A96SY | Esophagus - Mucosa | TCGA-ESCA | 18.29 | 7.42 |
| GTEX-1IDJD-1626-SM-C1YRC | Esophagus - Mucosa | TCGA-ESCA | 45.07 | 16.56 |
| GTEX-1IDJF-1226-SM-C1YPX | Esophagus - Mucosa | TCGA-ESCA | 40.79 | 14 |
| GTEX-1IDJH-2026-SM-CNNQX | Esophagus - Mucosa | TCGA-ESCA | 23.05 | 7.68 |
| GTEX-1IKK5-1326-SM-AHZ2X | Esophagus - Mucosa | TCGA-ESCA | 35.44 | 11.66 |
| GTEX-1IKOE-1526-SM-C1YSD | Esophagus - Mucosa | TCGA-ESCA | 42.08 | 13.79 |
| GTEX-1IY9M-1626-SM-AHZ39 | Esophagus - Mucosa | TCGA-ESCA | 43.41 | 14.22 |
| GTEX-1J1OQ-2526-SM-AHZ3Q | Esophagus - Mucosa | TCGA-ESCA | 41.05 | 13.54 |
| GTEX-1J1R8-0326-SM-AHZ3D | Esophagus - Mucosa | TCGA-ESCA | 31.87 | 7.32 |
| GTEX-1J8EW-1326-SM-AHZ3S | Esophagus - Mucosa | TCGA-ESCA | 32.57 | 12.04 |
| GTEX-1J8QM-1926-SM-AHZ3Z | Esophagus - Mucosa | TCGA-ESCA | 34.17 | 10.92 |
| GTEX-1JK1U-1726-SM-CNNPM | Esophagus - Mucosa | TCGA-ESCA | 43.13 | 15.74 |
| GTEX-1JKYN-1526-SM-CGQGA | Esophagus - Mucosa | TCGA-ESCA | 38.35 | 10.81 |

| | | | | |
|---|---|---|---|---|
| GTEX-1JKYR-2126-SM-CNNPC | Esophagus - Mucosa | TCGA-ESCA | 42.93 | 14.39 |
| GTEX-1JMLX-2326-SM-CXZJP | Esophagus - Mucosa | TCGA-ESCA | 29.31 | 7.07 |
| GTEX-1JMPY-0826-SM-AHZ4A | Esophagus - Mucosa | TCGA-ESCA | 31.47 | 5.54 |
| GTEX-1JMPZ-2026-SM-C1YR8 | Esophagus - Mucosa | TCGA-ESCA | 47.54 | 17.54 |
| GTEX-1JMQJ-1226-SM-D3L97 | Esophagus - Mucosa | TCGA-ESCA | 32.31 | 13.3 |
| GTEX-1JMQK-2026-SM-C1YQN | Esophagus - Mucosa | TCGA-ESCA | 58.77 | 20.86 |
| GTEX-1K2DA-0626-SM-CGQGO | Esophagus - Mucosa | TCGA-ESCA | 29.82 | 9.62 |
| GTEX-1K2DU-1726-SM-EV79S | Esophagus - Mucosa | TCGA-ESCA | 26.39 | 8.58 |
| GTEX-1K9T9-2026-SM-CXZK3 | Esophagus - Mucosa | TCGA-ESCA | 63 | 23.7 |
| GTEX-1KAFJ-0626-SM-D3L8X | Esophagus - Mucosa | TCGA-ESCA | 33.42 | 10.08 |
| GTEX-1KANA-0626-SM-D4P3I | Esophagus - Mucosa | TCGA-ESCA | 28.59 | 8.26 |
| GTEX-1KANB-1826-SM-DHXJU | Esophagus - Mucosa | TCGA-ESCA | 24.27 | 7.09 |
| GTEX-1KD4Q-1126-SM-CXZKK | Esophagus - Mucosa | TCGA-ESCA | 31.14 | 10.48 |
| GTEX-1KXAM-0826-SM-CXZK9 | Esophagus - Mucosa | TCGA-ESCA | 36.41 | 10.69 |
| GTEX-1L5NE-1426-SM-DHXKV | Esophagus - Mucosa | TCGA-ESCA | 36.61 | 11.18 |
| GTEX-1LB8K-1326-SM-CXZKB | Esophagus - Mucosa | TCGA-ESCA | 17.78 | 6.81 |
| GTEX-1LBAC-1226-SM-DHXK1 | Esophagus - Mucosa | TCGA-ESCA | 37.19 | 11.73 |
| GTEX-1LG7Y-1726-SM-EV7AR | Esophagus - Mucosa | TCGA-ESCA | 29.56 | 10.73 |
| GTEX-1LG7Z-0926-SM-CXKYK | Esophagus - Mucosa | TCGA-ESCA | 34.75 | 7.38 |
| GTEX-1LGOU-0526-SM-D3LAO | Esophagus - Mucosa | TCGA-ESCA | 39.88 | 12.96 |
| GTEX-1LGRB-1626-SM-CNNR7 | Esophagus - Mucosa | TCGA-ESCA | 38.51 | 10.35 |
| GTEX-1LNCM-0626-SM-EV7AI | Esophagus - Mucosa | TCGA-ESCA | 44.38 | 14.47 |
| GTEX-1LSNL-2226-SM-DHXKM | Esophagus - Mucosa | TCGA-ESCA | 26.64 | 8.59 |
| GTEX-1LSVX-1226-SM-DHXJN | Esophagus - Mucosa | TCGA-ESCA | 33.63 | 9.61 |
| GTEX-1LVA9-2126-SM-DLHBY | Esophagus - Mucosa | TCGA-ESCA | 13.91 | 4.09 |
| GTEX-1LVAM-1126-SM-DHXKP | Esophagus - Mucosa | TCGA-ESCA | 52.5 | 14.34 |
| GTEX-1LVAN-2326-SM-CNNPY | Esophagus - Mucosa | TCGA-ESCA | 36.88 | 11.59 |
| GTEX-1M5QR-0726-SM-DHXK7 | Esophagus - Mucosa | TCGA-ESCA | 56.76 | 22.11 |
| GTEX-1MA7W-0426-SM-E9TJ5 | Esophagus - Mucosa | TCGA-ESCA | 42.26 | 8.82 |
| GTEX-1MA7X-0726-SM-EV7AB | Esophagus - Mucosa | TCGA-ESCA | 23.74 | 8.76 |
| GTEX-1MCC2-1926-SM-DHXJR | Esophagus - Mucosa | TCGA-ESCA | 32.55 | 8 |

| | | | | |
|---|---|---|---|---|
| GTEX-1MCQQ-1026-SM-EV7AX | Esophagus - Mucosa | TCGA-ESCA | 10.29 | 3.03 |
| GTEX-1MCYP-1026-SM-DPRYB | Esophagus - Mucosa | TCGA-ESCA | 35.64 | 7.92 |
| GTEX-1MGNQ-1626-SM-EAZ4Z | Esophagus - Mucosa | TCGA-ESCA | 30.49 | 9.12 |
| GTEX-1MJIX-0826-SM-DTXEO | Esophagus - Mucosa | TCGA-ESCA | 34.72 | 8.19 |
| GTEX-1N2DV-2026-SM-EV7BL | Esophagus - Mucosa | TCGA-ESCA | 45.05 | 12.12 |
| GTEX-1N2DW-1426-SM-EV7AZ | Esophagus - Mucosa | TCGA-ESCA | 45.15 | 15.59 |
| GTEX-1N2EE-2426-SM-EXOIO | Esophagus - Mucosa | TCGA-ESCA | 39.03 | 11.89 |
| GTEX-1NSGN-2026-SM-DPRYM | Esophagus - Mucosa | TCGA-ESCA | 29.01 | 12.23 |
| GTEX-1NUQO-1126-SM-DTX98 | Esophagus - Mucosa | TCGA-ESCA | 34.03 | 9.44 |
| GTEX-1OJC3-0826-SM-DPRYU | Esophagus - Mucosa | TCGA-ESCA | 39.42 | 9.96 |
| GTEX-1OKEX-1326-SM-E9J2U | Esophagus - Mucosa | TCGA-ESCA | 41.13 | 13.53 |
| GTEX-1PFEY-0826-SM-E9U6A | Esophagus - Mucosa | TCGA-ESCA | 68.6 | 21.86 |
| GTEX-1PIGE-0926-SM-E9U5Z | Esophagus - Mucosa | TCGA-ESCA | 40.04 | 18.92 |
| GTEX-1PPGY-1726-SM-EVR3S | Esophagus - Mucosa | TCGA-ESCA | 29.59 | 9.23 |
| GTEX-1PPH8-1626-SM-E6CQP | Esophagus - Mucosa | TCGA-ESCA | 43.48 | 11.81 |
| GTEX-1QCLY-1526-SM-DPRXX | Esophagus - Mucosa | TCGA-ESCA | 40.1 | 10.44 |
| GTEX-1QCLZ-1626-SM-E9U5R | Esophagus - Mucosa | TCGA-ESCA | 46.48 | 15.75 |
| GTEX-1QEPI-1826-SM-EAZ3J | Esophagus - Mucosa | TCGA-ESCA | 27.05 | 8.25 |
| GTEX-1QP28-1826-SM-EAZ4L | Esophagus - Mucosa | TCGA-ESCA | 48.3 | 11.31 |
| GTEX-1QP29-2126-SM-E6CPZ | Esophagus - Mucosa | TCGA-ESCA | 57.95 | 17.17 |
| GTEX-1QP2A-1926-SM-EXOJ9 | Esophagus - Mucosa | TCGA-ESCA | 48.96 | 13.79 |
| GTEX-1QP67-0726-SM-DPRY1 | Esophagus - Mucosa | TCGA-ESCA | 55.03 | 14.99 |
| GTEX-1QP9N-0926-SM-E9U5S | Esophagus - Mucosa | TCGA-ESCA | 40.22 | 13.01 |
| GTEX-1QPFJ-1226-SM-EAZ3M | Esophagus - Mucosa | TCGA-ESCA | 36.14 | 12.35 |
| GTEX-1QW4Y-0526-SM-EAZ3V | Esophagus - Mucosa | TCGA-ESCA | 45.76 | 14.43 |
| GTEX-1R7EU-0826-SM-DPRZL | Esophagus - Mucosa | TCGA-ESCA | 48.55 | 13.17 |
| GTEX-1R9JW-1526-SM-DPRZK | Esophagus - Mucosa | TCGA-ESCA | 38.12 | 11.02 |
| GTEX-1R9K4-1926-SM-DPRZW | Esophagus - Mucosa | TCGA-ESCA | 51.16 | 18.68 |
| GTEX-1R9K5-1526-SM-DPRZA | Esophagus - Mucosa | TCGA-ESCA | 40.91 | 9.91 |
| GTEX-1R9PM-0926-SM-DPRYA | Esophagus - Mucosa | TCGA-ESCA | 34.45 | 9.63 |
| GTEX-1RAZA-1326-SM-DPS12 | Esophagus - Mucosa | TCGA-ESCA | 39.43 | 11.55 |

| | | | | |
|---|---|---|---|---|
| GTEX-1RAZQ-1726-SM-EAZ4T | Esophagus - Mucosa | TCGA-ESCA | 53.68 | 12.26 |
| GTEX-1RAZR-1226-SM-E6CRD | Esophagus - Mucosa | TCGA-ESCA | 52.32 | 18.45 |
| GTEX-1RAZS-2626-SM-EV79C | Esophagus - Mucosa | TCGA-ESCA | 23.31 | 8.74 |
| GTEX-1RDX4-2126-SM-EV7B3 | Esophagus - Mucosa | TCGA-ESCA | 21.95 | 7.68 |
| GTEX-1RLM8-0726-SM-E6CRH | Esophagus - Mucosa | TCGA-ESCA | 32.59 | 10 |
| GTEX-1S82Y-1226-SM-EV7BD | Esophagus - Mucosa | TCGA-ESCA | 32.21 | 10.89 |
| GTEX-1S83E-0826-SM-EV7B7 | Esophagus - Mucosa | TCGA-ESCA | 32.65 | 8.92 |
| GTEX-N7MS-1225-SM-3LK5G | Esophagus - Mucosa | TCGA-ESCA | 21.26 | 6.64 |
| GTEX-N7MT-0326-SM-48TDP | Esophagus - Mucosa | TCGA-ESCA | 29.49 | 9.52 |
| GTEX-NFK9-1126-SM-3LK78 | Esophagus - Mucosa | TCGA-ESCA | 36.46 | 9.76 |
| GTEX-NPJ8-1926-SM-3MJGB | Esophagus - Mucosa | TCGA-ESCA | 66.09 | 23.97 |
| GTEX-O5YV-1026-SM-3LK72 | Esophagus - Mucosa | TCGA-ESCA | 28.82 | 8.62 |
| GTEX-OHPN-1026-SM-E9TI8 | Esophagus - Mucosa | TCGA-ESCA | 37.43 | 11.92 |
| GTEX-OIZF-0626-SM-7MKHG | Esophagus - Mucosa | TCGA-ESCA | 39.05 | 11.46 |
| GTEX-OIZG-1026-SM-3LK5X | Esophagus - Mucosa | TCGA-ESCA | 26.57 | 7.76 |
| GTEX-OIZH-0626-SM-3NB1L | Esophagus - Mucosa | TCGA-ESCA | 39.9 | 9.8 |
| GTEX-OIZI-1426-SM-3NB1T | Esophagus - Mucosa | TCGA-ESCA | 37.14 | 12.49 |
| GTEX-OOBK-0625-SM-3LK5P | Esophagus - Mucosa | TCGA-ESCA | 37.94 | 13.36 |
| GTEX-OXRK-1326-SM-3NB1A | Esophagus - Mucosa | TCGA-ESCA | 47.82 | 14.69 |
| GTEX-OXRL-0626-SM-3NM9C | Esophagus - Mucosa | TCGA-ESCA | 52.33 | 18.85 |
| GTEX-OXRN-1726-SM-3NM9B | Esophagus - Mucosa | TCGA-ESCA | 24.1 | 6.4 |
| GTEX-OXRP-0926-SM-48TC1 | Esophagus - Mucosa | TCGA-ESCA | 30.27 | 10.89 |
| GTEX-P4PP-0626-SM-3NM9X | Esophagus - Mucosa | TCGA-ESCA | 33.78 | 8.5 |
| GTEX-P4PQ-0626-SM-3NMCU | Esophagus - Mucosa | TCGA-ESCA | 37.68 | 14.28 |
| GTEX-P4QS-0626-SM-3NMD1 | Esophagus - Mucosa | TCGA-ESCA | 47.49 | 16.05 |
| GTEX-P4QT-0626-SM-3NMCP | Esophagus - Mucosa | TCGA-ESCA | 30.05 | 6.04 |
| GTEX-P78B-1026-SM-3NMC4 | Esophagus - Mucosa | TCGA-ESCA | 59.67 | 18.89 |
| GTEX-PLZ4-1626-SM-3P618 | Esophagus - Mucosa | TCGA-ESCA | 32.04 | 12.09 |
| GTEX-PLZ5-0926-SM-3TW8L | Esophagus - Mucosa | TCGA-ESCA | 37.86 | 11.38 |
| GTEX-PLZ6-0526-SM-3P61C | Esophagus - Mucosa | TCGA-ESCA | 29.24 | 8.78 |
| GTEX-POYW-1326-SM-48TCG | Esophagus - Mucosa | TCGA-ESCA | 19.56 | 6.96 |

| | | | | |
|---|---|---|---|---|
| GTEX-PSDG-1326-SM-48TD2 | Esophagus - Mucosa | TCGA-ESCA | 40.04 | 17.73 |
| GTEX-PVOW-1626-SM-48TC9 | Esophagus - Mucosa | TCGA-ESCA | 39.51 | 13.13 |
| GTEX-PWCY-0726-SM-48TCS | Esophagus - Mucosa | TCGA-ESCA | 33.51 | 8.71 |
| GTEX-PWN1-0626-SM-48TDT | Esophagus - Mucosa | TCGA-ESCA | 29.1 | 9.49 |
| GTEX-Q2AG-1326-SM-43V8K | Esophagus - Mucosa | TCGA-ESCA | 19.53 | 3.18 |
| GTEX-Q2AH-0826-SM-48TZN | Esophagus - Mucosa | TCGA-ESCA | 39.98 | 15.29 |
| GTEX-Q2AI-0626-SM-48TZU | Esophagus - Mucosa | TCGA-ESCA | 33.99 | 10.43 |
| GTEX-QCQG-0626-SM-48U21 | Esophagus - Mucosa | TCGA-ESCA | 33.51 | 10.96 |
| GTEX-QDVJ-0726-SM-48U1W | Esophagus - Mucosa | TCGA-ESCA | 33.61 | 11.91 |
| GTEX-QDVN-1026-SM-48TZA | Esophagus - Mucosa | TCGA-ESCA | 43.12 | 12.29 |
| GTEX-QEG4-0726-SM-CNNRF | Esophagus - Mucosa | TCGA-ESCA | 49.74 | 15.83 |
| GTEX-QEG5-1426-SM-447AS | Esophagus - Mucosa | TCGA-ESCA | 52.41 | 22.96 |
| GTEX-QEL4-1426-SM-447AC | Esophagus - Mucosa | TCGA-ESCA | 48.15 | 13.63 |
| GTEX-QESD-0826-SM-47J4G | Esophagus - Mucosa | TCGA-ESCA | 42.8 | 14.31 |
| GTEX-QLQ7-0926-SM-447BC | Esophagus - Mucosa | TCGA-ESCA | 40.21 | 11.1 |
| GTEX-QLQW-0526-SM-4R1KS | Esophagus - Mucosa | TCGA-ESCA | 36.76 | 11.36 |
| GTEX-QV44-1126-SM-CL55C | Esophagus - Mucosa | TCGA-ESCA | 34.99 | 16.62 |
| GTEX-QVUS-0626-SM-447CC | Esophagus - Mucosa | TCGA-ESCA | 30.57 | 10.1 |
| GTEX-R3RS-1326-SM-48FE7 | Esophagus - Mucosa | TCGA-ESCA | 50.95 | 22.85 |
| GTEX-R45C-1226-SM-48FEE | Esophagus - Mucosa | TCGA-ESCA | 46.64 | 13.84 |
| GTEX-R53T-1126-SM-48FD4 | Esophagus - Mucosa | TCGA-ESCA | 35.3 | 7.99 |
| GTEX-R55C-0826-SM-48FCL | Esophagus - Mucosa | TCGA-ESCA | 33.77 | 6.68 |
| GTEX-R55D-1126-SM-48FEB | Esophagus - Mucosa | TCGA-ESCA | 49.4 | 14.27 |
| GTEX-R55E-1326-SM-48FCR | Esophagus - Mucosa | TCGA-ESCA | 51.94 | 20.33 |
| GTEX-R55G-1026-SM-48FDI | Esophagus - Mucosa | TCGA-ESCA | 31.02 | 9.13 |
| GTEX-REY6-1326-SM-48FDO | Esophagus - Mucosa | TCGA-ESCA | 34.9 | 14.46 |
| GTEX-RM2N-0726-SM-48FD5 | Esophagus - Mucosa | TCGA-ESCA | 42.19 | 13.19 |
| GTEX-RN64-1026-SM-48FDX | Esophagus - Mucosa | TCGA-ESCA | 26.86 | 10.14 |
| GTEX-RNOR-1226-SM-48FDQ | Esophagus - Mucosa | TCGA-ESCA | 42.22 | 15.64 |
| GTEX-RTLS-1126-SM-46MUQ | Esophagus - Mucosa | TCGA-ESCA | 29.66 | 9.72 |
| GTEX-RU1J-0326-SM-46MUM | Esophagus - Mucosa | TCGA-ESCA | 44.89 | 14.94 |

| | | | | |
|---|---|---|---|---|
| GTEX-RU72-0726-SM-46MUW | Esophagus - Mucosa | TCGA-ESCA | 61.92 | 26.07 |
| GTEX-RUSQ-0826-SM-47JWW | Esophagus - Mucosa | TCGA-ESCA | 86.07 | 39.49 |
| GTEX-RVPV-0526-SM-47JYL | Esophagus - Mucosa | TCGA-ESCA | 22.94 | 7.69 |
| GTEX-RWS6-0826-SM-47JXF | Esophagus - Mucosa | TCGA-ESCA | 25.72 | 9.9 |
| GTEX-S32W-0926-SM-4AD5X | Esophagus - Mucosa | TCGA-ESCA | 49.26 | 17.59 |
| GTEX-S33H-0826-SM-4AD5Y | Esophagus - Mucosa | TCGA-ESCA | 33.91 | 11.08 |
| GTEX-S341-0526-SM-4AD5U | Esophagus - Mucosa | TCGA-ESCA | 36.08 | 13.14 |
| GTEX-S3XE-0926-SM-4AD4S | Esophagus - Mucosa | TCGA-ESCA | 41.56 | 10.63 |
| GTEX-S4P3-0426-SM-4AD56 | Esophagus - Mucosa | TCGA-ESCA | 41.91 | 13.73 |
| GTEX-S4Q7-0626-SM-4AD5M | Esophagus - Mucosa | TCGA-ESCA | 28.54 | 4.62 |
| GTEX-S7SF-0926-SM-4AD4M | Esophagus - Mucosa | TCGA-ESCA | 33.49 | 14.73 |
| GTEX-S95S-0426-SM-4B64I | Esophagus - Mucosa | TCGA-ESCA | 29.4 | 9.29 |
| GTEX-SE5C-1226-SM-4BRWV | Esophagus - Mucosa | TCGA-ESCA | 27.64 | 8.09 |
| GTEX-SJXC-1426-SM-EYYVB | Esophagus - Mucosa | TCGA-ESCA | 38.79 | 11.81 |
| GTEX-SNMC-0526-SM-4DM69 | Esophagus - Mucosa | TCGA-ESCA | 35.33 | 11.6 |
| GTEX-SNOS-0626-SM-4DM5F | Esophagus - Mucosa | TCGA-ESCA | 40.67 | 8.87 |
| GTEX-SUCS-0726-SM-4DM7J | Esophagus - Mucosa | TCGA-ESCA | 30.2 | 10.01 |
| GTEX-T2IS-0826-SM-4DM6L | Esophagus - Mucosa | TCGA-ESCA | 38.98 | 13.14 |
| GTEX-T2YK-1326-SM-4DM6S | Esophagus - Mucosa | TCGA-ESCA | 23.13 | 6.71 |
| GTEX-T5JC-1726-SM-4DM6U | Esophagus - Mucosa | TCGA-ESCA | 30.84 | 12.37 |
| GTEX-T5JW-1326-SM-4DM5G | Esophagus - Mucosa | TCGA-ESCA | 58.82 | 18.07 |
| GTEX-T6MO-0526-SM-4DM6R | Esophagus - Mucosa | TCGA-ESCA | 29.74 | 7.29 |
| GTEX-T8EM-0526-SM-4DM64 | Esophagus - Mucosa | TCGA-ESCA | 32.61 | 10.81 |
| GTEX-TKQ1-0326-SM-4DXSM | Esophagus - Mucosa | TCGA-ESCA | 36.94 | 11.36 |
| GTEX-TML8-1526-SM-4DXUX | Esophagus - Mucosa | TCGA-ESCA | 32.82 | 6.35 |
| GTEX-TMMY-1126-SM-4DXSX | Esophagus - Mucosa | TCGA-ESCA | 76.39 | 23.97 |
| GTEX-TSE9-1226-SM-4DXTM | Esophagus - Mucosa | TCGA-ESCA | 53.23 | 17.71 |
| GTEX-U3ZH-1026-SM-4DXUZ | Esophagus - Mucosa | TCGA-ESCA | 34.38 | 12.67 |
| GTEX-U3ZM-0526-SM-4DXTB | Esophagus - Mucosa | TCGA-ESCA | 36.06 | 11.91 |
| GTEX-U3ZN-1726-SM-4DXUQ | Esophagus - Mucosa | TCGA-ESCA | 34.76 | 12.25 |
| GTEX-U4B1-0826-SM-4DXTW | Esophagus - Mucosa | TCGA-ESCA | 38.6 | 9.76 |

| | | | | |
|---|---|---|---|---|
| GTEX-U8XE-1726-SM-4E3IF | Esophagus - Mucosa | TCGA-ESCA | 42.17 | 12.02 |
| GTEX-UJHI-0826-SM-4IHLM | Esophagus - Mucosa | TCGA-ESCA | 35.89 | 9.74 |
| GTEX-UJMC-0926-SM-4IHLK | Esophagus - Mucosa | TCGA-ESCA | 28.3 | 9.64 |
| GTEX-UPJH-1026-SM-EAZB2 | Esophagus - Mucosa | TCGA-ESCA | 46.93 | 18.6 |
| GTEX-UPK5-1226-SM-4IHL8 | Esophagus - Mucosa | TCGA-ESCA | 45.09 | 14.44 |
| GTEX-V1D1-1026-SM-4JBHE | Esophagus - Mucosa | TCGA-ESCA | 28.96 | 10.13 |
| GTEX-V955-1126-SM-4JBH3 | Esophagus - Mucosa | TCGA-ESCA | 43.11 | 15.9 |
| GTEX-VJYA-0726-SM-4KL1T | Esophagus - Mucosa | TCGA-ESCA | 39.4 | 15.07 |
| GTEX-VUSG-1226-SM-4KKZF | Esophagus - Mucosa | TCGA-ESCA | 32.13 | 10.37 |
| GTEX-W5WG-1626-SM-4LMKG | Esophagus - Mucosa | TCGA-ESCA | 39.09 | 12.11 |
| GTEX-WEY5-0926-SM-4LMI4 | Esophagus - Mucosa | TCGA-ESCA | 39.36 | 10.64 |
| GTEX-WFG7-1126-SM-4LMK3 | Esophagus - Mucosa | TCGA-ESCA | 49.94 | 13.1 |
| GTEX-WFG8-1226-SM-4V6EI | Esophagus - Mucosa | TCGA-ESCA | 39.83 | 9.56 |
| GTEX-WFJO-0826-SM-4LVM5 | Esophagus - Mucosa | TCGA-ESCA | 44.52 | 16.09 |
| GTEX-WFON-0826-SM-4LVMI | Esophagus - Mucosa | TCGA-ESCA | 35.79 | 7.81 |
| GTEX-WH7G-0926-SM-4LVMJ | Esophagus - Mucosa | TCGA-ESCA | 33.91 | 6.46 |
| GTEX-WHPG-1126-SM-4M1XT | Esophagus - Mucosa | TCGA-ESCA | 32.48 | 10.56 |
| GTEX-WHSB-0426-SM-4M1XI | Esophagus - Mucosa | TCGA-ESCA | 31.1 | 9.46 |
| GTEX-WHSE-1526-SM-4M1XH | Esophagus - Mucosa | TCGA-ESCA | 37.2 | 13.15 |
| GTEX-WHWD-0926-SM-4OORY | Esophagus - Mucosa | TCGA-ESCA | 71.48 | 25.47 |
| GTEX-WK11-1026-SM-4OOS8 | Esophagus - Mucosa | TCGA-ESCA | 37.01 | 12.23 |
| GTEX-WOFM-0626-SM-4OOS3 | Esophagus - Mucosa | TCGA-ESCA | 41.48 | 12.95 |
| GTEX-WQUQ-1626-SM-4OOSH | Esophagus - Mucosa | TCGA-ESCA | 47.52 | 14.57 |
| GTEX-WRHK-0526-SM-4MVOF | Esophagus - Mucosa | TCGA-ESCA | 40.95 | 9.64 |
| GTEX-WVLH-1426-SM-4MVNX | Esophagus - Mucosa | TCGA-ESCA | 25.85 | 7.95 |
| GTEX-WXYG-1226-SM-4ONCJ | Esophagus - Mucosa | TCGA-ESCA | 40.66 | 11.52 |
| GTEX-WYJK-1926-SM-4ONDP | Esophagus - Mucosa | TCGA-ESCA | 44.85 | 9.72 |
| GTEX-WYVS-0726-SM-4ONDO | Esophagus - Mucosa | TCGA-ESCA | 30.82 | 9.38 |
| GTEX-X15G-0826-SM-4PQZD | Esophagus - Mucosa | TCGA-ESCA | 36.32 | 14.98 |
| GTEX-X3Y1-1326-SM-4PQZB | Esophagus - Mucosa | TCGA-ESCA | 31.38 | 8.81 |
| GTEX-X4EO-1226-SM-4QARR | Esophagus - Mucosa | TCGA-ESCA | 41.58 | 14.23 |

| | | | | |
|---|---|---|---|---|
| GTEX-X4EP-1626-SM-4QAS6 | Esophagus - Mucosa | TCGA-ESCA | 31.38 | 11.29 |
| GTEX-X4LF-0626-SM-4QASB | Esophagus - Mucosa | TCGA-ESCA | 42.03 | 8.46 |
| GTEX-X4XY-1726-SM-46MVO | Esophagus - Mucosa | TCGA-ESCA | 22.37 | 8.55 |
| GTEX-X585-1226-SM-46MW7 | Esophagus - Mucosa | TCGA-ESCA | 29.06 | 9.82 |
| GTEX-X62O-2226-SM-46MW3 | Esophagus - Mucosa | TCGA-ESCA | 20.8 | 7.55 |
| GTEX-XBED-1026-SM-48TCB | Esophagus - Mucosa | TCGA-ESCA | 31.34 | 9.64 |
| GTEX-XBEW-1326-SM-4AT63 | Esophagus - Mucosa | TCGA-ESCA | 34.71 | 6.09 |
| GTEX-XGQ4-1526-SM-4AT6H | Esophagus - Mucosa | TCGA-ESCA | 56.37 | 20.43 |
| GTEX-XK95-1026-SM-4GIDV | Esophagus - Mucosa | TCGA-ESCA | 27.52 | 10.3 |
| GTEX-XLM4-2326-SM-4AT5E | Esophagus - Mucosa | TCGA-ESCA | 67.86 | 15.63 |
| GTEX-XMD2-1326-SM-4YCET | Esophagus - Mucosa | TCGA-ESCA | 39.95 | 12.91 |
| GTEX-XMK1-1326-SM-4B65Q | Esophagus - Mucosa | TCGA-ESCA | 51.13 | 17.53 |
| GTEX-XQ3S-1126-SM-4BOPK | Esophagus - Mucosa | TCGA-ESCA | 38.22 | 10.71 |
| GTEX-XQ8I-1526-SM-4BOOH | Esophagus - Mucosa | TCGA-ESCA | 31.51 | 11.06 |
| GTEX-XUZC-0426-SM-4BOPE | Esophagus - Mucosa | TCGA-ESCA | 38.13 | 11.4 |
| GTEX-XV7Q-1126-SM-4BRVS | Esophagus - Mucosa | TCGA-ESCA | 37.84 | 11.06 |
| GTEX-XXEK-0326-SM-4BRVV | Esophagus - Mucosa | TCGA-ESCA | 36.48 | 13.99 |
| GTEX-XYKS-1526-SM-4BRUP | Esophagus - Mucosa | TCGA-ESCA | 36.46 | 12.25 |
| GTEX-Y111-1626-SM-57WCY | Esophagus - Mucosa | TCGA-ESCA | 73.69 | 21.78 |
| GTEX-Y114-0926-SM-4TT7J | Esophagus - Mucosa | TCGA-ESCA | 38.03 | 11.3 |
| GTEX-Y3I4-1226-SM-4TT8N | Esophagus - Mucosa | TCGA-ESCA | 45.7 | 15.99 |
| GTEX-Y3IK-1126-SM-51MSV | Esophagus - Mucosa | TCGA-ESCA | 32.92 | 10.57 |
| GTEX-Y5LM-1026-SM-4VDSQ | Esophagus - Mucosa | TCGA-ESCA | 36.05 | 10.78 |
| GTEX-Y5V5-1626-SM-4VDSG | Esophagus - Mucosa | TCGA-ESCA | 40.99 | 12.6 |
| GTEX-Y5V6-1026-SM-4VBS2 | Esophagus - Mucosa | TCGA-ESCA | 44.26 | 15.43 |
| GTEX-Y8E4-1726-SM-5LU9G | Esophagus - Mucosa | TCGA-ESCA | 48.49 | 16.78 |
| GTEX-Y8LW-0726-SM-5IFI1 | Esophagus - Mucosa | TCGA-ESCA | 52.11 | 13.1 |
| GTEX-Y9LG-0626-SM-4WWDH | Esophagus - Mucosa | TCGA-ESCA | 42.77 | 7.53 |
| GTEX-YB5E-0926-SM-4WWDQ | Esophagus - Mucosa | TCGA-ESCA | 32.37 | 6.87 |
| GTEX-YB5K-0826-SM-5YYB7 | Esophagus - Mucosa | TCGA-ESCA | 37.67 | 8.73 |
| GTEX-YEC3-2526-SM-CXZIH | Esophagus - Mucosa | TCGA-ESCA | 40.28 | 15.25 |

| | | | | |
|---|---|---|---|---|
| GTEX-YEC4-1026-SM-9WG7N | Esophagus - Mucosa | TCGA-ESCA | 25.02 | 5.68 |
| GTEX-YECK-1626-SM-5IFHQ | Esophagus - Mucosa | TCGA-ESCA | 30.92 | 10.08 |
| GTEX-YF7O-0926-SM-5IFIF | Esophagus - Mucosa | TCGA-ESCA | 42.36 | 12.33 |
| GTEX-YFC4-1226-SM-5LU9R | Esophagus - Mucosa | TCGA-ESCA | 52.69 | 19.42 |
| GTEX-YJ89-1526-SM-5P9IR | Esophagus - Mucosa | TCGA-ESCA | 26.43 | 7.49 |
| GTEX-Z9EW-0526-SM-5HL8S | Esophagus - Mucosa | TCGA-ESCA | 40.19 | 11.51 |
| GTEX-ZA64-0626-SM-5HL9O | Esophagus - Mucosa | TCGA-ESCA | 38.18 | 10.12 |
| GTEX-ZAB4-1126-SM-5HL86 | Esophagus - Mucosa | TCGA-ESCA | 41.41 | 12.9 |
| GTEX-ZAB5-0926-SM-5CVMV | Esophagus - Mucosa | TCGA-ESCA | 44.68 | 18.66 |
| GTEX-ZAJG-1326-SM-5S2NV | Esophagus - Mucosa | TCGA-ESCA | 74.76 | 20.38 |
| GTEX-ZAK1-1526-SM-5S2OV | Esophagus - Mucosa | TCGA-ESCA | 26.65 | 6.64 |
| GTEX-ZC5H-1326-SM-5HL7E | Esophagus - Mucosa | TCGA-ESCA | 26.22 | 7.49 |
| GTEX-ZDTT-1326-SM-4WKFH | Esophagus - Mucosa | TCGA-ESCA | 32.51 | 9.3 |
| GTEX-ZDYS-0826-SM-4WWCR | Esophagus - Mucosa | TCGA-ESCA | 48.35 | 14.5 |
| GTEX-ZE9C-1126-SM-4WWBD | Esophagus - Mucosa | TCGA-ESCA | 37.08 | 10.35 |
| GTEX-ZEX8-1426-SM-4WKHQ | Esophagus - Mucosa | TCGA-ESCA | 51.99 | 17.02 |
| GTEX-ZF29-1326-SM-DNZYV | Esophagus - Mucosa | TCGA-ESCA | 32.14 | 6.62 |
| GTEX-ZF2S-1226-SM-4WWAX | Esophagus - Mucosa | TCGA-ESCA | 31.96 | 10.16 |
| GTEX-ZF3C-1626-SM-4WWB4 | Esophagus - Mucosa | TCGA-ESCA | 32.46 | 11.99 |
| GTEX-ZG7Y-1126-SM-4WWEZ | Esophagus - Mucosa | TCGA-ESCA | 34.49 | 10.6 |
| GTEX-ZGAY-1126-SM-4WWBS | Esophagus - Mucosa | TCGA-ESCA | 37.34 | 10.69 |
| GTEX-ZLFU-1126-SM-DO11K | Esophagus - Mucosa | TCGA-ESCA | 41.45 | 12.68 |
| GTEX-ZLV1-0526-SM-4WWD4 | Esophagus - Mucosa | TCGA-ESCA | 31.77 | 6.95 |
| GTEX-ZLWG-1326-SM-DNZYZ | Esophagus - Mucosa | TCGA-ESCA | 30.75 | 8.58 |
| GTEX-ZPCL-1426-SM-57WEC | Esophagus - Mucosa | TCGA-ESCA | 36.39 | 8.39 |
| GTEX-ZPU1-1426-SM-4WWFU | Esophagus - Mucosa | TCGA-ESCA | 44.26 | 13.71 |
| GTEX-ZQUD-0426-SM-57WFR | Esophagus - Mucosa | TCGA-ESCA | 29.12 | 8.67 |
| GTEX-ZT9W-1326-SM-51MTB | Esophagus - Mucosa | TCGA-ESCA | 35 | 6.58 |
| GTEX-ZT9X-0526-SM-5GCOP | Esophagus - Mucosa | TCGA-ESCA | 54.19 | 16.71 |
| GTEX-ZTPG-1626-SM-5DUX2 | Esophagus - Mucosa | TCGA-ESCA | 50.03 | 14.92 |
| GTEX-ZTSS-0626-SM-59867 | Esophagus - Mucosa | TCGA-ESCA | 32.45 | 7.16 |

| | | | | |
|---|---|---|---|---|
| GTEX-ZTX8-0226-SM-5E44D | Esophagus - Mucosa | TCGA-ESCA | 67.12 | 22.39 |
| GTEX-ZUA1-1226-SM-5NQ6O | Esophagus - Mucosa | TCGA-ESCA | 35.63 | 12.67 |
| GTEX-ZV68-0926-SM-59HK7 | Esophagus - Mucosa | TCGA-ESCA | 22.72 | 7.78 |
| GTEX-ZVE2-1326-SM-5NQ94 | Esophagus - Mucosa | TCGA-ESCA | 52.03 | 14.61 |
| GTEX-ZVP2-1326-SM-57WCE | Esophagus - Mucosa | TCGA-ESCA | 35.85 | 6.92 |
| GTEX-ZVT2-1326-SM-5E45H | Esophagus - Mucosa | TCGA-ESCA | 34.63 | 7.65 |
| GTEX-ZXES-1126-SM-5GZXN | Esophagus - Mucosa | TCGA-ESCA | 44.67 | 11.78 |
| GTEX-ZY6K-0626-SM-59HJL | Esophagus - Mucosa | TCGA-ESCA | 55.85 | 16.27 |
| GTEX-ZYFC-1626-SM-5N9EU | Esophagus - Mucosa | TCGA-ESCA | 44.26 | 15.76 |
| GTEX-ZYFG-1126-SM-5GICY | Esophagus - Mucosa | TCGA-ESCA | 42.14 | 9.51 |
| GTEX-ZYT6-1126-SM-5GZYR | Esophagus - Mucosa | TCGA-ESCA | 54.29 | 13.37 |
| GTEX-ZYVF-2226-SM-5GIDD | Esophagus - Mucosa | TCGA-ESCA | 33.29 | 12.99 |
| GTEX-ZYW4-1326-SM-5E45N | Esophagus - Mucosa | TCGA-ESCA | 44.68 | 10.99 |
| GTEX-ZYY3-1526-SM-5SIA9 | Esophagus - Mucosa | TCGA-ESCA | 52.37 | 16.27 |
| GTEX-ZZ64-0226-SM-5E44X | Esophagus - Mucosa | TCGA-ESCA | 49.5 | 11.52 |
| GTEX-ZZPT-1426-SM-5N9C5 | Esophagus - Mucosa | TCGA-ESCA | 46.65 | 11.08 |
| GTEX-11GS4-2326-SM-5A5KS | Kidney - Cortex | TCGA-KIRP | 23.38 | 6.38 |
| GTEX-11OF3-1326-SM-5N9FJ | Kidney - Cortex | TCGA-KIRP | 16.1 | 4.92 |
| GTEX-11TTK-1926-SM-5PNW8 | Kidney - Cortex | TCGA-KIRP | 47.62 | 9.18 |
| GTEX-12696-0926-SM-5FQTV | Kidney - Cortex | TCGA-KIRP | 44.82 | 7.09 |
| GTEX-12WSG-0826-SM-5EQ5A | Kidney - Cortex | TCGA-KIRP | 29.56 | 6.51 |
| GTEX-13112-2126-SM-5GCO4 | Kidney - Cortex | TCGA-KIRP | 38.91 | 6.7 |
| GTEX-1399S-0526-SM-5IJG8 | Kidney - Cortex | TCGA-KIRP | 38.29 | 8.64 |
| GTEX-13NYB-1726-SM-5N9G2 | Kidney - Cortex | TCGA-KIRP | 51.12 | 12.42 |
| GTEX-13O1R-2526-SM-5N9FW | Kidney - Cortex | TCGA-KIRP | 30.56 | 6.01 |
| GTEX-13OVI-1126-SM-5KLZF | Kidney - Cortex | TCGA-KIRP | 37.32 | 7.05 |
| GTEX-13OVL-1826-SM-5KLZR | Kidney - Cortex | TCGA-KIRP | 26.5 | 6.43 |
| GTEX-13OW6-1826-SM-5N9F9 | Kidney - Cortex | TCGA-KIRP | 16.23 | 4.77 |
| GTEX-13RTJ-2226-SM-5S2Q1 | Kidney - Cortex | TCGA-KIRP | 33.36 | 8.09 |
| GTEX-145MN-0326-SM-5QGQI | Kidney - Cortex | TCGA-KIRP | 42.6 | 10.72 |
| GTEX-147F4-2626-SM-5Q5CS | Kidney - Cortex | TCGA-KIRP | 39.34 | 10.61 |

| | | | | |
|---|---|---|---|---|
| GTEX-1497J-0826-SM-5NQAJ | Kidney - Cortex | TCGA-KIRP | 27.27 | 6.84 |
| GTEX-14C39-2126-SM-664OH | Kidney - Cortex | TCGA-KIRP | 21.26 | 7.5 |
| GTEX-14C5O-2026-SM-5YYB1 | Kidney - Cortex | TCGA-KIRP | 19.93 | 4.84 |
| GTEX-14E6D-2526-SM-5YYA9 | Kidney - Cortex | TCGA-KIRP | 35.2 | 10.01 |
| GTEX-15CHQ-2126-SM-6871M | Kidney - Cortex | TCGA-KIRP | 28.53 | 8.19 |
| GTEX-15DYW-2026-SM-6AJBD | Kidney - Cortex | TCGA-KIRP | 14.07 | 2.51 |
| GTEX-16MT8-1926-SM-7EPGL | Kidney - Cortex | TCGA-KIRP | 20.46 | 4.37 |
| GTEX-17F96-1926-SM-7MGWY | Kidney - Cortex | TCGA-KIRP | 22.02 | 5.89 |
| GTEX-17F97-1926-SM-7IGM4 | Kidney - Cortex | TCGA-KIRP | 58.8 | 14.17 |
| GTEX-17HHY-2326-SM-7KFSR | Kidney - Cortex | TCGA-KIRP | 18.71 | 4.51 |
| GTEX-1A3MW-2226-SM-73KUX | Kidney - Cortex | TCGA-KIRP | 26.26 | 10.51 |
| GTEX-1A8FM-2026-SM-7P8RN | Kidney - Cortex | TCGA-KIRP | 16.19 | 4.96 |
| GTEX-1AX8Z-2326-SM-731CR | Kidney - Cortex | TCGA-KIRP | 41.34 | 15.21 |
| GTEX-1BAJH-1826-SM-731DA | Kidney - Cortex | TCGA-KIRP | 23.34 | 4.01 |
| GTEX-1C6VQ-2326-SM-7IGN1 | Kidney - Cortex | TCGA-KIRP | 23.5 | 6.65 |
| GTEX-1CB4J-1926-SM-7MGX8 | Kidney - Cortex | TCGA-KIRP | 32.47 | 9.11 |
| GTEX-1E2YA-1626-SM-7MGXS | Kidney - Cortex | TCGA-KIRP | 12.78 | 4.3 |
| GTEX-1EKGG-2226-SM-7IGOA | Kidney - Cortex | TCGA-KIRP | 21.1 | 4.64 |
| GTEX-1F48J-2226-SM-7MXTN | Kidney - Cortex | TCGA-KIRP | 20.18 | 5.36 |
| GTEX-1GF9V-1926-SM-9JGHE | Kidney - Cortex | TCGA-KIRP | 18.46 | 4.35 |
| GTEX-1GF9W-1326-SM-7P8PX | Kidney - Cortex | TCGA-KIRP | 28.05 | 8.05 |
| GTEX-1GMR3-2426-SM-7MXV2 | Kidney - Cortex | TCGA-KIRP | 14.1 | 4.16 |
| GTEX-1GN1U-2126-SM-AHZ4J | Kidney - Cortex | TCGA-KIRP | 12.58 | 2.8 |
| GTEX-1GTWX-1826-SM-9QEHG | Kidney - Cortex | TCGA-KIRP | 24.97 | 7.08 |
| GTEX-1H3NZ-2126-SM-9WPQ6 | Kidney - Cortex | TCGA-KIRP | 15.66 | 4.64 |
| GTEX-1H3VY-1926-SM-9JGHJ | Kidney - Cortex | TCGA-KIRP | 29.61 | 8.76 |
| GTEX-1HB9E-1826-SM-D4P33 | Kidney - Cortex | TCGA-KIRP | 36.26 | 11.04 |
| GTEX-1HBPI-2026-SM-CE6RZ | Kidney - Cortex | TCGA-KIRP | 14.37 | 4.3 |
| GTEX-1HCU7-1826-SM-A9G2A | Kidney - Cortex | TCGA-KIRP | 15.38 | 4.08 |
| GTEX-1I1GP-1926-SM-CNNQU | Kidney - Cortex | TCGA-KIRP | 19.44 | 5.22 |
| GTEX-1IKK5-0926-SM-A9G2U | Kidney - Cortex | TCGA-KIRP | 28.15 | 5.81 |

| | | | | |
|---|---|---|---|---|
| GTEX-1IL2V-1826-SM-ARU7R | Kidney - Cortex | TCGA-KIRP | 29.93 | 9.89 |
| GTEX-1J1OQ-1426-SM-AHZ3P | Kidney - Cortex | TCGA-KIRP | 30.05 | 7.28 |
| GTEX-1JJ6O-2526-SM-AHZ3J | Kidney - Cortex | TCGA-KIRP | 10.06 | 3.09 |
| GTEX-1JMLX-1526-SM-AHZ3N | Kidney - Cortex | TCGA-KIRP | 14.78 | 3.29 |
| GTEX-1JMPZ-1226-SM-ARU8Y | Kidney - Cortex | TCGA-KIRP | 40.91 | 12.17 |
| GTEX-1JMQJ-1726-SM-E9U4V | Kidney - Cortex | TCGA-KIRP | 38.19 | 9.03 |
| GTEX-1JMQK-1926-SM-CJI3B | Kidney - Cortex | TCGA-KIRP | 30.16 | 7.45 |
| GTEX-1K9T9-1826-SM-CXZK2 | Kidney - Cortex | TCGA-KIRP | 37.04 | 12.69 |
| GTEX-1KWVE-0826-SM-CXZKO | Kidney - Cortex | TCGA-KIRP | 22.27 | 4.81 |
| GTEX-1LG7Y-2726-SM-D5OVW | Kidney - Cortex | TCGA-KIRP | 30.53 | 6.04 |
| GTEX-1MUQO-2526-SM-E9TJN | Kidney - Cortex | TCGA-KIRP | 48.89 | 9.52 |
| GTEX-1N2DW-1826-SM-EAZ54 | Kidney - Cortex | TCGA-KIRP | 52.91 | 16.18 |
| GTEX-1N2EF-1426-SM-EXUSH | Kidney - Cortex | TCGA-KIRP | 25.99 | 4.35 |
| GTEX-1NV8Z-2226-SM-EXOJ2 | Kidney - Cortex | TCGA-KIRP | 26.98 | 6.89 |
| GTEX-1OJC4-2326-SM-DTXA5 | Kidney - Cortex | TCGA-KIRP | 33.47 | 10.65 |
| GTEX-1PWST-2226-SM-EVR3G | Kidney - Cortex | TCGA-KIRP | 36.03 | 9.07 |
| GTEX-1QP6S-1526-SM-EXUSG | Kidney - Cortex | TCGA-KIRP | 22.64 | 6.22 |
| GTEX-1R9PO-1626-SM-E6CPT | Kidney - Cortex | TCGA-KIRP | 58.54 | 15.17 |
| GTEX-NPJ8-2226-SM-3TW8D | Kidney - Cortex | TCGA-KIRP | 20.56 | 8.19 |
| GTEX-O5YV-1226-SM-EV795 | Kidney - Cortex | TCGA-KIRP | 21.56 | 5.99 |
| GTEX-P4QS-1126-SM-3NMD5 | Kidney - Cortex | TCGA-KIRP | 21.42 | 8.37 |
| GTEX-QDT8-1726-SM-EYYV6 | Kidney - Cortex | TCGA-KIRP | 34.91 | 4.81 |
| GTEX-QDVN-1626-SM-48TZC | Kidney - Cortex | TCGA-KIRP | 20.05 | 5.73 |
| GTEX-QLQW-1626-SM-CMKFE | Kidney - Cortex | TCGA-KIRP | 38.27 | 12.72 |
| GTEX-REY6-1826-SM-EAZAT | Kidney - Cortex | TCGA-KIRP | 14.28 | 3.64 |
| GTEX-RN64-1626-SM-48FD7 | Kidney - Cortex | TCGA-KIRP | 42.1 | 12.68 |
| GTEX-S33H-1426-SM-EZ6L7 | Kidney - Cortex | TCGA-KIRP | 19.41 | 4.94 |
| GTEX-T5JC-1526-SM-4DM68 | Kidney - Cortex | TCGA-KIRP | 23.07 | 5.1 |
| GTEX-T6MN-1826-SM-EXOHN | Kidney - Cortex | TCGA-KIRP | 42.39 | 9.91 |
| GTEX-XPVG-0526-SM-4B65N | Kidney - Cortex | TCGA-KIRP | 26.58 | 10.31 |
| GTEX-Y5V5-1226-SM-EYYVP | Kidney - Cortex | TCGA-KIRP | 18.15 | 4.49 |

| | | | | |
|---|---|---|---|---|
| GTEX-Y5V6-2026-SM-5IFHO | Kidney - Cortex | TCGA-KIRP | 17.37 | 4.96 |
| GTEX-ZC5H-1726-SM-5HL7X | Kidney - Cortex | TCGA-KIRP | 22.43 | 4.95 |
| GTEX-ZDXO-0226-SM-4WKH7 | Kidney - Cortex | TCGA-KIRP | 15.71 | 3.44 |
| GTEX-ZE9C-1426-SM-4WKGM | Kidney - Cortex | TCGA-KIRP | 46.49 | 16.61 |
| GTEX-ZLFU-0926-SM-5P9F8 | Kidney - Cortex | TCGA-KIRP | 38.75 | 8.3 |
| GTEX-ZVZP-0926-SM-5GIDB | Kidney - Cortex | TCGA-KIRP | 30.86 | 6.43 |
| GTEX-ZYFG-1626-SM-5GZYY | Kidney - Cortex | TCGA-KIRP | 35.83 | 7.77 |
| GTEX-ZYT6-2226-SM-5GIC9 | Kidney - Cortex | TCGA-KIRP | 44.09 | 8.73 |
| GTEX-11DXY-0526-SM-5EGGQ | Liver | TCGA-LIHC | 18.84 | 5.76 |
| GTEX-11DXZ-0126-SM-5EGGY | Liver | TCGA-LIHC | 18.14 | 7.67 |
| GTEX-11EQ9-0526-SM-5A5JZ | Liver | TCGA-LIHC | 24.58 | 6.51 |
| GTEX-11GSP-0626-SM-5986T | Liver | TCGA-LIHC | 11.2 | 5.23 |
| GTEX-11NUK-1226-SM-5P9GM | Liver | TCGA-LIHC | 11.64 | 4.09 |
| GTEX-11NV4-1326-SM-5HL6V | Liver | TCGA-LIHC | 19.46 | 4.69 |
| GTEX-11OF3-0726-SM-5BC4Z | Liver | TCGA-LIHC | 19.04 | 7.36 |
| GTEX-11TT1-1726-SM-5EQLJ | Liver | TCGA-LIHC | 55.17 | 29.42 |
| GTEX-11TUW-1726-SM-5BC5C | Liver | TCGA-LIHC | 29.5 | 7.15 |
| GTEX-11WQC-0726-SM-5EQMR | Liver | TCGA-LIHC | 29.12 | 9.01 |
| GTEX-11ZTS-1426-SM-5EQMM | Liver | TCGA-LIHC | 13.95 | 4.26 |
| GTEX-11ZUS-2526-SM-59872 | Liver | TCGA-LIHC | 14.67 | 3.07 |
| GTEX-11ZVC-0726-SM-5FQT9 | Liver | TCGA-LIHC | 21.66 | 6.67 |
| GTEX-1212Z-0226-SM-59HLF | Liver | TCGA-LIHC | 16.96 | 6.87 |
| GTEX-12696-0826-SM-5EGGE | Liver | TCGA-LIHC | 27.03 | 7.53 |
| GTEX-1269C-0626-SM-5FQSS | Liver | TCGA-LIHC | 27.93 | 12.35 |
| GTEX-12KS4-1326-SM-5LUB3 | Liver | TCGA-LIHC | 31 | 10.28 |
| GTEX-12WSD-1426-SM-5GCN9 | Liver | TCGA-LIHC | 18.71 | 7.12 |
| GTEX-12WSG-0626-SM-5FQTQ | Liver | TCGA-LIHC | 27.09 | 12.78 |
| GTEX-12WSI-0226-SM-5GCNA | Liver | TCGA-LIHC | 16.67 | 4.81 |
| GTEX-12WSL-0226-SM-5CVMJ | Liver | TCGA-LIHC | 18.64 | 9.41 |
| GTEX-12WSM-0726-SM-5GCOW | Liver | TCGA-LIHC | 21.62 | 6.42 |
| GTEX-12ZZZ-1326-SM-59HKW | Liver | TCGA-LIHC | 5.64 | 2.52 |

| | | | | |
|---|---|---|---|---|
| GTEX-13112-1426-SM-5EGH8 | Liver | TCGA-LIHC | 18.71 | 5.75 |
| GTEX-13113-1326-SM-5GCOI | Liver | TCGA-LIHC | 28.25 | 6.59 |
| GTEX-131XE-0326-SM-5LZVO | Liver | TCGA-LIHC | 38.09 | 10.57 |
| GTEX-131XH-0626-SM-5LZWH | Liver | TCGA-LIHC | 17.66 | 6.2 |
| GTEX-131YS-1626-SM-5HL6C | Liver | TCGA-LIHC | 20.2 | 7 |
| GTEX-132AR-0426-SM-5IFH8 | Liver | TCGA-LIHC | 53.82 | 14.12 |
| GTEX-132NY-0926-SM-5P9G3 | Liver | TCGA-LIHC | 24.61 | 11.12 |
| GTEX-1399R-1226-SM-5P9GF | Liver | TCGA-LIHC | 22.91 | 8.66 |
| GTEX-1399T-0826-SM-5IFES | Liver | TCGA-LIHC | 27.32 | 8.16 |
| GTEX-139TS-1426-SM-5IFJD | Liver | TCGA-LIHC | 22.9 | 8.53 |
| GTEX-139TU-0826-SM-5IJFG | Liver | TCGA-LIHC | 21.27 | 6.64 |
| GTEX-139YR-0226-SM-5IFEM | Liver | TCGA-LIHC | 32.21 | 9.99 |
| GTEX-13FLV-0326-SM-5N9DJ | Liver | TCGA-LIHC | 19.69 | 6.83 |
| GTEX-13FTW-1126-SM-5J2NV | Liver | TCGA-LIHC | 29.65 | 9.79 |
| GTEX-13FTZ-0726-SM-5IFFY | Liver | TCGA-LIHC | 19.73 | 7.64 |
| GTEX-13N11-0926-SM-5IJG2 | Liver | TCGA-LIHC | 38.6 | 15.83 |
| GTEX-13N2G-0926-SM-5IFGJ | Liver | TCGA-LIHC | 18.54 | 7.2 |
| GTEX-13NYB-1026-SM-5IFH3 | Liver | TCGA-LIHC | 28.14 | 9.6 |
| GTEX-13NZ9-1326-SM-5MR3V | Liver | TCGA-LIHC | 40.73 | 12.07 |
| GTEX-13NZB-0626-SM-5IFH6 | Liver | TCGA-LIHC | 15.6 | 5.72 |
| GTEX-13O1R-2026-SM-5KM3N | Liver | TCGA-LIHC | 58.93 | 13.58 |
| GTEX-13O3O-1826-SM-5IFGW | Liver | TCGA-LIHC | 19.07 | 5.86 |
| GTEX-13OW6-2626-SM-5IFF2 | Liver | TCGA-LIHC | 25.23 | 6.7 |
| GTEX-13PVQ-1526-SM-5IFEQ | Liver | TCGA-LIHC | 33.91 | 14.87 |
| GTEX-13PVR-0126-SM-5S2PY | Liver | TCGA-LIHC | 27.7 | 13.83 |
| GTEX-13QJC-0726-SM-5RQJK | Liver | TCGA-LIHC | 17.9 | 5.99 |
| GTEX-13SLX-1226-SM-5S2Q6 | Liver | TCGA-LIHC | 26.64 | 12.17 |
| GTEX-13VXU-0926-SM-5IFFH | Liver | TCGA-LIHC | 18.38 | 4.37 |
| GTEX-13X6J-1826-SM-5TDCT | Liver | TCGA-LIHC | 8.36 | 2.81 |
| GTEX-144GL-1426-SM-79OMA | Liver | TCGA-LIHC | 14.93 | 4.1 |
| GTEX-144GM-1326-SM-5LU5E | Liver | TCGA-LIHC | 26.39 | 7.89 |

| | | | | |
|---|---|---|---|---|
| GTEX-145LU-1326-SM-5LU9N | Liver | TCGA-LIHC | 15.9 | 3.73 |
| GTEX-145MF-0826-SM-5QGQA | Liver | TCGA-LIHC | 24.53 | 7.28 |
| GTEX-145MO-2326-SM-5NQ9K | Liver | TCGA-LIHC | 19.43 | 7.13 |
| GTEX-146FH-1526-SM-5NQBU | Liver | TCGA-LIHC | 31.46 | 9.49 |
| GTEX-14753-1626-SM-5NQ9L | Liver | TCGA-LIHC | 13.51 | 4.3 |
| GTEX-147F4-1426-SM-5LUA8 | Liver | TCGA-LIHC | 27.44 | 8.48 |
| GTEX-147GR-1326-SM-7IGLB | Liver | TCGA-LIHC | 16.5 | 4.86 |
| GTEX-147JS-1126-SM-5RQIW | Liver | TCGA-LIHC | 18.59 | 8.18 |
| GTEX-148VI-0626-SM-5TDDH | Liver | TCGA-LIHC | 30.65 | 11.47 |
| GTEX-1497J-0726-SM-5Q5D1 | Liver | TCGA-LIHC | 22.46 | 6.05 |
| GTEX-14A5I-1726-SM-5QGQ5 | Liver | TCGA-LIHC | 19.71 | 4.92 |
| GTEX-14AS3-0126-SM-5Q5F4 | Liver | TCGA-LIHC | 32.53 | 10.04 |
| GTEX-14BIL-1326-SM-79OMR | Liver | TCGA-LIHC | 21.1 | 8.88 |
| GTEX-14C38-1526-SM-5RQJ7 | Liver | TCGA-LIHC | 27.74 | 9.83 |
| GTEX-14DAQ-1726-SM-5S2R2 | Liver | TCGA-LIHC | 21.92 | 4.98 |
| GTEX-14E1K-0326-SM-5S2PE | Liver | TCGA-LIHC | 21.39 | 7.71 |
| GTEX-14E7W-1526-SM-6871X | Liver | TCGA-LIHC | 24.71 | 6.43 |
| GTEX-14JG1-1626-SM-664NH | Liver | TCGA-LIHC | 23.76 | 4.82 |
| GTEX-14PHX-0526-SM-664NW | Liver | TCGA-LIHC | 19.98 | 6.72 |
| GTEX-14PJO-1726-SM-68719 | Liver | TCGA-LIHC | 19.11 | 6.22 |
| GTEX-14XAO-0226-SM-68728 | Liver | TCGA-LIHC | 24.83 | 10.25 |
| GTEX-15DYW-1326-SM-6LPIV | Liver | TCGA-LIHC | 19.37 | 5.8 |
| GTEX-15ER7-1826-SM-6LLI7 | Liver | TCGA-LIHC | 25.49 | 6.52 |
| GTEX-15RJ7-2026-SM-6LPJ4 | Liver | TCGA-LIHC | 22.51 | 10.4 |
| GTEX-15RJE-1126-SM-6LPI5 | Liver | TCGA-LIHC | 11.41 | 3.72 |
| GTEX-15SHU-1826-SM-7KUKS | Liver | TCGA-LIHC | 20.38 | 6.59 |
| GTEX-15SZO-1226-SM-6LPIP | Liver | TCGA-LIHC | 20.26 | 5.65 |
| GTEX-17EVP-1026-SM-7IGOZ | Liver | TCGA-LIHC | 16.91 | 7.24 |
| GTEX-17F96-1226-SM-79OK2 | Liver | TCGA-LIHC | 21.01 | 6.32 |
| GTEX-17HGU-1826-SM-7IGQM | Liver | TCGA-LIHC | 20.09 | 4.92 |
| GTEX-17HHE-0126-SM-79398 | Liver | TCGA-LIHC | 23.58 | 10.05 |

| | | | | |
|---|---|---|---|---|
| GTEX-17HII-1026-SM-7IGLY | Liver | TCGA-LIHC | 19.38 | 6.66 |
| GTEX-17MF6-1126-SM-7DUFS | Liver | TCGA-LIHC | 19.22 | 5.97 |
| GTEX-18465-1626-SM-7LG6N | Liver | TCGA-LIHC | 13.24 | 3.74 |
| GTEX-18A66-2026-SM-7189C | Liver | TCGA-LIHC | 18.66 | 7.2 |
| GTEX-18A6Q-2026-SM-718AQ | Liver | TCGA-LIHC | 25.33 | 11.14 |
| GTEX-18A7A-1526-SM-72D69 | Liver | TCGA-LIHC | 24.7 | 6.82 |
| GTEX-18D9B-1526-SM-7KFRK | Liver | TCGA-LIHC | 27.57 | 13.03 |
| GTEX-1A32A-0826-SM-72D5C | Liver | TCGA-LIHC | 23.94 | 9.25 |
| GTEX-1A8FM-1226-SM-7IGMD | Liver | TCGA-LIHC | 19.72 | 5.87 |
| GTEX-1A8G7-1126-SM-731ED | Liver | TCGA-LIHC | 18.64 | 8.03 |
| GTEX-1AMFI-0826-SM-731DV | Liver | TCGA-LIHC | 26.26 | 11.38 |
| GTEX-1AX8Z-1026-SM-7189R | Liver | TCGA-LIHC | 27.76 | 9.47 |
| GTEX-1AX9I-1626-SM-72D5D | Liver | TCGA-LIHC | 12.37 | 5.97 |
| GTEX-1B8L1-0726-SM-7P8RT | Liver | TCGA-LIHC | 20.77 | 7.05 |
| GTEX-1B8SG-1326-SM-79ONT | Liver | TCGA-LIHC | 31.21 | 6.38 |
| GTEX-1B932-1426-SM-793AN | Liver | TCGA-LIHC | 19.34 | 9.34 |
| GTEX-1B933-1226-SM-731DW | Liver | TCGA-LIHC | 18.18 | 6.83 |
| GTEX-1B996-1426-SM-7EPIA | Liver | TCGA-LIHC | 26.55 | 11.04 |
| GTEX-1BAJH-1526-SM-7RHGI | Liver | TCGA-LIHC | 28.11 | 8.47 |
| GTEX-1C6VQ-1626-SM-79OO1 | Liver | TCGA-LIHC | 26.5 | 7.88 |
| GTEX-1CAMQ-1426-SM-7MKFL | Liver | TCGA-LIHC | 13.82 | 4.13 |
| GTEX-1CB4F-1726-SM-7MGXH | Liver | TCGA-LIHC | 21.57 | 8.36 |
| GTEX-1CB4J-0726-SM-7IGN8 | Liver | TCGA-LIHC | 24.75 | 9.47 |
| GTEX-1E2YA-1226-SM-7MGY3 | Liver | TCGA-LIHC | 16.79 | 6.09 |
| GTEX-1EH9U-1126-SM-7IGLU | Liver | TCGA-LIHC | 21.85 | 10.2 |
| GTEX-1EKGG-1226-SM-7IGNO | Liver | TCGA-LIHC | 13.22 | 3.95 |
| GTEX-1EN7A-2026-SM-7MGY4 | Liver | TCGA-LIHC | 19.42 | 4.74 |
| GTEX-1EWIQ-0726-SM-7MKFW | Liver | TCGA-LIHC | 19.07 | 5.69 |
| GTEX-1F75A-1626-SM-9JGGT | Liver | TCGA-LIHC | 10.55 | 3.79 |
| GTEX-1GF9V-1226-SM-7MKGY | Liver | TCGA-LIHC | 15.54 | 4.94 |
| GTEX-1GF9W-0826-SM-7MKHI | Liver | TCGA-LIHC | 31.66 | 7.02 |

| | | | | |
|---|---|---|---|---|
| GTEX-1GF9X-0426-SM-7MKHN | Liver | TCGA-LIHC | 35.28 | 9.5 |
| GTEX-1GN1U-0926-SM-9WPPY | Liver | TCGA-LIHC | 18.72 | 6.77 |
| GTEX-1GN2E-0926-SM-9WPOH | Liver | TCGA-LIHC | 22.76 | 8.31 |
| GTEX-1GN73-1126-SM-9OSW5 | Liver | TCGA-LIHC | 30.87 | 15.02 |
| GTEX-1GZ4I-1526-SM-9OSW9 | Liver | TCGA-LIHC | 19.74 | 6.6 |
| GTEX-1H11D-0826-SM-9OSWB | Liver | TCGA-LIHC | 27.4 | 10.3 |
| GTEX-1HB9E-0626-SM-AHZ4K | Liver | TCGA-LIHC | 21.55 | 7.36 |
| GTEX-1HBPH-2626-SM-A96TV | Liver | TCGA-LIHC | 16.66 | 6.42 |
| GTEX-1HBPI-1026-SM-AHZ4N | Liver | TCGA-LIHC | 23 | 5.88 |
| GTEX-1HCU7-1726-SM-B2LX9 | Liver | TCGA-LIHC | 19.17 | 8.34 |
| GTEX-1HCVE-1526-SM-B2LWW | Liver | TCGA-LIHC | 17.97 | 3.92 |
| GTEX-1HT8W-0826-SM-C1YRJ | Liver | TCGA-LIHC | 20.27 | 4.39 |
| GTEX-1I1GR-1526-SM-CNNQE | Liver | TCGA-LIHC | 12.02 | 4.27 |
| GTEX-1I1HK-1126-SM-AHZ2L | Liver | TCGA-LIHC | 13.61 | 4.4 |
| GTEX-1ICG6-1026-SM-B2LY6 | Liver | TCGA-LIHC | 19.69 | 8.05 |
| GTEX-1ICLY-1526-SM-C1YS4 | Liver | TCGA-LIHC | 24.57 | 10.58 |
| GTEX-1IKJJ-1526-SM-C1YSC | Liver | TCGA-LIHC | 21.38 | 7.44 |
| GTEX-1IKK5-0326-SM-C1YPY | Liver | TCGA-LIHC | 29.94 | 9.91 |
| GTEX-1J8EW-1626-SM-C1YQ8 | Liver | TCGA-LIHC | 14.66 | 5.77 |
| GTEX-1JJ6O-0826-SM-CXZJL | Liver | TCGA-LIHC | 19.81 | 10.87 |
| GTEX-1JJEA-1526-SM-CNNPK | Liver | TCGA-LIHC | 20.37 | 5.21 |
| GTEX-1JMLX-1326-SM-CXZJO | Liver | TCGA-LIHC | 31.64 | 8.04 |
| GTEX-1K9T9-0726-SM-CXZJZ | Liver | TCGA-LIHC | 31.29 | 14.9 |
| GTEX-1KAFJ-0726-SM-CXZKF | Liver | TCGA-LIHC | 18.47 | 8.86 |
| GTEX-1KANB-2226-SM-DHXJW | Liver | TCGA-LIHC | 18.17 | 5.97 |
| GTEX-1KD4Q-1526-SM-D4P3L | Liver | TCGA-LIHC | 14.41 | 4.85 |
| GTEX-1KD5A-1226-SM-CXZK7 | Liver | TCGA-LIHC | 29.57 | 11.59 |
| GTEX-1KWVE-0326-SM-EV7AF | Liver | TCGA-LIHC | 18.99 | 6.79 |
| GTEX-1LGRB-0626-SM-CNNQH | Liver | TCGA-LIHC | 27.62 | 9.27 |
| GTEX-1LH75-0926-SM-EAZ5C | Liver | TCGA-LIHC | 25.83 | 13.69 |
| GTEX-1MJIX-0726-SM-DPRYP | Liver | TCGA-LIHC | 12.66 | 4.48 |

| | | | | |
|---|---|---|---|---|
| GTEX-1NV8Z-1626-SM-DPRYT | Liver | TCGA-LIHC | 23.1 | 5.89 |
| GTEX-1PBJJ-0626-SM-DPRYO | Liver | TCGA-LIHC | 15.83 | 4.7 |
| GTEX-1PIIG-1326-SM-EXOIR | Liver | TCGA-LIHC | 13.13 | 4.61 |
| GTEX-1QCLZ-1226-SM-EAZ49 | Liver | TCGA-LIHC | 23.89 | 14.85 |
| GTEX-1QP66-0226-SM-DPRXS | Liver | TCGA-LIHC | 25.66 | 10.17 |
| GTEX-1QP6S-1326-SM-DPRZ8 | Liver | TCGA-LIHC | 25.56 | 7.6 |
| GTEX-1QPFJ-1126-SM-E9U5V | Liver | TCGA-LIHC | 29.12 | 10.75 |
| GTEX-1R9JW-0826-SM-DPRZJ | Liver | TCGA-LIHC | 23.3 | 7.9 |
| GTEX-1RQEC-1126-SM-EAZ5E | Liver | TCGA-LIHC | 15.73 | 5.11 |
| GTEX-1S5VW-1826-SM-EAZ53 | Liver | TCGA-LIHC | 18.88 | 8.38 |
| GTEX-1S82Y-1026-SM-EV7BJ | Liver | TCGA-LIHC | 13.53 | 5.81 |
| GTEX-O5YT-0826-SM-3TW8N | Liver | TCGA-LIHC | 50.49 | 23.22 |
| GTEX-O5YV-0726-SM-EV798 | Liver | TCGA-LIHC | 29.04 | 9.03 |
| GTEX-OIZF-0826-SM-3MJGO | Liver | TCGA-LIHC | 28.88 | 15.11 |
| GTEX-OOBJ-0826-SM-3NB2K | Liver | TCGA-LIHC | 20.46 | 8.01 |
| GTEX-P44G-1126-SM-3NM9D | Liver | TCGA-LIHC | 19.18 | 5.22 |
| GTEX-P78B-1326-SM-3P611 | Liver | TCGA-LIHC | 28.65 | 9.14 |
| GTEX-PX3G-0826-SM-48TZS | Liver | TCGA-LIHC | 21.48 | 6.32 |
| GTEX-Q2AG-1126-SM-48U1P | Liver | TCGA-LIHC | 18.34 | 5.52 |
| GTEX-Q734-0326-SM-48U15 | Liver | TCGA-LIHC | 23.93 | 6.16 |
| GTEX-QDVN-0826-SM-48TZ2 | Liver | TCGA-LIHC | 31.21 | 11.84 |
| GTEX-QEG4-1826-SM-CKZN9 | Liver | TCGA-LIHC | 20.59 | 6.17 |
| GTEX-QEL4-1226-SM-447A4 | Liver | TCGA-LIHC | 36.66 | 9.41 |
| GTEX-QESD-2026-SM-447BI | Liver | TCGA-LIHC | 26.17 | 7.56 |
| GTEX-QV44-0326-SM-C1YQX | Liver | TCGA-LIHC | 17.76 | 8.79 |
| GTEX-R53T-0326-SM-48FEC | Liver | TCGA-LIHC | 29 | 7.87 |
| GTEX-REY6-1226-SM-48FDR | Liver | TCGA-LIHC | 26.45 | 9.26 |
| GTEX-RM2N-1926-SM-48FCU | Liver | TCGA-LIHC | 23.73 | 12.93 |
| GTEX-RN64-1826-SM-48FDV | Liver | TCGA-LIHC | 20.45 | 5.42 |
| GTEX-RNOR-1426-SM-48FDJ | Liver | TCGA-LIHC | 19.65 | 5.73 |
| GTEX-RTLS-1326-SM-46MUN | Liver | TCGA-LIHC | 19.06 | 6.44 |

| | | | | |
|---|---|---|---|---|
| GTEX-RU72-1426-SM-46MUF | Liver | TCGA-LIHC | 27.15 | 9.81 |
| GTEX-RWSA-1426-SM-47JXA | Liver | TCGA-LIHC | 22.16 | 7.42 |
| GTEX-S32W-1926-SM-4AD63 | Liver | TCGA-LIHC | 31.13 | 16.21 |
| GTEX-S33H-1626-SM-4AD68 | Liver | TCGA-LIHC | 29.57 | 8.99 |
| GTEX-S4Z8-0526-SM-4AD4T | Liver | TCGA-LIHC | 19.27 | 6.96 |
| GTEX-SJXC-1226-SM-4DM78 | Liver | TCGA-LIHC | 14.37 | 5.26 |
| GTEX-T6MN-1226-SM-3NMA5 | Liver | TCGA-LIHC | 22.88 | 8.06 |
| GTEX-TKQ2-1726-SM-4DXUP | Liver | TCGA-LIHC | 26.87 | 10.83 |
| GTEX-U3ZN-0226-SM-3DB8D | Liver | TCGA-LIHC | 28.5 | 16.7 |
| GTEX-U8XE-1526-SM-4E3HT | Liver | TCGA-LIHC | 26.46 | 7.84 |
| GTEX-UPK5-1426-SM-4JBHH | Liver | TCGA-LIHC | 22.61 | 6.33 |
| GTEX-UTHO-2426-SM-4JBHD | Liver | TCGA-LIHC | 33.38 | 11.77 |
| GTEX-VUSG-0126-SM-4KL1X | Liver | TCGA-LIHC | 17.88 | 4 |
| GTEX-WFON-1726-SM-4LVMQ | Liver | TCGA-LIHC | 19.45 | 5.8 |
| GTEX-WK11-1326-SM-4OOSI | Liver | TCGA-LIHC | 6.95 | 1.81 |
| GTEX-WQUQ-1926-SM-4OOSA | Liver | TCGA-LIHC | 16.18 | 5.2 |
| GTEX-WVLH-2126-SM-EYYWD | Liver | TCGA-LIHC | 15.42 | 4.03 |
| GTEX-WY7C-0726-SM-4ONCB | Liver | TCGA-LIHC | 23.6 | 12.73 |
| GTEX-WYVS-1926-SM-4PQZ2 | Liver | TCGA-LIHC | 21.2 | 6.77 |
| GTEX-WZTO-0626-SM-4PQYY | Liver | TCGA-LIHC | 16.96 | 5.06 |
| GTEX-X261-1726-SM-4PQYT | Liver | TCGA-LIHC | 17.17 | 6.49 |
| GTEX-X3Y1-2726-SM-4PQZH | Liver | TCGA-LIHC | 21.5 | 8.67 |
| GTEX-X4EO-1126-SM-4QARQ | Liver | TCGA-LIHC | 21.49 | 6.57 |
| GTEX-X4EP-1026-SM-4QAS5 | Liver | TCGA-LIHC | 19.56 | 4.17 |
| GTEX-X4XY-1626-SM-46MVN | Liver | TCGA-LIHC | 28.98 | 13.58 |
| GTEX-XBEC-1526-SM-4AT68 | Liver | TCGA-LIHC | 15.07 | 5.01 |
| GTEX-XXEK-1126-SM-4BRUX | Liver | TCGA-LIHC | 39.6 | 10.25 |
| GTEX-Y111-0726-SM-EYYVM | Liver | TCGA-LIHC | 16.64 | 5.15 |
| GTEX-Y5LM-0426-SM-4VBRO | Liver | TCGA-LIHC | 33.8 | 18.34 |
| GTEX-Y5V5-0926-SM-4VBPZ | Liver | TCGA-LIHC | 18.64 | 6.1 |
| GTEX-YB5E-0326-SM-5IFHU | Liver | TCGA-LIHC | 38.87 | 12 |

| | | | | |
|---|---|---|---|---|
| GTEX-YEC4-0826-SM-5P9FV | Liver | TCGA-LIHC | 19.14 | 8.6 |
| GTEX-YECK-1926-SM-4W21H | Liver | TCGA-LIHC | 21.02 | 7.04 |
| GTEX-YFC4-1526-SM-5IFJS | Liver | TCGA-LIHC | 29.9 | 9.83 |
| GTEX-Z9EW-0426-SM-5CVM9 | Liver | TCGA-LIHC | 24.35 | 9.76 |
| GTEX-ZAB4-0826-SM-5LU9D | Liver | TCGA-LIHC | 18.65 | 6.42 |
| GTEX-ZAB5-0426-SM-5CVMI | Liver | TCGA-LIHC | 26.23 | 8.85 |
| GTEX-ZEX8-0826-SM-4WKHK | Liver | TCGA-LIHC | 26.31 | 8.78 |
| GTEX-ZF29-2026-SM-DNZYW | Liver | TCGA-LIHC | 24.84 | 8.24 |
| GTEX-ZF2S-3026-SM-4WWCH | Liver | TCGA-LIHC | 19.99 | 8.07 |
| GTEX-ZPU1-0826-SM-57WG2 | Liver | TCGA-LIHC | 25.2 | 9.96 |
| GTEX-ZTPG-1426-SM-51MT3 | Liver | TCGA-LIHC | 38.24 | 7.78 |
| GTEX-ZVP2-0626-SM-51MSO | Liver | TCGA-LIHC | 14.44 | 5.83 |
| GTEX-ZVT3-1626-SM-5GU66 | Liver | TCGA-LIHC | 23.05 | 5.43 |
| GTEX-ZVT4-0626-SM-5E45T | Liver | TCGA-LIHC | 19.26 | 6.63 |
| GTEX-ZYT6-0626-SM-5E45V | Liver | TCGA-LIHC | 19.96 | 4.09 |
| GTEX-ZYY3-0626-SM-5NQ6W | Liver | TCGA-LIHC | 38.58 | 8.97 |
| GTEX-ZZPU-0426-SM-5GZYH | Liver | TCGA-LIHC | 22.8 | 5.53 |
| GTEX-111CU-0326-SM-5GZXO | Lung | TCGA-LUSC | 77.77 | 17.7 |
| GTEX-111FC-1126-SM-5GZWU | Lung | TCGA-LUSC | 42.84 | 13.5 |
| GTEX-111VG-0726-SM-5GIDC | Lung | TCGA-LUSC | 36.88 | 12.38 |
| GTEX-111YS-0626-SM-5GZXV | Lung | TCGA-LUSC | 58.75 | 9.98 |
| GTEX-1122O-0126-SM-5GICA | Lung | TCGA-LUSC | 51.83 | 10.68 |
| GTEX-1128S-0726-SM-5N9D6 | Lung | TCGA-LUSC | 34.04 | 8.37 |
| GTEX-117YW-0526-SM-5H11C | Lung | TCGA-LUSC | 20 | 7.99 |
| GTEX-117YX-1326-SM-5H125 | Lung | TCGA-LUSC | 52.14 | 13.8 |
| GTEX-11DXX-0626-SM-5Q5AG | Lung | TCGA-LUSC | 55.12 | 14.77 |
| GTEX-11DXZ-0726-SM-5N9C4 | Lung | TCGA-LUSC | 38.24 | 15.92 |
| GTEX-11DZ1-0426-SM-5H11A | Lung | TCGA-LUSC | 25.98 | 7.09 |
| GTEX-11EI6-0826-SM-5985V | Lung | TCGA-LUSC | 23.22 | 4.68 |
| GTEX-11EMC-0126-SM-5EGKV | Lung | TCGA-LUSC | 33.05 | 6.97 |
| GTEX-11EQ9-0226-SM-5A5JX | Lung | TCGA-LUSC | 59.67 | 15.72 |

| | | | | |
|---|---|---|---|---|
| GTEX-11GSP-0726-SM-5986L | Lung | TCGA-LUSC | 53.76 | 9.06 |
| GTEX-11I78-0126-SM-5HL6F | Lung | TCGA-LUSC | 42.23 | 8.96 |
| GTEX-11LCK-0426-SM-5A5M8 | Lung | TCGA-LUSC | 55.94 | 17.52 |
| GTEX-11NSD-0326-SM-5A5LS | Lung | TCGA-LUSC | 50.23 | 11.57 |
| GTEX-11NUK-0826-SM-5HL4U | Lung | TCGA-LUSC | 30.53 | 7.3 |
| GTEX-11NV4-1126-SM-5HL6J | Lung | TCGA-LUSC | 22.72 | 3.5 |
| GTEX-11O72-1326-SM-5BC5A | Lung | TCGA-LUSC | 48.48 | 10.53 |
| GTEX-11OF3-1126-SM-5986C | Lung | TCGA-LUSC | 45.43 | 4.11 |
| GTEX-11P7K-0326-SM-59871 | Lung | TCGA-LUSC | 53.2 | 13.07 |
| GTEX-11P81-0226-SM-5HL5M | Lung | TCGA-LUSC | 47.2 | 15.22 |
| GTEX-11PRG-0926-SM-5EGI8 | Lung | TCGA-LUSC | 45.64 | 8.07 |
| GTEX-11TT1-1626-SM-5EQL7 | Lung | TCGA-LUSC | 63.62 | 23.23 |
| GTEX-11TUW-0526-SM-5LU9A | Lung | TCGA-LUSC | 44.28 | 8.92 |
| GTEX-11UD2-0726-SM-5EQ69 | Lung | TCGA-LUSC | 41.56 | 7.42 |
| GTEX-11WQC-0626-SM-5EQMF | Lung | TCGA-LUSC | 29.36 | 9.42 |
| GTEX-11WQK-1226-SM-5GU5Z | Lung | TCGA-LUSC | 46.31 | 10.71 |
| GTEX-11ZTS-1226-SM-5EQMQ | Lung | TCGA-LUSC | 29.57 | 5.55 |
| GTEX-11ZTT-0626-SM-5EQLM | Lung | TCGA-LUSC | 44.98 | 12.75 |
| GTEX-11ZUS-0126-SM-5EQM5 | Lung | TCGA-LUSC | 31.92 | 8.47 |
| GTEX-11ZVC-0226-SM-731E8 | Lung | TCGA-LUSC | 29.77 | 8.98 |
| GTEX-1211K-0826-SM-5FQUP | Lung | TCGA-LUSC | 55.91 | 18.42 |
| GTEX-1212Z-1026-SM-5EGJ8 | Lung | TCGA-LUSC | 28.77 | 11.04 |
| GTEX-12584-1426-SM-5EGJ9 | Lung | TCGA-LUSC | 44.53 | 7.51 |
| GTEX-12696-1026-SM-5FQUV | Lung | TCGA-LUSC | 45.26 | 7.9 |
| GTEX-1269C-0926-SM-5FQSR | Lung | TCGA-LUSC | 35.97 | 4.96 |
| GTEX-12BJ1-1026-SM-5EGJA | Lung | TCGA-LUSC | 62.82 | 9.17 |
| GTEX-12KS4-0726-SM-5FQSX | Lung | TCGA-LUSC | 62.97 | 24.11 |
| GTEX-12WSA-1026-SM-5EGHN | Lung | TCGA-LUSC | 35.46 | 5.9 |
| GTEX-12WSD-0826-SM-5GCNE | Lung | TCGA-LUSC | 52.19 | 20.79 |
| GTEX-12WSE-0826-SM-5S2VL | Lung | TCGA-LUSC | 40.79 | 11.47 |
| GTEX-12WSG-5004-SM-7EPG9 | Lung | TCGA-LUSC | 55.15 | 13.17 |

| | | | | |
|---|---|---|---|---|
| GTEX-12WSH-0126-SM-5GCO3 | Lung | TCGA-LUSC | 43.18 | 7.71 |
| GTEX-12WSI-0826-SM-5EGKD | Lung | TCGA-LUSC | 43.35 | 5.43 |
| GTEX-12WSJ-0226-SM-5GCP7 | Lung | TCGA-LUSC | 59.87 | 14.89 |
| GTEX-12WSK-0826-SM-5CVNP | Lung | TCGA-LUSC | 40.23 | 7.44 |
| GTEX-12WSL-1026-SM-5CVNJ | Lung | TCGA-LUSC | 48.84 | 13.27 |
| GTEX-12WSN-5004-SM-793CE | Lung | TCGA-LUSC | 69.96 | 18 |
| GTEX-12ZZW-0926-SM-5LZUD | Lung | TCGA-LUSC | 36.11 | 7.85 |
| GTEX-12ZZY-0926-SM-5EQ6I | Lung | TCGA-LUSC | 56.48 | 15.03 |
| GTEX-13111-0426-SM-5DUXR | Lung | TCGA-LUSC | 45.59 | 9.22 |
| GTEX-13113-5004-SM-79ONL | Lung | TCGA-LUSC | 65.33 | 14.69 |
| GTEX-1313W-0926-SM-5EQ56 | Lung | TCGA-LUSC | 42.2 | 9.09 |
| GTEX-131XE-0726-SM-5HL9K | Lung | TCGA-LUSC | 58.2 | 11.82 |
| GTEX-131XF-1026-SM-5BC6A | Lung | TCGA-LUSC | 42.34 | 7.89 |
| GTEX-131XH-0426-SM-5DUWU | Lung | TCGA-LUSC | 40.81 | 6.81 |
| GTEX-131XW-1126-SM-5EGK4 | Lung | TCGA-LUSC | 25 | 6.13 |
| GTEX-131YS-0926-SM-5IJB9 | Lung | TCGA-LUSC | 50.14 | 14.22 |
| GTEX-132NY-1226-SM-5PNVF | Lung | TCGA-LUSC | 52.01 | 17.07 |
| GTEX-132QS-0726-SM-5IJE9 | Lung | TCGA-LUSC | 62.65 | 10.65 |
| GTEX-1339X-0626-SM-5IJER | Lung | TCGA-LUSC | 40.28 | 8.37 |
| GTEX-133LE-0526-SM-5N9EJ | Lung | TCGA-LUSC | 51.58 | 11.17 |
| GTEX-1399S-1726-SM-5L3DI | Lung | TCGA-LUSC | 60 | 14.08 |
| GTEX-1399U-0826-SM-5KM1P | Lung | TCGA-LUSC | 40.22 | 6.76 |
| GTEX-139T6-0426-SM-5IJEM | Lung | TCGA-LUSC | 48.03 | 14.42 |
| GTEX-139TT-0726-SM-5K7XW | Lung | TCGA-LUSC | 35.19 | 6.54 |
| GTEX-139UW-0226-SM-5K7WU | Lung | TCGA-LUSC | 42.28 | 6.28 |
| GTEX-139YR-0926-SM-5LZYB | Lung | TCGA-LUSC | 50.2 | 18.33 |
| GTEX-13CF3-0426-SM-5IJEU | Lung | TCGA-LUSC | 67.12 | 20.08 |
| GTEX-13D11-0326-SM-5LZXX | Lung | TCGA-LUSC | 58.72 | 21.93 |
| GTEX-13FH7-1726-SM-5IJE7 | Lung | TCGA-LUSC | 61.71 | 21.04 |
| GTEX-13FHO-1026-SM-5KM1Q | Lung | TCGA-LUSC | 30.45 | 10.07 |
| GTEX-13FHP-0726-SM-5K7YI | Lung | TCGA-LUSC | 40.8 | 6.63 |

| | | | | |
|---|---|---|---|---|
| GTEX-13FLV-0426-SM-5KLZA | Lung | TCGA-LUSC | 39.74 | 5.9 |
| GTEX-13FTX-0326-SM-5J2NG | Lung | TCGA-LUSC | 49.72 | 8.71 |
| GTEX-13FTY-0126-SM-5J2NZ | Lung | TCGA-LUSC | 22.77 | 3.79 |
| GTEX-13FTZ-0526-SM-5IJCW | Lung | TCGA-LUSC | 50.87 | 13.2 |
| GTEX-13G51-0426-SM-5K7Z5 | Lung | TCGA-LUSC | 41.81 | 9.59 |
| GTEX-13JUV-0526-SM-5K7XE | Lung | TCGA-LUSC | 60.49 | 16.23 |
| GTEX-13JVG-1426-SM-5MR4W | Lung | TCGA-LUSC | 38.36 | 18.01 |
| GTEX-13N11-0326-SM-5LUA3 | Lung | TCGA-LUSC | 35.06 | 8.62 |
| GTEX-13N1W-0726-SM-5MR57 | Lung | TCGA-LUSC | 49.11 | 5.86 |
| GTEX-13N2G-0826-SM-5IJE6 | Lung | TCGA-LUSC | 27.31 | 5.83 |
| GTEX-13NYB-0626-SM-5MR47 | Lung | TCGA-LUSC | 32.55 | 7.78 |
| GTEX-13NYS-1626-SM-5J2MU | Lung | TCGA-LUSC | 32.44 | 7.4 |
| GTEX-13NZ8-0326-SM-5L3DF | Lung | TCGA-LUSC | 66.91 | 13.44 |
| GTEX-13NZ9-0926-SM-5KM12 | Lung | TCGA-LUSC | 48.73 | 13.35 |
| GTEX-13NZA-1426-SM-5KM4Y | Lung | TCGA-LUSC | 46.59 | 13.76 |
| GTEX-13O21-3026-SM-5J2NI | Lung | TCGA-LUSC | 42.21 | 11.75 |
| GTEX-13O3O-0726-SM-5J1N7 | Lung | TCGA-LUSC | 44.75 | 7.92 |
| GTEX-13O3P-1026-SM-5N9E7 | Lung | TCGA-LUSC | 44.61 | 9.24 |
| GTEX-13O3Q-0526-SM-5KM18 | Lung | TCGA-LUSC | 26.46 | 9.33 |
| GTEX-13O61-0726-SM-5J2MD | Lung | TCGA-LUSC | 55.55 | 12.52 |
| GTEX-13OVG-0326-SM-5KM57 | Lung | TCGA-LUSC | 46.65 | 5.21 |
| GTEX-13OVH-1026-SM-5J2NP | Lung | TCGA-LUSC | 40.5 | 6.1 |
| GTEX-13OVJ-0726-SM-5KM1W | Lung | TCGA-LUSC | 75.28 | 24.21 |
| GTEX-13OVL-0626-SM-5KM13 | Lung | TCGA-LUSC | 45 | 12.8 |
| GTEX-13OW5-0726-SM-5KLZK | Lung | TCGA-LUSC | 39.71 | 10.33 |
| GTEX-13OW6-0826-SM-5L3GA | Lung | TCGA-LUSC | 57.29 | 10.64 |
| GTEX-13OW7-0926-SM-5L3EX | Lung | TCGA-LUSC | 26.17 | 6.6 |
| GTEX-13OW8-1726-SM-5L3GO | Lung | TCGA-LUSC | 27.02 | 5.9 |
| GTEX-13PL7-1726-SM-5J2NX | Lung | TCGA-LUSC | 49.63 | 8.07 |
| GTEX-13PVQ-0926-SM-5IJFD | Lung | TCGA-LUSC | 34.49 | 8.94 |
| GTEX-13QBU-0726-SM-5J2OA | Lung | TCGA-LUSC | 66.13 | 13.66 |

| | | | | |
|---|---|---|---|---|
| GTEX-13QJ3-1026-SM-5QGQU | Lung | TCGA-LUSC | 45.95 | 13.6 |
| GTEX-13QJC-0526-SM-5RQKB | Lung | TCGA-LUSC | 20.09 | 4.01 |
| GTEX-13RTJ-1126-SM-5S2UJ | Lung | TCGA-LUSC | 34.75 | 10.1 |
| GTEX-13S7M-2126-SM-5S2QR | Lung | TCGA-LUSC | 33.43 | 7.79 |
| GTEX-13S86-0626-SM-5Q5E7 | Lung | TCGA-LUSC | 78.21 | 23.29 |
| GTEX-13SLW-1226-SM-5S2Q7 | Lung | TCGA-LUSC | 55.76 | 21.61 |
| GTEX-13U4I-1426-SM-5J2M3 | Lung | TCGA-LUSC | 40.37 | 8.54 |
| GTEX-13VXT-1426-SM-5LU4B | Lung | TCGA-LUSC | 58.46 | 13.1 |
| GTEX-13VXU-2726-SM-5LU4N | Lung | TCGA-LUSC | 29.14 | 6.33 |
| GTEX-13W3W-0326-SM-731DS | Lung | TCGA-LUSC | 43.82 | 17.68 |
| GTEX-13X6K-1626-SM-7EWCX | Lung | TCGA-LUSC | 44.45 | 11.36 |
| GTEX-13YAN-1026-SM-5O9CF | Lung | TCGA-LUSC | 27.07 | 5.5 |
| GTEX-144GM-0126-SM-5Q5AX | Lung | TCGA-LUSC | 66.36 | 16.56 |
| GTEX-144GN-0426-SM-5O9AP | Lung | TCGA-LUSC | 55.36 | 23.07 |
| GTEX-144GO-0226-SM-5LUB1 | Lung | TCGA-LUSC | 35.26 | 4.48 |
| GTEX-145LS-1226-SM-5Q5D9 | Lung | TCGA-LUSC | 39.29 | 7.01 |
| GTEX-145LT-0326-SM-5LUAD | Lung | TCGA-LUSC | 57.05 | 10.66 |
| GTEX-145LU-0526-SM-5O9AT | Lung | TCGA-LUSC | 35.21 | 8.57 |
| GTEX-145ME-0226-SM-5S2QN | Lung | TCGA-LUSC | 76.76 | 14.87 |
| GTEX-145MF-0726-SM-5Q5BT | Lung | TCGA-LUSC | 42.92 | 9.28 |
| GTEX-145MH-0626-SM-5NQAK | Lung | TCGA-LUSC | 54.39 | 14.44 |
| GTEX-145MN-0926-SM-5NQBT | Lung | TCGA-LUSC | 47.56 | 8.33 |
| GTEX-145MO-1326-SM-5Q5EF | Lung | TCGA-LUSC | 45.09 | 9.25 |
| GTEX-146FH-1226-SM-5NQB6 | Lung | TCGA-LUSC | 61.83 | 18.54 |
| GTEX-146FQ-0926-SM-5LUAV | Lung | TCGA-LUSC | 64.31 | 22.34 |
| GTEX-1477Z-0626-SM-5NQB7 | Lung | TCGA-LUSC | 37.61 | 7.12 |
| GTEX-147F3-0726-SM-5NQ9U | Lung | TCGA-LUSC | 46.02 | 10.68 |
| GTEX-147F4-0926-SM-5Q5EO | Lung | TCGA-LUSC | 34 | 5.09 |
| GTEX-147GR-1226-SM-5TDCL | Lung | TCGA-LUSC | 31.45 | 7.07 |
| GTEX-147JS-1226-SM-5RQK4 | Lung | TCGA-LUSC | 47.08 | 9.01 |
| GTEX-148VI-0226-SM-5RQKA | Lung | TCGA-LUSC | 50.25 | 8.65 |

| | | | | |
|---|---|---|---|---|
| GTEX-148VJ-0826-SM-5LU8V | Lung | TCGA-LUSC | 29.65 | 4.96 |
| GTEX-1497J-0326-SM-5Q5CN | Lung | TCGA-LUSC | 59.24 | 12.22 |
| GTEX-14A6H-0526-SM-5NQAZ | Lung | TCGA-LUSC | 45.33 | 17.82 |
| GTEX-14ABY-1126-SM-5Q5F8 | Lung | TCGA-LUSC | 33.97 | 6.63 |
| GTEX-14AS3-0926-SM-5TDD6 | Lung | TCGA-LUSC | 54.35 | 9.19 |
| GTEX-14BIL-1226-SM-79OME | Lung | TCGA-LUSC | 41.6 | 12.9 |
| GTEX-14BMU-0526-SM-73KW4 | Lung | TCGA-LUSC | 61.67 | 22.29 |
| GTEX-14BMV-0826-SM-73KXU | Lung | TCGA-LUSC | 32.54 | 7.58 |
| GTEX-14C39-0326-SM-5TDDX | Lung | TCGA-LUSC | 50.84 | 6.08 |
| GTEX-14C5O-1126-SM-5TDEH | Lung | TCGA-LUSC | 45.44 | 12.64 |
| GTEX-14DAQ-0926-SM-793AZ | Lung | TCGA-LUSC | 29.76 | 7.92 |
| GTEX-14DAR-0226-SM-5S2PR | Lung | TCGA-LUSC | 55.61 | 13.03 |
| GTEX-14E1K-0226-SM-62LDT | Lung | TCGA-LUSC | 54 | 11.21 |
| GTEX-14E6C-1426-SM-5ZZWH | Lung | TCGA-LUSC | 54.82 | 9.28 |
| GTEX-14E6D-1026-SM-5S2RS | Lung | TCGA-LUSC | 38.55 | 13.38 |
| GTEX-14E6E-0426-SM-73KUE | Lung | TCGA-LUSC | 43 | 9.93 |
| GTEX-14E7W-1326-SM-5RQIV | Lung | TCGA-LUSC | 33.57 | 7.15 |
| GTEX-14JG1-0926-SM-5YY8W | Lung | TCGA-LUSC | 41.76 | 9.09 |
| GTEX-14JG6-0326-SM-6AJBT | Lung | TCGA-LUSC | 49.41 | 11.6 |
| GTEX-14JIY-1326-SM-6AJB3 | Lung | TCGA-LUSC | 26.77 | 10.22 |
| GTEX-14LLW-0726-SM-5ZZVV | Lung | TCGA-LUSC | 37.4 | 8.55 |
| GTEX-14LZ3-0726-SM-5YYAB | Lung | TCGA-LUSC | 51.7 | 10.62 |
| GTEX-14PHY-0526-SM-664NM | Lung | TCGA-LUSC | 40.57 | 16.29 |
| GTEX-14PJ4-0626-SM-6AJBS | Lung | TCGA-LUSC | 56.68 | 12.48 |
| GTEX-14PJ5-0226-SM-5YY99 | Lung | TCGA-LUSC | 51.97 | 12.64 |
| GTEX-14PJM-0926-SM-6AJ9Y | Lung | TCGA-LUSC | 39.5 | 7.8 |
| GTEX-14PJO-0926-SM-686YT | Lung | TCGA-LUSC | 50.93 | 12.39 |
| GTEX-14PK6-0326-SM-6AJ9S | Lung | TCGA-LUSC | 55.6 | 14.31 |
| GTEX-14PQA-1126-SM-7KUM4 | Lung | TCGA-LUSC | 49.51 | 17.05 |
| GTEX-14XAO-0526-SM-6AJB7 | Lung | TCGA-LUSC | 65.1 | 19.56 |
| GTEX-15CHC-0226-SM-5YYBB | Lung | TCGA-LUSC | 36.12 | 8.2 |

| | | | | |
|---|---|---|---|---|
| GTEX-15CHR-0726-SM-7EPHG | Lung | TCGA-LUSC | 59.52 | 21.74 |
| GTEX-15D1Q-0526-SM-6AJAY | Lung | TCGA-LUSC | 33.99 | 9.94 |
| GTEX-15EOM-5010-SM-7P8PA | Lung | TCGA-LUSC | 81.94 | 34.73 |
| GTEX-15ER7-0926-SM-7KUMG | Lung | TCGA-LUSC | 42.09 | 5.29 |
| GTEX-15EU6-1226-SM-6AJBE | Lung | TCGA-LUSC | 52.82 | 11.28 |
| GTEX-15FZZ-0326-SM-6M48O | Lung | TCGA-LUSC | 58.57 | 20.13 |
| GTEX-15G1A-0426-SM-6M468 | Lung | TCGA-LUSC | 66.2 | 18.64 |
| GTEX-15RIE-0326-SM-6PAMC | Lung | TCGA-LUSC | 47.65 | 12.7 |
| GTEX-15RJ7-0626-SM-6M47V | Lung | TCGA-LUSC | 55.74 | 13.72 |
| GTEX-15RJE-1026-SM-6M46X | Lung | TCGA-LUSC | 33.79 | 8.95 |
| GTEX-15SB6-0426-SM-6LPJ5 | Lung | TCGA-LUSC | 38.99 | 9.37 |
| GTEX-15SHV-0326-SM-6M475 | Lung | TCGA-LUSC | 52.07 | 9.97 |
| GTEX-15SHW-0926-SM-6LPIO | Lung | TCGA-LUSC | 50.62 | 25.83 |
| GTEX-15SKB-0826-SM-7KUFB | Lung | TCGA-LUSC | 38.93 | 10.63 |
| GTEX-15UKP-1926-SM-6LPI9 | Lung | TCGA-LUSC | 68.63 | 14.49 |
| GTEX-169BO-0226-SM-79OL1 | Lung | TCGA-LUSC | 35.48 | 9.92 |
| GTEX-16AAH-0426-SM-7DUFM | Lung | TCGA-LUSC | 30.67 | 11.07 |
| GTEX-16GPK-1826-SM-7MGW5 | Lung | TCGA-LUSC | 42.81 | 15.69 |
| GTEX-16MT8-1026-SM-6LPK1 | Lung | TCGA-LUSC | 39.98 | 8.75 |
| GTEX-16MTA-1226-SM-7KULL | Lung | TCGA-LUSC | 52.37 | 10.36 |
| GTEX-16NGA-0226-SM-718AI | Lung | TCGA-LUSC | 42.53 | 10.06 |
| GTEX-16XZZ-0926-SM-7DHLI | Lung | TCGA-LUSC | 22.59 | 4.7 |
| GTEX-16Z82-1026-SM-6M48A | Lung | TCGA-LUSC | 33.14 | 8.92 |
| GTEX-178AV-0326-SM-6LPJF | Lung | TCGA-LUSC | 46.73 | 12.66 |
| GTEX-17EVP-0726-SM-7EWDX | Lung | TCGA-LUSC | 41.64 | 9.04 |
| GTEX-17EVQ-1526-SM-79ONG | Lung | TCGA-LUSC | 28.39 | 6.84 |
| GTEX-17F96-0626-SM-793CC | Lung | TCGA-LUSC | 51.56 | 12.07 |
| GTEX-17F98-0226-SM-793BV | Lung | TCGA-LUSC | 46.28 | 12.57 |
| GTEX-17F9Y-1026-SM-7IGO4 | Lung | TCGA-LUSC | 47.54 | 13.01 |
| GTEX-17GQL-0726-SM-731BL | Lung | TCGA-LUSC | 42.8 | 10.89 |
| GTEX-17HG3-0326-SM-7IGP4 | Lung | TCGA-LUSC | 40.22 | 9.07 |

| | | | | |
|---|---|---|---|---|
| GTEX-17HGU-0926-SM-79OKO | Lung | TCGA-LUSC | 61.93 | 26.21 |
| GTEX-17HHE-0626-SM-7DHL6 | Lung | TCGA-LUSC | 38.61 | 11.94 |
| GTEX-17HII-0926-SM-79ON7 | Lung | TCGA-LUSC | 37.64 | 10.05 |
| GTEX-17JCI-1526-SM-7IGOQ | Lung | TCGA-LUSC | 38.98 | 9.75 |
| GTEX-17KNJ-0926-SM-7IGP2 | Lung | TCGA-LUSC | 36.53 | 12.34 |
| GTEX-17MF6-0826-SM-7LT8F | Lung | TCGA-LUSC | 31.61 | 3.65 |
| GTEX-183FY-0726-SM-793C3 | Lung | TCGA-LUSC | 56.21 | 25.88 |
| GTEX-18465-0626-SM-7LT8X | Lung | TCGA-LUSC | 29.54 | 7.04 |
| GTEX-18A66-0926-SM-718BG | Lung | TCGA-LUSC | 54.43 | 28.43 |
| GTEX-18A67-1126-SM-7KFSB | Lung | TCGA-LUSC | 58.3 | 16.73 |
| GTEX-18A6Q-0826-SM-7KFRD | Lung | TCGA-LUSC | 57.1 | 14.36 |
| GTEX-18A7A-1026-SM-7LT8B | Lung | TCGA-LUSC | 40.58 | 6.93 |
| GTEX-18D9A-0226-SM-7KFSJ | Lung | TCGA-LUSC | 67.5 | 18.03 |
| GTEX-18D9B-1026-SM-CNPO5 | Lung | TCGA-LUSC | 35.98 | 13.19 |
| GTEX-18D9U-0526-SM-72D7C | Lung | TCGA-LUSC | 67.19 | 21.39 |
| GTEX-18QFQ-0926-SM-7LG4V | Lung | TCGA-LUSC | 37.6 | 7.09 |
| GTEX-1A32A-0726-SM-731D4 | Lung | TCGA-LUSC | 66.51 | 8.93 |
| GTEX-1A3MV-0526-SM-72D5A | Lung | TCGA-LUSC | 49.12 | 15.76 |
| GTEX-1A8FM-0826-SM-793D5 | Lung | TCGA-LUSC | 26.17 | 10.11 |
| GTEX-1A8G6-0726-SM-73KV5 | Lung | TCGA-LUSC | 33.58 | 4.99 |
| GTEX-1AX8Z-5010-SM-AHZ33 | Lung | TCGA-LUSC | 47.51 | 18.12 |
| GTEX-1AX9I-0826-SM-73KUT | Lung | TCGA-LUSC | 27.36 | 8.58 |
| GTEX-1AX9J-1626-SM-73KUI | Lung | TCGA-LUSC | 39.25 | 4.07 |
| GTEX-1AX9K-0826-SM-731F9 | Lung | TCGA-LUSC | 46.96 | 10.79 |
| GTEX-1AYCT-0726-SM-7IGNG | Lung | TCGA-LUSC | 54.43 | 17.97 |
| GTEX-1AYD5-1226-SM-7EWEP | Lung | TCGA-LUSC | 26.41 | 11.19 |
| GTEX-1B8KZ-0526-SM-73KW2 | Lung | TCGA-LUSC | 49.38 | 11.92 |
| GTEX-1B8L1-0626-SM-7EPHL | Lung | TCGA-LUSC | 39.78 | 5.6 |
| GTEX-1B8SF-0826-SM-73KW8 | Lung | TCGA-LUSC | 22.29 | 5.85 |
| GTEX-1B932-0726-SM-731EY | Lung | TCGA-LUSC | 43.27 | 13.29 |
| GTEX-1B933-0926-SM-9OSVH | Lung | TCGA-LUSC | 29.89 | 6.17 |

| | | | | |
|---|---|---|---|---|
| GTEX-1B97I-0226-SM-73KVO | Lung | TCGA-LUSC | 51.87 | 12.06 |
| GTEX-1B996-0726-SM-7IGPI | Lung | TCGA-LUSC | 72.81 | 8.69 |
| GTEX-1BAJH-1326-SM-7RHFS | Lung | TCGA-LUSC | 33.7 | 7.57 |
| GTEX-1C4CL-0826-SM-7EWEZ | Lung | TCGA-LUSC | 42.45 | 6.99 |
| GTEX-1C64O-1226-SM-79OOC | Lung | TCGA-LUSC | 28.64 | 4.65 |
| GTEX-1C6VR-0626-SM-79OOY | Lung | TCGA-LUSC | 26.85 | 7.09 |
| GTEX-1C6VS-1226-SM-79OO2 | Lung | TCGA-LUSC | 24.46 | 7.26 |
| GTEX-1C6WA-0726-SM-7IGQ5 | Lung | TCGA-LUSC | 32.25 | 5.98 |
| GTEX-1CAMQ-1026-SM-7EPIC | Lung | TCGA-LUSC | 41.23 | 9.56 |
| GTEX-1CAV2-0626-SM-7P8QE | Lung | TCGA-LUSC | 55.22 | 12.17 |
| GTEX-1CB4F-0926-SM-7DHMJ | Lung | TCGA-LUSC | 49.35 | 20.34 |
| GTEX-1CB4I-2126-SM-793AG | Lung | TCGA-LUSC | 40.73 | 11.26 |
| GTEX-1CB4J-1726-SM-79OOL | Lung | TCGA-LUSC | 43.59 | 12.97 |
| GTEX-1E1VI-0926-SM-7P8T3 | Lung | TCGA-LUSC | 26.68 | 5.85 |
| GTEX-1E2YA-0926-SM-7MXUK | Lung | TCGA-LUSC | 29.89 | 3.64 |
| GTEX-1EKGG-1326-SM-7MXUM | Lung | TCGA-LUSC | 38.82 | 9.63 |
| GTEX-1EMGI-1426-SM-7IGNQ | Lung | TCGA-LUSC | 50.66 | 8.64 |
| GTEX-1EN7A-0826-SM-7MKFV | Lung | TCGA-LUSC | 38.67 | 11.91 |
| GTEX-1EWIQ-0926-SM-7RHG7 | Lung | TCGA-LUSC | 43.25 | 11.19 |
| GTEX-1EX96-0226-SM-7RHHR | Lung | TCGA-LUSC | 38.62 | 9.88 |
| GTEX-1F48J-0826-SM-9WPOD | Lung | TCGA-LUSC | 37.16 | 9.86 |
| GTEX-1F5PK-0526-SM-7MXUA | Lung | TCGA-LUSC | 39.66 | 12.66 |
| GTEX-1F6I4-1026-SM-7P8PS | Lung | TCGA-LUSC | 15.87 | 4.28 |
| GTEX-1F6RS-0826-SM-7P8T9 | Lung | TCGA-LUSC | 62.81 | 13.92 |
| GTEX-1F75B-1026-SM-9JGFS | Lung | TCGA-LUSC | 33.88 | 9.06 |
| GTEX-1F75W-0626-SM-7PC13 | Lung | TCGA-LUSC | 28.39 | 9.47 |
| GTEX-1F7RK-0826-SM-7SB8V | Lung | TCGA-LUSC | 45.21 | 17.56 |
| GTEX-1F88F-0926-SM-7RHH8 | Lung | TCGA-LUSC | 26.73 | 5.27 |
| GTEX-1FIGZ-0326-SM-7RHGT | Lung | TCGA-LUSC | 67.25 | 16.08 |
| GTEX-1GF9V-1026-SM-9KNW3 | Lung | TCGA-LUSC | 42.59 | 8.34 |
| GTEX-1GF9X-0326-SM-7P8R9 | Lung | TCGA-LUSC | 52.56 | 13.09 |

| | | | | |
|---|---|---|---|---|
| GTEX-1GL5R-0526-SM-9WPOQ | Lung | TCGA-LUSC | 44.66 | 9.12 |
| GTEX-1GMR2-0126-SM-9JGHC | Lung | TCGA-LUSC | 46.98 | 7.9 |
| GTEX-1GMR3-0226-SM-7RHIB | Lung | TCGA-LUSC | 48.56 | 15.15 |
| GTEX-1GMR8-1426-SM-7RHHL | Lung | TCGA-LUSC | 51.58 | 16.37 |
| GTEX-1GMRU-1426-SM-9WPPL | Lung | TCGA-LUSC | 28.96 | 7.28 |
| GTEX-1GN1U-1026-SM-9WYUP | Lung | TCGA-LUSC | 39.13 | 12.54 |
| GTEX-1GN1W-1026-SM-9OSW2 | Lung | TCGA-LUSC | 61.71 | 15.65 |
| GTEX-1GN2E-1526-SM-7P8TD | Lung | TCGA-LUSC | 36.15 | 7.27 |
| GTEX-1GN73-1026-SM-9KNV9 | Lung | TCGA-LUSC | 59.66 | 7.34 |
| GTEX-1GPI7-1526-SM-9JGG8 | Lung | TCGA-LUSC | 61.21 | 12.56 |
| GTEX-1GTWX-0626-SM-9WYU1 | Lung | TCGA-LUSC | 53.63 | 20.17 |
| GTEX-1GZ2Q-0726-SM-9KNVV | Lung | TCGA-LUSC | 33.72 | 6.04 |
| GTEX-1GZ4H-0426-SM-7MXV5 | Lung | TCGA-LUSC | 36.98 | 6 |
| GTEX-1GZ4I-0726-SM-9QEHL | Lung | TCGA-LUSC | 34.72 | 7.51 |
| GTEX-1GZHY-0826-SM-9OSXT | Lung | TCGA-LUSC | 31.12 | 6.98 |
| GTEX-1H11D-0726-SM-9OSWA | Lung | TCGA-LUSC | 39.36 | 12.9 |
| GTEX-1H1CY-0626-SM-9KNVE | Lung | TCGA-LUSC | 35.57 | 9.53 |
| GTEX-1H1DG-1026-SM-9MQJS | Lung | TCGA-LUSC | 27.95 | 6.83 |
| GTEX-1H1E6-0726-SM-9WPQ4 | Lung | TCGA-LUSC | 38.39 | 9.37 |
| GTEX-1H1ZS-0826-SM-A9SMD | Lung | TCGA-LUSC | 37.41 | 10.42 |
| GTEX-1H23P-1026-SM-9QEHQ | Lung | TCGA-LUSC | 48 | 12.88 |
| GTEX-1H2FU-1226-SM-9KNVY | Lung | TCGA-LUSC | 41.3 | 10.81 |
| GTEX-1H3O1-0326-SM-9WPO8 | Lung | TCGA-LUSC | 36.17 | 5.32 |
| GTEX-1H3VY-1426-SM-9WPOB | Lung | TCGA-LUSC | 36.17 | 8.14 |
| GTEX-1HB9E-0726-SM-AHZ4L | Lung | TCGA-LUSC | 32 | 7.83 |
| GTEX-1HBPH-1326-SM-9WYSN | Lung | TCGA-LUSC | 44.3 | 10.62 |
| GTEX-1HBPM-0826-SM-9WYSP | Lung | TCGA-LUSC | 32.56 | 9.37 |
| GTEX-1HCU7-0926-SM-A96TK | Lung | TCGA-LUSC | 57.07 | 22.38 |
| GTEX-1HCUA-0226-SM-A8N88 | Lung | TCGA-LUSC | 43.62 | 9.12 |
| GTEX-1HCVE-0626-SM-A9SL2 | Lung | TCGA-LUSC | 37.54 | 10.79 |
| GTEX-1HFI6-0926-SM-9WPPV | Lung | TCGA-LUSC | 43.02 | 12.25 |

| | | | | |
|---|---|---|---|---|
| GTEX-1HSKV-1126-SM-ADEIA | Lung | TCGA-LUSC | 40.64 | 8.36 |
| GTEX-1HSMO-0926-SM-A96S7 | Lung | TCGA-LUSC | 35.88 | 8.72 |
| GTEX-1HSMP-1326-SM-A96TR | Lung | TCGA-LUSC | 42.38 | 17.74 |
| GTEX-1HSMQ-0726-SM-B2LXY | Lung | TCGA-LUSC | 45.14 | 11.19 |
| GTEX-1HT8W-1426-SM-CE6S2 | Lung | TCGA-LUSC | 45.09 | 11.75 |
| GTEX-1I19N-0526-SM-A9G27 | Lung | TCGA-LUSC | 36.77 | 9.34 |
| GTEX-1I1CD-0926-SM-CL55I | Lung | TCGA-LUSC | 28.68 | 10.46 |
| GTEX-1I1GP-0526-SM-ARL7G | Lung | TCGA-LUSC | 74.28 | 12.94 |
| GTEX-1I1GQ-1126-SM-CNNQT | Lung | TCGA-LUSC | 37.39 | 5.09 |
| GTEX-1I1GR-0726-SM-B2LWL | Lung | TCGA-LUSC | 25.44 | 7.13 |
| GTEX-1I1GS-1426-SM-COH3W | Lung | TCGA-LUSC | 42.49 | 14.24 |
| GTEX-1I1GU-0526-SM-B2LX2 | Lung | TCGA-LUSC | 58.65 | 14.42 |
| GTEX-1I1GV-0926-SM-B2LXM | Lung | TCGA-LUSC | 44.9 | 13 |
| GTEX-1I4MK-0326-SM-B2LWP | Lung | TCGA-LUSC | 42.52 | 10.24 |
| GTEX-1I6K6-0926-SM-AHZ2N | Lung | TCGA-LUSC | 49.45 | 11.09 |
| GTEX-1I6K7-1226-SM-B2LVS | Lung | TCGA-LUSC | 33.38 | 8.94 |
| GTEX-1ICLY-0926-SM-CM2RJ | Lung | TCGA-LUSC | 23.24 | 7.75 |
| GTEX-1ICLZ-0926-SM-CKZPQ | Lung | TCGA-LUSC | 40.2 | 12.49 |
| GTEX-1IDJH-0726-SM-ARU7U | Lung | TCGA-LUSC | 50.37 | 9.54 |
| GTEX-1IDJI-0626-SM-C1YQD | Lung | TCGA-LUSC | 48.61 | 22.8 |
| GTEX-1IDJU-0226-SM-CKZOF | Lung | TCGA-LUSC | 50.4 | 10.38 |
| GTEX-1IKJJ-0926-SM-A9G2L | Lung | TCGA-LUSC | 39.81 | 10.09 |
| GTEX-1IKK5-1926-SM-A9G2V | Lung | TCGA-LUSC | 33.87 | 4.76 |
| GTEX-1IL2U-0926-SM-CKZOB | Lung | TCGA-LUSC | 27.15 | 5.65 |
| GTEX-1IL2V-0726-SM-ARZMB | Lung | TCGA-LUSC | 23.31 | 6.19 |
| GTEX-1IOXB-1126-SM-CNNQ5 | Lung | TCGA-LUSC | 26.75 | 11.34 |
| GTEX-1IY9M-1126-SM-A9SLV | Lung | TCGA-LUSC | 45.43 | 13 |
| GTEX-1J1OQ-0926-SM-CJI34 | Lung | TCGA-LUSC | 49.41 | 15.05 |
| GTEX-1J1R8-0226-SM-AHZ3C | Lung | TCGA-LUSC | 40.95 | 8.88 |
| GTEX-1J8EW-1026-SM-A96T7 | Lung | TCGA-LUSC | 25.08 | 3.35 |
| GTEX-1J8JJ-1126-SM-CL55P | Lung | TCGA-LUSC | 29.44 | 7.69 |

| | | | | |
|---|---|---|---|---|
| GTEX-1J8QM-0526-SM-AHZ3Y | Lung | TCGA-LUSC | 41.4 | 11.37 |
| GTEX-1JJE9-0926-SM-CNNP9 | Lung | TCGA-LUSC | 56.99 | 25.53 |
| GTEX-1JJEA-1126-SM-CKZOC | Lung | TCGA-LUSC | 30.69 | 8.37 |
| GTEX-1JK1U-1126-SM-CY8HK | Lung | TCGA-LUSC | 49.87 | 11.47 |
| GTEX-1JKYN-2126-SM-ARU8W | Lung | TCGA-LUSC | 60.65 | 11.73 |
| GTEX-1JMLX-0926-SM-CNNPQ | Lung | TCGA-LUSC | 32.03 | 5.48 |
| GTEX-1JMOU-0526-SM-CNNPE | Lung | TCGA-LUSC | 50.82 | 22.13 |
| GTEX-1JMPZ-1126-SM-ARU8X | Lung | TCGA-LUSC | 47.08 | 19.5 |
| GTEX-1JMQI-1126-SM-ARZN2 | Lung | TCGA-LUSC | 34.98 | 11.99 |
| GTEX-1JMQJ-2126-SM-CNPPA | Lung | TCGA-LUSC | 20.35 | 7.1 |
| GTEX-1JMQL-1126-SM-CKZOP | Lung | TCGA-LUSC | 40.21 | 9.8 |
| GTEX-1JN6P-0726-SM-ARZN7 | Lung | TCGA-LUSC | 38.42 | 7.6 |
| GTEX-1JN76-0826-SM-C1YRK | Lung | TCGA-LUSC | 41.08 | 11.95 |
| GTEX-1K2DU-0526-SM-D4P2Y | Lung | TCGA-LUSC | 47.24 | 10.99 |
| GTEX-1K9T9-0926-SM-D3L9C | Lung | TCGA-LUSC | 60.25 | 22.62 |
| GTEX-1KAFJ-0326-SM-DHXJG | Lung | TCGA-LUSC | 41.14 | 7.9 |
| GTEX-1KANB-1026-SM-D3L9F | Lung | TCGA-LUSC | 36.63 | 9.36 |
| GTEX-1KD4Q-0926-SM-CXZKI | Lung | TCGA-LUSC | 30.66 | 8.82 |
| GTEX-1KD5A-1126-SM-CXZK6 | Lung | TCGA-LUSC | 31.62 | 11.31 |
| GTEX-1KWVE-0426-SM-DHXJX | Lung | TCGA-LUSC | 36.38 | 6.21 |
| GTEX-1KXAM-0426-SM-DHXKG | Lung | TCGA-LUSC | 44.42 | 12.78 |
| GTEX-1L5NE-0726-SM-CXZKQ | Lung | TCGA-LUSC | 46.57 | 14.46 |
| GTEX-1LB8K-0826-SM-D4P41 | Lung | TCGA-LUSC | 41.13 | 11.25 |
| GTEX-1LC47-1126-SM-D4P43 | Lung | TCGA-LUSC | 50.16 | 19.91 |
| GTEX-1LG7Y-0826-SM-E9TJX | Lung | TCGA-LUSC | 41.29 | 10.2 |
| GTEX-1LGRB-0226-SM-CNPQK | Lung | TCGA-LUSC | 62.95 | 19.63 |
| GTEX-1LSVX-0626-SM-E9U59 | Lung | TCGA-LUSC | 43.07 | 10.69 |
| GTEX-1LVAN-1126-SM-CNNRE | Lung | TCGA-LUSC | 30.96 | 7.34 |
| GTEX-1LVAO-1226-SM-DIPGB | Lung | TCGA-LUSC | 27.76 | 6.97 |
| GTEX-1M4P7-0826-SM-E9TJR | Lung | TCGA-LUSC | 28.93 | 6.82 |
| GTEX-1MA7X-0626-SM-E9TJI | Lung | TCGA-LUSC | 23.22 | 4.98 |

| | | | | |
|---|---|---|---|---|
| GTEX-1MCC2-0726-SM-EV7AL | Lung | TCGA-LUSC | 46.63 | 11.2 |
| GTEX-1MGNQ-0826-SM-EVR41 | Lung | TCGA-LUSC | 46.47 | 15.41 |
| GTEX-1MJIX-0626-SM-DTX8V | Lung | TCGA-LUSC | 36.3 | 7.14 |
| GTEX-1MJK2-1126-SM-E9TJ7 | Lung | TCGA-LUSC | 27.04 | 10.12 |
| GTEX-1N2DW-1126-SM-EVR59 | Lung | TCGA-LUSC | 34.94 | 7.82 |
| GTEX-1NSGN-1626-SM-EXOJF | Lung | TCGA-LUSC | 40.62 | 11.14 |
| GTEX-1NV5F-0926-SM-DTX8N | Lung | TCGA-LUSC | 63.22 | 20.54 |
| GTEX-1OJC3-0726-SM-DTX92 | Lung | TCGA-LUSC | 38.5 | 6.9 |
| GTEX-1OJC4-0626-SM-DTXEY | Lung | TCGA-LUSC | 18.79 | 4.01 |
| GTEX-1P4AB-1026-SM-DTXES | Lung | TCGA-LUSC | 31.63 | 8.19 |
| GTEX-1PIIG-1826-SM-DTX7N | Lung | TCGA-LUSC | 29.26 | 4.39 |
| GTEX-1PPGY-1826-SM-EXOJ8 | Lung | TCGA-LUSC | 29.66 | 8.95 |
| GTEX-1PPH7-0226-SM-E76Q1 | Lung | TCGA-LUSC | 35.04 | 9.48 |
| GTEX-1PWST-0826-SM-DTXF1 | Lung | TCGA-LUSC | 59.36 | 14.75 |
| GTEX-1QAET-0426-SM-EXUSM | Lung | TCGA-LUSC | 35.47 | 13.55 |
| GTEX-1QCLY-1026-SM-EWRNI | Lung | TCGA-LUSC | 44.17 | 11.2 |
| GTEX-1QEPI-0826-SM-DTX9B | Lung | TCGA-LUSC | 39.33 | 11.77 |
| GTEX-1QMI2-0926-SM-DTX7S | Lung | TCGA-LUSC | 43.1 | 10.22 |
| GTEX-1QP29-1226-SM-E6CQ1 | Lung | TCGA-LUSC | 68.73 | 19.56 |
| GTEX-1QP2A-1226-SM-EVR3I | Lung | TCGA-LUSC | 47.51 | 10.85 |
| GTEX-1R9PM-0426-SM-EVR3P | Lung | TCGA-LUSC | 47.23 | 12.13 |
| GTEX-1RAZQ-1026-SM-EAZ4R | Lung | TCGA-LUSC | 69.34 | 22.94 |
| GTEX-1RAZR-0726-SM-EV7BA | Lung | TCGA-LUSC | 51.47 | 13.64 |
| GTEX-1RAZS-2226-SM-E8VMU | Lung | TCGA-LUSC | 56.97 | 16.96 |
| GTEX-1RDX4-1626-SM-EVR53 | Lung | TCGA-LUSC | 45.84 | 11.96 |
| GTEX-1RLM8-0626-SM-E76OW | Lung | TCGA-LUSC | 36.6 | 11.25 |
| GTEX-1RNSC-1226-SM-E9U52 | Lung | TCGA-LUSC | 30.66 | 7.79 |
| GTEX-1RQED-1026-SM-EVYCT | Lung | TCGA-LUSC | 33.55 | 7.76 |
| GTEX-1S5VW-0926-SM-EWRNU | Lung | TCGA-LUSC | 77.56 | 21.91 |
| GTEX-1S82P-0826-SM-EVR4P | Lung | TCGA-LUSC | 56.7 | 9.05 |
| GTEX-N7MS-0926-SM-2HMIZ | Lung | TCGA-LUSC | 24.01 | 5.16 |

| | | | | |
|---|---|---|---|---|
| GTEX-N7MT-0126-SM-2D7VT | Lung | TCGA-LUSC | 35.43 | 12.47 |
| GTEX-NFK9-1026-SM-2HMK1 | Lung | TCGA-LUSC | 33.11 | 6.33 |
| GTEX-NPJ8-0326-SM-2D7VV | Lung | TCGA-LUSC | 47.62 | 10.99 |
| GTEX-O5YT-0526-SM-32PK8 | Lung | TCGA-LUSC | 53.99 | 27.87 |
| GTEX-O5YV-0526-SM-2I5GE | Lung | TCGA-LUSC | 45.35 | 16 |
| GTEX-O5YW-0526-SM-2YUMX | Lung | TCGA-LUSC | 36.75 | 7.87 |
| GTEX-OHPL-0526-SM-3NM8U | Lung | TCGA-LUSC | 52.12 | 17.24 |
| GTEX-OHPM-0526-SM-2YUMJ | Lung | TCGA-LUSC | 57.97 | 14.82 |
| GTEX-OIZF-0526-SM-7MXVM | Lung | TCGA-LUSC | 46.46 | 10.35 |
| GTEX-OIZG-0526-SM-2HMLF | Lung | TCGA-LUSC | 42.02 | 11.56 |
| GTEX-OIZH-0526-SM-2HMKV | Lung | TCGA-LUSC | 51.29 | 17.72 |
| GTEX-OIZI-1026-SM-3NB1K | Lung | TCGA-LUSC | 44.81 | 8.33 |
| GTEX-OOBJ-0526-SM-48TDK | Lung | TCGA-LUSC | 48.89 | 11.72 |
| GTEX-OOBK-0526-SM-2HMJJ | Lung | TCGA-LUSC | 58.08 | 20.63 |
| GTEX-OXRK-0926-SM-2HMKP | Lung | TCGA-LUSC | 44.95 | 9.81 |
| GTEX-OXRL-0526-SM-2I3EZ | Lung | TCGA-LUSC | 41.58 | 12.96 |
| GTEX-OXRN-0526-SM-2I5EN | Lung | TCGA-LUSC | 26.65 | 8.14 |
| GTEX-OXRO-0326-SM-33HBM | Lung | TCGA-LUSC | 46.27 | 17.23 |
| GTEX-OXRP-0526-SM-2I3EW | Lung | TCGA-LUSC | 47.05 | 16.46 |
| GTEX-P44H-1126-SM-48TBU | Lung | TCGA-LUSC | 29.61 | 6.04 |
| GTEX-P4PP-0526-SM-2HMKE | Lung | TCGA-LUSC | 48.5 | 12.68 |
| GTEX-P4PQ-0526-SM-2HMKR | Lung | TCGA-LUSC | 41.44 | 13.8 |
| GTEX-P4QS-0526-SM-2I3ET | Lung | TCGA-LUSC | 42.3 | 19.44 |
| GTEX-P4QT-0526-SM-2I3EX | Lung | TCGA-LUSC | 39.29 | 9.79 |
| GTEX-P78B-0926-SM-2I5FA | Lung | TCGA-LUSC | 40.81 | 12.82 |
| GTEX-PLZ4-0726-SM-2TC6Q | Lung | TCGA-LUSC | 38.31 | 19.5 |
| GTEX-PLZ5-0726-SM-2I5F9 | Lung | TCGA-LUSC | 44.71 | 14.1 |
| GTEX-PLZ6-0426-SM-5IJDW | Lung | TCGA-LUSC | 55.86 | 19.18 |
| GTEX-POMQ-0526-SM-3GADD | Lung | TCGA-LUSC | 44.97 | 18.12 |
| GTEX-POYW-1226-SM-2XCEP | Lung | TCGA-LUSC | 68.18 | 15.41 |
| GTEX-PSDG-1126-SM-2S1ON | Lung | TCGA-LUSC | 21.2 | 5.97 |

| | | | | |
|---|---|---|---|---|
| GTEX-PVOW-1026-SM-2XCF9 | Lung | TCGA-LUSC | 53.15 | 21.03 |
| GTEX-PW2O-0526-SM-2I3DX | Lung | TCGA-LUSC | 31.57 | 9.74 |
| GTEX-PX3G-0526-SM-2I3EM | Lung | TCGA-LUSC | 40.1 | 15.08 |
| GTEX-Q2AG-1026-SM-33HBW | Lung | TCGA-LUSC | 27.09 | 7.52 |
| GTEX-Q2AH-0426-SM-2I3EP | Lung | TCGA-LUSC | 46.34 | 16.52 |
| GTEX-Q734-0626-SM-2I3EF | Lung | TCGA-LUSC | 40.9 | 10.87 |
| GTEX-QCQG-0326-SM-2I3ES | Lung | TCGA-LUSC | 45.26 | 15.05 |
| GTEX-QDT8-0926-SM-32PL2 | Lung | TCGA-LUSC | 50.52 | 16.29 |
| GTEX-QDVJ-0926-SM-2I5FU | Lung | TCGA-LUSC | 48.63 | 17.85 |
| GTEX-QDVN-0726-SM-4B64L | Lung | TCGA-LUSC | 50.03 | 18.74 |
| GTEX-QEG4-0526-SM-48TZD | Lung | TCGA-LUSC | 27 | 7.39 |
| GTEX-QEG5-1126-SM-33HC2 | Lung | TCGA-LUSC | 32.62 | 12.56 |
| GTEX-QEL4-0826-SM-3GAF2 | Lung | TCGA-LUSC | 55.22 | 19.79 |
| GTEX-QESD-0626-SM-2I5G4 | Lung | TCGA-LUSC | 76.24 | 16.31 |
| GTEX-QMR6-1926-SM-32PL9 | Lung | TCGA-LUSC | 19.05 | 9.76 |
| GTEX-QMRM-0826-SM-3NB33 | Lung | TCGA-LUSC | 39.17 | 12.96 |
| GTEX-QV44-0926-SM-2S1RH | Lung | TCGA-LUSC | 45.23 | 20.11 |
| GTEX-QVUS-2026-SM-EYYW9 | Lung | TCGA-LUSC | 40.32 | 10.37 |
| GTEX-QXCU-0626-SM-2TC69 | Lung | TCGA-LUSC | 68.51 | 6.29 |
| GTEX-R3RS-1026-SM-3GADF | Lung | TCGA-LUSC | 44.08 | 13.08 |
| GTEX-R55C-0526-SM-3GIKA | Lung | TCGA-LUSC | 48.18 | 23.05 |
| GTEX-R55D-0926-SM-3GAEU | Lung | TCGA-LUSC | 49.94 | 13.07 |
| GTEX-R55G-0826-SM-2TC5U | Lung | TCGA-LUSC | 41.01 | 17.36 |
| GTEX-REY6-0426-SM-2TF5G | Lung | TCGA-LUSC | 33.95 | 14.74 |
| GTEX-RM2N-0426-SM-2TF4T | Lung | TCGA-LUSC | 42.71 | 15.23 |
| GTEX-RN64-1226-SM-2TC6E | Lung | TCGA-LUSC | 40.2 | 10.66 |
| GTEX-RNOR-0726-SM-2TF5I | Lung | TCGA-LUSC | 38.84 | 9.11 |
| GTEX-RTLS-0926-SM-2TF5X | Lung | TCGA-LUSC | 33.28 | 8.41 |
| GTEX-RU1J-0126-SM-2TF6Y | Lung | TCGA-LUSC | 41.68 | 13.55 |
| GTEX-RU72-0526-SM-2TF5Z | Lung | TCGA-LUSC | 27.96 | 6.14 |
| GTEX-RUSQ-0626-SM-2TF5V | Lung | TCGA-LUSC | 50.21 | 15.87 |

| | | | | |
|---|---|---|---|---|
| GTEX-RVPV-1726-SM-2NKAQ | Lung | TCGA-LUSC | 54.61 | 10.64 |
| GTEX-RWS6-0226-SM-2XCA9 | Lung | TCGA-LUSC | 43.42 | 15.44 |
| GTEX-RWSA-1126-SM-2XCAZ | Lung | TCGA-LUSC | 71.54 | 12.82 |
| GTEX-S32W-0326-SM-2XCBI | Lung | TCGA-LUSC | 59.5 | 18.7 |
| GTEX-S33H-0626-SM-2XCBJ | Lung | TCGA-LUSC | 64.44 | 26.8 |
| GTEX-S341-0326-SM-2XCAU | Lung | TCGA-LUSC | 40.38 | 12.4 |
| GTEX-S3LF-1126-SM-EVR33 | Lung | TCGA-LUSC | 69.68 | 24.85 |
| GTEX-S7SE-0926-SM-2XCD6 | Lung | TCGA-LUSC | 42.97 | 10.69 |
| GTEX-SE5C-0526-SM-2XCE1 | Lung | TCGA-LUSC | 40.04 | 10.03 |
| GTEX-SIU8-0926-SM-COH1X | Lung | TCGA-LUSC | 25.23 | 3.78 |
| GTEX-SJXC-1026-SM-EYYVA | Lung | TCGA-LUSC | 23.47 | 6.84 |
| GTEX-SN8G-0926-SM-4DM5I | Lung | TCGA-LUSC | 38.47 | 8.13 |
| GTEX-SNOS-0426-SM-32PMH | Lung | TCGA-LUSC | 54.92 | 12.1 |
| GTEX-SUCS-0626-SM-32PM5 | Lung | TCGA-LUSC | 36.39 | 8.88 |
| GTEX-T2IS-0526-SM-32QP9 | Lung | TCGA-LUSC | 50.46 | 12.49 |
| GTEX-T5JC-0826-SM-32PMC | Lung | TCGA-LUSC | 42.64 | 8.35 |
| GTEX-T6MN-0826-SM-32PM4 | Lung | TCGA-LUSC | 37.52 | 6.84 |
| GTEX-T6MO-0426-SM-32QOI | Lung | TCGA-LUSC | 43.4 | 22.5 |
| GTEX-T8EM-0326-SM-3DB7F | Lung | TCGA-LUSC | 50.64 | 16.82 |
| GTEX-TML8-0326-SM-4GICN | Lung | TCGA-LUSC | 48.51 | 18.28 |
| GTEX-TMMY-0926-SM-4TT1Z | Lung | TCGA-LUSC | 52.64 | 8.1 |
| GTEX-TSE9-0726-SM-3DB8C | Lung | TCGA-LUSC | 22.24 | 6.86 |
| GTEX-U3ZH-0526-SM-3DB75 | Lung | TCGA-LUSC | 31.31 | 12.66 |
| GTEX-U3ZM-0426-SM-3DB73 | Lung | TCGA-LUSC | 48.38 | 15.53 |
| GTEX-U3ZN-0626-SM-3DB7U | Lung | TCGA-LUSC | 51.23 | 15.31 |
| GTEX-U412-0826-SM-3DB9K | Lung | TCGA-LUSC | 47.1 | 8.44 |
| GTEX-U8T8-2226-SM-3DB95 | Lung | TCGA-LUSC | 29.02 | 5.91 |
| GTEX-U8XE-1426-SM-3DB8Q | Lung | TCGA-LUSC | 67.52 | 17.25 |
| GTEX-UJHI-0726-SM-3DB92 | Lung | TCGA-LUSC | 26.44 | 6.72 |
| GTEX-UJMC-0726-SM-3GADX | Lung | TCGA-LUSC | 47.98 | 10.17 |
| GTEX-UPJH-0826-SM-4WKFD | Lung | TCGA-LUSC | 34.35 | 8.98 |

| | | | | |
|---|---|---|---|---|
| GTEX-UPK5-1126-SM-3GAEJ | Lung | TCGA-LUSC | 60.44 | 21.59 |
| GTEX-V1D1-0826-SM-3P5ZA | Lung | TCGA-LUSC | 34.72 | 11.83 |
| GTEX-VJWN-1326-SM-EVR2Z | Lung | TCGA-LUSC | 21.29 | 6.29 |
| GTEX-VJYA-0326-SM-3GAEX | Lung | TCGA-LUSC | 62.88 | 18.5 |
| GTEX-VUSG-0926-SM-3GIK6 | Lung | TCGA-LUSC | 65.49 | 18.6 |
| GTEX-W5X1-0526-SM-3GILH | Lung | TCGA-LUSC | 43.86 | 12.02 |
| GTEX-WFG7-0526-SM-3GIKI | Lung | TCGA-LUSC | 51.75 | 15.78 |
| GTEX-WFG8-0926-SM-3GIKJ | Lung | TCGA-LUSC | 48.66 | 12.29 |
| GTEX-WFJO-0326-SM-3GIL3 | Lung | TCGA-LUSC | 50.5 | 12.19 |
| GTEX-WFON-0426-SM-3GIL4 | Lung | TCGA-LUSC | 46.33 | 16.39 |
| GTEX-WH7G-0726-SM-3NMBM | Lung | TCGA-LUSC | 46.24 | 10.39 |
| GTEX-WHPG-1426-SM-3NMBB | Lung | TCGA-LUSC | 54.01 | 22.14 |
| GTEX-WHSB-0326-SM-5FQSD | Lung | TCGA-LUSC | 41.38 | 10.34 |
| GTEX-WK11-0526-SM-3NB3O | Lung | TCGA-LUSC | 34.35 | 12.02 |
| GTEX-WOFM-0126-SM-3MJFE | Lung | TCGA-LUSC | 39.54 | 9.59 |
| GTEX-WRHU-0226-SM-3MJFV | Lung | TCGA-LUSC | 28.2 | 6.63 |
| GTEX-WVJS-0826-SM-4MVNR | Lung | TCGA-LUSC | 34.6 | 10.02 |
| GTEX-WWYW-0926-SM-3NB2Z | Lung | TCGA-LUSC | 26.48 | 6.09 |
| GTEX-WY7C-0426-SM-3NB3C | Lung | TCGA-LUSC | 55.58 | 18.18 |
| GTEX-WYBS-1126-SM-3NMAM | Lung | TCGA-LUSC | 27.95 | 11.01 |
| GTEX-WYJK-0826-SM-3NM8Y | Lung | TCGA-LUSC | 46.94 | 9.67 |
| GTEX-WYVS-0526-SM-3NM9W | Lung | TCGA-LUSC | 67.19 | 23.22 |
| GTEX-WZTO-0426-SM-3NM99 | Lung | TCGA-LUSC | 27.89 | 6.91 |
| GTEX-X15G-0626-SM-EVR32 | Lung | TCGA-LUSC | 45.71 | 13.58 |
| GTEX-X261-1026-SM-3NMDL | Lung | TCGA-LUSC | 60.4 | 18.11 |
| GTEX-X3Y1-0626-SM-3P5YS | Lung | TCGA-LUSC | 35.49 | 7.25 |
| GTEX-X4EO-0926-SM-3P5Z2 | Lung | TCGA-LUSC | 31.69 | 10.18 |
| GTEX-X4EP-0526-SM-3P5YW | Lung | TCGA-LUSC | 50.35 | 18.12 |
| GTEX-X4LF-0526-SM-3NMB6 | Lung | TCGA-LUSC | 35.76 | 8.56 |
| GTEX-X4XX-1026-SM-4QARO | Lung | TCGA-LUSC | 53.05 | 17.18 |
| GTEX-X4XY-1026-SM-46MVX | Lung | TCGA-LUSC | 37.24 | 15.94 |

| | | | | |
|---|---|---|---|---|
| GTEX-X585-1026-SM-46MW6 | Lung | TCGA-LUSC | 34.42 | 11.54 |
| GTEX-X5EB-0426-SM-46MVY | Lung | TCGA-LUSC | 32.24 | 6.64 |
| GTEX-XBEC-1026-SM-4QASM | Lung | TCGA-LUSC | 30.06 | 7.54 |
| GTEX-XBED-0826-SM-47JYC | Lung | TCGA-LUSC | 52.91 | 15.86 |
| GTEX-XBEW-0226-SM-4AT6A | Lung | TCGA-LUSC | 42.93 | 10.27 |
| GTEX-XGQ4-0826-SM-4AT4T | Lung | TCGA-LUSC | 51.5 | 7.82 |
| GTEX-XMD2-1026-SM-4WWE8 | Lung | TCGA-LUSC | 25.65 | 7.9 |
| GTEX-XOT4-1426-SM-4B65T | Lung | TCGA-LUSC | 25.69 | 6.06 |
| GTEX-XPVG-1026-SM-4B64Y | Lung | TCGA-LUSC | 49.57 | 10.36 |
| GTEX-XQ3S-0926-SM-4BOPI | Lung | TCGA-LUSC | 28.64 | 7.48 |
| GTEX-XQ8I-1126-SM-4BOO2 | Lung | TCGA-LUSC | 40.21 | 7.3 |
| GTEX-XV7Q-0426-SM-4BRVN | Lung | TCGA-LUSC | 57.5 | 23.91 |
| GTEX-XXEK-0626-SM-4BRWE | Lung | TCGA-LUSC | 69.53 | 22.69 |
| GTEX-XYKS-0526-SM-4BRW2 | Lung | TCGA-LUSC | 35.02 | 14.17 |
| GTEX-Y111-1026-SM-4TT22 | Lung | TCGA-LUSC | 44.22 | 12.03 |
| GTEX-Y3I4-0426-SM-4TT29 | Lung | TCGA-LUSC | 37.28 | 18.34 |
| GTEX-Y3IK-0626-SM-4WWE4 | Lung | TCGA-LUSC | 39.94 | 9.79 |
| GTEX-Y5LM-0726-SM-4VBRP | Lung | TCGA-LUSC | 54.59 | 15.29 |
| GTEX-Y5V5-0826-SM-4VBQD | Lung | TCGA-LUSC | 47.93 | 10.6 |
| GTEX-Y5V6-0226-SM-4V6G7 | Lung | TCGA-LUSC | 49.07 | 11.41 |
| GTEX-Y8E4-0526-SM-4V6GC | Lung | TCGA-LUSC | 50.91 | 17.68 |
| GTEX-Y8LW-0326-SM-4VBQ9 | Lung | TCGA-LUSC | 37.35 | 9.1 |
| GTEX-Y9LG-0526-SM-4VBRY | Lung | TCGA-LUSC | 56.41 | 18.18 |
| GTEX-YB5E-0726-SM-4VDSH | Lung | TCGA-LUSC | 40.05 | 7.01 |
| GTEX-YEC3-0226-SM-DO11B | Lung | TCGA-LUSC | 48.9 | 11.91 |
| GTEX-YEC4-0526-SM-4W21U | Lung | TCGA-LUSC | 51.55 | 12.35 |
| GTEX-YECK-0926-SM-4W214 | Lung | TCGA-LUSC | 26.36 | 8.24 |
| GTEX-YF7O-0626-SM-4W21R | Lung | TCGA-LUSC | 61.1 | 15.54 |
| GTEX-YFC4-1126-SM-5RQJN | Lung | TCGA-LUSC | 45.79 | 12.77 |
| GTEX-YFCO-0426-SM-4W1Z7 | Lung | TCGA-LUSC | 51.12 | 8.92 |
| GTEX-YJ8O-1826-SM-5HL82 | Lung | TCGA-LUSC | 47.22 | 15.48 |

| | | | | |
|---|---|---|---|---|
| GTEX-ZA64-0326-SM-5HL8T | Lung | TCGA-LUSC | 53.78 | 11.47 |
| GTEX-ZAB4-0626-SM-5CVN3 | Lung | TCGA-LUSC | 36.27 | 13.21 |
| GTEX-ZAB5-0626-SM-5PNVB | Lung | TCGA-LUSC | 48.92 | 10.69 |
| GTEX-ZC5H-0926-SM-5CVMZ | Lung | TCGA-LUSC | 43.91 | 11.48 |
| GTEX-ZDTS-1026-SM-4WAXS | Lung | TCGA-LUSC | 26.09 | 6.46 |
| GTEX-ZDTT-0926-SM-5J2MS | Lung | TCGA-LUSC | 54.85 | 12.5 |
| GTEX-ZDXO-1326-SM-57WBS | Lung | TCGA-LUSC | 25.65 | 6.92 |
| GTEX-ZDYS-0426-SM-5IJEQ | Lung | TCGA-LUSC | 38.89 | 13.15 |
| GTEX-ZE7O-0826-SM-57WCP | Lung | TCGA-LUSC | 41.58 | 14.54 |
| GTEX-ZEX8-0526-SM-DO115 | Lung | TCGA-LUSC | 47.19 | 12.95 |
| GTEX-ZF28-0726-SM-4WKFU | Lung | TCGA-LUSC | 42.44 | 11.01 |
| GTEX-ZF29-1026-SM-4WKGC | Lung | TCGA-LUSC | 53.85 | 10.85 |
| GTEX-ZF2S-0626-SM-4WKH2 | Lung | TCGA-LUSC | 46.09 | 11.63 |
| GTEX-ZF3C-0926-SM-DO92P | Lung | TCGA-LUSC | 39.31 | 15.66 |
| GTEX-ZG7Y-1026-SM-4WWDG | Lung | TCGA-LUSC | 49.23 | 18.41 |
| GTEX-ZLV1-0426-SM-4WWC2 | Lung | TCGA-LUSC | 55.84 | 16.72 |
| GTEX-ZLWG-0626-SM-4WWFR | Lung | TCGA-LUSC | 59.03 | 22.17 |
| GTEX-ZPCL-0926-SM-DNZZ8 | Lung | TCGA-LUSC | 52.39 | 10.71 |
| GTEX-ZPIC-0626-SM-57WDY | Lung | TCGA-LUSC | 49.71 | 19.58 |
| GTEX-ZPU1-0926-SM-57WDO | Lung | TCGA-LUSC | 50.54 | 12.38 |
| GTEX-ZQG8-0326-SM-51MSE | Lung | TCGA-LUSC | 45.77 | 8.79 |
| GTEX-ZT9W-0726-SM-9JGG2 | Lung | TCGA-LUSC | 61.57 | 9.53 |
| GTEX-ZT9X-0326-SM-51MTE | Lung | TCGA-LUSC | 59.36 | 24.53 |
| GTEX-ZTPG-0926-SM-5O99H | Lung | TCGA-LUSC | 69.59 | 15.91 |
| GTEX-ZTTD-1126-SM-51MRP | Lung | TCGA-LUSC | 43.24 | 14.56 |
| GTEX-ZUA1-1026-SM-4YCEA | Lung | TCGA-LUSC | 38.79 | 11.48 |
| GTEX-ZVT3-0926-SM-5GICK | Lung | TCGA-LUSC | 42 | 7.43 |
| GTEX-ZVZQ-1526-SM-5N9G6 | Lung | TCGA-LUSC | 29.57 | 10.17 |
| GTEX-ZXG5-0826-SM-5GID6 | Lung | TCGA-LUSC | 52.47 | 13.63 |
| GTEX-ZY6K-0326-SM-5SIBB | Lung | TCGA-LUSC | 44.39 | 14.31 |
| GTEX-ZYFG-0226-SM-5GIDT | Lung | TCGA-LUSC | 35.69 | 8.8 |

| | | | | |
|---|---|---|---|---|
| GTEX-ZYT6-0526-SM-5GIEA | Lung | TCGA-LUSC | 49.39 | 9.9 |
| GTEX-ZYVF-1726-SM-5E443 | Lung | TCGA-LUSC | 51.65 | 17.34 |
| GTEX-ZYW4-1526-SM-5SIBA | Lung | TCGA-LUSC | 40.15 | 9.49 |
| GTEX-ZYY3-0926-SM-5E454 | Lung | TCGA-LUSC | 43.48 | 8.35 |
| GTEX-ZZPT-1326-SM-5E43H | Lung | TCGA-LUSC | 46.22 | 9.24 |
| GTEX-ZZPU-0526-SM-5E44U | Lung | TCGA-LUSC | 57.87 | 14.85 |
| GTEX-111CU-0526-SM-5EGHK | Pancreas | TCGA-PAAD | 12.94 | 4.02 |
| GTEX-111YS-1226-SM-5EGGJ | Pancreas | TCGA-PAAD | 11.9 | 3.45 |
| GTEX-1122O-0726-SM-5GIEV | Pancreas | TCGA-PAAD | 12.56 | 4.27 |
| GTEX-1128S-0826-SM-5GZZI | Pancreas | TCGA-PAAD | 17.24 | 4.07 |
| GTEX-117YX-0226-SM-5EGH6 | Pancreas | TCGA-PAAD | 18.58 | 6.23 |
| GTEX-11DXX-0926-SM-5H112 | Pancreas | TCGA-PAAD | 12.61 | 2.55 |
| GTEX-11EQ9-1026-SM-5H134 | Pancreas | TCGA-PAAD | 13.39 | 3.88 |
| GTEX-11GSP-0426-SM-5A5KX | Pancreas | TCGA-PAAD | 5.98 | 2.25 |
| GTEX-11I78-0626-SM-5A5LZ | Pancreas | TCGA-PAAD | 11.93 | 2.97 |
| GTEX-11LCK-0226-SM-5A5M6 | Pancreas | TCGA-PAAD | 14.68 | 6.43 |
| GTEX-11NSD-0526-SM-5A5LT | Pancreas | TCGA-PAAD | 10.41 | 3.11 |
| GTEX-11ONC-0526-SM-5BC57 | Pancreas | TCGA-PAAD | 9.8 | 5.21 |
| GTEX-11P7K-0526-SM-5BC5I | Pancreas | TCGA-PAAD | 13.48 | 5.46 |
| GTEX-11TT1-0326-SM-5LUAY | Pancreas | TCGA-PAAD | 21.15 | 8.68 |
| GTEX-11VI4-0426-SM-5EGHZ | Pancreas | TCGA-PAAD | 13.35 | 4.96 |
| GTEX-11XUK-0626-SM-5N9ES | Pancreas | TCGA-PAAD | 13.29 | 4.96 |
| GTEX-1211K-1126-SM-5EGGB | Pancreas | TCGA-PAAD | 14.37 | 3.89 |
| GTEX-12BJ1-1126-SM-5EGGG | Pancreas | TCGA-PAAD | 9.26 | 1.82 |
| GTEX-12WSD-1626-SM-5GCNR | Pancreas | TCGA-PAAD | 6.31 | 3.59 |
| GTEX-12WSG-1026-SM-5EGII | Pancreas | TCGA-PAAD | 8.22 | 2.08 |
| GTEX-12WSK-0226-SM-5BC62 | Pancreas | TCGA-PAAD | 14.05 | 3.93 |
| GTEX-12WSL-0426-SM-5GCNX | Pancreas | TCGA-PAAD | 15.86 | 5.79 |
| GTEX-12WSN-0826-SM-5GCOF | Pancreas | TCGA-PAAD | 14.84 | 4.35 |
| GTEX-131XE-1926-SM-5IFER | Pancreas | TCGA-PAAD | 18.55 | 6.17 |
| GTEX-131XF-1426-SM-5BC68 | Pancreas | TCGA-PAAD | 15.07 | 5.72 |

| | | | | |
|---|---|---|---|---|
| GTEX-131XG-1426-SM-5GCMO | Pancreas | TCGA-PAAD | 11.25 | 3.38 |
| GTEX-132AR-1826-SM-5EGHR | Pancreas | TCGA-PAAD | 21.65 | 5.42 |
| GTEX-132QS-0926-SM-5P9GB | Pancreas | TCGA-PAAD | 21.86 | 4.17 |
| GTEX-1339X-1026-SM-5IFH5 | Pancreas | TCGA-PAAD | 22.07 | 6.26 |
| GTEX-1399R-0426-SM-5IJE3 | Pancreas | TCGA-PAAD | 13.54 | 3.22 |
| GTEX-1399S-0326-SM-5IFFS | Pancreas | TCGA-PAAD | 16.26 | 3.93 |
| GTEX-1399U-0926-SM-5IFHA | Pancreas | TCGA-PAAD | 16.63 | 4.68 |
| GTEX-139YR-1526-SM-5IFJ1 | Pancreas | TCGA-PAAD | 19.03 | 6.88 |
| GTEX-13CF3-1026-SM-5LZWY | Pancreas | TCGA-PAAD | 18.23 | 6.05 |
| GTEX-13D11-2226-SM-5IFEO | Pancreas | TCGA-PAAD | 20.12 | 6.65 |
| GTEX-13FH7-1426-SM-5IFIC | Pancreas | TCGA-PAAD | 18.05 | 6.98 |
| GTEX-13FLV-0626-SM-5IFEY | Pancreas | TCGA-PAAD | 8.96 | 3.43 |
| GTEX-13FTW-0526-SM-5IFIP | Pancreas | TCGA-PAAD | 12.44 | 3.53 |
| GTEX-13FTX-1226-SM-5IFGN | Pancreas | TCGA-PAAD | 14.8 | 3.9 |
| GTEX-13N11-0226-SM-5KM3C | Pancreas | TCGA-PAAD | 21.6 | 6.62 |
| GTEX-13O1R-1826-SM-5KM3B | Pancreas | TCGA-PAAD | 12.1 | 5.14 |
| GTEX-13O61-2126-SM-5IJEO | Pancreas | TCGA-PAAD | 10.23 | 3.09 |
| GTEX-13OVI-0526-SM-5IFFQ | Pancreas | TCGA-PAAD | 10.72 | 3.09 |
| GTEX-13PL7-2226-SM-5L3IC | Pancreas | TCGA-PAAD | 24.11 | 7.24 |
| GTEX-13PVR-0726-SM-5S2PX | Pancreas | TCGA-PAAD | 15.32 | 4.46 |
| GTEX-13SLX-1326-SM-5S2QS | Pancreas | TCGA-PAAD | 8.28 | 3.09 |
| GTEX-13U4I-1526-SM-5IFFF | Pancreas | TCGA-PAAD | 16.63 | 7.54 |
| GTEX-13VXT-1226-SM-5LU3M | Pancreas | TCGA-PAAD | 15 | 4.92 |
| GTEX-13W3W-1026-SM-5IFG4 | Pancreas | TCGA-PAAD | 18.61 | 5.27 |
| GTEX-13X6H-0626-SM-5LU53 | Pancreas | TCGA-PAAD | 18.75 | 5.03 |
| GTEX-13YAN-2126-SM-5Q5C4 | Pancreas | TCGA-PAAD | 20.65 | 10.29 |
| GTEX-144FL-1526-SM-5Q5CA | Pancreas | TCGA-PAAD | 22.69 | 7.6 |
| GTEX-144GM-0726-SM-79OJQ | Pancreas | TCGA-PAAD | 15.35 | 4.61 |
| GTEX-144GN-0826-SM-5LU5G | Pancreas | TCGA-PAAD | 20.9 | 5.25 |
| GTEX-145ME-0326-SM-5S2QO | Pancreas | TCGA-PAAD | 15.82 | 3.46 |
| GTEX-145MN-1426-SM-5SI9H | Pancreas | TCGA-PAAD | 17.82 | 4.75 |

| | | | | |
|---|---|---|---|---|
| GTEX-145MO-2126-SM-5Q5CZ | Pancreas | TCGA-PAAD | 13.48 | 5.89 |
| GTEX-146FQ-1526-SM-5Q5CX | Pancreas | TCGA-PAAD | 12.13 | 3.63 |
| GTEX-146FR-0126-SM-5Q5F3 | Pancreas | TCGA-PAAD | 14.94 | 3.96 |
| GTEX-148VI-0826-SM-5TDDI | Pancreas | TCGA-PAAD | 11.09 | 3.25 |
| GTEX-1497J-0426-SM-5Q5CO | Pancreas | TCGA-PAAD | 22.34 | 8.68 |
| GTEX-14AS3-0326-SM-5Q5DB | Pancreas | TCGA-PAAD | 13.42 | 4.25 |
| GTEX-14BMU-0726-SM-73KXS | Pancreas | TCGA-PAAD | 16.42 | 6.5 |
| GTEX-14C39-1026-SM-73KWG | Pancreas | TCGA-PAAD | 11.06 | 4.76 |
| GTEX-14DAR-0526-SM-664MX | Pancreas | TCGA-PAAD | 12.41 | 4.7 |
| GTEX-14E6E-1426-SM-5RQI2 | Pancreas | TCGA-PAAD | 18.38 | 6.01 |
| GTEX-14ICL-1126-SM-5S2RE | Pancreas | TCGA-PAAD | 15.3 | 4.84 |
| GTEX-14JG6-0426-SM-68715 | Pancreas | TCGA-PAAD | 12.34 | 3.96 |
| GTEX-14PHX-2026-SM-6872C | Pancreas | TCGA-PAAD | 18.02 | 4.72 |
| GTEX-14PHY-1126-SM-664OR | Pancreas | TCGA-PAAD | 13.25 | 3.84 |
| GTEX-14PJ2-1526-SM-686Z4 | Pancreas | TCGA-PAAD | 15.7 | 7.05 |
| GTEX-14PJ3-0426-SM-6LLHF | Pancreas | TCGA-PAAD | 13.88 | 4.1 |
| GTEX-14PJ4-0726-SM-6872E | Pancreas | TCGA-PAAD | 15.21 | 5.3 |
| GTEX-14PJ6-0926-SM-686ZA | Pancreas | TCGA-PAAD | 11.14 | 3.39 |
| GTEX-14PJM-2126-SM-6AJAS | Pancreas | TCGA-PAAD | 4.9 | 1.47 |
| GTEX-14PJN-1526-SM-6AJBP | Pancreas | TCGA-PAAD | 10.1 | 3.03 |
| GTEX-14PJO-1826-SM-69LPR | Pancreas | TCGA-PAAD | 16.36 | 8.08 |
| GTEX-14PK6-1026-SM-664NZ | Pancreas | TCGA-PAAD | 12.81 | 3.32 |
| GTEX-14PKU-0626-SM-6EU1Q | Pancreas | TCGA-PAAD | 13.39 | 5.36 |
| GTEX-14PKV-0926-SM-6AJB4 | Pancreas | TCGA-PAAD | 13.18 | 4.28 |
| GTEX-14PN3-0126-SM-6AJAU | Pancreas | TCGA-PAAD | 12.6 | 3.79 |
| GTEX-14PN4-1626-SM-6AJB5 | Pancreas | TCGA-PAAD | 25.01 | 10.64 |
| GTEX-15CHR-2026-SM-7MXU2 | Pancreas | TCGA-PAAD | 10.79 | 5.01 |
| GTEX-15DYW-1626-SM-6LLI1 | Pancreas | TCGA-PAAD | 6.51 | 3 |
| GTEX-15ER7-1326-SM-6LPK9 | Pancreas | TCGA-PAAD | 10.5 | 4.93 |
| GTEX-15EU6-1126-SM-68721 | Pancreas | TCGA-PAAD | 15.73 | 4.9 |
| GTEX-15FZZ-0626-SM-7KFTO | Pancreas | TCGA-PAAD | 23.25 | 7.04 |

| | | | | |
|---|---|---|---|---|
| GTEX-15RIF-0626-SM-6LLI3 | Pancreas | TCGA-PAAD | 12.85 | 3.36 |
| GTEX-15RJ7-0826-SM-6LLI8 | Pancreas | TCGA-PAAD | 12.33 | 4.06 |
| GTEX-15SB6-0226-SM-6LLIA | Pancreas | TCGA-PAAD | 10.3 | 3.7 |
| GTEX-15SDE-0826-SM-6M463 | Pancreas | TCGA-PAAD | 18.09 | 7.98 |
| GTEX-15SHV-1526-SM-6LLIB | Pancreas | TCGA-PAAD | 11.38 | 3.58 |
| GTEX-15UF6-1826-SM-6LLI9 | Pancreas | TCGA-PAAD | 19.92 | 9.37 |
| GTEX-15UF7-0826-SM-6M46E | Pancreas | TCGA-PAAD | 17.24 | 5.62 |
| GTEX-15UKP-1326-SM-6LPI8 | Pancreas | TCGA-PAAD | 14.07 | 6.33 |
| GTEX-16MT8-1726-SM-7EWE7 | Pancreas | TCGA-PAAD | 14.84 | 10.55 |
| GTEX-16Z82-1726-SM-7MGX3 | Pancreas | TCGA-PAAD | 5.78 | 2.96 |
| GTEX-178AV-0226-SM-793B7 | Pancreas | TCGA-PAAD | 12.47 | 5.14 |
| GTEX-17F98-0426-SM-79ONZ | Pancreas | TCGA-PAAD | 10.93 | 4.89 |
| GTEX-17F9E-1426-SM-793BG | Pancreas | TCGA-PAAD | 12.9 | 4.66 |
| GTEX-17HGU-2026-SM-79OKS | Pancreas | TCGA-PAAD | 13.52 | 6.49 |
| GTEX-17HHE-0526-SM-7DUGR | Pancreas | TCGA-PAAD | 10.51 | 3.64 |
| GTEX-17KNJ-1926-SM-7KFRV | Pancreas | TCGA-PAAD | 15.22 | 6.19 |
| GTEX-17MF6-1426-SM-7IGOX | Pancreas | TCGA-PAAD | 11.84 | 3.57 |
| GTEX-18A66-2126-SM-7189D | Pancreas | TCGA-PAAD | 18.67 | 6.26 |
| GTEX-18A67-1726-SM-7KFT9 | Pancreas | TCGA-PAAD | 22.04 | 7.16 |
| GTEX-18A6Q-1726-SM-7LT9A | Pancreas | TCGA-PAAD | 6.51 | 3.7 |
| GTEX-18A7A-1726-SM-7LT93 | Pancreas | TCGA-PAAD | 5.31 | 1.85 |
| GTEX-18D9U-1626-SM-7KFTW | Pancreas | TCGA-PAAD | 21.94 | 5.73 |
| GTEX-1A3MW-1926-SM-7IGM8 | Pancreas | TCGA-PAAD | 20.52 | 7.13 |
| GTEX-1A8FM-1826-SM-7MGXO | Pancreas | TCGA-PAAD | 8.02 | 4.98 |
| GTEX-1AMFI-0726-SM-731D9 | Pancreas | TCGA-PAAD | 11.05 | 4.8 |
| GTEX-1AX8Z-1726-SM-731DE | Pancreas | TCGA-PAAD | 22.14 | 6.86 |
| GTEX-1AX9J-1726-SM-731FF | Pancreas | TCGA-PAAD | 13.36 | 3.76 |
| GTEX-1AX9K-2026-SM-731D1 | Pancreas | TCGA-PAAD | 15.51 | 3.57 |
| GTEX-1AYCT-1226-SM-7EPGH | Pancreas | TCGA-PAAD | 12.71 | 3.1 |
| GTEX-1AYD5-0326-SM-7EWEO | Pancreas | TCGA-PAAD | 17.84 | 6.14 |
| GTEX-1B8KE-0726-SM-72D7G | Pancreas | TCGA-PAAD | 13.84 | 6.29 |

| | | | | |
|---|---|---|---|---|
| GTEX-1B8KZ-0926-SM-731D7 | Pancreas | TCGA-PAAD | 12.33 | 7.42 |
| GTEX-1B932-1626-SM-731EA | Pancreas | TCGA-PAAD | 18.01 | 8.62 |
| GTEX-1B933-1726-SM-731FC | Pancreas | TCGA-PAAD | 11.51 | 6.72 |
| GTEX-1B97J-1526-SM-79ONI | Pancreas | TCGA-PAAD | 11.34 | 3.47 |
| GTEX-1B996-1326-SM-731EO | Pancreas | TCGA-PAAD | 8.77 | 3.13 |
| GTEX-1C2JI-0926-SM-731F1 | Pancreas | TCGA-PAAD | 14.09 | 4.38 |
| GTEX-1C4CL-0926-SM-79OL2 | Pancreas | TCGA-PAAD | 14.94 | 5.95 |
| GTEX-1C64O-1426-SM-7EWER | Pancreas | TCGA-PAAD | 21.87 | 8.62 |
| GTEX-1C6VQ-1726-SM-7IGLQ | Pancreas | TCGA-PAAD | 15.83 | 6.32 |
| GTEX-1CAMQ-1626-SM-7MGXG | Pancreas | TCGA-PAAD | 7.32 | 2.48 |
| GTEX-1CAMR-1126-SM-7MGXF | Pancreas | TCGA-PAAD | 13.16 | 2.87 |
| GTEX-1CAMS-1926-SM-7DUEU | Pancreas | TCGA-PAAD | 15.09 | 4.49 |
| GTEX-1CB4F-1226-SM-7DHKU | Pancreas | TCGA-PAAD | 11.36 | 3.6 |
| GTEX-1CB4G-1826-SM-7MKGO | Pancreas | TCGA-PAAD | 18.06 | 6.49 |
| GTEX-1CB4I-1326-SM-79OLY | Pancreas | TCGA-PAAD | 12.94 | 4.08 |
| GTEX-1F5PL-0726-SM-7MKH8 | Pancreas | TCGA-PAAD | 14.17 | 4.19 |
| GTEX-1GF9X-1126-SM-7MKHC | Pancreas | TCGA-PAAD | 11.38 | 2.51 |
| GTEX-1GL5R-0926-SM-9JGFO | Pancreas | TCGA-PAAD | 9.79 | 3.45 |
| GTEX-1GMR3-1226-SM-7MGY1 | Pancreas | TCGA-PAAD | 4.94 | 1.71 |
| GTEX-1GN1U-1126-SM-9WPPZ | Pancreas | TCGA-PAAD | 12.45 | 4 |
| GTEX-1GN1W-1126-SM-9OSW3 | Pancreas | TCGA-PAAD | 12.5 | 3.57 |
| GTEX-1GZ4H-0626-SM-9OSXJ | Pancreas | TCGA-PAAD | 12.97 | 5.51 |
| GTEX-1H11D-0926-SM-9OSXO | Pancreas | TCGA-PAAD | 20.54 | 7.99 |
| GTEX-1H1DG-1426-SM-9OSXP | Pancreas | TCGA-PAAD | 17.45 | 9.16 |
| GTEX-1H1E6-0826-SM-9WG83 | Pancreas | TCGA-PAAD | 12.63 | 4.53 |
| GTEX-1H23P-2026-SM-9OSXY | Pancreas | TCGA-PAAD | 15.36 | 7.07 |
| GTEX-1HKZK-1126-SM-9WG82 | Pancreas | TCGA-PAAD | 22.51 | 11.48 |
| GTEX-1HSGN-1426-SM-A8NAA | Pancreas | TCGA-PAAD | 15.13 | 4.71 |
| GTEX-1HSMO-2026-SM-CNNQO | Pancreas | TCGA-PAAD | 28.4 | 7.83 |
| GTEX-1I1GQ-2226-SM-B2LX1 | Pancreas | TCGA-PAAD | 6.28 | 3.48 |
| GTEX-1I1GT-0926-SM-AHZ2G | Pancreas | TCGA-PAAD | 14.23 | 5.57 |

| | | | | |
|---|---|---|---|---|
| GTEX-1I1GU-0326-SM-AHZ2K | Pancreas | TCGA-PAAD | 18.58 | 6.17 |
| GTEX-1I4MK-1026-SM-AHZ2H | Pancreas | TCGA-PAAD | 13.34 | 4.11 |
| GTEX-1IDJC-0526-SM-AHZ2Q | Pancreas | TCGA-PAAD | 14.83 | 4.98 |
| GTEX-1IDJD-0126-SM-C1YQZ | Pancreas | TCGA-PAAD | 15.32 | 6.31 |
| GTEX-1IDJU-0326-SM-CGQEE | Pancreas | TCGA-PAAD | 9.63 | 3.05 |
| GTEX-1IL2U-2326-SM-CNNQV | Pancreas | TCGA-PAAD | 8.98 | 3.17 |
| GTEX-1J8EW-1926-SM-CGQGR | Pancreas | TCGA-PAAD | 5.45 | 3.84 |
| GTEX-1J8Q3-0626-SM-AHZ3V | Pancreas | TCGA-PAAD | 12.4 | 3.24 |
| GTEX-1JK1U-1026-SM-AHZ44 | Pancreas | TCGA-PAAD | 25.15 | 6.56 |
| GTEX-1JKYN-0426-SM-CGQG2 | Pancreas | TCGA-PAAD | 20.4 | 6.45 |
| GTEX-1JKYR-1026-SM-CGQGI | Pancreas | TCGA-PAAD | 16.79 | 5.05 |
| GTEX-1JMPZ-1026-SM-CXZJS | Pancreas | TCGA-PAAD | 28.82 | 10.45 |
| GTEX-1K2DA-0326-SM-CGQGX | Pancreas | TCGA-PAAD | 14.02 | 6.21 |
| GTEX-1K2DU-1026-SM-DHXKC | Pancreas | TCGA-PAAD | 25.03 | 9.2 |
| GTEX-1K9T9-1726-SM-CXZK1 | Pancreas | TCGA-PAAD | 17.78 | 7.18 |
| GTEX-1KANA-1526-SM-CXZKG | Pancreas | TCGA-PAAD | 12.93 | 3.68 |
| GTEX-1KXAM-0226-SM-EV7AP | Pancreas | TCGA-PAAD | 14.17 | 4.45 |
| GTEX-1L5NE-0826-SM-DHXKU | Pancreas | TCGA-PAAD | 9.86 | 3.14 |
| GTEX-1LG7Z-0326-SM-DHXK3 | Pancreas | TCGA-PAAD | 11.31 | 2.69 |
| GTEX-1LGRB-0426-SM-CNNRD | Pancreas | TCGA-PAAD | 11.88 | 3.38 |
| GTEX-1LSNM-0726-SM-DHXL1 | Pancreas | TCGA-PAAD | 10.84 | 2.54 |
| GTEX-1LVAN-1226-SM-CNNQY | Pancreas | TCGA-PAAD | 13.05 | 3.94 |
| GTEX-1M5QR-0926-SM-DHXK8 | Pancreas | TCGA-PAAD | 17.45 | 6.31 |
| GTEX-1MGNQ-1226-SM-EV7BM | Pancreas | TCGA-PAAD | 17.03 | 5.66 |
| GTEX-1MUQO-1426-SM-DPRYJ | Pancreas | TCGA-PAAD | 17.32 | 3.95 |
| GTEX-1N2DV-1726-SM-DPRYR | Pancreas | TCGA-PAAD | 10.32 | 3.65 |
| GTEX-1N2EF-2126-SM-DPRYL | Pancreas | TCGA-PAAD | 19.74 | 6.59 |
| GTEX-1NSGN-1726-SM-EXOJG | Pancreas | TCGA-PAAD | 22.38 | 9.38 |
| GTEX-1NUQO-0826-SM-DPS16 | Pancreas | TCGA-PAAD | 10.3 | 3.97 |
| GTEX-1O97I-1726-SM-EXOIP | Pancreas | TCGA-PAAD | 10.97 | 4.16 |
| GTEX-1OFPY-0626-SM-EXOJ4 | Pancreas | TCGA-PAAD | 14.07 | 4.15 |

| | | | | |
|---|---|---|---|---|
| GTEX-1OKEX-1726-SM-E9U4F | Pancreas | TCGA-PAAD | 10.11 | 4.6 |
| GTEX-1PBJI-2126-SM-DPRYG | Pancreas | TCGA-PAAD | 3.96 | 2.38 |
| GTEX-1PFEY-1126-SM-DPRZ3 | Pancreas | TCGA-PAAD | 14.87 | 6.77 |
| GTEX-1PPH8-2326-SM-EAZ48 | Pancreas | TCGA-PAAD | 20.81 | 6.85 |
| GTEX-1PWST-1526-SM-EXOIT | Pancreas | TCGA-PAAD | 24.95 | 8.85 |
| GTEX-1QAET-1326-SM-E9U64 | Pancreas | TCGA-PAAD | 17.11 | 5.62 |
| GTEX-1QEPI-1626-SM-EAZ3I | Pancreas | TCGA-PAAD | 8.63 | 3.38 |
| GTEX-1QP28-1026-SM-EAZ4K | Pancreas | TCGA-PAAD | 13.19 | 4.17 |
| GTEX-1QP29-1926-SM-DPRZO | Pancreas | TCGA-PAAD | 17.28 | 5.12 |
| GTEX-1QP2A-1326-SM-EAZ3K | Pancreas | TCGA-PAAD | 16.17 | 4.25 |
| GTEX-1QPFJ-2226-SM-EXOJW | Pancreas | TCGA-PAAD | 14.72 | 6.14 |
| GTEX-1QW4Y-0926-SM-DPRY2 | Pancreas | TCGA-PAAD | 8.4 | 2.34 |
| GTEX-1R46S-1426-SM-EXOJD | Pancreas | TCGA-PAAD | 24.86 | 10.26 |
| GTEX-1R7EU-0726-SM-DPRZM | Pancreas | TCGA-PAAD | 10.49 | 2.92 |
| GTEX-1R7EV-1126-SM-DPRXU | Pancreas | TCGA-PAAD | 14.22 | 3.2 |
| GTEX-1R9K5-1226-SM-E6CPQ | Pancreas | TCGA-PAAD | 18.21 | 6.24 |
| GTEX-1RAZA-1626-SM-EV79I | Pancreas | TCGA-PAAD | 12.89 | 4.21 |
| GTEX-1RAZQ-1426-SM-EV7B2 | Pancreas | TCGA-PAAD | 17.4 | 5.88 |
| GTEX-1S82P-1026-SM-EAZ4W | Pancreas | TCGA-PAAD | 11.68 | 3.91 |
| GTEX-1S831-0626-SM-EV79N | Pancreas | TCGA-PAAD | 15.64 | 4.83 |
| GTEX-1S83E-1426-SM-EV7B8 | Pancreas | TCGA-PAAD | 12.9 | 4.22 |
| GTEX-N7MT-1626-SM-3LK71 | Pancreas | TCGA-PAAD | 16.69 | 6.71 |
| GTEX-NFK9-1626-SM-3LK5J | Pancreas | TCGA-PAAD | 13.24 | 5.02 |
| GTEX-NPJ8-2126-SM-3MJGK | Pancreas | TCGA-PAAD | 26.42 | 8.08 |
| GTEX-O5YT-1026-SM-3MJGF | Pancreas | TCGA-PAAD | 16.74 | 9.21 |
| GTEX-OHPL-1026-SM-3MJGI | Pancreas | TCGA-PAAD | 13.54 | 4.25 |
| GTEX-OHPM-1026-SM-3LK74 | Pancreas | TCGA-PAAD | 16.79 | 4.03 |
| GTEX-OIZF-1026-SM-2HML5 | Pancreas | TCGA-PAAD | 21.87 | 7.2 |
| GTEX-OIZI-1526-SM-E6CHI | Pancreas | TCGA-PAAD | 9.85 | 4.33 |
| GTEX-OOBJ-1026-SM-3NB2L | Pancreas | TCGA-PAAD | 8.62 | 3.61 |
| GTEX-OOBK-1026-SM-48TC2 | Pancreas | TCGA-PAAD | 11.99 | 6.23 |

| | | | | |
|---|---|---|---|---|
| GTEX-OXRK-1526-SM-E6CHJ | Pancreas | TCGA-PAAD | 11 | 3.92 |
| GTEX-P4PP-1026-SM-3NM9O | Pancreas | TCGA-PAAD | 9.2 | 4.48 |
| GTEX-P4PQ-1026-SM-3NMCN | Pancreas | TCGA-PAAD | 15.41 | 4.53 |
| GTEX-P4QS-1026-SM-3NMCW | Pancreas | TCGA-PAAD | 16.64 | 6.95 |
| GTEX-PLZ6-0726-SM-3P619 | Pancreas | TCGA-PAAD | 10.92 | 4.21 |
| GTEX-PSDG-1526-SM-48TCY | Pancreas | TCGA-PAAD | 14.89 | 5.81 |
| GTEX-PW2O-0826-SM-48TC5 | Pancreas | TCGA-PAAD | 13.16 | 4.17 |
| GTEX-PX3G-1026-SM-48TZW | Pancreas | TCGA-PAAD | 10.51 | 3.09 |
| GTEX-Q2AH-0926-SM-48TZK | Pancreas | TCGA-PAAD | 23.09 | 11.2 |
| GTEX-Q2AI-0426-SM-48U13 | Pancreas | TCGA-PAAD | 16.73 | 5.63 |
| GTEX-Q734-0426-SM-48TZX | Pancreas | TCGA-PAAD | 9.68 | 3.24 |
| GTEX-QCQG-0426-SM-48U29 | Pancreas | TCGA-PAAD | 17.56 | 8.32 |
| GTEX-QDVJ-1226-SM-48U1V | Pancreas | TCGA-PAAD | 17.15 | 8.97 |
| GTEX-QDVN-0926-SM-2I5GL | Pancreas | TCGA-PAAD | 16.33 | 5.02 |
| GTEX-QEL4-1326-SM-447AD | Pancreas | TCGA-PAAD | 11.89 | 4.01 |
| GTEX-QESD-0226-SM-447BH | Pancreas | TCGA-PAAD | 8.29 | 1.68 |
| GTEX-QLQ7-0626-SM-CE6QX | Pancreas | TCGA-PAAD | 11.51 | 3.24 |
| GTEX-QLQW-0326-SM-447A8 | Pancreas | TCGA-PAAD | 10.41 | 3.65 |
| GTEX-QMRM-0326-SM-CKZNB | Pancreas | TCGA-PAAD | 12.67 | 4.77 |
| GTEX-QV31-0226-SM-447BO | Pancreas | TCGA-PAAD | 10.25 | 4.04 |
| GTEX-QV44-0426-SM-4R1KF | Pancreas | TCGA-PAAD | 9.73 | 4.88 |
| GTEX-R53T-0426-SM-48FEM | Pancreas | TCGA-PAAD | 11.92 | 4.32 |
| GTEX-R55D-1426-SM-48FEN | Pancreas | TCGA-PAAD | 11.54 | 4.47 |
| GTEX-R55G-0326-SM-48FDM | Pancreas | TCGA-PAAD | 16.71 | 4.62 |
| GTEX-RM2N-0326-SM-48FD8 | Pancreas | TCGA-PAAD | 9 | 4.66 |
| GTEX-RWS6-1126-SM-47JXC | Pancreas | TCGA-PAAD | 9.67 | 4.08 |
| GTEX-S32W-0826-SM-4AD5Z | Pancreas | TCGA-PAAD | 19.22 | 6.84 |
| GTEX-S33H-1226-SM-4AD69 | Pancreas | TCGA-PAAD | 16.63 | 6.11 |
| GTEX-S3XE-0526-SM-4AD4G | Pancreas | TCGA-PAAD | 10.94 | 5.25 |
| GTEX-S4P3-0626-SM-4AD59 | Pancreas | TCGA-PAAD | 16.85 | 6.17 |
| GTEX-S95S-0726-SM-4B64H | Pancreas | TCGA-PAAD | 11.6 | 4.73 |

| | | | | |
|---|---|---|---|---|
| GTEX-SE5C-0326-SM-4BRWX | Pancreas | TCGA-PAAD | 13.07 | 5.05 |
| GTEX-SNOS-0926-SM-4DM7A | Pancreas | TCGA-PAAD | 12.16 | 2.89 |
| GTEX-SUCS-1426-SM-4DM5W | Pancreas | TCGA-PAAD | 16.21 | 4.1 |
| GTEX-T5JW-0226-SM-4DM7I | Pancreas | TCGA-PAAD | 16.29 | 5.34 |
| GTEX-T8EM-0826-SM-4DM76 | Pancreas | TCGA-PAAD | 13.81 | 4.93 |
| GTEX-TKQ2-0426-SM-4DXUO | Pancreas | TCGA-PAAD | 11.29 | 3.36 |
| GTEX-TML8-0526-SM-32QOQ | Pancreas | TCGA-PAAD | 19.21 | 7.89 |
| GTEX-TMMY-1326-SM-4DXU9 | Pancreas | TCGA-PAAD | 11.64 | 3.66 |
| GTEX-U4B1-0726-SM-4DXUA | Pancreas | TCGA-PAAD | 16.34 | 4.41 |
| GTEX-U8XE-2026-SM-3DB8S | Pancreas | TCGA-PAAD | 17.2 | 7.13 |
| GTEX-UJHI-0626-SM-3DB8T | Pancreas | TCGA-PAAD | 12.99 | 6.06 |
| GTEX-UJMC-1126-SM-3GADP | Pancreas | TCGA-PAAD | 11.35 | 5.08 |
| GTEX-V1D1-0726-SM-4JBH7 | Pancreas | TCGA-PAAD | 16.34 | 5.72 |
| GTEX-V955-0326-SM-4JBGV | Pancreas | TCGA-PAAD | 14.72 | 4.87 |
| GTEX-VJYA-0826-SM-4KL1M | Pancreas | TCGA-PAAD | 16.97 | 6.24 |
| GTEX-VUSG-0326-SM-3GIJ7 | Pancreas | TCGA-PAAD | 7.89 | 5.32 |
| GTEX-W5WG-0826-SM-4RGNE | Pancreas | TCGA-PAAD | 12.43 | 5.06 |
| GTEX-WFG7-0426-SM-4LMK5 | Pancreas | TCGA-PAAD | 17.91 | 6.23 |
| GTEX-WFG8-0326-SM-4LVN4 | Pancreas | TCGA-PAAD | 15.05 | 5.14 |
| GTEX-WFJO-0626-SM-4LVMC | Pancreas | TCGA-PAAD | 12.64 | 6.2 |
| GTEX-WFON-0626-SM-4LVLX | Pancreas | TCGA-PAAD | 21.23 | 7.46 |
| GTEX-WH7G-0826-SM-4LVMR | Pancreas | TCGA-PAAD | 8.12 | 3.3 |
| GTEX-WHPG-0326-SM-4M1XV | Pancreas | TCGA-PAAD | 14.46 | 6.68 |
| GTEX-WHSB-0726-SM-4M1XQ | Pancreas | TCGA-PAAD | 14.93 | 5.43 |
| GTEX-WHWD-0726-SM-4OORX | Pancreas | TCGA-PAAD | 16.02 | 5.22 |
| GTEX-WI4N-1826-SM-4OOSF | Pancreas | TCGA-PAAD | 11.64 | 6.65 |
| GTEX-WQUQ-2126-SM-4OOSO | Pancreas | TCGA-PAAD | 11.45 | 4.45 |
| GTEX-WRHK-0226-SM-4MVOH | Pancreas | TCGA-PAAD | 8.64 | 3.4 |
| GTEX-WXYG-0826-SM-4ONC7 | Pancreas | TCGA-PAAD | 17.3 | 4.63 |
| GTEX-WY7C-1026-SM-4OND3 | Pancreas | TCGA-PAAD | 22.94 | 5.71 |
| GTEX-WYJK-2426-SM-4ONDQ | Pancreas | TCGA-PAAD | 17.7 | 5.91 |

| | | | | |
|---|---|---|---|---|
| GTEX-WYVS-0926-SM-4SOJV | Pancreas | TCGA-PAAD | 9.74 | 2.92 |
| GTEX-X15G-0726-SM-4PQZ5 | Pancreas | TCGA-PAAD | 13.4 | 5.64 |
| GTEX-X3Y1-0726-SM-3P5YU | Pancreas | TCGA-PAAD | 8.46 | 3.05 |
| GTEX-X4LF-0326-SM-4QAS9 | Pancreas | TCGA-PAAD | 14.39 | 4.69 |
| GTEX-X5EB-0526-SM-46MVP | Pancreas | TCGA-PAAD | 11.41 | 6.5 |
| GTEX-XBED-0226-SM-47JY8 | Pancreas | TCGA-PAAD | 18.99 | 6.18 |
| GTEX-XBEW-1626-SM-4SOJP | Pancreas | TCGA-PAAD | 8.79 | 2.82 |
| GTEX-XGQ4-0226-SM-4GIDS | Pancreas | TCGA-PAAD | 14.28 | 4.61 |
| GTEX-XMK1-0326-SM-4B652 | Pancreas | TCGA-PAAD | 10.83 | 2.83 |
| GTEX-XPVG-0326-SM-4B653 | Pancreas | TCGA-PAAD | 15.21 | 6.09 |
| GTEX-XQ8I-1926-SM-4BOOK | Pancreas | TCGA-PAAD | 12.08 | 4.57 |
| GTEX-XUW1-1726-SM-4BOOZ | Pancreas | TCGA-PAAD | 11.25 | 3.5 |
| GTEX-XV7Q-0926-SM-4BRVQ | Pancreas | TCGA-PAAD | 20.97 | 9.93 |
| GTEX-XXEK-1726-SM-4BRVB | Pancreas | TCGA-PAAD | 13.29 | 6.48 |
| GTEX-XYKS-1226-SM-4BRVI | Pancreas | TCGA-PAAD | 24.43 | 10.12 |
| GTEX-Y114-0826-SM-4TT77 | Pancreas | TCGA-PAAD | 14.38 | 4.2 |
| GTEX-Y3I4-0826-SM-4TT2A | Pancreas | TCGA-PAAD | 15.16 | 4.88 |
| GTEX-Y3IK-0426-SM-4WWE2 | Pancreas | TCGA-PAAD | 16.65 | 4.3 |
| GTEX-Y5LM-0526-SM-4V6G3 | Pancreas | TCGA-PAAD | 16.65 | 5.05 |
| GTEX-Y8E4-1326-SM-5IFIY | Pancreas | TCGA-PAAD | 9.96 | 3.49 |
| GTEX-Y8LW-1026-SM-5IFJY | Pancreas | TCGA-PAAD | 19.75 | 7.2 |
| GTEX-Y9LG-0726-SM-4VDS3 | Pancreas | TCGA-PAAD | 22.48 | 5.46 |
| GTEX-YB5E-0526-SM-4VDSD | Pancreas | TCGA-PAAD | 16.25 | 4.73 |
| GTEX-YB5K-1826-SM-5IFJC | Pancreas | TCGA-PAAD | 10.5 | 2.56 |
| GTEX-YEC3-0626-SM-CMKFG | Pancreas | TCGA-PAAD | 12.56 | 4.26 |
| GTEX-YEC4-1326-SM-5IFHG | Pancreas | TCGA-PAAD | 9.95 | 3.6 |
| GTEX-YF7O-1326-SM-4W1ZR | Pancreas | TCGA-PAAD | 11.87 | 4.1 |
| GTEX-YFCO-1026-SM-5LU9S | Pancreas | TCGA-PAAD | 17.45 | 4.17 |
| GTEX-ZAB4-1726-SM-5HL8C | Pancreas | TCGA-PAAD | 17.72 | 6.28 |
| GTEX-ZAB5-0826-SM-5P9FU | Pancreas | TCGA-PAAD | 15.59 | 5.57 |
| GTEX-ZAK1-2326-SM-5CVMY | Pancreas | TCGA-PAAD | 7.43 | 4.04 |

| | | | | |
|---|---|---|---|---|
| GTEX-ZC5H-0826-SM-5N9FH | Pancreas | TCGA-PAAD | 23.15 | 7.56 |
| GTEX-ZDTT-1126-SM-4WKFW | Pancreas | TCGA-PAAD | 14.57 | 4.67 |
| GTEX-ZDYS-2526-SM-4WKGU | Pancreas | TCGA-PAAD | 10.72 | 4.47 |
| GTEX-ZEX8-1026-SM-4WKHE | Pancreas | TCGA-PAAD | 8.1 | 3.71 |
| GTEX-ZF29-1126-SM-4WKGO | Pancreas | TCGA-PAAD | 11.92 | 3.24 |
| GTEX-ZF3C-2026-SM-4WWB5 | Pancreas | TCGA-PAAD | 15.49 | 5.74 |
| GTEX-ZG7Y-0326-SM-4WWEY | Pancreas | TCGA-PAAD | 14.46 | 5.11 |
| GTEX-ZLFU-0726-SM-57WF6 | Pancreas | TCGA-PAAD | 12.12 | 4.31 |
| GTEX-ZLWG-0326-SM-4WWC7 | Pancreas | TCGA-PAAD | 13.49 | 3.29 |
| GTEX-ZP4G-0426-SM-4YCER | Pancreas | TCGA-PAAD | 16.12 | 5.44 |
| GTEX-ZPCL-0726-SM-DO91S | Pancreas | TCGA-PAAD | 10.55 | 2.42 |
| GTEX-ZPIC-0926-SM-4WWFK | Pancreas | TCGA-PAAD | 15.09 | 4.5 |
| GTEX-ZPU1-0226-SM-4WWC9 | Pancreas | TCGA-PAAD | 9.75 | 2.26 |
| GTEX-ZT9W-0926-SM-57WFS | Pancreas | TCGA-PAAD | 13.47 | 3.16 |
| GTEX-ZTPG-1026-SM-5DUWP | Pancreas | TCGA-PAAD | 25.77 | 6.84 |
| GTEX-ZV7C-0726-SM-59HKH | Pancreas | TCGA-PAAD | 14.57 | 5.42 |
| GTEX-ZVP2-0726-SM-59HKY | Pancreas | TCGA-PAAD | 13.77 | 3.3 |
| GTEX-ZVT2-2026-SM-5NQ8Q | Pancreas | TCGA-PAAD | 10.61 | 4.63 |
| GTEX-ZVZP-0626-SM-59HL5 | Pancreas | TCGA-PAAD | 15.02 | 4.9 |
| GTEX-ZYFG-0826-SM-5BC5T | Pancreas | TCGA-PAAD | 15.85 | 4.63 |
| GTEX-ZYT6-1326-SM-5E453 | Pancreas | TCGA-PAAD | 22.07 | 4.75 |
| GTEX-ZYW4-2126-SM-59HJ9 | Pancreas | TCGA-PAAD | 11.51 | 4.57 |
| GTEX-ZYY3-0826-SM-5E44R | Pancreas | TCGA-PAAD | 16.63 | 6.43 |
| GTEX-ZZPU-0726-SM-5N9C8 | Pancreas | TCGA-PAAD | 16.43 | 4.44 |
| GTEX-111CU-1526-SM-5N9FS | Prostate | TCGA-PRAD | 32.88 | 14.48 |
| GTEX-111FC-2026-SM-5GZYO | Prostate | TCGA-PRAD | 26.87 | 12.47 |
| GTEX-111YS-1726-SM-5GIED | Prostate | TCGA-PRAD | 30.67 | 9.16 |
| GTEX-117YW-1426-SM-5EGGO | Prostate | TCGA-PRAD | 24.49 | 9.29 |
| GTEX-117YX-1526-SM-5H12T | Prostate | TCGA-PRAD | 36.94 | 13.56 |
| GTEX-11DXZ-1826-SM-5H12Y | Prostate | TCGA-PRAD | 34.42 | 12.8 |
| GTEX-11DYG-2526-SM-5N9BB | Prostate | TCGA-PRAD | 30.01 | 12.8 |

| | | | | |
|---|---|---|---|---|
| GTEX-11EQ9-1726-SM-5HL6U | Prostate | TCGA-PRAD | 43.73 | 8.47 |
| GTEX-11NSD-0826-SM-5986S | Prostate | TCGA-PRAD | 34.19 | 12.27 |
| GTEX-11O72-2726-SM-5HL6K | Prostate | TCGA-PRAD | 37.95 | 11.19 |
| GTEX-11OF3-1726-SM-5GU5Q | Prostate | TCGA-PRAD | 22.71 | 7.01 |
| GTEX-11P7K-0626-SM-5985Z | Prostate | TCGA-PRAD | 32.5 | 11.04 |
| GTEX-11P82-1126-SM-5BC5K | Prostate | TCGA-PRAD | 37.64 | 14.69 |
| GTEX-11TT1-2026-SM-5EQM8 | Prostate | TCGA-PRAD | 50.03 | 17.02 |
| GTEX-11TUW-2126-SM-5GU7B | Prostate | TCGA-PRAD | 29.44 | 10.56 |
| GTEX-11WQK-2726-SM-5EQMU | Prostate | TCGA-PRAD | 37.31 | 9.25 |
| GTEX-11ZUS-2126-SM-5N9E6 | Prostate | TCGA-PRAD | 31.81 | 12.87 |
| GTEX-12696-2226-SM-5EQ65 | Prostate | TCGA-PRAD | 45.65 | 12.2 |
| GTEX-12BJ1-1226-SM-5LUAE | Prostate | TCGA-PRAD | 40.83 | 13 |
| GTEX-12C56-1326-SM-5FQSV | Prostate | TCGA-PRAD | 28.45 | 12.56 |
| GTEX-12WSN-5007-SM-7MGWH | Prostate | TCGA-PRAD | 60.26 | 18.78 |
| GTEX-12ZZZ-2026-SM-5LZWJ | Prostate | TCGA-PRAD | 41.68 | 15.43 |
| GTEX-13111-1326-SM-5GCNO | Prostate | TCGA-PRAD | 34.71 | 13.16 |
| GTEX-131XF-5013-SM-7EWFI | Prostate | TCGA-PRAD | 48.94 | 15.75 |
| GTEX-132NY-2826-SM-5P9G2 | Prostate | TCGA-PRAD | 18.81 | 7.4 |
| GTEX-132QS-1126-SM-5P9GC | Prostate | TCGA-PRAD | 40.41 | 10.34 |
| GTEX-1399R-0526-SM-5IJEF | Prostate | TCGA-PRAD | 41.03 | 12.18 |
| GTEX-1399T-1226-SM-5P9J5 | Prostate | TCGA-PRAD | 38.56 | 14.41 |
| GTEX-139T6-1526-SM-5P9G6 | Prostate | TCGA-PRAD | 32.92 | 12.16 |
| GTEX-13FHO-2826-SM-5K7ZA | Prostate | TCGA-PRAD | 34.23 | 13.05 |
| GTEX-13FTW-1226-SM-5LZZ1 | Prostate | TCGA-PRAD | 43.05 | 15.36 |
| GTEX-13FXS-2826-SM-5LZYC | Prostate | TCGA-PRAD | 26.99 | 11.35 |
| GTEX-13G51-2326-SM-5LZXV | Prostate | TCGA-PRAD | 19.66 | 8.49 |
| GTEX-13N2G-1826-SM-5KM1I | Prostate | TCGA-PRAD | 36.17 | 13.92 |
| GTEX-13NYS-1026-SM-5KLZT | Prostate | TCGA-PRAD | 48.11 | 13.46 |
| GTEX-13O21-1126-SM-5N9EV | Prostate | TCGA-PRAD | 29.77 | 14.45 |
| GTEX-13O61-1726-SM-5KM46 | Prostate | TCGA-PRAD | 46.14 | 11.23 |
| GTEX-13OW8-0426-SM-5J2NR | Prostate | TCGA-PRAD | 36.56 | 9.25 |

| | | | | |
|---|---|---|---|---|
| GTEX-13RTJ-2126-SM-5S2PZ | Prostate | TCGA-PRAD | 36.94 | 15.17 |
| GTEX-144GM-0826-SM-5O98R | Prostate | TCGA-PRAD | 30.53 | 8.28 |
| GTEX-145LT-0626-SM-5O99E | Prostate | TCGA-PRAD | 21.93 | 9.77 |
| GTEX-145LU-2126-SM-5Q5E9 | Prostate | TCGA-PRAD | 35.96 | 13.91 |
| GTEX-145LV-1226-SM-5S2QH | Prostate | TCGA-PRAD | 34.51 | 15.12 |
| GTEX-146FQ-1726-SM-5QGPX | Prostate | TCGA-PRAD | 37.26 | 13.33 |
| GTEX-1477Z-2226-SM-5QGPG | Prostate | TCGA-PRAD | 39.26 | 15.56 |
| GTEX-147JS-2826-SM-5YYA7 | Prostate | TCGA-PRAD | 24.69 | 8.21 |
| GTEX-148VJ-1826-SM-5Q5DV | Prostate | TCGA-PRAD | 26.14 | 7.69 |
| GTEX-14ABY-0726-SM-5Q5DF | Prostate | TCGA-PRAD | 31.09 | 13.25 |
| GTEX-14BIL-2126-SM-73KW3 | Prostate | TCGA-PRAD | 15.3 | 7.97 |
| GTEX-14BMV-2326-SM-5RQJ4 | Prostate | TCGA-PRAD | 24.45 | 8.65 |
| GTEX-14C5O-2826-SM-5RQI6 | Prostate | TCGA-PRAD | 32.71 | 8.38 |
| GTEX-14DAR-1026-SM-73KV3 | Prostate | TCGA-PRAD | 43.57 | 12.18 |
| GTEX-14E6C-2826-SM-664NC | Prostate | TCGA-PRAD | 59.51 | 9.26 |
| GTEX-14E6E-1126-SM-5S2R4 | Prostate | TCGA-PRAD | 57.8 | 23.53 |
| GTEX-14ICK-1826-SM-69LOP | Prostate | TCGA-PRAD | 27.56 | 10.66 |
| GTEX-14PJ2-2126-SM-5YY96 | Prostate | TCGA-PRAD | 41.78 | 14.5 |
| GTEX-14PJ3-1726-SM-5ZZVH | Prostate | TCGA-PRAD | 37.79 | 13.96 |
| GTEX-14PJ4-1226-SM-5YY9U | Prostate | TCGA-PRAD | 44.5 | 11.68 |
| GTEX-14PJO-2826-SM-6AJA6 | Prostate | TCGA-PRAD | 30.12 | 14.03 |
| GTEX-15CHR-1226-SM-79OON | Prostate | TCGA-PRAD | 37.74 | 12.34 |
| GTEX-15RIF-1526-SM-6M46K | Prostate | TCGA-PRAD | 34.45 | 10.72 |
| GTEX-16GPK-2826-SM-6LPJ6 | Prostate | TCGA-PRAD | 21.77 | 9.36 |
| GTEX-16MTA-1426-SM-6PALY | Prostate | TCGA-PRAD | 49.36 | 14.84 |
| GTEX-16XZZ-2426-SM-7KULN | Prostate | TCGA-PRAD | 35.11 | 11.72 |
| GTEX-16YQH-2326-SM-7EWDS | Prostate | TCGA-PRAD | 19.7 | 8.89 |
| GTEX-178AV-0426-SM-6LPJG | Prostate | TCGA-PRAD | 35.12 | 11.29 |
| GTEX-17EUY-2026-SM-79OMV | Prostate | TCGA-PRAD | 37.14 | 11.31 |
| GTEX-17HGU-2226-SM-7IGLI | Prostate | TCGA-PRAD | 41.61 | 13.45 |
| GTEX-17HHE-1126-SM-793BS | Prostate | TCGA-PRAD | 32.26 | 13.38 |

| | | | | |
|---|---|---|---|---|
| GTEX-17HHY-2526-SM-7IGME | Prostate | TCGA-PRAD | 21.18 | 9.67 |
| GTEX-17HII-2526-SM-7LT8C | Prostate | TCGA-PRAD | 21.99 | 11.92 |
| GTEX-17MFQ-1326-SM-718BX | Prostate | TCGA-PRAD | 24.43 | 9.31 |
| GTEX-18A66-2326-SM-7LT8Q | Prostate | TCGA-PRAD | 33.05 | 11.32 |
| GTEX-18QFQ-2026-SM-72D7J | Prostate | TCGA-PRAD | 23.15 | 10.49 |
| GTEX-1A3MV-0826-SM-72D6J | Prostate | TCGA-PRAD | 23.42 | 10.28 |
| GTEX-1AX8Z-5019-SM-AHZ2C | Prostate | TCGA-PRAD | 58.79 | 18.27 |
| GTEX-1AX9I-2326-SM-7PBXV | Prostate | TCGA-PRAD | 41.42 | 16.92 |
| GTEX-1AX9J-1426-SM-731BV | Prostate | TCGA-PRAD | 37.65 | 10.89 |
| GTEX-1AX9K-1526-SM-73KVP | Prostate | TCGA-PRAD | 32.78 | 12.53 |
| GTEX-1B8KE-1026-SM-731EQ | Prostate | TCGA-PRAD | 44.91 | 10.93 |
| GTEX-1B8KZ-1626-SM-73KUU | Prostate | TCGA-PRAD | 30.45 | 16.17 |
| GTEX-1B97I-0826-SM-731DN | Prostate | TCGA-PRAD | 46.01 | 18.69 |
| GTEX-1BAJH-0626-SM-7P8OY | Prostate | TCGA-PRAD | 26.72 | 8.72 |
| GTEX-1C64O-2026-SM-7IGPA | Prostate | TCGA-PRAD | 40.45 | 12.16 |
| GTEX-1C6VQ-2426-SM-7EWEU | Prostate | TCGA-PRAD | 22.25 | 9.41 |
| GTEX-1C6WA-2326-SM-7P8OW | Prostate | TCGA-PRAD | 30.37 | 11.92 |
| GTEX-1CAMQ-2226-SM-7MGWS | Prostate | TCGA-PRAD | 27.29 | 12.15 |
| GTEX-1CAMR-1626-SM-79OLC | Prostate | TCGA-PRAD | 35.57 | 11.76 |
| GTEX-1CB4E-1726-SM-7IGMS | Prostate | TCGA-PRAD | 52.11 | 28.95 |
| GTEX-1CB4F-2326-SM-7MKFO | Prostate | TCGA-PRAD | 35.66 | 15.14 |
| GTEX-1CB4G-2526-SM-7MXTG | Prostate | TCGA-PRAD | 25.4 | 13.57 |
| GTEX-1CB4J-1526-SM-7MXVB | Prostate | TCGA-PRAD | 48.6 | 14.04 |
| GTEX-1E1VI-2726-SM-7MXVH | Prostate | TCGA-PRAD | 28.58 | 7.5 |
| GTEX-1EH9U-1826-SM-7P8QL | Prostate | TCGA-PRAD | 32.84 | 10.22 |
| GTEX-1EKGG-2826-SM-7MGXA | Prostate | TCGA-PRAD | 15.78 | 9.51 |
| GTEX-1EU9M-1526-SM-79OLI | Prostate | TCGA-PRAD | 38.15 | 17.95 |
| GTEX-1F6I4-1326-SM-7P8QQ | Prostate | TCGA-PRAD | 27.69 | 13.74 |
| GTEX-1GF9V-0726-SM-9KNU4 | Prostate | TCGA-PRAD | 33.84 | 11.92 |
| GTEX-1GF9W-2026-SM-7MXVO | Prostate | TCGA-PRAD | 46.59 | 11.56 |
| GTEX-1GL5R-1226-SM-7MXUY | Prostate | TCGA-PRAD | 41.94 | 10.12 |

| | | | | |
|---|---|---|---|---|
| GTEX-1GN2E-2026-SM-9JGGG | Prostate | TCGA-PRAD | 14.94 | 7.35 |
| GTEX-1GN73-1626-SM-9WYTS | Prostate | TCGA-PRAD | 43.8 | 13.26 |
| GTEX-1GPI7-1626-SM-7P8TN | Prostate | TCGA-PRAD | 35.48 | 11.58 |
| GTEX-1GZ2Q-2426-SM-9KNV8 | Prostate | TCGA-PRAD | 36.62 | 11.81 |
| GTEX-1GZ4I-2226-SM-9WPPO | Prostate | TCGA-PRAD | 30.92 | 13.64 |
| GTEX-1H1E6-0526-SM-9WPQ3 | Prostate | TCGA-PRAD | 35.83 | 15.72 |
| GTEX-1H3NZ-2826-SM-9WPQ7 | Prostate | TCGA-PRAD | 26.7 | 12.78 |
| GTEX-1H3VY-2726-SM-A96TL | Prostate | TCGA-PRAD | 41.79 | 14.76 |
| GTEX-1HBPI-0326-SM-A9G3N | Prostate | TCGA-PRAD | 13.36 | 8.95 |
| GTEX-1HBPN-2126-SM-9WPNQ | Prostate | TCGA-PRAD | 30.54 | 11.49 |
| GTEX-1HFI6-2326-SM-9WYUN | Prostate | TCGA-PRAD | 39.42 | 15.35 |
| GTEX-1HKZK-1626-SM-B2LWH | Prostate | TCGA-PRAD | 28.53 | 15.15 |
| GTEX-1HSKV-1726-SM-C1YRO | Prostate | TCGA-PRAD | 25.22 | 11.46 |
| GTEX-1HSMO-2326-SM-CGQF3 | Prostate | TCGA-PRAD | 45.72 | 14.64 |
| GTEX-1HSMQ-1826-SM-A9SMJ | Prostate | TCGA-PRAD | 19.6 | 8.04 |
| GTEX-1I19N-2226-SM-A96TS | Prostate | TCGA-PRAD | 28.97 | 15.03 |
| GTEX-1I1GP-2026-SM-CE6SZ | Prostate | TCGA-PRAD | 48.55 | 13.33 |
| GTEX-1I1GU-1126-SM-A96S9 | Prostate | TCGA-PRAD | 60.19 | 24.14 |
| GTEX-1I1GV-1926-SM-CNPQH | Prostate | TCGA-PRAD | 30.13 | 10.9 |
| GTEX-1I6K6-2126-SM-B2LX6 | Prostate | TCGA-PRAD | 28.81 | 9.53 |
| GTEX-1I6K7-2326-SM-ARZM2 | Prostate | TCGA-PRAD | 34.06 | 16.11 |
| GTEX-1ICLZ-2626-SM-A9G2K | Prostate | TCGA-PRAD | 47.52 | 16.23 |
| GTEX-1IDJD-2026-SM-A96TC | Prostate | TCGA-PRAD | 29.51 | 12.33 |
| GTEX-1IDJF-2126-SM-AHZ2S | Prostate | TCGA-PRAD | 39.96 | 14.65 |
| GTEX-1IDJH-2426-SM-CNNQW | Prostate | TCGA-PRAD | 36.98 | 14.68 |
| GTEX-1IL2U-2826-SM-A96SR | Prostate | TCGA-PRAD | 24.56 | 11.18 |
| GTEX-1J8Q3-0926-SM-A96SU | Prostate | TCGA-PRAD | 65.32 | 21.02 |
| GTEX-1JJ6O-0726-SM-CY8HS | Prostate | TCGA-PRAD | 26.36 | 11.59 |
| GTEX-1JJE9-0426-SM-D4P2P | Prostate | TCGA-PRAD | 26.79 | 10.39 |
| GTEX-1JK1U-0926-SM-AHZ43 | Prostate | TCGA-PRAD | 43.67 | 14.66 |
| GTEX-1JKYN-0926-SM-AHZ47 | Prostate | TCGA-PRAD | 34.21 | 11 |

| | | | | |
|---|---|---|---|---|
| GTEX-1JKYR-1726-SM-C1YR6 | Prostate | TCGA-PRAD | 46.16 | 15.16 |
| GTEX-1JMLX-1226-SM-CY8HW | Prostate | TCGA-PRAD | 20.21 | 6.99 |
| GTEX-1JMPZ-2326-SM-CE6RW | Prostate | TCGA-PRAD | 38.81 | 16.38 |
| GTEX-1JMQK-2226-SM-ARU8U | Prostate | TCGA-PRAD | 33.61 | 14.6 |
| GTEX-1JN6P-2626-SM-D4P3C | Prostate | TCGA-PRAD | 26.72 | 13.93 |
| GTEX-1K2DA-1226-SM-CGQGH | Prostate | TCGA-PRAD | 32.08 | 13.38 |
| GTEX-1KAFJ-1226-SM-D4P46 | Prostate | TCGA-PRAD | 28.22 | 14.45 |
| GTEX-1KANA-1026-SM-D3L9L | Prostate | TCGA-PRAD | 41.03 | 12.83 |
| GTEX-1KANB-2026-SM-DIPEY | Prostate | TCGA-PRAD | 34.49 | 13.5 |
| GTEX-1KWVE-1226-SM-EV7AG | Prostate | TCGA-PRAD | 21.04 | 7.78 |
| GTEX-1KXAM-1126-SM-E9TJU | Prostate | TCGA-PRAD | 34.94 | 14.91 |
| GTEX-1L5NE-1026-SM-D3LAP | Prostate | TCGA-PRAD | 45.33 | 14.96 |
| GTEX-1LBAC-1926-SM-DHXK2 | Prostate | TCGA-PRAD | 24.67 | 10.6 |
| GTEX-1LG7Y-2426-SM-EVR57 | Prostate | TCGA-PRAD | 18.53 | 8.81 |
| GTEX-1LG7Z-0826-SM-CXKYN | Prostate | TCGA-PRAD | 20.56 | 8.56 |
| GTEX-1LSNL-2326-SM-DHXKN | Prostate | TCGA-PRAD | 31.55 | 10.3 |
| GTEX-1MGNQ-2226-SM-EV7A2 | Prostate | TCGA-PRAD | 37.56 | 14.73 |
| GTEX-1MJK2-0726-SM-DTXE7 | Prostate | TCGA-PRAD | 38.19 | 14.71 |
| GTEX-1MUQO-1526-SM-DPRYK | Prostate | TCGA-PRAD | 36.51 | 12.09 |
| GTEX-1N2EE-1026-SM-EXUS8 | Prostate | TCGA-PRAD | 23.22 | 9.85 |
| GTEX-1OJC3-1926-SM-DPRYV | Prostate | TCGA-PRAD | 39.96 | 7.4 |
| GTEX-1OKEX-1826-SM-DTX8J | Prostate | TCGA-PRAD | 40.98 | 19.97 |
| GTEX-1P4AB-1926-SM-E6CQL | Prostate | TCGA-PRAD | 23.72 | 10.05 |
| GTEX-1PFEY-0626-SM-E76PH | Prostate | TCGA-PRAD | 43.57 | 11.85 |
| GTEX-1PIGE-1726-SM-E6CQH | Prostate | TCGA-PRAD | 35.77 | 14.01 |
| GTEX-1PPH8-1426-SM-DTXA3 | Prostate | TCGA-PRAD | 24.88 | 11.24 |
| GTEX-1PWST-2626-SM-E8VN8 | Prostate | TCGA-PRAD | 57.38 | 13.39 |
| GTEX-1QCLY-1826-SM-DTX89 | Prostate | TCGA-PRAD | 31.14 | 12.15 |
| GTEX-1QP29-0526-SM-EAZ3T | Prostate | TCGA-PRAD | 44.49 | 26.84 |
| GTEX-1QP2A-1526-SM-E6CPN | Prostate | TCGA-PRAD | 16.38 | 7.36 |
| GTEX-1QP6S-0726-SM-E76QS | Prostate | TCGA-PRAD | 39.09 | 14.87 |

| | | | | |
|---|---|---|---|---|
| GTEX-1R7EU-0126-SM-EVR4F | Prostate | TCGA-PRAD | 33.37 | 10.42 |
| GTEX-1RAZQ-0826-SM-EVR4L | Prostate | TCGA-PRAD | 33.05 | 11.89 |
| GTEX-1S3DN-1926-SM-E6CIR | Prostate | TCGA-PRAD | 26.78 | 11.87 |
| GTEX-1S5ZA-1626-SM-E9TKG | Prostate | TCGA-PRAD | 35.22 | 12.92 |
| GTEX-1S831-1226-SM-EVR5D | Prostate | TCGA-PRAD | 31.61 | 10.24 |
| GTEX-NFK9-2226-SM-3MJGP | Prostate | TCGA-PRAD | 18.5 | 10.1 |
| GTEX-NPJ8-2426-SM-3MJHL | Prostate | TCGA-PRAD | 44.99 | 14.77 |
| GTEX-OIZF-2026-SM-3MJH1 | Prostate | TCGA-PRAD | 35.68 | 13.78 |
| GTEX-OIZH-2026-SM-3NB1M | Prostate | TCGA-PRAD | 33.53 | 11.17 |
| GTEX-OOBJ-2026-SM-3NB1R | Prostate | TCGA-PRAD | 41.91 | 17.39 |
| GTEX-OOBK-2025-SM-3LK5S | Prostate | TCGA-PRAD | 38.48 | 14.67 |
| GTEX-OXRL-2026-SM-E6CHN | Prostate | TCGA-PRAD | 22.62 | 11.61 |
| GTEX-P4PQ-2026-SM-E9U4E | Prostate | TCGA-PRAD | 26.67 | 11.27 |
| GTEX-P4QS-2026-SM-3NMCG | Prostate | TCGA-PRAD | 38.28 | 14.5 |
| GTEX-PLZ6-1126-SM-3P5ZR | Prostate | TCGA-PRAD | 26.72 | 12.04 |
| GTEX-Q2AH-1426-SM-EXOHC | Prostate | TCGA-PRAD | 42.09 | 18.32 |
| GTEX-Q2AI-1126-SM-48U19 | Prostate | TCGA-PRAD | 42.99 | 12.71 |
| GTEX-QEG4-2226-SM-CXZJD | Prostate | TCGA-PRAD | 24.85 | 11.62 |
| GTEX-QEG5-2426-SM-EYYVC | Prostate | TCGA-PRAD | 46.48 | 20.9 |
| GTEX-QLQ7-1326-SM-CNNOR | Prostate | TCGA-PRAD | 32.13 | 10.72 |
| GTEX-QMRM-1426-SM-CM2U2 | Prostate | TCGA-PRAD | 33.18 | 12.21 |
| GTEX-QV31-1026-SM-CNNOS | Prostate | TCGA-PRAD | 37.57 | 14.14 |
| GTEX-QV44-1526-SM-DHXIP | Prostate | TCGA-PRAD | 30.23 | 9.29 |
| GTEX-REY6-2126-SM-48FD9 | Prostate | TCGA-PRAD | 30.44 | 12.72 |
| GTEX-RM2N-1126-SM-48FCY | Prostate | TCGA-PRAD | 35.35 | 13.71 |
| GTEX-RVPV-2526-SM-EYYVL | Prostate | TCGA-PRAD | 27.05 | 13.6 |
| GTEX-RWSA-2026-SM-47JX8 | Prostate | TCGA-PRAD | 31.49 | 10.73 |
| GTEX-S33H-1826-SM-4AD65 | Prostate | TCGA-PRAD | 38.11 | 19.34 |
| GTEX-S3XE-1326-SM-4AD4H | Prostate | TCGA-PRAD | 50.5 | 11.53 |
| GTEX-S4Q7-1026-SM-4AD75 | Prostate | TCGA-PRAD | 29.85 | 9.97 |
| GTEX-S95S-1026-SM-4B64M | Prostate | TCGA-PRAD | 34.15 | 16.67 |

| | | | | |
|---|---|---|---|---|
| GTEX-SIU8-2226-SM-EAZ2T | Prostate | TCGA-PRAD | 26.35 | 13.99 |
| GTEX-SJXC-2026-SM-4DM6N | Prostate | TCGA-PRAD | 33.31 | 12.88 |
| GTEX-SNMC-0926-SM-4DM5U | Prostate | TCGA-PRAD | 27.33 | 9.61 |
| GTEX-SNOS-1026-SM-4DM4Y | Prostate | TCGA-PRAD | 41.41 | 15.66 |
| GTEX-SUCS-1126-SM-4DM61 | Prostate | TCGA-PRAD | 48.5 | 16.76 |
| GTEX-T5JC-1226-SM-4DM7C | Prostate | TCGA-PRAD | 35.34 | 12.4 |
| GTEX-T6MN-2126-SM-EZ6LC | Prostate | TCGA-PRAD | 24.02 | 13.8 |
| GTEX-T8EM-1826-SM-4DM7F | Prostate | TCGA-PRAD | 22.08 | 8.36 |
| GTEX-U3ZH-0926-SM-4DXU4 | Prostate | TCGA-PRAD | 46.03 | 13.24 |
| GTEX-U3ZM-0926-SM-4DXSW | Prostate | TCGA-PRAD | 27.08 | 14.88 |
| GTEX-U412-2026-SM-4DXSI | Prostate | TCGA-PRAD | 30.94 | 13.14 |
| GTEX-U4B1-1426-SM-4DXTX | Prostate | TCGA-PRAD | 34.72 | 12.41 |
| GTEX-U8XE-2526-SM-4E3IT | Prostate | TCGA-PRAD | 53.24 | 18.67 |
| GTEX-UTHO-2726-SM-4JBH9 | Prostate | TCGA-PRAD | 31.47 | 9.98 |
| GTEX-V1D1-1926-SM-4JBGX | Prostate | TCGA-PRAD | 21.96 | 11.92 |
| GTEX-V955-1826-SM-4JBIL | Prostate | TCGA-PRAD | 42.87 | 12.69 |
| GTEX-VJYA-1226-SM-3GIJ6 | Prostate | TCGA-PRAD | 42.86 | 13.51 |
| GTEX-WFG7-1826-SM-3GIL1 | Prostate | TCGA-PRAD | 45.73 | 17.05 |
| GTEX-WFG8-1826-SM-4LVM4 | Prostate | TCGA-PRAD | 34.8 | 12.25 |
| GTEX-WFON-1626-SM-4LVMV | Prostate | TCGA-PRAD | 32.84 | 12.84 |
| GTEX-WH7G-1726-SM-4LVKY | Prostate | TCGA-PRAD | 35.32 | 14.75 |
| GTEX-WK11-2626-SM-4OOSK | Prostate | TCGA-PRAD | 43.31 | 16.32 |
| GTEX-WOFM-0326-SM-4OOSL | Prostate | TCGA-PRAD | 27.66 | 9.5 |
| GTEX-WY7C-2026-SM-4ONCM | Prostate | TCGA-PRAD | 33.35 | 11.45 |
| GTEX-WYJK-0626-SM-4ONCZ | Prostate | TCGA-PRAD | 38.59 | 16.44 |
| GTEX-WZTO-0926-SM-4PQZ3 | Prostate | TCGA-PRAD | 20.9 | 10.64 |
| GTEX-X5EB-1826-SM-4E3K8 | Prostate | TCGA-PRAD | 40.2 | 20.28 |
| GTEX-XAJ8-0626-SM-47JY4 | Prostate | TCGA-PRAD | 30.79 | 12.21 |
| GTEX-XGQ4-1826-SM-4AT6F | Prostate | TCGA-PRAD | 33.58 | 10.4 |
| GTEX-XMK1-1026-SM-4B65H | Prostate | TCGA-PRAD | 37.93 | 13.65 |
| GTEX-XPVG-2026-SM-4B65E | Prostate | TCGA-PRAD | 45.64 | 15.76 |

| | | | | |
|---|---|---|---|---|
| GTEX-XQ3S-2526-SM-4BOOG | Prostate | TCGA-PRAD | 28.19 | 13.11 |
| GTEX-XQ8I-2526-SM-EAZB3 | Prostate | TCGA-PRAD | 35.92 | 15.03 |
| GTEX-XXEK-1226-SM-4BRUY | Prostate | TCGA-PRAD | 69.17 | 19.94 |
| GTEX-Y8E4-2126-SM-5RQHO | Prostate | TCGA-PRAD | 33.25 | 11.13 |
| GTEX-Y9LG-1826-SM-4VBQC | Prostate | TCGA-PRAD | 24.81 | 11.55 |
| GTEX-YB5E-1826-SM-5IFI4 | Prostate | TCGA-PRAD | 23.4 | 10.11 |
| GTEX-YF7O-1926-SM-4W1YQ | Prostate | TCGA-PRAD | 47.81 | 15.32 |
| GTEX-YJ8A-1126-SM-5IFJU | Prostate | TCGA-PRAD | 51.53 | 8.34 |
| GTEX-ZDYS-1226-SM-5IJF3 | Prostate | TCGA-PRAD | 39.43 | 11.87 |
| GTEX-ZEX8-2026-SM-4WKF3 | Prostate | TCGA-PRAD | 50.33 | 15.19 |
| GTEX-ZG7Y-2226-SM-DO92I | Prostate | TCGA-PRAD | 46.3 | 19.05 |
| GTEX-ZLFU-1726-SM-4WWBU | Prostate | TCGA-PRAD | 39.29 | 9.54 |
| GTEX-ZPU1-2026-SM-57WFI | Prostate | TCGA-PRAD | 47 | 12.41 |
| GTEX-ZT9W-1626-SM-4YCDQ | Prostate | TCGA-PRAD | 38.42 | 13.99 |
| GTEX-ZTTD-2726-SM-57WFA | Prostate | TCGA-PRAD | 46 | 18.08 |
| GTEX-ZTX8-1026-SM-5EGHD | Prostate | TCGA-PRAD | 36.51 | 11.16 |
| GTEX-ZUA1-2826-SM-59HLE | Prostate | TCGA-PRAD | 22.23 | 9.41 |
| GTEX-ZVP2-0826-SM-59HJ6 | Prostate | TCGA-PRAD | 31.78 | 12.55 |
| GTEX-ZY6K-1526-SM-5GZXE | Prostate | TCGA-PRAD | 76.6 | 30.78 |
| GTEX-ZYFD-2526-SM-5E45L | Prostate | TCGA-PRAD | 35.64 | 12.59 |
| GTEX-ZZ64-0926-SM-5E44L | Prostate | TCGA-PRAD | 27.89 | 9.32 |
| GTEX-111CU-0926-SM-5EGIK | Stomach | TCGA-STAD | 22.5 | 7.55 |
| GTEX-111YS-1126-SM-5GZYQ | Stomach | TCGA-STAD | 17.81 | 5.46 |
| GTEX-1122O-1926-SM-5EGIQ | Stomach | TCGA-STAD | 20.42 | 5.03 |
| GTEX-117YW-2226-SM-5N9DB | Stomach | TCGA-STAD | 41.03 | 3.2 |
| GTEX-117YX-1026-SM-5H11V | Stomach | TCGA-STAD | 28.4 | 8.42 |
| GTEX-11DXX-1326-SM-5GIDZ | Stomach | TCGA-STAD | 27.78 | 6.25 |
| GTEX-11EI6-2426-SM-5PNVS | Stomach | TCGA-STAD | 26.66 | 9.55 |
| GTEX-11EM3-1026-SM-5A5KL | Stomach | TCGA-STAD | 20.73 | 6.45 |
| GTEX-11EQ9-1226-SM-5987E | Stomach | TCGA-STAD | 43.73 | 9.35 |
| GTEX-11GSP-2126-SM-5HL5E | Stomach | TCGA-STAD | 17.6 | 4.96 |

| | | | | |
|---|---|---|---|---|
| GTEX-11I78-1726-SM-5A5M3 | Stomach | TCGA-STAD | 22.27 | 6.87 |
| GTEX-11NSD-1426-SM-5HL67 | Stomach | TCGA-STAD | 16.49 | 5.48 |
| GTEX-11NUK-2426-SM-5BC4U | Stomach | TCGA-STAD | 29.7 | 6.28 |
| GTEX-11P7K-1726-SM-5GU6F | Stomach | TCGA-STAD | 52.78 | 17.08 |
| GTEX-11P82-0726-SM-5PNYL | Stomach | TCGA-STAD | 45.88 | 12.23 |
| GTEX-11TT1-0726-SM-5GU5A | Stomach | TCGA-STAD | 38.2 | 12.23 |
| GTEX-11VI4-0326-SM-5EQ6L | Stomach | TCGA-STAD | 38.6 | 12.52 |
| GTEX-11WQK-2626-SM-5EQ4K | Stomach | TCGA-STAD | 66.96 | 9.03 |
| GTEX-1211K-1426-SM-5FQTF | Stomach | TCGA-STAD | 39.38 | 9.07 |
| GTEX-12696-1726-SM-5EQLH | Stomach | TCGA-STAD | 15.25 | 4.69 |
| GTEX-12BJ1-1726-SM-5HL9B | Stomach | TCGA-STAD | 28.22 | 9.91 |
| GTEX-12C56-0526-SM-5FQST | Stomach | TCGA-STAD | 51.76 | 12.75 |
| GTEX-12WSD-2326-SM-59HKQ | Stomach | TCGA-STAD | 19.85 | 6.4 |
| GTEX-12WSG-2026-SM-5FQUU | Stomach | TCGA-STAD | 10.92 | 5.24 |
| GTEX-12WSJ-0926-SM-5P9JD | Stomach | TCGA-STAD | 43.55 | 13.54 |
| GTEX-12WSK-1426-SM-5CVNN | Stomach | TCGA-STAD | 19.71 | 4.32 |
| GTEX-13111-1226-SM-5GCNC | Stomach | TCGA-STAD | 44.27 | 10.57 |
| GTEX-13113-0726-SM-5LZUF | Stomach | TCGA-STAD | 22.02 | 4.97 |
| GTEX-131XE-2226-SM-5PNYX | Stomach | TCGA-STAD | 52.05 | 10.15 |
| GTEX-131XF-1226-SM-5HL8V | Stomach | TCGA-STAD | 23.86 | 7.78 |
| GTEX-131XG-1226-SM-5EGH9 | Stomach | TCGA-STAD | 33.34 | 13.37 |
| GTEX-132AR-2426-SM-5IFFD | Stomach | TCGA-STAD | 67.23 | 14.41 |
| GTEX-132QS-1826-SM-5IFFN | Stomach | TCGA-STAD | 26.83 | 4.98 |
| GTEX-1339X-1426-SM-5K7YO | Stomach | TCGA-STAD | 43.43 | 14.91 |
| GTEX-133LE-1326-SM-5IFGO | Stomach | TCGA-STAD | 36.42 | 10.23 |
| GTEX-1399R-1126-SM-5IFIO | Stomach | TCGA-STAD | 22.33 | 6.22 |
| GTEX-1399S-1626-SM-5P9GI | Stomach | TCGA-STAD | 28.95 | 5.27 |
| GTEX-1399U-1626-SM-5P9J3 | Stomach | TCGA-STAD | 52.02 | 11.75 |
| GTEX-139D8-2326-SM-5IFGE | Stomach | TCGA-STAD | 30.77 | 10.6 |
| GTEX-139YR-1926-SM-5LZXM | Stomach | TCGA-STAD | 25.02 | 6.69 |
| GTEX-13CF3-1926-SM-5K7WF | Stomach | TCGA-STAD | 22.79 | 7.22 |

| | | | | |
|---|---|---|---|---|
| GTEX-13D11-2126-SM-5IFH2 | Stomach | TCGA-STAD | 23.12 | 5.85 |
| GTEX-13FH7-1026-SM-5IJGF | Stomach | TCGA-STAD | 23.86 | 5.17 |
| GTEX-13FTW-1726-SM-5KM2B | Stomach | TCGA-STAD | 20.75 | 7.79 |
| GTEX-13FTX-0726-SM-5N9BI | Stomach | TCGA-STAD | 26.6 | 8.8 |
| GTEX-13N1W-2026-SM-5K7YU | Stomach | TCGA-STAD | 29.86 | 12.62 |
| GTEX-13O21-0926-SM-5IFGT | Stomach | TCGA-STAD | 21.67 | 10.02 |
| GTEX-13O3Q-2126-SM-5KM4C | Stomach | TCGA-STAD | 40.84 | 16.49 |
| GTEX-13O61-1126-SM-5L3FI | Stomach | TCGA-STAD | 53.21 | 10.83 |
| GTEX-13OVI-2426-SM-5KM4J | Stomach | TCGA-STAD | 35.03 | 5.69 |
| GTEX-13OVK-0926-SM-7KUFX | Stomach | TCGA-STAD | 14.99 | 6.6 |
| GTEX-13OW6-2526-SM-5IJEC | Stomach | TCGA-STAD | 17.73 | 5.06 |
| GTEX-13PVQ-2126-SM-5L3FW | Stomach | TCGA-STAD | 24.73 | 6.95 |
| GTEX-13PVR-2126-SM-73KVR | Stomach | TCGA-STAD | 14.36 | 5.02 |
| GTEX-13QBU-1126-SM-5LU44 | Stomach | TCGA-STAD | 19.83 | 6.95 |
| GTEX-13QJ3-2726-SM-5SI6L | Stomach | TCGA-STAD | 27.94 | 11.03 |
| GTEX-13RTK-0226-SM-5RQHR | Stomach | TCGA-STAD | 38.17 | 7.77 |
| GTEX-13S7M-1826-SM-5RQK6 | Stomach | TCGA-STAD | 23.1 | 6 |
| GTEX-13U4I-2526-SM-5SI8Z | Stomach | TCGA-STAD | 30.25 | 9.91 |
| GTEX-13X6H-1626-SM-5Q5CT | Stomach | TCGA-STAD | 40.97 | 12.03 |
| GTEX-13X6J-2126-SM-5TDCV | Stomach | TCGA-STAD | 17.5 | 6.36 |
| GTEX-144GM-1726-SM-5O9AS | Stomach | TCGA-STAD | 13.71 | 3.68 |
| GTEX-144GN-1226-SM-5O991 | Stomach | TCGA-STAD | 19.92 | 7.97 |
| GTEX-145LT-1426-SM-5O9B3 | Stomach | TCGA-STAD | 15.25 | 4.42 |
| GTEX-145ME-1026-SM-5O9B4 | Stomach | TCGA-STAD | 34.49 | 8.62 |
| GTEX-145MN-1326-SM-5NQ9S | Stomach | TCGA-STAD | 22.14 | 6.1 |
| GTEX-145MO-2226-SM-5Q5BN | Stomach | TCGA-STAD | 21.88 | 6.44 |
| GTEX-146FH-2126-SM-5SI9U | Stomach | TCGA-STAD | 50.95 | 16.81 |
| GTEX-146FR-2026-SM-5NQAI | Stomach | TCGA-STAD | 39.09 | 6.92 |
| GTEX-14B4R-0726-SM-5TDDR | Stomach | TCGA-STAD | 25.52 | 7.23 |
| GTEX-14BIL-1926-SM-73KYI | Stomach | TCGA-STAD | 12.12 | 6.24 |
| GTEX-14BMU-1126-SM-5RQJ8 | Stomach | TCGA-STAD | 26.02 | 8.71 |

| | | | | |
|---|---|---|---|---|
| GTEX-14BMV-2226-SM-5RQHX | Stomach | TCGA-STAD | 29.4 | 9.32 |
| GTEX-14DAQ-2826-SM-664MU | Stomach | TCGA-STAD | 16.03 | 6.12 |
| GTEX-14DAR-0926-SM-73KUQ | Stomach | TCGA-STAD | 13.99 | 5.1 |
| GTEX-14E1K-1526-SM-664MN | Stomach | TCGA-STAD | 62.13 | 12.42 |
| GTEX-14E6E-0926-SM-664N8 | Stomach | TCGA-STAD | 19.72 | 7.94 |
| GTEX-14ICL-0826-SM-5YY9W | Stomach | TCGA-STAD | 25.01 | 5.83 |
| GTEX-14JG6-1226-SM-664O6 | Stomach | TCGA-STAD | 30.13 | 9.3 |
| GTEX-14PHX-1026-SM-664OL | Stomach | TCGA-STAD | 26.55 | 6.03 |
| GTEX-14PHY-0926-SM-5ZZWL | Stomach | TCGA-STAD | 53.52 | 15.68 |
| GTEX-14PJ3-1426-SM-664O8 | Stomach | TCGA-STAD | 20.71 | 6.74 |
| GTEX-14PJ4-1626-SM-664OB | Stomach | TCGA-STAD | 18.39 | 4.89 |
| GTEX-14PJ5-0826-SM-6AJAR | Stomach | TCGA-STAD | 17.08 | 5.39 |
| GTEX-14PJ6-0826-SM-686Z9 | Stomach | TCGA-STAD | 16.65 | 4.03 |
| GTEX-14PKU-1126-SM-69LOU | Stomach | TCGA-STAD | 46.18 | 8.02 |
| GTEX-14PKV-1826-SM-69LO9 | Stomach | TCGA-STAD | 22.55 | 6.72 |
| GTEX-14PN3-1326-SM-6AJ9U | Stomach | TCGA-STAD | 48.12 | 12.91 |
| GTEX-15CHR-1326-SM-79OLT | Stomach | TCGA-STAD | 12.79 | 6.46 |
| GTEX-15DCE-1626-SM-6LPJM | Stomach | TCGA-STAD | 17.31 | 3.77 |
| GTEX-15DZA-1126-SM-6LPK8 | Stomach | TCGA-STAD | 15.91 | 5.5 |
| GTEX-15EU6-2526-SM-7KUGB | Stomach | TCGA-STAD | 50.73 | 14.08 |
| GTEX-15G19-1526-SM-6LPKP | Stomach | TCGA-STAD | 39.7 | 13.22 |
| GTEX-15G1A-1226-SM-7KUF3 | Stomach | TCGA-STAD | 40.73 | 13.16 |
| GTEX-15RIE-1026-SM-6LPJQ | Stomach | TCGA-STAD | 14.1 | 6.33 |
| GTEX-15RIF-1126-SM-7KUG2 | Stomach | TCGA-STAD | 53.26 | 15.5 |
| GTEX-15SHW-2126-SM-6M46Y | Stomach | TCGA-STAD | 39.34 | 13.34 |
| GTEX-15UF7-1226-SM-6M46G | Stomach | TCGA-STAD | 21.48 | 5.08 |
| GTEX-16AAH-1226-SM-79OO9 | Stomach | TCGA-STAD | 21.15 | 7.27 |
| GTEX-16NGA-1526-SM-72D56 | Stomach | TCGA-STAD | 35.71 | 11.46 |
| GTEX-17EUY-1526-SM-793CA | Stomach | TCGA-STAD | 50.2 | 16.07 |
| GTEX-17F98-1526-SM-7EWD8 | Stomach | TCGA-STAD | 26.79 | 9.93 |
| GTEX-17F9E-2226-SM-7IGNC | Stomach | TCGA-STAD | 22.39 | 6.15 |

| | | | | |
|---|---|---|---|---|
| GTEX-17GQL-1226-SM-7KFTS | Stomach | TCGA-STAD | 20.39 | 5.72 |
| GTEX-17HGU-2126-SM-7IGOV | Stomach | TCGA-STAD | 14.22 | 6.58 |
| GTEX-17HHE-1026-SM-79ON6 | Stomach | TCGA-STAD | 13.91 | 4.96 |
| GTEX-17HII-2426-SM-7KFTT | Stomach | TCGA-STAD | 18.62 | 7.29 |
| GTEX-17JCI-2226-SM-7LTA1 | Stomach | TCGA-STAD | 21.68 | 8.25 |
| GTEX-17KNJ-1526-SM-7LTA9 | Stomach | TCGA-STAD | 54.42 | 16.32 |
| GTEX-17MF6-2126-SM-7KFS7 | Stomach | TCGA-STAD | 27.49 | 8.66 |
| GTEX-18A66-2226-SM-7189E | Stomach | TCGA-STAD | 25.92 | 11.29 |
| GTEX-18A67-2626-SM-718AD | Stomach | TCGA-STAD | 36.26 | 15.91 |
| GTEX-18A7A-2726-SM-7LG6U | Stomach | TCGA-STAD | 13.2 | 3.4 |
| GTEX-18D9A-0626-SM-7KFTY | Stomach | TCGA-STAD | 35.91 | 8.85 |
| GTEX-18D9B-2226-SM-718BM | Stomach | TCGA-STAD | 33.29 | 11.25 |
| GTEX-18QFQ-1226-SM-718AE | Stomach | TCGA-STAD | 18.99 | 4.86 |
| GTEX-1A3MV-1226-SM-718BW | Stomach | TCGA-STAD | 33.95 | 7.99 |
| GTEX-1A3MX-1926-SM-72D7F | Stomach | TCGA-STAD | 26.8 | 9.03 |
| GTEX-1A8G6-1426-SM-731EZ | Stomach | TCGA-STAD | 14.63 | 4.51 |
| GTEX-1AMEY-1226-SM-731DU | Stomach | TCGA-STAD | 19.79 | 5.03 |
| GTEX-1AMFI-1226-SM-731CZ | Stomach | TCGA-STAD | 18.49 | 5.23 |
| GTEX-1AX9J-2626-SM-731EK | Stomach | TCGA-STAD | 26.47 | 7.16 |
| GTEX-1AYCT-1326-SM-7MKHB | Stomach | TCGA-STAD | 18.84 | 4.43 |
| GTEX-1B8KE-1326-SM-73KVI | Stomach | TCGA-STAD | 22.08 | 3.8 |
| GTEX-1B8KZ-1226-SM-793CP | Stomach | TCGA-STAD | 20.64 | 8.27 |
| GTEX-1B8L1-1526-SM-79OLK | Stomach | TCGA-STAD | 40.22 | 11.67 |
| GTEX-1B97I-1226-SM-7MGWQ | Stomach | TCGA-STAD | 17.67 | 5.21 |
| GTEX-1B97J-1826-SM-7IGPP | Stomach | TCGA-STAD | 22.74 | 5.4 |
| GTEX-1C475-1026-SM-7MKFJ | Stomach | TCGA-STAD | 19.64 | 4.78 |
| GTEX-1C64O-2226-SM-73KTV | Stomach | TCGA-STAD | 43.61 | 17.26 |
| GTEX-1CAMR-0726-SM-7EPIE | Stomach | TCGA-STAD | 31.18 | 9.91 |
| GTEX-1CB4E-1326-SM-7DUGD | Stomach | TCGA-STAD | 55.91 | 13.6 |
| GTEX-1CB4H-2426-SM-7PBYR | Stomach | TCGA-STAD | 37.49 | 13.66 |
| GTEX-1E1VI-2826-SM-7MGWU | Stomach | TCGA-STAD | 12.08 | 4.75 |

| | | | | |
|---|---|---|---|---|
| GTEX-1E2YA-2426-SM-7EPIG | Stomach | TCGA-STAD | 31.69 | 10.22 |
| GTEX-1EH9U-2226-SM-7PBXX | Stomach | TCGA-STAD | 35.07 | 16.18 |
| GTEX-1F5PK-1826-SM-7RHHX | Stomach | TCGA-STAD | 22.31 | 8.35 |
| GTEX-1F5PL-1426-SM-7MKFH | Stomach | TCGA-STAD | 31.12 | 7.17 |
| GTEX-1F75I-1826-SM-7MKGX | Stomach | TCGA-STAD | 43.25 | 16.12 |
| GTEX-1GF9U-1126-SM-AHZ4H | Stomach | TCGA-STAD | 16.86 | 7.1 |
| GTEX-1GL5R-1926-SM-9WG7U | Stomach | TCGA-STAD | 17.99 | 4.38 |
| GTEX-1GMRU-2526-SM-9WG7T | Stomach | TCGA-STAD | 19.49 | 4.66 |
| GTEX-1GN1W-2126-SM-9JGI7 | Stomach | TCGA-STAD | 25.75 | 8.76 |
| GTEX-1GPI7-1326-SM-9KNVA | Stomach | TCGA-STAD | 47.98 | 11.61 |
| GTEX-1GZ4H-1026-SM-9OSXL | Stomach | TCGA-STAD | 15.15 | 6.37 |
| GTEX-1H11D-2226-SM-9OSXN | Stomach | TCGA-STAD | 32.62 | 10.66 |
| GTEX-1H1DE-1226-SM-9JGHH | Stomach | TCGA-STAD | 37.16 | 13.34 |
| GTEX-1H1DG-2126-SM-9WPPB | Stomach | TCGA-STAD | 18.72 | 10.71 |
| GTEX-1H1E6-1626-SM-9OSWC | Stomach | TCGA-STAD | 15.51 | 3.84 |
| GTEX-1H23P-2726-SM-9JGHN | Stomach | TCGA-STAD | 28.03 | 9.48 |
| GTEX-1H3NZ-2226-SM-9WYUT | Stomach | TCGA-STAD | 46.66 | 12.32 |
| GTEX-1H3VY-2326-SM-9WG81 | Stomach | TCGA-STAD | 26.88 | 10.39 |
| GTEX-1HBPI-0726-SM-9WG7Z | Stomach | TCGA-STAD | 18.38 | 3.83 |
| GTEX-1HBPN-1626-SM-9WPNP | Stomach | TCGA-STAD | 15.42 | 4.36 |
| GTEX-1HCU8-1326-SM-C1YQQ | Stomach | TCGA-STAD | 28.37 | 6.77 |
| GTEX-1HCUA-1626-SM-A9SMG | Stomach | TCGA-STAD | 41.02 | 12.14 |
| GTEX-1HFI7-1526-SM-9WG7W | Stomach | TCGA-STAD | 27.98 | 12.06 |
| GTEX-1HKZK-1526-SM-B2LWG | Stomach | TCGA-STAD | 16.14 | 6.81 |
| GTEX-1HR9M-2326-SM-CNPPZ | Stomach | TCGA-STAD | 44.64 | 12.05 |
| GTEX-1HSEH-1526-SM-B2LXG | Stomach | TCGA-STAD | 24.3 | 7.08 |
| GTEX-1HSMQ-2426-SM-CE6RY | Stomach | TCGA-STAD | 37.85 | 8.91 |
| GTEX-1HT8W-2326-SM-A9SLA | Stomach | TCGA-STAD | 24.67 | 8.69 |
| GTEX-1I1GT-1726-SM-C1YS3 | Stomach | TCGA-STAD | 24.05 | 7.87 |
| GTEX-1I1HK-2026-SM-CNNRB | Stomach | TCGA-STAD | 10.97 | 4.58 |
| GTEX-1I4MK-0926-SM-A9SLD | Stomach | TCGA-STAD | 31.72 | 11.47 |

| | | | | |
|---|---|---|---|---|
| GTEX-1I6K7-1726-SM-B2LXQ | Stomach | TCGA-STAD | 21.95 | 11.29 |
| GTEX-1ICG6-1826-SM-AHZ2O | Stomach | TCGA-STAD | 16.27 | 5.36 |
| GTEX-1IDJD-2426-SM-C1YS6 | Stomach | TCGA-STAD | 27.12 | 7.89 |
| GTEX-1IDJF-1426-SM-ACKWK | Stomach | TCGA-STAD | 48.44 | 15.33 |
| GTEX-1IDJU-1526-SM-CNNR3 | Stomach | TCGA-STAD | 29.06 | 10.51 |
| GTEX-1IL2V-2026-SM-ARZMA | Stomach | TCGA-STAD | 30.77 | 11.62 |
| GTEX-1J8EW-2226-SM-AHZ3T | Stomach | TCGA-STAD | 18.67 | 6.59 |
| GTEX-1J8JJ-1726-SM-C1YQG | Stomach | TCGA-STAD | 10.8 | 4.48 |
| GTEX-1J8Q3-1926-SM-C1YSJ | Stomach | TCGA-STAD | 19.43 | 4.81 |
| GTEX-1J8QM-1126-SM-C1YQB | Stomach | TCGA-STAD | 18.37 | 6.28 |
| GTEX-1JK1U-1326-SM-AHZ45 | Stomach | TCGA-STAD | 19.74 | 7.12 |
| GTEX-1JKYN-1726-SM-AHZ48 | Stomach | TCGA-STAD | 33.03 | 9 |
| GTEX-1JMLX-2426-SM-CXZJQ | Stomach | TCGA-STAD | 25.13 | 3.41 |
| GTEX-1JMPY-0926-SM-ARL92 | Stomach | TCGA-STAD | 36.12 | 5.95 |
| GTEX-1JMPZ-1826-SM-CXZJT | Stomach | TCGA-STAD | 31.33 | 8.3 |
| GTEX-1JMQJ-1326-SM-D4P3B | Stomach | TCGA-STAD | 25.4 | 8.56 |
| GTEX-1JN1M-2926-SM-CNNPJ | Stomach | TCGA-STAD | 43.38 | 10.7 |
| GTEX-1JN76-0926-SM-ARZNH | Stomach | TCGA-STAD | 38.37 | 11.53 |
| GTEX-1K2DA-0826-SM-CGQGD | Stomach | TCGA-STAD | 37.22 | 12.62 |
| GTEX-1KANA-1126-SM-D3L9M | Stomach | TCGA-STAD | 28.46 | 10.65 |
| GTEX-1KD5A-1426-SM-D4P3V | Stomach | TCGA-STAD | 31.95 | 15.23 |
| GTEX-1KXAM-0926-SM-CXZKA | Stomach | TCGA-STAD | 23.64 | 7.93 |
| GTEX-1L5NE-1526-SM-DIPFW | Stomach | TCGA-STAD | 57.5 | 12.73 |
| GTEX-1LBAC-1426-SM-CXZKT | Stomach | TCGA-STAD | 21.75 | 6.91 |
| GTEX-1LG7Z-1226-SM-DHXK4 | Stomach | TCGA-STAD | 19.13 | 3.9 |
| GTEX-1LGOU-1526-SM-D3LAJ | Stomach | TCGA-STAD | 20.55 | 6.96 |
| GTEX-1LGRB-1726-SM-CNNRA | Stomach | TCGA-STAD | 21.16 | 5.08 |
| GTEX-1LSNM-1026-SM-DHXL2 | Stomach | TCGA-STAD | 13.56 | 3.78 |
| GTEX-1LVAM-1426-SM-E9TJ4 | Stomach | TCGA-STAD | 49.78 | 12.09 |
| GTEX-1LVAN-2426-SM-CNNPZ | Stomach | TCGA-STAD | 11.83 | 5.02 |
| GTEX-1MCYP-1126-SM-E9J2P | Stomach | TCGA-STAD | 37.78 | 8.22 |

| | | | | |
|---|---|---|---|---|
| GTEX-1MGNQ-1826-SM-DPRYI | Stomach | TCGA-STAD | 23.24 | 5.15 |
| GTEX-1MJIX-1126-SM-EV7AO | Stomach | TCGA-STAD | 28.37 | 9.13 |
| GTEX-1MUQO-1926-SM-EAZ59 | Stomach | TCGA-STAD | 27.37 | 10.95 |
| GTEX-1N2DV-0926-SM-E9TJT | Stomach | TCGA-STAD | 45.7 | 14.07 |
| GTEX-1NSGN-1926-SM-E9J36 | Stomach | TCGA-STAD | 33.56 | 15.28 |
| GTEX-1PIGE-1126-SM-EWRMV | Stomach | TCGA-STAD | 56.61 | 14.55 |
| GTEX-1PPH8-1326-SM-DTXFO | Stomach | TCGA-STAD | 34.95 | 7.83 |
| GTEX-1QCLY-1126-SM-EAZ4J | Stomach | TCGA-STAD | 27 | 7.64 |
| GTEX-1QCLZ-1826-SM-DPRXP | Stomach | TCGA-STAD | 18.13 | 4.69 |
| GTEX-1QMI2-2126-SM-DPRXQ | Stomach | TCGA-STAD | 17.13 | 3.62 |
| GTEX-1QP28-2126-SM-DTX8B | Stomach | TCGA-STAD | 36.65 | 7.86 |
| GTEX-1QP29-2326-SM-EXOJO | Stomach | TCGA-STAD | 29.68 | 10.05 |
| GTEX-1QP2A-2126-SM-DPRZ7 | Stomach | TCGA-STAD | 16.37 | 4.04 |
| GTEX-1QP67-1026-SM-EVR47 | Stomach | TCGA-STAD | 48.04 | 11.16 |
| GTEX-1QPFJ-1526-SM-DPRXT | Stomach | TCGA-STAD | 23.08 | 5.81 |
| GTEX-1QW4Y-0726-SM-DTX8E | Stomach | TCGA-STAD | 43.32 | 13.86 |
| GTEX-1R7EU-1126-SM-EAZ4M | Stomach | TCGA-STAD | 17.17 | 6.05 |
| GTEX-1R9K5-1026-SM-EAZ4F | Stomach | TCGA-STAD | 24.58 | 7.95 |
| GTEX-1RAZR-1526-SM-EAZ4U | Stomach | TCGA-STAD | 37.01 | 19.12 |
| GTEX-1RB15-1026-SM-E9U6V | Stomach | TCGA-STAD | 16.6 | 6.71 |
| GTEX-1RMOY-1126-SM-EV79E | Stomach | TCGA-STAD | 21.35 | 6.74 |
| GTEX-1RQEC-2126-SM-DPS15 | Stomach | TCGA-STAD | 13.06 | 4.4 |
| GTEX-1S3DN-1526-SM-EVYCB | Stomach | TCGA-STAD | 26.28 | 13.66 |
| GTEX-1S82Z-1726-SM-E6CHZ | Stomach | TCGA-STAD | 26.78 | 9.06 |
| GTEX-1S831-1126-SM-EWROI | Stomach | TCGA-STAD | 48.99 | 13.43 |
| GTEX-NFK9-1526-SM-3LK7B | Stomach | TCGA-STAD | 19.24 | 6.58 |
| GTEX-O5YV-1926-SM-EV797 | Stomach | TCGA-STAD | 32.14 | 7.29 |
| GTEX-O5YW-1526-SM-3MJGL | Stomach | TCGA-STAD | 18.1 | 6.52 |
| GTEX-OHPK-1526-SM-3MJGM | Stomach | TCGA-STAD | 25.59 | 10.48 |
| GTEX-OHPM-1526-SM-EV796 | Stomach | TCGA-STAD | 20.07 | 6.88 |
| GTEX-OIZF-1526-SM-2I5GR | Stomach | TCGA-STAD | 9.67 | 6.18 |

| | | | | |
|---|---|---|---|---|
| GTEX-OIZH-1526-SM-3NB1J | Stomach | TCGA-STAD | 39.36 | 14.44 |
| GTEX-OOBJ-1526-SM-3NB1Q | Stomach | TCGA-STAD | 25.51 | 8.97 |
| GTEX-OXRK-1626-SM-3NB17 | Stomach | TCGA-STAD | 17.19 | 6.06 |
| GTEX-OXRL-1526-SM-E6CHQ | Stomach | TCGA-STAD | 12.22 | 4.73 |
| GTEX-P4PP-1526-SM-3P61M | Stomach | TCGA-STAD | 23.79 | 6.69 |
| GTEX-P4PQ-1526-SM-3NMCK | Stomach | TCGA-STAD | 47.09 | 19.24 |
| GTEX-P4QT-1526-SM-3NMCT | Stomach | TCGA-STAD | 50.27 | 10.63 |
| GTEX-P78B-1826-SM-3P5YX | Stomach | TCGA-STAD | 28.42 | 9.28 |
| GTEX-PLZ5-1226-SM-EXOHA | Stomach | TCGA-STAD | 25.26 | 9.99 |
| GTEX-PLZ6-0826-SM-3P61K | Stomach | TCGA-STAD | 38.01 | 15.3 |
| GTEX-POMQ-0826-SM-3P61H | Stomach | TCGA-STAD | 31.55 | 10.67 |
| GTEX-PW2O-1226-SM-48TCH | Stomach | TCGA-STAD | 16.52 | 6.37 |
| GTEX-PWCY-0926-SM-48TD7 | Stomach | TCGA-STAD | 21.41 | 5.93 |
| GTEX-PWN1-1526-SM-48TDA | Stomach | TCGA-STAD | 27.64 | 12.13 |
| GTEX-PX3G-1526-SM-48U11 | Stomach | TCGA-STAD | 12.18 | 4.39 |
| GTEX-Q2AH-1126-SM-48TZM | Stomach | TCGA-STAD | 26.45 | 11.23 |
| GTEX-Q2AI-0826-SM-48TZO | Stomach | TCGA-STAD | 23.7 | 10.47 |
| GTEX-Q734-1026-SM-48U16 | Stomach | TCGA-STAD | 18.76 | 7.17 |
| GTEX-QCQG-0526-SM-48U2A | Stomach | TCGA-STAD | 30.98 | 10.28 |
| GTEX-QDT8-2026-SM-EZ6LS | Stomach | TCGA-STAD | 39.73 | 12.8 |
| GTEX-QDVJ-1426-SM-48U1Y | Stomach | TCGA-STAD | 25.47 | 10.68 |
| GTEX-QDVN-1226-SM-48TZ5 | Stomach | TCGA-STAD | 30.6 | 7.57 |
| GTEX-QLQ7-0826-SM-447B3 | Stomach | TCGA-STAD | 17.77 | 6 |
| GTEX-QLQW-0726-SM-447AA | Stomach | TCGA-STAD | 12.21 | 4.98 |
| GTEX-QMRM-1126-SM-447BN | Stomach | TCGA-STAD | 22.64 | 7.93 |
| GTEX-QV31-0626-SM-447C5 | Stomach | TCGA-STAD | 23.24 | 8.09 |
| GTEX-QV44-1226-SM-CJI1S | Stomach | TCGA-STAD | 15.19 | 5.1 |
| GTEX-QXCU-1926-SM-48FE4 | Stomach | TCGA-STAD | 23.86 | 6.11 |
| GTEX-R53T-1226-SM-48FCT | Stomach | TCGA-STAD | 19.76 | 8.09 |
| GTEX-R55C-1026-SM-48FCM | Stomach | TCGA-STAD | 45.4 | 14.49 |
| GTEX-R55D-1526-SM-48FEJ | Stomach | TCGA-STAD | 35.88 | 16.07 |

| | | | | |
|---|---|---|---|---|
| GTEX-R55G-1126-SM-48FDG | Stomach | TCGA-STAD | 49.34 | 17.59 |
| GTEX-RM2N-0826-SM-48FD3 | Stomach | TCGA-STAD | 17.39 | 6.61 |
| GTEX-RTLS-2626-SM-46MUJ | Stomach | TCGA-STAD | 20.41 | 6.72 |
| GTEX-RU1J-0526-SM-46MUT | Stomach | TCGA-STAD | 39.46 | 10.57 |
| GTEX-RUSQ-1126-SM-EAZAU | Stomach | TCGA-STAD | 68.5 | 22.57 |
| GTEX-RWS6-0926-SM-47JXE | Stomach | TCGA-STAD | 12.89 | 4.43 |
| GTEX-S341-0626-SM-4AD5T | Stomach | TCGA-STAD | 22.34 | 6.93 |
| GTEX-S3XE-1026-SM-4AD4O | Stomach | TCGA-STAD | 22.76 | 5.55 |
| GTEX-S4P3-0726-SM-4AD57 | Stomach | TCGA-STAD | 33.1 | 15.76 |
| GTEX-S4Q7-0726-SM-4AD5F | Stomach | TCGA-STAD | 23.68 | 6.91 |
| GTEX-S4UY-1626-SM-4AD55 | Stomach | TCGA-STAD | 23.96 | 5.96 |
| GTEX-S4Z8-1226-SM-4AD6W | Stomach | TCGA-STAD | 22 | 9.18 |
| GTEX-S7SF-0626-SM-4AD4V | Stomach | TCGA-STAD | 37.41 | 16.36 |
| GTEX-S95S-0826-SM-4B64N | Stomach | TCGA-STAD | 19 | 5.3 |
| GTEX-SNMC-0626-SM-4DM6H | Stomach | TCGA-STAD | 21.31 | 10.27 |
| GTEX-SNOS-0826-SM-4DM5N | Stomach | TCGA-STAD | 34.17 | 6.74 |
| GTEX-SUCS-0926-SM-4DM4Z | Stomach | TCGA-STAD | 21.87 | 8.27 |
| GTEX-T5JC-1926-SM-4DM6Q | Stomach | TCGA-STAD | 34.88 | 11.63 |
| GTEX-T5JW-0926-SM-4DM5K | Stomach | TCGA-STAD | 19.32 | 8.02 |
| GTEX-T6MO-0726-SM-4DM58 | Stomach | TCGA-STAD | 19.38 | 4.05 |
| GTEX-T8EM-1226-SM-4DM5J | Stomach | TCGA-STAD | 35.92 | 13.47 |
| GTEX-TKQ1-0526-SM-4DXTG | Stomach | TCGA-STAD | 44.66 | 13.4 |
| GTEX-TKQ2-0926-SM-4DXU5 | Stomach | TCGA-STAD | 42.78 | 7.55 |
| GTEX-TMMY-1626-SM-4DXTY | Stomach | TCGA-STAD | 31.18 | 7.43 |
| GTEX-U3ZN-2026-SM-4DXUC | Stomach | TCGA-STAD | 15.49 | 4.39 |
| GTEX-U4B1-1026-SM-4DXT1 | Stomach | TCGA-STAD | 39.43 | 18.54 |
| GTEX-UJHI-1026-SM-4IHJP | Stomach | TCGA-STAD | 41.45 | 9.08 |
| GTEX-UJMC-1226-SM-4IHLI | Stomach | TCGA-STAD | 16.16 | 4.26 |
| GTEX-UPK5-2126-SM-4JBJK | Stomach | TCGA-STAD | 42.47 | 12.23 |
| GTEX-V1D1-1726-SM-4JBHB | Stomach | TCGA-STAD | 15.67 | 5.7 |
| GTEX-V955-1326-SM-4JBHR | Stomach | TCGA-STAD | 21.39 | 5.09 |

| | | | | |
|---|---|---|---|---|
| GTEX-VJYA-1026-SM-4KL21 | Stomach | TCGA-STAD | 18.21 | 5.99 |
| GTEX-W5WG-1726-SM-4LMI5 | Stomach | TCGA-STAD | 13.61 | 4.89 |
| GTEX-WEY5-1226-SM-4LMIQ | Stomach | TCGA-STAD | 23.88 | 6.35 |
| GTEX-WFG7-1326-SM-4LMK1 | Stomach | TCGA-STAD | 26.92 | 10.91 |
| GTEX-WFG8-1326-SM-4LVN3 | Stomach | TCGA-STAD | 27.16 | 6.44 |
| GTEX-WFJO-1126-SM-4LVLZ | Stomach | TCGA-STAD | 45.67 | 10.69 |
| GTEX-WFON-1126-SM-4LVMA | Stomach | TCGA-STAD | 13.29 | 4.56 |
| GTEX-WH7G-1526-SM-4LVMX | Stomach | TCGA-STAD | 42.24 | 10.12 |
| GTEX-WHPG-0426-SM-4M1XW | Stomach | TCGA-STAD | 44.75 | 19.79 |
| GTEX-WHSB-1226-SM-4M1XR | Stomach | TCGA-STAD | 25.57 | 9.54 |
| GTEX-WHWD-1426-SM-4OORU | Stomach | TCGA-STAD | 20.51 | 5.9 |
| GTEX-WRHK-0626-SM-4MVOE | Stomach | TCGA-STAD | 21.53 | 5.65 |
| GTEX-WXYG-1626-SM-4ONCR | Stomach | TCGA-STAD | 18.26 | 7.24 |
| GTEX-WY7C-1726-SM-4ONCC | Stomach | TCGA-STAD | 28.13 | 7.92 |
| GTEX-WYJK-2526-SM-4ONDF | Stomach | TCGA-STAD | 23.42 | 9.53 |
| GTEX-WYVS-1326-SM-4ONCQ | Stomach | TCGA-STAD | 16.39 | 6.28 |
| GTEX-WZTO-2126-SM-4PQYW | Stomach | TCGA-STAD | 20.8 | 8.94 |
| GTEX-X15G-1126-SM-4PQZG | Stomach | TCGA-STAD | 14.56 | 6.84 |
| GTEX-XAJ8-0326-SM-47JYI | Stomach | TCGA-STAD | 39.83 | 10.63 |
| GTEX-XBED-1226-SM-4AT5V | Stomach | TCGA-STAD | 20.09 | 8.57 |
| GTEX-XBEW-1826-SM-4RTWX | Stomach | TCGA-STAD | 29.12 | 10.4 |
| GTEX-XMK1-1526-SM-EYYVO | Stomach | TCGA-STAD | 28.38 | 10.9 |
| GTEX-XPVG-1526-SM-4B66C | Stomach | TCGA-STAD | 39.22 | 11.85 |
| GTEX-XQ8I-2026-SM-4BOOL | Stomach | TCGA-STAD | 27.55 | 11.29 |
| GTEX-XUZC-0726-SM-4BOPH | Stomach | TCGA-STAD | 14.9 | 4.88 |
| GTEX-XV7Q-1326-SM-4BRWM | Stomach | TCGA-STAD | 29.75 | 10.46 |
| GTEX-XXEK-0826-SM-4BRWG | Stomach | TCGA-STAD | 14.51 | 4.37 |
| GTEX-Y114-1226-SM-4TT88 | Stomach | TCGA-STAD | 46.4 | 14.39 |
| GTEX-Y3I4-1626-SM-4TT7W | Stomach | TCGA-STAD | 18.19 | 5.81 |
| GTEX-Y3IK-1426-SM-4YCES | Stomach | TCGA-STAD | 28.87 | 9.48 |
| GTEX-Y5LM-1326-SM-5RQIS | Stomach | TCGA-STAD | 39.75 | 15.24 |

| | | | | |
|---|---|---|---|---|
| GTEX-Y5V6-1326-SM-4VDTF | Stomach | TCGA-STAD | 14.11 | 3.05 |
| GTEX-Y8E4-1526-SM-4WWDI | Stomach | TCGA-STAD | 27.2 | 8.73 |
| GTEX-Y8LW-0826-SM-4WWDO | Stomach | TCGA-STAD | 18.86 | 5.99 |
| GTEX-Y9LG-1026-SM-5IFJN | Stomach | TCGA-STAD | 30.13 | 7.45 |
| GTEX-YB5E-1126-SM-664NU | Stomach | TCGA-STAD | 18.26 | 5.2 |
| GTEX-YEC3-1426-SM-D4P1S | Stomach | TCGA-STAD | 38.04 | 14.1 |
| GTEX-YEC4-1426-SM-5IFHS | Stomach | TCGA-STAD | 20.58 | 6.42 |
| GTEX-YFCO-0826-SM-5LUAG | Stomach | TCGA-STAD | 19.68 | 6.65 |
| GTEX-YJ8A-1526-SM-5P9FT | Stomach | TCGA-STAD | 27.28 | 6.31 |
| GTEX-ZA64-0826-SM-5HL9U | Stomach | TCGA-STAD | 22.12 | 7.43 |
| GTEX-ZAB4-1526-SM-5CVN7 | Stomach | TCGA-STAD | 14.5 | 6.06 |
| GTEX-ZDYS-1926-SM-5HL59 | Stomach | TCGA-STAD | 18.99 | 6.53 |
| GTEX-ZE7O-3026-SM-51MS4 | Stomach | TCGA-STAD | 33.96 | 9.73 |
| GTEX-ZEX8-1626-SM-4WKG7 | Stomach | TCGA-STAD | 14.23 | 4.49 |
| GTEX-ZF29-1526-SM-4WKF7 | Stomach | TCGA-STAD | 18.11 | 4.58 |
| GTEX-ZF2S-1426-SM-57WET | Stomach | TCGA-STAD | 18.34 | 6.64 |
| GTEX-ZGAY-1626-SM-DNZZ3 | Stomach | TCGA-STAD | 19.57 | 4.11 |
| GTEX-ZLFU-1326-SM-DO11R | Stomach | TCGA-STAD | 20.63 | 5.03 |
| GTEX-ZLV1-0826-SM-DO11P | Stomach | TCGA-STAD | 19.06 | 4.81 |
| GTEX-ZLWG-1526-SM-DO12F | Stomach | TCGA-STAD | 27.09 | 6.66 |
| GTEX-ZP4G-1226-SM-4WWCJ | Stomach | TCGA-STAD | 23.73 | 6.63 |
| GTEX-ZPIC-2026-SM-57WG3 | Stomach | TCGA-STAD | 43.97 | 12.64 |
| GTEX-ZPU1-1626-SM-4WWB2 | Stomach | TCGA-STAD | 13.89 | 6.1 |
| GTEX-ZQUD-0826-SM-57WDQ | Stomach | TCGA-STAD | 18.74 | 7.6 |
| GTEX-ZT9W-1526-SM-4YCDE | Stomach | TCGA-STAD | 17.39 | 4.78 |
| GTEX-ZTX8-0426-SM-59HLG | Stomach | TCGA-STAD | 46.99 | 15.46 |
| GTEX-ZV6S-0926-SM-57WGB | Stomach | TCGA-STAD | 49.8 | 13.54 |
| GTEX-ZV7C-1626-SM-5NQ7E | Stomach | TCGA-STAD | 18.14 | 5.18 |
| GTEX-ZVP2-1626-SM-5GU5D | Stomach | TCGA-STAD | 18.41 | 4.14 |
| GTEX-ZVT2-1626-SM-51MRC | Stomach | TCGA-STAD | 14.29 | 3.71 |
| GTEX-ZVT3-2126-SM-59HL2 | Stomach | TCGA-STAD | 22.8 | 5.67 |

| | | | | |
|---|---|---|---|---|
| GTEX-ZVZP-1726-SM-5GZWY | Stomach | TCGA-STAD | 21.04 | 4.46 |
| GTEX-ZXES-1426-SM-5NQ8S | Stomach | TCGA-STAD | 21.6 | 4.19 |
| GTEX-ZY6K-0726-SM-5A5L8 | Stomach | TCGA-STAD | 21.68 | 7.9 |
| GTEX-ZYFG-1326-SM-5GICJ | Stomach | TCGA-STAD | 18.38 | 4.62 |
| GTEX-ZYVF-2726-SM-5GID4 | Stomach | TCGA-STAD | 57.21 | 16.81 |
| GTEX-ZYY3-1726-SM-5EGH3 | Stomach | TCGA-STAD | 18.26 | 4.5 |
| GTEX-ZZ64-0426-SM-5E43F | Stomach | TCGA-STAD | 19.23 | 3.91 |
| GTEX-ZZPU-1426-SM-5GZZ6 | Stomach | TCGA-STAD | 24.56 | 7.6 |
| GTEX-111CU-0226-SM-5GZXC | Thyroid | TCGA-THCA | 70.62 | 13.9 |
| GTEX-111FC-1026-SM-5GZX1 | Thyroid | TCGA-THCA | 60.53 | 13.12 |
| GTEX-111VG-0526-SM-5N9BW | Thyroid | TCGA-THCA | 30.31 | 8.23 |
| GTEX-111YS-0726-SM-5GZY8 | Thyroid | TCGA-THCA | 64.24 | 9.64 |
| GTEX-1122O-0226-SM-5N9DA | Thyroid | TCGA-THCA | 50.22 | 8.6 |
| GTEX-1128S-0126-SM-5H12S | Thyroid | TCGA-THCA | 72.67 | 12.45 |
| GTEX-113JC-0126-SM-5EGJW | Thyroid | TCGA-THCA | 49.44 | 9.62 |
| GTEX-117XS-0526-SM-5987Q | Thyroid | TCGA-THCA | 54.47 | 10.42 |
| GTEX-117YW-0126-SM-5EGGN | Thyroid | TCGA-THCA | 49.11 | 13.88 |
| GTEX-117YX-1226-SM-5H11S | Thyroid | TCGA-THCA | 61.98 | 14.25 |
| GTEX-1192W-0126-SM-5EGGS | Thyroid | TCGA-THCA | 52.87 | 14.03 |
| GTEX-1192X-1126-SM-5EGGU | Thyroid | TCGA-THCA | 62.21 | 9.83 |
| GTEX-11DXX-0226-SM-5P9HL | Thyroid | TCGA-THCA | 59.1 | 11.53 |
| GTEX-11DXY-0426-SM-5H12R | Thyroid | TCGA-THCA | 58.15 | 15.77 |
| GTEX-11DXZ-0926-SM-5N9CG | Thyroid | TCGA-THCA | 49.69 | 15.31 |
| GTEX-11DYG-0826-SM-5N9GH | Thyroid | TCGA-THCA | 62.95 | 14.3 |
| GTEX-11DZ1-2726-SM-5A5KH | Thyroid | TCGA-THCA | 65.89 | 9.93 |
| GTEX-11EI6-0726-SM-59866 | Thyroid | TCGA-THCA | 56.51 | 11.67 |
| GTEX-11EM3-0126-SM-5985K | Thyroid | TCGA-THCA | 58.41 | 10.56 |
| GTEX-11EMC-0226-SM-5EGLP | Thyroid | TCGA-THCA | 56.34 | 11.11 |
| GTEX-11EQ8-0826-SM-5N9FG | Thyroid | TCGA-THCA | 78.59 | 9.69 |
| GTEX-11EQ9-0626-SM-5A5K1 | Thyroid | TCGA-THCA | 51.53 | 10.08 |
| GTEX-11GS4-0826-SM-5986J | Thyroid | TCGA-THCA | 64.31 | 14.38 |

| | | | | |
|---|---|---|---|---|
| GTEX-11GSO-0626-SM-5A5LW | Thyroid | TCGA-THCA | 56.53 | 11.88 |
| GTEX-11GSP-0126-SM-5A5KU | Thyroid | TCGA-THCA | 32.24 | 9.14 |
| GTEX-11I78-0526-SM-5986A | Thyroid | TCGA-THCA | 57.46 | 9.39 |
| GTEX-11LCK-0526-SM-5A5M9 | Thyroid | TCGA-THCA | 54.32 | 14.77 |
| GTEX-11NSD-0126-SM-5987F | Thyroid | TCGA-THCA | 59.26 | 9.63 |
| GTEX-11NUK-1026-SM-5HL5J | Thyroid | TCGA-THCA | 52.65 | 10.32 |
| GTEX-11NV4-0626-SM-5N9BR | Thyroid | TCGA-THCA | 51.42 | 16.35 |
| GTEX-11O72-2326-SM-5BC7H | Thyroid | TCGA-THCA | 66.43 | 14.02 |
| GTEX-11OF3-0626-SM-5BC4Y | Thyroid | TCGA-THCA | 63.71 | 14.55 |
| GTEX-11P7K-0226-SM-5986Z | Thyroid | TCGA-THCA | 53.05 | 8.06 |
| GTEX-11P81-0126-SM-5HL5Y | Thyroid | TCGA-THCA | 59.84 | 16.68 |
| GTEX-11P82-0226-SM-5HL4O | Thyroid | TCGA-THCA | 56.9 | 11.27 |
| GTEX-11TT1-1126-SM-5P9GV | Thyroid | TCGA-THCA | 58.73 | 20.21 |
| GTEX-11TTK-0826-SM-5N9EG | Thyroid | TCGA-THCA | 44.96 | 12.71 |
| GTEX-11TUW-0226-SM-5LU8X | Thyroid | TCGA-THCA | 74.92 | 10.85 |
| GTEX-11UD2-0626-SM-5GU6L | Thyroid | TCGA-THCA | 81.47 | 12.82 |
| GTEX-11VI4-0226-SM-5GU6C | Thyroid | TCGA-THCA | 59.71 | 10.93 |
| GTEX-11XUK-0226-SM-5EQLW | Thyroid | TCGA-THCA | 40.54 | 8.69 |
| GTEX-11ZTS-1126-SM-5LU9X | Thyroid | TCGA-THCA | 35.77 | 12.2 |
| GTEX-11ZTT-1026-SM-5EQKF | Thyroid | TCGA-THCA | 64.24 | 13.44 |
| GTEX-11ZVC-0126-SM-5986G | Thyroid | TCGA-THCA | 56.43 | 11.18 |
| GTEX-1211K-0726-SM-5FQUW | Thyroid | TCGA-THCA | 78.28 | 8.35 |
| GTEX-1212Z-0426-SM-5FQT6 | Thyroid | TCGA-THCA | 49.41 | 12.1 |
| GTEX-12584-0826-SM-5FQSK | Thyroid | TCGA-THCA | 46.08 | 11.07 |
| GTEX-12696-0326-SM-5EGL4 | Thyroid | TCGA-THCA | 94.19 | 12.46 |
| GTEX-1269C-0226-SM-5EGKS | Thyroid | TCGA-THCA | 52.64 | 9.78 |
| GTEX-12BJ1-0426-SM-5FQSO | Thyroid | TCGA-THCA | 49.68 | 9.43 |
| GTEX-12WSC-0826-SM-5EQ5Q | Thyroid | TCGA-THCA | 57.31 | 12.17 |
| GTEX-12WSD-0926-SM-5GCNL | Thyroid | TCGA-THCA | 57.92 | 14.9 |
| GTEX-12WSE-1226-SM-73KUF | Thyroid | TCGA-THCA | 53.72 | 12.19 |
| GTEX-12WSG-0226-SM-5EGIF | Thyroid | TCGA-THCA | 38.28 | 12.99 |

| | | | | |
|---|---|---|---|---|
| GTEX-12WSH-0226-SM-5GCOG | Thyroid | TCGA-THCA | 86.65 | 16.32 |
| GTEX-12WSJ-0326-SM-5GCMT | Thyroid | TCGA-THCA | 52.83 | 14.88 |
| GTEX-12WSK-0926-SM-5CVNQ | Thyroid | TCGA-THCA | 44.83 | 9.98 |
| GTEX-12WSL-0626-SM-5GCOY | Thyroid | TCGA-THCA | 79.73 | 13.86 |
| GTEX-12WSN-0726-SM-5GCMS | Thyroid | TCGA-THCA | 62.94 | 13.34 |
| GTEX-12ZZX-1226-SM-5EGHS | Thyroid | TCGA-THCA | 46.26 | 9.67 |
| GTEX-12ZZY-0826-SM-5EQMT | Thyroid | TCGA-THCA | 57.46 | 11.97 |
| GTEX-12ZZZ-1226-SM-59HK1 | Thyroid | TCGA-THCA | 82.93 | 13.59 |
| GTEX-13111-0226-SM-5EQ55 | Thyroid | TCGA-THCA | 56.57 | 13.85 |
| GTEX-13112-0326-SM-5P9IW | Thyroid | TCGA-THCA | 58.45 | 13.94 |
| GTEX-13113-0126-SM-5LZVX | Thyroid | TCGA-THCA | 50.76 | 6.65 |
| GTEX-1313W-0726-SM-5EGK1 | Thyroid | TCGA-THCA | 65.8 | 13.38 |
| GTEX-131XE-0126-SM-5LZVC | Thyroid | TCGA-THCA | 67.55 | 16.45 |
| GTEX-131XF-1826-SM-5EGKG | Thyroid | TCGA-THCA | 54.63 | 8.43 |
| GTEX-131XG-0226-SM-5IFG1 | Thyroid | TCGA-THCA | 53.53 | 7.19 |
| GTEX-131XH-0526-SM-5DUX7 | Thyroid | TCGA-THCA | 72.14 | 15.02 |
| GTEX-131YS-0726-SM-5P9G9 | Thyroid | TCGA-THCA | 47.05 | 11.82 |
| GTEX-132AR-1126-SM-5P9GA | Thyroid | TCGA-THCA | 59.76 | 10.86 |
| GTEX-132NY-1026-SM-5P9IY | Thyroid | TCGA-THCA | 55.74 | 18.86 |
| GTEX-132QS-0326-SM-5IJFN | Thyroid | TCGA-THCA | 68.53 | 14.73 |
| GTEX-133LE-0326-SM-5P9G4 | Thyroid | TCGA-THCA | 45.39 | 12.7 |
| GTEX-1399R-0126-SM-5IFEV | Thyroid | TCGA-THCA | 60.45 | 15.3 |
| GTEX-1399T-0126-SM-5KM15 | Thyroid | TCGA-THCA | 49.93 | 10.71 |
| GTEX-1399U-0326-SM-5P9G5 | Thyroid | TCGA-THCA | 65.27 | 13.47 |
| GTEX-139T6-0326-SM-5J2LY | Thyroid | TCGA-THCA | 66.31 | 8.91 |
| GTEX-139TS-0126-SM-5K7XJ | Thyroid | TCGA-THCA | 67.34 | 15.65 |
| GTEX-139UW-0126-SM-5KM1B | Thyroid | TCGA-THCA | 50.61 | 9.86 |
| GTEX-139YR-1226-SM-5IFEU | Thyroid | TCGA-THCA | 67.73 | 13.93 |
| GTEX-13CF3-0926-SM-5LZZC | Thyroid | TCGA-THCA | 69.35 | 11.76 |
| GTEX-13D11-0226-SM-5LZXL | Thyroid | TCGA-THCA | 64.11 | 10.46 |
| GTEX-13FH7-0126-SM-5KLZ1 | Thyroid | TCGA-THCA | 63.56 | 15.18 |

| | | | | |
|---|---|---|---|---|
| GTEX-13FHO-0926-SM-5N9EW | Thyroid | TCGA-THCA | 50.77 | 12.73 |
| GTEX-13FHP-0926-SM-5L3EC | Thyroid | TCGA-THCA | 58.27 | 17.34 |
| GTEX-13FLV-0226-SM-5J2OF | Thyroid | TCGA-THCA | 52.98 | 12.46 |
| GTEX-13FLW-0326-SM-5J2M4 | Thyroid | TCGA-THCA | 73.05 | 13.93 |
| GTEX-13FTW-0626-SM-5IFEX | Thyroid | TCGA-THCA | 73.44 | 16.81 |
| GTEX-13FTY-0726-SM-5J2OH | Thyroid | TCGA-THCA | 67 | 14.78 |
| GTEX-13FXS-0726-SM-5LZXJ | Thyroid | TCGA-THCA | 59.68 | 11.57 |
| GTEX-13G51-1226-SM-5K7Z3 | Thyroid | TCGA-THCA | 57.14 | 12.31 |
| GTEX-13IVO-0926-SM-5KLZP | Thyroid | TCGA-THCA | 103.38 | 14.07 |
| GTEX-13JVG-0926-SM-5IJE1 | Thyroid | TCGA-THCA | 44.65 | 12.82 |
| GTEX-13N11-1026-SM-5K7XQ | Thyroid | TCGA-THCA | 72.92 | 14.44 |
| GTEX-13N1W-0826-SM-5MR5J | Thyroid | TCGA-THCA | 44.73 | 7.25 |
| GTEX-13N2G-0726-SM-5MR38 | Thyroid | TCGA-THCA | 63.93 | 10.92 |
| GTEX-13NYB-0726-SM-5MR4J | Thyroid | TCGA-THCA | 70.91 | 17.21 |
| GTEX-13NYC-2426-SM-5MR3K | Thyroid | TCGA-THCA | 55.07 | 14.95 |
| GTEX-13NZ8-0226-SM-5J2OK | Thyroid | TCGA-THCA | 71.87 | 11.75 |
| GTEX-13NZ9-1126-SM-5MR37 | Thyroid | TCGA-THCA | 53.96 | 9.58 |
| GTEX-13NZA-1026-SM-5MR48 | Thyroid | TCGA-THCA | 48.31 | 13.68 |
| GTEX-13O1R-0826-SM-5J2MB | Thyroid | TCGA-THCA | 98.15 | 13.46 |
| GTEX-13O21-2226-SM-5MR3L | Thyroid | TCGA-THCA | 52.03 | 11.65 |
| GTEX-13O3O-0926-SM-5KM1F | Thyroid | TCGA-THCA | 73.11 | 13.73 |
| GTEX-13O3P-0726-SM-5J2OM | Thyroid | TCGA-THCA | 47.14 | 12.34 |
| GTEX-13O3Q-0626-SM-5IJG1 | Thyroid | TCGA-THCA | 66.7 | 16.24 |
| GTEX-13O61-0226-SM-5KM52 | Thyroid | TCGA-THCA | 61.63 | 16.34 |
| GTEX-13OVG-0226-SM-5LU93 | Thyroid | TCGA-THCA | 57.21 | 11.21 |
| GTEX-13OVI-0826-SM-5KLZ8 | Thyroid | TCGA-THCA | 42 | 7.16 |
| GTEX-13OVJ-0626-SM-5J2O2 | Thyroid | TCGA-THCA | 66.29 | 7.58 |
| GTEX-13OVK-0226-SM-6M472 | Thyroid | TCGA-THCA | 69.92 | 11.98 |
| GTEX-13OW5-0626-SM-5J2N2 | Thyroid | TCGA-THCA | 57.33 | 9.37 |
| GTEX-13OW6-0726-SM-5L3FX | Thyroid | TCGA-THCA | 65.78 | 14.34 |
| GTEX-13OW7-0826-SM-5L3EL | Thyroid | TCGA-THCA | 50.33 | 12.78 |

| | | | | |
|---|---|---|---|---|
| GTEX-13OW8-0126-SM-5IJE5 | Thyroid | TCGA-THCA | 54.47 | 9.48 |
| GTEX-13PDP-1026-SM-5L3FA | Thyroid | TCGA-THCA | 80.63 | 12.41 |
| GTEX-13PL6-1026-SM-5L3E5 | Thyroid | TCGA-THCA | 68.83 | 16.84 |
| GTEX-13PVQ-0726-SM-5L3GI | Thyroid | TCGA-THCA | 61.22 | 11.51 |
| GTEX-13PVR-0626-SM-5S2RC | Thyroid | TCGA-THCA | 56.69 | 9.84 |
| GTEX-13QBU-0626-SM-5J2OG | Thyroid | TCGA-THCA | 62.83 | 7.68 |
| GTEX-13QJ3-0926-SM-73KX5 | Thyroid | TCGA-THCA | 49.87 | 9.82 |
| GTEX-13QJC-0826-SM-5RQKC | Thyroid | TCGA-THCA | 42.45 | 7.59 |
| GTEX-13RTJ-0326-SM-5YYAE | Thyroid | TCGA-THCA | 49.95 | 9.23 |
| GTEX-13RTK-0326-SM-5RQHS | Thyroid | TCGA-THCA | 60.28 | 10.42 |
| GTEX-13S86-1126-SM-5RQJX | Thyroid | TCGA-THCA | 51.67 | 10.78 |
| GTEX-13U4I-0526-SM-5LU59 | Thyroid | TCGA-THCA | 54.5 | 11.44 |
| GTEX-13VXT-0626-SM-5SIA1 | Thyroid | TCGA-THCA | 69.05 | 14.09 |
| GTEX-13VXU-0826-SM-5KLZ2 | Thyroid | TCGA-THCA | 70.13 | 17.71 |
| GTEX-13W46-0926-SM-5LU3T | Thyroid | TCGA-THCA | 53.17 | 9.46 |
| GTEX-13X6H-0526-SM-5LU4Q | Thyroid | TCGA-THCA | 52.66 | 9.96 |
| GTEX-13X6J-0826-SM-5LU32 | Thyroid | TCGA-THCA | 62.84 | 15.68 |
| GTEX-13YAN-0926-SM-5O9C3 | Thyroid | TCGA-THCA | 51.96 | 11.82 |
| GTEX-144GL-1226-SM-5O9A4 | Thyroid | TCGA-THCA | 52.74 | 11.33 |
| GTEX-144GM-0226-SM-5Q5CB | Thyroid | TCGA-THCA | 66.75 | 10.53 |
| GTEX-144GO-0126-SM-5LUAO | Thyroid | TCGA-THCA | 63.13 | 10.33 |
| GTEX-145LT-0226-SM-5S2QK | Thyroid | TCGA-THCA | 42.83 | 8.04 |
| GTEX-145LU-0426-SM-5O9AH | Thyroid | TCGA-THCA | 51.6 | 14.16 |
| GTEX-145ME-0126-SM-5S2QM | Thyroid | TCGA-THCA | 50.08 | 10.72 |
| GTEX-145MG-0826-SM-5Q5C2 | Thyroid | TCGA-THCA | 73.15 | 14.2 |
| GTEX-145MH-0426-SM-5LU8T | Thyroid | TCGA-THCA | 63.44 | 13.69 |
| GTEX-145MI-1126-SM-5O9AK | Thyroid | TCGA-THCA | 91.15 | 16.47 |
| GTEX-146FQ-0726-SM-5LUA7 | Thyroid | TCGA-THCA | 54.23 | 16.44 |
| GTEX-146FR-0326-SM-5SI8U | Thyroid | TCGA-THCA | 60.84 | 8.48 |
| GTEX-14753-0926-SM-5Q5BI | Thyroid | TCGA-THCA | 39.9 | 13.54 |
| GTEX-1477Z-0226-SM-5TDCI | Thyroid | TCGA-THCA | 45.14 | 13.7 |

| | | | | |
|---|---|---|---|---|
| GTEX-147F4-0826-SM-5QGRB | Thyroid | TCGA-THCA | 53.7 | 12.49 |
| GTEX-147GR-0726-SM-5S2PL | Thyroid | TCGA-THCA | 59.15 | 14.25 |
| GTEX-148VI-0526-SM-5TDDG | Thyroid | TCGA-THCA | 56.27 | 11.21 |
| GTEX-148VJ-0726-SM-5LU8J | Thyroid | TCGA-THCA | 58.96 | 11.23 |
| GTEX-1497J-0126-SM-5Q5BK | Thyroid | TCGA-THCA | 56.8 | 10.65 |
| GTEX-14A5H-0726-SM-5Q5DW | Thyroid | TCGA-THCA | 73.75 | 14.24 |
| GTEX-14A6H-2426-SM-5Q5BO | Thyroid | TCGA-THCA | 57.48 | 14.87 |
| GTEX-14ABY-0926-SM-5Q5DY | Thyroid | TCGA-THCA | 72.65 | 16.38 |
| GTEX-14AS3-0226-SM-5Q5B6 | Thyroid | TCGA-THCA | 52.82 | 11.71 |
| GTEX-14ASI-0726-SM-5Q5DC | Thyroid | TCGA-THCA | 74.61 | 19.58 |
| GTEX-14B4R-0126-SM-5TDE4 | Thyroid | TCGA-THCA | 68.28 | 14.9 |
| GTEX-14BIN-0126-SM-5TDCG | Thyroid | TCGA-THCA | 47.66 | 13.32 |
| GTEX-14BMU-0226-SM-5S2QA | Thyroid | TCGA-THCA | 62.15 | 10.04 |
| GTEX-14BMV-0726-SM-73KVE | Thyroid | TCGA-THCA | 47.42 | 13.14 |
| GTEX-14C38-0826-SM-5S2U8 | Thyroid | TCGA-THCA | 59.18 | 14.17 |
| GTEX-14C39-0226-SM-5TDDW | Thyroid | TCGA-THCA | 80.93 | 16.81 |
| GTEX-14C5O-0826-SM-5TDEG | Thyroid | TCGA-THCA | 40.51 | 7.68 |
| GTEX-14DAQ-0826-SM-73KWT | Thyroid | TCGA-THCA | 55.97 | 12.71 |
| GTEX-14E6C-2626-SM-5RQJP | Thyroid | TCGA-THCA | 69.05 | 12.04 |
| GTEX-14E6E-0326-SM-73KY6 | Thyroid | TCGA-THCA | 54.73 | 12.57 |
| GTEX-14E7W-0926-SM-5YYA4 | Thyroid | TCGA-THCA | 60.53 | 13.1 |
| GTEX-14ICK-1626-SM-6ETZX | Thyroid | TCGA-THCA | 40.69 | 11.84 |
| GTEX-14ICL-0426-SM-5RQJ3 | Thyroid | TCGA-THCA | 59.86 | 8.94 |
| GTEX-14JG6-1326-SM-9OSXU | Thyroid | TCGA-THCA | 69.43 | 15.03 |
| GTEX-14JIY-1226-SM-6871R | Thyroid | TCGA-THCA | 56.96 | 12.64 |
| GTEX-14PHW-2926-SM-6AJBA | Thyroid | TCGA-THCA | 48.32 | 9 |
| GTEX-14PII-0826-SM-6871S | Thyroid | TCGA-THCA | 51.39 | 14.81 |
| GTEX-14PJ3-0126-SM-69LQP | Thyroid | TCGA-THCA | 57.96 | 10.33 |
| GTEX-14PJ4-0326-SM-664OT | Thyroid | TCGA-THCA | 64.94 | 12.15 |
| GTEX-14PJ6-0326-SM-6871H | Thyroid | TCGA-THCA | 40.57 | 11.31 |
| GTEX-14PJM-1326-SM-664NX | Thyroid | TCGA-THCA | 44.06 | 10.13 |

| | | | | |
|---|---|---|---|---|
| GTEX-14PJO-0626-SM-6LLHH | Thyroid | TCGA-THCA | 56.33 | 14.49 |
| GTEX-14PK6-0426-SM-6EU1J | Thyroid | TCGA-THCA | 68.83 | 9.77 |
| GTEX-14PKU-0326-SM-6AJA7 | Thyroid | TCGA-THCA | 80.25 | 10.91 |
| GTEX-14PKV-0626-SM-6AJA2 | Thyroid | TCGA-THCA | 61.58 | 7.87 |
| GTEX-14PN3-0826-SM-69LOS | Thyroid | TCGA-THCA | 67.45 | 12.2 |
| GTEX-14PN4-1526-SM-6871V | Thyroid | TCGA-THCA | 58.77 | 12.94 |
| GTEX-14PQA-1226-SM-6M47A | Thyroid | TCGA-THCA | 50.21 | 10.86 |
| GTEX-14XAO-0426-SM-6AJB6 | Thyroid | TCGA-THCA | 62.12 | 17.34 |
| GTEX-15CHC-0126-SM-5YYBA | Thyroid | TCGA-THCA | 55.65 | 10.55 |
| GTEX-15CHQ-0826-SM-69LOT | Thyroid | TCGA-THCA | 70.7 | 11.97 |
| GTEX-15CHR-1726-SM-7DUGW | Thyroid | TCGA-THCA | 53.05 | 11.6 |
| GTEX-15D1Q-0626-SM-6AJAZ | Thyroid | TCGA-THCA | 56.44 | 11.55 |
| GTEX-15DDE-0626-SM-69LOK | Thyroid | TCGA-THCA | 94.94 | 13.49 |
| GTEX-15DZA-0226-SM-7KFS6 | Thyroid | TCGA-THCA | 65.38 | 12.82 |
| GTEX-15EO6-0126-SM-6LPKJ | Thyroid | TCGA-THCA | 65.76 | 11.73 |
| GTEX-15ER7-0726-SM-7KUMF | Thyroid | TCGA-THCA | 45.11 | 7.63 |
| GTEX-15ETS-0526-SM-6PAN3 | Thyroid | TCGA-THCA | 60.08 | 9.7 |
| GTEX-15EU6-1426-SM-6M48E | Thyroid | TCGA-THCA | 63.81 | 11.9 |
| GTEX-15FZZ-0226-SM-6LLI4 | Thyroid | TCGA-THCA | 37.21 | 9.88 |
| GTEX-15G19-0626-SM-6M474 | Thyroid | TCGA-THCA | 61.87 | 12.94 |
| GTEX-15G1A-0326-SM-6M467 | Thyroid | TCGA-THCA | 61.69 | 9.85 |
| GTEX-15RIE-0426-SM-7KUMH | Thyroid | TCGA-THCA | 53.71 | 11.19 |
| GTEX-15RIF-0326-SM-7KUFR | Thyroid | TCGA-THCA | 78.42 | 18.05 |
| GTEX-15RJ7-0326-SM-6M47H | Thyroid | TCGA-THCA | 55.7 | 17.12 |
| GTEX-15RJE-1326-SM-6LPI6 | Thyroid | TCGA-THCA | 37.46 | 7.79 |
| GTEX-15SB6-1526-SM-7KUMQ | Thyroid | TCGA-THCA | 60.36 | 10.73 |
| GTEX-15SHU-0726-SM-7KUFI | Thyroid | TCGA-THCA | 65.36 | 11.74 |
| GTEX-15SHV-0426-SM-6M476 | Thyroid | TCGA-THCA | 71.4 | 15.6 |
| GTEX-15UF6-1126-SM-6LPJ3 | Thyroid | TCGA-THCA | 59.6 | 16.47 |
| GTEX-169BO-0326-SM-7EPIM | Thyroid | TCGA-THCA | 50.95 | 15.41 |
| GTEX-16AAH-0326-SM-7DHML | Thyroid | TCGA-THCA | 51.1 | 16.42 |

| | | | | |
|---|---|---|---|---|
| GTEX-16BQI-0726-SM-6LPJZ | Thyroid | TCGA-THCA | 61.63 | 13.07 |
| GTEX-16GPK-0926-SM-6LPJ9 | Thyroid | TCGA-THCA | 65.94 | 14.57 |
| GTEX-16MT8-0626-SM-6M47Q | Thyroid | TCGA-THCA | 34.26 | 10.34 |
| GTEX-16MTA-0726-SM-7KUL4 | Thyroid | TCGA-THCA | 50.24 | 7.5 |
| GTEX-16NGA-0326-SM-718AJ | Thyroid | TCGA-THCA | 55.37 | 11.59 |
| GTEX-16NPX-1426-SM-6LPK3 | Thyroid | TCGA-THCA | 52.8 | 11.72 |
| GTEX-16XZY-0726-SM-79OMU | Thyroid | TCGA-THCA | 55.3 | 13.69 |
| GTEX-16XZZ-0826-SM-7IGM3 | Thyroid | TCGA-THCA | 61.54 | 10.64 |
| GTEX-16YQH-0326-SM-6LPJV | Thyroid | TCGA-THCA | 43.94 | 7.73 |
| GTEX-16Z82-0426-SM-7EPGX | Thyroid | TCGA-THCA | 49.76 | 9.72 |
| GTEX-178AV-0726-SM-6LPJI | Thyroid | TCGA-THCA | 51.91 | 13.24 |
| GTEX-17EVP-0126-SM-7EPHW | Thyroid | TCGA-THCA | 51.79 | 10.06 |
| GTEX-17EVQ-0526-SM-7KFSK | Thyroid | TCGA-THCA | 35.07 | 13.86 |
| GTEX-17F96-0526-SM-79OLE | Thyroid | TCGA-THCA | 45.4 | 9.28 |
| GTEX-17F97-0626-SM-7IGOH | Thyroid | TCGA-THCA | 58.3 | 12 |
| GTEX-17F9E-0626-SM-79ON5 | Thyroid | TCGA-THCA | 50.01 | 10.4 |
| GTEX-17F9Y-0526-SM-7EWDC | Thyroid | TCGA-THCA | 49.26 | 7.31 |
| GTEX-17GQL-0226-SM-7LTAK | Thyroid | TCGA-THCA | 71.84 | 13.94 |
| GTEX-17HG3-0226-SM-7EWEA | Thyroid | TCGA-THCA | 45.04 | 11.1 |
| GTEX-17HGU-0826-SM-7EWE5 | Thyroid | TCGA-THCA | 48.22 | 11.14 |
| GTEX-17HHE-0426-SM-79OK3 | Thyroid | TCGA-THCA | 45.19 | 12.81 |
| GTEX-17HHY-0826-SM-7EPID | Thyroid | TCGA-THCA | 57.75 | 10.25 |
| GTEX-17HII-1926-SM-79OLB | Thyroid | TCGA-THCA | 54.14 | 11.76 |
| GTEX-17JCI-0626-SM-7IGM7 | Thyroid | TCGA-THCA | 53.53 | 13.58 |
| GTEX-17KNJ-1026-SM-79ONK | Thyroid | TCGA-THCA | 54.78 | 11.4 |
| GTEX-17MF6-0626-SM-7LT8D | Thyroid | TCGA-THCA | 44.46 | 8.39 |
| GTEX-183FY-0626-SM-79OKR | Thyroid | TCGA-THCA | 49.38 | 10.76 |
| GTEX-183WM-2626-SM-7KFRY | Thyroid | TCGA-THCA | 58.84 | 13.06 |
| GTEX-18465-1426-SM-7KFTF | Thyroid | TCGA-THCA | 62.83 | 14.88 |
| GTEX-18A66-0826-SM-72D5Z | Thyroid | TCGA-THCA | 64.32 | 14.65 |
| GTEX-18A67-0826-SM-7KFTI | Thyroid | TCGA-THCA | 59.95 | 12.92 |

| | | | | |
|---|---|---|---|---|
| GTEX-18A6Q-0726-SM-7LT8Y | Thyroid | TCGA-THCA | 48.28 | 9.3 |
| GTEX-18A7A-0826-SM-7KFTJ | Thyroid | TCGA-THCA | 61.82 | 10.89 |
| GTEX-18D9A-0126-SM-7KFSI | Thyroid | TCGA-THCA | 69.19 | 13.69 |
| GTEX-18D9B-0726-SM-72D6K | Thyroid | TCGA-THCA | 54.1 | 15.66 |
| GTEX-18D9U-1026-SM-72D5R | Thyroid | TCGA-THCA | 60.87 | 12.21 |
| GTEX-18QFQ-0726-SM-7LG6D | Thyroid | TCGA-THCA | 52.96 | 6.88 |
| GTEX-1A3MV-0326-SM-73KW7 | Thyroid | TCGA-THCA | 65.22 | 12.88 |
| GTEX-1A8FM-0726-SM-7DUGK | Thyroid | TCGA-THCA | 53.76 | 13.13 |
| GTEX-1A8G6-0626-SM-7IGNB | Thyroid | TCGA-THCA | 47.26 | 11.73 |
| GTEX-1A8G7-1026-SM-73KVA | Thyroid | TCGA-THCA | 52.99 | 11.99 |
| GTEX-1AMEY-0126-SM-73KTX | Thyroid | TCGA-THCA | 36.93 | 8.41 |
| GTEX-1AMFI-0526-SM-7189M | Thyroid | TCGA-THCA | 54.3 | 15.95 |
| GTEX-1AX8Z-0826-SM-7DUFZ | Thyroid | TCGA-THCA | 51.85 | 15.52 |
| GTEX-1AX9I-0626-SM-72D54 | Thyroid | TCGA-THCA | 61.73 | 10.13 |
| GTEX-1AX9J-2126-SM-731DD | Thyroid | TCGA-THCA | 47.93 | 9.61 |
| GTEX-1AX9K-0626-SM-73KVD | Thyroid | TCGA-THCA | 53.63 | 15.92 |
| GTEX-1AYCT-0226-SM-73KVB | Thyroid | TCGA-THCA | 64.9 | 13.15 |
| GTEX-1B8KE-0626-SM-7189H | Thyroid | TCGA-THCA | 73.48 | 9.95 |
| GTEX-1B8KZ-0426-SM-731DP | Thyroid | TCGA-THCA | 72.92 | 16.96 |
| GTEX-1B8L1-1626-SM-7IGMH | Thyroid | TCGA-THCA | 53.31 | 8.24 |
| GTEX-1B8SF-0626-SM-73KVV | Thyroid | TCGA-THCA | 48.91 | 17.66 |
| GTEX-1B8SG-1126-SM-7IGMT | Thyroid | TCGA-THCA | 108.98 | 11.42 |
| GTEX-1B932-0926-SM-73KUP | Thyroid | TCGA-THCA | 63.64 | 16.01 |
| GTEX-1B933-1026-SM-7P8RO | Thyroid | TCGA-THCA | 67.16 | 15.71 |
| GTEX-1B97I-0326-SM-7DUGB | Thyroid | TCGA-THCA | 54.18 | 17.43 |
| GTEX-1B97J-0626-SM-7P8RU | Thyroid | TCGA-THCA | 64.56 | 10.42 |
| GTEX-1BAJH-0926-SM-79OO6 | Thyroid | TCGA-THCA | 48.46 | 10.04 |
| GTEX-1C2JI-0326-SM-7EWFD | Thyroid | TCGA-THCA | 49.11 | 7.68 |
| GTEX-1C475-0226-SM-7MGXM | Thyroid | TCGA-THCA | 44.96 | 7.97 |
| GTEX-1C4CL-0726-SM-7IGP9 | Thyroid | TCGA-THCA | 48.52 | 12.09 |
| GTEX-1C64N-1026-SM-79ONM | Thyroid | TCGA-THCA | 52.53 | 15.25 |

| | | | | |
|---|---|---|---|---|
| GTEX-1C6VQ-0526-SM-7MGWD | Thyroid | TCGA-THCA | 46.37 | 8.13 |
| GTEX-1C6VR-0426-SM-7IGN6 | Thyroid | TCGA-THCA | 50.7 | 12.12 |
| GTEX-1C6VS-0826-SM-7EWEI | Thyroid | TCGA-THCA | 64.67 | 10.33 |
| GTEX-1CAMQ-1126-SM-7EWFE | Thyroid | TCGA-THCA | 41.65 | 8.42 |
| GTEX-1CAMR-0226-SM-7DUGO | Thyroid | TCGA-THCA | 45.64 | 10.05 |
| GTEX-1CAMS-0826-SM-7P8P1 | Thyroid | TCGA-THCA | 48.27 | 7.8 |
| GTEX-1CB4E-0326-SM-7MXT3 | Thyroid | TCGA-THCA | 49.28 | 9.54 |
| GTEX-1CB4F-0826-SM-793CV | Thyroid | TCGA-THCA | 60.35 | 16.56 |
| GTEX-1CB4G-1526-SM-7P8QG | Thyroid | TCGA-THCA | 48.23 | 11.65 |
| GTEX-1CB4H-0126-SM-7IGN2 | Thyroid | TCGA-THCA | 51.46 | 11.43 |
| GTEX-1CB4I-0726-SM-7DUGS | Thyroid | TCGA-THCA | 63.33 | 13.49 |
| GTEX-1CB4J-1426-SM-7MKFR | Thyroid | TCGA-THCA | 61.29 | 11.3 |
| GTEX-1E1VI-0726-SM-7P8QJ | Thyroid | TCGA-THCA | 40.67 | 9.07 |
| GTEX-1EH9U-0926-SM-7EWF1 | Thyroid | TCGA-THCA | 38.83 | 8.76 |
| GTEX-1EKGG-0726-SM-7IGPX | Thyroid | TCGA-THCA | 39.72 | 11.05 |
| GTEX-1EMGI-0826-SM-7EPHY | Thyroid | TCGA-THCA | 55.58 | 10.25 |
| GTEX-1EN7A-1026-SM-7IGPZ | Thyroid | TCGA-THCA | 44.72 | 11.14 |
| GTEX-1EU9M-0626-SM-7MXUT | Thyroid | TCGA-THCA | 51.48 | 8.98 |
| GTEX-1EWIQ-0826-SM-7MXTH | Thyroid | TCGA-THCA | 59.42 | 17.29 |
| GTEX-1EX96-0126-SM-9KNVH | Thyroid | TCGA-THCA | 61.22 | 13.75 |
| GTEX-1F48J-0926-SM-7P8PZ | Thyroid | TCGA-THCA | 43 | 10.11 |
| GTEX-1F52S-0526-SM-7P8TP | Thyroid | TCGA-THCA | 63.15 | 13.06 |
| GTEX-1F5PL-0826-SM-7MXU7 | Thyroid | TCGA-THCA | 75.24 | 20.86 |
| GTEX-1F6IF-1126-SM-9KNV2 | Thyroid | TCGA-THCA | 58.22 | 12.8 |
| GTEX-1F75A-0326-SM-7MXTR | Thyroid | TCGA-THCA | 62.4 | 14.6 |
| GTEX-1F75B-0826-SM-9QEIT | Thyroid | TCGA-THCA | 54.09 | 12.84 |
| GTEX-1F75I-0926-SM-7MXUU | Thyroid | TCGA-THCA | 55.52 | 10.45 |
| GTEX-1F88F-0726-SM-7P8R7 | Thyroid | TCGA-THCA | 36.9 | 10.02 |
| GTEX-1FIGZ-0226-SM-7P8QR | Thyroid | TCGA-THCA | 58.21 | 14.73 |
| GTEX-1GF9U-0226-SM-7P8TY | Thyroid | TCGA-THCA | 58.76 | 13.82 |
| GTEX-1GF9V-0926-SM-9KNW7 | Thyroid | TCGA-THCA | 56.12 | 12.79 |

| | | | | |
|---|---|---|---|---|
| GTEX-1GF9W-0626-SM-7MXVJ | Thyroid | TCGA-THCA | 84.6 | 13.98 |
| GTEX-1GF9X-0126-SM-A9G38 | Thyroid | TCGA-THCA | 62.9 | 11.8 |
| GTEX-1GL5R-0626-SM-7PBZY | Thyroid | TCGA-THCA | 58.45 | 10.22 |
| GTEX-1GMR3-0726-SM-9WPOF | Thyroid | TCGA-THCA | 42.85 | 11.95 |
| GTEX-1GMR8-0826-SM-9KNUR | Thyroid | TCGA-THCA | 64.35 | 12.71 |
| GTEX-1GMRU-1126-SM-9KNV1 | Thyroid | TCGA-THCA | 47.76 | 12.02 |
| GTEX-1GN1U-0826-SM-9WPPX | Thyroid | TCGA-THCA | 46.5 | 12.83 |
| GTEX-1GN1V-0626-SM-7P8QU | Thyroid | TCGA-THCA | 70.81 | 16.03 |
| GTEX-1GN1W-0126-SM-7MXV7 | Thyroid | TCGA-THCA | 55.57 | 11.79 |
| GTEX-1GN2E-0826-SM-9WYT8 | Thyroid | TCGA-THCA | 43.27 | 9.61 |
| GTEX-1GN73-0826-SM-9OSW4 | Thyroid | TCGA-THCA | 64.38 | 11.6 |
| GTEX-1GPI7-0726-SM-9WG84 | Thyroid | TCGA-THCA | 68.67 | 14.59 |
| GTEX-1GTWX-0726-SM-9WPPA | Thyroid | TCGA-THCA | 45.65 | 11.81 |
| GTEX-1GZ2Q-0826-SM-7P8TK | Thyroid | TCGA-THCA | 46.65 | 7.65 |
| GTEX-1GZ4H-0226-SM-9JGGJ | Thyroid | TCGA-THCA | 27.76 | 4.95 |
| GTEX-1GZ4I-0626-SM-9WPPN | Thyroid | TCGA-THCA | 54.61 | 10.48 |
| GTEX-1GZHY-1426-SM-9WG5H | Thyroid | TCGA-THCA | 47.64 | 13.77 |
| GTEX-1H11D-0626-SM-9WPQ1 | Thyroid | TCGA-THCA | 54.25 | 15.13 |
| GTEX-1H1CY-0726-SM-9OSXX | Thyroid | TCGA-THCA | 40.24 | 11.98 |
| GTEX-1H1DE-0226-SM-ARZN8 | Thyroid | TCGA-THCA | 43.79 | 12.84 |
| GTEX-1H1DF-1026-SM-9QEJ1 | Thyroid | TCGA-THCA | 69.09 | 17.78 |
| GTEX-1H1DG-0726-SM-A96TN | Thyroid | TCGA-THCA | 72.69 | 20.17 |
| GTEX-1H1E6-0326-SM-9OSWE | Thyroid | TCGA-THCA | 61.06 | 13.34 |
| GTEX-1H1ZS-0726-SM-A9G3K | Thyroid | TCGA-THCA | 39.89 | 11.52 |
| GTEX-1H2FU-0626-SM-9WPPS | Thyroid | TCGA-THCA | 52.31 | 9.36 |
| GTEX-1H3VE-0826-SM-9KNVQ | Thyroid | TCGA-THCA | 48.66 | 8.34 |
| GTEX-1HB9E-0826-SM-CNNQJ | Thyroid | TCGA-THCA | 54.77 | 8.7 |
| GTEX-1HBPH-0926-SM-A9SME | Thyroid | TCGA-THCA | 51.65 | 11.98 |
| GTEX-1HBPI-0826-SM-9WYTX | Thyroid | TCGA-THCA | 59.36 | 18.49 |
| GTEX-1HBPM-0926-SM-ARL9C | Thyroid | TCGA-THCA | 50.26 | 9.03 |
| GTEX-1HBPN-1226-SM-B2LW9 | Thyroid | TCGA-THCA | 40.79 | 8.85 |

| | | | | |
|---|---|---|---|---|
| GTEX-1HC8U-1226-SM-9WYUW | Thyroid | TCGA-THCA | 39.48 | 13.27 |
| GTEX-1HCU7-0826-SM-B2LX7 | Thyroid | TCGA-THCA | 60.69 | 14.24 |
| GTEX-1HCU9-0226-SM-9WPNT | Thyroid | TCGA-THCA | 61.29 | 11.71 |
| GTEX-1HCUA-0126-SM-B2LWC | Thyroid | TCGA-THCA | 53.69 | 9.69 |
| GTEX-1HCVE-0726-SM-A96SH | Thyroid | TCGA-THCA | 45.93 | 10.49 |
| GTEX-1HFI6-1326-SM-A9SL8 | Thyroid | TCGA-THCA | 56.71 | 11.33 |
| GTEX-1HFI7-0326-SM-9WPOX | Thyroid | TCGA-THCA | 50.2 | 14.65 |
| GTEX-1HGF4-0726-SM-9WYU9 | Thyroid | TCGA-THCA | 37.48 | 12.79 |
| GTEX-1HKZK-0826-SM-A9G2C | Thyroid | TCGA-THCA | 61.54 | 13 |
| GTEX-1HR98-0326-SM-CGQEL | Thyroid | TCGA-THCA | 69.19 | 15.78 |
| GTEX-1HR9M-0626-SM-A96S6 | Thyroid | TCGA-THCA | 55.73 | 12.93 |
| GTEX-1HSEH-0826-SM-B2LXE | Thyroid | TCGA-THCA | 51.47 | 13.43 |
| GTEX-1HSGN-0726-SM-A9G2F | Thyroid | TCGA-THCA | 53.74 | 14.72 |
| GTEX-1HSKV-1026-SM-B2LWK | Thyroid | TCGA-THCA | 58.73 | 13.99 |
| GTEX-1HSMO-0726-SM-A96TM | Thyroid | TCGA-THCA | 66.63 | 12.56 |
| GTEX-1HSMQ-0526-SM-A96TY | Thyroid | TCGA-THCA | 68.47 | 18.38 |
| GTEX-1HT8W-1526-SM-CE6S3 | Thyroid | TCGA-THCA | 54.25 | 11.58 |
| GTEX-1HUB1-1226-SM-B2LWX | Thyroid | TCGA-THCA | 50.49 | 12.27 |
| GTEX-1I19N-0226-SM-CE6ST | Thyroid | TCGA-THCA | 43.82 | 12.07 |
| GTEX-1I1GP-0326-SM-CNPQ4 | Thyroid | TCGA-THCA | 105.03 | 15.59 |
| GTEX-1I1GQ-0826-SM-CNNQR | Thyroid | TCGA-THCA | 44.9 | 11.85 |
| GTEX-1I1GR-1126-SM-A9G2I | Thyroid | TCGA-THCA | 48.26 | 11.53 |
| GTEX-1I1GS-0726-SM-C1YS7 | Thyroid | TCGA-THCA | 55.76 | 12.44 |
| GTEX-1I1GT-0826-SM-B2LWO | Thyroid | TCGA-THCA | 42.96 | 8.94 |
| GTEX-1I1GU-0826-SM-ARZM5 | Thyroid | TCGA-THCA | 57.89 | 14.83 |
| GTEX-1I1GV-0726-SM-ARZLY | Thyroid | TCGA-THCA | 71.09 | 15.66 |
| GTEX-1I6K6-0726-SM-B2LX4 | Thyroid | TCGA-THCA | 42.18 | 9.51 |
| GTEX-1I6K7-1126-SM-A96SW | Thyroid | TCGA-THCA | 55.94 | 12.99 |
| GTEX-1ICG6-0426-SM-ARU7M | Thyroid | TCGA-THCA | 54.63 | 12 |
| GTEX-1ICLY-0726-SM-A96TB | Thyroid | TCGA-THCA | 46.75 | 11.2 |
| GTEX-1ICLZ-0526-SM-ARZM8 | Thyroid | TCGA-THCA | 41.33 | 8.97 |

| | | | | |
|---|---|---|---|---|
| GTEX-1IDJC-1026-SM-A96T2 | Thyroid | TCGA-THCA | 52.19 | 8.98 |
| GTEX-1IDJH-0826-SM-CE6SY | Thyroid | TCGA-THCA | 45.78 | 12.55 |
| GTEX-1IDJU-1626-SM-CJI4B | Thyroid | TCGA-THCA | 55.08 | 8.94 |
| GTEX-1IE54-0926-SM-A9G2Y | Thyroid | TCGA-THCA | 52.63 | 9.27 |
| GTEX-1IGQW-0626-SM-C1YR9 | Thyroid | TCGA-THCA | 70.75 | 15.28 |
| GTEX-1IKJJ-1026-SM-A96SQ | Thyroid | TCGA-THCA | 50.64 | 8.31 |
| GTEX-1IKOE-0926-SM-ARU7X | Thyroid | TCGA-THCA | 52.4 | 13.72 |
| GTEX-1IKOH-0626-SM-CJI35 | Thyroid | TCGA-THCA | 40.7 | 9.63 |
| GTEX-1IL2U-0826-SM-CL53Q | Thyroid | TCGA-THCA | 36.05 | 11.14 |
| GTEX-1IL2V-0626-SM-ARU7S | Thyroid | TCGA-THCA | 43.22 | 9.71 |
| GTEX-1IOXB-1026-SM-CJI4D | Thyroid | TCGA-THCA | 37.58 | 10.98 |
| GTEX-1IY9M-1026-SM-A96TG | Thyroid | TCGA-THCA | 46.65 | 12.05 |
| GTEX-1J1OQ-0526-SM-A9G2P | Thyroid | TCGA-THCA | 51.23 | 11.88 |
| GTEX-1J1R8-0126-SM-CJI36 | Thyroid | TCGA-THCA | 39.24 | 6.99 |
| GTEX-1J8EW-1126-SM-A96T8 | Thyroid | TCGA-THCA | 42.88 | 10.83 |
| GTEX-1J8JJ-0726-SM-AHZ3E | Thyroid | TCGA-THCA | 51.4 | 13.19 |
| GTEX-1J8Q3-0226-SM-C1YR4 | Thyroid | TCGA-THCA | 64.51 | 14.39 |
| GTEX-1J8QM-0726-SM-CE6RF | Thyroid | TCGA-THCA | 48.47 | 11.48 |
| GTEX-1JJ6O-0626-SM-AHZ3H | Thyroid | TCGA-THCA | 31.46 | 7.22 |
| GTEX-1JJE9-0226-SM-CY8HU | Thyroid | TCGA-THCA | 46.55 | 11.45 |
| GTEX-1JJEA-0626-SM-D3L96 | Thyroid | TCGA-THCA | 59.87 | 15.9 |
| GTEX-1JK1U-0326-SM-CXZJH | Thyroid | TCGA-THCA | 51.75 | 13.35 |
| GTEX-1JKYN-1126-SM-CGQG5 | Thyroid | TCGA-THCA | 60.14 | 9.65 |
| GTEX-1JKYR-0826-SM-ARU8L | Thyroid | TCGA-THCA | 54.1 | 11.52 |
| GTEX-1JMLX-0826-SM-CNNPP | Thyroid | TCGA-THCA | 54.68 | 11.07 |
| GTEX-1JMPY-0126-SM-CJI39 | Thyroid | TCGA-THCA | 45.21 | 8.41 |
| GTEX-1JMPZ-0626-SM-CY8HY | Thyroid | TCGA-THCA | 45.66 | 13.03 |
| GTEX-1JMQJ-2026-SM-CNPP9 | Thyroid | TCGA-THCA | 58.44 | 12.5 |
| GTEX-1JMQK-2126-SM-CY8I2 | Thyroid | TCGA-THCA | 46.67 | 13.43 |
| GTEX-1JMQL-0926-SM-CJI3F | Thyroid | TCGA-THCA | 41.52 | 11.86 |
| GTEX-1JN6P-0626-SM-ARZN6 | Thyroid | TCGA-THCA | 31.37 | 10.1 |

| | | | | |
|---|---|---|---|---|
| GTEX-1K2DA-1526-SM-CGQGQ | Thyroid | TCGA-THCA | 53.5 | 13.37 |
| GTEX-1K2DU-0126-SM-E9TIU | Thyroid | TCGA-THCA | 47.18 | 8.93 |
| GTEX-1K9T9-0826-SM-EAZ4X | Thyroid | TCGA-THCA | 52.49 | 16.81 |
| GTEX-1KAFJ-0426-SM-DHXJH | Thyroid | TCGA-THCA | 51.35 | 8.78 |
| GTEX-1KANC-0826-SM-D3LA8 | Thyroid | TCGA-THCA | 56.94 | 15.08 |
| GTEX-1KD4Q-1026-SM-CXZKJ | Thyroid | TCGA-THCA | 66.5 | 15.7 |
| GTEX-1KD5A-0426-SM-D4P3X | Thyroid | TCGA-THCA | 55.1 | 12.12 |
| GTEX-1KXAM-1726-SM-D3LAE | Thyroid | TCGA-THCA | 67.97 | 12.55 |
| GTEX-1LB8K-0626-SM-D4P3Z | Thyroid | TCGA-THCA | 49.99 | 11.72 |
| GTEX-1LBAC-0726-SM-D3L9U | Thyroid | TCGA-THCA | 54.66 | 13.82 |
| GTEX-1LG7Y-0626-SM-DHXJM | Thyroid | TCGA-THCA | 63.57 | 9.42 |
| GTEX-1LG7Z-0526-SM-D4P3Q | Thyroid | TCGA-THCA | 40.9 | 7.12 |
| GTEX-1LGOU-0126-SM-EV79W | Thyroid | TCGA-THCA | 70.17 | 12.87 |
| GTEX-1LGRB-1226-SM-CNNPS | Thyroid | TCGA-THCA | 50.92 | 9.13 |
| GTEX-1LH75-0626-SM-DIPG3 | Thyroid | TCGA-THCA | 53.6 | 10.05 |
| GTEX-1LSVX-0726-SM-E9TJG | Thyroid | TCGA-THCA | 49.01 | 11.26 |
| GTEX-1LVAM-0826-SM-EV79X | Thyroid | TCGA-THCA | 69.75 | 13.7 |
| GTEX-1LVAN-0726-SM-CNNPF | Thyroid | TCGA-THCA | 49 | 13.33 |
| GTEX-1M4P7-0626-SM-EVR51 | Thyroid | TCGA-THCA | 59.09 | 13.64 |
| GTEX-1MA7X-0926-SM-DIPEJ | Thyroid | TCGA-THCA | 50.72 | 11.8 |
| GTEX-1MCC2-0526-SM-DHXJS | Thyroid | TCGA-THCA | 60.46 | 11.98 |
| GTEX-1MGNQ-0726-SM-DPRYH | Thyroid | TCGA-THCA | 52.96 | 10.45 |
| GTEX-1MJK2-0826-SM-DTXE8 | Thyroid | TCGA-THCA | 66.9 | 13.19 |
| GTEX-1MJK3-0326-SM-DTX96 | Thyroid | TCGA-THCA | 60.26 | 9.75 |
| GTEX-1MUQO-0726-SM-E9TJM | Thyroid | TCGA-THCA | 44.07 | 11.18 |
| GTEX-1N5O9-1026-SM-DTX8Y | Thyroid | TCGA-THCA | 51.13 | 10.34 |
| GTEX-1NUQO-0926-SM-DTXEX | Thyroid | TCGA-THCA | 54.39 | 12.27 |
| GTEX-1NV5F-0726-SM-DTXEH | Thyroid | TCGA-THCA | 58.71 | 10.86 |
| GTEX-1O97I-1026-SM-E76PI | Thyroid | TCGA-THCA | 47.36 | 11.06 |
| GTEX-1O9I2-1226-SM-E6CQC | Thyroid | TCGA-THCA | 53.93 | 10.03 |
| GTEX-1OFPY-1526-SM-DTX8R | Thyroid | TCGA-THCA | 53.45 | 10.31 |

| | | | | |
|---|---|---|---|---|
| GTEX-1OJC3-0626-SM-E8VMO | Thyroid | TCGA-THCA | 48.66 | 7.61 |
| GTEX-1OJC4-0526-SM-E6CQV | Thyroid | TCGA-THCA | 59.26 | 12.31 |
| GTEX-1P4AB-1126-SM-DTX94 | Thyroid | TCGA-THCA | 56.75 | 8.89 |
| GTEX-1PBJI-0726-SM-EXUSQ | Thyroid | TCGA-THCA | 47.63 | 10.27 |
| GTEX-1PBJJ-1926-SM-DTX8T | Thyroid | TCGA-THCA | 48.75 | 12.44 |
| GTEX-1PDJ9-0826-SM-E76PS | Thyroid | TCGA-THCA | 58.39 | 11.36 |
| GTEX-1PFEY-0226-SM-EVR3M | Thyroid | TCGA-THCA | 72.69 | 12.02 |
| GTEX-1PIGE-0726-SM-DPRZD | Thyroid | TCGA-THCA | 68.65 | 15.4 |
| GTEX-1PIIG-2026-SM-EVR43 | Thyroid | TCGA-THCA | 43.39 | 9.47 |
| GTEX-1PPGY-1426-SM-EVR3R | Thyroid | TCGA-THCA | 69.7 | 17.59 |
| GTEX-1PPH8-0926-SM-DTX9U | Thyroid | TCGA-THCA | 50.96 | 7.73 |
| GTEX-1PWST-0726-SM-EWRMO | Thyroid | TCGA-THCA | 57.63 | 13 |
| GTEX-1QAET-0226-SM-EVR3T | Thyroid | TCGA-THCA | 47.24 | 14.99 |
| GTEX-1QCLY-0826-SM-DTX8A | Thyroid | TCGA-THCA | 41.05 | 7 |
| GTEX-1QCLZ-0626-SM-E6CR2 | Thyroid | TCGA-THCA | 57.12 | 12.92 |
| GTEX-1QEPI-0726-SM-E76QG | Thyroid | TCGA-THCA | 60.66 | 14.55 |
| GTEX-1QMI2-0826-SM-EXUT4 | Thyroid | TCGA-THCA | 44.1 | 12.03 |
| GTEX-1QP28-0626-SM-E76P5 | Thyroid | TCGA-THCA | 63.23 | 14.74 |
| GTEX-1QP2A-0626-SM-DTX9C | Thyroid | TCGA-THCA | 54.56 | 10.26 |
| GTEX-1QP66-0326-SM-DTX7V | Thyroid | TCGA-THCA | 60.61 | 12.59 |
| GTEX-1QP67-0326-SM-EVR45 | Thyroid | TCGA-THCA | 60.47 | 14.52 |
| GTEX-1QP6S-2526-SM-DTX9Z | Thyroid | TCGA-THCA | 46.62 | 12.23 |
| GTEX-1QP9N-0126-SM-DPRZS | Thyroid | TCGA-THCA | 67.98 | 10.51 |
| GTEX-1QPFJ-0626-SM-DTX7Y | Thyroid | TCGA-THCA | 56 | 7.77 |
| GTEX-1QW4Y-0226-SM-EAZ3U | Thyroid | TCGA-THCA | 46.68 | 14.91 |
| GTEX-1R46S-0426-SM-EVR48 | Thyroid | TCGA-THCA | 70.84 | 11.66 |
| GTEX-1R9JW-0226-SM-EVR4K | Thyroid | TCGA-THCA | 59.79 | 12.01 |
| GTEX-1R9K4-0726-SM-EWRN2 | Thyroid | TCGA-THCA | 61.01 | 11.83 |
| GTEX-1R9K5-0726-SM-EAZ4E | Thyroid | TCGA-THCA | 64.44 | 10.88 |
| GTEX-1R9PM-0226-SM-DPRY7 | Thyroid | TCGA-THCA | 64.97 | 11.53 |
| GTEX-1RAZA-0826-SM-EAZ4I | Thyroid | TCGA-THCA | 72.1 | 16.1 |

| | | | | |
|---|---|---|---|---|
| GTEX-1RAZQ-0926-SM-EAZ4Q | Thyroid | TCGA-THCA | 43.69 | 10.28 |
| GTEX-1RAZR-1126-SM-E6CRC | Thyroid | TCGA-THCA | 83.39 | 17.36 |
| GTEX-1RAZS-1026-SM-E9TID | Thyroid | TCGA-THCA | 49.8 | 11.32 |
| GTEX-1RDX4-0926-SM-E9U6D | Thyroid | TCGA-THCA | 49.37 | 18.76 |
| GTEX-1RMOY-0126-SM-EWRNN | Thyroid | TCGA-THCA | 55.99 | 11.42 |
| GTEX-1RNSC-1126-SM-E9TIP | Thyroid | TCGA-THCA | 36.78 | 12.94 |
| GTEX-1RQEC-1026-SM-EVR5B | Thyroid | TCGA-THCA | 64.68 | 18.49 |
| GTEX-1RQED-0926-SM-EWRNZ | Thyroid | TCGA-THCA | 58.46 | 11.07 |
| GTEX-1S5ZU-0726-SM-EWROH | Thyroid | TCGA-THCA | 54.09 | 8.04 |
| GTEX-1S83E-0726-SM-EV7B6 | Thyroid | TCGA-THCA | 62.38 | 10.48 |
| GTEX-N7MS-2326-SM-2HMLD | Thyroid | TCGA-THCA | 56.15 | 11.61 |
| GTEX-NFK9-0726-SM-2HMJW | Thyroid | TCGA-THCA | 43.65 | 8.7 |
| GTEX-O5YV-1526-SM-EV799 | Thyroid | TCGA-THCA | 62.81 | 14.75 |
| GTEX-OHPK-2626-SM-2HMK9 | Thyroid | TCGA-THCA | 49.16 | 10.23 |
| GTEX-OHPL-2626-SM-2HMJA | Thyroid | TCGA-THCA | 53.36 | 19.42 |
| GTEX-OHPM-2626-SM-33HC5 | Thyroid | TCGA-THCA | 51.45 | 13.94 |
| GTEX-OIZF-2626-SM-7P8QZ | Thyroid | TCGA-THCA | 61.16 | 11.7 |
| GTEX-OIZG-0226-SM-2TC5L | Thyroid | TCGA-THCA | 51.77 | 13.41 |
| GTEX-OIZI-0726-SM-2XCEI | Thyroid | TCGA-THCA | 66.13 | 15.2 |
| GTEX-OOBJ-2626-SM-2I3F6 | Thyroid | TCGA-THCA | 46.21 | 11.29 |
| GTEX-OXRK-0626-SM-2HMJ5 | Thyroid | TCGA-THCA | 52.17 | 9.69 |
| GTEX-OXRL-2626-SM-2I3F1 | Thyroid | TCGA-THCA | 49.59 | 12.68 |
| GTEX-OXRN-1526-SM-E6CHM | Thyroid | TCGA-THCA | 36.05 | 7.53 |
| GTEX-OXRO-1226-SM-48TDL | Thyroid | TCGA-THCA | 79.18 | 15.75 |
| GTEX-OXRP-0326-SM-33HBJ | Thyroid | TCGA-THCA | 59.13 | 12.14 |
| GTEX-P44G-0626-SM-2I5EO | Thyroid | TCGA-THCA | 54.13 | 11 |
| GTEX-P4PQ-2626-SM-33HC9 | Thyroid | TCGA-THCA | 53.09 | 14.63 |
| GTEX-P4QS-2626-SM-2I3EV | Thyroid | TCGA-THCA | 43.43 | 11.08 |
| GTEX-P4QT-2626-SM-2I3FM | Thyroid | TCGA-THCA | 47.8 | 9.4 |
| GTEX-P78B-0526-SM-2I5F7 | Thyroid | TCGA-THCA | 56.76 | 11.83 |
| GTEX-PLZ4-1226-SM-2I5FE | Thyroid | TCGA-THCA | 53.61 | 11.21 |

| | | | | |
|---|---|---|---|---|
| GTEX-POYW-0826-SM-2XCEM | Thyroid | TCGA-THCA | 79.94 | 21.11 |
| GTEX-PWCY-2326-SM-2I3EQ | Thyroid | TCGA-THCA | 48.84 | 11.54 |
| GTEX-PWN1-2626-SM-2I3FH | Thyroid | TCGA-THCA | 66.93 | 9.25 |
| GTEX-PX3G-2626-SM-2I3EG | Thyroid | TCGA-THCA | 59.67 | 14.66 |
| GTEX-Q2AG-0826-SM-2HMKF | Thyroid | TCGA-THCA | 64.55 | 12.38 |
| GTEX-Q2AH-0726-SM-2I3EA | Thyroid | TCGA-THCA | 65.86 | 12.35 |
| GTEX-Q2AI-0326-SM-2I3EK | Thyroid | TCGA-THCA | 54.1 | 16.3 |
| GTEX-Q734-0526-SM-2I3EH | Thyroid | TCGA-THCA | 62.75 | 10.85 |
| GTEX-QDT8-1226-SM-EVR35 | Thyroid | TCGA-THCA | 58.11 | 12.76 |
| GTEX-QDVJ-0226-SM-2I5FV | Thyroid | TCGA-THCA | 47.63 | 11.66 |
| GTEX-QDVN-0626-SM-2I3FP | Thyroid | TCGA-THCA | 49.47 | 9.88 |
| GTEX-QEG5-0826-SM-2I5GF | Thyroid | TCGA-THCA | 65.22 | 23.03 |
| GTEX-QEL4-0726-SM-3GIJ5 | Thyroid | TCGA-THCA | 52.36 | 14.86 |
| GTEX-QLQ7-0726-SM-2I5G2 | Thyroid | TCGA-THCA | 53.36 | 12.42 |
| GTEX-QV31-0726-SM-3GAEG | Thyroid | TCGA-THCA | 58.79 | 16.57 |
| GTEX-QV44-0826-SM-2S1RG | Thyroid | TCGA-THCA | 61.77 | 14 |
| GTEX-QXCU-0326-SM-2TC63 | Thyroid | TCGA-THCA | 58.98 | 10.59 |
| GTEX-R3RS-0726-SM-3GIJR | Thyroid | TCGA-THCA | 61.81 | 11.11 |
| GTEX-R53T-0526-SM-3GADL | Thyroid | TCGA-THCA | 55.9 | 9.14 |
| GTEX-R55C-0626-SM-2TF4Q | Thyroid | TCGA-THCA | 52.79 | 11.45 |
| GTEX-R55E-0826-SM-2TC5M | Thyroid | TCGA-THCA | 50.7 | 13.44 |
| GTEX-R55G-0726-SM-2TC6J | Thyroid | TCGA-THCA | 47.63 | 11.01 |
| GTEX-REY6-0526-SM-2TF5M | Thyroid | TCGA-THCA | 42.55 | 10.58 |
| GTEX-RM2N-0526-SM-2TF4N | Thyroid | TCGA-THCA | 49.89 | 13.62 |
| GTEX-RN5K-2826-SM-EZ6L3 | Thyroid | TCGA-THCA | 60.99 | 13.29 |
| GTEX-RN64-0626-SM-2TC5V | Thyroid | TCGA-THCA | 53.64 | 12.04 |
| GTEX-RNOR-0926-SM-2TF56 | Thyroid | TCGA-THCA | 39.11 | 12.97 |
| GTEX-RTLS-0626-SM-5SI7Z | Thyroid | TCGA-THCA | 33.29 | 9.02 |
| GTEX-RU1J-0226-SM-2TF5Y | Thyroid | TCGA-THCA | 49.47 | 7.26 |
| GTEX-RU72-0126-SM-2TF6Z | Thyroid | TCGA-THCA | 39.07 | 9.29 |
| GTEX-RUSQ-1026-SM-2TF6V | Thyroid | TCGA-THCA | 36.58 | 13.69 |

| | | | | |
|---|---|---|---|---|
| GTEX-RVPV-1226-SM-2TF73 | Thyroid | TCGA-THCA | 53.68 | 13.88 |
| GTEX-RWS6-0626-SM-2XCAS | Thyroid | TCGA-THCA | 44.44 | 10.14 |
| GTEX-RWSA-0826-SM-2XCBF | Thyroid | TCGA-THCA | 43.88 | 10.69 |
| GTEX-S32W-0726-SM-2XCBL | Thyroid | TCGA-THCA | 47.01 | 9.04 |
| GTEX-S341-0226-SM-D4P1Y | Thyroid | TCGA-THCA | 45.71 | 12.69 |
| GTEX-S7SE-0726-SM-2XCD7 | Thyroid | TCGA-THCA | 62.57 | 12.72 |
| GTEX-S7SF-0226-SM-5SI7H | Thyroid | TCGA-THCA | 53.51 | 9.09 |
| GTEX-SE5C-0726-SM-4BRWY | Thyroid | TCGA-THCA | 55.8 | 11.13 |
| GTEX-SIU8-0626-SM-2XCDN | Thyroid | TCGA-THCA | 47.06 | 17.39 |
| GTEX-SJXC-0726-SM-2XCFJ | Thyroid | TCGA-THCA | 70.04 | 21.67 |
| GTEX-SN8G-1526-SM-4DM79 | Thyroid | TCGA-THCA | 49.41 | 9.78 |
| GTEX-SNOS-0226-SM-32PLR | Thyroid | TCGA-THCA | 50.83 | 11.58 |
| GTEX-SUCS-0226-SM-32PLQ | Thyroid | TCGA-THCA | 66.7 | 11.57 |
| GTEX-T2IS-0626-SM-32QP6 | Thyroid | TCGA-THCA | 56.93 | 13.22 |
| GTEX-T5JW-1226-SM-3GACY | Thyroid | TCGA-THCA | 66.07 | 13.32 |
| GTEX-T6MN-0626-SM-32PM9 | Thyroid | TCGA-THCA | 53.28 | 13.78 |
| GTEX-T6MO-0226-SM-32QOL | Thyroid | TCGA-THCA | 53.32 | 11.96 |
| GTEX-T8EM-0226-SM-3DB7C | Thyroid | TCGA-THCA | 51.32 | 15.57 |
| GTEX-TKQ1-0126-SM-33HB3 | Thyroid | TCGA-THCA | 73.17 | 11.73 |
| GTEX-TMMY-0826-SM-33HB9 | Thyroid | TCGA-THCA | 38.27 | 8.23 |
| GTEX-TSE9-0626-SM-3DB8B | Thyroid | TCGA-THCA | 40.97 | 7.29 |
| GTEX-U3ZH-0226-SM-3DB7B | Thyroid | TCGA-THCA | 42.62 | 10.52 |
| GTEX-U3ZM-0126-SM-3DB8M | Thyroid | TCGA-THCA | 57.59 | 10.14 |
| GTEX-U3ZN-0326-SM-3DB86 | Thyroid | TCGA-THCA | 53.87 | 17.15 |
| GTEX-U412-0726-SM-EVR2Y | Thyroid | TCGA-THCA | 61.16 | 13.19 |
| GTEX-U4B1-0626-SM-3DB8L | Thyroid | TCGA-THCA | 57.59 | 10.56 |
| GTEX-U8T8-2326-SM-3DB96 | Thyroid | TCGA-THCA | 61.2 | 12.78 |
| GTEX-UJMC-0326-SM-3GAE2 | Thyroid | TCGA-THCA | 66.56 | 18.09 |
| GTEX-V1D1-0926-SM-4JBHQ | Thyroid | TCGA-THCA | 42.94 | 12.07 |
| GTEX-V955-0426-SM-3GAEL | Thyroid | TCGA-THCA | 53.76 | 15.02 |
| GTEX-VJYA-0426-SM-3GIJK | Thyroid | TCGA-THCA | 61.84 | 18.94 |

| | | | | |
|---|---|---|---|---|
| GTEX-VUSG-0426-SM-3GIKD | Thyroid | TCGA-THCA | 72.45 | 16.14 |
| GTEX-W5WG-1426-SM-4KKZP | Thyroid | TCGA-THCA | 51.94 | 10.75 |
| GTEX-W5X1-0426-SM-3GILB | Thyroid | TCGA-THCA | 47.54 | 12.78 |
| GTEX-WEY5-0526-SM-3GIKZ | Thyroid | TCGA-THCA | 60.06 | 11.53 |
| GTEX-WFG7-0326-SM-DKPO8 | Thyroid | TCGA-THCA | 58.35 | 10.75 |
| GTEX-WFG8-0426-SM-3GILD | Thyroid | TCGA-THCA | 65.01 | 10.49 |
| GTEX-WFJO-0226-SM-3GIKW | Thyroid | TCGA-THCA | 44.95 | 11.27 |
| GTEX-WH7G-0526-SM-3NMBI | Thyroid | TCGA-THCA | 48.68 | 8.69 |
| GTEX-WHPG-0226-SM-3NMB9 | Thyroid | TCGA-THCA | 55.01 | 18.49 |
| GTEX-WHSB-1626-SM-3LK6J | Thyroid | TCGA-THCA | 52.28 | 10.49 |
| GTEX-WHSE-0626-SM-4RGNF | Thyroid | TCGA-THCA | 51.67 | 10.84 |
| GTEX-WK11-0926-SM-3NMAU | Thyroid | TCGA-THCA | 62.61 | 16.12 |
| GTEX-WL46-0126-SM-3TW8I | Thyroid | TCGA-THCA | 45.43 | 8.2 |
| GTEX-WOFL-0726-SM-3MJG4 | Thyroid | TCGA-THCA | 43.77 | 14.28 |
| GTEX-WRHU-0926-SM-4E3IG | Thyroid | TCGA-THCA | 40.98 | 10.98 |
| GTEX-WVLH-0626-SM-3MJG7 | Thyroid | TCGA-THCA | 48.81 | 15.09 |
| GTEX-WWYW-0826-SM-3NB2X | Thyroid | TCGA-THCA | 61.59 | 13.14 |
| GTEX-WXYG-0226-SM-3NB2Y | Thyroid | TCGA-THCA | 40.59 | 11.75 |
| GTEX-WY7C-0226-SM-3NB37 | Thyroid | TCGA-THCA | 46 | 8.91 |
| GTEX-WYBS-1926-SM-3NM8N | Thyroid | TCGA-THCA | 50.52 | 12.87 |
| GTEX-WYJK-1626-SM-3NM9J | Thyroid | TCGA-THCA | 52.4 | 13.34 |
| GTEX-WYVS-0326-SM-3NM9V | Thyroid | TCGA-THCA | 56.42 | 13.6 |
| GTEX-X15G-0526-SM-3NMB7 | Thyroid | TCGA-THCA | 41.97 | 15.5 |
| GTEX-X261-0726-SM-EZ6MH | Thyroid | TCGA-THCA | 64.73 | 11.59 |
| GTEX-X4LF-0426-SM-3NMB5 | Thyroid | TCGA-THCA | 54.06 | 7.01 |
| GTEX-X4XX-0926-SM-46MV7 | Thyroid | TCGA-THCA | 46.98 | 11.53 |
| GTEX-X4XY-0826-SM-4E3JM | Thyroid | TCGA-THCA | 55.42 | 13.4 |
| GTEX-X5EB-0726-SM-46MVR | Thyroid | TCGA-THCA | 79.51 | 11.28 |
| GTEX-X8HC-0726-SM-46MWG | Thyroid | TCGA-THCA | 50.31 | 9.16 |
| GTEX-XBED-0126-SM-47JY7 | Thyroid | TCGA-THCA | 66.64 | 11.48 |
| GTEX-XBEW-0126-SM-4AT66 | Thyroid | TCGA-THCA | 54.76 | 11.97 |

| | | | | |
|---|---|---|---|---|
| GTEX-XGQ4-0426-SM-4AT4I | Thyroid | TCGA-THCA | 71.34 | 16.84 |
| GTEX-XLM4-0726-SM-4AT64 | Thyroid | TCGA-THCA | 52.48 | 12.16 |
| GTEX-XMK1-0626-SM-4B65A | Thyroid | TCGA-THCA | 56.55 | 11.73 |
| GTEX-XOT4-1526-SM-EAZ2W | Thyroid | TCGA-THCA | 50.54 | 14.02 |
| GTEX-XUW1-1026-SM-4BONY | Thyroid | TCGA-THCA | 57.58 | 8.18 |
| GTEX-XUZC-0126-SM-4BOO6 | Thyroid | TCGA-THCA | 63.7 | 21.47 |
| GTEX-XV7Q-0326-SM-4BRVM | Thyroid | TCGA-THCA | 52.95 | 12.85 |
| GTEX-XXEK-1326-SM-4BRV1 | Thyroid | TCGA-THCA | 59.41 | 12.06 |
| GTEX-XYKS-0826-SM-4BRVF | Thyroid | TCGA-THCA | 34.66 | 13.57 |
| GTEX-Y111-1926-SM-4SOIS | Thyroid | TCGA-THCA | 57.96 | 9.84 |
| GTEX-Y114-0626-SM-4TT98 | Thyroid | TCGA-THCA | 52.91 | 10.31 |
| GTEX-Y3I4-0226-SM-4TT27 | Thyroid | TCGA-THCA | 50.64 | 9.69 |
| GTEX-Y3IK-0526-SM-4WWE3 | Thyroid | TCGA-THCA | 51.36 | 10.94 |
| GTEX-Y5LM-0626-SM-4V6G4 | Thyroid | TCGA-THCA | 56.79 | 11.35 |
| GTEX-Y5V5-0326-SM-5RQJG | Thyroid | TCGA-THCA | 48.42 | 8.39 |
| GTEX-Y5V6-0526-SM-4VBRV | Thyroid | TCGA-THCA | 50.51 | 17.16 |
| GTEX-Y8E4-0126-SM-4VBQ2 | Thyroid | TCGA-THCA | 48.76 | 13.03 |
| GTEX-Y9LG-0426-SM-4VBRT | Thyroid | TCGA-THCA | 76.17 | 10.25 |
| GTEX-YB5E-0626-SM-4VDSE | Thyroid | TCGA-THCA | 55.7 | 13.2 |
| GTEX-YB5K-0526-SM-5LUAS | Thyroid | TCGA-THCA | 54.19 | 10 |
| GTEX-YEC3-0826-SM-4WWFP | Thyroid | TCGA-THCA | 48.38 | 12.27 |
| GTEX-YEC4-0626-SM-5CVLU | Thyroid | TCGA-THCA | 64.15 | 12.93 |
| GTEX-YF7O-0726-SM-4W213 | Thyroid | TCGA-THCA | 75.24 | 17.73 |
| GTEX-YFC4-2626-SM-5P9FQ | Thyroid | TCGA-THCA | 53.08 | 13.27 |
| GTEX-YFCO-0326-SM-4W1ZP | Thyroid | TCGA-THCA | 59.57 | 11.12 |
| GTEX-YJ89-0726-SM-5P9F7 | Thyroid | TCGA-THCA | 48.89 | 12.5 |
| GTEX-Z93S-0526-SM-EVYB4 | Thyroid | TCGA-THCA | 34.87 | 10.27 |
| GTEX-Z9EW-0226-SM-5CVM7 | Thyroid | TCGA-THCA | 62.21 | 12.42 |
| GTEX-ZA64-0426-SM-5HL96 | Thyroid | TCGA-THCA | 63.06 | 11.31 |
| GTEX-ZAB5-0726-SM-5P9JG | Thyroid | TCGA-THCA | 51.45 | 12.01 |
| GTEX-ZAJG-0726-SM-5HL9A | Thyroid | TCGA-THCA | 83.99 | 10.88 |

| | | | | |
|---|---|---|---|---|
| GTEX-ZAK1-0726-SM-5HL8Q | Thyroid | TCGA-THCA | 50.54 | 11.7 |
| GTEX-ZC5H-0626-SM-5LU9K | Thyroid | TCGA-THCA | 37.23 | 13.29 |
| GTEX-ZDTS-0926-SM-5YY9D | Thyroid | TCGA-THCA | 64.43 | 14.6 |
| GTEX-ZDXO-1726-SM-5HL6L | Thyroid | TCGA-THCA | 51.52 | 14.47 |
| GTEX-ZDYS-0626-SM-5J2N5 | Thyroid | TCGA-THCA | 62.63 | 13.07 |
| GTEX-ZE7O-1126-SM-57WC8 | Thyroid | TCGA-THCA | 63.9 | 17.98 |
| GTEX-ZF28-0826-SM-4WKGJ | Thyroid | TCGA-THCA | 50.85 | 8.25 |
| GTEX-ZF29-0726-SM-DO92H | Thyroid | TCGA-THCA | 56.7 | 10.21 |
| GTEX-ZF3C-0826-SM-DO11E | Thyroid | TCGA-THCA | 41.02 | 12.39 |
| GTEX-ZGAY-1026-SM-4WWBR | Thyroid | TCGA-THCA | 61.59 | 9.38 |
| GTEX-ZLFU-0626-SM-4WWBO | Thyroid | TCGA-THCA | 44.41 | 8.84 |
| GTEX-ZLV1-0126-SM-4WWBZ | Thyroid | TCGA-THCA | 38.54 | 10.7 |
| GTEX-ZLWG-0526-SM-4WWFB | Thyroid | TCGA-THCA | 74.12 | 14.23 |
| GTEX-ZPCL-0126-SM-4WWC8 | Thyroid | TCGA-THCA | 62.73 | 11.65 |
| GTEX-ZPIC-0226-SM-DO91T | Thyroid | TCGA-THCA | 62.02 | 11.75 |
| GTEX-ZPU1-0426-SM-4WWCA | Thyroid | TCGA-THCA | 54.99 | 12.02 |
| GTEX-ZQG8-0926-SM-57WFF | Thyroid | TCGA-THCA | 63.61 | 13.04 |
| GTEX-ZQUD-0126-SM-7EPIS | Thyroid | TCGA-THCA | 51.71 | 10.75 |
| GTEX-ZT9W-0226-SM-7SB6H | Thyroid | TCGA-THCA | 43.73 | 7.5 |
| GTEX-ZT9X-0226-SM-51MT2 | Thyroid | TCGA-THCA | 47.75 | 8.92 |
| GTEX-ZTPG-0826-SM-5DUVC | Thyroid | TCGA-THCA | 82.69 | 13.87 |
| GTEX-ZTSS-0226-SM-59877 | Thyroid | TCGA-THCA | 61.48 | 12.73 |
| GTEX-ZTX8-0626-SM-59HKC | Thyroid | TCGA-THCA | 65.49 | 10.7 |
| GTEX-ZUA1-0926-SM-4YCDX | Thyroid | TCGA-THCA | 66.57 | 15.17 |
| GTEX-ZV6S-0226-SM-59HJT | Thyroid | TCGA-THCA | 63.12 | 11.95 |
| GTEX-ZV7C-0126-SM-57WDE | Thyroid | TCGA-THCA | 56.28 | 14.72 |
| GTEX-ZVP2-0426-SM-57WC2 | Thyroid | TCGA-THCA | 52.83 | 8.97 |
| GTEX-ZVT3-0726-SM-5GICN | Thyroid | TCGA-THCA | 46.92 | 11.11 |
| GTEX-ZVZP-1026-SM-5GICI | Thyroid | TCGA-THCA | 51.83 | 12.54 |
| GTEX-ZVZQ-0626-SM-59HJU | Thyroid | TCGA-THCA | 49.92 | 12.11 |
| GTEX-ZXG5-0926-SM-5NQ8H | Thyroid | TCGA-THCA | 49.11 | 12.93 |

| | | | | |
|---|---|---|---|---|
| GTEX-ZY6K-0226-SM-5SIAY | Thyroid | TCGA-THCA | 55.76 | 17.94 |
| GTEX-ZYFC-0926-SM-5GZWW | Thyroid | TCGA-THCA | 52.32 | 14.99 |
| GTEX-ZYFD-0826-SM-5NQ9A | Thyroid | TCGA-THCA | 67.79 | 14.58 |
| GTEX-ZYFG-0626-SM-5GZYA | Thyroid | TCGA-THCA | 59.48 | 10.82 |
| GTEX-ZYT6-0426-SM-5GID3 | Thyroid | TCGA-THCA | 60.97 | 9.45 |
| GTEX-ZYVF-1126-SM-5E458 | Thyroid | TCGA-THCA | 70.59 | 16.95 |
| GTEX-ZYW4-1126-SM-5SI99 | Thyroid | TCGA-THCA | 36.45 | 10.15 |
| GTEX-ZYY3-1926-SM-5GZXS | Thyroid | TCGA-THCA | 34.83 | 8.74 |
| GTEX-ZZ64-0126-SM-5GZXA | Thyroid | TCGA-THCA | 67.11 | 11.06 |
| GTEX-ZZPU-1326-SM-5GZWS | Thyroid | TCGA-THCA | 56.57 | 9.47 |
| GTEX-1117F-2426-SM-5EGGH | Uterus | TCGA-UCS | 50.35 | 15.45 |
| GTEX-113JC-2226-SM-5EGJG | Uterus | TCGA-UCS | 65.31 | 16.55 |
| GTEX-11DXX-1526-SM-5H115 | Uterus | TCGA-UCS | 65.43 | 16.01 |
| GTEX-11EM3-1926-SM-5987U | Uterus | TCGA-UCS | 55.67 | 12.3 |
| GTEX-11EMC-1826-SM-5A5JT | Uterus | TCGA-UCS | 57.48 | 22.77 |
| GTEX-11GSP-2426-SM-5N9BD | Uterus | TCGA-UCS | 51.21 | 14.81 |
| GTEX-11I78-2126-SM-5A5K8 | Uterus | TCGA-UCS | 80.66 | 16.67 |
| GTEX-11P81-1626-SM-5BC52 | Uterus | TCGA-UCS | 64.59 | 18.15 |
| GTEX-11ZTS-2326-SM-5EQMY | Uterus | TCGA-UCS | 57.56 | 17.91 |
| GTEX-11ZTT-1726-SM-5EQL4 | Uterus | TCGA-UCS | 42.37 | 14.84 |
| GTEX-12WSD-2826-SM-59HKT | Uterus | TCGA-UCS | 51.39 | 18.81 |
| GTEX-12WSG-2126-SM-5EGJ7 | Uterus | TCGA-UCS | 54.7 | 13.41 |
| GTEX-12WSK-2026-SM-5CVNB | Uterus | TCGA-UCS | 56.9 | 10.41 |
| GTEX-12ZZX-2126-SM-5LZVL | Uterus | TCGA-UCS | 64.2 | 15.51 |
| GTEX-1313W-2826-SM-5P9G1 | Uterus | TCGA-UCS | 49.87 | 16.82 |
| GTEX-131XG-2026-SM-5GCN5 | Uterus | TCGA-UCS | 54.91 | 22.57 |
| GTEX-131YS-2326-SM-5IJFJ | Uterus | TCGA-UCS | 55.59 | 20.98 |
| GTEX-132AR-1526-SM-5KM1L | Uterus | TCGA-UCS | 66.77 | 11.73 |
| GTEX-1399S-2226-SM-5IFEW | Uterus | TCGA-UCS | 66.49 | 11.12 |
| GTEX-1399U-1326-SM-5IJET | Uterus | TCGA-UCS | 63.45 | 12.51 |
| GTEX-139D8-2526-SM-5N9G3 | Uterus | TCGA-UCS | 49.98 | 13.16 |

| | | | | |
|---|---|---|---|---|
| GTEX-13D11-1226-SM-5IFGA | Uterus | TCGA-UCS | 70.28 | 18.16 |
| GTEX-13FTX-1026-SM-5J2O5 | Uterus | TCGA-UCS | 57.16 | 10.37 |
| GTEX-13N11-1126-SM-5KM41 | Uterus | TCGA-UCS | 52.08 | 17.84 |
| GTEX-13OVI-1026-SM-5L3EM | Uterus | TCGA-UCS | 46.44 | 8.65 |
| GTEX-13OVJ-2326-SM-5IJGA | Uterus | TCGA-UCS | 41.31 | 14.36 |
| GTEX-13PL7-2026-SM-5IFGK | Uterus | TCGA-UCS | 47.71 | 12.98 |
| GTEX-13QBU-1626-SM-5LU4S | Uterus | TCGA-UCS | 43.18 | 10.99 |
| GTEX-13S7M-1726-SM-5RQK3 | Uterus | TCGA-UCS | 50.22 | 15.89 |
| GTEX-13U4I-1926-SM-5LU39 | Uterus | TCGA-UCS | 64.12 | 16.17 |
| GTEX-13VXT-1626-SM-5IJES | Uterus | TCGA-UCS | 56.63 | 14.42 |
| GTEX-13W3W-1426-SM-5LU56 | Uterus | TCGA-UCS | 50.44 | 15.04 |
| GTEX-145LS-2426-SM-5TDCZ | Uterus | TCGA-UCS | 57.27 | 20.76 |
| GTEX-145ME-1326-SM-5O98Q | Uterus | TCGA-UCS | 59.28 | 12.15 |
| GTEX-14AS3-1826-SM-5TDD8 | Uterus | TCGA-UCS | 66.73 | 15.37 |
| GTEX-14BIM-2226-SM-5SI8Y | Uterus | TCGA-UCS | 70.22 | 16.77 |
| GTEX-14BIN-1526-SM-73KZ7 | Uterus | TCGA-UCS | 47.35 | 19.39 |
| GTEX-14BMU-1826-SM-5RQII | Uterus | TCGA-UCS | 67.19 | 13.14 |
| GTEX-14JG6-0826-SM-6AJB9 | Uterus | TCGA-UCS | 65.49 | 12.54 |
| GTEX-14PHW-2426-SM-69LPU | Uterus | TCGA-UCS | 32.63 | 12.56 |
| GTEX-14PJ6-1526-SM-6AJ9M | Uterus | TCGA-UCS | 65.45 | 16.18 |
| GTEX-14PJM-2826-SM-69LPV | Uterus | TCGA-UCS | 49.64 | 15.04 |
| GTEX-14PKV-1426-SM-5YYB9 | Uterus | TCGA-UCS | 51.91 | 9.07 |
| GTEX-15CHC-2026-SM-6EU1Z | Uterus | TCGA-UCS | 47 | 16.28 |
| GTEX-15DZA-1326-SM-68722 | Uterus | TCGA-UCS | 58.87 | 15.39 |
| GTEX-15ER7-2426-SM-793B9 | Uterus | TCGA-UCS | 59.09 | 16.85 |
| GTEX-15ETS-1526-SM-6LPKM | Uterus | TCGA-UCS | 55.03 | 13.09 |
| GTEX-15FZZ-1026-SM-6AJBI | Uterus | TCGA-UCS | 63.19 | 13.24 |
| GTEX-16A39-0726-SM-7KUEB | Uterus | TCGA-UCS | 44.53 | 13.43 |
| GTEX-16NGA-0626-SM-72D72 | Uterus | TCGA-UCS | 65.58 | 17.45 |
| GTEX-1A32A-2826-SM-72D5S | Uterus | TCGA-UCS | 62.44 | 18.02 |
| GTEX-1A8FM-2326-SM-7MKGC | Uterus | TCGA-UCS | 59.57 | 17.94 |

| | | | | |
|---|---|---|---|---|
| GTEX-1AMEY-0626-SM-72D6B | Uterus | TCGA-UCS | 62.2 | 15.67 |
| GTEX-1B932-2726-SM-731ER | Uterus | TCGA-UCS | 63.06 | 22.28 |
| GTEX-1B97J-0226-SM-79392 | Uterus | TCGA-UCS | 45.83 | 14.48 |
| GTEX-1C475-0626-SM-73KV9 | Uterus | TCGA-UCS | 56.05 | 10.32 |
| GTEX-1CAMS-2026-SM-79395 | Uterus | TCGA-UCS | 43.63 | 17.77 |
| GTEX-1CAV2-1426-SM-7MGX6 | Uterus | TCGA-UCS | 44.15 | 8.86 |
| GTEX-1CB4H-2726-SM-7EWFF | Uterus | TCGA-UCS | 41.15 | 16.18 |
| GTEX-1GF9U-1826-SM-7P8T7 | Uterus | TCGA-UCS | 53.42 | 12.55 |
| GTEX-1GMR8-2426-SM-7MKH3 | Uterus | TCGA-UCS | 54.06 | 16.18 |
| GTEX-1GN1U-2526-SM-7MKH4 | Uterus | TCGA-UCS | 56.33 | 10.37 |
| GTEX-1GN1W-1326-SM-9JGHZ | Uterus | TCGA-UCS | 68.94 | 15.86 |
| GTEX-1GZ4H-0826-SM-9OSXK | Uterus | TCGA-UCS | 51.04 | 14.56 |
| GTEX-1GZHY-2526-SM-9OSW6 | Uterus | TCGA-UCS | 54.7 | 15.36 |
| GTEX-1H1CY-2126-SM-9OSXV | Uterus | TCGA-UCS | 56.94 | 19.22 |
| GTEX-1HBPH-2826-SM-9WYUU | Uterus | TCGA-UCS | 38.47 | 14.28 |
| GTEX-1HCUA-1226-SM-9WPP5 | Uterus | TCGA-UCS | 59.07 | 14.3 |
| GTEX-1HSGN-2326-SM-A9G1D | Uterus | TCGA-UCS | 47.18 | 13.88 |
| GTEX-1I4MK-1426-SM-B2LWU | Uterus | TCGA-UCS | 85.03 | 29.28 |
| GTEX-1ICG6-1426-SM-A96T1 | Uterus | TCGA-UCS | 43.76 | 21.08 |
| GTEX-1IDJI-1726-SM-C1YQE | Uterus | TCGA-UCS | 60.76 | 13.53 |
| GTEX-1IKOE-2426-SM-ARU7W | Uterus | TCGA-UCS | 57.74 | 12.98 |
| GTEX-1J1OQ-1726-SM-CL55N | Uterus | TCGA-UCS | 64.24 | 24.52 |
| GTEX-1JN76-1626-SM-CNNQ3 | Uterus | TCGA-UCS | 60.03 | 15.71 |
| GTEX-1K9T9-2526-SM-D3L9B | Uterus | TCGA-UCS | 97.68 | 21.8 |
| GTEX-1KANC-1126-SM-CY8I7 | Uterus | TCGA-UCS | 54.46 | 15.66 |
| GTEX-1LGOU-1226-SM-D3L8L | Uterus | TCGA-UCS | 61.78 | 14.41 |
| GTEX-1LGRB-2226-SM-CNNQI | Uterus | TCGA-UCS | 63.29 | 10.66 |
| GTEX-1LH75-1626-SM-EWROL | Uterus | TCGA-UCS | 56.2 | 14.23 |
| GTEX-1MA7W-1526-SM-DHXKS | Uterus | TCGA-UCS | 64.25 | 15.33 |
| GTEX-1MCYP-1226-SM-DTX8H | Uterus | TCGA-UCS | 64.3 | 11.73 |
| GTEX-1NV5F-1726-SM-EXOIZ | Uterus | TCGA-UCS | 51.91 | 17.98 |

| | | | | |
|---|---|---|---|---|
| GTEX-1PBJI-2326-SM-E6CP6 | Uterus | TCGA-UCS | 54.19 | 12.95 |
| GTEX-1PIEJ-1726-SM-EVYB6 | Uterus | TCGA-UCS | 47.06 | 8.57 |
| GTEX-1QP28-1526-SM-E76P6 | Uterus | TCGA-UCS | 96.78 | 19.24 |
| GTEX-1R9JW-1326-SM-E6CQ7 | Uterus | TCGA-UCS | 60.79 | 19.18 |
| GTEX-1R9PN-0526-SM-DTX86 | Uterus | TCGA-UCS | 46.18 | 14.79 |
| GTEX-1RDX4-0726-SM-E8VO8 | Uterus | TCGA-UCS | 43.54 | 16.95 |
| GTEX-1S82U-1226-SM-E9TKB | Uterus | TCGA-UCS | 74.61 | 20.47 |
| GTEX-N7MT-0726-SM-3TW8S | Uterus | TCGA-UCS | 44.98 | 16.74 |
| GTEX-OHPK-2026-SM-3MJH7 | Uterus | TCGA-UCS | 59.26 | 12.32 |
| GTEX-OHPL-2026-SM-3TW8R | Uterus | TCGA-UCS | 69.94 | 19.74 |
| GTEX-OHPN-1926-SM-E9TI9 | Uterus | TCGA-UCS | 41.54 | 11.51 |
| GTEX-P4PP-2026-SM-3P61N | Uterus | TCGA-UCS | 50.48 | 15.61 |
| GTEX-P4QT-2026-SM-3NMCJ | Uterus | TCGA-UCS | 55 | 11.53 |
| GTEX-P78B-2526-SM-3P5ZY | Uterus | TCGA-UCS | 49.93 | 13.6 |
| GTEX-POMQ-1226-SM-3P61F | Uterus | TCGA-UCS | 49.29 | 12.33 |
| GTEX-PWCY-1426-SM-48TCT | Uterus | TCGA-UCS | 48.46 | 11.25 |
| GTEX-PWN1-2026-SM-48TD9 | Uterus | TCGA-UCS | 53.23 | 14.02 |
| GTEX-PX3G-2026-SM-48U1H | Uterus | TCGA-UCS | 54.73 | 10.61 |
| GTEX-Q734-1626-SM-48U1B | Uterus | TCGA-UCS | 61.48 | 15.55 |
| GTEX-QCQG-1326-SM-48U24 | Uterus | TCGA-UCS | 65.78 | 28.34 |
| GTEX-R55G-1626-SM-48FF4 | Uterus | TCGA-UCS | 59.33 | 25.16 |
| GTEX-RTLS-2426-SM-46MUO | Uterus | TCGA-UCS | 36.16 | 20.36 |
| GTEX-RU1J-1026-SM-46MUR | Uterus | TCGA-UCS | 54.69 | 12.99 |
| GTEX-RU72-2626-SM-CYPS6 | Uterus | TCGA-UCS | 61.1 | 15.21 |
| GTEX-RWS6-1326-SM-47JXB | Uterus | TCGA-UCS | 68.19 | 16.97 |
| GTEX-S32W-1426-SM-4AD66 | Uterus | TCGA-UCS | 67.11 | 20.3 |
| GTEX-S341-1026-SM-4AD71 | Uterus | TCGA-UCS | 47.13 | 18.23 |
| GTEX-S4UY-1226-SM-4AD51 | Uterus | TCGA-UCS | 52.8 | 12.89 |
| GTEX-S7SF-0826-SM-4AD4W | Uterus | TCGA-UCS | 49.28 | 15.6 |
| GTEX-T2IS-2226-SM-4DM65 | Uterus | TCGA-UCS | 76.78 | 20.84 |
| GTEX-T5JW-1526-SM-4DM5E | Uterus | TCGA-UCS | 47.17 | 13.29 |

| | | | | |
|---|---|---|---|---|
| GTEX-T6MO-1526-SM-4DM57 | Uterus | TCGA-UCS | 48.19 | 10.76 |
| GTEX-TMMY-2226-SM-4DXTN | Uterus | TCGA-UCS | 59.6 | 10.53 |
| GTEX-TSE9-2626-SM-4DXV2 | Uterus | TCGA-UCS | 40.65 | 7.71 |
| GTEX-U3ZN-0726-SM-4DXT5 | Uterus | TCGA-UCS | 49.72 | 12.08 |
| GTEX-W5WG-1326-SM-4LMI9 | Uterus | TCGA-UCS | 58.16 | 19.37 |
| GTEX-WEY5-0726-SM-4LMID | Uterus | TCGA-UCS | 48.94 | 11.58 |
| GTEX-WI4N-2426-SM-4OOSC | Uterus | TCGA-UCS | 29.16 | 8.42 |
| GTEX-WWYW-2826-SM-4MVOQ | Uterus | TCGA-UCS | 63.91 | 11.49 |
| GTEX-WXYG-1326-SM-4ONCN | Uterus | TCGA-UCS | 45.84 | 15.21 |
| GTEX-WYVS-1226-SM-4ONCL | Uterus | TCGA-UCS | 48.72 | 12.88 |
| GTEX-XUW1-0226-SM-4BOOS | Uterus | TCGA-UCS | 67.13 | 15.94 |
| GTEX-XUZC-0926-SM-4BOQF | Uterus | TCGA-UCS | 45.65 | 14.41 |
| GTEX-XV7Q-1526-SM-4BRWB | Uterus | TCGA-UCS | 88.62 | 33.12 |
| GTEX-XYKS-1626-SM-4BRUQ | Uterus | TCGA-UCS | 57.55 | 21.71 |
| GTEX-Y114-1826-SM-4TT87 | Uterus | TCGA-UCS | 44.88 | 11.57 |
| GTEX-Y3IK-2126-SM-4YCDS | Uterus | TCGA-UCS | 47.33 | 11.88 |
| GTEX-YFC4-2826-SM-5P9FR | Uterus | TCGA-UCS | 47.44 | 13.92 |
| GTEX-YJ8O-1126-SM-5P9IS | Uterus | TCGA-UCS | 52.09 | 14.14 |
| GTEX-ZAJG-2826-SM-5HL8D | Uterus | TCGA-UCS | 115.63 | 19.34 |
| GTEX-ZAK1-2526-SM-5S2N7 | Uterus | TCGA-UCS | 36.09 | 12.1 |
| GTEX-ZLWG-1026-SM-4WWC4 | Uterus | TCGA-UCS | 70.88 | 16.1 |
| GTEX-ZP4G-0726-SM-4WWF2 | Uterus | TCGA-UCS | 59.09 | 12.72 |
| GTEX-ZPIC-1226-SM-DO91X | Uterus | TCGA-UCS | 70.11 | 23.65 |
| GTEX-ZTPG-2026-SM-5O999 | Uterus | TCGA-UCS | 82.14 | 15.85 |
| GTEX-ZV6S-1526-SM-5NQ7O | Uterus | TCGA-UCS | 81.27 | 19.13 |
| GTEX-ZVT2-0426-SM-5E44S | Uterus | TCGA-UCS | 75.91 | 15.51 |
| GTEX-ZXES-0726-SM-5E451 | Uterus | TCGA-UCS | 50.57 | 8.5 |
| GTEX-ZYVF-0526-SM-5E43E | Uterus | TCGA-UCS | 46.44 | 14.74 |

| Table S2. MAGOH and MAGOHB expression in tumor (TCGA) data. | | | |
|---|---|---|---|
| **sample_id** | **tcga_correspondence** | **MAGOH_tpm** | **MAGOHB_tpm** |
| TCGA-02-0047 | TCGA-GBM | 38.164676 | 13.6963106 |
| TCGA-02-0055 | TCGA-GBM | 85.0317744 | 27.5218395 |
| TCGA-02-2483 | TCGA-GBM | 95.283095 | 38.7765385 |
| TCGA-02-2485 | TCGA-GBM | 65.4858311 | 32.7891766 |
| TCGA-02-2486 | TCGA-GBM | 70.6058681 | 25.0475672 |
| TCGA-04-1331 | TCGA-OV | 126.564122 | 35.4333389 |
| TCGA-04-1332 | TCGA-OV | 12.0100047 | 9.77170248 |
| TCGA-04-1338 | TCGA-OV | 60.9859768 | 54.5501001 |
| TCGA-04-1341 | TCGA-OV | 39.4866892 | 19.6682365 |
| TCGA-04-1343 | TCGA-OV | 70.2157796 | 24.0159209 |
| TCGA-04-1347 | TCGA-OV | 87.1272849 | 37.5335245 |
| TCGA-04-1350 | TCGA-OV | 53.4836763 | 12.8195201 |
| TCGA-04-1356 | TCGA-OV | 122.690745 | 76.1543452 |
| TCGA-04-1357 | TCGA-OV | 40.741664 | 22.9933168 |
| TCGA-04-1361 | TCGA-OV | 87.9616068 | 17.14403 |
| TCGA-04-1362 | TCGA-OV | 81.8166223 | 21.9397379 |
| TCGA-04-1364 | TCGA-OV | 83.152035 | 22.3311211 |
| TCGA-04-1365 | TCGA-OV | 93.971582 | 44.3843988 |
| TCGA-04-1514 | TCGA-OV | 72.1815484 | 21.3386713 |
| TCGA-04-1519 | TCGA-OV | 96.5081859 | 16.0071888 |
| TCGA-04-1530 | TCGA-OV | 50.5100611 | 24.0428344 |
| TCGA-04-1536 | TCGA-OV | 91.4875074 | 93.936253 |
| TCGA-04-1542 | TCGA-OV | 40.1052462 | 42.8420149 |
| TCGA-04-1648 | TCGA-OV | 57.553833 | 30.5643098 |
| TCGA-04-1651 | TCGA-OV | 42.8800668 | 69.4760234 |
| TCGA-04-1655 | TCGA-OV | 51.2006695 | 41.0201986 |
| TCGA-05-4244 | TCGA-LUAD | 40.2926607 | 27.6895137 |
| TCGA-05-4249 | TCGA-LUAD | 45.4177003 | 16.2268823 |
| TCGA-05-4250 | TCGA-LUAD | 48.5417123 | 19.9120432 |
| TCGA-05-4382 | TCGA-LUAD | 42.5989764 | 16.6988656 |

| | | | |
|---|---|---|---|
| TCGA-05-4384 | TCGA-LUAD | 41.4718458 | 21.1373379 |
| TCGA-05-4389 | TCGA-LUAD | 42.8838906 | 27.0736933 |
| TCGA-05-4390 | TCGA-LUAD | 67.0930564 | 28.277685 |
| TCGA-05-4395 | TCGA-LUAD | 44.3114474 | 55.4509485 |
| TCGA-05-4396 | TCGA-LUAD | 28.0697995 | 51.8681417 |
| TCGA-05-4397 | TCGA-LUAD | 103.652843 | 44.2863043 |
| TCGA-05-4398 | TCGA-LUAD | 52.9504677 | 41.7208965 |
| TCGA-05-4402 | TCGA-LUAD | 53.7501131 | 22.0854303 |
| TCGA-05-4403 | TCGA-LUAD | 35.2479237 | 15.0332721 |
| TCGA-05-4405 | TCGA-LUAD | 39.5451558 | 14.1032903 |
| TCGA-05-4410 | TCGA-LUAD | 28.2864546 | 12.960147 |
| TCGA-05-4415 | TCGA-LUAD | 78.54505 | 73.1593091 |
| TCGA-05-4417 | TCGA-LUAD | 34.4550326 | 38.7634295 |
| TCGA-05-4418 | TCGA-LUAD | 37.1754082 | 23.1828067 |
| TCGA-05-4420 | TCGA-LUAD | 71.1374807 | 38.573245 |
| TCGA-05-4422 | TCGA-LUAD | 22.9186584 | 24.9773001 |
| TCGA-05-4424 | TCGA-LUAD | 35.9165692 | 21.4656824 |
| TCGA-05-4425 | TCGA-LUAD | 70.600488 | 16.7831792 |
| TCGA-05-4426 | TCGA-LUAD | 38.8845824 | 13.0508917 |
| TCGA-05-4427 | TCGA-LUAD | 113.370333 | 18.4452271 |
| TCGA-05-4430 | TCGA-LUAD | 42.0579622 | 28.1668256 |
| TCGA-05-4432 | TCGA-LUAD | 47.4219284 | 10.1319084 |
| TCGA-05-4433 | TCGA-LUAD | 37.3483166 | 22.0443518 |
| TCGA-05-4434 | TCGA-LUAD | 64.1615028 | 27.1505354 |
| TCGA-05-5420 | TCGA-LUAD | 28.1307883 | 35.0669144 |
| TCGA-05-5423 | TCGA-LUAD | 61.2045066 | 48.5491535 |
| TCGA-05-5425 | TCGA-LUAD | 58.1004252 | 38.6000214 |
| TCGA-05-5428 | TCGA-LUAD | 66.7153293 | 38.7976676 |
| TCGA-05-5429 | TCGA-LUAD | 90.0106403 | 24.9727039 |
| TCGA-05-5715 | TCGA-LUAD | 51.4310322 | 40.3463132 |
| TCGA-06-0125 | TCGA-GBM | 53.2694908 | 21.0828713 |

| | | | |
|---|---|---|---|
| TCGA-06-0129 | TCGA-GBM | 60.2484084 | 15.266424 |
| TCGA-06-0130 | TCGA-GBM | 91.1860283 | 45.6958519 |
| TCGA-06-0132 | TCGA-GBM | 32.4920213 | 15.8339589 |
| TCGA-06-0138 | TCGA-GBM | 40.1883036 | 29.1278319 |
| TCGA-06-0139 | TCGA-GBM | 45.9245285 | 18.5964452 |
| TCGA-06-0141 | TCGA-GBM | 51.8432049 | 14.3798288 |
| TCGA-06-0152 | TCGA-GBM | 56.724238 | 27.613946 |
| TCGA-06-0156 | TCGA-GBM | 53.5313797 | 30.9807802 |
| TCGA-06-0157 | TCGA-GBM | 46.6540555 | 25.8417248 |
| TCGA-06-0158 | TCGA-GBM | 42.695559 | 18.8541599 |
| TCGA-06-0168 | TCGA-GBM | 67.6937047 | 24.1376048 |
| TCGA-06-0171 | TCGA-GBM | 36.0278793 | 13.8291492 |
| TCGA-06-0174 | TCGA-GBM | 45.2782073 | 28.8940278 |
| TCGA-06-0178 | TCGA-GBM | 52.0706203 | 18.4201204 |
| TCGA-06-0184 | TCGA-GBM | 60.1623758 | 26.1188863 |
| TCGA-06-0187 | TCGA-GBM | 69.7697865 | 28.5453435 |
| TCGA-06-0190 | TCGA-GBM | 68.4952628 | 25.0811512 |
| TCGA-06-0210 | TCGA-GBM | 46.5464973 | 14.9165686 |
| TCGA-06-0211 | TCGA-GBM | 54.0845771 | 37.4958421 |
| TCGA-06-0219 | TCGA-GBM | 57.001031 | 37.91726 |
| TCGA-06-0221 | TCGA-GBM | 59.66346 | 11.6339325 |
| TCGA-06-0238 | TCGA-GBM | 53.1240651 | 39.4154313 |
| TCGA-06-0644 | TCGA-GBM | 78.1668688 | 43.9654821 |
| TCGA-06-0645 | TCGA-GBM | 48.9268766 | 21.1852275 |
| TCGA-06-0646 | TCGA-GBM | 46.884887 | 28.2835861 |
| TCGA-06-0649 | TCGA-GBM | 48.3241593 | 14.9086867 |
| TCGA-06-0686 | TCGA-GBM | 50.0832039 | 28.0329044 |
| TCGA-06-0743 | TCGA-GBM | 46.5326816 | 19.2970135 |
| TCGA-06-0744 | TCGA-GBM | 58.8120013 | 22.3272493 |
| TCGA-06-0745 | TCGA-GBM | 62.6999439 | 31.9047443 |
| TCGA-06-0747 | TCGA-GBM | 63.2713533 | 23.8135004 |

| | | | |
|---|---|---|---|
| TCGA-06-0749 | TCGA-GBM | 41.6658566 | 14.0575672 |
| TCGA-06-0750 | TCGA-GBM | 55.328238 | 20.2211493 |
| TCGA-06-0878 | TCGA-GBM | 68.3631983 | 29.3353905 |
| TCGA-06-0882 | TCGA-GBM | 40.9874896 | 15.0482666 |
| TCGA-06-1804 | TCGA-GBM | 52.9457874 | 18.8707102 |
| TCGA-06-2557 | TCGA-GBM | 64.8242667 | 23.3529138 |
| TCGA-06-2558 | TCGA-GBM | 60.7034649 | 35.1574292 |
| TCGA-06-2559 | TCGA-GBM | 64.947856 | 39.8193876 |
| TCGA-06-2561 | TCGA-GBM | 51.7995632 | 23.0059319 |
| TCGA-06-2562 | TCGA-GBM | 74.0437467 | 32.1517825 |
| TCGA-06-2563 | TCGA-GBM | 61.281203 | 31.4658185 |
| TCGA-06-2564 | TCGA-GBM | 50.3183519 | 28.5827286 |
| TCGA-06-2565 | TCGA-GBM | 63.997846 | 24.7216751 |
| TCGA-06-2567 | TCGA-GBM | 67.7719731 | 35.2215478 |
| TCGA-06-2569 | TCGA-GBM | 104.990983 | 31.5965976 |
| TCGA-06-2570 | TCGA-GBM | 45.8684398 | 12.5214438 |
| TCGA-06-5408 | TCGA-GBM | 99.6154389 | 42.5944325 |
| TCGA-06-5410 | TCGA-GBM | 52.8232832 | 19.7263275 |
| TCGA-06-5411 | TCGA-GBM | 50.5486184 | 31.5348416 |
| TCGA-06-5412 | TCGA-GBM | 64.7542591 | 32.8526475 |
| TCGA-06-5413 | TCGA-GBM | 89.1175619 | 46.7573893 |
| TCGA-06-5414 | TCGA-GBM | 58.8610421 | 27.9083257 |
| TCGA-06-5416 | TCGA-GBM | 61.389451 | 22.8688862 |
| TCGA-06-5417 | TCGA-GBM | 66.8142471 | 14.6673709 |
| TCGA-06-5418 | TCGA-GBM | 58.3025381 | 23.8448434 |
| TCGA-06-5856 | TCGA-GBM | 98.040666 | 56.920874 |
| TCGA-06-5858 | TCGA-GBM | 58.6861936 | 27.4910266 |
| TCGA-06-5859 | TCGA-GBM | 69.0273296 | 26.1927571 |
| TCGA-08-0386 | TCGA-GBM | 75.5535637 | 17.3527995 |
| TCGA-09-0364 | TCGA-OV | 70.5306965 | 80.8518177 |
| TCGA-09-0366 | TCGA-OV | 84.5397087 | 37.3878306 |

| | | | |
|---|---|---|---|
| TCGA-09-0367 | TCGA-OV | 47.1631307 | 22.5037466 |
| TCGA-09-0369 | TCGA-OV | 67.1144759 | 73.3384322 |
| TCGA-09-1659 | TCGA-OV | 55.6283815 | 29.013359 |
| TCGA-09-1661 | TCGA-OV | 29.3754686 | 49.1045497 |
| TCGA-09-1662 | TCGA-OV | 58.3612788 | 40.051043 |
| TCGA-09-1665 | TCGA-OV | 37.9100857 | 81.9714248 |
| TCGA-09-1666 | TCGA-OV | 93.000942 | 37.1880147 |
| TCGA-09-1667 | TCGA-OV | 74.5453355 | 21.0875767 |
| TCGA-09-1668 | TCGA-OV | 56.1819513 | 33.6864788 |
| TCGA-09-1669 | TCGA-OV | 76.2535381 | 51.3628662 |
| TCGA-09-1670 | TCGA-OV | 89.2875933 | 17.2883336 |
| TCGA-09-1673 | TCGA-OV | 56.3763463 | 6.28289848 |
| TCGA-09-2044 | TCGA-OV | 169.990592 | 91.6166687 |
| TCGA-09-2045 | TCGA-OV | 50.1298514 | 15.3609771 |
| TCGA-09-2048 | TCGA-OV | 69.7312614 | 47.4843099 |
| TCGA-09-2051 | TCGA-OV | 98.59849 | 59.0238981 |
| TCGA-09-2053 | TCGA-OV | 77.5207468 | 28.4001801 |
| TCGA-09-2054 | TCGA-OV | 88.2850132 | 26.0156675 |
| TCGA-09-2056 | TCGA-OV | 78.1738331 | 22.4561202 |
| TCGA-10-0927 | TCGA-OV | 46.7139782 | 16.3847727 |
| TCGA-10-0928 | TCGA-OV | 66.1637151 | 37.4989448 |
| TCGA-10-0931 | TCGA-OV | 105.416578 | 48.4158939 |
| TCGA-10-0933 | TCGA-OV | 63.2179847 | 79.3242638 |
| TCGA-10-0936 | TCGA-OV | 35.7820255 | 11.2302181 |
| TCGA-10-0937 | TCGA-OV | 53.7303281 | 35.7753765 |
| TCGA-10-0938 | TCGA-OV | 65.5018629 | 64.3305696 |
| TCGA-12-0616 | TCGA-GBM | 49.2203224 | 35.4081573 |
| TCGA-12-0618 | TCGA-GBM | 42.8697388 | 17.1830664 |
| TCGA-12-0619 | TCGA-GBM | 71.3736682 | 15.6560923 |
| TCGA-12-0821 | TCGA-GBM | 77.2098517 | 37.1754475 |
| TCGA-12-1597 | TCGA-GBM | 28.3498072 | 15.8086608 |

| | | | |
|---|---|---|---|
| TCGA-12-3650 | TCGA-GBM | 38.5346526 | 34.8907948 |
| TCGA-12-3652 | TCGA-GBM | 46.8211666 | 32.6790056 |
| TCGA-12-3653 | TCGA-GBM | 68.4439196 | 41.2080391 |
| TCGA-12-5295 | TCGA-GBM | 62.4605157 | 35.3252356 |
| TCGA-12-5299 | TCGA-GBM | 63.5955579 | 28.3887652 |
| TCGA-13-0714 | TCGA-OV | 12.5089722 | 3.15187263 |
| TCGA-13-0720 | TCGA-OV | 42.8442419 | 15.350436 |
| TCGA-13-0724 | TCGA-OV | 54.8561789 | 16.7839118 |
| TCGA-13-0725 | TCGA-OV | 84.225481 | 36.9231333 |
| TCGA-13-0726 | TCGA-OV | 63.6517044 | 21.7279077 |
| TCGA-13-0727 | TCGA-OV | 41.0833622 | 13.7993578 |
| TCGA-13-0730 | TCGA-OV | 49.3843305 | 53.175693 |
| TCGA-13-0762 | TCGA-OV | 60.3864768 | 34.1236348 |
| TCGA-13-0765 | TCGA-OV | 60.3781131 | 15.7842477 |
| TCGA-13-0766 | TCGA-OV | 39.0030106 | 15.4984798 |
| TCGA-13-0768 | TCGA-OV | 81.7012989 | 18.0086521 |
| TCGA-13-0795 | TCGA-OV | 50.452011 | 26.793374 |
| TCGA-13-0797 | TCGA-OV | 83.6948235 | 26.614113 |
| TCGA-13-0800 | TCGA-OV | 84.2103968 | 24.4385616 |
| TCGA-13-0804 | TCGA-OV | 69.8760683 | 29.678073 |
| TCGA-13-0883 | TCGA-OV | 31.2778323 | 30.5415547 |
| TCGA-13-0884 | TCGA-OV | 83.8718264 | 28.287259 |
| TCGA-13-0885 | TCGA-OV | 37.5478055 | 20.7354117 |
| TCGA-13-0886 | TCGA-OV | 63.1192718 | 21.5183001 |
| TCGA-13-0887 | TCGA-OV | 74.4675528 | 38.4205692 |
| TCGA-13-0888 | TCGA-OV | 58.9891348 | 22.2111176 |
| TCGA-13-0891 | TCGA-OV | 83.1698978 | 77.1940929 |
| TCGA-13-0893 | TCGA-OV | 151.985058 | 31.311576 |
| TCGA-13-0897 | TCGA-OV | 70.0885547 | 36.9951379 |
| TCGA-13-0900 | TCGA-OV | 61.1044257 | 57.9951744 |
| TCGA-13-0901 | TCGA-OV | 72.8578779 | 98.8430495 |

| | | | |
|---|---|---|---|
| TCGA-13-0905 | TCGA-OV | 51.797134 | 65.3815773 |
| TCGA-13-0906 | TCGA-OV | 169.374195 | 58.9860302 |
| TCGA-13-0908 | TCGA-OV | 51.8325645 | 18.0602115 |
| TCGA-13-0911 | TCGA-OV | 100.67157 | 27.4003479 |
| TCGA-13-0913 | TCGA-OV | 66.0636038 | 40.072821 |
| TCGA-13-0916 | TCGA-OV | 124.802541 | 54.9696349 |
| TCGA-13-0920 | TCGA-OV | 163.915046 | 79.575497 |
| TCGA-13-0923 | TCGA-OV | 78.9164466 | 40.936029 |
| TCGA-13-0924 | TCGA-OV | 105.47948 | 53.6333802 |
| TCGA-13-1403 | TCGA-OV | 36.2298582 | 13.0037283 |
| TCGA-13-1404 | TCGA-OV | 63.4359297 | 26.4850643 |
| TCGA-13-1405 | TCGA-OV | 72.2394285 | 29.1894593 |
| TCGA-13-1407 | TCGA-OV | 161.742118 | 40.6507036 |
| TCGA-13-1408 | TCGA-OV | 55.4568669 | 19.0595034 |
| TCGA-13-1409 | TCGA-OV | 76.8677792 | 23.7044208 |
| TCGA-13-1410 | TCGA-OV | 88.1664324 | 24.3864302 |
| TCGA-13-1411 | TCGA-OV | 44.795243 | 36.0199597 |
| TCGA-13-1477 | TCGA-OV | 25.3158142 | 25.778521 |
| TCGA-13-1483 | TCGA-OV | 76.5074127 | 23.8139118 |
| TCGA-13-1485 | TCGA-OV | 37.9641191 | 35.7687419 |
| TCGA-13-1487 | TCGA-OV | 81.0837758 | 24.7600367 |
| TCGA-13-1488 | TCGA-OV | 120.288672 | 36.3874056 |
| TCGA-13-1489 | TCGA-OV | 71.3279442 | 89.7964906 |
| TCGA-13-1492 | TCGA-OV | 142.453053 | 37.7115475 |
| TCGA-13-1495 | TCGA-OV | 74.2619383 | 31.9881505 |
| TCGA-13-1496 | TCGA-OV | 102.393763 | 42.8052777 |
| TCGA-13-1497 | TCGA-OV | 113.561891 | 39.1825892 |
| TCGA-13-1498 | TCGA-OV | 141.281837 | 45.3390064 |
| TCGA-13-1499 | TCGA-OV | 89.331003 | 50.2343809 |
| TCGA-13-1501 | TCGA-OV | 49.784989 | 73.9186194 |
| TCGA-13-1505 | TCGA-OV | 54.0984056 | 34.739415 |

| | | | |
|---|---|---|---|
| TCGA-13-1506 | TCGA-OV | 139.537539 | 61.9133403 |
| TCGA-13-1507 | TCGA-OV | 50.0897564 | 29.1550628 |
| TCGA-13-1509 | TCGA-OV | 93.2689292 | 28.2865864 |
| TCGA-13-1510 | TCGA-OV | 214.221823 | 43.8536581 |
| TCGA-13-1511 | TCGA-OV | 45.941419 | 29.3513908 |
| TCGA-13-1512 | TCGA-OV | 97.0835882 | 36.6676874 |
| TCGA-13-2060 | TCGA-OV | 77.8427855 | 41.3379677 |
| TCGA-13-A5FT | TCGA-OV | 139.669071 | 35.5867563 |
| TCGA-14-0736 | TCGA-GBM | 42.8473695 | 25.7044471 |
| TCGA-14-0781 | TCGA-GBM | 37.660733 | 18.088428 |
| TCGA-14-0787 | TCGA-GBM | 81.7128399 | 39.0070485 |
| TCGA-14-0789 | TCGA-GBM | 42.1229315 | 11.4573231 |
| TCGA-14-0790 | TCGA-GBM | 65.5844327 | 42.8295838 |
| TCGA-14-0817 | TCGA-GBM | 85.9827204 | 25.2874574 |
| TCGA-14-0871 | TCGA-GBM | 121.13329 | 20.9334582 |
| TCGA-14-1034 | TCGA-GBM | 127.877833 | 53.6319451 |
| TCGA-14-1402 | TCGA-GBM | 56.5976521 | 33.2564266 |
| TCGA-14-1823 | TCGA-GBM | 63.8286633 | 57.9685554 |
| TCGA-14-1825 | TCGA-GBM | 59.9553594 | 43.5735144 |
| TCGA-14-1829 | TCGA-GBM | 47.9334034 | 43.8666968 |
| TCGA-14-2554 | TCGA-GBM | 61.2894065 | 30.9660197 |
| TCGA-15-0742 | TCGA-GBM | 64.2954104 | 29.0554546 |
| TCGA-15-1444 | TCGA-GBM | 41.0221327 | 17.078489 |
| TCGA-16-0846 | TCGA-GBM | 43.7499135 | 18.0174279 |
| TCGA-16-1045 | TCGA-GBM | 45.0877319 | 19.951586 |
| TCGA-18-3406 | TCGA-LUSC | 33.7367576 | 31.377949 |
| TCGA-18-3407 | TCGA-LUSC | 14.4892596 | 3.18970703 |
| TCGA-18-3408 | TCGA-LUSC | 10.8604093 | 49.398043 |
| TCGA-18-3409 | TCGA-LUSC | 20.7848374 | 15.6778938 |
| TCGA-18-3410 | TCGA-LUSC | 86.3704039 | 74.5714898 |
| TCGA-18-3411 | TCGA-LUSC | 25.4212322 | 8.30644956 |

| | | | |
|---|---|---|---|
| TCGA-18-3412 | TCGA-LUSC | 45.1361793 | 35.0744873 |
| TCGA-18-3414 | TCGA-LUSC | 71.6866952 | 36.4719573 |
| TCGA-18-3415 | TCGA-LUSC | 15.9407533 | 9.15508528 |
| TCGA-18-3416 | TCGA-LUSC | 76.1192039 | 22.7941148 |
| TCGA-18-3417 | TCGA-LUSC | 27.9568003 | 18.6804541 |
| TCGA-18-3419 | TCGA-LUSC | 16.6210236 | 12.5289709 |
| TCGA-18-3421 | TCGA-LUSC | 67.3909919 | 49.8037951 |
| TCGA-18-4083 | TCGA-LUSC | 60.0357909 | 24.9498268 |
| TCGA-18-4086 | TCGA-LUSC | 33.6310914 | 12.4319087 |
| TCGA-18-4721 | TCGA-LUSC | 7.77954039 | 5.86598809 |
| TCGA-18-5592 | TCGA-LUSC | 32.2345229 | 23.1989608 |
| TCGA-18-5595 | TCGA-LUSC | 43.3195758 | 19.7507139 |
| TCGA-19-0957 | TCGA-GBM | 32.7149942 | 17.3123742 |
| TCGA-19-1389 | TCGA-GBM | 62.1669042 | 26.4008135 |
| TCGA-19-1390 | TCGA-GBM | 25.22206 | 25.650955 |
| TCGA-19-1787 | TCGA-GBM | 80.4077945 | 38.171558 |
| TCGA-19-2619 | TCGA-GBM | 48.9038435 | 24.8710914 |
| TCGA-19-2620 | TCGA-GBM | 45.3727042 | 16.0844768 |
| TCGA-19-2624 | TCGA-GBM | 55.912457 | 30.0841802 |
| TCGA-19-2625 | TCGA-GBM | 75.2406081 | 19.8231392 |
| TCGA-19-2629 | TCGA-GBM | 63.7929138 | 23.4888838 |
| TCGA-19-4065 | TCGA-GBM | 62.1065273 | 27.261933 |
| TCGA-19-5960 | TCGA-GBM | 78.4333842 | 20.7321267 |
| TCGA-20-0987 | TCGA-OV | 66.7846304 | 18.9135029 |
| TCGA-20-0991 | TCGA-OV | 38.2001971 | 26.772583 |
| TCGA-20-1682 | TCGA-OV | 90.9013753 | 100.847431 |
| TCGA-20-1683 | TCGA-OV | 58.1039337 | 12.9911826 |
| TCGA-20-1686 | TCGA-OV | 58.236095 | 47.5347295 |
| TCGA-20-1687 | TCGA-OV | 26.5857731 | 13.76224 |
| TCGA-21-1070 | TCGA-LUSC | 55.094374 | 47.5201355 |
| TCGA-21-1071 | TCGA-LUSC | 64.6550104 | 40.7361451 |

| | | | |
|---|---|---|---|
| TCGA-21-1072 | TCGA-LUSC | 64.338833 | 17.634192 |
| TCGA-21-1075 | TCGA-LUSC | 50.2371434 | 52.5407928 |
| TCGA-21-1076 | TCGA-LUSC | 56.456267 | 14.324862 |
| TCGA-21-1077 | TCGA-LUSC | 60.0679356 | 17.7152832 |
| TCGA-21-1078 | TCGA-LUSC | 55.8254856 | 13.8315419 |
| TCGA-21-1079 | TCGA-LUSC | 24.137689 | 8.7075311 |
| TCGA-21-1080 | TCGA-LUSC | 69.7936301 | 20.446556 |
| TCGA-21-1081 | TCGA-LUSC | 148.912593 | 7.91390627 |
| TCGA-21-1082 | TCGA-LUSC | 37.7727895 | 3.18183802 |
| TCGA-21-1083 | TCGA-LUSC | 72.25259 | 14.7519609 |
| TCGA-21-5782 | TCGA-LUSC | 35.4234487 | 16.2777624 |
| TCGA-21-5783 | TCGA-LUSC | 61.1726637 | 35.6823437 |
| TCGA-21-5784 | TCGA-LUSC | 38.0123274 | 14.1309226 |
| TCGA-21-5786 | TCGA-LUSC | 14.1988506 | 23.6798242 |
| TCGA-21-5787 | TCGA-LUSC | 67.6056541 | 45.6252381 |
| TCGA-21-A5DI | TCGA-LUSC | 70.5084088 | 115.820632 |
| TCGA-22-0940 | TCGA-LUSC | 45.3346009 | 43.0407447 |
| TCGA-22-0944 | TCGA-LUSC | 43.2580044 | 35.3807086 |
| TCGA-22-1000 | TCGA-LUSC | 39.2089975 | 16.3518613 |
| TCGA-22-1002 | TCGA-LUSC | 37.1591792 | 15.5593535 |
| TCGA-22-1005 | TCGA-LUSC | 20.9963024 | 8.21405988 |
| TCGA-22-1011 | TCGA-LUSC | 42.6669223 | 15.1827149 |
| TCGA-22-1012 | TCGA-LUSC | 52.5035849 | 36.2599608 |
| TCGA-22-1016 | TCGA-LUSC | 99.0134752 | 26.349492 |
| TCGA-22-1017 | TCGA-LUSC | 45.2593117 | 28.2891292 |
| TCGA-22-4591 | TCGA-LUSC | 35.8663377 | 18.7027556 |
| TCGA-22-4593 | TCGA-LUSC | 52.5980249 | 9.44163006 |
| TCGA-22-4594 | TCGA-LUSC | 30.911357 | 22.4044622 |
| TCGA-22-4595 | TCGA-LUSC | 25.392139 | 28.3314599 |
| TCGA-22-4596 | TCGA-LUSC | 27.0055369 | 20.1665882 |
| TCGA-22-4599 | TCGA-LUSC | 35.9261694 | 49.5783515 |

| | | | |
|---|---|---|---|
| TCGA-22-4601 | TCGA-LUSC | 85.8145796 | 22.9081854 |
| TCGA-22-4604 | TCGA-LUSC | 40.3062102 | 12.8479395 |
| TCGA-22-4605 | TCGA-LUSC | 38.4650074 | 21.3303466 |
| TCGA-22-4607 | TCGA-LUSC | 70.9462874 | 11.6205137 |
| TCGA-22-4609 | TCGA-LUSC | 23.5649003 | 11.3634438 |
| TCGA-22-4613 | TCGA-LUSC | 41.0086937 | 19.4046091 |
| TCGA-22-5471 | TCGA-LUSC | 38.8618796 | 51.5759077 |
| TCGA-22-5472 | TCGA-LUSC | 51.8004759 | 20.6007125 |
| TCGA-22-5473 | TCGA-LUSC | 27.8889252 | 11.1173769 |
| TCGA-22-5474 | TCGA-LUSC | 20.3074777 | 3.93679213 |
| TCGA-22-5477 | TCGA-LUSC | 45.4292143 | 57.6039884 |
| TCGA-22-5478 | TCGA-LUSC | 60.4814762 | 20.5061081 |
| TCGA-22-5479 | TCGA-LUSC | 39.3577693 | 28.6597507 |
| TCGA-22-5480 | TCGA-LUSC | 60.9590836 | 20.3237956 |
| TCGA-22-5481 | TCGA-LUSC | 75.8386502 | 20.2557014 |
| TCGA-22-5482 | TCGA-LUSC | 26.2160496 | 5.42673122 |
| TCGA-22-5483 | TCGA-LUSC | 99.0258318 | 35.465415 |
| TCGA-22-5485 | TCGA-LUSC | 71.367867 | 32.0534689 |
| TCGA-22-5489 | TCGA-LUSC | 19.5899464 | 9.78848543 |
| TCGA-22-5491 | TCGA-LUSC | 45.8468155 | 40.7810533 |
| TCGA-22-5492 | TCGA-LUSC | 25.791731 | 3.49616424 |
| TCGA-23-1021 | TCGA-OV | 34.8497989 | 24.5026461 |
| TCGA-23-1022 | TCGA-OV | 13.6139627 | 8.40919618 |
| TCGA-23-1023 | TCGA-OV | 60.1834846 | 24.734283 |
| TCGA-23-1024 | TCGA-OV | 54.9759354 | 34.6770315 |
| TCGA-23-1026 | TCGA-OV | 47.4369035 | 30.8745503 |
| TCGA-23-1027 | TCGA-OV | 90.394835 | 36.2269143 |
| TCGA-23-1028 | TCGA-OV | 63.0148372 | 21.4976068 |
| TCGA-23-1029 | TCGA-OV | 105.798131 | 29.0756865 |
| TCGA-23-1030 | TCGA-OV | 59.458719 | 25.9655764 |
| TCGA-23-1107 | TCGA-OV | 78.809178 | 63.2345461 |

| | | | |
|---|---|---|---|
| TCGA-23-1109 | TCGA-OV | 85.6934945 | 24.056156 |
| TCGA-23-1110 | TCGA-OV | 64.7657131 | 46.6032898 |
| TCGA-23-1111 | TCGA-OV | 58.5378571 | 22.7780193 |
| TCGA-23-1113 | TCGA-OV | 81.1407567 | 17.4942171 |
| TCGA-23-1114 | TCGA-OV | 31.8557925 | 32.6040009 |
| TCGA-23-1116 | TCGA-OV | 56.7394832 | 37.196116 |
| TCGA-23-1118 | TCGA-OV | 140.144609 | 52.4323407 |
| TCGA-23-1119 | TCGA-OV | 68.3314701 | 63.2756148 |
| TCGA-23-1120 | TCGA-OV | 89.1982913 | 26.3719978 |
| TCGA-23-1122 | TCGA-OV | 93.3310759 | 43.6699147 |
| TCGA-23-1123 | TCGA-OV | 100.464685 | 26.405827 |
| TCGA-23-1809 | TCGA-OV | 51.5873934 | 58.5147263 |
| TCGA-23-2077 | TCGA-OV | 54.165621 | 37.2075952 |
| TCGA-23-2078 | TCGA-OV | 65.2926061 | 24.7055737 |
| TCGA-23-2084 | TCGA-OV | 102.24991 | 29.6637843 |
| TCGA-24-0966 | TCGA-OV | 77.9788997 | 11.7976359 |
| TCGA-24-0968 | TCGA-OV | 56.0300934 | 24.134956 |
| TCGA-24-0970 | TCGA-OV | 81.2034968 | 31.5534754 |
| TCGA-24-0979 | TCGA-OV | 64.3160382 | 15.6829053 |
| TCGA-24-0982 | TCGA-OV | 47.0463649 | 51.0289535 |
| TCGA-24-1103 | TCGA-OV | 44.9839675 | 59.7893849 |
| TCGA-24-1104 | TCGA-OV | 64.6750675 | 23.6231468 |
| TCGA-24-1105 | TCGA-OV | 70.140011 | 19.4242558 |
| TCGA-24-1413 | TCGA-OV | 78.570552 | 30.0452707 |
| TCGA-24-1416 | TCGA-OV | 64.2052907 | 16.2747301 |
| TCGA-24-1417 | TCGA-OV | 121.361982 | 23.9745028 |
| TCGA-24-1418 | TCGA-OV | 94.2912134 | 37.4038837 |
| TCGA-24-1419 | TCGA-OV | 88.6157732 | 18.1091451 |
| TCGA-24-1422 | TCGA-OV | 84.4908812 | 35.1525407 |
| TCGA-24-1423 | TCGA-OV | 32.3598734 | 18.9822193 |
| TCGA-24-1424 | TCGA-OV | 70.7243798 | 54.0327799 |

| | | | |
|---|---|---|---|
| TCGA-24-1425 | TCGA-OV | 30.515041 | 13.2664953 |
| TCGA-24-1426 | TCGA-OV | 49.0661758 | 17.1379297 |
| TCGA-24-1427 | TCGA-OV | 31.5656889 | 19.4838281 |
| TCGA-24-1428 | TCGA-OV | 53.41467 | 19.2322903 |
| TCGA-24-1430 | TCGA-OV | 52.2717964 | 31.6505778 |
| TCGA-24-1431 | TCGA-OV | 39.590072 | 14.0881189 |
| TCGA-24-1434 | TCGA-OV | 52.1687919 | 30.8906675 |
| TCGA-24-1435 | TCGA-OV | 103.252846 | 18.6427231 |
| TCGA-24-1464 | TCGA-OV | 114.511492 | 48.0051937 |
| TCGA-24-1467 | TCGA-OV | 34.1436134 | 12.9232266 |
| TCGA-24-1469 | TCGA-OV | 91.4449727 | 49.1969867 |
| TCGA-24-1470 | TCGA-OV | 56.2868269 | 25.6326681 |
| TCGA-24-1471 | TCGA-OV | 54.0916118 | 6.96501638 |
| TCGA-24-1474 | TCGA-OV | 72.6230165 | 19.0372214 |
| TCGA-24-1544 | TCGA-OV | 67.3306035 | 43.7946568 |
| TCGA-24-1546 | TCGA-OV | 35.9450564 | 34.4830023 |
| TCGA-24-1549 | TCGA-OV | 48.4695983 | 24.968766 |
| TCGA-24-1550 | TCGA-OV | 71.2645096 | 28.3224682 |
| TCGA-24-1551 | TCGA-OV | 44.4203601 | 35.3898839 |
| TCGA-24-1552 | TCGA-OV | 64.0243995 | 18.287283 |
| TCGA-24-1553 | TCGA-OV | 30.9419779 | 17.7992343 |
| TCGA-24-1557 | TCGA-OV | 122.206411 | 30.1007723 |
| TCGA-24-1558 | TCGA-OV | 65.5360971 | 20.6177032 |
| TCGA-24-1560 | TCGA-OV | 59.9853988 | 20.4761199 |
| TCGA-24-1562 | TCGA-OV | 65.9860359 | 8.82136728 |
| TCGA-24-1563 | TCGA-OV | 49.1807258 | 44.1256326 |
| TCGA-24-1565 | TCGA-OV | 44.2919064 | 18.9130263 |
| TCGA-24-1567 | TCGA-OV | 91.6504602 | 45.568938 |
| TCGA-24-1603 | TCGA-OV | 64.0145345 | 43.2985822 |
| TCGA-24-1604 | TCGA-OV | 47.556606 | 41.7138801 |
| TCGA-24-1616 | TCGA-OV | 60.5147354 | 24.6178727 |

| | | | |
|---|---|---|---|
| TCGA-24-1842 | TCGA-OV | 48.064339 | 35.0844608 |
| TCGA-24-1843 | TCGA-OV | 106.099392 | 49.035945 |
| TCGA-24-1844 | TCGA-OV | 65.3831113 | 22.052457 |
| TCGA-24-1845 | TCGA-OV | 57.5789695 | 17.5753064 |
| TCGA-24-1846 | TCGA-OV | 64.7754167 | 22.2475293 |
| TCGA-24-1847 | TCGA-OV | 44.030827 | 14.4867779 |
| TCGA-24-1850 | TCGA-OV | 48.4928114 | 36.8482231 |
| TCGA-24-1923 | TCGA-OV | 55.3014107 | 20.7568989 |
| TCGA-24-1924 | TCGA-OV | 116.082745 | 32.5239169 |
| TCGA-24-1928 | TCGA-OV | 67.7742711 | 30.8980558 |
| TCGA-24-1930 | TCGA-OV | 45.1151402 | 23.4416199 |
| TCGA-24-2020 | TCGA-OV | 66.061346 | 51.6232992 |
| TCGA-24-2023 | TCGA-OV | 70.9898736 | 28.641278 |
| TCGA-24-2024 | TCGA-OV | 34.9661026 | 46.8771892 |
| TCGA-24-2026 | TCGA-OV | 49.07336 | 18.5261535 |
| TCGA-24-2027 | TCGA-OV | 88.2583264 | 31.453911 |
| TCGA-24-2033 | TCGA-OV | 37.2835674 | 22.0711173 |
| TCGA-24-2035 | TCGA-OV | 42.5270812 | 19.895856 |
| TCGA-24-2036 | TCGA-OV | 59.4265076 | 25.6440667 |
| TCGA-24-2038 | TCGA-OV | 37.3462204 | 21.1404615 |
| TCGA-24-2254 | TCGA-OV | 81.8561961 | 24.4563368 |
| TCGA-24-2261 | TCGA-OV | 47.854839 | 22.6543621 |
| TCGA-24-2262 | TCGA-OV | 82.568048 | 63.8257135 |
| TCGA-24-2267 | TCGA-OV | 44.1613998 | 20.6371657 |
| TCGA-24-2271 | TCGA-OV | 63.2970715 | 18.5818492 |
| TCGA-24-2280 | TCGA-OV | 108.918514 | 29.4852752 |
| TCGA-24-2281 | TCGA-OV | 51.5811882 | 30.7852201 |
| TCGA-24-2288 | TCGA-OV | 55.9423922 | 30.1796628 |
| TCGA-24-2289 | TCGA-OV | 28.3122025 | 29.8369625 |
| TCGA-24-2290 | TCGA-OV | 127.040413 | 27.4669918 |
| TCGA-24-2293 | TCGA-OV | 43.8740546 | 19.9839983 |

| | | | |
|---|---|---|---|
| TCGA-24-2297 | TCGA-OV | 47.6730496 | 20.8200397 |
| TCGA-24-2298 | TCGA-OV | 58.9942458 | 16.0334095 |
| TCGA-25-1312 | TCGA-OV | 62.154854 | 23.0067346 |
| TCGA-25-1313 | TCGA-OV | 96.1890964 | 29.8117743 |
| TCGA-25-1315 | TCGA-OV | 93.5719904 | 47.4942891 |
| TCGA-25-1316 | TCGA-OV | 71.8905177 | 36.2782573 |
| TCGA-25-1318 | TCGA-OV | 22.2018001 | 12.7356126 |
| TCGA-25-1319 | TCGA-OV | 67.6627512 | 12.3583481 |
| TCGA-25-1320 | TCGA-OV | 90.551816 | 25.2771609 |
| TCGA-25-1321 | TCGA-OV | 58.5447323 | 22.5434296 |
| TCGA-25-1322 | TCGA-OV | 62.4694726 | 24.5200911 |
| TCGA-25-1323 | TCGA-OV | 114.97468 | 38.3987439 |
| TCGA-25-1326 | TCGA-OV | 47.7982317 | 21.392039 |
| TCGA-25-1328 | TCGA-OV | 34.3624797 | 11.5878183 |
| TCGA-25-1329 | TCGA-OV | 59.0314583 | 14.683858 |
| TCGA-25-1623 | TCGA-OV | 35.7044807 | 16.8993795 |
| TCGA-25-1626 | TCGA-OV | 53.9097487 | 19.7050635 |
| TCGA-25-1627 | TCGA-OV | 125.492608 | 86.9664336 |
| TCGA-25-1628 | TCGA-OV | 67.2142297 | 25.5252115 |
| TCGA-25-1630 | TCGA-OV | 71.4506435 | 56.0978919 |
| TCGA-25-1631 | TCGA-OV | 80.9126461 | 24.7050096 |
| TCGA-25-1632 | TCGA-OV | 51.602569 | 14.7093556 |
| TCGA-25-1633 | TCGA-OV | 46.8014472 | 27.1275382 |
| TCGA-25-1634 | TCGA-OV | 64.1349225 | 59.0185875 |
| TCGA-25-1635 | TCGA-OV | 69.4052559 | 26.1414699 |
| TCGA-25-1870 | TCGA-OV | 56.7974742 | 26.2760045 |
| TCGA-25-1877 | TCGA-OV | 38.5924974 | 29.5190648 |
| TCGA-25-2042 | TCGA-OV | 30.6604112 | 23.3142858 |
| TCGA-25-2391 | TCGA-OV | 153.855225 | 29.6546305 |
| TCGA-25-2392 | TCGA-OV | 70.9552243 | 26.7395778 |
| TCGA-25-2393 | TCGA-OV | 112.917731 | 84.337884 |

| | | | |
|---|---|---|---|
| TCGA-25-2396 | TCGA-OV | 51.5923747 | 34.1345844 |
| TCGA-25-2398 | TCGA-OV | 58.9824798 | 26.403722 |
| TCGA-25-2399 | TCGA-OV | 69.3863945 | 24.6213153 |
| TCGA-25-2400 | TCGA-OV | 54.1555395 | 23.2880004 |
| TCGA-25-2401 | TCGA-OV | 86.6191438 | 29.6293237 |
| TCGA-25-2404 | TCGA-OV | 54.7257464 | 17.188659 |
| TCGA-25-2409 | TCGA-OV | 58.1399213 | 17.9011978 |
| TCGA-26-1442 | TCGA-GBM | 30.767252 | 25.7857277 |
| TCGA-26-5132 | TCGA-GBM | 76.8350962 | 15.5999388 |
| TCGA-26-5133 | TCGA-GBM | 77.4979313 | 22.213922 |
| TCGA-26-5134 | TCGA-GBM | 11.1291906 | 8.44316038 |
| TCGA-26-5135 | TCGA-GBM | 44.7868813 | 21.5527105 |
| TCGA-26-5136 | TCGA-GBM | 100.039219 | 24.5319163 |
| TCGA-26-5139 | TCGA-GBM | 63.3201439 | 23.2389394 |
| TCGA-27-1830 | TCGA-GBM | 50.8164894 | 24.0174014 |
| TCGA-27-1831 | TCGA-GBM | 51.2862714 | 43.1833906 |
| TCGA-27-1832 | TCGA-GBM | 67.5135126 | 36.739781 |
| TCGA-27-1834 | TCGA-GBM | 51.5389148 | 19.3744202 |
| TCGA-27-1835 | TCGA-GBM | 64.3715204 | 27.091633 |
| TCGA-27-1837 | TCGA-GBM | 43.0259043 | 25.1346762 |
| TCGA-27-2519 | TCGA-GBM | 73.0678526 | 35.8875647 |
| TCGA-27-2521 | TCGA-GBM | 72.8044107 | 20.2612195 |
| TCGA-27-2523 | TCGA-GBM | 72.7437879 | 44.707098 |
| TCGA-27-2524 | TCGA-GBM | 75.646846 | 34.0004831 |
| TCGA-27-2526 | TCGA-GBM | 56.6054906 | 38.3467202 |
| TCGA-27-2528 | TCGA-GBM | 52.3420973 | 26.3197689 |
| TCGA-28-1747 | TCGA-GBM | 59.3330031 | 26.5697951 |
| TCGA-28-1753 | TCGA-GBM | 49.6500946 | 26.2639571 |
| TCGA-28-2499 | TCGA-GBM | 67.7979756 | 20.3516272 |
| TCGA-28-2509 | TCGA-GBM | 86.6016559 | 24.7576966 |
| TCGA-28-2510 | TCGA-GBM | 37.451419 | 14.9753959 |

| | | | |
|---|---|---|---|
| TCGA-28-2513 | TCGA-GBM | 46.9360217 | 18.6040134 |
| TCGA-28-2514 | TCGA-GBM | 46.6053748 | 12.0369717 |
| TCGA-28-5204 | TCGA-GBM | 87.3933118 | 16.7484499 |
| TCGA-28-5207 | TCGA-GBM | 83.4782837 | 31.3423839 |
| TCGA-28-5208 | TCGA-GBM | 84.7626531 | 46.7552754 |
| TCGA-28-5209 | TCGA-GBM | 81.2298824 | 25.9216725 |
| TCGA-28-5213 | TCGA-GBM | 47.9389097 | 28.5942931 |
| TCGA-28-5215 | TCGA-GBM | 54.4488808 | 30.1855144 |
| TCGA-28-5216 | TCGA-GBM | 68.9102337 | 26.9711957 |
| TCGA-28-5218 | TCGA-GBM | 59.4779003 | 48.3274241 |
| TCGA-28-5220 | TCGA-GBM | 76.5058914 | 35.985228 |
| TCGA-29-1688 | TCGA-OV | 67.309058 | 30.7075322 |
| TCGA-29-1690 | TCGA-OV | 19.3823852 | 30.2262547 |
| TCGA-29-1691 | TCGA-OV | 106.883984 | 55.7964431 |
| TCGA-29-1693 | TCGA-OV | 51.522819 | 41.2263427 |
| TCGA-29-1694 | TCGA-OV | 45.5861485 | 22.3736963 |
| TCGA-29-1695 | TCGA-OV | 62.3804808 | 75.2938012 |
| TCGA-29-1696 | TCGA-OV | 60.5560699 | 26.3854841 |
| TCGA-29-1697 | TCGA-OV | 80.5924078 | 37.3983095 |
| TCGA-29-1701 | TCGA-OV | 96.1447553 | 60.6589697 |
| TCGA-29-1703 | TCGA-OV | 77.1136006 | 13.9653221 |
| TCGA-29-1705 | TCGA-OV | 73.8494007 | 33.4996793 |
| TCGA-29-1707 | TCGA-OV | 53.5077249 | 16.5541163 |
| TCGA-29-1710 | TCGA-OV | 44.3808416 | 25.4919396 |
| TCGA-29-1711 | TCGA-OV | 75.4170353 | 10.8708072 |
| TCGA-29-1761 | TCGA-OV | 87.3325045 | 73.2578658 |
| TCGA-29-1762 | TCGA-OV | 178.787351 | 59.8325327 |
| TCGA-29-1763 | TCGA-OV | 70.9914694 | 40.405009 |
| TCGA-29-1766 | TCGA-OV | 85.8664734 | 38.3161305 |
| TCGA-29-1768 | TCGA-OV | 63.304784 | 41.3729205 |
| TCGA-29-1769 | TCGA-OV | 78.4229289 | 143.15677 |

| | | | |
|---|---|---|---|
| TCGA-29-1770 | TCGA-OV | 76.7982712 | 42.3760836 |
| TCGA-29-1774 | TCGA-OV | 50.0484854 | 46.2633798 |
| TCGA-29-1776 | TCGA-OV | 49.1673864 | 13.8760066 |
| TCGA-29-1777 | TCGA-OV | 55.8876361 | 42.9491975 |
| TCGA-29-1778 | TCGA-OV | 45.1672885 | 26.9788776 |
| TCGA-29-1781 | TCGA-OV | 84.5583806 | 27.3079398 |
| TCGA-29-1783 | TCGA-OV | 74.0551431 | 20.2849109 |
| TCGA-29-1784 | TCGA-OV | 140.358111 | 28.9424111 |
| TCGA-29-1785 | TCGA-OV | 93.9292286 | 33.8793544 |
| TCGA-29-2414 | TCGA-OV | 42.1383781 | 27.1816752 |
| TCGA-29-2425 | TCGA-OV | 47.4826001 | 25.0299102 |
| TCGA-29-2427 | TCGA-OV | 37.3641043 | 18.2471701 |
| TCGA-29-2428 | TCGA-OV | 67.3114682 | 30.250544 |
| TCGA-29-A5NZ | TCGA-OV | 60.3852711 | 24.7588587 |
| TCGA-2A-A8VL | TCGA-PRAD | 61.0097292 | 21.8966466 |
| TCGA-2A-A8VO | TCGA-PRAD | 39.0636728 | 16.9889531 |
| TCGA-2A-A8VT | TCGA-PRAD | 64.1329662 | 12.9500488 |
| TCGA-2A-A8VV | TCGA-PRAD | 32.3166674 | 15.5277514 |
| TCGA-2A-A8VX | TCGA-PRAD | 47.3367562 | 20.9073872 |
| TCGA-2A-A8W1 | TCGA-PRAD | 42.6799121 | 23.1568424 |
| TCGA-2A-A8W3 | TCGA-PRAD | 41.8813416 | 8.87762067 |
| TCGA-2A-AAYF | TCGA-PRAD | 43.3429735 | 23.2102376 |
| TCGA-2A-AAYO | TCGA-PRAD | 48.841891 | 16.3783314 |
| TCGA-2A-AAYU | TCGA-PRAD | 63.0652329 | 13.5288569 |
| TCGA-2E-A9G8 | TCGA-UCEC | 47.9432274 | 24.3173865 |
| TCGA-2F-A9KO | TCGA-BLCA | 57.2514024 | 41.9890469 |
| TCGA-2F-A9KP | TCGA-BLCA | 83.9691821 | 33.7146784 |
| TCGA-2F-A9KQ | TCGA-BLCA | 81.6537555 | 17.1063727 |
| TCGA-2F-A9KR | TCGA-BLCA | 62.6256839 | 32.1088001 |
| TCGA-2F-A9KT | TCGA-BLCA | 94.0016739 | 21.3956084 |
| TCGA-2F-A9KW | TCGA-BLCA | 123.18377 | 40.3175571 |

| | | | |
|---|---|---|---|
| TCGA-2G-AAEW | TCGA-TGCT | 30.7220584 | 56.9228518 |
| TCGA-2G-AAEX | TCGA-TGCT | 48.4400266 | 88.7882045 |
| TCGA-2G-AAF1 | TCGA-TGCT | 45.2294271 | 44.2610086 |
| TCGA-2G-AAF6 | TCGA-TGCT | 50.1899463 | 40.3153143 |
| TCGA-2G-AAF8 | TCGA-TGCT | 58.084174 | 57.9024142 |
| TCGA-2G-AAFE | TCGA-TGCT | 41.9502096 | 63.622668 |
| TCGA-2G-AAFG | TCGA-TGCT | 70.8511215 | 69.6152344 |
| TCGA-2G-AAFH | TCGA-TGCT | 41.7765277 | 38.7936674 |
| TCGA-2G-AAFI | TCGA-TGCT | 102.860591 | 89.7623734 |
| TCGA-2G-AAFJ | TCGA-TGCT | 73.1084025 | 62.0629892 |
| TCGA-2G-AAFL | TCGA-TGCT | 63.8258089 | 14.2226516 |
| TCGA-2G-AAFM | TCGA-TGCT | 142.496592 | 131.412119 |
| TCGA-2G-AAFN | TCGA-TGCT | 81.1744917 | 60.6476529 |
| TCGA-2G-AAFO | TCGA-TGCT | 112.347575 | 20.7490269 |
| TCGA-2G-AAFV | TCGA-TGCT | 24.3952232 | 20.4728916 |
| TCGA-2G-AAFY | TCGA-TGCT | 189.983665 | 151.699572 |
| TCGA-2G-AAFZ | TCGA-TGCT | 69.283605 | 73.5531823 |
| TCGA-2G-AAG0 | TCGA-TGCT | 158.522438 | 132.810931 |
| TCGA-2G-AAG3 | TCGA-TGCT | 36.5240359 | 61.2246738 |
| TCGA-2G-AAG5 | TCGA-TGCT | 36.4498258 | 56.3050226 |
| TCGA-2G-AAG6 | TCGA-TGCT | 47.4952005 | 186.553629 |
| TCGA-2G-AAG7 | TCGA-TGCT | 104.6295 | 68.5300624 |
| TCGA-2G-AAG8 | TCGA-TGCT | 90.0067848 | 65.0138476 |
| TCGA-2G-AAG9 | TCGA-TGCT | 32.2272447 | 41.506136 |
| TCGA-2G-AAGA | TCGA-TGCT | 156.777416 | 137.321785 |
| TCGA-2G-AAGC | TCGA-TGCT | 135.853028 | 104.097351 |
| TCGA-2G-AAGE | TCGA-TGCT | 52.2087454 | 63.9761227 |
| TCGA-2G-AAGF | TCGA-TGCT | 100.787669 | 61.48767 |
| TCGA-2G-AAGG | TCGA-TGCT | 44.777208 | 36.7630481 |
| TCGA-2G-AAGI | TCGA-TGCT | 65.4081606 | 74.9736733 |
| TCGA-2G-AAGJ | TCGA-TGCT | 33.3988699 | 38.9040274 |

| | | | |
|---|---|---|---|
| TCGA-2G-AAGK | TCGA-TGCT | 112.298052 | 145.27529 |
| TCGA-2G-AAGM | TCGA-TGCT | 91.1329928 | 79.7202058 |
| TCGA-2G-AAGN | TCGA-TGCT | 149.053088 | 184.845658 |
| TCGA-2G-AAGC | TCGA-TGCT | 131.830771 | 102.796037 |
| TCGA-2G-AAGP | TCGA-TGCT | 71.3585017 | 58.6026436 |
| TCGA-2G-AAGS | TCGA-TGCT | 104.661456 | 76.6021302 |
| TCGA-2G-AAGT | TCGA-TGCT | 136.671503 | 24.4383373 |
| TCGA-2G-AAGV | TCGA-TGCT | 49.7646184 | 38.2216657 |
| TCGA-2G-AAGW | TCGA-TGCT | 117.364257 | 104.870288 |
| TCGA-2G-AAGX | TCGA-TGCT | 67.8319503 | 49.4289846 |
| TCGA-2G-AAGY | TCGA-TGCT | 95.4857765 | 18.2015453 |
| TCGA-2G-AAGZ | TCGA-TGCT | 90.2187647 | 17.8159132 |
| TCGA-2G-AAH0 | TCGA-TGCT | 59.5461151 | 90.2973286 |
| TCGA-2G-AAH2 | TCGA-TGCT | 50.4313853 | 37.6191744 |
| TCGA-2G-AAH3 | TCGA-TGCT | 34.9572439 | 65.478543 |
| TCGA-2G-AAH4 | TCGA-TGCT | 85.6722811 | 67.4244775 |
| TCGA-2G-AAH8 | TCGA-TGCT | 48.6465407 | 32.6997616 |
| TCGA-2G-AAHA | TCGA-TGCT | 56.0706419 | 51.1347426 |
| TCGA-2G-AAHC | TCGA-TGCT | 54.5824434 | 39.2698704 |
| TCGA-2G-AAHG | TCGA-TGCT | 87.7042387 | 71.2828968 |
| TCGA-2G-AAHL | TCGA-TGCT | 37.7501267 | 58.3551669 |
| TCGA-2G-AAHN | TCGA-TGCT | 49.1726721 | 92.4662209 |
| TCGA-2G-AAHP | TCGA-TGCT | 50.2121161 | 68.0051688 |
| TCGA-2G-AAHT | TCGA-TGCT | 53.4174991 | 60.5873497 |
| TCGA-2G-AAKD | TCGA-TGCT | 138.352204 | 154.001803 |
| TCGA-2G-AAKG | TCGA-TGCT | 70.8830951 | 239.214419 |
| TCGA-2G-AAKH | TCGA-TGCT | 83.3934574 | 56.8679274 |
| TCGA-2G-AAKL | TCGA-TGCT | 155.977948 | 39.8591686 |
| TCGA-2G-AAKM | TCGA-TGCT | 123.550134 | 156.256766 |
| TCGA-2G-AAKO | TCGA-TGCT | 51.9675367 | 71.1699812 |
| TCGA-2G-AAL5 | TCGA-TGCT | 86.232215 | 38.5976436 |

| | | | |
|---|---|---|---|
| TCGA-2G-AAL7 | TCGA-TGCT | 74.9323092 | 13.8071387 |
| TCGA-2G-AALF | TCGA-TGCT | 67.511628 | 66.4119524 |
| TCGA-2G-AALG | TCGA-TGCT | 60.2428946 | 57.4706651 |
| TCGA-2G-AALN | TCGA-TGCT | 122.560945 | 102.611898 |
| TCGA-2G-AALO | TCGA-TGCT | 69.5606436 | 56.7351656 |
| TCGA-2G-AALP | TCGA-TGCT | 87.3001335 | 16.4280852 |
| TCGA-2G-AALQ | TCGA-TGCT | 68.3173423 | 71.3200272 |
| TCGA-2G-AALR | TCGA-TGCT | 132.345366 | 62.0490994 |
| TCGA-2G-AALS | TCGA-TGCT | 85.9404676 | 72.9259214 |
| TCGA-2G-AALT | TCGA-TGCT | 75.7165511 | 61.7267578 |
| TCGA-2G-AALW | TCGA-TGCT | 17.5533277 | 78.877138 |
| TCGA-2G-AALX | TCGA-TGCT | 44.426457 | 35.1691867 |
| TCGA-2G-AALY | TCGA-TGCT | 62.835715 | 98.1937182 |
| TCGA-2G-AALZ | TCGA-TGCT | 26.1887391 | 40.5747819 |
| TCGA-2G-AAM2 | TCGA-TGCT | 158.49513 | 192.80236 |
| TCGA-2G-AAM3 | TCGA-TGCT | 21.8844788 | 16.5819851 |
| TCGA-2G-AAM4 | TCGA-TGCT | 54.8676178 | 145.086702 |
| TCGA-2H-A9GF | TCGA-ESCA | 45.0392976 | 15.6446316 |
| TCGA-2H-A9GG | TCGA-ESCA | 25.8330572 | 13.4629297 |
| TCGA-2H-A9GH | TCGA-ESCA | 59.1752012 | 36.1672883 |
| TCGA-2H-A9GI | TCGA-ESCA | 28.5662709 | 24.4737076 |
| TCGA-2H-A9GJ | TCGA-ESCA | 33.2459772 | 17.8997506 |
| TCGA-2H-A9GK | TCGA-ESCA | 28.0533575 | 13.9026459 |
| TCGA-2H-A9GL | TCGA-ESCA | 44.3320226 | 30.5388244 |
| TCGA-2H-A9GM | TCGA-ESCA | 38.0812794 | 20.3189347 |
| TCGA-2H-A9GN | TCGA-ESCA | 55.2935613 | 24.5410581 |
| TCGA-2H-A9GO | TCGA-ESCA | 26.2151448 | 102.168652 |
| TCGA-2H-A9GQ | TCGA-ESCA | 41.830111 | 43.0983816 |
| TCGA-2H-A9GR | TCGA-ESCA | 27.9342272 | 16.9813539 |
| TCGA-2J-AAB1 | TCGA-PAAD | 46.5160571 | 8.55919228 |
| TCGA-2J-AAB4 | TCGA-PAAD | 47.694533 | 11.3831309 |

| | | | |
|---|---|---|---|
| TCGA-2J-AAB6 | TCGA-PAAD | 59.8885395 | 23.1778615 |
| TCGA-2J-AAB8 | TCGA-PAAD | 44.454028 | 30.4603814 |
| TCGA-2J-AAB9 | TCGA-PAAD | 31.5220567 | 13.2397026 |
| TCGA-2J-AABA | TCGA-PAAD | 42.4829829 | 16.2102054 |
| TCGA-2J-AABE | TCGA-PAAD | 46.5594571 | 12.1977758 |
| TCGA-2J-AABF | TCGA-PAAD | 45.8497446 | 28.9488124 |
| TCGA-2J-AABH | TCGA-PAAD | 23.6527222 | 14.0120084 |
| TCGA-2J-AABI | TCGA-PAAD | 35.8798747 | 35.4114504 |
| TCGA-2J-AABK | TCGA-PAAD | 50.5353171 | 17.1876604 |
| TCGA-2J-AABO | TCGA-PAAD | 48.1834488 | 20.3417995 |
| TCGA-2J-AABP | TCGA-PAAD | 85.8005167 | 33.1150968 |
| TCGA-2J-AABR | TCGA-PAAD | 43.7490285 | 17.2885449 |
| TCGA-2J-AABT | TCGA-PAAD | 36.7913027 | 12.2708303 |
| TCGA-2J-AABU | TCGA-PAAD | 63.476677 | 18.2072944 |
| TCGA-2J-AABV | TCGA-PAAD | 10.8917572 | 3.92582941 |
| TCGA-2K-A9WE | TCGA-KIRP | 34.016841 | 20.9105919 |
| TCGA-2L-AAQA | TCGA-PAAD | 36.6130245 | 22.408827 |
| TCGA-2L-AAQE | TCGA-PAAD | 58.8018828 | 12.7926717 |
| TCGA-2L-AAQI | TCGA-PAAD | 38.8838789 | 19.3911615 |
| TCGA-2L-AAQJ | TCGA-PAAD | 51.5656569 | 12.8699185 |
| TCGA-2L-AAQL | TCGA-PAAD | 54.799091 | 19.6384665 |
| TCGA-2L-AAQM | TCGA-PAAD | 38.3097138 | 12.5772742 |
| TCGA-2V-A95S | TCGA-LIHC | 53.7327972 | 15.5463807 |
| TCGA-2X-A9D5 | TCGA-TGCT | 16.2720825 | 12.4498538 |
| TCGA-2X-A9D6 | TCGA-TGCT | 35.6882208 | 37.7245455 |
| TCGA-2Y-A9GS | TCGA-LIHC | 55.4690089 | 12.6327908 |
| TCGA-2Y-A9GT | TCGA-LIHC | 27.9484122 | 12.3276956 |
| TCGA-2Y-A9GU | TCGA-LIHC | 24.2515155 | 10.7176129 |
| TCGA-2Y-A9GV | TCGA-LIHC | 30.6981207 | 8.61281065 |
| TCGA-2Y-A9GW | TCGA-LIHC | 36.3716577 | 11.0800556 |
| TCGA-2Y-A9GX | TCGA-LIHC | 34.9137232 | 10.4455005 |

| | | | |
|---|---|---|---|
| TCGA-2Y-A9GY | TCGA-LIHC | 61.8648995 | 21.0602719 |
| TCGA-2Y-A9GZ | TCGA-LIHC | 35.8917284 | 12.4303458 |
| TCGA-2Y-A9H0 | TCGA-LIHC | 56.522006 | 19.7343957 |
| TCGA-2Y-A9H1 | TCGA-LIHC | 18.3087838 | 11.3301693 |
| TCGA-2Y-A9H2 | TCGA-LIHC | 71.9086954 | 23.0215173 |
| TCGA-2Y-A9H3 | TCGA-LIHC | 19.9506628 | 15.6663164 |
| TCGA-2Y-A9H4 | TCGA-LIHC | 57.9277083 | 32.1725078 |
| TCGA-2Y-A9H5 | TCGA-LIHC | 25.3375644 | 23.1694822 |
| TCGA-2Y-A9H6 | TCGA-LIHC | 20.76124 | 25.8877923 |
| TCGA-2Y-A9H7 | TCGA-LIHC | 31.6518976 | 25.7020744 |
| TCGA-2Y-A9H8 | TCGA-LIHC | 40.8178735 | 10.1944821 |
| TCGA-2Y-A9H9 | TCGA-LIHC | 32.6888674 | 31.2285701 |
| TCGA-2Y-A9HA | TCGA-LIHC | 52.2424828 | 23.0362933 |
| TCGA-2Y-A9HB | TCGA-LIHC | 34.712316 | 12.1015851 |
| TCGA-2Z-A9J1 | TCGA-KIRP | 16.6794868 | 2.41299204 |
| TCGA-2Z-A9J2 | TCGA-KIRP | 20.1460706 | 9.85443192 |
| TCGA-2Z-A9J3 | TCGA-KIRP | 46.94835 | 20.1657301 |
| TCGA-2Z-A9J5 | TCGA-KIRP | 23.0954836 | 2.76175828 |
| TCGA-2Z-A9J6 | TCGA-KIRP | 37.1558419 | 14.4943261 |
| TCGA-2Z-A9J7 | TCGA-KIRP | 34.9229943 | 7.18243697 |
| TCGA-2Z-A9J8 | TCGA-KIRP | 48.1839209 | 22.2447409 |
| TCGA-2Z-A9J9 | TCGA-KIRP | 60.2192187 | 25.8919131 |
| TCGA-2Z-A9JD | TCGA-KIRP | 31.2630325 | 8.71251737 |
| TCGA-2Z-A9JE | TCGA-KIRP | 62.7868322 | 20.9371179 |
| TCGA-2Z-A9JG | TCGA-KIRP | 29.6907233 | 27.6716985 |
| TCGA-2Z-A9JI | TCGA-KIRP | 21.6421768 | 31.6769374 |
| TCGA-2Z-A9JJ | TCGA-KIRP | 16.9072764 | 8.76265701 |
| TCGA-2Z-A9JK | TCGA-KIRP | 22.4166375 | 7.43297119 |
| TCGA-2Z-A9JL | TCGA-KIRP | 42.4670018 | 31.8692632 |
| TCGA-2Z-A9JM | TCGA-KIRP | 54.1861017 | 26.9477988 |
| TCGA-2Z-A9JN | TCGA-KIRP | 24.6421075 | 6.57729822 |

| | | | |
|---|---|---|---|
| TCGA-2Z-A9JO | TCGA-KIRP | 27.0737444 | 10.2348526 |
| TCGA-2Z-A9JP | TCGA-KIRP | 24.2361588 | 24.9732522 |
| TCGA-2Z-A9JQ | TCGA-KIRP | 23.0970137 | 10.9549759 |
| TCGA-2Z-A9JR | TCGA-KIRP | 41.1066719 | 5.41477768 |
| TCGA-2Z-A9JS | TCGA-KIRP | 44.981589 | 17.7634307 |
| TCGA-2Z-A9JT | TCGA-KIRP | 11.4108289 | 7.50322189 |
| TCGA-30-1714 | TCGA-OV | 58.1858595 | 17.2127091 |
| TCGA-30-1718 | TCGA-OV | 101.754172 | 20.840662 |
| TCGA-30-1853 | TCGA-OV | 39.5959289 | 14.7619862 |
| TCGA-30-1857 | TCGA-OV | 82.6130922 | 24.5492074 |
| TCGA-30-1860 | TCGA-OV | 209.533846 | 112.640799 |
| TCGA-30-1861 | TCGA-OV | 37.8609176 | 14.1647441 |
| TCGA-30-1862 | TCGA-OV | 76.5172482 | 23.8990925 |
| TCGA-30-1866 | TCGA-OV | 100.099865 | 39.1480627 |
| TCGA-30-1891 | TCGA-OV | 82.2365525 | 40.9626183 |
| TCGA-30-1892 | TCGA-OV | 60.4562631 | 12.3504208 |
| TCGA-31-1944 | TCGA-OV | 57.2935088 | 21.3214148 |
| TCGA-31-1946 | TCGA-OV | 34.244256 | 20.8546471 |
| TCGA-31-1950 | TCGA-OV | 64.6076359 | 19.5492915 |
| TCGA-31-1951 | TCGA-OV | 38.0679299 | 19.559169 |
| TCGA-31-1953 | TCGA-OV | 74.1080587 | 30.5019562 |
| TCGA-31-1956 | TCGA-OV | 60.8778552 | 32.2509367 |
| TCGA-31-1959 | TCGA-OV | 75.2972374 | 37.4655684 |
| TCGA-32-1970 | TCGA-GBM | 57.8057686 | 24.0601738 |
| TCGA-32-1980 | TCGA-GBM | 46.7232883 | 20.3571691 |
| TCGA-32-1982 | TCGA-GBM | 47.7859433 | 20.3005585 |
| TCGA-32-2615 | TCGA-GBM | 54.32427 | 20.0853909 |
| TCGA-32-2616 | TCGA-GBM | 50.6047721 | 22.1899594 |
| TCGA-32-2632 | TCGA-GBM | 58.4288906 | 31.1375877 |
| TCGA-32-2634 | TCGA-GBM | 76.0741312 | 42.4165921 |
| TCGA-32-2638 | TCGA-GBM | 52.6809512 | 21.6946721 |

| | | | |
|---|---|---|---|
| TCGA-32-4213 | TCGA-GBM | 13.9509742 | 11.9954525 |
| TCGA-32-5222 | TCGA-GBM | 81.4919257 | 40.8175545 |
| TCGA-33-4532 | TCGA-LUSC | 23.8339335 | 43.4139702 |
| TCGA-33-4538 | TCGA-LUSC | 38.2520138 | 34.0116766 |
| TCGA-33-4547 | TCGA-LUSC | 44.4983922 | 10.1954362 |
| TCGA-33-4566 | TCGA-LUSC | 27.3550033 | 63.7525608 |
| TCGA-33-4582 | TCGA-LUSC | 85.6564557 | 58.7065101 |
| TCGA-33-4583 | TCGA-LUSC | 44.1718707 | 35.7925095 |
| TCGA-33-4586 | TCGA-LUSC | 45.7506636 | 48.3022786 |
| TCGA-33-4587 | TCGA-LUSC | 147.651145 | 56.0139051 |
| TCGA-33-6737 | TCGA-LUSC | 31.775567 | 59.9338475 |
| TCGA-33-6738 | TCGA-LUSC | 52.4474715 | 4.81844266 |
| TCGA-33-A4WN | TCGA-LUSC | 29.0219991 | 22.558921 |
| TCGA-33-A5GW | TCGA-LUSC | 66.9428212 | 28.2724052 |
| TCGA-33-AAS8 | TCGA-LUSC | 61.4358267 | 48.4463922 |
| TCGA-33-AASB | TCGA-LUSC | 51.3927037 | 33.7276988 |
| TCGA-33-AASD | TCGA-LUSC | 83.0123808 | 64.2223743 |
| TCGA-33-AASI | TCGA-LUSC | 43.8013554 | 57.1867157 |
| TCGA-33-AASJ | TCGA-LUSC | 109.326409 | 31.7987372 |
| TCGA-33-AASL | TCGA-LUSC | 48.6476185 | 55.3692093 |
| TCGA-34-2596 | TCGA-LUSC | 39.2118016 | 25.4422362 |
| TCGA-34-2600 | TCGA-LUSC | 58.8978519 | 24.138517 |
| TCGA-34-2608 | TCGA-LUSC | 28.3188984 | 17.5869359 |
| TCGA-34-5231 | TCGA-LUSC | 47.2792456 | 32.0608525 |
| TCGA-34-5232 | TCGA-LUSC | 27.3221976 | 3.68925553 |
| TCGA-34-5234 | TCGA-LUSC | 17.5914101 | 8.32195995 |
| TCGA-34-5236 | TCGA-LUSC | 42.556881 | 8.19930415 |
| TCGA-34-5239 | TCGA-LUSC | 42.5564817 | 23.4981892 |
| TCGA-34-5240 | TCGA-LUSC | 30.5626725 | 14.0427172 |
| TCGA-34-5241 | TCGA-LUSC | 59.9826708 | 59.7055579 |
| TCGA-34-5927 | TCGA-LUSC | 74.5277942 | 26.1722471 |

| | | | |
|---|---|---|---|
| TCGA-34-5928 | TCGA-LUSC | 35.4463034 | 6.31205504 |
| TCGA-34-5929 | TCGA-LUSC | 35.3861211 | 14.1154303 |
| TCGA-34-7107 | TCGA-LUSC | 39.5948594 | 26.5687395 |
| TCGA-34-8454 | TCGA-LUSC | 69.8622809 | 28.1241451 |
| TCGA-34-8455 | TCGA-LUSC | 75.2244843 | 21.5217103 |
| TCGA-34-8456 | TCGA-LUSC | 41.0423671 | 13.3023197 |
| TCGA-34-A5IX | TCGA-LUSC | 94.3622945 | 16.3883798 |
| TCGA-35-3615 | TCGA-LUAD | 35.2299416 | 29.9316782 |
| TCGA-35-4122 | TCGA-LUAD | 70.631454 | 62.7517961 |
| TCGA-35-4123 | TCGA-LUAD | 74.9372499 | 29.7880302 |
| TCGA-35-5375 | TCGA-LUAD | 65.4251022 | 31.5466252 |
| TCGA-36-1568 | TCGA-OV | 49.9996194 | 9.02206603 |
| TCGA-36-1569 | TCGA-OV | 34.3202459 | 21.62559 |
| TCGA-36-1570 | TCGA-OV | 57.4640049 | 25.6495422 |
| TCGA-36-1571 | TCGA-OV | 53.3201958 | 55.9004383 |
| TCGA-36-1574 | TCGA-OV | 96.0814607 | 32.7950265 |
| TCGA-36-1576 | TCGA-OV | 34.9728909 | 39.6045423 |
| TCGA-36-1577 | TCGA-OV | 197.463419 | 39.6569213 |
| TCGA-36-1580 | TCGA-OV | 42.8282277 | 34.2372817 |
| TCGA-36-1581 | TCGA-OV | 59.9697866 | 43.2742756 |
| TCGA-37-3783 | TCGA-LUSC | 36.9019828 | 26.7483288 |
| TCGA-37-3789 | TCGA-LUSC | 77.6544073 | 34.6896247 |
| TCGA-37-3792 | TCGA-LUSC | 91.5322876 | 67.525929 |
| TCGA-37-4129 | TCGA-LUSC | 51.7913873 | 17.2541491 |
| TCGA-37-4130 | TCGA-LUSC | 38.7344219 | 11.2147846 |
| TCGA-37-4132 | TCGA-LUSC | 56.7351477 | 24.1431869 |
| TCGA-37-4133 | TCGA-LUSC | 15.4374737 | 14.2416982 |
| TCGA-37-4135 | TCGA-LUSC | 71.7400088 | 65.0971721 |
| TCGA-37-4141 | TCGA-LUSC | 36.1163406 | 7.63830686 |
| TCGA-37-5819 | TCGA-LUSC | 44.9748102 | 63.1982867 |
| TCGA-37-A5EL | TCGA-LUSC | 39.877944 | 8.97086344 |

| | | | |
|---|---|---|---|
| TCGA-37-A5EM | TCGA-LUSC | 109.848206 | 39.023831 |
| TCGA-37-A5EN | TCGA-LUSC | 112.239333 | 36.2841134 |
| TCGA-38-4625 | TCGA-LUAD | 35.7556008 | 13.7016893 |
| TCGA-38-4626 | TCGA-LUAD | 40.6974846 | 17.7321747 |
| TCGA-38-4627 | TCGA-LUAD | 36.7995696 | 20.5488222 |
| TCGA-38-4628 | TCGA-LUAD | 39.2259437 | 23.1572913 |
| TCGA-38-4629 | TCGA-LUAD | 40.0268762 | 23.3561648 |
| TCGA-38-4630 | TCGA-LUAD | 102.7685 | 30.6292299 |
| TCGA-38-4631 | TCGA-LUAD | 63.1517985 | 37.4039567 |
| TCGA-38-4632 | TCGA-LUAD | 75.8472863 | 48.6013779 |
| TCGA-38-6178 | TCGA-LUAD | 62.1749278 | 14.7852656 |
| TCGA-38-7271 | TCGA-LUAD | 33.2573132 | 16.7601059 |
| TCGA-38-A44F | TCGA-LUAD | 42.8008463 | 20.3764352 |
| TCGA-39-5011 | TCGA-LUSC | 61.5690384 | 22.7591115 |
| TCGA-39-5016 | TCGA-LUSC | 49.2763167 | 25.426378 |
| TCGA-39-5019 | TCGA-LUSC | 42.3441046 | 30.0762622 |
| TCGA-39-5021 | TCGA-LUSC | 43.0235499 | 52.1753886 |
| TCGA-39-5022 | TCGA-LUSC | 23.4627237 | 11.4783713 |
| TCGA-39-5024 | TCGA-LUSC | 48.0416959 | 22.9753656 |
| TCGA-39-5027 | TCGA-LUSC | 22.8586498 | 7.35938764 |
| TCGA-39-5028 | TCGA-LUSC | 45.4737124 | 16.8092672 |
| TCGA-39-5029 | TCGA-LUSC | 23.6928436 | 10.0796638 |
| TCGA-39-5030 | TCGA-LUSC | 9.38891678 | 1.80768064 |
| TCGA-39-5031 | TCGA-LUSC | 48.0317255 | 20.2122843 |
| TCGA-39-5034 | TCGA-LUSC | 9.24070681 | 8.41702032 |
| TCGA-39-5035 | TCGA-LUSC | 89.7659265 | 20.6216217 |
| TCGA-39-5036 | TCGA-LUSC | 58.2386524 | 65.4754227 |
| TCGA-39-5037 | TCGA-LUSC | 61.9985611 | 43.2784166 |
| TCGA-39-5039 | TCGA-LUSC | 10.7704745 | 3.43545355 |
| TCGA-39-5040 | TCGA-LUSC | 101.081183 | 5.31839532 |
| TCGA-3A-A9I5 | TCGA-PAAD | 36.4388424 | 31.4085673 |

| | | | |
|---|---|---|---|
| TCGA-3A-A9I7 | TCGA-PAAD | 43.9052601 | 15.9911561 |
| TCGA-3A-A9I9 | TCGA-PAAD | 39.2005846 | 17.2387184 |
| TCGA-3A-A9IB | TCGA-PAAD | 46.9513983 | 20.2722552 |
| TCGA-3A-A9IC | TCGA-PAAD | 57.3343528 | 17.4942236 |
| TCGA-3A-A9IH | TCGA-PAAD | 42.3088442 | 14.3776817 |
| TCGA-3A-A9IJ | TCGA-PAAD | 50.7452397 | 7.77174518 |
| TCGA-3A-A9IL | TCGA-PAAD | 29.8678453 | 16.2345052 |
| TCGA-3A-A9IN | TCGA-PAAD | 24.8706255 | 11.0583245 |
| TCGA-3A-A9IO | TCGA-PAAD | 40.245259 | 18.461615 |
| TCGA-3A-A9IR | TCGA-PAAD | 55.8159619 | 14.4825693 |
| TCGA-3A-A9IS | TCGA-PAAD | 41.3723125 | 27.2887793 |
| TCGA-3A-A9IU | TCGA-PAAD | 59.2071925 | 22.0339518 |
| TCGA-3A-A9IV | TCGA-PAAD | 33.1712959 | 16.2510188 |
| TCGA-3A-A9IX | TCGA-PAAD | 35.8457246 | 16.6533562 |
| TCGA-3A-A9IZ | TCGA-PAAD | 57.9873815 | 17.1947547 |
| TCGA-3A-A9J0 | TCGA-PAAD | 59.7138344 | 12.4511457 |
| TCGA-3C-AAAU | TCGA-BRCA | 53.1312722 | 40.2702976 |
| TCGA-3C-AALI | TCGA-BRCA | 33.7458151 | 26.4040957 |
| TCGA-3C-AALJ | TCGA-BRCA | 59.4271547 | 51.486805 |
| TCGA-3C-AALK | TCGA-BRCA | 53.1201688 | 25.302053 |
| TCGA-3E-AAAY | TCGA-PAAD | 59.8108768 | 16.0600717 |
| TCGA-3E-AAAZ | TCGA-PAAD | 36.508193 | 15.5342897 |
| TCGA-3K-AAZ8 | TCGA-LIHC | 42.1828736 | 21.4397949 |
| TCGA-3L-AA1B | TCGA-COAD | 65.0846853 | 26.9430002 |
| TCGA-3M-AB46 | TCGA-STAD | 71.5406421 | 23.452293 |
| TCGA-3M-AB47 | TCGA-STAD | 28.3779333 | 15.7210957 |
| TCGA-3N-A9WB | TCGA-SKCM | 78.9548351 | 51.2723308 |
| TCGA-3N-A9WC | TCGA-SKCM | 61.0446017 | 34.8698808 |
| TCGA-3N-A9WD | TCGA-SKCM | 42.5467564 | 22.5190223 |
| TCGA-3Z-A93Z | TCGA-KIRC | 59.4776783 | 22.3378461 |
| TCGA-41-2571 | TCGA-GBM | 37.5366635 | 18.4008837 |

| | | | |
|---|---|---|---|
| TCGA-41-2572 | TCGA-GBM | 37.2383593 | 21.4136483 |
| TCGA-41-3915 | TCGA-GBM | 36.6685321 | 16.5118833 |
| TCGA-41-4097 | TCGA-GBM | 40.7701634 | 14.3609181 |
| TCGA-41-5651 | TCGA-GBM | 73.0511948 | 29.3274706 |
| TCGA-43-2576 | TCGA-LUSC | 46.7758998 | 33.2625623 |
| TCGA-43-2578 | TCGA-LUSC | 80.9484386 | 52.1208865 |
| TCGA-43-2581 | TCGA-LUSC | 15.2491574 | 5.53539226 |
| TCGA-43-3394 | TCGA-LUSC | 39.9178843 | 68.3896094 |
| TCGA-43-3920 | TCGA-LUSC | 24.7282485 | 73.8518578 |
| TCGA-43-5668 | TCGA-LUSC | 19.9512604 | 30.4015747 |
| TCGA-43-5670 | TCGA-LUSC | 62.3176895 | 26.8727887 |
| TCGA-43-6143 | TCGA-LUSC | 18.5835941 | 49.2269384 |
| TCGA-43-6647 | TCGA-LUSC | 7.51544522 | 5.87558991 |
| TCGA-43-6770 | TCGA-LUSC | 66.0676181 | 32.0709226 |
| TCGA-43-6771 | TCGA-LUSC | 31.3843162 | 17.6291518 |
| TCGA-43-6773 | TCGA-LUSC | 15.1032503 | 10.6881275 |
| TCGA-43-7656 | TCGA-LUSC | 73.4493032 | 33.2139338 |
| TCGA-43-7657 | TCGA-LUSC | 45.7488477 | 17.7519811 |
| TCGA-43-7658 | TCGA-LUSC | 168.540235 | 74.0915598 |
| TCGA-43-8115 | TCGA-LUSC | 69.0031395 | 10.5166768 |
| TCGA-43-8116 | TCGA-LUSC | 79.0093856 | 47.9073364 |
| TCGA-43-8118 | TCGA-LUSC | 57.0754874 | 32.1409617 |
| TCGA-43-A474 | TCGA-LUSC | 49.755282 | 64.3645439 |
| TCGA-43-A475 | TCGA-LUSC | 23.9353966 | 11.7333124 |
| TCGA-43-A56U | TCGA-LUSC | 31.8185483 | 1.56063199 |
| TCGA-43-A56V | TCGA-LUSC | 58.7060822 | 100.288552 |
| TCGA-44-2655 | TCGA-LUAD | 52.1993752 | 23.0534856 |
| TCGA-44-2656 | TCGA-LUAD | 15.743381 | 7.10173194 |
| TCGA-44-2657 | TCGA-LUAD | 41.9015506 | 9.90137088 |
| TCGA-44-2659 | TCGA-LUAD | 36.632504 | 16.7235207 |
| TCGA-44-2661 | TCGA-LUAD | 35.9658335 | 10.1562485 |

| | | | |
|---|---|---|---|
| TCGA-44-2662 | TCGA-LUAD | 15.1983837 | 14.2507188 |
| TCGA-44-2665 | TCGA-LUAD | 36.60963 | 12.5125421 |
| TCGA-44-2666 | TCGA-LUAD | 43.3227097 | 16.3952354 |
| TCGA-44-2668 | TCGA-LUAD | 17.6046474 | 10.9285209 |
| TCGA-44-3396 | TCGA-LUAD | 36.0349231 | 11.4161294 |
| TCGA-44-3398 | TCGA-LUAD | 38.5739005 | 30.2575287 |
| TCGA-44-3917 | TCGA-LUAD | 19.7501933 | 10.7114939 |
| TCGA-44-3918 | TCGA-LUAD | 14.4517107 | 7.84308769 |
| TCGA-44-3919 | TCGA-LUAD | 52.9201037 | 17.0626015 |
| TCGA-44-4112 | TCGA-LUAD | 5.52720151 | 10.5088772 |
| TCGA-44-5643 | TCGA-LUAD | 48.1064372 | 77.4957075 |
| TCGA-44-5644 | TCGA-LUAD | 57.2339257 | 42.4833495 |
| TCGA-44-5645 | TCGA-LUAD | 45.7044939 | 21.9584149 |
| TCGA-44-6144 | TCGA-LUAD | 41.8421065 | 11.3219376 |
| TCGA-44-6145 | TCGA-LUAD | 62.8421618 | 35.6649064 |
| TCGA-44-6146 | TCGA-LUAD | 8.12068447 | 5.99411918 |
| TCGA-44-6147 | TCGA-LUAD | 60.2035869 | 14.6178274 |
| TCGA-44-6148 | TCGA-LUAD | 40.7111461 | 10.3021307 |
| TCGA-44-6774 | TCGA-LUAD | 37.2528191 | 13.8964376 |
| TCGA-44-6775 | TCGA-LUAD | 64.2498543 | 17.5119167 |
| TCGA-44-6776 | TCGA-LUAD | 46.774766 | 13.7512399 |
| TCGA-44-6777 | TCGA-LUAD | 36.2889004 | 16.0998064 |
| TCGA-44-6778 | TCGA-LUAD | 61.0347264 | 12.3846611 |
| TCGA-44-6779 | TCGA-LUAD | 60.8198457 | 24.5108441 |
| TCGA-44-7659 | TCGA-LUAD | 52.1950648 | 23.1495273 |
| TCGA-44-7660 | TCGA-LUAD | 385.398546 | 27.2393601 |
| TCGA-44-7661 | TCGA-LUAD | 54.3590552 | 26.3149713 |
| TCGA-44-7662 | TCGA-LUAD | 62.1794375 | 30.3931712 |
| TCGA-44-7667 | TCGA-LUAD | 126.719739 | 42.1487503 |
| TCGA-44-7669 | TCGA-LUAD | 88.6528484 | 59.8236005 |
| TCGA-44-7670 | TCGA-LUAD | 94.7995599 | 39.0992656 |

| | | | |
|---|---|---|---|
| TCGA-44-7671 | TCGA-LUAD | 36.912077 | 16.6314671 |
| TCGA-44-7672 | TCGA-LUAD | 28.891947 | 21.0014839 |
| TCGA-44-8117 | TCGA-LUAD | 46.7047427 | 13.3841167 |
| TCGA-44-8119 | TCGA-LUAD | 67.7685061 | 18.7967081 |
| TCGA-44-8120 | TCGA-LUAD | 63.0220511 | 21.2638555 |
| TCGA-44-A479 | TCGA-LUAD | 35.8184948 | 18.2271026 |
| TCGA-44-A47A | TCGA-LUAD | 54.9947554 | 26.5148179 |
| TCGA-44-A47B | TCGA-LUAD | 90.3147719 | 17.9138378 |
| TCGA-44-A47G | TCGA-LUAD | 30.5768682 | 14.7234763 |
| TCGA-44-A4SS | TCGA-LUAD | 42.3068786 | 23.3355899 |
| TCGA-44-A4SU | TCGA-LUAD | 43.8937631 | 10.8000146 |
| TCGA-46-3765 | TCGA-LUSC | 21.8197127 | 4.75712421 |
| TCGA-46-3766 | TCGA-LUSC | 25.2150546 | 15.632577 |
| TCGA-46-3767 | TCGA-LUSC | 36.4855329 | 9.44381785 |
| TCGA-46-3768 | TCGA-LUSC | 21.4856565 | 38.4687769 |
| TCGA-46-3769 | TCGA-LUSC | 55.9100276 | 56.2781994 |
| TCGA-46-6025 | TCGA-LUSC | 74.2305973 | 19.2161815 |
| TCGA-46-6026 | TCGA-LUSC | 42.387673 | 5.37694509 |
| TCGA-49-4486 | TCGA-LUAD | 37.9058917 | 24.2801172 |
| TCGA-49-4487 | TCGA-LUAD | 79.1053879 | 42.0431398 |
| TCGA-49-4488 | TCGA-LUAD | 79.0741357 | 24.0351486 |
| TCGA-49-4490 | TCGA-LUAD | 69.6202262 | 13.9649924 |
| TCGA-49-4494 | TCGA-LUAD | 43.8599881 | 39.3360755 |
| TCGA-49-4501 | TCGA-LUAD | 46.8491283 | 17.8954748 |
| TCGA-49-4505 | TCGA-LUAD | 48.5485787 | 27.1080357 |
| TCGA-49-4506 | TCGA-LUAD | 19.0953291 | 31.5720695 |
| TCGA-49-4507 | TCGA-LUAD | 68.8098882 | 41.9373904 |
| TCGA-49-4510 | TCGA-LUAD | 43.931962 | 19.8158069 |
| TCGA-49-4512 | TCGA-LUAD | 60.0881354 | 25.1013093 |
| TCGA-49-4514 | TCGA-LUAD | 68.7181809 | 41.6020449 |
| TCGA-49-6742 | TCGA-LUAD | 43.5615354 | 14.9979286 |

| | | | |
|---|---|---|---|
| TCGA-49-6743 | TCGA-LUAD | 43.1086858 | 14.9380423 |
| TCGA-49-6744 | TCGA-LUAD | 47.2990132 | 10.3020223 |
| TCGA-49-6745 | TCGA-LUAD | 53.8923214 | 16.0770562 |
| TCGA-49-6761 | TCGA-LUAD | 54.7182323 | 29.7611663 |
| TCGA-49-6767 | TCGA-LUAD | 105.884657 | 36.2347012 |
| TCGA-49-AAQV | TCGA-LUAD | 95.9333844 | 21.9078822 |
| TCGA-49-AAR0 | TCGA-LUAD | 82.3676741 | 18.5925373 |
| TCGA-49-AAR2 | TCGA-LUAD | 45.3488367 | 34.9516179 |
| TCGA-49-AAR3 | TCGA-LUAD | 51.5842745 | 27.6579906 |
| TCGA-49-AAR4 | TCGA-LUAD | 48.0683556 | 12.0297742 |
| TCGA-49-AAR9 | TCGA-LUAD | 74.1663096 | 31.1370759 |
| TCGA-49-AARE | TCGA-LUAD | 52.01656 | 27.3604709 |
| TCGA-49-AARN | TCGA-LUAD | 52.8231897 | 16.4269399 |
| TCGA-49-AARO | TCGA-LUAD | 47.9171555 | 18.85638 |
| TCGA-49-AARQ | TCGA-LUAD | 135.290449 | 15.5323574 |
| TCGA-49-AARR | TCGA-LUAD | 36.5625971 | 12.5464352 |
| TCGA-4A-A93W | TCGA-KIRP | 21.2054893 | 36.0176019 |
| TCGA-4A-A93X | TCGA-KIRP | 32.2104387 | 13.9413646 |
| TCGA-4A-A93Y | TCGA-KIRP | 21.0686888 | 9.19389076 |
| TCGA-4B-A93V | TCGA-LUAD | 41.2165889 | 23.9353081 |
| TCGA-4C-A93U | TCGA-THCA | 51.9890835 | 15.244276 |
| TCGA-4E-A92E | TCGA-UCEC | 36.3115774 | 13.4413467 |
| TCGA-4H-AAAK | TCGA-BRCA | 64.9189579 | 14.677463 |
| TCGA-4K-AA1G | TCGA-TGCT | 21.4228593 | 46.9263504 |
| TCGA-4K-AA1H | TCGA-TGCT | 51.5943109 | 23.5924171 |
| TCGA-4K-AA1I | TCGA-TGCT | 53.3823624 | 38.2433021 |
| TCGA-4K-AAAL | TCGA-TGCT | 46.6853216 | 56.203418 |
| TCGA-4L-AA1F | TCGA-PRAD | 43.2546519 | 23.1106627 |
| TCGA-4N-A93T | TCGA-COAD | 37.2469869 | 20.9211454 |
| TCGA-4R-AA8I | TCGA-LIHC | 45.7193141 | 18.6992196 |
| TCGA-4T-AA8H | TCGA-COAD | 43.5657276 | 20.5327506 |

| | | | |
|---|---|---|---|
| TCGA-4Z-AA7M | TCGA-BLCA | 92.043649 | 27.8361615 |
| TCGA-4Z-AA7N | TCGA-BLCA | 47.8844526 | 11.2063231 |
| TCGA-4Z-AA7O | TCGA-BLCA | 78.8362359 | 26.3844757 |
| TCGA-4Z-AA7Q | TCGA-BLCA | 40.4274646 | 14.2344871 |
| TCGA-4Z-AA7R | TCGA-BLCA | 84.3773265 | 36.8671418 |
| TCGA-4Z-AA7S | TCGA-BLCA | 77.4508214 | 23.3783138 |
| TCGA-4Z-AA7W | TCGA-BLCA | 92.4644745 | 56.4038392 |
| TCGA-4Z-AA7Y | TCGA-BLCA | 81.4888219 | 43.2166009 |
| TCGA-4Z-AA80 | TCGA-BLCA | 95.4247547 | 46.549189 |
| TCGA-4Z-AA81 | TCGA-BLCA | 53.4888416 | 80.5317981 |
| TCGA-4Z-AA82 | TCGA-BLCA | 80.1070911 | 24.1346072 |
| TCGA-4Z-AA83 | TCGA-BLCA | 52.8525842 | 22.2145198 |
| TCGA-4Z-AA84 | TCGA-BLCA | 70.265767 | 27.7060373 |
| TCGA-4Z-AA86 | TCGA-BLCA | 166.01637 | 78.6703946 |
| TCGA-4Z-AA87 | TCGA-BLCA | 90.4433394 | 24.8736414 |
| TCGA-4Z-AA89 | TCGA-BLCA | 62.2326513 | 22.7851121 |
| TCGA-50-5044 | TCGA-LUAD | 79.247901 | 33.6606138 |
| TCGA-50-5045 | TCGA-LUAD | 52.6372315 | 25.9023411 |
| TCGA-50-5049 | TCGA-LUAD | 62.2474371 | 16.0738264 |
| TCGA-50-5051 | TCGA-LUAD | 42.1598057 | 19.3089755 |
| TCGA-50-5055 | TCGA-LUAD | 43.317025 | 26.591041 |
| TCGA-50-5066 | TCGA-LUAD | 55.9013533 | 20.774107 |
| TCGA-50-5068 | TCGA-LUAD | 66.3885995 | 28.9565759 |
| TCGA-50-5072 | TCGA-LUAD | 62.7353786 | 61.5859895 |
| TCGA-50-5930 | TCGA-LUAD | 37.5612238 | 26.186219 |
| TCGA-50-5931 | TCGA-LUAD | 49.4512155 | 14.8356723 |
| TCGA-50-5932 | TCGA-LUAD | 60.0434933 | 31.403543 |
| TCGA-50-5933 | TCGA-LUAD | 67.5809166 | 35.9092155 |
| TCGA-50-5935 | TCGA-LUAD | 38.2140751 | 13.3195583 |
| TCGA-50-5936 | TCGA-LUAD | 39.6590299 | 11.0025143 |
| TCGA-50-5939 | TCGA-LUAD | 39.3389208 | 11.4054329 |

| | | | |
|---|---|---|---|
| TCGA-50-5941 | TCGA-LUAD | 63.500383 | 29.3605834 |
| TCGA-50-5942 | TCGA-LUAD | 39.1600107 | 6.2333229 |
| TCGA-50-5944 | TCGA-LUAD | 56.3568965 | 22.6456861 |
| TCGA-50-5946 | TCGA-LUAD | 58.0495245 | 19.4966019 |
| TCGA-50-6590 | TCGA-LUAD | 49.9077924 | 58.0333174 |
| TCGA-50-6591 | TCGA-LUAD | 93.266629 | 36.6860458 |
| TCGA-50-6592 | TCGA-LUAD | 53.8150738 | 22.7446464 |
| TCGA-50-6593 | TCGA-LUAD | 60.9793301 | 22.4053069 |
| TCGA-50-6594 | TCGA-LUAD | 58.9799619 | 21.8828849 |
| TCGA-50-6597 | TCGA-LUAD | 79.9180471 | 26.3642741 |
| TCGA-50-6673 | TCGA-LUAD | 38.861725 | 24.5303831 |
| TCGA-50-7109 | TCGA-LUAD | 49.337511 | 26.8432961 |
| TCGA-50-8457 | TCGA-LUAD | 29.9006147 | 6.91514507 |
| TCGA-50-8459 | TCGA-LUAD | 33.0228171 | 10.1175702 |
| TCGA-50-8460 | TCGA-LUAD | 39.6081145 | 17.012218 |
| TCGA-51-4079 | TCGA-LUSC | 63.9814587 | 27.7603817 |
| TCGA-51-4080 | TCGA-LUSC | 43.8847658 | 13.4295296 |
| TCGA-51-4081 | TCGA-LUSC | 5.14279874 | 19.3515194 |
| TCGA-51-6867 | TCGA-LUSC | 14.3429903 | 19.3044577 |
| TCGA-52-7622 | TCGA-LUSC | 39.0925605 | 27.7195782 |
| TCGA-52-7809 | TCGA-LUSC | 88.6087056 | 73.6371145 |
| TCGA-52-7810 | TCGA-LUSC | 86.1908748 | 40.0181241 |
| TCGA-52-7811 | TCGA-LUSC | 38.8512789 | 36.8095496 |
| TCGA-52-7812 | TCGA-LUSC | 127.956924 | 47.7146271 |
| TCGA-53-7624 | TCGA-LUAD | 114.659671 | 35.6276966 |
| TCGA-53-7626 | TCGA-LUAD | 41.6320206 | 19.4610728 |
| TCGA-53-7813 | TCGA-LUAD | 52.1567706 | 33.8010379 |
| TCGA-53-A4EZ | TCGA-LUAD | 77.406812 | 16.7874397 |
| TCGA-55-1592 | TCGA-LUAD | 62.3871612 | 26.0095864 |
| TCGA-55-1594 | TCGA-LUAD | 76.5903945 | 39.0500531 |
| TCGA-55-1596 | TCGA-LUAD | 88.7642817 | 41.9077671 |

| | | | |
|---|---|---|---|
| TCGA-55-5899 | TCGA-LUAD | 46.0210135 | 38.1197129 |
| TCGA-55-6543 | TCGA-LUAD | 41.3996345 | 23.1522031 |
| TCGA-55-6642 | TCGA-LUAD | 25.5230482 | 7.69637514 |
| TCGA-55-6712 | TCGA-LUAD | 51.2719469 | 20.6458416 |
| TCGA-55-6968 | TCGA-LUAD | 59.6299944 | 17.0409447 |
| TCGA-55-6969 | TCGA-LUAD | 34.932923 | 10.8252497 |
| TCGA-55-6970 | TCGA-LUAD | 34.0864971 | 32.8039764 |
| TCGA-55-6971 | TCGA-LUAD | 41.0370943 | 14.1498533 |
| TCGA-55-6972 | TCGA-LUAD | 70.6168938 | 39.7474501 |
| TCGA-55-6975 | TCGA-LUAD | 39.4388703 | 12.8810054 |
| TCGA-55-6978 | TCGA-LUAD | 46.6993161 | 15.9643423 |
| TCGA-55-6979 | TCGA-LUAD | 44.799511 | 15.5480021 |
| TCGA-55-6980 | TCGA-LUAD | 67.4667474 | 24.0898542 |
| TCGA-55-6981 | TCGA-LUAD | 40.9379639 | 27.3493327 |
| TCGA-55-6982 | TCGA-LUAD | 58.0878094 | 23.1297809 |
| TCGA-55-6983 | TCGA-LUAD | 42.7968563 | 26.8016149 |
| TCGA-55-6984 | TCGA-LUAD | 43.2688953 | 12.9152374 |
| TCGA-55-6985 | TCGA-LUAD | 41.2304883 | 14.9525039 |
| TCGA-55-6986 | TCGA-LUAD | 38.0694964 | 14.1038029 |
| TCGA-55-6987 | TCGA-LUAD | 77.5311884 | 22.1901186 |
| TCGA-55-7227 | TCGA-LUAD | 50.625544 | 12.1190225 |
| TCGA-55-7281 | TCGA-LUAD | 63.5471935 | 27.1472945 |
| TCGA-55-7283 | TCGA-LUAD | 70.0697684 | 13.7532458 |
| TCGA-55-7284 | TCGA-LUAD | 45.3197732 | 19.2180822 |
| TCGA-55-7570 | TCGA-LUAD | 101.778088 | 43.6899855 |
| TCGA-55-7573 | TCGA-LUAD | 64.3464899 | 16.0968876 |
| TCGA-55-7574 | TCGA-LUAD | 34.224918 | 17.4947826 |
| TCGA-55-7576 | TCGA-LUAD | 60.0259188 | 24.0483531 |
| TCGA-55-7724 | TCGA-LUAD | 42.6801172 | 24.8863431 |
| TCGA-55-7725 | TCGA-LUAD | 52.8922511 | 24.0819241 |
| TCGA-55-7726 | TCGA-LUAD | 72.0642045 | 49.0867004 |

| | | | |
|---|---|---|---|
| TCGA-55-7727 | TCGA-LUAD | 86.3274867 | 25.8366805 |
| TCGA-55-7728 | TCGA-LUAD | 33.7539141 | 7.70041078 |
| TCGA-55-7815 | TCGA-LUAD | 49.901607 | 33.3862691 |
| TCGA-55-7816 | TCGA-LUAD | 35.4324104 | 16.8844077 |
| TCGA-55-7903 | TCGA-LUAD | 62.1814599 | 31.119271 |
| TCGA-55-7907 | TCGA-LUAD | 40.8238559 | 19.6620348 |
| TCGA-55-7910 | TCGA-LUAD | 64.2823166 | 2.96161193 |
| TCGA-55-7911 | TCGA-LUAD | 57.1269824 | 39.2019368 |
| TCGA-55-7913 | TCGA-LUAD | 47.2764968 | 38.0430598 |
| TCGA-55-7914 | TCGA-LUAD | 58.7344854 | 10.5959323 |
| TCGA-55-7994 | TCGA-LUAD | 50.6261355 | 21.1702529 |
| TCGA-55-7995 | TCGA-LUAD | 47.2113309 | 36.3339585 |
| TCGA-55-8085 | TCGA-LUAD | 51.907023 | 24.9971839 |
| TCGA-55-8087 | TCGA-LUAD | 60.5449011 | 11.6899502 |
| TCGA-55-8089 | TCGA-LUAD | 49.8991952 | 20.843128 |
| TCGA-55-8090 | TCGA-LUAD | 42.1899332 | 21.8682479 |
| TCGA-55-8091 | TCGA-LUAD | 61.3368018 | 20.0136983 |
| TCGA-55-8092 | TCGA-LUAD | 28.5903233 | 23.6268816 |
| TCGA-55-8094 | TCGA-LUAD | 64.6727304 | 52.07221 |
| TCGA-55-8096 | TCGA-LUAD | 42.4745057 | 19.7325288 |
| TCGA-55-8097 | TCGA-LUAD | 54.3622749 | 24.8963047 |
| TCGA-55-8203 | TCGA-LUAD | 42.2464692 | 30.2408836 |
| TCGA-55-8204 | TCGA-LUAD | 49.1332404 | 23.3040695 |
| TCGA-55-8205 | TCGA-LUAD | 78.1639249 | 27.9326975 |
| TCGA-55-8206 | TCGA-LUAD | 64.9585795 | 19.4448522 |
| TCGA-55-8207 | TCGA-LUAD | 45.2850463 | 21.7402258 |
| TCGA-55-8208 | TCGA-LUAD | 43.3729246 | 21.344247 |
| TCGA-55-8299 | TCGA-LUAD | 53.7873319 | 24.0139501 |
| TCGA-55-8301 | TCGA-LUAD | 48.2667583 | 23.959175 |
| TCGA-55-8302 | TCGA-LUAD | 48.805247 | 21.4918395 |
| TCGA-55-8505 | TCGA-LUAD | 53.9783234 | 20.5458712 |

| | | | |
|---|---|---|---|
| TCGA-55-8506 | TCGA-LUAD | 57.2854835 | 26.6289 |
| TCGA-55-8507 | TCGA-LUAD | 71.8980136 | 13.6219231 |
| TCGA-55-8508 | TCGA-LUAD | 48.9716231 | 15.0910002 |
| TCGA-55-8510 | TCGA-LUAD | 44.2721648 | 13.3319054 |
| TCGA-55-8511 | TCGA-LUAD | 54.2179001 | 13.9016302 |
| TCGA-55-8512 | TCGA-LUAD | 41.1090823 | 11.8230268 |
| TCGA-55-8513 | TCGA-LUAD | 33.925122 | 11.6984559 |
| TCGA-55-8514 | TCGA-LUAD | 43.499665 | 10.7285355 |
| TCGA-55-8614 | TCGA-LUAD | 62.8626038 | 16.5568765 |
| TCGA-55-8615 | TCGA-LUAD | 37.5745788 | 31.4141383 |
| TCGA-55-8616 | TCGA-LUAD | 40.5549118 | 16.4221529 |
| TCGA-55-8619 | TCGA-LUAD | 28.5720131 | 8.25263924 |
| TCGA-55-8620 | TCGA-LUAD | 48.3325121 | 26.7501071 |
| TCGA-55-8621 | TCGA-LUAD | 37.2078504 | 12.282466 |
| TCGA-55-A48X | TCGA-LUAD | 46.153936 | 9.03375948 |
| TCGA-55-A48Y | TCGA-LUAD | 39.8450583 | 27.1890835 |
| TCGA-55-A48Z | TCGA-LUAD | 44.5572693 | 16.5804767 |
| TCGA-55-A490 | TCGA-LUAD | 40.9119925 | 13.3488623 |
| TCGA-55-A491 | TCGA-LUAD | 43.801079 | 11.7013268 |
| TCGA-55-A492 | TCGA-LUAD | 37.4574809 | 29.5394138 |
| TCGA-55-A493 | TCGA-LUAD | 63.6482966 | 33.7056351 |
| TCGA-55-A494 | TCGA-LUAD | 42.7511201 | 47.3709996 |
| TCGA-55-A4DF | TCGA-LUAD | 106.69907 | 40.6055242 |
| TCGA-55-A4DG | TCGA-LUAD | 40.6808383 | 12.0100258 |
| TCGA-55-A57B | TCGA-LUAD | 56.3895187 | 17.8643541 |
| TCGA-56-1622 | TCGA-LUSC | 45.8183858 | 61.470095 |
| TCGA-56-5897 | TCGA-LUSC | 27.3997438 | 15.3012682 |
| TCGA-56-5898 | TCGA-LUSC | 30.8084114 | 21.9392487 |
| TCGA-56-6545 | TCGA-LUSC | 36.2632818 | 25.3283408 |
| TCGA-56-6546 | TCGA-LUSC | 14.3609358 | 14.3512099 |
| TCGA-56-7221 | TCGA-LUSC | 26.2001755 | 7.28477148 |

| | | | |
|---|---|---|---|
| TCGA-56-7222 | TCGA-LUSC | 106.12297 | 65.6800033 |
| TCGA-56-7223 | TCGA-LUSC | 37.5723709 | 17.2401358 |
| TCGA-56-7579 | TCGA-LUSC | 57.1472624 | 7.46930263 |
| TCGA-56-7580 | TCGA-LUSC | 28.5846455 | 4.58897492 |
| TCGA-56-7582 | TCGA-LUSC | 31.2507499 | 36.5594404 |
| TCGA-56-7730 | TCGA-LUSC | 63.2389481 | 70.6656849 |
| TCGA-56-7731 | TCGA-LUSC | 52.0636314 | 25.0091796 |
| TCGA-56-7822 | TCGA-LUSC | 42.2267986 | 32.4929894 |
| TCGA-56-7823 | TCGA-LUSC | 45.7525539 | 45.1740877 |
| TCGA-56-8082 | TCGA-LUSC | 86.8569494 | 30.3522125 |
| TCGA-56-8083 | TCGA-LUSC | 139.220967 | 58.9573945 |
| TCGA-56-8201 | TCGA-LUSC | 63.3337565 | 21.7718863 |
| TCGA-56-8304 | TCGA-LUSC | 16.0840712 | 14.7601903 |
| TCGA-56-8305 | TCGA-LUSC | 60.0433419 | 22.3765119 |
| TCGA-56-8308 | TCGA-LUSC | 38.0990625 | 11.4258865 |
| TCGA-56-8309 | TCGA-LUSC | 27.6713478 | 14.9178316 |
| TCGA-56-8503 | TCGA-LUSC | 41.1646326 | 18.476182 |
| TCGA-56-8504 | TCGA-LUSC | 65.6464409 | 47.4610255 |
| TCGA-56-8622 | TCGA-LUSC | 13.1066535 | 9.01113305 |
| TCGA-56-8623 | TCGA-LUSC | 42.9335004 | 11.114468 |
| TCGA-56-8624 | TCGA-LUSC | 25.1084977 | 3.37335037 |
| TCGA-56-8625 | TCGA-LUSC | 29.419862 | 26.9720837 |
| TCGA-56-8626 | TCGA-LUSC | 24.9141398 | 9.02701833 |
| TCGA-56-8628 | TCGA-LUSC | 47.3876588 | 9.72827908 |
| TCGA-56-8629 | TCGA-LUSC | 66.1397524 | 22.6308142 |
| TCGA-56-A49D | TCGA-LUSC | 88.5640801 | 31.3021524 |
| TCGA-56-A4BW | TCGA-LUSC | 58.2947465 | 40.1906935 |
| TCGA-56-A4BX | TCGA-LUSC | 28.5574137 | 7.62072978 |
| TCGA-56-A4BY | TCGA-LUSC | 74.5547874 | 24.4762693 |
| TCGA-56-A4ZK | TCGA-LUSC | 17.3160852 | 4.13117149 |
| TCGA-56-A5DR | TCGA-LUSC | 62.3482274 | 81.5150471 |

| | | | |
|---|---|---|---|
| TCGA-56-A5DS | TCGA-LUSC | 54.9847172 | 15.1569826 |
| TCGA-56-A62T | TCGA-LUSC | 70.7776221 | 62.369462 |
| TCGA-57-1582 | TCGA-OV | 59.6018302 | 21.7619363 |
| TCGA-57-1583 | TCGA-OV | 30.2357153 | 19.4649716 |
| TCGA-57-1584 | TCGA-OV | 72.9950824 | 18.1999623 |
| TCGA-57-1585 | TCGA-OV | 61.2246476 | 13.019936 |
| TCGA-57-1586 | TCGA-OV | 42.6476473 | 20.9569629 |
| TCGA-57-1993 | TCGA-OV | 62.0035683 | 82.2631464 |
| TCGA-57-1994 | TCGA-OV | 57.1996985 | 48.0881436 |
| TCGA-58-8386 | TCGA-LUSC | 89.7381144 | 3.53587624 |
| TCGA-58-8387 | TCGA-LUSC | 17.1820575 | 8.26907142 |
| TCGA-58-8388 | TCGA-LUSC | 49.502235 | 44.3442091 |
| TCGA-58-8390 | TCGA-LUSC | 93.888941 | 39.3655472 |
| TCGA-58-8391 | TCGA-LUSC | 87.2183604 | 5.50137474 |
| TCGA-58-8392 | TCGA-LUSC | 45.4310691 | 23.5677017 |
| TCGA-58-8393 | TCGA-LUSC | 38.7240632 | 20.8772806 |
| TCGA-58-A46J | TCGA-LUSC | 107.116551 | 49.769759 |
| TCGA-58-A46K | TCGA-LUSC | 31.8169014 | 12.5938733 |
| TCGA-58-A46L | TCGA-LUSC | 50.0873584 | 18.3787391 |
| TCGA-58-A46M | TCGA-LUSC | 55.2698011 | 45.2807083 |
| TCGA-58-A46N | TCGA-LUSC | 50.1989906 | 13.1372777 |
| TCGA-59-2348 | TCGA-OV | 81.7366991 | 44.2852204 |
| TCGA-59-2350 | TCGA-OV | 121.484277 | 32.1911902 |
| TCGA-59-2351 | TCGA-OV | 31.4787673 | 32.9229821 |
| TCGA-59-2352 | TCGA-OV | 46.0382839 | 32.5394386 |
| TCGA-59-2354 | TCGA-OV | 68.1974245 | 25.6911508 |
| TCGA-59-2355 | TCGA-OV | 61.5832663 | 22.1765864 |
| TCGA-59-2363 | TCGA-OV | 88.4105596 | 57.9519124 |
| TCGA-59-A5PD | TCGA-OV | 55.0550525 | 26.8373159 |
| TCGA-5B-A90C | TCGA-UCEC | 122.615882 | 14.2978654 |
| TCGA-5C-A9VG | TCGA-LIHC | 47.3160178 | 50.5924645 |

| | | | |
|---|---|---|---|
| TCGA-5C-A9VH | TCGA-LIHC | 25.8500533 | 32.0551878 |
| TCGA-5C-AAPD | TCGA-LIHC | 63.1041601 | 28.1953876 |
| TCGA-5L-AAT0 | TCGA-BRCA | 55.6001998 | 16.8482369 |
| TCGA-5L-AAT1 | TCGA-BRCA | 63.8808463 | 22.2316536 |
| TCGA-5M-AAT4 | TCGA-COAD | 46.8933853 | 32.8193777 |
| TCGA-5M-AAT5 | TCGA-COAD | 105.321504 | 34.6929313 |
| TCGA-5M-AAT6 | TCGA-COAD | 77.9768662 | 47.481494 |
| TCGA-5M-AATA | TCGA-COAD | 55.588478 | 23.029164 |
| TCGA-5M-AATE | TCGA-COAD | 70.3129365 | 27.2413682 |
| TCGA-5N-A9KI | TCGA-BLCA | 91.3057507 | 33.7591781 |
| TCGA-5N-A9KM | TCGA-BLCA | 76.3947489 | 27.540776 |
| TCGA-5P-A9JU | TCGA-KIRP | 43.3453465 | 13.855049 |
| TCGA-5P-A9JV | TCGA-KIRP | 41.7597304 | 14.379761 |
| TCGA-5P-A9JY | TCGA-KIRP | 49.6444288 | 17.0961309 |
| TCGA-5P-A9JZ | TCGA-KIRP | 13.9096332 | 7.29678416 |
| TCGA-5P-A9K0 | TCGA-KIRP | 37.6384598 | 13.9176542 |
| TCGA-5P-A9K3 | TCGA-KIRP | 28.7940312 | 18.5381158 |
| TCGA-5P-A9K4 | TCGA-KIRP | 46.6449075 | 16.2578296 |
| TCGA-5P-A9K6 | TCGA-KIRP | 29.0005798 | 8.83735462 |
| TCGA-5P-A9K8 | TCGA-KIRP | 27.8480655 | 29.1294611 |
| TCGA-5P-A9K9 | TCGA-KIRP | 65.2651951 | 35.5781631 |
| TCGA-5P-A9KA | TCGA-KIRP | 24.7973012 | 25.4569621 |
| TCGA-5P-A9KC | TCGA-KIRP | 13.3919144 | 1.97498723 |
| TCGA-5P-A9KE | TCGA-KIRP | 7.14650745 | 1.24030889 |
| TCGA-5P-A9KF | TCGA-KIRP | 49.3157242 | 16.3165648 |
| TCGA-5P-A9KH | TCGA-KIRP | 23.5922235 | 26.4415342 |
| TCGA-5R-AA1C | TCGA-LIHC | 57.5900067 | 36.0368651 |
| TCGA-5R-AA1D | TCGA-LIHC | 34.1368377 | 16.8185641 |
| TCGA-5R-AAAM | TCGA-LIHC | 34.1402599 | 13.8708606 |
| TCGA-5S-A9Q8 | TCGA-UCEC | 9.25086836 | 5.08259054 |
| TCGA-5T-A9QA | TCGA-BRCA | 40.3338169 | 43.2639318 |

| | | | |
|---|---|---|---|
| TCGA-5X-AA5U | TCGA-OV | 77.8280345 | 14.0483083 |
| TCGA-60-2695 | TCGA-LUSC | 81.7696998 | 32.0913164 |
| TCGA-60-2696 | TCGA-LUSC | 63.6384011 | 29.074308 |
| TCGA-60-2697 | TCGA-LUSC | 5.26411382 | 12.6280671 |
| TCGA-60-2698 | TCGA-LUSC | 55.527499 | 29.2945602 |
| TCGA-60-2703 | TCGA-LUSC | 58.1952193 | 52.6091381 |
| TCGA-60-2704 | TCGA-LUSC | 31.864503 | 16.3109545 |
| TCGA-60-2706 | TCGA-LUSC | 21.9497771 | 39.7192752 |
| TCGA-60-2707 | TCGA-LUSC | 76.544648 | 7.18549885 |
| TCGA-60-2708 | TCGA-LUSC | 27.7859363 | 3.57459623 |
| TCGA-60-2709 | TCGA-LUSC | 8.7194256 | 7.15234132 |
| TCGA-60-2710 | TCGA-LUSC | 59.9664697 | 22.3083093 |
| TCGA-60-2711 | TCGA-LUSC | 55.3323277 | 10.4924639 |
| TCGA-60-2712 | TCGA-LUSC | 3.8217724 | 2.45976065 |
| TCGA-60-2713 | TCGA-LUSC | 52.933337 | 14.1601277 |
| TCGA-60-2714 | TCGA-LUSC | 9.81385911 | 5.80095655 |
| TCGA-60-2715 | TCGA-LUSC | 35.3683613 | 14.6904758 |
| TCGA-60-2716 | TCGA-LUSC | 43.9687042 | 12.5102063 |
| TCGA-60-2719 | TCGA-LUSC | 77.0009973 | 27.2062073 |
| TCGA-60-2720 | TCGA-LUSC | 30.8237951 | 8.74477746 |
| TCGA-60-2721 | TCGA-LUSC | 22.2393497 | 4.37485255 |
| TCGA-60-2722 | TCGA-LUSC | 79.1813577 | 2.73739842 |
| TCGA-60-2723 | TCGA-LUSC | 55.380459 | 33.6144427 |
| TCGA-60-2724 | TCGA-LUSC | 30.8911714 | 9.24946127 |
| TCGA-60-2725 | TCGA-LUSC | 33.1550821 | 17.8672585 |
| TCGA-60-2726 | TCGA-LUSC | 61.5948049 | 77.5433994 |
| TCGA-61-1721 | TCGA-OV | 42.0688872 | 18.3748426 |
| TCGA-61-1724 | TCGA-OV | 102.98126 | 13.9089527 |
| TCGA-61-1725 | TCGA-OV | 84.0403804 | 24.6487521 |
| TCGA-61-1728 | TCGA-OV | 69.5045901 | 20.5145768 |
| TCGA-61-1733 | TCGA-OV | 123.162944 | 36.8514842 |

| | | | |
|---|---|---|---|
| TCGA-61-1736 | TCGA-OV | 45.3300162 | 22.4876732 |
| TCGA-61-1737 | TCGA-OV | 133.00846 | 37.3595177 |
| TCGA-61-1738 | TCGA-OV | 79.5646929 | 18.9907049 |
| TCGA-61-1741 | TCGA-OV | 80.332602 | 33.6337286 |
| TCGA-61-1900 | TCGA-OV | 108.304531 | 29.8335903 |
| TCGA-61-1907 | TCGA-OV | 104.0463 | 30.138695 |
| TCGA-61-1910 | TCGA-OV | 55.4089221 | 11.5266768 |
| TCGA-61-1911 | TCGA-OV | 46.567893 | 22.2089597 |
| TCGA-61-1914 | TCGA-OV | 110.722578 | 46.7491074 |
| TCGA-61-1918 | TCGA-OV | 74.8388975 | 16.7004489 |
| TCGA-61-1919 | TCGA-OV | 66.7976148 | 48.6563123 |
| TCGA-61-1995 | TCGA-OV | 65.0139076 | 24.9193186 |
| TCGA-61-1998 | TCGA-OV | 69.0317662 | 18.0447165 |
| TCGA-61-2000 | TCGA-OV | 76.3161624 | 24.6981402 |
| TCGA-61-2002 | TCGA-OV | 64.1343254 | 31.4730844 |
| TCGA-61-2003 | TCGA-OV | 62.1996204 | 29.3435985 |
| TCGA-61-2008 | TCGA-OV | 61.2491174 | 34.8359751 |
| TCGA-61-2009 | TCGA-OV | 46.182391 | 35.6764421 |
| TCGA-61-2012 | TCGA-OV | 34.4796176 | 52.4309968 |
| TCGA-61-2088 | TCGA-OV | 71.7823028 | 11.0548022 |
| TCGA-61-2092 | TCGA-OV | 76.4041545 | 20.9644705 |
| TCGA-61-2097 | TCGA-OV | 50.2029773 | 20.656124 |
| TCGA-61-2098 | TCGA-OV | 45.9925273 | 31.8749058 |
| TCGA-61-2101 | TCGA-OV | 35.1367885 | 25.0736351 |
| TCGA-61-2102 | TCGA-OV | 125.132388 | 32.9610842 |
| TCGA-61-2104 | TCGA-OV | 66.6370681 | 41.6727321 |
| TCGA-61-2109 | TCGA-OV | 77.636625 | 19.4168359 |
| TCGA-61-2110 | TCGA-OV | 63.5301457 | 43.244 |
| TCGA-61-2111 | TCGA-OV | 69.9412005 | 32.0440295 |
| TCGA-61-2113 | TCGA-OV | 84.7230532 | 43.9109966 |
| TCGA-62-8394 | TCGA-LUAD | 132.733365 | 31.1718364 |

| | | | |
|---|---|---|---|
| TCGA-62-8395 | TCGA-LUAD | 50.8008202 | 19.0819319 |
| TCGA-62-8397 | TCGA-LUAD | 31.924138 | 9.29010045 |
| TCGA-62-8398 | TCGA-LUAD | 31.4380458 | 38.5399184 |
| TCGA-62-8399 | TCGA-LUAD | 71.2584525 | 33.732757 |
| TCGA-62-8402 | TCGA-LUAD | 70.9170007 | 22.8369153 |
| TCGA-62-A46O | TCGA-LUAD | 85.2809467 | 46.8256271 |
| TCGA-62-A46P | TCGA-LUAD | 52.2646788 | 37.2572318 |
| TCGA-62-A46R | TCGA-LUAD | 57.9778878 | 19.3603997 |
| TCGA-62-A46S | TCGA-LUAD | 32.7096539 | 23.8272425 |
| TCGA-62-A46V | TCGA-LUAD | 79.4392384 | 22.0826948 |
| TCGA-62-A46Y | TCGA-LUAD | 51.6663619 | 20.6053505 |
| TCGA-62-A470 | TCGA-LUAD | 59.3651285 | 33.304341 |
| TCGA-62-A471 | TCGA-LUAD | 57.8275242 | 47.5966019 |
| TCGA-62-A472 | TCGA-LUAD | 59.3267651 | 30.5005232 |
| TCGA-63-5128 | TCGA-LUSC | 42.9370349 | 103.334238 |
| TCGA-63-5131 | TCGA-LUSC | 49.1775343 | 33.2730877 |
| TCGA-63-6202 | TCGA-LUSC | 42.0365368 | 22.6873969 |
| TCGA-63-7020 | TCGA-LUSC | 39.4760216 | 24.5405662 |
| TCGA-63-7021 | TCGA-LUSC | 57.5937178 | 2.39886479 |
| TCGA-63-7022 | TCGA-LUSC | 21.0470084 | 7.23742686 |
| TCGA-63-7023 | TCGA-LUSC | 41.9900679 | 81.9268279 |
| TCGA-63-A5M9 | TCGA-LUSC | 49.0092526 | 19.1905182 |
| TCGA-63-A5MB | TCGA-LUSC | 65.8795799 | 31.7621982 |
| TCGA-63-A5MG | TCGA-LUSC | 31.7146144 | 16.86515 |
| TCGA-63-A5MH | TCGA-LUSC | 64.6591366 | 13.6846257 |
| TCGA-63-A5MI | TCGA-LUSC | 72.3541725 | 17.1072916 |
| TCGA-63-A5MJ | TCGA-LUSC | 52.0677239 | 34.9595654 |
| TCGA-63-A5ML | TCGA-LUSC | 80.6274692 | 34.4426952 |
| TCGA-63-A5MM | TCGA-LUSC | 8.38241017 | 7.58500908 |
| TCGA-63-A5MN | TCGA-LUSC | 7.15103867 | 7.57203373 |
| TCGA-63-A5MP | TCGA-LUSC | 67.1310352 | 20.7546093 |

| | | | |
|---|---|---|---|
| TCGA-63-A5MR | TCGA-LUSC | 48.0363601 | 36.1217225 |
| TCGA-63-A5MS | TCGA-LUSC | 25.835518 | 10.155039 |
| TCGA-63-A5MT | TCGA-LUSC | 30.7421657 | 7.85701412 |
| TCGA-63-A5MU | TCGA-LUSC | 75.34479 | 18.8180222 |
| TCGA-63-A5MV | TCGA-LUSC | 58.8876619 | 8.79266914 |
| TCGA-63-A5MW | TCGA-LUSC | 51.2954142 | 9.58625553 |
| TCGA-63-A5MY | TCGA-LUSC | 146.88468 | 7.99450632 |
| TCGA-64-1676 | TCGA-LUAD | 47.4490365 | 25.8392335 |
| TCGA-64-1677 | TCGA-LUAD | 54.4909062 | 19.9545087 |
| TCGA-64-1678 | TCGA-LUAD | 83.4995771 | 49.2216557 |
| TCGA-64-1679 | TCGA-LUAD | 62.8371754 | 29.722322 |
| TCGA-64-1680 | TCGA-LUAD | 43.6263392 | 22.4254454 |
| TCGA-64-1681 | TCGA-LUAD | 104.917744 | 22.8144722 |
| TCGA-64-5774 | TCGA-LUAD | 50.9567116 | 28.1078328 |
| TCGA-64-5775 | TCGA-LUAD | 64.6107474 | 40.9279188 |
| TCGA-64-5778 | TCGA-LUAD | 73.0744241 | 31.440584 |
| TCGA-64-5779 | TCGA-LUAD | 27.2161727 | 11.6485392 |
| TCGA-64-5781 | TCGA-LUAD | 56.9580446 | 34.6774789 |
| TCGA-64-5815 | TCGA-LUAD | 32.1526505 | 22.2243209 |
| TCGA-66-2727 | TCGA-LUSC | 83.7247925 | 37.5217747 |
| TCGA-66-2734 | TCGA-LUSC | 73.5594326 | 36.3607452 |
| TCGA-66-2737 | TCGA-LUSC | 29.1199578 | 27.8894997 |
| TCGA-66-2742 | TCGA-LUSC | 35.4281021 | 20.969646 |
| TCGA-66-2744 | TCGA-LUSC | 48.5640302 | 24.7434322 |
| TCGA-66-2754 | TCGA-LUSC | 76.6624177 | 49.1628021 |
| TCGA-66-2755 | TCGA-LUSC | 52.4480836 | 26.392172 |
| TCGA-66-2756 | TCGA-LUSC | 19.1491053 | 8.29549799 |
| TCGA-66-2757 | TCGA-LUSC | 43.2616757 | 41.4320927 |
| TCGA-66-2758 | TCGA-LUSC | 18.6786981 | 10.2133352 |
| TCGA-66-2759 | TCGA-LUSC | 73.6110047 | 11.7523582 |
| TCGA-66-2763 | TCGA-LUSC | 107.485211 | 5.70129696 |

| | | | |
|---|---|---|---|
| TCGA-66-2766 | TCGA-LUSC | 37.1845594 | 31.2254586 |
| TCGA-66-2767 | TCGA-LUSC | 14.6517507 | 54.1675514 |
| TCGA-66-2768 | TCGA-LUSC | 84.311526 | 32.9523628 |
| TCGA-66-2769 | TCGA-LUSC | 39.9243975 | 29.6888222 |
| TCGA-66-2770 | TCGA-LUSC | 59.0889229 | 40.2206289 |
| TCGA-66-2771 | TCGA-LUSC | 22.5842167 | 44.071211 |
| TCGA-66-2773 | TCGA-LUSC | 41.7074391 | 35.5734285 |
| TCGA-66-2777 | TCGA-LUSC | 39.1193998 | 19.7467658 |
| TCGA-66-2778 | TCGA-LUSC | 30.6078396 | 14.5573847 |
| TCGA-66-2780 | TCGA-LUSC | 72.1605735 | 32.4990379 |
| TCGA-66-2781 | TCGA-LUSC | 14.8038352 | 4.49692243 |
| TCGA-66-2782 | TCGA-LUSC | 57.3128262 | 33.6798149 |
| TCGA-66-2783 | TCGA-LUSC | 44.509296 | 21.7478708 |
| TCGA-66-2785 | TCGA-LUSC | 30.1133807 | 41.9700677 |
| TCGA-66-2786 | TCGA-LUSC | 30.8345781 | 7.62085159 |
| TCGA-66-2787 | TCGA-LUSC | 18.3707821 | 23.0362345 |
| TCGA-66-2788 | TCGA-LUSC | 38.4678231 | 18.7342752 |
| TCGA-66-2789 | TCGA-LUSC | 31.4458983 | 7.39800159 |
| TCGA-66-2791 | TCGA-LUSC | 38.5081101 | 29.1817143 |
| TCGA-66-2792 | TCGA-LUSC | 43.4987233 | 21.3968391 |
| TCGA-66-2793 | TCGA-LUSC | 145.298114 | 48.0606438 |
| TCGA-66-2794 | TCGA-LUSC | 33.4481379 | 50.894463 |
| TCGA-66-2795 | TCGA-LUSC | 14.2052357 | 12.2997008 |
| TCGA-66-2800 | TCGA-LUSC | 49.6087201 | 19.4582716 |
| TCGA-67-3770 | TCGA-LUAD | 23.9659513 | 11.8902019 |
| TCGA-67-3771 | TCGA-LUAD | 57.9271635 | 9.21067664 |
| TCGA-67-3772 | TCGA-LUAD | 23.6906109 | 22.6377228 |
| TCGA-67-3773 | TCGA-LUAD | 53.4029497 | 17.3576362 |
| TCGA-67-3774 | TCGA-LUAD | 40.7735999 | 11.9508236 |
| TCGA-67-4679 | TCGA-LUAD | 37.1221812 | 13.8216513 |
| TCGA-67-6215 | TCGA-LUAD | 37.9486147 | 25.7434608 |

| | | | |
|---|---|---|---|
| TCGA-67-6216 | TCGA-LUAD | 44.8448418 | 20.024218 |
| TCGA-67-6217 | TCGA-LUAD | 60.9181431 | 18.569408 |
| TCGA-68-7755 | TCGA-LUSC | 29.2999261 | 45.1645681 |
| TCGA-68-7756 | TCGA-LUSC | 32.7096943 | 8.92379072 |
| TCGA-68-7757 | TCGA-LUSC | 15.6033534 | 2.91549028 |
| TCGA-68-8250 | TCGA-LUSC | 44.3137433 | 31.8718016 |
| TCGA-68-8251 | TCGA-LUSC | 81.0463857 | 29.4791714 |
| TCGA-68-A59I | TCGA-LUSC | 33.0532177 | 14.8791929 |
| TCGA-68-A59J | TCGA-LUSC | 90.3224447 | 8.90146757 |
| TCGA-69-7760 | TCGA-LUAD | 59.1731446 | 29.9966016 |
| TCGA-69-7761 | TCGA-LUAD | 32.5602243 | 18.9917502 |
| TCGA-69-7763 | TCGA-LUAD | 52.9827734 | 22.9810045 |
| TCGA-69-7764 | TCGA-LUAD | 42.0894818 | 21.0777285 |
| TCGA-69-7765 | TCGA-LUAD | 43.1676212 | 24.4431054 |
| TCGA-69-7973 | TCGA-LUAD | 37.8495689 | 52.1147943 |
| TCGA-69-7974 | TCGA-LUAD | 43.5921141 | 18.9360367 |
| TCGA-69-7978 | TCGA-LUAD | 62.5482833 | 14.664267 |
| TCGA-69-7979 | TCGA-LUAD | 57.9945239 | 17.353315 |
| TCGA-69-7980 | TCGA-LUAD | 56.8286926 | 38.0339284 |
| TCGA-69-8253 | TCGA-LUAD | 58.5495157 | 23.2946257 |
| TCGA-69-8254 | TCGA-LUAD | 40.9149334 | 18.3160869 |
| TCGA-69-8255 | TCGA-LUAD | 72.3049308 | 30.8131447 |
| TCGA-69-8453 | TCGA-LUAD | 44.6855502 | 12.7364548 |
| TCGA-6D-AA2E | TCGA-KIRC | 44.1765246 | 11.9258384 |
| TCGA-70-6722 | TCGA-LUSC | 73.4614962 | 28.1849377 |
| TCGA-70-6723 | TCGA-LUSC | 5.39599951 | 8.60455874 |
| TCGA-71-6725 | TCGA-LUAD | 67.369038 | 23.0114761 |
| TCGA-71-8520 | TCGA-LUAD | 82.3523888 | 23.1507755 |
| TCGA-73-4658 | TCGA-LUAD | 45.4580188 | 24.189522 |
| TCGA-73-4659 | TCGA-LUAD | 38.5255753 | 32.5572486 |
| TCGA-73-4662 | TCGA-LUAD | 48.8385434 | 15.7804 |

| | | | |
|---|---|---|---|
| TCGA-73-4666 | TCGA-LUAD | 52.7210733 | 39.4791794 |
| TCGA-73-4668 | TCGA-LUAD | 47.9387786 | 19.7922716 |
| TCGA-73-4670 | TCGA-LUAD | 34.0029328 | 60.345569 |
| TCGA-73-4675 | TCGA-LUAD | 37.6037986 | 25.7083747 |
| TCGA-73-4676 | TCGA-LUAD | 38.3555437 | 18.1711444 |
| TCGA-73-4677 | TCGA-LUAD | 25.4828102 | 19.3637757 |
| TCGA-73-7498 | TCGA-LUAD | 43.3670858 | 19.4091918 |
| TCGA-73-7499 | TCGA-LUAD | 94.5558583 | 16.8607317 |
| TCGA-73-A9RS | TCGA-LUAD | 39.0098456 | 32.829721 |
| TCGA-75-5122 | TCGA-LUAD | 45.3790739 | 28.6939337 |
| TCGA-75-5125 | TCGA-LUAD | 90.8889146 | 47.2953669 |
| TCGA-75-5126 | TCGA-LUAD | 57.1590708 | 27.7069072 |
| TCGA-75-5146 | TCGA-LUAD | 68.3983694 | 30.5362496 |
| TCGA-75-5147 | TCGA-LUAD | 100.86081 | 17.2349074 |
| TCGA-75-6203 | TCGA-LUAD | 31.73334 | 15.3030427 |
| TCGA-75-6205 | TCGA-LUAD | 63.4785268 | 19.1359621 |
| TCGA-75-6206 | TCGA-LUAD | 26.1152756 | 28.8120818 |
| TCGA-75-6207 | TCGA-LUAD | 124.091201 | 20.9781531 |
| TCGA-75-6211 | TCGA-LUAD | 49.1803543 | 65.171454 |
| TCGA-75-6212 | TCGA-LUAD | 34.1179399 | 14.8382145 |
| TCGA-75-6214 | TCGA-LUAD | 53.0600657 | 20.6660058 |
| TCGA-75-7025 | TCGA-LUAD | 40.7886015 | 18.8185873 |
| TCGA-75-7027 | TCGA-LUAD | 43.1137888 | 53.0158038 |
| TCGA-75-7030 | TCGA-LUAD | 33.5325581 | 10.7550961 |
| TCGA-75-7031 | TCGA-LUAD | 60.6187922 | 36.599861 |
| TCGA-76-4925 | TCGA-GBM | 75.6680157 | 32.7779969 |
| TCGA-76-4926 | TCGA-GBM | 51.2066219 | 16.1960695 |
| TCGA-76-4927 | TCGA-GBM | 54.504116 | 26.8133122 |
| TCGA-76-4928 | TCGA-GBM | 61.2925942 | 30.6450145 |
| TCGA-76-4929 | TCGA-GBM | 64.2429187 | 23.33443 |
| TCGA-76-4931 | TCGA-GBM | 60.8130004 | 27.3569488 |

| | | | |
|---|---|---|---|
| TCGA-76-4932 | TCGA-GBM | 66.9195858 | 24.5926609 |
| TCGA-77-6842 | TCGA-LUSC | 43.3010539 | 26.6359389 |
| TCGA-77-6843 | TCGA-LUSC | 64.5883895 | 30.0906005 |
| TCGA-77-6844 | TCGA-LUSC | 92.0803996 | 2.53957864 |
| TCGA-77-6845 | TCGA-LUSC | 54.0385545 | 5.94964179 |
| TCGA-77-7138 | TCGA-LUSC | 88.3015343 | 52.3393015 |
| TCGA-77-7139 | TCGA-LUSC | 55.1352382 | 21.0240837 |
| TCGA-77-7140 | TCGA-LUSC | 36.6421994 | 20.5393051 |
| TCGA-77-7141 | TCGA-LUSC | 43.8706423 | 41.3092021 |
| TCGA-77-7142 | TCGA-LUSC | 118.01949 | 33.0158862 |
| TCGA-77-7335 | TCGA-LUSC | 76.1292061 | 20.5658689 |
| TCGA-77-7337 | TCGA-LUSC | 48.4383685 | 15.9501194 |
| TCGA-77-7338 | TCGA-LUSC | 62.320759 | 29.0477527 |
| TCGA-77-7463 | TCGA-LUSC | 44.7221597 | 63.1690629 |
| TCGA-77-7465 | TCGA-LUSC | 78.4331137 | 92.0395567 |
| TCGA-77-8007 | TCGA-LUSC | 38.2745055 | 21.6538661 |
| TCGA-77-8008 | TCGA-LUSC | 32.598575 | 17.0449157 |
| TCGA-77-8009 | TCGA-LUSC | 76.8957405 | 72.9735654 |
| TCGA-77-8128 | TCGA-LUSC | 54.1366288 | 32.9681557 |
| TCGA-77-8130 | TCGA-LUSC | 28.9754143 | 8.14688309 |
| TCGA-77-8131 | TCGA-LUSC | 47.1815493 | 16.8483905 |
| TCGA-77-8133 | TCGA-LUSC | 46.9538501 | 30.6165423 |
| TCGA-77-8136 | TCGA-LUSC | 42.5369342 | 7.36914231 |
| TCGA-77-8138 | TCGA-LUSC | 29.9741274 | 33.8488224 |
| TCGA-77-8139 | TCGA-LUSC | 82.315662 | 4.41469676 |
| TCGA-77-8140 | TCGA-LUSC | 33.0164003 | 10.3871753 |
| TCGA-77-8143 | TCGA-LUSC | 75.0397972 | 32.4948487 |
| TCGA-77-8145 | TCGA-LUSC | 45.0566971 | 23.2301354 |
| TCGA-77-8146 | TCGA-LUSC | 30.2540938 | 29.4452349 |
| TCGA-77-8148 | TCGA-LUSC | 64.1077098 | 24.8187887 |
| TCGA-77-8150 | TCGA-LUSC | 8.07989269 | 6.4958224 |

| | | | |
|---|---|---|---|
| TCGA-77-8153 | TCGA-LUSC | 86.7530823 | 124.0125 |
| TCGA-77-8154 | TCGA-LUSC | 87.2412945 | 27.3010723 |
| TCGA-77-8156 | TCGA-LUSC | 57.8490691 | 17.6485675 |
| TCGA-77-A5FZ | TCGA-LUSC | 22.5183789 | 3.22845742 |
| TCGA-77-A5G1 | TCGA-LUSC | 74.5683443 | 69.323385 |
| TCGA-77-A5G3 | TCGA-LUSC | 45.1176732 | 29.233982 |
| TCGA-77-A5G6 | TCGA-LUSC | 61.9394042 | 21.3917452 |
| TCGA-77-A5G8 | TCGA-LUSC | 17.4028145 | 5.96887253 |
| TCGA-77-A5GA | TCGA-LUSC | 52.2190928 | 30.9872323 |
| TCGA-77-A5GB | TCGA-LUSC | 62.1959768 | 15.6861234 |
| TCGA-77-A5GF | TCGA-LUSC | 29.7066159 | 12.0604577 |
| TCGA-77-A5GH | TCGA-LUSC | 54.4082372 | 52.3326349 |
| TCGA-78-7143 | TCGA-LUAD | 72.5891649 | 37.9933839 |
| TCGA-78-7145 | TCGA-LUAD | 126.43045 | 21.6509466 |
| TCGA-78-7146 | TCGA-LUAD | 81.7246291 | 49.2895302 |
| TCGA-78-7147 | TCGA-LUAD | 85.5353939 | 26.0294074 |
| TCGA-78-7148 | TCGA-LUAD | 33.6604109 | 33.3495386 |
| TCGA-78-7149 | TCGA-LUAD | 49.6368211 | 17.4119496 |
| TCGA-78-7150 | TCGA-LUAD | 64.6234678 | 54.3138536 |
| TCGA-78-7152 | TCGA-LUAD | 37.6630981 | 14.7008448 |
| TCGA-78-7153 | TCGA-LUAD | 36.1832406 | 14.3873259 |
| TCGA-78-7154 | TCGA-LUAD | 63.0425028 | 40.0900086 |
| TCGA-78-7155 | TCGA-LUAD | 94.7549884 | 35.9076289 |
| TCGA-78-7156 | TCGA-LUAD | 44.7130419 | 17.1414147 |
| TCGA-78-7158 | TCGA-LUAD | 45.8324955 | 16.1781677 |
| TCGA-78-7159 | TCGA-LUAD | 59.8659861 | 24.2770679 |
| TCGA-78-7160 | TCGA-LUAD | 32.3522067 | 30.0191558 |
| TCGA-78-7161 | TCGA-LUAD | 50.6354563 | 6.2868732 |
| TCGA-78-7162 | TCGA-LUAD | 36.4549898 | 16.0819966 |
| TCGA-78-7163 | TCGA-LUAD | 40.7582345 | 30.3127174 |
| TCGA-78-7166 | TCGA-LUAD | 41.8520153 | 34.4938866 |

| | | | |
|---|---|---|---|
| TCGA-78-7167 | TCGA-LUAD | 47.3212782 | 21.6276322 |
| TCGA-78-7220 | TCGA-LUAD | 63.6804678 | 67.6799883 |
| TCGA-78-7535 | TCGA-LUAD | 80.7268424 | 24.1155325 |
| TCGA-78-7536 | TCGA-LUAD | 54.1972245 | 72.1846843 |
| TCGA-78-7537 | TCGA-LUAD | 41.9355175 | 24.6870237 |
| TCGA-78-7539 | TCGA-LUAD | 52.8981646 | 19.7922364 |
| TCGA-78-7540 | TCGA-LUAD | 43.3951002 | 19.8513538 |
| TCGA-78-7542 | TCGA-LUAD | 82.2594057 | 29.5321013 |
| TCGA-78-7633 | TCGA-LUAD | 35.8984047 | 20.2952951 |
| TCGA-78-8640 | TCGA-LUAD | 55.0081678 | 18.2604046 |
| TCGA-78-8648 | TCGA-LUAD | 26.483342 | 11.2613242 |
| TCGA-78-8655 | TCGA-LUAD | 66.3287903 | 30.7002353 |
| TCGA-78-8660 | TCGA-LUAD | 45.7345315 | 24.4881507 |
| TCGA-78-8662 | TCGA-LUAD | 63.9908385 | 15.9106651 |
| TCGA-79-5596 | TCGA-LUSC | 16.2304958 | 5.77197068 |
| TCGA-80-5607 | TCGA-LUAD | 58.0521093 | 33.4633787 |
| TCGA-80-5608 | TCGA-LUAD | 36.4057823 | 32.4805579 |
| TCGA-80-5611 | TCGA-LUAD | 69.862703 | 29.7269891 |
| TCGA-83-5908 | TCGA-LUAD | 109.251706 | 15.9501417 |
| TCGA-85-6175 | TCGA-LUSC | 6.68863522 | 21.3482187 |
| TCGA-85-6560 | TCGA-LUSC | 100.098826 | 44.9675007 |
| TCGA-85-6561 | TCGA-LUSC | 24.0865535 | 15.4181084 |
| TCGA-85-6798 | TCGA-LUSC | 16.4668185 | 3.77162497 |
| TCGA-85-7696 | TCGA-LUSC | 41.3831798 | 27.2151715 |
| TCGA-85-7697 | TCGA-LUSC | 77.4138514 | 29.0950126 |
| TCGA-85-7698 | TCGA-LUSC | 63.7843788 | 29.7488589 |
| TCGA-85-7699 | TCGA-LUSC | 93.4660126 | 51.6446655 |
| TCGA-85-7710 | TCGA-LUSC | 188.473753 | 14.5075248 |
| TCGA-85-7843 | TCGA-LUSC | 47.1781723 | 15.0680403 |
| TCGA-85-7844 | TCGA-LUSC | 45.7469006 | 7.42058898 |
| TCGA-85-7950 | TCGA-LUSC | 82.1859806 | 28.7687782 |

| | | | |
|---|---|---|---|
| TCGA-85-8048 | TCGA-LUSC | 66.9906178 | 37.4211665 |
| TCGA-85-8049 | TCGA-LUSC | 21.4558818 | 13.3958949 |
| TCGA-85-8052 | TCGA-LUSC | 131.965418 | 5.89725443 |
| TCGA-85-8071 | TCGA-LUSC | 92.7620458 | 21.7544723 |
| TCGA-85-8072 | TCGA-LUSC | 9.53099139 | 35.9487328 |
| TCGA-85-8276 | TCGA-LUSC | 14.1621357 | 3.33024643 |
| TCGA-85-8277 | TCGA-LUSC | 59.3036457 | 38.2558653 |
| TCGA-85-8287 | TCGA-LUSC | 19.7304877 | 4.06818125 |
| TCGA-85-8288 | TCGA-LUSC | 84.1844186 | 21.5257613 |
| TCGA-85-8350 | TCGA-LUSC | 65.8966773 | 7.03130864 |
| TCGA-85-8351 | TCGA-LUSC | 79.1323271 | 9.51817368 |
| TCGA-85-8352 | TCGA-LUSC | 88.6197772 | 30.6492375 |
| TCGA-85-8353 | TCGA-LUSC | 35.2975407 | 10.80593 |
| TCGA-85-8354 | TCGA-LUSC | 61.6850351 | 19.7894157 |
| TCGA-85-8355 | TCGA-LUSC | 64.5299202 | 11.7573996 |
| TCGA-85-8479 | TCGA-LUSC | 95.7599294 | 27.2875746 |
| TCGA-85-8481 | TCGA-LUSC | 18.6225825 | 27.7949256 |
| TCGA-85-8580 | TCGA-LUSC | 62.9455693 | 35.439904 |
| TCGA-85-8582 | TCGA-LUSC | 41.4963025 | 22.3500321 |
| TCGA-85-8584 | TCGA-LUSC | 41.5219962 | 3.63820788 |
| TCGA-85-8664 | TCGA-LUSC | 50.6983018 | 135.218178 |
| TCGA-85-8666 | TCGA-LUSC | 61.9893701 | 66.4720033 |
| TCGA-85-A4CL | TCGA-LUSC | 48.3681583 | 28.4166797 |
| TCGA-85-A4CN | TCGA-LUSC | 39.7658779 | 26.1387196 |
| TCGA-85-A4JB | TCGA-LUSC | 73.312305 | 38.3254727 |
| TCGA-85-A4PA | TCGA-LUSC | 103.883941 | 5.60549435 |
| TCGA-85-A4QQ | TCGA-LUSC | 52.676006 | 14.7068614 |
| TCGA-85-A4QR | TCGA-LUSC | 86.6844489 | 42.8436732 |
| TCGA-85-A50M | TCGA-LUSC | 40.1765008 | 5.82932053 |
| TCGA-85-A50Z | TCGA-LUSC | 56.4656681 | 22.6441752 |
| TCGA-85-A510 | TCGA-LUSC | 56.5339768 | 7.42532873 |

| | | | |
|---|---|---|---|
| TCGA-85-A511 | TCGA-LUSC | 52.0388156 | 71.9431028 |
| TCGA-85-A513 | TCGA-LUSC | 24.9904879 | 8.15773726 |
| TCGA-85-A53L | TCGA-LUSC | 36.6544723 | 31.5033904 |
| TCGA-85-A5B5 | TCGA-LUSC | 70.4004574 | 15.2005761 |
| TCGA-86-6562 | TCGA-LUAD | 34.1876659 | 28.9346631 |
| TCGA-86-6851 | TCGA-LUAD | 29.0039289 | 12.5752098 |
| TCGA-86-7701 | TCGA-LUAD | 39.5692763 | 23.5289396 |
| TCGA-86-7711 | TCGA-LUAD | 48.9062551 | 31.5329308 |
| TCGA-86-7713 | TCGA-LUAD | 77.592743 | 50.1578532 |
| TCGA-86-7714 | TCGA-LUAD | 42.8806479 | 19.1719429 |
| TCGA-86-7953 | TCGA-LUAD | 71.9021292 | 37.1629605 |
| TCGA-86-7954 | TCGA-LUAD | 70.5211362 | 41.6446838 |
| TCGA-86-7955 | TCGA-LUAD | 43.0177136 | 11.7998207 |
| TCGA-86-8054 | TCGA-LUAD | 83.3971939 | 40.317628 |
| TCGA-86-8055 | TCGA-LUAD | 44.5932381 | 22.4176998 |
| TCGA-86-8056 | TCGA-LUAD | 49.8788363 | 28.8044457 |
| TCGA-86-8073 | TCGA-LUAD | 64.8557776 | 14.4475769 |
| TCGA-86-8074 | TCGA-LUAD | 92.6340237 | 25.6239831 |
| TCGA-86-8075 | TCGA-LUAD | 61.9997186 | 30.968364 |
| TCGA-86-8076 | TCGA-LUAD | 32.4254532 | 12.7707673 |
| TCGA-86-8278 | TCGA-LUAD | 32.5506472 | 23.549793 |
| TCGA-86-8279 | TCGA-LUAD | 81.7372661 | 25.2960216 |
| TCGA-86-8280 | TCGA-LUAD | 67.563873 | 23.059672 |
| TCGA-86-8281 | TCGA-LUAD | 55.3166879 | 24.1829704 |
| TCGA-86-8358 | TCGA-LUAD | 90.3688949 | 17.7958442 |
| TCGA-86-8359 | TCGA-LUAD | 51.4949847 | 31.6017306 |
| TCGA-86-8585 | TCGA-LUAD | 41.8123139 | 37.8490687 |
| TCGA-86-8668 | TCGA-LUAD | 37.5880317 | 14.1969567 |
| TCGA-86-8669 | TCGA-LUAD | 40.2610217 | 15.2041263 |
| TCGA-86-8671 | TCGA-LUAD | 32.900144 | 12.1974462 |
| TCGA-86-8672 | TCGA-LUAD | 65.831788 | 29.3715428 |

| | | | |
|---|---|---|---|
| TCGA-86-8673 | TCGA-LUAD | 64.9054956 | 52.8163766 |
| TCGA-86-8674 | TCGA-LUAD | 58.8384814 | 40.4185893 |
| TCGA-86-A456 | TCGA-LUAD | 60.1441273 | 27.5287838 |
| TCGA-86-A4D0 | TCGA-LUAD | 103.179514 | 47.0866886 |
| TCGA-86-A4JF | TCGA-LUAD | 57.3947448 | 42.1240212 |
| TCGA-86-A4P7 | TCGA-LUAD | 39.6824111 | 16.4005505 |
| TCGA-86-A4P8 | TCGA-LUAD | 36.5340642 | 11.8941965 |
| TCGA-90-6837 | TCGA-LUSC | 31.9219929 | 20.1510281 |
| TCGA-90-7766 | TCGA-LUSC | 66.9295162 | 27.6254382 |
| TCGA-90-7767 | TCGA-LUSC | 43.1004543 | 52.7098409 |
| TCGA-90-7769 | TCGA-LUSC | 47.6449817 | 41.1814648 |
| TCGA-90-7964 | TCGA-LUSC | 119.478072 | 31.7600043 |
| TCGA-90-A4ED | TCGA-LUSC | 41.8791529 | 14.8712871 |
| TCGA-90-A4EE | TCGA-LUSC | 40.9173012 | 38.5374457 |
| TCGA-91-6828 | TCGA-LUAD | 39.3052613 | 13.9947608 |
| TCGA-91-6829 | TCGA-LUAD | 54.1605371 | 35.6530474 |
| TCGA-91-6830 | TCGA-LUAD | 40.2677498 | 37.0416942 |
| TCGA-91-6831 | TCGA-LUAD | 59.5993307 | 20.4866769 |
| TCGA-91-6835 | TCGA-LUAD | 50.9591311 | 13.8549047 |
| TCGA-91-6836 | TCGA-LUAD | 44.4615925 | 9.58476397 |
| TCGA-91-6840 | TCGA-LUAD | 96.7452251 | 20.8994843 |
| TCGA-91-6847 | TCGA-LUAD | 119.796571 | 26.9080584 |
| TCGA-91-6848 | TCGA-LUAD | 108.637159 | 52.751924 |
| TCGA-91-6849 | TCGA-LUAD | 47.2398108 | 14.6183752 |
| TCGA-91-7771 | TCGA-LUAD | 44.9379969 | 27.8202364 |
| TCGA-91-8496 | TCGA-LUAD | 67.9355141 | 11.2338586 |
| TCGA-91-8499 | TCGA-LUAD | 75.1329007 | 57.7298146 |
| TCGA-91-A4BC | TCGA-LUAD | 30.5349554 | 14.8773332 |
| TCGA-91-A4BD | TCGA-LUAD | 39.4847904 | 19.007655 |
| TCGA-92-7340 | TCGA-LUSC | 66.9026901 | 20.3970209 |
| TCGA-92-7341 | TCGA-LUSC | 32.1556703 | 26.9704095 |

| | | | |
|---|---|---|---|
| TCGA-92-8063 | TCGA-LUSC | 95.0738076 | 30.0646995 |
| TCGA-92-8064 | TCGA-LUSC | 12.4445383 | 3.61931797 |
| TCGA-92-8065 | TCGA-LUSC | 12.4871747 | 1.34772358 |
| TCGA-93-7347 | TCGA-LUAD | 34.374106 | 16.2852217 |
| TCGA-93-7348 | TCGA-LUAD | 30.3323758 | 25.2561455 |
| TCGA-93-8067 | TCGA-LUAD | 51.1572437 | 26.6855419 |
| TCGA-93-A4JN | TCGA-LUAD | 61.372229 | 18.2824071 |
| TCGA-93-A4JO | TCGA-LUAD | 38.0520474 | 32.1987737 |
| TCGA-93-A4JP | TCGA-LUAD | 45.1264327 | 22.8328767 |
| TCGA-93-A4JQ | TCGA-LUAD | 53.4434404 | 24.2083382 |
| TCGA-94-7033 | TCGA-LUSC | 12.6728071 | 13.0269862 |
| TCGA-94-7557 | TCGA-LUSC | 82.4981889 | 69.9804528 |
| TCGA-94-7943 | TCGA-LUSC | 22.7851443 | 42.7373284 |
| TCGA-94-8035 | TCGA-LUSC | 145.335124 | 54.5894778 |
| TCGA-94-8490 | TCGA-LUSC | 53.0943399 | 23.0845158 |
| TCGA-94-8491 | TCGA-LUSC | 47.9020449 | 11.6553665 |
| TCGA-94-A4VJ | TCGA-LUSC | 49.476011 | 32.9172228 |
| TCGA-94-A5I4 | TCGA-LUSC | 42.6791856 | 19.9885009 |
| TCGA-94-A5I6 | TCGA-LUSC | 43.3323417 | 19.3435921 |
| TCGA-95-7039 | TCGA-LUAD | 61.7282796 | 45.981295 |
| TCGA-95-7043 | TCGA-LUAD | 35.450274 | 46.5420238 |
| TCGA-95-7562 | TCGA-LUAD | 99.4359898 | 27.2546054 |
| TCGA-95-7567 | TCGA-LUAD | 55.6728981 | 35.6763547 |
| TCGA-95-7944 | TCGA-LUAD | 65.8429375 | 28.9317238 |
| TCGA-95-7947 | TCGA-LUAD | 65.7744367 | 18.9256744 |
| TCGA-95-7948 | TCGA-LUAD | 31.4521921 | 28.0710982 |
| TCGA-95-8039 | TCGA-LUAD | 58.0967868 | 18.9994129 |
| TCGA-95-8494 | TCGA-LUAD | 56.0016155 | 16.9355354 |
| TCGA-95-A4VK | TCGA-LUAD | 44.8458015 | 29.064122 |
| TCGA-95-A4VN | TCGA-LUAD | 59.057682 | 21.6472712 |
| TCGA-95-A4VP | TCGA-LUAD | 38.9871513 | 33.6343379 |

| | | | |
|---|---|---|---|
| TCGA-96-7544 | TCGA-LUSC | 30.6680074 | 20.3041259 |
| TCGA-96-7545 | TCGA-LUSC | 25.2687319 | 11.0613825 |
| TCGA-96-8169 | TCGA-LUSC | 59.5017598 | 19.7297752 |
| TCGA-96-8170 | TCGA-LUSC | 47.7427348 | 25.9497287 |
| TCGA-96-A4JK | TCGA-LUSC | 43.0106489 | 19.0831904 |
| TCGA-96-A4JL | TCGA-LUSC | 28.1410036 | 34.9267491 |
| TCGA-97-7546 | TCGA-LUAD | 79.3090082 | 17.3648068 |
| TCGA-97-7547 | TCGA-LUAD | 33.3636205 | 13.8016321 |
| TCGA-97-7552 | TCGA-LUAD | 30.7753469 | 12.1514092 |
| TCGA-97-7553 | TCGA-LUAD | 54.1637714 | 18.3080321 |
| TCGA-97-7554 | TCGA-LUAD | 28.1836217 | 11.2957642 |
| TCGA-97-7937 | TCGA-LUAD | 72.0477804 | 23.3841377 |
| TCGA-97-7938 | TCGA-LUAD | 63.5845346 | 16.0971425 |
| TCGA-97-7941 | TCGA-LUAD | 41.6066368 | 19.1667652 |
| TCGA-97-8171 | TCGA-LUAD | 41.5567565 | 9.18516858 |
| TCGA-97-8172 | TCGA-LUAD | 38.8865286 | 16.3051674 |
| TCGA-97-8174 | TCGA-LUAD | 59.2563323 | 17.4203302 |
| TCGA-97-8175 | TCGA-LUAD | 50.1779666 | 20.3256121 |
| TCGA-97-8176 | TCGA-LUAD | 61.9155494 | 44.742023 |
| TCGA-97-8177 | TCGA-LUAD | 48.2192677 | 19.0401703 |
| TCGA-97-8179 | TCGA-LUAD | 47.222185 | 59.1637658 |
| TCGA-97-8547 | TCGA-LUAD | 51.0601779 | 16.5012476 |
| TCGA-97-8552 | TCGA-LUAD | 38.1607245 | 17.7470531 |
| TCGA-97-A4LX | TCGA-LUAD | 51.0265592 | 17.2705627 |
| TCGA-97-A4M0 | TCGA-LUAD | 51.1131502 | 22.8718117 |
| TCGA-97-A4M1 | TCGA-LUAD | 60.4615497 | 16.6808384 |
| TCGA-97-A4M2 | TCGA-LUAD | 32.203771 | 12.9102116 |
| TCGA-97-A4M3 | TCGA-LUAD | 45.8647399 | 11.0521112 |
| TCGA-97-A4M5 | TCGA-LUAD | 24.6075451 | 19.197102 |
| TCGA-97-A4M6 | TCGA-LUAD | 60.6030838 | 12.6663966 |
| TCGA-97-A4M7 | TCGA-LUAD | 38.2417092 | 19.5965354 |

| | | | |
|---|---|---|---|
| TCGA-98-7454 | TCGA-LUSC | 18.4725173 | 20.35395 |
| TCGA-98-8020 | TCGA-LUSC | 26.4844381 | 12.2068543 |
| TCGA-98-8021 | TCGA-LUSC | 95.0771291 | 8.96995491 |
| TCGA-98-8022 | TCGA-LUSC | 56.7782508 | 15.1290724 |
| TCGA-98-8023 | TCGA-LUSC | 36.1047352 | 31.2697192 |
| TCGA-98-A538 | TCGA-LUSC | 60.1282146 | 34.8621857 |
| TCGA-98-A539 | TCGA-LUSC | 75.7378509 | 6.60110425 |
| TCGA-98-A53A | TCGA-LUSC | 87.2847759 | 30.4158152 |
| TCGA-98-A53B | TCGA-LUSC | 81.8385443 | 8.64954569 |
| TCGA-98-A53C | TCGA-LUSC | 26.9176646 | 11.6022099 |
| TCGA-98-A53D | TCGA-LUSC | 25.7228433 | 3.49493098 |
| TCGA-98-A53H | TCGA-LUSC | 33.0764122 | 14.4675002 |
| TCGA-98-A53I | TCGA-LUSC | 19.8342526 | 4.6903462 |
| TCGA-98-A53J | TCGA-LUSC | 11.7946914 | 41.251491 |
| TCGA-99-7458 | TCGA-LUAD | 49.0362415 | 21.6804953 |
| TCGA-99-8025 | TCGA-LUAD | 46.262229 | 25.1536786 |
| TCGA-99-8028 | TCGA-LUAD | 38.2701076 | 13.4934829 |
| TCGA-99-8032 | TCGA-LUAD | 47.5783033 | 13.8258132 |
| TCGA-99-8033 | TCGA-LUAD | 66.6343783 | 60.4489417 |
| TCGA-99-AA5R | TCGA-LUAD | 37.1487728 | 10.3638679 |
| TCGA-A1-A0SB | TCGA-BRCA | 43.0969459 | 16.8404315 |
| TCGA-A1-A0SD | TCGA-BRCA | 43.7329119 | 29.3015223 |
| TCGA-A1-A0SE | TCGA-BRCA | 55.8589752 | 26.7655469 |
| TCGA-A1-A0SF | TCGA-BRCA | 70.8244136 | 32.3979083 |
| TCGA-A1-A0SG | TCGA-BRCA | 46.3767845 | 22.8393179 |
| TCGA-A1-A0SH | TCGA-BRCA | 40.3623541 | 19.1328075 |
| TCGA-A1-A0SI | TCGA-BRCA | 73.0557374 | 26.1925537 |
| TCGA-A1-A0SJ | TCGA-BRCA | 50.9364132 | 75.0784621 |
| TCGA-A1-A0SK | TCGA-BRCA | 107.774118 | 133.797979 |
| TCGA-A1-A0SM | TCGA-BRCA | 64.6761908 | 24.3306774 |
| TCGA-A1-A0SN | TCGA-BRCA | 41.1382699 | 50.4332754 |

| | | | |
|---|---|---|---|
| TCGA-A1-A0SO | TCGA-BRCA | 162.526154 | 72.5554417 |
| TCGA-A1-A0SP | TCGA-BRCA | 96.6456424 | 170.757541 |
| TCGA-A1-A0SQ | TCGA-BRCA | 38.505424 | 29.4280785 |
| TCGA-A2-A04N | TCGA-BRCA | 55.9947737 | 25.7005849 |
| TCGA-A2-A04P | TCGA-BRCA | 114.386297 | 49.3159684 |
| TCGA-A2-A04Q | TCGA-BRCA | 54.1200968 | 52.2053032 |
| TCGA-A2-A04R | TCGA-BRCA | 38.7797972 | 44.9095604 |
| TCGA-A2-A04T | TCGA-BRCA | 116.084946 | 86.5536633 |
| TCGA-A2-A04U | TCGA-BRCA | 164.699752 | 99.3657123 |
| TCGA-A2-A04V | TCGA-BRCA | 44.7949071 | 20.1666397 |
| TCGA-A2-A04W | TCGA-BRCA | 54.0297807 | 32.4677275 |
| TCGA-A2-A04X | TCGA-BRCA | 61.891977 | 39.9524254 |
| TCGA-A2-A04Y | TCGA-BRCA | 44.6133998 | 28.3290518 |
| TCGA-A2-A0CK | TCGA-BRCA | 50.9289966 | 19.7545338 |
| TCGA-A2-A0CL | TCGA-BRCA | 51.9347656 | 36.1986288 |
| TCGA-A2-A0CM | TCGA-BRCA | 174.385644 | 62.3290862 |
| TCGA-A2-A0CO | TCGA-BRCA | 47.8768746 | 13.7725359 |
| TCGA-A2-A0CP | TCGA-BRCA | 46.3199293 | 18.9592375 |
| TCGA-A2-A0CQ | TCGA-BRCA | 83.4766132 | 25.9997587 |
| TCGA-A2-A0CR | TCGA-BRCA | 54.3281019 | 15.3500622 |
| TCGA-A2-A0CS | TCGA-BRCA | 46.9232275 | 40.4359199 |
| TCGA-A2-A0CT | TCGA-BRCA | 49.4845585 | 48.7019564 |
| TCGA-A2-A0CU | TCGA-BRCA | 44.5769492 | 20.0986429 |
| TCGA-A2-A0CV | TCGA-BRCA | 61.947012 | 21.7609585 |
| TCGA-A2-A0CW | TCGA-BRCA | 63.1874662 | 34.6467964 |
| TCGA-A2-A0CX | TCGA-BRCA | 115.954134 | 38.2563785 |
| TCGA-A2-A0CY | TCGA-BRCA | 23.4611327 | 21.7723337 |
| TCGA-A2-A0CZ | TCGA-BRCA | 37.0240691 | 17.0424018 |
| TCGA-A2-A0D0 | TCGA-BRCA | 239.933733 | 51.4169561 |
| TCGA-A2-A0D1 | TCGA-BRCA | 52.1571968 | 18.4971821 |
| TCGA-A2-A0D2 | TCGA-BRCA | 209.165738 | 43.122013 |

| | | | |
|---|---|---|---|
| TCGA-A2-A0D3 | TCGA-BRCA | 54.476084 | 28.9348247 |
| TCGA-A2-A0D4 | TCGA-BRCA | 51.2379226 | 21.2365967 |
| TCGA-A2-A0EM | TCGA-BRCA | 49.4029653 | 28.1110464 |
| TCGA-A2-A0EN | TCGA-BRCA | 35.6687811 | 25.5081443 |
| TCGA-A2-A0EO | TCGA-BRCA | 47.2452989 | 24.835951 |
| TCGA-A2-A0EP | TCGA-BRCA | 44.9850754 | 21.2537618 |
| TCGA-A2-A0EQ | TCGA-BRCA | 49.659633 | 25.2672531 |
| TCGA-A2-A0ER | TCGA-BRCA | 43.8311797 | 25.5472057 |
| TCGA-A2-A0ES | TCGA-BRCA | 51.0861689 | 20.4416891 |
| TCGA-A2-A0ET | TCGA-BRCA | 59.2499332 | 39.7132591 |
| TCGA-A2-A0EU | TCGA-BRCA | 60.3314133 | 55.5586247 |
| TCGA-A2-A0EV | TCGA-BRCA | 51.7494652 | 21.2977577 |
| TCGA-A2-A0EW | TCGA-BRCA | 49.9533081 | 17.9522391 |
| TCGA-A2-A0EX | TCGA-BRCA | 99.8859657 | 40.1511951 |
| TCGA-A2-A0EY | TCGA-BRCA | 28.1118829 | 29.510021 |
| TCGA-A2-A0ST | TCGA-BRCA | 54.3963839 | 26.9582137 |
| TCGA-A2-A0SU | TCGA-BRCA | 48.6175967 | 25.5545847 |
| TCGA-A2-A0SV | TCGA-BRCA | 58.4155817 | 51.2467192 |
| TCGA-A2-A0SW | TCGA-BRCA | 44.8636683 | 28.2075684 |
| TCGA-A2-A0SX | TCGA-BRCA | 67.3786006 | 21.2793084 |
| TCGA-A2-A0SY | TCGA-BRCA | 34.4386802 | 24.1755247 |
| TCGA-A2-A0T0 | TCGA-BRCA | 128.079862 | 28.2855769 |
| TCGA-A2-A0T1 | TCGA-BRCA | 38.8379382 | 23.8621691 |
| TCGA-A2-A0T2 | TCGA-BRCA | 115.546088 | 47.2803709 |
| TCGA-A2-A0T3 | TCGA-BRCA | 71.4691977 | 39.2658579 |
| TCGA-A2-A0T4 | TCGA-BRCA | 39.8236177 | 42.3787445 |
| TCGA-A2-A0T5 | TCGA-BRCA | 47.9570737 | 25.7446906 |
| TCGA-A2-A0T6 | TCGA-BRCA | 48.0602272 | 16.6978958 |
| TCGA-A2-A0T7 | TCGA-BRCA | 69.2074747 | 22.1929628 |
| TCGA-A2-A0YC | TCGA-BRCA | 47.33702 | 24.4468193 |
| TCGA-A2-A0YD | TCGA-BRCA | 54.0419736 | 31.9844064 |

| | | | |
|---|---|---|---|
| TCGA-A2-A0YE | TCGA-BRCA | 81.9158233 | 47.5669015 |
| TCGA-A2-A0YF | TCGA-BRCA | 33.899328 | 27.0170339 |
| TCGA-A2-A0YG | TCGA-BRCA | 54.3941797 | 52.8121225 |
| TCGA-A2-A0YH | TCGA-BRCA | 40.8439183 | 48.2512961 |
| TCGA-A2-A0YI | TCGA-BRCA | 66.5860188 | 31.9326712 |
| TCGA-A2-A0YJ | TCGA-BRCA | 55.0021988 | 17.0217563 |
| TCGA-A2-A0YK | TCGA-BRCA | 47.6117058 | 18.8061943 |
| TCGA-A2-A0YL | TCGA-BRCA | 50.8761735 | 25.6525242 |
| TCGA-A2-A0YM | TCGA-BRCA | 80.3360524 | 81.1674613 |
| TCGA-A2-A0YT | TCGA-BRCA | 44.4125703 | 33.6627578 |
| TCGA-A2-A1FV | TCGA-BRCA | 33.6406109 | 24.8578902 |
| TCGA-A2-A1FW | TCGA-BRCA | 34.5143226 | 29.2601675 |
| TCGA-A2-A1FX | TCGA-BRCA | 59.3000248 | 34.6479244 |
| TCGA-A2-A1FZ | TCGA-BRCA | 43.3066087 | 21.3583138 |
| TCGA-A2-A1G0 | TCGA-BRCA | 51.7846637 | 13.0148982 |
| TCGA-A2-A1G1 | TCGA-BRCA | 80.6723945 | 83.3439532 |
| TCGA-A2-A1G4 | TCGA-BRCA | 46.7814303 | 13.9052403 |
| TCGA-A2-A1G6 | TCGA-BRCA | 65.9455826 | 23.4871615 |
| TCGA-A2-A259 | TCGA-BRCA | 52.182499 | 17.1942724 |
| TCGA-A2-A25A | TCGA-BRCA | 67.0585115 | 27.5546452 |
| TCGA-A2-A25B | TCGA-BRCA | 65.6975001 | 27.8006341 |
| TCGA-A2-A25C | TCGA-BRCA | 40.8859605 | 31.4635011 |
| TCGA-A2-A25D | TCGA-BRCA | 38.4423401 | 19.3557468 |
| TCGA-A2-A25E | TCGA-BRCA | 67.0576142 | 42.6701861 |
| TCGA-A2-A25F | TCGA-BRCA | 89.8751954 | 36.8923982 |
| TCGA-A2-A3KC | TCGA-BRCA | 35.997375 | 29.3502902 |
| TCGA-A2-A3KD | TCGA-BRCA | 39.0725625 | 22.6331551 |
| TCGA-A2-A3XS | TCGA-BRCA | 83.8510877 | 36.6004266 |
| TCGA-A2-A3XT | TCGA-BRCA | 86.1712649 | 23.3563826 |
| TCGA-A2-A3XU | TCGA-BRCA | 100.28281 | 37.3791193 |
| TCGA-A2-A3XV | TCGA-BRCA | 43.3743541 | 19.212657 |

| | | | |
|---|---|---|---|
| TCGA-A2-A3XW | TCGA-BRCA | 49.7556151 | 15.489461 |
| TCGA-A2-A3XX | TCGA-BRCA | 66.7803791 | 36.4450013 |
| TCGA-A2-A3XY | TCGA-BRCA | 76.5406663 | 50.9980033 |
| TCGA-A2-A3XZ | TCGA-BRCA | 54.0865585 | 27.3436533 |
| TCGA-A2-A3Y0 | TCGA-BRCA | 154.160201 | 61.9647961 |
| TCGA-A2-A4RW | TCGA-BRCA | 69.4569684 | 23.6662171 |
| TCGA-A2-A4RX | TCGA-BRCA | 39.2289839 | 16.5190566 |
| TCGA-A2-A4RY | TCGA-BRCA | 43.4900033 | 15.8517107 |
| TCGA-A2-A4S0 | TCGA-BRCA | 35.7210279 | 19.0602425 |
| TCGA-A2-A4S1 | TCGA-BRCA | 72.5609412 | 21.2338329 |
| TCGA-A2-A4S2 | TCGA-BRCA | 26.1139255 | 30.9513155 |
| TCGA-A2-A4S3 | TCGA-BRCA | 35.0797359 | 43.2941594 |
| TCGA-A3-3306 | TCGA-KIRC | 37.3006265 | 14.9335104 |
| TCGA-A3-3307 | TCGA-KIRC | 58.0884046 | 20.802836 |
| TCGA-A3-3308 | TCGA-KIRC | 37.0982755 | 14.6675269 |
| TCGA-A3-3311 | TCGA-KIRC | 27.8150894 | 17.9390418 |
| TCGA-A3-3313 | TCGA-KIRC | 14.0336332 | 9.41940281 |
| TCGA-A3-3316 | TCGA-KIRC | 31.7918392 | 17.476976 |
| TCGA-A3-3317 | TCGA-KIRC | 38.2577634 | 18.8687425 |
| TCGA-A3-3319 | TCGA-KIRC | 25.3242292 | 31.6679948 |
| TCGA-A3-3320 | TCGA-KIRC | 45.1719641 | 15.9690911 |
| TCGA-A3-3322 | TCGA-KIRC | 26.2629212 | 11.7690441 |
| TCGA-A3-3323 | TCGA-KIRC | 27.9040448 | 11.2230302 |
| TCGA-A3-3324 | TCGA-KIRC | 32.163425 | 13.5667703 |
| TCGA-A3-3325 | TCGA-KIRC | 43.738425 | 17.7416804 |
| TCGA-A3-3326 | TCGA-KIRC | 38.4261464 | 14.4991856 |
| TCGA-A3-3328 | TCGA-KIRC | 28.5528525 | 29.4314085 |
| TCGA-A3-3329 | TCGA-KIRC | 62.5751286 | 18.7650607 |
| TCGA-A3-3331 | TCGA-KIRC | 31.7412238 | 12.4237306 |
| TCGA-A3-3335 | TCGA-KIRC | 27.2853233 | 10.0029369 |
| TCGA-A3-3343 | TCGA-KIRC | 33.1550436 | 10.0972524 |

| | | | |
|---|---|---|---|
| TCGA-A3-3346 | TCGA-KIRC | 64.7025287 | 38.8041678 |
| TCGA-A3-3347 | TCGA-KIRC | 48.4169637 | 26.0357278 |
| TCGA-A3-3349 | TCGA-KIRC | 39.1184004 | 13.9699137 |
| TCGA-A3-3351 | TCGA-KIRC | 40.5127734 | 13.8060814 |
| TCGA-A3-3352 | TCGA-KIRC | 47.7026058 | 13.8352587 |
| TCGA-A3-3357 | TCGA-KIRC | 34.4334709 | 12.8407722 |
| TCGA-A3-3358 | TCGA-KIRC | 31.2223636 | 9.72859174 |
| TCGA-A3-3359 | TCGA-KIRC | 31.0537062 | 9.11187075 |
| TCGA-A3-3362 | TCGA-KIRC | 38.1467574 | 14.0740012 |
| TCGA-A3-3363 | TCGA-KIRC | 48.3658157 | 15.6603688 |
| TCGA-A3-3365 | TCGA-KIRC | 63.8291134 | 17.2881501 |
| TCGA-A3-3367 | TCGA-KIRC | 34.4262721 | 12.27546 |
| TCGA-A3-3370 | TCGA-KIRC | 39.0120783 | 12.6590154 |
| TCGA-A3-3372 | TCGA-KIRC | 42.9597104 | 18.4053195 |
| TCGA-A3-3373 | TCGA-KIRC | 28.3251508 | 17.2088526 |
| TCGA-A3-3374 | TCGA-KIRC | 12.8191742 | 12.0148378 |
| TCGA-A3-3376 | TCGA-KIRC | 40.6127851 | 11.6737781 |
| TCGA-A3-3378 | TCGA-KIRC | 25.0515392 | 14.7151479 |
| TCGA-A3-3380 | TCGA-KIRC | 34.9073841 | 12.2474773 |
| TCGA-A3-3382 | TCGA-KIRC | 34.8591731 | 24.1580717 |
| TCGA-A3-3383 | TCGA-KIRC | 54.2978259 | 16.9660494 |
| TCGA-A3-3385 | TCGA-KIRC | 28.2264952 | 9.65584161 |
| TCGA-A3-3387 | TCGA-KIRC | 46.1126508 | 25.9016314 |
| TCGA-A3-A6NI | TCGA-KIRC | 47.4332742 | 30.0902754 |
| TCGA-A3-A6NJ | TCGA-KIRC | 56.9312209 | 14.1339269 |
| TCGA-A3-A6NL | TCGA-KIRC | 54.7722927 | 13.4683736 |
| TCGA-A3-A6NN | TCGA-KIRC | 47.5090714 | 19.1265735 |
| TCGA-A3-A8CQ | TCGA-KIRC | 59.0867518 | 15.8376186 |
| TCGA-A3-A8OU | TCGA-KIRC | 59.5584394 | 17.7155609 |
| TCGA-A3-A8OV | TCGA-KIRC | 58.4969992 | 14.3349601 |
| TCGA-A3-A8OW | TCGA-KIRC | 68.2216985 | 19.2199056 |

| | | | |
|---|---|---|---|
| TCGA-A3-A8OX | TCGA-KIRC | 47.1146551 | 19.7637378 |
| TCGA-A4-7286 | TCGA-KIRP | 15.9710693 | 12.4178859 |
| TCGA-A4-7287 | TCGA-KIRP | 89.6140957 | 54.9626632 |
| TCGA-A4-7288 | TCGA-KIRP | 35.1684251 | 12.5482173 |
| TCGA-A4-7583 | TCGA-KIRP | 4.29167832 | 42.9482659 |
| TCGA-A4-7584 | TCGA-KIRP | 13.654451 | 3.51326654 |
| TCGA-A4-7585 | TCGA-KIRP | 26.6048288 | 11.6934518 |
| TCGA-A4-7732 | TCGA-KIRP | 46.4861104 | 18.1862978 |
| TCGA-A4-7734 | TCGA-KIRP | 51.6453933 | 20.0060604 |
| TCGA-A4-7915 | TCGA-KIRP | 13.3441234 | 7.01993082 |
| TCGA-A4-7996 | TCGA-KIRP | 43.2205388 | 18.6415342 |
| TCGA-A4-7997 | TCGA-KIRP | 39.1296798 | 35.1672121 |
| TCGA-A4-8098 | TCGA-KIRP | 13.3253989 | 21.4596936 |
| TCGA-A4-8310 | TCGA-KIRP | 11.1455455 | 28.2778733 |
| TCGA-A4-8311 | TCGA-KIRP | 23.1387377 | 9.18155902 |
| TCGA-A4-8312 | TCGA-KIRP | 35.7815357 | 13.8252644 |
| TCGA-A4-8515 | TCGA-KIRP | 16.1136873 | 34.4794526 |
| TCGA-A4-8516 | TCGA-KIRP | 25.4937381 | 6.94880778 |
| TCGA-A4-8517 | TCGA-KIRP | 9.47332561 | 3.21762251 |
| TCGA-A4-8518 | TCGA-KIRP | 13.3701192 | 4.91035065 |
| TCGA-A4-8630 | TCGA-KIRP | 6.76960908 | 5.45440379 |
| TCGA-A4-A4ZT | TCGA-KIRP | 25.444827 | 15.3626144 |
| TCGA-A4-A57E | TCGA-KIRP | 37.4438002 | 16.0363253 |
| TCGA-A4-A5DU | TCGA-KIRP | 28.4240162 | 13.965225 |
| TCGA-A4-A5XZ | TCGA-KIRP | 28.5269272 | 16.5026211 |
| TCGA-A4-A5Y0 | TCGA-KIRP | 22.8267096 | 6.90016579 |
| TCGA-A4-A5Y1 | TCGA-KIRP | 34.5825295 | 20.9066252 |
| TCGA-A4-A6HP | TCGA-KIRP | 34.5123558 | 15.6320027 |
| TCGA-A4-A772 | TCGA-KIRP | 41.098422 | 22.5728004 |
| TCGA-A4-A7UZ | TCGA-KIRP | 43.632924 | 26.6628665 |
| TCGA-A5-A1OH | TCGA-UCEC | 80.3171624 | 39.9723362 |

| | | | |
|---|---|---|---|
| TCGA-A5-A2K2 | TCGA-UCEC | 95.1204887 | 28.109018 |
| TCGA-A5-A2K3 | TCGA-UCEC | 28.8800758 | 22.3004801 |
| TCGA-A5-A2K4 | TCGA-UCEC | 64.7285321 | 65.3068483 |
| TCGA-A5-A2K5 | TCGA-UCEC | 83.2929412 | 9.82792975 |
| TCGA-A5-A2K7 | TCGA-UCEC | 38.3825727 | 18.720851 |
| TCGA-A5-A3LO | TCGA-UCEC | 68.9784762 | 38.8718252 |
| TCGA-A5-A3LP | TCGA-UCEC | 75.5029003 | 28.337141 |
| TCGA-A5-A7WJ | TCGA-UCEC | 36.1416256 | 42.147782 |
| TCGA-A5-A7WK | TCGA-UCEC | 86.3422416 | 31.3227052 |
| TCGA-A5-AB3J | TCGA-UCEC | 34.648976 | 18.4765454 |
| TCGA-A6-2672 | TCGA-COAD | 14.3146176 | 5.69728547 |
| TCGA-A6-2674 | TCGA-COAD | 11.6418006 | 15.3610924 |
| TCGA-A6-2675 | TCGA-COAD | 33.7562035 | 10.4279723 |
| TCGA-A6-2683 | TCGA-COAD | 40.2779894 | 13.2608843 |
| TCGA-A6-2686 | TCGA-COAD | 71.8964198 | 20.9522706 |
| TCGA-A6-3809 | TCGA-COAD | 26.404652 | 24.3705285 |
| TCGA-A6-3810 | TCGA-COAD | 5.24409983 | 4.41627211 |
| TCGA-A6-4105 | TCGA-COAD | 73.4684374 | 47.316215 |
| TCGA-A6-5656 | TCGA-COAD | 64.5260277 | 21.757573 |
| TCGA-A6-5657 | TCGA-COAD | 86.2610672 | 32.6369664 |
| TCGA-A6-5659 | TCGA-COAD | 108.730983 | 39.0563955 |
| TCGA-A6-5660 | TCGA-COAD | 76.0040299 | 31.3024569 |
| TCGA-A6-5661 | TCGA-COAD | 96.3632075 | 35.6957868 |
| TCGA-A6-5662 | TCGA-COAD | 44.9936561 | 14.620621 |
| TCGA-A6-5664 | TCGA-COAD | 62.7584091 | 25.1786119 |
| TCGA-A6-5665 | TCGA-COAD | 29.0660202 | 5.78137517 |
| TCGA-A6-5666 | TCGA-COAD | 88.243291 | 33.6539955 |
| TCGA-A6-5667 | TCGA-COAD | 38.6937222 | 11.0934004 |
| TCGA-A6-6137 | TCGA-COAD | 60.3779732 | 16.9021404 |
| TCGA-A6-6138 | TCGA-COAD | 58.4137306 | 29.7013919 |
| TCGA-A6-6140 | TCGA-COAD | 89.9520841 | 39.7072665 |

| | | | |
|---|---|---|---|
| TCGA-A6-6141 | TCGA-COAD | 46.4682843 | 18.6526523 |
| TCGA-A6-6142 | TCGA-COAD | 57.5115757 | 26.6399781 |
| TCGA-A6-6648 | TCGA-COAD | 103.741255 | 32.4176055 |
| TCGA-A6-6649 | TCGA-COAD | 49.5781429 | 15.1561647 |
| TCGA-A6-6650 | TCGA-COAD | 26.2754909 | 33.2307779 |
| TCGA-A6-6651 | TCGA-COAD | 51.6731797 | 12.5815934 |
| TCGA-A6-6652 | TCGA-COAD | 47.9801761 | 8.61716564 |
| TCGA-A6-6653 | TCGA-COAD | 55.7219459 | 33.1243294 |
| TCGA-A6-6654 | TCGA-COAD | 56.5202511 | 30.6795869 |
| TCGA-A6-6780 | TCGA-COAD | 70.2503938 | 36.4736177 |
| TCGA-A6-6781 | TCGA-COAD | 11.2162209 | 7.90655678 |
| TCGA-A6-6782 | TCGA-COAD | 57.2300743 | 20.0752186 |
| TCGA-A6-A565 | TCGA-COAD | 51.082226 | 19.413732 |
| TCGA-A6-A566 | TCGA-COAD | 54.8828995 | 40.5788609 |
| TCGA-A6-A567 | TCGA-COAD | 68.9406809 | 25.2740603 |
| TCGA-A6-A56B | TCGA-COAD | 106.612686 | 24.1112224 |
| TCGA-A6-A5ZU | TCGA-COAD | 51.0084591 | 16.8499467 |
| TCGA-A7-A0CD | TCGA-BRCA | 49.5129603 | 29.8031425 |
| TCGA-A7-A0CE | TCGA-BRCA | 56.9114846 | 20.8187143 |
| TCGA-A7-A0CG | TCGA-BRCA | 37.1563745 | 37.1692641 |
| TCGA-A7-A0CH | TCGA-BRCA | 34.2212097 | 17.5796255 |
| TCGA-A7-A0CJ | TCGA-BRCA | 53.3654177 | 23.9758549 |
| TCGA-A7-A0D9 | TCGA-BRCA | 60.213354 | 34.6898205 |
| TCGA-A7-A0DA | TCGA-BRCA | 79.5571191 | 19.7871137 |
| TCGA-A7-A0DB | TCGA-BRCA | 41.0684967 | 25.4345114 |
| TCGA-A7-A0DC | TCGA-BRCA | 35.2591512 | 15.702426 |
| TCGA-A7-A13D | TCGA-BRCA | 89.7896804 | 47.5669761 |
| TCGA-A7-A13E | TCGA-BRCA | 28.2622345 | 13.5763602 |
| TCGA-A7-A13F | TCGA-BRCA | 51.1066557 | 38.8762131 |
| TCGA-A7-A13G | TCGA-BRCA | 12.0109709 | 5.94378102 |
| TCGA-A7-A13H | TCGA-BRCA | 42.1924631 | 20.4182809 |

| | | | |
|---|---|---|---|
| TCGA-A7-A26E | TCGA-BRCA | 8.78496261 | 14.3754257 |
| TCGA-A7-A26F | TCGA-BRCA | 20.7666186 | 5.55671704 |
| TCGA-A7-A26G | TCGA-BRCA | 47.5138687 | 21.8446132 |
| TCGA-A7-A26H | TCGA-BRCA | 38.4278781 | 28.3777916 |
| TCGA-A7-A26I | TCGA-BRCA | 17.70665 | 6.84153827 |
| TCGA-A7-A26J | TCGA-BRCA | 64.8159224 | 49.9180378 |
| TCGA-A7-A2KD | TCGA-BRCA | 94.2848033 | 24.6321888 |
| TCGA-A7-A3IY | TCGA-BRCA | 53.193678 | 26.2135712 |
| TCGA-A7-A3IZ | TCGA-BRCA | 40.8025703 | 20.4767679 |
| TCGA-A7-A3J0 | TCGA-BRCA | 38.0220951 | 21.6209348 |
| TCGA-A7-A3J1 | TCGA-BRCA | 34.8243476 | 26.5701798 |
| TCGA-A7-A3RF | TCGA-BRCA | 35.0545818 | 19.5490905 |
| TCGA-A7-A425 | TCGA-BRCA | 57.7445275 | 59.7721396 |
| TCGA-A7-A426 | TCGA-BRCA | 54.5607812 | 20.5417857 |
| TCGA-A7-A4SA | TCGA-BRCA | 54.9993562 | 16.4297387 |
| TCGA-A7-A4SB | TCGA-BRCA | 41.692796 | 31.5220471 |
| TCGA-A7-A4SC | TCGA-BRCA | 29.2150699 | 34.2422148 |
| TCGA-A7-A4SD | TCGA-BRCA | 88.9169764 | 50.7982894 |
| TCGA-A7-A4SE | TCGA-BRCA | 113.870934 | 38.8906775 |
| TCGA-A7-A4SF | TCGA-BRCA | 49.2892788 | 41.036475 |
| TCGA-A7-A56D | TCGA-BRCA | 29.3327612 | 24.2751153 |
| TCGA-A7-A5ZV | TCGA-BRCA | 142.837442 | 48.8579181 |
| TCGA-A7-A5ZW | TCGA-BRCA | 62.9184582 | 27.7651616 |
| TCGA-A7-A5ZX | TCGA-BRCA | 69.5872722 | 21.8025028 |
| TCGA-A7-A6VV | TCGA-BRCA | 93.6195956 | 47.494862 |
| TCGA-A7-A6VW | TCGA-BRCA | 145.97987 | 97.3135565 |
| TCGA-A7-A6VX | TCGA-BRCA | 83.3553847 | 90.5334136 |
| TCGA-A7-A6VY | TCGA-BRCA | 128.753116 | 58.7188754 |
| TCGA-A8-A06N | TCGA-BRCA | 65.0040756 | 33.0811865 |
| TCGA-A8-A06O | TCGA-BRCA | 43.4766503 | 17.2717132 |
| TCGA-A8-A06P | TCGA-BRCA | 47.2206807 | 19.1536617 |

| | | | |
|---|---|---|---|
| TCGA-A8-A06Q | TCGA-BRCA | 61.5425932 | 45.3016696 |
| TCGA-A8-A06R | TCGA-BRCA | 90.5503362 | 20.323536 |
| TCGA-A8-A06T | TCGA-BRCA | 53.6044121 | 23.7169421 |
| TCGA-A8-A06U | TCGA-BRCA | 45.6111097 | 35.4151611 |
| TCGA-A8-A06X | TCGA-BRCA | 60.6606634 | 38.4718596 |
| TCGA-A8-A06Y | TCGA-BRCA | 32.6469417 | 47.1331372 |
| TCGA-A8-A06Z | TCGA-BRCA | 60.1790053 | 26.1194711 |
| TCGA-A8-A075 | TCGA-BRCA | 34.3159637 | 25.4375172 |
| TCGA-A8-A076 | TCGA-BRCA | 69.3736505 | 54.357747 |
| TCGA-A8-A079 | TCGA-BRCA | 58.6864355 | 20.1455702 |
| TCGA-A8-A07B | TCGA-BRCA | 53.2360345 | 35.5503665 |
| TCGA-A8-A07C | TCGA-BRCA | 98.7280178 | 90.6174836 |
| TCGA-A8-A07E | TCGA-BRCA | 61.5028292 | 33.8346382 |
| TCGA-A8-A07F | TCGA-BRCA | 44.5883086 | 54.7998884 |
| TCGA-A8-A07G | TCGA-BRCA | 40.2616941 | 26.2651021 |
| TCGA-A8-A07I | TCGA-BRCA | 51.1835906 | 45.3786687 |
| TCGA-A8-A07J | TCGA-BRCA | 46.2284991 | 17.5219483 |
| TCGA-A8-A07L | TCGA-BRCA | 89.0501818 | 54.4941633 |
| TCGA-A8-A07O | TCGA-BRCA | 89.2545241 | 42.7218103 |
| TCGA-A8-A07P | TCGA-BRCA | 45.2211303 | 26.3237237 |
| TCGA-A8-A07R | TCGA-BRCA | 122.215939 | 63.7186431 |
| TCGA-A8-A07S | TCGA-BRCA | 52.9514784 | 28.0597512 |
| TCGA-A8-A07U | TCGA-BRCA | 45.7569103 | 62.6521542 |
| TCGA-A8-A07W | TCGA-BRCA | 55.4562799 | 39.1964278 |
| TCGA-A8-A07Z | TCGA-BRCA | 63.8261957 | 62.3223522 |
| TCGA-A8-A081 | TCGA-BRCA | 52.0803078 | 55.4400665 |
| TCGA-A8-A082 | TCGA-BRCA | 62.28599 | 24.5832259 |
| TCGA-A8-A083 | TCGA-BRCA | 62.4969821 | 26.8347043 |
| TCGA-A8-A084 | TCGA-BRCA | 54.566397 | 50.5796186 |
| TCGA-A8-A085 | TCGA-BRCA | 49.9648622 | 11.8086162 |
| TCGA-A8-A086 | TCGA-BRCA | 48.0997216 | 26.0915713 |

| | | | |
|---|---|---|---|
| TCGA-A8-A08A | TCGA-BRCA | 47.7491146 | 27.581491 |
| TCGA-A8-A08B | TCGA-BRCA | 87.2191406 | 64.1781008 |
| TCGA-A8-A08C | TCGA-BRCA | 48.2807407 | 28.4964825 |
| TCGA-A8-A08F | TCGA-BRCA | 54.9831846 | 36.4695723 |
| TCGA-A8-A08G | TCGA-BRCA | 49.8400771 | 45.0488563 |
| TCGA-A8-A08H | TCGA-BRCA | 48.9741368 | 29.8370268 |
| TCGA-A8-A08I | TCGA-BRCA | 90.0309602 | 26.644976 |
| TCGA-A8-A08J | TCGA-BRCA | 43.949018 | 54.4921997 |
| TCGA-A8-A08L | TCGA-BRCA | 70.3712895 | 60.7065873 |
| TCGA-A8-A08O | TCGA-BRCA | 74.9191244 | 48.7394006 |
| TCGA-A8-A08P | TCGA-BRCA | 66.6704605 | 58.0964736 |
| TCGA-A8-A08R | TCGA-BRCA | 50.5870249 | 34.3398371 |
| TCGA-A8-A08S | TCGA-BRCA | 40.0891161 | 36.2522753 |
| TCGA-A8-A08T | TCGA-BRCA | 47.1235906 | 30.3858223 |
| TCGA-A8-A08X | TCGA-BRCA | 37.5801002 | 17.4313284 |
| TCGA-A8-A08Z | TCGA-BRCA | 62.5506832 | 17.4289103 |
| TCGA-A8-A090 | TCGA-BRCA | 47.6781627 | 39.2891113 |
| TCGA-A8-A091 | TCGA-BRCA | 36.4589894 | 16.0238918 |
| TCGA-A8-A092 | TCGA-BRCA | 49.723802 | 47.5365172 |
| TCGA-A8-A093 | TCGA-BRCA | 43.5883392 | 25.3963901 |
| TCGA-A8-A094 | TCGA-BRCA | 53.8330846 | 69.819673 |
| TCGA-A8-A095 | TCGA-BRCA | 65.6684761 | 50.8795007 |
| TCGA-A8-A096 | TCGA-BRCA | 53.9061233 | 27.5532097 |
| TCGA-A8-A097 | TCGA-BRCA | 43.9162758 | 22.1878644 |
| TCGA-A8-A099 | TCGA-BRCA | 36.1572097 | 54.8616841 |
| TCGA-A8-A09A | TCGA-BRCA | 45.3674527 | 39.3448023 |
| TCGA-A8-A09B | TCGA-BRCA | 43.1010485 | 28.1761081 |
| TCGA-A8-A09D | TCGA-BRCA | 46.2457839 | 27.4609198 |
| TCGA-A8-A09E | TCGA-BRCA | 46.5594516 | 36.592973 |
| TCGA-A8-A09G | TCGA-BRCA | 48.1463626 | 81.6312724 |
| TCGA-A8-A09I | TCGA-BRCA | 43.9138886 | 48.6662159 |

| | | | |
|---|---|---|---|
| TCGA-A8-A09K | TCGA-BRCA | 51.7775568 | 28.6319859 |
| TCGA-A8-A09M | TCGA-BRCA | 45.5105134 | 35.619544 |
| TCGA-A8-A09N | TCGA-BRCA | 64.2512913 | 65.6610057 |
| TCGA-A8-A09Q | TCGA-BRCA | 33.7701251 | 14.5974547 |
| TCGA-A8-A09R | TCGA-BRCA | 34.8390484 | 27.3474741 |
| TCGA-A8-A09T | TCGA-BRCA | 95.0950973 | 21.9902418 |
| TCGA-A8-A09V | TCGA-BRCA | 61.0390416 | 29.73854 |
| TCGA-A8-A09W | TCGA-BRCA | 53.5848425 | 34.010862 |
| TCGA-A8-A09X | TCGA-BRCA | 52.4692862 | 65.9284891 |
| TCGA-A8-A09Z | TCGA-BRCA | 48.8062913 | 17.6586408 |
| TCGA-A8-A0A1 | TCGA-BRCA | 45.3670406 | 25.5158703 |
| TCGA-A8-A0A2 | TCGA-BRCA | 42.054556 | 23.7803712 |
| TCGA-A8-A0A4 | TCGA-BRCA | 44.6126336 | 22.7956195 |
| TCGA-A8-A0A6 | TCGA-BRCA | 48.1319665 | 55.2801639 |
| TCGA-A8-A0A7 | TCGA-BRCA | 29.8617375 | 14.4903655 |
| TCGA-A8-A0A9 | TCGA-BRCA | 54.4856512 | 30.399224 |
| TCGA-A8-A0AB | TCGA-BRCA | 55.8800142 | 25.6532825 |
| TCGA-A8-A0AD | TCGA-BRCA | 53.8287546 | 31.0467341 |
| TCGA-AA-3489 | TCGA-COAD | 68.5434011 | 19.0548701 |
| TCGA-AA-3496 | TCGA-COAD | 42.4594528 | 14.1380539 |
| TCGA-AA-3511 | TCGA-COAD | 44.8525371 | 12.1616167 |
| TCGA-AA-3514 | TCGA-COAD | 49.5770534 | 18.827742 |
| TCGA-AA-3516 | TCGA-COAD | 28.7628132 | 7.78514057 |
| TCGA-AA-3520 | TCGA-COAD | 41.1945239 | 10.6323163 |
| TCGA-AA-3522 | TCGA-COAD | 41.4446224 | 9.51758015 |
| TCGA-AA-3526 | TCGA-COAD | 120.677757 | 49.9626468 |
| TCGA-AA-3531 | TCGA-COAD | 29.2250028 | 8.05312621 |
| TCGA-AA-3655 | TCGA-COAD | 77.8289224 | 21.7697206 |
| TCGA-AA-3660 | TCGA-COAD | 38.2329667 | 13.563413 |
| TCGA-AA-3662 | TCGA-COAD | 39.2185318 | 12.3674629 |
| TCGA-AA-3663 | TCGA-COAD | 33.0253581 | 9.17649668 |

| | | | |
|---|---|---|---|
| TCGA-AA-3675 | TCGA-COAD | 52.2720662 | 19.9118331 |
| TCGA-AA-3685 | TCGA-COAD | 48.7350049 | 44.9808712 |
| TCGA-AA-3697 | TCGA-COAD | 55.4052048 | 24.7580359 |
| TCGA-AA-3712 | TCGA-COAD | 34.083957 | 8.91876314 |
| TCGA-AA-3713 | TCGA-COAD | 54.3573797 | 30.4633683 |
| TCGA-AA-A02K | TCGA-COAD | 66.8902637 | 42.4037954 |
| TCGA-AA-A02Y | TCGA-COAD | 45.0160319 | 24.6768918 |
| TCGA-AC-A23C | TCGA-BRCA | 51.9585162 | 49.1799979 |
| TCGA-AC-A23E | TCGA-BRCA | 46.1552757 | 26.4659994 |
| TCGA-AC-A23G | TCGA-BRCA | 34.6716838 | 22.2361588 |
| TCGA-AC-A23H | TCGA-BRCA | 53.3408473 | 26.4497773 |
| TCGA-AC-A2B8 | TCGA-BRCA | 47.1263058 | 25.520685 |
| TCGA-AC-A2BK | TCGA-BRCA | 91.5419361 | 58.5935539 |
| TCGA-AC-A2BM | TCGA-BRCA | 45.7984497 | 51.8262678 |
| TCGA-AC-A2FB | TCGA-BRCA | 42.349878 | 18.0091529 |
| TCGA-AC-A2FE | TCGA-BRCA | 34.5869184 | 21.8563587 |
| TCGA-AC-A2FF | TCGA-BRCA | 45.7875908 | 19.5808048 |
| TCGA-AC-A2FG | TCGA-BRCA | 53.9542259 | 24.8252107 |
| TCGA-AC-A2FK | TCGA-BRCA | 39.581958 | 13.8240018 |
| TCGA-AC-A2FM | TCGA-BRCA | 36.6920963 | 30.8954422 |
| TCGA-AC-A2FO | TCGA-BRCA | 48.2847513 | 27.29982 |
| TCGA-AC-A2QH | TCGA-BRCA | 44.5797639 | 18.2289297 |
| TCGA-AC-A2QI | TCGA-BRCA | 51.3523264 | 14.9622931 |
| TCGA-AC-A2QJ | TCGA-BRCA | 37.5834444 | 20.2730662 |
| TCGA-AC-A3BB | TCGA-BRCA | 43.4318932 | 21.0704109 |
| TCGA-AC-A3EH | TCGA-BRCA | 43.1052157 | 31.7224069 |
| TCGA-AC-A3HN | TCGA-BRCA | 46.9361368 | 22.2673291 |
| TCGA-AC-A3OD | TCGA-BRCA | 21.4398334 | 3.2035707 |
| TCGA-AC-A3QP | TCGA-BRCA | 58.8562238 | 10.6780101 |
| TCGA-AC-A3QQ | TCGA-BRCA | 12.0400713 | 6.52832546 |
| TCGA-AC-A3TM | TCGA-BRCA | 45.1447279 | 20.7147581 |

| | | | |
|---|---|---|---|
| TCGA-AC-A3TN | TCGA-BRCA | 50.0915364 | 17.6495913 |
| TCGA-AC-A3W5 | TCGA-BRCA | 44.2336517 | 33.4868022 |
| TCGA-AC-A3W6 | TCGA-BRCA | 43.3438373 | 32.63977 |
| TCGA-AC-A3W7 | TCGA-BRCA | 46.5327466 | 17.5273962 |
| TCGA-AC-A3YI | TCGA-BRCA | 56.9051783 | 19.1475025 |
| TCGA-AC-A3YJ | TCGA-BRCA | 59.4026723 | 31.5534233 |
| TCGA-AC-A4ZE | TCGA-BRCA | 52.2566052 | 28.1659574 |
| TCGA-AC-A5EH | TCGA-BRCA | 61.402069 | 25.310412 |
| TCGA-AC-A5XS | TCGA-BRCA | 83.6314713 | 22.4422726 |
| TCGA-AC-A5XU | TCGA-BRCA | 32.6293359 | 26.2927032 |
| TCGA-AC-A62V | TCGA-BRCA | 52.4731907 | 34.798782 |
| TCGA-AC-A62X | TCGA-BRCA | 173.455799 | 65.0076603 |
| TCGA-AC-A62Y | TCGA-BRCA | 102.800276 | 57.5850051 |
| TCGA-AC-A6IV | TCGA-BRCA | 47.3324843 | 15.5439474 |
| TCGA-AC-A6IW | TCGA-BRCA | 115.156413 | 51.4596462 |
| TCGA-AC-A6IX | TCGA-BRCA | 50.8092598 | 45.995844 |
| TCGA-AC-A6NO | TCGA-BRCA | 60.8790536 | 34.4843821 |
| TCGA-AC-A7VB | TCGA-BRCA | 84.641115 | 64.0567366 |
| TCGA-AC-A7VC | TCGA-BRCA | 78.3322131 | 24.0866509 |
| TCGA-AC-A8OP | TCGA-BRCA | 41.1750662 | 18.7008274 |
| TCGA-AC-A8OQ | TCGA-BRCA | 58.3130624 | 17.9620059 |
| TCGA-AC-A8OR | TCGA-BRCA | 59.2533493 | 19.9008624 |
| TCGA-AC-A8OS | TCGA-BRCA | 34.359389 | 18.4903164 |
| TCGA-AD-5900 | TCGA-COAD | 66.8862472 | 54.7310016 |
| TCGA-AD-6548 | TCGA-COAD | 99.2194887 | 32.2612661 |
| TCGA-AD-6888 | TCGA-COAD | 62.5137782 | 19.0816946 |
| TCGA-AD-6889 | TCGA-COAD | 59.386076 | 16.4540569 |
| TCGA-AD-6890 | TCGA-COAD | 109.939701 | 29.4546884 |
| TCGA-AD-6895 | TCGA-COAD | 55.305751 | 27.4619756 |
| TCGA-AD-6899 | TCGA-COAD | 54.3852346 | 16.2788561 |
| TCGA-AD-6901 | TCGA-COAD | 52.9611008 | 20.5854546 |

| | | | |
|---|---|---|---|
| TCGA-AD-6963 | TCGA-COAD | 80.1846839 | 46.4655485 |
| TCGA-AD-6964 | TCGA-COAD | 56.8681172 | 20.4181974 |
| TCGA-AD-6965 | TCGA-COAD | 71.8060067 | 35.4654597 |
| TCGA-AD-A5EJ | TCGA-COAD | 57.9975658 | 31.7690375 |
| TCGA-AD-A5EK | TCGA-COAD | 68.0635534 | 17.8761415 |
| TCGA-AJ-A23N | TCGA-UCEC | 89.4281619 | 30.0655915 |
| TCGA-AJ-A2QM | TCGA-UCEC | 75.5732757 | 20.78897 |
| TCGA-AJ-A3BD | TCGA-UCEC | 130.118439 | 18.8773937 |
| TCGA-AJ-A3BF | TCGA-UCEC | 105.300097 | 19.4845882 |
| TCGA-AJ-A3BG | TCGA-UCEC | 14.7212668 | 9.99853458 |
| TCGA-AJ-A3BH | TCGA-UCEC | 49.6379508 | 20.9809996 |
| TCGA-AJ-A3BI | TCGA-UCEC | 8.8981976 | 6.04007926 |
| TCGA-AJ-A3BK | TCGA-UCEC | 30.6142701 | 7.15300191 |
| TCGA-AJ-A3EJ | TCGA-UCEC | 61.9763144 | 7.75472712 |
| TCGA-AJ-A3EK | TCGA-UCEC | 48.6711814 | 9.25381049 |
| TCGA-AJ-A3EL | TCGA-UCEC | 111.864523 | 52.638769 |
| TCGA-AJ-A3EM | TCGA-UCEC | 38.9645783 | 10.1165778 |
| TCGA-AJ-A3I9 | TCGA-UCEC | 87.3596595 | 33.8732372 |
| TCGA-AJ-A3IA | TCGA-UCEC | 89.8949561 | 56.737425 |
| TCGA-AJ-A3NC | TCGA-UCEC | 71.0053221 | 27.4682522 |
| TCGA-AJ-A3NE | TCGA-UCEC | 37.6548955 | 37.1964104 |
| TCGA-AJ-A3NF | TCGA-UCEC | 106.110244 | 6.32382321 |
| TCGA-AJ-A3NG | TCGA-UCEC | 70.9715539 | 28.338991 |
| TCGA-AJ-A3NH | TCGA-UCEC | 154.376841 | 26.2375706 |
| TCGA-AJ-A3OJ | TCGA-UCEC | 54.896045 | 23.9706619 |
| TCGA-AJ-A3OK | TCGA-UCEC | 79.07453 | 3.27962367 |
| TCGA-AJ-A3OL | TCGA-UCEC | 108.202992 | 42.9689292 |
| TCGA-AJ-A3QS | TCGA-UCEC | 115.847378 | 6.91578373 |
| TCGA-AJ-A3TW | TCGA-UCEC | 76.721838 | 39.7093417 |
| TCGA-AJ-A5DV | TCGA-UCEC | 62.4567478 | 22.1710922 |
| TCGA-AJ-A5DW | TCGA-UCEC | 72.9051951 | 109.192578 |

| | | | |
|---|---|---|---|
| TCGA-AJ-A8CT | TCGA-UCEC | 57.3254993 | 37.4090081 |
| TCGA-AJ-A8CV | TCGA-UCEC | 67.2841216 | 26.7281512 |
| TCGA-AJ-A8CW | TCGA-UCEC | 92.6478857 | 38.6841739 |
| TCGA-AK-3425 | TCGA-KIRC | 34.8637253 | 17.8821722 |
| TCGA-AK-3426 | TCGA-KIRC | 30.6058985 | 13.9446217 |
| TCGA-AK-3427 | TCGA-KIRC | 17.4033645 | 16.7164765 |
| TCGA-AK-3428 | TCGA-KIRC | 47.5910362 | 17.0173396 |
| TCGA-AK-3429 | TCGA-KIRC | 42.3143364 | 18.4250038 |
| TCGA-AK-3431 | TCGA-KIRC | 46.3796218 | 20.159612 |
| TCGA-AK-3433 | TCGA-KIRC | 19.6040426 | 21.5036501 |
| TCGA-AK-3434 | TCGA-KIRC | 47.550345 | 16.7764181 |
| TCGA-AK-3436 | TCGA-KIRC | 20.1172048 | 17.7294312 |
| TCGA-AK-3440 | TCGA-KIRC | 32.8055779 | 37.8401032 |
| TCGA-AK-3443 | TCGA-KIRC | 32.2504894 | 13.4940325 |
| TCGA-AK-3445 | TCGA-KIRC | 35.2158463 | 17.7856319 |
| TCGA-AK-3447 | TCGA-KIRC | 10.0987133 | 21.583286 |
| TCGA-AK-3450 | TCGA-KIRC | 49.7809344 | 29.4731736 |
| TCGA-AK-3451 | TCGA-KIRC | 32.0746795 | 24.2610432 |
| TCGA-AK-3453 | TCGA-KIRC | 40.6876018 | 16.0768196 |
| TCGA-AK-3454 | TCGA-KIRC | 28.7373162 | 10.0618874 |
| TCGA-AK-3455 | TCGA-KIRC | 52.2742266 | 27.3201545 |
| TCGA-AK-3456 | TCGA-KIRC | 23.4038707 | 8.81511314 |
| TCGA-AK-3458 | TCGA-KIRC | 34.5804277 | 14.7802843 |
| TCGA-AK-3460 | TCGA-KIRC | 39.9810868 | 14.86447 |
| TCGA-AK-3461 | TCGA-KIRC | 55.5881566 | 22.5759075 |
| TCGA-AK-3465 | TCGA-KIRC | 25.5714725 | 14.7510485 |
| TCGA-AL-3466 | TCGA-KIRP | 97.938804 | 1.02594448 |
| TCGA-AL-3468 | TCGA-KIRP | 58.6222085 | 10.4043987 |
| TCGA-AL-3471 | TCGA-KIRP | 60.0966287 | 25.5081266 |
| TCGA-AL-3472 | TCGA-KIRP | 47.3157513 | 17.19864 |
| TCGA-AL-3473 | TCGA-KIRP | 40.1132138 | 5.25227173 |

| | | | |
|---|---|---|---|
| TCGA-AL-7173 | TCGA-KIRP | 31.1457709 | 8.52413347 |
| TCGA-AL-A5DJ | TCGA-KIRP | 58.6965645 | 26.8689738 |
| TCGA-AM-5820 | TCGA-COAD | 89.1033659 | 25.8564393 |
| TCGA-AM-5821 | TCGA-COAD | 99.7959736 | 56.0121407 |
| TCGA-AN-A03X | TCGA-BRCA | 31.1739452 | 45.0130821 |
| TCGA-AN-A03Y | TCGA-BRCA | 44.5610845 | 47.4956307 |
| TCGA-AN-A041 | TCGA-BRCA | 41.1660618 | 30.6930033 |
| TCGA-AN-A046 | TCGA-BRCA | 43.1767207 | 13.5394543 |
| TCGA-AN-A049 | TCGA-BRCA | 54.2536249 | 23.8211671 |
| TCGA-AN-A04A | TCGA-BRCA | 40.3378925 | 22.2972162 |
| TCGA-AN-A04C | TCGA-BRCA | 61.2239204 | 36.7616914 |
| TCGA-AN-A04D | TCGA-BRCA | 291.703375 | 60.7131647 |
| TCGA-AN-A0AJ | TCGA-BRCA | 50.5573231 | 51.1793137 |
| TCGA-AN-A0AK | TCGA-BRCA | 45.2305993 | 45.6294725 |
| TCGA-AN-A0AL | TCGA-BRCA | 76.9441397 | 26.9122196 |
| TCGA-AN-A0AM | TCGA-BRCA | 166.791306 | 39.0264263 |
| TCGA-AN-A0AR | TCGA-BRCA | 43.8938818 | 63.2894761 |
| TCGA-AN-A0AS | TCGA-BRCA | 54.8185899 | 31.4091449 |
| TCGA-AN-A0AT | TCGA-BRCA | 92.6953663 | 45.1315855 |
| TCGA-AN-A0FD | TCGA-BRCA | 36.1692598 | 17.8965871 |
| TCGA-AN-A0FF | TCGA-BRCA | 66.7392352 | 28.9801968 |
| TCGA-AN-A0FJ | TCGA-BRCA | 83.770059 | 146.475978 |
| TCGA-AN-A0FK | TCGA-BRCA | 41.7008859 | 67.3315287 |
| TCGA-AN-A0FL | TCGA-BRCA | 80.8372621 | 15.0080721 |
| TCGA-AN-A0FN | TCGA-BRCA | 55.3755709 | 30.952585 |
| TCGA-AN-A0FS | TCGA-BRCA | 67.8406646 | 26.5243623 |
| TCGA-AN-A0FT | TCGA-BRCA | 58.5137899 | 53.4484316 |
| TCGA-AN-A0FV | TCGA-BRCA | 48.8449972 | 28.9972227 |
| TCGA-AN-A0FW | TCGA-BRCA | 47.4023139 | 44.3142464 |
| TCGA-AN-A0FX | TCGA-BRCA | 66.3796659 | 55.8706099 |
| TCGA-AN-A0FY | TCGA-BRCA | 93.3937937 | 60.8315135 |

| | | | |
|---|---|---|---|
| TCGA-AN-A0FZ | TCGA-BRCA | 82.0507343 | 32.0285202 |
| TCGA-AN-A0G0 | TCGA-BRCA | 46.1462163 | 36.277417 |
| TCGA-AN-A0XL | TCGA-BRCA | 55.7445989 | 38.6965153 |
| TCGA-AN-A0XN | TCGA-BRCA | 56.2643481 | 20.8458509 |
| TCGA-AN-A0XO | TCGA-BRCA | 48.7646777 | 28.842327 |
| TCGA-AN-A0XP | TCGA-BRCA | 45.0482162 | 50.061986 |
| TCGA-AN-A0XR | TCGA-BRCA | 81.9468423 | 44.6035291 |
| TCGA-AN-A0XS | TCGA-BRCA | 38.8756684 | 23.0106352 |
| TCGA-AN-A0XT | TCGA-BRCA | 75.1221773 | 24.027115 |
| TCGA-AN-A0XU | TCGA-BRCA | 124.885438 | 39.9147542 |
| TCGA-AN-A0XV | TCGA-BRCA | 45.0455073 | 35.5535381 |
| TCGA-AN-A0XW | TCGA-BRCA | 57.5740166 | 24.4342014 |
| TCGA-AO-A03L | TCGA-BRCA | 57.9001733 | 47.9105607 |
| TCGA-AO-A03M | TCGA-BRCA | 54.7632668 | 36.2221577 |
| TCGA-AO-A03N | TCGA-BRCA | 38.6135174 | 70.9828961 |
| TCGA-AO-A03O | TCGA-BRCA | 81.9384353 | 32.3237191 |
| TCGA-AO-A03P | TCGA-BRCA | 46.6182132 | 34.0705247 |
| TCGA-AO-A03R | TCGA-BRCA | 71.6485124 | 12.0993784 |
| TCGA-AO-A03T | TCGA-BRCA | 55.3980335 | 19.0131953 |
| TCGA-AO-A03U | TCGA-BRCA | 34.0489768 | 18.4805107 |
| TCGA-AO-A03V | TCGA-BRCA | 82.6810514 | 29.4148245 |
| TCGA-AO-A0J2 | TCGA-BRCA | 108.999085 | 66.5138329 |
| TCGA-AO-A0J3 | TCGA-BRCA | 51.2499902 | 65.9035415 |
| TCGA-AO-A0J4 | TCGA-BRCA | 67.5614121 | 31.1332253 |
| TCGA-AO-A0J5 | TCGA-BRCA | 34.1571757 | 24.8290216 |
| TCGA-AO-A0J6 | TCGA-BRCA | 122.278029 | 41.764945 |
| TCGA-AO-A0J7 | TCGA-BRCA | 48.8899661 | 32.2576675 |
| TCGA-AO-A0J8 | TCGA-BRCA | 57.5384964 | 27.9780058 |
| TCGA-AO-A0J9 | TCGA-BRCA | 38.3134127 | 40.2519097 |
| TCGA-AO-A0JA | TCGA-BRCA | 36.2826213 | 53.6873468 |
| TCGA-AO-A0JB | TCGA-BRCA | 27.163681 | 11.6559827 |

| | | | |
|---|---|---|---|
| TCGA-AO-A0JC | TCGA-BRCA | 51.8712844 | 16.9354342 |
| TCGA-AO-A0JD | TCGA-BRCA | 51.6154726 | 83.7151465 |
| TCGA-AO-A0JE | TCGA-BRCA | 68.5562448 | 34.8731122 |
| TCGA-AO-A0JF | TCGA-BRCA | 45.4385407 | 28.0543661 |
| TCGA-AO-A0JG | TCGA-BRCA | 48.3499857 | 22.9800972 |
| TCGA-AO-A0JI | TCGA-BRCA | 33.2352913 | 44.5873976 |
| TCGA-AO-A0JJ | TCGA-BRCA | 53.8475374 | 38.1915918 |
| TCGA-AO-A0JL | TCGA-BRCA | 54.4504716 | 33.2869623 |
| TCGA-AO-A0JM | TCGA-BRCA | 69.3078903 | 55.7926005 |
| TCGA-AO-A124 | TCGA-BRCA | 147.77678 | 101.058127 |
| TCGA-AO-A125 | TCGA-BRCA | 61.3362509 | 32.6605593 |
| TCGA-AO-A126 | TCGA-BRCA | 67.0807785 | 16.8228556 |
| TCGA-AO-A128 | TCGA-BRCA | 84.6797068 | 31.6626728 |
| TCGA-AO-A129 | TCGA-BRCA | 136.189405 | 79.2326059 |
| TCGA-AO-A12A | TCGA-BRCA | 53.5056348 | 42.4786931 |
| TCGA-AO-A12B | TCGA-BRCA | 42.5944564 | 29.3526878 |
| TCGA-AO-A12C | TCGA-BRCA | 38.8164704 | 24.9759296 |
| TCGA-AO-A12D | TCGA-BRCA | 24.5113361 | 12.5004053 |
| TCGA-AO-A12E | TCGA-BRCA | 47.1147614 | 31.604284 |
| TCGA-AO-A12F | TCGA-BRCA | 128.969117 | 117.709259 |
| TCGA-AO-A12G | TCGA-BRCA | 36.3310095 | 15.189312 |
| TCGA-AO-A12H | TCGA-BRCA | 46.7730615 | 26.9953409 |
| TCGA-AO-A1KO | TCGA-BRCA | 30.1051277 | 16.0448614 |
| TCGA-AO-A1KP | TCGA-BRCA | 33.7001754 | 51.0758973 |
| TCGA-AO-A1KQ | TCGA-BRCA | 64.5905181 | 82.1611211 |
| TCGA-AO-A1KR | TCGA-BRCA | 107.95439 | 35.7485731 |
| TCGA-AO-A1KS | TCGA-BRCA | 36.3503591 | 60.2856313 |
| TCGA-AO-A1KT | TCGA-BRCA | 64.8519934 | 50.7186207 |
| TCGA-AP-A3K1 | TCGA-UCEC | 100.391364 | 24.7819211 |
| TCGA-AP-A5FX | TCGA-UCEC | 79.3903048 | 18.9428354 |
| TCGA-AQ-A04H | TCGA-BRCA | 61.353244 | 33.2040252 |

| | | | |
|---|---|---|---|
| TCGA-AQ-A04J | TCGA-BRCA | 100.895956 | 58.6456914 |
| TCGA-AQ-A04L | TCGA-BRCA | 76.6529848 | 27.9585773 |
| TCGA-AQ-A0Y5 | TCGA-BRCA | 35.080729 | 36.5759871 |
| TCGA-AQ-A1H2 | TCGA-BRCA | 75.3991373 | 23.4056951 |
| TCGA-AQ-A1H3 | TCGA-BRCA | 52.4899879 | 26.4164637 |
| TCGA-AQ-A54N | TCGA-BRCA | 71.6410578 | 26.2160846 |
| TCGA-AQ-A54O | TCGA-BRCA | 48.4841092 | 25.6049484 |
| TCGA-AQ-A7U7 | TCGA-BRCA | 42.5221679 | 26.9625866 |
| TCGA-AR-A0TP | TCGA-BRCA | 56.1860535 | 36.4802795 |
| TCGA-AR-A0TQ | TCGA-BRCA | 40.5895196 | 41.0336272 |
| TCGA-AR-A0TR | TCGA-BRCA | 65.2216779 | 16.870823 |
| TCGA-AR-A0TS | TCGA-BRCA | 87.3527323 | 34.6527386 |
| TCGA-AR-A0TT | TCGA-BRCA | 41.6103125 | 28.2103261 |
| TCGA-AR-A0TU | TCGA-BRCA | 256.053441 | 97.1958844 |
| TCGA-AR-A0TV | TCGA-BRCA | 54.7485501 | 26.9914431 |
| TCGA-AR-A0TW | TCGA-BRCA | 29.5987797 | 17.8733791 |
| TCGA-AR-A0TX | TCGA-BRCA | 50.8565495 | 36.2672218 |
| TCGA-AR-A0TY | TCGA-BRCA | 74.1878839 | 75.5095891 |
| TCGA-AR-A0TZ | TCGA-BRCA | 77.0404623 | 27.9833457 |
| TCGA-AR-A0U0 | TCGA-BRCA | 63.924459 | 29.6298158 |
| TCGA-AR-A0U2 | TCGA-BRCA | 38.4012639 | 36.261614 |
| TCGA-AR-A0U3 | TCGA-BRCA | 54.8336981 | 48.087624 |
| TCGA-AR-A0U4 | TCGA-BRCA | 94.2605681 | 37.2065118 |
| TCGA-AR-A1AH | TCGA-BRCA | 339.411483 | 110.00876 |
| TCGA-AR-A1AI | TCGA-BRCA | 115.897649 | 34.987898 |
| TCGA-AR-A1AJ | TCGA-BRCA | 82.5674795 | 41.4162574 |
| TCGA-AR-A1AK | TCGA-BRCA | 55.3782295 | 29.7929747 |
| TCGA-AR-A1AL | TCGA-BRCA | 41.6189068 | 24.047936 |
| TCGA-AR-A1AM | TCGA-BRCA | 54.2411921 | 19.2236043 |
| TCGA-AR-A1AN | TCGA-BRCA | 52.0129416 | 26.9973883 |
| TCGA-AR-A1AO | TCGA-BRCA | 74.0865107 | 38.305034 |

| | | | |
|---|---|---|---|
| TCGA-AR-A1AP | TCGA-BRCA | 45.6505841 | 33.2537382 |
| TCGA-AR-A1AQ | TCGA-BRCA | 86.9777667 | 32.9528027 |
| TCGA-AR-A1AR | TCGA-BRCA | 112.229646 | 35.9124658 |
| TCGA-AR-A1AS | TCGA-BRCA | 67.6470437 | 35.4478421 |
| TCGA-AR-A1AT | TCGA-BRCA | 47.0381276 | 21.7655367 |
| TCGA-AR-A1AU | TCGA-BRCA | 36.6778103 | 18.7866516 |
| TCGA-AR-A1AV | TCGA-BRCA | 47.7669405 | 45.0341204 |
| TCGA-AR-A1AW | TCGA-BRCA | 36.7995891 | 13.525998 |
| TCGA-AR-A1AX | TCGA-BRCA | 44.7334395 | 22.7742222 |
| TCGA-AR-A1AY | TCGA-BRCA | 161.596029 | 87.0057232 |
| TCGA-AR-A24H | TCGA-BRCA | 118.449504 | 75.7294325 |
| TCGA-AR-A24K | TCGA-BRCA | 56.799321 | 21.8305327 |
| TCGA-AR-A24L | TCGA-BRCA | 66.6520718 | 43.2556466 |
| TCGA-AR-A24M | TCGA-BRCA | 62.1393837 | 17.2875484 |
| TCGA-AR-A24N | TCGA-BRCA | 45.7638864 | 28.540712 |
| TCGA-AR-A24O | TCGA-BRCA | 50.0336621 | 26.8387587 |
| TCGA-AR-A24P | TCGA-BRCA | 50.0151598 | 25.6792065 |
| TCGA-AR-A24Q | TCGA-BRCA | 80.8629912 | 58.8223674 |
| TCGA-AR-A24R | TCGA-BRCA | 52.1415379 | 33.9715503 |
| TCGA-AR-A24S | TCGA-BRCA | 91.9191671 | 24.5430487 |
| TCGA-AR-A24T | TCGA-BRCA | 42.0688351 | 28.3870053 |
| TCGA-AR-A24U | TCGA-BRCA | 31.0675024 | 21.7167105 |
| TCGA-AR-A24V | TCGA-BRCA | 45.1557561 | 29.87171 |
| TCGA-AR-A24W | TCGA-BRCA | 57.8558115 | 34.5877545 |
| TCGA-AR-A24X | TCGA-BRCA | 56.6438248 | 31.5167135 |
| TCGA-AR-A24Z | TCGA-BRCA | 138.164505 | 45.2527418 |
| TCGA-AR-A250 | TCGA-BRCA | 53.9607894 | 41.8985771 |
| TCGA-AR-A251 | TCGA-BRCA | 122.28458 | 29.0378743 |
| TCGA-AR-A252 | TCGA-BRCA | 34.6165745 | 24.4943823 |
| TCGA-AR-A255 | TCGA-BRCA | 54.6440247 | 35.1577548 |
| TCGA-AR-A256 | TCGA-BRCA | 125.805216 | 74.0363031 |

| | | | |
|---|---|---|---|
| TCGA-AR-A2LE | TCGA-BRCA | 20.5044354 | 9.31770378 |
| TCGA-AR-A2LH | TCGA-BRCA | 58.9218664 | 23.7253539 |
| TCGA-AR-A2LJ | TCGA-BRCA | 39.4131096 | 13.4710153 |
| TCGA-AR-A2LK | TCGA-BRCA | 51.3012863 | 41.3301632 |
| TCGA-AR-A2LL | TCGA-BRCA | 46.8627563 | 25.8590505 |
| TCGA-AR-A2LM | TCGA-BRCA | 50.5133272 | 14.7152121 |
| TCGA-AR-A2LN | TCGA-BRCA | 48.4588286 | 14.0082465 |
| TCGA-AR-A2LO | TCGA-BRCA | 39.4848327 | 26.7157662 |
| TCGA-AR-A2LQ | TCGA-BRCA | 35.483792 | 19.3406732 |
| TCGA-AR-A2LR | TCGA-BRCA | 95.8296689 | 46.3868972 |
| TCGA-AR-A5QM | TCGA-BRCA | 48.9479983 | 41.8812572 |
| TCGA-AR-A5QN | TCGA-BRCA | 38.6387297 | 47.9191626 |
| TCGA-AR-A5QP | TCGA-BRCA | 61.367602 | 29.1215249 |
| TCGA-AR-A5QQ | TCGA-BRCA | 93.3807709 | 36.2189299 |
| TCGA-AS-3777 | TCGA-KIRC | 21.6876792 | 18.4252453 |
| TCGA-AS-3778 | TCGA-KIRC | 41.1011976 | 10.7185319 |
| TCGA-AT-A5NU | TCGA-KIRP | 31.6989063 | 15.0227252 |
| TCGA-AU-3779 | TCGA-COAD | 37.8143844 | 26.2356379 |
| TCGA-AU-6004 | TCGA-COAD | 49.6655355 | 21.0707658 |
| TCGA-AX-A05W | TCGA-UCEC | 43.164277 | 13.1801384 |
| TCGA-AX-A1C7 | TCGA-UCEC | 21.2364817 | 2.23034403 |
| TCGA-AX-A2H4 | TCGA-UCEC | 53.463104 | 4.11021179 |
| TCGA-AX-A2HH | TCGA-UCEC | 90.7485153 | 16.9872446 |
| TCGA-AX-A2IN | TCGA-UCEC | 87.4840082 | 37.7107406 |
| TCGA-AX-A3FS | TCGA-UCEC | 91.3477451 | 18.8859437 |
| TCGA-AX-A3FT | TCGA-UCEC | 72.1903843 | 22.9923775 |
| TCGA-AX-A3FV | TCGA-UCEC | 73.9010655 | 94.3556229 |
| TCGA-AX-A3FW | TCGA-UCEC | 96.2406203 | 21.9847473 |
| TCGA-AX-A3FX | TCGA-UCEC | 70.7068819 | 19.0110078 |
| TCGA-AX-A3FZ | TCGA-UCEC | 115.930924 | 52.0981206 |
| TCGA-AX-A3G1 | TCGA-UCEC | 95.418537 | 39.6784577 |

| | | | |
|---|---|---|---|
| TCGA-AX-A3G3 | TCGA-UCEC | 97.9214198 | 56.7200384 |
| TCGA-AX-A3G4 | TCGA-UCEC | 30.9286882 | 28.3879037 |
| TCGA-AX-A3G6 | TCGA-UCEC | 91.0352036 | 6.29658956 |
| TCGA-AX-A3G7 | TCGA-UCEC | 65.5428786 | 23.8582623 |
| TCGA-AX-A3G8 | TCGA-UCEC | 6.39055319 | 9.07548966 |
| TCGA-AX-A3G9 | TCGA-UCEC | 79.5424846 | 27.0408356 |
| TCGA-AX-A3GB | TCGA-UCEC | 60.3844072 | 18.6168007 |
| TCGA-AX-A3GI | TCGA-UCEC | 76.9340426 | 40.5690278 |
| TCGA-AY-5543 | TCGA-COAD | 110.428338 | 50.978886 |
| TCGA-AY-6196 | TCGA-COAD | 27.9531734 | 11.795645 |
| TCGA-AY-6197 | TCGA-COAD | 59.7869913 | 16.2352102 |
| TCGA-AY-6386 | TCGA-COAD | 54.4142569 | 21.5760856 |
| TCGA-AY-A54L | TCGA-COAD | 80.5812912 | 32.9548626 |
| TCGA-AY-A69D | TCGA-COAD | 74.0241434 | 43.5575319 |
| TCGA-AY-A71X | TCGA-COAD | 67.1977548 | 21.5817285 |
| TCGA-AY-A8YK | TCGA-COAD | 59.5019551 | 27.2277189 |
| TCGA-AZ-4323 | TCGA-COAD | 62.6045359 | 26.4734582 |
| TCGA-AZ-4616 | TCGA-COAD | 72.8688461 | 37.6142267 |
| TCGA-AZ-4682 | TCGA-COAD | 174.002055 | 45.7037693 |
| TCGA-AZ-5403 | TCGA-COAD | 67.7067116 | 46.4570421 |
| TCGA-AZ-5407 | TCGA-COAD | 55.1866323 | 17.9783665 |
| TCGA-AZ-6598 | TCGA-COAD | 34.8361441 | 10.3177659 |
| TCGA-AZ-6599 | TCGA-COAD | 35.2508672 | 11.2634009 |
| TCGA-AZ-6600 | TCGA-COAD | 45.6879547 | 32.4938512 |
| TCGA-AZ-6601 | TCGA-COAD | 33.7022009 | 11.4235786 |
| TCGA-AZ-6603 | TCGA-COAD | 44.8731286 | 16.6048027 |
| TCGA-AZ-6605 | TCGA-COAD | 63.3613401 | 24.6756743 |
| TCGA-AZ-6606 | TCGA-COAD | 74.3202966 | 29.7026766 |
| TCGA-AZ-6607 | TCGA-COAD | 33.1710095 | 24.3234028 |
| TCGA-AZ-6608 | TCGA-COAD | 92.7327111 | 51.5234508 |
| TCGA-B0-4688 | TCGA-KIRC | 65.8746569 | 66.7361884 |

| | | | |
|---|---|---|---|
| TCGA-B0-4690 | TCGA-KIRC | 69.6482142 | 33.2435718 |
| TCGA-B0-4691 | TCGA-KIRC | 47.7905039 | 24.531742 |
| TCGA-B0-4693 | TCGA-KIRC | 38.597217 | 15.7777766 |
| TCGA-B0-4694 | TCGA-KIRC | 39.1005782 | 12.796607 |
| TCGA-B0-4696 | TCGA-KIRC | 86.2529322 | 100.780282 |
| TCGA-B0-4697 | TCGA-KIRC | 14.0785525 | 6.39839225 |
| TCGA-B0-4698 | TCGA-KIRC | 62.0966026 | 57.0043605 |
| TCGA-B0-4699 | TCGA-KIRC | 63.987977 | 35.4842598 |
| TCGA-B0-4700 | TCGA-KIRC | 19.6890392 | 5.79478649 |
| TCGA-B0-4701 | TCGA-KIRC | 41.38733 | 16.5195878 |
| TCGA-B0-4703 | TCGA-KIRC | 47.1206548 | 21.4308765 |
| TCGA-B0-4706 | TCGA-KIRC | 43.4662072 | 12.8386008 |
| TCGA-B0-4707 | TCGA-KIRC | 22.0690593 | 12.1636751 |
| TCGA-B0-4710 | TCGA-KIRC | 42.3701223 | 17.125404 |
| TCGA-B0-4712 | TCGA-KIRC | 49.4929316 | 34.8094845 |
| TCGA-B0-4713 | TCGA-KIRC | 42.6247162 | 25.0174394 |
| TCGA-B0-4714 | TCGA-KIRC | 41.6077536 | 13.4362419 |
| TCGA-B0-4718 | TCGA-KIRC | 32.0002476 | 16.2328198 |
| TCGA-B0-4810 | TCGA-KIRC | 44.2789192 | 20.2600419 |
| TCGA-B0-4811 | TCGA-KIRC | 31.4382474 | 12.6153361 |
| TCGA-B0-4813 | TCGA-KIRC | 34.8579453 | 11.6137865 |
| TCGA-B0-4814 | TCGA-KIRC | 37.2865579 | 14.5748457 |
| TCGA-B0-4815 | TCGA-KIRC | 30.360665 | 16.9793968 |
| TCGA-B0-4816 | TCGA-KIRC | 48.4780193 | 13.9958123 |
| TCGA-B0-4817 | TCGA-KIRC | 27.3612014 | 16.3041623 |
| TCGA-B0-4818 | TCGA-KIRC | 48.1870777 | 15.5728422 |
| TCGA-B0-4819 | TCGA-KIRC | 28.2412534 | 16.1381477 |
| TCGA-B0-4821 | TCGA-KIRC | 39.301898 | 25.5360327 |
| TCGA-B0-4822 | TCGA-KIRC | 33.8426078 | 13.5060812 |
| TCGA-B0-4823 | TCGA-KIRC | 35.2509641 | 16.8361419 |
| TCGA-B0-4824 | TCGA-KIRC | 51.1786994 | 16.4068174 |

| | | | |
|---|---|---|---|
| TCGA-B0-4827 | TCGA-KIRC | 33.0175458 | 10.4421743 |
| TCGA-B0-4828 | TCGA-KIRC | 39.1774087 | 14.5819259 |
| TCGA-B0-4833 | TCGA-KIRC | 45.1980682 | 14.3066166 |
| TCGA-B0-4834 | TCGA-KIRC | 14.260686 | 8.68182437 |
| TCGA-B0-4836 | TCGA-KIRC | 34.3357758 | 20.5130886 |
| TCGA-B0-4837 | TCGA-KIRC | 37.0153268 | 8.37232559 |
| TCGA-B0-4838 | TCGA-KIRC | 43.109541 | 20.4324412 |
| TCGA-B0-4839 | TCGA-KIRC | 33.6737104 | 13.2546043 |
| TCGA-B0-4841 | TCGA-KIRC | 25.3313269 | 8.78762066 |
| TCGA-B0-4842 | TCGA-KIRC | 36.9234238 | 18.0695633 |
| TCGA-B0-4843 | TCGA-KIRC | 28.8522358 | 15.3660728 |
| TCGA-B0-4844 | TCGA-KIRC | 43.8944775 | 15.1168035 |
| TCGA-B0-4845 | TCGA-KIRC | 44.7806042 | 15.5310036 |
| TCGA-B0-4846 | TCGA-KIRC | 58.9226917 | 18.2129554 |
| TCGA-B0-4847 | TCGA-KIRC | 41.0432716 | 16.054049 |
| TCGA-B0-4848 | TCGA-KIRC | 37.5437537 | 31.2493145 |
| TCGA-B0-4849 | TCGA-KIRC | 62.0291464 | 19.7380689 |
| TCGA-B0-4852 | TCGA-KIRC | 45.5230396 | 17.9420149 |
| TCGA-B0-4945 | TCGA-KIRC | 69.5914282 | 26.0439019 |
| TCGA-B0-5075 | TCGA-KIRC | 29.3927173 | 20.5518592 |
| TCGA-B0-5077 | TCGA-KIRC | 36.3249596 | 14.1142175 |
| TCGA-B0-5080 | TCGA-KIRC | 33.5671782 | 14.8831758 |
| TCGA-B0-5081 | TCGA-KIRC | 31.1022074 | 16.5427267 |
| TCGA-B0-5083 | TCGA-KIRC | 23.872277 | 17.7094009 |
| TCGA-B0-5084 | TCGA-KIRC | 34.638413 | 39.8428552 |
| TCGA-B0-5085 | TCGA-KIRC | 32.4605402 | 8.92505793 |
| TCGA-B0-5088 | TCGA-KIRC | 40.8993381 | 11.5344777 |
| TCGA-B0-5092 | TCGA-KIRC | 49.4194658 | 28.5917971 |
| TCGA-B0-5094 | TCGA-KIRC | 29.8196134 | 24.7694333 |
| TCGA-B0-5095 | TCGA-KIRC | 42.2322468 | 25.9798133 |
| TCGA-B0-5096 | TCGA-KIRC | 22.7778912 | 27.1273843 |

| | | | |
|---|---|---|---|
| TCGA-B0-5097 | TCGA-KIRC | 36.5382999 | 14.6304897 |
| TCGA-B0-5098 | TCGA-KIRC | 30.2074917 | 25.3023608 |
| TCGA-B0-5099 | TCGA-KIRC | 43.6787571 | 18.2657563 |
| TCGA-B0-5100 | TCGA-KIRC | 58.5713961 | 14.8207457 |
| TCGA-B0-5102 | TCGA-KIRC | 29.3565104 | 12.3328059 |
| TCGA-B0-5104 | TCGA-KIRC | 39.5606233 | 10.2719896 |
| TCGA-B0-5106 | TCGA-KIRC | 36.4532117 | 13.609119 |
| TCGA-B0-5107 | TCGA-KIRC | 57.9577067 | 24.7441622 |
| TCGA-B0-5108 | TCGA-KIRC | 40.5929323 | 12.9725373 |
| TCGA-B0-5109 | TCGA-KIRC | 37.7381019 | 22.1455918 |
| TCGA-B0-5110 | TCGA-KIRC | 51.9147228 | 18.3277308 |
| TCGA-B0-5113 | TCGA-KIRC | 35.53373 | 22.0359434 |
| TCGA-B0-5115 | TCGA-KIRC | 43.0981152 | 16.8520195 |
| TCGA-B0-5116 | TCGA-KIRC | 29.5702771 | 14.1013413 |
| TCGA-B0-5117 | TCGA-KIRC | 17.1813814 | 17.8588081 |
| TCGA-B0-5119 | TCGA-KIRC | 48.9142829 | 17.053096 |
| TCGA-B0-5120 | TCGA-KIRC | 37.749017 | 14.9241739 |
| TCGA-B0-5121 | TCGA-KIRC | 35.797514 | 12.7389946 |
| TCGA-B0-5399 | TCGA-KIRC | 52.0249861 | 16.6195027 |
| TCGA-B0-5400 | TCGA-KIRC | 36.061328 | 17.3809366 |
| TCGA-B0-5402 | TCGA-KIRC | 31.2067736 | 9.82978504 |
| TCGA-B0-5690 | TCGA-KIRC | 59.2947691 | 16.3604716 |
| TCGA-B0-5691 | TCGA-KIRC | 48.5844778 | 15.2126147 |
| TCGA-B0-5692 | TCGA-KIRC | 56.9514929 | 27.9385808 |
| TCGA-B0-5693 | TCGA-KIRC | 60.6721383 | 15.9397567 |
| TCGA-B0-5694 | TCGA-KIRC | 37.154854 | 14.7581194 |
| TCGA-B0-5695 | TCGA-KIRC | 54.1935662 | 15.7813849 |
| TCGA-B0-5696 | TCGA-KIRC | 32.4336079 | 8.45432925 |
| TCGA-B0-5697 | TCGA-KIRC | 34.6785453 | 10.4431354 |
| TCGA-B0-5698 | TCGA-KIRC | 49.0869113 | 18.7262818 |
| TCGA-B0-5699 | TCGA-KIRC | 29.5107912 | 8.62172429 |

| | | | |
|---|---|---|---|
| TCGA-B0-5700 | TCGA-KIRC | 47.5126834 | 14.4179966 |
| TCGA-B0-5701 | TCGA-KIRC | 29.9031855 | 8.51841906 |
| TCGA-B0-5702 | TCGA-KIRC | 3.55402505 | 3.98000032 |
| TCGA-B0-5703 | TCGA-KIRC | 52.8840844 | 17.7792803 |
| TCGA-B0-5705 | TCGA-KIRC | 52.3873453 | 15.98172 |
| TCGA-B0-5706 | TCGA-KIRC | 19.2498735 | 6.87788687 |
| TCGA-B0-5707 | TCGA-KIRC | 43.0957734 | 18.8798335 |
| TCGA-B0-5709 | TCGA-KIRC | 43.6922465 | 20.4081477 |
| TCGA-B0-5710 | TCGA-KIRC | 53.3948203 | 19.4747788 |
| TCGA-B0-5711 | TCGA-KIRC | 40.7220255 | 15.9231571 |
| TCGA-B0-5712 | TCGA-KIRC | 46.9407697 | 14.1214872 |
| TCGA-B0-5713 | TCGA-KIRC | 52.3671911 | 19.5293995 |
| TCGA-B0-5812 | TCGA-KIRC | 56.6422728 | 17.071362 |
| TCGA-B1-5398 | TCGA-KIRP | 5.35722722 | 2.77045863 |
| TCGA-B1-7332 | TCGA-KIRP | 28.4359532 | 22.429169 |
| TCGA-B1-A47M | TCGA-KIRP | 52.7879043 | 12.0576805 |
| TCGA-B1-A47N | TCGA-KIRP | 33.4572376 | 12.9096471 |
| TCGA-B1-A47O | TCGA-KIRP | 8.90426786 | 1.72232569 |
| TCGA-B1-A654 | TCGA-KIRP | 29.5773763 | 3.93871568 |
| TCGA-B1-A655 | TCGA-KIRP | 31.6234428 | 10.9916353 |
| TCGA-B1-A656 | TCGA-KIRP | 43.9816943 | 17.8179242 |
| TCGA-B1-A657 | TCGA-KIRP | 40.0425386 | 15.1281987 |
| TCGA-B2-3923 | TCGA-KIRC | 4.83953506 | 5.59817641 |
| TCGA-B2-3924 | TCGA-KIRC | 12.0964275 | 8.03637483 |
| TCGA-B2-4098 | TCGA-KIRC | 10.3809115 | 12.3735895 |
| TCGA-B2-4099 | TCGA-KIRC | 49.2109991 | 24.0020449 |
| TCGA-B2-4101 | TCGA-KIRC | 33.7679674 | 13.6638853 |
| TCGA-B2-4102 | TCGA-KIRC | 32.208337 | 17.7123369 |
| TCGA-B2-5633 | TCGA-KIRC | 36.703244 | 18.6585648 |
| TCGA-B2-5635 | TCGA-KIRC | 3.37378323 | 5.20455641 |
| TCGA-B2-5636 | TCGA-KIRC | 50.1311032 | 17.1711058 |

| | | | |
|---|---|---|---|
| TCGA-B2-5639 | TCGA-KIRC | 44.328855 | 21.4389749 |
| TCGA-B2-5641 | TCGA-KIRC | 37.1474631 | 8.71201001 |
| TCGA-B2-A4SR | TCGA-KIRC | 54.9689709 | 12.6632813 |
| TCGA-B3-3925 | TCGA-KIRP | 13.0110893 | 86.7717774 |
| TCGA-B3-3926 | TCGA-KIRP | 18.5581859 | 26.7329212 |
| TCGA-B3-4103 | TCGA-KIRP | 31.3614766 | 15.1109219 |
| TCGA-B3-4104 | TCGA-KIRP | 26.0502197 | 8.23200015 |
| TCGA-B3-8121 | TCGA-KIRP | 129.679192 | 5.90823728 |
| TCGA-B3-A6W5 | TCGA-KIRP | 36.7946852 | 29.6580502 |
| TCGA-B4-5377 | TCGA-KIRC | 67.2745197 | 18.4384545 |
| TCGA-B4-5378 | TCGA-KIRC | 58.671687 | 12.0599437 |
| TCGA-B4-5832 | TCGA-KIRC | 42.8284834 | 74.4454915 |
| TCGA-B4-5834 | TCGA-KIRC | 50.0766243 | 16.8963455 |
| TCGA-B4-5835 | TCGA-KIRC | 37.4508573 | 26.8981923 |
| TCGA-B4-5836 | TCGA-KIRC | 50.5079857 | 24.988255 |
| TCGA-B4-5838 | TCGA-KIRC | 28.0683648 | 20.307689 |
| TCGA-B4-5843 | TCGA-KIRC | 45.2122751 | 21.2969916 |
| TCGA-B5-A0JN | TCGA-UCEC | 108.002721 | 32.7507424 |
| TCGA-B5-A0JR | TCGA-UCEC | 69.6416571 | 25.0006246 |
| TCGA-B5-A0K9 | TCGA-UCEC | 57.6127272 | 31.2425793 |
| TCGA-B5-A11R | TCGA-UCEC | 53.2523416 | 9.94483593 |
| TCGA-B5-A1MS | TCGA-UCEC | 100.34543 | 7.99116024 |
| TCGA-B5-A1MW | TCGA-UCEC | 58.2638592 | 22.2061576 |
| TCGA-B5-A3F9 | TCGA-UCEC | 117.491425 | 13.6846141 |
| TCGA-B5-A3FA | TCGA-UCEC | 82.6870173 | 24.2164916 |
| TCGA-B5-A3FB | TCGA-UCEC | 57.7544216 | 21.2336596 |
| TCGA-B5-A3FC | TCGA-UCEC | 76.3264802 | 16.7355089 |
| TCGA-B5-A3FD | TCGA-UCEC | 112.277028 | 30.3707147 |
| TCGA-B5-A3FH | TCGA-UCEC | 53.3603672 | 15.6984806 |
| TCGA-B5-A3S1 | TCGA-UCEC | 125.030796 | 22.9294831 |
| TCGA-B5-A5OC | TCGA-UCEC | 57.1190287 | 67.0652075 |

| | | | |
|---|---|---|---|
| TCGA-B5-A5OD | TCGA-UCEC | 100.827938 | 28.6459977 |
| TCGA-B6-A0I1 | TCGA-BRCA | 148.380179 | 38.3778334 |
| TCGA-B6-A0I2 | TCGA-BRCA | 111.441745 | 57.7523954 |
| TCGA-B6-A0I5 | TCGA-BRCA | 48.7323302 | 24.1170371 |
| TCGA-B6-A0I6 | TCGA-BRCA | 112.035125 | 49.2981087 |
| TCGA-B6-A0I8 | TCGA-BRCA | 35.670922 | 28.8271024 |
| TCGA-B6-A0I9 | TCGA-BRCA | 31.9090492 | 40.0891194 |
| TCGA-B6-A0IA | TCGA-BRCA | 51.0057291 | 26.8659764 |
| TCGA-B6-A0IB | TCGA-BRCA | 37.3971978 | 48.9490773 |
| TCGA-B6-A0IC | TCGA-BRCA | 50.2558614 | 13.4105342 |
| TCGA-B6-A0IE | TCGA-BRCA | 77.2463131 | 39.4543003 |
| TCGA-B6-A0IG | TCGA-BRCA | 30.9975665 | 39.5651027 |
| TCGA-B6-A0IH | TCGA-BRCA | 36.7190042 | 26.1198769 |
| TCGA-B6-A0IJ | TCGA-BRCA | 155.4528 | 100.099267 |
| TCGA-B6-A0IK | TCGA-BRCA | 74.1451745 | 43.19291 |
| TCGA-B6-A0IM | TCGA-BRCA | 53.2185322 | 22.3071324 |
| TCGA-B6-A0IN | TCGA-BRCA | 30.3865658 | 19.4957929 |
| TCGA-B6-A0IO | TCGA-BRCA | 56.3993711 | 40.5192566 |
| TCGA-B6-A0IP | TCGA-BRCA | 51.8704017 | 25.8426191 |
| TCGA-B6-A0IQ | TCGA-BRCA | 165.765611 | 46.9622967 |
| TCGA-B6-A0RE | TCGA-BRCA | 119.527359 | 75.285161 |
| TCGA-B6-A0RG | TCGA-BRCA | 50.9172725 | 31.9333706 |
| TCGA-B6-A0RH | TCGA-BRCA | 69.624219 | 42.8713754 |
| TCGA-B6-A0RI | TCGA-BRCA | 56.076278 | 19.0166848 |
| TCGA-B6-A0RL | TCGA-BRCA | 74.0702372 | 57.6635803 |
| TCGA-B6-A0RM | TCGA-BRCA | 51.6952022 | 18.4547243 |
| TCGA-B6-A0RN | TCGA-BRCA | 51.4787893 | 26.2788909 |
| TCGA-B6-A0RO | TCGA-BRCA | 36.0152496 | 40.0021517 |
| TCGA-B6-A0RP | TCGA-BRCA | 47.6016707 | 20.0747816 |
| TCGA-B6-A0RQ | TCGA-BRCA | 44.6942599 | 16.7677992 |
| TCGA-B6-A0RS | TCGA-BRCA | 61.9586689 | 51.8966886 |

| | | | |
|---|---|---|---|
| TCGA-B6-A0RT | TCGA-BRCA | 67.7369072 | 34.6854836 |
| TCGA-B6-A0RU | TCGA-BRCA | 103.004468 | 25.9045219 |
| TCGA-B6-A0RV | TCGA-BRCA | 48.88301 | 16.1304831 |
| TCGA-B6-A0WS | TCGA-BRCA | 48.5910601 | 30.2105451 |
| TCGA-B6-A0WT | TCGA-BRCA | 47.4138504 | 35.1612181 |
| TCGA-B6-A0WV | TCGA-BRCA | 56.0739347 | 55.4258595 |
| TCGA-B6-A0WW | TCGA-BRCA | 37.3048332 | 44.6855961 |
| TCGA-B6-A0WX | TCGA-BRCA | 110.125843 | 41.880815 |
| TCGA-B6-A0WY | TCGA-BRCA | 47.8266584 | 27.3883268 |
| TCGA-B6-A0WZ | TCGA-BRCA | 40.0199505 | 21.5024253 |
| TCGA-B6-A0X0 | TCGA-BRCA | 40.9825094 | 20.5428501 |
| TCGA-B6-A0X1 | TCGA-BRCA | 225.138235 | 113.492739 |
| TCGA-B6-A0X4 | TCGA-BRCA | 40.623834 | 24.4921648 |
| TCGA-B6-A0X5 | TCGA-BRCA | 67.360672 | 50.4858484 |
| TCGA-B6-A0X7 | TCGA-BRCA | 29.6173463 | 22.6643388 |
| TCGA-B6-A1KC | TCGA-BRCA | 46.4401646 | 31.1758425 |
| TCGA-B6-A1KF | TCGA-BRCA | 55.9696233 | 45.5310671 |
| TCGA-B6-A1KI | TCGA-BRCA | 48.0040297 | 40.0113356 |
| TCGA-B6-A1KN | TCGA-BRCA | 36.6455628 | 24.4175453 |
| TCGA-B6-A2IU | TCGA-BRCA | 54.5242208 | 23.2824143 |
| TCGA-B6-A3ZX | TCGA-BRCA | 58.7088717 | 24.5676319 |
| TCGA-B6-A400 | TCGA-BRCA | 104.474089 | 54.8760019 |
| TCGA-B6-A401 | TCGA-BRCA | 58.1719905 | 35.4669802 |
| TCGA-B6-A402 | TCGA-BRCA | 52.9712754 | 33.6362946 |
| TCGA-B6-A408 | TCGA-BRCA | 59.0988262 | 26.4643276 |
| TCGA-B6-A409 | TCGA-BRCA | 143.671919 | 49.9622392 |
| TCGA-B6-A40B | TCGA-BRCA | 42.8846162 | 32.9070611 |
| TCGA-B6-A40C | TCGA-BRCA | 51.93319 | 22.3242574 |
| TCGA-B7-5818 | TCGA-STAD | 70.5510528 | 24.8355876 |
| TCGA-B7-A5TI | TCGA-STAD | 42.1410548 | 25.6734224 |
| TCGA-B7-A5TJ | TCGA-STAD | 22.8247657 | 23.0491927 |

| | | | |
|---|---|---|---|
| TCGA-B7-A5TK | TCGA-STAD | 46.938742 | 44.2475342 |
| TCGA-B7-A5TN | TCGA-STAD | 35.2180508 | 30.5896466 |
| TCGA-B8-4143 | TCGA-KIRC | 38.4449901 | 18.4505205 |
| TCGA-B8-4146 | TCGA-KIRC | 64.9646067 | 50.0185882 |
| TCGA-B8-4148 | TCGA-KIRC | 43.5159886 | 13.3013332 |
| TCGA-B8-4151 | TCGA-KIRC | 44.919983 | 14.7040245 |
| TCGA-B8-4153 | TCGA-KIRC | 46.918589 | 12.2230135 |
| TCGA-B8-4154 | TCGA-KIRC | 38.57215 | 16.7868662 |
| TCGA-B8-4619 | TCGA-KIRC | 25.7876597 | 10.464432 |
| TCGA-B8-4620 | TCGA-KIRC | 29.8191609 | 14.2245566 |
| TCGA-B8-4621 | TCGA-KIRC | 24.4569036 | 12.1765207 |
| TCGA-B8-4622 | TCGA-KIRC | 30.5836601 | 15.5542433 |
| TCGA-B8-5158 | TCGA-KIRC | 34.7771842 | 17.0802738 |
| TCGA-B8-5159 | TCGA-KIRC | 40.661652 | 17.8734268 |
| TCGA-B8-5162 | TCGA-KIRC | 19.7294965 | 10.8003045 |
| TCGA-B8-5163 | TCGA-KIRC | 39.2208665 | 24.0022772 |
| TCGA-B8-5164 | TCGA-KIRC | 47.4021868 | 19.6222045 |
| TCGA-B8-5165 | TCGA-KIRC | 39.8577282 | 12.096923 |
| TCGA-B8-5545 | TCGA-KIRC | 47.3661947 | 15.4462399 |
| TCGA-B8-5546 | TCGA-KIRC | 55.3663587 | 21.9498919 |
| TCGA-B8-5549 | TCGA-KIRC | 40.6489739 | 12.2551783 |
| TCGA-B8-5550 | TCGA-KIRC | 42.5028022 | 21.3349098 |
| TCGA-B8-5551 | TCGA-KIRC | 39.2660349 | 14.8273152 |
| TCGA-B8-5552 | TCGA-KIRC | 51.9274831 | 20.8558889 |
| TCGA-B8-5553 | TCGA-KIRC | 42.1184482 | 10.9298964 |
| TCGA-B8-A54D | TCGA-KIRC | 42.6724914 | 16.3019588 |
| TCGA-B8-A54E | TCGA-KIRC | 42.5621416 | 14.3520408 |
| TCGA-B8-A54F | TCGA-KIRC | 43.2584668 | 21.0864618 |
| TCGA-B8-A54G | TCGA-KIRC | 61.9567048 | 18.9763425 |
| TCGA-B8-A54H | TCGA-KIRC | 49.0357289 | 23.2972369 |
| TCGA-B8-A54I | TCGA-KIRC | 15.7643081 | 14.1748057 |

| | | | |
|---|---|---|---|
| TCGA-B8-A54J | TCGA-KIRC | 69.3807485 | 19.8226747 |
| TCGA-B8-A54K | TCGA-KIRC | 43.4941359 | 16.1228536 |
| TCGA-B8-A7U6 | TCGA-KIRC | 76.2159791 | 20.0749427 |
| TCGA-B8-A8YJ | TCGA-KIRC | 45.7210401 | 13.8064639 |
| TCGA-B9-4113 | TCGA-KIRP | 33.4969532 | 17.0014536 |
| TCGA-B9-4115 | TCGA-KIRP | 48.2464081 | 21.5933382 |
| TCGA-B9-4116 | TCGA-KIRP | 28.4512451 | 35.9432868 |
| TCGA-B9-4117 | TCGA-KIRP | 25.1486756 | 11.7061083 |
| TCGA-B9-4617 | TCGA-KIRP | 40.5341037 | 7.35080552 |
| TCGA-B9-5155 | TCGA-KIRP | 42.8517877 | 5.87438918 |
| TCGA-B9-5156 | TCGA-KIRP | 8.66265066 | 3.32928863 |
| TCGA-B9-7268 | TCGA-KIRP | 40.127379 | 26.0549746 |
| TCGA-B9-A44B | TCGA-KIRP | 4.66383826 | 1.30178654 |
| TCGA-B9-A5W7 | TCGA-KIRP | 41.3484445 | 18.7296949 |
| TCGA-B9-A5W8 | TCGA-KIRP | 20.9199376 | 17.581971 |
| TCGA-B9-A5W9 | TCGA-KIRP | 35.4608905 | 23.6036597 |
| TCGA-B9-A69E | TCGA-KIRP | 25.7091103 | 14.7028897 |
| TCGA-B9-A8YH | TCGA-KIRP | 44.8257987 | 19.4775503 |
| TCGA-B9-A8YI | TCGA-KIRP | 24.847167 | 2.64666599 |
| TCGA-BC-4072 | TCGA-LIHC | 49.8400575 | 14.8499564 |
| TCGA-BC-4073 | TCGA-LIHC | 42.2023155 | 12.3683142 |
| TCGA-BC-A10Q | TCGA-LIHC | 21.3511291 | 8.63709055 |
| TCGA-BC-A10R | TCGA-LIHC | 24.1025994 | 8.42433702 |
| TCGA-BC-A10S | TCGA-LIHC | 35.2332049 | 16.7440028 |
| TCGA-BC-A10T | TCGA-LIHC | 28.211217 | 12.1005524 |
| TCGA-BC-A10U | TCGA-LIHC | 23.075404 | 8.09276555 |
| TCGA-BC-A10W | TCGA-LIHC | 24.7483608 | 9.94402391 |
| TCGA-BC-A10X | TCGA-LIHC | 21.4485529 | 7.56494309 |
| TCGA-BC-A10Y | TCGA-LIHC | 53.4948061 | 13.9090214 |
| TCGA-BC-A10Z | TCGA-LIHC | 31.0454563 | 8.47814313 |
| TCGA-BC-A110 | TCGA-LIHC | 30.7009 | 8.4991672 |

| | | | |
|---|---|---|---|
| TCGA-BC-A112 | TCGA-LIHC | 45.1631109 | 19.5131167 |
| TCGA-BC-A216 | TCGA-LIHC | 20.0748457 | 8.77614659 |
| TCGA-BC-A217 | TCGA-LIHC | 69.9906172 | 21.2092558 |
| TCGA-BC-A3KF | TCGA-LIHC | 57.6743228 | 15.6903494 |
| TCGA-BC-A3KG | TCGA-LIHC | 88.3123506 | 41.3516683 |
| TCGA-BC-A5W4 | TCGA-LIHC | 44.7418497 | 9.27963849 |
| TCGA-BC-A69H | TCGA-LIHC | 30.2725312 | 13.7023437 |
| TCGA-BC-A69I | TCGA-LIHC | 36.1862123 | 14.2882509 |
| TCGA-BC-A8YO | TCGA-LIHC | 102.056586 | 31.3374023 |
| TCGA-BD-A2L6 | TCGA-LIHC | 49.375785 | 10.5991583 |
| TCGA-BD-A3EP | TCGA-LIHC | 32.6927386 | 9.26227015 |
| TCGA-BD-A3ER | TCGA-LIHC | 29.593005 | 10.7003572 |
| TCGA-BF-A1PU | TCGA-SKCM | 21.6299778 | 33.121948 |
| TCGA-BF-A1PV | TCGA-SKCM | 43.5691578 | 29.4163326 |
| TCGA-BF-A1PX | TCGA-SKCM | 39.0701288 | 17.8560058 |
| TCGA-BF-A1PZ | TCGA-SKCM | 53.1118305 | 21.1960585 |
| TCGA-BF-A1Q0 | TCGA-SKCM | 78.810172 | 28.4500783 |
| TCGA-BF-A3DJ | TCGA-SKCM | 53.1466291 | 31.7433772 |
| TCGA-BF-A3DL | TCGA-SKCM | 34.9371093 | 29.4358387 |
| TCGA-BF-A3DM | TCGA-SKCM | 36.0336556 | 28.1228758 |
| TCGA-BF-A3DN | TCGA-SKCM | 46.7500758 | 24.6408189 |
| TCGA-BF-A5EO | TCGA-SKCM | 70.7037859 | 14.3653292 |
| TCGA-BF-A5EP | TCGA-SKCM | 39.5639363 | 33.2289078 |
| TCGA-BF-A5EQ | TCGA-SKCM | 37.5242483 | 23.1142054 |
| TCGA-BF-A5ER | TCGA-SKCM | 25.7658624 | 10.343143 |
| TCGA-BF-A5ES | TCGA-SKCM | 28.3510665 | 16.383431 |
| TCGA-BF-A9VF | TCGA-SKCM | 59.9576021 | 21.2958063 |
| TCGA-BF-AAOU | TCGA-SKCM | 30.0634452 | 49.6152731 |
| TCGA-BF-AAOX | TCGA-SKCM | 38.0798852 | 22.488114 |
| TCGA-BF-AAP0 | TCGA-SKCM | 40.5464206 | 35.7538525 |
| TCGA-BF-AAP1 | TCGA-SKCM | 85.5049304 | 77.820223 |

| | | | |
|---|---|---|---|
| TCGA-BF-AAP2 | TCGA-SKCM | 65.5471546 | 21.7689045 |
| TCGA-BF-AAP4 | TCGA-SKCM | 88.2915527 | 42.2104513 |
| TCGA-BF-AAP6 | TCGA-SKCM | 63.920833 | 44.6107466 |
| TCGA-BF-AAP7 | TCGA-SKCM | 43.4704517 | 18.8381272 |
| TCGA-BF-AAP8 | TCGA-SKCM | 63.1950827 | 43.5290574 |
| TCGA-BG-A0MK | TCGA-UCEC | 54.1407468 | 17.4491241 |
| TCGA-BG-A3EW | TCGA-UCEC | 74.2322042 | 29.0384953 |
| TCGA-BG-A3PP | TCGA-UCEC | 80.6709647 | 12.5768006 |
| TCGA-BH-A0AU | TCGA-BRCA | 39.8943668 | 15.4796584 |
| TCGA-BH-A0AV | TCGA-BRCA | 81.1761683 | 57.5611374 |
| TCGA-BH-A0AW | TCGA-BRCA | 42.5838174 | 34.2872718 |
| TCGA-BH-A0AY | TCGA-BRCA | 41.1705667 | 13.2894973 |
| TCGA-BH-A0AZ | TCGA-BRCA | 49.3025177 | 20.1728214 |
| TCGA-BH-A0B0 | TCGA-BRCA | 61.2882328 | 35.8273487 |
| TCGA-BH-A0B1 | TCGA-BRCA | 39.9148638 | 44.1067556 |
| TCGA-BH-A0B2 | TCGA-BRCA | 60.613534 | 23.1626163 |
| TCGA-BH-A0B3 | TCGA-BRCA | 52.6079213 | 16.8668628 |
| TCGA-BH-A0B4 | TCGA-BRCA | 44.4574962 | 20.7632072 |
| TCGA-BH-A0B5 | TCGA-BRCA | 27.2248079 | 15.6508306 |
| TCGA-BH-A0B6 | TCGA-BRCA | 33.5222126 | 20.3638526 |
| TCGA-BH-A0B7 | TCGA-BRCA | 55.3672169 | 24.1327379 |
| TCGA-BH-A0B8 | TCGA-BRCA | 24.5013156 | 10.8308847 |
| TCGA-BH-A0B9 | TCGA-BRCA | 111.316583 | 146.309754 |
| TCGA-BH-A0BA | TCGA-BRCA | 56.627001 | 71.6432823 |
| TCGA-BH-A0BC | TCGA-BRCA | 45.9382936 | 14.0646965 |
| TCGA-BH-A0BD | TCGA-BRCA | 70.8836853 | 28.1461508 |
| TCGA-BH-A0BF | TCGA-BRCA | 37.485443 | 22.4712325 |
| TCGA-BH-A0BG | TCGA-BRCA | 74.8099523 | 53.4687436 |
| TCGA-BH-A0BJ | TCGA-BRCA | 54.3586703 | 13.3748412 |
| TCGA-BH-A0BL | TCGA-BRCA | 74.4527094 | 78.2394566 |
| TCGA-BH-A0BM | TCGA-BRCA | 53.1741561 | 20.7110148 |

| | | | |
|---|---|---|---|
| TCGA-BH-A0BO | TCGA-BRCA | 51.3828962 | 17.1362394 |
| TCGA-BH-A0BP | TCGA-BRCA | 45.9149521 | 16.9736255 |
| TCGA-BH-A0BQ | TCGA-BRCA | 38.5846429 | 16.0101888 |
| TCGA-BH-A0BR | TCGA-BRCA | 62.5480175 | 31.3492664 |
| TCGA-BH-A0BS | TCGA-BRCA | 49.2942193 | 15.9092455 |
| TCGA-BH-A0BT | TCGA-BRCA | 50.3996389 | 19.1054317 |
| TCGA-BH-A0BV | TCGA-BRCA | 40.5015011 | 16.9950277 |
| TCGA-BH-A0BW | TCGA-BRCA | 72.9440951 | 39.0001836 |
| TCGA-BH-A0BZ | TCGA-BRCA | 36.5155266 | 13.8724958 |
| TCGA-BH-A0C0 | TCGA-BRCA | 46.3674646 | 20.9805484 |
| TCGA-BH-A0C1 | TCGA-BRCA | 34.0142347 | 54.3008897 |
| TCGA-BH-A0C3 | TCGA-BRCA | 36.8890275 | 18.8239485 |
| TCGA-BH-A0C7 | TCGA-BRCA | 61.0786613 | 40.8871329 |
| TCGA-BH-A0DD | TCGA-BRCA | 66.0668671 | 33.8977397 |
| TCGA-BH-A0DE | TCGA-BRCA | 32.8516726 | 21.7941846 |
| TCGA-BH-A0DG | TCGA-BRCA | 49.5483541 | 15.5952889 |
| TCGA-BH-A0DH | TCGA-BRCA | 44.5262427 | 17.6912157 |
| TCGA-BH-A0DI | TCGA-BRCA | 45.0142769 | 18.7232699 |
| TCGA-BH-A0DK | TCGA-BRCA | 45.0864354 | 26.02011 |
| TCGA-BH-A0DL | TCGA-BRCA | 44.8484494 | 17.40474 |
| TCGA-BH-A0DO | TCGA-BRCA | 41.2747943 | 30.2302668 |
| TCGA-BH-A0DP | TCGA-BRCA | 53.8860995 | 31.0243915 |
| TCGA-BH-A0DQ | TCGA-BRCA | 44.8740755 | 15.399645 |
| TCGA-BH-A0DS | TCGA-BRCA | 41.8141573 | 25.4977374 |
| TCGA-BH-A0DT | TCGA-BRCA | 46.6973514 | 24.1187616 |
| TCGA-BH-A0DV | TCGA-BRCA | 63.9724897 | 20.7138646 |
| TCGA-BH-A0DX | TCGA-BRCA | 49.5852082 | 38.8706664 |
| TCGA-BH-A0DZ | TCGA-BRCA | 49.0598605 | 22.1866282 |
| TCGA-BH-A0E0 | TCGA-BRCA | 48.9976465 | 18.7922451 |
| TCGA-BH-A0E1 | TCGA-BRCA | 49.0911038 | 31.9638907 |
| TCGA-BH-A0E2 | TCGA-BRCA | 45.3330684 | 71.8180525 |

| | | | |
|---|---|---|---|
| TCGA-BH-A0E6 | TCGA-BRCA | 88.5691534 | 35.3494017 |
| TCGA-BH-A0E7 | TCGA-BRCA | 51.8772708 | 24.5945006 |
| TCGA-BH-A0E9 | TCGA-BRCA | 31.7536143 | 21.2190089 |
| TCGA-BH-A0EA | TCGA-BRCA | 39.0737117 | 19.8557414 |
| TCGA-BH-A0EB | TCGA-BRCA | 47.8261089 | 29.2249743 |
| TCGA-BH-A0EE | TCGA-BRCA | 99.1438042 | 21.8758507 |
| TCGA-BH-A0EI | TCGA-BRCA | 46.9815362 | 27.1419077 |
| TCGA-BH-A0GY | TCGA-BRCA | 124.414839 | 46.689223 |
| TCGA-BH-A0GZ | TCGA-BRCA | 42.3062392 | 45.3942919 |
| TCGA-BH-A0H0 | TCGA-BRCA | 74.0176893 | 34.0083322 |
| TCGA-BH-A0H3 | TCGA-BRCA | 53.5136061 | 26.2795438 |
| TCGA-BH-A0H5 | TCGA-BRCA | 46.6665166 | 17.2737767 |
| TCGA-BH-A0H6 | TCGA-BRCA | 56.5668316 | 29.2597924 |
| TCGA-BH-A0H7 | TCGA-BRCA | 54.2490035 | 20.462147 |
| TCGA-BH-A0H9 | TCGA-BRCA | 28.870512 | 13.9653253 |
| TCGA-BH-A0HA | TCGA-BRCA | 80.6209875 | 35.3866941 |
| TCGA-BH-A0HB | TCGA-BRCA | 61.7131081 | 32.1932869 |
| TCGA-BH-A0HF | TCGA-BRCA | 48.8130864 | 31.6578793 |
| TCGA-BH-A0HI | TCGA-BRCA | 51.5103674 | 26.7829584 |
| TCGA-BH-A0HK | TCGA-BRCA | 62.7693186 | 16.2692701 |
| TCGA-BH-A0HL | TCGA-BRCA | 57.3323811 | 29.5674404 |
| TCGA-BH-A0HN | TCGA-BRCA | 57.6618879 | 24.7107667 |
| TCGA-BH-A0HO | TCGA-BRCA | 45.7620689 | 33.0990366 |
| TCGA-BH-A0HP | TCGA-BRCA | 40.8552062 | 26.3442388 |
| TCGA-BH-A0HQ | TCGA-BRCA | 39.2566541 | 26.5914808 |
| TCGA-BH-A0HU | TCGA-BRCA | 233.852524 | 50.0181798 |
| TCGA-BH-A0HW | TCGA-BRCA | 39.443229 | 22.4779688 |
| TCGA-BH-A0HX | TCGA-BRCA | 42.9137473 | 40.8509251 |
| TCGA-BH-A0HY | TCGA-BRCA | 45.1951835 | 30.7761567 |
| TCGA-BH-A0RX | TCGA-BRCA | 123.943956 | 38.2112081 |
| TCGA-BH-A0W3 | TCGA-BRCA | 49.2377772 | 29.7153159 |

| | | | |
|---|---|---|---|
| TCGA-BH-A0W4 | TCGA-BRCA | 47.9738664 | 17.78488 |
| TCGA-BH-A0W5 | TCGA-BRCA | 44.8193891 | 18.6950444 |
| TCGA-BH-A0W7 | TCGA-BRCA | 60.7810276 | 39.5212492 |
| TCGA-BH-A0WA | TCGA-BRCA | 94.9533802 | 32.5792297 |
| TCGA-BH-A18F | TCGA-BRCA | 63.1116999 | 51.4545709 |
| TCGA-BH-A18G | TCGA-BRCA | 48.5998491 | 28.9622581 |
| TCGA-BH-A18H | TCGA-BRCA | 53.296542 | 19.1576832 |
| TCGA-BH-A18I | TCGA-BRCA | 67.2589063 | 28.6275926 |
| TCGA-BH-A18J | TCGA-BRCA | 51.3601232 | 31.9226273 |
| TCGA-BH-A18K | TCGA-BRCA | 37.3861083 | 14.5811696 |
| TCGA-BH-A18L | TCGA-BRCA | 52.7215028 | 28.5217872 |
| TCGA-BH-A18M | TCGA-BRCA | 48.0374617 | 15.9239674 |
| TCGA-BH-A18P | TCGA-BRCA | 28.0227893 | 9.38695715 |
| TCGA-BH-A18Q | TCGA-BRCA | 64.094226 | 70.2762241 |
| TCGA-BH-A18R | TCGA-BRCA | 45.4563337 | 35.3639574 |
| TCGA-BH-A18S | TCGA-BRCA | 63.1362319 | 36.6048744 |
| TCGA-BH-A18T | TCGA-BRCA | 144.487739 | 108.23476 |
| TCGA-BH-A18U | TCGA-BRCA | 42.5863459 | 27.1385041 |
| TCGA-BH-A18V | TCGA-BRCA | 52.9796091 | 17.3146223 |
| TCGA-BH-A1EN | TCGA-BRCA | 31.683395 | 50.007699 |
| TCGA-BH-A1EO | TCGA-BRCA | 42.313235 | 31.7390172 |
| TCGA-BH-A1ES | TCGA-BRCA | 88.7965868 | 51.6816206 |
| TCGA-BH-A1ET | TCGA-BRCA | 54.7444287 | 27.2978313 |
| TCGA-BH-A1EU | TCGA-BRCA | 40.2416436 | 35.5338739 |
| TCGA-BH-A1EV | TCGA-BRCA | 45.2286481 | 19.9053837 |
| TCGA-BH-A1EW | TCGA-BRCA | 60.5805978 | 45.3748577 |
| TCGA-BH-A1EX | TCGA-BRCA | 41.4741868 | 31.2091557 |
| TCGA-BH-A1EY | TCGA-BRCA | 74.4132091 | 31.2724825 |
| TCGA-BH-A1F0 | TCGA-BRCA | 47.55339 | 15.7189322 |
| TCGA-BH-A1F2 | TCGA-BRCA | 33.7080228 | 12.4228131 |
| TCGA-BH-A1F5 | TCGA-BRCA | 37.1112844 | 25.9966227 |

| | | | |
|---|---|---|---|
| TCGA-BH-A1F6 | TCGA-BRCA | 78.6959229 | 41.5683402 |
| TCGA-BH-A1F8 | TCGA-BRCA | 53.1418557 | 21.5166222 |
| TCGA-BH-A1FB | TCGA-BRCA | 48.8923645 | 18.0883816 |
| TCGA-BH-A1FC | TCGA-BRCA | 69.5751311 | 16.8672253 |
| TCGA-BH-A1FD | TCGA-BRCA | 40.0895514 | 13.7636919 |
| TCGA-BH-A1FE | TCGA-BRCA | 51.5703309 | 30.982953 |
| TCGA-BH-A1FG | TCGA-BRCA | 24.5838589 | 18.1231219 |
| TCGA-BH-A1FH | TCGA-BRCA | 59.3628325 | 20.4316933 |
| TCGA-BH-A1FJ | TCGA-BRCA | 84.7424696 | 68.284832 |
| TCGA-BH-A1FL | TCGA-BRCA | 71.1800751 | 29.6953434 |
| TCGA-BH-A1FM | TCGA-BRCA | 57.2257187 | 16.1762857 |
| TCGA-BH-A1FN | TCGA-BRCA | 103.438222 | 34.9319838 |
| TCGA-BH-A1FR | TCGA-BRCA | 48.0212079 | 20.7974377 |
| TCGA-BH-A1FU | TCGA-BRCA | 50.8101684 | 36.5312736 |
| TCGA-BH-A201 | TCGA-BRCA | 63.4871879 | 21.2691585 |
| TCGA-BH-A202 | TCGA-BRCA | 51.3336693 | 47.8059893 |
| TCGA-BH-A203 | TCGA-BRCA | 37.7043361 | 14.3146125 |
| TCGA-BH-A204 | TCGA-BRCA | 32.1757132 | 9.81726746 |
| TCGA-BH-A208 | TCGA-BRCA | 50.0708802 | 13.6938157 |
| TCGA-BH-A209 | TCGA-BRCA | 43.5766505 | 19.2502544 |
| TCGA-BH-A28O | TCGA-BRCA | 40.2085218 | 13.7522585 |
| TCGA-BH-A28Q | TCGA-BRCA | 54.0503777 | 34.8927295 |
| TCGA-BH-A2L8 | TCGA-BRCA | 55.7634958 | 19.7412871 |
| TCGA-BH-A42T | TCGA-BRCA | 45.3583222 | 34.3388418 |
| TCGA-BH-A42U | TCGA-BRCA | 55.1438432 | 15.1877983 |
| TCGA-BH-A42V | TCGA-BRCA | 47.2369551 | 17.8167015 |
| TCGA-BH-A5IZ | TCGA-BRCA | 90.8763218 | 79.1029271 |
| TCGA-BH-A5J0 | TCGA-BRCA | 58.9469036 | 34.7699849 |
| TCGA-BH-A6R8 | TCGA-BRCA | 68.1893479 | 52.1283516 |
| TCGA-BH-A6R9 | TCGA-BRCA | 63.3004531 | 28.0720323 |
| TCGA-BH-A8FY | TCGA-BRCA | 81.747123 | 51.4830614 |

| | | | |
|---|---|---|---|
| TCGA-BH-A8FZ | TCGA-BRCA | 72.6926726 | 18.4802022 |
| TCGA-BH-A8G0 | TCGA-BRCA | 51.3583741 | 18.988977 |
| TCGA-BH-AB28 | TCGA-BRCA | 58.7783664 | 29.4145854 |
| TCGA-BJ-A0YZ | TCGA-THCA | 43.9448318 | 13.5122288 |
| TCGA-BJ-A0Z0 | TCGA-THCA | 64.6000474 | 18.89503 |
| TCGA-BJ-A0Z2 | TCGA-THCA | 70.9203089 | 19.3436572 |
| TCGA-BJ-A0Z3 | TCGA-THCA | 58.3678551 | 14.0089558 |
| TCGA-BJ-A0Z5 | TCGA-THCA | 45.9157551 | 11.3056747 |
| TCGA-BJ-A0Z9 | TCGA-THCA | 40.5332984 | 16.6433144 |
| TCGA-BJ-A0ZA | TCGA-THCA | 48.6470241 | 16.2494721 |
| TCGA-BJ-A0ZB | TCGA-THCA | 43.4516577 | 9.05561968 |
| TCGA-BJ-A0ZC | TCGA-THCA | 49.85347 | 15.3718019 |
| TCGA-BJ-A0ZE | TCGA-THCA | 28.8332055 | 7.80629745 |
| TCGA-BJ-A0ZF | TCGA-THCA | 60.0304005 | 17.6275426 |
| TCGA-BJ-A0ZG | TCGA-THCA | 48.3910368 | 10.7413291 |
| TCGA-BJ-A0ZH | TCGA-THCA | 52.751193 | 14.1041361 |
| TCGA-BJ-A0ZJ | TCGA-THCA | 61.604793 | 13.2487897 |
| TCGA-BJ-A18Y | TCGA-THCA | 54.0679731 | 14.8812057 |
| TCGA-BJ-A18Z | TCGA-THCA | 43.3458296 | 12.7685566 |
| TCGA-BJ-A190 | TCGA-THCA | 62.3059408 | 24.1435332 |
| TCGA-BJ-A191 | TCGA-THCA | 75.6092626 | 15.4259005 |
| TCGA-BJ-A192 | TCGA-THCA | 46.0931565 | 19.5438157 |
| TCGA-BJ-A28R | TCGA-THCA | 48.414834 | 13.6233727 |
| TCGA-BJ-A28S | TCGA-THCA | 67.1954855 | 13.3559579 |
| TCGA-BJ-A28T | TCGA-THCA | 44.2219384 | 11.396574 |
| TCGA-BJ-A28V | TCGA-THCA | 60.9036289 | 18.5897105 |
| TCGA-BJ-A28W | TCGA-THCA | 19.4694441 | 10.6007252 |
| TCGA-BJ-A28X | TCGA-THCA | 44.0131059 | 10.7450457 |
| TCGA-BJ-A28Z | TCGA-THCA | 27.214523 | 10.6716767 |
| TCGA-BJ-A290 | TCGA-THCA | 65.4500307 | 15.0444747 |
| TCGA-BJ-A291 | TCGA-THCA | 26.051604 | 10.6025159 |

| | | | |
|---|---|---|---|
| TCGA-BJ-A2N7 | TCGA-THCA | 74.3908119 | 22.6154078 |
| TCGA-BJ-A2N8 | TCGA-THCA | 67.7221121 | 21.0707971 |
| TCGA-BJ-A2N9 | TCGA-THCA | 62.0846914 | 14.9951393 |
| TCGA-BJ-A2NA | TCGA-THCA | 45.0664579 | 15.2950725 |
| TCGA-BJ-A2P4 | TCGA-THCA | 66.5815255 | 16.6363221 |
| TCGA-BJ-A3EZ | TCGA-THCA | 53.5935823 | 17.3053699 |
| TCGA-BJ-A3F0 | TCGA-THCA | 47.3054531 | 14.7169777 |
| TCGA-BJ-A3PR | TCGA-THCA | 51.6948898 | 13.4966438 |
| TCGA-BJ-A3PT | TCGA-THCA | 66.7718156 | 13.5114313 |
| TCGA-BJ-A3PU | TCGA-THCA | 81.6174836 | 17.692818 |
| TCGA-BJ-A45C | TCGA-THCA | 49.9022669 | 14.463801 |
| TCGA-BJ-A45D | TCGA-THCA | 44.3642796 | 13.0684789 |
| TCGA-BJ-A45E | TCGA-THCA | 20.774964 | 4.75839355 |
| TCGA-BJ-A45F | TCGA-THCA | 29.2131389 | 8.54566179 |
| TCGA-BJ-A45G | TCGA-THCA | 58.5429898 | 18.2926193 |
| TCGA-BJ-A45H | TCGA-THCA | 72.1537556 | 18.7515693 |
| TCGA-BJ-A45I | TCGA-THCA | 49.5785394 | 16.554892 |
| TCGA-BJ-A45J | TCGA-THCA | 66.4760195 | 22.3136019 |
| TCGA-BJ-A45K | TCGA-THCA | 54.5157275 | 17.5833706 |
| TCGA-BJ-A4O8 | TCGA-THCA | 48.2594968 | 14.8643205 |
| TCGA-BJ-A4O9 | TCGA-THCA | 56.5737414 | 12.8854168 |
| TCGA-BK-A0CA | TCGA-UCEC | 16.253141 | 9.53141427 |
| TCGA-BK-A0CC | TCGA-UCEC | 12.7054069 | 2.00523959 |
| TCGA-BK-A139 | TCGA-UCEC | 19.6858995 | 16.2063661 |
| TCGA-BK-A26L | TCGA-UCEC | 14.1952114 | 11.8229908 |
| TCGA-BK-A4ZD | TCGA-UCEC | 67.1013916 | 9.51414226 |
| TCGA-BK-A56F | TCGA-UCEC | 62.9213252 | 21.2376552 |
| TCGA-BK-A6W3 | TCGA-UCEC | 70.5324213 | 34.9057572 |
| TCGA-BK-A6W4 | TCGA-UCEC | 42.6284003 | 10.0322736 |
| TCGA-BL-A0C8 | TCGA-BLCA | 54.0497657 | 19.256456 |
| TCGA-BL-A13I | TCGA-BLCA | 12.7687682 | 11.5566366 |

| | | | |
|---|---|---|---|
| TCGA-BL-A13J | TCGA-BLCA | 8.37295516 | 6.95396656 |
| TCGA-BL-A3JM | TCGA-BLCA | 88.6508842 | 50.183248 |
| TCGA-BL-A5ZZ | TCGA-BLCA | 57.5256877 | 35.466932 |
| TCGA-BP-4158 | TCGA-KIRC | 39.8187008 | 14.1360946 |
| TCGA-BP-4159 | TCGA-KIRC | 36.5595226 | 12.4522929 |
| TCGA-BP-4160 | TCGA-KIRC | 39.3824418 | 11.9468665 |
| TCGA-BP-4161 | TCGA-KIRC | 32.6812788 | 18.3293692 |
| TCGA-BP-4162 | TCGA-KIRC | 39.7854482 | 17.5940101 |
| TCGA-BP-4163 | TCGA-KIRC | 32.8343199 | 25.9061915 |
| TCGA-BP-4164 | TCGA-KIRC | 47.3723614 | 18.8271326 |
| TCGA-BP-4165 | TCGA-KIRC | 63.2412154 | 17.5187012 |
| TCGA-BP-4166 | TCGA-KIRC | 32.3876527 | 12.4789563 |
| TCGA-BP-4167 | TCGA-KIRC | 30.7143458 | 14.6713318 |
| TCGA-BP-4169 | TCGA-KIRC | 46.2136162 | 21.0609175 |
| TCGA-BP-4170 | TCGA-KIRC | 37.505772 | 15.9678599 |
| TCGA-BP-4173 | TCGA-KIRC | 44.1168926 | 14.8814807 |
| TCGA-BP-4174 | TCGA-KIRC | 34.4261057 | 48.137645 |
| TCGA-BP-4176 | TCGA-KIRC | 32.7707403 | 24.0933118 |
| TCGA-BP-4177 | TCGA-KIRC | 50.575361 | 14.005231 |
| TCGA-BP-4325 | TCGA-KIRC | 41.8290474 | 14.9313183 |
| TCGA-BP-4326 | TCGA-KIRC | 31.4447874 | 15.7711401 |
| TCGA-BP-4327 | TCGA-KIRC | 38.619458 | 11.40184 |
| TCGA-BP-4329 | TCGA-KIRC | 31.4061957 | 15.2618019 |
| TCGA-BP-4330 | TCGA-KIRC | 33.693066 | 11.0979616 |
| TCGA-BP-4331 | TCGA-KIRC | 43.6783495 | 18.6462296 |
| TCGA-BP-4332 | TCGA-KIRC | 46.6685929 | 15.4197242 |
| TCGA-BP-4334 | TCGA-KIRC | 17.1345141 | 14.1527106 |
| TCGA-BP-4335 | TCGA-KIRC | 37.4953514 | 12.0827006 |
| TCGA-BP-4337 | TCGA-KIRC | 27.1127377 | 19.1247134 |
| TCGA-BP-4338 | TCGA-KIRC | 27.1377464 | 13.1318296 |
| TCGA-BP-4340 | TCGA-KIRC | 32.5924814 | 16.3211332 |

| | | | |
|---|---|---|---|
| TCGA-BP-4341 | TCGA-KIRC | 42.152543 | 11.8890833 |
| TCGA-BP-4342 | TCGA-KIRC | 23.7360543 | 9.23784217 |
| TCGA-BP-4343 | TCGA-KIRC | 27.8035834 | 7.74224673 |
| TCGA-BP-4344 | TCGA-KIRC | 53.8943614 | 22.7717719 |
| TCGA-BP-4345 | TCGA-KIRC | 45.0557981 | 21.5394866 |
| TCGA-BP-4346 | TCGA-KIRC | 43.5550549 | 15.2977492 |
| TCGA-BP-4347 | TCGA-KIRC | 39.7110774 | 19.8434688 |
| TCGA-BP-4349 | TCGA-KIRC | 54.0160615 | 16.3824432 |
| TCGA-BP-4351 | TCGA-KIRC | 38.9512451 | 20.5547565 |
| TCGA-BP-4352 | TCGA-KIRC | 37.766594 | 26.3819302 |
| TCGA-BP-4353 | TCGA-KIRC | 29.0269281 | 13.4424223 |
| TCGA-BP-4354 | TCGA-KIRC | 69.8316379 | 32.0876483 |
| TCGA-BP-4355 | TCGA-KIRC | 51.6493188 | 17.9820027 |
| TCGA-BP-4756 | TCGA-KIRC | 40.5251035 | 10.1949154 |
| TCGA-BP-4758 | TCGA-KIRC | 48.0291055 | 14.1805478 |
| TCGA-BP-4759 | TCGA-KIRC | 28.1337994 | 10.427977 |
| TCGA-BP-4760 | TCGA-KIRC | 40.0996328 | 15.7985029 |
| TCGA-BP-4761 | TCGA-KIRC | 34.5709892 | 25.0677788 |
| TCGA-BP-4762 | TCGA-KIRC | 32.3335895 | 16.3454479 |
| TCGA-BP-4763 | TCGA-KIRC | 34.2952717 | 14.0229769 |
| TCGA-BP-4765 | TCGA-KIRC | 54.2489153 | 22.0342582 |
| TCGA-BP-4766 | TCGA-KIRC | 44.6731329 | 18.4965786 |
| TCGA-BP-4768 | TCGA-KIRC | 30.8488181 | 15.158466 |
| TCGA-BP-4769 | TCGA-KIRC | 54.3605466 | 18.730999 |
| TCGA-BP-4770 | TCGA-KIRC | 63.3861622 | 36.2767416 |
| TCGA-BP-4771 | TCGA-KIRC | 37.7788764 | 15.5512829 |
| TCGA-BP-4774 | TCGA-KIRC | 58.2816109 | 15.4046761 |
| TCGA-BP-4775 | TCGA-KIRC | 31.2093029 | 8.85581081 |
| TCGA-BP-4776 | TCGA-KIRC | 38.7964021 | 10.0131839 |
| TCGA-BP-4777 | TCGA-KIRC | 39.2360683 | 15.216274 |
| TCGA-BP-4781 | TCGA-KIRC | 25.0572078 | 15.2133407 |

| | | | |
|---|---|---|---|
| TCGA-BP-4782 | TCGA-KIRC | 37.9357549 | 18.5142354 |
| TCGA-BP-4784 | TCGA-KIRC | 47.3903451 | 16.8762009 |
| TCGA-BP-4787 | TCGA-KIRC | 30.646006 | 26.384271 |
| TCGA-BP-4789 | TCGA-KIRC | 37.0779586 | 13.2367122 |
| TCGA-BP-4790 | TCGA-KIRC | 18.6334157 | 8.97988608 |
| TCGA-BP-4795 | TCGA-KIRC | 45.4110359 | 16.816781 |
| TCGA-BP-4797 | TCGA-KIRC | 22.0774404 | 14.6110319 |
| TCGA-BP-4798 | TCGA-KIRC | 31.370925 | 12.2994468 |
| TCGA-BP-4799 | TCGA-KIRC | 16.798996 | 7.77172311 |
| TCGA-BP-4801 | TCGA-KIRC | 41.7375629 | 13.5124846 |
| TCGA-BP-4803 | TCGA-KIRC | 22.1178178 | 8.88377433 |
| TCGA-BP-4804 | TCGA-KIRC | 33.1232728 | 11.8314931 |
| TCGA-BP-4807 | TCGA-KIRC | 28.4471523 | 7.65819652 |
| TCGA-BP-4959 | TCGA-KIRC | 28.3669681 | 11.4971871 |
| TCGA-BP-4960 | TCGA-KIRC | 24.4323298 | 9.54774244 |
| TCGA-BP-4961 | TCGA-KIRC | 44.1260372 | 12.8965266 |
| TCGA-BP-4962 | TCGA-KIRC | 28.134631 | 11.9260649 |
| TCGA-BP-4963 | TCGA-KIRC | 30.3669023 | 13.4637041 |
| TCGA-BP-4964 | TCGA-KIRC | 32.2188558 | 15.3170104 |
| TCGA-BP-4965 | TCGA-KIRC | 48.1941408 | 18.7753811 |
| TCGA-BP-4967 | TCGA-KIRC | 32.9590566 | 14.4782653 |
| TCGA-BP-4968 | TCGA-KIRC | 49.7121216 | 18.4865022 |
| TCGA-BP-4969 | TCGA-KIRC | 37.4430702 | 12.9268559 |
| TCGA-BP-4970 | TCGA-KIRC | 36.6165266 | 15.5327447 |
| TCGA-BP-4971 | TCGA-KIRC | 29.7208294 | 11.5766394 |
| TCGA-BP-4972 | TCGA-KIRC | 34.2355711 | 13.9902053 |
| TCGA-BP-4973 | TCGA-KIRC | 31.434405 | 10.9752069 |
| TCGA-BP-4974 | TCGA-KIRC | 46.7621635 | 15.3186495 |
| TCGA-BP-4975 | TCGA-KIRC | 28.9916474 | 10.9421772 |
| TCGA-BP-4976 | TCGA-KIRC | 38.738527 | 15.3945559 |
| TCGA-BP-4977 | TCGA-KIRC | 28.7175756 | 13.7228488 |

| | | | |
|---|---|---|---|
| TCGA-BP-4981 | TCGA-KIRC | 41.847304 | 14.1094229 |
| TCGA-BP-4982 | TCGA-KIRC | 39.4698622 | 12.7914154 |
| TCGA-BP-4983 | TCGA-KIRC | 54.3140045 | 27.0137512 |
| TCGA-BP-4985 | TCGA-KIRC | 33.3307658 | 33.0650173 |
| TCGA-BP-4986 | TCGA-KIRC | 30.7979266 | 24.4426799 |
| TCGA-BP-4987 | TCGA-KIRC | 39.0995023 | 10.0410755 |
| TCGA-BP-4989 | TCGA-KIRC | 40.4365945 | 18.8076713 |
| TCGA-BP-4991 | TCGA-KIRC | 42.0434276 | 15.9260451 |
| TCGA-BP-4992 | TCGA-KIRC | 20.6581182 | 9.73333115 |
| TCGA-BP-4993 | TCGA-KIRC | 53.4235775 | 12.7745625 |
| TCGA-BP-4994 | TCGA-KIRC | 28.1532752 | 21.7077745 |
| TCGA-BP-4995 | TCGA-KIRC | 31.1258953 | 19.6412903 |
| TCGA-BP-4998 | TCGA-KIRC | 35.7563082 | 13.3788731 |
| TCGA-BP-4999 | TCGA-KIRC | 42.1493212 | 12.9188476 |
| TCGA-BP-5000 | TCGA-KIRC | 36.6307057 | 15.3745756 |
| TCGA-BP-5001 | TCGA-KIRC | 48.4972186 | 14.3807327 |
| TCGA-BP-5004 | TCGA-KIRC | 27.3375462 | 20.1809865 |
| TCGA-BP-5006 | TCGA-KIRC | 45.774287 | 18.6146852 |
| TCGA-BP-5007 | TCGA-KIRC | 50.3563083 | 14.5905216 |
| TCGA-BP-5008 | TCGA-KIRC | 51.2605022 | 20.7301352 |
| TCGA-BP-5009 | TCGA-KIRC | 29.3756043 | 14.857839 |
| TCGA-BP-5010 | TCGA-KIRC | 35.87369 | 15.9091429 |
| TCGA-BP-5168 | TCGA-KIRC | 49.6390452 | 19.9496792 |
| TCGA-BP-5169 | TCGA-KIRC | 20.4031633 | 6.9479695 |
| TCGA-BP-5170 | TCGA-KIRC | 41.9204889 | 15.6269421 |
| TCGA-BP-5173 | TCGA-KIRC | 44.4825116 | 29.7592781 |
| TCGA-BP-5174 | TCGA-KIRC | 41.2178538 | 20.9667313 |
| TCGA-BP-5175 | TCGA-KIRC | 39.2143195 | 17.5515402 |
| TCGA-BP-5176 | TCGA-KIRC | 45.9627171 | 14.422989 |
| TCGA-BP-5177 | TCGA-KIRC | 55.3538916 | 17.4003286 |
| TCGA-BP-5178 | TCGA-KIRC | 33.635611 | 13.7417982 |

| | | | |
|---|---|---|---|
| TCGA-BP-5180 | TCGA-KIRC | 44.1476669 | 15.1295153 |
| TCGA-BP-5181 | TCGA-KIRC | 47.5370606 | 15.5685975 |
| TCGA-BP-5182 | TCGA-KIRC | 42.3164116 | 12.3839719 |
| TCGA-BP-5183 | TCGA-KIRC | 38.2194796 | 14.7442533 |
| TCGA-BP-5184 | TCGA-KIRC | 37.4816318 | 14.3821838 |
| TCGA-BP-5185 | TCGA-KIRC | 21.0040209 | 7.63779599 |
| TCGA-BP-5186 | TCGA-KIRC | 42.5006564 | 13.0702897 |
| TCGA-BP-5187 | TCGA-KIRC | 31.0381541 | 9.01788936 |
| TCGA-BP-5189 | TCGA-KIRC | 37.2519526 | 14.6385818 |
| TCGA-BP-5190 | TCGA-KIRC | 40.4251868 | 19.7014177 |
| TCGA-BP-5191 | TCGA-KIRC | 25.6429707 | 10.0942944 |
| TCGA-BP-5192 | TCGA-KIRC | 62.0540394 | 13.1435842 |
| TCGA-BP-5194 | TCGA-KIRC | 46.2652056 | 12.8666422 |
| TCGA-BP-5195 | TCGA-KIRC | 43.2468596 | 16.2584526 |
| TCGA-BP-5196 | TCGA-KIRC | 63.4103894 | 28.047521 |
| TCGA-BP-5198 | TCGA-KIRC | 38.0087946 | 19.5169509 |
| TCGA-BP-5199 | TCGA-KIRC | 62.4893962 | 22.1246057 |
| TCGA-BP-5200 | TCGA-KIRC | 47.6330435 | 16.4551268 |
| TCGA-BP-5201 | TCGA-KIRC | 53.0984854 | 17.4802518 |
| TCGA-BP-5202 | TCGA-KIRC | 45.8888557 | 11.6994441 |
| TCGA-BQ-5875 | TCGA-KIRP | 15.96239 | 12.3512822 |
| TCGA-BQ-5876 | TCGA-KIRP | 42.6434988 | 27.962218 |
| TCGA-BQ-5877 | TCGA-KIRP | 23.4782778 | 22.7095648 |
| TCGA-BQ-5878 | TCGA-KIRP | 18.388083 | 5.23623048 |
| TCGA-BQ-5879 | TCGA-KIRP | 35.810122 | 12.2631934 |
| TCGA-BQ-5880 | TCGA-KIRP | 18.5287144 | 7.02596488 |
| TCGA-BQ-5881 | TCGA-KIRP | 7.68625885 | 3.06225372 |
| TCGA-BQ-5882 | TCGA-KIRP | 27.702512 | 23.6087334 |
| TCGA-BQ-5883 | TCGA-KIRP | 11.5632825 | 2.28975163 |
| TCGA-BQ-5884 | TCGA-KIRP | 29.5527377 | 11.900294 |
| TCGA-BQ-5885 | TCGA-KIRP | 37.0967874 | 21.736169 |

| | | | |
|---|---|---|---|
| TCGA-BQ-5886 | TCGA-KIRP | 13.23285 | 5.78110278 |
| TCGA-BQ-5887 | TCGA-KIRP | 58.0778182 | 6.53828264 |
| TCGA-BQ-5888 | TCGA-KIRP | 30.530691 | 2.54822138 |
| TCGA-BQ-5889 | TCGA-KIRP | 35.3938047 | 17.3736064 |
| TCGA-BQ-5890 | TCGA-KIRP | 91.1104474 | 27.1260371 |
| TCGA-BQ-5891 | TCGA-KIRP | 41.5857751 | 19.5825573 |
| TCGA-BQ-5892 | TCGA-KIRP | 12.6714553 | 3.75477262 |
| TCGA-BQ-5893 | TCGA-KIRP | 8.29306744 | 1.37143647 |
| TCGA-BQ-5894 | TCGA-KIRP | 31.3775761 | 17.0988868 |
| TCGA-BQ-7044 | TCGA-KIRP | 15.0851208 | 15.4172239 |
| TCGA-BQ-7045 | TCGA-KIRP | 17.7268117 | 5.48560397 |
| TCGA-BQ-7046 | TCGA-KIRP | 18.1663312 | 2.8620182 |
| TCGA-BQ-7048 | TCGA-KIRP | 23.1409811 | 7.0017953 |
| TCGA-BQ-7049 | TCGA-KIRP | 18.4850624 | 9.17742147 |
| TCGA-BQ-7050 | TCGA-KIRP | 31.2794985 | 10.2740373 |
| TCGA-BQ-7051 | TCGA-KIRP | 32.5047599 | 12.3976593 |
| TCGA-BQ-7053 | TCGA-KIRP | 26.8907726 | 9.15553924 |
| TCGA-BQ-7055 | TCGA-KIRP | 17.9652231 | 12.319279 |
| TCGA-BQ-7056 | TCGA-KIRP | 23.9211788 | 7.87625056 |
| TCGA-BQ-7058 | TCGA-KIRP | 22.7211791 | 4.69309149 |
| TCGA-BQ-7059 | TCGA-KIRP | 24.8609821 | 10.9767957 |
| TCGA-BQ-7060 | TCGA-KIRP | 12.906754 | 0.73686268 |
| TCGA-BQ-7061 | TCGA-KIRP | 25.4227296 | 8.80311592 |
| TCGA-BQ-7062 | TCGA-KIRP | 39.5592587 | 17.1769674 |
| TCGA-BR-4187 | TCGA-STAD | 34.6109464 | 25.8299645 |
| TCGA-BR-4191 | TCGA-STAD | 49.579158 | 27.566939 |
| TCGA-BR-4201 | TCGA-STAD | 59.7221928 | 25.8876016 |
| TCGA-BR-4253 | TCGA-STAD | 86.0288689 | 44.4381071 |
| TCGA-BR-4256 | TCGA-STAD | 40.1125558 | 27.8529687 |
| TCGA-BR-4257 | TCGA-STAD | 34.7814706 | 33.7647677 |
| TCGA-BR-4267 | TCGA-STAD | 41.9907113 | 19.2559249 |

| | | | |
|---|---|---|---|
| TCGA-BR-4279 | TCGA-STAD | 30.9588382 | 24.4840278 |
| TCGA-BR-4280 | TCGA-STAD | 55.988698 | 24.631321 |
| TCGA-BR-4294 | TCGA-STAD | 16.0863926 | 24.7601548 |
| TCGA-BR-4357 | TCGA-STAD | 38.4465098 | 18.5508707 |
| TCGA-BR-4361 | TCGA-STAD | 69.1789702 | 36.9270152 |
| TCGA-BR-4363 | TCGA-STAD | 61.8050066 | 22.8651494 |
| TCGA-BR-4366 | TCGA-STAD | 74.0302802 | 19.3939437 |
| TCGA-BR-4367 | TCGA-STAD | 51.8261375 | 31.8840917 |
| TCGA-BR-4368 | TCGA-STAD | 51.8355409 | 44.5359411 |
| TCGA-BR-4369 | TCGA-STAD | 69.0871467 | 27.6833076 |
| TCGA-BR-4370 | TCGA-STAD | 27.0707571 | 26.8440422 |
| TCGA-BR-4371 | TCGA-STAD | 58.2019565 | 29.313032 |
| TCGA-BR-6452 | TCGA-STAD | 38.5531507 | 23.3837438 |
| TCGA-BR-6453 | TCGA-STAD | 28.4560835 | 10.8502669 |
| TCGA-BR-6454 | TCGA-STAD | 14.0620082 | 9.15131361 |
| TCGA-BR-6455 | TCGA-STAD | 49.8526555 | 33.3321935 |
| TCGA-BR-6456 | TCGA-STAD | 36.6682112 | 26.572161 |
| TCGA-BR-6457 | TCGA-STAD | 12.3453279 | 6.13781648 |
| TCGA-BR-6458 | TCGA-STAD | 39.8633874 | 22.6153928 |
| TCGA-BR-6563 | TCGA-STAD | 28.7224164 | 15.9300066 |
| TCGA-BR-6564 | TCGA-STAD | 28.7214223 | 17.2544708 |
| TCGA-BR-6565 | TCGA-STAD | 46.0331835 | 28.7207429 |
| TCGA-BR-6566 | TCGA-STAD | 39.951523 | 27.7597495 |
| TCGA-BR-6705 | TCGA-STAD | 22.9403152 | 17.3025519 |
| TCGA-BR-6707 | TCGA-STAD | 85.6566312 | 39.3373054 |
| TCGA-BR-6709 | TCGA-STAD | 38.5244249 | 10.7114908 |
| TCGA-BR-6710 | TCGA-STAD | 10.6739502 | 4.99043909 |
| TCGA-BR-6801 | TCGA-STAD | 26.4530798 | 26.457324 |
| TCGA-BR-6802 | TCGA-STAD | 43.0426984 | 27.9697796 |
| TCGA-BR-6803 | TCGA-STAD | 27.1242867 | 23.9888082 |
| TCGA-BR-6852 | TCGA-STAD | 40.7802069 | 17.3507408 |

| | | | |
|---|---|---|---|
| TCGA-BR-7196 | TCGA-STAD | 38.1901439 | 21.0799229 |
| TCGA-BR-7197 | TCGA-STAD | 44.3672699 | 22.3428144 |
| TCGA-BR-7703 | TCGA-STAD | 15.9873371 | 10.0975837 |
| TCGA-BR-7704 | TCGA-STAD | 69.1586036 | 38.4849513 |
| TCGA-BR-7707 | TCGA-STAD | 64.4815928 | 52.4217123 |
| TCGA-BR-7715 | TCGA-STAD | 28.2452009 | 15.3128075 |
| TCGA-BR-7716 | TCGA-STAD | 28.7221147 | 12.5428823 |
| TCGA-BR-7717 | TCGA-STAD | 31.0771736 | 18.3064597 |
| TCGA-BR-7722 | TCGA-STAD | 70.4522029 | 13.2241266 |
| TCGA-BR-7723 | TCGA-STAD | 59.2057973 | 37.6077042 |
| TCGA-BR-7851 | TCGA-STAD | 59.6166989 | 20.129818 |
| TCGA-BR-7901 | TCGA-STAD | 48.5581011 | 5.19665222 |
| TCGA-BR-7957 | TCGA-STAD | 34.6921376 | 21.5084258 |
| TCGA-BR-7958 | TCGA-STAD | 94.692858 | 36.7522215 |
| TCGA-BR-7959 | TCGA-STAD | 40.9338444 | 24.9779156 |
| TCGA-BR-8058 | TCGA-STAD | 67.9789936 | 35.0945487 |
| TCGA-BR-8059 | TCGA-STAD | 33.5869731 | 30.9520057 |
| TCGA-BR-8060 | TCGA-STAD | 17.1297464 | 6.46141671 |
| TCGA-BR-8077 | TCGA-STAD | 51.7721382 | 32.2646718 |
| TCGA-BR-8080 | TCGA-STAD | 57.8200195 | 22.3264241 |
| TCGA-BR-8081 | TCGA-STAD | 56.0166877 | 21.1273906 |
| TCGA-BR-8284 | TCGA-STAD | 58.9149229 | 33.3279543 |
| TCGA-BR-8286 | TCGA-STAD | 79.9019072 | 36.300984 |
| TCGA-BR-8289 | TCGA-STAD | 128.91027 | 19.8642425 |
| TCGA-BR-8291 | TCGA-STAD | 64.4591758 | 30.7194808 |
| TCGA-BR-8295 | TCGA-STAD | 66.9592922 | 50.9849857 |
| TCGA-BR-8296 | TCGA-STAD | 34.7352666 | 15.2968612 |
| TCGA-BR-8297 | TCGA-STAD | 47.8030524 | 32.918542 |
| TCGA-BR-8361 | TCGA-STAD | 54.5020218 | 27.290928 |
| TCGA-BR-8364 | TCGA-STAD | 42.0923723 | 24.2822046 |
| TCGA-BR-8365 | TCGA-STAD | 36.6743854 | 19.7743956 |

| | | | |
|---|---|---|---|
| TCGA-BR-8366 | TCGA-STAD | 42.88875 | 25.9651813 |
| TCGA-BR-8367 | TCGA-STAD | 28.2945584 | 17.4370968 |
| TCGA-BR-8368 | TCGA-STAD | 43.9080847 | 23.1499865 |
| TCGA-BR-8369 | TCGA-STAD | 24.3021397 | 19.6227547 |
| TCGA-BR-8371 | TCGA-STAD | 26.1172319 | 14.8274802 |
| TCGA-BR-8372 | TCGA-STAD | 48.2309945 | 46.5625721 |
| TCGA-BR-8373 | TCGA-STAD | 45.502255 | 14.2132064 |
| TCGA-BR-8380 | TCGA-STAD | 30.7149455 | 18.7769737 |
| TCGA-BR-8381 | TCGA-STAD | 53.3952932 | 29.090383 |
| TCGA-BR-8382 | TCGA-STAD | 62.8121364 | 22.8573193 |
| TCGA-BR-8384 | TCGA-STAD | 36.1375249 | 18.060832 |
| TCGA-BR-8483 | TCGA-STAD | 44.1062474 | 26.8404002 |
| TCGA-BR-8484 | TCGA-STAD | 45.6193346 | 36.0152854 |
| TCGA-BR-8485 | TCGA-STAD | 55.6638497 | 42.3836889 |
| TCGA-BR-8486 | TCGA-STAD | 39.003491 | 31.5635547 |
| TCGA-BR-8487 | TCGA-STAD | 47.9013119 | 25.3180532 |
| TCGA-BR-8588 | TCGA-STAD | 40.818599 | 29.1355544 |
| TCGA-BR-8589 | TCGA-STAD | 109.719897 | 26.4954021 |
| TCGA-BR-8590 | TCGA-STAD | 28.9503535 | 18.7535015 |
| TCGA-BR-8591 | TCGA-STAD | 43.9103422 | 20.9858334 |
| TCGA-BR-8592 | TCGA-STAD | 28.8632945 | 14.7784429 |
| TCGA-BR-8676 | TCGA-STAD | 43.5616745 | 68.7871528 |
| TCGA-BR-8677 | TCGA-STAD | 67.2368789 | 28.7386302 |
| TCGA-BR-8678 | TCGA-STAD | 78.2806464 | 47.6011114 |
| TCGA-BR-8679 | TCGA-STAD | 31.9409554 | 42.4406583 |
| TCGA-BR-8680 | TCGA-STAD | 72.9525736 | 33.9184401 |
| TCGA-BR-8682 | TCGA-STAD | 30.0070477 | 9.65884187 |
| TCGA-BR-8683 | TCGA-STAD | 45.3792774 | 21.0134416 |
| TCGA-BR-8687 | TCGA-STAD | 26.632721 | 21.6753952 |
| TCGA-BR-8690 | TCGA-STAD | 35.8020927 | 30.9979385 |
| TCGA-BR-A44T | TCGA-STAD | 34.2283775 | 17.2348143 |

| | | | |
|---|---|---|---|
| TCGA-BR-A44U | TCGA-STAD | 38.3944384 | 25.7853929 |
| TCGA-BR-A4CR | TCGA-STAD | 49.694253 | 55.7671911 |
| TCGA-BR-A4CS | TCGA-STAD | 45.3344416 | 29.0157341 |
| TCGA-BR-A4IV | TCGA-STAD | 28.3657168 | 17.7471143 |
| TCGA-BR-A4J4 | TCGA-STAD | 64.6819632 | 21.0155349 |
| TCGA-BR-A4J5 | TCGA-STAD | 33.6609654 | 27.4305737 |
| TCGA-BR-A4J6 | TCGA-STAD | 44.1016102 | 64.5611852 |
| TCGA-BR-A4J7 | TCGA-STAD | 27.0121373 | 17.0867859 |
| TCGA-BR-A4J8 | TCGA-STAD | 56.4782658 | 19.9868708 |
| TCGA-BR-A4J9 | TCGA-STAD | 20.7692596 | 22.29479 |
| TCGA-BR-A4PF | TCGA-STAD | 56.0262855 | 33.3414412 |
| TCGA-BR-A4QL | TCGA-STAD | 52.0751924 | 23.0027794 |
| TCGA-BS-A0V4 | TCGA-UCEC | 5.58160186 | 4.64061671 |
| TCGA-BS-A0V7 | TCGA-UCEC | 6.58762287 | 7.32935641 |
| TCGA-BT-A0S7 | TCGA-BLCA | 50.2696452 | 25.9125169 |
| TCGA-BT-A0YX | TCGA-BLCA | 44.319021 | 16.0917671 |
| TCGA-BT-A20J | TCGA-BLCA | 54.4990799 | 42.2660688 |
| TCGA-BT-A20N | TCGA-BLCA | 90.5128887 | 55.3234664 |
| TCGA-BT-A20O | TCGA-BLCA | 58.4353159 | 33.6451709 |
| TCGA-BT-A20P | TCGA-BLCA | 99.128416 | 19.6114702 |
| TCGA-BT-A20Q | TCGA-BLCA | 65.954224 | 17.0615352 |
| TCGA-BT-A20R | TCGA-BLCA | 42.732615 | 18.6882762 |
| TCGA-BT-A20T | TCGA-BLCA | 78.1715691 | 25.9585186 |
| TCGA-BT-A20U | TCGA-BLCA | 65.8036225 | 27.0102545 |
| TCGA-BT-A20V | TCGA-BLCA | 112.899805 | 72.1547903 |
| TCGA-BT-A20W | TCGA-BLCA | 68.7280359 | 26.0804379 |
| TCGA-BT-A20X | TCGA-BLCA | 71.0164009 | 33.2908303 |
| TCGA-BT-A2LA | TCGA-BLCA | 133.125256 | 1.0265416 |
| TCGA-BT-A2LB | TCGA-BLCA | 39.4416766 | 14.9461552 |
| TCGA-BT-A2LD | TCGA-BLCA | 69.9684408 | 44.5216646 |
| TCGA-BT-A3PH | TCGA-BLCA | 72.3850369 | 17.3134265 |

| | | | |
|---|---|---|---|
| TCGA-BT-A3PJ | TCGA-BLCA | 103.669398 | 27.6281149 |
| TCGA-BT-A3PK | TCGA-BLCA | 91.0990412 | 42.1564241 |
| TCGA-BT-A42C | TCGA-BLCA | 113.821161 | 26.2218074 |
| TCGA-BT-A42E | TCGA-BLCA | 146.216847 | 42.3881472 |
| TCGA-BT-A42F | TCGA-BLCA | 62.4742063 | 22.7845706 |
| TCGA-BW-A5NC | TCGA-LIHC | 91.7291096 | 25.5220558 |
| TCGA-BW-A5NF | TCGA-LIHC | 46.2152577 | 15.986347 |
| TCGA-BW-A5NC | TCGA-LIHC | 83.3853125 | 51.7738078 |
| TCGA-C4-A0EZ | TCGA-BLCA | 109.761718 | 48.0645681 |
| TCGA-C4-A0F0 | TCGA-BLCA | 120.989718 | 42.4538047 |
| TCGA-C4-A0F1 | TCGA-BLCA | 34.5454747 | 14.4186766 |
| TCGA-C4-A0F6 | TCGA-BLCA | 59.5166577 | 18.5488375 |
| TCGA-C4-A0F7 | TCGA-BLCA | 77.6226052 | 8.70391165 |
| TCGA-C8-A12K | TCGA-BRCA | 99.8867456 | 23.2394705 |
| TCGA-C8-A12L | TCGA-BRCA | 56.511384 | 33.7582471 |
| TCGA-C8-A12M | TCGA-BRCA | 36.4434556 | 23.2032893 |
| TCGA-C8-A12N | TCGA-BRCA | 74.9689283 | 26.742175 |
| TCGA-C8-A12O | TCGA-BRCA | 47.3735022 | 46.5222424 |
| TCGA-C8-A12P | TCGA-BRCA | 64.9706129 | 31.7086369 |
| TCGA-C8-A12Q | TCGA-BRCA | 69.3518229 | 36.726938 |
| TCGA-C8-A12T | TCGA-BRCA | 47.6583155 | 29.5968422 |
| TCGA-C8-A12U | TCGA-BRCA | 56.6824117 | 38.132187 |
| TCGA-C8-A12V | TCGA-BRCA | 71.8646157 | 17.8942869 |
| TCGA-C8-A12W | TCGA-BRCA | 29.1377257 | 47.460902 |
| TCGA-C8-A12X | TCGA-BRCA | 19.4148419 | 17.0516761 |
| TCGA-C8-A12Y | TCGA-BRCA | 89.8815772 | 45.9342046 |
| TCGA-C8-A12Z | TCGA-BRCA | 64.3222002 | 29.4631572 |
| TCGA-C8-A130 | TCGA-BRCA | 85.4878229 | 35.681437 |
| TCGA-C8-A131 | TCGA-BRCA | 62.9755788 | 31.0758836 |
| TCGA-C8-A132 | TCGA-BRCA | 44.3325247 | 27.1798134 |
| TCGA-C8-A133 | TCGA-BRCA | 28.67158 | 26.5468293 |

| | | | |
|---|---|---|---|
| TCGA-C8-A134 | TCGA-BRCA | 237.496484 | 40.4999961 |
| TCGA-C8-A135 | TCGA-BRCA | 41.1082912 | 25.3119025 |
| TCGA-C8-A137 | TCGA-BRCA | 56.7387773 | 29.9501966 |
| TCGA-C8-A138 | TCGA-BRCA | 48.4811365 | 30.4672551 |
| TCGA-C8-A1HE | TCGA-BRCA | 57.5322661 | 27.3076658 |
| TCGA-C8-A1HF | TCGA-BRCA | 101.975886 | 48.9747834 |
| TCGA-C8-A1HG | TCGA-BRCA | 68.1078982 | 43.3994278 |
| TCGA-C8-A1HI | TCGA-BRCA | 50.9761116 | 33.3881317 |
| TCGA-C8-A1HJ | TCGA-BRCA | 78.8199553 | 123.342516 |
| TCGA-C8-A1HK | TCGA-BRCA | 53.7463905 | 52.1054773 |
| TCGA-C8-A1HL | TCGA-BRCA | 64.8981059 | 60.0192619 |
| TCGA-C8-A1HM | TCGA-BRCA | 60.0846016 | 81.5734403 |
| TCGA-C8-A1HN | TCGA-BRCA | 70.2112904 | 85.6418212 |
| TCGA-C8-A1HO | TCGA-BRCA | 46.5463204 | 34.7775353 |
| TCGA-C8-A26V | TCGA-BRCA | 70.5988876 | 33.6836668 |
| TCGA-C8-A26W | TCGA-BRCA | 103.578488 | 41.6918621 |
| TCGA-C8-A26X | TCGA-BRCA | 32.225889 | 14.0857096 |
| TCGA-C8-A26Y | TCGA-BRCA | 87.1595889 | 26.6250517 |
| TCGA-C8-A26Z | TCGA-BRCA | 42.6874195 | 63.8907258 |
| TCGA-C8-A273 | TCGA-BRCA | 61.396144 | 47.9987702 |
| TCGA-C8-A274 | TCGA-BRCA | 42.1792758 | 29.0641912 |
| TCGA-C8-A275 | TCGA-BRCA | 42.8335955 | 31.0906928 |
| TCGA-C8-A278 | TCGA-BRCA | 133.962348 | 22.4189412 |
| TCGA-C8-A27A | TCGA-BRCA | 45.3103675 | 32.3109489 |
| TCGA-C8-A27B | TCGA-BRCA | 228.162137 | 22.8285897 |
| TCGA-C8-A3M7 | TCGA-BRCA | 59.6705861 | 49.1994995 |
| TCGA-C8-A3M8 | TCGA-BRCA | 41.3237863 | 35.9809606 |
| TCGA-C8-A8HP | TCGA-BRCA | 45.3839671 | 29.8069657 |
| TCGA-C8-A8HQ | TCGA-BRCA | 76.3059866 | 53.3334354 |
| TCGA-C8-A8HR | TCGA-BRCA | 72.3352252 | 28.7311238 |
| TCGA-CA-5254 | TCGA-COAD | 71.5152464 | 45.7327433 |

| | | | |
|---|---|---|---|
| TCGA-CA-5255 | TCGA-COAD | 35.4962753 | 26.2824481 |
| TCGA-CA-5796 | TCGA-COAD | 53.058234 | 15.0123656 |
| TCGA-CA-5797 | TCGA-COAD | 94.7257821 | 32.7304018 |
| TCGA-CA-6715 | TCGA-COAD | 103.919882 | 14.5459704 |
| TCGA-CA-6716 | TCGA-COAD | 54.6396691 | 20.8412846 |
| TCGA-CA-6717 | TCGA-COAD | 71.2866488 | 21.0873628 |
| TCGA-CA-6718 | TCGA-COAD | 99.5268568 | 30.4125776 |
| TCGA-CA-6719 | TCGA-COAD | 123.236155 | 23.7632417 |
| TCGA-CC-5258 | TCGA-LIHC | 35.5263728 | 22.4095993 |
| TCGA-CC-5259 | TCGA-LIHC | 26.0059654 | 17.860084 |
| TCGA-CC-5260 | TCGA-LIHC | 53.3178085 | 24.0254428 |
| TCGA-CC-5261 | TCGA-LIHC | 42.2944905 | 22.0568608 |
| TCGA-CC-5262 | TCGA-LIHC | 43.5220962 | 17.1907792 |
| TCGA-CC-5263 | TCGA-LIHC | 51.141786 | 19.7533955 |
| TCGA-CC-5264 | TCGA-LIHC | 88.3800003 | 43.7960584 |
| TCGA-CC-A123 | TCGA-LIHC | 29.5286559 | 9.42680756 |
| TCGA-CC-A1HT | TCGA-LIHC | 94.7640778 | 34.4481822 |
| TCGA-CC-A3M9 | TCGA-LIHC | 126.054868 | 28.2295738 |
| TCGA-CC-A3MA | TCGA-LIHC | 84.3412408 | 29.0524121 |
| TCGA-CC-A3MB | TCGA-LIHC | 51.3032539 | 28.0429508 |
| TCGA-CC-A3MC | TCGA-LIHC | 29.2155384 | 20.5040358 |
| TCGA-CC-A5UC | TCGA-LIHC | 41.3705637 | 22.0062599 |
| TCGA-CC-A5UD | TCGA-LIHC | 74.2879607 | 45.0407808 |
| TCGA-CC-A5UE | TCGA-LIHC | 95.6529191 | 15.2656953 |
| TCGA-CC-A7IE | TCGA-LIHC | 37.7385537 | 17.7207852 |
| TCGA-CC-A7IF | TCGA-LIHC | 31.9804005 | 15.3753039 |
| TCGA-CC-A7IG | TCGA-LIHC | 48.1147986 | 17.9721322 |
| TCGA-CC-A7IH | TCGA-LIHC | 58.6531701 | 21.4906127 |
| TCGA-CC-A7II | TCGA-LIHC | 85.676501 | 24.946346 |
| TCGA-CC-A7IJ | TCGA-LIHC | 92.657117 | 57.7079628 |
| TCGA-CC-A7IK | TCGA-LIHC | 75.5903025 | 7.79844609 |

| | | | |
|---|---|---|---|
| TCGA-CC-A7IL | TCGA-LIHC | 22.9030166 | 15.0325122 |
| TCGA-CC-A8HS | TCGA-LIHC | 61.0775791 | 27.986017 |
| TCGA-CC-A8HT | TCGA-LIHC | 66.8864436 | 26.5764535 |
| TCGA-CC-A8HU | TCGA-LIHC | 92.8936507 | 36.668467 |
| TCGA-CC-A8HV | TCGA-LIHC | 42.9575657 | 12.3151344 |
| TCGA-CC-A9FS | TCGA-LIHC | 30.0746393 | 10.6986331 |
| TCGA-CC-A9FU | TCGA-LIHC | 106.613866 | 25.5581593 |
| TCGA-CC-A9FV | TCGA-LIHC | 38.1750718 | 10.9978157 |
| TCGA-CC-A9FW | TCGA-LIHC | 36.6728549 | 19.5296483 |
| TCGA-CD-5798 | TCGA-STAD | 37.1832439 | 29.9319621 |
| TCGA-CD-5799 | TCGA-STAD | 76.3218645 | 39.8278792 |
| TCGA-CD-5800 | TCGA-STAD | 75.7800219 | 30.9014964 |
| TCGA-CD-5801 | TCGA-STAD | 57.2825712 | 24.924772 |
| TCGA-CD-5803 | TCGA-STAD | 46.7092137 | 18.5989915 |
| TCGA-CD-5804 | TCGA-STAD | 42.7078315 | 34.4728554 |
| TCGA-CD-5813 | TCGA-STAD | 28.6537863 | 15.2534462 |
| TCGA-CD-8524 | TCGA-STAD | 90.5364747 | 26.3512369 |
| TCGA-CD-8525 | TCGA-STAD | 78.5874682 | 45.7766461 |
| TCGA-CD-8526 | TCGA-STAD | 81.0420409 | 76.1735598 |
| TCGA-CD-8527 | TCGA-STAD | 83.347849 | 40.4695669 |
| TCGA-CD-8528 | TCGA-STAD | 62.8634715 | 28.3357074 |
| TCGA-CD-8529 | TCGA-STAD | 46.9020975 | 21.5441754 |
| TCGA-CD-8530 | TCGA-STAD | 48.9568421 | 32.1834201 |
| TCGA-CD-8531 | TCGA-STAD | 64.0502994 | 43.9099466 |
| TCGA-CD-8532 | TCGA-STAD | 121.289942 | 20.8799229 |
| TCGA-CD-8533 | TCGA-STAD | 62.2109106 | 47.7149318 |
| TCGA-CD-8534 | TCGA-STAD | 28.5894346 | 30.595853 |
| TCGA-CD-8535 | TCGA-STAD | 56.2904845 | 17.2415336 |
| TCGA-CD-A486 | TCGA-STAD | 42.7378645 | 19.9485644 |
| TCGA-CD-A487 | TCGA-STAD | 49.8561349 | 47.080013 |
| TCGA-CD-A489 | TCGA-STAD | 32.6293562 | 24.4492769 |

| | | | |
|---|---|---|---|
| TCGA-CD-A48A | TCGA-STAD | 48.0544504 | 12.1090009 |
| TCGA-CD-A48C | TCGA-STAD | 69.0712842 | 37.1839239 |
| TCGA-CD-A4MG | TCGA-STAD | 44.3927097 | 24.4303929 |
| TCGA-CD-A4MH | TCGA-STAD | 38.8577535 | 21.3561401 |
| TCGA-CE-A13K | TCGA-THCA | 54.9415409 | 13.2370979 |
| TCGA-CE-A27D | TCGA-THCA | 71.4469555 | 18.0913466 |
| TCGA-CE-A3MD | TCGA-THCA | 74.3174681 | 16.3853945 |
| TCGA-CE-A3ME | TCGA-THCA | 57.9384819 | 14.9645219 |
| TCGA-CE-A481 | TCGA-THCA | 56.3050146 | 12.1677714 |
| TCGA-CE-A482 | TCGA-THCA | 64.9455859 | 15.8339771 |
| TCGA-CE-A483 | TCGA-THCA | 58.9284106 | 13.4984266 |
| TCGA-CE-A484 | TCGA-THCA | 71.4380516 | 21.5098324 |
| TCGA-CE-A485 | TCGA-THCA | 57.529345 | 18.4564753 |
| TCGA-CF-A1HR | TCGA-BLCA | 54.6129269 | 21.2051673 |
| TCGA-CF-A1HS | TCGA-BLCA | 155.053794 | 29.9970227 |
| TCGA-CF-A27C | TCGA-BLCA | 69.646391 | 14.2284484 |
| TCGA-CF-A3MF | TCGA-BLCA | 12.4604083 | 5.86293179 |
| TCGA-CF-A3MG | TCGA-BLCA | 71.6442335 | 22.3062821 |
| TCGA-CF-A3MH | TCGA-BLCA | 53.3566745 | 18.8931497 |
| TCGA-CF-A3MI | TCGA-BLCA | 36.1059202 | 9.01893973 |
| TCGA-CF-A47S | TCGA-BLCA | 59.2446795 | 15.1555101 |
| TCGA-CF-A47T | TCGA-BLCA | 79.7618632 | 19.0053222 |
| TCGA-CF-A47V | TCGA-BLCA | 51.217856 | 14.8723935 |
| TCGA-CF-A47W | TCGA-BLCA | 59.8673602 | 11.3718011 |
| TCGA-CF-A47X | TCGA-BLCA | 43.3903198 | 10.3393824 |
| TCGA-CF-A47Y | TCGA-BLCA | 34.3466315 | 8.02165624 |
| TCGA-CF-A5U8 | TCGA-BLCA | 63.4724211 | 14.4140539 |
| TCGA-CF-A5UA | TCGA-BLCA | 98.8642834 | 35.790317 |
| TCGA-CF-A7I0 | TCGA-BLCA | 64.6889469 | 7.08056461 |
| TCGA-CF-A8HX | TCGA-BLCA | 51.6388318 | 24.077499 |
| TCGA-CF-A8HY | TCGA-BLCA | 50.0691201 | 16.7643244 |

| | | | |
|---|---|---|---|
| TCGA-CF-A9FF | TCGA-BLCA | 53.1182718 | 14.7902704 |
| TCGA-CF-A9FH | TCGA-BLCA | 93.0645575 | 21.505389 |
| TCGA-CF-A9FL | TCGA-BLCA | 57.5272312 | 12.1823093 |
| TCGA-CF-A9FM | TCGA-BLCA | 60.8253994 | 13.9976724 |
| TCGA-CG-4301 | TCGA-STAD | 26.8189507 | 21.6375702 |
| TCGA-CG-4304 | TCGA-STAD | 31.9301786 | 13.0980178 |
| TCGA-CG-4305 | TCGA-STAD | 35.8870586 | 21.765407 |
| TCGA-CG-4306 | TCGA-STAD | 64.6166069 | 41.8086445 |
| TCGA-CG-4436 | TCGA-STAD | 53.3503497 | 25.8154805 |
| TCGA-CG-4437 | TCGA-STAD | 37.2120628 | 26.7196521 |
| TCGA-CG-4438 | TCGA-STAD | 42.6180558 | 47.0342298 |
| TCGA-CG-4440 | TCGA-STAD | 81.7669731 | 56.3609401 |
| TCGA-CG-4441 | TCGA-STAD | 41.8102902 | 30.3650518 |
| TCGA-CG-4442 | TCGA-STAD | 47.1784984 | 25.735071 |
| TCGA-CG-4443 | TCGA-STAD | 57.4503263 | 15.5765282 |
| TCGA-CG-4444 | TCGA-STAD | 68.2182553 | 40.5991683 |
| TCGA-CG-4460 | TCGA-STAD | 43.975008 | 33.3219048 |
| TCGA-CG-4462 | TCGA-STAD | 36.6996031 | 22.4635965 |
| TCGA-CG-4465 | TCGA-STAD | 62.7057444 | 45.0484573 |
| TCGA-CG-4466 | TCGA-STAD | 60.9688226 | 36.1547318 |
| TCGA-CG-4469 | TCGA-STAD | 59.4627256 | 27.6513864 |
| TCGA-CG-4475 | TCGA-STAD | 58.4716759 | 41.157034 |
| TCGA-CG-4476 | TCGA-STAD | 61.2009955 | 22.5562975 |
| TCGA-CG-4477 | TCGA-STAD | 65.0116081 | 26.9833498 |
| TCGA-CG-5716 | TCGA-STAD | 19.9835917 | 17.2276267 |
| TCGA-CG-5717 | TCGA-STAD | 27.1959288 | 14.2473466 |
| TCGA-CG-5718 | TCGA-STAD | 26.1237264 | 15.5302641 |
| TCGA-CG-5719 | TCGA-STAD | 30.2352404 | 24.953986 |
| TCGA-CG-5720 | TCGA-STAD | 41.2393047 | 16.4820485 |
| TCGA-CG-5721 | TCGA-STAD | 16.9919854 | 10.4998669 |
| TCGA-CG-5722 | TCGA-STAD | 51.6379301 | 47.5432691 |

| | | | |
|---|---|---|---|
| TCGA-CG-5723 | TCGA-STAD | 67.8449294 | 58.7687992 |
| TCGA-CG-5724 | TCGA-STAD | 37.9262791 | 36.5130295 |
| TCGA-CG-5725 | TCGA-STAD | 65.7741824 | 18.9621692 |
| TCGA-CG-5726 | TCGA-STAD | 39.9873292 | 17.5989084 |
| TCGA-CG-5728 | TCGA-STAD | 12.6502362 | 8.58107771 |
| TCGA-CG-5730 | TCGA-STAD | 20.2067533 | 8.82978128 |
| TCGA-CG-5732 | TCGA-STAD | 31.0751598 | 9.00380594 |
| TCGA-CG-5733 | TCGA-STAD | 14.4895885 | 8.05785313 |
| TCGA-CG-5734 | TCGA-STAD | 18.765995 | 11.0445784 |
| TCGA-CH-5737 | TCGA-PRAD | 26.8041613 | 17.2843714 |
| TCGA-CH-5738 | TCGA-PRAD | 40.5087984 | 18.7346546 |
| TCGA-CH-5739 | TCGA-PRAD | 34.2240577 | 12.9741467 |
| TCGA-CH-5740 | TCGA-PRAD | 38.0160318 | 15.1463304 |
| TCGA-CH-5741 | TCGA-PRAD | 44.2176366 | 27.2302964 |
| TCGA-CH-5743 | TCGA-PRAD | 50.5376961 | 17.0149447 |
| TCGA-CH-5744 | TCGA-PRAD | 32.0891561 | 8.8940918 |
| TCGA-CH-5745 | TCGA-PRAD | 33.5773123 | 11.8076071 |
| TCGA-CH-5746 | TCGA-PRAD | 37.7719228 | 16.6984318 |
| TCGA-CH-5748 | TCGA-PRAD | 31.8172237 | 17.5517996 |
| TCGA-CH-5750 | TCGA-PRAD | 37.5513992 | 16.6527089 |
| TCGA-CH-5751 | TCGA-PRAD | 37.4105711 | 12.7946539 |
| TCGA-CH-5752 | TCGA-PRAD | 35.6496319 | 7.57834964 |
| TCGA-CH-5753 | TCGA-PRAD | 23.1208504 | 23.8798613 |
| TCGA-CH-5754 | TCGA-PRAD | 40.1190131 | 6.64696951 |
| TCGA-CH-5761 | TCGA-PRAD | 52.6049555 | 11.4373139 |
| TCGA-CH-5762 | TCGA-PRAD | 40.1054227 | 22.5264428 |
| TCGA-CH-5763 | TCGA-PRAD | 41.3589159 | 14.1767408 |
| TCGA-CH-5764 | TCGA-PRAD | 38.7542547 | 19.1740826 |
| TCGA-CH-5765 | TCGA-PRAD | 35.9489613 | 14.6417692 |
| TCGA-CH-5766 | TCGA-PRAD | 47.0916607 | 9.31357156 |
| TCGA-CH-5767 | TCGA-PRAD | 36.4248693 | 12.3527967 |

| | | | |
|---|---|---|---|
| TCGA-CH-5768 | TCGA-PRAD | 32.1330344 | 14.0818601 |
| TCGA-CH-5769 | TCGA-PRAD | 22.8859861 | 21.7323141 |
| TCGA-CH-5771 | TCGA-PRAD | 30.4104355 | 13.7720196 |
| TCGA-CH-5772 | TCGA-PRAD | 40.0483567 | 16.0231749 |
| TCGA-CH-5788 | TCGA-PRAD | 34.6388189 | 16.8098033 |
| TCGA-CH-5789 | TCGA-PRAD | 45.8343227 | 18.2552977 |
| TCGA-CH-5790 | TCGA-PRAD | 40.217705 | 19.8223612 |
| TCGA-CH-5791 | TCGA-PRAD | 47.0228367 | 18.0755246 |
| TCGA-CH-5792 | TCGA-PRAD | 34.6716779 | 10.4994377 |
| TCGA-CH-5794 | TCGA-PRAD | 36.5526031 | 16.9417955 |
| TCGA-CJ-4634 | TCGA-KIRC | 51.2811916 | 19.1931277 |
| TCGA-CJ-4635 | TCGA-KIRC | 31.3679552 | 11.4774606 |
| TCGA-CJ-4636 | TCGA-KIRC | 30.9550984 | 19.3939612 |
| TCGA-CJ-4637 | TCGA-KIRC | 41.8647855 | 23.6217578 |
| TCGA-CJ-4638 | TCGA-KIRC | 33.801316 | 19.6444599 |
| TCGA-CJ-4639 | TCGA-KIRC | 42.6532155 | 14.3254843 |
| TCGA-CJ-4640 | TCGA-KIRC | 29.7271316 | 30.6093972 |
| TCGA-CJ-4641 | TCGA-KIRC | 41.6493273 | 18.9935987 |
| TCGA-CJ-4642 | TCGA-KIRC | 7.85116617 | 18.1249307 |
| TCGA-CJ-4643 | TCGA-KIRC | 43.1728417 | 13.9286165 |
| TCGA-CJ-4644 | TCGA-KIRC | 37.1029301 | 22.0036508 |
| TCGA-CJ-4868 | TCGA-KIRC | 46.1905727 | 18.0677367 |
| TCGA-CJ-4869 | TCGA-KIRC | 49.3888388 | 19.0738291 |
| TCGA-CJ-4870 | TCGA-KIRC | 40.9273181 | 12.8956537 |
| TCGA-CJ-4871 | TCGA-KIRC | 34.0109241 | 16.6933169 |
| TCGA-CJ-4872 | TCGA-KIRC | 41.6322867 | 14.0396012 |
| TCGA-CJ-4873 | TCGA-KIRC | 28.8997396 | 21.3457052 |
| TCGA-CJ-4874 | TCGA-KIRC | 38.8401093 | 18.1997353 |
| TCGA-CJ-4875 | TCGA-KIRC | 40.3996937 | 18.7222128 |
| TCGA-CJ-4876 | TCGA-KIRC | 41.2399558 | 39.0573749 |
| TCGA-CJ-4878 | TCGA-KIRC | 72.3592873 | 19.6726754 |

| | | | |
|---|---|---|---|
| TCGA-CJ-4881 | TCGA-KIRC | 49.3848276 | 27.4514769 |
| TCGA-CJ-4882 | TCGA-KIRC | 24.4265339 | 12.1235381 |
| TCGA-CJ-4884 | TCGA-KIRC | 36.5441199 | 21.4759454 |
| TCGA-CJ-4885 | TCGA-KIRC | 41.8687478 | 23.36606 |
| TCGA-CJ-4886 | TCGA-KIRC | 49.8349869 | 15.6149396 |
| TCGA-CJ-4887 | TCGA-KIRC | 41.1892128 | 10.8002613 |
| TCGA-CJ-4888 | TCGA-KIRC | 26.3218239 | 19.6650249 |
| TCGA-CJ-4889 | TCGA-KIRC | 31.0729819 | 13.2978636 |
| TCGA-CJ-4890 | TCGA-KIRC | 30.5281663 | 11.914193 |
| TCGA-CJ-4891 | TCGA-KIRC | 36.5688791 | 20.457078 |
| TCGA-CJ-4892 | TCGA-KIRC | 44.9748395 | 13.6225149 |
| TCGA-CJ-4893 | TCGA-KIRC | 45.457041 | 18.2241807 |
| TCGA-CJ-4894 | TCGA-KIRC | 39.2507577 | 22.0907615 |
| TCGA-CJ-4895 | TCGA-KIRC | 46.2131155 | 17.2597006 |
| TCGA-CJ-4897 | TCGA-KIRC | 31.1267343 | 12.1094702 |
| TCGA-CJ-4899 | TCGA-KIRC | 57.036201 | 14.157527 |
| TCGA-CJ-4900 | TCGA-KIRC | 31.5067654 | 11.2726729 |
| TCGA-CJ-4901 | TCGA-KIRC | 24.8171661 | 9.70610153 |
| TCGA-CJ-4902 | TCGA-KIRC | 44.5035488 | 15.5058332 |
| TCGA-CJ-4903 | TCGA-KIRC | 40.7111176 | 23.2505061 |
| TCGA-CJ-4904 | TCGA-KIRC | 54.851756 | 21.9255802 |
| TCGA-CJ-4905 | TCGA-KIRC | 49.7363114 | 16.8297054 |
| TCGA-CJ-4907 | TCGA-KIRC | 31.9621232 | 16.4800426 |
| TCGA-CJ-4908 | TCGA-KIRC | 47.531227 | 15.3521244 |
| TCGA-CJ-4912 | TCGA-KIRC | 27.6355811 | 18.6027566 |
| TCGA-CJ-4916 | TCGA-KIRC | 43.0136822 | 16.6569879 |
| TCGA-CJ-4918 | TCGA-KIRC | 31.8882007 | 9.84179137 |
| TCGA-CJ-4920 | TCGA-KIRC | 41.7173737 | 21.3794298 |
| TCGA-CJ-5671 | TCGA-KIRC | 31.3700966 | 18.7952611 |
| TCGA-CJ-5672 | TCGA-KIRC | 35.4939257 | 12.7482259 |
| TCGA-CJ-5675 | TCGA-KIRC | 57.9143056 | 25.2962363 |

| | | | |
|---|---|---|---|
| TCGA-CJ-5676 | TCGA-KIRC | 39.7140894 | 12.242795 |
| TCGA-CJ-5677 | TCGA-KIRC | 49.821671 | 27.9313621 |
| TCGA-CJ-5678 | TCGA-KIRC | 21.3058544 | 8.46932612 |
| TCGA-CJ-5679 | TCGA-KIRC | 43.4828084 | 13.8686123 |
| TCGA-CJ-5680 | TCGA-KIRC | 41.7659274 | 18.6543699 |
| TCGA-CJ-5681 | TCGA-KIRC | 35.7246356 | 13.7743405 |
| TCGA-CJ-5682 | TCGA-KIRC | 43.9791571 | 27.0603101 |
| TCGA-CJ-5683 | TCGA-KIRC | 53.4022813 | 17.4028681 |
| TCGA-CJ-5684 | TCGA-KIRC | 49.9530377 | 15.950849 |
| TCGA-CJ-5686 | TCGA-KIRC | 38.5333804 | 20.6816967 |
| TCGA-CJ-5689 | TCGA-KIRC | 45.5933389 | 18.1548217 |
| TCGA-CJ-6027 | TCGA-KIRC | 59.492312 | 31.1981915 |
| TCGA-CJ-6028 | TCGA-KIRC | 25.2249559 | 12.3644488 |
| TCGA-CJ-6030 | TCGA-KIRC | 50.4251022 | 21.560176 |
| TCGA-CJ-6031 | TCGA-KIRC | 33.3032566 | 10.4912362 |
| TCGA-CJ-6032 | TCGA-KIRC | 46.2122971 | 16.6373333 |
| TCGA-CJ-6033 | TCGA-KIRC | 55.80618 | 23.959359 |
| TCGA-CK-4947 | TCGA-COAD | 60.0159971 | 27.3734074 |
| TCGA-CK-4948 | TCGA-COAD | 75.5991153 | 45.0034214 |
| TCGA-CK-4950 | TCGA-COAD | 51.3678491 | 29.3128157 |
| TCGA-CK-4952 | TCGA-COAD | 74.4581183 | 45.7993538 |
| TCGA-CK-5912 | TCGA-COAD | 170.497533 | 25.3464258 |
| TCGA-CK-5913 | TCGA-COAD | 65.0938269 | 41.9876529 |
| TCGA-CK-5914 | TCGA-COAD | 107.148754 | 36.2422872 |
| TCGA-CK-5916 | TCGA-COAD | 73.7431565 | 17.7758635 |
| TCGA-CK-6746 | TCGA-COAD | 60.2259495 | 25.4415399 |
| TCGA-CK-6747 | TCGA-COAD | 60.7447038 | 22.1752322 |
| TCGA-CK-6748 | TCGA-COAD | 45.2029758 | 22.8872301 |
| TCGA-CK-6751 | TCGA-COAD | 58.3898874 | 28.3243658 |
| TCGA-CM-4743 | TCGA-COAD | 64.8074532 | 40.7332785 |
| TCGA-CM-4744 | TCGA-COAD | 92.1078794 | 60.8549888 |

| | | | |
|---|---|---|---|
| TCGA-CM-4751 | TCGA-COAD | 71.1254662 | 27.2612772 |
| TCGA-CM-5344 | TCGA-COAD | 67.9206042 | 31.8996767 |
| TCGA-CM-5348 | TCGA-COAD | 44.2664094 | 15.4458085 |
| TCGA-CM-5349 | TCGA-COAD | 92.8532282 | 30.0719338 |
| TCGA-CM-5860 | TCGA-COAD | 86.6134112 | 45.1389718 |
| TCGA-CM-5861 | TCGA-COAD | 44.721977 | 21.1012441 |
| TCGA-CM-5862 | TCGA-COAD | 77.9382782 | 21.466912 |
| TCGA-CM-5863 | TCGA-COAD | 35.5425481 | 10.6146936 |
| TCGA-CM-5864 | TCGA-COAD | 76.3260943 | 26.1751421 |
| TCGA-CM-5868 | TCGA-COAD | 103.206017 | 36.3271383 |
| TCGA-CM-6161 | TCGA-COAD | 73.643999 | 14.6885651 |
| TCGA-CM-6162 | TCGA-COAD | 60.3815345 | 28.5341181 |
| TCGA-CM-6163 | TCGA-COAD | 66.4123303 | 23.1122177 |
| TCGA-CM-6164 | TCGA-COAD | 75.7163186 | 23.1643744 |
| TCGA-CM-6165 | TCGA-COAD | 83.8539152 | 24.4125098 |
| TCGA-CM-6166 | TCGA-COAD | 138.857884 | 70.808772 |
| TCGA-CM-6167 | TCGA-COAD | 65.7530977 | 30.7465031 |
| TCGA-CM-6168 | TCGA-COAD | 66.4711865 | 25.0779295 |
| TCGA-CM-6169 | TCGA-COAD | 63.6613304 | 23.7810635 |
| TCGA-CM-6170 | TCGA-COAD | 74.4022901 | 23.040725 |
| TCGA-CM-6171 | TCGA-COAD | 58.7436496 | 31.521958 |
| TCGA-CM-6172 | TCGA-COAD | 98.4742448 | 18.5688164 |
| TCGA-CM-6674 | TCGA-COAD | 73.575964 | 23.9855929 |
| TCGA-CM-6675 | TCGA-COAD | 124.69024 | 38.1560621 |
| TCGA-CM-6676 | TCGA-COAD | 66.5668159 | 25.644744 |
| TCGA-CM-6677 | TCGA-COAD | 57.7455019 | 18.9105204 |
| TCGA-CM-6678 | TCGA-COAD | 53.6497915 | 15.3780914 |
| TCGA-CM-6679 | TCGA-COAD | 66.1585271 | 17.3939097 |
| TCGA-CM-6680 | TCGA-COAD | 63.4053735 | 15.4933806 |
| TCGA-CS-4938 | TCGA-LGG | 30.2472228 | 11.2290868 |
| TCGA-CS-4941 | TCGA-LGG | 54.2642617 | 20.8165127 |

| | | | |
|---|---|---|---|
| TCGA-CS-4942 | TCGA-LGG | 30.5601206 | 13.5263462 |
| TCGA-CS-4943 | TCGA-LGG | 41.7485738 | 76.9146813 |
| TCGA-CS-4944 | TCGA-LGG | 27.3161602 | 9.27134793 |
| TCGA-CS-5390 | TCGA-LGG | 17.0675127 | 17.5203901 |
| TCGA-CS-5393 | TCGA-LGG | 22.2253454 | 10.2409007 |
| TCGA-CS-5394 | TCGA-LGG | 17.4069958 | 16.6627174 |
| TCGA-CS-5395 | TCGA-LGG | 35.7419756 | 16.3573677 |
| TCGA-CS-5396 | TCGA-LGG | 16.1014763 | 19.4661983 |
| TCGA-CS-5397 | TCGA-LGG | 35.5262286 | 12.9278239 |
| TCGA-CS-6186 | TCGA-LGG | 56.0149162 | 21.1000864 |
| TCGA-CS-6188 | TCGA-LGG | 38.4615477 | 16.5039009 |
| TCGA-CS-6290 | TCGA-LGG | 37.022631 | 10.3444476 |
| TCGA-CS-6666 | TCGA-LGG | 36.3390157 | 12.6744661 |
| TCGA-CS-6667 | TCGA-LGG | 27.781261 | 8.97890244 |
| TCGA-CS-6668 | TCGA-LGG | 17.379091 | 19.7312492 |
| TCGA-CS-6669 | TCGA-LGG | 17.2394933 | 5.79589886 |
| TCGA-CS-6670 | TCGA-LGG | 17.509911 | 9.98507284 |
| TCGA-CU-A0YN | TCGA-BLCA | 68.131336 | 28.275407 |
| TCGA-CU-A0YO | TCGA-BLCA | 40.1417732 | 15.0338295 |
| TCGA-CU-A0YR | TCGA-BLCA | 47.1249035 | 33.8948487 |
| TCGA-CU-A3KJ | TCGA-BLCA | 77.6033091 | 43.7084379 |
| TCGA-CU-A3QU | TCGA-BLCA | 69.4380508 | 29.0579393 |
| TCGA-CU-A3YL | TCGA-BLCA | 44.1951614 | 9.64092322 |
| TCGA-CU-A5W6 | TCGA-BLCA | 74.2374279 | 40.8695817 |
| TCGA-CU-A72E | TCGA-BLCA | 57.7204076 | 20.4178108 |
| TCGA-CW-5580 | TCGA-KIRC | 42.7527438 | 18.5453181 |
| TCGA-CW-5581 | TCGA-KIRC | 37.007183 | 13.3708034 |
| TCGA-CW-5583 | TCGA-KIRC | 48.7612822 | 13.1305603 |
| TCGA-CW-5584 | TCGA-KIRC | 34.9007682 | 16.7503254 |
| TCGA-CW-5585 | TCGA-KIRC | 39.545076 | 11.23067 |
| TCGA-CW-5587 | TCGA-KIRC | 43.31507 | 19.7041748 |

| | | | |
|---|---|---|---|
| TCGA-CW-5588 | TCGA-KIRC | 26.5608589 | 19.6128064 |
| TCGA-CW-5589 | TCGA-KIRC | 43.6578083 | 13.146753 |
| TCGA-CW-5590 | TCGA-KIRC | 38.5434655 | 17.2721953 |
| TCGA-CW-5591 | TCGA-KIRC | 41.6232829 | 16.2663942 |
| TCGA-CW-6087 | TCGA-KIRC | 50.7449464 | 31.0862557 |
| TCGA-CW-6088 | TCGA-KIRC | 58.8289391 | 19.9472214 |
| TCGA-CW-6090 | TCGA-KIRC | 32.3492865 | 11.1075681 |
| TCGA-CW-6093 | TCGA-KIRC | 63.1965175 | 18.0848981 |
| TCGA-CW-6097 | TCGA-KIRC | 41.5307799 | 21.8273804 |
| TCGA-CZ-4853 | TCGA-KIRC | 42.4985238 | 12.6017682 |
| TCGA-CZ-4854 | TCGA-KIRC | 26.1163343 | 22.9415156 |
| TCGA-CZ-4856 | TCGA-KIRC | 36.561086 | 17.3360575 |
| TCGA-CZ-4857 | TCGA-KIRC | 33.5374373 | 18.0637426 |
| TCGA-CZ-4858 | TCGA-KIRC | 47.8075358 | 23.4653459 |
| TCGA-CZ-4859 | TCGA-KIRC | 59.706196 | 20.8240068 |
| TCGA-CZ-4860 | TCGA-KIRC | 42.3355871 | 27.8405266 |
| TCGA-CZ-4861 | TCGA-KIRC | 32.2996071 | 17.6165437 |
| TCGA-CZ-4862 | TCGA-KIRC | 45.4237706 | 17.9401241 |
| TCGA-CZ-4863 | TCGA-KIRC | 33.6599151 | 19.8972948 |
| TCGA-CZ-4864 | TCGA-KIRC | 51.2037044 | 21.0532796 |
| TCGA-CZ-4865 | TCGA-KIRC | 55.3350929 | 14.6062213 |
| TCGA-CZ-4866 | TCGA-KIRC | 48.2377491 | 29.7776863 |
| TCGA-CZ-5451 | TCGA-KIRC | 33.8588331 | 10.1357265 |
| TCGA-CZ-5452 | TCGA-KIRC | 49.2556608 | 18.1501 |
| TCGA-CZ-5453 | TCGA-KIRC | 56.6095994 | 12.8817561 |
| TCGA-CZ-5454 | TCGA-KIRC | 38.7872086 | 14.1191101 |
| TCGA-CZ-5455 | TCGA-KIRC | 52.8074999 | 19.9427402 |
| TCGA-CZ-5456 | TCGA-KIRC | 34.1719481 | 10.5476774 |
| TCGA-CZ-5457 | TCGA-KIRC | 31.5457775 | 12.6431331 |
| TCGA-CZ-5458 | TCGA-KIRC | 47.9076347 | 15.4758492 |
| TCGA-CZ-5459 | TCGA-KIRC | 40.2995569 | 10.8215819 |

| | | | |
|---|---|---|---|
| TCGA-CZ-5460 | TCGA-KIRC | 59.5414258 | 15.0317446 |
| TCGA-CZ-5461 | TCGA-KIRC | 44.7384158 | 20.9430609 |
| TCGA-CZ-5462 | TCGA-KIRC | 26.9722108 | 35.0179606 |
| TCGA-CZ-5463 | TCGA-KIRC | 41.2378403 | 21.5575331 |
| TCGA-CZ-5464 | TCGA-KIRC | 40.7763422 | 16.9628057 |
| TCGA-CZ-5465 | TCGA-KIRC | 45.3129049 | 12.0971633 |
| TCGA-CZ-5466 | TCGA-KIRC | 30.7479528 | 14.7557186 |
| TCGA-CZ-5467 | TCGA-KIRC | 37.3951346 | 14.8117492 |
| TCGA-CZ-5468 | TCGA-KIRC | 68.2336908 | 32.9666366 |
| TCGA-CZ-5469 | TCGA-KIRC | 54.196639 | 20.9728851 |
| TCGA-CZ-5470 | TCGA-KIRC | 40.5476549 | 11.5041048 |
| TCGA-CZ-5982 | TCGA-KIRC | 55.7510063 | 17.4097193 |
| TCGA-CZ-5984 | TCGA-KIRC | 36.9640172 | 26.0357892 |
| TCGA-CZ-5985 | TCGA-KIRC | 41.8415159 | 20.6123532 |
| TCGA-CZ-5986 | TCGA-KIRC | 36.1414246 | 10.1819849 |
| TCGA-CZ-5987 | TCGA-KIRC | 39.2438562 | 18.5194847 |
| TCGA-CZ-5988 | TCGA-KIRC | 38.7023731 | 11.7742119 |
| TCGA-CZ-5989 | TCGA-KIRC | 46.2325197 | 12.8938462 |
| TCGA-D1-A2G0 | TCGA-UCEC | 102.674012 | 8.76483712 |
| TCGA-D1-A3DA | TCGA-UCEC | 25.5723634 | 8.57795545 |
| TCGA-D1-A3DG | TCGA-UCEC | 186.516857 | 6.76363378 |
| TCGA-D1-A3DH | TCGA-UCEC | 82.728182 | 16.0463078 |
| TCGA-D1-A3JP | TCGA-UCEC | 84.1905077 | 31.6027513 |
| TCGA-D1-A3JQ | TCGA-UCEC | 75.3911455 | 21.4063702 |
| TCGA-D3-A1Q1 | TCGA-SKCM | 36.3925368 | 37.2236432 |
| TCGA-D3-A1Q3 | TCGA-SKCM | 49.0059053 | 24.2481122 |
| TCGA-D3-A1Q4 | TCGA-SKCM | 23.0010425 | 21.154258 |
| TCGA-D3-A1Q5 | TCGA-SKCM | 60.9133338 | 31.4957741 |
| TCGA-D3-A1Q6 | TCGA-SKCM | 59.5792418 | 66.0920758 |
| TCGA-D3-A1Q7 | TCGA-SKCM | 45.2126947 | 32.305912 |
| TCGA-D3-A1Q8 | TCGA-SKCM | 50.1980028 | 29.2468266 |

| | | | |
|---|---|---|---|
| TCGA-D3-A1Q9 | TCGA-SKCM | 31.9256787 | 18.4340885 |
| TCGA-D3-A1QA | TCGA-SKCM | 43.058349 | 19.3868 |
| TCGA-D3-A1QB | TCGA-SKCM | 45.0143823 | 18.674498 |
| TCGA-D3-A2J6 | TCGA-SKCM | 37.7759941 | 12.8675763 |
| TCGA-D3-A2J7 | TCGA-SKCM | 39.3479224 | 33.9613056 |
| TCGA-D3-A2J8 | TCGA-SKCM | 48.5345024 | 22.885695 |
| TCGA-D3-A2J9 | TCGA-SKCM | 41.8420718 | 16.991546 |
| TCGA-D3-A2JA | TCGA-SKCM | 74.4765818 | 29.6782072 |
| TCGA-D3-A2JB | TCGA-SKCM | 43.4000999 | 15.6919238 |
| TCGA-D3-A2JC | TCGA-SKCM | 42.3765798 | 25.2806552 |
| TCGA-D3-A2JD | TCGA-SKCM | 33.155247 | 9.93034585 |
| TCGA-D3-A2JE | TCGA-SKCM | 79.0698701 | 44.4516353 |
| TCGA-D3-A2JF | TCGA-SKCM | 41.7811488 | 19.090485 |
| TCGA-D3-A2JG | TCGA-SKCM | 26.6506926 | 19.1698426 |
| TCGA-D3-A2JH | TCGA-SKCM | 58.4881523 | 26.3384251 |
| TCGA-D3-A2JK | TCGA-SKCM | 27.9717088 | 9.37230361 |
| TCGA-D3-A2JL | TCGA-SKCM | 58.1767303 | 10.2814471 |
| TCGA-D3-A2JN | TCGA-SKCM | 61.6156141 | 45.9662975 |
| TCGA-D3-A2JO | TCGA-SKCM | 39.4732603 | 16.7187151 |
| TCGA-D3-A2JP | TCGA-SKCM | 34.659608 | 29.4085819 |
| TCGA-D3-A3BZ | TCGA-SKCM | 55.6664043 | 12.8022919 |
| TCGA-D3-A3C1 | TCGA-SKCM | 43.9909292 | 11.4639139 |
| TCGA-D3-A3C3 | TCGA-SKCM | 56.6676544 | 29.5428405 |
| TCGA-D3-A3C6 | TCGA-SKCM | 45.9693354 | 50.2411499 |
| TCGA-D3-A3C7 | TCGA-SKCM | 53.1113288 | 33.5215367 |
| TCGA-D3-A3C8 | TCGA-SKCM | 62.3214392 | 32.7421186 |
| TCGA-D3-A3CB | TCGA-SKCM | 22.179796 | 10.5577207 |
| TCGA-D3-A3CC | TCGA-SKCM | 33.5902819 | 39.9692533 |
| TCGA-D3-A3CE | TCGA-SKCM | 33.9553319 | 20.179426 |
| TCGA-D3-A3CF | TCGA-SKCM | 53.5578583 | 20.6037083 |
| TCGA-D3-A3ML | TCGA-SKCM | 28.3088134 | 18.8694829 |

| | | | |
|---|---|---|---|
| TCGA-D3-A3MO | TCGA-SKCM | 58.5873473 | 44.9164358 |
| TCGA-D3-A3MR | TCGA-SKCM | 46.1963535 | 26.217035 |
| TCGA-D3-A3MU | TCGA-SKCM | 37.5906944 | 20.7295589 |
| TCGA-D3-A3MV | TCGA-SKCM | 41.2842984 | 28.7692347 |
| TCGA-D3-A51E | TCGA-SKCM | 80.9903511 | 22.2604158 |
| TCGA-D3-A51F | TCGA-SKCM | 38.5935764 | 20.5005801 |
| TCGA-D3-A51G | TCGA-SKCM | 103.80808 | 28.1444055 |
| TCGA-D3-A51H | TCGA-SKCM | 54.7091452 | 19.5991725 |
| TCGA-D3-A51J | TCGA-SKCM | 47.2882033 | 33.0307999 |
| TCGA-D3-A51K | TCGA-SKCM | 63.906203 | 8.82072316 |
| TCGA-D3-A51N | TCGA-SKCM | 44.1551286 | 29.6221496 |
| TCGA-D3-A51R | TCGA-SKCM | 119.551323 | 58.9432855 |
| TCGA-D3-A51T | TCGA-SKCM | 57.8373546 | 21.3034784 |
| TCGA-D3-A5GL | TCGA-SKCM | 55.5865968 | 32.8399999 |
| TCGA-D3-A5GN | TCGA-SKCM | 51.390444 | 32.734202 |
| TCGA-D3-A5GO | TCGA-SKCM | 76.175542 | 26.6333932 |
| TCGA-D3-A5GR | TCGA-SKCM | 41.1745948 | 24.6712747 |
| TCGA-D3-A5GS | TCGA-SKCM | 39.5879284 | 19.0585036 |
| TCGA-D3-A5GT | TCGA-SKCM | 51.2741384 | 14.1669674 |
| TCGA-D3-A5GU | TCGA-SKCM | 51.6097191 | 22.7408117 |
| TCGA-D3-A8GB | TCGA-SKCM | 44.5793594 | 8.25335905 |
| TCGA-D3-A8GC | TCGA-SKCM | 71.6277325 | 36.2946347 |
| TCGA-D3-A8GD | TCGA-SKCM | 79.4063173 | 17.5163179 |
| TCGA-D3-A8GE | TCGA-SKCM | 49.9364513 | 25.6638787 |
| TCGA-D3-A8GI | TCGA-SKCM | 45.2978817 | 15.1318912 |
| TCGA-D3-A8GJ | TCGA-SKCM | 68.369665 | 23.442635 |
| TCGA-D3-A8GK | TCGA-SKCM | 38.1703796 | 28.455093 |
| TCGA-D3-A8GL | TCGA-SKCM | 71.9158515 | 38.0634478 |
| TCGA-D3-A8GM | TCGA-SKCM | 84.241476 | 24.9787771 |
| TCGA-D3-A8GN | TCGA-SKCM | 38.1599558 | 11.1565748 |
| TCGA-D3-A8GO | TCGA-SKCM | 40.913516 | 35.2799749 |

| | | | |
|---|---|---|---|
| TCGA-D3-A8GP | TCGA-SKCM | 65.340762 | 29.3985742 |
| TCGA-D3-A8GQ | TCGA-SKCM | 54.5235583 | 25.4451086 |
| TCGA-D3-A8GR | TCGA-SKCM | 51.6672349 | 27.8371994 |
| TCGA-D3-A8GS | TCGA-SKCM | 48.1206644 | 44.2299546 |
| TCGA-D3-A8GV | TCGA-SKCM | 88.0635924 | 38.6151558 |
| TCGA-D5-5537 | TCGA-COAD | 78.7033879 | 21.5212459 |
| TCGA-D5-5538 | TCGA-COAD | 72.6923641 | 31.3258533 |
| TCGA-D5-5539 | TCGA-COAD | 67.5339593 | 29.3431476 |
| TCGA-D5-5540 | TCGA-COAD | 65.3089706 | 69.8278384 |
| TCGA-D5-5541 | TCGA-COAD | 85.7200081 | 25.0403823 |
| TCGA-D5-6529 | TCGA-COAD | 60.0447022 | 36.2160548 |
| TCGA-D5-6530 | TCGA-COAD | 45.4028849 | 14.4690529 |
| TCGA-D5-6531 | TCGA-COAD | 55.8904486 | 29.0856091 |
| TCGA-D5-6532 | TCGA-COAD | 96.5687825 | 30.0880927 |
| TCGA-D5-6533 | TCGA-COAD | 46.4953171 | 39.2144589 |
| TCGA-D5-6534 | TCGA-COAD | 45.6713619 | 15.4237425 |
| TCGA-D5-6535 | TCGA-COAD | 34.1193406 | 19.4097821 |
| TCGA-D5-6536 | TCGA-COAD | 29.6289661 | 29.1320559 |
| TCGA-D5-6537 | TCGA-COAD | 74.8426225 | 43.1505564 |
| TCGA-D5-6538 | TCGA-COAD | 45.4562966 | 12.2584924 |
| TCGA-D5-6539 | TCGA-COAD | 48.8496023 | 21.3877824 |
| TCGA-D5-6540 | TCGA-COAD | 54.2928187 | 32.6131575 |
| TCGA-D5-6541 | TCGA-COAD | 75.57433 | 18.2398579 |
| TCGA-D5-6898 | TCGA-COAD | 65.1494728 | 17.8116604 |
| TCGA-D5-6920 | TCGA-COAD | 60.5379602 | 17.8639293 |
| TCGA-D5-6922 | TCGA-COAD | 112.452952 | 23.7849616 |
| TCGA-D5-6923 | TCGA-COAD | 98.6917643 | 35.0861169 |
| TCGA-D5-6924 | TCGA-COAD | 54.168422 | 19.8178114 |
| TCGA-D5-6926 | TCGA-COAD | 69.204077 | 15.8313952 |
| TCGA-D5-6927 | TCGA-COAD | 59.301455 | 30.1334889 |
| TCGA-D5-6928 | TCGA-COAD | 40.7141883 | 24.4239625 |

| | | | |
|---|---|---|---|
| TCGA-D5-6929 | TCGA-COAD | 43.9884044 | 15.974398 |
| TCGA-D5-6930 | TCGA-COAD | 54.5471607 | 32.8997706 |
| TCGA-D5-6931 | TCGA-COAD | 76.2766678 | 38.6346058 |
| TCGA-D5-6932 | TCGA-COAD | 65.2421608 | 23.6171405 |
| TCGA-D5-7000 | TCGA-COAD | 62.4211557 | 34.90045 |
| TCGA-D7-5577 | TCGA-STAD | 62.528173 | 32.4817042 |
| TCGA-D7-5578 | TCGA-STAD | 54.2492181 | 26.164887 |
| TCGA-D7-6519 | TCGA-STAD | 33.6976724 | 22.2945357 |
| TCGA-D7-6520 | TCGA-STAD | 37.0404233 | 26.7706367 |
| TCGA-D7-6521 | TCGA-STAD | 47.849602 | 24.7847232 |
| TCGA-D7-6522 | TCGA-STAD | 31.3107238 | 18.8783935 |
| TCGA-D7-6524 | TCGA-STAD | 37.0823329 | 17.4249846 |
| TCGA-D7-6525 | TCGA-STAD | 60.531781 | 28.2950113 |
| TCGA-D7-6526 | TCGA-STAD | 32.6606163 | 35.4576846 |
| TCGA-D7-6527 | TCGA-STAD | 29.7750491 | 24.1183821 |
| TCGA-D7-6528 | TCGA-STAD | 56.0729564 | 40.0594782 |
| TCGA-D7-6815 | TCGA-STAD | 47.237342 | 13.0325317 |
| TCGA-D7-6818 | TCGA-STAD | 25.1720418 | 19.8766774 |
| TCGA-D7-6822 | TCGA-STAD | 46.858555 | 19.1079899 |
| TCGA-D7-8570 | TCGA-STAD | 38.9378157 | 23.8374062 |
| TCGA-D7-8572 | TCGA-STAD | 59.9304077 | 24.2436041 |
| TCGA-D7-8573 | TCGA-STAD | 50.8531293 | 30.8134356 |
| TCGA-D7-8574 | TCGA-STAD | 39.4175226 | 18.4224153 |
| TCGA-D7-8575 | TCGA-STAD | 46.2708047 | 13.2681693 |
| TCGA-D7-8576 | TCGA-STAD | 53.9195541 | 33.1607806 |
| TCGA-D7-8578 | TCGA-STAD | 52.4317638 | 24.2792602 |
| TCGA-D7-8579 | TCGA-STAD | 28.5621265 | 14.3011072 |
| TCGA-D7-A4YU | TCGA-STAD | 75.5880684 | 42.3449378 |
| TCGA-D7-A4YX | TCGA-STAD | 58.2457388 | 33.4841349 |
| TCGA-D7-A4Z0 | TCGA-STAD | 24.5098848 | 12.9495639 |
| TCGA-D7-A6EV | TCGA-STAD | 44.5025398 | 45.4567283 |

| | | | |
|---|---|---|---|
| TCGA-D7-A6EX | TCGA-STAD | 26.1206477 | 32.9824676 |
| TCGA-D7-A6EY | TCGA-STAD | 29.5197171 | 16.4825267 |
| TCGA-D7-A6EZ | TCGA-STAD | 70.5647552 | 52.592766 |
| TCGA-D7-A6F0 | TCGA-STAD | 49.594155 | 68.5250099 |
| TCGA-D7-A6F2 | TCGA-STAD | 16.4674017 | 39.6291618 |
| TCGA-D7-A747 | TCGA-STAD | 21.5635489 | 13.2513505 |
| TCGA-D7-A748 | TCGA-STAD | 25.672637 | 15.6876054 |
| TCGA-D7-A74A | TCGA-STAD | 49.891396 | 42.7280773 |
| TCGA-D8-A13Y | TCGA-BRCA | 133.120418 | 55.22541 |
| TCGA-D8-A13Z | TCGA-BRCA | 106.784151 | 55.0037971 |
| TCGA-D8-A140 | TCGA-BRCA | 36.0876587 | 32.3506038 |
| TCGA-D8-A141 | TCGA-BRCA | 43.3838686 | 23.1425249 |
| TCGA-D8-A142 | TCGA-BRCA | 33.9285088 | 32.1096314 |
| TCGA-D8-A143 | TCGA-BRCA | 96.3385599 | 60.087165 |
| TCGA-D8-A145 | TCGA-BRCA | 34.9874772 | 17.2341147 |
| TCGA-D8-A146 | TCGA-BRCA | 50.7978959 | 42.6306864 |
| TCGA-D8-A147 | TCGA-BRCA | 105.561998 | 70.8093574 |
| TCGA-D8-A1J8 | TCGA-BRCA | 45.5695347 | 96.6707665 |
| TCGA-D8-A1J9 | TCGA-BRCA | 56.7893434 | 38.762536 |
| TCGA-D8-A1JA | TCGA-BRCA | 38.0262541 | 33.7100678 |
| TCGA-D8-A1JB | TCGA-BRCA | 48.8551495 | 23.6835984 |
| TCGA-D8-A1JC | TCGA-BRCA | 37.4669677 | 43.5535242 |
| TCGA-D8-A1JD | TCGA-BRCA | 67.6905016 | 35.0412338 |
| TCGA-D8-A1JE | TCGA-BRCA | 46.197442 | 26.2884884 |
| TCGA-D8-A1JF | TCGA-BRCA | 20.6737177 | 34.7999265 |
| TCGA-D8-A1JG | TCGA-BRCA | 62.1008767 | 32.4980368 |
| TCGA-D8-A1JH | TCGA-BRCA | 45.1712229 | 19.5455562 |
| TCGA-D8-A1JI | TCGA-BRCA | 52.9142859 | 36.1767941 |
| TCGA-D8-A1JJ | TCGA-BRCA | 56.3681221 | 69.8061687 |
| TCGA-D8-A1JK | TCGA-BRCA | 52.6562482 | 31.2278978 |
| TCGA-D8-A1JL | TCGA-BRCA | 90.2585752 | 37.1378276 |

| | | | |
|---|---|---|---|
| TCGA-D8-A1JM | TCGA-BRCA | 76.516363 | 152.497149 |
| TCGA-D8-A1JN | TCGA-BRCA | 58.7805529 | 51.4544275 |
| TCGA-D8-A1JP | TCGA-BRCA | 58.4312198 | 33.1854109 |
| TCGA-D8-A1JS | TCGA-BRCA | 66.8007206 | 26.9103892 |
| TCGA-D8-A1JT | TCGA-BRCA | 48.0519363 | 39.1385403 |
| TCGA-D8-A1JU | TCGA-BRCA | 53.8638592 | 18.1470473 |
| TCGA-D8-A1X5 | TCGA-BRCA | 34.2320969 | 27.3572322 |
| TCGA-D8-A1X6 | TCGA-BRCA | 71.6544097 | 34.6417794 |
| TCGA-D8-A1X7 | TCGA-BRCA | 64.7710366 | 48.13445 |
| TCGA-D8-A1X8 | TCGA-BRCA | 72.0198094 | 37.7031679 |
| TCGA-D8-A1X9 | TCGA-BRCA | 125.982445 | 22.0717526 |
| TCGA-D8-A1XA | TCGA-BRCA | 53.4208008 | 41.2303949 |
| TCGA-D8-A1XB | TCGA-BRCA | 47.9263065 | 29.3187361 |
| TCGA-D8-A1XC | TCGA-BRCA | 36.2310222 | 21.7859736 |
| TCGA-D8-A1XD | TCGA-BRCA | 45.2865877 | 33.3433253 |
| TCGA-D8-A1XF | TCGA-BRCA | 25.9320356 | 34.8639478 |
| TCGA-D8-A1XG | TCGA-BRCA | 30.4777212 | 39.2154592 |
| TCGA-D8-A1XJ | TCGA-BRCA | 74.5593826 | 52.3012528 |
| TCGA-D8-A1XK | TCGA-BRCA | 217.623714 | 45.9318952 |
| TCGA-D8-A1XL | TCGA-BRCA | 80.9101228 | 34.9038933 |
| TCGA-D8-A1XM | TCGA-BRCA | 58.1526536 | 17.7618706 |
| TCGA-D8-A1XO | TCGA-BRCA | 61.7734594 | 22.6640286 |
| TCGA-D8-A1XQ | TCGA-BRCA | 66.7801635 | 24.5910389 |
| TCGA-D8-A1XR | TCGA-BRCA | 44.9884573 | 60.7242478 |
| TCGA-D8-A1XS | TCGA-BRCA | 40.9521259 | 30.8512811 |
| TCGA-D8-A1XT | TCGA-BRCA | 128.51294 | 37.4789073 |
| TCGA-D8-A1XU | TCGA-BRCA | 33.724101 | 39.0336543 |
| TCGA-D8-A1XV | TCGA-BRCA | 33.0312319 | 27.7360967 |
| TCGA-D8-A1XW | TCGA-BRCA | 47.9344027 | 31.4945944 |
| TCGA-D8-A1XY | TCGA-BRCA | 36.9813991 | 28.8096395 |
| TCGA-D8-A1XZ | TCGA-BRCA | 52.6345014 | 32.4477797 |

| | | | |
|---|---|---|---|
| TCGA-D8-A1Y0 | TCGA-BRCA | 65.5016938 | 35.5179351 |
| TCGA-D8-A1Y1 | TCGA-BRCA | 50.2798707 | 43.9659049 |
| TCGA-D8-A1Y2 | TCGA-BRCA | 50.9681989 | 19.0121398 |
| TCGA-D8-A1Y3 | TCGA-BRCA | 104.230488 | 59.4768246 |
| TCGA-D8-A27E | TCGA-BRCA | 37.8960638 | 22.1903664 |
| TCGA-D8-A27F | TCGA-BRCA | 121.619711 | 43.9566713 |
| TCGA-D8-A27G | TCGA-BRCA | 46.6666615 | 25.2706254 |
| TCGA-D8-A27H | TCGA-BRCA | 46.9189167 | 24.1404308 |
| TCGA-D8-A27I | TCGA-BRCA | 43.5388111 | 31.5729164 |
| TCGA-D8-A27K | TCGA-BRCA | 39.7645872 | 23.9827274 |
| TCGA-D8-A27L | TCGA-BRCA | 53.1300908 | 21.0311981 |
| TCGA-D8-A27M | TCGA-BRCA | 57.8180783 | 28.04016 |
| TCGA-D8-A27N | TCGA-BRCA | 44.6518829 | 27.2847446 |
| TCGA-D8-A27P | TCGA-BRCA | 43.8365241 | 21.1333602 |
| TCGA-D8-A27R | TCGA-BRCA | 58.5186242 | 43.5692569 |
| TCGA-D8-A27T | TCGA-BRCA | 50.5306887 | 31.3205168 |
| TCGA-D8-A27V | TCGA-BRCA | 52.5646128 | 38.1840449 |
| TCGA-D8-A27W | TCGA-BRCA | 32.9780443 | 53.2666474 |
| TCGA-D8-A3Z5 | TCGA-BRCA | 46.4143649 | 17.1019833 |
| TCGA-D8-A3Z6 | TCGA-BRCA | 42.3606351 | 26.2385529 |
| TCGA-D8-A4Z1 | TCGA-BRCA | 41.4168594 | 19.4734166 |
| TCGA-D8-A73U | TCGA-BRCA | 29.5898754 | 27.3166041 |
| TCGA-D8-A73W | TCGA-BRCA | 36.6291743 | 32.7508009 |
| TCGA-D8-A73X | TCGA-BRCA | 63.1747647 | 27.9401134 |
| TCGA-D9-A148 | TCGA-SKCM | 40.8492804 | 16.5970954 |
| TCGA-D9-A149 | TCGA-SKCM | 54.3748897 | 27.9635204 |
| TCGA-D9-A1JW | TCGA-SKCM | 32.1346846 | 14.0269057 |
| TCGA-D9-A1JX | TCGA-SKCM | 49.8642142 | 20.2005735 |
| TCGA-D9-A1X3 | TCGA-SKCM | 60.1181567 | 17.7062151 |
| TCGA-D9-A3Z1 | TCGA-SKCM | 54.6918864 | 17.3695174 |
| TCGA-D9-A3Z3 | TCGA-SKCM | 31.9961408 | 13.528567 |

| | | | |
|---|---|---|---|
| TCGA-D9-A3Z4 | TCGA-SKCM | 62.7758956 | 23.8435574 |
| TCGA-D9-A4Z2 | TCGA-SKCM | 44.1617489 | 24.4078121 |
| TCGA-D9-A4Z3 | TCGA-SKCM | 30.3197592 | 11.4328111 |
| TCGA-D9-A4Z5 | TCGA-SKCM | 36.7986998 | 28.5005613 |
| TCGA-D9-A4Z6 | TCGA-SKCM | 29.5499463 | 25.3905591 |
| TCGA-D9-A6E9 | TCGA-SKCM | 40.2021138 | 18.690888 |
| TCGA-D9-A6EA | TCGA-SKCM | 75.411227 | 32.608877 |
| TCGA-D9-A6EC | TCGA-SKCM | 93.6621342 | 27.2729022 |
| TCGA-D9-A6EG | TCGA-SKCM | 68.7927363 | 45.5675046 |
| TCGA-DA-A1HV | TCGA-SKCM | 57.162156 | 17.4494494 |
| TCGA-DA-A1HW | TCGA-SKCM | 48.2261825 | 18.8249358 |
| TCGA-DA-A1HY | TCGA-SKCM | 68.2015563 | 51.5125248 |
| TCGA-DA-A1I0 | TCGA-SKCM | 23.6536856 | 7.31641534 |
| TCGA-DA-A1I1 | TCGA-SKCM | 35.918267 | 14.2354293 |
| TCGA-DA-A1I2 | TCGA-SKCM | 37.4277391 | 27.9371599 |
| TCGA-DA-A1I4 | TCGA-SKCM | 59.3279083 | 34.363449 |
| TCGA-DA-A1I5 | TCGA-SKCM | 48.0871711 | 24.3052597 |
| TCGA-DA-A1I7 | TCGA-SKCM | 23.6864772 | 60.4222019 |
| TCGA-DA-A1I8 | TCGA-SKCM | 32.3971685 | 42.7235619 |
| TCGA-DA-A1IA | TCGA-SKCM | 23.691848 | 14.5808479 |
| TCGA-DA-A1IB | TCGA-SKCM | 30.7195783 | 13.0099379 |
| TCGA-DA-A1IC | TCGA-SKCM | 49.3760098 | 22.882609 |
| TCGA-DA-A3F2 | TCGA-SKCM | 11.216157 | 3.05103652 |
| TCGA-DA-A3F3 | TCGA-SKCM | 36.2049391 | 12.4792258 |
| TCGA-DA-A3F5 | TCGA-SKCM | 28.9988612 | 49.5247551 |
| TCGA-DA-A3F8 | TCGA-SKCM | 38.8094543 | 19.6242103 |
| TCGA-DA-A95V | TCGA-SKCM | 29.5016445 | 40.3445412 |
| TCGA-DA-A95W | TCGA-SKCM | 55.7056036 | 26.3223641 |
| TCGA-DA-A95X | TCGA-SKCM | 77.3888098 | 24.2322895 |
| TCGA-DA-A95Y | TCGA-SKCM | 45.0221763 | 22.161708 |
| TCGA-DA-A95Z | TCGA-SKCM | 46.0082592 | 42.7474611 |

| | | | |
|---|---|---|---|
| TCGA-DA-A960 | TCGA-SKCM | 49.5148166 | 10.6721036 |
| TCGA-DB-5270 | TCGA-LGG | 30.145389 | 8.25156786 |
| TCGA-DB-5273 | TCGA-LGG | 30.4644628 | 17.1569199 |
| TCGA-DB-5274 | TCGA-LGG | 23.3493825 | 31.6300897 |
| TCGA-DB-5275 | TCGA-LGG | 46.622983 | 24.6927365 |
| TCGA-DB-5277 | TCGA-LGG | 57.9112386 | 56.7121126 |
| TCGA-DB-5278 | TCGA-LGG | 19.7854302 | 14.6898563 |
| TCGA-DB-5279 | TCGA-LGG | 18.8451489 | 19.9446021 |
| TCGA-DB-5280 | TCGA-LGG | 32.1389006 | 12.2276077 |
| TCGA-DB-5281 | TCGA-LGG | 31.3545944 | 8.22370567 |
| TCGA-DB-A4X9 | TCGA-LGG | 33.1451841 | 16.4226737 |
| TCGA-DB-A4XA | TCGA-LGG | 19.6243993 | 12.9934692 |
| TCGA-DB-A4XB | TCGA-LGG | 41.7393863 | 14.2085763 |
| TCGA-DB-A4XC | TCGA-LGG | 28.620056 | 13.2352156 |
| TCGA-DB-A4XD | TCGA-LGG | 28.5316824 | 10.7969436 |
| TCGA-DB-A4XE | TCGA-LGG | 33.4347992 | 24.9078721 |
| TCGA-DB-A4XF | TCGA-LGG | 23.0349273 | 9.89418376 |
| TCGA-DB-A4XG | TCGA-LGG | 21.7802297 | 17.3334399 |
| TCGA-DB-A4XH | TCGA-LGG | 10.6468931 | 8.33810961 |
| TCGA-DB-A64L | TCGA-LGG | 22.8283724 | 21.7360783 |
| TCGA-DB-A64O | TCGA-LGG | 42.4403279 | 15.7759659 |
| TCGA-DB-A64P | TCGA-LGG | 19.7076201 | 24.0300076 |
| TCGA-DB-A64Q | TCGA-LGG | 31.1548793 | 29.9009174 |
| TCGA-DB-A64R | TCGA-LGG | 21.8540046 | 17.3711758 |
| TCGA-DB-A64S | TCGA-LGG | 28.8949944 | 11.910554 |
| TCGA-DB-A64U | TCGA-LGG | 23.8667593 | 18.8915313 |
| TCGA-DB-A64V | TCGA-LGG | 23.3914904 | 17.4090927 |
| TCGA-DB-A64W | TCGA-LGG | 26.2047358 | 22.7667992 |
| TCGA-DB-A64X | TCGA-LGG | 30.3014788 | 9.93273878 |
| TCGA-DB-A75K | TCGA-LGG | 24.276313 | 18.6373851 |
| TCGA-DB-A75L | TCGA-LGG | 37.3163248 | 13.0380538 |

| | | | |
|---|---|---|---|
| TCGA-DB-A75M | TCGA-LGG | 35.7013984 | 11.0073102 |
| TCGA-DB-A75O | TCGA-LGG | 31.9009112 | 13.9113626 |
| TCGA-DB-A75P | TCGA-LGG | 27.6013761 | 8.54027209 |
| TCGA-DD-A113 | TCGA-LIHC | 17.3046667 | 18.799557 |
| TCGA-DD-A114 | TCGA-LIHC | 62.5650828 | 16.5974067 |
| TCGA-DD-A115 | TCGA-LIHC | 31.8925533 | 20.2210209 |
| TCGA-DD-A116 | TCGA-LIHC | 21.6393928 | 7.39612095 |
| TCGA-DD-A118 | TCGA-LIHC | 23.9861802 | 11.6264435 |
| TCGA-DD-A119 | TCGA-LIHC | 36.8382348 | 16.6777744 |
| TCGA-DD-A11A | TCGA-LIHC | 19.7290078 | 9.34119072 |
| TCGA-DD-A11B | TCGA-LIHC | 17.3472785 | 20.919936 |
| TCGA-DD-A11C | TCGA-LIHC | 16.9831394 | 6.64285076 |
| TCGA-DD-A11D | TCGA-LIHC | 23.9100084 | 8.76487787 |
| TCGA-DD-A1EA | TCGA-LIHC | 38.4242609 | 17.2711312 |
| TCGA-DD-A1EB | TCGA-LIHC | 27.0306627 | 9.45318331 |
| TCGA-DD-A1EC | TCGA-LIHC | 16.2597524 | 4.3787204 |
| TCGA-DD-A1ED | TCGA-LIHC | 25.6022549 | 10.0388307 |
| TCGA-DD-A1EE | TCGA-LIHC | 37.676894 | 22.6888544 |
| TCGA-DD-A1EF | TCGA-LIHC | 59.6127658 | 18.9343981 |
| TCGA-DD-A1EG | TCGA-LIHC | 26.6914813 | 7.31175052 |
| TCGA-DD-A1EH | TCGA-LIHC | 31.3032685 | 17.7014492 |
| TCGA-DD-A1EI | TCGA-LIHC | 32.3976543 | 19.827461 |
| TCGA-DD-A1EJ | TCGA-LIHC | 77.1372872 | 15.253778 |
| TCGA-DD-A1EK | TCGA-LIHC | 30.0393371 | 10.7264533 |
| TCGA-DD-A1EL | TCGA-LIHC | 22.300782 | 9.26772048 |
| TCGA-DD-A39V | TCGA-LIHC | 18.8988998 | 6.86603596 |
| TCGA-DD-A39W | TCGA-LIHC | 23.4540339 | 7.15862819 |
| TCGA-DD-A39X | TCGA-LIHC | 25.3127451 | 9.27106018 |
| TCGA-DD-A39Y | TCGA-LIHC | 91.0611074 | 7.94206966 |
| TCGA-DD-A39Z | TCGA-LIHC | 32.7559731 | 13.9979658 |
| TCGA-DD-A3A1 | TCGA-LIHC | 18.9077597 | 5.96620721 |

| | | | |
|---|---|---|---|
| TCGA-DD-A3A2 | TCGA-LIHC | 17.1589828 | 6.51957457 |
| TCGA-DD-A3A3 | TCGA-LIHC | 30.8659397 | 8.63781746 |
| TCGA-DD-A3A4 | TCGA-LIHC | 37.44599 | 17.2120636 |
| TCGA-DD-A3A5 | TCGA-LIHC | 47.5596931 | 19.725252 |
| TCGA-DD-A3A6 | TCGA-LIHC | 20.1257541 | 10.1529077 |
| TCGA-DD-A3A7 | TCGA-LIHC | 21.3535025 | 7.95730454 |
| TCGA-DD-A3A8 | TCGA-LIHC | 17.0581073 | 4.32781434 |
| TCGA-DD-A3A9 | TCGA-LIHC | 26.6829431 | 28.6679742 |
| TCGA-DD-A4NA | TCGA-LIHC | 54.7254618 | 11.8722563 |
| TCGA-DD-A4NB | TCGA-LIHC | 31.72397 | 13.351071 |
| TCGA-DD-A4ND | TCGA-LIHC | 44.2665883 | 10.1747499 |
| TCGA-DD-A4NE | TCGA-LIHC | 58.2386756 | 25.2610088 |
| TCGA-DD-A4NF | TCGA-LIHC | 38.7136459 | 19.8619483 |
| TCGA-DD-A4NG | TCGA-LIHC | 18.285014 | 13.0864996 |
| TCGA-DD-A4NH | TCGA-LIHC | 30.4799572 | 11.8003792 |
| TCGA-DD-A4NI | TCGA-LIHC | 27.6519287 | 12.8069881 |
| TCGA-DD-A4NJ | TCGA-LIHC | 88.5083261 | 11.598469 |
| TCGA-DD-A4NK | TCGA-LIHC | 30.6442752 | 16.3309257 |
| TCGA-DD-A4NL | TCGA-LIHC | 20.8507291 | 8.20054628 |
| TCGA-DD-A4NN | TCGA-LIHC | 41.8119441 | 28.3396463 |
| TCGA-DD-A4NO | TCGA-LIHC | 27.0151512 | 14.5207006 |
| TCGA-DD-A4NP | TCGA-LIHC | 25.845039 | 10.3948279 |
| TCGA-DD-A4NQ | TCGA-LIHC | 41.4273827 | 16.6390089 |
| TCGA-DD-A4NR | TCGA-LIHC | 34.5174033 | 20.9298369 |
| TCGA-DD-A4NS | TCGA-LIHC | 40.429562 | 12.6165465 |
| TCGA-DD-A4NV | TCGA-LIHC | 29.9540015 | 15.8835762 |
| TCGA-DD-A73A | TCGA-LIHC | 29.8053906 | 16.5558717 |
| TCGA-DD-A73B | TCGA-LIHC | 30.9114585 | 24.2822779 |
| TCGA-DD-A73C | TCGA-LIHC | 27.4058954 | 12.2322858 |
| TCGA-DD-A73D | TCGA-LIHC | 38.1330877 | 14.1313724 |
| TCGA-DD-A73E | TCGA-LIHC | 56.0340496 | 12.8261802 |

| | | | |
|---|---|---|---|
| TCGA-DD-A73F | TCGA-LIHC | 40.1555782 | 10.9869031 |
| TCGA-DD-A73G | TCGA-LIHC | 22.9413561 | 24.6771195 |
| TCGA-DD-AA3A | TCGA-LIHC | 76.7284417 | 38.4200821 |
| TCGA-DD-AAC8 | TCGA-LIHC | 41.8272375 | 28.4251423 |
| TCGA-DD-AAC9 | TCGA-LIHC | 27.0291755 | 13.2906716 |
| TCGA-DD-AACA | TCGA-LIHC | 55.019597 | 22.6734436 |
| TCGA-DD-AACB | TCGA-LIHC | 64.7379613 | 10.8276386 |
| TCGA-DD-AACC | TCGA-LIHC | 40.072887 | 20.0418099 |
| TCGA-DD-AACD | TCGA-LIHC | 25.8056908 | 14.9141558 |
| TCGA-DD-AACE | TCGA-LIHC | 35.3663512 | 9.49313177 |
| TCGA-DD-AACF | TCGA-LIHC | 41.0896134 | 26.4332685 |
| TCGA-DD-AACG | TCGA-LIHC | 36.2312294 | 20.0138286 |
| TCGA-DD-AACH | TCGA-LIHC | 59.8927917 | 19.5067133 |
| TCGA-DD-AACI | TCGA-LIHC | 33.8324679 | 30.4253791 |
| TCGA-DD-AACJ | TCGA-LIHC | 58.4304138 | 15.2673423 |
| TCGA-DD-AACK | TCGA-LIHC | 58.572375 | 21.8272294 |
| TCGA-DD-AACL | TCGA-LIHC | 60.9530727 | 21.0608771 |
| TCGA-DD-AACN | TCGA-LIHC | 38.5035357 | 14.9375364 |
| TCGA-DD-AACO | TCGA-LIHC | 33.0919718 | 26.5086744 |
| TCGA-DD-AACP | TCGA-LIHC | 46.2078912 | 26.2394378 |
| TCGA-DD-AACQ | TCGA-LIHC | 28.0999272 | 7.16410484 |
| TCGA-DD-AACS | TCGA-LIHC | 46.210551 | 21.5809481 |
| TCGA-DD-AACT | TCGA-LIHC | 33.9243071 | 20.3588803 |
| TCGA-DD-AACU | TCGA-LIHC | 60.1736234 | 23.6012508 |
| TCGA-DD-AACV | TCGA-LIHC | 61.7812858 | 12.9581036 |
| TCGA-DD-AACW | TCGA-LIHC | 37.5705017 | 16.5413285 |
| TCGA-DD-AACX | TCGA-LIHC | 49.9945715 | 19.7903056 |
| TCGA-DD-AACY | TCGA-LIHC | 34.817431 | 7.48200927 |
| TCGA-DD-AACZ | TCGA-LIHC | 55.3978734 | 34.262363 |
| TCGA-DD-AAD0 | TCGA-LIHC | 65.5541406 | 19.3564506 |
| TCGA-DD-AAD1 | TCGA-LIHC | 35.9621122 | 16.2825445 |

| | | | |
|---|---|---|---|
| TCGA-DD-AAD2 | TCGA-LIHC | 30.346319 | 16.8904644 |
| TCGA-DD-AAD3 | TCGA-LIHC | 29.2359409 | 11.9611401 |
| TCGA-DD-AAD5 | TCGA-LIHC | 55.9999991 | 20.2503529 |
| TCGA-DD-AAD6 | TCGA-LIHC | 49.4930463 | 11.0248028 |
| TCGA-DD-AAD8 | TCGA-LIHC | 59.820093 | 29.5975185 |
| TCGA-DD-AADA | TCGA-LIHC | 30.5537582 | 25.2644953 |
| TCGA-DD-AADB | TCGA-LIHC | 36.4090546 | 18.0799711 |
| TCGA-DD-AADC | TCGA-LIHC | 78.6520693 | 34.3238726 |
| TCGA-DD-AADD | TCGA-LIHC | 94.2366239 | 25.8973604 |
| TCGA-DD-AADF | TCGA-LIHC | 52.5737985 | 31.9362708 |
| TCGA-DD-AADG | TCGA-LIHC | 44.7920778 | 18.045663 |
| TCGA-DD-AADI | TCGA-LIHC | 37.8512116 | 25.4490304 |
| TCGA-DD-AADJ | TCGA-LIHC | 21.6775409 | 13.0370559 |
| TCGA-DD-AADK | TCGA-LIHC | 30.2435428 | 13.0455033 |
| TCGA-DD-AADL | TCGA-LIHC | 50.1337663 | 14.7911277 |
| TCGA-DD-AADM | TCGA-LIHC | 32.248919 | 15.3517003 |
| TCGA-DD-AADN | TCGA-LIHC | 26.27697 | 13.3908053 |
| TCGA-DD-AADO | TCGA-LIHC | 20.321296 | 7.99536135 |
| TCGA-DD-AADP | TCGA-LIHC | 46.3941585 | 19.6928526 |
| TCGA-DD-AADQ | TCGA-LIHC | 42.7818521 | 20.0605534 |
| TCGA-DD-AADR | TCGA-LIHC | 60.4807384 | 23.2722648 |
| TCGA-DD-AADS | TCGA-LIHC | 18.3993275 | 9.75219317 |
| TCGA-DD-AADU | TCGA-LIHC | 30.593909 | 9.59946023 |
| TCGA-DD-AADV | TCGA-LIHC | 58.6229647 | 26.65652 |
| TCGA-DD-AADW | TCGA-LIHC | 47.8947003 | 14.3366137 |
| TCGA-DD-AADY | TCGA-LIHC | 28.7240713 | 29.5210763 |
| TCGA-DD-AAE0 | TCGA-LIHC | 50.1914562 | 35.9739399 |
| TCGA-DD-AAE1 | TCGA-LIHC | 27.5615061 | 15.5602394 |
| TCGA-DD-AAE2 | TCGA-LIHC | 51.127075 | 16.5750912 |
| TCGA-DD-AAE3 | TCGA-LIHC | 33.4740055 | 10.7233802 |
| TCGA-DD-AAE4 | TCGA-LIHC | 38.9795051 | 20.9976521 |

| | | | |
|---|---|---|---|
| TCGA-DD-AAE6 | TCGA-LIHC | 62.8593479 | 18.865049 |
| TCGA-DD-AAE7 | TCGA-LIHC | 27.4415075 | 10.2831472 |
| TCGA-DD-AAE9 | TCGA-LIHC | 53.3220788 | 20.6948798 |
| TCGA-DD-AAEA | TCGA-LIHC | 53.8378176 | 25.9854878 |
| TCGA-DD-AAEB | TCGA-LIHC | 27.5871756 | 14.6969269 |
| TCGA-DD-AAED | TCGA-LIHC | 51.0766715 | 17.9025177 |
| TCGA-DD-AAEE | TCGA-LIHC | 43.6590684 | 10.8836975 |
| TCGA-DD-AAEG | TCGA-LIHC | 44.4901366 | 32.2337551 |
| TCGA-DD-AAEH | TCGA-LIHC | 52.7784591 | 7.70823074 |
| TCGA-DD-AAEI | TCGA-LIHC | 31.5044973 | 6.89905061 |
| TCGA-DD-AAEK | TCGA-LIHC | 28.9009836 | 14.7290437 |
| TCGA-DD-AAVP | TCGA-LIHC | 54.6505428 | 9.64998795 |
| TCGA-DD-AAVQ | TCGA-LIHC | 33.5548522 | 20.8192568 |
| TCGA-DD-AAVR | TCGA-LIHC | 45.3867644 | 18.3486673 |
| TCGA-DD-AAVS | TCGA-LIHC | 34.9893715 | 17.8887097 |
| TCGA-DD-AAVU | TCGA-LIHC | 20.9050467 | 9.84004495 |
| TCGA-DD-AAVV | TCGA-LIHC | 65.2673652 | 17.5737648 |
| TCGA-DD-AAVW | TCGA-LIHC | 22.2753868 | 17.3651556 |
| TCGA-DD-AAVX | TCGA-LIHC | 38.2045807 | 14.7554036 |
| TCGA-DD-AAVY | TCGA-LIHC | 44.0674944 | 17.5828502 |
| TCGA-DD-AAVZ | TCGA-LIHC | 29.8906784 | 24.4203067 |
| TCGA-DD-AAW0 | TCGA-LIHC | 39.6391455 | 14.1730347 |
| TCGA-DD-AAW1 | TCGA-LIHC | 41.8335058 | 14.2102561 |
| TCGA-DD-AAW2 | TCGA-LIHC | 40.4436981 | 15.4446088 |
| TCGA-DD-AAW3 | TCGA-LIHC | 50.6814736 | 23.2717917 |
| TCGA-DE-A0XZ | TCGA-THCA | 56.5982267 | 13.7487957 |
| TCGA-DE-A0Y2 | TCGA-THCA | 39.2961412 | 11.6263131 |
| TCGA-DE-A0Y3 | TCGA-THCA | 48.1003166 | 9.49656134 |
| TCGA-DE-A2OL | TCGA-THCA | 56.2723046 | 17.1558794 |
| TCGA-DE-A3KN | TCGA-THCA | 67.0756874 | 14.8966959 |
| TCGA-DE-A4M8 | TCGA-THCA | 30.4122039 | 7.86299287 |

| | | | |
|---|---|---|---|
| TCGA-DE-A4M9 | TCGA-THCA | 63.0648416 | 18.8738051 |
| TCGA-DE-A4MA | TCGA-THCA | 39.4608412 | 17.4150282 |
| TCGA-DE-A4MB | TCGA-THCA | 32.8482126 | 15.8668521 |
| TCGA-DE-A4MC | TCGA-THCA | 38.728986 | 13.9744807 |
| TCGA-DE-A4MD | TCGA-THCA | 54.4601721 | 16.1763373 |
| TCGA-DE-A69J | TCGA-THCA | 67.0901634 | 24.4583635 |
| TCGA-DE-A69K | TCGA-THCA | 66.2089959 | 18.4450905 |
| TCGA-DE-A7U5 | TCGA-THCA | 39.5745155 | 16.8729609 |
| TCGA-DF-A2KN | TCGA-UCEC | 43.0656278 | 35.6599544 |
| TCGA-DF-A2KR | TCGA-UCEC | 35.3496185 | 4.71111264 |
| TCGA-DF-A2KV | TCGA-UCEC | 70.9167775 | 25.5612809 |
| TCGA-DF-A2KY | TCGA-UCEC | 65.7709944 | 18.173333 |
| TCGA-DF-A2KZ | TCGA-UCEC | 41.2875921 | 28.5992009 |
| TCGA-DF-A2L0 | TCGA-UCEC | 20.5470915 | 17.8405782 |
| TCGA-DH-5140 | TCGA-LGG | 57.8830209 | 28.8740161 |
| TCGA-DH-5141 | TCGA-LGG | 15.4394617 | 13.2293161 |
| TCGA-DH-5142 | TCGA-LGG | 40.799857 | 14.224446 |
| TCGA-DH-5143 | TCGA-LGG | 26.3242269 | 13.9478204 |
| TCGA-DH-5144 | TCGA-LGG | 24.056128 | 20.1760876 |
| TCGA-DH-A669 | TCGA-LGG | 45.3126974 | 39.5949487 |
| TCGA-DH-A66B | TCGA-LGG | 51.6046006 | 17.8188463 |
| TCGA-DH-A66D | TCGA-LGG | 34.2107581 | 10.3555885 |
| TCGA-DH-A66F | TCGA-LGG | 17.0834673 | 22.3258613 |
| TCGA-DH-A66G | TCGA-LGG | 31.8162897 | 8.73600984 |
| TCGA-DH-A7UR | TCGA-LGG | 27.5801544 | 24.8836041 |
| TCGA-DH-A7US | TCGA-LGG | 21.8354361 | 22.9278101 |
| TCGA-DH-A7UT | TCGA-LGG | 30.0102146 | 12.7349709 |
| TCGA-DH-A7UU | TCGA-LGG | 36.9576803 | 16.6609048 |
| TCGA-DH-A7UV | TCGA-LGG | 28.0919934 | 9.59913856 |
| TCGA-DI-A1BU | TCGA-UCEC | 88.1470786 | 45.1743488 |
| TCGA-DI-A2QT | TCGA-UCEC | 29.811971 | 15.6038117 |

| | | | |
|---|---|---|---|
| TCGA-DI-A2QY | TCGA-UCEC | 54.4689909 | 15.1827359 |
| TCGA-DJ-A13L | TCGA-THCA | 52.4888989 | 19.105248 |
| TCGA-DJ-A13M | TCGA-THCA | 66.7458835 | 18.8238714 |
| TCGA-DJ-A13O | TCGA-THCA | 52.6872757 | 12.6880466 |
| TCGA-DJ-A13P | TCGA-THCA | 68.0727823 | 17.0018746 |
| TCGA-DJ-A13R | TCGA-THCA | 28.291322 | 6.46806069 |
| TCGA-DJ-A13S | TCGA-THCA | 36.0326168 | 10.4539927 |
| TCGA-DJ-A13T | TCGA-THCA | 71.5332902 | 15.4697409 |
| TCGA-DJ-A13U | TCGA-THCA | 40.320766 | 10.6463309 |
| TCGA-DJ-A13V | TCGA-THCA | 61.167627 | 14.7778248 |
| TCGA-DJ-A13W | TCGA-THCA | 15.9185832 | 2.78667764 |
| TCGA-DJ-A13X | TCGA-THCA | 63.0138072 | 12.7285224 |
| TCGA-DJ-A1QD | TCGA-THCA | 43.1401405 | 11.0943286 |
| TCGA-DJ-A1QE | TCGA-THCA | 43.8629613 | 13.9118999 |
| TCGA-DJ-A1QF | TCGA-THCA | 46.2279243 | 10.686525 |
| TCGA-DJ-A1QG | TCGA-THCA | 45.6914915 | 14.7007638 |
| TCGA-DJ-A1QH | TCGA-THCA | 47.3740226 | 15.7101717 |
| TCGA-DJ-A1QI | TCGA-THCA | 68.2453483 | 21.3153621 |
| TCGA-DJ-A1QL | TCGA-THCA | 58.8751643 | 19.8345477 |
| TCGA-DJ-A1QM | TCGA-THCA | 43.4242897 | 14.4555262 |
| TCGA-DJ-A1QN | TCGA-THCA | 54.6773536 | 11.0852757 |
| TCGA-DJ-A1QO | TCGA-THCA | 37.6960932 | 10.4170594 |
| TCGA-DJ-A1QQ | TCGA-THCA | 49.6129558 | 14.0412421 |
| TCGA-DJ-A2PN | TCGA-THCA | 46.8166858 | 15.0353816 |
| TCGA-DJ-A2PO | TCGA-THCA | 42.6213002 | 13.1799111 |
| TCGA-DJ-A2PP | TCGA-THCA | 42.1315373 | 16.0125358 |
| TCGA-DJ-A2PQ | TCGA-THCA | 73.1728208 | 23.0543603 |
| TCGA-DJ-A2PR | TCGA-THCA | 48.0874652 | 14.3966251 |
| TCGA-DJ-A2PS | TCGA-THCA | 64.1010375 | 19.3665432 |
| TCGA-DJ-A2PT | TCGA-THCA | 47.1697447 | 14.959244 |
| TCGA-DJ-A2PU | TCGA-THCA | 52.4286966 | 15.2041552 |

| | | | |
|---|---|---|---|
| TCGA-DJ-A2PV | TCGA-THCA | 51.3146924 | 14.1336454 |
| TCGA-DJ-A2PW | TCGA-THCA | 54.2115831 | 16.3093919 |
| TCGA-DJ-A2PX | TCGA-THCA | 57.5826672 | 14.2605664 |
| TCGA-DJ-A2PY | TCGA-THCA | 61.2179755 | 21.6061078 |
| TCGA-DJ-A2PZ | TCGA-THCA | 43.8533116 | 10.2355786 |
| TCGA-DJ-A2Q0 | TCGA-THCA | 38.6107645 | 16.0211718 |
| TCGA-DJ-A2Q1 | TCGA-THCA | 39.2846786 | 19.7745771 |
| TCGA-DJ-A2Q2 | TCGA-THCA | 60.2119284 | 14.9289079 |
| TCGA-DJ-A2Q3 | TCGA-THCA | 41.0426883 | 12.6320733 |
| TCGA-DJ-A2Q4 | TCGA-THCA | 53.4482651 | 21.4065922 |
| TCGA-DJ-A2Q5 | TCGA-THCA | 53.8273893 | 14.9159045 |
| TCGA-DJ-A2Q6 | TCGA-THCA | 66.7085396 | 18.0961718 |
| TCGA-DJ-A2Q7 | TCGA-THCA | 72.0381793 | 13.5212972 |
| TCGA-DJ-A2Q9 | TCGA-THCA | 48.4149766 | 16.4909197 |
| TCGA-DJ-A2QA | TCGA-THCA | 67.6411138 | 13.34784 |
| TCGA-DJ-A2QB | TCGA-THCA | 34.0353075 | 10.9021114 |
| TCGA-DJ-A2QC | TCGA-THCA | 78.7108666 | 15.7276405 |
| TCGA-DJ-A3UK | TCGA-THCA | 65.7697759 | 14.6887081 |
| TCGA-DJ-A3UM | TCGA-THCA | 50.0792282 | 12.4527545 |
| TCGA-DJ-A3UN | TCGA-THCA | 63.419822 | 13.9150247 |
| TCGA-DJ-A3UO | TCGA-THCA | 47.5681349 | 13.2724299 |
| TCGA-DJ-A3UP | TCGA-THCA | 55.9528048 | 12.0465557 |
| TCGA-DJ-A3UQ | TCGA-THCA | 51.7732123 | 11.0048197 |
| TCGA-DJ-A3UR | TCGA-THCA | 71.8045935 | 13.3890166 |
| TCGA-DJ-A3US | TCGA-THCA | 61.0917843 | 11.8357677 |
| TCGA-DJ-A3UT | TCGA-THCA | 60.0392285 | 15.7099498 |
| TCGA-DJ-A3UU | TCGA-THCA | 55.8773337 | 15.9536786 |
| TCGA-DJ-A3UV | TCGA-THCA | 65.3768169 | 18.142788 |
| TCGA-DJ-A3UW | TCGA-THCA | 37.2857944 | 11.0358798 |
| TCGA-DJ-A3UX | TCGA-THCA | 58.2322594 | 11.8344553 |
| TCGA-DJ-A3UY | TCGA-THCA | 47.3649518 | 12.651655 |

| | | | |
|---|---|---|---|
| TCGA-DJ-A3UZ | TCGA-THCA | 62.2378806 | 13.9943224 |
| TCGA-DJ-A3V0 | TCGA-THCA | 56.838172 | 15.2716906 |
| TCGA-DJ-A3V2 | TCGA-THCA | 71.4329748 | 18.0450899 |
| TCGA-DJ-A3V3 | TCGA-THCA | 54.4826733 | 13.3381118 |
| TCGA-DJ-A3V4 | TCGA-THCA | 57.3817004 | 15.5538316 |
| TCGA-DJ-A3V5 | TCGA-THCA | 52.3367759 | 15.0595193 |
| TCGA-DJ-A3V6 | TCGA-THCA | 44.5604635 | 14.9104735 |
| TCGA-DJ-A3V7 | TCGA-THCA | 55.7037295 | 11.8674119 |
| TCGA-DJ-A3V8 | TCGA-THCA | 64.1695876 | 17.1575345 |
| TCGA-DJ-A3V9 | TCGA-THCA | 66.3714072 | 15.3451748 |
| TCGA-DJ-A3VA | TCGA-THCA | 58.9079551 | 15.7807803 |
| TCGA-DJ-A3VB | TCGA-THCA | 58.1292206 | 17.2860966 |
| TCGA-DJ-A3VD | TCGA-THCA | 43.3099904 | 15.195571 |
| TCGA-DJ-A3VE | TCGA-THCA | 64.9700819 | 16.7637896 |
| TCGA-DJ-A3VF | TCGA-THCA | 43.1981073 | 14.8188467 |
| TCGA-DJ-A3VG | TCGA-THCA | 74.7189987 | 17.4388158 |
| TCGA-DJ-A3VI | TCGA-THCA | 53.0065308 | 11.6748789 |
| TCGA-DJ-A3VJ | TCGA-THCA | 72.9290353 | 18.424247 |
| TCGA-DJ-A3VK | TCGA-THCA | 76.1377701 | 16.1107152 |
| TCGA-DJ-A3VL | TCGA-THCA | 56.4474081 | 13.9025738 |
| TCGA-DJ-A3VM | TCGA-THCA | 39.9524351 | 8.62579393 |
| TCGA-DJ-A4UL | TCGA-THCA | 68.8961364 | 23.409264 |
| TCGA-DJ-A4UP | TCGA-THCA | 58.4461354 | 16.1850039 |
| TCGA-DJ-A4UQ | TCGA-THCA | 57.4725145 | 15.9538319 |
| TCGA-DJ-A4UR | TCGA-THCA | 47.2058048 | 15.8990559 |
| TCGA-DJ-A4UT | TCGA-THCA | 45.3330239 | 14.381571 |
| TCGA-DJ-A4UW | TCGA-THCA | 74.2690877 | 19.9781739 |
| TCGA-DJ-A4V0 | TCGA-THCA | 44.7908114 | 12.9565279 |
| TCGA-DJ-A4V2 | TCGA-THCA | 46.1560004 | 16.7904931 |
| TCGA-DJ-A4V4 | TCGA-THCA | 51.2854423 | 13.8824509 |
| TCGA-DJ-A4V5 | TCGA-THCA | 60.0461622 | 17.2093728 |

| | | | |
|---|---|---|---|
| TCGA-DK-A1A3 | TCGA-BLCA | 51.6243416 | 27.324403 |
| TCGA-DK-A1A5 | TCGA-BLCA | 108.841039 | 52.8144324 |
| TCGA-DK-A1A6 | TCGA-BLCA | 83.4071012 | 67.5014763 |
| TCGA-DK-A1A7 | TCGA-BLCA | 54.4900072 | 13.9959751 |
| TCGA-DK-A1AA | TCGA-BLCA | 69.8508631 | 20.8509616 |
| TCGA-DK-A1AB | TCGA-BLCA | 69.6878733 | 21.4644246 |
| TCGA-DK-A1AC | TCGA-BLCA | 94.0661994 | 33.4508796 |
| TCGA-DK-A1AD | TCGA-BLCA | 76.577113 | 22.6205935 |
| TCGA-DK-A1AE | TCGA-BLCA | 71.2395075 | 22.5788808 |
| TCGA-DK-A1AF | TCGA-BLCA | 41.7893063 | 28.5533532 |
| TCGA-DK-A1AG | TCGA-BLCA | 103.401289 | 22.8461623 |
| TCGA-DK-A2HX | TCGA-BLCA | 70.622694 | 27.4216914 |
| TCGA-DK-A2I1 | TCGA-BLCA | 76.11866 | 22.8851354 |
| TCGA-DK-A2I2 | TCGA-BLCA | 115.817731 | 39.809962 |
| TCGA-DK-A2I4 | TCGA-BLCA | 85.6499284 | 22.6491818 |
| TCGA-DK-A2I6 | TCGA-BLCA | 62.4748411 | 25.6667582 |
| TCGA-DK-A3IK | TCGA-BLCA | 54.1914628 | 18.6621242 |
| TCGA-DK-A3IL | TCGA-BLCA | 67.997696 | 18.7781663 |
| TCGA-DK-A3IM | TCGA-BLCA | 113.088443 | 57.7939821 |
| TCGA-DK-A3IN | TCGA-BLCA | 64.4200703 | 21.6713308 |
| TCGA-DK-A3IQ | TCGA-BLCA | 40.4549179 | 25.5272667 |
| TCGA-DK-A3IS | TCGA-BLCA | 102.327984 | 51.1609582 |
| TCGA-DK-A3IT | TCGA-BLCA | 55.3806655 | 36.243381 |
| TCGA-DK-A3IU | TCGA-BLCA | 69.966516 | 22.5035098 |
| TCGA-DK-A3IV | TCGA-BLCA | 93.2330138 | 42.3499299 |
| TCGA-DK-A3WV | TCGA-BLCA | 86.7115991 | 28.5308558 |
| TCGA-DK-A3WX | TCGA-BLCA | 73.9209916 | 34.618432 |
| TCGA-DK-A3WY | TCGA-BLCA | 66.6618229 | 17.2294524 |
| TCGA-DK-A3X1 | TCGA-BLCA | 55.4841522 | 29.6154851 |
| TCGA-DK-A3X2 | TCGA-BLCA | 112.917618 | 24.9678591 |
| TCGA-DK-A6AV | TCGA-BLCA | 156.673522 | 42.9743564 |

| | | | |
|---|---|---|---|
| TCGA-DK-A6AW | TCGA-BLCA | 53.9959925 | 14.2692038 |
| TCGA-DK-A6B0 | TCGA-BLCA | 60.3837778 | 17.8182432 |
| TCGA-DK-A6B1 | TCGA-BLCA | 66.7254437 | 31.3790385 |
| TCGA-DK-A6B2 | TCGA-BLCA | 57.0805406 | 39.4617365 |
| TCGA-DK-A6B5 | TCGA-BLCA | 99.3944048 | 38.0539798 |
| TCGA-DK-A6B6 | TCGA-BLCA | 53.514016 | 17.6788887 |
| TCGA-DK-AA6L | TCGA-BLCA | 159.514325 | 32.0617697 |
| TCGA-DK-AA6M | TCGA-BLCA | 76.1023729 | 29.9412072 |
| TCGA-DK-AA6P | TCGA-BLCA | 58.0487813 | 26.2502132 |
| TCGA-DK-AA6Q | TCGA-BLCA | 78.8163332 | 36.7944043 |
| TCGA-DK-AA6R | TCGA-BLCA | 118.308108 | 40.0306131 |
| TCGA-DK-AA6S | TCGA-BLCA | 49.1584631 | 27.8350498 |
| TCGA-DK-AA6T | TCGA-BLCA | 109.882192 | 61.651758 |
| TCGA-DK-AA6U | TCGA-BLCA | 86.6194885 | 68.5972815 |
| TCGA-DK-AA6W | TCGA-BLCA | 102.047897 | 42.756909 |
| TCGA-DK-AA6X | TCGA-BLCA | 75.4646527 | 25.4899606 |
| TCGA-DK-AA71 | TCGA-BLCA | 70.1955246 | 21.1814541 |
| TCGA-DK-AA74 | TCGA-BLCA | 68.7021837 | 30.9964213 |
| TCGA-DK-AA75 | TCGA-BLCA | 139.135803 | 54.6865091 |
| TCGA-DK-AA76 | TCGA-BLCA | 71.6604047 | 25.2151788 |
| TCGA-DK-AA77 | TCGA-BLCA | 90.2866181 | 32.4910207 |
| TCGA-DM-A0X9 | TCGA-COAD | 72.7389568 | 37.5517456 |
| TCGA-DM-A0XD | TCGA-COAD | 75.7270954 | 25.3245581 |
| TCGA-DM-A0XF | TCGA-COAD | 68.0806585 | 60.1179464 |
| TCGA-DM-A1D0 | TCGA-COAD | 88.3658842 | 26.2543667 |
| TCGA-DM-A1D4 | TCGA-COAD | 43.5178489 | 32.8463761 |
| TCGA-DM-A1D6 | TCGA-COAD | 45.7137581 | 41.5788906 |
| TCGA-DM-A1D7 | TCGA-COAD | 90.1900843 | 13.7387639 |
| TCGA-DM-A1D8 | TCGA-COAD | 68.8453705 | 18.0869727 |
| TCGA-DM-A1D9 | TCGA-COAD | 87.5060142 | 30.4646375 |
| TCGA-DM-A1DA | TCGA-COAD | 72.5063038 | 24.6920941 |

| | | | |
|---|---|---|---|
| TCGA-DM-A1DB | TCGA-COAD | 72.211974 | 64.3585241 |
| TCGA-DM-A1HA | TCGA-COAD | 109.487841 | 31.1110419 |
| TCGA-DM-A280 | TCGA-COAD | 60.0032965 | 26.4487806 |
| TCGA-DM-A282 | TCGA-COAD | 106.762118 | 25.2829438 |
| TCGA-DM-A285 | TCGA-COAD | 83.9484784 | 31.4834016 |
| TCGA-DM-A288 | TCGA-COAD | 50.811802 | 18.0914409 |
| TCGA-DM-A28A | TCGA-COAD | 62.8871971 | 45.2793974 |
| TCGA-DM-A28C | TCGA-COAD | 47.2841337 | 13.9677333 |
| TCGA-DM-A28E | TCGA-COAD | 66.4508131 | 21.5399795 |
| TCGA-DM-A28F | TCGA-COAD | 66.5154295 | 30.8615747 |
| TCGA-DM-A28G | TCGA-COAD | 48.5726559 | 9.88644254 |
| TCGA-DM-A28H | TCGA-COAD | 150.098671 | 52.424813 |
| TCGA-DM-A28K | TCGA-COAD | 61.1637091 | 26.7693186 |
| TCGA-DM-A28M | TCGA-COAD | 84.5828992 | 21.3782975 |
| TCGA-DO-A1JZ | TCGA-THCA | 50.6521046 | 15.147279 |
| TCGA-DO-A1K0 | TCGA-THCA | 43.3012832 | 12.9109376 |
| TCGA-DO-A2HM | TCGA-THCA | 38.3432155 | 15.2044535 |
| TCGA-DU-5847 | TCGA-LGG | 68.2182656 | 21.0268508 |
| TCGA-DU-5849 | TCGA-LGG | 18.0830848 | 11.5981131 |
| TCGA-DU-5852 | TCGA-LGG | 38.7129084 | 14.7906792 |
| TCGA-DU-5853 | TCGA-LGG | 27.2882948 | 11.012934 |
| TCGA-DU-5854 | TCGA-LGG | 33.2341376 | 12.3724731 |
| TCGA-DU-5855 | TCGA-LGG | 39.1446363 | 11.2040269 |
| TCGA-DU-5870 | TCGA-LGG | 21.4318443 | 25.8438155 |
| TCGA-DU-5871 | TCGA-LGG | 28.6580418 | 8.69665208 |
| TCGA-DU-5872 | TCGA-LGG | 23.4241582 | 7.85537728 |
| TCGA-DU-5874 | TCGA-LGG | 22.7457904 | 20.1733808 |
| TCGA-DU-6392 | TCGA-LGG | 34.089941 | 12.1823517 |
| TCGA-DU-6393 | TCGA-LGG | 20.0752697 | 17.8134973 |
| TCGA-DU-6394 | TCGA-LGG | 21.6440222 | 23.0890368 |
| TCGA-DU-6395 | TCGA-LGG | 28.2830848 | 14.6236783 |

| | | | |
|---|---|---|---|
| TCGA-DU-6396 | TCGA-LGG | 33.1371487 | 13.2830894 |
| TCGA-DU-6397 | TCGA-LGG | 18.1322352 | 17.1200595 |
| TCGA-DU-6399 | TCGA-LGG | 34.852514 | 12.4272243 |
| TCGA-DU-6400 | TCGA-LGG | 24.158509 | 20.7090756 |
| TCGA-DU-6401 | TCGA-LGG | 22.5973516 | 6.05942991 |
| TCGA-DU-6402 | TCGA-LGG | 60.0952003 | 14.9091016 |
| TCGA-DU-6403 | TCGA-LGG | 42.4346655 | 13.2616506 |
| TCGA-DU-6404 | TCGA-LGG | 62.9151662 | 37.5706563 |
| TCGA-DU-6405 | TCGA-LGG | 48.1046078 | 16.1402104 |
| TCGA-DU-6406 | TCGA-LGG | 48.3002808 | 20.5133327 |
| TCGA-DU-6407 | TCGA-LGG | 29.0482015 | 8.02900172 |
| TCGA-DU-6408 | TCGA-LGG | 31.6951502 | 15.5001428 |
| TCGA-DU-6410 | TCGA-LGG | 15.2190003 | 15.6964091 |
| TCGA-DU-6542 | TCGA-LGG | 36.7346452 | 10.2100015 |
| TCGA-DU-7006 | TCGA-LGG | 55.452836 | 23.8830551 |
| TCGA-DU-7007 | TCGA-LGG | 45.0809611 | 98.5849037 |
| TCGA-DU-7008 | TCGA-LGG | 33.7056152 | 16.9745471 |
| TCGA-DU-7009 | TCGA-LGG | 18.8839732 | 19.6514726 |
| TCGA-DU-7010 | TCGA-LGG | 71.052954 | 20.7308244 |
| TCGA-DU-7011 | TCGA-LGG | 23.4095583 | 7.9965042 |
| TCGA-DU-7012 | TCGA-LGG | 47.4172574 | 14.1666484 |
| TCGA-DU-7013 | TCGA-LGG | 45.3705404 | 17.4093892 |
| TCGA-DU-7014 | TCGA-LGG | 24.1835424 | 11.2548345 |
| TCGA-DU-7015 | TCGA-LGG | 30.7030351 | 12.0809508 |
| TCGA-DU-7018 | TCGA-LGG | 14.6453907 | 21.2305831 |
| TCGA-DU-7019 | TCGA-LGG | 31.6357992 | 11.9772425 |
| TCGA-DU-7290 | TCGA-LGG | 80.5558534 | 19.1450647 |
| TCGA-DU-7292 | TCGA-LGG | 33.1771737 | 15.5598373 |
| TCGA-DU-7294 | TCGA-LGG | 18.0919744 | 21.6535934 |
| TCGA-DU-7298 | TCGA-LGG | 43.1280562 | 21.5087134 |
| TCGA-DU-7299 | TCGA-LGG | 35.817709 | 11.9718985 |

| | | | |
|---|---|---|---|
| TCGA-DU-7300 | TCGA-LGG | 21.4777734 | 16.6577954 |
| TCGA-DU-7301 | TCGA-LGG | 29.0376654 | 11.2965391 |
| TCGA-DU-7302 | TCGA-LGG | 21.0295343 | 19.4080258 |
| TCGA-DU-7304 | TCGA-LGG | 28.3568238 | 10.6911558 |
| TCGA-DU-7306 | TCGA-LGG | 30.0038863 | 7.68026127 |
| TCGA-DU-7309 | TCGA-LGG | 19.949619 | 6.28486341 |
| TCGA-DU-8158 | TCGA-LGG | 38.7911246 | 14.9644356 |
| TCGA-DU-8161 | TCGA-LGG | 45.9523646 | 19.4428125 |
| TCGA-DU-8162 | TCGA-LGG | 27.7083322 | 11.6413937 |
| TCGA-DU-8163 | TCGA-LGG | 30.0333681 | 10.7330651 |
| TCGA-DU-8164 | TCGA-LGG | 20.2190381 | 17.9079361 |
| TCGA-DU-8165 | TCGA-LGG | 99.6054801 | 32.5256973 |
| TCGA-DU-8166 | TCGA-LGG | 38.460255 | 10.9438977 |
| TCGA-DU-8167 | TCGA-LGG | 37.582196 | 14.3635125 |
| TCGA-DU-8168 | TCGA-LGG | 21.5017357 | 27.9744689 |
| TCGA-DU-A5TP | TCGA-LGG | 66.324411 | 23.7323049 |
| TCGA-DU-A5TR | TCGA-LGG | 26.3771909 | 12.871046 |
| TCGA-DU-A5TS | TCGA-LGG | 20.0388254 | 13.4419328 |
| TCGA-DU-A5TT | TCGA-LGG | 60.8342999 | 18.0224354 |
| TCGA-DU-A5TU | TCGA-LGG | 43.2637769 | 37.0218412 |
| TCGA-DU-A5TW | TCGA-LGG | 38.3188076 | 12.4075736 |
| TCGA-DU-A5TY | TCGA-LGG | 49.9813202 | 24.9396941 |
| TCGA-DU-A6S2 | TCGA-LGG | 21.2683111 | 12.0377535 |
| TCGA-DU-A6S3 | TCGA-LGG | 16.5104973 | 11.9635656 |
| TCGA-DU-A6S6 | TCGA-LGG | 19.1728099 | 12.319475 |
| TCGA-DU-A6S7 | TCGA-LGG | 36.7904107 | 10.081031 |
| TCGA-DU-A6S8 | TCGA-LGG | 25.6095972 | 33.2845566 |
| TCGA-DU-A76K | TCGA-LGG | 26.2817151 | 12.1002025 |
| TCGA-DU-A76L | TCGA-LGG | 71.3749987 | 30.8120082 |
| TCGA-DU-A76O | TCGA-LGG | 32.4674503 | 12.0726313 |
| TCGA-DU-A76R | TCGA-LGG | 22.9320919 | 17.5092835 |

| | | | |
|---|---|---|---|
| TCGA-DU-A7T6 | TCGA-LGG | 25.714968 | 32.2292869 |
| TCGA-DU-A7T8 | TCGA-LGG | 32.4710741 | 15.742124 |
| TCGA-DU-A7TA | TCGA-LGG | 43.6599704 | 23.6368763 |
| TCGA-DU-A7TB | TCGA-LGG | 31.5898555 | 17.0808695 |
| TCGA-DU-A7TC | TCGA-LGG | 31.9626992 | 12.6507454 |
| TCGA-DU-A7TG | TCGA-LGG | 28.9691045 | 11.6785701 |
| TCGA-DU-A7TI | TCGA-LGG | 39.8334501 | 25.2546668 |
| TCGA-DU-A7TJ | TCGA-LGG | 46.6413199 | 25.1557483 |
| TCGA-DV-5565 | TCGA-KIRC | 66.582061 | 19.721205 |
| TCGA-DV-5566 | TCGA-KIRC | 53.7503986 | 15.0081123 |
| TCGA-DV-5567 | TCGA-KIRC | 46.7128037 | 12.6653491 |
| TCGA-DV-5568 | TCGA-KIRC | 51.7266947 | 17.3419617 |
| TCGA-DV-5569 | TCGA-KIRC | 50.0807653 | 14.1605697 |
| TCGA-DV-5573 | TCGA-KIRC | 63.9257735 | 25.8267956 |
| TCGA-DV-5574 | TCGA-KIRC | 43.9780827 | 12.8412446 |
| TCGA-DV-5575 | TCGA-KIRC | 49.7844701 | 16.7128704 |
| TCGA-DV-5576 | TCGA-KIRC | 50.1567142 | 12.0146195 |
| TCGA-DV-A4VX | TCGA-KIRC | 43.5485472 | 26.7360781 |
| TCGA-DV-A4VZ | TCGA-KIRC | 51.0643564 | 12.1972379 |
| TCGA-DV-A4W0 | TCGA-KIRC | 54.0957662 | 13.1449804 |
| TCGA-DW-5560 | TCGA-KIRP | 26.1704205 | 10.3872202 |
| TCGA-DW-5561 | TCGA-KIRP | 44.7504296 | 10.3851164 |
| TCGA-DW-7834 | TCGA-KIRP | 172.154662 | 14.2043518 |
| TCGA-DW-7836 | TCGA-KIRP | 65.9197739 | 17.9876165 |
| TCGA-DW-7837 | TCGA-KIRP | 42.5132606 | 14.8594433 |
| TCGA-DW-7838 | TCGA-KIRP | 15.2960636 | 7.00986918 |
| TCGA-DW-7839 | TCGA-KIRP | 86.5856915 | 17.7840604 |
| TCGA-DW-7840 | TCGA-KIRP | 35.0590949 | 12.3533925 |
| TCGA-DW-7841 | TCGA-KIRP | 46.6009557 | 7.47554774 |
| TCGA-DW-7842 | TCGA-KIRP | 8.08040364 | 2.20177869 |
| TCGA-DW-7963 | TCGA-KIRP | 13.3891061 | 8.68662857 |

| | | | |
|---|---|---|---|
| TCGA-DZ-6132 | TCGA-KIRP | 22.1804333 | 10.0321721 |
| TCGA-DZ-6133 | TCGA-KIRP | 26.0575029 | 12.3341736 |
| TCGA-DZ-6134 | TCGA-KIRP | 16.819519 | 7.2104742 |
| TCGA-DZ-6135 | TCGA-KIRP | 25.6203994 | 7.07093558 |
| TCGA-E1-5303 | TCGA-LGG | 28.1110304 | 10.061596 |
| TCGA-E1-5304 | TCGA-LGG | 82.332642 | 25.2706047 |
| TCGA-E1-5307 | TCGA-LGG | 31.5201286 | 11.1678982 |
| TCGA-E1-5311 | TCGA-LGG | 20.8155499 | 15.8464133 |
| TCGA-E1-5318 | TCGA-LGG | 16.9158504 | 14.4997536 |
| TCGA-E1-5319 | TCGA-LGG | 19.6715103 | 18.303681 |
| TCGA-E1-5322 | TCGA-LGG | 43.5091887 | 12.6559558 |
| TCGA-E1-A7YD | TCGA-LGG | 69.2108493 | 34.4916782 |
| TCGA-E1-A7YE | TCGA-LGG | 47.4129407 | 15.7009333 |
| TCGA-E1-A7YH | TCGA-LGG | 35.850172 | 22.6803502 |
| TCGA-E1-A7YI | TCGA-LGG | 75.7250825 | 24.7311135 |
| TCGA-E1-A7YJ | TCGA-LGG | 71.864852 | 29.5017688 |
| TCGA-E1-A7YK | TCGA-LGG | 37.4170507 | 16.8186854 |
| TCGA-E1-A7YL | TCGA-LGG | 44.4932439 | 26.5283458 |
| TCGA-E1-A7YM | TCGA-LGG | 42.9350031 | 25.4242826 |
| TCGA-E1-A7YN | TCGA-LGG | 51.2252075 | 22.198704 |
| TCGA-E1-A7YO | TCGA-LGG | 19.8339769 | 17.7734771 |
| TCGA-E1-A7YQ | TCGA-LGG | 66.9305061 | 40.0492323 |
| TCGA-E1-A7YS | TCGA-LGG | 23.4143985 | 21.4621026 |
| TCGA-E1-A7YU | TCGA-LGG | 32.9460003 | 9.09670792 |
| TCGA-E1-A7YV | TCGA-LGG | 43.8648004 | 15.8000727 |
| TCGA-E1-A7YW | TCGA-LGG | 38.4290058 | 15.383023 |
| TCGA-E1-A7Z2 | TCGA-LGG | 41.3306061 | 19.0679298 |
| TCGA-E1-A7Z3 | TCGA-LGG | 28.8475554 | 8.18566533 |
| TCGA-E1-A7Z4 | TCGA-LGG | 45.7580524 | 16.4257664 |
| TCGA-E1-A7Z6 | TCGA-LGG | 30.1763714 | 13.5919913 |
| TCGA-E2-A105 | TCGA-BRCA | 55.644333 | 31.1991424 |

| | | | |
|---|---|---|---|
| TCGA-E2-A106 | TCGA-BRCA | 28.6606806 | 21.7664163 |
| TCGA-E2-A107 | TCGA-BRCA | 51.9558556 | 56.1743745 |
| TCGA-E2-A108 | TCGA-BRCA | 51.2438589 | 22.5167916 |
| TCGA-E2-A109 | TCGA-BRCA | 44.6635582 | 46.2929669 |
| TCGA-E2-A10A | TCGA-BRCA | 78.4816813 | 28.6655772 |
| TCGA-E2-A10B | TCGA-BRCA | 53.0813541 | 61.1788272 |
| TCGA-E2-A10C | TCGA-BRCA | 58.3340553 | 69.5423334 |
| TCGA-E2-A10E | TCGA-BRCA | 36.7573972 | 35.1583826 |
| TCGA-E2-A10F | TCGA-BRCA | 49.2067889 | 33.8763575 |
| TCGA-E2-A14N | TCGA-BRCA | 186.989446 | 79.5215957 |
| TCGA-E2-A14O | TCGA-BRCA | 44.2741523 | 42.5271439 |
| TCGA-E2-A14P | TCGA-BRCA | 102.730996 | 30.8161587 |
| TCGA-E2-A14Q | TCGA-BRCA | 62.6686654 | 17.3472319 |
| TCGA-E2-A14R | TCGA-BRCA | 79.3803646 | 21.5320176 |
| TCGA-E2-A14T | TCGA-BRCA | 23.7068953 | 21.233503 |
| TCGA-E2-A14U | TCGA-BRCA | 50.4591853 | 12.6008129 |
| TCGA-E2-A14V | TCGA-BRCA | 38.9539901 | 55.7406656 |
| TCGA-E2-A14W | TCGA-BRCA | 66.9011083 | 29.4700208 |
| TCGA-E2-A14X | TCGA-BRCA | 46.6356553 | 26.7200919 |
| TCGA-E2-A14Y | TCGA-BRCA | 46.9083546 | 44.9785483 |
| TCGA-E2-A14Z | TCGA-BRCA | 41.4538948 | 19.5802513 |
| TCGA-E2-A150 | TCGA-BRCA | 78.7564933 | 48.4836416 |
| TCGA-E2-A152 | TCGA-BRCA | 56.4846918 | 39.3459647 |
| TCGA-E2-A153 | TCGA-BRCA | 38.6351013 | 19.92845 |
| TCGA-E2-A154 | TCGA-BRCA | 32.0225982 | 26.2837797 |
| TCGA-E2-A155 | TCGA-BRCA | 38.7124886 | 16.7205742 |
| TCGA-E2-A156 | TCGA-BRCA | 53.9577731 | 27.076167 |
| TCGA-E2-A158 | TCGA-BRCA | 72.7466027 | 26.44879 |
| TCGA-E2-A159 | TCGA-BRCA | 101.121738 | 29.0795488 |
| TCGA-E2-A15A | TCGA-BRCA | 33.079693 | 45.2561187 |
| TCGA-E2-A15C | TCGA-BRCA | 36.0708264 | 17.7901864 |

| | | | |
|---|---|---|---|
| TCGA-E2-A15D | TCGA-BRCA | 47.155618 | 23.7297046 |
| TCGA-E2-A15E | TCGA-BRCA | 59.6025265 | 20.4882856 |
| TCGA-E2-A15F | TCGA-BRCA | 41.8039149 | 33.4434239 |
| TCGA-E2-A15G | TCGA-BRCA | 58.9178871 | 35.049045 |
| TCGA-E2-A15H | TCGA-BRCA | 61.2169776 | 41.636972 |
| TCGA-E2-A15I | TCGA-BRCA | 51.838766 | 22.3314533 |
| TCGA-E2-A15J | TCGA-BRCA | 40.0650931 | 22.8519407 |
| TCGA-E2-A15K | TCGA-BRCA | 43.8292486 | 15.1220905 |
| TCGA-E2-A15L | TCGA-BRCA | 50.5785106 | 30.2794903 |
| TCGA-E2-A15M | TCGA-BRCA | 58.2030795 | 40.9983521 |
| TCGA-E2-A15O | TCGA-BRCA | 80.3671491 | 56.2043601 |
| TCGA-E2-A15P | TCGA-BRCA | 55.8367775 | 18.6777736 |
| TCGA-E2-A15R | TCGA-BRCA | 26.1722578 | 38.8853939 |
| TCGA-E2-A15S | TCGA-BRCA | 70.5842746 | 34.8472984 |
| TCGA-E2-A15T | TCGA-BRCA | 39.939519 | 18.7905826 |
| TCGA-E2-A1AZ | TCGA-BRCA | 111.656135 | 128.593811 |
| TCGA-E2-A1B0 | TCGA-BRCA | 38.6794226 | 29.4934007 |
| TCGA-E2-A1B1 | TCGA-BRCA | 47.5669037 | 26.9044147 |
| TCGA-E2-A1B4 | TCGA-BRCA | 71.648496 | 16.1960895 |
| TCGA-E2-A1B5 | TCGA-BRCA | 46.0399157 | 17.6457694 |
| TCGA-E2-A1B6 | TCGA-BRCA | 120.268221 | 73.268733 |
| TCGA-E2-A1BC | TCGA-BRCA | 37.3320845 | 13.5337166 |
| TCGA-E2-A1BD | TCGA-BRCA | 65.4278673 | 27.1930324 |
| TCGA-E2-A1IE | TCGA-BRCA | 58.5640739 | 29.8629095 |
| TCGA-E2-A1IF | TCGA-BRCA | 57.9351518 | 34.5794579 |
| TCGA-E2-A1IG | TCGA-BRCA | 39.8969269 | 16.0194986 |
| TCGA-E2-A1IH | TCGA-BRCA | 31.9269774 | 39.3789357 |
| TCGA-E2-A1II | TCGA-BRCA | 138.944649 | 71.7280679 |
| TCGA-E2-A1IJ | TCGA-BRCA | 39.5523218 | 23.1934552 |
| TCGA-E2-A1IK | TCGA-BRCA | 49.6362953 | 27.4773125 |
| TCGA-E2-A1IL | TCGA-BRCA | 48.8399876 | 24.7331109 |

| | | | |
|---|---|---|---|
| TCGA-E2-A1IN | TCGA-BRCA | 57.7303363 | 38.2417857 |
| TCGA-E2-A1IO | TCGA-BRCA | 50.3398917 | 27.2475807 |
| TCGA-E2-A1IU | TCGA-BRCA | 51.4340279 | 22.2588931 |
| TCGA-E2-A1L6 | TCGA-BRCA | 51.2750973 | 28.1346467 |
| TCGA-E2-A1L7 | TCGA-BRCA | 52.994835 | 18.738895 |
| TCGA-E2-A1L8 | TCGA-BRCA | 45.1432798 | 29.236785 |
| TCGA-E2-A1L9 | TCGA-BRCA | 61.568602 | 33.2237026 |
| TCGA-E2-A1LA | TCGA-BRCA | 64.119367 | 31.8344417 |
| TCGA-E2-A1LB | TCGA-BRCA | 36.2784792 | 14.2313777 |
| TCGA-E2-A1LE | TCGA-BRCA | 59.3832836 | 32.9319234 |
| TCGA-E2-A1LG | TCGA-BRCA | 44.0139569 | 40.2993851 |
| TCGA-E2-A1LH | TCGA-BRCA | 52.2728625 | 17.4266249 |
| TCGA-E2-A1LI | TCGA-BRCA | 194.963456 | 76.1340029 |
| TCGA-E2-A1LK | TCGA-BRCA | 68.9791902 | 26.2045766 |
| TCGA-E2-A1LL | TCGA-BRCA | 61.7122654 | 53.4774912 |
| TCGA-E2-A1LS | TCGA-BRCA | 26.86431 | 33.5212105 |
| TCGA-E2-A2P5 | TCGA-BRCA | 40.9917584 | 54.092532 |
| TCGA-E2-A2P6 | TCGA-BRCA | 55.2415463 | 36.9354809 |
| TCGA-E2-A3DX | TCGA-BRCA | 35.6067599 | 14.6793092 |
| TCGA-E2-A56Z | TCGA-BRCA | 65.9341639 | 51.7219808 |
| TCGA-E2-A570 | TCGA-BRCA | 66.5254332 | 41.898563 |
| TCGA-E2-A572 | TCGA-BRCA | 43.5076144 | 26.3351098 |
| TCGA-E2-A573 | TCGA-BRCA | 126.074511 | 65.4281282 |
| TCGA-E2-A574 | TCGA-BRCA | 85.1560438 | 52.6800826 |
| TCGA-E2-A576 | TCGA-BRCA | 29.4619337 | 41.4147101 |
| TCGA-E2-A9RU | TCGA-BRCA | 57.5832804 | 37.7725818 |
| TCGA-E3-A3DY | TCGA-THCA | 66.664513 | 15.2080138 |
| TCGA-E3-A3DZ | TCGA-THCA | 83.1072233 | 22.7332304 |
| TCGA-E3-A3E0 | TCGA-THCA | 76.5241637 | 16.6921086 |
| TCGA-E3-A3E1 | TCGA-THCA | 55.9837856 | 17.9522412 |
| TCGA-E3-A3E2 | TCGA-THCA | 90.7716176 | 17.7488743 |

| | | | |
|---|---|---|---|
| TCGA-E3-A3E3 | TCGA-THCA | 55.9798572 | 13.6896376 |
| TCGA-E3-A3E5 | TCGA-THCA | 72.8304468 | 15.3849003 |
| TCGA-E5-A2PC | TCGA-BLCA | 65.8236577 | 29.1487997 |
| TCGA-E5-A4TZ | TCGA-BLCA | 66.0600713 | 39.5485402 |
| TCGA-E5-A4U1 | TCGA-BLCA | 59.4457985 | 35.4829487 |
| TCGA-E6-A2P8 | TCGA-UCEC | 61.8395564 | 29.8821632 |
| TCGA-E6-A2P9 | TCGA-UCEC | 55.5823999 | 23.7729992 |
| TCGA-E6-A8L9 | TCGA-UCEC | 79.2720586 | 6.00876031 |
| TCGA-E7-A3X6 | TCGA-BLCA | 87.888314 | 22.1313588 |
| TCGA-E7-A3Y1 | TCGA-BLCA | 50.4538827 | 11.4138314 |
| TCGA-E7-A4IJ | TCGA-BLCA | 70.6353267 | 58.521223 |
| TCGA-E7-A4XJ | TCGA-BLCA | 32.5105009 | 6.94296315 |
| TCGA-E7-A519 | TCGA-BLCA | 20.7742307 | 7.73941183 |
| TCGA-E7-A541 | TCGA-BLCA | 86.0200054 | 44.9265746 |
| TCGA-E7-A5KE | TCGA-BLCA | 109.171654 | 33.8021473 |
| TCGA-E7-A5KF | TCGA-BLCA | 66.9831017 | 20.9490989 |
| TCGA-E7-A677 | TCGA-BLCA | 70.0804113 | 19.8311777 |
| TCGA-E7-A678 | TCGA-BLCA | 46.5223688 | 11.1989785 |
| TCGA-E7-A6MD | TCGA-BLCA | 82.6225579 | 29.6968179 |
| TCGA-E7-A6ME | TCGA-BLCA | 111.325861 | 39.9481645 |
| TCGA-E7-A6MF | TCGA-BLCA | 51.2164198 | 11.7434449 |
| TCGA-E7-A7DU | TCGA-BLCA | 91.6874895 | 15.7026682 |
| TCGA-E7-A7DV | TCGA-BLCA | 105.116354 | 74.4760077 |
| TCGA-E7-A7PW | TCGA-BLCA | 98.6406268 | 31.4545197 |
| TCGA-E7-A7XN | TCGA-BLCA | 92.2656466 | 36.1833567 |
| TCGA-E7-A85H | TCGA-BLCA | 79.4758901 | 37.1592464 |
| TCGA-E7-A8O7 | TCGA-BLCA | 60.9598871 | 21.6443371 |
| TCGA-E7-A8O8 | TCGA-BLCA | 80.3232869 | 19.4272704 |
| TCGA-E7-A97P | TCGA-BLCA | 100.555095 | 34.01421 |
| TCGA-E7-A97Q | TCGA-BLCA | 62.9955491 | 13.9115491 |
| TCGA-E8-A242 | TCGA-THCA | 34.3349942 | 11.1692137 |

| | | | |
|---|---|---|---|
| TCGA-E8-A2EA | TCGA-THCA | 45.7903263 | 13.4707335 |
| TCGA-E8-A2JQ | TCGA-THCA | 30.4186351 | 14.2270669 |
| TCGA-E8-A3X7 | TCGA-THCA | 49.3657676 | 10.9979951 |
| TCGA-E8-A413 | TCGA-THCA | 63.1317988 | 15.8895092 |
| TCGA-E8-A414 | TCGA-THCA | 55.2647724 | 15.0519105 |
| TCGA-E8-A415 | TCGA-THCA | 58.569542 | 14.8413097 |
| TCGA-E8-A416 | TCGA-THCA | 75.7407278 | 15.8343184 |
| TCGA-E8-A417 | TCGA-THCA | 47.4210612 | 10.153782 |
| TCGA-E8-A418 | TCGA-THCA | 57.8148012 | 16.129265 |
| TCGA-E8-A419 | TCGA-THCA | 38.4652351 | 12.1931926 |
| TCGA-E8-A432 | TCGA-THCA | 57.6473311 | 24.1882321 |
| TCGA-E8-A433 | TCGA-THCA | 65.5346575 | 14.9760711 |
| TCGA-E8-A434 | TCGA-THCA | 70.1772042 | 17.6619481 |
| TCGA-E8-A436 | TCGA-THCA | 48.3196688 | 12.1067948 |
| TCGA-E8-A437 | TCGA-THCA | 55.6992634 | 12.5864754 |
| TCGA-E8-A438 | TCGA-THCA | 48.4991308 | 10.8906059 |
| TCGA-E8-A44K | TCGA-THCA | 38.532536 | 9.67062449 |
| TCGA-E8-A44M | TCGA-THCA | 44.3732382 | 10.4420724 |
| TCGA-E9-A1N3 | TCGA-BRCA | 35.134762 | 22.1266884 |
| TCGA-E9-A1N4 | TCGA-BRCA | 55.7448206 | 33.733465 |
| TCGA-E9-A1N5 | TCGA-BRCA | 35.8795559 | 16.7286644 |
| TCGA-E9-A1N6 | TCGA-BRCA | 39.6095928 | 16.0150163 |
| TCGA-E9-A1N8 | TCGA-BRCA | 113.056478 | 271.018524 |
| TCGA-E9-A1N9 | TCGA-BRCA | 66.3159094 | 71.6333402 |
| TCGA-E9-A1NA | TCGA-BRCA | 53.2224401 | 36.1255457 |
| TCGA-E9-A1NC | TCGA-BRCA | 71.5756099 | 23.2381268 |
| TCGA-E9-A1ND | TCGA-BRCA | 90.975759 | 23.7378252 |
| TCGA-E9-A1NE | TCGA-BRCA | 51.4484846 | 39.2253222 |
| TCGA-E9-A1NF | TCGA-BRCA | 34.5872596 | 11.769215 |
| TCGA-E9-A1NG | TCGA-BRCA | 41.7364451 | 20.5259323 |
| TCGA-E9-A1NH | TCGA-BRCA | 45.1606853 | 21.5519593 |

| | | | |
|---|---|---|---|
| TCGA-E9-A1NI | TCGA-BRCA | 68.6089433 | 52.4611343 |
| TCGA-E9-A1QZ | TCGA-BRCA | 55.5791631 | 37.9322926 |
| TCGA-E9-A1R0 | TCGA-BRCA | 47.5076541 | 22.9529553 |
| TCGA-E9-A1R2 | TCGA-BRCA | 62.2527504 | 28.8898767 |
| TCGA-E9-A1R3 | TCGA-BRCA | 52.2379603 | 34.7403225 |
| TCGA-E9-A1R4 | TCGA-BRCA | 35.3678262 | 12.8947278 |
| TCGA-E9-A1R5 | TCGA-BRCA | 28.5997686 | 29.0023989 |
| TCGA-E9-A1R6 | TCGA-BRCA | 45.8049684 | 40.8009729 |
| TCGA-E9-A1R7 | TCGA-BRCA | 94.0846456 | 31.7546517 |
| TCGA-E9-A1RA | TCGA-BRCA | 50.4853843 | 36.1550095 |
| TCGA-E9-A1RB | TCGA-BRCA | 37.0278808 | 15.4186959 |
| TCGA-E9-A1RC | TCGA-BRCA | 28.8592763 | 11.0778158 |
| TCGA-E9-A1RD | TCGA-BRCA | 35.6281842 | 13.9364068 |
| TCGA-E9-A1RE | TCGA-BRCA | 47.7045884 | 49.6813258 |
| TCGA-E9-A1RF | TCGA-BRCA | 29.2542384 | 12.0432528 |
| TCGA-E9-A1RG | TCGA-BRCA | 65.4056079 | 66.3577705 |
| TCGA-E9-A1RH | TCGA-BRCA | 55.2968676 | 19.2662461 |
| TCGA-E9-A1RI | TCGA-BRCA | 25.4226218 | 16.5437782 |
| TCGA-E9-A226 | TCGA-BRCA | 55.298757 | 36.337287 |
| TCGA-E9-A227 | TCGA-BRCA | 49.8298487 | 28.8160972 |
| TCGA-E9-A228 | TCGA-BRCA | 53.3177231 | 34.4505263 |
| TCGA-E9-A229 | TCGA-BRCA | 60.7761265 | 23.8228014 |
| TCGA-E9-A22A | TCGA-BRCA | 33.9122149 | 33.0061979 |
| TCGA-E9-A22B | TCGA-BRCA | 37.6412613 | 42.4034873 |
| TCGA-E9-A22D | TCGA-BRCA | 81.6927153 | 34.3334138 |
| TCGA-E9-A22E | TCGA-BRCA | 59.4334886 | 32.2230812 |
| TCGA-E9-A22G | TCGA-BRCA | 184.534656 | 72.028807 |
| TCGA-E9-A22H | TCGA-BRCA | 61.7802642 | 47.1129562 |
| TCGA-E9-A243 | TCGA-BRCA | 40.318206 | 52.7385963 |
| TCGA-E9-A244 | TCGA-BRCA | 64.8705419 | 57.9796842 |
| TCGA-E9-A245 | TCGA-BRCA | 44.2306609 | 32.6177706 |

| | | | |
|---|---|---|---|
| TCGA-E9-A247 | TCGA-BRCA | 73.5776917 | 83.0731248 |
| TCGA-E9-A248 | TCGA-BRCA | 52.01102 | 19.4354528 |
| TCGA-E9-A249 | TCGA-BRCA | 45.3115574 | 42.0232204 |
| TCGA-E9-A24A | TCGA-BRCA | 54.7798719 | 29.9131747 |
| TCGA-E9-A295 | TCGA-BRCA | 48.0242611 | 44.0698189 |
| TCGA-E9-A2JS | TCGA-BRCA | 33.5834936 | 52.5013824 |
| TCGA-E9-A2JT | TCGA-BRCA | 48.9482233 | 21.5416387 |
| TCGA-E9-A3HO | TCGA-BRCA | 41.1287956 | 43.0368091 |
| TCGA-E9-A3Q9 | TCGA-BRCA | 30.9523754 | 12.4081269 |
| TCGA-E9-A3QA | TCGA-BRCA | 89.9455051 | 32.0559974 |
| TCGA-E9-A3X8 | TCGA-BRCA | 32.464931 | 20.7889005 |
| TCGA-E9-A54X | TCGA-BRCA | 41.7549592 | 27.7431119 |
| TCGA-E9-A54Y | TCGA-BRCA | 41.2950223 | 32.275146 |
| TCGA-E9-A5FK | TCGA-BRCA | 36.9328356 | 20.6945452 |
| TCGA-E9-A5FL | TCGA-BRCA | 82.0652433 | 43.2965717 |
| TCGA-E9-A5UO | TCGA-BRCA | 44.8282258 | 13.521123 |
| TCGA-E9-A5UP | TCGA-BRCA | 27.0904337 | 16.9466954 |
| TCGA-E9-A6HE | TCGA-BRCA | 58.4849813 | 39.4705759 |
| TCGA-EB-A1NK | TCGA-SKCM | 30.9470947 | 16.0299944 |
| TCGA-EB-A24C | TCGA-SKCM | 25.1099987 | 25.382964 |
| TCGA-EB-A24D | TCGA-SKCM | 32.0276685 | 25.7661743 |
| TCGA-EB-A299 | TCGA-SKCM | 48.3728394 | 16.95411 |
| TCGA-EB-A3HV | TCGA-SKCM | 26.9763024 | 11.5518748 |
| TCGA-EB-A3XB | TCGA-SKCM | 56.5645164 | 25.4684042 |
| TCGA-EB-A3XC | TCGA-SKCM | 108.605323 | 46.0051147 |
| TCGA-EB-A3XD | TCGA-SKCM | 40.384606 | 6.24525194 |
| TCGA-EB-A3XE | TCGA-SKCM | 49.3093581 | 17.9230174 |
| TCGA-EB-A3XF | TCGA-SKCM | 60.7690503 | 30.0730094 |
| TCGA-EB-A3Y6 | TCGA-SKCM | 51.5401248 | 21.5176194 |
| TCGA-EB-A3Y7 | TCGA-SKCM | 44.7924571 | 12.4721241 |
| TCGA-EB-A41A | TCGA-SKCM | 103.315867 | 19.7863191 |

| | | | |
|---|---|---|---|
| TCGA-EB-A41B | TCGA-SKCM | 48.1742102 | 14.3517086 |
| TCGA-EB-A42Y | TCGA-SKCM | 53.3057525 | 22.7388757 |
| TCGA-EB-A42Z | TCGA-SKCM | 66.038491 | 22.9164105 |
| TCGA-EB-A430 | TCGA-SKCM | 59.3835263 | 20.9427671 |
| TCGA-EB-A431 | TCGA-SKCM | 59.1348772 | 24.2758302 |
| TCGA-EB-A44N | TCGA-SKCM | 52.5245868 | 22.9989855 |
| TCGA-EB-A44O | TCGA-SKCM | 74.5272376 | 34.5825808 |
| TCGA-EB-A44P | TCGA-SKCM | 63.2209363 | 23.9216564 |
| TCGA-EB-A44Q | TCGA-SKCM | 48.1000758 | 20.7528323 |
| TCGA-EB-A44R | TCGA-SKCM | 47.8222816 | 32.4111144 |
| TCGA-EB-A4IQ | TCGA-SKCM | 43.6044477 | 20.612326 |
| TCGA-EB-A4IS | TCGA-SKCM | 35.8143789 | 21.2720087 |
| TCGA-EB-A4OY | TCGA-SKCM | 17.368622 | 12.0418934 |
| TCGA-EB-A4OZ | TCGA-SKCM | 39.7581993 | 7.68596841 |
| TCGA-EB-A4P0 | TCGA-SKCM | 48.4816391 | 24.7781643 |
| TCGA-EB-A4XL | TCGA-SKCM | 38.8439473 | 11.846434 |
| TCGA-EB-A51B | TCGA-SKCM | 54.2623634 | 41.7117462 |
| TCGA-EB-A550 | TCGA-SKCM | 36.8331373 | 22.6636979 |
| TCGA-EB-A551 | TCGA-SKCM | 41.5481615 | 33.5313705 |
| TCGA-EB-A553 | TCGA-SKCM | 24.8945182 | 6.63676639 |
| TCGA-EB-A57M | TCGA-SKCM | 24.91192 | 8.38150993 |
| TCGA-EB-A5FP | TCGA-SKCM | 68.3142262 | 31.1130815 |
| TCGA-EB-A5KH | TCGA-SKCM | 30.1589412 | 15.0220544 |
| TCGA-EB-A5SE | TCGA-SKCM | 51.2313411 | 27.794645 |
| TCGA-EB-A5SF | TCGA-SKCM | 14.3666921 | 8.99627742 |
| TCGA-EB-A5SG | TCGA-SKCM | 29.9287721 | 11.3966926 |
| TCGA-EB-A5SH | TCGA-SKCM | 26.5290075 | 13.6301246 |
| TCGA-EB-A5UL | TCGA-SKCM | 68.0006052 | 26.5552494 |
| TCGA-EB-A5UM | TCGA-SKCM | 22.5600249 | 23.3580713 |
| TCGA-EB-A5UN | TCGA-SKCM | 46.1110266 | 35.954164 |
| TCGA-EB-A5VU | TCGA-SKCM | 56.6130539 | 48.17519 |

| | | | |
|---|---|---|---|
| TCGA-EB-A5VV | TCGA-SKCM | 59.1165204 | 12.5298673 |
| TCGA-EB-A6L9 | TCGA-SKCM | 49.3699213 | 19.136086 |
| TCGA-EB-A6QY | TCGA-SKCM | 71.4716255 | 41.9434384 |
| TCGA-EB-A6QZ | TCGA-SKCM | 77.5599303 | 26.2546556 |
| TCGA-EB-A6R0 | TCGA-SKCM | 56.9027935 | 43.4599775 |
| TCGA-EB-A82B | TCGA-SKCM | 56.3398421 | 26.718509 |
| TCGA-EB-A82C | TCGA-SKCM | 60.1410761 | 19.7876172 |
| TCGA-EB-A85I | TCGA-SKCM | 53.6597147 | 16.521006 |
| TCGA-EB-A85J | TCGA-SKCM | 61.2547918 | 10.7697189 |
| TCGA-EB-A97M | TCGA-SKCM | 25.2521828 | 22.1949363 |
| TCGA-ED-A459 | TCGA-LIHC | 74.7080755 | 23.2762645 |
| TCGA-ED-A4XI | TCGA-LIHC | 29.4448572 | 8.08577276 |
| TCGA-ED-A5KG | TCGA-LIHC | 53.6475889 | 18.0898002 |
| TCGA-ED-A627 | TCGA-LIHC | 32.5995868 | 13.169994 |
| TCGA-ED-A66X | TCGA-LIHC | 26.1553032 | 13.6313158 |
| TCGA-ED-A66Y | TCGA-LIHC | 40.0453328 | 12.8478238 |
| TCGA-ED-A7PX | TCGA-LIHC | 77.3299483 | 47.8381632 |
| TCGA-ED-A7PY | TCGA-LIHC | 37.022394 | 22.7390862 |
| TCGA-ED-A7PZ | TCGA-LIHC | 45.0796234 | 40.3379147 |
| TCGA-ED-A7XO | TCGA-LIHC | 61.4385237 | 16.9320208 |
| TCGA-ED-A7XP | TCGA-LIHC | 46.4213202 | 13.1570157 |
| TCGA-ED-A82E | TCGA-LIHC | 37.9337686 | 28.2156785 |
| TCGA-ED-A8O5 | TCGA-LIHC | 24.6286775 | 14.5539299 |
| TCGA-ED-A8O6 | TCGA-LIHC | 48.9227745 | 12.8030555 |
| TCGA-ED-A97K | TCGA-LIHC | 88.6843675 | 15.1543919 |
| TCGA-EE-A17X | TCGA-SKCM | 50.7249821 | 12.6553065 |
| TCGA-EE-A17Y | TCGA-SKCM | 26.941213 | 33.931309 |
| TCGA-EE-A17Z | TCGA-SKCM | 76.9249842 | 22.3692215 |
| TCGA-EE-A180 | TCGA-SKCM | 31.3327734 | 33.6992823 |
| TCGA-EE-A181 | TCGA-SKCM | 55.8157677 | 24.0970115 |
| TCGA-EE-A182 | TCGA-SKCM | 21.7623542 | 8.17843541 |

| | | | |
|---|---|---|---|
| TCGA-EE-A183 | TCGA-SKCM | 46.2435243 | 14.7215699 |
| TCGA-EE-A184 | TCGA-SKCM | 30.6518 | 14.3459636 |
| TCGA-EE-A185 | TCGA-SKCM | 29.8157421 | 15.5974717 |
| TCGA-EE-A20B | TCGA-SKCM | 45.9660496 | 20.9019418 |
| TCGA-EE-A20C | TCGA-SKCM | 32.496886 | 15.710472 |
| TCGA-EE-A20F | TCGA-SKCM | 51.4329792 | 22.7904421 |
| TCGA-EE-A20H | TCGA-SKCM | 50.0478392 | 54.7566184 |
| TCGA-EE-A20I | TCGA-SKCM | 54.8422447 | 15.4907648 |
| TCGA-EE-A29A | TCGA-SKCM | 44.6817866 | 19.0456383 |
| TCGA-EE-A29B | TCGA-SKCM | 36.0895478 | 15.487438 |
| TCGA-EE-A29C | TCGA-SKCM | 47.9136156 | 24.6477981 |
| TCGA-EE-A29D | TCGA-SKCM | 37.5293218 | 30.556344 |
| TCGA-EE-A29E | TCGA-SKCM | 58.1508498 | 30.864435 |
| TCGA-EE-A29G | TCGA-SKCM | 37.8458497 | 11.3246396 |
| TCGA-EE-A29H | TCGA-SKCM | 37.3093425 | 28.9178671 |
| TCGA-EE-A29L | TCGA-SKCM | 65.967757 | 17.1026674 |
| TCGA-EE-A29M | TCGA-SKCM | 85.7604049 | 25.014294 |
| TCGA-EE-A29N | TCGA-SKCM | 46.9934508 | 25.1730206 |
| TCGA-EE-A29P | TCGA-SKCM | 39.4772048 | 23.7812922 |
| TCGA-EE-A29Q | TCGA-SKCM | 43.6339608 | 12.433686 |
| TCGA-EE-A29R | TCGA-SKCM | 37.2895921 | 17.9498152 |
| TCGA-EE-A29S | TCGA-SKCM | 57.1962889 | 27.9775415 |
| TCGA-EE-A29T | TCGA-SKCM | 42.9498209 | 26.0154896 |
| TCGA-EE-A29V | TCGA-SKCM | 48.5663151 | 6.25327209 |
| TCGA-EE-A29W | TCGA-SKCM | 57.0308974 | 19.7922081 |
| TCGA-EE-A29X | TCGA-SKCM | 54.6886208 | 21.9515267 |
| TCGA-EE-A2A0 | TCGA-SKCM | 36.2565481 | 16.5850866 |
| TCGA-EE-A2A1 | TCGA-SKCM | 43.5296651 | 14.2166067 |
| TCGA-EE-A2A2 | TCGA-SKCM | 71.11522 | 21.4502373 |
| TCGA-EE-A2A5 | TCGA-SKCM | 45.6255613 | 26.4777596 |
| TCGA-EE-A2A6 | TCGA-SKCM | 42.4024743 | 32.3454801 |

| | | | |
|---|---|---|---|
| TCGA-EE-A2GB | TCGA-SKCM | 30.7457129 | 29.5884247 |
| TCGA-EE-A2GC | TCGA-SKCM | 56.0058833 | 23.2333049 |
| TCGA-EE-A2GD | TCGA-SKCM | 55.2873354 | 26.1193461 |
| TCGA-EE-A2GE | TCGA-SKCM | 27.8643916 | 18.2068166 |
| TCGA-EE-A2GH | TCGA-SKCM | 35.5162725 | 23.5358853 |
| TCGA-EE-A2GI | TCGA-SKCM | 87.6979145 | 22.5640083 |
| TCGA-EE-A2GJ | TCGA-SKCM | 48.0906769 | 29.9341577 |
| TCGA-EE-A2GK | TCGA-SKCM | 51.5244208 | 12.9148664 |
| TCGA-EE-A2GL | TCGA-SKCM | 68.9617609 | 21.7735991 |
| TCGA-EE-A2GM | TCGA-SKCM | 30.2252176 | 13.6979313 |
| TCGA-EE-A2GN | TCGA-SKCM | 34.6119426 | 16.080363 |
| TCGA-EE-A2GO | TCGA-SKCM | 27.6684667 | 24.8768558 |
| TCGA-EE-A2GP | TCGA-SKCM | 98.9878487 | 25.7523967 |
| TCGA-EE-A2GR | TCGA-SKCM | 34.7218784 | 16.0801935 |
| TCGA-EE-A2GS | TCGA-SKCM | 58.4822491 | 30.7623031 |
| TCGA-EE-A2GT | TCGA-SKCM | 39.1497445 | 20.0505342 |
| TCGA-EE-A2GU | TCGA-SKCM | 62.1123299 | 32.1985194 |
| TCGA-EE-A2M5 | TCGA-SKCM | 77.8126805 | 53.8728151 |
| TCGA-EE-A2M6 | TCGA-SKCM | 87.1426979 | 43.694006 |
| TCGA-EE-A2M7 | TCGA-SKCM | 37.5821112 | 19.2026912 |
| TCGA-EE-A2M8 | TCGA-SKCM | 27.8307089 | 21.1213822 |
| TCGA-EE-A2MC | TCGA-SKCM | 53.7396254 | 30.441316 |
| TCGA-EE-A2MD | TCGA-SKCM | 68.032712 | 15.7787041 |
| TCGA-EE-A2ME | TCGA-SKCM | 31.3957201 | 12.4955821 |
| TCGA-EE-A2MF | TCGA-SKCM | 61.2653107 | 37.5584306 |
| TCGA-EE-A2MG | TCGA-SKCM | 49.7272089 | 15.9278546 |
| TCGA-EE-A2MH | TCGA-SKCM | 51.148105 | 41.6869407 |
| TCGA-EE-A2MI | TCGA-SKCM | 46.4596021 | 23.1351632 |
| TCGA-EE-A2MJ | TCGA-SKCM | 56.0636149 | 29.07312 |
| TCGA-EE-A2MK | TCGA-SKCM | 39.0518692 | 20.8140815 |
| TCGA-EE-A2ML | TCGA-SKCM | 43.8602584 | 18.7179166 |

| | | | |
|---|---|---|---|
| TCGA-EE-A2MM | TCGA-SKCM | 73.7961633 | 8.35743232 |
| TCGA-EE-A2MN | TCGA-SKCM | 31.4577381 | 12.8708034 |
| TCGA-EE-A2MP | TCGA-SKCM | 41.1248509 | 23.1906552 |
| TCGA-EE-A2MQ | TCGA-SKCM | 64.298208 | 35.3487393 |
| TCGA-EE-A2MR | TCGA-SKCM | 111.589849 | 23.3522684 |
| TCGA-EE-A2MS | TCGA-SKCM | 32.7134525 | 25.760655 |
| TCGA-EE-A2MT | TCGA-SKCM | 48.4183535 | 24.5866393 |
| TCGA-EE-A2MU | TCGA-SKCM | 68.0559979 | 18.7633009 |
| TCGA-EE-A3AA | TCGA-SKCM | 47.869485 | 69.3498192 |
| TCGA-EE-A3AB | TCGA-SKCM | 37.0195972 | 23.259795 |
| TCGA-EE-A3AC | TCGA-SKCM | 44.305237 | 20.8699046 |
| TCGA-EE-A3AD | TCGA-SKCM | 40.8238138 | 32.3223102 |
| TCGA-EE-A3AF | TCGA-SKCM | 62.8222135 | 23.3138732 |
| TCGA-EE-A3AG | TCGA-SKCM | 104.352108 | 49.4462307 |
| TCGA-EE-A3AH | TCGA-SKCM | 70.2912838 | 39.0038601 |
| TCGA-EE-A3J3 | TCGA-SKCM | 44.5451665 | 29.6547011 |
| TCGA-EE-A3J4 | TCGA-SKCM | 42.527062 | 25.4038973 |
| TCGA-EE-A3J5 | TCGA-SKCM | 62.1603071 | 27.9528999 |
| TCGA-EE-A3J7 | TCGA-SKCM | 65.7386356 | 17.5282296 |
| TCGA-EE-A3J8 | TCGA-SKCM | 71.508976 | 19.7017741 |
| TCGA-EE-A3JA | TCGA-SKCM | 56.6310703 | 16.6081447 |
| TCGA-EE-A3JB | TCGA-SKCM | 78.0798411 | 48.4173632 |
| TCGA-EE-A3JD | TCGA-SKCM | 75.5825099 | 33.8067856 |
| TCGA-EE-A3JE | TCGA-SKCM | 37.3432543 | 23.2717155 |
| TCGA-EE-A3JH | TCGA-SKCM | 27.8804091 | 7.26998628 |
| TCGA-EE-A3JI | TCGA-SKCM | 50.8900403 | 22.5899217 |
| TCGA-EJ-5494 | TCGA-PRAD | 32.3052588 | 15.324609 |
| TCGA-EJ-5495 | TCGA-PRAD | 37.1707079 | 13.9776128 |
| TCGA-EJ-5496 | TCGA-PRAD | 44.3018943 | 18.3644145 |
| TCGA-EJ-5497 | TCGA-PRAD | 43.4081083 | 19.3097011 |
| TCGA-EJ-5498 | TCGA-PRAD | 34.6184892 | 15.7488665 |

| | | | |
|---|---|---|---|
| TCGA-EJ-5499 | TCGA-PRAD | 47.6569068 | 18.6274567 |
| TCGA-EJ-5501 | TCGA-PRAD | 35.9150103 | 20.8369411 |
| TCGA-EJ-5502 | TCGA-PRAD | 33.7486552 | 15.0512194 |
| TCGA-EJ-5503 | TCGA-PRAD | 38.9578806 | 15.8076174 |
| TCGA-EJ-5504 | TCGA-PRAD | 30.9533109 | 21.7024402 |
| TCGA-EJ-5505 | TCGA-PRAD | 38.7469707 | 21.0158837 |
| TCGA-EJ-5506 | TCGA-PRAD | 40.6626062 | 20.0839476 |
| TCGA-EJ-5507 | TCGA-PRAD | 68.0947122 | 22.3495051 |
| TCGA-EJ-5508 | TCGA-PRAD | 41.4254944 | 17.3528481 |
| TCGA-EJ-5509 | TCGA-PRAD | 33.3405615 | 14.4767138 |
| TCGA-EJ-5510 | TCGA-PRAD | 33.1146265 | 10.4557379 |
| TCGA-EJ-5511 | TCGA-PRAD | 37.1255161 | 15.3975083 |
| TCGA-EJ-5512 | TCGA-PRAD | 39.2280882 | 20.5714798 |
| TCGA-EJ-5514 | TCGA-PRAD | 50.9273742 | 4.16436327 |
| TCGA-EJ-5515 | TCGA-PRAD | 39.9782478 | 17.4382876 |
| TCGA-EJ-5516 | TCGA-PRAD | 45.3453028 | 26.9246338 |
| TCGA-EJ-5517 | TCGA-PRAD | 41.5223132 | 19.7262013 |
| TCGA-EJ-5518 | TCGA-PRAD | 33.2844001 | 13.9013805 |
| TCGA-EJ-5519 | TCGA-PRAD | 33.4363708 | 10.3508035 |
| TCGA-EJ-5521 | TCGA-PRAD | 35.2686582 | 18.990633 |
| TCGA-EJ-5522 | TCGA-PRAD | 48.4801762 | 13.6218541 |
| TCGA-EJ-5524 | TCGA-PRAD | 50.3710712 | 21.6488269 |
| TCGA-EJ-5525 | TCGA-PRAD | 43.3918643 | 20.6185478 |
| TCGA-EJ-5526 | TCGA-PRAD | 53.1444967 | 26.3140079 |
| TCGA-EJ-5527 | TCGA-PRAD | 42.7943923 | 18.3069783 |
| TCGA-EJ-5530 | TCGA-PRAD | 37.018784 | 17.0963431 |
| TCGA-EJ-5531 | TCGA-PRAD | 33.3296216 | 12.8237458 |
| TCGA-EJ-5532 | TCGA-PRAD | 32.8499691 | 19.0853275 |
| TCGA-EJ-5542 | TCGA-PRAD | 42.6517238 | 23.2611266 |
| TCGA-EJ-7115 | TCGA-PRAD | 27.4176403 | 10.8533737 |
| TCGA-EJ-7123 | TCGA-PRAD | 29.0765086 | 11.2048506 |

| | | | |
|---|---|---|---|
| TCGA-EJ-7125 | TCGA-PRAD | 21.674942 | 12.5357106 |
| TCGA-EJ-7218 | TCGA-PRAD | 27.0962816 | 12.5590725 |
| TCGA-EJ-7312 | TCGA-PRAD | 44.2675812 | 21.5231521 |
| TCGA-EJ-7314 | TCGA-PRAD | 44.0008388 | 17.4997831 |
| TCGA-EJ-7315 | TCGA-PRAD | 50.5342051 | 17.1590666 |
| TCGA-EJ-7317 | TCGA-PRAD | 41.1483279 | 14.0639847 |
| TCGA-EJ-7318 | TCGA-PRAD | 39.3882103 | 19.0356226 |
| TCGA-EJ-7321 | TCGA-PRAD | 32.3538175 | 11.6818476 |
| TCGA-EJ-7325 | TCGA-PRAD | 24.2822185 | 8.4450941 |
| TCGA-EJ-7327 | TCGA-PRAD | 41.7369819 | 15.9095551 |
| TCGA-EJ-7328 | TCGA-PRAD | 45.4181181 | 16.8978309 |
| TCGA-EJ-7330 | TCGA-PRAD | 43.8738648 | 22.2901869 |
| TCGA-EJ-7331 | TCGA-PRAD | 35.489113 | 15.0808206 |
| TCGA-EJ-7781 | TCGA-PRAD | 33.8797624 | 12.2090327 |
| TCGA-EJ-7782 | TCGA-PRAD | 53.1121437 | 20.7968286 |
| TCGA-EJ-7783 | TCGA-PRAD | 33.2159351 | 11.2957593 |
| TCGA-EJ-7784 | TCGA-PRAD | 51.1787389 | 16.8604007 |
| TCGA-EJ-7785 | TCGA-PRAD | 48.9237817 | 17.1624943 |
| TCGA-EJ-7786 | TCGA-PRAD | 40.2517782 | 14.7721345 |
| TCGA-EJ-7788 | TCGA-PRAD | 30.2552219 | 12.6789022 |
| TCGA-EJ-7789 | TCGA-PRAD | 31.6621559 | 20.9124679 |
| TCGA-EJ-7791 | TCGA-PRAD | 35.2033762 | 16.1550706 |
| TCGA-EJ-7792 | TCGA-PRAD | 38.0698247 | 16.2108912 |
| TCGA-EJ-7793 | TCGA-PRAD | 48.7325256 | 19.2903462 |
| TCGA-EJ-7794 | TCGA-PRAD | 48.6383643 | 14.441146 |
| TCGA-EJ-7797 | TCGA-PRAD | 44.9406169 | 16.4187405 |
| TCGA-EJ-8468 | TCGA-PRAD | 48.5792032 | 11.8809063 |
| TCGA-EJ-8469 | TCGA-PRAD | 48.6781971 | 20.2511693 |
| TCGA-EJ-8470 | TCGA-PRAD | 52.5543896 | 13.4856567 |
| TCGA-EJ-8472 | TCGA-PRAD | 64.8614174 | 13.3245597 |
| TCGA-EJ-8474 | TCGA-PRAD | 36.1257933 | 20.1946794 |

| | | | |
|---|---|---|---|
| TCGA-EJ-A46B | TCGA-PRAD | 33.0440151 | 14.5221378 |
| TCGA-EJ-A46D | TCGA-PRAD | 39.3000327 | 15.524394 |
| TCGA-EJ-A46E | TCGA-PRAD | 50.1679729 | 13.1202726 |
| TCGA-EJ-A46F | TCGA-PRAD | 45.9709078 | 18.7402554 |
| TCGA-EJ-A46G | TCGA-PRAD | 25.6797333 | 14.4614768 |
| TCGA-EJ-A46H | TCGA-PRAD | 39.0094943 | 12.6128736 |
| TCGA-EJ-A46I | TCGA-PRAD | 28.774932 | 9.92091179 |
| TCGA-EJ-A65B | TCGA-PRAD | 42.1024067 | 20.5283435 |
| TCGA-EJ-A65D | TCGA-PRAD | 38.7413219 | 17.8708209 |
| TCGA-EJ-A65E | TCGA-PRAD | 53.924431 | 12.4850679 |
| TCGA-EJ-A65F | TCGA-PRAD | 38.8546481 | 14.6350955 |
| TCGA-EJ-A65G | TCGA-PRAD | 27.8663289 | 16.3530282 |
| TCGA-EJ-A65J | TCGA-PRAD | 38.3209176 | 29.981686 |
| TCGA-EJ-A6RA | TCGA-PRAD | 49.5980554 | 22.83903 |
| TCGA-EJ-A6RC | TCGA-PRAD | 35.1219007 | 13.4606536 |
| TCGA-EJ-A7NF | TCGA-PRAD | 45.1862748 | 16.1887122 |
| TCGA-EJ-A7NG | TCGA-PRAD | 47.3122284 | 20.4078843 |
| TCGA-EJ-A7NH | TCGA-PRAD | 33.6202242 | 11.2990929 |
| TCGA-EJ-A7NJ | TCGA-PRAD | 43.503894 | 19.7461558 |
| TCGA-EJ-A7NK | TCGA-PRAD | 36.4316178 | 16.842713 |
| TCGA-EJ-A7NM | TCGA-PRAD | 62.5963044 | 25.7470424 |
| TCGA-EJ-A7NN | TCGA-PRAD | 42.7768536 | 13.0948408 |
| TCGA-EJ-A8FN | TCGA-PRAD | 38.8813225 | 22.9044145 |
| TCGA-EJ-A8FO | TCGA-PRAD | 53.0507818 | 21.1821269 |
| TCGA-EJ-A8FP | TCGA-PRAD | 41.4052947 | 17.7514603 |
| TCGA-EJ-A8FS | TCGA-PRAD | 46.8271398 | 9.16933893 |
| TCGA-EJ-A8FU | TCGA-PRAD | 47.7260267 | 18.538678 |
| TCGA-EJ-AB20 | TCGA-PRAD | 65.2312182 | 16.0865389 |
| TCGA-EJ-AB27 | TCGA-PRAD | 46.0343818 | 22.5901971 |
| TCGA-EL-A3CL | TCGA-THCA | 51.2242501 | 14.7209512 |
| TCGA-EL-A3CM | TCGA-THCA | 35.0579012 | 8.29212315 |

| | | | |
|---|---|---|---|
| TCGA-EL-A3CN | TCGA-THCA | 38.3850075 | 13.5889883 |
| TCGA-EL-A3CO | TCGA-THCA | 52.4620738 | 13.8922237 |
| TCGA-EL-A3CP | TCGA-THCA | 68.5912963 | 17.6862333 |
| TCGA-EL-A3CR | TCGA-THCA | 56.5137776 | 14.2915151 |
| TCGA-EL-A3CS | TCGA-THCA | 40.9845643 | 11.2318916 |
| TCGA-EL-A3CT | TCGA-THCA | 47.6479728 | 15.8353141 |
| TCGA-EL-A3CU | TCGA-THCA | 48.8753911 | 11.856715 |
| TCGA-EL-A3CV | TCGA-THCA | 48.6032569 | 13.6561491 |
| TCGA-EL-A3CW | TCGA-THCA | 50.9384411 | 12.5952731 |
| TCGA-EL-A3CX | TCGA-THCA | 83.9094934 | 18.6614281 |
| TCGA-EL-A3CY | TCGA-THCA | 40.9235274 | 14.1966507 |
| TCGA-EL-A3CZ | TCGA-THCA | 49.5359922 | 12.7664429 |
| TCGA-EL-A3D0 | TCGA-THCA | 81.9678696 | 17.5200653 |
| TCGA-EL-A3D1 | TCGA-THCA | 53.9315198 | 11.3317319 |
| TCGA-EL-A3D4 | TCGA-THCA | 66.8480782 | 15.1956055 |
| TCGA-EL-A3D5 | TCGA-THCA | 106.477388 | 21.7261759 |
| TCGA-EL-A3D6 | TCGA-THCA | 44.6100793 | 14.003463 |
| TCGA-EL-A3GO | TCGA-THCA | 89.8009431 | 21.3992783 |
| TCGA-EL-A3GP | TCGA-THCA | 53.8344383 | 16.8748004 |
| TCGA-EL-A3GQ | TCGA-THCA | 70.1425057 | 19.7065843 |
| TCGA-EL-A3GR | TCGA-THCA | 60.8643599 | 14.8699506 |
| TCGA-EL-A3GS | TCGA-THCA | 46.0929087 | 13.4381134 |
| TCGA-EL-A3GU | TCGA-THCA | 51.0602897 | 20.2329197 |
| TCGA-EL-A3GV | TCGA-THCA | 54.1935249 | 20.2503327 |
| TCGA-EL-A3GW | TCGA-THCA | 63.4672401 | 23.1587071 |
| TCGA-EL-A3GX | TCGA-THCA | 55.8828332 | 17.6054935 |
| TCGA-EL-A3GY | TCGA-THCA | 58.4413399 | 16.69159 |
| TCGA-EL-A3GZ | TCGA-THCA | 70.1407611 | 17.4197846 |
| TCGA-EL-A3H1 | TCGA-THCA | 57.1987863 | 19.42671 |
| TCGA-EL-A3H2 | TCGA-THCA | 88.6477074 | 18.1089489 |
| TCGA-EL-A3H3 | TCGA-THCA | 78.7590821 | 25.0890532 |

| | | | |
|---|---|---|---|
| TCGA-EL-A3H4 | TCGA-THCA | 53.1996378 | 12.6149875 |
| TCGA-EL-A3H5 | TCGA-THCA | 34.2505719 | 14.3086334 |
| TCGA-EL-A3H7 | TCGA-THCA | 37.8077128 | 17.4248298 |
| TCGA-EL-A3H8 | TCGA-THCA | 55.7573645 | 14.3123631 |
| TCGA-EL-A3MW | TCGA-THCA | 82.2906139 | 16.8595384 |
| TCGA-EL-A3MX | TCGA-THCA | 61.3625548 | 18.624451 |
| TCGA-EL-A3MY | TCGA-THCA | 64.5205222 | 18.2869389 |
| TCGA-EL-A3MZ | TCGA-THCA | 41.7714629 | 16.4017676 |
| TCGA-EL-A3N2 | TCGA-THCA | 52.3334236 | 13.2067437 |
| TCGA-EL-A3N3 | TCGA-THCA | 71.3870817 | 17.3543429 |
| TCGA-EL-A3T0 | TCGA-THCA | 62.6793591 | 16.5457958 |
| TCGA-EL-A3T1 | TCGA-THCA | 58.5206522 | 12.6365723 |
| TCGA-EL-A3T2 | TCGA-THCA | 67.4798118 | 11.5931385 |
| TCGA-EL-A3T3 | TCGA-THCA | 42.6528606 | 13.4866221 |
| TCGA-EL-A3T6 | TCGA-THCA | 71.7132142 | 14.5677776 |
| TCGA-EL-A3T7 | TCGA-THCA | 60.8687871 | 14.0789834 |
| TCGA-EL-A3T8 | TCGA-THCA | 63.8906019 | 13.7289558 |
| TCGA-EL-A3T9 | TCGA-THCA | 46.8118037 | 26.8894969 |
| TCGA-EL-A3TA | TCGA-THCA | 70.8128155 | 14.7029218 |
| TCGA-EL-A3TB | TCGA-THCA | 71.4770916 | 16.62679 |
| TCGA-EL-A3ZG | TCGA-THCA | 65.7353309 | 15.5418845 |
| TCGA-EL-A3ZH | TCGA-THCA | 89.1072339 | 22.6646648 |
| TCGA-EL-A3ZK | TCGA-THCA | 37.5272826 | 11.9849469 |
| TCGA-EL-A3ZL | TCGA-THCA | 80.64837 | 16.4629636 |
| TCGA-EL-A3ZM | TCGA-THCA | 85.653387 | 23.9643669 |
| TCGA-EL-A3ZN | TCGA-THCA | 77.0051799 | 12.7140271 |
| TCGA-EL-A3ZO | TCGA-THCA | 71.5051 | 15.9446925 |
| TCGA-EL-A3ZP | TCGA-THCA | 69.3189272 | 18.743992 |
| TCGA-EL-A3ZQ | TCGA-THCA | 42.4284122 | 10.3854987 |
| TCGA-EL-A3ZR | TCGA-THCA | 64.0104496 | 11.0835728 |
| TCGA-EL-A3ZS | TCGA-THCA | 77.3641091 | 14.3396224 |

| | | | |
|---|---|---|---|
| TCGA-EL-A3ZT | TCGA-THCA | 51.824572 | 13.1182925 |
| TCGA-EL-A4JV | TCGA-THCA | 61.5227666 | 15.6491027 |
| TCGA-EL-A4JW | TCGA-THCA | 46.5836487 | 16.0615125 |
| TCGA-EL-A4JX | TCGA-THCA | 19.7205528 | 9.1646647 |
| TCGA-EL-A4JZ | TCGA-THCA | 60.779826 | 16.2128559 |
| TCGA-EL-A4K0 | TCGA-THCA | 45.1179341 | 13.7578101 |
| TCGA-EL-A4K1 | TCGA-THCA | 29.584727 | 9.21640888 |
| TCGA-EL-A4K2 | TCGA-THCA | 55.0295644 | 14.2251217 |
| TCGA-EL-A4K4 | TCGA-THCA | 39.315539 | 12.1098719 |
| TCGA-EL-A4K6 | TCGA-THCA | 48.9186568 | 11.4134353 |
| TCGA-EL-A4K7 | TCGA-THCA | 63.8453113 | 14.8675448 |
| TCGA-EL-A4K9 | TCGA-THCA | 60.8102358 | 12.6335702 |
| TCGA-EL-A4KD | TCGA-THCA | 47.9160752 | 14.0875913 |
| TCGA-EL-A4KG | TCGA-THCA | 50.0476319 | 18.2050851 |
| TCGA-EL-A4KH | TCGA-THCA | 48.7012141 | 12.0845018 |
| TCGA-EM-A1CT | TCGA-THCA | 66.5993652 | 20.0447506 |
| TCGA-EM-A1CU | TCGA-THCA | 62.3960282 | 17.3651073 |
| TCGA-EM-A1CV | TCGA-THCA | 62.8964946 | 15.5269028 |
| TCGA-EM-A1CW | TCGA-THCA | 37.796832 | 11.9930621 |
| TCGA-EM-A1YA | TCGA-THCA | 34.784246 | 10.6633139 |
| TCGA-EM-A1YB | TCGA-THCA | 26.2715072 | 7.79010061 |
| TCGA-EM-A1YC | TCGA-THCA | 71.4738298 | 17.2692705 |
| TCGA-EM-A1YD | TCGA-THCA | 53.6247165 | 15.9674102 |
| TCGA-EM-A1YE | TCGA-THCA | 57.7481129 | 23.1163574 |
| TCGA-EM-A22I | TCGA-THCA | 37.3367664 | 11.3485178 |
| TCGA-EM-A22J | TCGA-THCA | 29.6659644 | 10.7474718 |
| TCGA-EM-A22K | TCGA-THCA | 56.9611289 | 14.8915037 |
| TCGA-EM-A22L | TCGA-THCA | 60.5467993 | 20.3118378 |
| TCGA-EM-A22M | TCGA-THCA | 29.9306512 | 10.9079387 |
| TCGA-EM-A22N | TCGA-THCA | 30.6171287 | 11.0233218 |
| TCGA-EM-A22O | TCGA-THCA | 45.2503849 | 9.68738807 |

| | | | |
|---|---|---|---|
| TCGA-EM-A22P | TCGA-THCA | 45.2587389 | 13.9446481 |
| TCGA-EM-A22Q | TCGA-THCA | 46.8528333 | 7.78836762 |
| TCGA-EM-A2CJ | TCGA-THCA | 65.651154 | 17.4563114 |
| TCGA-EM-A2CK | TCGA-THCA | 54.6103236 | 13.1952616 |
| TCGA-EM-A2CL | TCGA-THCA | 54.7577114 | 16.6804144 |
| TCGA-EM-A2CM | TCGA-THCA | 25.9963219 | 6.88763837 |
| TCGA-EM-A2CN | TCGA-THCA | 56.0707996 | 21.19021 |
| TCGA-EM-A2CC | TCGA-THCA | 26.651754 | 10.9827494 |
| TCGA-EM-A2CP | TCGA-THCA | 43.7185498 | 12.4498406 |
| TCGA-EM-A2CC | TCGA-THCA | 44.1204939 | 12.5555275 |
| TCGA-EM-A2CR | TCGA-THCA | 31.3880582 | 15.0002687 |
| TCGA-EM-A2CS | TCGA-THCA | 50.3327412 | 11.9755239 |
| TCGA-EM-A2CT | TCGA-THCA | 51.5127029 | 16.7871078 |
| TCGA-EM-A2CU | TCGA-THCA | 62.1500994 | 12.7087358 |
| TCGA-EM-A2OV | TCGA-THCA | 34.1327703 | 11.8600273 |
| TCGA-EM-A2OW | TCGA-THCA | 38.2985252 | 10.7920113 |
| TCGA-EM-A2OX | TCGA-THCA | 55.9384458 | 21.1420857 |
| TCGA-EM-A2OY | TCGA-THCA | 34.3401291 | 13.4018866 |
| TCGA-EM-A2OZ | TCGA-THCA | 57.9064916 | 19.0082157 |
| TCGA-EM-A2P0 | TCGA-THCA | 57.9907643 | 15.7644422 |
| TCGA-EM-A2P1 | TCGA-THCA | 46.2438683 | 21.8583278 |
| TCGA-EM-A2P2 | TCGA-THCA | 50.1799306 | 20.6697732 |
| TCGA-EM-A2P3 | TCGA-THCA | 73.4471916 | 18.1061738 |
| TCGA-EM-A3AI | TCGA-THCA | 67.6269236 | 18.8413115 |
| TCGA-EM-A3AJ | TCGA-THCA | 56.0622029 | 16.9326707 |
| TCGA-EM-A3AK | TCGA-THCA | 52.476447 | 15.5440975 |
| TCGA-EM-A3AL | TCGA-THCA | 70.8414147 | 20.1758728 |
| TCGA-EM-A3AN | TCGA-THCA | 68.0522037 | 20.595099 |
| TCGA-EM-A3AC | TCGA-THCA | 88.2968021 | 17.8623577 |
| TCGA-EM-A3AP | TCGA-THCA | 32.7906002 | 14.598362 |
| TCGA-EM-A3AQ | TCGA-THCA | 71.829292 | 14.7783653 |

| | | | |
|---|---|---|---|
| TCGA-EM-A3AR | TCGA-THCA | 75.5038749 | 19.716989 |
| TCGA-EM-A3FJ | TCGA-THCA | 94.4685546 | 22.275186 |
| TCGA-EM-A3FK | TCGA-THCA | 60.5453631 | 22.3718124 |
| TCGA-EM-A3FL | TCGA-THCA | 50.9766843 | 16.2892072 |
| TCGA-EM-A3FM | TCGA-THCA | 41.9238293 | 13.4374525 |
| TCGA-EM-A3FN | TCGA-THCA | 61.0896384 | 24.701827 |
| TCGA-EM-A3FO | TCGA-THCA | 76.3815543 | 27.4196424 |
| TCGA-EM-A3FP | TCGA-THCA | 52.1913216 | 21.8539379 |
| TCGA-EM-A3FQ | TCGA-THCA | 69.5254081 | 35.3412488 |
| TCGA-EM-A3FR | TCGA-THCA | 80.3593083 | 15.3145662 |
| TCGA-EM-A3O3 | TCGA-THCA | 48.3404277 | 27.2804115 |
| TCGA-EM-A3O6 | TCGA-THCA | 77.406984 | 18.8285891 |
| TCGA-EM-A3O7 | TCGA-THCA | 80.6784918 | 15.0782702 |
| TCGA-EM-A3O8 | TCGA-THCA | 70.9520851 | 15.9132241 |
| TCGA-EM-A3OA | TCGA-THCA | 76.0749599 | 18.6258499 |
| TCGA-EM-A3OB | TCGA-THCA | 70.8410658 | 14.3186704 |
| TCGA-EM-A3ST | TCGA-THCA | 86.9701711 | 17.8412169 |
| TCGA-EM-A3SU | TCGA-THCA | 41.914987 | 12.7362524 |
| TCGA-EM-A3SX | TCGA-THCA | 74.382122 | 14.0991367 |
| TCGA-EM-A3SY | TCGA-THCA | 28.1866894 | 25.1667436 |
| TCGA-EM-A3SZ | TCGA-THCA | 43.4660976 | 17.0676508 |
| TCGA-EM-A4FF | TCGA-THCA | 42.81306 | 11.0532397 |
| TCGA-EM-A4FH | TCGA-THCA | 64.3621295 | 25.3921949 |
| TCGA-EM-A4FK | TCGA-THCA | 58.8855642 | 18.1398615 |
| TCGA-EM-A4FM | TCGA-THCA | 55.6080182 | 14.5811274 |
| TCGA-EM-A4FN | TCGA-THCA | 41.8681764 | 13.8385103 |
| TCGA-EM-A4FO | TCGA-THCA | 39.1480111 | 14.0928208 |
| TCGA-EM-A4FQ | TCGA-THCA | 34.340909 | 10.6827694 |
| TCGA-EM-A4FR | TCGA-THCA | 50.9652282 | 12.9563024 |
| TCGA-EM-A4FU | TCGA-THCA | 38.8269552 | 10.8980872 |
| TCGA-EM-A4FV | TCGA-THCA | 60.4184916 | 15.3552591 |

| | | | |
|---|---|---|---|
| TCGA-EM-A4G1 | TCGA-THCA | 48.4145321 | 12.3949454 |
| TCGA-EO-A1Y7 | TCGA-UCEC | 97.1427137 | 12.375901 |
| TCGA-EO-A22U | TCGA-UCEC | 89.4155808 | 76.0504496 |
| TCGA-EO-A22X | TCGA-UCEC | 116.952771 | 33.6045685 |
| TCGA-EO-A22Y | TCGA-UCEC | 61.2883517 | 16.3072567 |
| TCGA-EO-A3AS | TCGA-UCEC | 65.2884892 | 29.6298418 |
| TCGA-EO-A3AU | TCGA-UCEC | 38.0376704 | 9.43664784 |
| TCGA-EO-A3AV | TCGA-UCEC | 39.4458457 | 10.4161027 |
| TCGA-EO-A3AY | TCGA-UCEC | 15.3631921 | 4.27009407 |
| TCGA-EO-A3AZ | TCGA-UCEC | 5.14097829 | 2.57928347 |
| TCGA-EO-A3B0 | TCGA-UCEC | 82.2242987 | 30.18692 |
| TCGA-EO-A3B1 | TCGA-UCEC | 48.9368517 | 16.3427853 |
| TCGA-EO-A3KU | TCGA-UCEC | 143.919218 | 26.7709757 |
| TCGA-EO-A3KW | TCGA-UCEC | 84.4083826 | 28.7321474 |
| TCGA-EO-A3KX | TCGA-UCEC | 93.6845275 | 18.4549793 |
| TCGA-EO-A3L0 | TCGA-UCEC | 85.65677 | 31.7988806 |
| TCGA-EP-A12J | TCGA-LIHC | 46.6522915 | 18.6386498 |
| TCGA-EP-A26S | TCGA-LIHC | 45.4155901 | 13.7674593 |
| TCGA-EP-A2KA | TCGA-LIHC | 30.8624344 | 32.0968513 |
| TCGA-EP-A2KB | TCGA-LIHC | 28.3133284 | 11.2406575 |
| TCGA-EP-A2KC | TCGA-LIHC | 29.1798468 | 11.5623187 |
| TCGA-EP-A3JL | TCGA-LIHC | 31.7155736 | 27.1763738 |
| TCGA-EP-A3RK | TCGA-LIHC | 17.8742591 | 6.43984802 |
| TCGA-EQ-8122 | TCGA-STAD | 44.7307017 | 29.8329815 |
| TCGA-ER-A193 | TCGA-SKCM | 45.4365399 | 9.38628854 |
| TCGA-ER-A194 | TCGA-SKCM | 41.0607615 | 9.15484596 |
| TCGA-ER-A195 | TCGA-SKCM | 51.7650712 | 19.370267 |
| TCGA-ER-A196 | TCGA-SKCM | 30.5809024 | 14.1258483 |
| TCGA-ER-A197 | TCGA-SKCM | 45.4309534 | 24.0877801 |
| TCGA-ER-A198 | TCGA-SKCM | 54.6791863 | 33.8531982 |
| TCGA-ER-A199 | TCGA-SKCM | 41.7930094 | 20.1308982 |

| | | | |
|---|---|---|---|
| TCGA-ER-A19A | TCGA-SKCM | 55.7981928 | 16.8681504 |
| TCGA-ER-A19B | TCGA-SKCM | 34.357652 | 33.9547881 |
| TCGA-ER-A19C | TCGA-SKCM | 31.4430438 | 17.8614038 |
| TCGA-ER-A19D | TCGA-SKCM | 39.9381044 | 22.1540833 |
| TCGA-ER-A19E | TCGA-SKCM | 53.0010443 | 20.6378291 |
| TCGA-ER-A19F | TCGA-SKCM | 39.5397289 | 17.9801122 |
| TCGA-ER-A19G | TCGA-SKCM | 67.8187982 | 25.0896505 |
| TCGA-ER-A19H | TCGA-SKCM | 43.5973014 | 25.517284 |
| TCGA-ER-A19J | TCGA-SKCM | 40.5387594 | 24.7683383 |
| TCGA-ER-A19K | TCGA-SKCM | 39.5910022 | 32.1412538 |
| TCGA-ER-A19L | TCGA-SKCM | 13.7679518 | 39.0332787 |
| TCGA-ER-A19M | TCGA-SKCM | 58.4778683 | 45.0227798 |
| TCGA-ER-A19N | TCGA-SKCM | 32.0091045 | 13.0279945 |
| TCGA-ER-A19O | TCGA-SKCM | 15.6237237 | 5.95382517 |
| TCGA-ER-A19P | TCGA-SKCM | 35.9975891 | 21.947946 |
| TCGA-ER-A19Q | TCGA-SKCM | 43.0559621 | 23.1246208 |
| TCGA-ER-A19S | TCGA-SKCM | 19.2427392 | 16.3514489 |
| TCGA-ER-A19T | TCGA-SKCM | 93.7017099 | 11.4128124 |
| TCGA-ER-A19W | TCGA-SKCM | 72.5985674 | 28.1940977 |
| TCGA-ER-A1A1 | TCGA-SKCM | 53.8732667 | 15.5580671 |
| TCGA-ER-A2NB | TCGA-SKCM | 20.8312189 | 5.99995398 |
| TCGA-ER-A2NC | TCGA-SKCM | 54.4014217 | 34.3080118 |
| TCGA-ER-A2ND | TCGA-SKCM | 39.448999 | 29.2463612 |
| TCGA-ER-A2NE | TCGA-SKCM | 64.3047995 | 28.138961 |
| TCGA-ER-A2NF | TCGA-SKCM | 25.6690072 | 12.0489982 |
| TCGA-ER-A2NG | TCGA-SKCM | 46.9813718 | 28.8301909 |
| TCGA-ER-A2NH | TCGA-SKCM | 35.1571219 | 30.7726382 |
| TCGA-ER-A3ES | TCGA-SKCM | 48.4408404 | 13.9343042 |
| TCGA-ER-A3ET | TCGA-SKCM | 46.7976137 | 38.6533082 |
| TCGA-ER-A3EV | TCGA-SKCM | 37.7084117 | 14.2957311 |
| TCGA-ER-A3PL | TCGA-SKCM | 43.2248087 | 40.2326752 |

| | | | |
|---|---|---|---|
| TCGA-ER-A42H | TCGA-SKCM | 66.7846238 | 35.5397596 |
| TCGA-ER-A42K | TCGA-SKCM | 46.1491156 | 38.4146846 |
| TCGA-ER-A42L | TCGA-SKCM | 45.2056605 | 20.8010578 |
| TCGA-ES-A2HS | TCGA-LIHC | 21.0205771 | 7.44824543 |
| TCGA-ES-A2HT | TCGA-LIHC | 25.4374838 | 14.9478992 |
| TCGA-ET-A25G | TCGA-THCA | 35.0197923 | 15.9336799 |
| TCGA-ET-A25I | TCGA-THCA | 33.7101691 | 10.3722608 |
| TCGA-ET-A25K | TCGA-THCA | 53.6064012 | 12.928601 |
| TCGA-ET-A25L | TCGA-THCA | 41.7989312 | 12.9949898 |
| TCGA-ET-A25M | TCGA-THCA | 45.3716825 | 8.13408048 |
| TCGA-ET-A25N | TCGA-THCA | 53.4728399 | 14.396665 |
| TCGA-ET-A25O | TCGA-THCA | 65.6178272 | 12.5211606 |
| TCGA-ET-A25P | TCGA-THCA | 63.4015286 | 12.545168 |
| TCGA-ET-A25R | TCGA-THCA | 50.9820602 | 10.3596943 |
| TCGA-ET-A2MX | TCGA-THCA | 45.5297591 | 10.7162913 |
| TCGA-ET-A2MY | TCGA-THCA | 44.5933174 | 11.0527034 |
| TCGA-ET-A2MZ | TCGA-THCA | 52.9476855 | 14.0970941 |
| TCGA-ET-A2N0 | TCGA-THCA | 47.7824418 | 11.0924276 |
| TCGA-ET-A2N3 | TCGA-THCA | 69.6301225 | 17.1040053 |
| TCGA-ET-A2N4 | TCGA-THCA | 55.4193025 | 14.2887443 |
| TCGA-ET-A2N5 | TCGA-THCA | 79.5249976 | 24.8557607 |
| TCGA-ET-A39I | TCGA-THCA | 48.6218775 | 17.0172171 |
| TCGA-ET-A39J | TCGA-THCA | 56.2696863 | 15.4812802 |
| TCGA-ET-A39K | TCGA-THCA | 60.2527134 | 14.2542383 |
| TCGA-ET-A39L | TCGA-THCA | 67.0980484 | 18.1482716 |
| TCGA-ET-A39M | TCGA-THCA | 38.7850408 | 8.55884236 |
| TCGA-ET-A39N | TCGA-THCA | 51.307521 | 27.9440839 |
| TCGA-ET-A39O | TCGA-THCA | 35.7395717 | 13.3390803 |
| TCGA-ET-A39P | TCGA-THCA | 42.1973138 | 15.3458273 |
| TCGA-ET-A39R | TCGA-THCA | 55.130029 | 15.3550062 |
| TCGA-ET-A39S | TCGA-THCA | 64.8999315 | 18.09307 |

| | | | |
|---|---|---|---|
| TCGA-ET-A39T | TCGA-THCA | 50.1100567 | 13.1756227 |
| TCGA-ET-A3BN | TCGA-THCA | 44.656344 | 11.0226302 |
| TCGA-ET-A3BO | TCGA-THCA | 41.1865823 | 14.0234994 |
| TCGA-ET-A3BP | TCGA-THCA | 60.248928 | 13.396356 |
| TCGA-ET-A3BQ | TCGA-THCA | 62.9994293 | 17.802108 |
| TCGA-ET-A3BS | TCGA-THCA | 58.0945828 | 15.1872079 |
| TCGA-ET-A3BT | TCGA-THCA | 50.8620939 | 14.1006519 |
| TCGA-ET-A3BU | TCGA-THCA | 44.9126742 | 14.2534173 |
| TCGA-ET-A3BV | TCGA-THCA | 49.2856163 | 20.0136701 |
| TCGA-ET-A3BW | TCGA-THCA | 62.4142554 | 14.9631656 |
| TCGA-ET-A3BX | TCGA-THCA | 41.5784657 | 15.0387607 |
| TCGA-ET-A3DO | TCGA-THCA | 31.6515879 | 11.8901356 |
| TCGA-ET-A3DP | TCGA-THCA | 87.7998928 | 19.0429987 |
| TCGA-ET-A3DQ | TCGA-THCA | 46.0884389 | 11.0902577 |
| TCGA-ET-A3DR | TCGA-THCA | 60.4823049 | 17.1734771 |
| TCGA-ET-A3DS | TCGA-THCA | 50.6614691 | 14.9022096 |
| TCGA-ET-A3DU | TCGA-THCA | 26.6532766 | 8.82814655 |
| TCGA-ET-A3DV | TCGA-THCA | 69.4783026 | 22.6135072 |
| TCGA-ET-A3DW | TCGA-THCA | 62.3524898 | 16.3420507 |
| TCGA-ET-A40P | TCGA-THCA | 106.188645 | 21.4286663 |
| TCGA-ET-A40Q | TCGA-THCA | 47.2765926 | 14.2926079 |
| TCGA-ET-A40R | TCGA-THCA | 95.1739958 | 18.7103452 |
| TCGA-ET-A40S | TCGA-THCA | 69.4688438 | 17.72075 |
| TCGA-ET-A40T | TCGA-THCA | 63.0634829 | 16.6799274 |
| TCGA-ET-A4KN | TCGA-THCA | 6.88652819 | 7.79669039 |
| TCGA-ET-A4KQ | TCGA-THCA | 45.7490932 | 14.7142303 |
| TCGA-EU-5904 | TCGA-KIRC | 46.698328 | 13.1932994 |
| TCGA-EU-5905 | TCGA-KIRC | 33.9315789 | 18.0799649 |
| TCGA-EU-5906 | TCGA-KIRC | 47.9338457 | 16.0915404 |
| TCGA-EU-5907 | TCGA-KIRC | 27.2777929 | 30.1701013 |
| TCGA-EV-5901 | TCGA-KIRP | 10.0073716 | 3.66840733 |

| | | | |
|---|---|---|---|
| TCGA-EV-5902 | TCGA-KIRP | 45.3223933 | 18.429168 |
| TCGA-EV-5903 | TCGA-KIRP | 27.6025485 | 11.914346 |
| TCGA-EW-A1IW | TCGA-BRCA | 48.4204534 | 37.8062487 |
| TCGA-EW-A1IX | TCGA-BRCA | 41.5940742 | 14.8666744 |
| TCGA-EW-A1IY | TCGA-BRCA | 40.2593117 | 35.1116425 |
| TCGA-EW-A1IZ | TCGA-BRCA | 63.4106883 | 24.3373046 |
| TCGA-EW-A1J1 | TCGA-BRCA | 46.4547567 | 33.63073 |
| TCGA-EW-A1J2 | TCGA-BRCA | 49.0502546 | 22.2787643 |
| TCGA-EW-A1J3 | TCGA-BRCA | 50.3048529 | 29.6676184 |
| TCGA-EW-A1J5 | TCGA-BRCA | 50.9074138 | 32.5248096 |
| TCGA-EW-A1J6 | TCGA-BRCA | 40.5300471 | 58.2907656 |
| TCGA-EW-A1OV | TCGA-BRCA | 63.0266995 | 22.3876763 |
| TCGA-EW-A1OW | TCGA-BRCA | 244.934305 | 76.8576611 |
| TCGA-EW-A1OX | TCGA-BRCA | 27.0415776 | 38.7622896 |
| TCGA-EW-A1OY | TCGA-BRCA | 81.2603103 | 16.8294063 |
| TCGA-EW-A1OZ | TCGA-BRCA | 27.4540036 | 29.0147366 |
| TCGA-EW-A1P0 | TCGA-BRCA | 45.0074533 | 24.2966147 |
| TCGA-EW-A1P1 | TCGA-BRCA | 61.5500499 | 24.8980019 |
| TCGA-EW-A1P3 | TCGA-BRCA | 53.9328616 | 35.0090001 |
| TCGA-EW-A1P4 | TCGA-BRCA | 36.5496219 | 46.8846999 |
| TCGA-EW-A1P5 | TCGA-BRCA | 51.7843492 | 32.5367176 |
| TCGA-EW-A1P6 | TCGA-BRCA | 43.2555172 | 23.1620559 |
| TCGA-EW-A1P7 | TCGA-BRCA | 45.1990967 | 19.3625337 |
| TCGA-EW-A1P8 | TCGA-BRCA | 142.549482 | 94.128341 |
| TCGA-EW-A1PA | TCGA-BRCA | 51.675745 | 32.385966 |
| TCGA-EW-A1PB | TCGA-BRCA | 115.88563 | 47.609987 |
| TCGA-EW-A1PC | TCGA-BRCA | 143.197974 | 46.1292437 |
| TCGA-EW-A1PD | TCGA-BRCA | 86.1551524 | 22.7678905 |
| TCGA-EW-A1PE | TCGA-BRCA | 38.1597687 | 35.58513 |
| TCGA-EW-A1PF | TCGA-BRCA | 46.2393578 | 29.03618 |
| TCGA-EW-A1PG | TCGA-BRCA | 69.1081887 | 24.3152867 |

| | | | |
|---|---|---|---|
| TCGA-EW-A1PH | TCGA-BRCA | 81.9156011 | 50.1120935 |
| TCGA-EW-A2FR | TCGA-BRCA | 55.4592087 | 63.6308455 |
| TCGA-EW-A2FS | TCGA-BRCA | 57.6888353 | 44.3697724 |
| TCGA-EW-A2FV | TCGA-BRCA | 52.5951213 | 39.0552778 |
| TCGA-EW-A2FW | TCGA-BRCA | 45.8992778 | 69.6028856 |
| TCGA-EW-A3E8 | TCGA-BRCA | 55.0177536 | 23.723028 |
| TCGA-EW-A3U0 | TCGA-BRCA | 54.5664455 | 13.2643271 |
| TCGA-EW-A423 | TCGA-BRCA | 106.293314 | 49.8795787 |
| TCGA-EW-A424 | TCGA-BRCA | 63.0217981 | 27.4305019 |
| TCGA-EW-A6S9 | TCGA-BRCA | 66.5256404 | 39.7334964 |
| TCGA-EW-A6SA | TCGA-BRCA | 77.2040851 | 37.7074486 |
| TCGA-EW-A6SB | TCGA-BRCA | 70.2961829 | 61.1628226 |
| TCGA-EW-A6SC | TCGA-BRCA | 62.1524721 | 91.0439492 |
| TCGA-EW-A6SD | TCGA-BRCA | 49.7099839 | 16.4485506 |
| TCGA-EY-A1GL | TCGA-UCEC | 50.1105468 | 18.638974 |
| TCGA-EY-A1GO | TCGA-UCEC | 58.2167478 | 40.7336539 |
| TCGA-EY-A1GP | TCGA-UCEC | 58.9681235 | 46.7112324 |
| TCGA-EY-A1GX | TCGA-UCEC | 33.338368 | 19.0467628 |
| TCGA-EY-A210 | TCGA-UCEC | 127.59602 | 24.4370973 |
| TCGA-EY-A2ON | TCGA-UCEC | 86.8713499 | 27.5282351 |
| TCGA-EY-A2OO | TCGA-UCEC | 20.352026 | 7.13358217 |
| TCGA-EY-A2OP | TCGA-UCEC | 74.6429997 | 30.3150497 |
| TCGA-EY-A2OQ | TCGA-UCEC | 22.5499116 | 14.6257896 |
| TCGA-EY-A3L3 | TCGA-UCEC | 108.078764 | 47.7133095 |
| TCGA-EY-A3QX | TCGA-UCEC | 147.216394 | 25.778346 |
| TCGA-EY-A547 | TCGA-UCEC | 53.3887924 | 5.09117522 |
| TCGA-EY-A548 | TCGA-UCEC | 61.3342943 | 16.0290308 |
| TCGA-EY-A549 | TCGA-UCEC | 54.7693799 | 23.2342995 |
| TCGA-EY-A54A | TCGA-UCEC | 106.430054 | 43.0083814 |
| TCGA-EY-A5W2 | TCGA-UCEC | 52.3591253 | 41.4474602 |
| TCGA-EY-A72D | TCGA-UCEC | 67.4208098 | 36.8572869 |

| | | | |
|---|---|---|---|
| TCGA-EZ-7264 | TCGA-LGG | 19.3741989 | 19.7444533 |
| TCGA-F1-6177 | TCGA-STAD | 44.0057 | 30.5699654 |
| TCGA-F1-6874 | TCGA-STAD | 25.9439106 | 13.8886645 |
| TCGA-F1-6875 | TCGA-STAD | 45.1757942 | 12.1934198 |
| TCGA-F1-A448 | TCGA-STAD | 36.3296606 | 21.2356815 |
| TCGA-F1-A72C | TCGA-STAD | 59.8251212 | 30.9666316 |
| TCGA-F2-6879 | TCGA-PAAD | 73.1250896 | 14.357423 |
| TCGA-F2-6880 | TCGA-PAAD | 8.42189405 | 3.05416912 |
| TCGA-F2-7273 | TCGA-PAAD | 37.6214983 | 11.2630654 |
| TCGA-F2-7276 | TCGA-PAAD | 51.0318679 | 12.6027635 |
| TCGA-F2-A44G | TCGA-PAAD | 49.0664293 | 11.2515317 |
| TCGA-F2-A44H | TCGA-PAAD | 20.8794655 | 9.47790341 |
| TCGA-F2-A7TX | TCGA-PAAD | 67.072307 | 42.1012517 |
| TCGA-F2-A8YN | TCGA-PAAD | 53.2403923 | 14.0870499 |
| TCGA-F4-6459 | TCGA-COAD | 59.613687 | 34.1595826 |
| TCGA-F4-6460 | TCGA-COAD | 36.2193149 | 32.7001618 |
| TCGA-F4-6461 | TCGA-COAD | 42.0592995 | 8.48269101 |
| TCGA-F4-6463 | TCGA-COAD | 59.7514151 | 19.0122215 |
| TCGA-F4-6569 | TCGA-COAD | 53.7737943 | 21.5775434 |
| TCGA-F4-6570 | TCGA-COAD | 58.430472 | 18.7202453 |
| TCGA-F4-6703 | TCGA-COAD | 45.1695627 | 33.3833514 |
| TCGA-F4-6704 | TCGA-COAD | 51.8499138 | 28.281217 |
| TCGA-F4-6805 | TCGA-COAD | 80.4423935 | 27.4017012 |
| TCGA-F4-6806 | TCGA-COAD | 69.1617407 | 23.079356 |
| TCGA-F4-6807 | TCGA-COAD | 71.4685336 | 16.4774444 |
| TCGA-F4-6808 | TCGA-COAD | 110.831546 | 34.1047931 |
| TCGA-F4-6809 | TCGA-COAD | 55.4541043 | 26.6156298 |
| TCGA-F4-6854 | TCGA-COAD | 82.6836784 | 39.6746825 |
| TCGA-F4-6855 | TCGA-COAD | 39.7351424 | 20.9526847 |
| TCGA-F4-6856 | TCGA-COAD | 59.0276216 | 30.1657153 |
| TCGA-F6-A8O3 | TCGA-LGG | 27.2006235 | 29.1921981 |

| | | | |
|---|---|---|---|
| TCGA-F6-A8O4 | TCGA-LGG | 43.1359958 | 14.3362604 |
| TCGA-F9-A4JJ | TCGA-KIRP | 24.5688969 | 32.7183044 |
| TCGA-F9-A7Q0 | TCGA-KIRP | 6.27622845 | 41.0992537 |
| TCGA-F9-A7VF | TCGA-KIRP | 39.1008708 | 11.3869241 |
| TCGA-F9-A8NY | TCGA-KIRP | 37.6387951 | 11.2766861 |
| TCGA-F9-A97G | TCGA-KIRP | 19.1093129 | 4.6323993 |
| TCGA-FB-A4P5 | TCGA-PAAD | 37.4546082 | 10.2330712 |
| TCGA-FB-A4P6 | TCGA-PAAD | 34.8449099 | 11.2406332 |
| TCGA-FB-A545 | TCGA-PAAD | 39.8969213 | 15.6768814 |
| TCGA-FB-A5VM | TCGA-PAAD | 90.2179563 | 25.1878896 |
| TCGA-FB-A78T | TCGA-PAAD | 47.2867375 | 9.09521973 |
| TCGA-FB-A7DR | TCGA-PAAD | 42.5109487 | 26.3286903 |
| TCGA-FB-AAPP | TCGA-PAAD | 46.7585755 | 25.6265037 |
| TCGA-FB-AAPQ | TCGA-PAAD | 33.4552812 | 10.3187228 |
| TCGA-FB-AAPS | TCGA-PAAD | 46.7899289 | 19.4501164 |
| TCGA-FB-AAPU | TCGA-PAAD | 57.6953408 | 16.7449341 |
| TCGA-FB-AAPY | TCGA-PAAD | 42.3980345 | 10.3125346 |
| TCGA-FB-AAPZ | TCGA-PAAD | 41.6753504 | 12.7986587 |
| TCGA-FB-AAQ0 | TCGA-PAAD | 49.7681544 | 23.6879718 |
| TCGA-FB-AAQ1 | TCGA-PAAD | 54.0435951 | 34.0807932 |
| TCGA-FB-AAQ2 | TCGA-PAAD | 85.059801 | 24.58619 |
| TCGA-FB-AAQ3 | TCGA-PAAD | 51.7570977 | 28.0088662 |
| TCGA-FB-AAQ6 | TCGA-PAAD | 42.1502524 | 20.679961 |
| TCGA-FC-7708 | TCGA-PRAD | 43.3472665 | 12.2321734 |
| TCGA-FC-7961 | TCGA-PRAD | 45.5282303 | 41.2449408 |
| TCGA-FC-A4JI | TCGA-PRAD | 52.1035451 | 26.9120344 |
| TCGA-FC-A5OB | TCGA-PRAD | 61.5116388 | 37.7743983 |
| TCGA-FC-A66V | TCGA-PRAD | 36.5109276 | 16.1900782 |
| TCGA-FC-A6HD | TCGA-PRAD | 32.7523202 | 14.7777105 |
| TCGA-FC-A8O0 | TCGA-PRAD | 44.0511882 | 16.6181867 |
| TCGA-FD-A3B3 | TCGA-BLCA | 88.5402974 | 56.5681017 |

| | | | |
|---|---|---|---|
| TCGA-FD-A3B4 | TCGA-BLCA | 140.34882 | 50.9193123 |
| TCGA-FD-A3B5 | TCGA-BLCA | 114.599628 | 23.8031413 |
| TCGA-FD-A3B6 | TCGA-BLCA | 98.1911253 | 74.7092685 |
| TCGA-FD-A3B7 | TCGA-BLCA | 69.582092 | 26.0026566 |
| TCGA-FD-A3B8 | TCGA-BLCA | 60.5073167 | 43.4746994 |
| TCGA-FD-A3N5 | TCGA-BLCA | 64.6261332 | 32.9495727 |
| TCGA-FD-A3N6 | TCGA-BLCA | 62.9047211 | 26.2557215 |
| TCGA-FD-A3NA | TCGA-BLCA | 92.0637755 | 32.6451007 |
| TCGA-FD-A3SJ | TCGA-BLCA | 138.785437 | 29.8819782 |
| TCGA-FD-A3SL | TCGA-BLCA | 69.4666022 | 17.1721352 |
| TCGA-FD-A3SM | TCGA-BLCA | 59.0932326 | 22.6156623 |
| TCGA-FD-A3SN | TCGA-BLCA | 88.9847137 | 32.2529286 |
| TCGA-FD-A3SO | TCGA-BLCA | 93.600313 | 30.2394765 |
| TCGA-FD-A3SP | TCGA-BLCA | 71.6896248 | 34.4857582 |
| TCGA-FD-A3SQ | TCGA-BLCA | 38.9336259 | 13.596313 |
| TCGA-FD-A3SR | TCGA-BLCA | 96.6945091 | 19.8719223 |
| TCGA-FD-A3SS | TCGA-BLCA | 110.878716 | 35.2502695 |
| TCGA-FD-A43N | TCGA-BLCA | 45.1701328 | 16.6029981 |
| TCGA-FD-A43P | TCGA-BLCA | 82.10069 | 56.1220228 |
| TCGA-FD-A43S | TCGA-BLCA | 43.3810023 | 14.4478808 |
| TCGA-FD-A43U | TCGA-BLCA | 37.8439012 | 15.4257651 |
| TCGA-FD-A43X | TCGA-BLCA | 74.8560778 | 22.6400867 |
| TCGA-FD-A43Y | TCGA-BLCA | 415.417414 | 26.8825243 |
| TCGA-FD-A5BR | TCGA-BLCA | 47.7955672 | 15.9147353 |
| TCGA-FD-A5BS | TCGA-BLCA | 23.3974381 | 26.5690017 |
| TCGA-FD-A5BT | TCGA-BLCA | 39.6636381 | 19.0185343 |
| TCGA-FD-A5BU | TCGA-BLCA | 109.447344 | 19.9762498 |
| TCGA-FD-A5BV | TCGA-BLCA | 64.0995059 | 27.4708444 |
| TCGA-FD-A5BX | TCGA-BLCA | 49.1175965 | 17.2031545 |
| TCGA-FD-A5BY | TCGA-BLCA | 151.9639 | 45.9515446 |
| TCGA-FD-A5BZ | TCGA-BLCA | 62.4988886 | 15.9872416 |

| | | | |
|---|---|---|---|
| TCGA-FD-A5C0 | TCGA-BLCA | 130.380461 | 18.0225082 |
| TCGA-FD-A5C1 | TCGA-BLCA | 97.8484978 | 34.0799213 |
| TCGA-FD-A62N | TCGA-BLCA | 59.1063842 | 20.1383426 |
| TCGA-FD-A62O | TCGA-BLCA | 41.9586932 | 12.9626991 |
| TCGA-FD-A62P | TCGA-BLCA | 53.1141229 | 36.98153 |
| TCGA-FD-A62S | TCGA-BLCA | 76.7558261 | 32.2516577 |
| TCGA-FD-A6TA | TCGA-BLCA | 72.3294872 | 21.6354767 |
| TCGA-FD-A6TB | TCGA-BLCA | 64.006149 | 48.4510058 |
| TCGA-FD-A6TC | TCGA-BLCA | 102.838369 | 24.960383 |
| TCGA-FD-A6TD | TCGA-BLCA | 101.951323 | 42.1732833 |
| TCGA-FD-A6TE | TCGA-BLCA | 96.8485038 | 27.629013 |
| TCGA-FD-A6TF | TCGA-BLCA | 98.1635969 | 56.2925763 |
| TCGA-FD-A6TG | TCGA-BLCA | 51.3208016 | 25.0317375 |
| TCGA-FD-A6TH | TCGA-BLCA | 83.5668003 | 25.4544925 |
| TCGA-FD-A6TI | TCGA-BLCA | 39.2886278 | 16.7529013 |
| TCGA-FD-A6TK | TCGA-BLCA | 78.8188183 | 45.6416932 |
| TCGA-FE-A22Z | TCGA-THCA | 43.2624203 | 11.5120548 |
| TCGA-FE-A230 | TCGA-THCA | 24.591371 | 6.18703067 |
| TCGA-FE-A231 | TCGA-THCA | 38.5204475 | 12.7170166 |
| TCGA-FE-A232 | TCGA-THCA | 49.8248255 | 14.3635628 |
| TCGA-FE-A233 | TCGA-THCA | 41.5733239 | 11.7252861 |
| TCGA-FE-A234 | TCGA-THCA | 58.3760806 | 21.7190689 |
| TCGA-FE-A235 | TCGA-THCA | 76.3890313 | 17.6250015 |
| TCGA-FE-A236 | TCGA-THCA | 42.6788445 | 12.0063492 |
| TCGA-FE-A237 | TCGA-THCA | 58.6532282 | 13.0297484 |
| TCGA-FE-A238 | TCGA-THCA | 54.7930971 | 16.307146 |
| TCGA-FE-A239 | TCGA-THCA | 86.0287577 | 14.9414324 |
| TCGA-FE-A23A | TCGA-THCA | 31.1906689 | 10.4793193 |
| TCGA-FE-A3PA | TCGA-THCA | 78.779601 | 17.6434856 |
| TCGA-FE-A3PB | TCGA-THCA | 56.5559266 | 14.3460052 |
| TCGA-FE-A3PD | TCGA-THCA | 96.1281806 | 16.9706963 |

| | | | |
|---|---|---|---|
| TCGA-FG-5962 | TCGA-LGG | 22.8538978 | 19.7593643 |
| TCGA-FG-5963 | TCGA-LGG | 36.4385671 | 15.0821899 |
| TCGA-FG-5964 | TCGA-LGG | 25.3534359 | 10.8636755 |
| TCGA-FG-5965 | TCGA-LGG | 34.1281101 | 12.4054676 |
| TCGA-FG-6688 | TCGA-LGG | 38.8046847 | 15.0395673 |
| TCGA-FG-6689 | TCGA-LGG | 31.6487914 | 9.06472733 |
| TCGA-FG-6690 | TCGA-LGG | 25.5646758 | 10.1089184 |
| TCGA-FG-6691 | TCGA-LGG | 29.3424243 | 16.4503415 |
| TCGA-FG-6692 | TCGA-LGG | 42.1922786 | 23.8200494 |
| TCGA-FG-7634 | TCGA-LGG | 23.8208754 | 21.8643617 |
| TCGA-FG-7636 | TCGA-LGG | 26.3975805 | 11.0334801 |
| TCGA-FG-7637 | TCGA-LGG | 18.4082441 | 16.9452791 |
| TCGA-FG-7638 | TCGA-LGG | 18.8999129 | 12.6655702 |
| TCGA-FG-7641 | TCGA-LGG | 22.0984097 | 18.9538968 |
| TCGA-FG-7643 | TCGA-LGG | 30.8141305 | 9.89762665 |
| TCGA-FG-8181 | TCGA-LGG | 30.9463501 | 7.8834469 |
| TCGA-FG-8182 | TCGA-LGG | 33.2034204 | 14.6399362 |
| TCGA-FG-8185 | TCGA-LGG | 29.788699 | 14.1057678 |
| TCGA-FG-8186 | TCGA-LGG | 24.9565754 | 15.9868006 |
| TCGA-FG-8187 | TCGA-LGG | 19.6063802 | 17.4435452 |
| TCGA-FG-8188 | TCGA-LGG | 30.3604467 | 10.3139445 |
| TCGA-FG-8189 | TCGA-LGG | 25.8450777 | 7.89394801 |
| TCGA-FG-8191 | TCGA-LGG | 32.7946962 | 12.0346866 |
| TCGA-FG-A4MT | TCGA-LGG | 37.2174905 | 12.6992263 |
| TCGA-FG-A4MU | TCGA-LGG | 44.8205278 | 18.7403677 |
| TCGA-FG-A4MW | TCGA-LGG | 28.7562874 | 12.7270117 |
| TCGA-FG-A4MX | TCGA-LGG | 20.427179 | 8.48813326 |
| TCGA-FG-A4MY | TCGA-LGG | 34.2686863 | 11.999142 |
| TCGA-FG-A60J | TCGA-LGG | 28.6679567 | 13.7872156 |
| TCGA-FG-A60K | TCGA-LGG | 18.6917932 | 19.7091327 |
| TCGA-FG-A60L | TCGA-LGG | 29.7375751 | 14.4465974 |

| | | | |
|---|---|---|---|
| TCGA-FG-A6IZ | TCGA-LGG | 22.8824339 | 29.4906046 |
| TCGA-FG-A6J1 | TCGA-LGG | 18.6599641 | 21.2494039 |
| TCGA-FG-A6J3 | TCGA-LGG | 57.4399072 | 54.5232108 |
| TCGA-FG-A70Y | TCGA-LGG | 26.7521209 | 9.7789736 |
| TCGA-FG-A70Z | TCGA-LGG | 45.720811 | 25.8609964 |
| TCGA-FG-A710 | TCGA-LGG | 29.0320154 | 22.3737782 |
| TCGA-FG-A711 | TCGA-LGG | 44.5238873 | 9.02979603 |
| TCGA-FG-A713 | TCGA-LGG | 21.6785539 | 6.81563663 |
| TCGA-FG-A87N | TCGA-LGG | 49.4297655 | 15.1547727 |
| TCGA-FG-A87Q | TCGA-LGG | 39.8826292 | 16.6818507 |
| TCGA-FI-A2EY | TCGA-UCEC | 22.7456806 | 15.9791982 |
| TCGA-FI-A3PV | TCGA-UCEC | 123.169991 | 17.6255497 |
| TCGA-FI-A3PX | TCGA-UCEC | 112.477573 | 64.5605234 |
| TCGA-FJ-A3Z7 | TCGA-BLCA | 28.9732894 | 10.874893 |
| TCGA-FJ-A3Z9 | TCGA-BLCA | 47.6676592 | 19.597956 |
| TCGA-FJ-A3ZE | TCGA-BLCA | 77.727115 | 39.2022144 |
| TCGA-FJ-A3ZF | TCGA-BLCA | 78.3421212 | 40.8083222 |
| TCGA-FJ-A871 | TCGA-BLCA | 76.5756702 | 42.2517028 |
| TCGA-FK-A3S3 | TCGA-THCA | 65.0690906 | 12.8892927 |
| TCGA-FK-A3SB | TCGA-THCA | 67.9864512 | 15.4203886 |
| TCGA-FK-A3SD | TCGA-THCA | 63.719974 | 16.4072837 |
| TCGA-FK-A3SE | TCGA-THCA | 71.4262243 | 17.7977362 |
| TCGA-FK-A3SG | TCGA-THCA | 81.0798232 | 19.3434592 |
| TCGA-FK-A3SH | TCGA-THCA | 66.9200527 | 15.6974897 |
| TCGA-FK-A4UB | TCGA-THCA | 53.201137 | 14.8189689 |
| TCGA-FN-7833 | TCGA-LGG | 38.1070724 | 15.3595896 |
| TCGA-FP-7735 | TCGA-STAD | 28.5812449 | 15.9000422 |
| TCGA-FP-7829 | TCGA-STAD | 35.8540019 | 45.6264717 |
| TCGA-FP-7916 | TCGA-STAD | 35.9466058 | 17.0589171 |
| TCGA-FP-7998 | TCGA-STAD | 35.9231485 | 14.6189118 |
| TCGA-FP-8099 | TCGA-STAD | 73.7239913 | 37.4520621 |

| | | | |
|---|---|---|---|
| TCGA-FP-8209 | TCGA-STAD | 33.9163893 | 13.6909962 |
| TCGA-FP-8210 | TCGA-STAD | 46.8160507 | 18.7704431 |
| TCGA-FP-8211 | TCGA-STAD | 79.7679394 | 37.0877456 |
| TCGA-FP-8631 | TCGA-STAD | 47.7203986 | 22.7249929 |
| TCGA-FP-A8CX | TCGA-STAD | 31.0918131 | 24.3647289 |
| TCGA-FP-A9TM | TCGA-STAD | 37.131303 | 24.1872105 |
| TCGA-FR-A2OS | TCGA-SKCM | 56.2789945 | 25.8660444 |
| TCGA-FR-A3R1 | TCGA-SKCM | 48.9045145 | 26.3165838 |
| TCGA-FR-A3YN | TCGA-SKCM | 62.2768679 | 13.3447918 |
| TCGA-FR-A3YO | TCGA-SKCM | 37.136001 | 13.792819 |
| TCGA-FR-A44A | TCGA-SKCM | 55.3408077 | 20.8050904 |
| TCGA-FR-A69P | TCGA-SKCM | 36.6114496 | 31.1032519 |
| TCGA-FR-A726 | TCGA-SKCM | 65.756813 | 18.8585867 |
| TCGA-FR-A728 | TCGA-SKCM | 33.4615676 | 9.94010406 |
| TCGA-FR-A729 | TCGA-SKCM | 45.9445021 | 22.0358631 |
| TCGA-FR-A7U8 | TCGA-SKCM | 66.6197542 | 19.8550858 |
| TCGA-FR-A7U9 | TCGA-SKCM | 115.922311 | 51.3584799 |
| TCGA-FR-A7UA | TCGA-SKCM | 54.8641447 | 19.2543675 |
| TCGA-FR-A8YC | TCGA-SKCM | 34.6656477 | 18.5499051 |
| TCGA-FR-A8YD | TCGA-SKCM | 39.78942 | 9.47958216 |
| TCGA-FR-A8YE | TCGA-SKCM | 46.7139694 | 21.3014855 |
| TCGA-FS-A1YW | TCGA-SKCM | 36.1585458 | 13.1442922 |
| TCGA-FS-A1YX | TCGA-SKCM | 50.8572998 | 27.490248 |
| TCGA-FS-A1YY | TCGA-SKCM | 43.9766571 | 33.3590072 |
| TCGA-FS-A1Z0 | TCGA-SKCM | 61.6428382 | 36.1610786 |
| TCGA-FS-A1Z3 | TCGA-SKCM | 49.93767 | 22.9360126 |
| TCGA-FS-A1Z4 | TCGA-SKCM | 58.123603 | 23.1572748 |
| TCGA-FS-A1Z7 | TCGA-SKCM | 40.7830567 | 28.4603455 |
| TCGA-FS-A1ZA | TCGA-SKCM | 43.1114257 | 20.5581919 |
| TCGA-FS-A1ZB | TCGA-SKCM | 43.8802545 | 13.0714421 |
| TCGA-FS-A1ZD | TCGA-SKCM | 66.6714169 | 26.0947177 |

| | | | |
|---|---|---|---|
| TCGA-FS-A1ZE | TCGA-SKCM | 31.6743469 | 18.2632533 |
| TCGA-FS-A1ZG | TCGA-SKCM | 37.8725915 | 19.2781582 |
| TCGA-FS-A1ZJ | TCGA-SKCM | 89.8116926 | 24.0188394 |
| TCGA-FS-A1ZK | TCGA-SKCM | 33.5830306 | 20.4298009 |
| TCGA-FS-A1ZM | TCGA-SKCM | 35.9677574 | 53.5374508 |
| TCGA-FS-A1ZN | TCGA-SKCM | 46.6149455 | 26.4451509 |
| TCGA-FS-A1ZP | TCGA-SKCM | 44.0883633 | 12.4575787 |
| TCGA-FS-A1ZQ | TCGA-SKCM | 36.8139489 | 21.5865768 |
| TCGA-FS-A1ZR | TCGA-SKCM | 44.6261664 | 24.3971977 |
| TCGA-FS-A1ZS | TCGA-SKCM | 36.0470954 | 20.0416384 |
| TCGA-FS-A1ZT | TCGA-SKCM | 31.0719291 | 8.57632924 |
| TCGA-FS-A1ZU | TCGA-SKCM | 56.7833095 | 20.0405853 |
| TCGA-FS-A1ZW | TCGA-SKCM | 25.7436353 | 12.884379 |
| TCGA-FS-A1ZY | TCGA-SKCM | 30.0208587 | 14.1239492 |
| TCGA-FS-A1ZZ | TCGA-SKCM | 52.7197652 | 29.4535376 |
| TCGA-FS-A4F0 | TCGA-SKCM | 75.1927599 | 22.5047066 |
| TCGA-FS-A4F2 | TCGA-SKCM | 66.0673818 | 31.018533 |
| TCGA-FS-A4F4 | TCGA-SKCM | 40.6941582 | 14.7671668 |
| TCGA-FS-A4F5 | TCGA-SKCM | 68.0584348 | 21.6489459 |
| TCGA-FS-A4F8 | TCGA-SKCM | 39.204765 | 17.115638 |
| TCGA-FS-A4F9 | TCGA-SKCM | 46.2343757 | 29.0129144 |
| TCGA-FS-A4FB | TCGA-SKCM | 50.6881356 | 19.9891686 |
| TCGA-FS-A4FC | TCGA-SKCM | 47.7830933 | 18.0306115 |
| TCGA-FS-A4FD | TCGA-SKCM | 44.1567303 | 20.8388971 |
| TCGA-FT-A3EE | TCGA-BLCA | 37.8812788 | 20.7630598 |
| TCGA-FT-A61P | TCGA-BLCA | 69.416759 | 31.0800961 |
| TCGA-FV-A23B | TCGA-LIHC | 27.4736656 | 7.91257192 |
| TCGA-FV-A2QQ | TCGA-LIHC | 23.0482037 | 22.6268618 |
| TCGA-FV-A2QR | TCGA-LIHC | 44.1542131 | 28.2854811 |
| TCGA-FV-A3I0 | TCGA-LIHC | 35.6518592 | 32.9059203 |
| TCGA-FV-A3I1 | TCGA-LIHC | 35.4168161 | 16.1131566 |

| | | | |
|---|---|---|---|
| TCGA-FV-A3R2 | TCGA-LIHC | 60.4374426 | 11.060333 |
| TCGA-FV-A3R3 | TCGA-LIHC | 21.1179683 | 6.39068632 |
| TCGA-FV-A495 | TCGA-LIHC | 36.5065538 | 20.7811535 |
| TCGA-FV-A496 | TCGA-LIHC | 28.2902873 | 11.0236317 |
| TCGA-FV-A4ZP | TCGA-LIHC | 10.8723444 | 13.7603895 |
| TCGA-FV-A4ZQ | TCGA-LIHC | 61.8978411 | 23.5819952 |
| TCGA-FW-A3I3 | TCGA-SKCM | 101.272859 | 20.203871 |
| TCGA-FW-A3R5 | TCGA-SKCM | 78.0080203 | 42.8606073 |
| TCGA-FW-A3TU | TCGA-SKCM | 23.7280486 | 9.02153931 |
| TCGA-FW-A3TV | TCGA-SKCM | 22.5485318 | 25.0997283 |
| TCGA-FW-A5DX | TCGA-SKCM | 69.4731049 | 28.9763807 |
| TCGA-FW-A5DY | TCGA-SKCM | 38.2355549 | 23.5174319 |
| TCGA-FY-A2QD | TCGA-THCA | 97.4746958 | 30.5791481 |
| TCGA-FY-A3BL | TCGA-THCA | 43.3085428 | 8.87889687 |
| TCGA-FY-A3I4 | TCGA-THCA | 49.3003934 | 11.846263 |
| TCGA-FY-A3I5 | TCGA-THCA | 46.7460094 | 12.8683646 |
| TCGA-FY-A3NM | TCGA-THCA | 87.7911199 | 25.0513238 |
| TCGA-FY-A3NN | TCGA-THCA | 41.3453476 | 15.4463639 |
| TCGA-FY-A3NP | TCGA-THCA | 55.7410661 | 18.1574629 |
| TCGA-FY-A3ON | TCGA-THCA | 75.7993657 | 20.8486626 |
| TCGA-FY-A3R6 | TCGA-THCA | 52.087491 | 11.8597034 |
| TCGA-FY-A3R7 | TCGA-THCA | 73.7355769 | 18.7884454 |
| TCGA-FY-A3R8 | TCGA-THCA | 64.9400324 | 18.101754 |
| TCGA-FY-A3R9 | TCGA-THCA | 40.9154878 | 10.5269311 |
| TCGA-FY-A3RA | TCGA-THCA | 64.6901258 | 13.9671478 |
| TCGA-FY-A3TY | TCGA-THCA | 73.238281 | 14.658071 |
| TCGA-FY-A3W9 | TCGA-THCA | 72.2676111 | 19.6265466 |
| TCGA-FY-A3WA | TCGA-THCA | 76.0344864 | 9.63709606 |
| TCGA-FY-A3YR | TCGA-THCA | 44.2564064 | 12.3602809 |
| TCGA-FY-A40K | TCGA-THCA | 66.228782 | 12.6702453 |
| TCGA-FY-A40L | TCGA-THCA | 45.4570578 | 13.2856105 |

| | | | |
|---|---|---|---|
| TCGA-FY-A40M | TCGA-THCA | 101.619534 | 18.2524081 |
| TCGA-FY-A40N | TCGA-THCA | 73.9044032 | 20.5985711 |
| TCGA-FY-A4B0 | TCGA-THCA | 57.4675447 | 18.2117851 |
| TCGA-FY-A4B3 | TCGA-THCA | 49.0618813 | 15.4988107 |
| TCGA-FY-A4B4 | TCGA-THCA | 46.0986614 | 12.1444886 |
| TCGA-FY-A76V | TCGA-THCA | 58.7156779 | 13.752707 |
| TCGA-G2-A2EC | TCGA-BLCA | 43.2067134 | 14.8167754 |
| TCGA-G2-A2EF | TCGA-BLCA | 175.844482 | 50.4317781 |
| TCGA-G2-A2EJ | TCGA-BLCA | 81.0028802 | 36.5666407 |
| TCGA-G2-A2EK | TCGA-BLCA | 45.6024693 | 11.3162242 |
| TCGA-G2-A2EL | TCGA-BLCA | 227.066891 | 60.3716955 |
| TCGA-G2-A2EO | TCGA-BLCA | 125.89058 | 39.2098773 |
| TCGA-G2-A2ES | TCGA-BLCA | 94.2154887 | 21.2847336 |
| TCGA-G2-A3IB | TCGA-BLCA | 58.8923163 | 30.6245452 |
| TCGA-G2-A3IE | TCGA-BLCA | 70.9482699 | 19.5716366 |
| TCGA-G2-A3VY | TCGA-BLCA | 52.9803358 | 30.7797919 |
| TCGA-G2-AA3B | TCGA-BLCA | 67.7812385 | 63.4980947 |
| TCGA-G2-AA3C | TCGA-BLCA | 61.8470622 | 43.0584297 |
| TCGA-G2-AA3D | TCGA-BLCA | 65.5864859 | 15.0576101 |
| TCGA-G2-AA3F | TCGA-BLCA | 37.3379918 | 22.5528179 |
| TCGA-G3-A25S | TCGA-LIHC | 40.503379 | 25.9476187 |
| TCGA-G3-A25T | TCGA-LIHC | 46.8674378 | 19.1580212 |
| TCGA-G3-A25U | TCGA-LIHC | 48.9276206 | 14.2002503 |
| TCGA-G3-A25V | TCGA-LIHC | 36.5367357 | 14.7956681 |
| TCGA-G3-A25X | TCGA-LIHC | 59.5173017 | 32.8972647 |
| TCGA-G3-A25Y | TCGA-LIHC | 55.0001018 | 26.4117547 |
| TCGA-G3-A25Z | TCGA-LIHC | 41.8537424 | 7.09569171 |
| TCGA-G3-A3CG | TCGA-LIHC | 35.6954217 | 19.0129018 |
| TCGA-G3-A3CH | TCGA-LIHC | 20.6061891 | 7.93999441 |
| TCGA-G3-A3CI | TCGA-LIHC | 22.8423611 | 5.73981968 |
| TCGA-G3-A3CJ | TCGA-LIHC | 35.0641734 | 12.3911338 |

| | | | |
|---|---|---|---|
| TCGA-G3-A3CK | TCGA-LIHC | 38.0988954 | 12.438984 |
| TCGA-G3-A5SI | TCGA-LIHC | 45.6590049 | 19.85306 |
| TCGA-G3-A5SJ | TCGA-LIHC | 96.7686013 | 14.840779 |
| TCGA-G3-A5SK | TCGA-LIHC | 23.588584 | 15.2944171 |
| TCGA-G3-A5SL | TCGA-LIHC | 35.223855 | 14.9443673 |
| TCGA-G3-A5SM | TCGA-LIHC | 25.1872764 | 16.5859533 |
| TCGA-G3-A6UC | TCGA-LIHC | 47.5523491 | 20.5766144 |
| TCGA-G3-A7M5 | TCGA-LIHC | 31.0808119 | 12.5506622 |
| TCGA-G3-A7M6 | TCGA-LIHC | 76.1971087 | 17.0646816 |
| TCGA-G3-A7M7 | TCGA-LIHC | 33.0565057 | 13.1867943 |
| TCGA-G3-A7M8 | TCGA-LIHC | 29.6462609 | 12.7941299 |
| TCGA-G3-A7M9 | TCGA-LIHC | 157.215088 | 41.2888566 |
| TCGA-G3-AAUZ | TCGA-LIHC | 28.2991989 | 10.6279279 |
| TCGA-G3-AAV0 | TCGA-LIHC | 28.6214129 | 13.8639607 |
| TCGA-G3-AAV1 | TCGA-LIHC | 49.7097243 | 17.9045316 |
| TCGA-G3-AAV2 | TCGA-LIHC | 49.3985256 | 17.7859271 |
| TCGA-G3-AAV3 | TCGA-LIHC | 44.8212218 | 14.8076733 |
| TCGA-G3-AAV4 | TCGA-LIHC | 75.876597 | 13.4949835 |
| TCGA-G3-AAV5 | TCGA-LIHC | 47.7683735 | 25.3048663 |
| TCGA-G3-AAV6 | TCGA-LIHC | 45.3934309 | 33.2081895 |
| TCGA-G3-AAV7 | TCGA-LIHC | 84.320003 | 21.6958893 |
| TCGA-G4-6293 | TCGA-COAD | 47.1275989 | 14.0652526 |
| TCGA-G4-6294 | TCGA-COAD | 65.2866096 | 23.7147922 |
| TCGA-G4-6295 | TCGA-COAD | 86.3210053 | 28.9972432 |
| TCGA-G4-6297 | TCGA-COAD | 56.2215787 | 35.26527 |
| TCGA-G4-6298 | TCGA-COAD | 29.7911747 | 22.9057912 |
| TCGA-G4-6299 | TCGA-COAD | 51.4028157 | 21.5801171 |
| TCGA-G4-6302 | TCGA-COAD | 41.0645592 | 17.418006 |
| TCGA-G4-6303 | TCGA-COAD | 40.2303415 | 26.4654381 |
| TCGA-G4-6304 | TCGA-COAD | 55.0295284 | 12.874224 |
| TCGA-G4-6306 | TCGA-COAD | 67.5105535 | 27.9961189 |

| | | | |
|---|---|---|---|
| TCGA-G4-6307 | TCGA-COAD | 77.004121 | 26.477548 |
| TCGA-G4-6309 | TCGA-COAD | 107.441741 | 37.9745956 |
| TCGA-G4-6310 | TCGA-COAD | 73.7585305 | 20.6526182 |
| TCGA-G4-6311 | TCGA-COAD | 59.1642222 | 21.6112103 |
| TCGA-G4-6314 | TCGA-COAD | 40.1025349 | 29.7684131 |
| TCGA-G4-6315 | TCGA-COAD | 69.3467867 | 25.1513852 |
| TCGA-G4-6317 | TCGA-COAD | 77.217945 | 19.3645469 |
| TCGA-G4-6320 | TCGA-COAD | 53.3895604 | 16.5648546 |
| TCGA-G4-6321 | TCGA-COAD | 26.0598059 | 7.25546314 |
| TCGA-G4-6322 | TCGA-COAD | 62.0002539 | 23.1876617 |
| TCGA-G4-6323 | TCGA-COAD | 67.8957449 | 17.5821044 |
| TCGA-G4-6586 | TCGA-COAD | 57.894914 | 22.2215463 |
| TCGA-G4-6588 | TCGA-COAD | 63.6454414 | 20.4891562 |
| TCGA-G4-6625 | TCGA-COAD | 61.6424687 | 32.2394371 |
| TCGA-G4-6626 | TCGA-COAD | 60.1231132 | 39.2049217 |
| TCGA-G4-6627 | TCGA-COAD | 48.2157619 | 22.6819076 |
| TCGA-G4-6628 | TCGA-COAD | 73.5900911 | 27.039657 |
| TCGA-G6-A5PC | TCGA-KIRC | 31.9430907 | 14.4445359 |
| TCGA-G6-A8L6 | TCGA-KIRC | 58.1201137 | 20.4690434 |
| TCGA-G6-A8L7 | TCGA-KIRC | 38.0150218 | 20.4880291 |
| TCGA-G6-A8L8 | TCGA-KIRC | 44.5193194 | 21.6636752 |
| TCGA-G7-6789 | TCGA-KIRP | 11.2463792 | 9.49789478 |
| TCGA-G7-6790 | TCGA-KIRP | 4.54739955 | 2.14038882 |
| TCGA-G7-6792 | TCGA-KIRP | 17.6058564 | 5.66374527 |
| TCGA-G7-6793 | TCGA-KIRP | 14.9286974 | 1.1637127 |
| TCGA-G7-6795 | TCGA-KIRP | 3.28331549 | 0.65527124 |
| TCGA-G7-6796 | TCGA-KIRP | 4.56454903 | 4.71782145 |
| TCGA-G7-6797 | TCGA-KIRP | 15.7511074 | 13.5751593 |
| TCGA-G7-7501 | TCGA-KIRP | 44.082436 | 6.15249507 |
| TCGA-G7-7502 | TCGA-KIRP | 35.8622971 | 21.8291656 |
| TCGA-G7-A4TM | TCGA-KIRP | 12.2393573 | 23.9065846 |

| | | | |
|---|---|---|---|
| TCGA-G7-A8LB | TCGA-KIRP | 32.8739977 | 2.04836485 |
| TCGA-G7-A8LC | TCGA-KIRP | 20.9252587 | 21.5219849 |
| TCGA-G7-A8LD | TCGA-KIRP | 52.3537223 | 46.7282756 |
| TCGA-G7-A8LE | TCGA-KIRP | 51.1586164 | 4.42888324 |
| TCGA-G9-6329 | TCGA-PRAD | 44.8576473 | 13.8064472 |
| TCGA-G9-6332 | TCGA-PRAD | 49.4250191 | 25.0342461 |
| TCGA-G9-6333 | TCGA-PRAD | 34.1573713 | 10.5661321 |
| TCGA-G9-6336 | TCGA-PRAD | 41.9190491 | 17.1689414 |
| TCGA-G9-6338 | TCGA-PRAD | 23.200864 | 11.8653704 |
| TCGA-G9-6339 | TCGA-PRAD | 45.1389134 | 18.8783638 |
| TCGA-G9-6342 | TCGA-PRAD | 34.8692539 | 14.4188186 |
| TCGA-G9-6343 | TCGA-PRAD | 21.5591305 | 9.52722144 |
| TCGA-G9-6347 | TCGA-PRAD | 30.4588397 | 12.3720786 |
| TCGA-G9-6348 | TCGA-PRAD | 32.4860715 | 11.4756641 |
| TCGA-G9-6351 | TCGA-PRAD | 34.2523691 | 16.8537634 |
| TCGA-G9-6353 | TCGA-PRAD | 29.8674787 | 12.6559711 |
| TCGA-G9-6354 | TCGA-PRAD | 44.8597782 | 12.2784571 |
| TCGA-G9-6356 | TCGA-PRAD | 45.2476037 | 15.075723 |
| TCGA-G9-6361 | TCGA-PRAD | 34.0991221 | 14.7626456 |
| TCGA-G9-6362 | TCGA-PRAD | 55.8608598 | 24.541683 |
| TCGA-G9-6363 | TCGA-PRAD | 35.398361 | 15.6611749 |
| TCGA-G9-6364 | TCGA-PRAD | 19.8316206 | 5.99594542 |
| TCGA-G9-6365 | TCGA-PRAD | 41.8170449 | 13.4460145 |
| TCGA-G9-6366 | TCGA-PRAD | 21.8285206 | 11.8357562 |
| TCGA-G9-6367 | TCGA-PRAD | 30.2586007 | 14.1376996 |
| TCGA-G9-6369 | TCGA-PRAD | 32.6106741 | 23.192227 |
| TCGA-G9-6370 | TCGA-PRAD | 33.691033 | 12.5871834 |
| TCGA-G9-6371 | TCGA-PRAD | 29.0882818 | 15.7891473 |
| TCGA-G9-6373 | TCGA-PRAD | 50.9113202 | 14.8070557 |
| TCGA-G9-6377 | TCGA-PRAD | 40.0704753 | 11.7420681 |
| TCGA-G9-6378 | TCGA-PRAD | 30.2183745 | 13.4407405 |

| | | | |
|---|---|---|---|
| TCGA-G9-6379 | TCGA-PRAD | 29.4453438 | 11.5797402 |
| TCGA-G9-6384 | TCGA-PRAD | 33.909524 | 7.27016892 |
| TCGA-G9-6385 | TCGA-PRAD | 29.358247 | 11.6702159 |
| TCGA-G9-6494 | TCGA-PRAD | 38.4388605 | 21.062013 |
| TCGA-G9-6496 | TCGA-PRAD | 40.514471 | 20.2961816 |
| TCGA-G9-6498 | TCGA-PRAD | 31.0000841 | 12.9980981 |
| TCGA-G9-6499 | TCGA-PRAD | 37.3420533 | 17.7686417 |
| TCGA-G9-7509 | TCGA-PRAD | 41.3197339 | 19.8377897 |
| TCGA-G9-7510 | TCGA-PRAD | 32.5582848 | 12.7212256 |
| TCGA-G9-7519 | TCGA-PRAD | 32.5105543 | 17.4353982 |
| TCGA-G9-7521 | TCGA-PRAD | 54.1159956 | 11.1204704 |
| TCGA-G9-7522 | TCGA-PRAD | 46.8031841 | 13.912188 |
| TCGA-G9-7523 | TCGA-PRAD | 32.9902108 | 14.0504986 |
| TCGA-G9-7525 | TCGA-PRAD | 36.1623455 | 12.6622029 |
| TCGA-G9-A9S0 | TCGA-PRAD | 63.0516603 | 21.2282819 |
| TCGA-G9-A9S4 | TCGA-PRAD | 62.0662591 | 10.0273705 |
| TCGA-G9-A9S7 | TCGA-PRAD | 74.2634496 | 22.6802551 |
| TCGA-GC-A3BM | TCGA-BLCA | 83.315744 | 25.4867025 |
| TCGA-GC-A3I6 | TCGA-BLCA | 74.0750933 | 40.0246255 |
| TCGA-GC-A3OO | TCGA-BLCA | 71.9593123 | 38.4781953 |
| TCGA-GC-A3RB | TCGA-BLCA | 87.2856486 | 35.4597282 |
| TCGA-GC-A3RC | TCGA-BLCA | 77.0905999 | 42.7170529 |
| TCGA-GC-A3RD | TCGA-BLCA | 110.983655 | 20.9108415 |
| TCGA-GC-A3W( | TCGA-BLCA | 57.8021345 | 24.2166159 |
| TCGA-GC-A3YS | TCGA-BLCA | 145.215421 | 55.9302869 |
| TCGA-GC-A4ZW | TCGA-BLCA | 147.550774 | 37.1743807 |
| TCGA-GC-A6I1 | TCGA-BLCA | 60.8776411 | 24.4610022 |
| TCGA-GC-A6I3 | TCGA-BLCA | 66.7573746 | 28.8107895 |
| TCGA-GD-A2C5 | TCGA-BLCA | 73.9226458 | 13.4815116 |
| TCGA-GD-A3OP | TCGA-BLCA | 48.6939338 | 13.9001772 |
| TCGA-GD-A3OQ | TCGA-BLCA | 39.9033281 | 13.9229108 |

| | | | |
|---|---|---|---|
| TCGA-GD-A3OS | TCGA-BLCA | 105.805686 | 13.9262482 |
| TCGA-GD-A6C6 | TCGA-BLCA | 31.4305201 | 19.5419106 |
| TCGA-GD-A76B | TCGA-BLCA | 81.7370036 | 18.9034077 |
| TCGA-GE-A2C6 | TCGA-THCA | 47.1183913 | 15.0904182 |
| TCGA-GF-A2C7 | TCGA-SKCM | 46.8761267 | 22.0069628 |
| TCGA-GF-A3OT | TCGA-SKCM | 54.195698 | 37.2448567 |
| TCGA-GF-A4EO | TCGA-SKCM | 21.1185437 | 22.1635247 |
| TCGA-GF-A6C8 | TCGA-SKCM | 64.3121929 | 37.0028668 |
| TCGA-GF-A6C9 | TCGA-SKCM | 69.8031065 | 15.2373445 |
| TCGA-GF-A769 | TCGA-SKCM | 43.1134386 | 19.9191559 |
| TCGA-GI-A2C8 | TCGA-BRCA | 33.9807233 | 32.1099385 |
| TCGA-GI-A2C9 | TCGA-BRCA | 83.9246257 | 76.7304438 |
| TCGA-GJ-A3OU | TCGA-LIHC | 32.3076349 | 26.5231783 |
| TCGA-GJ-A6C0 | TCGA-LIHC | 50.9515166 | 10.9436956 |
| TCGA-GJ-A9DB | TCGA-LIHC | 42.2100997 | 20.1754908 |
| TCGA-GK-A6C7 | TCGA-KIRC | 60.7145444 | 22.6955738 |
| TCGA-GL-7773 | TCGA-KIRP | 63.5810973 | 5.20692732 |
| TCGA-GL-8500 | TCGA-KIRP | 19.1113114 | 26.0501734 |
| TCGA-GL-A4EM | TCGA-KIRP | 3.0762576 | 1.22424895 |
| TCGA-GL-A59R | TCGA-KIRP | 34.3167321 | 16.450179 |
| TCGA-GL-A59T | TCGA-KIRP | 29.6814612 | 17.9542553 |
| TCGA-GL-A9DC | TCGA-KIRP | 38.8280061 | 25.7810962 |
| TCGA-GL-A9DD | TCGA-KIRP | 9.11262088 | 5.48729423 |
| TCGA-GL-A9DE | TCGA-KIRP | 42.0070619 | 10.8986561 |
| TCGA-GM-A2D9 | TCGA-BRCA | 65.2940745 | 28.4995864 |
| TCGA-GM-A2DA | TCGA-BRCA | 41.7594507 | 21.5567227 |
| TCGA-GM-A2DB | TCGA-BRCA | 67.5447755 | 40.0653018 |
| TCGA-GM-A2DC | TCGA-BRCA | 58.3731577 | 32.0015703 |
| TCGA-GM-A2DD | TCGA-BRCA | 65.4153778 | 41.3476069 |
| TCGA-GM-A2DF | TCGA-BRCA | 64.7132894 | 19.674595 |
| TCGA-GM-A2DH | TCGA-BRCA | 126.752458 | 39.1470607 |

| | | | |
|---|---|---|---|
| TCGA-GM-A2DI | TCGA-BRCA | 54.2034884 | 12.1660294 |
| TCGA-GM-A2DK | TCGA-BRCA | 80.0919879 | 23.3800555 |
| TCGA-GM-A2DL | TCGA-BRCA | 66.917566 | 20.2261462 |
| TCGA-GM-A2DM | TCGA-BRCA | 37.2185775 | 21.3250587 |
| TCGA-GM-A2DN | TCGA-BRCA | 45.6284985 | 22.4256688 |
| TCGA-GM-A2DC | TCGA-BRCA | 54.960216 | 19.1857742 |
| TCGA-GM-A3NW | TCGA-BRCA | 40.1643424 | 28.0857555 |
| TCGA-GM-A3NY | TCGA-BRCA | 83.0714445 | 60.6655352 |
| TCGA-GM-A3XG | TCGA-BRCA | 60.8825227 | 15.8332048 |
| TCGA-GM-A3XL | TCGA-BRCA | 157.291129 | 82.465237 |
| TCGA-GM-A3XN | TCGA-BRCA | 51.8014525 | 18.5533179 |
| TCGA-GM-A4E0 | TCGA-BRCA | 35.1926224 | 15.4281709 |
| TCGA-GM-A5PV | TCGA-BRCA | 75.0973725 | 20.102786 |
| TCGA-GM-A5PX | TCGA-BRCA | 69.6443821 | 34.9590545 |
| TCGA-GN-A262 | TCGA-SKCM | 40.2523929 | 15.0721193 |
| TCGA-GN-A263 | TCGA-SKCM | 57.7905095 | 34.0239545 |
| TCGA-GN-A264 | TCGA-SKCM | 41.9806216 | 21.8674136 |
| TCGA-GN-A266 | TCGA-SKCM | 75.4825202 | 22.6560152 |
| TCGA-GN-A267 | TCGA-SKCM | 42.6930271 | 26.2208295 |
| TCGA-GN-A268 | TCGA-SKCM | 63.9855582 | 26.6442119 |
| TCGA-GN-A26A | TCGA-SKCM | 51.4410749 | 40.6338315 |
| TCGA-GN-A26C | TCGA-SKCM | 75.3004579 | 14.0517709 |
| TCGA-GN-A26D | TCGA-SKCM | 22.3233219 | 13.3245442 |
| TCGA-GN-A4U3 | TCGA-SKCM | 97.3405325 | 45.1818917 |
| TCGA-GN-A4U4 | TCGA-SKCM | 45.2631745 | 18.0364394 |
| TCGA-GN-A4U5 | TCGA-SKCM | 51.6877079 | 20.6360719 |
| TCGA-GN-A4U7 | TCGA-SKCM | 58.4278061 | 18.8267344 |
| TCGA-GN-A4U8 | TCGA-SKCM | 50.6072424 | 19.9102685 |
| TCGA-GN-A4U9 | TCGA-SKCM | 154.58852 | 57.180721 |
| TCGA-GN-A8LK | TCGA-SKCM | 87.4891021 | 21.6821521 |
| TCGA-GN-A8LL | TCGA-SKCM | 45.9513786 | 42.579705 |

| | | | |
|---|---|---|---|
| TCGA-GN-A8LN | TCGA-SKCM | 101.068801 | 34.9466931 |
| TCGA-GN-A9SD | TCGA-SKCM | 59.0304576 | 29.645052 |
| TCGA-GU-A42P | TCGA-BLCA | 98.7004099 | 17.5845548 |
| TCGA-GU-A42Q | TCGA-BLCA | 60.6707852 | 27.7469596 |
| TCGA-GU-A42R | TCGA-BLCA | 64.513865 | 22.9873129 |
| TCGA-GU-A762 | TCGA-BLCA | 90.6309365 | 55.6029941 |
| TCGA-GU-A763 | TCGA-BLCA | 80.1575679 | 15.6960308 |
| TCGA-GU-A764 | TCGA-BLCA | 73.9025744 | 25.1326863 |
| TCGA-GU-A766 | TCGA-BLCA | 84.2942368 | 49.5822533 |
| TCGA-GU-A767 | TCGA-BLCA | 68.3568491 | 22.992776 |
| TCGA-GU-AATO | TCGA-BLCA | 62.2369557 | 37.4505843 |
| TCGA-GU-AATP | TCGA-BLCA | 59.6199396 | 45.1521261 |
| TCGA-GU-AATQ | TCGA-BLCA | 78.9035819 | 25.4626177 |
| TCGA-GV-A3JV | TCGA-BLCA | 62.3715235 | 27.7599687 |
| TCGA-GV-A3JW | TCGA-BLCA | 79.0096322 | 8.89731658 |
| TCGA-GV-A3JX | TCGA-BLCA | 89.5811038 | 28.3189373 |
| TCGA-GV-A3JZ | TCGA-BLCA | 81.0921166 | 61.0901337 |
| TCGA-GV-A3QF | TCGA-BLCA | 57.2714885 | 23.7133405 |
| TCGA-GV-A3QG | TCGA-BLCA | 54.689111 | 26.415157 |
| TCGA-GV-A3QH | TCGA-BLCA | 71.1244818 | 23.9598021 |
| TCGA-GV-A3QI | TCGA-BLCA | 61.162943 | 50.3472852 |
| TCGA-GV-A40E | TCGA-BLCA | 102.217657 | 77.3111055 |
| TCGA-GV-A40G | TCGA-BLCA | 132.250654 | 52.2061254 |
| TCGA-GV-A6ZA | TCGA-BLCA | 132.670032 | 39.7712311 |
| TCGA-H2-A26U | TCGA-THCA | 40.9088106 | 12.7613681 |
| TCGA-H2-A2K9 | TCGA-THCA | 69.4101384 | 18.7214048 |
| TCGA-H2-A3RH | TCGA-THCA | 77.8786941 | 17.8509413 |
| TCGA-H2-A3RI | TCGA-THCA | 57.6684684 | 11.6222275 |
| TCGA-H2-A421 | TCGA-THCA | 45.4450966 | 11.836139 |
| TCGA-H2-A422 | TCGA-THCA | 38.6307809 | 13.6959968 |
| TCGA-H4-A2HO | TCGA-BLCA | 37.1155878 | 13.0943431 |

| | | | |
|---|---|---|---|
| TCGA-H4-A2HQ | TCGA-BLCA | 95.4163792 | 39.2816732 |
| TCGA-H5-A2HR | TCGA-UCEC | 45.4725468 | 5.07059558 |
| TCGA-H6-8124 | TCGA-PAAD | 68.4159071 | 28.874552 |
| TCGA-H6-A45N | TCGA-PAAD | 35.9414382 | 14.2639753 |
| TCGA-H8-A6C1 | TCGA-PAAD | 46.3281698 | 15.4325447 |
| TCGA-H9-7775 | TCGA-PRAD | 43.8983025 | 19.8587574 |
| TCGA-H9-A6BX | TCGA-PRAD | 42.0002558 | 14.7643179 |
| TCGA-H9-A6BY | TCGA-PRAD | 34.3232069 | 13.285045 |
| TCGA-HC-7075 | TCGA-PRAD | 28.7012095 | 17.4177768 |
| TCGA-HC-7077 | TCGA-PRAD | 31.4577303 | 12.3063441 |
| TCGA-HC-7078 | TCGA-PRAD | 18.9443753 | 12.2674414 |
| TCGA-HC-7079 | TCGA-PRAD | 31.0229856 | 9.75839784 |
| TCGA-HC-7080 | TCGA-PRAD | 48.6801786 | 19.1219744 |
| TCGA-HC-7081 | TCGA-PRAD | 28.2627428 | 7.70827921 |
| TCGA-HC-7209 | TCGA-PRAD | 43.0789729 | 16.0368319 |
| TCGA-HC-7210 | TCGA-PRAD | 29.6392028 | 21.1607358 |
| TCGA-HC-7211 | TCGA-PRAD | 34.7027801 | 16.4920938 |
| TCGA-HC-7212 | TCGA-PRAD | 33.115354 | 15.5805484 |
| TCGA-HC-7213 | TCGA-PRAD | 41.3749853 | 28.1753049 |
| TCGA-HC-7230 | TCGA-PRAD | 38.3770105 | 9.02637655 |
| TCGA-HC-7231 | TCGA-PRAD | 39.7515719 | 23.9543524 |
| TCGA-HC-7232 | TCGA-PRAD | 44.3257563 | 23.6607813 |
| TCGA-HC-7233 | TCGA-PRAD | 27.5641803 | 17.4075446 |
| TCGA-HC-7736 | TCGA-PRAD | 36.5446415 | 8.58976546 |
| TCGA-HC-7737 | TCGA-PRAD | 33.3189992 | 13.951069 |
| TCGA-HC-7738 | TCGA-PRAD | 33.0727533 | 16.1233649 |
| TCGA-HC-7740 | TCGA-PRAD | 39.4527436 | 14.9048688 |
| TCGA-HC-7742 | TCGA-PRAD | 33.1890941 | 15.9348177 |
| TCGA-HC-7744 | TCGA-PRAD | 41.0222549 | 17.3516443 |
| TCGA-HC-7745 | TCGA-PRAD | 32.0115003 | 11.3786792 |
| TCGA-HC-7747 | TCGA-PRAD | 30.2960684 | 14.6731475 |

| | | | |
|---|---|---|---|
| TCGA-HC-7748 | TCGA-PRAD | 39.6423881 | 19.774572 |
| TCGA-HC-7749 | TCGA-PRAD | 37.6817951 | 20.0738856 |
| TCGA-HC-7750 | TCGA-PRAD | 26.6511271 | 12.4005971 |
| TCGA-HC-7752 | TCGA-PRAD | 9.3756674 | 4.19567137 |
| TCGA-HC-7817 | TCGA-PRAD | 47.5398581 | 14.1476211 |
| TCGA-HC-7818 | TCGA-PRAD | 32.6229887 | 11.8202525 |
| TCGA-HC-7819 | TCGA-PRAD | 44.8365843 | 19.2259496 |
| TCGA-HC-7820 | TCGA-PRAD | 33.0718014 | 13.9081431 |
| TCGA-HC-7821 | TCGA-PRAD | 43.7382141 | 23.0591034 |
| TCGA-HC-8213 | TCGA-PRAD | 39.0848684 | 13.9179152 |
| TCGA-HC-8216 | TCGA-PRAD | 47.3445711 | 10.1151133 |
| TCGA-HC-8256 | TCGA-PRAD | 33.0901629 | 14.7230304 |
| TCGA-HC-8257 | TCGA-PRAD | 23.3730246 | 11.1803405 |
| TCGA-HC-8258 | TCGA-PRAD | 38.7469115 | 13.2394581 |
| TCGA-HC-8259 | TCGA-PRAD | 43.4560933 | 16.1421876 |
| TCGA-HC-8260 | TCGA-PRAD | 42.7993526 | 16.493608 |
| TCGA-HC-8262 | TCGA-PRAD | 38.5846762 | 11.3512286 |
| TCGA-HC-8264 | TCGA-PRAD | 37.2747407 | 10.6517695 |
| TCGA-HC-8265 | TCGA-PRAD | 18.2061578 | 3.2855155 |
| TCGA-HC-8266 | TCGA-PRAD | 43.6947102 | 16.9574417 |
| TCGA-HC-A48F | TCGA-PRAD | 29.9099624 | 15.473354 |
| TCGA-HC-A4ZV | TCGA-PRAD | 19.1577444 | 15.585798 |
| TCGA-HC-A631 | TCGA-PRAD | 46.2988134 | 12.4307038 |
| TCGA-HC-A632 | TCGA-PRAD | 33.463159 | 5.83265072 |
| TCGA-HC-A6AL | TCGA-PRAD | 46.2121279 | 15.7883919 |
| TCGA-HC-A6AN | TCGA-PRAD | 46.4116535 | 19.3233559 |
| TCGA-HC-A6AO | TCGA-PRAD | 35.9556773 | 21.0732617 |
| TCGA-HC-A6AP | TCGA-PRAD | 42.9901548 | 23.7651091 |
| TCGA-HC-A6AQ | TCGA-PRAD | 38.0593326 | 13.3868953 |
| TCGA-HC-A6AS | TCGA-PRAD | 43.1157746 | 17.1995527 |
| TCGA-HC-A6HX | TCGA-PRAD | 33.4569846 | 14.3125069 |

| | | | |
|---|---|---|---|
| TCGA-HC-A6HY | TCGA-PRAD | 48.4329296 | 14.6899564 |
| TCGA-HC-A76W | TCGA-PRAD | 34.5027086 | 13.7830702 |
| TCGA-HC-A76X | TCGA-PRAD | 40.6397354 | 23.1882439 |
| TCGA-HC-A8CY | TCGA-PRAD | 38.8242666 | 18.8534013 |
| TCGA-HC-A8D0 | TCGA-PRAD | 37.8101364 | 11.6505803 |
| TCGA-HC-A8D1 | TCGA-PRAD | 44.0723386 | 13.6469121 |
| TCGA-HC-A9TE | TCGA-PRAD | 43.5754687 | 24.3331202 |
| TCGA-HC-A9TH | TCGA-PRAD | 51.793505 | 30.6496215 |
| TCGA-HE-7128 | TCGA-KIRP | 20.8658403 | 13.5225253 |
| TCGA-HE-7130 | TCGA-KIRP | 12.7938809 | 27.5344439 |
| TCGA-HE-A5NF | TCGA-KIRP | 2.62698121 | 1.93270582 |
| TCGA-HE-A5NH | TCGA-KIRP | 37.5859223 | 15.6653102 |
| TCGA-HE-A5NI | TCGA-KIRP | 18.3325981 | 8.11533711 |
| TCGA-HE-A5NJ | TCGA-KIRP | 28.2265399 | 12.4076326 |
| TCGA-HF-7131 | TCGA-STAD | 46.6524659 | 27.7405229 |
| TCGA-HF-7132 | TCGA-STAD | 34.642957 | 20.8153145 |
| TCGA-HF-7133 | TCGA-STAD | 67.3468184 | 47.8146579 |
| TCGA-HF-7134 | TCGA-STAD | 35.3706938 | 20.5312306 |
| TCGA-HF-A5NB | TCGA-STAD | 35.0036499 | 27.7670915 |
| TCGA-HI-7168 | TCGA-PRAD | 40.7912167 | 19.4950693 |
| TCGA-HI-7169 | TCGA-PRAD | 19.7901832 | 5.45387632 |
| TCGA-HI-7170 | TCGA-PRAD | 34.3220306 | 10.7341237 |
| TCGA-HI-7171 | TCGA-PRAD | 39.8793761 | 21.8382229 |
| TCGA-HJ-7597 | TCGA-STAD | 66.1042 | 26.7275436 |
| TCGA-HN-A2NL | TCGA-BRCA | 120.892068 | 35.8085918 |
| TCGA-HN-A2OB | TCGA-BRCA | 45.7640975 | 18.7182146 |
| TCGA-HP-A5MZ | TCGA-LIHC | 32.6360677 | 15.6925233 |
| TCGA-HP-A5N0 | TCGA-LIHC | 37.0648559 | 13.5784287 |
| TCGA-HQ-A2OE | TCGA-BLCA | 104.905868 | 48.0410485 |
| TCGA-HQ-A2OF | TCGA-BLCA | 53.6667163 | 33.7241815 |
| TCGA-HQ-A5ND | TCGA-BLCA | 120.554273 | 72.1816025 |

| | | | |
|---|---|---|---|
| TCGA-HQ-A5NE | TCGA-BLCA | 75.2812653 | 28.3351422 |
| TCGA-HR-A2OG | TCGA-SKCM | 33.8871936 | 23.2826213 |
| TCGA-HR-A2OH | TCGA-SKCM | 49.6780009 | 19.3362994 |
| TCGA-HR-A5NC | TCGA-SKCM | 20.2889562 | 27.2427294 |
| TCGA-HT-7467 | TCGA-LGG | 17.8281082 | 10.1192966 |
| TCGA-HT-7468 | TCGA-LGG | 22.4346442 | 21.2660472 |
| TCGA-HT-7469 | TCGA-LGG | 61.7150138 | 22.5483168 |
| TCGA-HT-7470 | TCGA-LGG | 28.2121115 | 8.28482231 |
| TCGA-HT-7471 | TCGA-LGG | 29.2286055 | 34.1619096 |
| TCGA-HT-7472 | TCGA-LGG | 32.3625574 | 10.9495136 |
| TCGA-HT-7473 | TCGA-LGG | 35.495505 | 16.5718233 |
| TCGA-HT-7474 | TCGA-LGG | 25.3668905 | 9.48040004 |
| TCGA-HT-7475 | TCGA-LGG | 35.7331225 | 11.3842202 |
| TCGA-HT-7476 | TCGA-LGG | 22.7976621 | 9.63871247 |
| TCGA-HT-7477 | TCGA-LGG | 51.5020338 | 35.7726595 |
| TCGA-HT-7478 | TCGA-LGG | 31.2395215 | 14.8134038 |
| TCGA-HT-7479 | TCGA-LGG | 28.325311 | 13.9848012 |
| TCGA-HT-7480 | TCGA-LGG | 19.8688211 | 13.0712765 |
| TCGA-HT-7481 | TCGA-LGG | 17.5990241 | 18.0473554 |
| TCGA-HT-7482 | TCGA-LGG | 35.4999115 | 14.778876 |
| TCGA-HT-7483 | TCGA-LGG | 34.8826704 | 11.7530369 |
| TCGA-HT-7485 | TCGA-LGG | 33.9303532 | 12.5839056 |
| TCGA-HT-7601 | TCGA-LGG | 31.6619761 | 10.9656648 |
| TCGA-HT-7602 | TCGA-LGG | 22.6886326 | 12.3319028 |
| TCGA-HT-7603 | TCGA-LGG | 41.1248359 | 12.782185 |
| TCGA-HT-7604 | TCGA-LGG | 34.8735788 | 40.3009914 |
| TCGA-HT-7605 | TCGA-LGG | 21.4204196 | 13.3157192 |
| TCGA-HT-7606 | TCGA-LGG | 56.5592596 | 18.8433281 |
| TCGA-HT-7607 | TCGA-LGG | 21.8111236 | 9.1962665 |
| TCGA-HT-7608 | TCGA-LGG | 14.7148723 | 13.8612202 |
| TCGA-HT-7609 | TCGA-LGG | 26.8536399 | 11.0018091 |

| | | | |
|---|---|---|---|
| TCGA-HT-7610 | TCGA-LGG | 26.4970875 | 8.84121305 |
| TCGA-HT-7611 | TCGA-LGG | 38.9949216 | 14.2884902 |
| TCGA-HT-7616 | TCGA-LGG | 27.8879547 | 28.5212732 |
| TCGA-HT-7620 | TCGA-LGG | 20.5237214 | 16.0268423 |
| TCGA-HT-7676 | TCGA-LGG | 29.4598189 | 11.2352993 |
| TCGA-HT-7677 | TCGA-LGG | 19.9301762 | 22.0328516 |
| TCGA-HT-7680 | TCGA-LGG | 35.8911236 | 12.5096052 |
| TCGA-HT-7681 | TCGA-LGG | 28.1089898 | 15.5040602 |
| TCGA-HT-7684 | TCGA-LGG | 34.6798806 | 12.7224365 |
| TCGA-HT-7686 | TCGA-LGG | 43.8045675 | 11.0399258 |
| TCGA-HT-7687 | TCGA-LGG | 22.5320416 | 20.1420148 |
| TCGA-HT-7688 | TCGA-LGG | 37.1853669 | 14.5484335 |
| TCGA-HT-7689 | TCGA-LGG | 39.3325506 | 15.1619038 |
| TCGA-HT-7690 | TCGA-LGG | 44.6145864 | 19.875233 |
| TCGA-HT-7691 | TCGA-LGG | 40.8400849 | 16.0475923 |
| TCGA-HT-7692 | TCGA-LGG | 20.0616829 | 18.9244846 |
| TCGA-HT-7693 | TCGA-LGG | 63.647439 | 20.624294 |
| TCGA-HT-7694 | TCGA-LGG | 28.2804711 | 10.3294395 |
| TCGA-HT-7695 | TCGA-LGG | 20.1051268 | 13.4896319 |
| TCGA-HT-7854 | TCGA-LGG | 33.5243512 | 11.4175757 |
| TCGA-HT-7855 | TCGA-LGG | 30.0461671 | 9.95748576 |
| TCGA-HT-7856 | TCGA-LGG | 19.8793125 | 9.67259902 |
| TCGA-HT-7857 | TCGA-LGG | 34.7737752 | 14.0240733 |
| TCGA-HT-7858 | TCGA-LGG | 33.163321 | 15.1066669 |
| TCGA-HT-7860 | TCGA-LGG | 31.6128778 | 20.7039665 |
| TCGA-HT-7873 | TCGA-LGG | 33.7119483 | 18.7632362 |
| TCGA-HT-7874 | TCGA-LGG | 22.1951218 | 12.9344556 |
| TCGA-HT-7875 | TCGA-LGG | 18.1442277 | 16.0827524 |
| TCGA-HT-7877 | TCGA-LGG | 18.5111089 | 16.1889163 |
| TCGA-HT-7879 | TCGA-LGG | 30.0119097 | 13.105883 |
| TCGA-HT-7880 | TCGA-LGG | 21.8638085 | 7.96510404 |

| | | | |
|---|---|---|---|
| TCGA-HT-7881 | TCGA-LGG | 26.2025322 | 12.9966861 |
| TCGA-HT-7882 | TCGA-LGG | 51.8754723 | 15.7579983 |
| TCGA-HT-7884 | TCGA-LGG | 27.836668 | 14.6170745 |
| TCGA-HT-7902 | TCGA-LGG | 28.4133082 | 13.1432844 |
| TCGA-HT-8010 | TCGA-LGG | 16.2909825 | 10.0118773 |
| TCGA-HT-8011 | TCGA-LGG | 70.9640692 | 32.2525877 |
| TCGA-HT-8012 | TCGA-LGG | 19.4761948 | 12.0051702 |
| TCGA-HT-8013 | TCGA-LGG | 39.2427116 | 15.3678791 |
| TCGA-HT-8015 | TCGA-LGG | 21.8453768 | 9.56707498 |
| TCGA-HT-8018 | TCGA-LGG | 36.7741091 | 9.79493528 |
| TCGA-HT-8019 | TCGA-LGG | 22.1486969 | 6.32150376 |
| TCGA-HT-8104 | TCGA-LGG | 54.2378467 | 23.0549791 |
| TCGA-HT-8105 | TCGA-LGG | 23.9168492 | 18.3463852 |
| TCGA-HT-8106 | TCGA-LGG | 34.7092512 | 14.9212571 |
| TCGA-HT-8107 | TCGA-LGG | 20.2210477 | 6.13254946 |
| TCGA-HT-8108 | TCGA-LGG | 27.5746637 | 16.4340605 |
| TCGA-HT-8109 | TCGA-LGG | 17.4418952 | 12.3507747 |
| TCGA-HT-8110 | TCGA-LGG | 33.2080443 | 15.8455266 |
| TCGA-HT-8111 | TCGA-LGG | 33.1857636 | 11.98299 |
| TCGA-HT-8113 | TCGA-LGG | 25.2531219 | 9.21014242 |
| TCGA-HT-8114 | TCGA-LGG | 31.3671224 | 10.5744176 |
| TCGA-HT-8558 | TCGA-LGG | 29.7583637 | 9.70543975 |
| TCGA-HT-8563 | TCGA-LGG | 40.7336955 | 16.1751588 |
| TCGA-HT-8564 | TCGA-LGG | 33.8115631 | 15.2269751 |
| TCGA-HT-A4DS | TCGA-LGG | 52.0859353 | 15.6623109 |
| TCGA-HT-A4DV | TCGA-LGG | 23.6131371 | 17.5718367 |
| TCGA-HT-A5R5 | TCGA-LGG | 31.7885208 | 9.65781755 |
| TCGA-HT-A5R7 | TCGA-LGG | 25.9712941 | 10.0261231 |
| TCGA-HT-A5R9 | TCGA-LGG | 15.4922309 | 16.0980074 |
| TCGA-HT-A5RA | TCGA-LGG | 60.5256005 | 25.4420335 |
| TCGA-HT-A5RB | TCGA-LGG | 25.8867166 | 11.9209377 |

| | | | |
|---|---|---|---|
| TCGA-HT-A5RC | TCGA-LGG | 38.8619962 | 14.8880252 |
| TCGA-HT-A614 | TCGA-LGG | 39.9090799 | 31.8694445 |
| TCGA-HT-A615 | TCGA-LGG | 27.4659116 | 22.7128404 |
| TCGA-HT-A616 | TCGA-LGG | 29.7442184 | 9.42099963 |
| TCGA-HT-A617 | TCGA-LGG | 57.6531193 | 15.237286 |
| TCGA-HT-A618 | TCGA-LGG | 35.0007146 | 12.5662869 |
| TCGA-HT-A619 | TCGA-LGG | 21.3200837 | 21.5029424 |
| TCGA-HT-A61A | TCGA-LGG | 23.2091327 | 9.18911898 |
| TCGA-HT-A61B | TCGA-LGG | 63.1552275 | 25.9559048 |
| TCGA-HT-A61C | TCGA-LGG | 43.1972859 | 45.8955811 |
| TCGA-HT-A74H | TCGA-LGG | 41.1315892 | 9.95642448 |
| TCGA-HT-A74J | TCGA-LGG | 29.9759145 | 8.58792551 |
| TCGA-HT-A74K | TCGA-LGG | 25.9680202 | 17.670427 |
| TCGA-HT-A74L | TCGA-LGG | 21.4024021 | 15.1074994 |
| TCGA-HT-A74O | TCGA-LGG | 43.6621112 | 8.31902144 |
| TCGA-HU-8238 | TCGA-STAD | 56.1604875 | 28.5333242 |
| TCGA-HU-8244 | TCGA-STAD | 31.3088813 | 16.7528789 |
| TCGA-HU-8602 | TCGA-STAD | 82.7185346 | 49.2729542 |
| TCGA-HU-8604 | TCGA-STAD | 29.2501623 | 34.8339567 |
| TCGA-HU-8610 | TCGA-STAD | 38.0456804 | 18.5182014 |
| TCGA-HU-A4G2 | TCGA-STAD | 37.6836777 | 64.0565881 |
| TCGA-HU-A4G3 | TCGA-STAD | 22.226478 | 22.3376429 |
| TCGA-HU-A4G8 | TCGA-STAD | 55.2108065 | 25.257268 |
| TCGA-HU-A4G9 | TCGA-STAD | 27.986688 | 16.3390501 |
| TCGA-HU-A4GC | TCGA-STAD | 32.3571985 | 20.5159998 |
| TCGA-HU-A4GD | TCGA-STAD | 45.1555026 | 36.737189 |
| TCGA-HU-A4GF | TCGA-STAD | 95.0925077 | 29.7911997 |
| TCGA-HU-A4GH | TCGA-STAD | 66.0938732 | 23.1036866 |
| TCGA-HU-A4GJ | TCGA-STAD | 33.2143895 | 15.3326333 |
| TCGA-HU-A4GN | TCGA-STAD | 36.2034746 | 14.5341323 |
| TCGA-HU-A4GP | TCGA-STAD | 30.0811763 | 14.3865893 |

| | | | |
|---|---|---|---|
| TCGA-HU-A4GC | TCGA-STAD | 33.7653211 | 40.4310764 |
| TCGA-HU-A4GT | TCGA-STAD | 79.6137562 | 57.5066147 |
| TCGA-HU-A4GU | TCGA-STAD | 67.0801254 | 23.1119403 |
| TCGA-HU-A4GX | TCGA-STAD | 57.7191923 | 31.802843 |
| TCGA-HU-A4GY | TCGA-STAD | 31.9247036 | 19.5399709 |
| TCGA-HU-A4H0 | TCGA-STAD | 78.3939209 | 40.3265213 |
| TCGA-HU-A4H2 | TCGA-STAD | 40.0808689 | 21.8145972 |
| TCGA-HU-A4H3 | TCGA-STAD | 68.353451 | 31.6322384 |
| TCGA-HU-A4H4 | TCGA-STAD | 66.6707616 | 26.8408516 |
| TCGA-HU-A4H5 | TCGA-STAD | 57.6369452 | 17.4496734 |
| TCGA-HU-A4H6 | TCGA-STAD | 43.2512168 | 33.6416386 |
| TCGA-HU-A4H8 | TCGA-STAD | 42.6764895 | 15.4534845 |
| TCGA-HU-A4HB | TCGA-STAD | 32.2253149 | 16.6583795 |
| TCGA-HU-A4HD | TCGA-STAD | 42.7485129 | 29.355453 |
| TCGA-HV-A5A3 | TCGA-PAAD | 38.9091043 | 14.3903134 |
| TCGA-HV-A5A4 | TCGA-PAAD | 44.6435252 | 8.26749303 |
| TCGA-HV-A5A5 | TCGA-PAAD | 32.2978935 | 10.6094562 |
| TCGA-HV-A5A6 | TCGA-PAAD | 49.272158 | 15.406363 |
| TCGA-HV-A7OL | TCGA-PAAD | 59.0728723 | 13.9475844 |
| TCGA-HV-A7OP | TCGA-PAAD | 12.796565 | 9.67025015 |
| TCGA-HV-AA8V | TCGA-PAAD | 42.0598871 | 13.9313419 |
| TCGA-HV-AA8X | TCGA-PAAD | 74.1990849 | 12.8906565 |
| TCGA-HW-7486 | TCGA-LGG | 21.5885993 | 15.3774471 |
| TCGA-HW-7487 | TCGA-LGG | 22.6683318 | 17.4782569 |
| TCGA-HW-7489 | TCGA-LGG | 29.9225329 | 11.9375707 |
| TCGA-HW-7490 | TCGA-LGG | 27.3354559 | 13.2421929 |
| TCGA-HW-7491 | TCGA-LGG | 18.1006321 | 19.5933009 |
| TCGA-HW-7493 | TCGA-LGG | 33.1033635 | 14.7323227 |
| TCGA-HW-7495 | TCGA-LGG | 20.1378857 | 13.6620069 |
| TCGA-HW-8319 | TCGA-LGG | 35.5050209 | 10.8659544 |
| TCGA-HW-8320 | TCGA-LGG | 34.5900768 | 8.84714202 |

| | | | |
|---|---|---|---|
| TCGA-HW-8321 | TCGA-LGG | 26.5756575 | 9.95234815 |
| TCGA-HW-8322 | TCGA-LGG | 20.1093707 | 18.623926 |
| TCGA-HW-A5KJ | TCGA-LGG | 21.6358314 | 14.5682109 |
| TCGA-HW-A5KK | TCGA-LGG | 38.17539 | 15.4219677 |
| TCGA-HW-A5KL | TCGA-LGG | 19.4306393 | 8.00569319 |
| TCGA-HW-A5KM | TCGA-LGG | 39.1215333 | 24.5411503 |
| TCGA-HZ-7289 | TCGA-PAAD | 41.9016796 | 12.5962365 |
| TCGA-HZ-7918 | TCGA-PAAD | 53.0048182 | 22.6342389 |
| TCGA-HZ-7919 | TCGA-PAAD | 73.3299315 | 17.0493748 |
| TCGA-HZ-7920 | TCGA-PAAD | 60.8286933 | 19.8897813 |
| TCGA-HZ-7922 | TCGA-PAAD | 55.125144 | 22.9447248 |
| TCGA-HZ-7923 | TCGA-PAAD | 34.6559873 | 10.4888758 |
| TCGA-HZ-7924 | TCGA-PAAD | 50.16807 | 11.6598788 |
| TCGA-HZ-7925 | TCGA-PAAD | 66.7097449 | 15.4539532 |
| TCGA-HZ-7926 | TCGA-PAAD | 48.4317765 | 10.7488253 |
| TCGA-HZ-8001 | TCGA-PAAD | 45.8717116 | 22.8738726 |
| TCGA-HZ-8002 | TCGA-PAAD | 60.9139145 | 22.8910436 |
| TCGA-HZ-8003 | TCGA-PAAD | 21.5595929 | 6.05348393 |
| TCGA-HZ-8005 | TCGA-PAAD | 56.0291914 | 24.7690277 |
| TCGA-HZ-8315 | TCGA-PAAD | 50.3160725 | 13.5135934 |
| TCGA-HZ-8317 | TCGA-PAAD | 30.3043366 | 13.9493321 |
| TCGA-HZ-8519 | TCGA-PAAD | 39.2845588 | 15.2811457 |
| TCGA-HZ-8636 | TCGA-PAAD | 63.7363597 | 20.9237557 |
| TCGA-HZ-8637 | TCGA-PAAD | 61.1483507 | 25.9303148 |
| TCGA-HZ-8638 | TCGA-PAAD | 46.7260195 | 18.213993 |
| TCGA-HZ-A49G | TCGA-PAAD | 36.1416857 | 10.5531758 |
| TCGA-HZ-A49H | TCGA-PAAD | 30.0841455 | 10.1947738 |
| TCGA-HZ-A49I | TCGA-PAAD | 39.0915171 | 9.12424839 |
| TCGA-HZ-A4BH | TCGA-PAAD | 49.2784333 | 10.2002065 |
| TCGA-HZ-A4BK | TCGA-PAAD | 57.0264115 | 13.414039 |
| TCGA-HZ-A77O | TCGA-PAAD | 33.5248279 | 19.4691755 |

| | | | |
|---|---|---|---|
| TCGA-HZ-A77P | TCGA-PAAD | 52.0149396 | 17.3877745 |
| TCGA-HZ-A77Q | TCGA-PAAD | 52.8277119 | 18.9552749 |
| TCGA-HZ-A8P0 | TCGA-PAAD | 45.656782 | 21.6279025 |
| TCGA-HZ-A8P1 | TCGA-PAAD | 38.4209101 | 27.4270314 |
| TCGA-HZ-A9TJ | TCGA-PAAD | 57.2484861 | 22.5280333 |
| TCGA-IA-A40Y | TCGA-KIRP | 37.566278 | 48.8570533 |
| TCGA-IA-A83S | TCGA-KIRP | 28.405905 | 22.8302033 |
| TCGA-IA-A83T | TCGA-KIRP | 42.2966003 | 11.5517201 |
| TCGA-IA-A83V | TCGA-KIRP | 36.8479434 | 12.3984564 |
| TCGA-IA-A83W | TCGA-KIRP | 43.7718203 | 24.8537485 |
| TCGA-IB-7644 | TCGA-PAAD | 67.6117709 | 14.4186494 |
| TCGA-IB-7645 | TCGA-PAAD | 56.0621584 | 16.072149 |
| TCGA-IB-7646 | TCGA-PAAD | 58.8998588 | 46.0143353 |
| TCGA-IB-7649 | TCGA-PAAD | 23.5632904 | 8.81968868 |
| TCGA-IB-7651 | TCGA-PAAD | 37.8139778 | 14.1882276 |
| TCGA-IB-7652 | TCGA-PAAD | 45.9175162 | 18.0857267 |
| TCGA-IB-7654 | TCGA-PAAD | 38.7228772 | 16.6467089 |
| TCGA-IB-7885 | TCGA-PAAD | 54.1742865 | 19.383213 |
| TCGA-IB-7886 | TCGA-PAAD | 49.5929121 | 19.5166212 |
| TCGA-IB-7887 | TCGA-PAAD | 79.9573807 | 15.718439 |
| TCGA-IB-7888 | TCGA-PAAD | 48.1903514 | 20.4422177 |
| TCGA-IB-7889 | TCGA-PAAD | 43.7756116 | 20.7779766 |
| TCGA-IB-7890 | TCGA-PAAD | 40.9283051 | 20.5368153 |
| TCGA-IB-7891 | TCGA-PAAD | 36.4753805 | 14.964858 |
| TCGA-IB-7893 | TCGA-PAAD | 57.0371759 | 20.1232971 |
| TCGA-IB-7897 | TCGA-PAAD | 50.6337762 | 17.7518387 |
| TCGA-IB-8126 | TCGA-PAAD | 20.799128 | 5.54204871 |
| TCGA-IB-8127 | TCGA-PAAD | 57.2042063 | 20.2038663 |
| TCGA-IB-A5SO | TCGA-PAAD | 43.9480072 | 11.5586055 |
| TCGA-IB-A5SP | TCGA-PAAD | 47.019757 | 10.4241322 |
| TCGA-IB-A5SQ | TCGA-PAAD | 53.6820188 | 23.8028963 |

| | | | |
|---|---|---|---|
| TCGA-IB-A5SS | TCGA-PAAD | 42.8468277 | 27.4026534 |
| TCGA-IB-A5ST | TCGA-PAAD | 48.458976 | 15.7812464 |
| TCGA-IB-A6UF | TCGA-PAAD | 81.341091 | 15.3902413 |
| TCGA-IB-A6UG | TCGA-PAAD | 26.7514595 | 14.1813238 |
| TCGA-IB-A7LX | TCGA-PAAD | 35.1216592 | 28.779729 |
| TCGA-IB-A7M4 | TCGA-PAAD | 56.9475634 | 25.2940843 |
| TCGA-IB-AAUM | TCGA-PAAD | 23.1724374 | 7.1412496 |
| TCGA-IB-AAUN | TCGA-PAAD | 39.4335934 | 24.8833009 |
| TCGA-IB-AAUO | TCGA-PAAD | 75.5939746 | 23.642362 |
| TCGA-IB-AAUP | TCGA-PAAD | 55.5656209 | 15.567553 |
| TCGA-IB-AAUQ | TCGA-PAAD | 44.3674279 | 13.7287631 |
| TCGA-IB-AAUR | TCGA-PAAD | 47.8268148 | 13.9583172 |
| TCGA-IB-AAUS | TCGA-PAAD | 51.7271857 | 19.8046355 |
| TCGA-IB-AAUT | TCGA-PAAD | 37.5482581 | 10.748353 |
| TCGA-IB-AAUU | TCGA-PAAD | 69.0184371 | 14.8239672 |
| TCGA-IB-AAUV | TCGA-PAAD | 50.2574687 | 16.8084063 |
| TCGA-IB-AAUW | TCGA-PAAD | 51.0389731 | 13.3673443 |
| TCGA-IC-A6RE | TCGA-ESCA | 34.2547726 | 20.4106665 |
| TCGA-IC-A6RF | TCGA-ESCA | 37.8615541 | 13.941766 |
| TCGA-IG-A3I8 | TCGA-ESCA | 45.6453519 | 24.7100999 |
| TCGA-IG-A3QL | TCGA-ESCA | 45.330661 | 76.8049393 |
| TCGA-IG-A3YA | TCGA-ESCA | 59.657089 | 28.0850593 |
| TCGA-IG-A3YB | TCGA-ESCA | 24.2085389 | 17.3010015 |
| TCGA-IG-A4P3 | TCGA-ESCA | 27.8600319 | 20.70201 |
| TCGA-IG-A4QS | TCGA-ESCA | 30.2336928 | 25.4739466 |
| TCGA-IG-A50L | TCGA-ESCA | 40.2588476 | 35.5710368 |
| TCGA-IG-A51D | TCGA-ESCA | 81.8501149 | 38.3650665 |
| TCGA-IG-A5B8 | TCGA-ESCA | 40.6953039 | 31.9340895 |
| TCGA-IG-A5S3 | TCGA-ESCA | 42.9137172 | 74.4161839 |
| TCGA-IG-A625 | TCGA-ESCA | 34.2404055 | 17.0986816 |
| TCGA-IG-A6QS | TCGA-ESCA | 100.191239 | 26.8394783 |

| | | | |
|---|---|---|---|
| TCGA-IG-A7DP | TCGA-ESCA | 25.6622925 | 11.0708519 |
| TCGA-IG-A8O2 | TCGA-ESCA | 77.4855109 | 20.9851063 |
| TCGA-IG-A97H | TCGA-ESCA | 78.5253992 | 69.1693314 |
| TCGA-IG-A97I | TCGA-ESCA | 44.3105871 | 19.2660379 |
| TCGA-IH-A3EA | TCGA-SKCM | 46.1220494 | 18.9347497 |
| TCGA-IK-7675 | TCGA-LGG | 44.915186 | 16.1681719 |
| TCGA-IK-8125 | TCGA-LGG | 16.1212029 | 12.3494797 |
| TCGA-IM-A3EB | TCGA-THCA | 41.0676377 | 17.3596079 |
| TCGA-IM-A3ED | TCGA-THCA | 49.3443718 | 13.075033 |
| TCGA-IM-A3U2 | TCGA-THCA | 46.4138257 | 14.6546748 |
| TCGA-IM-A3U3 | TCGA-THCA | 47.8528227 | 10.5031608 |
| TCGA-IM-A41Y | TCGA-THCA | 37.1021165 | 10.7529263 |
| TCGA-IM-A41Z | TCGA-THCA | 49.9092795 | 14.2182344 |
| TCGA-IM-A420 | TCGA-THCA | 46.1352801 | 11.8605288 |
| TCGA-IM-A4EB | TCGA-THCA | 41.791378 | 15.1942199 |
| TCGA-IN-7806 | TCGA-STAD | 22.3909192 | 12.2799225 |
| TCGA-IN-7808 | TCGA-STAD | 50.3229562 | 43.920882 |
| TCGA-IN-8462 | TCGA-STAD | 39.8086467 | 22.7309944 |
| TCGA-IN-8663 | TCGA-STAD | 60.0335399 | 22.2543944 |
| TCGA-IN-A6RI | TCGA-STAD | 77.2768571 | 18.5900695 |
| TCGA-IN-A6RJ | TCGA-STAD | 20.1398395 | 10.1594911 |
| TCGA-IN-A6RL | TCGA-STAD | 50.5248294 | 18.4976317 |
| TCGA-IN-A6RN | TCGA-STAD | 28.7982686 | 4.02457701 |
| TCGA-IN-A6RO | TCGA-STAD | 56.0043591 | 24.5684798 |
| TCGA-IN-A6RR | TCGA-STAD | 55.0649962 | 21.2122511 |
| TCGA-IN-A6RS | TCGA-STAD | 44.8323466 | 20.192123 |
| TCGA-IN-A7NR | TCGA-STAD | 24.8156455 | 19.6497714 |
| TCGA-IN-A7NT | TCGA-STAD | 57.3273097 | 46.1439297 |
| TCGA-IN-A7NU | TCGA-STAD | 46.319619 | 16.6531111 |
| TCGA-IN-AB1V | TCGA-STAD | 20.5836246 | 16.9447264 |
| TCGA-IN-AB1X | TCGA-STAD | 17.1608814 | 9.5139186 |

| | | | |
|---|---|---|---|
| TCGA-IP-7968 | TCGA-STAD | 58.4838036 | 13.4295126 |
| TCGA-IZ-8196 | TCGA-KIRP | 35.8795429 | 10.5453101 |
| TCGA-IZ-A6M8 | TCGA-KIRP | 12.5567866 | 5.66354019 |
| TCGA-IZ-A6M9 | TCGA-KIRP | 34.404061 | 13.5853008 |
| TCGA-J1-A4AH | TCGA-LUSC | 46.6828301 | 11.7262942 |
| TCGA-J2-8192 | TCGA-LUAD | 54.6784044 | 22.2306942 |
| TCGA-J2-8194 | TCGA-LUAD | 42.3033961 | 27.5347938 |
| TCGA-J2-A4AD | TCGA-LUAD | 60.6591818 | 41.0651858 |
| TCGA-J2-A4AE | TCGA-LUAD | 46.0620811 | 21.3328495 |
| TCGA-J2-A4AG | TCGA-LUAD | 52.5779521 | 30.8793065 |
| TCGA-J4-8198 | TCGA-PRAD | 32.9602265 | 10.6341879 |
| TCGA-J4-8200 | TCGA-PRAD | 36.5962301 | 13.6613294 |
| TCGA-J4-A67K | TCGA-PRAD | 31.5799873 | 12.9612796 |
| TCGA-J4-A67L | TCGA-PRAD | 48.3468284 | 28.9618483 |
| TCGA-J4-A67M | TCGA-PRAD | 35.2665293 | 13.302461 |
| TCGA-J4-A67N | TCGA-PRAD | 40.9161081 | 11.4129432 |
| TCGA-J4-A67O | TCGA-PRAD | 26.1139364 | 13.4571623 |
| TCGA-J4-A67Q | TCGA-PRAD | 32.0023205 | 12.2116821 |
| TCGA-J4-A67R | TCGA-PRAD | 39.1765056 | 17.1251265 |
| TCGA-J4-A67S | TCGA-PRAD | 39.7908481 | 20.3130055 |
| TCGA-J4-A67T | TCGA-PRAD | 32.8987235 | 14.3752451 |
| TCGA-J4-A6G1 | TCGA-PRAD | 45.6418293 | 11.8053072 |
| TCGA-J4-A6G3 | TCGA-PRAD | 49.4643077 | 21.0377906 |
| TCGA-J4-A6M7 | TCGA-PRAD | 41.0861174 | 15.1540082 |
| TCGA-J4-A83I | TCGA-PRAD | 50.6403393 | 22.9132614 |
| TCGA-J4-A83J | TCGA-PRAD | 46.2775397 | 13.5233888 |
| TCGA-J4-A83K | TCGA-PRAD | 43.5771415 | 14.3396236 |
| TCGA-J4-A83L | TCGA-PRAD | 40.450253 | 13.6751101 |
| TCGA-J4-A83M | TCGA-PRAD | 32.8216157 | 12.6604915 |
| TCGA-J4-A83N | TCGA-PRAD | 49.2664689 | 16.9154326 |
| TCGA-J4-AATV | TCGA-PRAD | 43.4991061 | 13.779222 |

| | | | |
|---|---|---|---|
| TCGA-J4-AATZ | TCGA-PRAD | 41.8497637 | 19.1347704 |
| TCGA-J7-6720 | TCGA-KIRP | 45.5326815 | 14.9462737 |
| TCGA-J7-8537 | TCGA-KIRP | 67.735723 | 14.7494074 |
| TCGA-J7-A8I2 | TCGA-KIRP | 13.151914 | 2.03795695 |
| TCGA-J8-A3NZ | TCGA-THCA | 65.7481876 | 20.7267432 |
| TCGA-J8-A3O0 | TCGA-THCA | 62.3765914 | 18.7800392 |
| TCGA-J8-A3O1 | TCGA-THCA | 50.7351029 | 18.1069422 |
| TCGA-J8-A3O2 | TCGA-THCA | 72.1659691 | 16.0364019 |
| TCGA-J8-A3YD | TCGA-THCA | 59.1268009 | 15.7400001 |
| TCGA-J8-A3YE | TCGA-THCA | 60.049768 | 13.0536612 |
| TCGA-J8-A3YF | TCGA-THCA | 49.3189778 | 14.5109092 |
| TCGA-J8-A3YG | TCGA-THCA | 44.7327527 | 14.5225536 |
| TCGA-J8-A3YH | TCGA-THCA | 68.8798138 | 14.7130479 |
| TCGA-J8-A42S | TCGA-THCA | 75.554439 | 13.6357864 |
| TCGA-J8-A4HW | TCGA-THCA | 45.4265689 | 17.8862334 |
| TCGA-J8-A4HY | TCGA-THCA | 50.3416138 | 12.0178537 |
| TCGA-J9-A52C | TCGA-PRAD | 34.0977094 | 16.0161715 |
| TCGA-J9-A52D | TCGA-PRAD | 28.4980219 | 15.9538808 |
| TCGA-J9-A52E | TCGA-PRAD | 39.1758056 | 18.0166735 |
| TCGA-J9-A8CK | TCGA-PRAD | 40.8978395 | 17.4723572 |
| TCGA-J9-A8CL | TCGA-PRAD | 33.7910484 | 15.4027396 |
| TCGA-J9-A8CM | TCGA-PRAD | 35.6322164 | 16.1286663 |
| TCGA-J9-A8CN | TCGA-PRAD | 29.2588935 | 10.2274983 |
| TCGA-J9-A8CP | TCGA-PRAD | 48.8385708 | 10.0192082 |
| TCGA-JL-A3YW | TCGA-BRCA | 57.8146345 | 18.5986628 |
| TCGA-JL-A3YX | TCGA-BRCA | 49.6095656 | 32.7062141 |
| TCGA-JU-AAVI | TCGA-UCEC | 70.3086287 | 35.3589026 |
| TCGA-JY-A6FA | TCGA-ESCA | 43.1608114 | 19.5899822 |
| TCGA-JY-A6FB | TCGA-ESCA | 29.3540713 | 23.3734026 |
| TCGA-JY-A6FD | TCGA-ESCA | 40.9934325 | 27.438278 |
| TCGA-JY-A6FE | TCGA-ESCA | 49.4985987 | 37.9257247 |

| | | | |
|---|---|---|---|
| TCGA-JY-A6FG | TCGA-ESCA | 19.7249042 | 13.1822733 |
| TCGA-JY-A6FH | TCGA-ESCA | 68.9840424 | 19.0306426 |
| TCGA-JY-A939 | TCGA-ESCA | 15.1847866 | 10.982073 |
| TCGA-JY-A93C | TCGA-ESCA | 24.8882253 | 15.9594113 |
| TCGA-JY-A93D | TCGA-ESCA | 27.4575178 | 13.670804 |
| TCGA-JY-A93E | TCGA-ESCA | 42.7516504 | 15.018027 |
| TCGA-JY-A93F | TCGA-ESCA | 40.1281305 | 25.5828652 |
| TCGA-K4-A3WS | TCGA-BLCA | 48.6486632 | 26.798252 |
| TCGA-K4-A3WU | TCGA-BLCA | 51.0392317 | 26.1009874 |
| TCGA-K4-A3WV | TCGA-BLCA | 49.4390313 | 9.82897957 |
| TCGA-K4-A4AB | TCGA-BLCA | 57.9225223 | 15.6746858 |
| TCGA-K4-A4AC | TCGA-BLCA | 126.743942 | 39.834222 |
| TCGA-K4-A5RH | TCGA-BLCA | 85.4855147 | 38.0950292 |
| TCGA-K4-A5RI | TCGA-BLCA | 54.5202834 | 11.8522534 |
| TCGA-K4-A5RJ | TCGA-BLCA | 77.5088573 | 35.3919115 |
| TCGA-K4-A6FZ | TCGA-BLCA | 67.9326601 | 13.7916157 |
| TCGA-K4-A6MB | TCGA-BLCA | 97.9779495 | 25.4975228 |
| TCGA-K4-A83P | TCGA-BLCA | 47.6883617 | 23.3601149 |
| TCGA-K4-AAQO | TCGA-BLCA | 59.2835366 | 22.9741781 |
| TCGA-K6-A3WQ | TCGA-UCEC | 61.8535411 | 23.7548626 |
| TCGA-K7-A5RF | TCGA-LIHC | 27.5895499 | 11.7952343 |
| TCGA-K7-A5RG | TCGA-LIHC | 31.0299481 | 18.07338 |
| TCGA-K7-A6G5 | TCGA-LIHC | 28.2607472 | 11.8635562 |
| TCGA-K7-AAU7 | TCGA-LIHC | 46.5172769 | 11.2825502 |
| TCGA-KB-A6F7 | TCGA-STAD | 71.9316188 | 31.1682665 |
| TCGA-KB-A93G | TCGA-STAD | 24.4474456 | 18.6003879 |
| TCGA-KB-A93H | TCGA-STAD | 49.5756173 | 42.192652 |
| TCGA-KB-A93J | TCGA-STAD | 36.8457945 | 18.2321968 |
| TCGA-KC-A4BL | TCGA-PRAD | 42.512801 | 16.5036724 |
| TCGA-KC-A4BN | TCGA-PRAD | 37.342363 | 15.132938 |
| TCGA-KC-A4BR | TCGA-PRAD | 31.1724399 | 18.1157423 |

| | | | |
|---|---|---|---|
| TCGA-KC-A4BV | TCGA-PRAD | 33.2377733 | 15.2505756 |
| TCGA-KC-A7F3 | TCGA-PRAD | 44.8759682 | 18.944436 |
| TCGA-KC-A7F5 | TCGA-PRAD | 42.7623772 | 33.6935401 |
| TCGA-KC-A7F6 | TCGA-PRAD | 52.170267 | 21.8027561 |
| TCGA-KC-A7FA | TCGA-PRAD | 52.9544198 | 14.3388108 |
| TCGA-KC-A7FD | TCGA-PRAD | 55.6880201 | 31.1219901 |
| TCGA-KC-A7FE | TCGA-PRAD | 41.8424494 | 15.0724882 |
| TCGA-KH-A6WC | TCGA-ESCA | 14.7866502 | 9.46089633 |
| TCGA-KJ-A3U4 | TCGA-UCEC | 90.1002319 | 26.3480914 |
| TCGA-KK-A59V | TCGA-PRAD | 75.8625704 | 28.1621573 |
| TCGA-KK-A59X | TCGA-PRAD | 57.4145992 | 31.7715856 |
| TCGA-KK-A59Y | TCGA-PRAD | 49.8628909 | 35.7430233 |
| TCGA-KK-A59Z | TCGA-PRAD | 19.6505906 | 11.7770391 |
| TCGA-KK-A5A1 | TCGA-PRAD | 34.9198895 | 20.5412839 |
| TCGA-KK-A6DY | TCGA-PRAD | 45.8192979 | 15.9912542 |
| TCGA-KK-A6E0 | TCGA-PRAD | 26.829424 | 27.24111 |
| TCGA-KK-A6E1 | TCGA-PRAD | 37.8931492 | 17.9722634 |
| TCGA-KK-A6E2 | TCGA-PRAD | 43.0372571 | 14.4040209 |
| TCGA-KK-A6E3 | TCGA-PRAD | 41.4603499 | 15.3098678 |
| TCGA-KK-A6E4 | TCGA-PRAD | 35.686715 | 22.7714553 |
| TCGA-KK-A6E5 | TCGA-PRAD | 32.2432631 | 9.52088576 |
| TCGA-KK-A6E6 | TCGA-PRAD | 47.3117202 | 25.3863822 |
| TCGA-KK-A6E7 | TCGA-PRAD | 50.258767 | 20.0442378 |
| TCGA-KK-A6E8 | TCGA-PRAD | 32.2678967 | 25.9432774 |
| TCGA-KK-A7AP | TCGA-PRAD | 46.3305223 | 21.1682042 |
| TCGA-KK-A7AQ | TCGA-PRAD | 30.9853097 | 15.9281191 |
| TCGA-KK-A7AU | TCGA-PRAD | 54.0239895 | 16.822255 |
| TCGA-KK-A7AV | TCGA-PRAD | 46.6411643 | 13.8408962 |
| TCGA-KK-A7AW | TCGA-PRAD | 20.8837946 | 13.1840382 |
| TCGA-KK-A7AY | TCGA-PRAD | 44.6023418 | 13.9208691 |
| TCGA-KK-A7AZ | TCGA-PRAD | 44.0161741 | 10.4121431 |

| | | | |
|---|---|---|---|
| TCGA-KK-A7B0 | TCGA-PRAD | 32.5713745 | 12.5249112 |
| TCGA-KK-A7B1 | TCGA-PRAD | 52.9773916 | 16.6093735 |
| TCGA-KK-A7B2 | TCGA-PRAD | 49.5578956 | 18.4952478 |
| TCGA-KK-A7B3 | TCGA-PRAD | 43.7659925 | 23.2142307 |
| TCGA-KK-A7B4 | TCGA-PRAD | 56.7834997 | 24.5867278 |
| TCGA-KK-A8I4 | TCGA-PRAD | 76.7259485 | 17.1337534 |
| TCGA-KK-A8I5 | TCGA-PRAD | 34.3114473 | 16.8549557 |
| TCGA-KK-A8I6 | TCGA-PRAD | 53.0426653 | 12.115721 |
| TCGA-KK-A8I7 | TCGA-PRAD | 42.7199023 | 13.4073349 |
| TCGA-KK-A8I8 | TCGA-PRAD | 64.7877447 | 11.3337577 |
| TCGA-KK-A8I9 | TCGA-PRAD | 57.4537449 | 18.8482485 |
| TCGA-KK-A8IA | TCGA-PRAD | 52.9486428 | 26.133956 |
| TCGA-KK-A8IB | TCGA-PRAD | 48.9627575 | 14.7888004 |
| TCGA-KK-A8IC | TCGA-PRAD | 62.6348769 | 12.2302002 |
| TCGA-KK-A8ID | TCGA-PRAD | 40.8024826 | 26.1760649 |
| TCGA-KK-A8IF | TCGA-PRAD | 48.1910884 | 23.1337742 |
| TCGA-KK-A8IG | TCGA-PRAD | 35.0306722 | 11.048249 |
| TCGA-KK-A8IH | TCGA-PRAD | 50.0255951 | 20.3662598 |
| TCGA-KK-A8II | TCGA-PRAD | 46.4510288 | 11.1748473 |
| TCGA-KK-A8IJ | TCGA-PRAD | 49.8193645 | 20.596283 |
| TCGA-KK-A8IK | TCGA-PRAD | 48.5259232 | 29.6149661 |
| TCGA-KK-A8IL | TCGA-PRAD | 51.1626546 | 15.6073472 |
| TCGA-KK-A8IM | TCGA-PRAD | 48.7179713 | 21.0716598 |
| TCGA-KP-A3VZ | TCGA-UCEC | 127.728287 | 35.3204899 |
| TCGA-KP-A3W0 | TCGA-UCEC | 69.8566256 | 2.23374209 |
| TCGA-KP-A3W1 | TCGA-UCEC | 53.9385646 | 10.6761109 |
| TCGA-KP-A3W3 | TCGA-UCEC | 163.37412 | 35.0844414 |
| TCGA-KP-A3W4 | TCGA-UCEC | 89.1421439 | 18.8728756 |
| TCGA-KQ-A41N | TCGA-BLCA | 53.3426272 | 22.6598417 |
| TCGA-KQ-A41O | TCGA-BLCA | 75.8902324 | 19.5158537 |
| TCGA-KQ-A41P | TCGA-BLCA | 42.5132668 | 24.9022806 |

| | | | |
|---|---|---|---|
| TCGA-KQ-A41Q | TCGA-BLCA | 104.149129 | 234.694169 |
| TCGA-KQ-A41R | TCGA-BLCA | 63.0954546 | 13.8863711 |
| TCGA-KQ-A41S | TCGA-BLCA | 55.9203694 | 17.5494534 |
| TCGA-KR-A7K0 | TCGA-LIHC | 49.8053442 | 26.0407068 |
| TCGA-KR-A7K2 | TCGA-LIHC | 35.1135724 | 28.8157059 |
| TCGA-KR-A7K7 | TCGA-LIHC | 45.3231415 | 26.3836757 |
| TCGA-KR-A7K8 | TCGA-LIHC | 34.819222 | 19.8962655 |
| TCGA-KS-A41F | TCGA-THCA | 24.8533363 | 8.87613264 |
| TCGA-KS-A41I | TCGA-THCA | 29.0114737 | 5.67977333 |
| TCGA-KS-A41J | TCGA-THCA | 66.7503824 | 13.149448 |
| TCGA-KS-A41L | TCGA-THCA | 76.629048 | 16.2538621 |
| TCGA-KS-A4I1 | TCGA-THCA | 71.8885646 | 17.5581395 |
| TCGA-KS-A4I3 | TCGA-THCA | 49.0048063 | 13.1489017 |
| TCGA-KS-A4I5 | TCGA-THCA | 70.0194086 | 22.857329 |
| TCGA-KS-A4I7 | TCGA-THCA | 65.9024614 | 17.363574 |
| TCGA-KS-A4I9 | TCGA-THCA | 44.3574155 | 13.9900648 |
| TCGA-KS-A4IB | TCGA-THCA | 42.8289763 | 11.6623536 |
| TCGA-KS-A4IC | TCGA-THCA | 55.6199987 | 16.2163305 |
| TCGA-KS-A4ID | TCGA-THCA | 35.1067657 | 10.8157021 |
| TCGA-KT-A74X | TCGA-LGG | 24.1685876 | 26.2379859 |
| TCGA-KT-A7W1 | TCGA-LGG | 59.9391433 | 43.7642377 |
| TCGA-KV-A6GD | TCGA-KIRP | 35.2370281 | 6.47695868 |
| TCGA-KV-A6GE | TCGA-KIRP | 33.0609119 | 15.969589 |
| TCGA-KV-A74V | TCGA-KIRP | 39.8246602 | 10.0677475 |
| TCGA-L1-A7W4 | TCGA-PAAD | 93.7773019 | 41.2975864 |
| TCGA-L3-A4E7 | TCGA-LUSC | 45.109263 | 26.4965663 |
| TCGA-L3-A524 | TCGA-LUSC | 50.5854127 | 23.4286154 |
| TCGA-L4-A4E5 | TCGA-LUAD | 66.0210972 | 25.398965 |
| TCGA-L4-A4E6 | TCGA-LUAD | 31.3906815 | 12.9844409 |
| TCGA-L5-A43C | TCGA-ESCA | 28.5622123 | 14.9320022 |
| TCGA-L5-A43E | TCGA-ESCA | 42.1056471 | 12.3293837 |

| | | | |
|---|---|---|---|
| TCGA-L5-A43J | TCGA-ESCA | 35.7808522 | 17.3884879 |
| TCGA-L5-A4OE | TCGA-ESCA | 32.2731412 | 17.7584599 |
| TCGA-L5-A4OG | TCGA-ESCA | 23.7536978 | 15.9360692 |
| TCGA-L5-A4OH | TCGA-ESCA | 42.3097053 | 21.4872226 |
| TCGA-L5-A4OI | TCGA-ESCA | 27.8237274 | 35.5305228 |
| TCGA-L5-A4OJ | TCGA-ESCA | 40.9102025 | 23.0011702 |
| TCGA-L5-A4OM | TCGA-ESCA | 46.7261739 | 28.8408194 |
| TCGA-L5-A4OO | TCGA-ESCA | 25.7479334 | 14.1030093 |
| TCGA-L5-A4OP | TCGA-ESCA | 37.4043937 | 9.9699112 |
| TCGA-L5-A4OS | TCGA-ESCA | 45.4949214 | 27.0057875 |
| TCGA-L5-A4OT | TCGA-ESCA | 23.3340932 | 20.4999601 |
| TCGA-L5-A4OU | TCGA-ESCA | 42.2315964 | 29.9447559 |
| TCGA-L5-A4OW | TCGA-ESCA | 22.1448501 | 25.8059914 |
| TCGA-L5-A4OX | TCGA-ESCA | 45.1616423 | 32.3167684 |
| TCGA-L5-A88S | TCGA-ESCA | 34.8847294 | 27.2266626 |
| TCGA-L5-A88T | TCGA-ESCA | 16.6365397 | 11.371647 |
| TCGA-L5-A88V | TCGA-ESCA | 28.2010456 | 23.5349662 |
| TCGA-L5-A88W | TCGA-ESCA | 33.7238991 | 15.9392226 |
| TCGA-L5-A88Y | TCGA-ESCA | 14.0098837 | 15.9131549 |
| TCGA-L5-A88Z | TCGA-ESCA | 27.5788184 | 27.4484688 |
| TCGA-L5-A891 | TCGA-ESCA | 66.4297614 | 12.7360667 |
| TCGA-L5-A893 | TCGA-ESCA | 48.414426 | 15.2917723 |
| TCGA-L5-A8NE | TCGA-ESCA | 42.4533211 | 20.4832311 |
| TCGA-L5-A8NF | TCGA-ESCA | 44.0597395 | 50.2757873 |
| TCGA-L5-A8NG | TCGA-ESCA | 25.6345256 | 27.5283708 |
| TCGA-L5-A8NH | TCGA-ESCA | 33.8996996 | 22.3974757 |
| TCGA-L5-A8NI | TCGA-ESCA | 57.7153021 | 49.9022877 |
| TCGA-L5-A8NJ | TCGA-ESCA | 41.8630308 | 37.9384808 |
| TCGA-L5-A8NK | TCGA-ESCA | 32.8002929 | 17.8909086 |
| TCGA-L5-A8NL | TCGA-ESCA | 61.1372019 | 47.4286019 |
| TCGA-L5-A8NM | TCGA-ESCA | 37.1742269 | 28.7963939 |

| | | | |
|---|---|---|---|
| TCGA-L5-A8NN | TCGA-ESCA | 58.7392778 | 39.6815923 |
| TCGA-L5-A8NQ | TCGA-ESCA | 35.454796 | 18.4549182 |
| TCGA-L5-A8NR | TCGA-ESCA | 38.9068743 | 19.254343 |
| TCGA-L5-A8NS | TCGA-ESCA | 42.9173308 | 20.7060961 |
| TCGA-L5-A8NT | TCGA-ESCA | 30.3079601 | 30.6636513 |
| TCGA-L5-A8NU | TCGA-ESCA | 17.9575171 | 10.3246092 |
| TCGA-L5-A8NV | TCGA-ESCA | 39.0643708 | 16.4193881 |
| TCGA-L5-A8NW | TCGA-ESCA | 24.7587025 | 20.7115495 |
| TCGA-L6-A4EP | TCGA-THCA | 69.2946192 | 27.0089547 |
| TCGA-L6-A4EQ | TCGA-THCA | 62.8559473 | 10.4184758 |
| TCGA-L6-A4ET | TCGA-THCA | 63.2194576 | 13.2019467 |
| TCGA-L6-A4EU | TCGA-THCA | 40.9762409 | 18.0936885 |
| TCGA-L7-A56G | TCGA-ESCA | 33.274146 | 44.3588878 |
| TCGA-L7-A6VZ | TCGA-ESCA | 42.6999329 | 33.9136933 |
| TCGA-L9-A443 | TCGA-LUAD | 51.2366734 | 26.6056942 |
| TCGA-L9-A444 | TCGA-LUAD | 31.9442498 | 11.0568929 |
| TCGA-L9-A50W | TCGA-LUAD | 67.8757829 | 18.0822323 |
| TCGA-L9-A5IP | TCGA-LUAD | 129.966411 | 38.281773 |
| TCGA-L9-A743 | TCGA-LUAD | 40.6953498 | 16.5456816 |
| TCGA-L9-A7SV | TCGA-LUAD | 61.7014099 | 8.69540112 |
| TCGA-L9-A8F4 | TCGA-LUAD | 48.1160417 | 16.760977 |
| TCGA-LA-A446 | TCGA-LUSC | 17.7244058 | 21.229286 |
| TCGA-LA-A7SW | TCGA-LUSC | 117.2647 | 89.8263035 |
| TCGA-LB-A7SX | TCGA-PAAD | 78.9585622 | 17.3931042 |
| TCGA-LB-A8F3 | TCGA-PAAD | 37.5429394 | 36.6510755 |
| TCGA-LB-A9Q5 | TCGA-PAAD | 22.524005 | 9.46222362 |
| TCGA-LC-A66R | TCGA-BLCA | 61.0128357 | 34.5129578 |
| TCGA-LD-A66U | TCGA-BRCA | 54.034192 | 34.5664167 |
| TCGA-LD-A74U | TCGA-BRCA | 59.7163728 | 33.4636345 |
| TCGA-LD-A7W6 | TCGA-BRCA | 67.9644967 | 23.9063905 |
| TCGA-LD-A9QF | TCGA-BRCA | 52.5104544 | 15.3621677 |

| | | | |
|---|---|---|---|
| TCGA-LG-A6GG | TCGA-LIHC | 28.4391337 | 17.0517372 |
| TCGA-LG-A9QC | TCGA-LIHC | 37.0310346 | 32.2610402 |
| TCGA-LG-A9QD | TCGA-LIHC | 32.6155277 | 15.8959614 |
| TCGA-LH-A9QB | TCGA-SKCM | 39.0368299 | 15.4416282 |
| TCGA-LL-A440 | TCGA-BRCA | 40.8367232 | 27.030295 |
| TCGA-LL-A441 | TCGA-BRCA | 110.895171 | 39.0615413 |
| TCGA-LL-A442 | TCGA-BRCA | 39.0470562 | 13.7837531 |
| TCGA-LL-A50Y | TCGA-BRCA | 59.544262 | 22.8822386 |
| TCGA-LL-A5YL | TCGA-BRCA | 35.0272579 | 62.1134793 |
| TCGA-LL-A5YM | TCGA-BRCA | 24.6901709 | 15.2215976 |
| TCGA-LL-A5YN | TCGA-BRCA | 56.2200424 | 29.1752385 |
| TCGA-LL-A5YO | TCGA-BRCA | 62.1296866 | 20.2556464 |
| TCGA-LL-A5YP | TCGA-BRCA | 65.6155059 | 31.7505744 |
| TCGA-LL-A6FP | TCGA-BRCA | 46.1250754 | 69.2886728 |
| TCGA-LL-A6FQ | TCGA-BRCA | 42.6624464 | 36.824137 |
| TCGA-LL-A6FR | TCGA-BRCA | 86.9805269 | 27.5833817 |
| TCGA-LL-A73Y | TCGA-BRCA | 66.4364844 | 20.1804329 |
| TCGA-LL-A73Z | TCGA-BRCA | 67.7508321 | 32.6823119 |
| TCGA-LL-A740 | TCGA-BRCA | 57.7735049 | 33.7334432 |
| TCGA-LL-A7SZ | TCGA-BRCA | 28.8744286 | 22.79059 |
| TCGA-LL-A7T0 | TCGA-BRCA | 81.0113662 | 28.3963194 |
| TCGA-LL-A8F5 | TCGA-BRCA | 132.303128 | 24.0246277 |
| TCGA-LL-A9Q3 | TCGA-BRCA | 36.586156 | 45.4417465 |
| TCGA-LN-A49M | TCGA-ESCA | 50.3724299 | 16.0264305 |
| TCGA-LN-A49O | TCGA-ESCA | 53.588542 | 49.7861899 |
| TCGA-LN-A49P | TCGA-ESCA | 54.7418492 | 25.2916582 |
| TCGA-LN-A49S | TCGA-ESCA | 56.564544 | 41.052579 |
| TCGA-LN-A49U | TCGA-ESCA | 49.8120348 | 32.90408 |
| TCGA-LN-A49W | TCGA-ESCA | 28.8135144 | 19.8148845 |
| TCGA-LN-A49X | TCGA-ESCA | 48.9075198 | 17.1175534 |
| TCGA-LN-A49Y | TCGA-ESCA | 63.69385 | 34.4339711 |

| | | | |
|---|---|---|---|
| TCGA-LN-A4A1 | TCGA-ESCA | 36.1236245 | 28.3443164 |
| TCGA-LN-A4A3 | TCGA-ESCA | 34.5521286 | 33.9921199 |
| TCGA-LN-A4A4 | TCGA-ESCA | 29.8003079 | 34.3569058 |
| TCGA-LN-A4A5 | TCGA-ESCA | 34.0079802 | 13.2219861 |
| TCGA-LN-A4A8 | TCGA-ESCA | 62.1199826 | 48.5819693 |
| TCGA-LN-A4MQ | TCGA-ESCA | 36.6412386 | 16.9318087 |
| TCGA-LN-A5U5 | TCGA-ESCA | 47.5823521 | 47.5204586 |
| TCGA-LN-A5U6 | TCGA-ESCA | 41.4866485 | 36.4411084 |
| TCGA-LN-A5U7 | TCGA-ESCA | 25.0426593 | 34.8243619 |
| TCGA-LN-A7HV | TCGA-ESCA | 47.5514079 | 55.5746771 |
| TCGA-LN-A7HW | TCGA-ESCA | 30.7516555 | 22.7208346 |
| TCGA-LN-A7HX | TCGA-ESCA | 52.8772426 | 29.6114692 |
| TCGA-LN-A7HY | TCGA-ESCA | 57.4867271 | 25.2159229 |
| TCGA-LN-A7HZ | TCGA-ESCA | 46.0926063 | 28.4149818 |
| TCGA-LN-A8I0 | TCGA-ESCA | 49.1443878 | 27.6189328 |
| TCGA-LN-A8I1 | TCGA-ESCA | 45.9915525 | 30.1655187 |
| TCGA-LN-A9FO | TCGA-ESCA | 86.9041382 | 34.4026756 |
| TCGA-LN-A9FP | TCGA-ESCA | 24.1761049 | 12.8659283 |
| TCGA-LN-A9FR | TCGA-ESCA | 36.3046761 | 24.7319637 |
| TCGA-LQ-A4E4 | TCGA-BRCA | 72.029343 | 36.79852 |
| TCGA-LT-A5Z6 | TCGA-BLCA | 60.9953228 | 23.7049392 |
| TCGA-LT-A8JT | TCGA-BLCA | 61.7088085 | 18.1221622 |
| TCGA-M7-A71Y | TCGA-PRAD | 52.2117155 | 12.2818111 |
| TCGA-M7-A71Z | TCGA-PRAD | 43.4950484 | 11.6774577 |
| TCGA-M7-A720 | TCGA-PRAD | 40.5032052 | 14.4983788 |
| TCGA-M7-A721 | TCGA-PRAD | 36.7083156 | 20.9722592 |
| TCGA-M7-A722 | TCGA-PRAD | 20.7194946 | 9.07669345 |
| TCGA-M7-A723 | TCGA-PRAD | 28.4946616 | 16.3793232 |
| TCGA-M7-A724 | TCGA-PRAD | 42.5685342 | 19.7424883 |
| TCGA-M7-A725 | TCGA-PRAD | 33.6860584 | 17.8532545 |
| TCGA-M8-A5N4 | TCGA-PAAD | 41.0423802 | 13.7577187 |

| | | | |
|---|---|---|---|
| TCGA-M9-A5M8 | TCGA-ESCA | 39.4611317 | 12.8651734 |
| TCGA-MF-A522 | TCGA-LUSC | 54.8511649 | 80.7105298 |
| TCGA-MG-AAM( | TCGA-PRAD | 55.5824029 | 20.7332172 |
| TCGA-MH-A55W | TCGA-KIRP | 25.1739707 | 12.6282457 |
| TCGA-MH-A55Z | TCGA-KIRP | 24.5180984 | 7.9083495 |
| TCGA-MH-A560 | TCGA-KIRP | 21.7943536 | 12.0625765 |
| TCGA-MH-A561 | TCGA-KIRP | 31.6548794 | 6.62194113 |
| TCGA-MH-A562 | TCGA-KIRP | 29.8197072 | 6.07282819 |
| TCGA-MH-A854 | TCGA-KIRP | 41.5606018 | 10.4290296 |
| TCGA-MH-A855 | TCGA-KIRP | 33.8208067 | 11.1355562 |
| TCGA-MH-A856 | TCGA-KIRP | 30.4522863 | 2.04217468 |
| TCGA-MH-A857 | TCGA-KIRP | 21.7666368 | 7.58127788 |
| TCGA-MI-A75C | TCGA-LIHC | 44.0253974 | 32.6284412 |
| TCGA-MI-A75E | TCGA-LIHC | 40.6728755 | 16.9906532 |
| TCGA-MI-A75G | TCGA-LIHC | 44.7889514 | 16.288294 |
| TCGA-MI-A75H | TCGA-LIHC | 61.0185557 | 24.4658245 |
| TCGA-MI-A75I | TCGA-LIHC | 47.8268567 | 4.60231311 |
| TCGA-MK-A4N6 | TCGA-THCA | 42.3097189 | 11.9560635 |
| TCGA-MK-A4N7 | TCGA-THCA | 66.7021929 | 17.9043586 |
| TCGA-MK-A4N9 | TCGA-THCA | 67.1305511 | 17.9401641 |
| TCGA-MK-A84Z | TCGA-THCA | 59.8258726 | 18.5382682 |
| TCGA-MM-A563 | TCGA-KIRC | 54.0552158 | 20.0726218 |
| TCGA-MM-A564 | TCGA-KIRC | 49.1263515 | 12.6515406 |
| TCGA-MM-A84U | TCGA-KIRC | 35.4095846 | 11.7501084 |
| TCGA-MN-A4N1 | TCGA-LUAD | 56.0851726 | 45.5427749 |
| TCGA-MN-A4N4 | TCGA-LUAD | 49.5669062 | 29.9081887 |
| TCGA-MN-A4N5 | TCGA-LUAD | 53.8875271 | 13.601008 |
| TCGA-MP-A4SV | TCGA-LUAD | 57.2283605 | 30.0586048 |
| TCGA-MP-A4SW | TCGA-LUAD | 45.3298933 | 21.5535112 |
| TCGA-MP-A4SY | TCGA-LUAD | 41.2468159 | 21.1070754 |
| TCGA-MP-A4T4 | TCGA-LUAD | 37.2625666 | 13.0834214 |

| | | | |
|---|---|---|---|
| TCGA-MP-A4T6 | TCGA-LUAD | 66.7808569 | 31.9573988 |
| TCGA-MP-A4T7 | TCGA-LUAD | 27.6049955 | 36.5367805 |
| TCGA-MP-A4T8 | TCGA-LUAD | 45.8401165 | 24.1608747 |
| TCGA-MP-A4T9 | TCGA-LUAD | 44.002642 | 18.8196174 |
| TCGA-MP-A4TA | TCGA-LUAD | 50.6973189 | 26.5512033 |
| TCGA-MP-A4TC | TCGA-LUAD | 70.6385949 | 19.033004 |
| TCGA-MP-A4TD | TCGA-LUAD | 22.7026291 | 17.7676463 |
| TCGA-MP-A4TE | TCGA-LUAD | 41.5342952 | 37.1812412 |
| TCGA-MP-A4TF | TCGA-LUAD | 46.8790609 | 31.6227642 |
| TCGA-MP-A4TH | TCGA-LUAD | 23.3568242 | 9.90215189 |
| TCGA-MP-A4TI | TCGA-LUAD | 71.2658916 | 28.8914609 |
| TCGA-MP-A4TJ | TCGA-LUAD | 37.2263304 | 17.032955 |
| TCGA-MP-A4TK | TCGA-LUAD | 28.5210977 | 14.2178063 |
| TCGA-MP-A5C7 | TCGA-LUAD | 52.4165864 | 22.0498287 |
| TCGA-MR-A520 | TCGA-LIHC | 19.9806526 | 7.14818179 |
| TCGA-MR-A8JO | TCGA-LIHC | 27.3689711 | 14.7308502 |
| TCGA-MS-A51U | TCGA-BRCA | 45.1659716 | 14.3497847 |
| TCGA-MV-A51V | TCGA-BLCA | 100.556846 | 16.7379878 |
| TCGA-MW-A4E( | TCGA-KIRC | 64.0049093 | 14.922867 |
| TCGA-MX-A5UG | TCGA-STAD | 18.9927974 | 11.61253 |
| TCGA-MX-A5UJ | TCGA-STAD | 35.0691713 | 26.6319469 |
| TCGA-MX-A663 | TCGA-STAD | 25.3655098 | 35.6226484 |
| TCGA-MX-A666 | TCGA-STAD | 51.9315998 | 19.6040196 |
| TCGA-N5-A4R8 | TCGA-UCS | 123.93779 | 92.1611179 |
| TCGA-N5-A4RA | TCGA-UCS | 76.0754893 | 31.488572 |
| TCGA-N5-A4RD | TCGA-UCS | 81.4803403 | 28.0599122 |
| TCGA-N5-A4RF | TCGA-UCS | 70.2756666 | 20.2447242 |
| TCGA-N5-A4RJ | TCGA-UCS | 80.683575 | 24.6140374 |
| TCGA-N5-A4RM | TCGA-UCS | 176.431999 | 34.7755249 |
| TCGA-N5-A4RN | TCGA-UCS | 52.5808449 | 32.8292938 |
| TCGA-N5-A4RO | TCGA-UCS | 57.1698085 | 53.411531 |

| | | | |
|---|---|---|---|
| TCGA-N5-A4RS | TCGA-UCS | 178.392494 | 9.18754342 |
| TCGA-N5-A4RT | TCGA-UCS | 89.8137326 | 55.5387748 |
| TCGA-N5-A4RU | TCGA-UCS | 103.717145 | 11.8732213 |
| TCGA-N5-A4RV | TCGA-UCS | 90.9169031 | 31.9896454 |
| TCGA-N5-A59E | TCGA-UCS | 155.241957 | 117.921048 |
| TCGA-N5-A59F | TCGA-UCS | 106.704815 | 35.8606971 |
| TCGA-N6-A4V9 | TCGA-UCS | 78.0356858 | 10.4723652 |
| TCGA-N6-A4VC | TCGA-UCS | 89.6014072 | 36.5229389 |
| TCGA-N6-A4VD | TCGA-UCS | 67.5481087 | 6.69537201 |
| TCGA-N6-A4VE | TCGA-UCS | 83.8315045 | 45.4615984 |
| TCGA-N6-A4VF | TCGA-UCS | 79.062575 | 41.9231183 |
| TCGA-N6-A4VG | TCGA-UCS | 145.986754 | 10.63157 |
| TCGA-N7-A4Y0 | TCGA-UCS | 63.8455761 | 53.2308815 |
| TCGA-N7-A4Y5 | TCGA-UCS | 72.57903 | 20.7203698 |
| TCGA-N7-A4Y8 | TCGA-UCS | 120.049443 | 25.1996084 |
| TCGA-N7-A59B | TCGA-UCS | 74.4547452 | 20.0996614 |
| TCGA-N8-A4PI | TCGA-UCS | 62.9644529 | 14.0115404 |
| TCGA-N8-A4PL | TCGA-UCS | 104.266641 | 46.3990793 |
| TCGA-N8-A4PM | TCGA-UCS | 72.1605576 | 29.5117232 |
| TCGA-N8-A4PN | TCGA-UCS | 34.6360618 | 3.47468744 |
| TCGA-N8-A4PO | TCGA-UCS | 66.1263435 | 47.9875216 |
| TCGA-N8-A4PP | TCGA-UCS | 72.6095724 | 26.1856887 |
| TCGA-N8-A4PQ | TCGA-UCS | 33.2406349 | 33.8535392 |
| TCGA-N8-A56S | TCGA-UCS | 99.8508375 | 31.8128011 |
| TCGA-N9-A4PZ | TCGA-UCS | 53.7282424 | 15.792139 |
| TCGA-N9-A4Q1 | TCGA-UCS | 85.3625542 | 34.5607644 |
| TCGA-N9-A4Q3 | TCGA-UCS | 115.695753 | 37.0047686 |
| TCGA-N9-A4Q4 | TCGA-UCS | 107.148372 | 46.4288414 |
| TCGA-N9-A4Q7 | TCGA-UCS | 84.0891412 | 34.0416278 |
| TCGA-NA-A4QV | TCGA-UCS | 82.5006177 | 30.9617663 |
| TCGA-NA-A4QW | TCGA-UCS | 26.2628141 | 20.2908079 |

| | | | |
|---|---|---|---|
| TCGA-NA-A4QX | TCGA-UCS | 90.2302223 | 35.9118951 |
| TCGA-NA-A4QY | TCGA-UCS | 99.2818931 | 16.875212 |
| TCGA-NA-A4R0 | TCGA-UCS | 83.8566368 | 29.8703002 |
| TCGA-NA-A4R1 | TCGA-UCS | 114.426439 | 35.8041714 |
| TCGA-NA-A5I1 | TCGA-UCS | 46.5723569 | 18.6338921 |
| TCGA-NC-A5HD | TCGA-LUSC | 103.402421 | 52.400775 |
| TCGA-NC-A5HE | TCGA-LUSC | 52.4498236 | 41.5974342 |
| TCGA-NC-A5HF | TCGA-LUSC | 24.491588 | 5.20817513 |
| TCGA-NC-A5HG | TCGA-LUSC | 58.9200222 | 10.7653015 |
| TCGA-NC-A5HH | TCGA-LUSC | 98.8384826 | 40.0805733 |
| TCGA-NC-A5HI | TCGA-LUSC | 66.6607465 | 14.5715125 |
| TCGA-NC-A5HJ | TCGA-LUSC | 161.152552 | 36.7952066 |
| TCGA-NC-A5HK | TCGA-LUSC | 106.615287 | 60.6693203 |
| TCGA-NC-A5HL | TCGA-LUSC | 58.2366969 | 24.7843127 |
| TCGA-NC-A5HM | TCGA-LUSC | 101.250412 | 48.2941302 |
| TCGA-NC-A5HN | TCGA-LUSC | 77.7423765 | 32.5327689 |
| TCGA-NC-A5HO | TCGA-LUSC | 55.124395 | 70.8299693 |
| TCGA-NC-A5HP | TCGA-LUSC | 103.919589 | 6.22246256 |
| TCGA-NC-A5HR | TCGA-LUSC | 71.0925778 | 29.4220619 |
| TCGA-NC-A5HT | TCGA-LUSC | 88.6796693 | 34.3924619 |
| TCGA-ND-A4W6 | TCGA-UCS | 44.2667509 | 18.9729062 |
| TCGA-ND-A4WA | TCGA-UCS | 70.0938048 | 13.1303993 |
| TCGA-ND-A4WC | TCGA-UCS | 47.2958122 | 10.4048966 |
| TCGA-ND-A4WF | TCGA-UCS | 70.8794429 | 54.5729552 |
| TCGA-NF-A4WU | TCGA-UCS | 63.1312575 | 27.6192275 |
| TCGA-NF-A4WX | TCGA-UCS | 63.6607566 | 24.7583028 |
| TCGA-NF-A4X2 | TCGA-UCS | 75.545247 | 42.1185129 |
| TCGA-NF-A5CP | TCGA-UCS | 66.1679926 | 19.1628805 |
| TCGA-NG-A4VU | TCGA-UCS | 40.9592559 | 18.7788107 |
| TCGA-NG-A4VW | TCGA-UCS | 64.1950574 | 32.3034017 |
| TCGA-NH-A50T | TCGA-COAD | 68.1979157 | 34.0357649 |

| | | | |
|---|---|---|---|
| TCGA-NH-A50U | TCGA-COAD | 62.645392 | 24.5036339 |
| TCGA-NH-A50V | TCGA-COAD | 45.9640056 | 20.522907 |
| TCGA-NH-A5IV | TCGA-COAD | 63.5924319 | 27.4353625 |
| TCGA-NH-A6GA | TCGA-COAD | 51.4558197 | 41.8285155 |
| TCGA-NH-A6GB | TCGA-COAD | 67.1780837 | 44.6418722 |
| TCGA-NH-A6GC | TCGA-COAD | 43.9298783 | 18.6139707 |
| TCGA-NH-A8F7 | TCGA-COAD | 79.8810711 | 31.6221368 |
| TCGA-NH-A8F8 | TCGA-COAD | 30.9324346 | 30.4516153 |
| TCGA-NI-A4U2 | TCGA-LIHC | 18.8495764 | 6.70017263 |
| TCGA-NI-A8LF | TCGA-LIHC | 30.7946999 | 12.1442532 |
| TCGA-NJ-A4YF | TCGA-LUAD | 41.245177 | 18.1583072 |
| TCGA-NJ-A4YG | TCGA-LUAD | 38.0500491 | 16.0694219 |
| TCGA-NJ-A4YI | TCGA-LUAD | 63.9144377 | 24.6196717 |
| TCGA-NJ-A4YP | TCGA-LUAD | 43.4646652 | 20.8172845 |
| TCGA-NJ-A4YQ | TCGA-LUAD | 33.832441 | 12.0419184 |
| TCGA-NJ-A55A | TCGA-LUAD | 52.168322 | 17.6057389 |
| TCGA-NJ-A55O | TCGA-LUAD | 33.4858202 | 33.9443276 |
| TCGA-NJ-A55R | TCGA-LUAD | 31.0326561 | 15.6447201 |
| TCGA-NJ-A7XG | TCGA-LUAD | 39.1844161 | 22.7820585 |
| TCGA-NK-A5CR | TCGA-LUSC | 132.539368 | 25.2550835 |
| TCGA-NK-A5CT | TCGA-LUSC | 80.5525639 | 34.3596549 |
| TCGA-NK-A5CX | TCGA-LUSC | 62.3934657 | 20.2354581 |
| TCGA-NK-A7XE | TCGA-LUSC | 57.7718682 | 83.7024587 |
| TCGA-O1-A52J | TCGA-LUAD | 39.1748513 | 14.326567 |
| TCGA-O2-A52N | TCGA-LUSC | 52.6683995 | 21.9837775 |
| TCGA-O2-A52Q | TCGA-LUSC | 140.557028 | 28.8303025 |
| TCGA-O2-A52S | TCGA-LUSC | 47.5052683 | 39.3260191 |
| TCGA-O2-A52V | TCGA-LUSC | 58.9968918 | 26.4101096 |
| TCGA-O2-A52W | TCGA-LUSC | 67.4439691 | 60.9258588 |
| TCGA-O2-A5IB | TCGA-LUSC | 84.630571 | 45.4652337 |
| TCGA-O8-A75V | TCGA-LIHC | 39.1599205 | 13.7451346 |

| | | | |
|---|---|---|---|
| TCGA-O9-A75Z | TCGA-KIRP | 33.273885 | 19.7671821 |
| TCGA-OD-A75X | TCGA-SKCM | 60.0506229 | 40.0339924 |
| TCGA-OE-A75W | TCGA-PAAD | 39.9830075 | 22.6381701 |
| TCGA-OK-A5Q2 | TCGA-BRCA | 47.7011534 | 27.2795326 |
| TCGA-OL-A5D6 | TCGA-BRCA | 33.9079465 | 14.6758745 |
| TCGA-OL-A5D7 | TCGA-BRCA | 109.606574 | 139.21718 |
| TCGA-OL-A5D8 | TCGA-BRCA | 50.7305303 | 30.8393297 |
| TCGA-OL-A5DA | TCGA-BRCA | 57.0488111 | 22.983638 |
| TCGA-OL-A5RU | TCGA-BRCA | 45.0727411 | 11.4141512 |
| TCGA-OL-A5RV | TCGA-BRCA | 52.2209848 | 26.1885639 |
| TCGA-OL-A5RW | TCGA-BRCA | 45.6486725 | 32.5821976 |
| TCGA-OL-A5RX | TCGA-BRCA | 62.5292385 | 16.6706848 |
| TCGA-OL-A5RY | TCGA-BRCA | 56.7550056 | 15.6050748 |
| TCGA-OL-A5RZ | TCGA-BRCA | 40.1295416 | 35.5205808 |
| TCGA-OL-A5S0 | TCGA-BRCA | 152.28659 | 56.9271725 |
| TCGA-OL-A66H | TCGA-BRCA | 59.5048898 | 30.0229418 |
| TCGA-OL-A66I | TCGA-BRCA | 113.062596 | 48.067071 |
| TCGA-OL-A66J | TCGA-BRCA | 61.3315253 | 18.2616933 |
| TCGA-OL-A66K | TCGA-BRCA | 46.0558731 | 24.8708806 |
| TCGA-OL-A66L | TCGA-BRCA | 60.207471 | 25.4897025 |
| TCGA-OL-A66N | TCGA-BRCA | 38.4716787 | 28.9114809 |
| TCGA-OL-A66O | TCGA-BRCA | 50.0421891 | 34.2723982 |
| TCGA-OL-A66P | TCGA-BRCA | 26.4059728 | 11.5536282 |
| TCGA-OL-A6VO | TCGA-BRCA | 120.417817 | 41.58336 |
| TCGA-OL-A6VQ | TCGA-BRCA | 52.6451478 | 21.541348 |
| TCGA-OL-A6VR | TCGA-BRCA | 58.5892994 | 16.2414187 |
| TCGA-OL-A97C | TCGA-BRCA | 51.5046292 | 16.8359041 |
| TCGA-OY-A56Q | TCGA-OV | 56.7699206 | 28.0124764 |
| TCGA-P4-A5E6 | TCGA-KIRP | 52.128902 | 1.186139 |
| TCGA-P4-A5E7 | TCGA-KIRP | 34.7083034 | 19.5631392 |
| TCGA-P4-A5E8 | TCGA-KIRP | 4.12605308 | 29.4127554 |

| | | | |
|---|---|---|---|
| TCGA-P4-A5EA | TCGA-KIRP | 34.9285003 | 14.4391198 |
| TCGA-P4-A5EB | TCGA-KIRP | 11.6285302 | 50.4215991 |
| TCGA-P4-A5ED | TCGA-KIRP | 21.3645044 | 7.305358 |
| TCGA-P4-AAVK | TCGA-KIRP | 38.6089791 | 25.3198929 |
| TCGA-P4-AAVL | TCGA-KIRP | 66.292262 | 24.9381737 |
| TCGA-P4-AAVM | TCGA-KIRP | 33.5640949 | 3.39600385 |
| TCGA-P4-AAVO | TCGA-KIRP | 35.856115 | 13.0991188 |
| TCGA-P5-A5ET | TCGA-LGG | 19.1662096 | 17.3026869 |
| TCGA-P5-A5EU | TCGA-LGG | 57.3027803 | 13.3181124 |
| TCGA-P5-A5EV | TCGA-LGG | 35.2722887 | 13.9389941 |
| TCGA-P5-A5EW | TCGA-LGG | 32.0756119 | 12.2009606 |
| TCGA-P5-A5EX | TCGA-LGG | 29.7799277 | 16.4998757 |
| TCGA-P5-A5EY | TCGA-LGG | 20.5671237 | 8.84443512 |
| TCGA-P5-A5EZ | TCGA-LGG | 36.4914102 | 10.9985912 |
| TCGA-P5-A5F0 | TCGA-LGG | 18.2549561 | 13.1965625 |
| TCGA-P5-A5F1 | TCGA-LGG | 42.7185519 | 14.4805401 |
| TCGA-P5-A5F2 | TCGA-LGG | 41.6781699 | 14.0376714 |
| TCGA-P5-A5F4 | TCGA-LGG | 35.6994433 | 12.9515462 |
| TCGA-P5-A5F6 | TCGA-LGG | 34.2111117 | 8.00697639 |
| TCGA-P5-A72U | TCGA-LGG | 45.0740383 | 12.5447491 |
| TCGA-P5-A72W | TCGA-LGG | 47.9224094 | 15.6554272 |
| TCGA-P5-A72X | TCGA-LGG | 32.4659411 | 6.52660084 |
| TCGA-P5-A72Z | TCGA-LGG | 20.6172374 | 21.1771625 |
| TCGA-P5-A731 | TCGA-LGG | 41.4845093 | 13.4975304 |
| TCGA-P5-A733 | TCGA-LGG | 23.7361432 | 8.62156778 |
| TCGA-P5-A735 | TCGA-LGG | 42.843004 | 11.0083484 |
| TCGA-P5-A736 | TCGA-LGG | 40.8984067 | 8.95043384 |
| TCGA-P5-A737 | TCGA-LGG | 17.9065969 | 17.5534067 |
| TCGA-P5-A77W | TCGA-LGG | 17.9803793 | 24.9525828 |
| TCGA-P5-A77X | TCGA-LGG | 21.4510167 | 24.7377256 |
| TCGA-P5-A780 | TCGA-LGG | 35.7842526 | 9.97089599 |

| | | | |
|---|---|---|---|
| TCGA-P5-A781 | TCGA-LGG | 24.8842876 | 16.719951 |
| TCGA-PD-A5DF | TCGA-LIHC | 81.252268 | 31.7829967 |
| TCGA-PE-A5DC | TCGA-BRCA | 75.273488 | 26.4021636 |
| TCGA-PE-A5DD | TCGA-BRCA | 70.3103786 | 25.8451062 |
| TCGA-PE-A5DE | TCGA-BRCA | 58.5464556 | 28.2424475 |
| TCGA-PG-A5BC | TCGA-UCEC | 34.2983629 | 15.6502104 |
| TCGA-PG-A6IB | TCGA-UCEC | 101.652448 | 43.8204955 |
| TCGA-PG-A7D5 | TCGA-UCEC | 144.429197 | 11.0983843 |
| TCGA-PG-A914 | TCGA-UCEC | 58.758572 | 21.7294355 |
| TCGA-PG-A915 | TCGA-UCEC | 102.518194 | 34.2840013 |
| TCGA-PG-A916 | TCGA-UCEC | 71.3971329 | 20.4264924 |
| TCGA-PG-A917 | TCGA-UCEC | 86.8896093 | 17.9864342 |
| TCGA-PJ-A5Z8 | TCGA-KIRP | 19.8058927 | 26.4777271 |
| TCGA-PJ-A5Z9 | TCGA-KIRP | 37.5967752 | 18.0389978 |
| TCGA-PL-A8LV | TCGA-BRCA | 87.7617549 | 16.6268303 |
| TCGA-PL-A8LX | TCGA-BRCA | 36.7162224 | 37.6786302 |
| TCGA-PL-A8LY | TCGA-BRCA | 53.5589575 | 18.3270174 |
| TCGA-PL-A8LZ | TCGA-BRCA | 137.179665 | 68.5513285 |
| TCGA-PQ-A6FI | TCGA-BLCA | 81.6242375 | 43.0922145 |
| TCGA-PQ-A6FN | TCGA-BLCA | 97.0848879 | 17.8349794 |
| TCGA-PZ-A5RE | TCGA-PAAD | 58.1412742 | 14.3318715 |
| TCGA-Q2-A5QZ | TCGA-KIRP | 21.1101569 | 19.1542557 |
| TCGA-Q3-A5QY | TCGA-PAAD | 47.8497496 | 14.1890445 |
| TCGA-Q3-AA2A | TCGA-PAAD | 61.0941538 | 18.0151017 |
| TCGA-QA-A7B7 | TCGA-LIHC | 68.0287802 | 34.9765587 |
| TCGA-QB-A6FS | TCGA-SKCM | 34.3433319 | 10.8053518 |
| TCGA-QB-AA9O | TCGA-SKCM | 51.7082613 | 31.4387753 |
| TCGA-QD-A8IV | TCGA-THCA | 56.2320291 | 16.620663 |
| TCGA-QF-A5YS | TCGA-UCEC | 86.296539 | 23.9151474 |
| TCGA-QF-A5YT | TCGA-UCEC | 54.4434771 | 81.6188374 |
| TCGA-QG-A5YV | TCGA-COAD | 96.8677491 | 50.1967528 |

| | | | |
|---|---|---|---|
| TCGA-QG-A5YW | TCGA-COAD | 57.9738468 | 17.5909922 |
| TCGA-QG-A5YX | TCGA-COAD | 71.6305833 | 22.27031 |
| TCGA-QG-A5Z1 | TCGA-COAD | 59.1124257 | 20.8438143 |
| TCGA-QG-A5Z2 | TCGA-COAD | 69.9196637 | 24.1998948 |
| TCGA-QH-A65R | TCGA-LGG | 18.0725305 | 13.2123902 |
| TCGA-QH-A65S | TCGA-LGG | 29.7167417 | 10.8372128 |
| TCGA-QH-A65V | TCGA-LGG | 25.2394508 | 18.3830828 |
| TCGA-QH-A65X | TCGA-LGG | 27.369649 | 18.4152415 |
| TCGA-QH-A65Z | TCGA-LGG | 24.6757357 | 21.32758 |
| TCGA-QH-A6CS | TCGA-LGG | 36.2938749 | 13.5490839 |
| TCGA-QH-A6CU | TCGA-LGG | 23.2370455 | 16.3256719 |
| TCGA-QH-A6CV | TCGA-LGG | 66.0320053 | 37.6126812 |
| TCGA-QH-A6CW | TCGA-LGG | 36.0809507 | 6.76987973 |
| TCGA-QH-A6CX | TCGA-LGG | 35.1670411 | 11.7687303 |
| TCGA-QH-A6CY | TCGA-LGG | 27.9810034 | 21.7366796 |
| TCGA-QH-A6CZ | TCGA-LGG | 23.0487036 | 25.02377 |
| TCGA-QH-A6X3 | TCGA-LGG | 42.6932625 | 16.4496977 |
| TCGA-QH-A6X4 | TCGA-LGG | 22.2815494 | 17.9401402 |
| TCGA-QH-A6X5 | TCGA-LGG | 23.3745746 | 19.1572078 |
| TCGA-QH-A6X8 | TCGA-LGG | 27.0195536 | 15.9519241 |
| TCGA-QH-A6X9 | TCGA-LGG | 34.3832546 | 11.1933623 |
| TCGA-QH-A6XA | TCGA-LGG | 38.3778618 | 13.6501395 |
| TCGA-QH-A6XC | TCGA-LGG | 53.6668836 | 18.6491586 |
| TCGA-QH-A86X | TCGA-LGG | 20.4054107 | 17.1119632 |
| TCGA-QH-A870 | TCGA-LGG | 47.4231656 | 14.4887289 |
| TCGA-QL-A97D | TCGA-COAD | 57.228236 | 18.2197922 |
| TCGA-QM-A5NM | TCGA-UCS | 87.2262708 | 31.8166437 |
| TCGA-QN-A5NN | TCGA-UCS | 43.1418146 | 44.5695426 |
| TCGA-QS-A5YQ | TCGA-UCEC | 50.7879564 | 17.4870015 |
| TCGA-QS-A5YR | TCGA-UCEC | 42.5201977 | 8.28293664 |
| TCGA-QS-A744 | TCGA-UCEC | 103.575539 | 30.6266834 |

| | | | |
|---|---|---|---|
| TCGA-QS-A8F1 | TCGA-UCEC | 8.71584569 | 9.53538012 |
| TCGA-QU-A6IL | TCGA-PRAD | 23.2647199 | 13.2061652 |
| TCGA-QU-A6IM | TCGA-PRAD | 20.655179 | 9.77559071 |
| TCGA-QU-A6IN | TCGA-PRAD | 44.7619382 | 16.1046065 |
| TCGA-QU-A6IO | TCGA-PRAD | 22.1912482 | 7.87163614 |
| TCGA-QU-A6IP | TCGA-PRAD | 39.048169 | 16.4199564 |
| TCGA-R3-A69X | TCGA-BLCA | 51.6916326 | 29.1330201 |
| TCGA-R5-A7O7 | TCGA-STAD | 31.5555694 | 22.0102934 |
| TCGA-R5-A7ZE | TCGA-STAD | 33.3136929 | 28.8504705 |
| TCGA-R5-A7ZF | TCGA-STAD | 49.3139135 | 61.4015551 |
| TCGA-R5-A7ZI | TCGA-STAD | 19.331657 | 58.2771923 |
| TCGA-R5-A7ZR | TCGA-STAD | 32.2064985 | 41.5582011 |
| TCGA-R5-A805 | TCGA-STAD | 25.4776389 | 19.2444423 |
| TCGA-R6-A6DN | TCGA-ESCA | 63.6429274 | 59.6706803 |
| TCGA-R6-A6DQ | TCGA-ESCA | 35.8023928 | 38.0557721 |
| TCGA-R6-A6KZ | TCGA-ESCA | 37.542402 | 38.7746951 |
| TCGA-R6-A6XG | TCGA-ESCA | 61.9999377 | 19.7097682 |
| TCGA-R6-A6XQ | TCGA-ESCA | 54.4041135 | 35.8188486 |
| TCGA-R6-A6Y0 | TCGA-ESCA | 27.567877 | 13.2558539 |
| TCGA-R6-A8W8 | TCGA-ESCA | 31.7438866 | 32.909033 |
| TCGA-R6-A8WC | TCGA-ESCA | 51.9766939 | 35.6989839 |
| TCGA-R6-A8WG | TCGA-ESCA | 61.0252048 | 27.7763248 |
| TCGA-R8-A6MK | TCGA-LGG | 26.2264901 | 23.5102952 |
| TCGA-R8-A6ML | TCGA-LGG | 20.1133745 | 21.5003129 |
| TCGA-R8-A6MO | TCGA-LGG | 28.7734972 | 27.2430052 |
| TCGA-R8-A6YH | TCGA-LGG | 38.2097647 | 11.3924562 |
| TCGA-R8-A73M | TCGA-LGG | 27.6858005 | 22.8868203 |
| TCGA-RB-A7B8 | TCGA-PAAD | 57.0741995 | 23.2770954 |
| TCGA-RB-AA9M | TCGA-PAAD | 55.7701478 | 27.8260951 |
| TCGA-RC-A6M3 | TCGA-LIHC | 43.2309797 | 23.7516006 |
| TCGA-RC-A6M4 | TCGA-LIHC | 44.9575655 | 15.4537612 |

| | | | |
|---|---|---|---|
| TCGA-RC-A6M5 | TCGA-LIHC | 28.2160266 | 10.0181059 |
| TCGA-RC-A6M6 | TCGA-LIHC | 58.8318893 | 26.7248944 |
| TCGA-RC-A7S9 | TCGA-LIHC | 51.4674512 | 25.2020468 |
| TCGA-RC-A7SB | TCGA-LIHC | 37.7917654 | 9.82527868 |
| TCGA-RC-A7SF | TCGA-LIHC | 38.0293487 | 12.6859344 |
| TCGA-RC-A7SH | TCGA-LIHC | 23.5564293 | 29.3810567 |
| TCGA-RC-A7SK | TCGA-LIHC | 25.0914553 | 11.1386554 |
| TCGA-RD-A7BS | TCGA-STAD | 24.225211 | 18.1101383 |
| TCGA-RD-A7BT | TCGA-STAD | 50.7854332 | 53.2831031 |
| TCGA-RD-A7BW | TCGA-STAD | 28.3709461 | 25.9359889 |
| TCGA-RD-A7C1 | TCGA-STAD | 48.9893683 | 53.9536003 |
| TCGA-RD-A8MV | TCGA-STAD | 35.9202225 | 23.0972948 |
| TCGA-RD-A8MW | TCGA-STAD | 33.0748508 | 13.6686878 |
| TCGA-RD-A8N0 | TCGA-STAD | 19.601257 | 14.2409367 |
| TCGA-RD-A8N1 | TCGA-STAD | 36.9493548 | 20.1216532 |
| TCGA-RD-A8N2 | TCGA-STAD | 21.3118791 | 13.707811 |
| TCGA-RD-A8N4 | TCGA-STAD | 45.5626564 | 21.5722013 |
| TCGA-RD-A8N5 | TCGA-STAD | 22.0798379 | 34.1163864 |
| TCGA-RD-A8N6 | TCGA-STAD | 42.5247104 | 48.3309884 |
| TCGA-RD-A8N9 | TCGA-STAD | 30.7056732 | 10.3182893 |
| TCGA-RD-A8NB | TCGA-STAD | 44.6330998 | 34.7743298 |
| TCGA-RE-A7BO | TCGA-ESCA | 51.3126233 | 31.7187385 |
| TCGA-RG-A7D4 | TCGA-LIHC | 53.7006886 | 38.3332024 |
| TCGA-RL-AAAS | TCGA-PAAD | 60.1933532 | 18.4716539 |
| TCGA-RP-A690 | TCGA-SKCM | 24.356394 | 7.91377911 |
| TCGA-RP-A693 | TCGA-SKCM | 47.0527955 | 20.4252202 |
| TCGA-RP-A694 | TCGA-SKCM | 67.2738004 | 19.3600168 |
| TCGA-RP-A695 | TCGA-SKCM | 31.0669941 | 9.99210571 |
| TCGA-RP-A6K9 | TCGA-SKCM | 56.1093051 | 23.1028153 |
| TCGA-RU-A8FL | TCGA-COAD | 91.1708163 | 23.3359169 |
| TCGA-RY-A83X | TCGA-LGG | 21.4285377 | 17.5653648 |

| | | | |
|---|---|---|---|
| TCGA-RY-A83Y | TCGA-LGG | 19.9266165 | 16.7829804 |
| TCGA-RY-A83Z | TCGA-LGG | 85.6073916 | 22.6375716 |
| TCGA-RY-A840 | TCGA-LGG | 17.8104228 | 23.7259564 |
| TCGA-RY-A843 | TCGA-LGG | 30.6389701 | 11.3645102 |
| TCGA-RY-A845 | TCGA-LGG | 32.9889052 | 14.8618061 |
| TCGA-RY-A847 | TCGA-LGG | 22.0579772 | 16.5687824 |
| TCGA-S2-AA1A | TCGA-LUAD | 25.2448776 | 22.6358328 |
| TCGA-S3-A6ZF | TCGA-BRCA | 50.0431247 | 73.0813015 |
| TCGA-S3-A6ZG | TCGA-BRCA | 66.7089802 | 42.876517 |
| TCGA-S3-A6ZH | TCGA-BRCA | 85.727595 | 63.743574 |
| TCGA-S3-AA0Z | TCGA-BRCA | 69.4976158 | 50.3164504 |
| TCGA-S3-AA10 | TCGA-BRCA | 78.1321271 | 46.3593913 |
| TCGA-S3-AA11 | TCGA-BRCA | 53.1754055 | 26.540903 |
| TCGA-S3-AA12 | TCGA-BRCA | 48.8013345 | 24.0713075 |
| TCGA-S3-AA14 | TCGA-BRCA | 38.7019782 | 15.7867892 |
| TCGA-S3-AA15 | TCGA-BRCA | 24.7123012 | 17.6800989 |
| TCGA-S3-AA17 | TCGA-BRCA | 36.8958238 | 20.180293 |
| TCGA-S4-A8RM | TCGA-PAAD | 37.0228657 | 16.7298737 |
| TCGA-S4-A8RO | TCGA-PAAD | 47.9292021 | 34.13698 |
| TCGA-S4-A8RP | TCGA-PAAD | 57.8998989 | 18.8505139 |
| TCGA-S5-A6DX | TCGA-BLCA | 40.7024969 | 34.4767644 |
| TCGA-S5-AA26 | TCGA-BLCA | 82.5642417 | 26.1064758 |
| TCGA-S6-A8JW | TCGA-TGCT | 44.8010046 | 82.5307303 |
| TCGA-S6-A8JX | TCGA-TGCT | 14.8536732 | 48.6118286 |
| TCGA-S6-A8JY | TCGA-TGCT | 53.7210316 | 96.4862292 |
| TCGA-S8-A6BV | TCGA-ESCA | 35.7137911 | 9.9447778 |
| TCGA-S8-A6BW | TCGA-ESCA | 36.1284779 | 29.2348024 |
| TCGA-S9-A6TS | TCGA-LGG | 33.9465154 | 12.135383 |
| TCGA-S9-A6TU | TCGA-LGG | 32.3970805 | 5.90851615 |
| TCGA-S9-A6TV | TCGA-LGG | 69.1099598 | 22.5990618 |
| TCGA-S9-A6TW | TCGA-LGG | 18.5032545 | 13.2958982 |

| | | | |
|---|---|---|---|
| TCGA-S9-A6TX | TCGA-LGG | 21.0430923 | 14.1408859 |
| TCGA-S9-A6TY | TCGA-LGG | 19.8312273 | 21.4692804 |
| TCGA-S9-A6TZ | TCGA-LGG | 45.6996724 | 15.0209201 |
| TCGA-S9-A6U0 | TCGA-LGG | 49.7358665 | 16.9772013 |
| TCGA-S9-A6U1 | TCGA-LGG | 40.7196753 | 18.3877588 |
| TCGA-S9-A6U2 | TCGA-LGG | 22.9117773 | 23.7857838 |
| TCGA-S9-A6U5 | TCGA-LGG | 23.8131237 | 21.8856456 |
| TCGA-S9-A6U6 | TCGA-LGG | 40.5785484 | 15.9536714 |
| TCGA-S9-A6U8 | TCGA-LGG | 37.3873741 | 17.9624947 |
| TCGA-S9-A6U9 | TCGA-LGG | 40.8325999 | 9.67809066 |
| TCGA-S9-A6UA | TCGA-LGG | 41.1456635 | 16.0652067 |
| TCGA-S9-A6UB | TCGA-LGG | 23.7103854 | 36.7373054 |
| TCGA-S9-A6WD | TCGA-LGG | 22.164928 | 20.5664426 |
| TCGA-S9-A6WE | TCGA-LGG | 21.5589545 | 20.0846152 |
| TCGA-S9-A6WG | TCGA-LGG | 38.2929574 | 15.6320302 |
| TCGA-S9-A6WH | TCGA-LGG | 25.8937676 | 26.1791319 |
| TCGA-S9-A6WI | TCGA-LGG | 28.5892228 | 13.4312868 |
| TCGA-S9-A6WL | TCGA-LGG | 21.4358496 | 15.5426326 |
| TCGA-S9-A6WM | TCGA-LGG | 79.6430987 | 24.0651051 |
| TCGA-S9-A6WN | TCGA-LGG | 25.2927024 | 14.9170281 |
| TCGA-S9-A6WO | TCGA-LGG | 32.2477758 | 15.8606267 |
| TCGA-S9-A6WP | TCGA-LGG | 25.7689447 | 24.0796554 |
| TCGA-S9-A6WQ | TCGA-LGG | 31.4566037 | 13.7850601 |
| TCGA-S9-A7IQ | TCGA-LGG | 28.7444987 | 13.7172639 |
| TCGA-S9-A7IS | TCGA-LGG | 66.9158883 | 28.0574344 |
| TCGA-S9-A7IX | TCGA-LGG | 42.4232142 | 19.0079387 |
| TCGA-S9-A7IY | TCGA-LGG | 18.3442422 | 19.0027212 |
| TCGA-S9-A7IZ | TCGA-LGG | 31.8280461 | 12.1552679 |
| TCGA-S9-A7J0 | TCGA-LGG | 86.8560275 | 83.7868366 |
| TCGA-S9-A7J1 | TCGA-LGG | 19.714072 | 26.1563499 |
| TCGA-S9-A7J2 | TCGA-LGG | 20.4036008 | 21.5668004 |

| | | | |
|---|---|---|---|
| TCGA-S9-A7J3 | TCGA-LGG | 19.6231531 | 22.0254493 |
| TCGA-S9-A7QW | TCGA-LGG | 37.471513 | 15.1632566 |
| TCGA-S9-A7QX | TCGA-LGG | 28.0050498 | 12.6044506 |
| TCGA-S9-A7QY | TCGA-LGG | 19.9454789 | 16.1291726 |
| TCGA-S9-A7QZ | TCGA-LGG | 20.0682666 | 19.3236567 |
| TCGA-S9-A7R1 | TCGA-LGG | 21.5846858 | 13.782367 |
| TCGA-S9-A7R2 | TCGA-LGG | 47.4400501 | 14.4487985 |
| TCGA-S9-A7R3 | TCGA-LGG | 35.5249824 | 14.3406792 |
| TCGA-S9-A7R4 | TCGA-LGG | 31.5724937 | 10.2125798 |
| TCGA-S9-A7R7 | TCGA-LGG | 38.2762277 | 18.5981686 |
| TCGA-S9-A7R8 | TCGA-LGG | 38.5023937 | 10.7402261 |
| TCGA-S9-A89V | TCGA-LGG | 36.1460395 | 12.5144182 |
| TCGA-S9-A89Z | TCGA-LGG | 57.3656395 | 20.4632626 |
| TCGA-SB-A6J6 | TCGA-TGCT | 50.1057741 | 40.3027128 |
| TCGA-SB-A76C | TCGA-TGCT | 36.2106032 | 44.9153562 |
| TCGA-SJ-A6ZI | TCGA-UCEC | 76.0143178 | 26.8599062 |
| TCGA-SJ-A6ZJ | TCGA-UCEC | 26.8013123 | 5.86892175 |
| TCGA-SL-A6J9 | TCGA-UCEC | 93.7902741 | 25.2112481 |
| TCGA-SL-A6JA | TCGA-UCEC | 48.2005315 | 31.3054145 |
| TCGA-SN-A6IS | TCGA-TGCT | 51.1907885 | 53.3655165 |
| TCGA-SN-A84W | TCGA-TGCT | 48.9600352 | 56.7030594 |
| TCGA-SN-A84X | TCGA-TGCT | 43.8838082 | 58.9707276 |
| TCGA-SN-A84Y | TCGA-TGCT | 161.032608 | 155.638816 |
| TCGA-SO-A8JP | TCGA-TGCT | 26.4957276 | 89.4954343 |
| TCGA-SS-A7HO | TCGA-COAD | 82.8480898 | 26.4332646 |
| TCGA-SU-A7E7 | TCGA-PRAD | 42.8410583 | 15.4098845 |
| TCGA-SW-A7EA | TCGA-STAD | 42.0236779 | 31.4607077 |
| TCGA-SW-A7EB | TCGA-STAD | 33.3193296 | 40.492293 |
| TCGA-SX-A71R | TCGA-KIRP | 30.1161074 | 10.6834686 |
| TCGA-SX-A71S | TCGA-KIRP | 61.0577314 | 4.39042821 |
| TCGA-SX-A71U | TCGA-KIRP | 30.2391509 | 3.38991657 |

| | | | |
|---|---|---|---|
| TCGA-SX-A71V | TCGA-KIRP | 29.0308231 | 7.0868638 |
| TCGA-SX-A71W | TCGA-KIRP | 46.1156909 | 23.3454802 |
| TCGA-SX-A7SL | TCGA-KIRP | 37.1200168 | 13.6278883 |
| TCGA-SX-A7SM | TCGA-KIRP | 52.3408764 | 15.9192938 |
| TCGA-SX-A7SN | TCGA-KIRP | 44.1618735 | 14.71016 |
| TCGA-SX-A7SP | TCGA-KIRP | 30.6428053 | 14.120691 |
| TCGA-SX-A7SQ | TCGA-KIRP | 18.0104664 | 5.37132917 |
| TCGA-SX-A7SR | TCGA-KIRP | 3.25622771 | 6.81795371 |
| TCGA-SX-A7SS | TCGA-KIRP | 20.2491448 | 9.77821718 |
| TCGA-SX-A7SU | TCGA-KIRP | 8.02323325 | 22.0346582 |
| TCGA-SY-A9G0 | TCGA-BLCA | 44.6864153 | 15.1051182 |
| TCGA-SY-A9G5 | TCGA-BLCA | 109.37753 | 26.2325412 |
| TCGA-T1-A6J8 | TCGA-LIHC | 34.7605072 | 8.97087984 |
| TCGA-T7-A92I | TCGA-KIRC | 35.4788077 | 12.7119362 |
| TCGA-T9-A92H | TCGA-COAD | 68.5710328 | 57.2633184 |
| TCGA-TK-A8OK | TCGA-PRAD | 40.0623777 | 12.9929547 |
| TCGA-TM-A7C3 | TCGA-LGG | 73.6814568 | 22.8521932 |
| TCGA-TM-A7C4 | TCGA-LGG | 33.8612226 | 11.8059264 |
| TCGA-TM-A7C5 | TCGA-LGG | 22.5233308 | 18.8592219 |
| TCGA-TM-A7CA | TCGA-LGG | 30.0251448 | 13.1472209 |
| TCGA-TM-A7CF | TCGA-LGG | 32.7140985 | 12.369695 |
| TCGA-TM-A84B | TCGA-LGG | 58.1964443 | 23.4452075 |
| TCGA-TM-A84C | TCGA-LGG | 38.1435949 | 16.794356 |
| TCGA-TM-A84F | TCGA-LGG | 32.8826813 | 14.8474341 |
| TCGA-TM-A84G | TCGA-LGG | 24.0614504 | 25.9143459 |
| TCGA-TM-A84H | TCGA-LGG | 33.3505347 | 15.6803328 |
| TCGA-TM-A84I | TCGA-LGG | 69.0135342 | 27.5922957 |
| TCGA-TM-A84J | TCGA-LGG | 49.0698326 | 24.9196051 |
| TCGA-TM-A84L | TCGA-LGG | 28.9370355 | 11.017391 |
| TCGA-TM-A84M | TCGA-LGG | 18.6965058 | 29.7515868 |
| TCGA-TM-A84O | TCGA-LGG | 23.4433957 | 19.8919343 |

| | | | |
|---|---|---|---|
| TCGA-TM-A84Q | TCGA-LGG | 41.8349735 | 12.6462585 |
| TCGA-TM-A84R | TCGA-LGG | 31.0150614 | 11.882248 |
| TCGA-TM-A84S | TCGA-LGG | 20.585086 | 20.5906713 |
| TCGA-TM-A84T | TCGA-LGG | 29.6029583 | 11.093596 |
| TCGA-TP-A8TT | TCGA-PRAD | 47.9877324 | 10.870259 |
| TCGA-TP-A8TV | TCGA-PRAD | 33.2444736 | 18.0423463 |
| TCGA-TQ-A7RF | TCGA-LGG | 60.4594819 | 13.1229631 |
| TCGA-TQ-A7RG | TCGA-LGG | 23.4114718 | 20.1884858 |
| TCGA-TQ-A7RH | TCGA-LGG | 31.0983127 | 14.9652186 |
| TCGA-TQ-A7RI | TCGA-LGG | 19.518259 | 23.2167267 |
| TCGA-TQ-A7RJ | TCGA-LGG | 33.625406 | 13.9387361 |
| TCGA-TQ-A7RK | TCGA-LGG | 47.4642383 | 20.3100035 |
| TCGA-TQ-A7RM | TCGA-LGG | 73.2157195 | 40.70367 |
| TCGA-TQ-A7RN | TCGA-LGG | 19.5045702 | 22.1531447 |
| TCGA-TQ-A7RO | TCGA-LGG | 20.0571376 | 16.7037269 |
| TCGA-TQ-A7RP | TCGA-LGG | 34.2926449 | 14.3504366 |
| TCGA-TQ-A7RQ | TCGA-LGG | 18.8280978 | 17.6111188 |
| TCGA-TQ-A7RR | TCGA-LGG | 39.3020057 | 18.7204568 |
| TCGA-TQ-A7RS | TCGA-LGG | 24.7100133 | 23.6428073 |
| TCGA-TQ-A7RV | TCGA-LGG | 31.326387 | 15.7558543 |
| TCGA-TQ-A7RW | TCGA-LGG | 35.5992324 | 13.4501073 |
| TCGA-TQ-A8XE | TCGA-LGG | 45.4499052 | 15.4100332 |
| TCGA-UB-A7MA | TCGA-LIHC | 56.3775447 | 8.98036354 |
| TCGA-UB-A7MB | TCGA-LIHC | 67.8394315 | 32.9356428 |
| TCGA-UB-A7MD | TCGA-LIHC | 49.0980322 | 22.9764179 |
| TCGA-UB-A7ME | TCGA-LIHC | 41.3545776 | 31.0771943 |
| TCGA-UB-A7MF | TCGA-LIHC | 67.9753831 | 31.2945846 |
| TCGA-UB-AA0U | TCGA-LIHC | 45.5713061 | 15.3392871 |
| TCGA-UB-AA0V | TCGA-LIHC | 33.6739691 | 11.7250166 |
| TCGA-UL-AAZ6 | TCGA-BRCA | 52.895073 | 40.1863164 |
| TCGA-UN-AAZ9 | TCGA-KIRP | 34.2222557 | 38.0878054 |

| | | | |
|---|---|---|---|
| TCGA-US-A774 | TCGA-PAAD | 43.6633304 | 12.0418248 |
| TCGA-US-A776 | TCGA-PAAD | 89.4484597 | 26.2416048 |
| TCGA-US-A779 | TCGA-PAAD | 61.6703891 | 10.5412239 |
| TCGA-US-A77E | TCGA-PAAD | 49.9610829 | 16.6036437 |
| TCGA-US-A77G | TCGA-PAAD | 61.6049442 | 16.5807069 |
| TCGA-US-A77J | TCGA-PAAD | 48.3840303 | 13.8401122 |
| TCGA-UU-A93S | TCGA-BRCA | 57.6417563 | 69.0787626 |
| TCGA-UY-A78K | TCGA-BLCA | 84.8668636 | 30.2773724 |
| TCGA-UY-A78L | TCGA-BLCA | 76.6866397 | 36.9237672 |
| TCGA-UY-A78M | TCGA-BLCA | 89.4241115 | 32.2121201 |
| TCGA-UY-A78N | TCGA-BLCA | 108.64277 | 33.1300736 |
| TCGA-UY-A78O | TCGA-BLCA | 58.6650591 | 23.6596753 |
| TCGA-UY-A78P | TCGA-BLCA | 122.503394 | 87.4599613 |
| TCGA-UY-A8OB | TCGA-BLCA | 149.397827 | 54.3313458 |
| TCGA-UY-A8OC | TCGA-BLCA | 70.2194076 | 22.5888208 |
| TCGA-UY-A8OD | TCGA-BLCA | 42.7415827 | 15.1804749 |
| TCGA-UY-A9PA | TCGA-BLCA | 80.8963806 | 31.6951315 |
| TCGA-UY-A9PB | TCGA-BLCA | 110.423409 | 31.3986444 |
| TCGA-UY-A9PD | TCGA-BLCA | 87.6750087 | 20.6772584 |
| TCGA-UY-A9PE | TCGA-BLCA | 99.6572406 | 25.7896287 |
| TCGA-UY-A9PF | TCGA-BLCA | 50.9893429 | 19.1438723 |
| TCGA-UY-A9PH | TCGA-BLCA | 94.474473 | 51.7819253 |
| TCGA-UZ-A9PJ | TCGA-KIRP | 80.6944595 | 7.65091563 |
| TCGA-UZ-A9PK | TCGA-KIRP | 43.893565 | 13.8452637 |
| TCGA-UZ-A9PL | TCGA-KIRP | 34.1347192 | 15.9735902 |
| TCGA-UZ-A9PM | TCGA-KIRP | 8.26302289 | 5.73943049 |
| TCGA-UZ-A9PN | TCGA-KIRP | 56.0812327 | 25.6839995 |
| TCGA-UZ-A9PO | TCGA-KIRP | 4.02954437 | 1.61588459 |
| TCGA-UZ-A9PP | TCGA-KIRP | 26.9616487 | 27.085 |
| TCGA-UZ-A9PQ | TCGA-KIRP | 27.2848306 | 23.7960428 |
| TCGA-UZ-A9PR | TCGA-KIRP | 64.7559268 | 16.7477237 |

| | | | |
|---|---|---|---|
| TCGA-UZ-A9PS | TCGA-KIRP | 33.4216773 | 19.2457097 |
| TCGA-UZ-A9PV | TCGA-KIRP | 59.4073655 | 18.561297 |
| TCGA-UZ-A9PX | TCGA-KIRP | 50.3921343 | 20.331925 |
| TCGA-UZ-A9PZ | TCGA-KIRP | 73.4198394 | 15.0897644 |
| TCGA-UZ-A9Q0 | TCGA-KIRP | 0.67398684 | 0.36268507 |
| TCGA-UZ-A9Q1 | TCGA-KIRP | 29.4117406 | 13.2250488 |
| TCGA-V1-A8MF | TCGA-PRAD | 21.6308303 | 5.71824701 |
| TCGA-V1-A8MG | TCGA-PRAD | 32.7202408 | 13.1028688 |
| TCGA-V1-A8MJ | TCGA-PRAD | 60.5983089 | 26.1952135 |
| TCGA-V1-A8MK | TCGA-PRAD | 42.3175374 | 18.2103321 |
| TCGA-V1-A8ML | TCGA-PRAD | 42.1533173 | 17.9102754 |
| TCGA-V1-A8MM | TCGA-PRAD | 41.0358658 | 17.147927 |
| TCGA-V1-A8MU | TCGA-PRAD | 51.1732034 | 20.59195 |
| TCGA-V1-A8WL | TCGA-PRAD | 45.9738442 | 17.6640583 |
| TCGA-V1-A8WN | TCGA-PRAD | 48.0337237 | 17.010527 |
| TCGA-V1-A8WS | TCGA-PRAD | 47.4759014 | 14.9184308 |
| TCGA-V1-A8WV | TCGA-PRAD | 44.093601 | 23.1895587 |
| TCGA-V1-A8WW | TCGA-PRAD | 34.7204437 | 24.9034217 |
| TCGA-V1-A8X3 | TCGA-PRAD | 48.1324063 | 18.1613983 |
| TCGA-V1-A9O5 | TCGA-PRAD | 67.7879546 | 38.1138479 |
| TCGA-V1-A9O7 | TCGA-PRAD | 42.4372663 | 17.3005125 |
| TCGA-V1-A9O9 | TCGA-PRAD | 41.6754773 | 10.7404824 |
| TCGA-V1-A9OA | TCGA-PRAD | 31.4594485 | 18.6475728 |
| TCGA-V1-A9OF | TCGA-PRAD | 43.9278724 | 20.4072914 |
| TCGA-V1-A9OH | TCGA-PRAD | 42.0933149 | 17.7902007 |
| TCGA-V1-A9OL | TCGA-PRAD | 70.0181821 | 28.9441176 |
| TCGA-V1-A9OQ | TCGA-PRAD | 36.881423 | 19.5844959 |
| TCGA-V1-A9OT | TCGA-PRAD | 19.4756913 | 14.0915522 |
| TCGA-V1-A9OX | TCGA-PRAD | 43.1513568 | 14.1896522 |
| TCGA-V1-A9OY | TCGA-PRAD | 54.9185074 | 22.8005865 |
| TCGA-V1-A9Z7 | TCGA-PRAD | 37.0885731 | 18.3426985 |

| | | | |
|---|---|---|---|
| TCGA-V1-A9Z8 | TCGA-PRAD | 58.255861 | 20.0141315 |
| TCGA-V1-A9Z9 | TCGA-PRAD | 42.2203926 | 24.6236952 |
| TCGA-V1-A9ZG | TCGA-PRAD | 37.272788 | 16.8115991 |
| TCGA-V1-A9ZI | TCGA-PRAD | 42.9078659 | 14.4213106 |
| TCGA-V1-A9ZK | TCGA-PRAD | 38.6631826 | 23.9291218 |
| TCGA-V1-A9ZR | TCGA-PRAD | 45.4642994 | 14.962247 |
| TCGA-V5-A7RB | TCGA-ESCA | 38.6019912 | 10.747209 |
| TCGA-V5-A7RC | TCGA-ESCA | 32.2142252 | 20.7017147 |
| TCGA-V5-A7RE | TCGA-ESCA | 38.4422796 | 30.0903536 |
| TCGA-V5-AASV | TCGA-ESCA | 44.3008105 | 24.1970602 |
| TCGA-V5-AASW | TCGA-ESCA | 33.1423683 | 36.7970911 |
| TCGA-V5-AASX | TCGA-ESCA | 27.6978572 | 23.2195701 |
| TCGA-V7-A7HQ | TCGA-BRCA | 43.5592302 | 18.3136279 |
| TCGA-V9-A7HT | TCGA-KIRP | 86.4229405 | 8.84370273 |
| TCGA-VF-A8A8 | TCGA-TGCT | 169.515302 | 126.999061 |
| TCGA-VF-A8A9 | TCGA-TGCT | 58.1092946 | 76.4959202 |
| TCGA-VF-A8AA | TCGA-TGCT | 46.8017313 | 56.9423936 |
| TCGA-VF-A8AB | TCGA-TGCT | 50.7824498 | 51.4294105 |
| TCGA-VF-A8AD | TCGA-TGCT | 81.4040041 | 475.944347 |
| TCGA-VF-A8AE | TCGA-TGCT | 36.8978811 | 16.3755736 |
| TCGA-VG-A8LO | TCGA-OV | 56.6762118 | 13.7397944 |
| TCGA-VM-A8C8 | TCGA-LGG | 44.5259539 | 15.6807662 |
| TCGA-VM-A8C9 | TCGA-LGG | 37.9611844 | 16.774803 |
| TCGA-VM-A8CA | TCGA-LGG | 33.9434123 | 18.3005831 |
| TCGA-VM-A8CB | TCGA-LGG | 26.8832491 | 42.3758194 |
| TCGA-VM-A8CD | TCGA-LGG | 46.4578499 | 25.4003394 |
| TCGA-VM-A8CE | TCGA-LGG | 22.3478377 | 14.778874 |
| TCGA-VM-A8CF | TCGA-LGG | 42.2324074 | 15.3758878 |
| TCGA-VM-A8CH | TCGA-LGG | 32.4688469 | 10.0674496 |
| TCGA-VN-A88I | TCGA-PRAD | 47.1268422 | 15.7853252 |
| TCGA-VN-A88K | TCGA-PRAD | 50.6364722 | 17.9570107 |

| | | | |
|---|---|---|---|
| TCGA-VN-A88L | TCGA-PRAD | 45.5787259 | 19.7321903 |
| TCGA-VN-A88M | TCGA-PRAD | 42.3748094 | 19.5542724 |
| TCGA-VN-A88N | TCGA-PRAD | 54.1052049 | 26.0135912 |
| TCGA-VN-A88O | TCGA-PRAD | 37.9421953 | 16.6629287 |
| TCGA-VN-A88P | TCGA-PRAD | 26.4600497 | 16.0515823 |
| TCGA-VN-A88Q | TCGA-PRAD | 45.0706398 | 19.1450551 |
| TCGA-VN-A88R | TCGA-PRAD | 58.0563703 | 19.0183268 |
| TCGA-VN-A943 | TCGA-PRAD | 39.0888464 | 19.0094175 |
| TCGA-VP-A872 | TCGA-PRAD | 53.374084 | 17.7471875 |
| TCGA-VP-A875 | TCGA-PRAD | 31.2700762 | 20.9109639 |
| TCGA-VP-A876 | TCGA-PRAD | 56.8200801 | 16.2027098 |
| TCGA-VP-A878 | TCGA-PRAD | 50.5486669 | 19.9555243 |
| TCGA-VP-A879 | TCGA-PRAD | 41.0410824 | 23.5558819 |
| TCGA-VP-A87B | TCGA-PRAD | 21.0098348 | 10.533263 |
| TCGA-VP-A87C | TCGA-PRAD | 55.2466968 | 15.0832197 |
| TCGA-VP-A87D | TCGA-PRAD | 45.982735 | 21.7337953 |
| TCGA-VP-A87E | TCGA-PRAD | 31.0521766 | 13.644618 |
| TCGA-VP-A87H | TCGA-PRAD | 34.9668894 | 16.0817462 |
| TCGA-VP-A87J | TCGA-PRAD | 52.2429187 | 14.2048705 |
| TCGA-VP-A87K | TCGA-PRAD | 64.8374802 | 11.241685 |
| TCGA-VP-AA1N | TCGA-PRAD | 52.9335489 | 27.4120101 |
| TCGA-VQ-A8DT | TCGA-STAD | 31.7611366 | 30.6023363 |
| TCGA-VQ-A8DU | TCGA-STAD | 21.9687491 | 19.9228621 |
| TCGA-VQ-A8DV | TCGA-STAD | 41.4719885 | 67.6902355 |
| TCGA-VQ-A8DZ | TCGA-STAD | 38.3680467 | 26.7187627 |
| TCGA-VQ-A8E2 | TCGA-STAD | 59.6372799 | 30.15374 |
| TCGA-VQ-A8E3 | TCGA-STAD | 56.4894756 | 40.4754234 |
| TCGA-VQ-A8E7 | TCGA-STAD | 34.0483607 | 15.0062505 |
| TCGA-VQ-A8P2 | TCGA-STAD | 48.2892905 | 31.2354141 |
| TCGA-VQ-A8P3 | TCGA-STAD | 29.5290658 | 35.1415522 |
| TCGA-VQ-A8P5 | TCGA-STAD | 46.3086919 | 71.8468385 |

| | | | |
|---|---|---|---|
| TCGA-VQ-A8P8 | TCGA-STAD | 21.3822271 | 21.6809961 |
| TCGA-VQ-A8PB | TCGA-STAD | 39.8058942 | 27.0948824 |
| TCGA-VQ-A8PC | TCGA-STAD | 35.4995973 | 20.3348819 |
| TCGA-VQ-A8PD | TCGA-STAD | 24.8920035 | 12.9382372 |
| TCGA-VQ-A8PE | TCGA-STAD | 67.5851806 | 30.7240211 |
| TCGA-VQ-A8PF | TCGA-STAD | 54.5203149 | 21.453722 |
| TCGA-VQ-A8PH | TCGA-STAD | 63.8213644 | 25.9446808 |
| TCGA-VQ-A8PJ | TCGA-STAD | 37.9184339 | 26.7569129 |
| TCGA-VQ-A8PK | TCGA-STAD | 30.2254271 | 50.1252243 |
| TCGA-VQ-A8PM | TCGA-STAD | 15.2775106 | 12.6811153 |
| TCGA-VQ-A8PO | TCGA-STAD | 40.6971182 | 28.7935834 |
| TCGA-VQ-A8PP | TCGA-STAD | 39.0352789 | 20.1210347 |
| TCGA-VQ-A8PQ | TCGA-STAD | 24.6263541 | 21.4727838 |
| TCGA-VQ-A8PU | TCGA-STAD | 51.1117333 | 40.2684323 |
| TCGA-VQ-A8PX | TCGA-STAD | 33.6960513 | 25.1157854 |
| TCGA-VQ-A91A | TCGA-STAD | 25.2341845 | 49.8849842 |
| TCGA-VQ-A91D | TCGA-STAD | 40.1090714 | 22.4771239 |
| TCGA-VQ-A91E | TCGA-STAD | 31.9422341 | 31.1042245 |
| TCGA-VQ-A91K | TCGA-STAD | 43.5925786 | 18.3645902 |
| TCGA-VQ-A91N | TCGA-STAD | 50.7150093 | 16.4692601 |
| TCGA-VQ-A91Q | TCGA-STAD | 35.760617 | 24.5493245 |
| TCGA-VQ-A91S | TCGA-STAD | 49.5615001 | 25.050815 |
| TCGA-VQ-A91U | TCGA-STAD | 40.2872326 | 12.5597606 |
| TCGA-VQ-A91V | TCGA-STAD | 44.4601067 | 20.7948046 |
| TCGA-VQ-A91X | TCGA-STAD | 51.2783308 | 16.1341321 |
| TCGA-VQ-A91Y | TCGA-STAD | 34.4186857 | 20.8540515 |
| TCGA-VQ-A91Z | TCGA-STAD | 29.8407137 | 14.6231529 |
| TCGA-VQ-A922 | TCGA-STAD | 45.0617597 | 14.2065026 |
| TCGA-VQ-A923 | TCGA-STAD | 43.3559204 | 18.7107292 |
| TCGA-VQ-A924 | TCGA-STAD | 52.1619395 | 39.8666829 |
| TCGA-VQ-A925 | TCGA-STAD | 20.4904612 | 19.9738688 |

| | | | |
|---|---|---|---|
| TCGA-VQ-A927 | TCGA-STAD | 20.0459206 | 15.3956743 |
| TCGA-VQ-A928 | TCGA-STAD | 26.445523 | 21.9373556 |
| TCGA-VQ-A92D | TCGA-STAD | 37.7656469 | 23.2278917 |
| TCGA-VQ-A94O | TCGA-STAD | 43.1209368 | 38.6355215 |
| TCGA-VQ-A94P | TCGA-STAD | 24.6476423 | 18.2019549 |
| TCGA-VQ-A94R | TCGA-STAD | 40.2522476 | 15.6900372 |
| TCGA-VQ-A94T | TCGA-STAD | 38.7126987 | 71.7107596 |
| TCGA-VQ-A94U | TCGA-STAD | 52.7657781 | 43.8831401 |
| TCGA-VQ-AA64 | TCGA-STAD | 51.8495234 | 46.7133291 |
| TCGA-VQ-AA68 | TCGA-STAD | 37.2953861 | 21.5483642 |
| TCGA-VQ-AA69 | TCGA-STAD | 75.2810956 | 30.8864913 |
| TCGA-VQ-AA6A | TCGA-STAD | 58.4633696 | 29.1767984 |
| TCGA-VQ-AA6D | TCGA-STAD | 65.5668624 | 20.668137 |
| TCGA-VQ-AA6F | TCGA-STAD | 30.4942785 | 21.4313189 |
| TCGA-VQ-AA6G | TCGA-STAD | 34.3296338 | 17.3503747 |
| TCGA-VQ-AA6J | TCGA-STAD | 32.1220325 | 34.6135087 |
| TCGA-VQ-AA6K | TCGA-STAD | 24.3642398 | 65.1046169 |
| TCGA-VR-A8EO | TCGA-ESCA | 83.7900574 | 38.5721621 |
| TCGA-VR-A8EP | TCGA-ESCA | 38.1367754 | 20.5362168 |
| TCGA-VR-A8EQ | TCGA-ESCA | 80.3442284 | 22.4573363 |
| TCGA-VR-A8ER | TCGA-ESCA | 29.8912486 | 28.8666836 |
| TCGA-VR-A8ET | TCGA-ESCA | 19.140146 | 22.4954657 |
| TCGA-VR-A8EU | TCGA-ESCA | 47.4815229 | 41.2969211 |
| TCGA-VR-A8EW | TCGA-ESCA | 47.9942781 | 32.5664801 |
| TCGA-VR-A8EX | TCGA-ESCA | 43.6560369 | 32.6039143 |
| TCGA-VR-A8EY | TCGA-ESCA | 44.1712536 | 12.528237 |
| TCGA-VR-A8EZ | TCGA-ESCA | 69.1093837 | 37.2207692 |
| TCGA-VR-A8Q7 | TCGA-ESCA | 49.1311952 | 49.4367705 |
| TCGA-VR-AA4D | TCGA-ESCA | 53.7466166 | 29.5500345 |
| TCGA-VR-AA4G | TCGA-ESCA | 54.5575426 | 38.8099582 |
| TCGA-VR-AA7I | TCGA-ESCA | 43.735942 | 24.2326274 |

| | | | |
|---|---|---|---|
| TCGA-VV-A829 | TCGA-LGG | 20.6146998 | 19.7793874 |
| TCGA-VV-A86M | TCGA-LGG | 44.0663098 | 20.9263302 |
| TCGA-VW-A7QS | TCGA-LGG | 31.6143916 | 29.8747318 |
| TCGA-VW-A8FI | TCGA-LGG | 48.018556 | 23.9786402 |
| TCGA-W3-A824 | TCGA-SKCM | 46.6985835 | 20.7164248 |
| TCGA-W3-A825 | TCGA-SKCM | 73.9663634 | 42.0253574 |
| TCGA-W3-A828 | TCGA-SKCM | 71.5775933 | 23.998226 |
| TCGA-W3-AA1O | TCGA-SKCM | 77.9483574 | 61.6042738 |
| TCGA-W3-AA1Q | TCGA-SKCM | 24.5623461 | 16.7449258 |
| TCGA-W3-AA1R | TCGA-SKCM | 110.418532 | 74.8876454 |
| TCGA-W3-AA1V | TCGA-SKCM | 116.14263 | 99.3865094 |
| TCGA-W3-AA1W | TCGA-SKCM | 58.1281965 | 38.9155137 |
| TCGA-W3-AA21 | TCGA-SKCM | 69.951243 | 49.1978804 |
| TCGA-W4-A7U2 | TCGA-TGCT | 62.3278711 | 65.1547845 |
| TCGA-W4-A7U3 | TCGA-TGCT | 72.05132 | 76.2543763 |
| TCGA-W4-A7U4 | TCGA-TGCT | 114.725522 | 129.372287 |
| TCGA-W8-A86G | TCGA-BRCA | 54.5470804 | 17.6308128 |
| TCGA-W9-A837 | TCGA-LGG | 25.3676089 | 17.925015 |
| TCGA-WE-A8JZ | TCGA-SKCM | 52.5301989 | 26.9299636 |
| TCGA-WE-A8K1 | TCGA-SKCM | 53.1960592 | 23.6906671 |
| TCGA-WE-A8K4 | TCGA-SKCM | 56.6513832 | 29.717181 |
| TCGA-WE-A8K5 | TCGA-SKCM | 55.0614954 | 35.4084673 |
| TCGA-WE-A8K6 | TCGA-SKCM | 26.9136701 | 38.718567 |
| TCGA-WE-A8ZM | TCGA-SKCM | 35.4031375 | 24.0195782 |
| TCGA-WE-A8ZN | TCGA-SKCM | 38.3031147 | 15.2018913 |
| TCGA-WE-A8ZO | TCGA-SKCM | 54.2710637 | 26.6572728 |
| TCGA-WE-A8ZQ | TCGA-SKCM | 77.5954137 | 53.8099211 |
| TCGA-WE-A8ZR | TCGA-SKCM | 75.5094307 | 17.7229204 |
| TCGA-WE-A8ZT | TCGA-SKCM | 54.1051666 | 30.5936253 |
| TCGA-WE-A8ZX | TCGA-SKCM | 58.4378357 | 51.3430129 |
| TCGA-WE-A8ZY | TCGA-SKCM | 106.274987 | 41.0894338 |

| | | | |
|---|---|---|---|
| TCGA-WE-AA9Y | TCGA-SKCM | 33.4363861 | 28.2705967 |
| TCGA-WE-AAA0 | TCGA-SKCM | 75.3168686 | 23.4966788 |
| TCGA-WE-AAA3 | TCGA-SKCM | 34.0317343 | 21.2384195 |
| TCGA-WE-AAA4 | TCGA-SKCM | 29.5248106 | 12.2519216 |
| TCGA-WH-A86K | TCGA-LGG | 35.905896 | 13.5996533 |
| TCGA-WJ-A86L | TCGA-LIHC | 19.3886632 | 9.585734 |
| TCGA-WN-A9G9 | TCGA-KIRP | 39.6861356 | 14.7539808 |
| TCGA-WN-AB4C | TCGA-KIRP | 33.7997059 | 11.5558128 |
| TCGA-WQ-A9G7 | TCGA-LIHC | 35.3589013 | 35.6850595 |
| TCGA-WQ-AB4B | TCGA-LIHC | 36.8818867 | 31.3248847 |
| TCGA-WR-A838 | TCGA-OV | 75.1860467 | 37.0969421 |
| TCGA-WS-AB45 | TCGA-COAD | 41.8540067 | 25.133908 |
| TCGA-WT-AB41 | TCGA-BRCA | 27.7028103 | 31.8257737 |
| TCGA-WT-AB44 | TCGA-BRCA | 37.7888939 | 30.827655 |
| TCGA-WW-A8ZI | TCGA-PRAD | 30.8257435 | 13.9376012 |
| TCGA-WX-AA44 | TCGA-LIHC | 50.2699867 | 27.4031092 |
| TCGA-WX-AA46 | TCGA-LIHC | 38.3696637 | 18.6205049 |
| TCGA-WX-AA47 | TCGA-LIHC | 28.2550598 | 10.1693228 |
| TCGA-WY-A858 | TCGA-LGG | 30.0926001 | 9.68398255 |
| TCGA-WY-A859 | TCGA-LGG | 50.1893762 | 17.9341407 |
| TCGA-WY-A85A | TCGA-LGG | 30.5549601 | 12.6786829 |
| TCGA-WY-A85B | TCGA-LGG | 44.7251649 | 22.0448926 |
| TCGA-WY-A85C | TCGA-LGG | 29.0328302 | 9.70418012 |
| TCGA-WY-A85D | TCGA-LGG | 50.8747776 | 17.3610108 |
| TCGA-WY-A85E | TCGA-LGG | 38.4201144 | 18.9927782 |
| TCGA-WZ-A7V3 | TCGA-TGCT | 50.6802915 | 48.5109617 |
| TCGA-WZ-A7V4 | TCGA-TGCT | 31.9362562 | 34.4994448 |
| TCGA-WZ-A7V5 | TCGA-TGCT | 35.5418565 | 50.1504997 |
| TCGA-WZ-A8D5 | TCGA-TGCT | 40.5214887 | 59.2483149 |
| TCGA-X3-A8G4 | TCGA-TGCT | 69.0391525 | 176.969156 |
| TCGA-X4-A8KQ | TCGA-PRAD | 33.7921197 | 13.5551301 |

| | | | |
|---|---|---|---|
| TCGA-X4-A8KS | TCGA-PRAD | 24.5300847 | 11.2215413 |
| TCGA-XA-A8JR | TCGA-PRAD | 38.6191693 | 17.0347304 |
| TCGA-XC-AA0X | TCGA-LUSC | 52.4780472 | 16.9409906 |
| TCGA-XD-AAUG | TCGA-PAAD | 41.0161111 | 14.3120267 |
| TCGA-XD-AAUH | TCGA-PAAD | 35.1213056 | 15.204484 |
| TCGA-XD-AAUI | TCGA-PAAD | 51.2683746 | 13.7279437 |
| TCGA-XD-AAUL | TCGA-PAAD | 54.4767282 | 15.2287375 |
| TCGA-XE-A8H1 | TCGA-TGCT | 137.837295 | 199.503553 |
| TCGA-XE-A8H4 | TCGA-TGCT | 59.1849964 | 76.511341 |
| TCGA-XE-A8H5 | TCGA-TGCT | 25.266929 | 60.5152627 |
| TCGA-XE-A9SE | TCGA-TGCT | 50.6508959 | 82.6726595 |
| TCGA-XE-AANI | TCGA-TGCT | 54.6475249 | 45.4537193 |
| TCGA-XE-AANJ | TCGA-TGCT | 65.8471959 | 62.2964057 |
| TCGA-XE-AANR | TCGA-TGCT | 34.6397852 | 49.8949294 |
| TCGA-XE-AANV | TCGA-TGCT | 42.2642403 | 42.3882335 |
| TCGA-XE-AAO3 | TCGA-TGCT | 75.515376 | 105.689793 |
| TCGA-XE-AAO4 | TCGA-TGCT | 57.4922619 | 71.0125077 |
| TCGA-XE-AAO6 | TCGA-TGCT | 134.909965 | 19.9500916 |
| TCGA-XE-AAOB | TCGA-TGCT | 15.5268761 | 15.8938047 |
| TCGA-XE-AAOC | TCGA-TGCT | 64.6601382 | 22.4395094 |
| TCGA-XE-AAOD | TCGA-TGCT | 33.6886336 | 29.0713842 |
| TCGA-XE-AAOF | TCGA-TGCT | 48.1054246 | 40.3444849 |
| TCGA-XE-AAOJ | TCGA-TGCT | 121.6077 | 89.1858964 |
| TCGA-XE-AAOL | TCGA-TGCT | 24.4985466 | 189.004329 |
| TCGA-XF-A8HB | TCGA-BLCA | 93.5172896 | 28.3776787 |
| TCGA-XF-A8HC | TCGA-BLCA | 75.928923 | 39.0611408 |
| TCGA-XF-A8HD | TCGA-BLCA | 85.4306609 | 21.7439595 |
| TCGA-XF-A8HE | TCGA-BLCA | 82.4495652 | 31.965626 |
| TCGA-XF-A8HF | TCGA-BLCA | 51.267947 | 24.9012477 |
| TCGA-XF-A8HG | TCGA-BLCA | 30.1784681 | 13.7361239 |
| TCGA-XF-A8HH | TCGA-BLCA | 55.9160577 | 16.3926434 |

| | | | |
|---|---|---|---|
| TCGA-XF-A8HI | TCGA-BLCA | 70.5591137 | 18.8642219 |
| TCGA-XF-A9SH | TCGA-BLCA | 49.53118 | 11.291991 |
| TCGA-XF-A9SI | TCGA-BLCA | 82.5635363 | 52.9642842 |
| TCGA-XF-A9SJ | TCGA-BLCA | 47.7071173 | 27.4211211 |
| TCGA-XF-A9SK | TCGA-BLCA | 81.3860098 | 45.1589898 |
| TCGA-XF-A9SL | TCGA-BLCA | 45.9095863 | 24.2484718 |
| TCGA-XF-A9SM | TCGA-BLCA | 79.9312131 | 40.4716846 |
| TCGA-XF-A9SP | TCGA-BLCA | 51.0307835 | 15.750985 |
| TCGA-XF-A9ST | TCGA-BLCA | 182.688225 | 41.8807291 |
| TCGA-XF-A9SU | TCGA-BLCA | 65.6262207 | 32.605333 |
| TCGA-XF-A9SV | TCGA-BLCA | 57.308273 | 17.5861174 |
| TCGA-XF-A9SW | TCGA-BLCA | 45.1977462 | 32.2675628 |
| TCGA-XF-A9SX | TCGA-BLCA | 95.7310107 | 34.1395495 |
| TCGA-XF-A9SY | TCGA-BLCA | 95.9574742 | 28.0341406 |
| TCGA-XF-A9SZ | TCGA-BLCA | 102.367553 | 25.7043817 |
| TCGA-XF-A9T0 | TCGA-BLCA | 71.4212459 | 31.9007085 |
| TCGA-XF-A9T2 | TCGA-BLCA | 101.756719 | 26.1524328 |
| TCGA-XF-A9T3 | TCGA-BLCA | 72.6705266 | 26.2178654 |
| TCGA-XF-A9T4 | TCGA-BLCA | 66.8337106 | 31.9268978 |
| TCGA-XF-A9T5 | TCGA-BLCA | 91.2773934 | 31.6821645 |
| TCGA-XF-A9T6 | TCGA-BLCA | 119.572228 | 42.9481035 |
| TCGA-XF-A9T8 | TCGA-BLCA | 74.5789585 | 28.9443297 |
| TCGA-XF-AAME | TCGA-BLCA | 70.898266 | 23.9021209 |
| TCGA-XF-AAMG | TCGA-BLCA | 105.451564 | 39.937194 |
| TCGA-XF-AAMH | TCGA-BLCA | 58.836202 | 19.4502678 |
| TCGA-XF-AAMJ | TCGA-BLCA | 68.4674103 | 28.1198222 |
| TCGA-XF-AAML | TCGA-BLCA | 158.101543 | 26.9112872 |
| TCGA-XF-AAMQ | TCGA-BLCA | 81.5809705 | 36.9087611 |
| TCGA-XF-AAMR | TCGA-BLCA | 56.7450832 | 18.6292586 |
| TCGA-XF-AAMT | TCGA-BLCA | 86.6138874 | 23.322796 |
| TCGA-XF-AAMW | TCGA-BLCA | 66.7713917 | 26.4730557 |

| | | | |
|---|---|---|---|
| TCGA-XF-AAMX | TCGA-BLCA | 73.2825093 | 37.8736195 |
| TCGA-XF-AAMY | TCGA-BLCA | 51.2848617 | 25.7840537 |
| TCGA-XF-AAMZ | TCGA-BLCA | 129.564396 | 39.6499412 |
| TCGA-XF-AAN0 | TCGA-BLCA | 44.6825334 | 17.1376996 |
| TCGA-XF-AAN1 | TCGA-BLCA | 58.1620396 | 27.7353565 |
| TCGA-XF-AAN2 | TCGA-BLCA | 76.1366694 | 44.3881274 |
| TCGA-XF-AAN3 | TCGA-BLCA | 59.3255701 | 11.2238997 |
| TCGA-XF-AAN4 | TCGA-BLCA | 52.5545701 | 26.7584258 |
| TCGA-XF-AAN5 | TCGA-BLCA | 54.9311493 | 55.7255756 |
| TCGA-XF-AAN7 | TCGA-BLCA | 106.82428 | 53.8146343 |
| TCGA-XF-AAN8 | TCGA-BLCA | 43.9072293 | 14.3163359 |
| TCGA-XJ-A83F | TCGA-PRAD | 43.6106861 | 9.52596391 |
| TCGA-XJ-A83G | TCGA-PRAD | 34.3515532 | 19.2110859 |
| TCGA-XJ-A83H | TCGA-PRAD | 38.1966968 | 8.31853946 |
| TCGA-XJ-A9DK | TCGA-PRAD | 58.9784817 | 24.6315729 |
| TCGA-XJ-A9DQ | TCGA-PRAD | 43.5816115 | 14.9593381 |
| TCGA-XJ-A9DX | TCGA-PRAD | 36.5641867 | 11.351729 |
| TCGA-XK-AAIR | TCGA-PRAD | 38.994817 | 13.3464795 |
| TCGA-XK-AAIV | TCGA-PRAD | 49.8745538 | 15.547686 |
| TCGA-XK-AAIW | TCGA-PRAD | 51.3079313 | 27.7357444 |
| TCGA-XK-AAJ3 | TCGA-PRAD | 39.7409418 | 18.9636795 |
| TCGA-XK-AAJA | TCGA-PRAD | 46.0458793 | 16.0833082 |
| TCGA-XK-AAJP | TCGA-PRAD | 35.9768854 | 14.1260237 |
| TCGA-XK-AAJR | TCGA-PRAD | 41.5443799 | 19.2146747 |
| TCGA-XK-AAJT | TCGA-PRAD | 35.9574326 | 16.3217758 |
| TCGA-XK-AAJU | TCGA-PRAD | 47.3257482 | 17.7161749 |
| TCGA-XK-AAK1 | TCGA-PRAD | 40.3232176 | 17.8943647 |
| TCGA-XN-A8T3 | TCGA-PAAD | 61.5833039 | 13.0376731 |
| TCGA-XN-A8T5 | TCGA-PAAD | 50.4219519 | 17.2920604 |
| TCGA-XP-A8T6 | TCGA-ESCA | 47.3898188 | 18.39341 |
| TCGA-XP-A8T8 | TCGA-ESCA | 39.2062465 | 51.2906432 |

| | | | |
|---|---|---|---|
| TCGA-XQ-A8TA | TCGA-PRAD | 58.079442 | 8.85967339 |
| TCGA-XQ-A8TB | TCGA-PRAD | 30.1563224 | 17.8302327 |
| TCGA-XR-A8TC | TCGA-LIHC | 23.5213264 | 22.9483242 |
| TCGA-XR-A8TD | TCGA-LIHC | 44.1726098 | 18.9221343 |
| TCGA-XR-A8TE | TCGA-LIHC | 39.228709 | 13.1074186 |
| TCGA-XR-A8TF | TCGA-LIHC | 49.438477 | 19.3252489 |
| TCGA-XR-A8TG | TCGA-LIHC | 50.5948422 | 12.5660889 |
| TCGA-XV-A9VZ | TCGA-SKCM | 77.8294992 | 16.3698239 |
| TCGA-XV-A9W2 | TCGA-SKCM | 56.8158123 | 28.7521091 |
| TCGA-XV-A9W5 | TCGA-SKCM | 62.4057789 | 29.2021511 |
| TCGA-XV-AAZV | TCGA-SKCM | 45.9486744 | 41.7274118 |
| TCGA-XV-AAZW | TCGA-SKCM | 63.7541642 | 27.5964794 |
| TCGA-XV-AAZY | TCGA-SKCM | 30.04691 | 16.4996676 |
| TCGA-XV-AB01 | TCGA-SKCM | 16.6689599 | 9.23249627 |
| TCGA-XX-A899 | TCGA-BRCA | 45.9783971 | 20.6297335 |
| TCGA-XX-A89A | TCGA-BRCA | 50.4564193 | 44.1256595 |
| TCGA-XY-A89B | TCGA-TGCT | 38.7108027 | 30.4092821 |
| TCGA-XY-A8S2 | TCGA-TGCT | 275.905896 | 32.2151793 |
| TCGA-XY-A8S3 | TCGA-TGCT | 193.50816 | 129.828259 |
| TCGA-XY-A9T9 | TCGA-TGCT | 23.6504974 | 5.43082982 |
| TCGA-Y6-A8TL | TCGA-PRAD | 37.5091936 | 16.4559434 |
| TCGA-Y6-A9XI | TCGA-PRAD | 62.5258942 | 19.6952364 |
| TCGA-Y8-A894 | TCGA-KIRP | 55.5218211 | 17.074965 |
| TCGA-Y8-A896 | TCGA-KIRP | 41.5250741 | 17.1371371 |
| TCGA-Y8-A897 | TCGA-KIRP | 14.9065433 | 15.0772158 |
| TCGA-Y8-A898 | TCGA-KIRP | 12.3609376 | 7.23426948 |
| TCGA-Y8-A8RY | TCGA-KIRP | 32.9412506 | 15.6832655 |
| TCGA-Y8-A8RZ | TCGA-KIRP | 39.4011138 | 10.9857386 |
| TCGA-Y8-A8S0 | TCGA-KIRP | 11.0063796 | 13.5825426 |
| TCGA-Y8-A8S1 | TCGA-KIRP | 22.5681671 | 4.60549483 |
| TCGA-YA-A8S7 | TCGA-LIHC | 76.8749746 | 42.6676419 |

| | | | |
|---|---|---|---|
| TCGA-YB-A89D | TCGA-PAAD | 45.5663457 | 15.6471187 |
| TCGA-YC-A89H | TCGA-BLCA | 64.3543827 | 29.1951233 |
| TCGA-YC-A8S6 | TCGA-BLCA | 66.9583481 | 27.6226072 |
| TCGA-YC-A9TC | TCGA-BLCA | 66.146229 | 29.7272475 |
| TCGA-YD-A89C | TCGA-SKCM | 47.4162262 | 22.9052684 |
| TCGA-YD-A9TA | TCGA-SKCM | 104.876752 | 17.5563791 |
| TCGA-YD-A9TB | TCGA-SKCM | 54.6687582 | 35.2486506 |
| TCGA-YF-AA3L | TCGA-BLCA | 98.2586828 | 15.4809868 |
| TCGA-YF-AA3M | TCGA-BLCA | 114.491709 | 35.8550876 |
| TCGA-YG-AA3N | TCGA-SKCM | 34.4809149 | 11.577466 |
| TCGA-YG-AA3O | TCGA-SKCM | 50.6424678 | 33.8478264 |
| TCGA-YG-AA3P | TCGA-SKCM | 54.7751749 | 46.171898 |
| TCGA-YH-A8SY | TCGA-PAAD | 56.5076915 | 36.4802981 |
| TCGA-YL-A8HJ | TCGA-PRAD | 32.8531112 | 13.2181564 |
| TCGA-YL-A8HK | TCGA-PRAD | 52.524829 | 18.8005645 |
| TCGA-YL-A8HL | TCGA-PRAD | 58.5794499 | 12.5374389 |
| TCGA-YL-A8HM | TCGA-PRAD | 48.4222993 | 39.9638421 |
| TCGA-YL-A8HO | TCGA-PRAD | 33.5164031 | 16.4037489 |
| TCGA-YL-A8S8 | TCGA-PRAD | 34.7042655 | 16.8930859 |
| TCGA-YL-A8S9 | TCGA-PRAD | 33.1344927 | 21.1690929 |
| TCGA-YL-A8SA | TCGA-PRAD | 49.254395 | 18.7282572 |
| TCGA-YL-A8SB | TCGA-PRAD | 34.355305 | 16.2887221 |
| TCGA-YL-A8SC | TCGA-PRAD | 46.1489797 | 12.8376501 |
| TCGA-YL-A8SH | TCGA-PRAD | 33.6974016 | 14.3157467 |
| TCGA-YL-A8SI | TCGA-PRAD | 35.9807151 | 15.9790955 |
| TCGA-YL-A8SJ | TCGA-PRAD | 36.6737377 | 15.6612104 |
| TCGA-YL-A8SK | TCGA-PRAD | 54.3120294 | 16.8642977 |
| TCGA-YL-A8SL | TCGA-PRAD | 65.2589193 | 32.5116984 |
| TCGA-YL-A8SO | TCGA-PRAD | 36.218081 | 15.900374 |
| TCGA-YL-A8SP | TCGA-PRAD | 46.6373311 | 14.0238974 |
| TCGA-YL-A8SQ | TCGA-PRAD | 28.3324975 | 20.2631564 |

| | | | |
|---|---|---|---|
| TCGA-YL-A8SR | TCGA-PRAD | 54.479591 | 19.6145435 |
| TCGA-YL-A9WH | TCGA-PRAD | 40.0274243 | 36.7307573 |
| TCGA-YL-A9WI | TCGA-PRAD | 37.2625206 | 17.039115 |
| TCGA-YL-A9WJ | TCGA-PRAD | 52.0492323 | 13.7798583 |
| TCGA-YL-A9WK | TCGA-PRAD | 40.0083301 | 11.4043166 |
| TCGA-YL-A9WL | TCGA-PRAD | 59.3935094 | 21.4219063 |
| TCGA-YL-A9WX | TCGA-PRAD | 56.7179113 | 18.4310126 |
| TCGA-YL-A9WY | TCGA-PRAD | 53.4317028 | 16.0174835 |
| TCGA-YU-A90P | TCGA-TGCT | 136.687007 | 254.028954 |
| TCGA-YU-A90Q | TCGA-TGCT | 29.0245723 | 42.6204763 |
| TCGA-YU-A90S | TCGA-TGCT | 7.05082967 | 4.2415591 |
| TCGA-YU-A90W | TCGA-TGCT | 38.226016 | 110.1775 |
| TCGA-YU-A90Y | TCGA-TGCT | 78.8091117 | 39.2986095 |
| TCGA-YU-A912 | TCGA-TGCT | 40.9861109 | 181.716496 |
| TCGA-YU-A94D | TCGA-TGCT | 53.1160114 | 15.1759562 |
| TCGA-YU-A94I | TCGA-TGCT | 66.3381143 | 94.6412293 |
| TCGA-YU-AA4L | TCGA-TGCT | 110.155041 | 33.2260304 |
| TCGA-YU-AA61 | TCGA-TGCT | 85.26456 | 67.5805125 |
| TCGA-YY-A8LH | TCGA-PAAD | 48.5761837 | 20.5336362 |
| TCGA-Z2-A8RT | TCGA-SKCM | 32.7218988 | 33.4618107 |
| TCGA-Z2-AA3S | TCGA-SKCM | 65.7267962 | 29.509627 |
| TCGA-Z2-AA3V | TCGA-SKCM | 28.5772757 | 7.73233861 |
| TCGA-Z5-AAPL | TCGA-PAAD | 60.0242867 | 19.5084431 |
| TCGA-Z6-A8JD | TCGA-ESCA | 50.9996007 | 24.1009338 |
| TCGA-Z6-A8JE | TCGA-ESCA | 60.1314291 | 22.7225713 |
| TCGA-Z6-A9VB | TCGA-ESCA | 38.4243242 | 38.9515026 |
| TCGA-Z6-AAPN | TCGA-ESCA | 68.5955913 | 48.447436 |
| TCGA-Z7-A8R5 | TCGA-BRCA | 52.3275269 | 26.9434117 |
| TCGA-Z7-A8R6 | TCGA-BRCA | 79.8889966 | 39.829607 |
| TCGA-ZA-A8F6 | TCGA-STAD | 30.2822897 | 14.396507 |
| TCGA-ZF-A9R0 | TCGA-BLCA | 54.6409956 | 6.67935846 |

| | | | |
|---|---|---|---|
| TCGA-ZF-A9R1 | TCGA-BLCA | 81.4181766 | 28.8914772 |
| TCGA-ZF-A9R2 | TCGA-BLCA | 86.2146715 | 33.6021287 |
| TCGA-ZF-A9R3 | TCGA-BLCA | 38.1321153 | 23.1281024 |
| TCGA-ZF-A9R4 | TCGA-BLCA | 161.406442 | 28.6664694 |
| TCGA-ZF-A9R5 | TCGA-BLCA | 71.4420495 | 21.1850687 |
| TCGA-ZF-A9R7 | TCGA-BLCA | 58.5225981 | 22.922487 |
| TCGA-ZF-A9R9 | TCGA-BLCA | 50.9526743 | 14.3506841 |
| TCGA-ZF-A9RC | TCGA-BLCA | 95.7713264 | 30.1701893 |
| TCGA-ZF-A9RD | TCGA-BLCA | 100.251918 | 47.7508083 |
| TCGA-ZF-A9RE | TCGA-BLCA | 70.012672 | 34.9645664 |
| TCGA-ZF-A9RF | TCGA-BLCA | 121.56807 | 49.9822072 |
| TCGA-ZF-A9RL | TCGA-BLCA | 81.3021371 | 33.7555291 |
| TCGA-ZF-A9RM | TCGA-BLCA | 93.0762873 | 36.6617876 |
| TCGA-ZF-A9RN | TCGA-BLCA | 82.1703118 | 44.508129 |
| TCGA-ZF-AA4N | TCGA-BLCA | 80.9205757 | 68.1266675 |
| TCGA-ZF-AA4R | TCGA-BLCA | 95.9714768 | 32.6321909 |
| TCGA-ZF-AA4T | TCGA-BLCA | 140.803505 | 42.6764163 |
| TCGA-ZF-AA4U | TCGA-BLCA | 111.618148 | 10.3755427 |
| TCGA-ZF-AA4V | TCGA-BLCA | 104.594219 | 82.2880072 |
| TCGA-ZF-AA4W | TCGA-BLCA | 94.3684653 | 15.2030056 |
| TCGA-ZF-AA4X | TCGA-BLCA | 76.5515604 | 30.8221011 |
| TCGA-ZF-AA51 | TCGA-BLCA | 93.6801158 | 23.5306941 |
| TCGA-ZF-AA52 | TCGA-BLCA | 72.1606809 | 17.4486556 |
| TCGA-ZF-AA53 | TCGA-BLCA | 65.1845014 | 23.8794439 |
| TCGA-ZF-AA54 | TCGA-BLCA | 142.811059 | 76.8614312 |
| TCGA-ZF-AA56 | TCGA-BLCA | 56.3502661 | 27.1172096 |
| TCGA-ZF-AA58 | TCGA-BLCA | 165.20593 | 43.4212666 |
| TCGA-ZF-AA5H | TCGA-BLCA | 91.903562 | 49.5604015 |
| TCGA-ZF-AA5N | TCGA-BLCA | 48.9539581 | 28.7495818 |
| TCGA-ZF-AA5P | TCGA-BLCA | 38.4012104 | 18.6746601 |
| TCGA-ZG-A8QW | TCGA-PRAD | 50.5748804 | 23.2866611 |

| | | | |
|---|---|---|---|
| TCGA-ZG-A8QX | TCGA-PRAD | 38.2570332 | 16.0695502 |
| TCGA-ZG-A8QY | TCGA-PRAD | 29.2866124 | 15.8290451 |
| TCGA-ZG-A8QZ | TCGA-PRAD | 53.2455888 | 25.3483939 |
| TCGA-ZG-A9KY | TCGA-PRAD | 40.5273381 | 29.7042859 |
| TCGA-ZG-A9L0 | TCGA-PRAD | 48.3986445 | 15.4676016 |
| TCGA-ZG-A9L1 | TCGA-PRAD | 52.8784255 | 20.7132352 |
| TCGA-ZG-A9L2 | TCGA-PRAD | 51.9658548 | 18.2139911 |
| TCGA-ZG-A9L4 | TCGA-PRAD | 57.6346325 | 20.7497266 |
| TCGA-ZG-A9L5 | TCGA-PRAD | 43.7102351 | 20.5707432 |
| TCGA-ZG-A9L6 | TCGA-PRAD | 44.1801153 | 15.0689596 |
| TCGA-ZG-A9L9 | TCGA-PRAD | 48.0613097 | 32.5235987 |
| TCGA-ZG-A9LB | TCGA-PRAD | 42.9438284 | 13.914112 |
| TCGA-ZG-A9LM | TCGA-PRAD | 54.1639711 | 20.0628093 |
| TCGA-ZG-A9LN | TCGA-PRAD | 38.5962102 | 15.4489837 |
| TCGA-ZG-A9LS | TCGA-PRAD | 53.6576946 | 17.2478999 |
| TCGA-ZG-A9LU | TCGA-PRAD | 57.4677534 | 14.2418754 |
| TCGA-ZG-A9LY | TCGA-PRAD | 50.6441414 | 19.4625057 |
| TCGA-ZG-A9LZ | TCGA-PRAD | 45.1901412 | 22.7184735 |
| TCGA-ZG-A9M4 | TCGA-PRAD | 34.7862627 | 15.3783304 |
| TCGA-ZG-A9MC | TCGA-PRAD | 58.7295813 | 14.1886778 |
| TCGA-ZG-A9N3 | TCGA-PRAD | 57.6540273 | 24.5392136 |
| TCGA-ZG-A9ND | TCGA-PRAD | 45.0960099 | 23.522613 |
| TCGA-ZG-A9NI | TCGA-PRAD | 34.0689636 | 17.9073787 |
| TCGA-ZM-AA05 | TCGA-TGCT | 1.56263509 | 6.84862297 |
| TCGA-ZM-AA06 | TCGA-TGCT | 30.6017642 | 38.9361991 |
| TCGA-ZM-AA0B | TCGA-TGCT | 83.6338365 | 74.8767854 |
| TCGA-ZM-AA0D | TCGA-TGCT | 65.701963 | 44.4400596 |
| TCGA-ZM-AA0E | TCGA-TGCT | 21.925757 | 33.0866158 |
| TCGA-ZM-AA0F | TCGA-TGCT | 34.7995067 | 46.1890375 |
| TCGA-ZM-AA0H | TCGA-TGCT | 30.6453766 | 31.8304372 |
| TCGA-ZM-AA0N | TCGA-TGCT | 51.9104622 | 49.3969332 |

| | | | |
|---|---|---|---|
| TCGA-ZP-A9CV | TCGA-LIHC | 36.7691323 | 22.0277244 |
| TCGA-ZP-A9CY | TCGA-LIHC | 30.425466 | 17.4437887 |
| TCGA-ZP-A9CZ | TCGA-LIHC | 43.0565819 | 21.8170299 |
| TCGA-ZP-A9D0 | TCGA-LIHC | 18.5735553 | 11.6146904 |
| TCGA-ZP-A9D1 | TCGA-LIHC | 41.2935907 | 24.895892 |
| TCGA-ZP-A9D2 | TCGA-LIHC | 61.1556487 | 43.6872637 |
| TCGA-ZP-A9D4 | TCGA-LIHC | 46.9511522 | 12.4323221 |
| TCGA-ZQ-A9CR | TCGA-STAD | 30.2611468 | 28.3704173 |
| TCGA-ZR-A9CJ | TCGA-ESCA | 46.2682273 | 29.8530851 |
| TCGA-ZS-A9CD | TCGA-LIHC | 36.5172174 | 11.5856447 |
| TCGA-ZS-A9CE | TCGA-LIHC | 43.2880936 | 28.0425032 |
| TCGA-ZS-A9CF | TCGA-LIHC | 45.3328045 | 10.1772772 |
| TCGA-ZS-A9CG | TCGA-LIHC | 42.3552367 | 17.3303194 |

| Table S3. MAGOH and MAOGHB expression in gliomas (grades 2 to 4). | | | | | | | | |
|---|---|---|---|---|---|---|---|---|
| **sample_id** | **tcga_correspondence** | **from** | **cancer_type** | **grade** | **tissue** | **MAGOHB_tpm** | **MAGOH_tpm** | **MAGOH+MAGOHB_tpm** |
| TCGA-02-0047 | TCGA-GBM | Tumor TCGA | GBM | GBM | Brain (Cortex) Tumor | 13.6963106 | 38.164676 | 51.8609866 |
| TCGA-02-0055 | TCGA-GBM | Tumor TCGA | GBM | GBM | Brain (Cortex) Tumor | 27.5218395 | 85.0317744 | 112.553614 |
| TCGA-02-2483 | TCGA-GBM | Tumor TCGA | GBM | GBM | Brain (Cortex) Tumor | 38.7765385 | 95.283095 | 134.059634 |
| TCGA-02-2485 | TCGA-GBM | Tumor TCGA | GBM | GBM | Brain (Cortex) Tumor | 32.7891766 | 65.4858311 | 98.2750077 |
| TCGA-02-2486 | TCGA-GBM | Tumor TCGA | GBM | GBM | Brain (Cortex) Tumor | 25.0475672 | 70.6058681 | 95.6534353 |
| TCGA-06-0125 | TCGA-GBM | Tumor TCGA | GBM | GBM | Brain (Cortex) Tumor | 21.0828713 | 53.2694908 | 74.3523622 |
| TCGA-06-0129 | TCGA-GBM | Tumor TCGA | GBM | GBM | Brain (Cortex) Tumor | 15.266424 | 60.2484084 | 75.5148324 |
| TCGA-06-0130 | TCGA-GBM | Tumor TCGA | GBM | GBM | Brain (Cortex) Tumor | 45.6958519 | 91.1860283 | 136.88188 |
| TCGA-06-0132 | TCGA-GBM | Tumor TCGA | GBM | GBM | Brain (Cortex) Tumor | 15.8339589 | 32.4920213 | 48.3259802 |
| TCGA-06-0138 | TCGA-GBM | Tumor TCGA | GBM | GBM | Brain (Cortex) Tumor | 29.1278319 | 40.1883036 | 69.3161354 |
| TCGA-06-0139 | TCGA-GBM | Tumor TCGA | GBM | GBM | Brain (Cortex) Tumor | 18.5964452 | 45.9245285 | 64.5209737 |
| TCGA-06-0141 | TCGA-GBM | Tumor TCGA | GBM | GBM | Brain (Cortex) Tumor | 14.3798288 | 51.8432049 | 66.2230336 |
| TCGA-06-0152 | TCGA-GBM | Tumor TCGA | GBM | GBM | Brain (Cortex) Tumor | 27.613946 | 56.724238 | 84.3381841 |
| TCGA-06-0156 | TCGA-GBM | Tumor TCGA | GBM | GBM | Brain (Cortex) Tumor | 30.9807802 | 53.5313797 | 84.5121598 |
| TCGA-06-0157 | TCGA-GBM | Tumor TCGA | GBM | GBM | Brain (Cortex) Tumor | 25.8417248 | 46.6540555 | 72.4957803 |
| TCGA-06-0158 | TCGA-GBM | Tumor TCGA | GBM | GBM | Brain (Cortex) Tumor | 18.8541599 | 42.695559 | 61.5497188 |
| TCGA-06-0168 | TCGA-GBM | Tumor TCGA | GBM | GBM | Brain (Cortex) Tumor | 24.1376048 | 67.6937047 | 91.8313095 |
| TCGA-06-0171 | TCGA-GBM | Tumor TCGA | GBM | GBM | Brain (Cortex) Tumor | 13.8291492 | 36.0278793 | 49.8570285 |
| TCGA-06-0174 | TCGA-GBM | Tumor TCGA | GBM | GBM | Brain (Cortex) Tumor | 28.8940278 | 45.2782073 | 74.1722351 |
| TCGA-06-0178 | TCGA-GBM | Tumor TCGA | GBM | GBM | Brain (Cortex) Tumor | 18.4201204 | 52.0706203 | 70.4907407 |
| TCGA-06-0184 | TCGA-GBM | Tumor TCGA | GBM | GBM | Brain (Cortex) Tumor | 26.1188863 | 60.1623758 | 86.2812621 |
| TCGA-06-0187 | TCGA-GBM | Tumor TCGA | GBM | GBM | Brain (Cortex) Tumor | 28.5453435 | 69.7697865 | 98.3151299 |
| TCGA-06-0190 | TCGA-GBM | Tumor TCGA | GBM | GBM | Brain (Cortex) Tumor | 25.0811512 | 68.4952628 | 93.5764139 |
| TCGA-06-0210 | TCGA-GBM | Tumor TCGA | GBM | GBM | Brain (Cortex) Tumor | 14.9165686 | 46.5464973 | 61.4630659 |
| TCGA-06-0211 | TCGA-GBM | Tumor TCGA | GBM | GBM | Brain (Cortex) Tumor | 37.4958421 | 54.0845771 | 91.5804191 |
| TCGA-06-0219 | TCGA-GBM | Tumor TCGA | GBM | GBM | Brain (Cortex) Tumor | 37.91726 | 57.001031 | 94.9182911 |
| TCGA-06-0221 | TCGA-GBM | Tumor TCGA | GBM | GBM | Brain (Cortex) Tumor | 11.6339325 | 59.66346 | 71.2973925 |
| TCGA-06-0238 | TCGA-GBM | Tumor TCGA | GBM | GBM | Brain (Cortex) Tumor | 39.4154313 | 53.1240651 | 92.5394964 |
| TCGA-06-0644 | TCGA-GBM | Tumor TCGA | GBM | GBM | Brain (Cortex) Tumor | 43.9654821 | 78.1668688 | 122.132351 |
| TCGA-06-0645 | TCGA-GBM | Tumor TCGA | GBM | GBM | Brain (Cortex) Tumor | 21.1852275 | 48.9268766 | 70.1121041 |
| TCGA-06-0646 | TCGA-GBM | Tumor TCGA | GBM | GBM | Brain (Cortex) Tumor | 28.2835861 | 46.884887 | 75.1684731 |
| TCGA-06-0649 | TCGA-GBM | Tumor TCGA | GBM | GBM | Brain (Cortex) Tumor | 14.9086867 | 48.3241593 | 63.232846 |
| TCGA-06-0686 | TCGA-GBM | Tumor TCGA | GBM | GBM | Brain (Cortex) Tumor | 28.0329044 | 50.0832039 | 78.1161083 |
| TCGA-06-0743 | TCGA-GBM | Tumor TCGA | GBM | GBM | Brain (Cortex) Tumor | 19.2970135 | 46.5326816 | 65.8296951 |
| TCGA-06-0744 | TCGA-GBM | Tumor TCGA | GBM | GBM | Brain (Cortex) Tumor | 22.3272493 | 58.8120013 | 81.1392506 |
| TCGA-06-0745 | TCGA-GBM | Tumor TCGA | GBM | GBM | Brain (Cortex) Tumor | 31.9047443 | 62.6999439 | 94.6046881 |

| | | | | | | | | |
|---|---|---|---|---|---|---|---|---|
| TCGA-06-0747 | TCGA-GBM | Tumor TCGA | GBM | GBM | Brain (Cortex) Tumor | 23.8135004 | 63.2713533 | 87.0848536 |
| TCGA-06-0749 | TCGA-GBM | Tumor TCGA | GBM | GBM | Brain (Cortex) Tumor | 14.0575672 | 41.6658566 | 55.7234238 |
| TCGA-06-0750 | TCGA-GBM | Tumor TCGA | GBM | GBM | Brain (Cortex) Tumor | 20.2211493 | 55.328238 | 75.5493873 |
| TCGA-06-0878 | TCGA-GBM | Tumor TCGA | GBM | GBM | Brain (Cortex) Tumor | 29.3353905 | 68.3631983 | 97.6985888 |
| TCGA-06-0882 | TCGA-GBM | Tumor TCGA | GBM | GBM | Brain (Cortex) Tumor | 15.0482666 | 40.9874896 | 56.0357562 |
| TCGA-06-1804 | TCGA-GBM | Tumor TCGA | GBM | GBM | Brain (Cortex) Tumor | 18.8707102 | 52.9457874 | 71.8164975 |
| TCGA-06-2557 | TCGA-GBM | Tumor TCGA | GBM | GBM | Brain (Cortex) Tumor | 23.3529138 | 64.8242667 | 88.1771805 |
| TCGA-06-2558 | TCGA-GBM | Tumor TCGA | GBM | GBM | Brain (Cortex) Tumor | 35.1574292 | 60.7034649 | 95.8608941 |
| TCGA-06-2559 | TCGA-GBM | Tumor TCGA | GBM | GBM | Brain (Cortex) Tumor | 39.8193876 | 64.947856 | 104.767244 |
| TCGA-06-2561 | TCGA-GBM | Tumor TCGA | GBM | GBM | Brain (Cortex) Tumor | 23.0059319 | 51.7995632 | 74.8054951 |
| TCGA-06-2562 | TCGA-GBM | Tumor TCGA | GBM | GBM | Brain (Cortex) Tumor | 32.1517825 | 74.0437467 | 106.195529 |
| TCGA-06-2563 | TCGA-GBM | Tumor TCGA | GBM | GBM | Brain (Cortex) Tumor | 31.4658185 | 61.281203 | 92.7470215 |
| TCGA-06-2564 | TCGA-GBM | Tumor TCGA | GBM | GBM | Brain (Cortex) Tumor | 28.5827286 | 50.3183519 | 78.9010806 |
| TCGA-06-2565 | TCGA-GBM | Tumor TCGA | GBM | GBM | Brain (Cortex) Tumor | 24.7216751 | 63.997846 | 88.7195212 |
| TCGA-06-2567 | TCGA-GBM | Tumor TCGA | GBM | GBM | Brain (Cortex) Tumor | 35.2215478 | 67.7719731 | 102.993521 |
| TCGA-06-2569 | TCGA-GBM | Tumor TCGA | GBM | GBM | Brain (Cortex) Tumor | 31.5965976 | 104.990983 | 136.58758 |
| TCGA-06-2570 | TCGA-GBM | Tumor TCGA | GBM | GBM | Brain (Cortex) Tumor | 12.5214438 | 45.8684398 | 58.3898836 |
| TCGA-06-5408 | TCGA-GBM | Tumor TCGA | GBM | GBM | Brain (Cortex) Tumor | 42.5944325 | 99.6154389 | 142.209871 |
| TCGA-06-5410 | TCGA-GBM | Tumor TCGA | GBM | GBM | Brain (Cortex) Tumor | 19.7263275 | 52.8232832 | 72.5496107 |
| TCGA-06-5411 | TCGA-GBM | Tumor TCGA | GBM | GBM | Brain (Cortex) Tumor | 31.5348416 | 50.5486184 | 82.08346 |
| TCGA-06-5412 | TCGA-GBM | Tumor TCGA | GBM | GBM | Brain (Cortex) Tumor | 32.8526475 | 64.7542591 | 97.6069066 |
| TCGA-06-5413 | TCGA-GBM | Tumor TCGA | GBM | GBM | Brain (Cortex) Tumor | 46.7573893 | 89.1175619 | 135.874951 |
| TCGA-06-5414 | TCGA-GBM | Tumor TCGA | GBM | GBM | Brain (Cortex) Tumor | 27.9083257 | 58.8610421 | 86.7693678 |
| TCGA-06-5416 | TCGA-GBM | Tumor TCGA | GBM | GBM | Brain (Cortex) Tumor | 22.8688862 | 61.389451 | 84.2583372 |
| TCGA-06-5417 | TCGA-GBM | Tumor TCGA | GBM | GBM | Brain (Cortex) Tumor | 14.6673709 | 66.8142471 | 81.481618 |
| TCGA-06-5418 | TCGA-GBM | Tumor TCGA | GBM | GBM | Brain (Cortex) Tumor | 23.8448434 | 58.3025381 | 82.1473815 |
| TCGA-06-5856 | TCGA-GBM | Tumor TCGA | GBM | GBM | Brain (Cortex) Tumor | 56.920874 | 98.040666 | 154.96154 |
| TCGA-06-5858 | TCGA-GBM | Tumor TCGA | GBM | GBM | Brain (Cortex) Tumor | 27.4910266 | 58.6861936 | 86.1772202 |
| TCGA-06-5859 | TCGA-GBM | Tumor TCGA | GBM | GBM | Brain (Cortex) Tumor | 26.1927571 | 69.0273296 | 95.2200867 |
| TCGA-08-0386 | TCGA-GBM | Tumor TCGA | GBM | GBM | Brain (Cortex) Tumor | 17.3527995 | 75.5535637 | 92.9063633 |
| TCGA-12-0616 | TCGA-GBM | Tumor TCGA | GBM | GBM | Brain (Cortex) Tumor | 35.4081573 | 49.2203224 | 84.6284797 |
| TCGA-12-0618 | TCGA-GBM | Tumor TCGA | GBM | GBM | Brain (Cortex) Tumor | 17.1830664 | 42.8697388 | 60.0528051 |
| TCGA-12-0619 | TCGA-GBM | Tumor TCGA | GBM | GBM | Brain (Cortex) Tumor | 15.6560923 | 71.3736682 | 87.0297605 |
| TCGA-12-0821 | TCGA-GBM | Tumor TCGA | GBM | GBM | Brain (Cortex) Tumor | 37.1754475 | 77.2098517 | 114.385299 |
| TCGA-12-1597 | TCGA-GBM | Tumor TCGA | GBM | GBM | Brain (Cortex) Tumor | 15.8086608 | 28.3498072 | 44.158468 |
| TCGA-12-3650 | TCGA-GBM | Tumor TCGA | GBM | GBM | Brain (Cortex) Tumor | 34.8907948 | 38.5346526 | 73.4254474 |
| TCGA-12-3652 | TCGA-GBM | Tumor TCGA | GBM | GBM | Brain (Cortex) Tumor | 32.6790056 | 46.8211666 | 79.5001723 |
| TCGA-12-3653 | TCGA-GBM | Tumor TCGA | GBM | GBM | Brain (Cortex) Tumor | 41.2080391 | 68.4439196 | 109.651959 |

| | | | | | | | | |
|---|---|---|---|---|---|---|---|---|
| TCGA-12-5295 | TCGA-GBM | Tumor TCGA | GBM | GBM | Brain (Cortex) Tumor | 35.3252356 | 62.4605157 | 97.7857512 |
| TCGA-12-5299 | TCGA-GBM | Tumor TCGA | GBM | GBM | Brain (Cortex) Tumor | 28.3887652 | 63.5955579 | 91.9843231 |
| TCGA-14-0736 | TCGA-GBM | Tumor TCGA | GBM | GBM | Brain (Cortex) Tumor | 25.7044471 | 42.8473695 | 68.5518166 |
| TCGA-14-0781 | TCGA-GBM | Tumor TCGA | GBM | GBM | Brain (Cortex) Tumor | 18.088428 | 37.660733 | 55.7491611 |
| TCGA-14-0787 | TCGA-GBM | Tumor TCGA | GBM | GBM | Brain (Cortex) Tumor | 39.0070485 | 81.7128399 | 120.719888 |
| TCGA-14-0789 | TCGA-GBM | Tumor TCGA | GBM | GBM | Brain (Cortex) Tumor | 11.4573231 | 42.1229315 | 53.5802546 |
| TCGA-14-0790 | TCGA-GBM | Tumor TCGA | GBM | GBM | Brain (Cortex) Tumor | 42.8295838 | 65.5844327 | 108.414016 |
| TCGA-14-0817 | TCGA-GBM | Tumor TCGA | GBM | GBM | Brain (Cortex) Tumor | 25.2874574 | 85.9827204 | 111.270178 |
| TCGA-14-0871 | TCGA-GBM | Tumor TCGA | GBM | GBM | Brain (Cortex) Tumor | 20.9334582 | 121.13329 | 142.066748 |
| TCGA-14-1034 | TCGA-GBM | Tumor TCGA | GBM | GBM | Brain (Cortex) Tumor | 53.6319451 | 127.877833 | 181.509778 |
| TCGA-14-1402 | TCGA-GBM | Tumor TCGA | GBM | GBM | Brain (Cortex) Tumor | 33.2564266 | 56.5976521 | 89.8540787 |
| TCGA-14-1823 | TCGA-GBM | Tumor TCGA | GBM | GBM | Brain (Cortex) Tumor | 57.9685554 | 63.8286633 | 121.797219 |
| TCGA-14-1825 | TCGA-GBM | Tumor TCGA | GBM | GBM | Brain (Cortex) Tumor | 43.5735144 | 59.9553594 | 103.528874 |
| TCGA-14-1829 | TCGA-GBM | Tumor TCGA | GBM | GBM | Brain (Cortex) Tumor | 43.8666968 | 47.9334034 | 91.8001002 |
| TCGA-14-2554 | TCGA-GBM | Tumor TCGA | GBM | GBM | Brain (Cortex) Tumor | 30.9660197 | 61.2894065 | 92.2554262 |
| TCGA-15-0742 | TCGA-GBM | Tumor TCGA | GBM | GBM | Brain (Cortex) Tumor | 29.0554546 | 64.2954104 | 93.350865 |
| TCGA-15-1444 | TCGA-GBM | Tumor TCGA | GBM | GBM | Brain (Cortex) Tumor | 17.078489 | 41.0221327 | 58.1006217 |
| TCGA-16-0846 | TCGA-GBM | Tumor TCGA | GBM | GBM | Brain (Cortex) Tumor | 18.0174279 | 43.7499135 | 61.7673414 |
| TCGA-16-1045 | TCGA-GBM | Tumor TCGA | GBM | GBM | Brain (Cortex) Tumor | 19.951586 | 45.0877319 | 65.0393179 |
| TCGA-19-0957 | TCGA-GBM | Tumor TCGA | GBM | GBM | Brain (Cortex) Tumor | 17.3123742 | 32.7149942 | 50.0273684 |
| TCGA-19-1389 | TCGA-GBM | Tumor TCGA | GBM | GBM | Brain (Cortex) Tumor | 26.4008135 | 62.1669042 | 88.5677176 |
| TCGA-19-1390 | TCGA-GBM | Tumor TCGA | GBM | GBM | Brain (Cortex) Tumor | 25.650955 | 25.22206 | 50.873015 |
| TCGA-19-1787 | TCGA-GBM | Tumor TCGA | GBM | GBM | Brain (Cortex) Tumor | 38.171558 | 80.4077945 | 118.579352 |
| TCGA-19-2619 | TCGA-GBM | Tumor TCGA | GBM | GBM | Brain (Cortex) Tumor | 24.8710914 | 48.9038435 | 73.7749348 |
| TCGA-19-2620 | TCGA-GBM | Tumor TCGA | GBM | GBM | Brain (Cortex) Tumor | 16.0844768 | 45.3727042 | 61.457181 |
| TCGA-19-2624 | TCGA-GBM | Tumor TCGA | GBM | GBM | Brain (Cortex) Tumor | 30.0841802 | 55.912457 | 85.9966372 |
| TCGA-19-2625 | TCGA-GBM | Tumor TCGA | GBM | GBM | Brain (Cortex) Tumor | 19.8231392 | 75.2406081 | 95.0637474 |
| TCGA-19-2629 | TCGA-GBM | Tumor TCGA | GBM | GBM | Brain (Cortex) Tumor | 23.4888838 | 63.7929138 | 87.2817975 |
| TCGA-19-4065 | TCGA-GBM | Tumor TCGA | GBM | GBM | Brain (Cortex) Tumor | 27.261933 | 62.1065273 | 89.3684603 |
| TCGA-19-5960 | TCGA-GBM | Tumor TCGA | GBM | GBM | Brain (Cortex) Tumor | 20.7321267 | 78.4333842 | 99.165511 |
| TCGA-26-1442 | TCGA-GBM | Tumor TCGA | GBM | GBM | Brain (Cortex) Tumor | 25.7857277 | 30.767252 | 56.5529798 |
| TCGA-26-5132 | TCGA-GBM | Tumor TCGA | GBM | GBM | Brain (Cortex) Tumor | 15.5999388 | 76.8350962 | 92.435035 |
| TCGA-26-5133 | TCGA-GBM | Tumor TCGA | GBM | GBM | Brain (Cortex) Tumor | 22.213922 | 77.4979313 | 99.7118533 |
| TCGA-26-5134 | TCGA-GBM | Tumor TCGA | GBM | GBM | Brain (Cortex) Tumor | 8.44316038 | 11.1291906 | 19.572351 |
| TCGA-26-5135 | TCGA-GBM | Tumor TCGA | GBM | GBM | Brain (Cortex) Tumor | 21.5527105 | 44.7868813 | 66.3395918 |
| TCGA-26-5136 | TCGA-GBM | Tumor TCGA | GBM | GBM | Brain (Cortex) Tumor | 24.5319163 | 100.039219 | 124.571135 |
| TCGA-26-5139 | TCGA-GBM | Tumor TCGA | GBM | GBM | Brain (Cortex) Tumor | 23.2389394 | 63.3201439 | 86.5590833 |
| TCGA-27-1830 | TCGA-GBM | Tumor TCGA | GBM | GBM | Brain (Cortex) Tumor | 24.0174014 | 50.8164894 | 74.8338908 |

| | | | | | | | | |
|---|---|---|---|---|---|---|---|---|
| TCGA-27-1831 | TCGA-GBM | Tumor TCGA | GBM | GBM | Brain (Cortex) Tumor | 43.1833906 | 51.2862714 | 94.469662 |
| TCGA-27-1832 | TCGA-GBM | Tumor TCGA | GBM | GBM | Brain (Cortex) Tumor | 36.739781 | 67.5135126 | 104.253294 |
| TCGA-27-1834 | TCGA-GBM | Tumor TCGA | GBM | GBM | Brain (Cortex) Tumor | 19.3744202 | 51.5389148 | 70.913335 |
| TCGA-27-1835 | TCGA-GBM | Tumor TCGA | GBM | GBM | Brain (Cortex) Tumor | 27.091633 | 64.3715204 | 91.4631534 |
| TCGA-27-1837 | TCGA-GBM | Tumor TCGA | GBM | GBM | Brain (Cortex) Tumor | 25.1346762 | 43.0259043 | 68.1605804 |
| TCGA-27-2519 | TCGA-GBM | Tumor TCGA | GBM | GBM | Brain (Cortex) Tumor | 35.8875647 | 73.0678526 | 108.955417 |
| TCGA-27-2521 | TCGA-GBM | Tumor TCGA | GBM | GBM | Brain (Cortex) Tumor | 20.2612195 | 72.8044107 | 93.0656302 |
| TCGA-27-2523 | TCGA-GBM | Tumor TCGA | GBM | GBM | Brain (Cortex) Tumor | 44.707098 | 72.7437879 | 117.450886 |
| TCGA-27-2524 | TCGA-GBM | Tumor TCGA | GBM | GBM | Brain (Cortex) Tumor | 34.0004831 | 75.646846 | 109.647329 |
| TCGA-27-2526 | TCGA-GBM | Tumor TCGA | GBM | GBM | Brain (Cortex) Tumor | 38.3467202 | 56.6054906 | 94.9522108 |
| TCGA-27-2528 | TCGA-GBM | Tumor TCGA | GBM | GBM | Brain (Cortex) Tumor | 26.3197689 | 52.3420973 | 78.6618662 |
| TCGA-28-1747 | TCGA-GBM | Tumor TCGA | GBM | GBM | Brain (Cortex) Tumor | 26.5697951 | 59.3330031 | 85.9027982 |
| TCGA-28-1753 | TCGA-GBM | Tumor TCGA | GBM | GBM | Brain (Cortex) Tumor | 26.2639571 | 49.6500946 | 75.9140517 |
| TCGA-28-2499 | TCGA-GBM | Tumor TCGA | GBM | GBM | Brain (Cortex) Tumor | 20.3516272 | 67.7979756 | 88.1496029 |
| TCGA-28-2509 | TCGA-GBM | Tumor TCGA | GBM | GBM | Brain (Cortex) Tumor | 24.7576966 | 86.6016559 | 111.359353 |
| TCGA-28-2510 | TCGA-GBM | Tumor TCGA | GBM | GBM | Brain (Cortex) Tumor | 14.9753959 | 37.451419 | 52.4268149 |
| TCGA-28-2513 | TCGA-GBM | Tumor TCGA | GBM | GBM | Brain (Cortex) Tumor | 18.6040134 | 46.9360217 | 65.5400351 |
| TCGA-28-2514 | TCGA-GBM | Tumor TCGA | GBM | GBM | Brain (Cortex) Tumor | 12.0369717 | 46.6053748 | 58.6423465 |
| TCGA-28-5204 | TCGA-GBM | Tumor TCGA | GBM | GBM | Brain (Cortex) Tumor | 16.7484499 | 87.3933118 | 104.141762 |
| TCGA-28-5207 | TCGA-GBM | Tumor TCGA | GBM | GBM | Brain (Cortex) Tumor | 31.3423839 | 83.4782837 | 114.820668 |
| TCGA-28-5208 | TCGA-GBM | Tumor TCGA | GBM | GBM | Brain (Cortex) Tumor | 46.7552754 | 84.7626531 | 131.517928 |
| TCGA-28-5209 | TCGA-GBM | Tumor TCGA | GBM | GBM | Brain (Cortex) Tumor | 25.9216725 | 81.2298824 | 107.151555 |
| TCGA-28-5213 | TCGA-GBM | Tumor TCGA | GBM | GBM | Brain (Cortex) Tumor | 28.5942931 | 47.9389097 | 76.5332029 |
| TCGA-28-5215 | TCGA-GBM | Tumor TCGA | GBM | GBM | Brain (Cortex) Tumor | 30.1855144 | 54.4488808 | 84.6343952 |
| TCGA-28-5216 | TCGA-GBM | Tumor TCGA | GBM | GBM | Brain (Cortex) Tumor | 26.9711957 | 68.9102337 | 95.8814294 |
| TCGA-28-5218 | TCGA-GBM | Tumor TCGA | GBM | GBM | Brain (Cortex) Tumor | 48.3274241 | 59.4779003 | 107.805324 |
| TCGA-28-5220 | TCGA-GBM | Tumor TCGA | GBM | GBM | Brain (Cortex) Tumor | 35.985228 | 76.5058914 | 112.491119 |
| TCGA-32-1970 | TCGA-GBM | Tumor TCGA | GBM | GBM | Brain (Cortex) Tumor | 24.0601738 | 57.8057686 | 81.8659424 |
| TCGA-32-1980 | TCGA-GBM | Tumor TCGA | GBM | GBM | Brain (Cortex) Tumor | 20.3571691 | 46.7232883 | 67.0804574 |
| TCGA-32-1982 | TCGA-GBM | Tumor TCGA | GBM | GBM | Brain (Cortex) Tumor | 20.3005585 | 47.7859433 | 68.0865017 |
| TCGA-32-2615 | TCGA-GBM | Tumor TCGA | GBM | GBM | Brain (Cortex) Tumor | 20.0853909 | 54.32427 | 74.4096609 |
| TCGA-32-2616 | TCGA-GBM | Tumor TCGA | GBM | GBM | Brain (Cortex) Tumor | 22.1899594 | 50.6047721 | 72.7947315 |
| TCGA-32-2632 | TCGA-GBM | Tumor TCGA | GBM | GBM | Brain (Cortex) Tumor | 31.1375877 | 58.4288906 | 89.5664782 |
| TCGA-32-2634 | TCGA-GBM | Tumor TCGA | GBM | GBM | Brain (Cortex) Tumor | 42.4165921 | 76.0741312 | 118.490723 |
| TCGA-32-2638 | TCGA-GBM | Tumor TCGA | GBM | GBM | Brain (Cortex) Tumor | 21.6946721 | 52.6809512 | 74.3756233 |
| TCGA-32-4213 | TCGA-GBM | Tumor TCGA | GBM | GBM | Brain (Cortex) Tumor | 11.9954525 | 13.9509742 | 25.9464267 |
| TCGA-32-5222 | TCGA-GBM | Tumor TCGA | GBM | GBM | Brain (Cortex) Tumor | 40.8175545 | 81.4919257 | 122.30948 |
| TCGA-41-2571 | TCGA-GBM | Tumor TCGA | GBM | GBM | Brain (Cortex) Tumor | 18.4008837 | 37.5366635 | 55.9375472 |

| | | | | | | | | |
|---|---|---|---|---|---|---|---|---|
| TCGA-41-2572 | TCGA-GBM | Tumor TCGA | GBM | GBM | Brain (Cortex) Tumor | 21.4136483 | 37.2383593 | 58.6520076 |
| TCGA-41-3915 | TCGA-GBM | Tumor TCGA | GBM | GBM | Brain (Cortex) Tumor | 16.5118833 | 36.6685321 | 53.1804154 |
| TCGA-41-4097 | TCGA-GBM | Tumor TCGA | GBM | GBM | Brain (Cortex) Tumor | 14.3609181 | 40.7701634 | 55.1310815 |
| TCGA-41-5651 | TCGA-GBM | Tumor TCGA | GBM | GBM | Brain (Cortex) Tumor | 29.3274706 | 73.0511948 | 102.378665 |
| TCGA-76-4925 | TCGA-GBM | Tumor TCGA | GBM | GBM | Brain (Cortex) Tumor | 32.7779969 | 75.6680157 | 108.446013 |
| TCGA-76-4926 | TCGA-GBM | Tumor TCGA | GBM | GBM | Brain (Cortex) Tumor | 16.1960695 | 51.2066219 | 67.4026914 |
| TCGA-76-4927 | TCGA-GBM | Tumor TCGA | GBM | GBM | Brain (Cortex) Tumor | 26.8133122 | 54.504116 | 81.3174282 |
| TCGA-76-4928 | TCGA-GBM | Tumor TCGA | GBM | GBM | Brain (Cortex) Tumor | 30.6450145 | 61.2925942 | 91.9376088 |
| TCGA-76-4929 | TCGA-GBM | Tumor TCGA | GBM | GBM | Brain (Cortex) Tumor | 23.33443 | 64.2429187 | 87.5773488 |
| TCGA-76-4931 | TCGA-GBM | Tumor TCGA | GBM | GBM | Brain (Cortex) Tumor | 27.3569488 | 60.8130004 | 88.1699491 |
| TCGA-76-4932 | TCGA-GBM | Tumor TCGA | GBM | GBM | Brain (Cortex) Tumor | 24.5926609 | 66.9195858 | 91.5122467 |
| TCGA-CS-4938 | TCGA-LGG | Tumor TCGA | Astrocytoma | Grade II | Brain (Cortex) Tumor | 11.2290868 | 30.2472228 | 41.4763097 |
| TCGA-CS-4941 | TCGA-LGG | Tumor TCGA | Astrocytoma | Grade III | Brain (Cortex) Tumor | 20.8165127 | 54.2642617 | 75.0807744 |
| TCGA-CS-4942 | TCGA-LGG | Tumor TCGA | Astrocytoma | Grade III | Brain (Cortex) Tumor | 13.5263462 | 30.5601206 | 44.0864668 |
| TCGA-CS-4943 | TCGA-LGG | Tumor TCGA | Astrocytoma | Grade III | Brain (Cortex) Tumor | 76.9146813 | 41.7485738 | 118.663255 |
| TCGA-CS-4944 | TCGA-LGG | Tumor TCGA | Astrocytoma | Grade II | Brain (Cortex) Tumor | 9.27134793 | 27.3161602 | 36.5875081 |
| TCGA-CS-5390 | TCGA-LGG | Tumor TCGA | Oligodendroglion | Grade II | Brain (Cortex) Tumor | 17.5203901 | 17.0675127 | 34.5879029 |
| TCGA-CS-5393 | TCGA-LGG | Tumor TCGA | Astrocytoma | Grade III | Brain (Cortex) Tumor | 10.2409007 | 22.2253454 | 32.4662461 |
| TCGA-CS-5394 | TCGA-LGG | Tumor TCGA | Astrocytoma | Grade III | Brain (Cortex) Tumor | 16.6627174 | 17.4069958 | 34.0697132 |
| TCGA-CS-5395 | TCGA-LGG | Tumor TCGA | Oligodendroglion | Grade II | Brain (Cortex) Tumor | 16.3573677 | 35.7419756 | 52.0993433 |
| TCGA-CS-5396 | TCGA-LGG | Tumor TCGA | Oligodendroglion | Grade III | Brain (Cortex) Tumor | 19.4661983 | 16.1014763 | 35.5676746 |
| TCGA-CS-5397 | TCGA-LGG | Tumor TCGA | Astrocytoma | Grade III | Brain (Cortex) Tumor | 12.9278239 | 35.5262286 | 48.4540526 |
| TCGA-CS-6186 | TCGA-LGG | Tumor TCGA | Oligoastrocytoma | Grade III | Brain (Cortex) Tumor | 21.1000864 | 56.0149162 | 77.1150027 |
| TCGA-CS-6188 | TCGA-LGG | Tumor TCGA | Astrocytoma | Grade III | Brain (Cortex) Tumor | 16.5039009 | 38.4615477 | 54.9654485 |
| TCGA-CS-6290 | TCGA-LGG | Tumor TCGA | Astrocytoma | Grade III | Brain (Cortex) Tumor | 10.3444476 | 37.022631 | 47.3670786 |
| TCGA-CS-6666 | TCGA-LGG | Tumor TCGA | Astrocytoma | Grade III | Brain (Cortex) Tumor | 12.6744661 | 36.3390157 | 49.0134818 |
| TCGA-CS-6667 | TCGA-LGG | Tumor TCGA | Astrocytoma | Grade II | Brain (Cortex) Tumor | 8.97890244 | 27.781261 | 36.7601634 |
| TCGA-CS-6668 | TCGA-LGG | Tumor TCGA | Oligodendroglion | Grade II | Brain (Cortex) Tumor | 19.7312492 | 17.379091 | 37.1103402 |
| TCGA-CS-6669 | TCGA-LGG | Tumor TCGA | Oligodendroglion | Grade II | Brain (Cortex) Tumor | 5.79589886 | 17.2394933 | 23.0353922 |
| TCGA-CS-6670 | TCGA-LGG | Tumor TCGA | Oligodendroglion | Grade III | Brain (Cortex) Tumor | 9.98507284 | 17.509911 | 27.4949838 |
| TCGA-DB-5270 | TCGA-LGG | Tumor TCGA | Oligoastrocytoma | Grade III | Brain (Cortex) Tumor | 8.25156786 | 30.145389 | 38.3969568 |
| TCGA-DB-5273 | TCGA-LGG | Tumor TCGA | Astrocytoma | Grade III | Brain (Cortex) Tumor | 17.1569199 | 30.4644628 | 47.6213828 |
| TCGA-DB-5274 | TCGA-LGG | Tumor TCGA | Oligoastrocytoma | Grade III | Brain (Cortex) Tumor | 31.6300897 | 23.3493825 | 54.9794722 |
| TCGA-DB-5275 | TCGA-LGG | Tumor TCGA | Oligoastrocytoma | Grade III | Brain (Cortex) Tumor | 24.6927365 | 46.622983 | 71.3157195 |
| TCGA-DB-5277 | TCGA-LGG | Tumor TCGA | Astrocytoma | Grade III | Brain (Cortex) Tumor | 56.7121126 | 57.9112386 | 114.623351 |
| TCGA-DB-5278 | TCGA-LGG | Tumor TCGA | Oligodendroglion | Grade II | Brain (Cortex) Tumor | 14.6898563 | 19.7854302 | 34.4752865 |
| TCGA-DB-5279 | TCGA-LGG | Tumor TCGA | Oligodendroglion | Grade II | Brain (Cortex) Tumor | 19.9446021 | 18.8451489 | 38.789751 |
| TCGA-DB-5280 | TCGA-LGG | Tumor TCGA | Oligoastrocytoma | Grade II | Brain (Cortex) Tumor | 12.2276077 | 32.1389006 | 44.3665084 |

| | | | | | | | | |
|---|---|---|---|---|---|---|---|---|
| TCGA-DB-5281 | TCGA-LGG | Tumor TCGA | Oligoastrocytoma | Grade III | Brain (Cortex) Tumor | 8.22370567 | 31.3545944 | 39.5783001 |
| TCGA-DB-A4X9 | TCGA-LGG | Tumor TCGA | Oligoastrocytoma | Grade II | Brain (Cortex) Tumor | 16.4226737 | 33.1451841 | 49.5678578 |
| TCGA-DB-A4XA | TCGA-LGG | Tumor TCGA | Oligoastrocytoma | Grade II | Brain (Cortex) Tumor | 12.9934692 | 19.6243993 | 32.6178684 |
| TCGA-DB-A4XB | TCGA-LGG | Tumor TCGA | Astrocytoma | Grade III | Brain (Cortex) Tumor | 14.2085763 | 41.7393863 | 55.9479627 |
| TCGA-DB-A4XC | TCGA-LGG | Tumor TCGA | Oligoastrocytoma | Grade II | Brain (Cortex) Tumor | 13.2352156 | 28.620056 | 41.8552716 |
| TCGA-DB-A4XD | TCGA-LGG | Tumor TCGA | Astrocytoma | Grade III | Brain (Cortex) Tumor | 10.7969436 | 28.5316824 | 39.328626 |
| TCGA-DB-A4XE | TCGA-LGG | Tumor TCGA | Oligoastrocytoma | Grade III | Brain (Cortex) Tumor | 24.9078721 | 33.4347992 | 58.3426713 |
| TCGA-DB-A4XF | TCGA-LGG | Tumor TCGA | Astrocytoma | Grade III | Brain (Cortex) Tumor | 9.89418376 | 23.0349273 | 32.929111 |
| TCGA-DB-A4XG | TCGA-LGG | Tumor TCGA | Oligodendroglion | Grade III | Brain (Cortex) Tumor | 17.3334399 | 21.7802297 | 39.1136696 |
| TCGA-DB-A4XH | TCGA-LGG | Tumor TCGA | Oligoastrocytoma | Grade II | Brain (Cortex) Tumor | 8.33810961 | 10.6468931 | 18.9850027 |
| TCGA-DB-A64L | TCGA-LGG | Tumor TCGA | Oligodendroglion | Grade II | Brain (Cortex) Tumor | 21.7360783 | 22.8283724 | 44.5644508 |
| TCGA-DB-A64O | TCGA-LGG | Tumor TCGA | Oligoastrocytoma | Grade II | Brain (Cortex) Tumor | 15.7759659 | 42.4403279 | 58.2162938 |
| TCGA-DB-A64P | TCGA-LGG | Tumor TCGA | Oligodendroglion | Grade III | Brain (Cortex) Tumor | 24.0300076 | 19.7076201 | 43.7376276 |
| TCGA-DB-A64Q | TCGA-LGG | Tumor TCGA | Oligoastrocytoma | Grade II | Brain (Cortex) Tumor | 29.9009174 | 31.1548793 | 61.0557967 |
| TCGA-DB-A64R | TCGA-LGG | Tumor TCGA | Oligodendroglion | Grade II | Brain (Cortex) Tumor | 17.3711758 | 21.8540046 | 39.2251804 |
| TCGA-DB-A64S | TCGA-LGG | Tumor TCGA | Oligoastrocytoma | Grade II | Brain (Cortex) Tumor | 11.910554 | 28.8949944 | 40.8055484 |
| TCGA-DB-A64U | TCGA-LGG | Tumor TCGA | Oligoastrocytoma | Grade II | Brain (Cortex) Tumor | 18.8915313 | 23.8667593 | 42.7582906 |
| TCGA-DB-A64V | TCGA-LGG | Tumor TCGA | Oligodendroglion | Grade II | Brain (Cortex) Tumor | 17.4090927 | 23.3914904 | 40.8005831 |
| TCGA-DB-A64W | TCGA-LGG | Tumor TCGA | Oligoastrocytoma | Grade III | Brain (Cortex) Tumor | 22.7667992 | 26.2047358 | 48.971535 |
| TCGA-DB-A64X | TCGA-LGG | Tumor TCGA | Astrocytoma | Grade III | Brain (Cortex) Tumor | 9.93273878 | 30.3014788 | 40.2342176 |
| TCGA-DB-A75K | TCGA-LGG | Tumor TCGA | Oligoastrocytoma | Grade III | Brain (Cortex) Tumor | 18.6373851 | 24.276313 | 42.9136981 |
| TCGA-DB-A75L | TCGA-LGG | Tumor TCGA | Astrocytoma | Grade III | Brain (Cortex) Tumor | 13.0380538 | 37.3163248 | 50.3543786 |
| TCGA-DB-A75M | TCGA-LGG | Tumor TCGA | Astrocytoma | Grade II | Brain (Cortex) Tumor | 11.0073102 | 35.7013984 | 46.7087086 |
| TCGA-DB-A75O | TCGA-LGG | Tumor TCGA | Astrocytoma | Grade III | Brain (Cortex) Tumor | 13.9113626 | 31.9009112 | 45.8122737 |
| TCGA-DB-A75P | TCGA-LGG | Tumor TCGA | Astrocytoma | Grade II | Brain (Cortex) Tumor | 8.54027209 | 27.6013761 | 36.1416481 |
| TCGA-DH-5140 | TCGA-LGG | Tumor TCGA | Oligoastrocytoma | Grade III | Brain (Cortex) Tumor | 28.8740161 | 57.8830209 | 86.757037 |
| TCGA-DH-5141 | TCGA-LGG | Tumor TCGA | Oligodendroglion | Grade III | Brain (Cortex) Tumor | 13.2293161 | 15.4394617 | 28.6687778 |
| TCGA-DH-5142 | TCGA-LGG | Tumor TCGA | Astrocytoma | Grade III | Brain (Cortex) Tumor | 14.224446 | 40.799857 | 55.024303 |
| TCGA-DH-5143 | TCGA-LGG | Tumor TCGA | Oligoastrocytoma | Grade III | Brain (Cortex) Tumor | 13.9478204 | 26.3242269 | 40.2720473 |
| TCGA-DH-5144 | TCGA-LGG | Tumor TCGA | Oligodendroglion | Grade III | Brain (Cortex) Tumor | 20.1760876 | 24.056128 | 44.2322157 |
| TCGA-DH-A669 | TCGA-LGG | Tumor TCGA | Oligodendroglion | Grade III | Brain (Cortex) Tumor | 39.5949487 | 45.3126974 | 84.907646 |
| TCGA-DH-A66B | TCGA-LGG | Tumor TCGA | Astrocytoma | Grade III | Brain (Cortex) Tumor | 17.8188463 | 51.6046006 | 69.4234469 |
| TCGA-DH-A66D | TCGA-LGG | Tumor TCGA | Astrocytoma | Grade III | Brain (Cortex) Tumor | 10.3555885 | 34.2107581 | 44.5663466 |
| TCGA-DH-A66F | TCGA-LGG | Tumor TCGA | Oligodendroglion | Grade II | Brain (Cortex) Tumor | 22.3258613 | 17.0834673 | 39.4093286 |
| TCGA-DH-A66G | TCGA-LGG | Tumor TCGA | Oligodendroglion | Grade III | Brain (Cortex) Tumor | 8.73600984 | 31.8162897 | 40.5522995 |
| TCGA-DH-A7UR | TCGA-LGG | Tumor TCGA | Oligodendroglion | Grade III | Brain (Cortex) Tumor | 24.8836041 | 27.5801544 | 52.4637586 |
| TCGA-DH-A7US | TCGA-LGG | Tumor TCGA | Oligodendroglion | Grade II | Brain (Cortex) Tumor | 22.9278101 | 21.8354361 | 44.7632462 |
| TCGA-DH-A7UT | TCGA-LGG | Tumor TCGA | Astrocytoma | Grade III | Brain (Cortex) Tumor | 12.7349709 | 30.0102146 | 42.7451855 |

| | | | | | | | | |
|---|---|---|---|---|---|---|---|---|
| TCGA-DH-A7UU | TCGA-LGG | Tumor TCGA | Astrocytoma | Grade III | Brain (Cortex) Tumor | 16.6609048 | 36.9576803 | 53.6185851 |
| TCGA-DH-A7UV | TCGA-LGG | Tumor TCGA | Astrocytoma | Grade III | Brain (Cortex) Tumor | 9.59913856 | 28.0919934 | 37.691132 |
| TCGA-DU-5847 | TCGA-LGG | Tumor TCGA | Astrocytoma | Grade III | Brain (Cortex) Tumor | 21.0268508 | 68.2182656 | 89.2451165 |
| TCGA-DU-5849 | TCGA-LGG | Tumor TCGA | Oligodendroglion | Grade II | Brain (Cortex) Tumor | 11.5981131 | 18.0830848 | 29.6811979 |
| TCGA-DU-5852 | TCGA-LGG | Tumor TCGA | Oligoastrocytoma | Grade III | Brain (Cortex) Tumor | 14.7906792 | 38.7129084 | 53.5035876 |
| TCGA-DU-5853 | TCGA-LGG | Tumor TCGA | Oligoastrocytoma | Grade II | Brain (Cortex) Tumor | 11.012934 | 27.2882948 | 38.3012288 |
| TCGA-DU-5854 | TCGA-LGG | Tumor TCGA | Astrocytoma | Grade III | Brain (Cortex) Tumor | 12.3724731 | 33.2341376 | 45.6066107 |
| TCGA-DU-5855 | TCGA-LGG | Tumor TCGA | Oligoastrocytoma | Grade III | Brain (Cortex) Tumor | 11.2040269 | 39.1446363 | 50.3486632 |
| TCGA-DU-5870 | TCGA-LGG | Tumor TCGA | Oligodendroglion | Grade II | Brain (Cortex) Tumor | 25.8438155 | 21.4318443 | 47.2756599 |
| TCGA-DU-5871 | TCGA-LGG | Tumor TCGA | Oligoastrocytoma | Grade II | Brain (Cortex) Tumor | 8.69665208 | 28.6580418 | 37.3546939 |
| TCGA-DU-5872 | TCGA-LGG | Tumor TCGA | Oligoastrocytoma | Grade II | Brain (Cortex) Tumor | 7.85537728 | 23.4241582 | 31.2795355 |
| TCGA-DU-5874 | TCGA-LGG | Tumor TCGA | Oligodendroglion | Grade II | Brain (Cortex) Tumor | 20.1733808 | 22.7457904 | 42.9191712 |
| TCGA-DU-6392 | TCGA-LGG | Tumor TCGA | Astrocytoma | Grade III | Brain (Cortex) Tumor | 12.1823517 | 34.089941 | 46.2722926 |
| TCGA-DU-6393 | TCGA-LGG | Tumor TCGA | Oligodendroglion | Grade III | Brain (Cortex) Tumor | 17.8134973 | 20.0752697 | 37.8887669 |
| TCGA-DU-6394 | TCGA-LGG | Tumor TCGA | Oligodendroglion | Grade III | Brain (Cortex) Tumor | 23.0890368 | 21.6440222 | 44.733059 |
| TCGA-DU-6395 | TCGA-LGG | Tumor TCGA | Oligoastrocytoma | Grade II | Brain (Cortex) Tumor | 14.6236783 | 28.2830848 | 42.9067631 |
| TCGA-DU-6396 | TCGA-LGG | Tumor TCGA | Oligoastrocytoma | Grade III | Brain (Cortex) Tumor | 13.2830894 | 33.1371487 | 46.4202381 |
| TCGA-DU-6397 | TCGA-LGG | Tumor TCGA | Oligodendroglion | Grade III | Brain (Cortex) Tumor | 17.1200595 | 18.1322352 | 35.2522947 |
| TCGA-DU-6399 | TCGA-LGG | Tumor TCGA | Oligodendroglion | Grade II | Brain (Cortex) Tumor | 12.4272243 | 34.852514 | 47.2797383 |
| TCGA-DU-6400 | TCGA-LGG | Tumor TCGA | Oligodendroglion | Grade II | Brain (Cortex) Tumor | 20.7090756 | 24.158509 | 44.8675846 |
| TCGA-DU-6401 | TCGA-LGG | Tumor TCGA | Oligodendroglion | Grade II | Brain (Cortex) Tumor | 6.05942991 | 22.5973516 | 28.6567815 |
| TCGA-DU-6402 | TCGA-LGG | Tumor TCGA | Astrocytoma | Grade III | Brain (Cortex) Tumor | 14.9091016 | 60.0952003 | 75.0043019 |
| TCGA-DU-6403 | TCGA-LGG | Tumor TCGA | Oligoastrocytoma | Grade III | Brain (Cortex) Tumor | 13.2616506 | 42.4346655 | 55.6963162 |
| TCGA-DU-6404 | TCGA-LGG | Tumor TCGA | Oligodendroglion | Grade III | Brain (Cortex) Tumor | 37.5706563 | 62.9151662 | 100.485823 |
| TCGA-DU-6405 | TCGA-LGG | Tumor TCGA | Astrocytoma | Grade III | Brain (Cortex) Tumor | 16.1402104 | 48.1046078 | 64.2448181 |
| TCGA-DU-6406 | TCGA-LGG | Tumor TCGA | Oligoastrocytoma | Grade III | Brain (Cortex) Tumor | 20.5133327 | 48.3002808 | 68.8136134 |
| TCGA-DU-6407 | TCGA-LGG | Tumor TCGA | Oligodendroglion | Grade II | Brain (Cortex) Tumor | 8.02900172 | 29.0482015 | 37.0772032 |
| TCGA-DU-6408 | TCGA-LGG | Tumor TCGA | Oligodendroglion | Grade III | Brain (Cortex) Tumor | 15.5001428 | 31.6951502 | 47.195293 |
| TCGA-DU-6410 | TCGA-LGG | Tumor TCGA | Oligodendroglion | Grade III | Brain (Cortex) Tumor | 15.6964091 | 15.2190003 | 30.9154094 |
| TCGA-DU-6542 | TCGA-LGG | Tumor TCGA | Oligoastrocytoma | Grade III | Brain (Cortex) Tumor | 10.2100015 | 36.7346452 | 46.9446467 |
| TCGA-DU-7006 | TCGA-LGG | Tumor TCGA | Astrocytoma | Grade III | Brain (Cortex) Tumor | 23.8830551 | 55.452836 | 79.3358911 |
| TCGA-DU-7007 | TCGA-LGG | Tumor TCGA | Astrocytoma | Grade II | Brain (Cortex) Tumor | 98.5849037 | 45.0809611 | 143.665865 |
| TCGA-DU-7008 | TCGA-LGG | Tumor TCGA | Oligodendroglion | Grade II | Brain (Cortex) Tumor | 16.9745471 | 33.7056152 | 50.6801623 |
| TCGA-DU-7009 | TCGA-LGG | Tumor TCGA | Oligodendroglion | Grade II | Brain (Cortex) Tumor | 19.6514726 | 18.8839732 | 38.5354457 |
| TCGA-DU-7010 | TCGA-LGG | Tumor TCGA | Astrocytoma | Grade III | Brain (Cortex) Tumor | 20.7308244 | 71.052954 | 91.7837783 |
| TCGA-DU-7011 | TCGA-LGG | Tumor TCGA | Oligoastrocytoma | Grade II | Brain (Cortex) Tumor | 7.9965042 | 23.4095583 | 31.4060625 |
| TCGA-DU-7012 | TCGA-LGG | Tumor TCGA | Astrocytoma | Grade III | Brain (Cortex) Tumor | 14.1666484 | 47.4172574 | 61.5839058 |
| TCGA-DU-7013 | TCGA-LGG | Tumor TCGA | Astrocytoma | Grade III | Brain (Cortex) Tumor | 17.4093892 | 45.3705404 | 62.7799296 |

| | | | | | | | | |
|---|---|---|---|---|---|---|---|---|
| TCGA-DU-7014 | TCGA-LGG | Tumor TCGA | Oligodendroglion | Grade II | Brain (Cortex) Tumor | 11.2548345 | 24.1835424 | 35.4383769 |
| TCGA-DU-7015 | TCGA-LGG | Tumor TCGA | Oligodendroglion | Grade II | Brain (Cortex) Tumor | 12.0809508 | 30.7030351 | 42.7839858 |
| TCGA-DU-7018 | TCGA-LGG | Tumor TCGA | Oligodendroglion | Grade III | Brain (Cortex) Tumor | 21.2305831 | 14.6453907 | 35.8759738 |
| TCGA-DU-7290 | TCGA-LGG | Tumor TCGA | Astrocytoma | Grade III | Brain (Cortex) Tumor | 19.1450647 | 80.5558534 | 99.7009181 |
| TCGA-DU-7292 | TCGA-LGG | Tumor TCGA | Astrocytoma | Grade III | Brain (Cortex) Tumor | 15.5598373 | 33.1771737 | 48.737011 |
| TCGA-DU-7294 | TCGA-LGG | Tumor TCGA | Oligodendroglion | Grade II | Brain (Cortex) Tumor | 21.6535934 | 18.0919744 | 39.7455679 |
| TCGA-DU-7298 | TCGA-LGG | Tumor TCGA | Astrocytoma | Grade III | Brain (Cortex) Tumor | 21.5087134 | 43.1280562 | 64.6367696 |
| TCGA-DU-7299 | TCGA-LGG | Tumor TCGA | Astrocytoma | Grade III | Brain (Cortex) Tumor | 11.9718985 | 35.817709 | 47.7896075 |
| TCGA-DU-7300 | TCGA-LGG | Tumor TCGA | Oligodendroglion | Grade III | Brain (Cortex) Tumor | 16.6577954 | 21.4777734 | 38.1355688 |
| TCGA-DU-7301 | TCGA-LGG | Tumor TCGA | Oligodendroglion | Grade II | Brain (Cortex) Tumor | 11.2965391 | 29.0376654 | 40.3342045 |
| TCGA-DU-7302 | TCGA-LGG | Tumor TCGA | Oligodendroglion | Grade III | Brain (Cortex) Tumor | 19.4080258 | 21.0295343 | 40.4375602 |
| TCGA-DU-7304 | TCGA-LGG | Tumor TCGA | Oligoastrocytoma | Grade III | Brain (Cortex) Tumor | 10.6911558 | 28.3568238 | 39.0479795 |
| TCGA-DU-7306 | TCGA-LGG | Tumor TCGA | Oligoastrocytoma | Grade II | Brain (Cortex) Tumor | 7.68026127 | 30.0038863 | 37.6841475 |
| TCGA-DU-7309 | TCGA-LGG | Tumor TCGA | Oligodendroglion | Grade III | Brain (Cortex) Tumor | 6.28486341 | 19.949619 | 26.2344824 |
| TCGA-DU-8158 | TCGA-LGG | Tumor TCGA | Astrocytoma | Grade III | Brain (Cortex) Tumor | 14.9644356 | 38.7911246 | 53.7555603 |
| TCGA-DU-8161 | TCGA-LGG | Tumor TCGA | Oligoastrocytoma | Grade III | Brain (Cortex) Tumor | 19.4428125 | 45.9523646 | 65.3951771 |
| TCGA-DU-8162 | TCGA-LGG | Tumor TCGA | Oligoastrocytoma | Grade III | Brain (Cortex) Tumor | 11.6413937 | 27.7083322 | 39.3497259 |
| TCGA-DU-8163 | TCGA-LGG | Tumor TCGA | Oligoastrocytoma | Grade III | Brain (Cortex) Tumor | 10.7330651 | 30.0333681 | 40.7664331 |
| TCGA-DU-8164 | TCGA-LGG | Tumor TCGA | Oligodendroglion | Grade II | Brain (Cortex) Tumor | 17.9079361 | 20.2190381 | 38.1269742 |
| TCGA-DU-8165 | TCGA-LGG | Tumor TCGA | Oligodendroglion | Grade III | Brain (Cortex) Tumor | 32.5256973 | 99.6054801 | 132.131177 |
| TCGA-DU-8166 | TCGA-LGG | Tumor TCGA | Oligoastrocytoma | Grade II | Brain (Cortex) Tumor | 10.9438977 | 38.460255 | 49.4041528 |
| TCGA-DU-8167 | TCGA-LGG | Tumor TCGA | Oligoastrocytoma | Grade II | Brain (Cortex) Tumor | 14.3635125 | 37.582196 | 51.9457085 |
| TCGA-DU-8168 | TCGA-LGG | Tumor TCGA | Oligodendroglion | Grade III | Brain (Cortex) Tumor | 27.9744689 | 21.5017357 | 49.4762046 |
| TCGA-DU-A5TP | TCGA-LGG | Tumor TCGA | Astrocytoma | Grade III | Brain (Cortex) Tumor | 23.7323049 | 66.324411 | 90.0567158 |
| TCGA-DU-A5TR | TCGA-LGG | Tumor TCGA | Oligoastrocytoma | Grade II | Brain (Cortex) Tumor | 12.871046 | 26.3771909 | 39.2482369 |
| TCGA-DU-A5TS | TCGA-LGG | Tumor TCGA | Oligodendroglion | Grade II | Brain (Cortex) Tumor | 13.4419328 | 20.0388254 | 33.4807582 |
| TCGA-DU-A5TT | TCGA-LGG | Tumor TCGA | Oligodendroglion | Grade III | Brain (Cortex) Tumor | 18.0224354 | 60.8342999 | 78.8567353 |
| TCGA-DU-A5TU | TCGA-LGG | Tumor TCGA | Astrocytoma | Grade II | Brain (Cortex) Tumor | 37.0218412 | 43.2637769 | 80.2856182 |
| TCGA-DU-A5TW | TCGA-LGG | Tumor TCGA | Astrocytoma | Grade III | Brain (Cortex) Tumor | 12.4075736 | 38.3188076 | 50.7263812 |
| TCGA-DU-A5TY | TCGA-LGG | Tumor TCGA | Astrocytoma | Grade III | Brain (Cortex) Tumor | 24.9396941 | 49.9813202 | 74.9210142 |
| TCGA-DU-A6S2 | TCGA-LGG | Tumor TCGA | Oligodendroglion | Grade II | Brain (Cortex) Tumor | 12.0377535 | 21.2683111 | 33.3060646 |
| TCGA-DU-A6S3 | TCGA-LGG | Tumor TCGA | Oligodendroglion | Grade II | Brain (Cortex) Tumor | 11.9635656 | 16.5104973 | 28.4740629 |
| TCGA-DU-A6S6 | TCGA-LGG | Tumor TCGA | Oligoastrocytoma | Grade II | Brain (Cortex) Tumor | 12.319475 | 19.1728099 | 31.4922849 |
| TCGA-DU-A6S7 | TCGA-LGG | Tumor TCGA | Astrocytoma | Grade III | Brain (Cortex) Tumor | 10.081031 | 36.7904107 | 46.8714417 |
| TCGA-DU-A6S8 | TCGA-LGG | Tumor TCGA | Oligodendroglion | Grade III | Brain (Cortex) Tumor | 33.2845566 | 25.6095972 | 58.8941538 |
| TCGA-DU-A76K | TCGA-LGG | Tumor TCGA | Oligodendroglion | Grade II | Brain (Cortex) Tumor | 12.1002025 | 26.2817151 | 38.3819176 |
| TCGA-DU-A76L | TCGA-LGG | Tumor TCGA | Oligodendroglion | Grade III | Brain (Cortex) Tumor | 30.8120082 | 71.3749987 | 102.187007 |
| TCGA-DU-A76O | TCGA-LGG | Tumor TCGA | Astrocytoma | Grade II | Brain (Cortex) Tumor | 12.0726313 | 32.4674503 | 44.5400816 |

| | | | | | | | | |
|---|---|---|---|---|---|---|---|---|
| TCGA-DU-A76R | TCGA-LGG | Tumor TCGA | Oligodendroglion | Grade III | Brain (Cortex) Tumor | 17.5092835 | 22.9320919 | 40.4413755 |
| TCGA-DU-A7T6 | TCGA-LGG | Tumor TCGA | Oligodendroglion | Grade III | Brain (Cortex) Tumor | 32.2292869 | 25.714968 | 57.9442549 |
| TCGA-DU-A7T8 | TCGA-LGG | Tumor TCGA | Oligoastrocytoma | Grade III | Brain (Cortex) Tumor | 15.742124 | 32.4710741 | 48.2131981 |
| TCGA-DU-A7TA | TCGA-LGG | Tumor TCGA | Oligodendroglion | Grade II | Brain (Cortex) Tumor | 23.6368763 | 43.6599704 | 67.2968467 |
| TCGA-DU-A7TB | TCGA-LGG | Tumor TCGA | Oligodendroglion | Grade II | Brain (Cortex) Tumor | 17.0808695 | 31.5898555 | 48.670725 |
| TCGA-DU-A7TC | TCGA-LGG | Tumor TCGA | Astrocytoma | Grade II | Brain (Cortex) Tumor | 12.6507454 | 31.9626992 | 44.6134446 |
| TCGA-DU-A7TG | TCGA-LGG | Tumor TCGA | Oligodendroglion | Grade II | Brain (Cortex) Tumor | 11.6785701 | 28.9691045 | 40.6476745 |
| TCGA-DU-A7TI | TCGA-LGG | Tumor TCGA | Astrocytoma | Grade III | Brain (Cortex) Tumor | 25.2546668 | 39.8334501 | 65.0881169 |
| TCGA-DU-A7TJ | TCGA-LGG | Tumor TCGA | Astrocytoma | Grade III | Brain (Cortex) Tumor | 25.1557483 | 46.6413199 | 71.7970682 |
| TCGA-E1-5303 | TCGA-LGG | Tumor TCGA | Astrocytoma | Grade III | Brain (Cortex) Tumor | 10.061596 | 28.1110304 | 38.1726264 |
| TCGA-E1-5304 | TCGA-LGG | Tumor TCGA | Astrocytoma | Grade III | Brain (Cortex) Tumor | 25.2706047 | 82.332642 | 107.603247 |
| TCGA-E1-5307 | TCGA-LGG | Tumor TCGA | Astrocytoma | Grade III | Brain (Cortex) Tumor | 11.1678982 | 31.5201286 | 42.6880268 |
| TCGA-E1-5311 | TCGA-LGG | Tumor TCGA | Oligodendroglion | Grade III | Brain (Cortex) Tumor | 15.8464133 | 20.8155499 | 36.6619632 |
| TCGA-E1-5318 | TCGA-LGG | Tumor TCGA | Oligodendroglion | Grade II | Brain (Cortex) Tumor | 14.4997536 | 16.9158504 | 31.4156041 |
| TCGA-E1-5319 | TCGA-LGG | Tumor TCGA | Oligodendroglion | Grade II | Brain (Cortex) Tumor | 18.303681 | 19.6715103 | 37.9751913 |
| TCGA-E1-5322 | TCGA-LGG | Tumor TCGA | Oligoastrocytoma | Grade II | Brain (Cortex) Tumor | 12.6559558 | 43.5091887 | 56.1651446 |
| TCGA-E1-A7YD | TCGA-LGG | Tumor TCGA | Astrocytoma | Grade III | Brain (Cortex) Tumor | 34.4916782 | 69.2108493 | 103.702528 |
| TCGA-E1-A7YE | TCGA-LGG | Tumor TCGA | Astrocytoma | Grade III | Brain (Cortex) Tumor | 15.7009333 | 47.4129407 | 63.113874 |
| TCGA-E1-A7YH | TCGA-LGG | Tumor TCGA | Astrocytoma | Grade III | Brain (Cortex) Tumor | 22.6803502 | 35.850172 | 58.5305222 |
| TCGA-E1-A7YI | TCGA-LGG | Tumor TCGA | Astrocytoma | Grade III | Brain (Cortex) Tumor | 24.7311135 | 75.7250825 | 100.456196 |
| TCGA-E1-A7YJ | TCGA-LGG | Tumor TCGA | Astrocytoma | Grade III | Brain (Cortex) Tumor | 29.5017688 | 71.864852 | 101.366621 |
| TCGA-E1-A7YK | TCGA-LGG | Tumor TCGA | Astrocytoma | Grade III | Brain (Cortex) Tumor | 16.8186854 | 37.4170507 | 54.2357361 |
| TCGA-E1-A7YL | TCGA-LGG | Tumor TCGA | Astrocytoma | Grade III | Brain (Cortex) Tumor | 26.5283458 | 44.4932439 | 71.0215897 |
| TCGA-E1-A7YM | TCGA-LGG | Tumor TCGA | Astrocytoma | Grade III | Brain (Cortex) Tumor | 25.4242826 | 42.9350031 | 68.3592857 |
| TCGA-E1-A7YN | TCGA-LGG | Tumor TCGA | Astrocytoma | Grade III | Brain (Cortex) Tumor | 22.198704 | 51.2252075 | 73.4239115 |
| TCGA-E1-A7YO | TCGA-LGG | Tumor TCGA | Oligodendroglion | Grade III | Brain (Cortex) Tumor | 17.7734771 | 19.8339769 | 37.607454 |
| TCGA-E1-A7YQ | TCGA-LGG | Tumor TCGA | Oligodendroglion | Grade III | Brain (Cortex) Tumor | 40.0492323 | 66.9305061 | 106.979738 |
| TCGA-E1-A7YS | TCGA-LGG | Tumor TCGA | Oligodendroglion | Grade III | Brain (Cortex) Tumor | 21.4621026 | 23.4143985 | 44.876501 |
| TCGA-E1-A7YU | TCGA-LGG | Tumor TCGA | Oligoastrocytoma | Grade III | Brain (Cortex) Tumor | 9.09670792 | 32.9460003 | 42.0427082 |
| TCGA-E1-A7YV | TCGA-LGG | Tumor TCGA | Oligoastrocytoma | Grade III | Brain (Cortex) Tumor | 15.8000727 | 43.8648004 | 59.6648731 |
| TCGA-E1-A7YW | TCGA-LGG | Tumor TCGA | Oligoastrocytoma | Grade II | Brain (Cortex) Tumor | 15.383023 | 38.4290058 | 53.8120288 |
| TCGA-E1-A7Z2 | TCGA-LGG | Tumor TCGA | Oligodendroglion | Grade II | Brain (Cortex) Tumor | 19.0679298 | 41.3306061 | 60.3985359 |
| TCGA-E1-A7Z3 | TCGA-LGG | Tumor TCGA | Astrocytoma | Grade II | Brain (Cortex) Tumor | 8.18566533 | 28.8475554 | 37.0332207 |
| TCGA-E1-A7Z4 | TCGA-LGG | Tumor TCGA | Astrocytoma | Grade II | Brain (Cortex) Tumor | 16.4257664 | 45.7580524 | 62.1838188 |
| TCGA-E1-A7Z6 | TCGA-LGG | Tumor TCGA | Astrocytoma | Grade II | Brain (Cortex) Tumor | 13.5919913 | 30.1763714 | 43.7683628 |
| TCGA-EZ-7264 | TCGA-LGG | Tumor TCGA | Oligodendroglion | Grade II | Brain (Cortex) Tumor | 19.7444533 | 19.3741989 | 39.1186522 |
| TCGA-F6-A8O3 | TCGA-LGG | Tumor TCGA | Oligodendroglion | Grade II | Brain (Cortex) Tumor | 29.1921981 | 27.2006235 | 56.3928217 |
| TCGA-F6-A8O4 | TCGA-LGG | Tumor TCGA | Astrocytoma | Grade II | Brain (Cortex) Tumor | 14.3362604 | 43.1359958 | 57.4722562 |

| | | | | | | | | |
|---|---|---|---|---|---|---|---|---|
| TCGA-FG-5962 | TCGA-LGG | Tumor TCGA | Oligodendroglion | Grade III | Brain (Cortex) Tumor | 19.7593643 | 22.8538978 | 42.6132621 |
| TCGA-FG-5963 | TCGA-LGG | Tumor TCGA | Astrocytoma | Grade III | Brain (Cortex) Tumor | 15.0821899 | 36.4385671 | 51.520757 |
| TCGA-FG-5964 | TCGA-LGG | Tumor TCGA | Oligodendroglion | Grade II | Brain (Cortex) Tumor | 10.8636755 | 25.3534359 | 36.2171114 |
| TCGA-FG-5965 | TCGA-LGG | Tumor TCGA | Oligoastrocytoma | Grade II | Brain (Cortex) Tumor | 12.4054676 | 34.1281101 | 46.5335777 |
| TCGA-FG-6688 | TCGA-LGG | Tumor TCGA | Astrocytoma | Grade III | Brain (Cortex) Tumor | 15.0395673 | 38.8046847 | 53.844252 |
| TCGA-FG-6689 | TCGA-LGG | Tumor TCGA | Astrocytoma | Grade II | Brain (Cortex) Tumor | 9.06472733 | 31.6487914 | 40.7135188 |
| TCGA-FG-6690 | TCGA-LGG | Tumor TCGA | Oligodendroglion | Grade II | Brain (Cortex) Tumor | 10.1089184 | 25.5646758 | 35.6735942 |
| TCGA-FG-6691 | TCGA-LGG | Tumor TCGA | Astrocytoma | Grade II | Brain (Cortex) Tumor | 16.4503415 | 29.3424243 | 45.7927658 |
| TCGA-FG-6692 | TCGA-LGG | Tumor TCGA | Oligodendroglion | Grade III | Brain (Cortex) Tumor | 23.8200494 | 42.1922786 | 66.0123279 |
| TCGA-FG-7634 | TCGA-LGG | Tumor TCGA | Oligodendroglion | Grade II | Brain (Cortex) Tumor | 21.8643617 | 23.8208754 | 45.6852371 |
| TCGA-FG-7636 | TCGA-LGG | Tumor TCGA | Astrocytoma | Grade III | Brain (Cortex) Tumor | 11.0334801 | 26.3975805 | 37.4310606 |
| TCGA-FG-7637 | TCGA-LGG | Tumor TCGA | Oligoastrocytoma | Grade II | Brain (Cortex) Tumor | 16.9452791 | 18.4082441 | 35.3535233 |
| TCGA-FG-7638 | TCGA-LGG | Tumor TCGA | Oligodendroglion | Grade III | Brain (Cortex) Tumor | 12.6655702 | 18.8999129 | 31.5654831 |
| TCGA-FG-7641 | TCGA-LGG | Tumor TCGA | Oligodendroglion | Grade II | Brain (Cortex) Tumor | 18.9538968 | 22.0984097 | 41.0523065 |
| TCGA-FG-7643 | TCGA-LGG | Tumor TCGA | Oligoastrocytoma | Grade II | Brain (Cortex) Tumor | 9.89762665 | 30.8141305 | 40.7117571 |
| TCGA-FG-8181 | TCGA-LGG | Tumor TCGA | Oligoastrocytoma | Grade III | Brain (Cortex) Tumor | 7.8834469 | 30.9463501 | 38.829797 |
| TCGA-FG-8182 | TCGA-LGG | Tumor TCGA | Oligodendroglion | Grade II | Brain (Cortex) Tumor | 14.6399362 | 33.2034204 | 47.8433565 |
| TCGA-FG-8185 | TCGA-LGG | Tumor TCGA | Astrocytoma | Grade III | Brain (Cortex) Tumor | 14.1057678 | 29.788699 | 43.8944668 |
| TCGA-FG-8186 | TCGA-LGG | Tumor TCGA | Oligoastrocytoma | Grade III | Brain (Cortex) Tumor | 15.9868006 | 24.9565754 | 40.943376 |
| TCGA-FG-8187 | TCGA-LGG | Tumor TCGA | Oligoastrocytoma | Grade II | Brain (Cortex) Tumor | 17.4435452 | 19.6063802 | 37.0499253 |
| TCGA-FG-8188 | TCGA-LGG | Tumor TCGA | Oligoastrocytoma | Grade II | Brain (Cortex) Tumor | 10.3139445 | 30.3604467 | 40.6743912 |
| TCGA-FG-8189 | TCGA-LGG | Tumor TCGA | Oligodendroglion | Grade II | Brain (Cortex) Tumor | 7.89394801 | 25.8450777 | 33.7390257 |
| TCGA-FG-8191 | TCGA-LGG | Tumor TCGA | Oligodendroglion | Grade III | Brain (Cortex) Tumor | 12.0346866 | 32.7946962 | 44.8293827 |
| TCGA-FG-A4MT | TCGA-LGG | Tumor TCGA | Oligodendroglion | Grade II | Brain (Cortex) Tumor | 12.6992263 | 37.2174905 | 49.9167168 |
| TCGA-FG-A4MU | TCGA-LGG | Tumor TCGA | Oligoastrocytoma | Grade III | Brain (Cortex) Tumor | 18.7403677 | 44.8205278 | 63.5608955 |
| TCGA-FG-A4MW | TCGA-LGG | Tumor TCGA | Oligoastrocytoma | Grade III | Brain (Cortex) Tumor | 12.7270117 | 28.7562874 | 41.483299 |
| TCGA-FG-A4MX | TCGA-LGG | Tumor TCGA | Astrocytoma | Grade II | Brain (Cortex) Tumor | 8.48813326 | 20.427179 | 28.9153123 |
| TCGA-FG-A4MY | TCGA-LGG | Tumor TCGA | Oligoastrocytoma | Grade II | Brain (Cortex) Tumor | 11.999142 | 34.2686863 | 46.2678283 |
| TCGA-FG-A60J | TCGA-LGG | Tumor TCGA | Oligoastrocytoma | Grade II | Brain (Cortex) Tumor | 13.7872156 | 28.6679567 | 42.4551723 |
| TCGA-FG-A60K | TCGA-LGG | Tumor TCGA | Oligoastrocytoma | Grade II | Brain (Cortex) Tumor | 19.7091327 | 18.6917932 | 38.400926 |
| TCGA-FG-A60L | TCGA-LGG | Tumor TCGA | Astrocytoma | Grade II | Brain (Cortex) Tumor | 14.4465974 | 29.7375751 | 44.1841725 |
| TCGA-FG-A6IZ | TCGA-LGG | Tumor TCGA | Oligodendroglion | Grade II | Brain (Cortex) Tumor | 29.4906046 | 22.8824339 | 52.3730385 |
| TCGA-FG-A6J1 | TCGA-LGG | Tumor TCGA | Oligodendroglion | Grade II | Brain (Cortex) Tumor | 21.2494039 | 18.6599641 | 39.909368 |
| TCGA-FG-A6J3 | TCGA-LGG | Tumor TCGA | Astrocytoma | Grade III | Brain (Cortex) Tumor | 54.5232108 | 57.4399072 | 111.963118 |
| TCGA-FG-A70Y | TCGA-LGG | Tumor TCGA | Oligodendroglion | Grade II | Brain (Cortex) Tumor | 9.7789736 | 26.7521209 | 36.5310945 |
| TCGA-FG-A70Z | TCGA-LGG | Tumor TCGA | Oligoastrocytoma | Grade III | Brain (Cortex) Tumor | 25.8609964 | 45.720811 | 71.5818074 |
| TCGA-FG-A710 | TCGA-LGG | Tumor TCGA | Oligodendroglion | Grade II | Brain (Cortex) Tumor | 22.3737782 | 29.0320154 | 51.4057935 |
| TCGA-FG-A711 | TCGA-LGG | Tumor TCGA | Oligodendroglion | Grade II | Brain (Cortex) Tumor | 9.02979603 | 44.5238873 | 53.5536834 |

| | | | | | | | | |
|---|---|---|---|---|---|---|---|---|
| TCGA-FG-A713 | TCGA-LGG | Tumor TCGA | Oligoastrocytoma | Grade II | Brain (Cortex) Tumor | 6.81563663 | 21.6785539 | 28.4941905 |
| TCGA-FG-A87N | TCGA-LGG | Tumor TCGA | Astrocytoma | Grade III | Brain (Cortex) Tumor | 15.1547727 | 49.4297655 | 64.5845382 |
| TCGA-FG-A87Q | TCGA-LGG | Tumor TCGA | Astrocytoma | Grade III | Brain (Cortex) Tumor | 16.6818507 | 39.8826292 | 56.5644798 |
| TCGA-FN-7833 | TCGA-LGG | Tumor TCGA | Oligoastrocytoma | Grade III | Brain (Cortex) Tumor | 15.3595896 | 38.1070724 | 53.466662 |
| TCGA-HT-7467 | TCGA-LGG | Tumor TCGA | Oligodendroglion | Grade II | Brain (Cortex) Tumor | 10.1192966 | 17.8281082 | 27.9474047 |
| TCGA-HT-7468 | TCGA-LGG | Tumor TCGA | Oligodendroglion | Grade III | Brain (Cortex) Tumor | 21.2660472 | 22.4346442 | 43.7006914 |
| TCGA-HT-7469 | TCGA-LGG | Tumor TCGA | Oligodendroglion | Grade III | Brain (Cortex) Tumor | 22.5483168 | 61.7150138 | 84.2633306 |
| TCGA-HT-7470 | TCGA-LGG | Tumor TCGA | Oligodendroglion | Grade III | Brain (Cortex) Tumor | 8.28482231 | 28.2121115 | 36.4969338 |
| TCGA-HT-7471 | TCGA-LGG | Tumor TCGA | Oligodendroglion | Grade III | Brain (Cortex) Tumor | 34.1619096 | 29.2286055 | 63.3905152 |
| TCGA-HT-7472 | TCGA-LGG | Tumor TCGA | Oligodendroglion | Grade II | Brain (Cortex) Tumor | 10.9495136 | 32.3625574 | 43.3120709 |
| TCGA-HT-7473 | TCGA-LGG | Tumor TCGA | Oligoastrocytoma | Grade II | Brain (Cortex) Tumor | 16.5718233 | 35.495505 | 52.0673284 |
| TCGA-HT-7474 | TCGA-LGG | Tumor TCGA | Oligoastrocytoma | Grade II | Brain (Cortex) Tumor | 9.48040004 | 25.3668905 | 34.8472906 |
| TCGA-HT-7475 | TCGA-LGG | Tumor TCGA | Oligoastrocytoma | Grade III | Brain (Cortex) Tumor | 11.3842202 | 35.7331225 | 47.1173426 |
| TCGA-HT-7476 | TCGA-LGG | Tumor TCGA | Astrocytoma | Grade II | Brain (Cortex) Tumor | 9.63871247 | 22.7976621 | 32.4363746 |
| TCGA-HT-7477 | TCGA-LGG | Tumor TCGA | Astrocytoma | Grade III | Brain (Cortex) Tumor | 35.7726595 | 51.5020338 | 87.2746933 |
| TCGA-HT-7478 | TCGA-LGG | Tumor TCGA | Astrocytoma | Grade II | Brain (Cortex) Tumor | 14.8134038 | 31.2395215 | 46.0529254 |
| TCGA-HT-7479 | TCGA-LGG | Tumor TCGA | Astrocytoma | Grade III | Brain (Cortex) Tumor | 13.9848012 | 28.325311 | 42.3101122 |
| TCGA-HT-7480 | TCGA-LGG | Tumor TCGA | Oligodendroglion | Grade II | Brain (Cortex) Tumor | 13.0712765 | 19.8688211 | 32.9400975 |
| TCGA-HT-7481 | TCGA-LGG | Tumor TCGA | Oligodendroglion | Grade II | Brain (Cortex) Tumor | 18.0473554 | 17.5990241 | 35.6463794 |
| TCGA-HT-7482 | TCGA-LGG | Tumor TCGA | Oligoastrocytoma | Grade II | Brain (Cortex) Tumor | 14.778876 | 35.4999115 | 50.2787874 |
| TCGA-HT-7483 | TCGA-LGG | Tumor TCGA | Oligoastrocytoma | Grade II | Brain (Cortex) Tumor | 11.7530369 | 34.8826704 | 46.6357074 |
| TCGA-HT-7485 | TCGA-LGG | Tumor TCGA | Astrocytoma | Grade II | Brain (Cortex) Tumor | 12.5839056 | 33.9303532 | 46.5142588 |
| TCGA-HT-7601 | TCGA-LGG | Tumor TCGA | Astrocytoma | Grade III | Brain (Cortex) Tumor | 10.9656648 | 31.6619761 | 42.6276409 |
| TCGA-HT-7602 | TCGA-LGG | Tumor TCGA | Oligodendroglion | Grade II | Brain (Cortex) Tumor | 12.3319028 | 22.6886326 | 35.0205354 |
| TCGA-HT-7603 | TCGA-LGG | Tumor TCGA | Oligodendroglion | Grade II | Brain (Cortex) Tumor | 12.782185 | 41.1248359 | 53.9070209 |
| TCGA-HT-7604 | TCGA-LGG | Tumor TCGA | Astrocytoma | Grade II | Brain (Cortex) Tumor | 40.3009914 | 34.8735788 | 75.1745702 |
| TCGA-HT-7605 | TCGA-LGG | Tumor TCGA | Oligodendroglion | Grade II | Brain (Cortex) Tumor | 13.3157192 | 21.4204196 | 34.7361388 |
| TCGA-HT-7606 | TCGA-LGG | Tumor TCGA | Astrocytoma | Grade II | Brain (Cortex) Tumor | 18.8433281 | 56.5592596 | 75.4025877 |
| TCGA-HT-7607 | TCGA-LGG | Tumor TCGA | Astrocytoma | Grade II | Brain (Cortex) Tumor | 9.1962665 | 21.8111236 | 31.0073901 |
| TCGA-HT-7608 | TCGA-LGG | Tumor TCGA | Oligoastrocytoma | Grade II | Brain (Cortex) Tumor | 13.8612202 | 14.7148723 | 28.5760924 |
| TCGA-HT-7609 | TCGA-LGG | Tumor TCGA | Oligoastrocytoma | Grade III | Brain (Cortex) Tumor | 11.0018091 | 26.8536399 | 37.8554491 |
| TCGA-HT-7610 | TCGA-LGG | Tumor TCGA | Oligoastrocytoma | Grade II | Brain (Cortex) Tumor | 8.84121305 | 26.4970875 | 35.3383005 |
| TCGA-HT-7611 | TCGA-LGG | Tumor TCGA | Oligoastrocytoma | Grade II | Brain (Cortex) Tumor | 14.2884902 | 38.9949216 | 53.2834118 |
| TCGA-HT-7616 | TCGA-LGG | Tumor TCGA | Oligodendroglion | Grade III | Brain (Cortex) Tumor | 28.5212732 | 27.8879547 | 56.4092279 |
| TCGA-HT-7620 | TCGA-LGG | Tumor TCGA | Oligodendroglion | Grade III | Brain (Cortex) Tumor | 16.0268423 | 20.5237214 | 36.5505637 |
| TCGA-HT-7676 | TCGA-LGG | Tumor TCGA | Oligodendroglion | Grade II | Brain (Cortex) Tumor | 11.2352993 | 29.4598189 | 40.6951182 |
| TCGA-HT-7677 | TCGA-LGG | Tumor TCGA | Oligodendroglion | Grade III | Brain (Cortex) Tumor | 22.0328516 | 19.9301762 | 41.9630278 |
| TCGA-HT-7680 | TCGA-LGG | Tumor TCGA | Astrocytoma | Grade II | Brain (Cortex) Tumor | 12.5096052 | 35.8911236 | 48.4007287 |

| | | | | | | | | |
|---|---|---|---|---|---|---|---|---|
| TCGA-HT-7681 | TCGA-LGG | Tumor TCGA | Oligoastrocytoma | Grade II | Brain (Cortex) Tumor | 15.5040602 | 28.1089898 | 43.61305 |
| TCGA-HT-7684 | TCGA-LGG | Tumor TCGA | Oligoastrocytoma | Grade III | Brain (Cortex) Tumor | 12.7224365 | 34.6798806 | 47.4023172 |
| TCGA-HT-7686 | TCGA-LGG | Tumor TCGA | Astrocytoma | Grade III | Brain (Cortex) Tumor | 11.0399258 | 43.8045675 | 54.8444933 |
| TCGA-HT-7687 | TCGA-LGG | Tumor TCGA | Oligodendroglion | Grade III | Brain (Cortex) Tumor | 20.1420148 | 22.5320416 | 42.6740564 |
| TCGA-HT-7688 | TCGA-LGG | Tumor TCGA | Oligodendroglion | Grade III | Brain (Cortex) Tumor | 14.5484335 | 37.1853669 | 51.7338003 |
| TCGA-HT-7689 | TCGA-LGG | Tumor TCGA | Oligodendroglion | Grade II | Brain (Cortex) Tumor | 15.1619038 | 39.3325506 | 54.4944545 |
| TCGA-HT-7690 | TCGA-LGG | Tumor TCGA | Oligoastrocytoma | Grade III | Brain (Cortex) Tumor | 19.875233 | 44.6145864 | 64.4898194 |
| TCGA-HT-7691 | TCGA-LGG | Tumor TCGA | Astrocytoma | Grade II | Brain (Cortex) Tumor | 16.0475923 | 40.8400849 | 56.8876772 |
| TCGA-HT-7692 | TCGA-LGG | Tumor TCGA | Oligoastrocytoma | Grade II | Brain (Cortex) Tumor | 18.9244846 | 20.0616829 | 38.9861675 |
| TCGA-HT-7693 | TCGA-LGG | Tumor TCGA | Oligodendroglion | Grade II | Brain (Cortex) Tumor | 20.624294 | 63.647439 | 84.2717329 |
| TCGA-HT-7694 | TCGA-LGG | Tumor TCGA | Oligodendroglion | Grade III | Brain (Cortex) Tumor | 10.3294395 | 28.2804711 | 38.6099106 |
| TCGA-HT-7695 | TCGA-LGG | Tumor TCGA | Oligodendroglion | Grade II | Brain (Cortex) Tumor | 13.4896319 | 20.1051268 | 33.5947587 |
| TCGA-HT-7854 | TCGA-LGG | Tumor TCGA | Astrocytoma | Grade II | Brain (Cortex) Tumor | 11.4175757 | 33.5243512 | 44.9419269 |
| TCGA-HT-7855 | TCGA-LGG | Tumor TCGA | Astrocytoma | Grade III | Brain (Cortex) Tumor | 9.95748576 | 30.0461671 | 40.0036529 |
| TCGA-HT-7856 | TCGA-LGG | Tumor TCGA | Oligodendroglion | Grade III | Brain (Cortex) Tumor | 9.67259902 | 19.8793125 | 29.5519115 |
| TCGA-HT-7857 | TCGA-LGG | Tumor TCGA | Astrocytoma | Grade III | Brain (Cortex) Tumor | 14.0240733 | 34.7737752 | 48.7978485 |
| TCGA-HT-7858 | TCGA-LGG | Tumor TCGA | Astrocytoma | Grade II | Brain (Cortex) Tumor | 15.1066669 | 33.163321 | 48.2699878 |
| TCGA-HT-7860 | TCGA-LGG | Tumor TCGA | Astrocytoma | Grade III | Brain (Cortex) Tumor | 20.7039665 | 31.6128778 | 52.3168443 |
| TCGA-HT-7873 | TCGA-LGG | Tumor TCGA | Oligoastrocytoma | Grade II | Brain (Cortex) Tumor | 18.7632362 | 33.7119483 | 52.4751845 |
| TCGA-HT-7874 | TCGA-LGG | Tumor TCGA | Oligodendroglion | Grade III | Brain (Cortex) Tumor | 12.9344556 | 22.1951218 | 35.1295774 |
| TCGA-HT-7875 | TCGA-LGG | Tumor TCGA | Oligodendroglion | Grade II | Brain (Cortex) Tumor | 16.0827524 | 18.1442277 | 34.2269801 |
| TCGA-HT-7877 | TCGA-LGG | Tumor TCGA | Oligodendroglion | Grade II | Brain (Cortex) Tumor | 16.1889163 | 18.5111089 | 34.7000252 |
| TCGA-HT-7879 | TCGA-LGG | Tumor TCGA | Oligoastrocytoma | Grade III | Brain (Cortex) Tumor | 13.105883 | 30.0119097 | 43.1177927 |
| TCGA-HT-7880 | TCGA-LGG | Tumor TCGA | Oligoastrocytoma | Grade II | Brain (Cortex) Tumor | 7.96510404 | 21.8638085 | 29.8289125 |
| TCGA-HT-7881 | TCGA-LGG | Tumor TCGA | Oligodendroglion | Grade II | Brain (Cortex) Tumor | 12.9966861 | 26.2025322 | 39.1992183 |
| TCGA-HT-7882 | TCGA-LGG | Tumor TCGA | Oligodendroglion | Grade III | Brain (Cortex) Tumor | 15.7579983 | 51.8754723 | 67.6334706 |
| TCGA-HT-7884 | TCGA-LGG | Tumor TCGA | Astrocytoma | Grade II | Brain (Cortex) Tumor | 14.6170745 | 27.836668 | 42.4537425 |
| TCGA-HT-7902 | TCGA-LGG | Tumor TCGA | Oligoastrocytoma | Grade II | Brain (Cortex) Tumor | 13.1432844 | 28.4133082 | 41.5565926 |
| TCGA-HT-8010 | TCGA-LGG | Tumor TCGA | Oligodendroglion | Grade II | Brain (Cortex) Tumor | 10.0118773 | 16.2909825 | 26.3028597 |
| TCGA-HT-8011 | TCGA-LGG | Tumor TCGA | Astrocytoma | Grade III | Brain (Cortex) Tumor | 32.2525877 | 70.9640692 | 103.216657 |
| TCGA-HT-8012 | TCGA-LGG | Tumor TCGA | Oligodendroglion | Grade II | Brain (Cortex) Tumor | 12.0051702 | 19.4761948 | 31.481365 |
| TCGA-HT-8013 | TCGA-LGG | Tumor TCGA | Oligoastrocytoma | Grade II | Brain (Cortex) Tumor | 15.3678791 | 39.2427116 | 54.6105906 |
| TCGA-HT-8015 | TCGA-LGG | Tumor TCGA | Astrocytoma | Grade II | Brain (Cortex) Tumor | 9.56707498 | 21.8453768 | 31.4124518 |
| TCGA-HT-8018 | TCGA-LGG | Tumor TCGA | Oligoastrocytoma | Grade II | Brain (Cortex) Tumor | 9.79493528 | 36.7741091 | 46.5690444 |
| TCGA-HT-8019 | TCGA-LGG | Tumor TCGA | Oligodendroglion | Grade III | Brain (Cortex) Tumor | 6.32150376 | 22.1486969 | 28.4702007 |
| TCGA-HT-8104 | TCGA-LGG | Tumor TCGA | Astrocytoma | Grade III | Brain (Cortex) Tumor | 23.0549791 | 54.2378467 | 77.2928259 |
| TCGA-HT-8105 | TCGA-LGG | Tumor TCGA | Oligodendroglion | Grade III | Brain (Cortex) Tumor | 18.3463852 | 23.9168492 | 42.2632344 |
| TCGA-HT-8106 | TCGA-LGG | Tumor TCGA | Astrocytoma | Grade III | Brain (Cortex) Tumor | 14.9212571 | 34.7092512 | 49.6305083 |

| | | | | | | | | |
|---|---|---|---|---|---|---|---|---|
| TCGA-HT-8107 | TCGA-LGG | Tumor TCGA | Oligodendroglion | Grade II | Brain (Cortex) Tumor | 6.13254946 | 20.2210477 | 26.3535972 |
| TCGA-HT-8108 | TCGA-LGG | Tumor TCGA | Oligodendroglion | Grade II | Brain (Cortex) Tumor | 16.4340605 | 27.5746637 | 44.0087242 |
| TCGA-HT-8109 | TCGA-LGG | Tumor TCGA | Oligodendroglion | Grade III | Brain (Cortex) Tumor | 12.3507747 | 17.4418952 | 29.7926699 |
| TCGA-HT-8110 | TCGA-LGG | Tumor TCGA | Astrocytoma | Grade III | Brain (Cortex) Tumor | 15.8455266 | 33.2080443 | 49.0535709 |
| TCGA-HT-8111 | TCGA-LGG | Tumor TCGA | Oligoastrocytoma | Grade III | Brain (Cortex) Tumor | 11.98299 | 33.1857636 | 45.1687536 |
| TCGA-HT-8113 | TCGA-LGG | Tumor TCGA | Oligodendroglion | Grade II | Brain (Cortex) Tumor | 9.21014242 | 25.2531219 | 34.4632644 |
| TCGA-HT-8114 | TCGA-LGG | Tumor TCGA | Oligoastrocytoma | Grade III | Brain (Cortex) Tumor | 10.5744176 | 31.3671224 | 41.94154 |
| TCGA-HT-8558 | TCGA-LGG | Tumor TCGA | Oligodendroglion | Grade II | Brain (Cortex) Tumor | 9.70543975 | 29.7583637 | 39.4638035 |
| TCGA-HT-8563 | TCGA-LGG | Tumor TCGA | Astrocytoma | Grade III | Brain (Cortex) Tumor | 16.1751588 | 40.7336955 | 56.9088543 |
| TCGA-HT-8564 | TCGA-LGG | Tumor TCGA | Astrocytoma | Grade III | Brain (Cortex) Tumor | 15.2269751 | 33.8115631 | 49.0385382 |
| TCGA-HT-A4DS | TCGA-LGG | Tumor TCGA | Astrocytoma | Grade III | Brain (Cortex) Tumor | 15.6623109 | 52.0859353 | 67.7482462 |
| TCGA-HT-A4DV | TCGA-LGG | Tumor TCGA | Oligodendroglion | Grade III | Brain (Cortex) Tumor | 17.5718367 | 23.6131371 | 41.1849738 |
| TCGA-HT-A5R5 | TCGA-LGG | Tumor TCGA | Oligodendroglion | Grade II | Brain (Cortex) Tumor | 9.65781755 | 31.7885208 | 41.4463383 |
| TCGA-HT-A5R7 | TCGA-LGG | Tumor TCGA | Astrocytoma | Grade III | Brain (Cortex) Tumor | 10.0261231 | 25.9712941 | 35.9974171 |
| TCGA-HT-A5R9 | TCGA-LGG | Tumor TCGA | Oligodendroglion | Grade III | Brain (Cortex) Tumor | 16.0980074 | 15.4922309 | 31.5902384 |
| TCGA-HT-A5RA | TCGA-LGG | Tumor TCGA | Astrocytoma | Grade III | Brain (Cortex) Tumor | 25.4420335 | 60.5256005 | 85.967634 |
| TCGA-HT-A5RB | TCGA-LGG | Tumor TCGA | Astrocytoma | Grade II | Brain (Cortex) Tumor | 11.9209377 | 25.8867166 | 37.8076543 |
| TCGA-HT-A5RC | TCGA-LGG | Tumor TCGA | Astrocytoma | Grade III | Brain (Cortex) Tumor | 14.8880252 | 38.8619962 | 53.7500214 |
| TCGA-HT-A614 | TCGA-LGG | Tumor TCGA | Oligoastrocytoma | Grade II | Brain (Cortex) Tumor | 31.8694445 | 39.9090799 | 71.7785244 |
| TCGA-HT-A615 | TCGA-LGG | Tumor TCGA | Oligodendroglion | Grade II | Brain (Cortex) Tumor | 22.7128404 | 27.4659116 | 50.1787519 |
| TCGA-HT-A616 | TCGA-LGG | Tumor TCGA | Astrocytoma | Grade II | Brain (Cortex) Tumor | 9.42099963 | 29.7442184 | 39.1652181 |
| TCGA-HT-A617 | TCGA-LGG | Tumor TCGA | Oligodendroglion | Grade II | Brain (Cortex) Tumor | 15.237286 | 57.6531193 | 72.8904052 |
| TCGA-HT-A618 | TCGA-LGG | Tumor TCGA | Astrocytoma | Grade III | Brain (Cortex) Tumor | 12.5662869 | 35.0007146 | 47.5670014 |
| TCGA-HT-A61A | TCGA-LGG | Tumor TCGA | Oligodendroglion | Grade II | Brain (Cortex) Tumor | 9.18911898 | 23.2091327 | 32.3982516 |
| TCGA-HT-A61B | TCGA-LGG | Tumor TCGA | Astrocytoma | Grade III | Brain (Cortex) Tumor | 25.9559048 | 63.1552275 | 89.1111322 |
| TCGA-HT-A61C | TCGA-LGG | Tumor TCGA | Oligodendroglion | Grade III | Brain (Cortex) Tumor | 45.8955811 | 43.1972859 | 89.092867 |
| TCGA-HT-A74H | TCGA-LGG | Tumor TCGA | Astrocytoma | Grade III | Brain (Cortex) Tumor | 9.95642448 | 41.1315892 | 51.0880136 |
| TCGA-HT-A74J | TCGA-LGG | Tumor TCGA | Oligoastrocytoma | Grade II | Brain (Cortex) Tumor | 8.58792551 | 29.9759145 | 38.56384 |
| TCGA-HT-A74K | TCGA-LGG | Tumor TCGA | Oligodendroglion | Grade III | Brain (Cortex) Tumor | 17.670427 | 25.9680202 | 43.6384472 |
| TCGA-HT-A74L | TCGA-LGG | Tumor TCGA | Oligoastrocytoma | Grade II | Brain (Cortex) Tumor | 15.1074994 | 21.4024021 | 36.5099015 |
| TCGA-HT-A74O | TCGA-LGG | Tumor TCGA | Astrocytoma | Grade III | Brain (Cortex) Tumor | 8.31902144 | 43.6621112 | 51.9811327 |
| TCGA-HW-7486 | TCGA-LGG | Tumor TCGA | Oligodendroglion | Grade II | Brain (Cortex) Tumor | 15.3774471 | 21.5885993 | 36.9660464 |
| TCGA-HW-7487 | TCGA-LGG | Tumor TCGA | Oligodendroglion | Grade II | Brain (Cortex) Tumor | 17.4782569 | 22.6683318 | 40.1465887 |
| TCGA-HW-7489 | TCGA-LGG | Tumor TCGA | Oligoastrocytoma | Grade II | Brain (Cortex) Tumor | 11.9375707 | 29.9225329 | 41.8601036 |
| TCGA-HW-7490 | TCGA-LGG | Tumor TCGA | Astrocytoma | Grade II | Brain (Cortex) Tumor | 13.2421929 | 27.3354559 | 40.5776488 |
| TCGA-HW-7491 | TCGA-LGG | Tumor TCGA | Oligodendroglion | Grade II | Brain (Cortex) Tumor | 19.5933009 | 18.1006321 | 37.693933 |
| TCGA-HW-7493 | TCGA-LGG | Tumor TCGA | Astrocytoma | Grade II | Brain (Cortex) Tumor | 14.7323227 | 33.1033635 | 47.8356862 |
| TCGA-HW-7495 | TCGA-LGG | Tumor TCGA | Oligodendroglion | Grade II | Brain (Cortex) Tumor | 13.6620069 | 20.1378857 | 33.7998926 |

| | | | | | | | | |
|---|---|---|---|---|---|---|---|---|
| TCGA-HW-8319 | TCGA-LGG | Tumor TCGA | Astrocytoma | Grade III | Brain (Cortex) Tumor | 10.8659544 | 35.5050209 | 46.3709753 |
| TCGA-HW-8320 | TCGA-LGG | Tumor TCGA | Astrocytoma | Grade III | Brain (Cortex) Tumor | 8.84714202 | 34.5900768 | 43.4372189 |
| TCGA-HW-8321 | TCGA-LGG | Tumor TCGA | Astrocytoma | Grade III | Brain (Cortex) Tumor | 9.95234815 | 26.5756575 | 36.5280057 |
| TCGA-HW-8322 | TCGA-LGG | Tumor TCGA | Oligodendroglion | Grade II | Brain (Cortex) Tumor | 18.623926 | 20.1093707 | 38.7332967 |
| TCGA-HW-A5KJ | TCGA-LGG | Tumor TCGA | Oligodendroglion | Grade III | Brain (Cortex) Tumor | 14.5682109 | 21.6358314 | 36.2040423 |
| TCGA-HW-A5KK | TCGA-LGG | Tumor TCGA | Astrocytoma | Grade III | Brain (Cortex) Tumor | 15.4219677 | 38.17539 | 53.5973578 |
| TCGA-HW-A5KL | TCGA-LGG | Tumor TCGA | Astrocytoma | Grade II | Brain (Cortex) Tumor | 8.00569319 | 19.4306393 | 27.4363325 |
| TCGA-HW-A5KM | TCGA-LGG | Tumor TCGA | Astrocytoma | Grade II | Brain (Cortex) Tumor | 24.5411503 | 39.1215333 | 63.6626836 |
| TCGA-IK-7675 | TCGA-LGG | Tumor TCGA | Oligodendroglion | Grade II | Brain (Cortex) Tumor | 16.1681719 | 44.915186 | 61.0833579 |
| TCGA-IK-8125 | TCGA-LGG | Tumor TCGA | Oligoastrocytoma | Grade III | Brain (Cortex) Tumor | 12.3494797 | 16.1212029 | 28.4706826 |
| TCGA-KT-A74X | TCGA-LGG | Tumor TCGA | Oligoastrocytoma | Grade III | Brain (Cortex) Tumor | 26.2379859 | 24.1685876 | 50.4065734 |
| TCGA-KT-A7W1 | TCGA-LGG | Tumor TCGA | Astrocytoma | Grade III | Brain (Cortex) Tumor | 43.7642377 | 59.9391433 | 103.703381 |
| TCGA-P5-A5ET | TCGA-LGG | Tumor TCGA | Oligodendroglion | Grade II | Brain (Cortex) Tumor | 17.3026869 | 19.1662096 | 36.4688965 |
| TCGA-P5-A5EU | TCGA-LGG | Tumor TCGA | Astrocytoma | Grade III | Brain (Cortex) Tumor | 13.3181124 | 57.3027803 | 70.6208927 |
| TCGA-P5-A5EV | TCGA-LGG | Tumor TCGA | Astrocytoma | Grade II | Brain (Cortex) Tumor | 13.9389941 | 35.2722887 | 49.2112828 |
| TCGA-P5-A5EW | TCGA-LGG | Tumor TCGA | Astrocytoma | Grade II | Brain (Cortex) Tumor | 12.2009606 | 32.0756119 | 44.2765725 |
| TCGA-P5-A5EX | TCGA-LGG | Tumor TCGA | Oligodendroglion | Grade III | Brain (Cortex) Tumor | 16.4998757 | 29.7799277 | 46.2798034 |
| TCGA-P5-A5EY | TCGA-LGG | Tumor TCGA | Astrocytoma | Grade II | Brain (Cortex) Tumor | 8.84443512 | 20.5671237 | 29.4115588 |
| TCGA-P5-A5EZ | TCGA-LGG | Tumor TCGA | Astrocytoma | Grade III | Brain (Cortex) Tumor | 10.9985912 | 36.4914102 | 47.4900014 |
| TCGA-P5-A5F0 | TCGA-LGG | Tumor TCGA | Oligodendroglion | Grade II | Brain (Cortex) Tumor | 13.1965625 | 18.2549561 | 31.4515186 |
| TCGA-P5-A5F1 | TCGA-LGG | Tumor TCGA | Astrocytoma | Grade II | Brain (Cortex) Tumor | 14.4805401 | 42.7185519 | 57.199092 |
| TCGA-P5-A5F2 | TCGA-LGG | Tumor TCGA | Astrocytoma | Grade II | Brain (Cortex) Tumor | 14.0376714 | 41.6781699 | 55.7158413 |
| TCGA-P5-A5F4 | TCGA-LGG | Tumor TCGA | Oligodendroglion | Grade III | Brain (Cortex) Tumor | 12.9515462 | 35.6994433 | 48.6509896 |
| TCGA-P5-A5F6 | TCGA-LGG | Tumor TCGA | Oligodendroglion | Grade II | Brain (Cortex) Tumor | 8.00697639 | 34.2111117 | 42.2180881 |
| TCGA-P5-A72U | TCGA-LGG | Tumor TCGA | Oligodendroglion | Grade III | Brain (Cortex) Tumor | 12.5447491 | 45.0740383 | 57.6187874 |
| TCGA-P5-A72W | TCGA-LGG | Tumor TCGA | Astrocytoma | Grade III | Brain (Cortex) Tumor | 15.6554272 | 47.9224094 | 63.5778365 |
| TCGA-P5-A72X | TCGA-LGG | Tumor TCGA | Astrocytoma | Grade III | Brain (Cortex) Tumor | 6.52660084 | 32.4659411 | 38.9925419 |
| TCGA-P5-A72Z | TCGA-LGG | Tumor TCGA | Oligodendroglion | Grade III | Brain (Cortex) Tumor | 21.1771625 | 20.6172374 | 41.7943998 |
| TCGA-P5-A731 | TCGA-LGG | Tumor TCGA | Oligoastrocytoma | Grade II | Brain (Cortex) Tumor | 13.4975304 | 41.4845093 | 54.9820397 |
| TCGA-P5-A733 | TCGA-LGG | Tumor TCGA | Astrocytoma | Grade II | Brain (Cortex) Tumor | 8.62156778 | 23.7361432 | 32.357711 |
| TCGA-P5-A735 | TCGA-LGG | Tumor TCGA | Astrocytoma | Grade II | Brain (Cortex) Tumor | 11.0083484 | 42.843004 | 53.8513524 |
| TCGA-P5-A736 | TCGA-LGG | Tumor TCGA | Astrocytoma | Grade III | Brain (Cortex) Tumor | 8.95043384 | 40.8984067 | 49.8488406 |
| TCGA-P5-A737 | TCGA-LGG | Tumor TCGA | Oligoastrocytoma | Grade II | Brain (Cortex) Tumor | 17.5534067 | 17.9065969 | 35.4600036 |
| TCGA-P5-A77W | TCGA-LGG | Tumor TCGA | Oligoastrocytoma | Grade III | Brain (Cortex) Tumor | 24.9525828 | 17.9803793 | 42.9329621 |
| TCGA-P5-A77X | TCGA-LGG | Tumor TCGA | Oligoastrocytoma | Grade II | Brain (Cortex) Tumor | 24.7377256 | 21.4510167 | 46.1887423 |
| TCGA-P5-A780 | TCGA-LGG | Tumor TCGA | Astrocytoma | Grade III | Brain (Cortex) Tumor | 9.97089599 | 35.7842526 | 45.7551486 |
| TCGA-P5-A781 | TCGA-LGG | Tumor TCGA | Astrocytoma | Grade III | Brain (Cortex) Tumor | 16.719951 | 24.8842876 | 41.6042385 |
| TCGA-QH-A65R | TCGA-LGG | Tumor TCGA | Oligodendroglion | Grade III | Brain (Cortex) Tumor | 13.2123902 | 18.0725305 | 31.2849206 |

| | | | | | | | | |
|---|---|---|---|---|---|---|---|---|
| TCGA-QH-A65S | TCGA-LGG | Tumor TCGA | Oligoastrocytoma | Grade II | Brain (Cortex) Tumor | 10.8372128 | 29.7167417 | 40.5539545 |
| TCGA-QH-A65V | TCGA-LGG | Tumor TCGA | Oligodendroglion | Grade II | Brain (Cortex) Tumor | 18.3830828 | 25.2394508 | 43.6225336 |
| TCGA-QH-A65X | TCGA-LGG | Tumor TCGA | Oligoastrocytoma | Grade III | Brain (Cortex) Tumor | 18.4152415 | 27.369649 | 45.7848904 |
| TCGA-QH-A65Z | TCGA-LGG | Tumor TCGA | Oligodendroglion | Grade II | Brain (Cortex) Tumor | 21.32758 | 24.6757357 | 46.0033156 |
| TCGA-QH-A6CS | TCGA-LGG | Tumor TCGA | Astrocytoma | Grade III | Brain (Cortex) Tumor | 13.5490839 | 36.2938749 | 49.8429588 |
| TCGA-QH-A6CU | TCGA-LGG | Tumor TCGA | Oligodendroglion | Grade III | Brain (Cortex) Tumor | 16.3256719 | 23.2370455 | 39.5627174 |
| TCGA-QH-A6CV | TCGA-LGG | Tumor TCGA | Oligoastrocytoma | Grade III | Brain (Cortex) Tumor | 37.6126812 | 66.0320053 | 103.644686 |
| TCGA-QH-A6CW | TCGA-LGG | Tumor TCGA | Oligoastrocytoma | Grade III | Brain (Cortex) Tumor | 6.76987973 | 36.0809507 | 42.8508305 |
| TCGA-QH-A6CX | TCGA-LGG | Tumor TCGA | Astrocytoma | Grade II | Brain (Cortex) Tumor | 11.7687303 | 35.1670411 | 46.9357714 |
| TCGA-QH-A6CY | TCGA-LGG | Tumor TCGA | Oligoastrocytoma | Grade III | Brain (Cortex) Tumor | 21.7366796 | 27.9810034 | 49.717683 |
| TCGA-QH-A6CZ | TCGA-LGG | Tumor TCGA | Oligoastrocytoma | Grade II | Brain (Cortex) Tumor | 25.02377 | 23.0487036 | 48.0724736 |
| TCGA-QH-A6X3 | TCGA-LGG | Tumor TCGA | Oligoastrocytoma | Grade II | Brain (Cortex) Tumor | 16.4496977 | 42.6932625 | 59.1429602 |
| TCGA-QH-A6X4 | TCGA-LGG | Tumor TCGA | Oligoastrocytoma | Grade III | Brain (Cortex) Tumor | 17.9401402 | 22.2815494 | 40.2216895 |
| TCGA-QH-A6X5 | TCGA-LGG | Tumor TCGA | Oligoastrocytoma | Grade II | Brain (Cortex) Tumor | 19.1572078 | 23.3745746 | 42.5317824 |
| TCGA-QH-A6X8 | TCGA-LGG | Tumor TCGA | Oligodendroglion | Grade III | Brain (Cortex) Tumor | 15.9519241 | 27.0195536 | 42.9714777 |
| TCGA-QH-A6X9 | TCGA-LGG | Tumor TCGA | Oligodendroglion | Grade II | Brain (Cortex) Tumor | 11.1933623 | 34.3832546 | 45.5766168 |
| TCGA-QH-A6XA | TCGA-LGG | Tumor TCGA | Oligoastrocytoma | Grade II | Brain (Cortex) Tumor | 13.6501395 | 38.3778618 | 52.0280013 |
| TCGA-QH-A6XC | TCGA-LGG | Tumor TCGA | Astrocytoma | Grade III | Brain (Cortex) Tumor | 18.6491586 | 53.6668836 | 72.3160421 |
| TCGA-QH-A86X | TCGA-LGG | Tumor TCGA | Oligodendroglion | Grade II | Brain (Cortex) Tumor | 17.1119632 | 20.4054107 | 37.5173739 |
| TCGA-QH-A870 | TCGA-LGG | Tumor TCGA | Oligoastrocytoma | Grade III | Brain (Cortex) Tumor | 14.4887289 | 47.4231656 | 61.9118945 |
| TCGA-R8-A6MK | TCGA-LGG | Tumor TCGA | Oligodendroglion | Grade II | Brain (Cortex) Tumor | 23.5102952 | 26.2264901 | 49.7367853 |
| TCGA-R8-A6ML | TCGA-LGG | Tumor TCGA | Oligodendroglion | Grade III | Brain (Cortex) Tumor | 21.5003129 | 20.1133745 | 41.6136874 |
| TCGA-R8-A6MO | TCGA-LGG | Tumor TCGA | Oligodendroglion | Grade II | Brain (Cortex) Tumor | 27.2430052 | 28.7734972 | 56.0165024 |
| TCGA-RY-A83X | TCGA-LGG | Tumor TCGA | Oligodendroglion | Grade II | Brain (Cortex) Tumor | 17.5653648 | 21.4285377 | 38.9939025 |
| TCGA-RY-A83Y | TCGA-LGG | Tumor TCGA | Oligodendroglion | Grade II | Brain (Cortex) Tumor | 16.7829804 | 19.9266165 | 36.7095969 |
| TCGA-RY-A83Z | TCGA-LGG | Tumor TCGA | Astrocytoma | Grade III | Brain (Cortex) Tumor | 22.6375716 | 85.6073916 | 108.244963 |
| TCGA-RY-A840 | TCGA-LGG | Tumor TCGA | Oligodendroglion | Grade III | Brain (Cortex) Tumor | 23.7259564 | 17.8104228 | 41.5363793 |
| TCGA-RY-A843 | TCGA-LGG | Tumor TCGA | Astrocytoma | Grade III | Brain (Cortex) Tumor | 11.3645102 | 30.6389701 | 42.0034802 |
| TCGA-RY-A845 | TCGA-LGG | Tumor TCGA | Oligoastrocytoma | Grade II | Brain (Cortex) Tumor | 14.8618061 | 32.9889052 | 47.8507113 |
| TCGA-RY-A847 | TCGA-LGG | Tumor TCGA | Oligodendroglion | Grade II | Brain (Cortex) Tumor | 16.5687824 | 22.0579772 | 38.6267596 |
| TCGA-S9-A6TS | TCGA-LGG | Tumor TCGA | Astrocytoma | Grade III | Brain (Cortex) Tumor | 12.135383 | 33.9465154 | 46.0818984 |
| TCGA-S9-A6TU | TCGA-LGG | Tumor TCGA | Astrocytoma | Grade II | Brain (Cortex) Tumor | 5.90851615 | 32.3970805 | 38.3055966 |
| TCGA-S9-A6TV | TCGA-LGG | Tumor TCGA | Oligoastrocytoma | Grade III | Brain (Cortex) Tumor | 22.5990618 | 69.1099598 | 91.7090216 |
| TCGA-S9-A6TW | TCGA-LGG | Tumor TCGA | Oligodendroglion | Grade III | Brain (Cortex) Tumor | 13.2958982 | 18.5032545 | 31.7991527 |
| TCGA-S9-A6TX | TCGA-LGG | Tumor TCGA | Oligodendroglion | Grade III | Brain (Cortex) Tumor | 14.1408859 | 21.0430923 | 35.1839782 |
| TCGA-S9-A6TY | TCGA-LGG | Tumor TCGA | Oligodendroglion | Grade II | Brain (Cortex) Tumor | 21.4692804 | 19.8312273 | 41.3005077 |
| TCGA-S9-A6TZ | TCGA-LGG | Tumor TCGA | Astrocytoma | Grade II | Brain (Cortex) Tumor | 15.0209201 | 45.6996724 | 60.7205924 |
| TCGA-S9-A6U0 | TCGA-LGG | Tumor TCGA | Astrocytoma | Grade III | Brain (Cortex) Tumor | 16.9772013 | 49.7358665 | 66.7130678 |

| | | | | | | | | |
|---|---|---|---|---|---|---|---|---|
| TCGA-S9-A6U1 | TCGA-LGG | Tumor TCGA | Astrocytoma | Grade III | Brain (Cortex) Tumor | 18.3877588 | 40.7196753 | 59.1074341 |
| TCGA-S9-A6U2 | TCGA-LGG | Tumor TCGA | Oligodendroglion | Grade II | Brain (Cortex) Tumor | 23.7857838 | 22.9117773 | 46.6975611 |
| TCGA-S9-A6U5 | TCGA-LGG | Tumor TCGA | Astrocytoma | Grade II | Brain (Cortex) Tumor | 21.8856456 | 23.8131237 | 45.6987694 |
| TCGA-S9-A6U6 | TCGA-LGG | Tumor TCGA | Astrocytoma | Grade III | Brain (Cortex) Tumor | 15.9536714 | 40.5785484 | 56.5322198 |
| TCGA-S9-A6U8 | TCGA-LGG | Tumor TCGA | Astrocytoma | Grade II | Brain (Cortex) Tumor | 17.9624947 | 37.3873741 | 55.3498688 |
| TCGA-S9-A6U9 | TCGA-LGG | Tumor TCGA | Astrocytoma | Grade III | Brain (Cortex) Tumor | 9.67809066 | 40.8325999 | 50.5106906 |
| TCGA-S9-A6UA | TCGA-LGG | Tumor TCGA | Astrocytoma | Grade III | Brain (Cortex) Tumor | 16.0652067 | 41.1456635 | 57.2108703 |
| TCGA-S9-A6UB | TCGA-LGG | Tumor TCGA | Oligodendroglion | Grade II | Brain (Cortex) Tumor | 36.7373054 | 23.7103854 | 60.4476908 |
| TCGA-S9-A6WD | TCGA-LGG | Tumor TCGA | Oligodendroglion | Grade III | Brain (Cortex) Tumor | 20.5664426 | 22.164928 | 42.7313707 |
| TCGA-S9-A6WE | TCGA-LGG | Tumor TCGA | Oligodendroglion | Grade II | Brain (Cortex) Tumor | 20.0846152 | 21.5589545 | 41.6435697 |
| TCGA-S9-A6WG | TCGA-LGG | Tumor TCGA | Astrocytoma | Grade III | Brain (Cortex) Tumor | 15.6320302 | 38.2929574 | 53.9249875 |
| TCGA-S9-A6WH | TCGA-LGG | Tumor TCGA | Oligoastrocytoma | Grade II | Brain (Cortex) Tumor | 26.1791319 | 25.8937676 | 52.0728995 |
| TCGA-S9-A6WI | TCGA-LGG | Tumor TCGA | Oligoastrocytoma | Grade II | Brain (Cortex) Tumor | 13.4312868 | 28.5892228 | 42.0205096 |
| TCGA-S9-A6WL | TCGA-LGG | Tumor TCGA | Astrocytoma | Grade III | Brain (Cortex) Tumor | 15.5426326 | 21.4358496 | 36.9784822 |
| TCGA-S9-A6WM | TCGA-LGG | Tumor TCGA | Astrocytoma | Grade III | Brain (Cortex) Tumor | 24.0651051 | 79.6430987 | 103.708204 |
| TCGA-S9-A6WN | TCGA-LGG | Tumor TCGA | Astrocytoma | Grade III | Brain (Cortex) Tumor | 14.9170281 | 25.2927024 | 40.2097306 |
| TCGA-S9-A6WO | TCGA-LGG | Tumor TCGA | Astrocytoma | Grade II | Brain (Cortex) Tumor | 15.8606267 | 32.2477758 | 48.1084025 |
| TCGA-S9-A6WP | TCGA-LGG | Tumor TCGA | Oligoastrocytoma | Grade III | Brain (Cortex) Tumor | 24.0796554 | 25.7689447 | 49.8486001 |
| TCGA-S9-A6WQ | TCGA-LGG | Tumor TCGA | Oligoastrocytoma | Grade II | Brain (Cortex) Tumor | 13.7850601 | 31.4566037 | 45.2416638 |
| TCGA-S9-A7IQ | TCGA-LGG | Tumor TCGA | Oligoastrocytoma | Grade II | Brain (Cortex) Tumor | 13.7172639 | 28.7444987 | 42.4617626 |
| TCGA-S9-A7IS | TCGA-LGG | Tumor TCGA | Astrocytoma | Grade III | Brain (Cortex) Tumor | 28.0574344 | 66.9158883 | 94.9733227 |
| TCGA-S9-A7IX | TCGA-LGG | Tumor TCGA | Astrocytoma | Grade III | Brain (Cortex) Tumor | 19.0079387 | 42.4232142 | 61.4311529 |
| TCGA-S9-A7IY | TCGA-LGG | Tumor TCGA | Oligoastrocytoma | Grade III | Brain (Cortex) Tumor | 19.0027212 | 18.3442422 | 37.3469634 |
| TCGA-S9-A7IZ | TCGA-LGG | Tumor TCGA | Astrocytoma | Grade III | Brain (Cortex) Tumor | 12.1552679 | 31.8280461 | 43.983314 |
| TCGA-S9-A7J0 | TCGA-LGG | Tumor TCGA | Oligodendroglion | Grade III | Brain (Cortex) Tumor | 83.7868366 | 86.8560275 | 170.642864 |
| TCGA-S9-A7J1 | TCGA-LGG | Tumor TCGA | Oligodendroglion | Grade II | Brain (Cortex) Tumor | 26.1563499 | 19.714072 | 45.8704219 |
| TCGA-S9-A7J2 | TCGA-LGG | Tumor TCGA | Oligodendroglion | Grade III | Brain (Cortex) Tumor | 21.5668004 | 20.4036008 | 41.9704011 |
| TCGA-S9-A7J3 | TCGA-LGG | Tumor TCGA | Oligodendroglion | Grade III | Brain (Cortex) Tumor | 22.0254493 | 19.6231531 | 41.6486025 |
| TCGA-S9-A7QW | TCGA-LGG | Tumor TCGA | Astrocytoma | Grade III | Brain (Cortex) Tumor | 15.1632566 | 37.471513 | 52.6347696 |
| TCGA-S9-A7QX | TCGA-LGG | Tumor TCGA | Astrocytoma | Grade III | Brain (Cortex) Tumor | 12.6044506 | 28.0050498 | 40.6095005 |
| TCGA-S9-A7QY | TCGA-LGG | Tumor TCGA | Oligoastrocytoma | Grade II | Brain (Cortex) Tumor | 16.1291726 | 19.9454789 | 36.0746515 |
| TCGA-S9-A7QZ | TCGA-LGG | Tumor TCGA | Oligodendroglion | Grade II | Brain (Cortex) Tumor | 19.3236567 | 20.0682666 | 39.3919234 |
| TCGA-S9-A7R1 | TCGA-LGG | Tumor TCGA | Oligodendroglion | Grade II | Brain (Cortex) Tumor | 13.782367 | 21.5846858 | 35.3670528 |
| TCGA-S9-A7R2 | TCGA-LGG | Tumor TCGA | Astrocytoma | Grade III | Brain (Cortex) Tumor | 14.4487985 | 47.4400501 | 61.8888486 |
| TCGA-S9-A7R3 | TCGA-LGG | Tumor TCGA | Astrocytoma | Grade II | Brain (Cortex) Tumor | 14.3406792 | 35.5249824 | 49.8656616 |
| TCGA-S9-A7R4 | TCGA-LGG | Tumor TCGA | Astrocytoma | Grade III | Brain (Cortex) Tumor | 10.2125798 | 31.5724937 | 41.7850735 |
| TCGA-S9-A7R7 | TCGA-LGG | Tumor TCGA | Astrocytoma | Grade II | Brain (Cortex) Tumor | 18.5981686 | 38.2762277 | 56.8743963 |
| TCGA-S9-A7R8 | TCGA-LGG | Tumor TCGA | Astrocytoma | Grade III | Brain (Cortex) Tumor | 10.7402261 | 38.5023937 | 49.2426198 |

| | | | | | | | | |
|---|---|---|---|---|---|---|---|---|
| TCGA-S9-A89V | TCGA-LGG | Tumor TCGA | Astrocytoma | Grade III | Brain (Cortex) Tumor | 12.5144182 | 36.1460395 | 48.6604577 |
| TCGA-S9-A89Z | TCGA-LGG | Tumor TCGA | Astrocytoma | Grade III | Brain (Cortex) Tumor | 20.4632626 | 57.3656395 | 77.828902 |
| TCGA-TM-A7C3 | TCGA-LGG | Tumor TCGA | Astrocytoma | Grade III | Brain (Cortex) Tumor | 22.8521932 | 73.6814568 | 96.53365 |
| TCGA-TM-A7C4 | TCGA-LGG | Tumor TCGA | Astrocytoma | Grade II | Brain (Cortex) Tumor | 11.8059264 | 33.8612226 | 45.6671489 |
| TCGA-TM-A7C5 | TCGA-LGG | Tumor TCGA | Oligoastrocytoma | Grade II | Brain (Cortex) Tumor | 18.8592219 | 22.5233308 | 41.3825526 |
| TCGA-TM-A7CA | TCGA-LGG | Tumor TCGA | Astrocytoma | Grade II | Brain (Cortex) Tumor | 13.1472209 | 30.0251448 | 43.1723657 |
| TCGA-TM-A7CF | TCGA-LGG | Tumor TCGA | Astrocytoma | Grade II | Brain (Cortex) Tumor | 12.369695 | 32.7140985 | 45.0837935 |
| TCGA-TM-A84B | TCGA-LGG | Tumor TCGA | Astrocytoma | Grade III | Brain (Cortex) Tumor | 23.4452075 | 58.1964443 | 81.6416518 |
| TCGA-TM-A84C | TCGA-LGG | Tumor TCGA | Astrocytoma | Grade II | Brain (Cortex) Tumor | 16.794356 | 38.1435949 | 54.9379509 |
| TCGA-TM-A84F | TCGA-LGG | Tumor TCGA | Astrocytoma | Grade III | Brain (Cortex) Tumor | 14.8474341 | 32.8826813 | 47.7301154 |
| TCGA-TM-A84G | TCGA-LGG | Tumor TCGA | Oligodendroglion | Grade III | Brain (Cortex) Tumor | 25.9143459 | 24.0614504 | 49.9757964 |
| TCGA-TM-A84H | TCGA-LGG | Tumor TCGA | Oligoastrocytoma | Grade III | Brain (Cortex) Tumor | 15.6803328 | 33.3505347 | 49.0308675 |
| TCGA-TM-A84I | TCGA-LGG | Tumor TCGA | Astrocytoma | Grade III | Brain (Cortex) Tumor | 27.5922957 | 69.0135342 | 96.6058299 |
| TCGA-TM-A84J | TCGA-LGG | Tumor TCGA | Oligodendroglion | Grade III | Brain (Cortex) Tumor | 24.9196051 | 49.0698326 | 73.9894377 |
| TCGA-TM-A84L | TCGA-LGG | Tumor TCGA | Oligoastrocytoma | Grade II | Brain (Cortex) Tumor | 11.017391 | 28.9370355 | 39.9544265 |
| TCGA-TM-A84M | TCGA-LGG | Tumor TCGA | Oligodendroglion | Grade III | Brain (Cortex) Tumor | 29.7515868 | 18.6965058 | 48.4480926 |
| TCGA-TM-A84O | TCGA-LGG | Tumor TCGA | Oligodendroglion | Grade III | Brain (Cortex) Tumor | 19.8919343 | 23.4433957 | 43.3353299 |
| TCGA-TM-A84Q | TCGA-LGG | Tumor TCGA | Astrocytoma | Grade II | Brain (Cortex) Tumor | 12.6462585 | 41.8349735 | 54.481232 |
| TCGA-TM-A84R | TCGA-LGG | Tumor TCGA | Oligodendroglion | Grade II | Brain (Cortex) Tumor | 11.882248 | 31.0150614 | 42.8973094 |
| TCGA-TM-A84S | TCGA-LGG | Tumor TCGA | Oligodendroglion | Grade III | Brain (Cortex) Tumor | 20.5906713 | 20.585086 | 41.1757574 |
| TCGA-TM-A84T | TCGA-LGG | Tumor TCGA | Oligoastrocytoma | Grade II | Brain (Cortex) Tumor | 11.093596 | 29.6029583 | 40.6965544 |
| TCGA-TQ-A7RF | TCGA-LGG | Tumor TCGA | Oligodendroglion | Grade III | Brain (Cortex) Tumor | 13.1229631 | 60.4594819 | 73.582445 |
| TCGA-TQ-A7RG | TCGA-LGG | Tumor TCGA | Oligoastrocytoma | Grade II | Brain (Cortex) Tumor | 20.1884858 | 23.4114718 | 43.5999576 |
| TCGA-TQ-A7RH | TCGA-LGG | Tumor TCGA | Oligoastrocytoma | Grade II | Brain (Cortex) Tumor | 14.9652186 | 31.0983127 | 46.0635312 |
| TCGA-TQ-A7RI | TCGA-LGG | Tumor TCGA | Oligodendroglion | Grade II | Brain (Cortex) Tumor | 23.2167267 | 19.518259 | 42.7349856 |
| TCGA-TQ-A7RJ | TCGA-LGG | Tumor TCGA | Oligoastrocytoma | Grade II | Brain (Cortex) Tumor | 13.9387361 | 33.625406 | 47.5641421 |
| TCGA-TQ-A7RK | TCGA-LGG | Tumor TCGA | Oligoastrocytoma | Grade II | Brain (Cortex) Tumor | 20.3100035 | 47.4642383 | 67.7742418 |
| TCGA-TQ-A7RM | TCGA-LGG | Tumor TCGA | Oligoastrocytoma | Grade III | Brain (Cortex) Tumor | 40.70367 | 73.2157195 | 113.919389 |
| TCGA-TQ-A7RN | TCGA-LGG | Tumor TCGA | Oligodendroglion | Grade II | Brain (Cortex) Tumor | 22.1531447 | 19.5045702 | 41.6577148 |
| TCGA-TQ-A7RO | TCGA-LGG | Tumor TCGA | Oligoastrocytoma | Grade II | Brain (Cortex) Tumor | 16.7037269 | 20.0571376 | 36.7608644 |
| TCGA-TQ-A7RP | TCGA-LGG | Tumor TCGA | Oligoastrocytoma | Grade II | Brain (Cortex) Tumor | 14.3504366 | 34.2926449 | 48.6430815 |
| TCGA-TQ-A7RQ | TCGA-LGG | Tumor TCGA | Oligodendroglion | Grade II | Brain (Cortex) Tumor | 17.6111188 | 18.8280978 | 36.4392167 |
| TCGA-TQ-A7RR | TCGA-LGG | Tumor TCGA | Oligoastrocytoma | Grade II | Brain (Cortex) Tumor | 18.7204568 | 39.3020057 | 58.0224625 |
| TCGA-TQ-A7RS | TCGA-LGG | Tumor TCGA | Oligodendroglion | Grade II | Brain (Cortex) Tumor | 23.6428073 | 24.7100133 | 48.3528205 |
| TCGA-TQ-A7RV | TCGA-LGG | Tumor TCGA | Astrocytoma | Grade II | Brain (Cortex) Tumor | 15.7558543 | 31.326387 | 47.0822414 |
| TCGA-TQ-A7RW | TCGA-LGG | Tumor TCGA | Oligodendroglion | Grade II | Brain (Cortex) Tumor | 13.4501073 | 35.5992324 | 49.0493398 |
| TCGA-TQ-A8XE | TCGA-LGG | Tumor TCGA | Oligodendroglion | Grade II | Brain (Cortex) Tumor | 15.4100332 | 45.4499052 | 60.8599384 |
| TCGA-VM-A8C8 | TCGA-LGG | Tumor TCGA | Oligodendroglion | Grade II | Brain (Cortex) Tumor | 15.6807662 | 44.5259539 | 60.2067201 |

| | | | | | | | | |
|---|---|---|---|---|---|---|---|---|
| TCGA-VM-A8C9 | TCGA-LGG | Tumor TCGA | Astrocytoma | Grade II | Brain (Cortex) Tumor | 16.774803 | 37.9611844 | 54.7359874 |
| TCGA-VM-A8CA | TCGA-LGG | Tumor TCGA | Oligodendroglion | Grade II | Brain (Cortex) Tumor | 18.3005831 | 33.9434123 | 52.2439954 |
| TCGA-VM-A8CB | TCGA-LGG | Tumor TCGA | Oligodendroglion | Grade III | Brain (Cortex) Tumor | 42.3758194 | 26.8832491 | 69.2590686 |
| TCGA-VM-A8CD | TCGA-LGG | Tumor TCGA | Astrocytoma | Grade III | Brain (Cortex) Tumor | 25.4003394 | 46.4578499 | 71.8581893 |
| TCGA-VM-A8CE | TCGA-LGG | Tumor TCGA | Oligodendroglion | Grade II | Brain (Cortex) Tumor | 14.778874 | 22.3478377 | 37.1267117 |
| TCGA-VM-A8CF | TCGA-LGG | Tumor TCGA | Astrocytoma | Grade III | Brain (Cortex) Tumor | 15.3758878 | 42.2324074 | 57.6082952 |
| TCGA-VM-A8CH | TCGA-LGG | Tumor TCGA | Astrocytoma | Grade II | Brain (Cortex) Tumor | 10.0674496 | 32.4688469 | 42.5362965 |
| TCGA-VV-A829 | TCGA-LGG | Tumor TCGA | Oligoastrocytoma | Grade III | Brain (Cortex) Tumor | 19.7793874 | 20.6146998 | 40.3940872 |
| TCGA-VV-A86M | TCGA-LGG | Tumor TCGA | Astrocytoma | Grade III | Brain (Cortex) Tumor | 20.9263302 | 44.0663098 | 64.9926399 |
| TCGA-VW-A7QS | TCGA-LGG | Tumor TCGA | Oligodendroglion | Grade III | Brain (Cortex) Tumor | 29.8747318 | 31.6143916 | 61.4891233 |
| TCGA-VW-A8FI | TCGA-LGG | Tumor TCGA | Astrocytoma | Grade III | Brain (Cortex) Tumor | 23.9786402 | 48.018556 | 71.9971962 |
| TCGA-W9-A837 | TCGA-LGG | Tumor TCGA | Oligodendroglion | Grade II | Brain (Cortex) Tumor | 17.925015 | 25.3676089 | 43.2926239 |
| TCGA-WH-A86K | TCGA-LGG | Tumor TCGA | Astrocytoma | Grade II | Brain (Cortex) Tumor | 13.5996533 | 35.905896 | 49.5055493 |
| TCGA-WY-A858 | TCGA-LGG | Tumor TCGA | Astrocytoma | Grade III | Brain (Cortex) Tumor | 9.68398255 | 30.0926001 | 39.7765827 |
| TCGA-WY-A859 | TCGA-LGG | Tumor TCGA | Astrocytoma | Grade II | Brain (Cortex) Tumor | 17.9341407 | 50.1893762 | 68.1235169 |
| TCGA-WY-A85A | TCGA-LGG | Tumor TCGA | Astrocytoma | Grade II | Brain (Cortex) Tumor | 12.6786829 | 30.5549601 | 43.233643 |
| TCGA-WY-A85B | TCGA-LGG | Tumor TCGA | Astrocytoma | Grade II | Brain (Cortex) Tumor | 22.0448926 | 44.7251649 | 66.7700575 |
| TCGA-WY-A85C | TCGA-LGG | Tumor TCGA | Astrocytoma | Grade II | Brain (Cortex) Tumor | 9.70418012 | 29.0328302 | 38.7370103 |
| TCGA-WY-A85D | TCGA-LGG | Tumor TCGA | Oligoastrocytoma | Grade II | Brain (Cortex) Tumor | 17.3610108 | 50.8747776 | 68.2357885 |
| TCGA-WY-A85E | TCGA-LGG | Tumor TCGA | Oligoastrocytoma | Grade II | Brain (Cortex) Tumor | 18.9927782 | 38.4201144 | 57.4128926 |
| GTEX-1117F-3226-SI | TCGA-LGG | GTEx | Normal | Cortex | Brain (Cortex) | 4.25 | 15.76 | 20.01 |
| GTEX-111FC-3126-S | TCGA-LGG | GTEx | Normal | Cortex | Brain (Cortex) | 6.9 | 21.71 | 28.61 |
| GTEX-1128S-2726-S | TCGA-LGG | GTEx | Normal | Cortex | Brain (Cortex) | 5.25 | 21.08 | 26.33 |
| GTEX-117XS-3026-S | TCGA-LGG | GTEx | Normal | Cortex | Brain (Cortex) | 5.51 | 25.59 | 31.1 |
| GTEX-1192X-3126-S | TCGA-LGG | GTEx | Normal | Cortex | Brain (Cortex) | 8.67 | 30.48 | 39.15 |
| GTEX-11DXW-1126-S | TCGA-LGG | GTEx | Normal | Cortex | Brain (Cortex) | 4.94 | 18.96 | 23.9 |
| GTEX-11DXY-3226-S | TCGA-LGG | GTEx | Normal | Cortex | Brain (Cortex) | 4.97 | 27.54 | 32.51 |
| GTEX-11EI6-3026-SI | TCGA-LGG | GTEx | Normal | Cortex | Brain (Cortex) | 4.44 | 20.1 | 24.54 |
| GTEX-11EMC-3226-S | TCGA-LGG | GTEx | Normal | Cortex | Brain (Cortex) | 5.06 | 14.14 | 19.2 |
| GTEX-11GS4-3126-S | TCGA-LGG | GTEx | Normal | Cortex | Brain (Cortex) | 6.1 | 15.96 | 22.06 |
| GTEX-11GSO-2926-S | TCGA-LGG | GTEx | Normal | Cortex | Brain (Cortex) | 6.84 | 24.7 | 31.54 |
| GTEX-11GSP-3226-S | TCGA-LGG | GTEx | Normal | Cortex | Brain (Cortex) | 5.64 | 14.82 | 20.46 |
| GTEX-11NUK-2926-S | TCGA-LGG | GTEx | Normal | Cortex | Brain (Cortex) | 7 | 23.28 | 30.28 |
| GTEX-11NV4-2126-S | TCGA-LGG | GTEx | Normal | Cortex | Brain (Cortex) | 4.26 | 9.67 | 13.93 |
| GTEX-11O72-2926-S | TCGA-LGG | GTEx | Normal | Cortex | Brain (Cortex) | 5.02 | 21.28 | 26.3 |
| GTEX-11OC5-0726-S | TCGA-LGG | GTEx | Normal | Cortex | Brain (Cortex) | 7.96 | 32.86 | 40.82 |
| GTEX-11OF3-2926-S | TCGA-LGG | GTEx | Normal | Cortex | Brain (Cortex) | 7.48 | 23.01 | 30.49 |
| GTEX-11ONC-2926-S | TCGA-LGG | GTEx | Normal | Cortex | Brain (Cortex) | 6.04 | 23.73 | 29.77 |

| | | | | | | | | |
|---|---|---|---|---|---|---|---|---|
| GTEX-11PRG-2926-S | TCGA-LGG | GTEx | Normal | Cortex | Brain (Cortex) | 6.62 | 18.79 | 25.41 |
| GTEX-11TTK-2926-S | TCGA-LGG | GTEx | Normal | Cortex | Brain (Cortex) | 4.1 | 15.14 | 19.24 |
| GTEX-11WQK-3026- | TCGA-LGG | GTEx | Normal | Cortex | Brain (Cortex) | 6.48 | 19.68 | 26.16 |
| GTEX-11ZUS-2926-S | TCGA-LGG | GTEx | Normal | Cortex | Brain (Cortex) | 4.5 | 20.48 | 24.98 |
| GTEX-12126-1026-S | TCGA-LGG | GTEx | Normal | Cortex | Brain (Cortex) | 5.63 | 18.74 | 24.37 |
| GTEX-12WSA-2926- | TCGA-LGG | GTEx | Normal | Cortex | Brain (Cortex) | 3.3 | 9.83 | 13.13 |
| GTEX-12WSC-3026- | TCGA-LGG | GTEx | Normal | Cortex | Brain (Cortex) | 5.99 | 23.89 | 29.88 |
| GTEX-12WSD-3126- | TCGA-LGG | GTEx | Normal | Cortex | Brain (Cortex) | 6.32 | 15.67 | 21.99 |
| GTEX-12WSF-3126- | TCGA-LGG | GTEx | Normal | Cortex | Brain (Cortex) | 5.05 | 19.37 | 24.42 |
| GTEX-12WSH-3026- | TCGA-LGG | GTEx | Normal | Cortex | Brain (Cortex) | 6.27 | 23.93 | 30.2 |
| GTEX-12ZZW-2926-S | TCGA-LGG | GTEx | Normal | Cortex | Brain (Cortex) | 3.75 | 19.84 | 23.59 |
| GTEX-12ZZX-2926-S | TCGA-LGG | GTEx | Normal | Cortex | Brain (Cortex) | 7.59 | 26.9 | 34.49 |
| GTEX-12ZZY-3026-S | TCGA-LGG | GTEx | Normal | Cortex | Brain (Cortex) | 8.06 | 25.15 | 33.21 |
| GTEX-12ZZZ-3026-S | TCGA-LGG | GTEx | Normal | Cortex | Brain (Cortex) | 6.4 | 22.58 | 28.98 |
| GTEX-1313W-3126-S | TCGA-LGG | GTEx | Normal | Cortex | Brain (Cortex) | 7.62 | 20.34 | 27.96 |
| GTEX-131XW-3126-S | TCGA-LGG | GTEx | Normal | Cortex | Brain (Cortex) | 3.81 | 14 | 17.81 |
| GTEX-131YS-3126-S | TCGA-LGG | GTEx | Normal | Cortex | Brain (Cortex) | 7.71 | 25.1 | 32.81 |
| GTEX-132Q8-3026-S | TCGA-LGG | GTEx | Normal | Cortex | Brain (Cortex) | 6.31 | 26.42 | 32.73 |
| GTEX-1399T-3026-S | TCGA-LGG | GTEx | Normal | Cortex | Brain (Cortex) | 6.86 | 22.1 | 28.96 |
| GTEX-139T8-1026-S | TCGA-LGG | GTEx | Normal | Cortex | Brain (Cortex) | 3.29 | 18.63 | 21.92 |
| GTEX-13FHO-3026-S | TCGA-LGG | GTEx | Normal | Cortex | Brain (Cortex) | 7.9 | 26.15 | 34.05 |
| GTEX-13FHP-3026-S | TCGA-LGG | GTEx | Normal | Cortex | Brain (Cortex) | 6.75 | 17.72 | 24.47 |
| GTEX-13FLW-1426-S | TCGA-LGG | GTEx | Normal | Cortex | Brain (Cortex) | 5.84 | 25.71 | 31.55 |
| GTEX-13FXS-3126-S | TCGA-LGG | GTEx | Normal | Cortex | Brain (Cortex) | 7.55 | 24.6 | 32.15 |
| GTEX-13G51-3026-S | TCGA-LGG | GTEx | Normal | Cortex | Brain (Cortex) | 8.53 | 18.11 | 26.64 |
| GTEX-13IVO-2926-S | TCGA-LGG | GTEx | Normal | Cortex | Brain (Cortex) | 5.45 | 19.55 | 25 |
| GTEX-13JUV-2926-S | TCGA-LGG | GTEx | Normal | Cortex | Brain (Cortex) | 4.63 | 14.94 | 19.57 |
| GTEX-13JVG-3126-S | TCGA-LGG | GTEx | Normal | Cortex | Brain (Cortex) | 7.78 | 19.05 | 26.83 |
| GTEX-13NYB-3026-S | TCGA-LGG | GTEx | Normal | Cortex | Brain (Cortex) | 7.3 | 25.19 | 32.49 |
| GTEX-13NYC-2826-S | TCGA-LGG | GTEx | Normal | Cortex | Brain (Cortex) | 10.16 | 34.28 | 44.44 |
| GTEX-13NYS-3126-S | TCGA-LGG | GTEx | Normal | Cortex | Brain (Cortex) | 8.29 | 30.07 | 38.36 |
| GTEX-13O3O-3126-S | TCGA-LGG | GTEx | Normal | Cortex | Brain (Cortex) | 4.9 | 23.19 | 28.09 |
| GTEX-13O3Q-2926-S | TCGA-LGG | GTEx | Normal | Cortex | Brain (Cortex) | 7.07 | 19.16 | 26.23 |
| GTEX-13OVH-3026-S | TCGA-LGG | GTEx | Normal | Cortex | Brain (Cortex) | 5.14 | 25.56 | 30.7 |
| GTEX-13OVJ-2826-S | TCGA-LGG | GTEx | Normal | Cortex | Brain (Cortex) | 8.67 | 27.45 | 36.12 |
| GTEX-13OW6-3026- | TCGA-LGG | GTEx | Normal | Cortex | Brain (Cortex) | 8.62 | 28 | 36.62 |
| GTEX-13OW7-3026- | TCGA-LGG | GTEx | Normal | Cortex | Brain (Cortex) | 5.41 | 15.23 | 20.64 |
| GTEX-13OW8-2826- | TCGA-LGG | GTEx | Normal | Cortex | Brain (Cortex) | 5.8 | 20.06 | 25.86 |

| | | | | | | | | |
|---|---|---|---|---|---|---|---|---|
| GTEX-13PL6-3126-S | TCGA-LGG | GTEx | Normal | Cortex | Brain (Cortex) | 8.09 | 22.3 | 30.39 |
| GTEX-13PVQ-3026-S | TCGA-LGG | GTEx | Normal | Cortex | Brain (Cortex) | 7.25 | 20.61 | 27.86 |
| GTEX-13QIC-2926-S | TCGA-LGG | GTEx | Normal | Cortex | Brain (Cortex) | 6.28 | 22.18 | 28.46 |
| GTEX-13S7M-3126-S | TCGA-LGG | GTEx | Normal | Cortex | Brain (Cortex) | 3.93 | 20.78 | 24.71 |
| GTEX-13SLX-3126-S | TCGA-LGG | GTEx | Normal | Cortex | Brain (Cortex) | 6.53 | 27.75 | 34.28 |
| GTEX-13VXU-2926-S | TCGA-LGG | GTEx | Normal | Cortex | Brain (Cortex) | 6.63 | 19.82 | 26.45 |
| GTEX-13X6J-3026-S | TCGA-LGG | GTEx | Normal | Cortex | Brain (Cortex) | 4.31 | 16.01 | 20.32 |
| GTEX-13X6K-2926-S | TCGA-LGG | GTEx | Normal | Cortex | Brain (Cortex) | 5.31 | 16.92 | 22.23 |
| GTEX-1445S-3026-S | TCGA-LGG | GTEx | Normal | Cortex | Brain (Cortex) | 4.74 | 13.51 | 18.25 |
| GTEX-144FL-3026-S | TCGA-LGG | GTEx | Normal | Cortex | Brain (Cortex) | 7.68 | 24.07 | 31.75 |
| GTEX-144GL-3026-S | TCGA-LGG | GTEx | Normal | Cortex | Brain (Cortex) | 5.08 | 22.66 | 27.74 |
| GTEX-145LS-3126-S | TCGA-LGG | GTEx | Normal | Cortex | Brain (Cortex) | 7.14 | 22.17 | 29.31 |
| GTEX-145MF-2726-S | TCGA-LGG | GTEx | Normal | Cortex | Brain (Cortex) | 5.71 | 17.92 | 23.63 |
| GTEX-145MG-3026-S | TCGA-LGG | GTEx | Normal | Cortex | Brain (Cortex) | 7 | 25.16 | 32.16 |
| GTEX-145MH-3026-S | TCGA-LGG | GTEx | Normal | Cortex | Brain (Cortex) | 7.67 | 25.1 | 32.77 |
| GTEX-147GR-3026-S | TCGA-LGG | GTEx | Normal | Cortex | Brain (Cortex) | 6.78 | 10.72 | 17.5 |
| GTEX-14A5I-2926-SI | TCGA-LGG | GTEx | Normal | Cortex | Brain (Cortex) | 5.75 | 23.81 | 29.56 |
| GTEX-14ASI-3026-S | TCGA-LGG | GTEx | Normal | Cortex | Brain (Cortex) | 6.45 | 23.64 | 30.09 |
| GTEX-14BIL-3026-SI | TCGA-LGG | GTEx | Normal | Cortex | Brain (Cortex) | 5.92 | 15.85 | 21.77 |
| GTEX-14BIM-3026-S | TCGA-LGG | GTEx | Normal | Cortex | Brain (Cortex) | 5.55 | 23.45 | 29 |
| GTEX-14BIN-3226-S | TCGA-LGG | GTEx | Normal | Cortex | Brain (Cortex) | 6.35 | 21.25 | 27.6 |
| GTEX-14BMV-3026-S | TCGA-LGG | GTEx | Normal | Cortex | Brain (Cortex) | 8.48 | 29.03 | 37.51 |
| GTEX-14C39-3126-S | TCGA-LGG | GTEx | Normal | Cortex | Brain (Cortex) | 5.99 | 21.54 | 27.53 |
| GTEX-14C5O-3026-S | TCGA-LGG | GTEx | Normal | Cortex | Brain (Cortex) | 3.76 | 21.96 | 25.72 |
| GTEX-14DAQ-3126-S | TCGA-LGG | GTEx | Normal | Cortex | Brain (Cortex) | 6.01 | 27.54 | 33.55 |
| GTEX-14JG1-3026-S | TCGA-LGG | GTEx | Normal | Cortex | Brain (Cortex) | 5.87 | 21.85 | 27.72 |
| GTEX-14JIY-2926-SM | TCGA-LGG | GTEx | Normal | Cortex | Brain (Cortex) | 7.7 | 23.77 | 31.47 |
| GTEX-14LZ3-3026-S | TCGA-LGG | GTEx | Normal | Cortex | Brain (Cortex) | 5.5 | 21.95 | 27.45 |
| GTEX-14PJM-3126-S | TCGA-LGG | GTEx | Normal | Cortex | Brain (Cortex) | 6.69 | 22.68 | 29.37 |
| GTEX-15DCD-2926-S | TCGA-LGG | GTEx | Normal | Cortex | Brain (Cortex) | 7.31 | 23.38 | 30.69 |
| GTEX-15EO6-3026-S | TCGA-LGG | GTEx | Normal | Cortex | Brain (Cortex) | 6.28 | 30.05 | 36.33 |
| GTEX-15ER7-3126-S | TCGA-LGG | GTEx | Normal | Cortex | Brain (Cortex) | 6.29 | 23.23 | 29.52 |
| GTEX-15G19-2926-S | TCGA-LGG | GTEx | Normal | Cortex | Brain (Cortex) | 10.26 | 29.63 | 39.89 |
| GTEX-15RJE-3026-S | TCGA-LGG | GTEx | Normal | Cortex | Brain (Cortex) | 4.72 | 17.37 | 22.09 |
| GTEX-15SHU-3126-S | TCGA-LGG | GTEx | Normal | Cortex | Brain (Cortex) | 7.37 | 17.89 | 25.26 |
| GTEX-15UF6-3026-S | TCGA-LGG | GTEx | Normal | Cortex | Brain (Cortex) | 4.34 | 13.54 | 17.88 |
| GTEX-16GPK-3026-S | TCGA-LGG | GTEx | Normal | Cortex | Brain (Cortex) | 4.57 | 12.38 | 16.95 |
| GTEX-16XZZ-3026-S | TCGA-LGG | GTEx | Normal | Cortex | Brain (Cortex) | 5.25 | 17.72 | 22.97 |

| | | | | | | | | |
|---|---|---|---|---|---|---|---|---|
| GTEX-16YQH-3026-S | TCGA-LGG | GTEx | Normal | Cortex | Brain (Cortex) | 4.36 | 16.14 | 20.5 |
| GTEX-16Z82-3026-S | TCGA-LGG | GTEx | Normal | Cortex | Brain (Cortex) | 6.18 | 17.1 | 23.28 |
| GTEX-17EVP-3026-S | TCGA-LGG | GTEx | Normal | Cortex | Brain (Cortex) | 4.54 | 13.35 | 17.89 |
| GTEX-17EVQ-2826-S | TCGA-LGG | GTEx | Normal | Cortex | Brain (Cortex) | 5.9 | 20.67 | 26.57 |
| GTEX-17F97-3026-S | TCGA-LGG | GTEx | Normal | Cortex | Brain (Cortex) | 7.11 | 23.99 | 31.1 |
| GTEX-17HG3-2926-S | TCGA-LGG | GTEx | Normal | Cortex | Brain (Cortex) | 5.26 | 15.63 | 20.89 |
| GTEX-17HHY-3026-S | TCGA-LGG | GTEx | Normal | Cortex | Brain (Cortex) | 6.62 | 23.06 | 29.68 |
| GTEX-17HII-3026-SM | TCGA-LGG | GTEx | Normal | Cortex | Brain (Cortex) | 5.24 | 14.9 | 20.14 |
| GTEX-17JCI-3126-SI | TCGA-LGG | GTEx | Normal | Cortex | Brain (Cortex) | 7.24 | 20.46 | 27.7 |
| GTEX-17MF6-2926-S | TCGA-LGG | GTEx | Normal | Cortex | Brain (Cortex) | 2.94 | 9.46 | 12.4 |
| GTEX-183WM-2826- | TCGA-LGG | GTEx | Normal | Cortex | Brain (Cortex) | 6.91 | 15.7 | 22.61 |
| GTEX-18465-3026-S | TCGA-LGG | GTEx | Normal | Cortex | Brain (Cortex) | 5.22 | 24.07 | 29.29 |
| GTEX-18A6Q-2926-S | TCGA-LGG | GTEx | Normal | Cortex | Brain (Cortex) | 6.21 | 19.13 | 25.34 |
| GTEX-1A3MX-3026-S | TCGA-LGG | GTEx | Normal | Cortex | Brain (Cortex) | 5.29 | 20.68 | 25.97 |
| GTEX-1A8G6-2926-S | TCGA-LGG | GTEx | Normal | Cortex | Brain (Cortex) | 5.07 | 14.6 | 19.67 |
| GTEX-1A8G7-3126-S | TCGA-LGG | GTEx | Normal | Cortex | Brain (Cortex) | 5.87 | 13.74 | 19.61 |
| GTEX-1AX9I-3026-S | TCGA-LGG | GTEx | Normal | Cortex | Brain (Cortex) | 4.27 | 16.19 | 20.46 |
| GTEX-1B8L1-3026-S | TCGA-LGG | GTEx | Normal | Cortex | Brain (Cortex) | 4.84 | 17.58 | 22.42 |
| GTEX-1B8SF-3126-S | TCGA-LGG | GTEx | Normal | Cortex | Brain (Cortex) | 5.98 | 19.25 | 25.23 |
| GTEX-1B8SG-1726-S | TCGA-LGG | GTEx | Normal | Cortex | Brain (Cortex) | 4.4 | 8.66 | 13.06 |
| GTEX-1B933-3126-S | TCGA-LGG | GTEx | Normal | Cortex | Brain (Cortex) | 5.25 | 21.43 | 26.68 |
| GTEX-1C64N-3026-S | TCGA-LGG | GTEx | Normal | Cortex | Brain (Cortex) | 2.87 | 8.28 | 11.15 |
| GTEX-1C6VQ-3026-S | TCGA-LGG | GTEx | Normal | Cortex | Brain (Cortex) | 6.66 | 14.67 | 21.33 |
| GTEX-1C6VS-3026-S | TCGA-LGG | GTEx | Normal | Cortex | Brain (Cortex) | 7.98 | 22.83 | 30.81 |
| GTEX-1C6WA-3026- | TCGA-LGG | GTEx | Normal | Cortex | Brain (Cortex) | 4.23 | 15.81 | 20.04 |
| GTEX-1CAV2-2726-S | TCGA-LGG | GTEx | Normal | Cortex | Brain (Cortex) | 2.24 | 5.81 | 8.05 |
| GTEX-1CB4H-3126-S | TCGA-LGG | GTEx | Normal | Cortex | Brain (Cortex) | 6.69 | 21.67 | 28.36 |
| GTEX-1E1VI-3026-S | TCGA-LGG | GTEx | Normal | Cortex | Brain (Cortex) | 6.47 | 21.41 | 27.88 |
| GTEX-1E2YA-3026-S | TCGA-LGG | GTEx | Normal | Cortex | Brain (Cortex) | 3.21 | 10.11 | 13.32 |
| GTEX-1EKGG-2926- | TCGA-LGG | GTEx | Normal | Cortex | Brain (Cortex) | 5.03 | 19.2 | 24.23 |
| GTEX-1EMGI-3026-S | TCGA-LGG | GTEx | Normal | Cortex | Brain (Cortex) | 7.11 | 24.18 | 31.29 |
| GTEX-1EN7A-3026-S | TCGA-LGG | GTEx | Normal | Cortex | Brain (Cortex) | 3.89 | 9.48 | 13.37 |
| GTEX-1EWIQ-3126-S | TCGA-LGG | GTEx | Normal | Cortex | Brain (Cortex) | 6.3 | 19.63 | 25.93 |
| GTEX-1EX96-3026-S | TCGA-LGG | GTEx | Normal | Cortex | Brain (Cortex) | 6.57 | 21.38 | 27.95 |
| GTEX-1F48J-2826-S | TCGA-LGG | GTEx | Normal | Cortex | Brain (Cortex) | 6.89 | 24.67 | 31.56 |
| GTEX-1F52S-3126-S | TCGA-LGG | GTEx | Normal | Cortex | Brain (Cortex) | 4.03 | 12.65 | 16.68 |
| GTEX-1F6I4-3126-SI | TCGA-LGG | GTEx | Normal | Cortex | Brain (Cortex) | 3.54 | 10.82 | 14.36 |
| GTEX-1F6RS-3126-S | TCGA-LGG | GTEx | Normal | Cortex | Brain (Cortex) | 6.91 | 24.41 | 31.32 |

| | | | | | | | | |
|---|---|---|---|---|---|---|---|---|
| GTEX-1F75A-3026-S | TCGA-LGG | GTEx | Normal | Cortex | Brain (Cortex) | 5.78 | 22.91 | 28.69 |
| GTEX-1F75B-3026-S | TCGA-LGG | GTEx | Normal | Cortex | Brain (Cortex) | 5.93 | 21.15 | 27.08 |
| GTEX-1F7RK-1826-S | TCGA-LGG | GTEx | Normal | Cortex | Brain (Cortex) | 7.77 | 15.71 | 23.48 |
| GTEX-1F88E-2826-S | TCGA-LGG | GTEx | Normal | Cortex | Brain (Cortex) | 4.75 | 12.39 | 17.14 |
| GTEX-1GF9V-2926-S | TCGA-LGG | GTEx | Normal | Cortex | Brain (Cortex) | 5.47 | 17.21 | 22.68 |
| GTEX-1GF9W-2626- | TCGA-LGG | GTEx | Normal | Cortex | Brain (Cortex) | 4.41 | 22.43 | 26.84 |
| GTEX-1GMR8-3026- | TCGA-LGG | GTEx | Normal | Cortex | Brain (Cortex) | 4.42 | 22.61 | 27.03 |
| GTEX-1GMRU-3026- | TCGA-LGG | GTEx | Normal | Cortex | Brain (Cortex) | 7.91 | 27.13 | 35.04 |
| GTEX-1GN1V-3126-S | TCGA-LGG | GTEx | Normal | Cortex | Brain (Cortex) | 3.23 | 7.69 | 10.92 |
| GTEX-1GN2E-3026-S | TCGA-LGG | GTEx | Normal | Cortex | Brain (Cortex) | 3.31 | 13.86 | 17.17 |
| GTEX-1GN73-3126-S | TCGA-LGG | GTEx | Normal | Cortex | Brain (Cortex) | 5.76 | 22.75 | 28.51 |
| GTEX-1GZ2Q-3226-S | TCGA-LGG | GTEx | Normal | Cortex | Brain (Cortex) | 5.52 | 22.52 | 28.04 |
| GTEX-1GZ4I-3026-S | TCGA-LGG | GTEx | Normal | Cortex | Brain (Cortex) | 6.97 | 27.2 | 34.17 |
| GTEX-1GZHY-3126-S | TCGA-LGG | GTEx | Normal | Cortex | Brain (Cortex) | 3.5 | 7.85 | 11.35 |
| GTEX-1H1DG-2926-S | TCGA-LGG | GTEx | Normal | Cortex | Brain (Cortex) | 9.44 | 19.83 | 29.27 |
| GTEX-1H1ZS-3026-S | TCGA-LGG | GTEx | Normal | Cortex | Brain (Cortex) | 6.56 | 24.78 | 31.34 |
| GTEX-1H23P-2926-S | TCGA-LGG | GTEx | Normal | Cortex | Brain (Cortex) | 4.25 | 16.63 | 20.88 |
| GTEX-1H2FU-3126-S | TCGA-LGG | GTEx | Normal | Cortex | Brain (Cortex) | 6.24 | 24.13 | 30.37 |
| GTEX-1H3NZ-3026-S | TCGA-LGG | GTEx | Normal | Cortex | Brain (Cortex) | 7.87 | 25.09 | 32.96 |
| GTEX-1H3O1-1726-S | TCGA-LGG | GTEx | Normal | Cortex | Brain (Cortex) | 5.96 | 13.16 | 19.12 |
| GTEX-1H3VE-2726-S | TCGA-LGG | GTEx | Normal | Cortex | Brain (Cortex) | 4.07 | 11.53 | 15.6 |
| GTEX-1HB9E-2926-S | TCGA-LGG | GTEx | Normal | Cortex | Brain (Cortex) | 7.77 | 17.39 | 25.16 |
| GTEX-1HBPH-3126-S | TCGA-LGG | GTEx | Normal | Cortex | Brain (Cortex) | 5.22 | 18.65 | 23.87 |
| GTEX-1HBPI-2826-S | TCGA-LGG | GTEx | Normal | Cortex | Brain (Cortex) | 5.87 | 15.62 | 21.49 |
| GTEX-1HBPM-2926- | TCGA-LGG | GTEx | Normal | Cortex | Brain (Cortex) | 2.75 | 6.53 | 9.28 |
| GTEX-1HCU6-3126-S | TCGA-LGG | GTEx | Normal | Cortex | Brain (Cortex) | 7.38 | 30.36 | 37.74 |
| GTEX-1HCVE-3026-S | TCGA-LGG | GTEx | Normal | Cortex | Brain (Cortex) | 7.55 | 21.07 | 28.62 |
| GTEX-1HFI6-3026-S | TCGA-LGG | GTEx | Normal | Cortex | Brain (Cortex) | 6.87 | 15.25 | 22.12 |
| GTEX-1HGF4-3126-S | TCGA-LGG | GTEx | Normal | Cortex | Brain (Cortex) | 6.31 | 15.95 | 22.26 |
| GTEX-1HR98-3026-S | TCGA-LGG | GTEx | Normal | Cortex | Brain (Cortex) | 6.81 | 21.75 | 28.56 |
| GTEX-1HSKV-3026-S | TCGA-LGG | GTEx | Normal | Cortex | Brain (Cortex) | 3.68 | 16.22 | 19.9 |
| GTEX-1HSMO-3026- | TCGA-LGG | GTEx | Normal | Cortex | Brain (Cortex) | 6.92 | 23.81 | 30.73 |
| GTEX-1HT8W-3026- | TCGA-LGG | GTEx | Normal | Cortex | Brain (Cortex) | 6.21 | 24.97 | 31.18 |
| GTEX-1HUB1-2926-S | TCGA-LGG | GTEx | Normal | Cortex | Brain (Cortex) | 4.2 | 15.38 | 19.58 |
| GTEX-1I19N-3026-SI | TCGA-LGG | GTEx | Normal | Cortex | Brain (Cortex) | 4.88 | 18.7 | 23.58 |
| GTEX-1I1CD-3026-S | TCGA-LGG | GTEx | Normal | Cortex | Brain (Cortex) | 6.7 | 17.68 | 24.38 |
| GTEX-1I1GQ-3026-S | TCGA-LGG | GTEx | Normal | Cortex | Brain (Cortex) | 5.32 | 15.67 | 20.99 |
| GTEX-1I1GR-2926-S | TCGA-LGG | GTEx | Normal | Cortex | Brain (Cortex) | 6.06 | 22.82 | 28.88 |

| | | | | | | | | |
|---|---|---|---|---|---|---|---|---|
| GTEX-1I1GV-3126-S | TCGA-LGG | GTEx | Normal | Cortex | Brain (Cortex) | 3.77 | 15.12 | 18.89 |
| GTEX-1I1HK-2826-S | TCGA-LGG | GTEx | Normal | Cortex | Brain (Cortex) | 5.94 | 18.79 | 24.73 |
| GTEX-1ICLZ-2826-S | TCGA-LGG | GTEx | Normal | Cortex | Brain (Cortex) | 6.06 | 16.34 | 22.4 |
| GTEX-1IDFM-2826-S | TCGA-LGG | GTEx | Normal | Cortex | Brain (Cortex) | 3.77 | 8.2 | 11.97 |
| GTEX-1IDJH-3026-S | TCGA-LGG | GTEx | Normal | Cortex | Brain (Cortex) | 7.6 | 23.22 | 30.82 |
| GTEX-1IKJJ-2926-SI | TCGA-LGG | GTEx | Normal | Cortex | Brain (Cortex) | 3.32 | 10.79 | 14.11 |
| GTEX-1IKK5-2926-S | TCGA-LGG | GTEx | Normal | Cortex | Brain (Cortex) | 7.41 | 23.26 | 30.67 |
| GTEX-1IL2U-3026-SI | TCGA-LGG | GTEx | Normal | Cortex | Brain (Cortex) | 6.28 | 20.16 | 26.44 |
| GTEX-1IOXB-3126-S | TCGA-LGG | GTEx | Normal | Cortex | Brain (Cortex) | 2.47 | 8.22 | 10.69 |
| GTEX-1IY9M-3026-S | TCGA-LGG | GTEx | Normal | Cortex | Brain (Cortex) | 5.61 | 17.92 | 23.53 |
| GTEX-1J1OQ-3226-S | TCGA-LGG | GTEx | Normal | Cortex | Brain (Cortex) | 5.24 | 20.13 | 25.37 |
| GTEX-1J8Q2-3026-S | TCGA-LGG | GTEx | Normal | Cortex | Brain (Cortex) | 5.77 | 22.91 | 28.68 |
| GTEX-1JJ6O-3126-S | TCGA-LGG | GTEx | Normal | Cortex | Brain (Cortex) | 5.3 | 22.5 | 27.8 |
| GTEX-1JJE9-2926-S | TCGA-LGG | GTEx | Normal | Cortex | Brain (Cortex) | 2.45 | 6.12 | 8.57 |
| GTEX-1JJEA-2826-S | TCGA-LGG | GTEx | Normal | Cortex | Brain (Cortex) | 3.84 | 15.66 | 19.5 |
| GTEX-1JMLX-2926-S | TCGA-LGG | GTEx | Normal | Cortex | Brain (Cortex) | 5.02 | 16.56 | 21.58 |
| GTEX-1JMOU-2726- | TCGA-LGG | GTEx | Normal | Cortex | Brain (Cortex) | 7.98 | 21.98 | 29.96 |
| GTEX-1JMPZ-2726-S | TCGA-LGG | GTEx | Normal | Cortex | Brain (Cortex) | 5.67 | 21.98 | 27.65 |
| GTEX-1JMQK-3026-S | TCGA-LGG | GTEx | Normal | Cortex | Brain (Cortex) | 5 | 19.72 | 24.72 |
| GTEX-1JMQL-2826-S | TCGA-LGG | GTEx | Normal | Cortex | Brain (Cortex) | 7.32 | 24.8 | 32.12 |
| GTEX-1KD4Q-2926-S | TCGA-LGG | GTEx | Normal | Cortex | Brain (Cortex) | 7.16 | 23.85 | 31.01 |
| GTEX-1KWVE-2326- | TCGA-LGG | GTEx | Normal | Cortex | Brain (Cortex) | 5.35 | 22.73 | 28.08 |
| GTEX-1LC47-2926-S | TCGA-LGG | GTEx | Normal | Cortex | Brain (Cortex) | 5.54 | 20.14 | 25.68 |
| GTEX-1LG7Y-3026-S | TCGA-LGG | GTEx | Normal | Cortex | Brain (Cortex) | 5.94 | 23.77 | 29.71 |
| GTEX-1LKK1-3126-S | TCGA-LGG | GTEx | Normal | Cortex | Brain (Cortex) | 5.44 | 20.33 | 25.77 |
| GTEX-1M4P7-2726-S | TCGA-LGG | GTEx | Normal | Cortex | Brain (Cortex) | 4.14 | 11.91 | 16.05 |
| GTEX-1N2DW-2826- | TCGA-LGG | GTEx | Normal | Cortex | Brain (Cortex) | 4.67 | 13.11 | 17.78 |
| GTEX-1N5O9-2926-S | TCGA-LGG | GTEx | Normal | Cortex | Brain (Cortex) | 4.27 | 16.67 | 20.94 |
| GTEX-1NV5F-3226-S | TCGA-LGG | GTEx | Normal | Cortex | Brain (Cortex) | 3.53 | 14.57 | 18.1 |
| GTEX-1NV8Z-3126-S | TCGA-LGG | GTEx | Normal | Cortex | Brain (Cortex) | 5.66 | 19.48 | 25.14 |
| GTEX-1O9I2-3026-S | TCGA-LGG | GTEx | Normal | Cortex | Brain (Cortex) | 5.89 | 17.31 | 23.2 |
| GTEX-1OJC4-2826-S | TCGA-LGG | GTEx | Normal | Cortex | Brain (Cortex) | 4.97 | 23.69 | 28.66 |
| GTEX-1PIIG-2726-SI | TCGA-LGG | GTEx | Normal | Cortex | Brain (Cortex) | 6.95 | 26.66 | 33.61 |
| GTEX-1PPGY-3126-S | TCGA-LGG | GTEx | Normal | Cortex | Brain (Cortex) | 5.31 | 22.75 | 28.06 |
| GTEX-1PWST-3126- | TCGA-LGG | GTEx | Normal | Cortex | Brain (Cortex) | 6.35 | 21.64 | 27.99 |
| GTEX-1QMI2-3126-S | TCGA-LGG | GTEx | Normal | Cortex | Brain (Cortex) | 7.21 | 25.04 | 32.25 |
| GTEX-1QP6S-2926-S | TCGA-LGG | GTEx | Normal | Cortex | Brain (Cortex) | 8.64 | 15.77 | 24.41 |
| GTEX-1R9K4-3126-S | TCGA-LGG | GTEx | Normal | Cortex | Brain (Cortex) | 5.22 | 20.88 | 26.1 |

| | | | | | | | | |
|---|---|---|---|---|---|---|---|---|
| GTEX-1R9PO-2926-S | TCGA-LGG | GTEx | Normal | Cortex | Brain (Cortex) | 8.75 | 29.88 | 38.63 |
| GTEX-1RQEC-2926- | TCGA-LGG | GTEx | Normal | Cortex | Brain (Cortex) | 4.81 | 18.1 | 22.91 |
| GTEX-N7MT-1126-SI | TCGA-LGG | GTEx | Normal | Cortex | Brain (Cortex) | 7.01 | 13.84 | 20.85 |
| GTEX-NPJ7-2726-SM | TCGA-LGG | GTEx | Normal | Cortex | Brain (Cortex) | 5.27 | 16.46 | 21.73 |
| GTEX-NPJ8-1526-SM | TCGA-LGG | GTEx | Normal | Cortex | Brain (Cortex) | 8.62 | 27.45 | 36.07 |
| GTEX-OHPN-2526-S | TCGA-LGG | GTEx | Normal | Cortex | Brain (Cortex) | 8.54 | 30.61 | 39.15 |
| GTEX-OXRN-2426-S | TCGA-LGG | GTEx | Normal | Cortex | Brain (Cortex) | 3.88 | 17.93 | 21.81 |
| GTEX-P44G-2426-SI | TCGA-LGG | GTEx | Normal | Cortex | Brain (Cortex) | 7.12 | 20.86 | 27.98 |
| GTEX-PVOW-2526-S | TCGA-LGG | GTEx | Normal | Cortex | Brain (Cortex) | 8.18 | 25.06 | 33.24 |
| GTEX-PWO3-0926-S | TCGA-LGG | GTEx | Normal | Cortex | Brain (Cortex) | 3.98 | 12.21 | 16.19 |
| GTEX-Q2AG-2926-S | TCGA-LGG | GTEx | Normal | Cortex | Brain (Cortex) | 4.8 | 13.96 | 18.76 |
| GTEX-QDT8-2926-SI | TCGA-LGG | GTEx | Normal | Cortex | Brain (Cortex) | 7.94 | 21.73 | 29.67 |
| GTEX-QMR6-1426-S | TCGA-LGG | GTEx | Normal | Cortex | Brain (Cortex) | 5.51 | 20.04 | 25.55 |
| GTEX-QVJO-1426-S | TCGA-LGG | GTEx | Normal | Cortex | Brain (Cortex) | 6.08 | 21.01 | 27.09 |
| GTEX-QVUS-2826-S | TCGA-LGG | GTEx | Normal | Cortex | Brain (Cortex) | 6 | 24.87 | 30.87 |
| GTEX-RNOR-2326-S | TCGA-LGG | GTEx | Normal | Cortex | Brain (Cortex) | 6.82 | 23.5 | 30.32 |
| GTEX-RU72-3026-SI | TCGA-LGG | GTEx | Normal | Cortex | Brain (Cortex) | 5.62 | 22.22 | 27.84 |
| GTEX-T2IS-3026-SM | TCGA-LGG | GTEx | Normal | Cortex | Brain (Cortex) | 6.41 | 19.66 | 26.07 |
| GTEX-T5JC-2426-SM | TCGA-LGG | GTEx | Normal | Cortex | Brain (Cortex) | 7.09 | 22.66 | 29.75 |
| GTEX-T6MN-2626-S | TCGA-LGG | GTEx | Normal | Cortex | Brain (Cortex) | 5.29 | 24.96 | 30.25 |
| GTEX-TSE9-3026-SM | TCGA-LGG | GTEx | Normal | Cortex | Brain (Cortex) | 5.29 | 19.9 | 25.19 |
| GTEX-UTHO-3026-S | TCGA-LGG | GTEx | Normal | Cortex | Brain (Cortex) | 8.68 | 26.71 | 35.39 |
| GTEX-WHSE-3026-S | TCGA-LGG | GTEx | Normal | Cortex | Brain (Cortex) | 6.51 | 22.28 | 28.79 |
| GTEX-WL46-2926-SI | TCGA-LGG | GTEx | Normal | Cortex | Brain (Cortex) | 5.45 | 22.27 | 27.72 |
| GTEX-WVLH-3026-S | TCGA-LGG | GTEx | Normal | Cortex | Brain (Cortex) | 5.78 | 23.1 | 28.88 |
| GTEX-WWYW-3126- | TCGA-LGG | GTEx | Normal | Cortex | Brain (Cortex) | 7.88 | 24.09 | 31.97 |
| GTEX-WZTO-2926-S | TCGA-LGG | GTEx | Normal | Cortex | Brain (Cortex) | 7.94 | 26.32 | 34.26 |
| GTEX-X4XX-3026-SI | TCGA-LGG | GTEx | Normal | Cortex | Brain (Cortex) | 6.48 | 28.92 | 35.4 |
| GTEX-X4XY-3026-SM | TCGA-LGG | GTEx | Normal | Cortex | Brain (Cortex) | 5.92 | 18.82 | 24.74 |
| GTEX-X585-3026-SM | TCGA-LGG | GTEx | Normal | Cortex | Brain (Cortex) | 10.35 | 34.45 | 44.8 |
| GTEX-XLM4-3026-SI | TCGA-LGG | GTEx | Normal | Cortex | Brain (Cortex) | 6.72 | 24.05 | 30.77 |
| GTEX-Y111-2726-SM | TCGA-LGG | GTEx | Normal | Cortex | Brain (Cortex) | 5.81 | 14.2 | 20.01 |
| GTEX-Y8DK-0826-SI | TCGA-LGG | GTEx | Normal | Cortex | Brain (Cortex) | 6.38 | 21.65 | 28.03 |
| GTEX-YFC4-3126-SI | TCGA-LGG | GTEx | Normal | Cortex | Brain (Cortex) | 7.44 | 24.01 | 31.45 |
| GTEX-YJ89-3026-SM | TCGA-LGG | GTEx | Normal | Cortex | Brain (Cortex) | 5.45 | 19.4 | 24.85 |
| GTEX-Z93S-2926-SM | TCGA-LGG | GTEx | Normal | Cortex | Brain (Cortex) | 4.46 | 15.57 | 20.03 |
| GTEX-ZAJG-3126-SI | TCGA-LGG | GTEx | Normal | Cortex | Brain (Cortex) | 5.51 | 28.02 | 33.53 |
| GTEX-ZAK1-3026-SM | TCGA-LGG | GTEx | Normal | Cortex | Brain (Cortex) | 7.11 | 24.39 | 31.5 |

| | | | | | | | | |
|---|---|---|---|---|---|---|---|---|
| GTEX-ZDXO-2126-S | TCGA-LGG | GTEx | Normal | Cortex | Brain (Cortex) | 5.14 | 16.03 | 21.17 |
| GTEX-ZE7O-3126-SI | TCGA-LGG | GTEx | Normal | Cortex | Brain (Cortex) | 6.25 | 21.09 | 27.34 |
| GTEX-ZE9C-3026-SI | TCGA-LGG | GTEx | Normal | Cortex | Brain (Cortex) | 6.03 | 24.8 | 30.83 |
| GTEX-ZF28-3026-SM | TCGA-LGG | GTEx | Normal | Cortex | Brain (Cortex) | 6.68 | 20.95 | 27.63 |
| GTEX-ZUA1-3026-SI | TCGA-LGG | GTEx | Normal | Cortex | Brain (Cortex) | 4.65 | 21.44 | 26.09 |
| GTEX-ZVT3-3026-SM | TCGA-LGG | GTEx | Normal | Cortex | Brain (Cortex) | 8.9 | 26.35 | 35.25 |
| GTEX-ZYFD-3026-SI | TCGA-LGG | GTEx | Normal | Cortex | Brain (Cortex) | 6.24 | 23.04 | 29.28 |
| GTEX-ZYY3-3126-SM | TCGA-LGG | GTEx | Normal | Cortex | Brain (Cortex) | 8.34 | 27.85 | 36.19 |
| GTEX-ZZPT-3026-SM | TCGA-LGG | GTEx | Normal | Cortex | Brain (Cortex) | 4.48 | 18.54 | 23.02 |
| GTEX-1192X-0011-R | TCGA-LGG | GTEx | Normal | Cortex | Brain (Cortex) | 9.75 | 41.07 | 50.82 |
| GTEX-11DXY-0011-R | TCGA-LGG | GTEx | Normal | Cortex | Brain (Cortex) | 6.95 | 30.44 | 37.39 |
| GTEX-11DYG-0011-F | TCGA-LGG | GTEx | Normal | Cortex | Brain (Cortex) | 7.24 | 27.08 | 34.32 |
| GTEX-11DZ1-0011-R | TCGA-LGG | GTEx | Normal | Cortex | Brain (Cortex) | 7.99 | 32.88 | 40.87 |
| GTEX-11EI6-0011-R1 | TCGA-LGG | GTEx | Normal | Cortex | Brain (Cortex) | 6.02 | 16.89 | 22.91 |
| GTEX-11EMC-0011-F | TCGA-LGG | GTEx | Normal | Cortex | Brain (Cortex) | 7.94 | 29.38 | 37.32 |
| GTEX-11GSP-0011-F | TCGA-LGG | GTEx | Normal | Cortex | Brain (Cortex) | 5.45 | 18.28 | 23.73 |
| GTEX-11H98-0011-R | TCGA-LGG | GTEx | Normal | Cortex | Brain (Cortex) | 9.07 | 33.18 | 42.25 |
| GTEX-11PRG-0011-F | TCGA-LGG | GTEx | Normal | Cortex | Brain (Cortex) | 7.05 | 27.01 | 34.06 |
| GTEX-11TTK-0011-R | TCGA-LGG | GTEx | Normal | Cortex | Brain (Cortex) | 8.53 | 15.89 | 24.42 |
| GTEX-11TUW-0011-F | TCGA-LGG | GTEx | Normal | Cortex | Brain (Cortex) | 13.28 | 27.07 | 40.35 |
| GTEX-11WQC-0011- | TCGA-LGG | GTEx | Normal | Cortex | Brain (Cortex) | 6.69 | 17.41 | 24.1 |
| GTEX-11WQK-0011-I | TCGA-LGG | GTEx | Normal | Cortex | Brain (Cortex) | 5.64 | 19.31 | 24.95 |
| GTEX-11ZTS-0011-R | TCGA-LGG | GTEx | Normal | Cortex | Brain (Cortex) | 6.4 | 22.2 | 28.6 |
| GTEX-12126-0011-R | TCGA-LGG | GTEx | Normal | Cortex | Brain (Cortex) | 7.5 | 20.55 | 28.05 |
| GTEX-12WSA-0011-I | TCGA-LGG | GTEx | Normal | Cortex | Brain (Cortex) | 3.72 | 19.09 | 22.81 |
| GTEX-12WSC-0011- | TCGA-LGG | GTEx | Normal | Cortex | Brain (Cortex) | 7.21 | 32.07 | 39.28 |
| GTEX-12WSE-0011-I | TCGA-LGG | GTEx | Normal | Cortex | Brain (Cortex) | 8.18 | 27.97 | 36.15 |
| GTEX-12WSF-0011-I | TCGA-LGG | GTEx | Normal | Cortex | Brain (Cortex) | 9.57 | 30.56 | 40.13 |
| GTEX-12WSH-0011- | TCGA-LGG | GTEx | Normal | Cortex | Brain (Cortex) | 7.86 | 27.94 | 35.8 |
| GTEX-12WSM-0011- | TCGA-LGG | GTEx | Normal | Cortex | Brain (Cortex) | 8.18 | 27.11 | 35.29 |
| GTEX-12ZZW-0011-F | TCGA-LGG | GTEx | Normal | Cortex | Brain (Cortex) | 4.7 | 16.02 | 20.72 |
| GTEX-12ZZX-0011-R | TCGA-LGG | GTEx | Normal | Cortex | Brain (Cortex) | 9.66 | 35.8 | 45.46 |
| GTEX-12ZZY-0011-R | TCGA-LGG | GTEx | Normal | Cortex | Brain (Cortex) | 11.28 | 38.45 | 49.73 |
| GTEX-12ZZZ-0011-R | TCGA-LGG | GTEx | Normal | Cortex | Brain (Cortex) | 6.31 | 24.38 | 30.69 |
| GTEX-1313W-0011-F | TCGA-LGG | GTEx | Normal | Cortex | Brain (Cortex) | 7.96 | 25.79 | 33.75 |
| GTEX-131XH-0011-F | TCGA-LGG | GTEx | Normal | Cortex | Brain (Cortex) | 7.59 | 26.94 | 34.53 |
| GTEX-131XW-0011-F | TCGA-LGG | GTEx | Normal | Cortex | Brain (Cortex) | 3.28 | 8.9 | 12.18 |
| GTEX-131YS-0011-R | TCGA-LGG | GTEx | Normal | Cortex | Brain (Cortex) | 6.05 | 27 | 33.05 |

| | | | | | | | | |
|---|---|---|---|---|---|---|---|---|
| GTEX-132Q8-0011-R | TCGA-LGG | GTEx | Normal | Cortex | Brain (Cortex) | 5.04 | 27.94 | 32.98 |
| GTEX-139TS-0011-R | TCGA-LGG | GTEx | Normal | Cortex | Brain (Cortex) | 7.09 | 17.44 | 24.53 |
| GTEX-13CZV-0011-R | TCGA-LGG | GTEx | Normal | Cortex | Brain (Cortex) | 7.25 | 30.03 | 37.28 |
| GTEX-13FHO-0011-R | TCGA-LGG | GTEx | Normal | Cortex | Brain (Cortex) | 6.36 | 30 | 36.36 |
| GTEX-13FLV-0011-R | TCGA-LGG | GTEx | Normal | Cortex | Brain (Cortex) | 7.85 | 23.61 | 31.46 |
| GTEX-13FLW-0011-R | TCGA-LGG | GTEx | Normal | Cortex | Brain (Cortex) | 6.42 | 29.28 | 35.7 |
| GTEX-13FTZ-0011-R | TCGA-LGG | GTEx | Normal | Cortex | Brain (Cortex) | 5.19 | 20.07 | 25.26 |
| GTEX-13FXS-0011-R | TCGA-LGG | GTEx | Normal | Cortex | Brain (Cortex) | 5.2 | 23.88 | 29.08 |
| GTEX-13G51-0011-R | TCGA-LGG | GTEx | Normal | Cortex | Brain (Cortex) | 4.63 | 16.64 | 21.27 |
| GTEX-13IVO-0011-R | TCGA-LGG | GTEx | Normal | Cortex | Brain (Cortex) | 2.66 | 13.78 | 16.44 |
| GTEX-13JUV-0011-R | TCGA-LGG | GTEx | Normal | Cortex | Brain (Cortex) | 7.02 | 19.86 | 26.88 |
| GTEX-13JVG-0011-R | TCGA-LGG | GTEx | Normal | Cortex | Brain (Cortex) | 8.1 | 13.2 | 21.3 |
| GTEX-13N1W-0011-R | TCGA-LGG | GTEx | Normal | Cortex | Brain (Cortex) | 8.03 | 30.78 | 38.81 |
| GTEX-13N2G-0011-R | TCGA-LGG | GTEx | Normal | Cortex | Brain (Cortex) | 20.02 | 61.29 | 81.31 |
| GTEX-13NYB-0011-R | TCGA-LGG | GTEx | Normal | Cortex | Brain (Cortex) | 6.56 | 26.84 | 33.4 |
| GTEX-13NYS-0011-R | TCGA-LGG | GTEx | Normal | Cortex | Brain (Cortex) | 7.7 | 43.03 | 50.73 |
| GTEX-13NZA-0011-R | TCGA-LGG | GTEx | Normal | Cortex | Brain (Cortex) | 4.3 | 12.43 | 16.73 |
| GTEX-13O3O-0011-R | TCGA-LGG | GTEx | Normal | Cortex | Brain (Cortex) | 7.11 | 33.7 | 40.81 |
| GTEX-13O3Q-0011-R | TCGA-LGG | GTEx | Normal | Cortex | Brain (Cortex) | 9.74 | 33.13 | 42.87 |
| GTEX-13OVJ-0011-R | TCGA-LGG | GTEx | Normal | Cortex | Brain (Cortex) | 9.61 | 24.82 | 34.43 |
| GTEX-13OVL-0011-R | TCGA-LGG | GTEx | Normal | Cortex | Brain (Cortex) | 8.26 | 22.66 | 30.92 |
| GTEX-13OW6-0011-R | TCGA-LGG | GTEx | Normal | Cortex | Brain (Cortex) | 10.97 | 31.31 | 42.28 |
| GTEX-13OW7-0011-R | TCGA-LGG | GTEx | Normal | Cortex | Brain (Cortex) | 8.22 | 27.76 | 35.98 |
| GTEX-13OW8-0011-R | TCGA-LGG | GTEx | Normal | Cortex | Brain (Cortex) | 5.08 | 19.36 | 24.44 |
| GTEX-13QIC-0011-R | TCGA-LGG | GTEx | Normal | Cortex | Brain (Cortex) | 10.57 | 30.26 | 40.83 |
| GTEX-13QJC-0011-R | TCGA-LGG | GTEx | Normal | Cortex | Brain (Cortex) | 6.56 | 39.69 | 46.25 |
| GTEX-13RTJ-0011-R | TCGA-LGG | GTEx | Normal | Cortex | Brain (Cortex) | 6.63 | 35.17 | 41.8 |
| GTEX-13S7M-0011-R | TCGA-LGG | GTEx | Normal | Cortex | Brain (Cortex) | 8.51 | 32.32 | 40.83 |
| GTEX-13SLW-0011-R | TCGA-LGG | GTEx | Normal | Cortex | Brain (Cortex) | 5.23 | 23.95 | 29.18 |
| GTEX-13SLX-0011-R | TCGA-LGG | GTEx | Normal | Cortex | Brain (Cortex) | 8.91 | 39.06 | 47.97 |
| GTEX-13X6I-0011-R1 | TCGA-LGG | GTEx | Normal | Cortex | Brain (Cortex) | 8.27 | 29.43 | 37.7 |
| GTEX-13X6J-0011-R | TCGA-LGG | GTEx | Normal | Cortex | Brain (Cortex) | 6.19 | 24.65 | 30.84 |
| GTEX-13X6K-0011-R | TCGA-LGG | GTEx | Normal | Cortex | Brain (Cortex) | 8.77 | 31.54 | 40.31 |
| GTEX-1445S-0011-R | TCGA-LGG | GTEx | Normal | Cortex | Brain (Cortex) | 5.57 | 17.61 | 23.18 |
| GTEX-145LS-0011-R | TCGA-LGG | GTEx | Normal | Cortex | Brain (Cortex) | 8.81 | 26.93 | 35.74 |
| GTEX-145LU-0011-R | TCGA-LGG | GTEx | Normal | Cortex | Brain (Cortex) | 7.72 | 23.17 | 30.89 |
| GTEX-145MG-0011-R | TCGA-LGG | GTEx | Normal | Cortex | Brain (Cortex) | 9.15 | 36.24 | 45.39 |
| GTEX-145MH-0011-R | TCGA-LGG | GTEx | Normal | Cortex | Brain (Cortex) | 9.38 | 30.51 | 39.89 |

| | | | | | | | | |
|---|---|---|---|---|---|---|---|---|
| GTEX-145MI-0011-R | TCGA-LGG | GTEx | Normal | Cortex | Brain (Cortex) | 8.01 | 27.82 | 35.83 |
| GTEX-14753-0011-R | TCGA-LGG | GTEx | Normal | Cortex | Brain (Cortex) | 2.68 | 8.98 | 11.66 |
| GTEX-1477Z-0011-R | TCGA-LGG | GTEx | Normal | Cortex | Brain (Cortex) | 4.06 | 11.48 | 15.54 |
| GTEX-147F4-0011-R | TCGA-LGG | GTEx | Normal | Cortex | Brain (Cortex) | 6.3 | 17.77 | 24.07 |
| GTEX-147GR-0011-F | TCGA-LGG | GTEx | Normal | Cortex | Brain (Cortex) | 8.6 | 25.68 | 34.28 |
| GTEX-14ASI-0011-R | TCGA-LGG | GTEx | Normal | Cortex | Brain (Cortex) | 11.08 | 38.24 | 49.32 |
| GTEX-14BIL-0011-R1 | TCGA-LGG | GTEx | Normal | Cortex | Brain (Cortex) | 9.86 | 35.82 | 45.68 |
| GTEX-14BIN-0011-R | TCGA-LGG | GTEx | Normal | Cortex | Brain (Cortex) | 7.23 | 29.87 | 37.1 |
| GTEX-14C5O-0011-F | TCGA-LGG | GTEx | Normal | Cortex | Brain (Cortex) | 3.56 | 16.96 | 20.52 |
| GTEX-14E6D-0011-R | TCGA-LGG | GTEx | Normal | Cortex | Brain (Cortex) | 11.16 | 33.23 | 44.39 |
| GTEX-14E7W-0011-F | TCGA-LGG | GTEx | Normal | Cortex | Brain (Cortex) | 10.76 | 38.05 | 48.81 |
| GTEX-14JIY-0011-R1 | TCGA-LGG | GTEx | Normal | Cortex | Brain (Cortex) | 9.2 | 29.57 | 38.77 |
| GTEX-14LZ3-0011-R | TCGA-LGG | GTEx | Normal | Cortex | Brain (Cortex) | 8.32 | 19.93 | 28.25 |
| GTEX-14PJM-0011-F | TCGA-LGG | GTEx | Normal | Cortex | Brain (Cortex) | 8.62 | 27.97 | 36.59 |
| GTEX-14PJO-0011-R | TCGA-LGG | GTEx | Normal | Cortex | Brain (Cortex) | 6.67 | 21.49 | 28.16 |
| GTEX-14PN4-0011-R | TCGA-LGG | GTEx | Normal | Cortex | Brain (Cortex) | 8.07 | 31.22 | 39.29 |
| GTEX-14PQA-0011-F | TCGA-LGG | GTEx | Normal | Cortex | Brain (Cortex) | 6.91 | 28.39 | 35.3 |
| GTEX-15CHQ-0011-I | TCGA-LGG | GTEx | Normal | Cortex | Brain (Cortex) | 9.01 | 30.28 | 39.29 |
| GTEX-15DCD-0011-F | TCGA-LGG | GTEx | Normal | Cortex | Brain (Cortex) | 8.97 | 29.38 | 38.35 |
| GTEX-15DDE-0011-F | TCGA-LGG | GTEx | Normal | Cortex | Brain (Cortex) | 3.87 | 19.32 | 23.19 |
| GTEX-15G19-0011-R | TCGA-LGG | GTEx | Normal | Cortex | Brain (Cortex) | 13.15 | 39.44 | 52.59 |
| GTEX-15UF6-0011-R | TCGA-LGG | GTEx | Normal | Cortex | Brain (Cortex) | 3.76 | 9.65 | 13.41 |
| GTEX-16GPK-0011-F | TCGA-LGG | GTEx | Normal | Cortex | Brain (Cortex) | 3.47 | 16.37 | 19.84 |
| GTEX-16NPV-0011-F | TCGA-LGG | GTEx | Normal | Cortex | Brain (Cortex) | 5.44 | 16.64 | 22.08 |
| GTEX-16XZZ-0011-R | TCGA-LGG | GTEx | Normal | Cortex | Brain (Cortex) | 8.46 | 26.85 | 35.31 |
| GTEX-16YQH-0011-F | TCGA-LGG | GTEx | Normal | Cortex | Brain (Cortex) | 6.01 | 22.91 | 28.92 |
| GTEX-16Z82-0011-R | TCGA-LGG | GTEx | Normal | Cortex | Brain (Cortex) | 6.17 | 19.18 | 25.35 |
| GTEX-17EVP-0011-F | TCGA-LGG | GTEx | Normal | Cortex | Brain (Cortex) | 12.92 | 34.51 | 47.43 |
| GTEX-17F97-0011-R | TCGA-LGG | GTEx | Normal | Cortex | Brain (Cortex) | 7.52 | 27.09 | 34.61 |
| GTEX-17HG3-0011-F | TCGA-LGG | GTEx | Normal | Cortex | Brain (Cortex) | 9.34 | 36.77 | 46.11 |
| GTEX-17HII-0011-R1 | TCGA-LGG | GTEx | Normal | Cortex | Brain (Cortex) | 4.02 | 21.04 | 25.06 |
| GTEX-17JCI-0011-R1 | TCGA-LGG | GTEx | Normal | Cortex | Brain (Cortex) | 8.59 | 24.71 | 33.3 |
| GTEX-18464-0011-R | TCGA-LGG | GTEx | Normal | Cortex | Brain (Cortex) | 8.03 | 29.14 | 37.17 |
| GTEX-18A6Q-0011-F | TCGA-LGG | GTEx | Normal | Cortex | Brain (Cortex) | 9.4 | 31.14 | 40.54 |
| GTEX-1A3MX-0011-F | TCGA-LGG | GTEx | Normal | Cortex | Brain (Cortex) | 7.37 | 27.25 | 34.62 |
| GTEX-1A8G6-0011-F | TCGA-LGG | GTEx | Normal | Cortex | Brain (Cortex) | 6.07 | 17.87 | 23.94 |
| GTEX-1B8L1-0011-R | TCGA-LGG | GTEx | Normal | Cortex | Brain (Cortex) | 7.27 | 31.4 | 38.67 |
| GTEX-1B8SF-0011-F | TCGA-LGG | GTEx | Normal | Cortex | Brain (Cortex) | 7.86 | 31.19 | 39.05 |

| | | | | | | | | |
|---|---|---|---|---|---|---|---|---|
| GTEX-1B933-0011-R | TCGA-LGG | GTEx | Normal | Cortex | Brain (Cortex) | 10.2 | 26.6 | 36.8 |
| GTEX-1B996-0011-R | TCGA-LGG | GTEx | Normal | Cortex | Brain (Cortex) | 12.35 | 45 | 57.35 |
| GTEX-1BAJH-0011-F | TCGA-LGG | GTEx | Normal | Cortex | Brain (Cortex) | 4.28 | 19.99 | 24.27 |
| GTEX-1C6VQ-0011-I | TCGA-LGG | GTEx | Normal | Cortex | Brain (Cortex) | 6.83 | 13.91 | 20.74 |
| GTEX-1C6VR-0011-I | TCGA-LGG | GTEx | Normal | Cortex | Brain (Cortex) | 6.1 | 20.29 | 26.39 |
| GTEX-1CB4G-0011-I | TCGA-LGG | GTEx | Normal | Cortex | Brain (Cortex) | 7.78 | 34.66 | 42.44 |
| GTEX-1CB4H-0011-I | TCGA-LGG | GTEx | Normal | Cortex | Brain (Cortex) | 8.51 | 29.89 | 38.4 |
| GTEX-1CB4J-0011-R | TCGA-LGG | GTEx | Normal | Cortex | Brain (Cortex) | 18.39 | 59.3 | 77.69 |
| GTEX-1E1VI-0011-R | TCGA-LGG | GTEx | Normal | Cortex | Brain (Cortex) | 7.73 | 22.47 | 30.2 |
| GTEX-1EH9U-0011-I | TCGA-LGG | GTEx | Normal | Cortex | Brain (Cortex) | 4.78 | 17.61 | 22.39 |
| GTEX-1EMGI-0011-F | TCGA-LGG | GTEx | Normal | Cortex | Brain (Cortex) | 6.6 | 23.63 | 30.23 |
| GTEX-1EX96-0011-R | TCGA-LGG | GTEx | Normal | Cortex | Brain (Cortex) | 10.18 | 35.26 | 45.44 |
| GTEX-1F48J-0011-R | TCGA-LGG | GTEx | Normal | Cortex | Brain (Cortex) | 9.89 | 31.8 | 41.69 |
| GTEX-1F6I4-0011-R1 | TCGA-LGG | GTEx | Normal | Cortex | Brain (Cortex) | 8.28 | 27.85 | 36.13 |
| GTEX-1F6IF-0011-R1 | TCGA-LGG | GTEx | Normal | Cortex | Brain (Cortex) | 8.04 | 24.64 | 32.68 |
| GTEX-1F75W-0011-F | TCGA-LGG | GTEx | Normal | Cortex | Brain (Cortex) | 5.81 | 19.23 | 25.04 |
| GTEX-1F7RK-0011-F | TCGA-LGG | GTEx | Normal | Cortex | Brain (Cortex) | 6.59 | 22.33 | 28.92 |
| GTEX-1GF9V-0011-F | TCGA-LGG | GTEx | Normal | Cortex | Brain (Cortex) | 9.08 | 39.89 | 48.97 |
| GTEX-1GF9W-0011-I | TCGA-LGG | GTEx | Normal | Cortex | Brain (Cortex) | 8.61 | 33.63 | 42.24 |
| GTEX-1GMR8-0011- | TCGA-LGG | GTEx | Normal | Cortex | Brain (Cortex) | 7.1 | 28.14 | 35.24 |
| GTEX-1GMRU-0011- | TCGA-LGG | GTEx | Normal | Cortex | Brain (Cortex) | 7.78 | 31.17 | 38.95 |
| GTEX-1GN1U-0011-I | TCGA-LGG | GTEx | Normal | Cortex | Brain (Cortex) | 11.99 | 39.09 | 51.08 |
| GTEX-1GN1V-0011-F | TCGA-LGG | GTEx | Normal | Cortex | Brain (Cortex) | 2.37 | 10.45 | 12.82 |
| GTEX-1GN2E-0011-I | TCGA-LGG | GTEx | Normal | Cortex | Brain (Cortex) | 5.14 | 18.94 | 24.08 |
| GTEX-1GN73-0011-F | TCGA-LGG | GTEx | Normal | Cortex | Brain (Cortex) | 10.75 | 35.04 | 45.79 |
| GTEX-1GTWX-0011- | TCGA-LGG | GTEx | Normal | Cortex | Brain (Cortex) | 5.77 | 22.52 | 28.29 |
| GTEX-1GZ2Q-0011-I | TCGA-LGG | GTEx | Normal | Cortex | Brain (Cortex) | 6.44 | 27.32 | 33.76 |
| GTEX-1GZ4I-0011-R | TCGA-LGG | GTEx | Normal | Cortex | Brain (Cortex) | 8.73 | 27.81 | 36.54 |
| GTEX-1H1CY-0011-F | TCGA-LGG | GTEx | Normal | Cortex | Brain (Cortex) | 5.37 | 16.12 | 21.49 |
| GTEX-1H1DG-0011-I | TCGA-LGG | GTEx | Normal | Cortex | Brain (Cortex) | 7.99 | 28.06 | 36.05 |
| GTEX-1H1ZS-0011-F | TCGA-LGG | GTEx | Normal | Cortex | Brain (Cortex) | 7.97 | 28.08 | 36.05 |
| GTEX-1H23P-0011-R | TCGA-LGG | GTEx | Normal | Cortex | Brain (Cortex) | 4.65 | 21.39 | 26.04 |
| GTEX-1H3O1-0011-F | TCGA-LGG | GTEx | Normal | Cortex | Brain (Cortex) | 5.56 | 20.98 | 26.54 |
| GTEX-1H3VE-0011-F | TCGA-LGG | GTEx | Normal | Cortex | Brain (Cortex) | 7.76 | 19.57 | 27.33 |
| GTEX-1H4P4-0011-R | TCGA-LGG | GTEx | Normal | Cortex | Brain (Cortex) | 5.81 | 17.03 | 22.84 |
| GTEX-1HB9E-0011-F | TCGA-LGG | GTEx | Normal | Cortex | Brain (Cortex) | 9.08 | 31.47 | 40.55 |
| GTEX-1HBPH-0011-I | TCGA-LGG | GTEx | Normal | Cortex | Brain (Cortex) | 6.04 | 16.63 | 22.67 |
| GTEX-1HBPI-0011-R | TCGA-LGG | GTEx | Normal | Cortex | Brain (Cortex) | 7.54 | 28 | 35.54 |

| | | | | | | | | |
|---|---|---|---|---|---|---|---|---|
| GTEX-1HBPM-0011- | TCGA-LGG | GTEx | Normal | Cortex | Brain (Cortex) | 2.5 | 7.86 | 10.36 |
| GTEX-1HCU6-0011- | TCGA-LGG | GTEx | Normal | Cortex | Brain (Cortex) | 9.52 | 24.75 | 34.27 |
| GTEX-1HCVE-0011- | TCGA-LGG | GTEx | Normal | Cortex | Brain (Cortex) | 7.2 | 27.78 | 34.98 |
| GTEX-1HFI6-0011-R | TCGA-LGG | GTEx | Normal | Cortex | Brain (Cortex) | 11.15 | 30.45 | 41.6 |
| GTEX-1HGF4-0011- | TCGA-LGG | GTEx | Normal | Cortex | Brain (Cortex) | 8.73 | 23.17 | 31.9 |
| GTEX-1HR9M-0011- | TCGA-LGG | GTEx | Normal | Cortex | Brain (Cortex) | 4.99 | 14.22 | 19.21 |
| GTEX-1HSKV-0011- | TCGA-LGG | GTEx | Normal | Cortex | Brain (Cortex) | 6.12 | 22.03 | 28.15 |
| GTEX-1HSMO-0011- | TCGA-LGG | GTEx | Normal | Cortex | Brain (Cortex) | 6.34 | 19.02 | 25.36 |
| GTEX-1I1CD-0011-R | TCGA-LGG | GTEx | Normal | Cortex | Brain (Cortex) | 4.47 | 15.23 | 19.7 |
| GTEX-1I1GV-0011-R | TCGA-LGG | GTEx | Normal | Cortex | Brain (Cortex) | 5.09 | 25.7 | 30.79 |
| GTEX-1I1HK-0011-R | TCGA-LGG | GTEx | Normal | Cortex | Brain (Cortex) | 6.09 | 23.56 | 29.65 |
| GTEX-1ICLZ-0011-R | TCGA-LGG | GTEx | Normal | Cortex | Brain (Cortex) | 7.48 | 26.2 | 33.68 |
| GTEX-1IDJH-0011-R | TCGA-LGG | GTEx | Normal | Cortex | Brain (Cortex) | 6.59 | 27.76 | 34.35 |
| GTEX-1IKK5-0011-R | TCGA-LGG | GTEx | Normal | Cortex | Brain (Cortex) | 6.6 | 16.24 | 22.84 |
| GTEX-1IL2V-0011-R | TCGA-LGG | GTEx | Normal | Cortex | Brain (Cortex) | 4.47 | 6.02 | 10.49 |
| GTEX-1IOXB-0011-R | TCGA-LGG | GTEx | Normal | Cortex | Brain (Cortex) | 8.6 | 31.55 | 40.15 |
| GTEX-1J1OQ-0011- | TCGA-LGG | GTEx | Normal | Cortex | Brain (Cortex) | 7.25 | 26.92 | 34.17 |
| GTEX-1J8Q2-0011-R | TCGA-LGG | GTEx | Normal | Cortex | Brain (Cortex) | 7.14 | 27.12 | 34.26 |
| GTEX-1LC47-0011-R | TCGA-LGG | GTEx | Normal | Cortex | Brain (Cortex) | 6.28 | 21.89 | 28.17 |
| GTEX-N7MS-0011-R | TCGA-LGG | GTEx | Normal | Cortex | Brain (Cortex) | 7.96 | 23.58 | 31.54 |
| GTEX-N7MT-0011-R | TCGA-LGG | GTEx | Normal | Cortex | Brain (Cortex) | 8.07 | 27.41 | 35.48 |
| GTEX-NL3H-0011-R | TCGA-LGG | GTEx | Normal | Cortex | Brain (Cortex) | 7.56 | 31.11 | 38.67 |
| GTEX-NL4W-0011-R | TCGA-LGG | GTEx | Normal | Cortex | Brain (Cortex) | 8.32 | 23.13 | 31.45 |
| GTEX-NPJ7-0011-R1 | TCGA-LGG | GTEx | Normal | Cortex | Brain (Cortex) | 7.42 | 32.09 | 39.51 |
| GTEX-NPJ8-0011-R1 | TCGA-LGG | GTEx | Normal | Cortex | Brain (Cortex) | 11.36 | 39.79 | 51.15 |
| GTEX-OHPN-0011-R | TCGA-LGG | GTEx | Normal | Cortex | Brain (Cortex) | 4.67 | 13.34 | 18.01 |
| GTEX-OXRN-0011-R | TCGA-LGG | GTEx | Normal | Cortex | Brain (Cortex) | 5.93 | 20.05 | 25.98 |
| GTEX-OXRO-0011-R | TCGA-LGG | GTEx | Normal | Cortex | Brain (Cortex) | 5.69 | 15.98 | 21.67 |
| GTEX-P44G-0011-R | TCGA-LGG | GTEx | Normal | Cortex | Brain (Cortex) | 8.55 | 24.96 | 33.51 |
| GTEX-P44H-0011-R1 | TCGA-LGG | GTEx | Normal | Cortex | Brain (Cortex) | 7.32 | 32.58 | 39.9 |
| GTEX-PVOW-0011- | TCGA-LGG | GTEx | Normal | Cortex | Brain (Cortex) | 11.46 | 37.65 | 49.11 |
| GTEX-Q2AG-0011-R | TCGA-LGG | GTEx | Normal | Cortex | Brain (Cortex) | 8.27 | 26.02 | 34.29 |
| GTEX-QDT8-0011-R | TCGA-LGG | GTEx | Normal | Cortex | Brain (Cortex) | 13.24 | 35.49 | 48.73 |
| GTEX-QMR6-0011-R | TCGA-LGG | GTEx | Normal | Cortex | Brain (Cortex) | 9.9 | 33.15 | 43.05 |
| GTEX-QVJO-0011-R | TCGA-LGG | GTEx | Normal | Cortex | Brain (Cortex) | 8.72 | 28.17 | 36.89 |
| GTEX-QVUS-0011-R | TCGA-LGG | GTEx | Normal | Cortex | Brain (Cortex) | 8.37 | 35.78 | 44.15 |
| GTEX-S7SE-0011-R | TCGA-LGG | GTEx | Normal | Cortex | Brain (Cortex) | 7.13 | 23.29 | 30.42 |
| GTEX-T5JC-0011-R1 | TCGA-LGG | GTEx | Normal | Cortex | Brain (Cortex) | 8.54 | 25.68 | 34.22 |

| | | | | | | | | |
|---|---|---|---|---|---|---|---|---|
| GTEX-T6MN-0011-R | TCGA-LGG | GTEx | Normal | Cortex | Brain (Cortex) | 9.42 | 36.52 | 45.94 |
| GTEX-TSE9-0011-R1 | TCGA-LGG | GTEx | Normal | Cortex | Brain (Cortex) | 5.99 | 17.92 | 23.91 |
| GTEX-UTHO-0011-R | TCGA-LGG | GTEx | Normal | Cortex | Brain (Cortex) | 15.1 | 34.34 | 49.44 |
| GTEX-WHSE-0011-R | TCGA-LGG | GTEx | Normal | Cortex | Brain (Cortex) | 8.39 | 19.04 | 27.43 |
| GTEX-WL46-0011-R1 | TCGA-LGG | GTEx | Normal | Cortex | Brain (Cortex) | 7.55 | 33.85 | 41.4 |
| GTEX-WVLH-0011-R | TCGA-LGG | GTEx | Normal | Cortex | Brain (Cortex) | 11.45 | 37.03 | 48.48 |
| GTEX-WWYW-0011- | TCGA-LGG | GTEx | Normal | Cortex | Brain (Cortex) | 11.41 | 41.59 | 53 |
| GTEX-WZTO-0011-R | TCGA-LGG | GTEx | Normal | Cortex | Brain (Cortex) | 8.45 | 26.55 | 35 |
| GTEX-X261-0011-R1 | TCGA-LGG | GTEx | Normal | Cortex | Brain (Cortex) | 8.32 | 17.78 | 26.1 |
| GTEX-X4EP-0011-R1 | TCGA-LGG | GTEx | Normal | Cortex | Brain (Cortex) | 18.12 | 48.96 | 67.08 |
| GTEX-X4XX-0011-R1 | TCGA-LGG | GTEx | Normal | Cortex | Brain (Cortex) | 9.58 | 37.88 | 47.46 |
| GTEX-X4XY-0011-R1 | TCGA-LGG | GTEx | Normal | Cortex | Brain (Cortex) | 3.4 | 4.24 | 7.64 |
| GTEX-X585-0011-R1 | TCGA-LGG | GTEx | Normal | Cortex | Brain (Cortex) | 12.18 | 34.85 | 47.03 |
| GTEX-XLM4-0011-R1 | TCGA-LGG | GTEx | Normal | Cortex | Brain (Cortex) | 6.48 | 27.94 | 34.42 |
| GTEX-XMD1-0011-R | TCGA-LGG | GTEx | Normal | Cortex | Brain (Cortex) | 6.78 | 18.36 | 25.14 |
| GTEX-Y8DK-0011-R1 | TCGA-LGG | GTEx | Normal | Cortex | Brain (Cortex) | 6.47 | 27.95 | 34.42 |
| GTEX-YFC4-0011-R1 | TCGA-LGG | GTEx | Normal | Cortex | Brain (Cortex) | 8.36 | 27.09 | 35.45 |
| GTEX-YJ89-0011-R1 | TCGA-LGG | GTEx | Normal | Cortex | Brain (Cortex) | 7.68 | 24.56 | 32.24 |
| GTEX-Z93S-0011-R1 | TCGA-LGG | GTEx | Normal | Cortex | Brain (Cortex) | 6.45 | 23.76 | 30.21 |
| GTEX-ZAB4-0011-R1 | TCGA-LGG | GTEx | Normal | Cortex | Brain (Cortex) | 7.07 | 22.26 | 29.33 |
| GTEX-ZDXO-0011-R | TCGA-LGG | GTEx | Normal | Cortex | Brain (Cortex) | 9.93 | 24.61 | 34.54 |
| GTEX-ZE7O-0011-R1 | TCGA-LGG | GTEx | Normal | Cortex | Brain (Cortex) | 11.92 | 17.1 | 29.02 |
| GTEX-ZF28-0011-R1 | TCGA-LGG | GTEx | Normal | Cortex | Brain (Cortex) | 7.15 | 27.45 | 34.6 |
| GTEX-ZUA1-0011-R1 | TCGA-LGG | GTEx | Normal | Cortex | Brain (Cortex) | 5.17 | 21.51 | 26.68 |
| GTEX-ZV68-0011-R1 | TCGA-LGG | GTEx | Normal | Cortex | Brain (Cortex) | 3.92 | 11.82 | 15.74 |
| GTEX-ZVT3-0011-R1 | TCGA-LGG | GTEx | Normal | Cortex | Brain (Cortex) | 6.07 | 20.35 | 26.42 |
| GTEX-ZVZQ-0011-R1 | TCGA-LGG | GTEx | Normal | Cortex | Brain (Cortex) | 10.49 | 28.14 | 38.63 |
| GTEX-ZXG5-0011-R1 | TCGA-LGG | GTEx | Normal | Cortex | Brain (Cortex) | 4.97 | 11.7 | 16.67 |

| Table S4. MAGOH/MAGOHB evaluation by immunostaining in glioma cohort from the Shanghai ChangZheng Hospital | | | | | | | |
|---|---|---|---|---|---|---|---|
| **who grade** | **low/high grade** | **ihc staing** | **ihc staing grade** | **dead/live** | **months** | **chemotherapy** | **radiotherapy** |
| 1 | 1 | 1 | 1 | 0 | 27 | no | no |
| 1 | 1 | 2 | 1 | 0 | 28 | no | no |
| 1 | 1 | 1 | 1 | 0 | 54 | no | no |
| 1 | 1 | 1 | 1 | 0 | 29 | no | yes |
| 1 | 1 | 1 | 1 | 0 | 28 | no | yes |
| 1 | 1 | 1 | 1 | 1 | 34 | no | yes |
| 1 | 1 | 1 | 1 | 0 | 35 | no | yes |
| 1 | 1 | 2 | 1 | 1 | 36 | no | yes |
| 1 | 1 | 1 | 1 | 0 | 47 | no | yes |
| 1 | 1 | 2 | 1 | 0 | 46 | yes | yes |
| 1 | 1 | 3 | 2 | 0 | 35 | no | yes |
| 1 | 1 | 3 | 2 | 0 | 45 | no | yes |
| 1 | 1 | 3 | 2 | 0 | 45 | no | yes |
| 1 | 1 | 3 | 2 | 0 | 37 | yes | yes |
| 2 | 1 | 1 | 1 | 1 | 13 | yes | no |
| 2 | 1 | 1 | 1 | 1 | 30 | no | yes |
| 2 | 1 | 1 | 1 | 0 | 6 | no | yes |
| 2 | 1 | 1 | 1 | 0 | 7 | no | yes |
| 2 | 1 | 1 | 1 | 0 | 9 | no | yes |
| 2 | 1 | 1 | 1 | 0 | 45 | no | yes |
| 2 | 1 | 1 | 1 | 0 | 21 | no | yes |
| 2 | 1 | 1 | 1 | 0 | 33 | no | yes |
| 2 | 1 | 1 | 1 | 0 | 32 | no | yes |
| 2 | 1 | 1 | 1 | 0 | 25 | yes | yes |
| 2 | 1 | 1 | 1 | 0 | 57 | yes | yes |
| 2 | 1 | 1 | 1 | 1 | 53 | yes | yes |
| 2 | 1 | 2 | 1 | 0 | 49 | yes | yes |
| 2 | 1 | 1 | 1 | 0 | 32 | yes | yes |

| | | | | | | | |
|---|---|---|---|---|---|---|---|
| 2 | 1 | 2 | 1 | 0 | 45 | yes | yes |
| 2 | 1 | 2 | 1 | 0 | 26 | yes | yes |
| 2 | 1 | 1 | 1 | 0 | 48 | yes | yes |
| 2 | 1 | 1 | 1 | 0 | 23 | yes | yes |
| 2 | 1 | 2 | 1 | 0 | 13 | yes | yes |
| 2 | 1 | 3 | 2 | 1 | 1 | no | no |
| 2 | 1 | 4 | 2 | 0 | 3 | no | no |
| 2 | 1 | 4 | 2 | 1 | 1 | no | no |
| 2 | 1 | 3 | 2 | 1 | 21 | yes | no |
| 2 | 1 | 3 | 2 | 1 | 33 | yes | no |
| 2 | 1 | 3 | 2 | 0 | 57 | no | yes |
| 2 | 1 | 4 | 2 | 0 | 38 | yes | yes |
| 2 | 1 | 3 | 2 | 0 | 46 | yes | yes |
| 3 | 2 | 2 | 1 | 1 | 18 | no | no |
| 3 | 2 | 2 | 1 | 1 | 22 | yes | no |
| 3 | 2 | 1 | 1 | 0 | 27 | yes | yes |
| 3 | 2 | 2 | 1 | 0 | 18 | yes | yes |
| 3 | 2 | 1 | 1 | 0 | 30 | yes | yes |
| 3 | 2 | 2 | 1 | 0 | 6 | yes | yes |
| 3 | 2 | 1 | 1 | 0 | 12 | yes | yes |
| 3 | 2 | 3 | 2 | 1 | 18 | no | no |
| 3 | 2 | 3 | 2 | 1 | 4 | no | no |
| 3 | 2 | 3 | 2 | 1 | 11 | no | no |
| 3 | 2 | 4 | 2 | 1 | 13 | yes | no |
| 3 | 2 | 3 | 2 | 1 | 29 | yes | no |
| 3 | 2 | 3 | 2 | 1 | 16 | yes | no |
| 3 | 2 | 4 | 2 | 1 | 12 | yes | no |
| 3 | 2 | 3 | 2 | 1 | 12 | yes | no |
| 3 | 2 | 3 | 2 | 1 | 18 | no | yes |
| 3 | 2 | 3 | 2 | 0 | 18 | no | yes |

| | | | | | | | |
|---|---|---|---|---|---|---|---|
| 3 | 2 | 3 | 2 | 0 | 12 | yes | yes |
| 4 | 2 | 2 | 1 | 1 | 6 | no | no |
| 4 | 2 | 2 | 1 | 1 | 10 | no | no |
| 4 | 2 | 2 | 1 | 1 | 4 | no | no |
| 4 | 2 | 2 | 1 | 0 | 1 | no | no |
| 4 | 2 | 2 | 1 | 1 | 5 | no | no |
| 4 | 2 | 1 | 1 | 1 | 13 | no | no |
| 4 | 2 | 2 | 1 | 1 | 1 | no | no |
| 4 | 2 | 2 | 1 | 0 | 8 | no | no |
| 4 | 2 | 2 | 1 | 1 | 9 | no | no |
| 4 | 2 | 2 | 1 | 1 | 18 | yes | no |
| 4 | 2 | 2 | 1 | 0 | 12 | yes | no |
| 4 | 2 | 2 | 1 | 1 | 3 | yes | no |
| 4 | 2 | 2 | 1 | 1 | 26 | no | yes |
| 4 | 2 | 2 | 1 | 0 | 12 | no | yes |
| 4 | 2 | 2 | 1 | 1 | 13 | no | yes |
| 4 | 2 | 2 | 1 | 1 | 15 | no | yes |
| 4 | 2 | 2 | 1 | 1 | 2 | no | yes |
| 4 | 2 | 1 | 1 | 1 | 37 | no | yes |
| 4 | 2 | 2 | 1 | 1 | 14 | no | yes |
| 4 | 2 | 2 | 1 | 1 | 12 | no | yes |
| 4 | 2 | 2 | 1 | 1 | 4 | no | yes |
| 4 | 2 | 2 | 1 | 1 | 5 | no | yes |
| 4 | 2 | 1 | 1 | 1 | 12 | no | yes |
| 4 | 2 | 1 | 1 | 1 | 24 | no | yes |
| 4 | 2 | 1 | 1 | 1 | 25 | no | yes |
| 4 | 2 | 2 | 1 | 1 | 11 | no | yes |
| 4 | 2 | 2 | 1 | 0 | 1 | no | yes |
| 4 | 2 | 1 | 1 | 0 | 25 | yes | yes |
| 4 | 2 | 2 | 1 | 0 | 45 | yes | yes |

| | | | | | | | |
|---|---|---|---|---|---|---|---|
| 4 | 2 | 2 | 1 | 0 | 9 | yes | yes |
| 4 | 2 | 2 | 1 | 0 | 27 | yes | yes |
| 4 | 2 | 2 | 1 | 0 | 14 | yes | yes |
| 4 | 2 | 2 | 1 | 0 | 25 | yes | yes |
| 4 | 2 | 2 | 1 | 0 | 9 | yes | yes |
| 4 | 2 | 2 | 1 | 0 | 3 | yes | yes |
| 4 | 2 | 2 | 1 | 1 | 14 | yes | yes |
| 4 | 2 | 2 | 1 | 1 | 28 | yes | yes |
| 4 | 2 | 2 | 1 | 1 | 11 | yes | yes |
| 4 | 2 | 2 | 1 | 1 | 2 | yes | yes |
| 4 | 2 | 2 | 1 | 1 | 9 | yes | yes |
| 4 | 2 | 2 | 1 | 0 | 3 | yes | yes |
| 4 | 2 | 1 | 1 | 0 | 3 | yes | yes |
| 4 | 2 | 1 | 1 | 1 | 10 | yes | yes |
| 4 | 2 | 2 | 1 | 0 | 10 | yes | yes |
| 4 | 2 | 2 | 1 | 1 | 2 | yes | yes |
| 4 | 2 | 2 | 1 | 1 | 4 | yes | yes |
| 4 | 2 | 3 | 2 | 1 | 18 | no | no |
| 4 | 2 | 3 | 2 | 1 | 14 | no | no |
| 4 | 2 | 3 | 2 | 1 | 10 | no | no |
| 4 | 2 | 3 | 2 | 1 | 2 | no | no |
| 4 | 2 | 3 | 2 | 1 | 10 | no | no |
| 4 | 2 | 3 | 2 | 1 | 14 | no | no |
| 4 | 2 | 4 | 2 | 1 | 31 | no | no |
| 4 | 2 | 3 | 2 | 1 | 4 | no | no |
| 4 | 2 | 3 | 2 | 1 | 2 | no | no |
| 4 | 2 | 4 | 2 | 0 | 2 | no | no |
| 4 | 2 | 4 | 2 | 0 | 14 | no | no |
| 4 | 2 | 3 | 2 | 0 | 1 | no | no |
| 4 | 2 | 4 | 2 | 0 | 6 | yes | no |

| | | | | | | | |
|---|---|---|---|---|---|---|---|
| 4 | 2 | 4 | 2 | 1 | 23 | yes | no |
| 4 | 2 | 4 | 2 | 1 | 15 | yes | no |
| 4 | 2 | 3 | 2 | 1 | 17 | yes | no |
| 4 | 2 | 3 | 2 | 1 | 9 | yes | no |
| 4 | 2 | 3 | 2 | 1 | 14 | yes | no |
| 4 | 2 | 3 | 2 | 1 | 10 | yes | no |
| 4 | 2 | 3 | 2 | 1 | 5 | yes | no |
| 4 | 2 | 4 | 2 | 0 | 17 | yes | no |
| 4 | 2 | 4 | 2 | 0 | 8 | yes | no |
| 4 | 2 | 3 | 2 | 1 | 2 | yes | no |
| 4 | 2 | 4 | 2 | 1 | 12 | no | yes |
| 4 | 2 | 4 | 2 | 1 | 34 | no | yes |
| 4 | 2 | 3 | 2 | 0 | 36 | no | yes |
| 4 | 2 | 3 | 2 | 1 | 14 | no | yes |
| 4 | 2 | 3 | 2 | 1 | 38 | no | yes |
| 4 | 2 | 3 | 2 | 1 | 31 | no | yes |
| 4 | 2 | 3 | 2 | 0 | 3 | no | yes |
| 4 | 2 | 3 | 2 | 1 | 13 | no | yes |
| 4 | 2 | 3 | 2 | 1 | 12 | no | yes |
| 4 | 2 | 3 | 2 | 1 | 9 | no | yes |
| 4 | 2 | 3 | 2 | 1 | 10 | no | yes |
| 4 | 2 | 3 | 2 | 0 | 9 | no | yes |
| 4 | 2 | 3 | 2 | 0 | 1 | no | yes |
| 4 | 2 | 3 | 2 | 0 | 12 | no | yes |
| 4 | 2 | 4 | 2 | 1 | 26 | no | yes |
| 4 | 2 | 4 | 2 | 0 | 45 | no | yes |
| 4 | 2 | 4 | 2 | 1 | 12 | no | yes |
| 4 | 2 | 3 | 2 | 1 | 13 | no | yes |
| 4 | 2 | 3 | 2 | 1 | 6 | no | yes |
| 4 | 2 | 3 | 2 | 1 | 5 | no | yes |

| | | | | | | | |
|---|---|---|---|---|---|---|---|
| 4 | 2 | 4 | 2 | 0 | 3 | no | yes |
| 4 | 2 | 3 | 2 | 1 | 13 | no | yes |
| 4 | 2 | 3 | 2 | 1 | 12 | no | yes |
| 4 | 2 | 4 | 2 | 1 | 37 | no | yes |
| 4 | 2 | 4 | 2 | 1 | 3 | no | yes |
| 4 | 2 | 4 | 2 | 1 | 7 | no | yes |
| 4 | 2 | 3 | 2 | 1 | 10 | no | yes |
| 4 | 2 | 3 | 2 | 0 | 6 | no | yes |
| 4 | 2 | 4 | 2 | 1 | 28 | yes | yes |
| 4 | 2 | 3 | 2 | 0 | 3 | yes | yes |
| 4 | 2 | 3 | 2 | 0 | 8 | yes | yes |
| 4 | 2 | 3 | 2 | 1 | 5 | yes | yes |
| 4 | 2 | 3 | 2 | 1 | 25 | yes | yes |
| 4 | 2 | 3 | 2 | 1 | 7 | yes | yes |
| 4 | 2 | 3 | 2 | 0 | 8 | yes | yes |
| 4 | 2 | 3 | 2 | 1 | 2 | yes | yes |
| 4 | 2 | 4 | 2 | 1 | 38 | yes | yes |
| 4 | 2 | 4 | 2 | 0 | 5 | yes | yes |
| 4 | 2 | 4 | 2 | 1 | 12 | yes | yes |
| 4 | 2 | 3 | 2 | 1 | 15 | yes | yes |
| 4 | 2 | 3 | 2 | 1 | 15 | yes | yes |
| 4 | 2 | 3 | 2 | 1 | 1 | yes | yes |
| 4 | 2 | 3 | 2 | 1 | 9 | yes | yes |
| 4 | 2 | 4 | 2 | 1 | 13 | yes | yes |
| 4 | 2 | 4 | 2 | 1 | 12 | yes | yes |
| 4 | 2 | 4 | 2 | 1 | 5 | yes | yes |
| 4 | 2 | 4 | 2 | 0 | 9 | yes | yes |
| 4 | 2 | 3 | 2 | 1 | 9 | yes | yes |
| 4 | 2 | 3 | 2 | 1 | 11 | yes | yes |

| Table S5. Gene Ontology (GO) enrichment analyses of genes displaying splicing alterations in MAGOH/MAGOHB KD cells | | | | | | | |
|---|---|---|---|---|---|---|---|
| **GO biological process complete** | **Homo sapiens - REFLIST (20589)** | **upload_1 (155)** | **upload_1 (expected)** | **upload_1 (over/under)** | **upload_1 (fold Enrichment)** | **upload_1 (raw P-value)** | **upload_1 (FDR)** |
| meiotic chromosome condensation (GO: 0010032) | 6 | 3 | 0.05 | + | 66.42 | 3.33E-05 | 2.08E-02 |
| regulation of chromosome condensation (GO: 0060623) | 10 | 3 | 0.08 | + | 39.85 | 1.11E-04 | 4.34E-02 |
| chromosome segregation (GO:0007059) | 286 | 11 | 2.15 | + | 5.11 | 1.50E-05 | 1.18E-02 |
| regulation of chromosome separation (GO:1905818) | 109 | 8 | 0.82 | + | 9.75 | 2.78E-06 | 6.22E-03 |
| mitotic sister chromatid segregation (GO: 0000070) | 119 | 7 | 0.9 | + | 7.81 | 4.56E-05 | 2.65E-02 |
| cell division (GO:0051301) | 508 | 15 | 3.82 | + | 3.92 | 9.24E-06 | 9.65E-03 |
| positive regulation of cellular amide metabolic process (GO:0034250) | 168 | 8 | 1.26 | + | 6.33 | 5.46E-05 | 2.76E-02 |
| regulation of cellular amide metabolic process (GO:0034248) | 460 | 14 | 3.46 | + | 4.04 | 1.34E-05 | 1.11E-02 |
| peptide biosynthetic process (GO:0043043) | 404 | 12 | 3.04 | + | 3.95 | 7.10E-05 | 3.27E-02 |
| regulation of cell cycle process (GO:0010564) | 727 | 19 | 5.47 | + | 3.47 | 3.18E-06 | 4.52E-03 |
| regulation of cell cycle (GO:0051726) | 1122 | 25 | 8.45 | + | 2.96 | 1.35E-06 | 4.22E-03 |
| regulation of organelle organization (GO: 0033043) | 1185 | 24 | 8.92 | + | 2.69 | 1.10E-05 | 1.01E-02 |
| organelle organization (GO:0006996) | 3424 | 52 | 25.78 | + | 2.02 | 3.61E-07 | 1.89E-03 |
| RNA metabolic process (GO:0016070) | 1442 | 25 | 10.86 | + | 2.3 | 9.96E-05 | 4.11E-02 |
| RNA processing (GO: 0006396) | 858 | 18 | 6.46 | + | 2.79 | 9.92E-05 | 4.20E-02 |
| translational initiation (GO: 0006413) | 70 | 7 | 0.53 | + | 13.28 | 1.76E-06 | 4.60E-03 |
| regulation of translation (GO:0006417) | 402 | 14 | 3.03 | + | 4.63 | 3.04E-06 | 5.30E-03 |
| translation (GO:0006412) | 377 | 12 | 2.84 | + | 4.23 | 3.73E-05 | 2.25E-02 |
| positive regulation of translation (GO:0045727) | 142 | 8 | 1.07 | + | 7.48 | 1.74E-05 | 1.30E-02 |

| Table S6. Exon-skipping eventis MAGOH/MAGOHB KD cells. | | | | | | | | | | | | | | | | | | | | | | |
|---|---|---|---|---|---|---|---|---|---|---|---|---|---|---|---|---|---|---|---|---|---|---|
| GeneID | geneSymbol | chr | exonStart_0base | exonEnd | upstreamES | upstreamEE | downstreamES | downstreamEE | IJC_SAMPLE_1 | SJC_SAMPLE_1 | IJC_SAMPLE_2 | SJC_SAMPLE_2 | IncFormLen | SkipFormLen | PValue | FDR | IncLevel1 | IncLevel2 | IncLevelDifferenc | gene_id | exon_list | skipped_exons |
| ENSG00000198 | EIF1AY | chrY | 20579607 | 20579691 | 20575724 | 20575887 | 20584473 | 20584524 | 35,34,38 | 0,0,0 | 32,92,45 | 13,10,5 | 234 | 150 | 8.16E-09 | 3.70E-07 | 1.0,1.0,1.0 | 0.612,0.855,0.85 | 0.227 | ENSG00000198 | [[[[20575776, 20 | 2 |
| ENSG00000198 | EIF1AY | chrY | 20582589 | 20582693 | 20575724 | 20575887 | 20584473 | 20584524 | 71,63,82 | 0,0,0 | 51,90,38 | 13,10,5 | 254 | 150 | 1.09E-11 | 8.29E-10 | 1.0,1.0,1.0 | 0.699,0.842,0.81 | 0.214 | ENSG00000198 | [[[[20575776, 20 | 2 |
| ENSG00000067 | DDX3Y | chrY | 12914563 | 12914649 | 12912726 | 12912882 | 12914883 | 12914982 | 42,24,67 | 0,0,0 | 31,45,29 | 1,4,3 | 236 | 150 | 0.000622734 | 0.007751124 | 1.0,1.0,1.0 | 0.952,0.877,0.86 | 0.104 | ENSG00000067 | [[[[12904868, 12 | 3 |
| ENSG00000067 | DDX3Y | chrY | 12916260 | 12916442 | 12915066 | 12915227 | 12916516 | 12916634 | 70,63,106 | 0,0,0 | 43,138,74 | 6,14,8 | 300 | 150 | 9.55E-11 | 6.29E-09 | 1.0,1.0,1.0 | 0.782,0.831,0.82 | 0.188 | ENSG00000067 | [[[[12904868, 12 | 3 |
| ENSG00000067 | IDH3G | chrX | 153786800 | 153786947 | 153785766 | 153786449 | 153787463 | 153787553 | 77,85,148 | 0,1,4 | 115,243,114 | 7,6,14 | 297 | 150 | 0.00042135 | 0.005541252 | 1.0,0.977,0.949 | 0.892,0.953,0.80 | 0.092 | ENSG00000067 | [[[[153794246, 1 | 2 |
| ENSG00000067 | IDH3G | chrX | 153787050 | 153787153 | 153785766 | 153786449 | 153787463 | 153787553 | 32,90,154 | 0,1,4 | 115,195,138 | 7,6,14 | 253 | 150 | 0.000640507 | 0.007934536 | 1.0,0.982,0.958 | 0.907,0.951,0.85 | 0.076 | ENSG00000067 | [[[[153794246, 1 | 2 |
| ENSG00000156 | PHF6 | chrX | 134377571 | 134377755 | 134373252 | 134373467 | 134393500 | 134393634 | 82,59,91 | 0,0,0 | 44,121,72 | 10,17,9 | 300 | 150 | 3.27E-13 | 3.15E-11 | 1.0,1.0,1.0 | 0.688,0.781,0.8 | 0.244 | ENSG00000156 | [[[[134373312, 1 | 2 |
| ENSG00000158 | CUL4B | chrX | 120532421 | 120532594 | 120530101 | 120530254 | 120536926 | 120537034 | 85,60,90 | 0,0,0 | 46,126,71 | 15,13,7 | 300 | 150 | 9.23E-12 | 7.09E-10 | 1.0,1.0,1.0 | 0.605,0.829,0.83 | 0.244 | ENSG00000158 | [[[[120560083, 1 | 3 |
| ENSG00000158 | CUL4B | chrX | 120535829 | 120535943 | 120532335 | 120532594 | 120536926 | 120537034 | 61,63,69 | 0,0,0 | 49,85,54 | 4,2,4 | 264 | 150 | 0.000280168 | 0.00389069 | 1.0,1.0,1.0 | 0.874,0.96,0.885 | 0.094 | ENSG00000158 | [[[[120560083, 1 | 2 |
| ENSG00000125 | SEPT6 | chrX | 119625334 | 119625379 | 119615723 | 119616742 | 119629317 | 119629508 | 0,0,0 | 2,6,3 | 0,3,3 | 1,0,1 | 195 | 150 | 6.09E-06 | 0.000138244 | 0.0,0.0,0.0 | 0.0,1.0,0.698 | -0.566 | ENSG00000125 | [[[[119693076, 1 | 2 |
| ENSG00000131 | IL13RA1 | chrX | 118776426 | 118776511 | 118766843 | 118766976 | 118791761 | 118794539 | 13,10,13 | 0,0,0 | 7,20,15 | 2,13,9 | 235 | 150 | 3.48E-10 | 2.10E-08 | 1.0,1.0,1.0 | 0.691,0.495,0.51 | 0.433 | ENSG00000131 | [[[[118727606, 1 | 2 |
| ENSG00000068 | ACSL4 | chrX | 109659353 | 109659511 | 109641336 | 109644186 | 109663210 | 109663402 | 79,43,102 | 0,0,0 | 30,58,31 | 5,12,6 | 300 | 150 | 5.81E-13 | 5.45E-11 | 1.0,1.0,1.0 | 0.75,0.707,0.721 | 0.274 | ENSG00000068 | [[[[109733139, 1 | 2 |
| ENSG00000147 | OGT | chrX | 71555953 | 71556094 | 71554512 | 71554592 | 71557492 | 71557672 | 21,7,30 | 0,0,0 | 13,36,18 | 1,5,2 | 291 | 150 | 7.64E-05 | 0.00125887 | 1.0,1.0,1.0 | 0.87,0.788,0.823 | 0.173 | ENSG00000147 | [[[[71533104, 71 | 5 |
| ENSG00000102 | RBM3 | chrX | 48575814 | 48576083 | 48575560 | 48575667 | 48577025 | 48577109 | 1,2,3 | 0,0,0 | 17,23,16 | 2,6,8 | 300 | 150 | 3.98E-07 | 1.20E-05 | 1.0,1.0,1.0 | 0.81,0.657,0.5 | 0.344 | ENSG00000102 | [[[[48574484, 48 | 2 |
| ENSG00000102 | CDK16 | chrX | 47226582 | 47226703 | 47226095 | 47226402 | 47226993 | 47227099 | 100,113,234 | 0,0,0 | 181,358,215 | 4,11,6 | 271 | 150 | 5.32E-05 | 0.000922096 | 1.0,1.0,1.0 | 0.962,0.947,0.95 | 0.046 | ENSG00000102 | [[[[47218681, 47 | 2 |
| ENSG00000196 | WDR45 | chrX | 49076644 | 49076750 | 49075865 | 49075945 | 49077642 | 49077746 | 12,19,23 | 0,0,0 | 11,22,19 | 1,4,3 | 256 | 150 | 4.15E-05 | 0.000743223 | 1.0,1.0,1.0 | 0.866,0.763,0.78 | 0.194 | ENSG00000196 | [[[[49079751, 49 | 2 |
| ENSG00000196 | WDR45 | chrX | 49076644 | 49076750 | 49075865 | 49075945 | 49077836 | 49077911 | 3,2,2 | 0,0,0 | 8,9,8 | 3,2,1 | 256 | 150 | 0.000240475 | 0.00340653 | 1.0,1.0,1.0 | 0.61,0.725,0.824 | 0.28 | ENSG00000196 | [[[[49079751, 49 | 3 |
| ENSG00000215 | DDX3X | chrX | 41342736 | 41342836 | 41340011 | 41342653 | 41343736 | 41343822 | 91,60,167 | 0,0,0 | 87,147,119 | 3,17,10 | 250 | 150 | 5.29E-10 | 3.09E-08 | 1.0,1.0,1.0 | 0.946,0.838,0.87 | 0.113 | ENSG00000215 | [[[[41334162, 41 | 2 |
| ENSG00000215 | DDX3X | chrX | 41343215 | 41343351 | 41340005 | 41342836 | 41344029 | 41344128 | 174,95,224 | 0,0,0 | 91,300,255 | 3,7,7 | 286 | 150 | 0.000135911 | 0.002076209 | 1.0,1.0,1.0 | 0.941,0.957,0.95 | 0.051 | ENSG00000215 | [[[[41334162, 41 | 2 |
| ENSG00000215 | DDX3X | chrX | 41343736 | 41343822 | 41342736 | 41342836 | 41344029 | 41344109 | 76,63,176 | 0,0,0 | 71,207,113 | 3,7,7 | 236 | 150 | 2.32E-05 | 0.000449325 | 1.0,1.0,1.0 | 0.938,0.949,0.91 | 0.067 | ENSG00000215 | [[[[41334162, 41 | 2 |
| ENSG00000124 | USP9X | chrX | 41225048 | 41225137 | 41223216 | 41223402 | 41229252 | 41229409 | 36,24,31 | 0,0,0 | 20,23,13 | 2,9,3 | 239 | 150 | 2.10E-08 | 8.71E-07 | 1.0,1.0,1.0 | 0.863,0.616,0.73 | 0.263 | ENSG00000124 | [[[[41085445, 41 | 2 |
| ENSG00000182 | ATP6AP2 | chrX | 40597526 | 40597664 | 40591233 | 40591365 | 40598680 | 40598734 | 586,430,731 | 5,3,7 | 5,171,038,648 | 9,35,18 | 288 | 150 | 5.36E-06 | 0.000123408 | 0.984,0.987,0.98 | 0.968,0.939,0.94 | 0.032 | ENSG00000182 | [[[[40580970, 40 | 2 |
| ENSG00000130 | EIF2S3 | chrX | 24057632 | 24057754 | 24055614 | 24055678 | 24060087 | 24060182 | 158,134,234 | 0,0,0 | 210,356,277 | 11,20,12 | 272 | 150 | 2.58E-10 | 1.59E-08 | 1.0,1.0,1.0 | 0.913,0.908,0.92 | 0.084 | ENSG00000130 | [[[[24054956, 24 | 2 |
| ENSG00000130 | EIF2S3 | chrX | 24064200 | 24064335 | 24062415 | 24062574 | 24076721 | 24077971 | 277,188,343 | 0,0,0 | 200,424,306 | 35,63,49 | 285 | 150 | 0 | 0 | 1.0,1.0,1.0 | 0.75,0.78,0.767 | 0.234 | ENSG00000130 | [[[[24054956, 24 | 5 |
| ENSG00000130 | EIF2S3 | chrX | 24065997 | 24066092 | 24064200 | 24064335 | 24076721 | 24077971 | 141,114,167 | 0,0,0 | 145,267,181 | 2,7,11 | 245 | 150 | 2.52E-05 | 0.000481344 | 1.0,1.0,1.0 | 0.978,0.959,0.91 | 0.051 | ENSG00000130 | [[[[24054956, 24 | 4 |
| ENSG00000130 | EIF2S3 | chrX | 24067963 | 24068108 | 24062415 | 24062574 | 24071557 | 24071727 | 323,272,434 | 0,0,0 | 132,309,157 | 2,9,6 | 295 | 150 | 1.22E-05 | 0.000254394 | 1.0,1.0,1.0 | 0.971,0.946,0.93 | 0.051 | ENSG00000130 | [[[[24054956, 24 | 3 |
| ENSG00000130 | EIF2S3 | chrX | 24067963 | 24068108 | 24065997 | 24066092 | 24076721 | 24077971 | 263,190,283 | 0,0,0 | 104,256,138 | 60,86,71 | 295 | 150 | 0 | 0 | 1.0,1.0,1.0 | 0.468,0.602,0.49 | 0.478 | ENSG00000130 | [[[[24054956, 24 | 3 |
| ENSG00000130 | EIF2S3 | chrX | 24071557 | 24071727 | 24060087 | 24060182 | 24076721 | 24077971 | 10,2,7 | 0,0,0 | 13,25,20 | 4,1,4 | 300 | 150 | 7.90E-06 | 0.000173804 | 1.0,1.0,1.0 | 0.619,0.926,0.71 | 0.247 | ENSG00000130 | [[[[24054956, 24 | 6 |
| ENSG00000130 | EIF2S3 | chrX | 24071557 | 24071727 | 24062415 | 24062574 | 24076721 | 24077971 | 10,2,7 | 0,0,0 | 15,34,25 | 35,63,49 | 300 | 150 | 7.69E-14 | 8.09E-12 | 1.0,1.0,1.0 | 0.176,0.212,0.20 | 0.803 | ENSG00000130 | [[[[24054956, 24 | 5 |
| ENSG00000130 | EIF2S3 | chrX | 24071557 | 24071727 | 24064200 | 24064335 | 24076721 | 24077971 | 10,2,7 | 0,0,0 | 13,28,19 | 2,7,11 | 300 | 150 | 1.21E-07 | 4.13E-06 | 1.0,1.0,1.0 | 0.765,0.667,0.46 | 0.368 | ENSG00000130 | [[[[24054956, 24 | 4 |
| ENSG00000130 | EIF2S3 | chrX | 24071557 | 24071727 | 24065997 | 24066092 | 24076721 | 24077971 | 10,4,7 | 0,0,0 | 14,27,20 | 60,86,71 | 300 | 150 | 0 | 0 | 1.0,1.0,1.0 | 0.104,0.136,0.12 | 0.879 | ENSG00000130 | [[[[24054956, 24 | 3 |
| ENSG00000130 | EIF2S3 | chrX | 24071557 | 24071727 | 24067963 | 24068108 | 24076721 | 24077971 | 344,282,447 | 0,0,0 | 146,339,183 | 14,41,18 | 300 | 150 | 0 | 0 | 1.0,1.0,1.0 | 0.839,0.805,0.83 | 0.173 | ENSG00000130 | [[[[24054956, 24 | 2 |
| ENSG00000123 | ACOT9 | chrX | 23707876 | 23707944 | 23706627 | 23706739 | 23730526 | 23730564 | 35,33,35 | 0,0,0 | 11,31,31 | 8,7,2 | 218 | 150 | 4.58E-08 | 1.74E-06 | 1.0,1.0,1.0 | 0.486,0.753,0.91 | 0.282 | ENSG00000123 | [[[[23743125, 23 | 4 |
| ENSG00000177 | RPS6KA3 | chrX | 20164898 | 20164961 | 20162963 | 20163040 | 20169401 | 20169491 | 31,34,37 | 0,0,0 | 18,51,17 | 5,8,10 | 213 | 150 | 3.20E-10 | 1.94E-08 | 1.0,1.0,1.0 | 0.717,0.818,0.54 | 0.307 | ENSG00000177 | [[[[20266564, 20 | 2 |
| ENSG00000173 | EIF1AX | chrX | 20135737 | 20135841 | 20133956 | 20134007 | 20141624 | 20141826 | 265,206,322 | 0,0,0 | 231,487,249 | 16,33,25 | 254 | 150 | 0 | 0 | 1.0,1.0,1.0 | 0.895,0.897,0.85 | 0.118 | ENSG00000173 | [[[[20141625, 20 | 2 |
| ENSG00000044 | PHKA2 | chrX | 18901484 | 18901603 | 18897162 | 18897333 | 18905757 | 18905859 | 19,14,11 | 0,0,0 | 10,20,9 | 3,4,0 | 269 | 150 | 0.000141675 | 0.002153048 | 1.0,1.0,1.0 | 0.65,0.736,1.0 | 0.205 | ENSG00000044 | [[[[18983855, 18 | 3 |
| ENSG00000044 | PHKA2 | chrX | 18948743 | 18948826 | 18945077 | 18945158 | 18952493 | 18952541 | 20,3,6 | 0,0,0 | 6,16,7 | 2,5,0 | 233 | 150 | 0.000133646 | 0.002048599 | 1.0,1.0,1.0 | 0.659,0.673,1.0 | 0.223 | ENSG00000044 | [[[[18983855, 18 | 2 |
| ENSG00000102 | RBBP7 | chrX | 16852550 | 16852628 | 16852045 | 16852122 | 16853681 | 16853842 | 101,76,104 | 0,0,0 | 48,117,74 | 4,18,6 | 228 | 150 | 4.45E-11 | 3.07E-09 | 1.0,1.0,1.0 | 0.888,0.81,0.89 | 0.137 | ENSG00000102 | [[[[16870038, 16 | 2 |
| ENSG00000102 | RBBP7 | chrX | 16852748 | 16852875 | 16852045 | 16852122 | 16853681 | 16853842 | 290,237,328 | 0,0,0 | 239,476,250 | 4,18,6 | 277 | 150 | 3.97E-07 | 1.20E-05 | 1.0,1.0,1.0 | 0.97,0.935,0.958 | 0.046 | ENSG00000102 | [[[[16870038, 16 | 2 |
| ENSG00000047 | CTPS2 | chrX | 16609540 | 16609685 | 16590751 | 16590862 | 16620276 | 16620332 | 14,9,14 | 0,0,0 | 7,23,6 | 2,0,6 | 295 | 150 | 3.64E-05 | 0.000663823 | 1.0,1.0,1.0 | 0.64,1.0,0.337 | 0.341 | ENSG00000047 | [[[[16712335, 16 | 2 |
| ENSG00000047 | CTPS2 | chrX | 16617149 | 16617246 | 16590751 | 16590862 | 16620276 | 16620332 | 7,2,15 | 0,0,0 | 8,19,6 | 2,0,6 | 247 | 150 | 5.84E-05 | 0.000999195 | 1.0,1.0,1.0 | 0.708,1.0,0.378 | 0.305 | ENSG00000047 | [[[[16712335, 16 | 2 |
| ENSG00000025 | NCAPH2 | chr22 | 50518648 | 50518732 | 50518132 | 50518278 | 50520964 | 50521036 | 15,32,76 | 0,0,0 | 44,81,76 | 4,15,4 | 234 | 150 | 1.24E-07 | 4.22E-06 | 1.0,1.0,1.0 | 0.876,0.776,0.92 | 0.141 | ENSG00000025 | [[[[50508224, 50 | 2 |
| ENSG00000025 | NCAPH2 | chr22 | 50519189 | 50519320 | 50518132 | 50518278 | 50520964 | 50521036 | 4,7,31 | 0,0,0 | 9,14,14 | 4,15,4 | 281 | 150 | 1.76E-10 | 1.11E-08 | 1.0,1.0,1.0 | 0.546,0.333,0.65 | 0.49 | ENSG00000025 | [[[[50508224, 50 | 2 |
| ENSG00000025 | NCAPH2 | chr22 | 50521985 | 50522039 | 50521542 | 50521609 | 50522180 | 50522251 | 18,31,29 | 0,0,0 | 17,53,28 | 4,12,8 | 204 | 150 | 1.27E-09 | 6.90E-08 | 1.0,1.0,1.0 | 0.758,0.765,0.72 | 0.252 | ENSG00000025 | [[[[50508224, 50 | 2 |
| ENSG00000188 | MAPK12 | chr22 | 50257081 | 50257193 | 50256598 | 50256646 | 50258242 | 50258301 | 12,19,37 | 0,0,1 | 25,66,58 | 2,9,1 | 262 | 150 | 0.000281123 | 0.003902539 | 1.0,1.0,0.955 | 0.877,0.808,0.97 | 0.1 | ENSG00000188 | [[[[50261385, 50 | 2 |
| ENSG00000100 | L3MBTL2 | chr22 | 41213892 | 41214026 | 41209695 | 41209933 | 41219418 | 41219536 | 32,16,64 | 0,0,0 | 35,72,51 | 4,7,5 | 284 | 150 | 5.91E-07 | 1.72E-05 | 1.0,1.0,1.0 | 0.822,0.845,0.84 | 0.163 | ENSG00000100 | [[[[41205312, 41 | 3 |
| ENSG00000100 | L3MBTL2 | chr22 | 41216138 | 41216262 | 41209695 | 41209933 | 41219418 | 41219536 | 6,2,3 | 0,0,0 | 5,3,2 | 4,7,5 | 274 | 150 | 1.33E-12 | 1.19E-10 | 1.0,1.0,1.0 | 0.406,0.19,0.18 | 0.741 | ENSG00000100 | [[[[41205312, 41 | 3 |
| ENSG00000100 | L3MBTL2 | chr22 | 41216138 | 41216262 | 41213850 | 41214026 | 41219418 | 41219536 | 37,19,50 | 0,0,0 | 20,40,31 | 2,7,2 | 274 | 150 | 2.29E-06 | 5.77E-05 | 1.0,1.0,1.0 | 0.846,0.758,0.89 | 0.167 | ENSG00000100 | [[[[41205312, 41 | 2 |
| ENSG00000100 | SLC25A17 | chr22 | 40792524 | 40792676 | 40779008 | 40779125 | 40819194 | 40819350 | 37,38,41 | 3,0,6 | 31,96,61 | 13,21,14 | 300 | 150 | 0.000645982 | 0.007995864 | 0.86,1.0,0.774 | 0.544,0.696,0.68 | 0.236 | ENSG00000100 | [[[[40819195, 40 | 3 |
| ENSG00000100 | SLC25A17 | chr22 | 40799022 | 40799083 | 40779008 | 40779125 | 40819194 | 40819326 | 20,32,63 | 3,0,6 | 32,77,51 | 13,21,14 | 211 | 150 | 7.13E-05 | 0.001188203 | 0.826,1.0,0.882 | 0.636,0.723,0.72 | 0.209 | ENSG00000100 | [[[[40819195, 40 | 3 |
| ENSG00000186 | NF2 | chr22 | 29655593 | 29655676 | 29654656 | 29654725 | 29661204 | 29661339 | 59,58,74 | 0,0,0 | 25,94,73 | 2,4,6 | 233 | 150 | 5.26E-05 | 0.000912051 | 1.0,1.0,1.0 | 0.889,0.938,0.88 | 0.095 | ENSG00000186 | [[[[29603633, 29 | 2 |
| ENSG00000182 | EWSR1 | chr22 | 29273737 | 29273864 | 29272379 | 29272431 | 29288605 | 29288687 | 14,21,25 | 0,0,0 | 27,57,42 | 2,2,8 | 277 | 150 | 1.84E-05 | 0.000365385 | 1.0,1.0,1.0 | 0.88,0.939,0.74 | 0.147 | ENSG00000182 | [[[[29268268, 29 | 4 |
| ENSG00000182 | EWSR1 | chr22 | 29278029 | 29278216 | 29273737 | 29273864 | 29288605 | 29288679 | 394,370,658 | 0,0,0 | 360,717,498 | 8,9,17 | 300 | 150 | 1.45E-07 | 4.88E-06 | 1.0,1.0,1.0 | 0.957,0.976,0.93 | 0.044 | ENSG00000182 | [[[[29268268, 29 | 3 |
| ENSG00000182 | EWSR1 | chr22 | 29278029 | 29278216 | 29273737 | 29273864 | 29292136 | 29292169 | 394,370,658 | 0,0,0 | 358,713,498 | 10,38,16 | 300 | 150 | 8.88E-16 | 1.21E-13 | 1.0,1.0,1.0 | 0.947,0.904,0.94 | 0.07 | ENSG00000182 | [[[[29268268, 29 | 5 |
| ENSG00000182 | EWSR1 | chr22 | 29286922 | 29287134 | 29278029 | 29278216 | 29288605 | 29288679 | 8,376,601,138 | 0,0,0 | 4,661,181,741 | 6,14,11 | 300 | 150 | 0.000257876 | 0.003616051 | 1.0,1.0,1.0 | 0.975,0.977,0.97 | 0.026 | ENSG00000182 | [[[[29268268, 29 | 2 |
| ENSG00000182 | EWSR1 | chr22 | 29286922 | 29287134 | 29278029 | 29278216 | 29291561 | 29291599 | 9,8,22 | 0,0,0 | 6,13,7 | 7,22,18 | 300 | 150 | 0 | 0 | 1.0,1.0,1.0 | 0.3,0.228,0.163 | 0.77 | ENSG00000182 | [[[[29268268, 29 | 3 |
| ENSG00000182 | EWSR1 | chr22 | 29286922 | 29287134 | 29278029 | 29278216 | 29292136 | 29292169 | 12,11,28 | 0,0,0 | 6,19,11 | 4,9,11 | 300 | 150 | 1.03E-12 | 9.33E-11 | 1.0,1.0,1.0 | 0.429,0.514,0.33 | 0.575 | ENSG00000182 | [[[[29268268, 29 | 4 |
| ENSG00000182 | EWSR1 | chr22 | 29288605 | 29288786 | 29272379 | 29272431 | 29291561 | 29291599 | 47,41,95 | 0,0,0 | 31,49,57 | 1,5,3 | 300 | 150 | 0.000154108 | 0.002317566 | 1.0,1.0,1.0 | 0.939,0.831,0.90 | 0.108 | ENSG00000182 | [[[[29268268, 29 | 5 |
| ENSG00000182 | EWSR1 | chr22 | 29288605 | 29288786 | 29278029 | 29278216 | 29291561 | 29291599 | 47,41,95 | 0,0,0 | 35,61,60 | 7,22,18 | 300 | 150 | 0 | 0 | 1.0,1.0,1.0 | 0.714,0.581,0.62 | 0.36 | ENSG00000182 | [[[[29268268, 29 | 3 |
| ENSG00000182 | EWSR1 | chr22 | 29288605 | 29288786 | 29278029 | 29278216 | 29291564 | 29291599 | 65,60,68 | 0,0,0 | 62,96,71 | 8,15,5 | 300 | 150 | 2.67E-10 | 1.64E-08 | 1.0,1.0,1.0 | 0.795,0.762,0.87 | 0.189 | ENSG00000182 | [[[[29268268, 29 | 3 |
| ENSG00000182 | EWSR1 | chr22 | 29288605 | 29291599 | 29275628 | 29278216 | 29292136 | 29292169 | 163,140,234 | 0,0,0 | 142,260,166 | 4,9,11 | 300 | 150 | 6.88E-08 | 2.52E-06 | 1.0,1.0,1.0 | 0.947,0.935,0.88 | 0.078 | ENSG00000182 | [[[[29268268, 29 | 4 |
| ENSG00000182 | EWSR1 | chr22 | 29291561 | 29291599 | 29273740 | 29273864 | 29292136 | 29292169 | 151,125,217 | 0,0,0 | 119,200,133 | 10,38,16 | 188 | 150 | 0 | 0 | 1.0,1.0,1.0 | 0.905,0.808,0.86 | 0.139 | ENSG00000182 | [[[[29268268, 29 | 5 |
| ENSG00000182 | EWSR1 | chr22 | 29291561 | 29291599 | 29278029 | 29278216 | 29292136 | 29292169 | 151,125,217 | 0,0,0 | 126,220,151 | 4,9,11 | 188 | 150 | 5.90E-07 | 1.71E-05 | 1.0,1.0,1.0 | 0.962,0.951,0.91 | 0.057 | ENSG00000182 | [[[[29268268, 29 | 4 |
| ENSG00000182 | EWSR1 | chr22 | 29291564 | 29291599 | 29273740 | 29273864 | 29292136 | 29292169 | 149,124,216 | 0,0,0 | 118,196,133 | 10,38,16 | 185 | 150 | 0 | 0 | 1.0,1.0,1.0 | 0.905,0.807,0.87 | 0.139 | ENSG00000182 | [[[[29268268, 29 | 5 |
| ENSG00000182 | EWSR1 | chr22 | 29291564 | 29291599 | 29278029 | 29278216 | 29292136 | 29292169 | 149,124,216 | 0,0,0 | 126,211,137 | 4,9,11 | 185 | 150 | 3.89E-07 | 1.18E-05 | 1.0,1.0,1.0 | 0.962,0.95,0.91 | 0.059 | ENSG00000182 | [[[[29268268, 29 | 4 |
| ENSG00000160 | PRMT2 | chr21 | 46658744 | 46658920 | 46649574 | 46649739 | 46664294 | 46665117 | 56,90,144 | 2,3,3 | 44,82,38 | 9,7,13 | 300 | 150 | 1.61E-06 | 4.22E-05 | 0.933,0.938,0.96 | 0.71,0.854,0.594 | 0.224 | ENSG00000160 | [[[[46635674, 46 | 4 |
| ENSG00000160 | PRMT2 | chr21 | 46663382 | 46663554 | 46649574 | 46649739 | 46664294 | 46664951 | 63,45,76 | 2,3,3 | 31,64,42 | 9,7,13 | 300 | 150 | 3.08E-05 | 0.000572896 | 0.94,0.882,0.927 | 0.633,0.821,0.61 | 0.226 | ENSG00000160 | [[[[46635674, 46 | 4 |
| ENSG00000185 | BRWD1 | chr21 | 39218528 | 39218660 | 39215236 | 39215362 | 39224407 | 39224469 | 8,1,7 | 0,0,0 | 1,6,6 | 0,4,2 | 282 | 150 | 0.000106492 | 0.001685255 | 1.0,1.0,1.0 | 1.0,0.444,0.615 | 0.314 | ENSG00000185 | [[[[39313443, 39 | 2 |
| ENSG00000183 | PSMG1 | chr21 | 39180284 | 39180436 | 39179923 | 39179986 | 39183251 | 39183385 | 97,84,146 | 0,0,0 | 118,162,101 | 7,5,7 | 300 | 150 | 7.79E-07 | 2.20E-05 | 1.0,1.0,1.0 | 0.894,0.942,0.87 | 0.095 | ENSG00000183 | [[[[39183252, 39 | 2 |
| ENSG00000185 | SETD4 | chr21 | 36058815 | 36058924 | 36053582 | 36053620 | 36060346 | 36060385 | 4,5,9 | 0,0,0 | 8,16,10 | 1,2,3 | 259 | 150 | 0.000639232 | 0.007923892 | 1.0,1.0,1.0 | 0.822,0.822,0.65 | 0.232 | ENSG00000185 | [[[[36060347, 36 | 2 |
| ENSG00000185 | SETD4 | chr21 | 36058815 | 36058973 | 36053582 | 36053620 | 36060346 | 36060363 | 1,6,7 | 0,0,0 | 7,19,19 | 1,2,3 | 300 | 150 | 0.000525579 | 0.00670597 | 1.0,1.0,1.0 | 0.778,0.826,0.76 | 0.212 | ENSG00000185 | [[[[36060347, 36 | 2 |
| ENSG00000159 | SON | chr21 | 33572550 | 33572630 | 33568970 | 33569087 | 33576417 | 33576447 | 5,4,9 | 0,0,0 | 30,59,24 | 3,2,9 | 230 | 150 | 2.92E-05 | 0.000546863 | 1.0,1.0,1.0 | 0.867,0.951,0.63 | 0.182 | ENSG00000159 | [[[[33543038, 33 | 3 |
| ENSG00000159 | SON | chr21 | 33573307 | 33573455 | 33572212 | 33572630 | 33576417 | 33576447 | 20,15,18 | 0,0,0 | 46,87,48 | 11,22,9 | 298 | 150 | 2.39E-09 | 1.23E-07 | 1.0,1.0,1.0 | 0.678,0.666,0.72 | 0.309 | ENSG00000159 | [[[[33543038, 33 | 3 |
| ENSG00000159 | SON | chr21 | 33575595 | 33575667 | 33573024 | 33573455 | 33576417 | 33576447 | 77,73,114 | 0,0,0 | 124,224,134 | 6,6,5 | 222 | 150 | 0.000241105 | 0.003413544 | 1.0,1.0,1.0 | 0.933,0.962,0.94 | 0.052 | ENSG00000159 | [[[[33543038, 33 | 2 |
| ENSG00000156 | SCAF4 | chr21 | 31691816 | 31691930 | 31685567 | 31685733 | 31692348 | 31692449 | 37,17,33 | 0,0,0 | 27,100,54 | 6,8,2 | 264 | 150 | 6.91E-06 | 0.000153932 | 1.0,1.0,1.0 | 0.719,0.877,0.93 | 0.155 | ENSG00000156 | [[[[31731663, 31 | 3 |
| ENSG00000124 | CSE1L | chr20 | 49085282 | 49085386 | 49084025 | 49084162 | 49089246 | 49089397 | 115,136,167 | 0,0,0 | 53,83,74 | 1,9,2 | 254 | 150 | 2.41E-05 | 0.000463155 | 1.0,1.0,1.0 | 0.969,0.845,0.95 | 0.077 | ENSG00000124 | [[[[49046312, 49 | 2 |
| ENSG00000124 | CSE1L | chr20 | 49088008 | 49088106 | 49085282 | 49085386 | 49090741 | 49090839 | 91,75,105 | 0,0,0 | 58,124,48 | 6,3,4 | 248 | 150 | 2.12E-05 | 0.000413854 | 1.0,1.0,1.0 | 0.854,0.962,0.87 | 0.102 | ENSG00000124 | [[[[49046312, 49 | 3 |
| ENSG00000124 | CSE1L | chr20 | 49089246 | 49089397 | 49085282 | 49085386 | 49090741 | 49090839 | 1,1,2 | 0,0,0 | 3,3,6 | 6,3,4 | 300 | 150 | 6.95E-12 | 5.50E-10 | 1.0,1.0,1.0 | 0.2,0.333,0.429 | 0.679 | ENSG00000124 | [[[[49046312, 49 | 3 |

| | | | | | | | | | | | | | | | | | | | | | | |
|---|---|---|---|---|---|---|---|---|---|---|---|---|---|---|---|---|---|---|---|---|---|---|
| ENSG000001242 | CSE1L | chr20 | 49089246 | 49089397 | 49088008 | 49088106 | 49090741 | 49090839 | 156,129,199 | 0,0,0 | 79,130,72 | 17,16,12 | 300 | 150 | 0 | 0 | 1.0,1.0,1.0 | 0.699,0.802,0.75 | 0.25 | ENSG000001242 | [[[[49046312, 490 | 2 |
| ENSG000001509 | SEC24D | chr4 | 118817263 | 118817412 | 118815450 | 118815726 | 118833578 | 118833737 | 38,33,66 | 0,0,0 | 39,72,83 | 11,23,9 | 299 | 150 | 3.12E-12 | 2.64E-10 | 1.0,1.0,1.0 | 0.64,0.611,0.822 | 0.309 | ENSG000001509 | [[[[118835941, 11 | 2 |
| ENSG000001509 | SEC24D | chr4 | 118824619 | 118824749 | 118815450 | 118815726 | 118833578 | 118833737 | 38,34,80 | 0,0,0 | 36,64,53 | 11,23,9 | 280 | 150 | 8.66E-15 | 1.05E-12 | 1.0,1.0,1.0 | 0.637,0.599,0.75 | 0.335 | ENSG000001509 | [[[[118835941, 11 | 2 |
| ENSG000001070 | RIC1 | chr9 | 5747301 | 5747505 | 5743688 | 5743737 | 5753199 | 5753238 | 26,20,41 | 0,0,0 | 19,80,45 | 0,4,7 | 300 | 150 | 4.75E-05 | 0.000835217 | 1.0,1.0,1.0 | 1.0,0.909,0.763 | 0.109 | ENSG000001070 | [[[[5629107, 5629 | 2 |
| ENSG000000460 | OFD1 | chrX | 13746318 | 13746455 | 13744414 | 13744519 | 13749426 | 13749533 | 6,7,11 | 0,0,0 | 9,25,19 | 2,8,0 | 287 | 150 | 1.35E-05 | 0.000277368 | 1.0,1.0,1.0 | 0.702,0.62,1.0 | 0.226 | ENSG000000460 | [[[[13734748, 137 | 2 |
| ENSG000001775 | ATOX1 | chr5 | 151751703 | 151751779 | 151742821 | 151742959 | 151758545 | 151758649 | 5,14,23 | 0,1,2 | 5,13,8 | 2,6,6 | 226 | 150 | 3.18E-05 | 0.000589285 | 1.0,0.903,0.884 | 0.624,0.59,0.469 | 0.368 | ENSG000001775 | [[[[151758546, 15 | 2 |
| ENSG000001440 | CIAO1 | chr2 | 96267624 | 96267736 | 96266240 | 96266489 | 96267835 | 96267924 | 77,59,107 | 0,0,0 | 70,132,83 | 4,10,12 | 262 | 150 | 3.21E-09 | 1.61E-07 | 1.0,1.0,1.0 | 0.909,0.883,0.79 | 0.137 | ENSG000001440 | [[[[96266225, 962 | 2 |
| ENSG000001153 | EVA1A | chr2 | 75522364 | 75522487 | 75518055 | 75518208 | 75561256 | 75561298 | 52,43,100 | 0,0,2 | 76,139,65 | 3,5,6 | 273 | 150 | 0.000477925 | 0.006192761 | 1.0,1.0,0.965 | 0.933,0.939,0.85 | 0.079 | ENSG000001153 | [[[[75560680, 755 | 2 |
| ENSG000001153 | EVA1A | chr2 | 75526111 | 75526199 | 75518055 | 75518208 | 75561256 | 75561329 | 2,3,3 | 0,0,2 | 4,11,1 | 3,5,6 | 238 | 150 | 0.000744314 | 0.009021037 | 1.0,1.0,0.486 | 0.457,0.581,0.09 | 0.451 | ENSG000001153 | [[[[75560680, 755 | 2 |
| ENSG000001045 | UBR5 | chr8 | 102285512 | 102285781 | 102285126 | 102285338 | 102288154 | 102288266 | 30,22,33 | 0,0,0 | 12,46,21 | 1,1,5 | 300 | 150 | 0.000794271 | 0.00954002 | 1.0,1.0,1.0 | 0.857,0.958,0.67 | 0.169 | ENSG000001045 | [[[[102412173, 10 | 3 |
| ENSG000001042 | ZDHHC2 | chr8 | 17209931 | 17210058 | 17198380 | 17198413 | 17210387 | 17210480 | 69,51,101 | 0,0,0 | 68,106,70 | 8,9,4 | 277 | 150 | 2.55E-08 | 1.04E-06 | 1.0,1.0,1.0 | 0.822,0.864,0.90 | 0.136 | ENSG000001042 | [[[[17156482, 171 | 3 |
| ENSG000001005 | PSMA3 | chr14 | 58247749 | 58247832 | 58244830 | 58244941 | 58257744 | 58257846 | 48,58,95 | 0,0,0 | 56,144,68 | 43,77,31 | 233 | 150 | 0 | 0 | 1.0,1.0,1.0 | 0.456,0.546,0.58 | 0.471 | ENSG000001005 | [[[[58244843, 582 | 2 |
| ENSG000001005 | PSMA3 | chr14 | 58252118 | 58252242 | 58244903 | 58244941 | 58257744 | 58257846 | 214,191,259 | 0,0,0 | 262,566,309 | 43,77,31 | 274 | 150 | 0 | 0 | 1.0,1.0,1.0 | 0.769,0.801,0.84 | 0.195 | ENSG000001005 | [[[[58244843, 582 | 2 |
| ENSG000001005 | PSMA3 | chr14 | 58257744 | 58257846 | 58244830 | 58244941 | 58257924 | 58257998 | 238,227,344 | 0,0,0 | 399,834,507 | 28,41,37 | 252 | 150 | 0 | 0 | 1.0,1.0,1.0 | 0.895,0.924,0.89 | 0.097 | ENSG000001005 | [[[[58244843, 582 | 3 |
| ENSG000001545 | ELOC | chr8 | 73964486 | 73964625 | 73955910 | 73956054 | 73972244 | 73972287 | 2,2,0 | 1,5,3 | 3,9,7 | 0,3,0 | 289 | 150 | 1.36E-05 | 0.000280053 | 0.509,0.172,0.0 | 1.0,0.609,1.0 | -0.643 | ENSG000001545 | [[[[73972077, 739 | 2 |
| ENSG000000403 | STAU2 | chr8 | 73613743 | 73613956 | 73603725 | 73603863 | 73617291 | 73617451 | 10,15,26 | 0,0,0 | 15,29,12 | 2,3,2 | 300 | 150 | 0.000134053 | 0.002053184 | 1.0,1.0,1.0 | 0.789,0.829,0.75 | 0.211 | ENSG000000403 | [[[[73746783, 737 | 2 |
| ENSG000001131 | HMGCR | chr5 | 75344244 | 75344332 | 75343852 | 75343964 | 75350041 | 75350148 | 28,28,59 | 0,0,0 | 15,47,25 | 0,6,4 | 238 | 150 | 1.44E-05 | 0.000294005 | 1.0,1.0,1.0 | 1.0,0.832,0.798 | 0.123 | ENSG000001131 | [[[[75337220, 753 | 3 |
| ENSG000001131 | HMGCR | chr5 | 75350788 | 75350949 | 75350257 | 75350374 | 75355364 | 75355522 | 41,45,86 | 0,0,0 | 33,76,47 | 4,6,7 | 300 | 150 | 3.12E-08 | 1.24E-06 | 1.0,1.0,1.0 | 0.805,0.864,0.77 | 0.187 | ENSG000001131 | [[[[75337220, 753 | 5 |
| ENSG000001131 | HMGCR | chr5 | 75354502 | 75354697 | 75350257 | 75350374 | 75355364 | 75355522 | 4,5,16 | 0,0,0 | 6,19,20 | 4,6,7 | 300 | 150 | 8.32E-09 | 3.76E-07 | 1.0,1.0,1.0 | 0.429,0.613,0.58 | 0.457 | ENSG000001131 | [[[[75337220, 753 | 5 |
| ENSG000000356 | NSMAF | chr8 | 58590866 | 58590934 | 58589451 | 58589558 | 58594231 | 58594290 | 13,17,18 | 0,0,0 | 9,8,6 | 11,11,4 | 218 | 150 | 1.64E-14 | 1.93E-12 | 1.0,1.0,1.0 | 0.36,0.334,0.508 | 0.599 | ENSG000000356 | [[[[58659573, 586 | 2 |
| ENSG000000356 | NSMAF | chr8 | 58607768 | 58607840 | 58603209 | 58603386 | 58609603 | 58609733 | 18,13,31 | 0,0,0 | 8,17,13 | 3,17,5 | 222 | 150 | 7.68E-12 | 6.02E-10 | 1.0,1.0,1.0 | 0.643,0.403,0.63 | 0.439 | ENSG000000356 | [[[[58659573, 586 | 2 |
| ENSG000000472 | ATP6V1H | chr8 | 53801798 | 53801896 | 53769653 | 53769743 | 53811163 | 53811217 | 33,37,63 | 0,0,0 | 25,73,44 | 1,2,6 | 248 | 150 | 0.000544024 | 0.006913528 | 1.0,1.0,1.0 | 0.938,0.957,0.81 | 0.096 | ENSG000000472 | [[[[53843034, 538 | 3 |
| ENSG000001701 | NIPA1 | chr15 | 22810748 | 22810796 | 22786631 | 22786834 | 22820312 | 22820473 | 2,0,3 | 0,1,0 | 2,2,2 | 2,8,1 | 198 | 150 | 0.000322957 | 0.004403846 | 1.0,0.0,1.0 | 0.431,0.159,0.60 | 0.269 | ENSG000001701 | [[[[22786644, 227 | 2 |
| ENSG000000786 | VDAC3 | chr8 | 42398711 | 42398864 | 42395083 | 42395133 | 42401787 | 42402015 | 369,321,522 | 0,0,0 | 226,457,305 | 11,8,2 | 300 | 150 | 2.08E-05 | 0.000407195 | 1.0,1.0,1.0 | 0.911,0.966,0.98 | 0.045 | ENSG000000786 | [[[[42391880, 423 | 2 |
| ENSG000000786 | VDAC3 | chr8 | 42401787 | 42402015 | 42395083 | 42395133 | 42403310 | 42403441 | 283,295,381 | 0,0,0 | 230,457,260 | 11,20,13 | 300 | 150 | 4.73E-13 | 4.50E-11 | 1.0,1.0,1.0 | 0.913,0.92,0.909 | 0.086 | ENSG000000786 | [[[[42391880, 423 | 3 |
| ENSG000001746 | ANGEL2 | chr1 | 213008209 | 213008466 | 213005032 | 213005454 | 213013092 | 213013418 | 17,9,11 | 0,0,0 | 9,48,32 | 3,2,3 | 300 | 150 | 0.000137857 | 0.002102987 | 1.0,1.0,1.0 | 0.6,0.923,0.842 | 0.212 | ENSG000001746 | [[[[213015613, 21 | 2 |
| ENSG000000650 | CYB5R4 | chr6 | 83921081 | 83921175 | 83914415 | 83914448 | 83922437 | 83922470 | 8,9,11 | 0,0,0 | 14,28,20 | 5,5,5 | 244 | 150 | 2.81E-07 | 8.84E-06 | 1.0,1.0,1.0 | 0.633,0.775,0.71 | 0.294 | ENSG000000650 | [[[[83859689, 838 | 3 |
| ENSG000000650 | CYB5R4 | chr6 | 83934594 | 83934735 | 83924469 | 83924592 | 83940055 | 83940206 | 41,46,39 | 0,0,0 | 13,34,26 | 7,11,10 | 291 | 150 | 6.44E-15 | 7.90E-13 | 1.0,1.0,1.0 | 0.489,0.614,0.57 | 0.441 | ENSG000000650 | [[[[83859689, 838 | 2 |
| ENSG000001091 | RAB34 | chr17 | 28715799 | 28715808 | 28715640 | 28715705 | 28716902 | 28716994 | 10,8,9 | 0,0,2 | 9,33,15 | 16,21,27 | 159 | 150 | 9.28E-09 | 4.15E-07 | 1.0,1.0,0.809 | 0.347,0.597,0.34 | 0.507 | ENSG000001091 | [[[[28717213, 287 | 2 |
| ENSG000001091 | RAB34 | chr17 | 28715799 | 28715901 | 28715194 | 28715214 | 28716902 | 28716994 | 2,5,18 | 0,0,0 | 8,20,11 | 2,12,6 | 252 | 150 | 5.03E-09 | 2.39E-07 | 1.0,1.0,1.0 | 0.704,0.498,0.52 | 0.425 | ENSG000001091 | [[[[28717213, 287 | 4 |
| ENSG000001091 | RAB34 | chr17 | 28715799 | 28715901 | 28715640 | 28715705 | 28716902 | 28716994 | 221,186,318 | 0,0,2 | 239,410,245 | 16,21,27 | 252 | 150 | 4.77E-15 | 5.95E-13 | 1.0,1.0,0.99 | 0.899,0.921,0.84 | 0.109 | ENSG000001091 | [[[[28717213, 287 | 2 |
| ENSG000001396 | LMBR1L | chr12 | 49105923 | 49105957 | 49103686 | 49103813 | 49106960 | 49107045 | 1,6,3 | 0,0,0 | 10,17,9 | 5,3,3 | 184 | 150 | 1.33E-06 | 3.55E-05 | 1.0,1.0,1.0 | 0.62,0.822,0.71 | 0.283 | ENSG000001396 | [[[[49110484, 491 | 3 |
| ENSG000001870 | ENTPD5 | chr14 | 73971851 | 73971908 | 73970009 | 73970125 | 73973876 | 73973978 | 5,5,2 | 0,0,0 | 3,6,1 | 1,3,2 | 207 | 150 | 2.87E-05 | 0.000537534 | 1.0,1.0,1.0 | 0.685,0.592,0.26 | 0.486 | ENSG000001870 | [[[[74019250, 740 | 2 |
| ENSG000001141 | CEP70 | chr3 | 138537177 | 138537347 | 138532513 | 138532570 | 138591853 | 138591956 | 16,5,30 | 0,0,0 | 13,25,5 | 7,3,4 | 300 | 150 | 7.81E-09 | 3.56E-07 | 1.0,1.0,1.0 | 0.481,0.806,0.38 | 0.443 | ENSG000001141 | [[[[138594198, 13 | 5 |
| ENSG000001551 | AGPAT5 | chr8 | 6741660 | 6741751 | 6732560 | 6732650 | 6755050 | 6755174 | 46,45,70 | 0,2,1 | 62,112,77 | 23,22,9 | 241 | 150 | 1.52E-09 | 8.15E-08 | 1.0,0.933,0.978 | 0.627,0.76,0.842 | 0.227 | ENSG000001551 | [[[[6708642, 6708 | 2 |
| ENSG000001551 | AGPAT5 | chr8 | 6747669 | 6747828 | 6732560 | 6732650 | 6755050 | 6755174 | 79,69,109 | 0,2,1 | 105,214,149 | 23,22,9 | 300 | 150 | 5.35E-09 | 2.53E-07 | 1.0,0.945,0.982 | 0.695,0.829,0.89 | 0.17 | ENSG000001551 | [[[[6708642, 6708 | 2 |
| ENSG000001140 | KPNA1 | chr3 | 122451975 | 122452064 | 122442037 | 122442116 | 122453869 | 122454001 | 34,31,58 | 0,0,0 | 15,51,38 | 1,9,0 | 239 | 150 | 4.54E-05 | 0.000802982 | 1.0,1.0,1.0 | 0.904,0.781,1.0 | 0.105 | ENSG000001140 | [[[[122514757, 12 | 3 |
| ENSG000001140 | KPNA1 | chr3 | 122453869 | 122454001 | 122433660 | 122433788 | 122461223 | 122461318 | 39,27,57 | 0,0,0 | 27,71,46 | 3,1,5 | 282 | 150 | 0.000291365 | 0.00402637 | 1.0,1.0,1.0 | 0.827,0.974,0.83 | 0.123 | ENSG000001140 | [[[[122514757, 12 | 6 |
| ENSG000000746 | DPP8 | chr15 | 65485098 | 65485160 | 65474208 | 65474288 | 65487689 | 65487818 | 9,10,18 | 0,0,0 | 13,22,8 | 1,9,4 | 212 | 150 | 1.85E-07 | 6.05E-06 | 1.0,1.0,1.0 | 0.902,0.634,0.58 | 0.293 | ENSG000000746 | [[[[65517486, 655 | 4 |
| ENSG000002136 | PPP1CB | chr2 | 28781737 | 28781842 | 28776850 | 28776982 | 28783906 | 28783978 | 242,206,361 | 0,0,0 | 151,333,229 | 3,17,4 | 255 | 150 | 5.58E-07 | 1.63E-05 | 1.0,1.0,1.0 | 0.967,0.92,0.971 | 0.047 | ENSG000002136 | [[[[28751830, 287 | 2 |
| ENSG000002136 | PPP1CB | chr2 | 28788657 | 28788809 | 28783906 | 28783978 | 28799198 | 28801648 | 302,306,474 | 11,17,21 | 105,279,214 | 112,157,104 | 300 | 150 | 0 | 0 | 0.932,0.9,0.919 | 0.319,0.47,0.507 | 0.485 | ENSG000002136 | [[[[28751830, 287 | 2 |
| ENSG000002136 | PPP1CB | chr2 | 28793862 | 28793997 | 28783906 | 28783978 | 28799198 | 28801648 | 327,296,494 | 11,17,21 | 141,272,195 | 112,157,104 | 285 | 150 | 0 | 0 | 0.94,0.902,0.925 | 0.399,0.477,0.49 | 0.465 | ENSG000002136 | [[[[28751830, 287 | 2 |
| ENSG000001496 | DSN1 | chr20 | 36770872 | 36771024 | 36762460 | 36762548 | 36771424 | 36771473 | 1,3,1 | 0,0,0 | 1,2,6 | 1,2,2 | 300 | 150 | 9.11E-05 | 0.001469804 | 1.0,1.0,1.0 | 0.333,0.333,0.6 | 0.578 | ENSG000001496 | [[[[36773662, 367 | 3 |
| ENSG000001496 | DSN1 | chr20 | 36771424 | 36771473 | 36767968 | 36768042 | 36773661 | 36773720 | 2,2,11 | 0,0,0 | 4,4,2 | 0,2,4 | 199 | 150 | 0.000268977 | 0.003755126 | 1.0,1.0,1.0 | 1.0,0.601,0.274 | 0.375 | ENSG000001496 | [[[[36773662, 367 | 2 |
| ENSG000001511 | UEVLD | chr11 | 18566346 | 18566482 | 18564891 | 18565010 | 18575346 | 18575412 | 17,13,23 | 0,0,0 | 13,42,32 | 1,3,3 | 286 | 150 | 0.000800593 | 0.009609892 | 1.0,1.0,1.0 | 0.872,0.88,0.848 | 0.133 | ENSG000001511 | [[[[18588613, 185 | 2 |
| ENSG000001301 | ECSIT | chr19 | 11510616 | 11510660 | 11507990 | 11508048 | 11519074 | 11519193 | 0,0,1 | 3,5,8 | 2,5,0 | 3,7,2 | 194 | 150 | 0.000440564 | 0.005764093 | 0.0,0.0,0.088 | 0.34,0.356,0.0 | -0.203 | ENSG000001301 | [[[[11529062, 115 | 2 |
| ENSG000001468 | TMEM209 | chr7 | 130201849 | 130202091 | 130192621 | 130192823 | 130203787 | 130203846 | 35,25,35 | 0,0,0 | 63,152,132 | 6,5,17 | 300 | 150 | 1.22E-06 | 3.28E-05 | 1.0,1.0,1.0 | 0.84,0.938,0.795 | 0.142 | ENSG000001468 | [[[[130205373, 13 | 2 |
| ENSG000001126 | PTK7 | chr6 | 43130229 | 43130420 | 43129726 | 43129829 | 43138848 | 43138982 | 77,90,209 | 0,0,0 | 70,100,108 | 0,5,6 | 300 | 150 | 0.00025963 | 0.003638644 | 1.0,1.0,1.0 | 1.0,0.909,0.9 | 0.064 | ENSG000001126 | [[[[43076314, 430 | 4 |
| ENSG000001126 | PTK7 | chr6 | 43130510 | 43130661 | 43129726 | 43129829 | 43132015 | 43132164 | 41,74,217 | 0,0,0 | 38,83,91 | 0,2,10 | 300 | 150 | 1.77E-05 | 0.000352636 | 1.0,1.0,1.0 | 1.0,0.954,0.82 | 0.075 | ENSG000001126 | [[[[43076314, 430 | 2 |
| ENSG000001126 | PTK7 | chr6 | 43130590 | 43130661 | 43129726 | 43129829 | 43132015 | 43132164 | 13,32,89 | 0,0,0 | 12,36,37 | 0,2,10 | 221 | 150 | 5.68E-06 | 0.000129864 | 1.0,1.0,1.0 | 1.0,0.924,0.715 | 0.12 | ENSG000001126 | [[[[43076314, 430 | 2 |
| ENSG000001265 | BECN1 | chr17 | 42818220 | 42818415 | 42814598 | 42814673 | 42819547 | 42819609 | 9,3,7 | 0,0,0 | 19,32,15 | 2,6,4 | 300 | 150 | 1.04E-06 | 2.86E-05 | 1.0,1.0,1.0 | 0.826,0.727,0.65 | 0.265 | ENSG000001265 | [[[[42824155, 428 | 4 |
| ENSG000001265 | BECN1 | chr17 | 42818543 | 42818680 | 42818220 | 42818415 | 42819547 | 42819609 | 105,74,142 | 0,1,2 | 93,226,141 | 7,18,6 | 287 | 150 | 6.80E-07 | 1.95E-05 | 1.0,0.975,0.974 | 0.874,0.868,0.92 | 0.094 | ENSG000001265 | [[[[42824155, 428 | 2 |
| ENSG000001265 | BECN1 | chr17 | 42818786 | 42818877 | 42814523 | 42814673 | 42819547 | 42819609 | 56,48,85 | 0,0,0 | 50,95,51 | 2,6,4 | 241 | 150 | 5.64E-05 | 0.000968729 | 1.0,1.0,1.0 | 0.94,0.908,0.888 | 0.088 | ENSG000001265 | [[[[42824155, 428 | 4 |
| ENSG000001265 | BECN1 | chr17 | 42818786 | 42818877 | 42818220 | 42818415 | 42819547 | 42819609 | 56,48,85 | 0,1,2 | 51,98,53 | 7,18,6 | 241 | 150 | 1.81E-08 | 7.63E-07 | 1.0,0.968,0.964 | 0.819,0.772,0.84 | 0.165 | ENSG000001265 | [[[[42824155, 428 | 2 |
| ENSG000001184 | ANKRD13C | chr1 | 70315480 | 70315566 | 70306223 | 70306290 | 70324852 | 70324957 | 16,6,12 | 0,0,0 | 5,35,17 | 8,18,8 | 236 | 150 | 1.92E-10 | 1.21E-08 | 1.0,1.0,1.0 | 0.284,0.553,0.57 | 0.529 | ENSG000001184 | [[[[70353979, 703 | 2 |
| ENSG000001403 | ETFA | chr15 | 76274411 | 76274494 | 76230503 | 76231398 | 76287845 | 76287945 | 195,140,198 | 0,0,0 | 186,361,197 | 12,13,8 | 233 | 150 | 2.22E-08 | 9.14E-07 | 1.0,1.0,1.0 | 0.909,0.947,0.94 | 0.068 | ENSG000001403 | [[[[76311350, 763 | 4 |
| ENSG000001449 | TRPC1 | chr3 | 142791018 | 142791158 | 142784703 | 142785040 | 142802168 | 142802344 | 5,8,7 | 2,0,0 | 3,4,7 | 1,6,3 | 290 | 150 | 0.000142115 | 0.002158003 | 0.564,1.0,1.0 | 0.608,0.256,0.54 | 0.384 | ENSG000001449 | [[[[142724034, 14 | 2 |
| ENSG000001058 | HBP1 | chr7 | 107189291 | 107189448 | 107186568 | 107186681 | 107200159 | 107200301 | 24,8,40 | 0,0,0 | 32,63,49 | 1,3,5 | 300 | 150 | 0.000369544 | 0.004946162 | 1.0,1.0,1.0 | 0.941,0.913,0.83 | 0.105 | ENSG000001058 | [[[[107169003, 10 | 3 |
| ENSG000000082 | SYPL1 | chr7 | 106097689 | 106097897 | 106092948 | 106093137 | 106112139 | 106112142 | 130,100,174 | 0,2,0 | 69,137,104 | 9,24,19 | 300 | 150 | 0 | 0 | 1.0,0.962,1.0 | 0.793,0.741,0.73 | 0.232 | ENSG000000082 | [[[[106112140, 10 | 2 |
| ENSG000001058 | PMPCB | chr7 | 103307595 | 103307708 | 103304410 | 103304490 | 103311807 | 103311896 | 57,74,72 | 0,0,0 | 45,133,62 | 6,11,5 | 263 | 150 | 2.08E-08 | 8.65E-07 | 1.0,1.0,1.0 | 0.811,0.873,0.87 | 0.147 | ENSG000001058 | [[[[103297435, 10 | 4 |
| ENSG000001058 | PMPCB | chr7 | 103308951 | 103309095 | 103304410 | 103304490 | 103311807 | 103311896 | 13,6,8 | 0,0,0 | 16,33,16 | 6,11,5 | 294 | 150 | 1.93E-08 | 8.11E-07 | 1.0,1.0,1.0 | 0.576,0.605,0.62 | 0.4 | ENSG000001058 | [[[[103297435, 10 | 4 |
| ENSG000001058 | PMPCB | chr7 | 103308951 | 103309095 | 103307595 | 103307708 | 103311807 | 103311896 | 108,105,159 | 0,0,0 | 110,216,116 | 3,9,3 | 294 | 150 | 6.99E-05 | 0.001167773 | 1.0,1.0,1.0 | 0.949,0.924,0.95 | 0.058 | ENSG000001058 | [[[[103297435, 10 | 3 |
| ENSG000001058 | PMPCB | chr7 | 103311642 | 103311728 | 103304410 | 103304490 | 103311807 | 103311896 | 80,74,91 | 0,0,0 | 81,175,108 | 6,11,5 | 236 | 150 | 6.86E-07 | 1.96E-05 | 1.0,1.0,1.0 | 0.896,0.91,0.932 | 0.087 | ENSG000001058 | [[[[103297435, 10 | 4 |
| ENSG000001058 | PMPCB | chr7 | 103311642 | 103311728 | 103307595 | 103307708 | 103311807 | 103311896 | 80,74,91 | 0,0,0 | 78,181,110 | 3,9,3 | 236 | 150 | 0.000152578 | 0.002296746 | 1.0,1.0,1.0 | 0.943,0.927,0.95 | 0.057 | ENSG000001058 | [[[[103297435, 10 | 3 |
| ENSG000001648 | ORC5 | chr7 | 104161071 | 104161182 | 104136780 | 104136893 | 104168472 | 104168525 | 21,29,33 | 0,0,0 | 30,80,36 | 5,7,0 | 261 | 150 | 4.18E-05 | 0.000748214 | 1.0,1.0,1.0 | 0.775,0.868,1.0 | 0.119 | ENSG000001648 | [[[[104207833, 10 | 3 |
| ENSG000001648 | ORC5 | chr7 | 104195142 | 104195254 | 104188250 | 104188381 | 104207832 | 104207957 | 16,19,23 | 0,0,0 | 30,49,33 | 4,4,1 | 262 | 150 | 0.000185057 | 0.002726939 | 1.0,1.0,1.0 | 0.811,0.875,0.95 | 0.121 | ENSG000001648 | [[[[104207833, 10 | 4 |
| ENSG000001648 | ORC5 | chr7 | 104197724 | 104197799 | 104195142 | 104195254 | 104207832 | 104207957 | 10,15,18 | 0,0,0 | 28,54,25 | 8,0,2 | 225 | 150 | 0.000158147 | 0.002372603 | 1.0,1.0,1.0 | 0.7,1.0,0.893 | 0.136 | ENSG000001648 | [[[[104207833, 10 | 3 |
| ENSG000001648 | ORC5 | chr7 | 104204141 | 104204234 | 104188250 | 104188381 | 104207832 | 104207957 | 14,11,18 | 0,0,0 | 21,30,23 | 4,4,1 | 243 | 150 | 2.71E-05 | 0.000511596 | 1.0,1.0,1.0 | 0.764,0.822,0.93 | 0.16 | ENSG000001648 | [[[[104207833, 10 | 4 |
| ENSG000001810 | NQO1 | chr16 | 69713027 | 69713129 | 69709400 | 69711281 | 69718369 | 69718534 | 515,341,635 | 0,0,0 | 8,561,391,814 | 17,14,12 | 252 | 150 | 0.000155762 | 0.002340048 | 1.0,1.0,1.0 | 0.968,0.983,0.97 | 0.024 | ENSG000001810 | [[[[69726433, 697 | 3 |
| ENSG000001810 | NQO1 | chr16 | 69714963 | 69715077 | 69709400 | 69711281 | 69718369 | 69718534 | 18,20,27 | 0,0,0 | 52,82,48 | 17,14,12 | 264 | 150 | 6.03E-10 | 3.47E-08 | 1.0,1.0,1.0 | 0.635,0.769,0.69 | 0.301 | ENSG000001810 | [[[[69726433, 697 | 3 |
| ENSG000001810 | NQO1 | chr16 | 69714963 | 69715077 | 69713027 | 69713129 | 69718369 | 69718534 | 417,329,489 | 0,0,0 | 6,451,180,730 | 15,24,22 | 264 | 150 | 4.85E-09 | 2.32E-07 | 1.0,1.0,1.0 | 0.961,0.965,0.95 | 0.041 | ENSG000001810 | [[[[69726433, 697 | 2 |
| ENSG000001810 | NQO1 | chr16 | 69718122 | 69718253 | 69710781 | 69711281 | 69718369 | 69718534 | 583,490,718 | 0,0,0 | 9,581,643,951 | 17,14,12 | 281 | 150 | 0.000234888 | 0.00333855 | 1.0,1.0,1.0 | 0.968,0.984,0.97 | 0.024 | ENSG000001810 | [[[[69726433, 697 | 3 |
| ENSG000001810 | NQO1 | chr16 | 69718122 | 69718253 | 69713027 | 69713129 | 69718369 | 69718534 | 626,532,780 | 0,0,0 | 100,617,581,025 | 15,24,22 | 281 | 150 | 1.60E-07 | 5.32E-06 | 1.0,1.0,1.0 | 0.973,0.975,0.96 | 0.03 | ENSG000001810 | [[[[69726433, 697 | 2 |
| ENSG000000840 | ZMPSTE24 | chr1 | 40271893 | 40272035 | 40260838 | 40260985 | 40281342 | 40281527 | 74,56,109 | 0,0,0 | 75,153,101 | 1,6,7 | 292 | 150 | 4.79E-05 | 0.000841553 | 1.0,1.0,1.0 | 0.975,0.929,0.88 | 0.072 | ENSG000000840 | [[[[40258236, 402 | 4 |
| ENSG000001058 | PON2 | chr7 | 95406986 | 95407068 | 95406118 | 95406247 | 95416241 | 95416297 | 87,79,110 | 0,0,0 | 67,164,109 | 8,10,6 | 232 | 150 | 5.07E-08 | 1.91E-06 | 1.0,1.0,1.0 | 0.844,0.914,0.92 | 0.107 | ENSG000001058 | [[[[95434878, 954 | 4 |
| ENSG000001058 | PON2 | chr7 | 95412311 | 95412477 | 95406118 | 95406247 | 95416241 | 95416297 | 117,76,131 | 0,0,0 | 75,175,112 | 8,10,6 | 300 | 150 | 1.05E-08 | 4.67E-07 | 1.0,1.0,1.0 | 0.824,0.897,0.90 | 0.125 | ENSG000001058 | [[[[95434878, 954 | 4 |
| ENSG000001058 | PON2 | chr7 | 95412311 | 95412477 | 95406986 | 95407068 | 95416241 | 95416297 | 117,76,131 | 0,0,0 | 76,175,111 | 3,9,4 | 300 | 150 | 8.42E-06 | 0.000183549 | 1.0,1.0,1.0 | 0.927,0.907,0.93 | 0.078 | ENSG000001058 | [[[[95434878, 954 | 3 |
| ENSG000001519 | DST | chr6 | 56463564 | 56463764 | 56460130 | 56460254 | 56466077 | 56466195 | 118,148,173 | 0,0,0 | 141,285,135 | 4,16,6 | 300 | 150 | 5.34E-08 | 2.00E-06 | 1.0,1.0,1.0 | 0.946,0.899,0.91 | 0.079 | ENSG000001519 | [[[[56642622, 566 | 3 |
| ENSG000001568 | NSMCE2 | chr8 | 125102316 | 125102487 | 125091869 | 125091958 | 125151170 | 125151277 | 51,35,61 | 0,0,0 | 49,115,53 | 7,8,8 | 300 | 150 | 1.89E-08 | 7.93E-07 | 1.0,1.0,1.0 | 0.778,0.878,0.76 | 0.192 | ENSG000001568 | [[[[125091860, 12 | 2 |
| ENSG000001842 | TSPYL2 | chrX | 53084754 | 53084866 | 53084531 | 53084622 | 53085226 | 53085321 | 3,1,8 | 0,0,0 | 37,33,20 | 5,0,3 | 262 | 150 | 0.000384367 | 0.005114018 | 1.0,1.0,1.0 | 0.809,1.0,0.792 | 0.133 | ENSG000001842 | [[[[53082367, 530 | 2 |

| | | | | | | | | | | | | | | | | | | | | | | |
|---|---|---|---|---|---|---|---|---|---|---|---|---|---|---|---|---|---|---|---|---|---|---|
| ENSG00000134 | EMC7 | chr15 | 34088052 | 34088133 | 34084016 | 34084486 | 34095894 | 34096014 | 203,155,223 | 3,0,0 | 295,583,265 | 9,12,15 | 231 | 150 | 1.52E-05 | 0.000307332 | 0.978,1.0,1.0 | 0.955,0.969,0.92 | 0.045 | ENSG00000134 | [[[[34101604, 34 | 2 |
| ENSG00000134 | EMC7 | chr15 | 34090316 | 34090455 | 34084016 | 34084486 | 34095894 | 34096014 | 195,163,226 | 3,0,0 | 290,525,323 | 9,12,15 | 289 | 150 | 6.81E-06 | 0.000151902 | 0.971,1.0,1.0 | 0.944,0.958,0.91 | 0.05 | ENSG00000134 | [[[[34101604, 34 | 2 |
| ENSG00000136 | WASHC4 | chr12 | 105164110 | 105164307 | 105160000 | 105160148 | 105164640 | 105164740 | 23,23,41 | 0,0,0 | 18,45,28 | 2,7,3 | 300 | 150 | 8.70E-07 | 2.43E-05 | 1.0,1.0,1.0 | 0.818,0.763,0.82 | 0.198 | ENSG00000136 | [[[[105107731, 10 | 2 |
| ENSG00000065 | IPO5 | chr13 | 98016728 | 98016851 | 98015725 | 98015781 | 98019580 | 98019809 | 63,61,68 | 0,0,0 | 20,41,12 | 5,2,1 | 273 | 150 | 7.72E-05 | 0.001269717 | 1.0,1.0,1.0 | 0.687,0.918,0.86 | 0.176 | ENSG00000065 | [[[[97953675, 97 | 2 |
| ENSG00000077 | GPR137B | chr1 | 236168705 | 236168755 | 236142531 | 236143036 | 236179878 | 236180028 | 10,10,6 | 0,0,0 | 7,31,8 | 3,1,4 | 200 | 150 | 3.59E-05 | 0.000656413 | 1.0,1.0,1.0 | 0.636,0.959,0.6 | 0.268 | ENSG00000077 | [[[[236142539, 23 | 2 |
| ENSG00000113 | RARS | chr5 | 168500590 | 168500720 | 168497011 | 168497348 | 168506721 | 168506831 | 110,86,160 | 0,0,0 | 91,147,101 | 5,13,3 | 280 | 150 | 4.85E-08 | 1.84E-06 | 1.0,1.0,1.0 | 0.907,0.858,0.94 | 0.096 | ENSG00000113 | [[[[168486471, 16 | 3 |
| ENSG00000146 | PSPH | chr7 | 56015022 | 56015171 | 56011066 | 56011869 | 56019599 | 56019734 | 194,135,193 | 0,0,0 | 181,333,194 | 26,68,30 | 299 | 150 | 0 | 0 | 1.0,1.0,1.0 | 0.777,0.711,0.76 | 0.249 | ENSG00000146 | [[[[56051138, 560 | 2 |
| ENSG00000146 | PSPH | chr7 | 56033960 | 56034154 | 56021072 | 56021231 | 56051137 | 56051288 | 9,23,29 | 0,0,0 | 5,17,11 | 0,6,4 | 300 | 150 | 7.95E-07 | 2.24E-05 | 1.0,1.0,1.0 | 1.0,0.586,0.579 | 0.278 | ENSG00000146 | [[[[56051138, 560 | 2 |
| ENSG00000146 | PSPH | chr7 | 56043147 | 56043261 | 56021205 | 56021231 | 56051137 | 56051604 | 1,6,5 | 0,0,0 | 4,16,8 | 0,6,4 | 264 | 150 | 1.83E-06 | 4.72E-05 | 1.0,1.0,1.0 | 1.0,0.602,0.532 | 0.289 | ENSG00000146 | [[[[56051138, 560 | 2 |
| ENSG00000183 | UPP1 | chr7 | 48101823 | 48101870 | 48094762 | 48094827 | 48106872 | 48107082 | 14,15,22 | 72,98,121 | 71,187,92 | 193,319,242 | 197 | 150 | 1.33E-07 | 4.51E-06 | 0.129,0.104,0.12 | 0.219,0.309,0.22 | -0.132 | ENSG00000183 | [[[[48089265, 48 | 3 |
| ENSG00000183 | UPP1 | chr7 | 48103296 | 48103411 | 48094762 | 48094827 | 48106872 | 48107082 | 98,77,104 | 72,98,121 | 93,190,114 | 193,319,242 | 265 | 150 | 8.81E-05 | 0.001425799 | 0.435,0.308,0.32 | 0.214,0.252,0.21 | 0.131 | ENSG00000183 | [[[[48089265, 48 | 3 |
| ENSG00000136 | CCM2 | chr7 | 45064462 | 45064646 | 45038252 | 45038426 | 45069825 | 45069961 | 2,3,4 | 2,2,1 | 1,8,4 | 0,0,1 | 300 | 150 | 0.000118458 | 0.00184847 | 0.333,0.429,0.66 | 1.0,1.0,0.667 | -0.413 | ENSG00000136 | [[[[45000201, 45 | 3 |
| ENSG00000136 | CCM2 | chr7 | 45068442 | 45068579 | 45064462 | 45064646 | 45072725 | 45072783 | 21,45,84 | 2,5,1 | 32,75,50 | 5,20,7 | 287 | 150 | 0.000441771 | 0.005777905 | 0.846,0.825,0.97 | 0.77,0.662,0.789 | 0.143 | ENSG00000136 | [[[[45000201, 45 | 2 |
| ENSG00000136 | CCM2 | chr7 | 45069825 | 45069961 | 45064462 | 45064646 | 45072725 | 45072783 | 53,63,64 | 2,5,1 | 46,81,57 | 5,20,7 | 286 | 150 | 0.000215867 | 0.003115278 | 0.933,0.869,0.97 | 0.828,0.68,0.81 | 0.152 | ENSG00000136 | [[[[45000201, 45 | 2 |
| ENSG00000136 | CCM2 | chr7 | 45069825 | 45069961 | 45064462 | 45064646 | 45073459 | 45073571 | 2,4,3 | 0,0,0 | 4,12,8 | 0,6,1 | 286 | 150 | 0.000142482 | 0.002163147 | 1.0,1.0,1.0 | 1.0,0.512,0.808 | 0.227 | ENSG00000136 | [[[[45000201, 45 | 3 |
| ENSG00000106 | POLD2 | chr7 | 44115765 | 44115893 | 44115294 | 44115354 | 44116429 | 44116510 | 2,6,5 | 0,0,0 | 24,39,13 | 2,2,4 | 278 | 150 | 9.70E-05 | 0.001555548 | 1.0,1.0,1.0 | 0.866,0.913,0.63 | 0.195 | ENSG00000106 | [[[[44123511, 441 | 2 |
| ENSG00000106 | POLD2 | chr7 | 44116429 | 44116510 | 44116114 | 44116272 | 44117132 | 44117247 | 70,145,285 | 0,0,0 | 153,220,164 | 4,19,3 | 231 | 150 | 9.73E-08 | 3.43E-06 | 1.0,1.0,1.0 | 0.961,0.883,0.97 | 0.061 | ENSG00000106 | [[[[44123511, 441 | 2 |
| ENSG00000106 | POLD2 | chr7 | 44117583 | 44117742 | 44117148 | 44117247 | 44121833 | 44122109 | 3,7,2 | 0,0,0 | 6,15,5 | 14,34,18 | 300 | 150 | 2.01E-11 | 1.46E-09 | 1.0,1.0,1.0 | 0.176,0.181,0.12 | 0.84 | ENSG00000106 | [[[[44123511, 441 | 2 |
| ENSG00000106 | POLD2 | chr7 | 44117618 | 44117742 | 44117062 | 44117247 | 44121833 | 44122109 | 294,267,371 | 0,0,0 | 205,383,240 | 14,34,18 | 274 | 150 | 0 | 0 | 1.0,1.0,1.0 | 0.889,0.86,0.88 | 0.124 | ENSG00000106 | [[[[44123511, 441 | 2 |
| ENSG00000137 | DCUN1D5 | chr11 | 103083255 | 103083326 | 103066458 | 103066567 | 103091786 | 103092154 | 2,5,3 | 0,0,0 | 0,2,4 | 8,8,8 | 221 | 150 | 5.98E-13 | 5.60E-11 | 1.0,1.0,1.0 | 0.0,0.145,0.253 | 0.867 | ENSG00000137 | [[[[103091787, 10 | 3 |
| ENSG00000173 | DPY19L1 | chr7 | 34954697 | 34954778 | 34949796 | 34949898 | 34966893 | 34966971 | 33,22,43 | 0,0,0 | 11,23,23 | 4,2,8 | 231 | 150 | 2.09E-08 | 8.69E-07 | 1.0,1.0,1.0 | 0.641,0.882,0.65 | 0.275 | ENSG00000173 | [[[[35037097, 350 | 3 |
| ENSG00000173 | DPY19L1 | chr7 | 34957983 | 34958070 | 34949796 | 34949898 | 34966893 | 34966971 | 12,13,23 | 0,0,0 | 9,17,18 | 4,2,8 | 237 | 150 | 2.97E-08 | 1.19E-06 | 1.0,1.0,1.0 | 0.587,0.843,0.58 | 0.328 | ENSG00000173 | [[[[35037097, 350 | 3 |
| ENSG00000131 | PPFIA1 | chr11 | 70343668 | 70343892 | 70339170 | 70339306 | 70354300 | 70354452 | 19,26,41 | 0,0,0 | 15,47,23 | 2,9,7 | 300 | 150 | 4.62E-09 | 2.22E-07 | 1.0,1.0,1.0 | 0.789,0.723,0.62 | 0.289 | ENSG00000131 | [[[[70270690, 702 | 2 |
| ENSG00000131 | PPFIA1 | chr11 | 70349922 | 70350039 | 70339170 | 70339306 | 70354300 | 70354452 | 2,2,4 | 0,0,0 | 9,15,23 | 2,9,7 | 267 | 150 | 2.10E-07 | 6.80E-06 | 1.0,1.0,1.0 | 0.717,0.484,0.64 | 0.383 | ENSG00000131 | [[[[70270690, 702 | 2 |
| ENSG00000109 | KLHL2 | chr4 | 165317825 | 165317969 | 165299506 | 165299656 | 165322031 | 165322458 | 19,17,24 | 0,0,0 | 24,30,13 | 1,7,0 | 294 | 150 | 0.000159717 | 0.002393811 | 1.0,1.0,1.0 | 0.924,0.686,1.0 | 0.13 | ENSG00000109 | [[[[165207561, 16 | 6 |
| ENSG00000070 | OSBPL3 | chr7 | 24863199 | 24863292 | 24849068 | 24849176 | 24863495 | 24863599 | 25,7,25 | 0,0,0 | 6,32,19 | 0,7,1 | 243 | 150 | 0.000140332 | 0.002135607 | 1.0,1.0,1.0 | 1.0,0.738,0.921 | 0.114 | ENSG00000070 | [[[[24979886, 24 | 3 |
| ENSG00000136 | IGF2BP3 | chr7 | 23343717 | 23343853 | 23342063 | 23342189 | 23347648 | 23347803 | 40,47,81 | 0,0,0 | 26,83,58 | 14,48,27 | 286 | 150 | 0 | 0 | 1.0,1.0,1.0 | 0.493,0.476,0.53 | 0.5 | ENSG00000136 | [[[[23469936, 234 | 2 |
| ENSG00000136 | IGF2BP3 | chr7 | 23345939 | 23346062 | 23342063 | 23342189 | 23347648 | 23347803 | 4,5,17 | 0,0,0 | 5,28,9 | 14,48,27 | 273 | 150 | 1.11E-16 | 1.65E-14 | 1.0,1.0,1.0 | 0.164,0.243,0.15 | 0.813 | ENSG00000136 | [[[[23469936, 234 | 2 |
| ENSG00000168 | LMBRD1 | chr6 | 69701888 | 69701953 | 69676449 | 69676541 | 69713644 | 69713797 | 9,9,13 | 0,0,0 | 6,7,6 | 4,5,2 | 215 | 150 | 3.33E-09 | 1.66E-07 | 1.0,1.0,1.0 | 0.511,0.494,0.67 | 0.439 | ENSG00000168 | [[[[69796813, 697 | 5 |
| ENSG00000165 | KDM1B | chr6 | 18208131 | 18208206 | 18205536 | 18205664 | 18212487 | 18212604 | 14,9,17 | 0,0,0 | 6,20,11 | 2,4,3 | 225 | 150 | 3.12E-06 | 7.62E-05 | 1.0,1.0,1.0 | 0.667,0.769,0.71 | 0.285 | ENSG00000165 | [[[[18155422, 18 | 2 |
| ENSG00000162 | ZFPL1 | chr11 | 65086414 | 65086608 | 65085114 | 65085226 | 65087315 | 65087433 | 20,37,70 | 0,0,0 | 32,52,34 | 6,13,9 | 300 | 150 | 2.43E-12 | 2.09E-10 | 1.0,1.0,1.0 | 0.727,0.667,0.65 | 0.317 | ENSG00000162 | [[[[65084222, 650 | 3 |
| ENSG00000159 | NPR2 | chr9 | 35800021 | 35800157 | 35799617 | 35799731 | 35800708 | 35800841 | 8,4,17 | 0,0,0 | 15,32,25 | 3,3,4 | 286 | 150 | 1.01E-05 | 0.000215515 | 1.0,1.0,1.0 | 0.724,0.848,0.76 | 0.221 | ENSG00000159 | [[[[35791591, 357 | 2 |
| ENSG00000159 | NPR2 | chr9 | 35800388 | 35800483 | 35799617 | 35799731 | 35800708 | 35800841 | 9,12,6 | 0,0,0 | 8,18,16 | 3,3,4 | 245 | 150 | 1.06E-06 | 2.91E-05 | 1.0,1.0,1.0 | 0.62,0.786,0.71 | 0.295 | ENSG00000159 | [[[[35791591, 357 | 2 |
| ENSG00000164 | SUN1 | chr7 | 849922 | 850033 | 849519 | 849603 | 852608 | 852667 | 36,41,94 | 0,0,0 | 18,35,41 | 1,5,3 | 261 | 150 | 4.35E-05 | 0.000773471 | 1.0,1.0,1.0 | 0.912,0.801,0.88 | 0.133 | ENSG00000164 | [[[[832476, 8326 | 2 |
| ENSG00000164 | SUN1 | chr7 | 851383 | 851482 | 849519 | 849603 | 852608 | 852667 | 13,14,22 | 0,0,0 | 13,51,44 | 1,5,3 | 249 | 150 | 0.000353643 | 0.004754217 | 1.0,1.0,1.0 | 0.887,0.86,0.898 | 0.118 | ENSG00000164 | [[[[832476, 8326 | 2 |
| ENSG00000164 | SUN1 | chr7 | 851383 | 851482 | 849519 | 849603 | 852812 | 852907 | 13,14,22 | 0,0,0 | 12,49,43 | 3,4,3 | 249 | 150 | 5.53E-05 | 0.000953283 | 1.0,1.0,1.0 | 0.707,0.881,0.89 | 0.172 | ENSG00000164 | [[[[832476, 8326 | 3 |
| ENSG00000164 | SUN1 | chr7 | 852608 | 852667 | 849519 | 849603 | 852812 | 852907 | 17,17,20 | 0,0,0 | 8,29,21 | 3,4,3 | 209 | 150 | 6.03E-06 | 0.000137055 | 1.0,1.0,1.0 | 0.657,0.839,0.83 | 0.223 | ENSG00000164 | [[[[832476, 8326 | 3 |
| ENSG00000104 | KCNN4 | chr19 | 43774191 | 43774619 | 43771999 | 43772103 | 43780702 | 43780973 | 1,3,1 | 0,0,1 | 7,7,5 | 1,2,4 | 300 | 150 | 0.000560647 | 0.007097554 | 1.0,1.0,0.333 | 0.778,0.636,0.38 | 0.178 | ENSG00000104 | [[[[43780703, 43 | 2 |
| ENSG00000270 | TAF15 | chr17 | 35820163 | 35820239 | 35820023 | 35820075 | 35834565 | 35834598 | 165,158,316 | 0,0,0 | 66,153,121 | 7,8,13 | 226 | 150 | 2.46E-12 | 2.11E-10 | 1.0,1.0,1.0 | 0.862,0.927,0.86 | 0.117 | ENSG00000270 | [[[[35809484, 35 | 5 |
| ENSG00000270 | TAF15 | chr17 | 35820163 | 35820248 | 35820023 | 35820075 | 35834565 | 35834598 | 177,170,346 | 0,0,0 | 71,160,129 | 7,8,13 | 235 | 150 | 2.15E-12 | 1.87E-10 | 1.0,1.0,1.0 | 0.866,0.927,0.86 | 0.114 | ENSG00000270 | [[[[35809484, 35 | 5 |
| ENSG00000270 | TAF15 | chr17 | 35820331 | 35820437 | 35820023 | 35820075 | 35833906 | 35833941 | 1,1,3 | 0,0,0 | 5,24,8 | 2,14,7 | 256 | 150 | 2.12E-07 | 6.85E-06 | 1.0,1.0,1.0 | 0.594,0.501,0.40 | 0.501 | ENSG00000270 | [[[[35809484, 35 | 4 |
| ENSG00000270 | TAF15 | chr17 | 35820331 | 35820437 | 35820023 | 35820075 | 35834565 | 35834598 | 1,1,3 | 0,0,0 | 11,34,14 | 7,8,13 | 256 | 150 | 6.78E-07 | 1.95E-05 | 1.0,1.0,1.0 | 0.479,0.713,0.38 | 0.474 | ENSG00000270 | [[[[35809484, 35 | 5 |
| ENSG00000270 | TAF15 | chr17 | 35822639 | 35822833 | 35820023 | 35820437 | 35834565 | 35834598 | 262,199,310 | 0,0,0 | 70,199,124 | 7,13,6 | 300 | 150 | 5.42E-12 | 4.41E-10 | 1.0,1.0,1.0 | 0.833,0.884,0.91 | 0.124 | ENSG00000270 | [[[[35809484, 35 | 3 |
| ENSG00000270 | TAF15 | chr17 | 35833906 | 35833941 | 35820023 | 35820075 | 35834565 | 35834598 | 62,69,102 | 0,0,0 | 31,43,30 | 7,8,13 | 185 | 150 | 1.35E-14 | 1.60E-12 | 1.0,1.0,1.0 | 0.782,0.813,0.65 | 0.251 | ENSG00000270 | [[[[35809484, 35 | 5 |
| ENSG00000270 | TAF15 | chr17 | 35833906 | 35833941 | 35820331 | 35820437 | 35834565 | 35834598 | 62,69,102 | 0,0,0 | 29,30,23 | 7,13,6 | 185 | 150 | 1.44E-15 | 1.91E-13 | 1.0,1.0,1.0 | 0.771,0.652,0.75 | 0.273 | ENSG00000270 | [[[[35809484, 35 | 3 |
| ENSG00000075 | TIPIN | chr15 | 66349314 | 66349437 | 66341149 | 66341356 | 66351524 | 66351600 | 38,45,25 | 0,0,0 | 34,99,52 | 2,15,0 | 273 | 150 | 6.84E-06 | 0.000152567 | 1.0,1.0,1.0 | 0.903,0.784,1.0 | 0.104 | ENSG00000075 | [[[[66356639, 66 | 2 |
| ENSG00000112 | MTRF1L | chr6 | 152994512 | 152994676 | 152991184 | 152991321 | 152995243 | 152995319 | 8,4,6 | 0,0,0 | 3,11,9 | 4,2,2 | 300 | 150 | 1.96E-07 | 6.36E-06 | 1.0,1.0,1.0 | 0.273,0.733,0.69 | 0.434 | ENSG00000112 | [[[[153002427, 15 | 2 |
| ENSG00000127 | SIN3B | chr19 | 16853068 | 16853158 | 16851411 | 16851534 | 16862351 | 16862559 | 1,7,25 | 0,0,0 | 12,17,8 | 2,2,4 | 240 | 150 | 6.19E-06 | 0.000140455 | 1.0,1.0,1.0 | 0.789,0.842,0.55 | 0.271 | ENSG00000127 | [[[[16829398, 16 | 2 |
| ENSG00000165 | SUGT1 | chr13 | 52666811 | 52666919 | 52665636 | 52665733 | 52687733 | 52687872 | 76,60,71 | 0,0,0 | 84,152,80 | 2,10,2 | 258 | 150 | 0.000109085 | 0.00172092 | 1.0,1.0,1.0 | 0.961,0.898,0.95 | 0.061 | ENSG00000165 | [[[[52652836, 52 | 3 |
| ENSG00000001 | FUCA2 | chr6 | 143497388 | 143497497 | 143494810 | 143495847 | 143503912 | 143504252 | 169,156,192 | 0,0,0 | 157,352,184 | 5,13,10 | 259 | 150 | 1.35E-07 | 4.56E-06 | 1.0,1.0,1.0 | 0.948,0.94,0.914 | 0.066 | ENSG00000001 | [[[[143511411, 14 | 3 |
| ENSG00000087 | SRRT | chr7 | 100888041 | 100888143 | 100887702 | 100887859 | 100888473 | 100888660 | 54,84,117 | 0,0,0 | 71,161,104 | 6,13,0 | 252 | 150 | 2.95E-06 | 7.22E-05 | 1.0,1.0,1.0 | 0.876,0.881,1.0 | 0.081 | ENSG00000087 | [[[[100875103, 10 | 2 |
| ENSG00000112 | EYA4 | chr6 | 133515320 | 133515435 | 133512877 | 133513038 | 133525153 | 133525254 | 33,15,35 | 0,0,0 | 26,49,32 | 3,3,2 | 265 | 150 | 0.000486652 | 0.006286141 | 1.0,1.0,1.0 | 0.831,0.902,0.90 | 0.122 | ENSG00000112 | [[[[133241357, 13 | 2 |
| ENSG00000168 | SETD5 | chr3 | 9434589 | 9434630 | 9434333 | 9434485 | 9441592 | 9441741 | 6,1,11 | 0,0,0 | 0,8,6 | 2,2,5 | 191 | 150 | 3.99E-07 | 1.21E-05 | 1.0,1.0,1.0 | 0.0,0.759,0.485 | 0.585 | ENSG00000168 | [[[[9397615, 939 | 3 |
| ENSG00000111 | TPD52L1 | chr6 | 125253716 | 125253755 | 125248281 | 125248383 | 125262833 | 125263497 | 20,25,35 | 7,11,20 | 27,36,23 | 34,60,39 | 189 | 150 | 2.56E-06 | 6.35E-05 | 0.694,0.643,0.58 | 0.387,0.323,0.31 | 0.296 | ENSG00000111 | [[[[125153773, 12 | 2 |
| ENSG00000172 | SMPDL3A | chr6 | 122801309 | 122801406 | 122796823 | 122796968 | 122804908 | 122805089 | 7,5,8 | 0,0,2 | 2,9,4 | 0,9,2 | 247 | 150 | 0.00075521 | 0.009134206 | 1.0,1.0,0.708 | 1.0,0.378,0.548 | 0.261 | ENSG00000172 | [[[[122789258, 12 | 2 |
| ENSG00000170 | GSTA4 | chr6 | 52985450 | 52985583 | 52984463 | 52984605 | 52994156 | 52994261 | 58,29,63 | 0,0,0 | 33,47,21 | 32,55,42 | 283 | 150 | 0 | 0 | 1.0,1.0,1.0 | 0.353,0.312,0.20 | 0.709 | ENSG00000170 | [[[[52995249, 52 | 2 |
| ENSG00000170 | GSTA4 | chr6 | 52987356 | 52987408 | 52984463 | 52984605 | 52994156 | 52994261 | 24,23,19 | 0,0,0 | 31,36,31 | 32,55,42 | 202 | 150 | 0 | 0 | 1.0,1.0,1.0 | 0.418,0.327,0.35 | 0.634 | ENSG00000170 | [[[[52995249, 52 | 2 |
| ENSG00000153 | ZUP1 | chr6 | 116647458 | 116647610 | 116645713 | 116645934 | 116652003 | 116652192 | 7,7,14 | 0,0,0 | 23,37,10 | 6,10,2 | 300 | 150 | 9.93E-08 | 3.48E-06 | 1.0,1.0,1.0 | 0.657,0.649,0.71 | 0.327 | ENSG00000153 | [[[[116668566, 11 | 2 |
| ENSG00000153 | ZUP1 | chr6 | 116651571 | 116651737 | 116645713 | 116645934 | 116652003 | 116652192 | 15,21,18 | 0,0,0 | 13,32,15 | 6,10,2 | 300 | 150 | 6.11E-09 | 2.85E-07 | 1.0,1.0,1.0 | 0.52,0.615,0.789 | 0.359 | ENSG00000153 | [[[[116668566, 11 | 2 |
| ENSG00000157 | SLC37A3 | chr7 | 140343411 | 140343563 | 140337283 | 140337349 | 140345868 | 140345970 | 27,32,44 | 0,2,12 | 37,81,45 | 0,0,1 | 300 | 150 | 9.74E-06 | 0.000208972 | 1.0,0.889,0.647 | 1.0,1.0,0.957 | -0.14 | ENSG00000157 | [[[[140398416, 14 | 2 |
| ENSG00000157 | SLC37A3 | chr7 | 140345215 | 140345263 | 140337283 | 140337349 | 140345868 | 140345907 | 16,0,9 | 0,2,12 | 12,27,27 | 0,0,1 | 198 | 150 | 0.000190113 | 0.002791954 | 1.0,0.0,0.362 | 1.0,1.0,0.953 | -0.53 | ENSG00000157 | [[[[140398416, 14 | 2 |
| ENSG00000157 | SLC37A3 | chr7 | 140345868 | 140345970 | 140343411 | 140343563 | 140348625 | 140348767 | 15,7,14 | 0,0,0 | 18,31,21 | 1,8,2 | 252 | 150 | 6.20E-06 | 0.000140568 | 1.0,1.0,1.0 | 0.915,0.698,0.86 | 0.175 | ENSG00000157 | [[[[140398416, 14 | 2 |
| ENSG00000157 | SLC37A3 | chr7 | 140358639 | 140358785 | 140355667 | 140355764 | 140369589 | 140369682 | 26,20,49 | 0,3,6 | 53,92,69 | 0,1,0 | 296 | 150 | 5.85E-05 | 0.001001192 | 1.0,0.772,0.805 | 1.0,0.979,1.0 | -0.134 | ENSG00000157 | [[[[140398416, 14 | 2 |
| ENSG00000157 | SLC37A3 | chr7 | 140364407 | 140364491 | 140355667 | 140355764 | 140369589 | 140369654 | 18,20,32 | 0,3,6 | 26,85,59 | 0,1,0 | 234 | 150 | 3.91E-05 | 0.000706779 | 1.0,0.81,0.774 | 1.0,0.982,1.0 | -0.133 | ENSG00000157 | [[[[140398416, 14 | 2 |
| ENSG00000058 | SEC61A1 | chr3 | 128067420 | 128067612 | 128064876 | 128065037 | 128067982 | 128068059 | 180,233,360 | 0,0,0 | 199,416,279 | 6,15,5 | 300 | 150 | 5.55E-07 | 1.63E-05 | 1.0,1.0,1.0 | 0.943,0.933,0.96 | 0.053 | ENSG00000058 | [[[[128052437, 12 | 2 |
| ENSG00000025 | SEC63 | chr6 | 107921796 | 107921909 | 107913365 | 107913427 | 107929414 | 107929514 | 32,38,39 | 0,0,0 | 22,104,57 | 5,9,7 | 263 | 150 | 1.36E-07 | 4.60E-06 | 1.0,1.0,1.0 | 0.715,0.868,0.82 | 0.198 | ENSG00000025 | [[[[107957886, 10 | 2 |
| ENSG00000182 | CEP63 | chr3 | 134559149 | 134559429 | 134537154 | 134537211 | 134561376 | 134561895 | 4,2,2 | 0,0,0 | 3,11,4 | 4,1,1 | 300 | 150 | 7.68E-05 | 0.001265564 | 1.0,1.0,1.0 | 0.273,0.846,0.66 | 0.405 | ENSG00000182 | [[[[134486057, 13 | 8 |
| ENSG00000182 | CEP63 | chr3 | 134559149 | 134559429 | 134537154 | 134537268 | 134561376 | 134561895 | 4,2,2 | 0,0,0 | 2,11,4 | 3,1,4 | 300 | 150 | 1.07E-07 | 3.71E-06 | 1.0,1.0,1.0 | 0.25,0.846,0.333 | 0.524 | ENSG00000182 | [[[[134486057, 13 | 8 |
| ENSG00000117 | VAMP4 | chr1 | 171709664 | 171709744 | 171701091 | 171702084 | 171710713 | 171710814 | 11,7,17 | 0,0,0 | 8,10,29 | 2,3,2 | 230 | 150 | 0.000276667 | 0.003849106 | 1.0,1.0,1.0 | 0.723,0.685,0.90 | 0.229 | ENSG00000117 | [[[[171741910, 17 | 2 |
| ENSG00000111 | CNOT2 | chr12 | 70329422 | 70329570 | 70319297 | 70319364 | 70332766 | 70332846 | 32,28,64 | 0,0,1 | 15,59,33 | 8,9,0 | 298 | 150 | 5.24E-06 | 0.00012093 | 1.0,1.0,0.97 | 0.486,0.767,1.0 | 0.239 | ENSG00000111 | [[[[70243433, 70 | 2 |
| ENSG00000136 | ZDHHC4 | chr7 | 6580554 | 6580678 | 6578500 | 6578720 | 6585015 | 6585260 | 156,140,206 | 0,0,0 | 145,265,165 | 5,12,4 | 274 | 150 | 2.55E-06 | 6.33E-05 | 1.0,1.0,1.0 | 0.941,0.924,0.95 | 0.059 | ENSG00000136 | [[[[6577454, 657 | 4 |
| ENSG00000136 | ZDHHC4 | chr7 | 6582072 | 6582251 | 6581606 | 6581680 | 6585015 | 6585260 | 208,182,270 | 0,0,0 | 208,281,209 | 15,16,4 | 300 | 150 | 9.54E-11 | 6.28E-09 | 1.0,1.0,1.0 | 0.874,0.898,0.96 | 0.088 | ENSG00000136 | [[[[6577454, 657 | 2 |
| ENSG00000156 | CD109 | chr6 | 73766947 | 73767010 | 73766758 | 73766860 | 73780423 | 73780498 | 26,13,34 | 0,0,0 | 17,56,30 | 2,8,0 | 213 | 150 | 8.67E-05 | 0.001406114 | 1.0,1.0,1.0 | 0.857,0.831,1.0 | 0.104 | ENSG00000156 | [[[[73696203, 73 | 3 |
| ENSG00000156 | CD109 | chr6 | 73809983 | 73810174 | 73806843 | 73807072 | 73810991 | 73811096 | 72,38,116 | 0,0,0 | 27,48,59 | 6,19,13 | 300 | 150 | 0 | 0 | 1.0,1.0,1.0 | 0.692,0.558,0.69 | 0.352 | ENSG00000156 | [[[[73696203, 73 | 2 |
| ENSG00000146 | PRIM2 | chr6 | 57507386 | 57507454 | 57382030 | 57382168 | 57601092 | 57601219 | 12,10,22 | 0,0,0 | 26,36,22 | 2,3,3 | 218 | 150 | 0.000602711 | 0.00753987 | 1.0,1.0,1.0 | 0.899,0.892,0.83 | 0.125 | ENSG00000146 | [[[[57317630, 57 | 3 |
| ENSG00000146 | PRIM2 | chr6 | 57537439 | 57537625 | 57382030 | 57382168 | 57601092 | 57601219 | 33,35,46 | 0,0,0 | 36,58,28 | 2,3,3 | 300 | 150 | 0.000462525 | 0.006018353 | 1.0,1.0,1.0 | 0.9,0.906,0.824 | 0.123 | ENSG00000146 | [[[[57317630, 57 | 3 |
| ENSG00000146 | PRIM2 | chr6 | 57537439 | 57537625 | 57507386 | 57507454 | 57601092 | 57601219 | 33,35,44 | 0,0,0 | 36,55,27 | 4,9,0 | 300 | 150 | 2.89E-06 | 7.11E-05 | 1.0,1.0,1.0 | 0.818,0.753,1.0 | 0.143 | ENSG00000146 | [[[[57317630, 57 | 2 |
| ENSG00000003 | ICA1 | chr7 | 8138839 | 8138881 | 8127872 | 8128142 | 8141764 | 8141817 | 2,2,5 | 0,0,0 | 4,13,4 | 2,2,2 | 192 | 150 | 0.000411717 | 0.005433485 | 1.0,1.0,1.0 | 0.61,0.835,0.61 | 0.315 | ENSG00000003 | [[[[8262094, 826 | 2 |
| ENSG00000112 | FBXO9 | chr6 | 53078798 | 53078898 | 53076485 | 53076543 | 53082503 | 53082618 | 36,36,56 | 4,1,4 | 43,116,81 | 0,1,1 | 250 | 150 | 0.000675789 | 0.008306782 | 0.844,0.956,0.89 | 1.0,0.986,0.98 | -0.091 | ENSG00000112 | [[[[53065404, 53 | 2 |
| ENSG00000075 | ACTR6 | chr12 | 100207662 | 100207786 | 100205675 | 100205744 | 100218414 | 100218545 | 24,13,26 | 0,0,0 | 27,46,33 | 1,5,2 | 274 | 150 | 0.000647942 | 0.008012362 | 1.0,1.0,1.0 | 0.937,0.834,0.9 | 0.11 | ENSG00000075 | [[[[100200815, 10 | 5 |

| | | | | | | | | | | | | | | | | | | | | | | |
|---|---|---|---|---|---|---|---|---|---|---|---|---|---|---|---|---|---|---|---|---|---|---|
| ENSG00000075( | ACTR6 | chr12 | 100210072 | 100210208 | 100207662 | 100207786 | 100212255 | 100212354 | 18,25,34 | 0,0,0 | 25,53,25 | 3,3,4 | 286 | 150 | 1.47E-05 | 0.00029978 | 1.0,1.0,1.0 | 0.814,0.903,0.76 | 0.172 | ENSG00000075( | [[[[100200815, 1( | 2 |
| ENSG00000075( | ACTR6 | chr12 | 100210294 | 100210351 | 100207662 | 100207786 | 100212255 | 100212354 | 32,22,34 | 0,0,0 | 21,35,23 | 3,3,4 | 207 | 150 | 2.46E-05 | 0.000470767 | 1.0,1.0,1.0 | 0.835,0.894,0.80 | 0.155 | ENSG00000075( | [[[[100200815, 1( | 2 |
| ENSG00000184( | DIABLO | chr12 | 122218265 | 122218397 | 122216758 | 122216869 | 122225964 | 122226568 | 100,96,103 | 0,0,0 | 101,217,142 | 4,7,5 | 282 | 150 | 8.42E-05 | 0.001370171 | 1.0,1.0,1.0 | 0.931,0.943,0.93 | 0.063 | ENSG00000184( | [[[[122225965, 12 | 2 |
| ENSG00000184( | DIABLO | chr12 | 122224511 | 122224644 | 122216758 | 122216869 | 122225964 | 122226009 | 56,53,63 | 0,0,0 | 50,102,43 | 4,7,5 | 283 | 150 | 5.55E-07 | 1.63E-05 | 1.0,1.0,1.0 | 0.869,0.885,0.82 | 0.142 | ENSG00000184( | [[[[122225965, 12 | 2 |
| ENSG00000184( | DIABLO | chr12 | 122224644 | 122224748 | 122216758 | 122216869 | 122225964 | 122226039 | 4,8,5 | 0,0,0 | 17,31,37 | 4,7,5 | 254 | 150 | 2.73E-06 | 6.74E-05 | 1.0,1.0,1.0 | 0.715,0.723,0.81 | 0.249 | ENSG00000184( | [[[[122225965, 12 | 2 |
| ENSG00000124 | KLHDC3 | chr6 | 43017133 | 43017346 | 43014102 | 43014348 | 43018144 | 43018216 | 3,13,59 | 0,0,0 | 10,23,35 | 1,6,1 | 300 | 150 | 5.49E-05 | 0.000947775 | 1.0,1.0,1.0 | 0.833,0.657,0.94 | 0.188 | ENSG00000124 | [[[[43014249, 43( | 3 |
| ENSG00000124 | KLHDC3 | chr6 | 43017852 | 43017968 | 43017518 | 43017695 | 43018342 | 43018556 | 82,88,160 | 0,0,0 | 74,174,94 | 1,10,1 | 266 | 150 | 0.000619033 | 0.00771423 | 1.0,1.0,1.0 | 0.977,0.908,0.98 | 0.045 | ENSG00000124 | [[[[43014249, 43( | 2 |
| ENSG00000050( | BCAR1 | chr16 | 75237182 | 75237344 | 75235667 | 75235986 | 75242469 | 75242702 | 5,25,60 | 1,1,2 | 9,20,25 | 26,37,22 | 300 | 150 | 4.11E-15 | 5.17E-13 | 0.714,0.926,0.93 | 0.148,0.213,0.36 | 0.618 | ENSG00000050( | [[[[75251471, 752 | 2 |
| ENSG00000137 | CMTR1 | chr6 | 37462828 | 37463008 | 37461969 | 37462102 | 37471846 | 37471904 | 32,28,36 | 0,0,0 | 18,30,25 | 0,5,3 | 300 | 150 | 7.81E-05 | 0.001282804 | 1.0,1.0,1.0 | 1.0,0.75,0.806 | 0.148 | ENSG00000137 | [[[[37433221, 374 | 2 |
| ENSG00000112' | RNF8 | chr6 | 37371511 | 37371574 | 37368377 | 37369218 | 37376925 | 37377033 | 29,18,20 | 0,0,0 | 28,63,24 | 1,5,3 | 213 | 150 | 0.000640624 | 0.007934698 | 1.0,1.0,1.0 | 0.952,0.899,0.84 | 0.1 | ENSG00000112' | [[[[37353983, 373 | 2 |
| ENSG00000112' | RNF8 | chr6 | 37374619 | 37374709 | 37368377 | 37369218 | 37376925 | 37377033 | 19,6,7 | 0,0,0 | 16,26,10 | 1,5,3 | 240 | 150 | 1.07E-05 | 0.000226344 | 1.0,1.0,1.0 | 0.909,0.765,0.67 | 0.217 | ENSG00000112' | [[[[37353983, 373 | 2 |
| ENSG00000137 | MTCH1 | chr6 | 36977198 | 36977250 | 36975657 | 36975717 | 36978077 | 36978155 | 132,121,184 | 0,0,0 | 177,284,169 | 5,18,0 | 202 | 150 | 1.72E-05 | 0.000343171 | 1.0,1.0,1.0 | 0.963,0.921,1.0 | 0.039 | ENSG00000137 | [[[[36985853, 369 | 2 |
| ENSG00000136 | DDX56 | chr7 | 44571491 | 44571736 | 44570757 | 44570877 | 44572573 | 44572744 | 177,139,199 | 0,0,0 | 254,413,236 | 44,96,59 | 300 | 150 | 0 | 0 | 1.0,1.0,1.0 | 0.743,0.683,0.66 | 0.302 | ENSG00000136 | [[[[44573836, 445 | 2 |
| ENSG00000138 | EGF | chr4 | 109940945 | 109941145 | 109912883 | 109913462 | 109943841 | 109944069 | 8,5,14 | 0,0,1 | 13,20,12 | 2,2,3 | 300 | 150 | 0.000541089 | 0.006884115 | 1.0,1.0,0.875 | 0.765,0.833,0.66 | 0.203 | ENSG00000138 | [[[[109912883, 1( | 2 |
| ENSG00000138 | EGF | chr4 | 109943253 | 109943435 | 109912883 | 109913462 | 109943841 | 109943972 | 16,5,20 | 0,0,1 | 9,20,21 | 2,2,3 | 300 | 150 | 0.000515102 | 0.006592165 | 1.0,1.0,0.909 | 0.692,0.833,0.77 | 0.202 | ENSG00000138 | [[[[109912883, 1( | 2 |
| ENSG00000138 | EGF | chr4 | 109959311 | 109959437 | 109912890 | 109913462 | 109960866 | 109960989 | 1,3,16 | 0,0,0 | 6,22,28 | 2,6,0 | 276 | 150 | 9.18E-05 | 0.001479469 | 1.0,1.0,1.0 | 0.62,0.666,1.0 | 0.238 | ENSG00000138 | [[[[109912883, 1( | 5 |
| ENSG00000197' | ABCB8 | chr7 | 151033604 | 151033917 | 151028422 | 151028610 | 151034504 | 151034551 | 22,40,109 | 0,1,0 | 33,66,59 | 8,24,16 | 300 | 150 | 3.33E-16 | 4.71E-14 | 1.0,0.952,1.0 | 0.673,0.579,0.64 | 0.351 | ENSG00000197' | [[[[151028450, 15 | 2 |
| ENSG00000197' | ABCB8 | chr7 | 151034272 | 151034428 | 151028421 | 151028610 | 151034504 | 151034547 | 27,66,85 | 0,1,0 | 51,108,73 | 8,24,16 | 300 | 150 | 1.24E-13 | 1.28E-11 | 1.0,0.971,1.0 | 0.761,0.692,0.69 | 0.274 | ENSG00000197' | [[[[151028450, 15 | 2 |
| ENSG00000204 | VARS | chr6 | 31781018 | 31781123 | 31780869 | 31780938 | 31781690 | 31781761 | 135,119,189 | 0,0,0 | 71,146,73 | 1,7,4 | 255 | 150 | 0.00013524 | 0.002068455 | 1.0,1.0,1.0 | 0.977,0.925,0.91 | 0.061 | ENSG00000204 | [[[[31795546, 317 | 2 |
| ENSG00000204 | VARS | chr6 | 31781480 | 31781606 | 31780869 | 31780938 | 31781690 | 31781761 | 80,118,173 | 0,0,0 | 60,126,84 | 1,7,4 | 276 | 150 | 6.50E-05 | 0.001098289 | 1.0,1.0,1.0 | 0.97,0.907,0.919 | 0.068 | ENSG00000204 | [[[[31795546, 317 | 2 |
| ENSG00000100( | SLC25A1 | chr22 | 19177726 | 19178041 | 19177119 | 19177204 | 19178132 | 19178240 | 24,39,93 | 0,2,3 | 35,70,44 | 9,8,9 | 300 | 150 | 1.63E-05 | 0.000327951 | 1.0,0.907,0.939 | 0.66,0.814,0.71 | 0.221 | ENSG00000100( | [[[[19178580, 19' | 2 |
| ENSG00000198 | DDX39B | chr6 | 31540321 | 31540534 | 31532779 | 31532911 | 31541949 | 31541989 | 134,135,240 | 0,0,0 | 57,160,95 | 5,14,3 | 300 | 150 | 6.29E-10 | 3.61E-08 | 1.0,1.0,1.0 | 0.851,0.851,0.94 | 0.119 | ENSG00000198 | [[[[31541950, 315 | 5 |
| ENSG00000198 | DDX39B | chr6 | 31540321 | 31540664 | 31532779 | 31532911 | 31541949 | 31541981 | 95,105,165 | 0,0,0 | 171,321,195 | 5,14,3 | 300 | 150 | 1.20E-05 | 0.000250324 | 1.0,1.0,1.0 | 0.945,0.92,0.97 | 0.055 | ENSG00000198 | [[[[31541950, 315 | 5 |
| ENSG00000221 | PPP2R2A | chr8 | 26354467 | 26354633 | 26338888 | 26338987 | 26361100 | 26361151 | 48,49,51 | 0,0,0 | 46,110,76 | 2,5,3 | 300 | 150 | 0.000616099 | 0.007685901 | 1.0,1.0,1.0 | 0.92,0.917,0.927 | 0.079 | ENSG00000221 | [[[[26291508, 262 | 2 |
| ENSG00000137 | FLOT1 | chr6 | 30741613 | 30741704 | 30740678 | 30740798 | 30742146 | 30742203 | 2,0,4 | 2,1,2 | 8,16,8 | 0,2,0 | 241 | 150 | 0.000146875 | 0.002223205 | 0.384,0.0,0.555 | 1.0,0.833,1.0 | -0.631 | ENSG00000137 | [[[[30742527, 307 | 3 |
| ENSG00000137 | FLOT1 | chr6 | 30741613 | 30741704 | 30741189 | 30741333 | 30742146 | 30742203 | 301,255,393 | 17,12,14 | 132,258,162 | 21,60,28 | 241 | 150 | 7.03E-14 | 7.44E-12 | 0.917,0.93,0.946 | 0.796,0.728,0.78 | 0.162 | ENSG00000137 | [[[[30742527, 307 | 2 |
| ENSG00000101 | NOP56 | chr20 | 2655933 | 2656034 | 2655594 | 2655746 | 2656773 | 2656895 | 53,97,126 | 0,0,0 | 72,212,160 | 7,6,2 | 251 | 150 | 0.000219954 | 0.003164072 | 1.0,1.0,1.0 | 0.86,0.955,0.98 | 0.068 | ENSG00000101 | [[[[2652632, 2652 | 2 |
| ENSG00000085 | EPS15 | chr1 | 51448045 | 51448135 | 51421785 | 51421858 | 51461090 | 51461150 | 13,13,33 | 0,0,0 | 21,30,40 | 0,6,6 | 240 | 150 | 8.83E-06 | 0.00019162 | 1.0,1.0,1.0 | 1.0,0.758,0.806 | 0.145 | ENSG00000085 | [[[[51519199, 515 | 4 |
| ENSG00000063( | MTMR1 | chrX | 150755688 | 150755865 | 150744360 | 150744453 | 150762564 | 150762737 | 5,2,20 | 0,0,0 | 7,7,10 | 1,1,7 | 300 | 150 | 4.75E-07 | 1.42E-05 | 1.0,1.0,1.0 | 0.778,0.778,0.41 | 0.342 | ENSG00000063( | [[[[150693396, 15 | 2 |
| ENSG00000253 | PRKDC | chr8 | 47779003 | 47779093 | 47778458 | 47778660 | 47782377 | 47782598 | 7,2,3 | 0,0,0 | 10,12,11 | 3,6,3 | 240 | 150 | 9.80E-08 | 3.45E-06 | 1.0,1.0,1.0 | 0.676,0.556,0.69 | 0.357 | ENSG00000253 | [[[[47959973, 479 | 3 |
| ENSG00000092 | TRPM7 | chr15 | 50604865 | 50605144 | 50599121 | 50599296 | 50609580 | 50609724 | 17,15,31 | 0,0,0 | 21,57,34 | 14,13,5 | 300 | 150 | 2.05E-09 | 1.07E-07 | 1.0,1.0,1.0 | 0.429,0.687,0.77 | 0.37 | ENSG00000092 | [[[[50686531, 50( | 2 |
| ENSG00000092 | TRPM7 | chr15 | 50613706 | 50613841 | 50609805 | 50609961 | 50614122 | 50614263 | 19,16,27 | 0,0,0 | 17,54,45 | 1,7,2 | 285 | 150 | 3.71E-05 | 0.000674761 | 1.0,1.0,1.0 | 0.899,0.802,0.92 | 0.126 | ENSG00000092 | [[[[50686531, 50( | 3 |
| ENSG00000073( | PVR | chr19 | 44657761 | 44657910 | 44653899 | 44654017 | 44661291 | 44661323 | 52,46,78 | 2,3,5 | 41,124,70 | 59,88,73 | 299 | 150 | 0 | 0 | 0.929,0.885,0.88 | 0.259,0.414,0.32 | 0.568 | ENSG00000073( | [[[[44643910, 44( | 2 |
| ENSG00000073( | PVR | chr19 | 44658741 | 44658900 | 44653899 | 44654017 | 44661291 | 44661323 | 36,46,58 | 2,3,5 | 32,47,32 | 59,88,73 | 300 | 150 | 0 | 0 | 0.9,0.885,0.853 | 0.213,0.211,0.18 | 0.678 | ENSG00000073( | [[[[44643910, 44( | 2 |
| ENSG00000112 | C6orf62 | chr6 | 24716147 | 24716324 | 24708776 | 24708911 | 24719859 | 24719998 | 7,6,5 | 0,0,0 | 5,16,4 | 17,27,30 | 300 | 150 | 1.20E-14 | 1.42E-12 | 1.0,1.0,1.0 | 0.128,0.229,0.06 | 0.86 | ENSG00000112 | [[[[24718540, 247 | 3 |
| ENSG00000118 | TDP2 | chr6 | 24658560 | 24658734 | 24654411 | 24654530 | 24666525 | 24666611 | 78,70,106 | 0,0,2 | 57,177,128 | 2,10,6 | 300 | 150 | 5.56E-05 | 0.000956995 | 1.0,1.0,0.964 | 0.934,0.898,0.91 | 0.073 | ENSG00000118 | [[[[24666698, 24( | 2 |
| ENSG00000137' | KIF13A | chr6 | 17851954 | 17852042 | 17837471 | 17837583 | 17855436 | 17855617 | 4,4,13 | 0,0,0 | 7,19,21 | 2,7,3 | 238 | 150 | 4.62E-07 | 1.38E-05 | 1.0,1.0,1.0 | 0.688,0.631,0.81 | 0.289 | ENSG00000137' | [[[[17987409, 179 | 3 |
| ENSG00000069 | NUP133 | chr1 | 229458160 | 229458296 | 229452524 | 229452643 | 229463542 | 229463676 | 41,37,59 | 0,0,0 | 29,50,15 | 5,7,6 | 286 | 150 | 1.23E-09 | 6.68E-08 | 1.0,1.0,1.0 | 0.753,0.789,0.56 | 0.297 | ENSG00000069 | [[[[229508068, 2 | 2 |
| ENSG00000110( | CARS | chr11 | 3040895 | 3040984 | 3039193 | 3039292 | 3042164 | 3042256 | 39,51,157 | 0,0,0 | 89,280,189 | 3,11,11 | 239 | 150 | 5.63E-06 | 0.000128882 | 1.0,1.0,1.0 | 0.949,0.941,0.91 | 0.065 | ENSG00000110( | [[[[3057343, 3057 | 2 |
| ENSG00000110( | CARS | chr11 | 3040895 | 3041031 | 3039193 | 3039292 | 3042164 | 3042256 | 3,5,14 | 0,0,0 | 18,18,29 | 3,11,11 | 286 | 150 | 5.57E-08 | 2.08E-06 | 1.0,1.0,1.0 | 0.759,0.462,0.58 | 0.4 | ENSG00000110( | [[[[3057343, 3057 | 2 |
| ENSG00000147( | VLDLR | chr9 | 2648668 | 2648810 | 2647473 | 2647592 | 2651414 | 2651498 | 1,5,2 | 0,0,0 | 37,65,40 | 6,8,2 | 292 | 150 | 2.84E-05 | 0.000532659 | 1.0,1.0,1.0 | 0.76,0.807,0.911 | 0.174 | ENSG00000147( | [[[[2621787, 2622 | 3 |
| ENSG00000147( | VLDLR | chr9 | 2648668 | 2648810 | 2647473 | 2647592 | 2651873 | 2651954 | 1,5,2 | 0,0,0 | 38,63,46 | 2,3,5 | 292 | 150 | 0.000249016 | 0.003509246 | 1.0,1.0,1.0 | 0.907,0.915,0.82 | 0.118 | ENSG00000147( | [[[[2621787, 2622 | 4 |
| ENSG00000147( | VLDLR | chr9 | 2650369 | 2650516 | 2647473 | 2647592 | 2651414 | 2651498 | 120,95,132 | 0,0,0 | 70,155,70 | 6,8,2 | 297 | 150 | 1.12E-06 | 3.04E-05 | 1.0,1.0,1.0 | 0.855,0.907,0.94 | 0.097 | ENSG00000147( | [[[[2621787, 2622 | 3 |
| ENSG00000147( | VLDLR | chr9 | 2650369 | 2650516 | 2647473 | 2647592 | 2651873 | 2651954 | 91,70,101 | 0,0,0 | 52,92,53 | 2,3,5 | 297 | 150 | 0.000161054 | 0.002410998 | 1.0,1.0,1.0 | 0.929,0.939,0.84 | 0.096 | ENSG00000147( | [[[[2621787, 2622 | 4 |
| ENSG00000147( | VLDLR | chr9 | 2650369 | 2650516 | 2648668 | 2648810 | 2651873 | 2651954 | 340,231,364 | 0,0,0 | 203,342,228 | 5,7,10 | 297 | 150 | 1.09E-06 | 2.97E-05 | 1.0,1.0,1.0 | 0.953,0.961,0.92 | 0.055 | ENSG00000147( | [[[[2621787, 2622 | 2 |
| ENSG00000147( | VLDLR | chr9 | 2651414 | 2651498 | 2648668 | 2648810 | 2651873 | 2651954 | 66,61,93 | 0,0,0 | 67,98,55 | 5,7,10 | 234 | 150 | 1.18E-08 | 5.19E-07 | 1.0,1.0,1.0 | 0.896,0.9,0.779 | 0.142 | ENSG00000147( | [[[[2621787, 2622 | 2 |
| ENSG00000143 | INTS7 | chr1 | 211975165 | 211975372 | 211968512 | 211968707 | 211978271 | 211978511 | 27,25,52 | 0,0,0 | 20,46,28 | 15,23,8 | 300 | 150 | 1.22E-15 | 1.64E-13 | 1.0,1.0,1.0 | 0.4,0.5,0.636 | 0.488 | ENSG00000143 | [[[[212035344, 2' | 2 |
| ENSG00000143 | INTS7 | chr1 | 211976581 | 211976719 | 211968512 | 211968707 | 211978271 | 211978511 | 26,33,49 | 0,0,0 | 16,47,23 | 15,23,8 | 288 | 150 | 2.22E-16 | 3.17E-14 | 1.0,1.0,1.0 | 0.357,0.516,0.6 | 0.509 | ENSG00000143 | [[[[212035344, 2' | 2 |
| ENSG00000124 | NQO2 | chr6 | 3016883 | 3016985 | 3012543 | 3012674 | 3019478 | 3019755 | 24,35,49 | 0,0,1 | 79,121,63 | 8,12,10 | 252 | 150 | 4.42E-07 | 1.33E-05 | 1.0,1.0,0.967 | 0.855,0.857,0.78 | 0.155 | ENSG00000124 | [[[[2999894, 300( | 2 |
| ENSG00000113 | CNOT6 | chr5 | 180569109 | 180569340 | 180565819 | 180565977 | 180573987 | 180578349 | 2,4,1 | 0,0,0 | 4,6,4 | 5,0,4 | 300 | 150 | 2.07E-06 | 5.28E-05 | 1.0,1.0,1.0 | 0.286,1.0,0.333 | 0.46 | ENSG00000113 | [[[[180494379, 18 | 4 |
| ENSG00000122 | KIAA1191 | chr5 | 176352621 | 176352748 | 176348249 | 176348356 | 176359480 | 176359567 | 0,0,0 | 14,2,16 | 2,8,6 | 20,60,33 | 277 | 150 | 0.000211308 | 0.003056713 | 0.0,0.0,0.0 | 0.051,0.067,0.09 | -0.069 | ENSG00000122 | [[[[176361602, 17 | 3 |
| ENSG00000122 | KIAA1191 | chr5 | 176359480 | 176359567 | 176350612 | 176350737 | 176361601 | 176361761 | 0,0,1 | 1,1,2 | 2,7,9 | 1,3,4 | 237 | 150 | 6.98E-06 | 0.000155315 | 0.0,0.0,0.24 | 0.559,0.596,0.58 | -0.501 | ENSG00000122 | [[[[176361602, 17 | 4 |
| ENSG00000122 | KIAA1191 | chr5 | 176359480 | 176359567 | 176352621 | 176352748 | 176361601 | 176361761 | 0,0,1 | 1,3,6 | 2,5,13 | 2,7,7 | 237 | 150 | 1.52E-06 | 4.00E-05 | 0.0,0.0,0.095 | 0.388,0.311,0.54 | -0.381 | ENSG00000122 | [[[[176361602, 17 | 3 |
| ENSG00000174 | SH3PXD2B | chr5 | 172382035 | 172382127 | 172373789 | 172373815 | 172406276 | 172406352 | 20,9,43 | 0,0,0 | 43,153,146 | 0,5,12 | 242 | 150 | 0.000249911 | 0.00352056 | 1.0,1.0,1.0 | 1.0,0.95,0.883 | 0.056 | ENSG00000174 | [[[[172454278, 17 | 2 |
| ENSG00000174 | SH3PXD2B | chr5 | 172394562 | 172394639 | 172373789 | 172373815 | 172406276 | 172406352 | 12,1,21 | 0,0,0 | 11,19,32 | 0,5,12 | 227 | 150 | 1.09E-06 | 2.96E-05 | 1.0,1.0,1.0 | 1.0,0.715,0.638 | 0.216 | ENSG00000174 | [[[[172454278, 17 | 2 |
| ENSG00000144( | NXPE3 | chr3 | 101807052 | 101807126 | 101801234 | 101801989 | 101821403 | 101822823 | 26,29,45 | 0,0,0 | 44,55,31 | 12,18,16 | 224 | 150 | 5.04E-14 | 5.46E-12 | 1.0,1.0,1.0 | 0.711,0.672,0.56 | 0.351 | ENSG00000144( | [[[[101779211, 1( | 2 |
| ENSG00000144( | NXPE3 | chr3 | 101816794 | 101817001 | 101801234 | 101801989 | 101821403 | 101822823 | 73,60,102 | 0,0,0 | 50,105,59 | 12,18,16 | 300 | 150 | 0 | 0 | 1.0,1.0,1.0 | 0.676,0.745,0.64 | 0.31 | ENSG00000144( | [[[[101779211, 1( | 2 |
| ENSG00000135( | ADAM19 | chr5 | 157502802 | 157502980 | 157499572 | 157499662 | 157507055 | 157507140 | 169,135,441 | 1,0,0 | 111,372,282 | 4,21,13 | 300 | 150 | 6.32E-11 | 4.26E-09 | 0.988,1.0,1.0 | 0.933,0.899,0.91 | 0.08 | ENSG00000135( | [[[[157575603, 15 | 2 |
| ENSG00000135( | ADAM19 | chr5 | 157505668 | 157505808 | 157499572 | 157499662 | 157507055 | 157507140 | 100,162,332 | 1,0,0 | 132,327,233 | 4,21,13 | 290 | 150 | 1.92E-10 | 1.21E-08 | 0.981,1.0,1.0 | 0.945,0.89,0.903 | 0.081 | ENSG00000135( | [[[[157575603, 15 | 2 |
| ENSG00000155 | CNOT8 | chr5 | 154863206 | 154863395 | 154858545 | 154858768 | 154870660 | 154870822 | 8,7,17 | 0,0,0 | 14,22,21 | 7,1,3 | 300 | 150 | 1.78E-06 | 4.60E-05 | 1.0,1.0,1.0 | 0.5,0.917,0.778 | 0.268 | ENSG00000155 | [[[[154858627, 15 | 2 |
| ENSG00000155 | CNOT8 | chr5 | 154865191 | 154865385 | 154858538 | 154858768 | 154870660 | 154870704 | 52,51,99 | 0,0,0 | 35,67,57 | 7,1,3 | 300 | 150 | 1.65E-05 | 0.000330704 | 1.0,1.0,1.0 | 0.714,0.971,0.90 | 0.137 | ENSG00000155 | [[[[154858627, 15 | 2 |
| ENSG00000143 | EIF2D | chr1 | 206599462 | 206599612 | 206599002 | 206599092 | 206602262 | 206600308 | 63,63,78 | 2,3,1 | 40,87,61 | 17,37,19 | 300 | 150 | 1.22E-15 | 1.64E-13 | 0.94,0.913,0.975 | 0.541,0.54,0.616 | 0.377 | ENSG00000143 | [[[[206612287, 2( | 2 |
| ENSG00000143 | EIF2D | chr1 | 206599732 | 206599836 | 206599002 | 206599092 | 206602262 | 206600308 | 35,23,31 | 2,3,1 | 24,45,44 | 17,37,19 | 254 | 150 | 1.01E-09 | 5.57E-08 | 0.912,0.819,0.94 | 0.455,0.418,0.57 | 0.409 | ENSG00000143 | [[[[206612287, 2( | 2 |
| ENSG00000143 | EIF2D | chr1 | 206602950 | 206603204 | 206600262 | 206600308 | 206605399 | 206605507 | 32,43,95 | 0,0,0 | 31,71,45 | 3,6,2 | 300 | 150 | 1.58E-05 | 0.00031863 | 1.0,1.0,1.0 | 0.838,0.855,0.91 | 0.13 | ENSG00000143 | [[[[206612287, 2( | 2 |
| ENSG00000143 | EIF2D | chr1 | 206608235 | 206608326 | 206605399 | 206605507 | 206611183 | 206611374 | 27,29,49 | 0,0,0 | 26,43,26 | 3,6,3 | 241 | 150 | 2.38E-06 | 5.96E-05 | 1.0,1.0,1.0 | 0.844,0.817,0.84 | 0.165 | ENSG00000143 | [[[[206612287, 2( | 2 |
| ENSG00000178 | DTX3 | chr12 | 57606864 | 57607613 | 57606442 | 57606518 | 57609076 | 57609800 | 35,28,49 | 0,0,0 | 28,52,33 | 2,3,3 | 300 | 150 | 0.000313579 | 0.004292094 | 1.0,1.0,1.0 | 0.875,0.897,0.84 | 0.127 | ENSG00000178 | [[[[57604824, 57( | 2 |
| ENSG00000178 | DTX3 | chr12 | 57606883 | 57607613 | 57606442 | 57606518 | 57609076 | 57609804 | 4,5,9 | 0,0,0 | 6,9,5 | 2,3,3 | 300 | 150 | 7.77E-08 | 2.81E-06 | 1.0,1.0,1.0 | 0.6,0.6,0.455 | 0.448 | ENSG00000178 | [[[[57604824, 57( | 2 |
| ENSG00000113 | CSNK1A1 | chr5 | 149505482 | 149505595 | 149492196 | 149496860 | 149520289 | 149520388 | 57,44,65 | 1,0,0 | 77,80,67 | 5,7,7 | 263 | 150 | 1.67E-06 | 4.35E-05 | 0.97,1.0,1.0 | 0.898,0.867,0.84 | 0.12 | ENSG00000113 | [[[[149550842, 14 | 5 |
| ENSG00000166 | AKIP1 | chr11 | 8914825 | 8914930 | 8912452 | 8912533 | 8919336 | 8919537 | 39,46,54 | 0,0,0 | 80,113,55 | 7,9,2 | 255 | 150 | 2.72E-06 | 6.73E-05 | 1.0,1.0,1.0 | 0.871,0.881,0.94 | 0.102 | ENSG00000166 | [[[[8911189, 8911 | 2 |
| ENSG00000166 | AP1G1 | chr16 | 71764349 | 71764448 | 71761511 | 71761567 | 71765488 | 71765584 | 25,14,27 | 0,0,0 | 3,15,10 | 6,14,12 | 249 | 150 | 0 | 0 | 1.0,1.0,1.0 | 0.231,0.392,0.33 | 0.681 | ENSG00000166 | [[[[71808763, 718 | 2 |
| ENSG00000113( | TCERG1 | chr5 | 146503374 | 146503539 | 146498535 | 146498686 | 146507027 | 146507207 | 49,30,59 | 0,0,0 | 23,82,44 | 9,22,7 | 300 | 150 | 4.36E-13 | 4.16E-11 | 1.0,1.0,1.0 | 0.561,0.651,0.75 | 0.343 | ENSG00000113( | [[[[146447333, 14 | 2 |
| ENSG00000197( | ZNF138 | chr7 | 64815571 | 64815653 | 64794427 | 64794571 | 64831450 | 64833327 | 4,3,5 | 0,2,0 | 2,7,1 | 1,2,5 | 232 | 150 | 0.000808684 | 0.00969632 | 1.0,0.492,1.0 | 0.564,0.694,0.11 | 0.373 | ENSG00000197( | [[[[64794428, 64( | 2 |
| ENSG00000086 | HUWE1 | chrX | 53543885 | 53543968 | 53542442 | 53542539 | 53545028 | 53545161 | 17,22,40 | 0,0,0 | 34,72,49 | 1,12,3 | 233 | 150 | 6.98E-06 | 0.000155315 | 1.0,1.0,1.0 | 0.956,0.794,0.91 | 0.112 | ENSG00000086 | [[[[53686588, 53( | 2 |
| ENSG00000086 | HUWE1 | chrX | 53544559 | 53544762 | 53542442 | 53542539 | 53545028 | 53545161 | 106,79,152 | 0,0,0 | 73,182,81 | 1,12,3 | 300 | 150 | 3.69E-06 | 8.83E-05 | 1.0,1.0,1.0 | 0.973,0.883,0.93 | 0.071 | ENSG00000086 | [[[[53686588, 53( | 2 |
| ENSG00000163 | GYG1 | chr3 | 149024052 | 149024272 | 149009275 | 149009402 | 149026759 | 149027632 | 110,106,134 | 3,0,5 | 97,182,114 | 9,16,8 | 300 | 150 | 8.14E-05 | 0.001329364 | 0.948,1.0,0.931 | 0.843,0.85,0.877 | 0.103 | ENSG00000163 | [[[[148991540, 14 | 2 |
| ENSG00000186 | PRELID2 | chr5 | 145817893 | 145818054 | 145764930 | 145765000 | 145819944 | 145820018 | 15,8,29 | 0,0,0 | 13,27,6 | 2,6,2 | 300 | 150 | 3.57E-07 | 1.10E-05 | 1.0,1.0,1.0 | 0.765,0.692,0.6 | 0.314 | ENSG00000186 | [[[[145835177, 14 | 2 |
| ENSG00000197( | FCHSD1 | chr5 | 141646602 | 141646722 | 141646086 | 141646191 | 141647397 | 141647520 | 4,7,6 | 0,0,0 | 16,24,21 | 3,4,3 | 270 | 150 | 1.19E-05 | 0.000249232 | 1.0,1.0,1.0 | 0.748,0.769,0.79 | 0.229 | ENSG00000197( | [[[[141651348, 14 | 2 |
| ENSG00000132 | GPS2 | chr17 | 7313211 | 7313291 | 7313028 | 7313097 | 7313905 | 7313988 | 1,2,3 | 0,0,0 | 15,44,23 | 3,7,4 | 230 | 150 | 2.21E-05 | 0.0004297 | 1.0,1.0,1.0 | 0.765,0.804,0.78 | 0.214 | ENSG00000132 | [[[[7315331, 731 | 3 |
| ENSG00000170 | HARS | chr5 | 140677654 | 140677753 | 140675016 | 140675133 | 140677907 | 140678015 | 143,118,161 | 0,0,0 | 73,140,79 | 14,16,12 | 249 | 150 | 0 | 0 | 1.0,1.0,1.0 | 0.759,0.841,0.79 | 0.2 | ENSG00000170 | [[[[140691215, 14 | 4 |
| ENSG00000170 | HARS | chr5 | 140683099 | 140683219 | 140677907 | 140678015 | 140690854 | 140690944 | 83,69,127 | 0,0,0 | 40,76,42 | 11,13,10 | 270 | 150 | 1.11E-16 | 1.65E-14 | 1.0,1.0,1.0 | 0.669,0.765,0.7 | 0.289 | ENSG00000170 | [[[[140691215, 14 | 3 |

| | | | | | | | | | | | | | | | | | | | | | | | |
|---|---|---|---|---|---|---|---|---|---|---|---|---|---|---|---|---|---|---|---|---|---|---|---|
| ENSG00000138 | ACTR1A | chr10 | 102481836 | 102481898 | 102481131 | 102481172 | 102483010 | 102483103 | 64,54,82 | 6,4,4 | 43,89,39 | 10,28,18 | 212 | 150 | 3.02E-07 | 9.46E-06 | 0.883,0.905,0.93 | 0.753,0.692,0.60 | 0.225 | ENSG00000138 | [[[[102502600, 1 | | 2 |
| ENSG00000138 | ACTR1A | chr10 | 102482000 | 102482175 | 102481131 | 102481172 | 102483010 | 102483103 | 204,184,279 | 6,4,4 | 169,329,206 | 10,28,18 | 300 | 150 | 1.00E-06 | 2.76E-05 | 0.944,0.958,0.97 | 0.894,0.855,0.85 | 0.091 | ENSG00000138 | [[[[102502600, 1 | | 2 |
| ENSG00000154 | MATR3 | chr5 | 139314674 | 139314736 | 139293673 | 139293772 | 139315696 | 139315738 | 118,76,168 | 0,0,0 | 114,274,202 | 4,13,14 | 212 | 150 | 1.05E-07 | 3.65E-06 | 1.0,1.0,1.0 | 0.953,0.937,0.91 | 0.066 | ENSG00000154 | [[[[139293740, 1 | | 2 |
| ENSG00000106 | ANKMY2 | chr7 | 16624982 | 16625081 | 16615743 | 16615904 | 16645506 | 16645755 | 25,25,47 | 0,0,0 | 29,64,56 | 2,8,5 | 249 | 150 | 2.10E-06 | 5.34E-05 | 1.0,1.0,1.0 | 0.897,0.828,0.87 | 0.135 | ENSG00000106 | [[[[16645507, 16 | | 3 |
| ENSG00000106 | ANKMY2 | chr7 | 16636390 | 16636455 | 16615743 | 16615904 | 16645506 | 16645755 | 20,9,22 | 0,0,0 | 15,42,32 | 2,8,5 | 215 | 150 | 1.01E-06 | 2.78E-05 | 1.0,1.0,1.0 | 0.84,0.786,0.817 | 0.186 | ENSG00000106 | [[[[16645507, 16 | | 3 |
| ENSG00000158 | CDC25C | chr5 | 138289500 | 138289563 | 138287168 | 138287267 | 138291969 | 138292116 | 19,12,14 | 1,0,0 | 5,14,6 | 2,4,1 | 213 | 150 | 0.000100956 | 0.001609674 | 0.93,1.0,1.0 | 0.638,0.711,0.80 | 0.257 | ENSG00000158 | [[[[138331595, 1 | | 2 |
| ENSG00000158 | CDC25C | chr5 | 138329552 | 138329647 | 138328483 | 138328529 | 138331594 | 138331835 | 31,19,25 | 0,0,2 | 6,32,17 | 3,8,1 | 245 | 150 | 6.91E-05 | 0.001157504 | 1.0,1.0,0.884 | 0.55,0.71,0.912 | 0.237 | ENSG00000158 | [[[[138331595, 1 | | 2 |
| ENSG00000158 | CDC25C | chr5 | 138330986 | 138331214 | 138328483 | 138328529 | 138331594 | 138331794 | 23,15,21 | 0,0,2 | 12,38,11 | 3,8,1 | 300 | 150 | 9.91E-05 | 0.001586173 | 1.0,1.0,0.84 | 0.667,0.704,0.84 | 0.208 | ENSG00000158 | [[[[138331595, 1 | | 2 |
| ENSG00000158 | CDC25C | chr5 | 138330986 | 138331218 | 138328483 | 138328529 | 138331594 | 138331835 | 22,15,14 | 0,0,2 | 14,40,17 | 3,8,1 | 300 | 150 | 0.000398262 | 0.005276684 | 1.0,1.0,0.778 | 0.7,0.714,0.895 | 0.156 | ENSG00000158 | [[[[138331595, 1 | | 2 |
| ENSG00000111 | UST | chr6 | 148941278 | 148941434 | 148886985 | 148887029 | 148964409 | 148964563 | 16,12,29 | 0,0,0 | 10,45,10 | 1,5,7 | 300 | 150 | 4.32E-07 | 1.30E-05 | 1.0,1.0,1.0 | 0.833,0.818,0.41 | 0.311 | ENSG00000111 | [[[[148747030, 14 | | 2 |
| ENSG00000111 | UST | chr6 | 148953871 | 148953951 | 148886985 | 148887029 | 148964409 | 148964563 | 6,6,17 | 0,0,0 | 11,14,5 | 1,5,7 | 230 | 150 | 1.54E-07 | 5.13E-06 | 1.0,1.0,1.0 | 0.878,0.646,0.31 | 0.386 | ENSG00000111 | [[[[148747030, 14 | | 2 |
| ENSG00000161 | LARP4 | chr12 | 50454313 | 50454417 | 50441589 | 50441643 | 50461134 | 50461347 | 23,13,42 | 0,0,1 | 33,67,42 | 5,5,0 | 254 | 150 | 0.000647946 | 0.008012362 | 1.0,1.0,0.961 | 0.796,0.888,1.0 | 0.092 | ENSG00000161 | [[[[50400885, 504 | | 2 |
| ENSG00000196 | PPIA | chr7 | 44799701 | 44799874 | 44799246 | 44799277 | 44801286 | 44801630 | 783,371,159,454 | 7,0,5 | 6,152,126,636,7 | 77,141,70 | 300 | 150 | 4.12E-14 | 4.52E-12 | 0.998,1.0,0.999 | 0.976,0.978,0.98 | 0.021 | ENSG00000196 | [[[[44796681, 447 | | 2 |
| ENSG00000196 | PPIA | chr7 | 44800423 | 44800543 | 44799246 | 44799277 | 44801286 | 44801624 | 7,25,26 | 7,0,5 | 37,78,35 | 77,141,70 | 270 | 150 | 5.74E-05 | 0.000983389 | 0.357,1.0,0.743 | 0.211,0.235,0.21 | 0.479 | ENSG00000196 | [[[[44796681, 447 | | 2 |
| ENSG00000148 | PARD3 | chr10 | 34374873 | 34375002 | 34360070 | 34360259 | 34377966 | 34378106 | 19,14,20 | 0,2,0 | 4,9,7 | 2,5,3 | 279 | 150 | 1.99E-06 | 5.10E-05 | 1.0,0.79,1.0 | 0.518,0.492,0.55 | 0.408 | ENSG00000148 | [[[[34814876, 348 | | 2 |
| ENSG00000119 | UBE2B | chr5 | 134380718 | 134380808 | 134376668 | 134376694 | 134390224 | 134390321 | 86,87,118 | 0,0,0 | 129,250,176 | 6,5,6 | 240 | 150 | 0.000352553 | 0.004741231 | 1.0,1.0,1.0 | 0.931,0.969,0.94 | 0.051 | ENSG00000119 | [[[[134371569, 1 | | 2 |
| ENSG00000119 | UBE2B | chr5 | 134388324 | 134388413 | 134376668 | 134376694 | 134390224 | 134390298 | 113,93,144 | 0,0,0 | 164,263,198 | 6,5,6 | 239 | 150 | 0.00057028 | 0.007192017 | 1.0,1.0,1.0 | 0.945,0.971,0.95 | 0.043 | ENSG00000119 | [[[[134371569, 1 | | 2 |
| ENSG00000119 | UBE2B | chr5 | 134388351 | 134388413 | 134376668 | 134376694 | 134390224 | 134390573 | 80,60,107 | 0,0,0 | 126,189,131 | 6,5,6 | 212 | 150 | 0.000204397 | 0.002968854 | 1.0,1.0,1.0 | 0.937,0.964,0.93 | 0.053 | ENSG00000119 | [[[[134371569, 1 | | 2 |
| ENSG00000121 | TMEM131L | chr4 | 153556965 | 153557082 | 153555786 | 153555910 | 153580825 | 153580903 | 17,4,17 | 0,0,0 | 7,23,19 | 0,1,7 | 267 | 150 | 0.000165705 | 0.002470428 | 1.0,1.0,1.0 | 1.0,0.928,0.604 | 0.156 | ENSG00000121 | [[[[153466360, 1 | | 2 |
| ENSG00000189 | RNFT1 | chr17 | 59960067 | 59960168 | 59956487 | 59956553 | 59962539 | 59962599 | 20,15,13 | 0,0,0 | 7,15,7 | 2,3,2 | 251 | 150 | 2.00E-05 | 0.000392544 | 1.0,1.0,1.0 | 0.677,0.749,0.67 | 0.299 | ENSG00000189 | [[[[59964608, 599 | | 3 |
| ENSG00000136 | TBRG4 | chr7 | 45104098 | 45104256 | 45103030 | 45103443 | 45105440 | 45105669 | 45,103,124 | 1,0,3 | 50,118,62 | 26,44,29 | 300 | 150 | 0 | 0 | 0.957,1.0,0.954 | 0.49,0.573,0.517 | 0.444 | ENSG00000136 | [[[[45111643, 451 | | 2 |
| ENSG00000136 | TBRG4 | chr7 | 45104537 | 45104709 | 45103030 | 45103443 | 45105440 | 45105478 | 49,39,97 | 1,0,3 | 43,145,104 | 26,44,29 | 300 | 150 | 3.38E-14 | 3.76E-12 | 0.961,1.0,0.942 | 0.453,0.622,0.64 | 0.395 | ENSG00000136 | [[[[45111643, 451 | | 2 |
| ENSG00000170 | NFXL1 | chr4 | 47875126 | 47875293 | 47862845 | 47862915 | 47879095 | 47879117 | 8,8,25 | 0,0,0 | 4,22,8 | 0,6,1 | 300 | 150 | 0.000125495 | 0.00194162 | 1.0,1.0,1.0 | 1.0,0.647,0.8 | 0.184 | ENSG00000170 | [[[[47914365, 479 | | 2 |
| ENSG00000088 | CRLS1 | chr20 | 6015360 | 6015490 | 6009774 | 6009912 | 6032011 | 6032080 | 47,40,67 | 0,3,7 | 67,81,71 | 2,0,0 | 280 | 150 | 0.000475419 | 0.006165072 | 1.0,0.877,0.837 | 0.947,1.0,1.0 | -0.078 | ENSG00000088 | [[[[6006123, 600 | | 2 |
| ENSG00000127 | SGCE | chr7 | 94603289 | 94603452 | 94599696 | 94599723 | 94618757 | 94618932 | 153,145,191 | 0,0,0 | 106,195,98 | 24,41,29 | 300 | 150 | 0 | 0 | 1.0,1.0,1.0 | 0.688,0.704,0.62 | 0.327 | ENSG00000127 | [[[[94655990, 94 | | 2 |
| ENSG00000127 | SGCE | chr7 | 94628201 | 94628359 | 94623324 | 94623397 | 94655989 | 94656045 | 83,83,114 | 0,0,0 | 62,105,71 | 16,24,42 | 300 | 150 | 0 | 0 | 1.0,1.0,1.0 | 0.66,0.686,0.458 | 0.399 | ENSG00000127 | [[[[94655990, 94 | | 2 |
| ENSG00000127 | SGCE | chr7 | 94629718 | 94629841 | 94623324 | 94623397 | 94655989 | 94655993 | 41,64,106 | 0,0,0 | 46,68,51 | 16,24,42 | 273 | 150 | 0 | 0 | 1.0,1.0,1.0 | 0.612,0.609,0.4 | 0.46 | ENSG00000127 | [[[[94655990, 94 | | 2 |
| ENSG00000127 | SGCE | chr7 | 94639340 | 94639448 | 94623324 | 94623397 | 94655989 | 94656079 | 2,6,8 | 0,0,0 | 2,6,12 | 16,24,42 | 258 | 150 | 2.78E-15 | 3.55E-13 | 1.0,1.0,1.0 | 0.068,0.127,0.14 | 0.888 | ENSG00000127 | [[[[94655990, 94 | | 2 |
| ENSG00000164 | ERAP2 | chr5 | 96896731 | 96896863 | 96896372 | 96896504 | 96901505 | 96901681 | 15,7,19 | 0,0,0 | 7,21,14 | 2,8,6 | 282 | 150 | 2.11E-09 | 1.10E-07 | 1.0,1.0,1.0 | 0.651,0.583,0.55 | 0.404 | ENSG00000164 | [[[[96876500, 968 | | 2 |
| ENSG00000153 | CAST | chr5 | 96727488 | 96727530 | 96722638 | 96722698 | 96729152 | 96729209 | 195,141,331 | 0,0,0 | 136,425,234 | 6,21,12 | 192 | 150 | 1.24E-09 | 6.73E-08 | 1.0,1.0,1.0 | 0.947,0.941,0.93 | 0.058 | ENSG00000153 | [[[[96662376, 96 | | 2 |
| ENSG00000153 | CAST | chr5 | 96727488 | 96727530 | 96722638 | 96722698 | 96729611 | 96729725 | 73,42,129 | 0,1,0 | 69,185,105 | 3,18,6 | 192 | 150 | 3.91E-07 | 1.19E-05 | 1.0,0.97,1.0 | 0.947,0.889,0.93 | 0.067 | ENSG00000153 | [[[[96662376, 96 | | 3 |
| ENSG00000153 | CAST | chr5 | 96729152 | 96729209 | 96722638 | 96722698 | 96729611 | 96729691 | 69,47,121 | 0,1,0 | 46,162,123 | 3,18,6 | 207 | 150 | 1.46E-07 | 4.90E-06 | 1.0,0.971,1.0 | 0.917,0.867,0.93 | 0.083 | ENSG00000153 | [[[[96662376, 96 | | 3 |
| ENSG00000153 | CAST | chr5 | 96740744 | 96740783 | 96740037 | 96740118 | 96741493 | 96741580 | 12,15,20 | 0,0,0 | 12,25,26 | 4,3,3 | 189 | 150 | 1.42E-05 | 0.000290485 | 1.0,1.0,1.0 | 0.704,0.869,0.86 | 0.186 | ENSG00000153 | [[[[96662376, 96 | | 2 |
| ENSG00000109 | SLAIN2 | chr4 | 48377895 | 48378060 | 48369848 | 48369997 | 48382567 | 48382927 | 35,25,52 | 0,0,0 | 34,53,38 | 7,10,7 | 300 | 150 | 2.37E-10 | 1.47E-08 | 1.0,1.0,1.0 | 0.708,0.726,0.73 | 0.278 | ENSG00000109 | [[[[48341529, 48 | | 2 |
| ENSG00000164 | ADGRV1 | chr5 | 90848636 | 90848821 | 90840577 | 90840985 | 90965414 | 90965531 | 19,9,18 | 0,0,0 | 16,35,13 | 3,5,1 | 300 | 150 | 5.40E-06 | 0.000124301 | 1.0,1.0,1.0 | 0.727,0.778,0.86 | 0.209 | ENSG00000164 | [[[[90558797, 90 | | 5 |
| ENSG00000164 | ADGRV1 | chr5 | 90854061 | 90854201 | 90853283 | 90853533 | 90965414 | 90965531 | 26,10,17 | 0,0,0 | 9,33,13 | 2,3,2 | 290 | 150 | 0.000200091 | 0.002912984 | 1.0,1.0,1.0 | 0.699,0.851,0.77 | 0.226 | ENSG00000164 | [[[[90558797, 90 | | 3 |
| ENSG00000214 | FIS1 | chr7 | 101244006 | 101244139 | 101240829 | 101240906 | 101251894 | 101252133 | 120,149,172 | 3,11,19 | 146,248,128 | 46,96,49 | 283 | 150 | 1.65E-14 | 1.94E-12 | 0.955,0.878,0.82 | 0.627,0.578,0.58 | 0.292 | ENSG00000214 | [[[[101244960, 1 | | 2 |
| ENSG00000078 | PIK3C3 | chr18 | 42057882 | 42058051 | 42049530 | 42049605 | 42067387 | 42067513 | 10,15,21 | 0,0,0 | 26,22,16 | 0,8,1 | 300 | 150 | 7.96E-05 | 0.001304433 | 1.0,1.0,1.0 | 1.0,0.579,0.889 | 0.177 | ENSG00000078 | [[[[41955234, 41 | | 2 |
| ENSG00000078 | PIK3C3 | chr18 | 42064739 | 42064830 | 42049291 | 42049605 | 42067387 | 42067410 | 14,5,8 | 0,0,0 | 5,11,3 | 0,8,1 | 241 | 150 | 2.21E-05 | 0.0004297 | 1.0,1.0,1.0 | 1.0,0.461,0.651 | 0.296 | ENSG00000078 | [[[[41955234, 41 | | 2 |
| ENSG00000064 | SLC12A2 | chr5 | 128138596 | 128138724 | 128135699 | 128135808 | 128141829 | 128141981 | 19,15,34 | 0,0,0 | 6,29,29 | 3,7,6 | 278 | 150 | 8.60E-09 | 3.87E-07 | 1.0,1.0,1.0 | 0.519,0.691,0.72 | 0.356 | ENSG00000064 | [[[[128083766, 1 | | 2 |
| ENSG00000132 | AP3B1 | chr5 | 78141142 | 78141319 | 78129120 | 78129307 | 78165069 | 78165672 | 37,25,56 | 0,0,0 | 23,61,46 | 4,5,3 | 300 | 150 | 3.71E-06 | 8.89E-05 | 1.0,1.0,1.0 | 0.742,0.859,0.88 | 0.171 | ENSG00000132 | [[[[78294452, 78 | | 3 |
| ENSG00000088 | XRN2 | chr20 | 21333703 | 21333837 | 21333543 | 21333618 | 21339043 | 21339088 | 116,94,184 | 0,0,0 | 58,165,97 | 4,7,5 | 284 | 150 | 1.85E-06 | 4.76E-05 | 1.0,1.0,1.0 | 0.885,0.926,0.91 | 0.093 | ENSG00000088 | [[[[21303331, 21 | | 3 |
| ENSG00000088 | XRN2 | chr20 | 21334077 | 21334185 | 21333543 | 21333618 | 21339043 | 21339088 | 48,56,108 | 0,0,0 | 59,111,68 | 4,7,5 | 258 | 150 | 2.34E-06 | 5.88E-05 | 1.0,1.0,1.0 | 0.896,0.902,0.88 | 0.105 | ENSG00000088 | [[[[21303331, 21 | | 3 |
| ENSG00000164 | WDR41 | chr5 | 77449759 | 77449860 | 77440812 | 77440997 | 77453816 | 77453928 | 52,32,58 | 1,1,1 | 35,64,40 | 15,38,23 | 251 | 150 | 0 | 0 | 0.969,0.95,0.972 | 0.582,0.502,0.51 | 0.432 | ENSG00000164 | [[[[77492170, 77 | | 2 |
| ENSG00000164 | WDR41 | chr5 | 77449759 | 77449870 | 77440812 | 77440997 | 77453816 | 77453928 | 55,34,61 | 1,1,1 | 34,70,44 | 15,38,23 | 261 | 150 | 0 | 0 | 0.969,0.951,0.97 | 0.566,0.514,0.52 | 0.429 | ENSG00000164 | [[[[77492170, 77 | | 2 |
| ENSG00000172 | LAMB2 | chr3 | 49124394 | 49124612 | 49123446 | 49123631 | 49124700 | 49124925 | 20,59,131 | 0,0,0 | 43,83,79 | 4,2,4 | 300 | 150 | 0.000254738 | 0.003580063 | 1.0,1.0,1.0 | 0.843,0.954,0.90 | 0.098 | ENSG00000172 | [[[[49132792, 49 | | 3 |
| ENSG00000152 | POC5 | chr5 | 75694654 | 75694831 | 75692393 | 75692500 | 75705703 | 75705787 | 6,12,12 | 0,0,0 | 29,41,20 | 4,9,3 | 300 | 150 | 7.35E-07 | 2.08E-05 | 1.0,1.0,1.0 | 0.784,0.695,0.76 | 0.251 | ENSG00000152 | [[[[75717306, 75 | | 2 |
| ENSG00000164 | UTP15 | chr5 | 73569496 | 73569675 | 73567261 | 73567434 | 73570585 | 73570711 | 30,19,17 | 0,0,0 | 12,31,19 | 4,7,0 | 300 | 150 | 3.00E-06 | 7.34E-05 | 1.0,1.0,1.0 | 0.6,0.689,1.0 | 0.237 | ENSG00000164 | [[[[73565743, 73 | | 3 |
| ENSG00000134 | CDK7 | chr5 | 69258042 | 69258153 | 69252417 | 69252451 | 69262204 | 69262304 | 4,2,2 | 0,0,0 | 3,13,5 | 5,8,0 | 261 | 150 | 1.46E-06 | 3.86E-05 | 1.0,1.0,1.0 | 0.256,0.483,1.0 | 0.42 | ENSG00000134 | [[[[69234886, 69 | | 4 |
| ENSG00000137 | TMEM14B | chr6 | 10755141 | 10755232 | 10747758 | 10747881 | 10770076 | 10770204 | 1,1,0 | 3,7,3 | 4,3,4 | 1,1,8 | 241 | 150 | 7.55E-05 | 0.001247272 | 0.172,0.082,0.0 | 0.713,0.651,0.23 | -0.449 | ENSG00000137 | [[[[10747805, 10 | | 5 |
| ENSG00000164 | MOCS2 | chr5 | 53107076 | 53107221 | 53100410 | 53100534 | 53108521 | 53108643 | 90,64,115 | 0,0,0 | 60,112,81 | 9,3,10 | 295 | 150 | 7.39E-09 | 3.39E-07 | 1.0,1.0,1.0 | 0.772,0.95,0.805 | 0.158 | ENSG00000164 | [[[[53109251, 53 | | 3 |
| ENSG00000112 | HMGCS1 | chr5 | 43297001 | 43297166 | 43292847 | 43292973 | 43298008 | 43298134 | 64,71,138 | 0,0,0 | 48,101,45 | 1,6,4 | 300 | 150 | 3.77E-05 | 0.000684478 | 1.0,1.0,1.0 | 0.96,0.894,0.849 | 0.099 | ENSG00000112 | [[[[43313356, 43 | | 4 |
| ENSG00000168 | ADAM9 | chr8 | 39082967 | 39083073 | 39082640 | 39082721 | 39091258 | 39091346 | 298,214,388 | 0,0,0 | 193,433,240 | 4,15,6 | 256 | 150 | 1.42E-06 | 3.75E-05 | 1.0,1.0,1.0 | 0.966,0.944,0.95 | 0.044 | ENSG00000168 | [[[[38996973, 38 | | 2 |
| ENSG00000168 | ADAM9 | chr8 | 39090046 | 39090188 | 39082640 | 39082721 | 39091258 | 39091346 | 295,281,406 | 0,0,0 | 232,535,308 | 4,15,6 | 292 | 150 | 4.68E-06 | 0.000108801 | 1.0,1.0,1.0 | 0.968,0.948,0.96 | 0.04 | ENSG00000168 | [[[[38996973, 38 | | 2 |
| ENSG00000146 | CCT6A | chr7 | 56058361 | 56058521 | 56057992 | 56058103 | 56062682 | 56062755 | 571,469,658 | 0,0,0 | 475,896,606 | 11,14,11 | 300 | 150 | 6.04E-07 | 1.75E-05 | 1.0,1.0,1.0 | 0.956,0.97,0.965 | 0.036 | ENSG00000146 | [[[[56051765, 56 | | 6 |
| ENSG00000146 | CCT6A | chr7 | 56058619 | 56058702 | 56057992 | 56058103 | 56059543 | 56059640 | 485,499,687 | 0,0,4 | 338,743,454 | 184,351,162 | 233 | 150 | 0 | 0 | 1.0,1.0,0.991 | 0.542,0.577,0.64 | 0.41 | ENSG00000146 | [[[[56051765, 56 | | 2 |
| ENSG00000146 | CCT6A | chr7 | 56060268 | 56060416 | 56059543 | 56059640 | 56061746 | 56061849 | 7,197,081,016 | 0,0,0 | 5,521,151,671 | 87,168,74 | 298 | 150 | 0 | 0 | 1.0,1.0,1.0 | 0.762,0.775,0.82 | 0.214 | ENSG00000146 | [[[[56051765, 56 | | 2 |
| ENSG00000146 | CCT6A | chr7 | 56060806 | 56060940 | 56057992 | 56058103 | 56062682 | 56062755 | 4,2,1 | 0,0,0 | 12,19,21 | 11,14,11 | 284 | 150 | 2.68E-07 | 8.47E-06 | 1.0,1.0,1.0 | 0.366,0.418,0.50 | 0.571 | ENSG00000146 | [[[[56051765, 56 | | 6 |
| ENSG00000146 | CCT6A | chr7 | 56060806 | 56060940 | 56058619 | 56058702 | 56062682 | 56062755 | 4,2,1 | 0,0,0 | 18,20,21 | 8,7,7 | 284 | 150 | 6.25E-07 | 1.81E-05 | 1.0,1.0,1.0 | 0.543,0.601,0.61 | 0.414 | ENSG00000146 | [[[[56051765, 56 | | 4 |
| ENSG00000146 | CCT6A | chr7 | 56060806 | 56060940 | 56059543 | 56059640 | 56061746 | 56061849 | 8,528,381,033 | 0,0,0 | 8,181,454,893 | 87,168,74 | 284 | 150 | 0 | 0 | 1.0,1.0,1.0 | 0.832,0.821,0.86 | 0.161 | ENSG00000146 | [[[[56051765, 56 | | 2 |
| ENSG00000146 | CCT6A | chr7 | 56060806 | 56060940 | 56059543 | 56059640 | 56062682 | 56062755 | 4,2,1 | 0,0,0 | 12,17,20 | 26,37,28 | 284 | 150 | 1.72E-08 | 7.27E-07 | 1.0,1.0,1.0 | 0.196,0.195,0.27 | 0.778 | ENSG00000146 | [[[[56051765, 56 | | 3 |
| ENSG00000146 | CCT6A | chr7 | 56061746 | 56061849 | 56057992 | 56058103 | 56062682 | 56062755 | 597,490,715 | 0,0,0 | 5,871,115,629 | 11,14,11 | 253 | 150 | 6.95E-05 | 0.001161218 | 1.0,1.0,1.0 | 0.969,0.979,0.97 | 0.027 | ENSG00000146 | [[[[56051765, 56 | | 6 |
| ENSG00000146 | CCT6A | chr7 | 56061746 | 56061849 | 56059543 | 56059640 | 56062682 | 56062755 | 597,490,715 | 0,0,0 | 6,611,264,686 | 26,37,28 | 253 | 150 | 1.11E-16 | 1.65E-14 | 1.0,1.0,1.0 | 0.938,0.953,0.93 | 0.058 | ENSG00000146 | [[[[56051765, 56 | | 3 |
| ENSG00000181 | TMEM45A | chr3 | 100519461 | 100519608 | 100492650 | 100492928 | 100558404 | 100558589 | 84,77,93 | 0,0,1 | 76,151,87 | 3,8,1 | 297 | 150 | 0.000564722 | 0.007139648 | 1.0,1.0,0.979 | 0.928,0.905,0.97 | 0.056 | ENSG00000181 | [[[[100492619, 1 | | 2 |
| ENSG00000181 | TMEM45A | chr3 | 100519532 | 100519608 | 100492650 | 100492928 | 100558404 | 100558527 | 3,7,9 | 0,0,1 | 7,15,7 | 3,8,1 | 226 | 150 | 1.68E-05 | 0.000337055 | 1.0,1.0,0.857 | 0.608,0.554,0.82 | 0.291 | ENSG00000181 | [[[[100492619, 1 | | 2 |
| ENSG00000145 | SKP2 | chr5 | 36176964 | 36177016 | 36171602 | 36171733 | 36183839 | 36184029 | 24,11,22 | 0,0,0 | 18,37,26 | 2,9,2 | 202 | 150 | 3.10E-06 | 7.58E-05 | 1.0,1.0,1.0 | 0.87,0.753,0.906 | 0.157 | ENSG00000145 | [[[[36152111, 36 | | 2 |
| ENSG00000167 | SPRYD3 | chr12 | 53073285 | 53073471 | 53068154 | 53068304 | 53075094 | 53075219 | 21,17,58 | 0,0,0 | 28,62,46 | 3,2,5 | 300 | 150 | 4.22E-05 | 0.000754111 | 1.0,1.0,1.0 | 0.824,0.939,0.82 | 0.139 | ENSG00000167 | [[[[53079311, 53 | | 2 |
| ENSG00000167 | SPRYD3 | chr12 | 53074648 | 53074784 | 53068154 | 53068304 | 53075094 | 53075219 | 36,35,31 | 0,0,0 | 42,65,47 | 3,2,5 | 286 | 150 | 0.000132637 | 0.002038016 | 1.0,1.0,1.0 | 0.88,0.945,0.831 | 0.115 | ENSG00000167 | [[[[53079311, 53 | | 2 |
| ENSG00000091 | DLD | chr7 | 107906266 | 107906368 | 107904957 | 107905058 | 107915505 | 107915696 | 74,47,116 | 0,0,0 | 23,53,26 | 1,4,5 | 252 | 150 | 9.68E-06 | 0.000207922 | 1.0,1.0,1.0 | 0.932,0.887,0.75 | 0.142 | ENSG00000091 | [[[[107891183, 1 | | 2 |
| ENSG00000056 | ZFR | chr5 | 32406773 | 32407021 | 32403905 | 32404097 | 32444228 | 32444328 | 31,23,61 | 0,0,0 | 39,98,81 | 3,4,3 | 300 | 150 | 0.00061897 | 0.00771423 | 1.0,1.0,1.0 | 0.867,0.925,0.93 | 0.092 | ENSG00000056 | [[[[32444622, 32 | | 4 |
| ENSG00000056 | ZFR | chr5 | 32414968 | 32415187 | 32406726 | 32407021 | 32444228 | 32444328 | 35,30,69 | 0,0,1 | 56,108,96 | 2,6,12 | 300 | 150 | 8.15E-06 | 0.000178695 | 1.0,1.0,0.972 | 0.933,0.9,0.8 | 0.113 | ENSG00000056 | [[[[32444622, 32 | | 3 |
| ENSG00000056 | ZFR | chr5 | 32419820 | 32420103 | 32403905 | 32404097 | 32444228 | 32444328 | 15,23,53 | 0,0,0 | 27,55,38 | 3,4,3 | 300 | 150 | 4.85E-05 | 0.000849913 | 1.0,1.0,1.0 | 0.818,0.873,0.86 | 0.148 | ENSG00000056 | [[[[32444622, 32 | | 4 |
| ENSG00000182 | CADM1 | chr11 | 115190887 | 115190941 | 115178643 | 115178775 | 115209573 | 115209657 | 0,0,1 | 85,56,125 | 2,13,2 | 53,113,77 | 204 | 150 | 0.000130269 | 0.002006106 | 0.0,0.0,0.006 | 0.027,0.078,0.01 | -0.039 | ENSG00000182 | [[[[115504271, 11 | | 2 |
| ENSG00000112 | HBS1L | chr6 | 134982457 | 134982562 | 134979177 | 134979268 | 134986735 | 134986810 | 42,19,49 | 0,0,0 | 41,60,40 | 1,12,3 | 255 | 150 | 1.53E-06 | 4.01E-05 | 1.0,1.0,1.0 | 0.96,0.746,0.887 | 0.136 | ENSG00000112 | [[[[135054649, 1 | | 3 |
| ENSG00000160 | LSS | chr21 | 46195675 | 46195756 | 46191880 | 46191959 | 46196201 | 46196267 | 23,54,92 | 0,0,0 | 23,48,26 | 9,9,5 | 231 | 150 | 5.76E-12 | 4.66E-10 | 1.0,1.0,1.0 | 0.624,0.776,0.77 | 0.276 | ENSG00000160 | [[[[46228732, 46 | | 2 |
| ENSG00000160 | LSS | chr21 | 46195675 | 46195756 | 46191880 | 46191959 | 46205835 | 46205941 | 2,6,5 | 0,0,0 | 1,2,0 | 13,23,16 | 231 | 150 | 0 | 0 | 1.0,1.0,1.0 | 0.048,0.053,0.0 | 0.966 | ENSG00000160 | [[[[46228732, 46 | | 3 |
| ENSG00000160 | LSS | chr21 | 46196201 | 46196267 | 46191880 | 46191959 | 46205835 | 46205941 | 38,66,86 | 0,0,0 | 41,42,30 | 13,23,16 | 216 | 150 | 0 | 0 | 1.0,1.0,1.0 | 0.687,0.559,0.56 | 0.396 | ENSG00000160 | [[[[46228732, 46 | | 3 |
| ENSG00000163 | RYK | chr3 | 134175929 | 134176039 | 134175608 | 134175768 | 134183001 | 134183071 | 62,59,92 | 0,0,0 | 31,83,44 | 4,19,7 | 260 | 150 | 3.03E-13 | 2.95E-11 | 1.0,1.0,1.0 | 0.817,0.716,0.78 | 0.228 | ENSG00000163 | [[[[134250423, 1 | | 2 |
| ENSG00000163 | RYK | chr3 | 134202729 | 134202874 | 134195081 | 134195182 | 134209694 | 134209829 | 41,52,100 | 0,0,0 | 27,53,23 | 4,5,6 | 295 | 150 | 5.87E-09 | 2.75E-07 | 1.0,1.0,1.0 | 0.774,0.844,0.66 | 0.24 | ENSG00000163 | [[[[134250423, 1 | | 2 |
| ENSG00000107 | FUBP3 | chr9 | 130623610 | 130623711 | 130620353 | 130620458 | 130626363 | 130626505 | 26,48,120 | 0,0,0 | 35,147,98 | 5,7,4 | 251 | 150 | 1.10E-05 | 0.000232314 | 1.0,1.0,1.0 | 0.807,0.926,0.93 | 0.11 | ENSG00000107 | [[[[130579577, 1 | | 2 |

| | | | | | | | | | | | | | | | | | | | | | | |
|---|---|---|---|---|---|---|---|---|---|---|---|---|---|---|---|---|---|---|---|---|---|---|
| ENSG00000204 | PRRC2A | chr6 | 31626019 | 31626162 | 31625459 | 31625611 | 31626771 | 31626862 | 74,61,113 | 0,0,0 | 55,141,94 | 1,5,6 | 293 | 150 | 0.00015517 | 0.002333006 | 1.0,1.0,1.0 | 0.966,0.935,0.88 | 0.07 | ENSG00000204 | [[[[31620715, 316 | 2 |
| ENSG00000014 | UFL1 | chr6 | 96524381 | 96524410 | 96523145 | 96523291 | 96528501 | 96528632 | 1,3,2 | 0,0,0 | 7,10,8 | 2,2,2 | 179 | 150 | 0.000541526 | 0.006887527 | 1.0,1.0,1.0 | 0.746,0.807,0.77 | 0.226 | ENSG00000014 | [[[[96521806, 96 | 3 |
| ENSG00000014 | UFL1 | chr6 | 96526320 | 96526435 | 96524381 | 96524410 | 96528501 | 96528632 | 16,13,27 | 0,0,0 | 6,19,16 | 7,8,5 | 265 | 150 | 1.14E-10 | 7.40E-09 | 1.0,1.0,1.0 | 0.327,0.573,0.64 | 0.485 | ENSG00000014 | [[[[96521806, 96 | 2 |
| ENSG00000145 | MAR6 | chr5 | 10397292 | 10397344 | 10394752 | 10394785 | 10402058 | 10402139 | 22,8,31 | 0,0,0 | 26,27,17 | 4,18,13 | 202 | 150 | 1.41E-10 | 9.02E-09 | 1.0,1.0,1.0 | 0.828,0.527,0.49 | 0.384 | ENSG00000145 | [[[[10353695, 10 | 2 |
| ENSG00000145 | MAR6 | chr5 | 10400783 | 10400842 | 10394752 | 10394785 | 10402058 | 10402139 | 10,8,30 | 0,0,0 | 6,15,13 | 4,18,13 | 209 | 150 | 5.00E-15 | 6.22E-13 | 1.0,1.0,1.0 | 0.518,0.374,0.41 | 0.563 | ENSG00000145 | [[[[10353695, 10 | 2 |
| ENSG00000145 | MAR6 | chr5 | 10423734 | 10423824 | 10417269 | 10417404 | 10429892 | 10430028 | 25,36,49 | 1,0,0 | 34,73,35 | 7,19,10 | 240 | 150 | 4.24E-11 | 2.94E-09 | 0.94,1.0,1.0 | 0.752,0.706,0.68 | 0.265 | ENSG00000145 | [[[[10353695, 10 | 2 |
| ENSG00000164 | DCBLD1 | chr6 | 117540917 | 117541025 | 117538619 | 117540815 | 117544527 | 117544577 | 55,47,91 | 0,0,0 | 23,43,48 | 3,15,3 | 258 | 150 | 1.25E-09 | 6.78E-08 | 1.0,1.0,1.0 | 0.817,0.625,0.90 | 0.218 | ENSG00000164 | [[[[117482674, 11 | 2 |
| ENSG00000164 | DCBLD1 | chr6 | 117543123 | 117543211 | 117538619 | 117540815 | 117544527 | 117544577 | 16,17,20 | 0,0,0 | 15,48,27 | 3,15,3 | 238 | 150 | 5.86E-08 | 2.18E-06 | 1.0,1.0,1.0 | 0.759,0.669,0.85 | 0.241 | ENSG00000164 | [[[[117482674, 11 | 2 |
| ENSG00000110 | KMT5B | chr11 | 68166981 | 68167178 | 68154862 | 68159171 | 68173803 | 68173913 | 10,6,8 | 0,0,0 | 8,19,12 | 5,2,2 | 300 | 150 | 1.10E-06 | 2.99E-05 | 1.0,1.0,1.0 | 0.444,0.826,0.75 | 0.327 | ENSG00000110 | [[[[68213138, 68 | 4 |
| ENSG00000131 | ANKHD1 | chr5 | 140507784 | 140507998 | 140506834 | 140506977 | 140510018 | 140510181 | 27,8,31 | 0,0,0 | 25,50,40 | 1,5,2 | 300 | 150 | 0.000629937 | 0.007827779 | 1.0,1.0,1.0 | 0.926,0.833,0.90 | 0.111 | ENSG00000131 | [[[[140401833, 1 | 2 |
| ENSG00000152 | NADK2 | chr5 | 36217747 | 36217884 | 36211843 | 36211922 | 36225541 | 36225612 | 19,25,36 | 0,0,0 | 14,75,46 | 8,4,1 | 287 | 150 | 2.33E-05 | 0.000449631 | 1.0,1.0,1.0 | 0.478,0.907,0.96 | 0.218 | ENSG00000152 | [[[[36241499, 36 | 2 |
| ENSG00000109 | UFSP2 | chr4 | 185403493 | 185403618 | 185399539 | 185400478 | 185407935 | 185408060 | 24,22,20 | 0,0,0 | 7,15,12 | 1,12,8 | 275 | 150 | 1.40E-10 | 8.99E-09 | 1.0,1.0,1.0 | 0.792,0.405,0.45 | 0.451 | ENSG00000109 | [[[[185425866, 1 | 2 |
| ENSG00000109 | UFSP2 | chr4 | 185415154 | 185415347 | 185413725 | 185413872 | 185418440 | 185418507 | 19,17,24 | 0,0,0 | 17,11,15 | 2,2,3 | 300 | 150 | 5.39E-05 | 0.000932903 | 1.0,1.0,1.0 | 0.81,0.733,0.714 | 0.248 | ENSG00000109 | [[[[185425866, 1 | 2 |
| ENSG00000125 | TM9SF2 | chr13 | 99539445 | 99539557 | 99537738 | 99537863 | 99543862 | 99543995 | 101,82,145 | 0,0,0 | 76,169,98 | 5,9,5 | 262 | 150 | 6.69E-07 | 1.92E-05 | 1.0,1.0,1.0 | 0.897,0.915,0.91 | 0.09 | ENSG00000125 | [[[[99501472, 99 | 3 |
| ENSG00000125 | TM9SF2 | chr13 | 99539445 | 99539557 | 99537738 | 99537863 | 99546984 | 99547104 | 101,82,145 | 0,0,0 | 76,169,98 | 4,11,0 | 262 | 150 | 2.50E-05 | 0.000478073 | 1.0,1.0,1.0 | 0.916,0.898,1.0 | 0.062 | ENSG00000125 | [[[[99501472, 99 | 4 |
| ENSG00000125 | TM9SF2 | chr13 | 99543862 | 99543995 | 99537738 | 99537863 | 99546984 | 99547104 | 137,130,164 | 0,0,0 | 127,291,191 | 4,11,0 | 283 | 150 | 0.000652075 | 0.008051687 | 1.0,1.0,1.0 | 0.944,0.933,1.0 | 0.041 | ENSG00000125 | [[[[99501472, 99 | 4 |
| ENSG00000125 | TM9SF2 | chr13 | 99549164 | 99549222 | 99546984 | 99547104 | 99554303 | 99554455 | 46,42,59 | 0,0,0 | 33,96,60 | 6,14,8 | 208 | 150 | 8.00E-10 | 4.52E-08 | 1.0,1.0,1.0 | 0.799,0.832,0.84 | 0.175 | ENSG00000125 | [[[[99501472, 99 | 2 |
| ENSG00000109 | AADAT | chr4 | 170064718 | 170064825 | 170061891 | 170061993 | 170070586 | 170070652 | 12,14,16 | 0,0,0 | 9,13,6 | 3,5,4 | 257 | 150 | 2.66E-09 | 1.36E-07 | 1.0,1.0,1.0 | 0.636,0.603,0.46 | 0.431 | ENSG00000109 | [[[[170089624, 1 | 5 |
| ENSG00000109 | AADAT | chr4 | 170069147 | 170069230 | 170061891 | 170061993 | 170070586 | 170070652 | 12,14,13 | 0,0,0 | 9,32,18 | 3,5,4 | 233 | 150 | 9.38E-07 | 2.60E-05 | 1.0,1.0,1.0 | 0.659,0.805,0.74 | 0.264 | ENSG00000109 | [[[[170089624, 1 | 5 |
| ENSG00000137 | NEK1 | chr4 | 169599099 | 169599197 | 169590725 | 169590809 | 169612019 | 169612143 | 3,1,3 | 0,0,0 | 1,2,3 | 3,3,3 | 248 | 150 | 2.89E-11 | 2.05E-09 | 1.0,1.0,1.0 | 0.168,0.287,0.37 | 0.723 | ENSG00000137 | [[[[169612222, 1 | 3 |
| ENSG00000162 | CLHC1 | chr2 | 55180509 | 55180712 | 55177601 | 55177781 | 55206269 | 55206376 | 11,10,8 | 0,0,0 | 2,6,8 | 1,4,2 | 300 | 150 | 5.32E-07 | 1.57E-05 | 1.0,1.0,1.0 | 0.5,0.429,0.667 | 0.468 | ENSG00000162 | [[[[55232223, 55 | 2 |
| ENSG00000162 | CLHC1 | chr2 | 55181569 | 55181744 | 55177601 | 55177781 | 55206269 | 55206376 | 1,7,1 | 0,0,0 | 2,4,2 | 1,4,2 | 300 | 150 | 1.09E-08 | 4.83E-07 | 1.0,1.0,1.0 | 0.5,0.333,0.333 | 0.611 | ENSG00000162 | [[[[55232223, 55 | 2 |
| ENSG00000162 | CLHC1 | chr2 | 55208625 | 55208710 | 55206269 | 55206376 | 55209629 | 55209831 | 0,0,1 | 4,1,2 | 0,6,5 | 1,4,0 | 235 | 150 | 0.000141105 | 0.002145238 | 0.0,0.0,0.242 | 0.0,0.489,1.0 | -0.416 | ENSG00000162 | [[[[55232223, 55 | 2 |
| ENSG00000162 | CLHC1 | chr2 | 55209403 | 55209516 | 55206269 | 55206376 | 55209629 | 55209652 | 0,0,2 | 4,1,2 | 2,6,3 | 1,4,0 | 263 | 150 | 0.00010393 | 0.00164984 | 0.0,0.0,0.363 | 0.533,0.461,1.0 | -0.544 | ENSG00000162 | [[[[55232223, 55 | 2 |
| ENSG00000138 | DNA2 | chr10 | 68442916 | 68443111 | 68432393 | 68432510 | 68444920 | 68445083 | 14,14,14 | 0,0,0 | 9,28,12 | 3,4,1 | 300 | 150 | 8.83E-06 | 0.00019162 | 1.0,1.0,1.0 | 0.6,0.778,0.857 | 0.255 | ENSG00000138 | [[[[68471791, 68 | 2 |
| ENSG00000171 | PLRG1 | chr4 | 154540593 | 154540695 | 154539950 | 154540053 | 154542186 | 154542279 | 79,49,77 | 0,0,0 | 42,112,56 | 2,8,4 | 252 | 150 | 8.32E-06 | 0.000181656 | 1.0,1.0,1.0 | 0.926,0.893,0.89 | 0.096 | ENSG00000171 | [[[[154550300, 1 | 2 |
| ENSG00000171 | PLRG1 | chr4 | 154540784 | 154540934 | 154539950 | 154540053 | 154542186 | 154542279 | 103,96,146 | 0,0,0 | 69,156,111 | 2,8,4 | 300 | 150 | 2.56E-05 | 0.000487736 | 1.0,1.0,1.0 | 0.945,0.907,0.93 | 0.072 | ENSG00000171 | [[[[154550300, 1 | 2 |
| ENSG00000112 | ERBIN | chr5 | 66048666 | 66048781 | 66038382 | 66038482 | 66050782 | 66050966 | 33,15,56 | 0,0,0 | 13,43,30 | 1,9,9 | 265 | 150 | 3.77E-09 | 1.85E-07 | 1.0,1.0,1.0 | 0.88,0.73,0.654 | 0.245 | ENSG00000112 | [[[[65926575, 65 | 4 |
| ENSG00000112 | ERBIN | chr5 | 66075023 | 66075230 | 66053405 | 66054951 | 66076315 | 66076408 | 77,60,72 | 33,28,42 | 16,44,29 | 23,61,51 | 300 | 150 | 7.95E-09 | 3.61E-07 | 0.538,0.517,0.46 | 0.258,0.265,0.22 | 0.258 | ENSG00000112 | [[[[65926575, 65 | 2 |
| ENSG00000089 | GCN1 | chr12 | 120144638 | 120144835 | 120144305 | 120144448 | 120145261 | 120145330 | 54,70,143 | 0,0,0 | 58,83,62 | 14,36,21 | 300 | 150 | 0 | 0 | 1.0,1.0,1.0 | 0.674,0.535,0.59 | 0.398 | ENSG00000089 | [[[[120194680, 1 | 2 |
| ENSG00000089 | GCN1 | chr12 | 120144922 | 120145061 | 120144305 | 120144448 | 120145261 | 120145330 | 15,39,99 | 0,0,0 | 29,71,60 | 14,36,21 | 289 | 150 | 0 | 0 | 1.0,1.0,1.0 | 0.518,0.506,0.59 | 0.46 | ENSG00000089 | [[[[120194680, 1 | 2 |
| ENSG00000118 | ELL2 | chr5 | 95913770 | 95913934 | 95906522 | 95906782 | 95943001 | 95943049 | 44,52,58 | 0,3,2 | 41,122,80 | 10,30,12 | 300 | 150 | 1.42E-07 | 4.78E-06 | 1.0,0.897,0.935 | 0.672,0.67,0.769 | 0.24 | ENSG00000118 | [[[[95961575, 95 | 2 |
| ENSG00000118 | ELL2 | chr5 | 95919423 | 95919545 | 95906522 | 95906782 | 95943001 | 95943013 | 27,14,48 | 0,3,2 | 19,53,50 | 10,30,12 | 272 | 150 | 5.60E-06 | 0.000128264 | 1.0,0.72,0.93 | 0.512,0.493,0.69 | 0.316 | ENSG00000118 | [[[[95961575, 95 | 2 |
| ENSG00000164 | ABCE1 | chr4 | 145124989 | 145125101 | 145123214 | 145123357 | 145127525 | 145127573 | 69,58,74 | 0,0,0 | 75,156,64 | 3,7,5 | 262 | 150 | 3.07E-05 | 0.000571852 | 1.0,1.0,1.0 | 0.935,0.927,0.88 | 0.086 | ENSG00000164 | [[[[145098311, 1 | 2 |
| ENSG00000091 | SEL1L3 | chr4 | 25830097 | 25830156 | 25821995 | 25822128 | 25835196 | 25835323 | 41,30,60 | 0,0,0 | 14,34,29 | 2,11,4 | 209 | 150 | 2.69E-09 | 1.37E-07 | 1.0,1.0,1.0 | 0.834,0.689,0.83 | 0.213 | ENSG00000091 | [[[[25862675, 25 | 3 |
| ENSG00000101 | NDUFAF5 | chr20 | 13794837 | 13794941 | 13793159 | 13793227 | 13801507 | 13801683 | 7,6,11 | 0,0,0 | 50,60,33 | 6,5,0 | 254 | 150 | 0.000216881 | 0.003126665 | 1.0,1.0,1.0 | 0.831,0.876,1.0 | 0.098 | ENSG00000101 | [[[[13785028, 13 | 2 |
| ENSG00000101 | NDUFAF5 | chr20 | 13794837 | 13794941 | 13793159 | 13793227 | 13808841 | 13808865 | 7,6,11 | 0,0,0 | 50,61,33 | 4,5,4 | 254 | 150 | 6.42E-05 | 0.001086243 | 1.0,1.0,1.0 | 0.881,0.878,0.83 | 0.137 | ENSG00000101 | [[[[13785028, 13 | 3 |
| ENSG00000101 | NDUFAF5 | chr20 | 13801485 | 13801683 | 13793159 | 13793227 | 13808841 | 13808902 | 15,21,45 | 0,0,0 | 54,134,91 | 4,5,4 | 300 | 150 | 0.000244604 | 0.003454746 | 1.0,1.0,1.0 | 0.871,0.931,0.91 | 0.093 | ENSG00000101 | [[[[13785028, 13 | 3 |
| ENSG00000101 | NDUFAF5 | chr20 | 13801507 | 13801683 | 13793179 | 13793227 | 13808841 | 13808865 | 15,21,45 | 0,0,0 | 60,138,91 | 4,5,4 | 300 | 150 | 0.00029271 | 0.004043493 | 1.0,1.0,1.0 | 0.882,0.932,0.91 | 0.089 | ENSG00000101 | [[[[13785028, 13 | 3 |
| ENSG00000154 | PDE1C | chr7 | 31837179 | 31837300 | 31828291 | 31828373 | 31847967 | 31848096 | 85,42,86 | 0,0,1 | 30,74,44 | 2,5,2 | 271 | 150 | 0.000490316 | 0.006325962 | 1.0,1.0,0.979 | 0.893,0.891,0.92 | 0.09 | ENSG00000154 | [[[[32070293, 32 | 2 |
| ENSG00000164 | MFSD8 | chr4 | 127930682 | 127930817 | 127921523 | 127921963 | 127938782 | 127938836 | 9,3,13 | 0,0,0 | 7,19,10 | 4,2,1 | 285 | 150 | 4.79E-05 | 0.000840513 | 1.0,1.0,1.0 | 0.479,0.833,0.84 | 0.283 | ENSG00000164 | [[[[127965072, 1 | 2 |
| ENSG00000145 | METTL14 | chr4 | 118694435 | 118694526 | 118691980 | 118692068 | 118700549 | 118700642 | 19,8,10 | 0,0,0 | 5,20,16 | 3,1,3 | 241 | 150 | 0.000159844 | 0.002394299 | 1.0,1.0,1.0 | 0.509,0.926,0.76 | 0.266 | ENSG00000145 | [[[[118685392, 11 | 2 |
| ENSG00000213 | NAXD | chr13 | 110624233 | 110624279 | 110622215 | 110622366 | 110627438 | 110627547 | 10,10,21 | 0,0,0 | 15,43,22 | 2,6,1 | 196 | 150 | 0.000222072 | 0.003191518 | 1.0,1.0,1.0 | 0.852,0.846,0.94 | 0.119 | ENSG00000213 | [[[[110615545, 11 | 2 |
| ENSG00000164 | PRSS12 | chr4 | 118316181 | 118316323 | 118312945 | 118313397 | 118331715 | 118331866 | 123,56,139 | 1,0,0 | 37,67,50 | 12,43,46 | 292 | 150 | 0 | 0 | 0.984,1.0,1.0 | 0.613,0.445,0.35 | 0.523 | ENSG00000164 | [[[[118352219, 11 | 2 |
| ENSG00000138 | NDST4 | chr4 | 114977186 | 114977274 | 114970429 | 114970584 | 115113443 | 115113849 | 3,1,1 | 0,0,1 | 6,12,2 | 0,6,3 | 238 | 150 | 0.000690905 | 0.008473453 | 1.0,1.0,0.387 | 1.0,0.558,0.296 | 0.178 | ENSG00000138 | [[[[115113444, 11 | 2 |
| ENSG00000138 | NDST4 | chr4 | 115076058 | 115077282 | 114970429 | 114970584 | 115113443 | 115113876 | 2,3,3 | 0,0,1 | 11,5,7 | 0,6,3 | 300 | 150 | 0.00026321 | 0.003682715 | 1.0,1.0,0.6 | 1.0,0.294,0.538 | 0.256 | ENSG00000138 | [[[[115113444, 11 | 2 |
| ENSG00000145 | CAMK2D | chr4 | 113500462 | 113500508 | 113465528 | 113465604 | 113509637 | 113509672 | 11,8,14 | 0,0,0 | 7,12,7 | 0,5,5 | 196 | 150 | 2.82E-06 | 6.95E-05 | 1.0,1.0,1.0 | 1.0,0.647,0.517 | 0.279 | ENSG00000145 | [[[[113761004, 11 | 3 |
| ENSG00000133 | ATP13A3 | chr3 | 194413758 | 194413839 | 194402671 | 194406116 | 194425341 | 194425529 | 4,3,9 | 0,0,1 | 4,16,4 | 2,12,1 | 231 | 150 | 3.94E-06 | 9.36E-05 | 1.0,1.0,0.854 | 0.565,0.464,0.72 | 0.368 | ENSG00000133 | [[[[194486566, 1 | 3 |
| ENSG00000133 | ATP13A3 | chr3 | 194419878 | 194419967 | 194402671 | 194406116 | 194425341 | 194425529 | 18,14,34 | 0,0,1 | 20,28,25 | 2,12,1 | 239 | 150 | 1.38E-05 | 0.000282913 | 1.0,1.0,0.955 | 0.863,0.594,0.94 | 0.186 | ENSG00000133 | [[[[194486566, 1 | 3 |
| ENSG00000133 | ATP13A3 | chr3 | 194428844 | 194428917 | 194425341 | 194425529 | 194429677 | 194429774 | 25,22,34 | 0,0,0 | 13,40,33 | 12,14,5 | 223 | 150 | 2.47E-10 | 1.53E-08 | 1.0,1.0,1.0 | 0.422,0.658,0.81 | 0.368 | ENSG00000133 | [[[[194486566, 1 | 2 |
| ENSG00000133 | ATP13A3 | chr3 | 194459470 | 194459541 | 194457093 | 194457174 | 194462139 | 194462212 | 7,2,12 | 0,0,0 | 5,6,12 | 2,4,1 | 221 | 150 | 2.26E-05 | 0.000438994 | 1.0,1.0,1.0 | 0.629,0.504,0.89 | 0.325 | ENSG00000133 | [[[[194486566, 1 | 3 |
| ENSG00000177 | TALDO1 | chr11 | 763343 | 763519 | 760121 | 760253 | 764287 | 764433 | 378,401,752 | 0,0,0 | 488,915,591 | 4,23,9 | 300 | 150 | 1.60E-06 | 4.19E-05 | 1.0,1.0,1.0 | 0.984,0.952,0.97 | 0.031 | ENSG00000177 | [[[[747464, 7475 | 2 |
| ENSG00000177 | TALDO1 | chr11 | 763746 | 763940 | 760121 | 760253 | 764287 | 764433 | 3,6,11 | 0,0,0 | 5,20,4 | 4,23,9 | 300 | 150 | 1.98E-11 | 1.44E-09 | 1.0,1.0,1.0 | 0.385,0.303,0.18 | 0.71 | ENSG00000177 | [[[[747464, 7475 | 2 |
| ENSG00000177 | TALDO1 | chr11 | 763746 | 763944 | 760121 | 760253 | 764287 | 764433 | 228,432,545 | 0,0,0 | 327,705,399 | 4,23,9 | 300 | 150 | 3.03E-08 | 1.21E-06 | 1.0,1.0,1.0 | 0.976,0.939,0.95 | 0.043 | ENSG00000177 | [[[[747464, 7475 | 2 |
| ENSG00000138 | ZGRF1 | chr4 | 112547284 | 112547408 | 112541091 | 112541268 | 112553834 | 112553982 | 17,9,16 | 0,0,0 | 4,25,8 | 3,6,1 | 274 | 150 | 5.43E-07 | 1.60E-05 | 1.0,1.0,1.0 | 0.422,0.695,0.81 | 0.356 | ENSG00000138 | [[[[112636851, 11 | 2 |
| ENSG00000136 | DRAM1 | chr12 | 101901290 | 101901433 | 101877350 | 101877920 | 101908185 | 101908327 | 17,14,48 | 0,0,0 | 20,41,26 | 5,5,4 | 293 | 150 | 1.40E-07 | 4.73E-06 | 1.0,1.0,1.0 | 0.672,0.808,0.76 | 0.25 | ENSG00000136 | [[[[101877580, 1 | 2 |
| ENSG00000136 | DRAM1 | chr12 | 101901290 | 101901433 | 101877350 | 101877920 | 101920108 | 101920201 | 4,8,5 | 0,0,0 | 0,6,4 | 2,4,1 | 293 | 150 | 2.98E-07 | 9.34E-06 | 1.0,1.0,1.0 | 0.0,0.434,0.672 | 0.631 | ENSG00000136 | [[[[101877580, 1 | 4 |
| ENSG00000147 | HAUS6 | chr9 | 19082872 | 19083043 | 19076601 | 19076704 | 19086733 | 19086782 | 33,25,60 | 0,0,0 | 40,99,60 | 11,8,2 | 300 | 150 | 3.82E-07 | 1.16E-05 | 1.0,1.0,1.0 | 0.645,0.861,0.93 | 0.185 | ENSG00000147 | [[[[19102524, 19 | 3 |
| ENSG00000074 | LMAN1 | chr18 | 59338759 | 59338953 | 59338556 | 59338627 | 59347512 | 59347571 | 46,38,97 | 0,0,0 | 39,156,88 | 3,16,1 | 300 | 150 | 1.06E-06 | 2.90E-05 | 1.0,1.0,1.0 | 0.867,0.83,0.978 | 0.108 | ENSG00000074 | [[[[59359031, 59 | 2 |
| ENSG00000044 | EPHA3 | chr3 | 89449224 | 89449374 | 89431149 | 89431359 | 89472463 | 89472522 | 70,44,122 | 0,0,0 | 47,141,89 | 0,10,3 | 300 | 150 | 4.36E-05 | 0.000775821 | 1.0,1.0,1.0 | 1.0,0.876,0.937 | 0.062 | ENSG00000044 | [[[[89107621, 89 | 2 |
| ENSG00000197 | ERO1A | chr14 | 52682324 | 52682408 | 52678318 | 52678472 | 52695367 | 52695529 | 50,36,94 | 0,0,0 | 43,86,57 | 8,47,19 | 234 | 150 | 0 | 0 | 1.0,1.0,1.0 | 0.775,0.54,0.658 | 0.342 | ENSG00000197 | [[[[52695368, 52 | 2 |
| ENSG00000138 | SEC31A | chr4 | 82880798 | 82880922 | 82878729 | 82878928 | 82891087 | 82891178 | 39,30,80 | 0,0,0 | 30,67,65 | 3,4,4 | 274 | 150 | 4.86E-05 | 0.000850119 | 1.0,1.0,1.0 | 0.846,0.902,0.89 | 0.118 | ENSG00000138 | [[[[82891088, 82 | 2 |
| ENSG00000138 | SEC31A | chr4 | 82881857 | 82881940 | 82878729 | 82878928 | 82891087 | 82891120 | 4,2,16 | 0,0,0 | 8,28,29 | 3,4,4 | 233 | 150 | 5.86E-06 | 0.000133487 | 1.0,1.0,1.0 | 0.632,0.818,0.82 | 0.242 | ENSG00000138 | [[[[82891088, 82 | 2 |
| ENSG00000122 | ZC3H7A | chr16 | 11776319 | 11776358 | 11774979 | 11775013 | 11779165 | 11779363 | 14,12,14 | 0,0,0 | 6,29,19 | 1,4,3 | 189 | 150 | 0.000186936 | 0.002751429 | 1.0,1.0,1.0 | 0.826,0.852,0.83 | 0.163 | ENSG00000122 | [[[[11797124, 11 | 3 |
| ENSG00000165 | WRN | chr8 | 31111614 | 31111799 | 31100848 | 31100955 | 31120242 | 31120424 | 6,7,22 | 0,0,0 | 6,20,12 | 1,5,3 | 300 | 150 | 8.89E-07 | 2.47E-05 | 1.0,1.0,1.0 | 0.75,0.667,0.667 | 0.305 | ENSG00000165 | [[[[31033810, 31 | 2 |
| ENSG00000118 | CCNI | chr4 | 77055961 | 77056100 | 77055322 | 77055380 | 77066248 | 77066405 | 244,235,359 | 1,0,0 | 232,445,312 | 16,32,33 | 289 | 150 | 0 | 0 | 0.992,1.0,1.0 | 0.883,0.878,0.83 | 0.133 | ENSG00000118 | [[[[77075472, 77 | 3 |
| ENSG00000118 | CCNI | chr4 | 77055961 | 77056102 | 77055149 | 77055380 | 77066248 | 77066405 | 245,239,360 | 1,0,0 | 240,456,316 | 16,32,33 | 291 | 150 | 0 | 0 | 0.992,1.0,1.0 | 0.885,0.88,0.832 | 0.132 | ENSG00000118 | [[[[77075472, 77 | 3 |
| ENSG00000118 | CCNI | chr4 | 77055961 | 77056102 | 77055149 | 77055380 | 77075471 | 77076005 | 245,239,360 | 0,0,0 | 237,453,313 | 14,8,12 | 291 | 150 | 4.94E-09 | 2.36E-07 | 1.0,1.0,1.0 | 0.897,0.967,0.93 | 0.068 | ENSG00000118 | [[[[77075472, 77 | 4 |
| ENSG00000118 | CCNI | chr4 | 77058506 | 77058635 | 77055149 | 77055380 | 77066248 | 77066405 | 276,220,301 | 1,0,0 | 268,493,343 | 16,32,33 | 279 | 150 | 0 | 0 | 0.993,1.0,1.0 | 0.9,0.892,0.848 | 0.118 | ENSG00000118 | [[[[77075472, 77 | 3 |
| ENSG00000118 | CCNI | chr4 | 77066248 | 77066405 | 77055149 | 77055380 | 77075471 | 77075693 | 94,96,221 | 0,0,0 | 157,325,242 | 14,8,12 | 300 | 150 | 1.58E-08 | 6.74E-07 | 1.0,1.0,1.0 | 0.849,0.953,0.91 | 0.096 | ENSG00000118 | [[[[77075472, 77 | 4 |
| ENSG00000138 | NUP54 | chr4 | 76131229 | 76131284 | 76130655 | 76130749 | 76136185 | 76136412 | 14,10,15 | 0,0,0 | 11,38,16 | 6,2,7 | 205 | 150 | 5.56E-07 | 1.63E-05 | 1.0,1.0,1.0 | 0.573,0.933,0.62 | 0.289 | ENSG00000138 | [[[[76148308, 76 | 3 |
| ENSG00000138 | NUP54 | chr4 | 76131229 | 76131284 | 76130655 | 76130749 | 76144148 | 76144292 | 14,10,15 | 0,0,0 | 12,42,19 | 0,10,4 | 205 | 150 | 4.28E-06 | 0.00010067 | 1.0,1.0,1.0 | 1.0,0.754,0.777 | 0.156 | ENSG00000138 | [[[[76148308, 76 | 4 |
| ENSG00000138 | NUP54 | chr4 | 76132522 | 76132719 | 76131229 | 76131284 | 76144148 | 76144292 | 43,41,68 | 0,0,0 | 33,90,52 | 1,7,3 | 300 | 150 | 5.14E-05 | 0.000894756 | 1.0,1.0,1.0 | 0.943,0.865,0.89 | 0.098 | ENSG00000138 | [[[[76148308, 76 | 3 |
| ENSG00000138 | NUP54 | chr4 | 76134174 | 76134362 | 76132522 | 76132719 | 76144148 | 76144292 | 51,47,109 | 0,0,0 | 37,59,57 | 2,9,5 | 300 | 150 | 4.01E-08 | 1.55E-06 | 1.0,1.0,1.0 | 0.902,0.766,0.85 | 0.16 | ENSG00000138 | [[[[76148308, 76 | 2 |
| ENSG00000138 | NUP54 | chr4 | 76136185 | 76136412 | 76130655 | 76130749 | 76144148 | 76144292 | 36,33,56 | 0,0,0 | 51,95,62 | 0,10,4 | 300 | 150 | 1.79E-05 | 0.000356015 | 1.0,1.0,1.0 | 1.0,0.826,0.886 | 0.096 | ENSG00000138 | [[[[76148308, 76 | 4 |
| ENSG00000138 | NUP54 | chr4 | 76136185 | 76136412 | 76131229 | 76131284 | 76144148 | 76144292 | 36,33,56 | 0,0,0 | 45,96,55 | 1,7,3 | 300 | 150 | 0.000117903 | 0.001841355 | 1.0,1.0,1.0 | 0.957,0.873,0.90 | 0.089 | ENSG00000138 | [[[[76148308, 76 | 3 |
| ENSG00000006 | DBF4 | chr7 | 87886843 | 87886894 | 87878052 | 87878133 | 87887328 | 87887398 | 11,8,12 | 0,0,0 | 8,23,19 | 3,5,0 | 201 | 150 | 0.000132841 | 0.002039131 | 1.0,1.0,1.0 | 0.666,0.774,1.0 | 0.187 | ENSG00000006 | [[[[87876493, 87 | 2 |
| ENSG00000132 | ANKRD17 | chr4 | 73097116 | 73097272 | 73090666 | 73092300 | 73098072 | 73098520 | 21,30,34 | 0,0,0 | 27,91,34 | 1,11,3 | 300 | 150 | 2.81E-06 | 6.92E-05 | 1.0,1.0,1.0 | 0.931,0.805,0.85 | 0.138 | ENSG00000132 | [[[[73258276, 73 | 2 |
| ENSG00000170 | ARL6IP1 | chr16 | 18797924 | 18797952 | 18795463 | 18795581 | 18801430 | 18801553 | 53,63,91 | 0,0,0 | 33,78,49 | 5,5,5 | 178 | 150 | 1.77E-06 | 4.60E-05 | 1.0,1.0,1.0 | 0.848,0.929,0.89 | 0.11 | ENSG00000170 | [[[[18801431, 18 | 2 |
| ENSG00000170 | ARL6IP1 | chr16 | 18797924 | 18798044 | 18795175 | 18795581 | 18801430 | 18801547 | 306,243,384 | 0,0,0 | 134,283,189 | 5,5,5 | 270 | 150 | 0.000375244 | 0.005013647 | 1.0,1.0,1.0 | 0.937,0.969,0.95 | 0.046 | ENSG00000170 | [[[[18801431, 18 | 2 |

| | | | | | | | | | | | | | | | | | | | | | | |
|---|---|---|---|---|---|---|---|---|---|---|---|---|---|---|---|---|---|---|---|---|---|---|
| ENSG00000136 | KDELR2 | chr7 | 6474183 | 6474284 | 6466070 | 6466323 | 6483966 | 6484057 | 194,285,550 | 1,0,0 | 404,771,628 | 13,13,16 | 251 | 150 | 8.41E-07 | 2.35E-05 | 0.991,1.0,1.0 | 0.949,0.973,0.95 | 0.037 | ENSG00000136 | [[[[6483967, 648 | 2 |
| ENSG00000126 | EMG1 | chr12 | 6975228 | 6975378 | 6970892 | 6971091 | 6975695 | 6979941 | 63,89,98 | 0,0,0 | 58,137,76 | 7,2,2 | 300 | 150 | 0.000230836 | 0.00329017 | 1.0,1.0,1.0 | 0.806,0.972,0.95 | 0.091 | ENSG00000126 | [[[[6970913, 697 | 4 |
| ENSG00000126 | EMG1 | chr12 | 6975228 | 6975378 | 6975089 | 6975148 | 6983001 | 6983206 | 149,133,191 | 0,0,0 | 126,240,151 | 3,8,2 | 300 | 150 | 0.00082219 | 0.009835047 | 1.0,1.0,1.0 | 0.955,0.938,0.97 | 0.044 | ENSG00000126 | [[[[6970913, 697 | 2 |
| ENSG00000126 | EMG1 | chr12 | 6983001 | 6983206 | 6975089 | 6975148 | 6987781 | 6987838 | 3,2,5 | 0,0,0 | 9,14,14 | 1,6,0 | 300 | 150 | 0.000269221 | 0.003757851 | 1.0,1.0,1.0 | 0.818,0.538,1.0 | 0.215 | ENSG00000126 | [[[[6970913, 697 | 2 |
| ENSG00000143 | PRUNE1 | chr1 | 151024610 | 151024795 | 151018466 | 151018669 | 151027232 | 151027327 | 11,10,8 | 0,0,0 | 6,31,12 | 3,3,1 | 300 | 150 | 0.000140307 | 0.002135607 | 1.0,1.0,1.0 | 0.5,0.838,0.857 | 0.268 | ENSG00000143 | [[[[151008449, 1 | 2 |
| ENSG00000198 | CAPZA2 | chr7 | 116888126 | 116888190 | 116862472 | 116862650 | 116898771 | 116898786 | 9,28,40 | 0,0,0 | 15,21,19 | 10,34,18 | 214 | 150 | 0 | 0 | 1.0,1.0,1.0 | 0.513,0.302,0.42 | 0.587 | ENSG00000198 | [[[[116862587, 11 | 2 |
| ENSG00000198 | CAPZA2 | chr7 | 116892993 | 116893045 | 116862472 | 116862650 | 116898771 | 116898786 | 27,26,36 | 0,0,0 | 16,43,41 | 10,34,18 | 202 | 150 | 0 | 0 | 1.0,1.0,1.0 | 0.543,0.484,0.62 | 0.448 | ENSG00000198 | [[[[116862587, 11 | 2 |
| ENSG00000198 | CAPZA2 | chr7 | 116893030 | 116893045 | 116862529 | 116862650 | 116898771 | 116898835 | 7,6,13 | 0,0,0 | 25,33,45 | 10,34,18 | 165 | 150 | 1.64E-09 | 8.73E-08 | 1.0,1.0,1.0 | 0.694,0.469,0.69 | 0.381 | ENSG00000198 | [[[[116862587, 11 | 2 |
| ENSG00000164 | FASTK | chr7 | 151076912 | 151077027 | 151076623 | 151076832 | 151077309 | 151077401 | 2,26,58 | 0,0,6 | 23,58,34 | 16,24,15 | 265 | 150 | 3.41E-07 | 1.05E-05 | 1.0,1.0,0.845 | 0.449,0.578,0.56 | 0.419 | ENSG00000164 | [[[[151080685, 1 | 2 |
| ENSG00000164 | FASTK | chr7 | 151076912 | 151077236 | 151076625 | 151076832 | 151077309 | 151077401 | 42,142,240 | 0,0,6 | 146,206,152 | 16,24,15 | 300 | 150 | 2.70E-10 | 1.66E-08 | 1.0,1.0,0.952 | 0.82,0.811,0.835 | 0.162 | ENSG00000164 | [[[[151080685, 1 | 2 |
| ENSG00000164 | FASTK | chr7 | 151077100 | 151077236 | 151076623 | 151076832 | 151077309 | 151077401 | 39,114,191 | 0,0,6 | 123,155,124 | 16,24,15 | 286 | 150 | 2.28E-10 | 1.41E-08 | 1.0,1.0,0.943 | 0.801,0.772,0.81 | 0.186 | ENSG00000164 | [[[[151080685, 1 | 2 |
| ENSG00000112 | CAP2 | chr6 | 17539268 | 17539458 | 17513848 | 17513954 | 17543060 | 17543143 | 52,46,81 | 0,0,0 | 35,32,37 | 10,12,3 | 300 | 150 | 1.09E-11 | 8.29E-10 | 1.0,1.0,1.0 | 0.636,0.571,0.86 | 0.311 | ENSG00000112 | [[[[17393595, 17 | 3 |
| ENSG00000112 | CAP2 | chr6 | 17542836 | 17542960 | 17513848 | 17513954 | 17543060 | 17543143 | 38,30,68 | 0,0,0 | 23,48,33 | 10,12,3 | 274 | 150 | 1.15E-10 | 7.47E-09 | 1.0,1.0,1.0 | 0.557,0.686,0.85 | 0.3 | ENSG00000112 | [[[[17393595, 17 | 3 |
| ENSG00000103 | WDR59 | chr16 | 74888168 | 74888319 | 74887682 | 74887755 | 74892483 | 74892565 | 23,16,54 | 0,0,0 | 19,57,26 | 3,12,3 | 300 | 150 | 1.16E-08 | 5.10E-07 | 1.0,1.0,1.0 | 0.76,0.704,0.813 | 0.241 | ENSG00000103 | [[[[74984964, 74 | 2 |
| ENSG00000164 | UBLCP1 | chr5 | 159270359 | 159270445 | 159269907 | 159269999 | 159274584 | 159274622 | 42,33,50 | 0,0,0 | 18,47,51 | 0,6,4 | 236 | 150 | 4.83E-05 | 0.00084632 | 1.0,1.0,1.0 | 1.0,0.833,0.89 | 0.092 | ENSG00000164 | [[[[159263290, 1 | 3 |
| ENSG00000164 | EDIL3 | chr5 | 84180392 | 84180521 | 84137240 | 84137354 | 84384307 | 84384386 | 231,134,314 | 0,0,0 | 125,366,240 | 10,14,9 | 279 | 150 | 7.05E-10 | 4.02E-08 | 1.0,1.0,1.0 | 0.87,0.934,0.935 | 0.087 | ENSG00000164 | [[[[84384308, 84 | 3 |
| ENSG00000164 | EDIL3 | chr5 | 84229854 | 84229884 | 84137240 | 84137354 | 84254083 | 84254212 | 14,10,22 | 0,0,1 | 7,21,22 | 4,3,3 | 180 | 150 | 5.27E-05 | 0.000915131 | 1.0,1.0,0.948 | 0.593,0.854,0.85 | 0.214 | ENSG00000164 | [[[[84384308, 84 | 2 |
| ENSG00000164 | EDIL3 | chr5 | 84254083 | 84254212 | 84137240 | 84137354 | 84384307 | 84384386 | 81,80,203 | 0,0,0 | 62,196,147 | 10,14,9 | 279 | 150 | 7.35E-11 | 4.92E-09 | 1.0,1.0,1.0 | 0.769,0.883,0.89 | 0.15 | ENSG00000164 | [[[[84384308, 84 | 3 |
| ENSG00000065 | TMEM206 | chr1 | 212385273 | 212385425 | 212379894 | 212380037 | 212410424 | 212410521 | 43,39,43 | 0,1,0 | 25,35,14 | 2,3,2 | 300 | 150 | 0.000569572 | 0.007185653 | 1.0,0.951,1.0 | 0.862,0.854,0.77 | 0.152 | ENSG00000065 | [[[[212414722, 2 | 2 |
| ENSG00000117 | DPH5 | chr1 | 101013709 | 101013818 | 101001466 | 101001544 | 101025308 | 101025351 | 6,3,4 | 0,0,0 | 7,38,12 | 10,33,18 | 259 | 150 | 1.30E-09 | 7.05E-08 | 1.0,1.0,1.0 | 0.288,0.4,0.279 | 0.678 | ENSG00000117 | [[[[101025683, 1 | 2 |
| ENSG00000117 | DPH5 | chr1 | 101013709 | 101013818 | 101001466 | 101001587 | 101025308 | 101025351 | 31,28,31 | 0,0,0 | 32,89,48 | 11,15,10 | 259 | 150 | 1.93E-10 | 1.21E-08 | 1.0,1.0,1.0 | 0.628,0.775,0.73 | 0.287 | ENSG00000117 | [[[[101025683, 1 | 2 |
| ENSG00000117 | DPH5 | chr1 | 101021640 | 101021765 | 101001466 | 101001544 | 101025308 | 101025351 | 63,63,89 | 0,0,0 | 102,187,126 | 10,33,18 | 275 | 150 | 8.65E-14 | 9.04E-12 | 1.0,1.0,1.0 | 0.848,0.756,0.79 | 0.201 | ENSG00000117 | [[[[101025683, 1 | 2 |
| ENSG00000117 | DPH5 | chr1 | 101021640 | 101021765 | 101001466 | 101001544 | 101025682 | 101025815 | 12,8,6 | 0,0,0 | 9,10,14 | 0,3,4 | 275 | 150 | 7.02E-05 | 0.001170853 | 1.0,1.0,1.0 | 1.0,0.645,0.656 | 0.233 | ENSG00000117 | [[[[101025683, 1 | 3 |
| ENSG00000117 | DPH5 | chr1 | 101021640 | 101021765 | 101001466 | 101001587 | 101025308 | 101025351 | 53,58,84 | 0,0,0 | 96,189,119 | 11,15,10 | 275 | 150 | 2.87E-09 | 1.45E-07 | 1.0,1.0,1.0 | 0.826,0.873,0.86 | 0.145 | ENSG00000117 | [[[[101025683, 1 | 2 |
| ENSG00000237 | C2orf74 | chr2 | 61157861 | 61158012 | 61145129 | 61145196 | 61162841 | 61162955 | 8,0,4 | 42,28,40 | 18,28,19 | 28,61,40 | 300 | 150 | 2.01E-07 | 6.51E-06 | 0.087,0.0,0.048 | 0.243,0.187,0.19 | -0.162 | ENSG00000237 | [[[[61162188, 61 | 2 |
| ENSG00000115 | FANCL | chr2 | 58162865 | 58162947 | 58161521 | 58161638 | 58163433 | 58163517 | 18,14,29 | 2,0,2 | 10,24,10 | 2,12,5 | 232 | 150 | 7.25E-05 | 0.001204058 | 0.853,1.0,0.904 | 0.764,0.564,0.56 | 0.288 | ENSG00000115 | [[[[58241218, 58 | 2 |
| ENSG00000275 | PPP4R3B | chr2 | 55588878 | 55588956 | 55579678 | 55579781 | 55598415 | 55599039 | 12,8,23 | 0,0,0 | 6,28,21 | 2,4,3 | 228 | 150 | 9.21E-06 | 0.000199355 | 1.0,1.0,1.0 | 0.664,0.822,0.82 | 0.231 | ENSG00000275 | [[[[55617144, 55 | 4 |
| ENSG00000170 | POLH | chr6 | 43583006 | 43583141 | 43576373 | 43576440 | 43587271 | 43587400 | 23,14,19 | 0,0,0 | 6,36,11 | 3,7,2 | 285 | 150 | 1.08E-07 | 3.75E-06 | 1.0,1.0,1.0 | 0.513,0.73,0.743 | 0.338 | ENSG00000170 | [[[[43576185, 43 | 2 |
| ENSG00000170 | POLH | chr6 | 43605253 | 43605319 | 43600987 | 43601091 | 43610553 | 43610723 | 6,5,10 | 0,0,0 | 9,21,8 | 3,3,8 | 216 | 150 | 1.42E-07 | 4.78E-06 | 1.0,1.0,1.0 | 0.676,0.829,0.41 | 0.362 | ENSG00000170 | [[[[43576185, 43 | 3 |
| ENSG00000182 | CEP97 | chr3 | 101728835 | 101728937 | 101723924 | 101724719 | 101731839 | 101731953 | 4,4,2 | 0,0,0 | 8,22,17 | 2,3,2 | 252 | 150 | 0.000250815 | 0.00353199 | 1.0,1.0,1.0 | 0.704,0.814,0.83 | 0.216 | ENSG00000182 | [[[[101724614, 1 | 3 |
| ENSG00000115 | SOS1 | chr2 | 39051143 | 39051287 | 39035211 | 39035310 | 39054613 | 39054823 | 14,4,18 | 0,0,0 | 3,11,7 | 3,3,5 | 294 | 150 | 5.54E-10 | 3.22E-08 | 1.0,1.0,1.0 | 0.338,0.652,0.41 | 0.531 | ENSG00000115 | [[[[39120336, 39 | 2 |
| ENSG00000178 | ZFAND2A | chr7 | 1157655 | 1157750 | 1155452 | 1155584 | 1159963 | 1160153 | 29,34,51 | 0,0,0 | 157,250,126 | 81,132,69 | 245 | 150 | 0 | 0 | 1.0,1.0,1.0 | 0.543,0.537,0.52 | 0.464 | ENSG00000178 | [[[[1159964, 116 | 2 |
| ENSG00000178 | ZFAND2A | chr7 | 1158157 | 1158257 | 1155452 | 1155584 | 1159963 | 1160153 | 40,66,95 | 0,0,0 | 210,427,210 | 81,132,69 | 250 | 150 | 0 | 0 | 1.0,1.0,1.0 | 0.609,0.66,0.646 | 0.362 | ENSG00000178 | [[[[1159964, 116 | 2 |
| ENSG00000138 | METTL5 | chr2 | 169821942 | 169822057 | 169819560 | 169819643 | 169824488 | 169824558 | 80,83,95 | 0,0,0 | 78,170,105 | 24,46,16 | 265 | 150 | 0 | 0 | 1.0,1.0,1.0 | 0.648,0.677,0.78 | 0.296 | ENSG00000138 | [[[[169824489, 1 | 2 |
| ENSG00000138 | METTL5 | chr2 | 169824488 | 169824640 | 169819560 | 169819643 | 169824781 | 169824814 | 6,16,51 | 0,0,0 | 55,95,46 | 1,5,4 | 300 | 150 | 0.000429733 | 0.005633917 | 1.0,1.0,1.0 | 0.965,0.905,0.85 | 0.093 | ENSG00000138 | [[[[169824489, 1 | 2 |
| ENSG00000138 | LRPPRC | chr2 | 43960540 | 43960634 | 43957384 | 43957451 | 43973606 | 43973714 | 39,38,67 | 0,0,0 | 49,70,70 | 10,10,16 | 244 | 150 | 5.21E-12 | 4.24E-10 | 1.0,1.0,1.0 | 0.751,0.811,0.72 | 0.236 | ENSG00000138 | [[[[43995799, 43 | 2 |
| ENSG00000018 | AGPS | chr2 | 177505505 | 177505575 | 177499617 | 177499730 | 177521268 | 177521368 | 88,58,99 | 0,0,0 | 65,111,91 | 3,10,5 | 220 | 150 | 1.26E-06 | 3.38E-05 | 1.0,1.0,1.0 | 0.937,0.883,0.92 | 0.085 | ENSG00000018 | [[[[177392773, 1 | 3 |
| ENSG00000059 | ALDH18A1 | chr10 | 95628367 | 95628492 | 95626702 | 95626776 | 95632958 | 95633049 | 62,75,146 | 0,0,0 | 68,147,97 | 3,10,9 | 275 | 150 | 3.32E-08 | 1.31E-06 | 1.0,1.0,1.0 | 0.925,0.889,0.85 | 0.11 | ENSG00000059 | [[[[95656597, 95 | 2 |
| ENSG00000174 | SLC25A30 | chr13 | 45408926 | 45409074 | 45405882 | 45405977 | 45418299 | 45418380 | 27,21,38 | 0,0,0 | 36,57,32 | 1,3,4 | 298 | 150 | 0.000764657 | 0.009227965 | 1.0,1.0,1.0 | 0.948,0.905,0.80 | 0.115 | ENSG00000174 | [[[[45418300, 45 | 2 |
| ENSG00000174 | SLC25A30 | chr13 | 45411361 | 45411480 | 45405882 | 45405977 | 45418299 | 45418380 | 14,4,27 | 0,0,0 | 14,18,33 | 1,3,4 | 269 | 150 | 9.48E-05 | 0.00152495 | 1.0,1.0,1.0 | 0.886,0.77,0.821 | 0.174 | ENSG00000174 | [[[[45418300, 45 | 2 |
| ENSG00000112 | HARS2 | chr5 | 140693934 | 140694054 | 140691608 | 140691756 | 140695737 | 140695845 | 1,1,2 | 0,0,0 | 2,14,2 | 3,2,2 | 270 | 150 | 1.25E-06 | 3.36E-05 | 1.0,1.0,1.0 | 0.27,0.795,0.357 | 0.526 | ENSG00000112 | [[[[140691455, 1 | 4 |
| ENSG00000109 | OCIAD1 | chr4 | 48849946 | 48850082 | 48848398 | 48848446 | 48857212 | 48857252 | 300,258,358 | 0,0,1 | 208,385,215 | 53,102,51 | 286 | 150 | 0 | 0 | 1.0,1.0,0.995 | 0.673,0.664,0.68 | 0.323 | ENSG00000109 | [[[[48831088, 48 | 2 |
| ENSG00000109 | OCIAD1 | chr4 | 48857212 | 48857365 | 48848398 | 48848446 | 48860724 | 48861200 | 238,228,284 | 0,0,0 | 276,601,329 | 9,17,6 | 300 | 150 | 4.96E-07 | 1.47E-05 | 1.0,1.0,1.0 | 0.939,0.946,0.96 | 0.05 | ENSG00000109 | [[[[48831088, 48 | 2 |
| ENSG00000153 | TMEM87B | chr2 | 112077191 | 112077282 | 112066935 | 112067067 | 112081056 | 112081118 | 17,19,23 | 0,0,0 | 11,17,13 | 5,2,1 | 241 | 150 | 1.41E-05 | 0.000288014 | 1.0,1.0,1.0 | 0.578,0.841,0.89 | 0.23 | ENSG00000153 | [[[[112055269, 1 | 2 |
| ENSG00000171 | EXOSC1 | chr10 | 97438669 | 97438703 | 97437699 | 97437750 | 97445731 | 97445847 | 28,13,35 | 0,0,0 | 57,106,73 | 22,28,14 | 184 | 150 | 3.13E-11 | 2.20E-09 | 1.0,1.0,1.0 | 0.679,0.755,0.81 | 0.252 | ENSG00000171 | [[[[97445955, 97 | 3 |
| ENSG00000171 | EXOSC1 | chr10 | 97441170 | 97441259 | 97437699 | 97437750 | 97445731 | 97445847 | 7,10,15 | 0,0,0 | 12,15,3 | 22,28,14 | 239 | 150 | 0 | 0 | 1.0,1.0,1.0 | 0.255,0.252,0.11 | 0.791 | ENSG00000171 | [[[[97445955, 97 | 3 |
| ENSG00000171 | EXOSC1 | chr10 | 97441170 | 97441259 | 97438662 | 97438703 | 97445731 | 97445847 | 70,58,110 | 0,0,0 | 88,167,91 | 27,55,51 | 239 | 150 | 0 | 0 | 1.0,1.0,1.0 | 0.672,0.656,0.52 | 0.381 | ENSG00000171 | [[[[97445955, 97 | 2 |
| ENSG00000171 | EXOSC1 | chr10 | 97443236 | 97443311 | 97437699 | 97437750 | 97445731 | 97445847 | 66,132,177 | 0,0,0 | 109,272,146 | 22,28,14 | 225 | 150 | 6.66E-16 | 9.20E-14 | 1.0,1.0,1.0 | 0.768,0.866,0.87 | 0.164 | ENSG00000171 | [[[[97445955, 97 | 3 |
| ENSG00000163 | RCHY1 | chr4 | 75508819 | 75508935 | 75494100 | 75494179 | 75514196 | 75514286 | 15,10,28 | 0,0,0 | 46,91,32 | 2,12,6 | 266 | 150 | 2.90E-06 | 7.11E-05 | 1.0,1.0,1.0 | 0.928,0.81,0.75 | 0.171 | ENSG00000163 | [[[[75514197, 75 | 2 |
| ENSG00000143 | SCCPDH | chr1 | 246759033 | 246759151 | 246758225 | 246758356 | 246766057 | 246766139 | 92,78,95 | 0,0,0 | 42,90,54 | 4,8,5 | 268 | 150 | 3.50E-08 | 1.38E-06 | 1.0,1.0,1.0 | 0.855,0.863,0.85 | 0.141 | ENSG00000143 | [[[[246724409, 2 | 4 |
| ENSG00000143 | SCCPDH | chr1 | 246764245 | 246764357 | 246758225 | 246758356 | 246766057 | 246766139 | 100,79,85 | 0,0,0 | 50,87,58 | 4,8,5 | 262 | 150 | 6.22E-08 | 2.30E-06 | 1.0,1.0,1.0 | 0.877,0.862,0.86 | 0.131 | ENSG00000143 | [[[[246724409, 2 | 4 |
| ENSG00000159 | GART | chr21 | 33505560 | 33505702 | 33504411 | 33504527 | 33509782 | 33509920 | 44,45,64 | 0,0,0 | 38,70,37 | 3,5,4 | 292 | 150 | 6.26E-06 | 0.00014169 | 1.0,1.0,1.0 | 0.867,0.878,0.82 | 0.143 | ENSG00000159 | [[[[33542065, 33 | 2 |
| ENSG00000159 | GART | chr21 | 33528166 | 33528335 | 33524008 | 33525000 | 33530758 | 33530884 | 74,69,108 | 0,0,0 | 88,224,161 | 4,7,4 | 300 | 150 | 0.000233624 | 0.003323692 | 1.0,1.0,1.0 | 0.917,0.941,0.95 | 0.063 | ENSG00000159 | [[[[33542065, 33 | 3 |
| ENSG00000159 | GART | chr21 | 33528849 | 33528937 | 33524008 | 33525000 | 33530758 | 33530884 | 32,42,50 | 0,0,0 | 27,46,28 | 4,7,4 | 238 | 150 | 6.20E-08 | 2.30E-06 | 1.0,1.0,1.0 | 0.81,0.806,0.815 | 0.19 | ENSG00000159 | [[[[33542065, 33 | 3 |
| ENSG00000159 | GART | chr21 | 33528849 | 33528937 | 33528166 | 33528335 | 33530758 | 33530884 | 32,42,50 | 0,0,0 | 27,43,28 | 19,47,54 | 238 | 150 | 0 | 0 | 1.0,1.0,1.0 | 0.472,0.366,0.24 | 0.639 | ENSG00000159 | [[[[33542065, 33 | 2 |
| ENSG00000159 | GART | chr21 | 33531488 | 33531557 | 33528166 | 33528335 | 33532344 | 33532456 | 48,32,78 | 0,0,0 | 36,134,67 | 4,3,5 | 219 | 150 | 0.000341205 | 0.004606515 | 1.0,1.0,1.0 | 0.86,0.968,0.902 | 0.09 | ENSG00000159 | [[[[33542065, 33 | 4 |
| ENSG00000144 | MRPS5 | chr2 | 95100473 | 95100536 | 95090385 | 95090522 | 95106422 | 95106457 | 42,35,67 | 0,0,0 | 53,132,58 | 19,29,14 | 213 | 150 | 4.33E-15 | 5.42E-13 | 1.0,1.0,1.0 | 0.663,0.762,0.74 | 0.277 | ENSG00000144 | [[[[95121734, 95 | 4 |
| ENSG00000144 | MRPS5 | chr2 | 95101676 | 95101723 | 95100836 | 95100894 | 95106422 | 95106457 | 22,19,44 | 0,0,0 | 87,173,93 | 11,16,8 | 197 | 150 | 8.52E-07 | 2.38E-05 | 1.0,1.0,1.0 | 0.858,0.892,0.89 | 0.117 | ENSG00000144 | [[[[95121734, 95 | 2 |
| ENSG00000144 | MRPS5 | chr2 | 95104639 | 95104730 | 95100836 | 95100894 | 95106422 | 95106457 | 100,72,130 | 0,0,0 | 100,193,107 | 11,16,8 | 241 | 150 | 1.36E-10 | 8.74E-09 | 1.0,1.0,1.0 | 0.85,0.882,0.893 | 0.125 | ENSG00000144 | [[[[95121734, 95 | 2 |
| ENSG00000153 | CDYL | chr6 | 4937564 | 4937737 | 4935514 | 4935771 | 4952265 | 4952409 | 12,14,38 | 0,0,0 | 15,78,38 | 4,10,3 | 300 | 150 | 8.45E-07 | 2.36E-05 | 1.0,1.0,1.0 | 0.652,0.796,0.86 | 0.229 | ENSG00000153 | [[[[4776404, 477 | 2 |
| ENSG00000171 | RPS7 | chr2 | 3575591 | 3575684 | 3575204 | 3575350 | 3576558 | 3576630 | 19,37,43 | 0,0,0 | 33,82,48 | 7,8,0 | 243 | 150 | 1.03E-05 | 0.00022014 | 1.0,1.0,1.0 | 0.744,0.864,1.0 | 0.131 | ENSG00000171 | [[[[3575260, 357 | 2 |
| ENSG00000171 | ZNF692 | chr1 | 248855379 | 248855458 | 248853936 | 248854051 | 248855724 | 248855943 | 18,4,16 | 0,0,0 | 23,39,14 | 3,8,4 | 229 | 150 | 6.97E-07 | 1.99E-05 | 1.0,1.0,1.0 | 0.834,0.762,0.69 | 0.236 | ENSG00000171 | [[[[248858918, 2 | 2 |
| ENSG00000103 | SETD6 | chr16 | 58518400 | 58518543 | 58516807 | 58516928 | 58518723 | 58518872 | 15,22,18 | 0,0,0 | 34,53,29 | 0,3,7 | 293 | 150 | 8.46E-05 | 0.001376168 | 1.0,1.0,1.0 | 1.0,0.9,0.68 | 0.14 | ENSG00000103 | [[[[58515517, 58 | 2 |
| ENSG00000106 | PHF14 | chr7 | 11035639 | 11035786 | 11028680 | 11028818 | 11036984 | 11037091 | 97,77,83 | 0,0,0 | 90,174,110 | 8,10,1 | 297 | 150 | 2.61E-06 | 6.47E-05 | 1.0,1.0,1.0 | 0.85,0.898,0.982 | 0.09 | ENSG00000106 | [[[[10973872, 10 | 2 |
| ENSG00000106 | PHF14 | chr7 | 11038759 | 11038855 | 11028680 | 11028818 | 11040671 | 11040775 | 59,39,71 | 0,0,0 | 70,140,77 | 10,26,8 | 246 | 150 | 7.92E-12 | 6.18E-10 | 1.0,1.0,1.0 | 0.81,0.767,0.854 | 0.19 | ENSG00000106 | [[[[10973872, 10 | 4 |
| ENSG00000106 | PHF14 | chr7 | 11061790 | 11061841 | 11051611 | 11051780 | 11111349 | 11111467 | 32,34,38 | 0,1,0 | 31,77,43 | 7,8,7 | 201 | 150 | 3.76E-07 | 1.15E-05 | 1.0,0.962,1.0 | 0.768,0.878,0.82 | 0.165 | ENSG00000106 | [[[[10973872, 10 | 2 |
| ENSG00000106 | PHF14 | chr7 | 11061963 | 11062085 | 11051611 | 11051780 | 11111349 | 11111467 | 127,96,104 | 0,1,0 | 142,278,163 | 7,8,7 | 272 | 150 | 2.30E-05 | 0.000445102 | 1.0,0.981,1.0 | 0.918,0.95,0.928 | 0.062 | ENSG00000106 | [[[[10973872, 10 | 2 |
| ENSG00000278 | MRPL45 | chr17 | 38299350 | 38299468 | 38297022 | 38297249 | 38306532 | 38306631 | 66,57,73 | 0,0,0 | 59,128,97 | 9,37,14 | 268 | 150 | 5.22E-15 | 6.48E-13 | 1.0,1.0,1.0 | 0.786,0.659,0.79 | 0.253 | ENSG00000278 | [[[[38297168, 38 | 2 |
| ENSG00000203 | COX20 | chr1 | 244842194 | 244842258 | 244835655 | 244835756 | 244843040 | 244843252 | 172,166,240 | 0,0,0 | 145,238,163 | 2,12,4 | 214 | 150 | 6.94E-05 | 0.001160539 | 1.0,1.0,1.0 | 0.981,0.933,0.96 | 0.04 | ENSG00000203 | [[[[244835658, 2 | 2 |
| ENSG00000196 | CASP4 | chr11 | 104944747 | 104944851 | 104942865 | 104942973 | 104948532 | 104948676 | 18,21,11 | 0,0,0 | 17,34,28 | 5,12,2 | 254 | 150 | 1.04E-07 | 3.62E-06 | 1.0,1.0,1.0 | 0.668,0.626,0.89 | 0.271 | ENSG00000196 | [[[[104968519, 1 | 2 |
| ENSG00000196 | CASP4 | chr11 | 104947082 | 104947192 | 104942868 | 104942973 | 104948532 | 104948676 | 59,51,89 | 0,0,0 | 87,131,95 | 5,12,2 | 260 | 150 | 2.58E-06 | 6.41E-05 | 1.0,1.0,1.0 | 0.909,0.863,0.96 | 0.088 | ENSG00000196 | [[[[104968519, 1 | 2 |
| ENSG00000122 | HERPUD2 | chr7 | 35673200 | 35673278 | 35667433 | 35667588 | 35694183 | 35694627 | 17,14,13 | 0,3,9 | 21,52,19 | 1,1,0 | 228 | 150 | 6.33E-05 | 0.001072614 | 1.0,0.754,0.487 | 0.933,0.972,1.0 | -0.221 | ENSG00000122 | [[[[35694801, 35 | 2 |
| ENSG00000054 | SYNE2 | chr14 | 64216247 | 64216387 | 64214143 | 64214470 | 64218397 | 64218512 | 56,60,64 | 0,0,0 | 60,135,74 | 2,3,6 | 290 | 150 | 0.000492548 | 0.006351252 | 1.0,1.0,1.0 | 0.939,0.959,0.86 | 0.079 | ENSG00000054 | [[[[63853004, 63 | 2 |
| ENSG00000162 | ACP6 | chr1 | 147659395 | 147659526 | 147658959 | 147659039 | 147669829 | 147670278 | 82,51,120 | 0,0,0 | 20,41,42 | 5,8,4 | 281 | 150 | 1.45E-10 | 9.27E-09 | 1.0,1.0,1.0 | 0.681,0.732,0.84 | 0.246 | ENSG00000162 | [[[[147669830, 1 | 2 |
| ENSG00000119 | HEATR1 | chr1 | 236574201 | 236574333 | 236572724 | 236572828 | 236576218 | 236576377 | 20,15,31 | 0,0,0 | 4,5,10 | 0,4,4 | 282 | 150 | 3.02E-06 | 7.39E-05 | 1.0,1.0,1.0 | 1.0,0.399,0.571 | 0.343 | ENSG00000119 | [[[[236604422, 2 | 2 |
| ENSG00000188 | RBM34 | chr1 | 235154880 | 235155112 | 235152487 | 235152765 | 235160892 | 235161067 | 48,66,77 | 0,0,0 | 86,181,116 | 3,10,4 | 300 | 150 | 2.08E-05 | 0.000407195 | 1.0,1.0,1.0 | 0.935,0.9,0.935 | 0.077 | ENSG00000188 | [[[[235161174, 2 | 2 |
| ENSG00000188 | RBM34 | chr1 | 235160510 | 235160647 | 235152487 | 235152765 | 235160892 | 235161067 | 50,61,89 | 0,0,0 | 102,169,121 | 3,10,4 | 287 | 150 | 2.52E-05 | 0.000480747 | 1.0,1.0,1.0 | 0.947,0.898,0.94 | 0.071 | ENSG00000188 | [[[[235161174, 2 | 2 |
| ENSG00000154 | TSEN2 | chr3 | 12519058 | 12519197 | 12505153 | 12505231 | 12528887 | 12528924 | 8,19,15 | 0,0,0 | 9,25,7 | 0,9,0 | 289 | 150 | 0.000285335 | 0.003953225 | 1.0,1.0,1.0 | 1.0,0.59,1.0 | 0.137 | ENSG00000154 | [[[[12484480, 12 | 2 |
| ENSG00000010 | ZNF207 | chr17 | 32366664 | 32366757 | 32362913 | 32362984 | 32367771 | 32368014 | 71,47,87 | 0,0,0 | 64,115,85 | 2,4,6 | 243 | 150 | 0.000271275 | 0.003784435 | 1.0,1.0,1.0 | 0.952,0.947,0.89 | 0.068 | ENSG00000010 | [[[[32350157, 32 | 2 |
| ENSG00000010 | ZNF207 | chr17 | 32367771 | 32368014 | 32362913 | 32362984 | 32369294 | 32369454 | 316,295,406 | 0,0,0 | 239,524,332 | 10,19,14 | 300 | 150 | 8.41E-12 | 6.51E-10 | 1.0,1.0,1.0 | 0.923,0.932,0.92 | 0.074 | ENSG00000010 | [[[[32350157, 32 | 3 |

| | | | | | | | | | | | | | | | | | | | | | | |
|---|---|---|---|---|---|---|---|---|---|---|---|---|---|---|---|---|---|---|---|---|---|---|
| ENSG000001682 | RBPJ | chr4 | 26428719 | 26428860 | 26424630 | 26424743 | 26430418 | 26430522 | 16,12,31 | 0,0,0 | 13,35,29 | 0,15,0 | 291 | 150 | 0.000213588 | 0.003085896 | 1.0,1.0,1.0 | 1.0,0.546,1.0 | 0.151 | ENSG000001682 | [[[[26321014, 263 | 2 |
| ENSG000001682 | RBPJ | chr4 | 26428719 | 26428860 | 26424630 | 26424743 | 26430691 | 26430851 | 16,13,31 | 0,0,0 | 13,35,29 | 3,4,3 | 291 | 150 | 5.21E-06 | 0.000120303 | 1.0,1.0,1.0 | 0.691,0.819,0.83 | 0.219 | ENSG000001682 | [[[[26321014, 263 | 3 |
| ENSG000001682 | RBPJ | chr4 | 26429897 | 26430053 | 26424630 | 26424743 | 26430691 | 26431156 | 3,4,3 | 0,0,0 | 1,2,1 | 3,4,3 | 300 | 150 | 1.66E-11 | 1.23E-09 | 1.0,1.0,1.0 | 0.143,0.2,0.143 | 0.838 | ENSG000001682 | [[[[26321014, 263 | 3 |
| ENSG000001682 | RBPJ | chr4 | 26430418 | 26430522 | 26424630 | 26424743 | 26430691 | 26430851 | 10,21,29 | 0,0,0 | 6,49,29 | 3,4,3 | 254 | 150 | 1.93E-05 | 0.000381783 | 1.0,1.0,1.0 | 0.542,0.879,0.85 | 0.243 | ENSG000001682 | [[[[26321014, 263 | 3 |
| ENSG000000518 | MPHOSPH9 | chr12 | 123218375 | 123218499 | 123214743 | 123214834 | 123227462 | 123227616 | 22,6,33 | 0,0,0 | 9,29,17 | 2,5,0 | 274 | 150 | 0.00050148 | 0.006442716 | 1.0,1.0,1.0 | 0.711,0.76,1.0 | 0.176 | ENSG000000518 | [[[[123233075, 12 | 3 |
| ENSG000000518 | MPHOSPH9 | chr12 | 123223037 | 123223127 | 123221371 | 123221723 | 123230260 | 123230522 | 4,3,4 | 0,0,0 | 1,5,1 | 2,4,2 | 240 | 150 | 1.29E-09 | 7.01E-08 | 1.0,1.0,1.0 | 0.238,0.439,0.23 | 0.695 | ENSG000000518 | [[[[123233075, 12 | 2 |
| ENSG000000518 | MPHOSPH9 | chr12 | 123227462 | 123227616 | 123221692 | 123221723 | 123230260 | 123230446 | 5,3,11 | 0,0,0 | 7,17,19 | 2,4,2 | 300 | 150 | 4.66E-06 | 0.00010842 | 1.0,1.0,1.0 | 0.636,0.68,0.826 | 0.286 | ENSG000000518 | [[[[123233075, 12 | 2 |
| ENSG000001683 | ATG4B | chr2 | 241654545 | 241654647 | 241651009 | 241651335 | 241666644 | 241666838 | 4,2,1 | 0,0,0 | 10,10,3 | 1,4,1 | 252 | 150 | 0.000244375 | 0.003452142 | 1.0,1.0,1.0 | 0.856,0.598,0.64 | 0.302 | ENSG000001683 | [[[[241637693, 24 | 4 |
| ENSG000001154 | CCT4 | chr2 | 61876919 | 61877052 | 61876094 | 61876234 | 61883458 | 61883548 | 503,422,566 | 0,0,0 | 293,692,438 | 17,13,11 | 283 | 150 | 1.64E-10 | 1.04E-08 | 1.0,1.0,1.0 | 0.901,0.966,0.95 | 0.059 | ENSG000001154 | [[[[61888381, 618 | 4 |
| ENSG000001365 | ACTL6A | chr3 | 179583352 | 179583448 | 179580639 | 179580701 | 179586545 | 179586632 | 153,150,205 | 0,0,0 | 154,439,200 | 23,48,23 | 246 | 150 | 0 | 0 | 1.0,1.0,1.0 | 0.803,0.848,0.84 | 0.169 | ENSG000001365 | [[[[179562926, 17 | 3 |
| ENSG000001358 | DHX9 | chr1 | 182883319 | 182883368 | 182881284 | 182881425 | 182883519 | 182883635 | 81,77,89 | 0,0,0 | 50,151,75 | 0,6,12 | 199 | 150 | 4.98E-06 | 0.000115496 | 1.0,1.0,1.0 | 1.0,0.95,0.825 | 0.075 | ENSG000001358 | [[[[182839347, 18 | 2 |
| ENSG000001358 | DHX9 | chr1 | 182883519 | 182883635 | 182881263 | 182881425 | 182884612 | 182884813 | 284,277,315 | 0,0,0 | 185,462,226 | 5,9,4 | 266 | 150 | 0.000518227 | 0.006625735 | 1.0,1.0,1.0 | 0.954,0.967,0.97 | 0.036 | ENSG000001358 | [[[[182839347, 18 | 3 |
| ENSG000000627 | APPBP2 | chr17 | 60460662 | 60460787 | 60456295 | 60456381 | 60461993 | 60462061 | 35,16,51 | 2,0,1 | 30,39,30 | 6,17,22 | 275 | 150 | 2.21E-09 | 1.15E-07 | 0.905,1.0,0.965 | 0.732,0.556,0.42 | 0.385 | ENSG000000627 | [[[[60525794, 605 | 2 |
| ENSG000000627 | APPBP2 | chr17 | 60461809 | 60461915 | 60456295 | 60456381 | 60461993 | 60462061 | 17,11,21 | 2,0,1 | 11,26,11 | 6,17,22 | 256 | 150 | 1.12E-08 | 4.95E-07 | 0.833,1.0,0.925 | 0.518,0.473,0.22 | 0.513 | ENSG000000627 | [[[[60525794, 605 | 2 |
| ENSG000001982 | HELZ | chr17 | 67193966 | 67194042 | 67190156 | 67190355 | 67201128 | 67201185 | 1,4,11 | 0,0,0 | 2,3,2 | 2,3,2 | 226 | 150 | 9.88E-08 | 3.47E-06 | 1.0,1.0,1.0 | 0.399,0.399,0.39 | 0.601 | ENSG000001982 | [[[[67245148, 672 | 2 |
| ENSG000001327 | NASP | chr1 | 45604916 | 45605016 | 45591222 | 45591270 | 45606481 | 45606591 | 126,120,219 | 2,1,5 | 80,191,125 | 5,13,11 | 250 | 150 | 2.57E-05 | 0.000488744 | 0.974,0.986,0.96 | 0.906,0.898,0.87 | 0.082 | ENSG000001327 | [[[[45584041, 455 | 2 |
| ENSG000001327 | NASP | chr1 | 45604935 | 45605016 | 45591222 | 45591270 | 45606481 | 45606532 | 127,133,230 | 2,1,5 | 87,208,137 | 5,13,11 | 231 | 150 | 3.80E-05 | 0.000688745 | 0.976,0.989,0.96 | 0.919,0.912,0.89 | 0.071 | ENSG000001327 | [[[[45584041, 455 | 2 |
| ENSG000001327 | NASP | chr1 | 45606481 | 45606591 | 45591222 | 45591270 | 45613168 | 45613248 | 130,112,179 | 0,3,0 | 129,240,102 | 1,18,7 | 260 | 150 | 2.31E-06 | 5.82E-05 | 1.0,0.956,1.0 | 0.987,0.885,0.89 | 0.063 | ENSG000001327 | [[[[45584041, 455 | 4 |
| ENSG000001327 | NASP | chr1 | 45614095 | 45614181 | 45610553 | 45613248 | 45615012 | 45615131 | 231,183,278 | 0,0,0 | 158,328,145 | 5,10,4 | 236 | 150 | 3.41E-05 | 0.000626523 | 1.0,1.0,1.0 | 0.953,0.954,0.95 | 0.045 | ENSG000001327 | [[[[45584041, 455 | 2 |
| ENSG000001377 | MAP2K5 | chr15 | 67748230 | 67748257 | 67703336 | 67703408 | 67748568 | 67748601 | 8,6,12 | 0,0,2 | 5,10,16 | 7,11,7 | 177 | 150 | 7.37E-07 | 2.09E-05 | 1.0,1.0,0.836 | 0.377,0.435,0.66 | 0.455 | ENSG000001377 | [[[[67542703, 675 | 2 |
| ENSG000000766 | NT5C2 | chr10 | 103111744 | 103111805 | 103101044 | 103101102 | 103139405 | 103139479 | 1,3,4 | 0,0,0 | 3,3,3 | 3,9,0 | 211 | 150 | 2.02E-06 | 5.17E-05 | 1.0,1.0,1.0 | 0.416,0.192,1.0 | 0.464 | ENSG000000766 | [[[[103193236, 10 | 3 |
| ENSG000001109 | ATP5F1B | chr12 | 56642672 | 56642831 | 56639979 | 56640192 | 56643402 | 56643587 | 125,411,141,577 | 0,0,0 | 465,914,589 | 12,25,8 | 300 | 150 | 4.55E-12 | 3.76E-10 | 1.0,1.0,1.0 | 0.951,0.948,0.97 | 0.042 | ENSG000001109 | [[[[56645837, 566 | 2 |
| ENSG000001109 | ATP5F1B | chr12 | 56645170 | 56645353 | 56642672 | 56642684 | 56645836 | 56645928 | 42,147,410 | 0,0,0 | 111,230,224 | 4,1,9 | 300 | 150 | 0.000403273 | 0.005334829 | 1.0,1.0,1.0 | 0.933,0.991,0.92 | 0.05 | ENSG000001109 | [[[[56645837, 566 | 4 |
| ENSG000001109 | ATP5F1B | chr12 | 56645170 | 56645353 | 56643402 | 56643587 | 56645836 | 56645928 | 42,147,412 | 0,0,0 | 360,691,547 | 1,17,13 | 300 | 150 | 6.32E-05 | 0.001071921 | 1.0,1.0,1.0 | 0.994,0.953,0.95 | 0.033 | ENSG000001109 | [[[[56645837, 566 | 3 |
| ENSG000001173 | CD46 | chr1 | 207761248 | 207761446 | 207759638 | 207759724 | 207783291 | 207783330 | 124,90,180 | 0,0,0 | 44,97,68 | 4,9,3 | 300 | 150 | 3.15E-08 | 1.25E-06 | 1.0,1.0,1.0 | 0.846,0.843,0.91 | 0.131 | ENSG000001173 | [[[[207752054, 20 | 4 |
| ENSG000001173 | CD46 | chr1 | 207767778 | 207767823 | 207759638 | 207759724 | 207770320 | 207770362 | 12,24,21 | 0,0,0 | 35,81,41 | 20,31,16 | 195 | 150 | 1.39E-12 | 1.24E-10 | 1.0,1.0,1.0 | 0.574,0.668,0.66 | 0.365 | ENSG000001173 | [[[[207752054, 20 | 3 |
| ENSG000001173 | CD46 | chr1 | 207767778 | 207767823 | 207767012 | 207767195 | 207785070 | 207785106 | 8,9,18 | 0,0,0 | 3,12,10 | 2,8,3 | 195 | 150 | 5.20E-09 | 2.47E-07 | 1.0,1.0,1.0 | 0.536,0.536,0.71 | 0.403 | ENSG000001173 | [[[[207752054, 20 | 3 |
| ENSG000001173 | CD46 | chr1 | 207767778 | 207767823 | 207767012 | 207767195 | 207785655 | 207785682 | 8,9,18 | 0,0,0 | 6,11,11 | 3,14,7 | 195 | 150 | 3.16E-11 | 2.22E-09 | 1.0,1.0,1.0 | 0.606,0.377,0.54 | 0.49 | ENSG000001173 | [[[[207752054, 20 | 4 |
| ENSG000001173 | CD46 | chr1 | 207770320 | 207770362 | 207759638 | 207759724 | 207783291 | 207783330 | 26,30,51 | 0,0,0 | 37,119,53 | 4,9,3 | 192 | 150 | 2.21E-05 | 0.000429133 | 1.0,1.0,1.0 | 0.878,0.912,0.93 | 0.093 | ENSG000001173 | [[[[207752054, 20 | 4 |
| ENSG000001173 | CD46 | chr1 | 207770320 | 207770362 | 207767012 | 207767195 | 207783291 | 207783330 | 135,140,222 | 6,5,8 | 71,208,120 | 9,17,15 | 192 | 150 | 0.000333589 | 0.00452373 | 0.946,0.956,0.95 | 0.86,0.905,0.862 | 0.077 | ENSG000001173 | [[[[207752054, 20 | 2 |
| ENSG000001173 | CD46 | chr1 | 207770320 | 207770362 | 207767012 | 207767195 | 207785070 | 207785106 | 24,25,48 | 0,0,0 | 13,37,22 | 2,8,3 | 192 | 150 | 3.67E-07 | 1.12E-05 | 1.0,1.0,1.0 | 0.835,0.783,0.85 | 0.177 | ENSG000001173 | [[[[207752054, 20 | 3 |
| ENSG000001173 | CD46 | chr1 | 207783291 | 207783330 | 207767012 | 207767195 | 207785070 | 207785106 | 64,47,93 | 0,0,0 | 39,79,76 | 2,8,3 | 189 | 150 | 4.65E-05 | 0.000819087 | 1.0,1.0,1.0 | 0.939,0.887,0.95 | 0.074 | ENSG000001173 | [[[[207752054, 20 | 3 |
| ENSG000001448 | BOC | chr3 | 113274374 | 113274682 | 113269381 | 113272703 | 113278094 | 113278257 | 8,4,17 | 0,0,0 | 10,20,20 | 2,6,2 | 300 | 150 | 8.42E-07 | 2.35E-05 | 1.0,1.0,1.0 | 0.714,0.625,0.83 | 0.276 | ENSG000001448 | [[[[113211557, 11 | 2 |
| ENSG000000781 | UBE2K | chr4 | 39737419 | 39737513 | 39698043 | 39698390 | 39755656 | 39755739 | 57,34,94 | 4,5,1 | 38,63,41 | 14,27,16 | 244 | 150 | 1.82E-09 | 9.57E-08 | 0.898,0.807,0.98 | 0.625,0.589,0.61 | 0.287 | ENSG000000781 | [[[[39698136, 396 | 2 |
| ENSG000000781 | UBE2K | chr4 | 39745751 | 39745810 | 39698138 | 39698390 | 39755656 | 39755739 | 35,44,62 | 4,5,1 | 37,53,28 | 14,27,16 | 209 | 150 | 3.60E-09 | 1.77E-07 | 0.863,0.863,0.97 | 0.655,0.585,0.55 | 0.302 | ENSG000000781 | [[[[39698136, 396 | 2 |
| ENSG000000781 | UBE2K | chr4 | 39755656 | 39755739 | 39698162 | 39698390 | 39774833 | 39774933 | 105,81,106 | 0,2,0 | 53,131,80 | 11,18,10 | 233 | 150 | 1.81E-12 | 1.59E-10 | 1.0,0.963,1.0 | 0.756,0.824,0.83 | 0.182 | ENSG000000781 | [[[[39698136, 396 | 3 |
| ENSG000001988 | SELENOT | chr3 | 150623042 | 150623169 | 150603278 | 150603499 | 150624811 | 150624899 | 113,118,189 | 0,0,0 | 75,129,82 | 5,15,5 | 277 | 150 | 5.30E-11 | 3.62E-09 | 1.0,1.0,1.0 | 0.89,0.823,0.899 | 0.129 | ENSG000001988 | [[[[150603321, 15 | 2 |
| ENSG000000541 | TRAPPC3 | chr1 | 36139719 | 36139819 | 36136569 | 36137322 | 36149336 | 36149469 | 0,0,0 | 2,4,3 | 3,3,4 | 3,15,7 | 250 | 150 | 2.32E-05 | 0.000449314 | 0.0,0.0,0.0 | 0.375,0.107,0.25 | -0.246 | ENSG000000541 | [[[[36149337, 361 | 3 |
| ENSG000000541 | TRAPPC3 | chr1 | 36139719 | 36139819 | 36136580 | 36137978 | 36149336 | 36149469 | 275,192,243 | 4,5,6 | 288,602,331 | 8,39,20 | 250 | 150 | 0.000215411 | 0.003109277 | 0.976,0.958,0.96 | 0.956,0.903,0.90 | 0.042 | ENSG000000541 | [[[[36149337, 361 | 2 |
| ENSG000000541 | TRAPPC3 | chr1 | 36140068 | 36140166 | 36137795 | 36137978 | 36149091 | 36149287 | 10,8,13 | 0,0,1 | 15,40,20 | 7,9,5 | 248 | 150 | 2.71E-06 | 6.71E-05 | 1.0,1.0,0.887 | 0.564,0.729,0.70 | 0.295 | ENSG000000541 | [[[[36149337, 361 | 2 |
| ENSG000000993 | BCL7C | chr16 | 30892585 | 30892714 | 30888859 | 30888945 | 30893211 | 30893290 | 48,103,157 | 0,0,0 | 86,150,97 | 13,26,18 | 279 | 150 | 0 | 0 | 1.0,1.0,1.0 | 0.781,0.756,0.74 | 0.24 | ENSG000000993 | [[[[30893853, 308 | 2 |
| ENSG000000993 | BCL7C | chr16 | 30892585 | 30892747 | 30888859 | 30888945 | 30893211 | 30893290 | 52,118,197 | 0,0,0 | 95,164,108 | 13,26,18 | 300 | 150 | 0 | 0 | 1.0,1.0,1.0 | 0.785,0.759,0.75 | 0.235 | ENSG000000993 | [[[[30893853, 308 | 2 |
| ENSG000001593 | ADIPOR1 | chr1 | 202944982 | 202945169 | 202943757 | 202943945 | 202948303 | 202948420 | 175,140,220 | 0,0,0 | 144,280,189 | 19,25,27 | 300 | 150 | 0 | 0 | 1.0,1.0,1.0 | 0.791,0.848,0.77 | 0.194 | ENSG000001593 | [[[[202958185, 20 | 2 |
| ENSG000001593 | ADIPOR1 | chr1 | 202946438 | 202946610 | 202943757 | 202943863 | 202948303 | 202948420 | 164,126,261 | 0,0,0 | 152,265,184 | 29,24,25 | 300 | 150 | 0 | 0 | 1.0,1.0,1.0 | 0.724,0.847,0.78 | 0.214 | ENSG000001593 | [[[[202958185, 20 | 2 |
| ENSG000001593 | ADIPOR1 | chr1 | 202946438 | 202946610 | 202943757 | 202943945 | 202948303 | 202948420 | 162,127,261 | 0,0,0 | 152,268,181 | 19,25,27 | 300 | 150 | 0 | 0 | 1.0,1.0,1.0 | 0.8,0.843,0.77 | 0.196 | ENSG000001593 | [[[[202958185, 20 | 2 |
| ENSG000001328 | SERINC3 | chr20 | 44512800 | 44512994 | 44509890 | 44510028 | 44513878 | 44514040 | 74,62,112 | 0,0,0 | 53,128,93 | 6,22,4 | 300 | 150 | 1.19E-10 | 7.72E-09 | 1.0,1.0,1.0 | 0.815,0.744,0.92 | 0.173 | ENSG000001328 | [[[[44521913, 445 | 2 |
| ENSG000000698 | ATP1B3 | chr3 | 141903619 | 141903748 | 141876613 | 141876910 | 141913651 | 141913706 | 142,131,173 | 0,0,0 | 181,455,294 | 6,11,2 | 279 | 150 | 0.000837398 | 0.009990296 | 1.0,1.0,1.0 | 0.942,0.957,0.98 | 0.038 | ENSG000000698 | [[[[141876643, 14 | 2 |
| ENSG000000698 | ATP1B3 | chr3 | 141913651 | 141913836 | 141907166 | 141907274 | 141921976 | 141922063 | 227,204,300 | 0,1,0 | 408,800,556 | 6,13,9 | 300 | 150 | 0.000595676 | 0.007469035 | 1.0,0.99,1.0 | 0.971,0.969,0.96 | 0.027 | ENSG000000698 | [[[[141876643, 14 | 2 |
| ENSG000001987 | IPO9 | chr1 | 201847278 | 201847340 | 201829140 | 201829372 | 201852103 | 201852192 | 24,33,71 | 0,0,0 | 16,39,22 | 3,3,2 | 212 | 150 | 0.000254746 | 0.003580063 | 1.0,1.0,1.0 | 0.791,0.902,0.88 | 0.14 | ENSG000001987 | [[[[201829157, 20 | 3 |
| ENSG000001987 | IPO9 | chr1 | 201866732 | 201866959 | 201858854 | 201858994 | 201868647 | 201868796 | 123,104,182 | 0,0,0 | 28,46,31 | 17,36,23 | 300 | 150 | 0 | 0 | 1.0,1.0,1.0 | 0.452,0.39,0.403 | 0.585 | ENSG000001987 | [[[[201829157, 20 | 2 |
| ENSG000001987 | IPO9 | chr1 | 201870582 | 201870858 | 201858854 | 201858994 | 201871160 | 201871327 | 130,111,224 | 0,0,0 | 77,173,89 | 2,5,14 | 300 | 150 | 2.77E-08 | 1.12E-06 | 1.0,1.0,1.0 | 0.951,0.945,0.76 | 0.114 | ENSG000001987 | [[[[201829157, 20 | 5 |
| ENSG000001652 | CARD19 | chr9 | 93110567 | 93110721 | 93107673 | 93107827 | 93112217 | 93112289 | 14,29,60 | 0,0,0 | 57,84,68 | 3,3,5 | 300 | 150 | 0.000238983 | 0.003388549 | 1.0,1.0,1.0 | 0.905,0.933,0.87 | 0.097 | ENSG000001652 | [[[[93096217, 930 | 2 |
| ENSG000001652 | CARD19 | chr9 | 93111878 | 93111938 | 93107673 | 93107816 | 93112217 | 93112289 | 3,7,20 | 0,0,0 | 3,17,21 | 1,7,0 | 210 | 150 | 0.000155799 | 0.002340145 | 1.0,1.0,1.0 | 0.682,0.634,1.0 | 0.228 | ENSG000001652 | [[[[93096217, 930 | 2 |
| ENSG000001652 | CARD19 | chr9 | 93111878 | 93111938 | 93107673 | 93107827 | 93112217 | 93112289 | 3,8,20 | 0,0,0 | 3,17,26 | 3,3,5 | 210 | 150 | 1.89E-06 | 4.87E-05 | 1.0,1.0,1.0 | 0.417,0.802,0.78 | 0.331 | ENSG000001652 | [[[[93096217, 930 | 2 |
| ENSG000001287 | SNRPN | chr15 | 24974386 | 24974456 | 24967931 | 24968082 | 24976876 | 24977029 | 1,1,1 | 0,0,0 | 2,8,6 | 3,3,2 | 220 | 150 | 4.05E-07 | 1.22E-05 | 1.0,1.0,1.0 | 0.313,0.645,0.67 | 0.457 | ENSG000001287 | [[[[24954987, 249 | 3 |
| ENSG000001181 | KIF14 | chr1 | 200560721 | 200560880 | 200559329 | 200559452 | 200569910 | 200570005 | 35,23,38 | 0,0,0 | 11,30,28 | 3,2,5 | 300 | 150 | 1.49E-06 | 3.92E-05 | 1.0,1.0,1.0 | 0.647,0.882,0.73 | 0.245 | ENSG000001181 | [[[[200620411, 20 | 3 |
| ENSG000001181 | KIF14 | chr1 | 200565444 | 200565669 | 200559329 | 200559452 | 200569910 | 200570005 | 18,18,29 | 0,0,0 | 14,28,22 | 3,2,5 | 300 | 150 | 1.56E-06 | 4.09E-05 | 1.0,1.0,1.0 | 0.7,0.875,0.688 | 0.246 | ENSG000001181 | [[[[200620411, 20 | 3 |
| ENSG000002042 | COL5A2 | chr2 | 189066728 | 189066782 | 189066389 | 189066497 | 189068785 | 189068884 | 59,42,119 | 0,0,0 | 53,84,66 | 0,2,9 | 204 | 150 | 0.00052371 | 0.006684368 | 1.0,1.0,1.0 | 1.0,0.969,0.844 | 0.062 | ENSG000002042 | [[[[189179508, 18 | 3 |
| ENSG000001167 | UCHL5 | chr1 | 193029178 | 193029309 | 193023843 | 193023946 | 193029387 | 193029449 | 71,38,77 | 0,0,0 | 44,118,61 | 2,2,8 | 281 | 150 | 4.79E-05 | 0.000840513 | 1.0,1.0,1.0 | 0.922,0.969,0.80 | 0.102 | ENSG000001167 | [[[[193059185, 19 | 2 |
| ENSG000001326 | ARHGEF11 | chr1 | 156979228 | 156979286 | 156978203 | 156978382 | 156984338 | 156984437 | 2,2,7 | 0,0,0 | 8,23,12 | 0,6,1 | 208 | 150 | 0.000737619 | 0.008949262 | 1.0,1.0,1.0 | 1.0,0.734,0.896 | 0.123 | ENSG000001326 | [[[[157044299, 15 | 2 |
| ENSG000001619 | SENP3 | chr17 | 7566926 | 7567004 | 7565694 | 7565764 | 7571372 | 7571461 | 94,56,112 | 0,0,0 | 71,124,82 | 4,12,4 | 228 | 150 | 1.86E-07 | 6.07E-06 | 1.0,1.0,1.0 | 0.921,0.872,0.93 | 0.092 | ENSG000001619 | [[[[7561919, 7562 | 4 |
| ENSG000001619 | SENP3 | chr17 | 7570680 | 7570764 | 7570355 | 7570493 | 7571372 | 7571686 | 104,93,122 | 0,0,6 | 109,210,125 | 29,36,18 | 234 | 150 | 3.01E-13 | 2.93E-11 | 1.0,1.0,0.929 | 0.707,0.789,0.81 | 0.205 | ENSG000001619 | [[[[7561919, 7562 | 2 |
| ENSG000001211 | NCAPH | chr2 | 96365875 | 96366058 | 96364480 | 96364591 | 96368971 | 96369063 | 48,56,78 | 0,0,0 | 36,57,35 | 2,5,4 | 300 | 150 | 5.76E-06 | 0.000131507 | 1.0,1.0,1.0 | 0.9,0.851,0.814 | 0.145 | ENSG000001211 | [[[[96335766, 963 | 2 |
| ENSG000001211 | NCAPH | chr2 | 96365968 | 96366058 | 96364479 | 96364591 | 96368971 | 96369063 | 4,5,15 | 0,0,0 | 13,22,12 | 2,5,4 | 240 | 150 | 9.61E-07 | 2.66E-05 | 1.0,1.0,1.0 | 0.802,0.733,0.65 | 0.271 | ENSG000001211 | [[[[96335766, 963 | 2 |
| ENSG000002661 | STRADA | chr17 | 63713405 | 63713527 | 63710490 | 63710836 | 63728333 | 63728413 | 39,22,49 | 0,0,0 | 25,36,39 | 3,3,2 | 272 | 150 | 0.000296815 | 0.004093517 | 1.0,1.0,1.0 | 0.821,0.869,0.91 | 0.132 | ENSG000002661 | [[[[63741741, 637 | 4 |
| ENSG000002661 | STRADA | chr17 | 63714005 | 63714108 | 63710490 | 63710836 | 63728333 | 63728404 | 7,1,15 | 0,0,0 | 1,7,7 | 3,3,2 | 253 | 150 | 4.63E-08 | 1.76E-06 | 1.0,1.0,1.0 | 0.165,0.58,0.675 | 0.527 | ENSG000002661 | [[[[63741741, 637 | 4 |
| ENSG000001152 | EIF2B4 | chr2 | 27367456 | 27367559 | 27367073 | 27367201 | 27368024 | 27368113 | 55,52,62 | 0,0,0 | 47,91,62 | 4,6,1 | 253 | 150 | 0.000188133 | 0.002767975 | 1.0,1.0,1.0 | 0.874,0.9,0.974 | 0.084 | ENSG000001152 | [[[[27370284, 273 | 2 |
| ENSG000001152 | EIF2B4 | chr2 | 27369413 | 27369549 | 27368371 | 27368463 | 27369875 | 27369919 | 24,30,82 | 0,0,0 | 32,59,79 | 2,3,7 | 286 | 150 | 2.37E-05 | 0.000455838 | 1.0,1.0,1.0 | 0.894,0.912,0.85 | 0.113 | ENSG000001152 | [[[[27370284, 273 | 3 |
| ENSG000001763 | RNPEP | chr1 | 201997318 | 201997554 | 201996146 | 201996263 | 202001375 | 202001488 | 198,181,333 | 0,0,0 | 90,156,94 | 3,7,2 | 300 | 150 | 0.00010831 | 0.001710108 | 1.0,1.0,1.0 | 0.938,0.918,0.95 | 0.062 | ENSG000001763 | [[[[201982648, 20 | 2 |
| ENSG000002133 | CHUK | chr10 | 100209594 | 100209789 | 100207229 | 100207332 | 100219269 | 100219359 | 36,22,35 | 0,0,0 | 17,24,24 | 2,6,1 | 300 | 150 | 4.23E-06 | 9.97E-05 | 1.0,1.0,1.0 | 0.81,0.667,0.923 | 0.2 | ENSG000002133 | [[[[100229428, 10 | 4 |
| ENSG000000572 | SOAT1 | chr1 | 179347599 | 179347697 | 179344946 | 179345076 | 179353584 | 179358677 | 44,28,55 | 0,0,0 | 25,58,22 | 0,3,9 | 248 | 150 | 2.10E-05 | 0.000410799 | 1.0,1.0,1.0 | 1.0,0.921,0.597 | 0.161 | ENSG000000572 | [[[[179293797, 17 | 4 |
| ENSG000001649 | INTS8 | chr8 | 94838462 | 94838618 | 94836523 | 94836631 | 94849461 | 94849532 | 31,21,35 | 0,0,0 | 29,56,17 | 3,6,1 | 300 | 150 | 2.15E-05 | 0.00041902 | 1.0,1.0,1.0 | 0.829,0.824,0.89 | 0.151 | ENSG000001649 | [[[[94823287, 948 | 3 |
| ENSG000001649 | INTS8 | chr8 | 94841490 | 94841591 | 94838462 | 94838618 | 94849461 | 94849532 | 24,22,32 | 0,0,0 | 3,31,13 | 2,8,0 | 251 | 150 | 5.09E-05 | 0.000886114 | 1.0,1.0,1.0 | 0.473,0.698,1.0 | 0.276 | ENSG000001649 | [[[[94823287, 948 | 2 |
| ENSG000001876 | B3GLCT | chr13 | 31286719 | 31286819 | 31276701 | 31276771 | 31317565 | 31317685 | 19,22,39 | 0,0,0 | 15,16,9 | 7,18,8 | 250 | 150 | 3.33E-16 | 4.71E-14 | 1.0,1.0,1.0 | 0.563,0.348,0.40 | 0.562 | ENSG000001876 | [[[[31199975, 312 | 2 |
| ENSG000001138 | SMC4 | chr3 | 160400821 | 160400965 | 160399303 | 160399749 | 160411919 | 160412084 | 1,4,5 | 0,0,0 | 0,9,6 | 10,39,17 | 294 | 150 | 9.88E-12 | 7.57E-10 | 1.0,1.0,1.0 | 0.0,0.105,0.153 | 0.914 | ENSG000001138 | [[[[160399650, 16 | 4 |
| ENSG000001138 | SMC4 | chr3 | 160402675 | 160402867 | 160401914 | 160402093 | 160411919 | 160411964 | 78,42,133 | 0,0,0 | 13,36,26 | 7,22,11 | 300 | 150 | 0 | 0 | 1.0,1.0,1.0 | 0.481,0.45,0.542 | 0.509 | ENSG000001138 | [[[[160399650, 16 | 2 |
| ENSG000001138 | SMC4 | chr3 | 160404327 | 160404504 | 160399303 | 160399749 | 160411919 | 160411964 | 94,57,172 | 0,0,0 | 21,73,72 | 10,39,17 | 300 | 150 | 0 | 0 | 1.0,1.0,1.0 | 0.512,0.483,0.67 | 0.442 | ENSG000001138 | [[[[160399650, 16 | 4 |
| ENSG000001138 | SMC4 | chr3 | 160404327 | 160404504 | 160401914 | 160402093 | 160411919 | 160411964 | 95,57,172 | 0,0,0 | 18,73,72 | 7,22,11 | 300 | 150 | 0 | 0 | 1.0,1.0,1.0 | 0.563,0.624,0.76 | 0.349 | ENSG000001138 | [[[[160399650, 16 | 2 |
| ENSG000001138 | SMC4 | chr3 | 160404377 | 160404504 | 160399642 | 160399697 | 160411919 | 160412084 | 89,51,149 | 0,2,0 | 19,70,65 | 2,4,7 | 277 | 150 | 6.18E-06 | 0.000140339 | 1.0,0.932,1.0 | 0.837,0.905,0.83 | 0.119 | ENSG000001138 | [[[[160399650, 16 | 4 |
| ENSG000001138 | SMC4 | chr3 | 160404377 | 160404504 | 160399668 | 160399749 | 160411919 | 160412084 | 89,51,149 | 0,0,0 | 17,76,68 | 10,39,17 | 277 | 150 | 0 | 0 | 1.0,1.0,1.0 | 0.479,0.513,0.68 | 0.441 | ENSG000001138 | [[[[160399650, 16 | 4 |

| | | | | | | | | | | | | | | | | | | | | | | |
|---|---|---|---|---|---|---|---|---|---|---|---|---|---|---|---|---|---|---|---|---|---|---|
| ENSG00000113 | SMC4 | chr3 | 160404377 | 160404504 | 160401989 | 160402093 | 160411919 | 160412084 | 89,51,149 | 0,0,0 | 19,84,73 | 7,22,11 | 277 | 150 | 1.11E-16 | 1.65E-14 | 1.0,1.0,1.0 | 0.595,0.674,0.78 | 0.316 | ENSG00000113 | [[[[160399650, 1 | 2 |
| ENSG00000113 | SMC4 | chr3 | 160412325 | 160412453 | 160411919 | 160411977 | 160414366 | 160414515 | 2,3,1 | 0,0,0 | 3,6,19 | 3,9,6 | 278 | 150 | 5.77E-09 | 2.71E-07 | 1.0,1.0,1.0 | 0.35,0.265,0.631 | 0.585 | ENSG00000113 | [[[[160399650, 1 | 2 |
| ENSG00000113 | SMC4 | chr3 | 160413472 | 160413613 | 160399303 | 160399749 | 160414366 | 160414500 | 62,47,78 | 0,0,0 | 19,60,18 | 5,4,5 | 291 | 150 | 1.63E-08 | 6.94E-07 | 1.0,1.0,1.0 | 0.662,0.885,0.65 | 0.268 | ENSG00000113 | [[[[160399650, 1 | 7 |
| ENSG00000113 | SMC4 | chr3 | 160413472 | 160413613 | 160411919 | 160411977 | 160414366 | 160414500 | 62,47,78 | 0,0,0 | 19,59,20 | 3,9,6 | 291 | 150 | 1.50E-10 | 9.55E-09 | 1.0,1.0,1.0 | 0.766,0.772,0.63 | 0.277 | ENSG00000113 | [[[[160399650, 1 | 2 |
| ENSG00000113 | SMC4 | chr3 | 160419357 | 160419543 | 160416250 | 160416415 | 160420739 | 160420901 | 179,150,276 | 0,0,0 | 38,183,84 | 5,3,3 | 300 | 150 | 0.000252131 | 0.003547894 | 1.0,1.0,1.0 | 0.792,0.968,0.93 | 0.102 | ENSG00000113 | [[[[160399650, 1 | 2 |
| ENSG00000076 | MCM6 | chr2 | 135866131 | 135866277 | 135865012 | 135865163 | 135868610 | 135868860 | 113,80,148 | 0,0,0 | 47,82,53 | 2,8,4 | 296 | 150 | 2.35E-07 | 7.50E-06 | 1.0,1.0,1.0 | 0.923,0.839,0.87 | 0.123 | ENSG00000076 | [[[[135876259, 1 | 2 |
| ENSG00000164 | SHROOM1 | chr5 | 132823849 | 132824419 | 132823622 | 132823764 | 132825017 | 132825073 | 3,14,37 | 0,0,0 | 23,46,24 | 6,11,5 | 300 | 150 | 7.77E-09 | 3.54E-07 | 1.0,1.0,1.0 | 0.657,0.676,0.70 | 0.32 | ENSG00000164 | [[[[132830594, 1 | 2 |
| ENSG00000164 | SHROOM1 | chr5 | 132824614 | 132824821 | 132823622 | 132823764 | 132825017 | 132825073 | 18,14,29 | 0,0,0 | 11,43,25 | 6,11,5 | 300 | 150 | 2.23E-09 | 1.15E-07 | 1.0,1.0,1.0 | 0.478,0.662,0.71 | 0.382 | ENSG00000164 | [[[[132830594, 1 | 2 |
| ENSG00000107 | CREB3 | chr9 | 35735305 | 35735374 | 35732995 | 35733143 | 35736047 | 35736132 | 99,62,114 | 0,0,0 | 122,199,109 | 8,8,0 | 219 | 150 | 0.000228954 | 0.003269457 | 1.0,1.0,1.0 | 0.913,0.945,1.0 | 0.047 | ENSG00000107 | [[[[35732666, 35 | 4 |
| ENSG00000136 | GPNMB | chr7 | 23256891 | 23257065 | 23246562 | 23246927 | 23259979 | 23260053 | 105,79,100 | 0,0,0 | 269,352,295 | 8,14,6 | 300 | 150 | 1.98E-05 | 0.000389816 | 1.0,1.0,1.0 | 0.944,0.926,0.96 | 0.056 | ENSG00000136 | [[[[23246775, 23 | 3 |
| ENSG00000117 | PRRC2C | chr1 | 171579353 | 171579466 | 171577433 | 171577637 | 171587002 | 171587221 | 71,75,107 | 0,0,0 | 100,193,135 | 5,6,4 | 263 | 150 | 0.000216763 | 0.003125829 | 1.0,1.0,1.0 | 0.919,0.948,0.95 | 0.061 | ENSG00000117 | [[[[171485530, 1 | 4 |
| ENSG00000117 | PRRC2C | chr1 | 171584418 | 171584526 | 171579353 | 171579466 | 171587002 | 171587221 | 60,42,77 | 0,0,0 | 53,126,80 | 9,7,6 | 258 | 150 | 1.20E-07 | 4.11E-06 | 1.0,1.0,1.0 | 0.774,0.913,0.88 | 0.142 | ENSG00000117 | [[[[171485530, 1 | 3 |
| ENSG00000117 | PRRC2C | chr1 | 171584418 | 171584541 | 171579353 | 171579466 | 171587002 | 171587221 | 17,21,33 | 0,0,0 | 30,42,19 | 9,7,6 | 273 | 150 | 2.04E-09 | 1.07E-07 | 1.0,1.0,1.0 | 0.647,0.767,0.63 | 0.317 | ENSG00000117 | [[[[171485530, 1 | 3 |
| ENSG00000117 | PRRC2C | chr1 | 171587002 | 171587221 | 171579353 | 171579466 | 171587647 | 171587751 | 92,91,125 | 0,0,0 | 97,220,130 | 5,10,1 | 300 | 150 | 4.05E-05 | 0.000728375 | 1.0,1.0,1.0 | 0.907,0.917,0.98 | 0.064 | ENSG00000117 | [[[[171485530, 1 | 4 |
| ENSG00000105 | DNAJB6 | chr7 | 157382245 | 157382377 | 157367372 | 157367483 | 157416015 | 157417433 | 180,129,286 | 0,0,0 | 269,596,333 | 8,14,4 | 282 | 150 | 5.98E-05 | 0.001020926 | 1.0,1.0,1.0 | 0.947,0.958,0.97 | 0.039 | ENSG00000105 | [[[[157337004, 1 | 4 |
| ENSG00000105 | DNAJB6 | chr7 | 157409794 | 157410001 | 157367372 | 157367483 | 157416015 | 157416475 | 11,12,31 | 0,0,0 | 21,53,57 | 8,14,4 | 300 | 150 | 7.36E-08 | 2.67E-06 | 1.0,1.0,1.0 | 0.568,0.654,0.87 | 0.3 | ENSG00000105 | [[[[157337004, 1 | 4 |
| ENSG00000105 | DUS4L | chr7 | 107571144 | 107571266 | 107567049 | 107567186 | 107575187 | 107575310 | 12,20,11 | 0,0,1 | 11,21,18 | 3,7,4 | 272 | 150 | 1.49E-06 | 3.92E-05 | 1.0,1.0,0.858 | 0.669,0.623,0.71 | 0.284 | ENSG00000105 | [[[[107563971, 1 | 2 |
| ENSG00000198 | OPA1 | chr3 | 193617177 | 193617285 | 193615673 | 193615770 | 193618868 | 193618936 | 93,81,145 | 1,1,6 | 37,69,74 | 4,26,20 | 258 | 150 | 1.50E-10 | 9.55E-09 | 0.982,0.979,0.93 | 0.843,0.607,0.68 | 0.254 | ENSG00000198 | [[[[193593208, 1 | 2 |
| ENSG00000106 | SSBP1 | chr7 | 141745495 | 141745584 | 141742168 | 141742229 | 141750310 | 141750314 | 129,117,129 | 0,0,0 | 139,296,121 | 5,13,9 | 239 | 150 | 2.11E-07 | 6.82E-06 | 1.0,1.0,1.0 | 0.946,0.935,0.89 | 0.075 | ENSG00000106 | [[[[141738358, 1 | 3 |
| ENSG00000143 | TIPRL | chr1 | 168183901 | 168184081 | 168178932 | 168179181 | 168196546 | 168196642 | 92,102,152 | 0,0,0 | 98,246,184 | 14,12,6 | 300 | 150 | 5.99E-09 | 2.80E-07 | 1.0,1.0,1.0 | 0.778,0.911,0.93 | 0.124 | ENSG00000143 | [[[[168178962, 1 | 3 |
| ENSG00000143 | TIPRL | chr1 | 168191368 | 168191500 | 168178932 | 168179181 | 168196546 | 168196642 | 102,99,131 | 0,0,0 | 99,243,100 | 14,12,6 | 282 | 150 | 8.63E-10 | 4.85E-08 | 1.0,1.0,1.0 | 0.79,0.915,0.899 | 0.132 | ENSG00000143 | [[[[168178962, 1 | 3 |
| ENSG00000058 | NDC1 | chr1 | 53787157 | 53787258 | 53772328 | 53772489 | 53796688 | 53796804 | 50,53,87 | 0,0,0 | 78,182,87 | 7,9,5 | 251 | 150 | 1.92E-06 | 4.93E-05 | 1.0,1.0,1.0 | 0.869,0.924,0.91 | 0.098 | ENSG00000058 | [[[[53838205, 53 | 3 |
| ENSG00000058 | NDC1 | chr1 | 53796898 | 53797144 | 53772328 | 53772489 | 53800692 | 53800848 | 79,52,95 | 0,0,0 | 84,175,89 | 7,6,3 | 300 | 150 | 1.31E-05 | 0.000270189 | 1.0,1.0,1.0 | 0.857,0.936,0.93 | 0.09 | ENSG00000058 | [[[[53838205, 53 | 5 |
| ENSG00000058 | NDC1 | chr1 | 53818970 | 53819079 | 53809694 | 53809746 | 53832489 | 53832591 | 30,26,51 | 0,0,0 | 22,99,66 | 3,11,2 | 259 | 150 | 4.83E-06 | 0.000112076 | 1.0,1.0,1.0 | 0.809,0.839,0.95 | 0.134 | ENSG00000058 | [[[[53838205, 53 | 3 |
| ENSG00000058 | NDC1 | chr1 | 53825797 | 53825936 | 53818969 | 53819079 | 53832489 | 53832591 | 49,53,77 | 0,0,0 | 25,83,60 | 3,15,7 | 289 | 150 | 8.69E-11 | 5.76E-09 | 1.0,1.0,1.0 | 0.812,0.742,0.81 | 0.21 | ENSG00000058 | [[[[53838205, 53 | 2 |
| ENSG00000058 | NDC1 | chr1 | 53827998 | 53828173 | 53809694 | 53809746 | 53832489 | 53832591 | 33,27,63 | 0,0,0 | 36,57,44 | 3,11,2 | 300 | 150 | 3.14E-07 | 9.76E-06 | 1.0,1.0,1.0 | 0.857,0.722,0.91 | 0.168 | ENSG00000058 | [[[[53838205, 53 | 3 |
| ENSG00000118 | SPP1 | chr4 | 87980392 | 87980434 | 87977058 | 87977097 | 87981474 | 87981535 | 196,119,257 | 2,2,1 | 243,435,330 | 8,14,18 | 192 | 150 | 1.74E-05 | 0.000347008 | 0.987,0.979,0.99 | 0.96,0.96,0.935 | 0.035 | ENSG00000118 | [[[[87975714, 87 | 2 |
| ENSG00000249 | PDCD6 | chr5 | 313700 | 313850 | 304176 | 304221 | 314416 | 314954 | 2,7,16 | 3,4,8 | 1,3,1 | 15,28,8 | 300 | 150 | 4.78E-08 | 1.82E-06 | 0.25,0.467,0.5 | 0.032,0.051,0.05 | 0.358 | ENSG00000249 | [[[[271646, 2718 | 2 |
| ENSG00000089 | ANAPC5 | chr12 | 121320384 | 121320459 | 121318276 | 121318424 | 121327095 | 121327231 | 27,59,95 | 0,0,0 | 58,90,59 | 10,22,8 | 225 | 150 | 2.75E-13 | 2.70E-11 | 1.0,1.0,1.0 | 0.795,0.732,0.83 | 0.214 | ENSG00000089 | [[[[121352134, 1 | 3 |
| ENSG00000138 | SLC5A6 | chr2 | 27202812 | 27202880 | 27201987 | 27202074 | 27204460 | 27204590 | 13,13,23 | 0,0,0 | 26,61,36 | 6,7,8 | 218 | 150 | 4.27E-07 | 1.29E-05 | 1.0,1.0,1.0 | 0.749,0.857,0.75 | 0.213 | ENSG00000138 | [[[[27212020, 27 | 3 |
| ENSG00000138 | SLC5A6 | chr2 | 27203778 | 27203867 | 27202812 | 27202880 | 27204460 | 27204590 | 23,19,40 | 0,0,0 | 18,47,34 | 1,21,13 | 239 | 150 | 1.16E-09 | 6.36E-08 | 1.0,1.0,1.0 | 0.919,0.584,0.62 | 0.292 | ENSG00000138 | [[[[27212020, 27 | 2 |
| ENSG00000138 | SLC5A6 | chr2 | 27206482 | 27206534 | 27204790 | 27204931 | 27206876 | 27206942 | 13,13,16 | 0,0,0 | 8,18,16 | 0,6,9 | 202 | 150 | 5.06E-07 | 1.50E-05 | 1.0,1.0,1.0 | 1.0,0.69,0.569 | 0.247 | ENSG00000138 | [[[[27212020, 27 | 3 |
| ENSG00000158 | NIT1 | chr1 | 161120106 | 161120232 | 161119133 | 161119388 | 161120498 | 161120568 | 39,30,50 | 0,0,0 | 54,108,44 | 4,4,2 | 276 | 150 | 0.000618063 | 0.007706623 | 1.0,1.0,1.0 | 0.88,0.936,0.923 | 0.087 | ENSG00000158 | [[[[161118105, 1 | 3 |
| ENSG00000109 | CLCN3 | chr4 | 169692113 | 169692320 | 169690529 | 169690652 | 169697360 | 169697734 | 37,20,44 | 0,0,0 | 29,37,32 | 2,7,6 | 300 | 150 | 6.94E-08 | 2.54E-06 | 1.0,1.0,1.0 | 0.879,0.725,0.72 | 0.223 | ENSG00000109 | [[[[169620578, 1 | 2 |
| ENSG00000103 | NPRL3 | chr16 | 98144 | 98301 | 93218 | 93325 | 110524 | 110606 | 47,37,61 | 0,0,0 | 56,66,50 | 2,8,1 | 300 | 150 | 3.96E-05 | 0.00071379 | 1.0,1.0,1.0 | 0.933,0.805,0.96 | 0.1 | ENSG00000103 | [[[[138642, 1386 | 2 |
| ENSG00000122 | COPA | chr1 | 160313084 | 160313167 | 160311867 | 160312018 | 160323430 | 160323530 | 86,43,89 | 0,0,0 | 33,74,57 | 6,4,4 | 233 | 150 | 1.26E-06 | 3.37E-05 | 1.0,1.0,1.0 | 0.78,0.923,0.902 | 0.132 | ENSG00000122 | [[[[160343131, 1 | 2 |
| ENSG00000122 | COPA | chr1 | 160333602 | 160333679 | 160332447 | 160332557 | 160339908 | 160339982 | 81,57,115 | 0,0,0 | 26,65,71 | 12,15,4 | 227 | 150 | 9.20E-12 | 7.08E-10 | 1.0,1.0,1.0 | 0.589,0.741,0.92 | 0.25 | ENSG00000122 | [[[[160343131, 1 | 2 |
| ENSG00000122 | COPA | chr1 | 160335241 | 160335322 | 160332447 | 160332557 | 160339908 | 160339982 | 58,43,105 | 0,0,0 | 42,77,81 | 12,15,4 | 231 | 150 | 8.06E-11 | 5.37E-09 | 1.0,1.0,1.0 | 0.694,0.769,0.92 | 0.203 | ENSG00000122 | [[[[160343131, 1 | 2 |
| ENSG00000157 | BRAF | chr7 | 140753274 | 140754233 | 140749286 | 140749418 | 140776911 | 140777088 | 8,4,8 | 0,0,0 | 10,17,20 | 1,2,4 | 300 | 150 | 0.000119258 | 0.00185833 | 1.0,1.0,1.0 | 0.833,0.81,0.714 | 0.214 | ENSG00000157 | [[[[140924566, 1 | 2 |
| ENSG00000065 | OAT | chr10 | 124400839 | 124400984 | 124397302 | 124398102 | 124403797 | 124403920 | 184,197,307 | 0,0,0 | 194,352,248 | 3,8,5 | 295 | 150 | 0.000815646 | 0.00976596 | 1.0,1.0,1.0 | 0.97,0.957,0.962 | 0.037 | ENSG00000065 | [[[[124418873, 1 | 3 |
| ENSG00000065 | OAT | chr10 | 124408541 | 124408637 | 124405435 | 124405563 | 124418872 | 124418936 | 148,127,229 | 0,0,0 | 121,228,156 | 10,7,6 | 246 | 150 | 2.19E-07 | 7.04E-06 | 1.0,1.0,1.0 | 0.881,0.952,0.94 | 0.075 | ENSG00000065 | [[[[124418873, 1 | 3 |
| ENSG00000065 | OAT | chr10 | 124408740 | 124408965 | 124405435 | 124405563 | 124418872 | 124418976 | 4,5,2 | 0,0,0 | 5,13,9 | 10,7,6 | 300 | 150 | 3.78E-09 | 1.85E-07 | 1.0,1.0,1.0 | 0.2,0.481,0.429 | 0.63 | ENSG00000065 | [[[[124418873, 1 | 3 |
| ENSG00000065 | OAT | chr10 | 124411972 | 124412200 | 124405435 | 124405563 | 124418872 | 124418936 | 172,126,299 | 0,0,0 | 148,311,222 | 10,7,6 | 300 | 150 | 7.07E-07 | 2.02E-05 | 1.0,1.0,1.0 | 0.881,0.957,0.94 | 0.071 | ENSG00000065 | [[[[124418873, 1 | 3 |
| ENSG00000008 | PSMB1 | chr6 | 170546102 | 170546184 | 170537233 | 170537340 | 170553129 | 170553341 | 4,5,4 | 0,0,0 | 7,12,13 | 3,8,9 | 232 | 150 | 1.42E-08 | 6.14E-07 | 1.0,1.0,1.0 | 0.601,0.492,0.48 | 0.475 | ENSG00000008 | [[[[170553130, 1 | 3 |
| ENSG00000158 | USF1 | chr1 | 161041650 | 161041846 | 161041323 | 161041411 | 161042585 | 161042670 | 73,97,143 | 0,0,0 | 177,314,182 | 8,6,2 | 300 | 150 | 0.000736859 | 0.008943486 | 1.0,1.0,1.0 | 0.917,0.963,0.97 | 0.047 | ENSG00000158 | [[[[161045858, 1 | 2 |
| ENSG00000114 | NKTR | chr3 | 42636867 | 42639750 | 42632600 | 42632823 | 42642500 | 42642596 | 39,30,58 | 0,0,0 | 29,70,58 | 5,11,1 | 300 | 150 | 4.98E-07 | 1.48E-05 | 1.0,1.0,1.0 | 0.744,0.761,0.96 | 0.176 | ENSG00000114 | [[[[42600686, 42 | 4 |
| ENSG00000168 | LRRC28 | chr15 | 99361335 | 99361511 | 99352368 | 99352471 | 99386029 | 99386296 | 37,24,28 | 0,0,0 | 26,41,23 | 8,2,3 | 300 | 150 | 5.19E-07 | 1.53E-05 | 1.0,1.0,1.0 | 0.619,0.911,0.79 | 0.226 | ENSG00000168 | [[[[99251481, 99 | 2 |
| ENSG00000082 | RNF13 | chr3 | 149911977 | 149912083 | 149895472 | 149895560 | 149921133 | 149921184 | 11,11,16 | 0,0,0 | 17,21,35 | 11,13,10 | 256 | 150 | 2.70E-10 | 1.65E-08 | 1.0,1.0,1.0 | 0.475,0.486,0.67 | 0.456 | ENSG00000082 | [[[[149813215, 1 | 2 |
| ENSG00000082 | RNF13 | chr3 | 149911977 | 149912083 | 149895472 | 149895570 | 149921133 | 149921184 | 11,11,16 | 0,0,0 | 16,23,35 | 5,11,8 | 256 | 150 | 8.92E-09 | 4.00E-07 | 1.0,1.0,1.0 | 0.652,0.551,0.71 | 0.359 | ENSG00000082 | [[[[149813215, 1 | 2 |
| ENSG00000132 | DAP3 | chr1 | 155709772 | 155709824 | 155689057 | 155689174 | 155727607 | 155727738 | 3,2,3 | 0,0,0 | 2,5,5 | 1,6,0 | 202 | 150 | 0.000262574 | 0.003674499 | 1.0,1.0,1.0 | 0.598,0.382,1.0 | 0.34 | ENSG00000132 | [[[[155689091, 1 | 5 |
| ENSG00000132 | DAP3 | chr1 | 155717005 | 155717128 | 155689057 | 155689174 | 155725381 | 155725490 | 4,12,13 | 0,0,4 | 31,40,16 | 18,30,16 | 273 | 150 | 0.000447885 | 0.005846833 | 1.0,1.0,0.641 | 0.486,0.423,0.35 | 0.459 | ENSG00000132 | [[[[155689091, 1 | 3 |
| ENSG00000132 | DAP3 | chr1 | 155721516 | 155721618 | 155689057 | 155689174 | 155725381 | 155725411 | 135,146,260 | 0,0,4 | 234,391,245 | 18,30,16 | 252 | 150 | 3.55E-11 | 2.48E-09 | 1.0,1.0,0.975 | 0.886,0.886,0.90 | 0.101 | ENSG00000132 | [[[[155689091, 1 | 3 |
| ENSG00000132 | DAP3 | chr1 | 155725381 | 155725411 | 155709772 | 155709824 | 155727607 | 155727738 | 1,1,2 | 0,0,0 | 7,6,5 | 2,0,5 | 180 | 150 | 0.000146798 | 0.002222914 | 1.0,1.0,1.0 | 0.745,1.0,0.455 | 0.267 | ENSG00000132 | [[[[155689091, 1 | 4 |
| ENSG00000132 | DAP3 | chr1 | 155731355 | 155731415 | 155729207 | 155729366 | 155736945 | 155737063 | 117,98,169 | 0,0,0 | 138,258,162 | 7,25,13 | 210 | 150 | 1.72E-11 | 1.27E-09 | 1.0,1.0,1.0 | 0.934,0.881,0.89 | 0.095 | ENSG00000132 | [[[[155689091, 1 | 2 |
| ENSG00000166 | POLL | chr10 | 101582762 | 101582891 | 101580247 | 101580416 | 101587245 | 101587406 | 25,16,35 | 0,0,1 | 36,40,47 | 0,10,6 | 279 | 150 | 2.33E-05 | 0.000449695 | 1.0,1.0,0.95 | 1.0,0.683,0.808 | 0.153 | ENSG00000166 | [[[[101587822, 1 | 5 |
| ENSG00000166 | POLL | chr10 | 101584877 | 101584919 | 101582762 | 101582891 | 101587245 | 101587406 | 0,0,2 | 4,1,3 | 5,8,5 | 3,2,2 | 192 | 150 | 1.08E-05 | 0.000228692 | 0.0,0.0,0.342 | 0.566,0.758,0.66 | -0.548 | ENSG00000166 | [[[[101587822, 1 | 4 |
| ENSG00000166 | POLL | chr10 | 101584877 | 101584919 | 101583507 | 101583681 | 101587245 | 101587406 | 1,1,3 | 1,0,0 | 2,7,10 | 0,4,3 | 192 | 150 | 0.000552594 | 0.007011911 | 0.439,1.0,1.0 | 1.0,0.578,0.723 | 0.046 | ENSG00000166 | [[[[101587822, 1 | 3 |
| ENSG00000160 | FDPS | chr1 | 155310042 | 155310205 | 155308793 | 155308965 | 155312254 | 155312395 | 9,727,841,318 | 17,10,29 | 5,421,127,667 | 20,52,24 | 300 | 150 | 1.21E-05 | 0.000253078 | 0.966,0.975,0.95 | 0.931,0.916,0.93 | 0.04 | ENSG00000160 | [[[[155308866, 1 | 2 |
| ENSG00000160 | FDPS | chr1 | 155310063 | 155310205 | 155308754 | 155308965 | 155312254 | 155312395 | 9,227,361,183 | 17,10,29 | 407,863,518 | 20,52,24 | 292 | 150 | 7.85E-08 | 2.83E-06 | 0.965,0.974,0.95 | 0.913,0.895,0.91 | 0.056 | ENSG00000160 | [[[[155308866, 1 | 2 |
| ENSG00000160 | FDPS | chr1 | 155318855 | 155318928 | 155312254 | 155312395 | 155319610 | 155319688 | 289,246,341 | 0,0,0 | 130,308,145 | 4,9,7 | 223 | 150 | 4.81E-06 | 0.000111611 | 1.0,1.0,1.0 | 0.956,0.958,0.93 | 0.051 | ENSG00000160 | [[[[155308866, 1 | 4 |
| ENSG00000160 | FLAD1 | chr1 | 154990158 | 154990257 | 154983372 | 154984066 | 154992712 | 154992786 | 0,0,0 | 2,1,4 | 6,4,2 | 6,14,5 | 249 | 150 | 6.27E-06 | 0.000141905 | 0.0,0.0,0.0 | 0.376,0.147,0.19 | -0.239 | ENSG00000160 | [[[[154983344, 1 | 4 |
| ENSG00000107 | LIPA | chr10 | 89245675 | 89245793 | 89228199 | 89228398 | 89251736 | 89252058 | 134,65,169 | 0,1,2 | 53,165,93 | 15,18,15 | 268 | 150 | 8.50E-14 | 8.90E-12 | 1.0,0.973,0.979 | 0.664,0.837,0.77 | 0.225 | ENSG00000107 | [[[[89251737, 89 | 2 |
| ENSG00000107 | LIPA | chr10 | 89247537 | 89247649 | 89228199 | 89228398 | 89251736 | 89252058 | 34,48,132 | 0,1,2 | 31,61,68 | 15,18,15 | 262 | 150 | 1.82E-14 | 2.11E-12 | 1.0,0.965,0.974 | 0.542,0.66,0.722 | 0.338 | ENSG00000107 | [[[[89251737, 89 | 2 |
| ENSG00000187 | SESTD1 | chr2 | 179149296 | 179149394 | 179124363 | 179124558 | 179151277 | 179151391 | 21,10,26 | 0,0,0 | 14,32,20 | 2,6,7 | 248 | 150 | 6.38E-08 | 2.35E-06 | 1.0,1.0,1.0 | 0.809,0.763,0.63 | 0.265 | ENSG00000187 | [[[[179264499, 1 | 4 |
| ENSG00000080 | RAB21 | chr12 | 71782030 | 71782085 | 71770535 | 71770699 | 71782569 | 71782658 | 44,36,33 | 0,0,0 | 29,65,50 | 7,7,6 | 205 | 150 | 6.84E-08 | 2.50E-06 | 1.0,1.0,1.0 | 0.752,0.872,0.85 | 0.172 | ENSG00000080 | [[[[71754863, 71 | 2 |
| ENSG00000125 | ATG4C | chr1 | 62821138 | 62821209 | 62819004 | 62819335 | 62834775 | 62834852 | 7,7,12 | 0,0,0 | 3,25,8 | 2,3,3 | 221 | 150 | 8.72E-06 | 0.000189676 | 1.0,1.0,1.0 | 0.504,0.85,0.644 | 0.334 | ENSG00000125 | [[[[62784132, 62 | 3 |
| ENSG00000186 | LEMD1 | chr1 | 205419229 | 205419352 | 205381377 | 205381856 | 205420454 | 205420574 | 5,4,5 | 46,20,35 | 7,13,8 | 18,19,18 | 273 | 150 | 0.000310727 | 0.004257651 | 0.056,0.099,0.07 | 0.176,0.273,0.19 | -0.139 | ENSG00000186 | [[[[205421990, 2 | 3 |
| ENSG00000186 | LEMD1 | chr1 | 205419229 | 205419352 | 205384287 | 205384364 | 205420454 | 205420574 | 1,4,2 | 20,7,18 | 4,15,6 | 7,13,10 | 273 | 150 | 0.000294456 | 0.004063193 | 0.027,0.239,0.05 | 0.239,0.388,0.24 | -0.184 | ENSG00000186 | [[[[205421990, 2 | 2 |
| ENSG00000161 | SQSTM1 | chr5 | 179822957 | 179823053 | 179820758 | 179821141 | 179833586 | 179833782 | 3,23,52 | 0,0,0 | 38,72,63 | 4,4,2 | 246 | 150 | 0.000645507 | 0.007991288 | 1.0,1.0,1.0 | 0.853,0.916,0.95 | 0.093 | ENSG00000161 | [[[[179820905, 1 | 5 |
| ENSG00000090 | USP48 | chr1 | 21736445 | 21736625 | 21721035 | 21721166 | 21747066 | 21747149 | 55,56,89 | 0,0,0 | 17,40,23 | 1,5,2 | 300 | 150 | 4.86E-05 | 0.000850509 | 1.0,1.0,1.0 | 0.895,0.8,0.852 | 0.151 | ENSG00000090 | [[[[21782824, 21 | 5 |
| ENSG00000090 | USP48 | chr1 | 21736445 | 21736625 | 21723897 | 21724095 | 21747066 | 21747149 | 55,56,89 | 0,0,0 | 28,41,26 | 7,7,6 | 300 | 150 | 2.71E-12 | 2.31E-10 | 1.0,1.0,1.0 | 0.667,0.745,0.68 | 0.301 | ENSG00000090 | [[[[21782824, 21 | 3 |
| ENSG00000183 | ZNF530 | chr19 | 57604276 | 57604406 | 57599884 | 57600134 | 57612502 | 57612722 | 4,3,1 | 1,0,0 | 1,4,3 | 8,8,2 | 280 | 150 | 8.67E-07 | 2.42E-05 | 0.682,1.0,1.0 | 0.063,0.211,0.44 | 0.654 | ENSG00000183 | [[[[57599885, 57 | 3 |
| ENSG00000106 | PTPRZ1 | chr7 | 122019123 | 122019268 | 122010333 | 122011309 | 122031473 | 122031559 | 17,16,43 | 0,0,0 | 14,32,13 | 3,13,12 | 295 | 150 | 1.33E-11 | 1.00E-09 | 1.0,1.0,1.0 | 0.704,0.556,0.35 | 0.462 | ENSG00000106 | [[[[121873161, 1 | 2 |
| ENSG00000132 | DPH2 | chr1 | 43970965 | 43971189 | 43970595 | 43970708 | 43972157 | 43972334 | 30,41,62 | 0,0,0 | 28,56,45 | 4,21,8 | 300 | 150 | 4.83E-12 | 3.98E-10 | 1.0,1.0,1.0 | 0.778,0.571,0.73 | 0.304 | ENSG00000132 | [[[[43970010, 43 | 2 |
| ENSG00000132 | DPH2 | chr1 | 43971386 | 43972070 | 43970595 | 43970708 | 43972157 | 43972334 | 31,67,101 | 0,0,0 | 93,187,92 | 4,21,8 | 300 | 150 | 4.52E-09 | 2.18E-07 | 1.0,1.0,1.0 | 0.921,0.817,0.85 | 0.137 | ENSG00000132 | [[[[43970010, 43 | 2 |
| ENSG00000132 | DPH2 | chr1 | 43971659 | 43972070 | 43970595 | 43970708 | 43972157 | 43972334 | 31,67,101 | 0,0,0 | 100,195,96 | 4,21,8 | 300 | 150 | 7.80E-09 | 3.56E-07 | 1.0,1.0,1.0 | 0.926,0.823,0.85 | 0.131 | ENSG00000132 | [[[[43970010, 43 | 2 |
| ENSG00000168 | MFSD2A | chr1 | 39967085 | 39967169 | 39966810 | 39966932 | 39967803 | 39967916 | 72,62,101 | 0,0,0 | 88,153,85 | 1,4,10 | 234 | 150 | 6.42E-05 | 0.001085623 | 1.0,1.0,1.0 | 0.983,0.961,0.84 | 0.07 | ENSG00000168 | [[[[39955145, 39 | 2 |
| ENSG00000085 | SLC25A24 | chr1 | 108148278 | 108148386 | 108139057 | 108139208 | 108154982 | 108155084 | 54,32,56 | 0,0,0 | 34,108,68 | 2,3,9 | 258 | 150 | 1.54E-05 | 0.000311654 | 1.0,1.0,1.0 | 0.908,0.954,0.81 | 0.108 | ENSG00000085 | [[[[108199956, 1 | 2 |
| ENSG00000164 | EXO5 | chr1 | 40509450 | 40509625 | 40508789 | 40508908 | 40514514 | 40514846 | 0,0,0 | 1,1,2 | 1,8,4 | 1,1,2 | 300 | 150 | 1.17E-08 | 5.14E-07 | 0.0,0.0,0.0 | 0.333,0.8,0.5 | -0.544 | ENSG00000164 | [[[[40508767, 40 | 2 |
| ENSG00000091 | NLRP1 | chr17 | 5515472 | 5515517 | 5501813 | 5501872 | 5517745 | 5517887 | 0,0,1 | 1,2,0 | 5,4,7 | 2,1,2 | 195 | 150 | 0.000190387 | 0.002794668 | 0.0,0.0,1.0 | 0.658,0.755,0.72 | -0.381 | ENSG00000091 | [[[[5583687, 558 | 2 |

| | | | | | | | | | | | | | | | | | | | | | | |
|---|---|---|---|---|---|---|---|---|---|---|---|---|---|---|---|---|---|---|---|---|---|---|
| ENSG00000198 | GFPT1 | chr2 | 69329683 | 69329798 | 69329296 | 69329424 | 69338444 | 69338565 | 84,65,112 | 0,0,0 | 91,169,94 | 10,23,10 | 265 | 150 | 8.34E-13 | 7.66E-11 | 1.0,1.0,1.0 | 0.837,0.806,0.84 | 0.172 | ENSG00000198 | [[[[69387065, 693 | 2 |
| ENSG00000086 | B4GALT1 | chr9 | 33120418 | 33120606 | 33113773 | 33113878 | 33135188 | 33135424 | 95,75,121 | 0,0,0 | 143,347,261 | 5,15,5 | 300 | 150 | 1.26E-05 | 0.000260791 | 1.0,1.0,1.0 | 0.935,0.92,0.963 | 0.061 | ENSG00000086 | [[[[33166758, 331 | 2 |
| ENSG00000119 | YLPM1 | chr14 | 74816930 | 74817107 | 74816570 | 74816690 | 74818230 | 74818314 | 27,31,58 | 0,0,0 | 27,77,49 | 2,5,4 | 300 | 150 | 4.05E-05 | 0.000728147 | 1.0,1.0,1.0 | 0.871,0.885,0.86 | 0.128 | ENSG00000119 | [[[[74763316, 747 | 2 |
| ENSG00000119 | YLPM1 | chr14 | 74817193 | 74817277 | 74816570 | 74816690 | 74818230 | 74818314 | 23,25,29 | 0,0,0 | 33,66,26 | 2,5,4 | 234 | 150 | 3.64E-05 | 0.000663962 | 1.0,1.0,1.0 | 0.914,0.894,0.80 | 0.129 | ENSG00000119 | [[[[74763316, 747 | 2 |
| ENSG00000100 | RANBP9 | chr6 | 13637807 | 13637955 | 13632369 | 13632521 | 13639562 | 13639753 | 46,42,59 | 0,0,0 | 45,93,68 | 16,25,6 | 298 | 150 | 3.88E-12 | 3.23E-10 | 1.0,1.0,1.0 | 0.586,0.652,0.85 | 0.304 | ENSG00000100 | [[[[13710935, 137 | 2 |
| ENSG00000100 | RANBP9 | chr6 | 13652658 | 13652681 | 13632369 | 13632521 | 13657108 | 13657276 | 5,6,16 | 0,0,0 | 10,18,13 | 8,0,1 | 173 | 150 | 0.000352425 | 0.004740355 | 1.0,1.0,1.0 | 0.52,1.0,0.919 | 0.187 | ENSG00000100 | [[[[13710935, 137 | 7 |
| ENSG00000115 | AAK1 | chr2 | 69476879 | 69476990 | 69465845 | 69466551 | 69480859 | 69480961 | 0,0,4 | 1,1,2 | 2,1,1 | 0,1,0 | 261 | 150 | 2.83E-05 | 0.000531561 | 0.0,0.0,0.535 | 1.0,0.365,1.0 | -0.61 | ENSG00000115 | [[[[69643575, 696 | 2 |
| ENSG00000175 | CCNE2 | chr8 | 94885066 | 94885201 | 94882780 | 94882892 | 94890414 | 94890458 | 23,26,24 | 0,0,0 | 14,36,23 | 3,7,6 | 285 | 150 | 1.13E-08 | 4.98E-07 | 1.0,1.0,1.0 | 0.711,0.73,0.669 | 0.297 | ENSG00000175 | [[[[94895177, 948 | 3 |
| ENSG00000160 | FAM189B | chr1 | 155251094 | 155251164 | 155248404 | 155248473 | 155253624 | 155253732 | 1,2,3 | 0,1,0 | 8,15,13 | 5,6,6 | 220 | 150 | 0.000109716 | 0.001729081 | 1.0,0.577,1.0 | 0.522,0.63,0.596 | 0.276 | ENSG00000160 | [[[[155254653, 15 | 4 |
| ENSG00000160 | FAM189B | chr1 | 155251501 | 155251601 | 155251094 | 155251164 | 155253624 | 155253732 | 22,28,41 | 0,0,1 | 35,80,47 | 10,16,8 | 250 | 150 | 4.10E-09 | 1.99E-07 | 1.0,1.0,0.961 | 0.677,0.75,0.779 | 0.252 | ENSG00000160 | [[[[155254653, 15 | 2 |
| ENSG00000171 | PPIH | chr1 | 42659521 | 42659566 | 42659227 | 42659251 | 42664862 | 42664955 | 105,88,175 | 0,0,0 | 175,345,202 | 8,11,11 | 195 | 150 | 2.50E-06 | 6.23E-05 | 1.0,1.0,1.0 | 0.944,0.96,0.934 | 0.054 | ENSG00000171 | [[[[42658423, 426 | 2 |
| ENSG00000171 | PPIH | chr1 | 42660861 | 42660904 | 42659227 | 42659251 | 42664862 | 42664955 | 59,45,71 | 0,0,0 | 81,126,90 | 8,11,11 | 193 | 150 | 1.52E-08 | 6.51E-07 | 1.0,1.0,1.0 | 0.887,0.899,0.86 | 0.117 | ENSG00000171 | [[[[42658423, 426 | 2 |
| ENSG00000117 | CENPF | chr1 | 214648674 | 214648827 | 214644556 | 214647400 | 214652827 | 214652989 | 119,104,106 | 0,0,0 | 67,211,109 | 8,45,20 | 300 | 150 | 0 | 0 | 1.0,1.0,1.0 | 0.807,0.701,0.73 | 0.253 | ENSG00000117 | [[[[214603195, 21 | 2 |
| ENSG00000075 | SLC25A3 | chr12 | 98597855 | 98598035 | 98595432 | 98595557 | 98598521 | 98598703 | 755,742,976 | 0,0,0 | 505,959,625 | 5,22,13 | 300 | 150 | 1.89E-08 | 7.94E-07 | 1.0,1.0,1.0 | 0.981,0.956,0.96 | 0.034 | ENSG00000075 | [[[[98593686, 985 | 2 |
| ENSG00000075 | SLC25A3 | chr12 | 98597855 | 98598035 | 98595661 | 98595661 | 98598521 | 98598703 | 758,746,979 | 0,0,0 | 5,331,038,653 | 15,33,14 | 300 | 150 | 6.44E-15 | 7.90E-13 | 1.0,1.0,1.0 | 0.947,0.94,0.959 | 0.051 | ENSG00000075 | [[[[98593686, 985 | 2 |
| ENSG00000122 | SHLD2 | chr10 | 87175888 | 87176095 | 87158047 | 87158155 | 87180074 | 87180303 | 39,36,34 | 0,0,0 | 15,29,25 | 0,4,3 | 300 | 150 | 0.000555118 | 0.007040422 | 1.0,1.0,1.0 | 1.0,0.784,0.806 | 0.137 | ENSG00000122 | [[[[87095206, 870 | 3 |
| ENSG00000117 | RPA2 | chr1 | 27894011 | 27894106 | 27891523 | 27892247 | 27906927 | 27907041 | 111,95,125 | 0,0,0 | 114,218,145 | 3,9,5 | 245 | 150 | 8.29E-05 | 0.001350051 | 1.0,1.0,1.0 | 0.959,0.937,0.94 | 0.052 | ENSG00000117 | [[[[27914434, 279 | 4 |
| ENSG00000198 | HIBCH | chr2 | 190212955 | 190213075 | 190203920 | 190205232 | 190244886 | 190244968 | 30,10,34 | 0,3,0 | 11,37,11 | 3,9,4 | 270 | 150 | 0.000151428 | 0.00228305 | 1.0,0.649,1.0 | 0.671,0.695,0.60 | 0.226 | ENSG00000198 | [[[[190319716, 19 | 2 |
| ENSG00000198 | HIBCH | chr2 | 190244886 | 190244968 | 190212955 | 190213075 | 190252161 | 190252307 | 17,3,13 | 0,0,0 | 4,20,7 | 3,2,2 | 232 | 150 | 4.67E-05 | 0.00082285 | 1.0,1.0,1.0 | 0.463,0.866,0.69 | 0.326 | ENSG00000198 | [[[[190319716, 19 | 3 |
| ENSG00000197 | PELI1 | chr2 | 64100397 | 64100499 | 64096412 | 64096610 | 64108239 | 64108379 | 14,4,20 | 0,1,0 | 6,32,19 | 1,4,3 | 252 | 150 | 0.000676993 | 0.008318906 | 1.0,0.704,1.0 | 0.781,0.826,0.79 | 0.102 | ENSG00000197 | [[[[64144081, 641 | 2 |
| ENSG00000204 | ABHD16A | chr6 | 31701273 | 31701340 | 31700941 | 31701028 | 31703149 | 31703444 | 14,16,34 | 1,3,4 | 28,36,29 | 11,16,13 | 217 | 150 | 0.000494828 | 0.006374456 | 0.906,0.787,0.85 | 0.638,0.609,0.60 | 0.231 | ENSG00000204 | [[[[31703150, 317 | 2 |
| ENSG00000204 | ABHD16A | chr6 | 31702073 | 31702130 | 31700941 | 31701028 | 31703149 | 31703311 | 11,31,56 | 1,3,4 | 33,52,57 | 11,16,13 | 207 | 150 | 0.000224224 | 0.003216383 | 0.889,0.882,0.91 | 0.685,0.702,0.76 | 0.178 | ENSG00000204 | [[[[31703150, 317 | 2 |
| ENSG00000166 | IVNS1ABP | chr1 | 185306484 | 185306625 | 185305535 | 185305643 | 185308799 | 185308875 | 15,6,21 | 1,0,0 | 17,54,25 | 12,88,59 | 291 | 150 | 1.78E-15 | 2.33E-13 | 0.885,1.0,1.0 | 0.422,0.24,0.179 | 0.681 | ENSG00000166 | [[[[185316953, 18 | 2 |
| ENSG00000166 | IVNS1ABP | chr1 | 185307013 | 185307139 | 185305535 | 185305643 | 185308799 | 185308875 | 86,86,150 | 1,0,0 | 17,43,34 | 12,88,59 | 276 | 150 | 0 | 0 | 0.979,1.0,1.0 | 0.435,0.21,0.238 | 0.699 | ENSG00000166 | [[[[185316953, 18 | 2 |
| ENSG00000166 | IVNS1ABP | chr1 | 185307013 | 185307139 | 185306484 | 185306625 | 185308799 | 185308875 | 20,8,24 | 0,0,0 | 8,30,17 | 6,14,4 | 276 | 150 | 2.72E-10 | 1.66E-08 | 1.0,1.0,1.0 | 0.42,0.538,0.698 | 0.448 | ENSG00000166 | [[[[185316953, 18 | 2 |
| ENSG00000137 | KIF23 | chr15 | 69440767 | 69441079 | 69440307 | 69440487 | 69446282 | 69446364 | 35,21,42 | 0,0,0 | 58,105,45 | 7,17,5 | 300 | 150 | 7.85E-09 | 3.57E-07 | 1.0,1.0,1.0 | 0.806,0.755,0.81 | 0.207 | ENSG00000137 | [[[[69414349, 694 | 3 |
| ENSG00000137 | KIF23 | chr15 | 69446008 | 69446091 | 69440767 | 69441079 | 69446282 | 69446364 | 108,87,112 | 0,0,0 | 43,99,48 | 8,15,0 | 233 | 150 | 2.22E-09 | 1.15E-07 | 1.0,1.0,1.0 | 0.776,0.809,1.0 | 0.138 | ENSG00000137 | [[[[69414349, 694 | 2 |
| ENSG00000126 | PCID2 | chr13 | 113179916 | 113180042 | 113177576 | 113178287 | 113180157 | 113180231 | 21,33,54 | 0,0,0 | 25,69,43 | 1,8,1 | 276 | 150 | 6.91E-05 | 0.001156792 | 1.0,1.0,1.0 | 0.931,0.824,0.95 | 0.095 | ENSG00000126 | [[[[113208599, 11 | 2 |
| ENSG00000138 | SELENOI | chr2 | 26364301 | 26364370 | 26346085 | 26346289 | 26373366 | 26373506 | 2,11,41 | 0,0,0 | 14,50,42 | 2,2,7 | 219 | 150 | 4.05E-05 | 0.000728147 | 1.0,1.0,1.0 | 0.827,0.945,0.80 | 0.141 | ENSG00000138 | [[[[26346143, 263 | 3 |
| ENSG00000138 | SELENOI | chr2 | 26367145 | 26367220 | 26346085 | 26346289 | 26373366 | 26373506 | 4,1,8 | 0,0,0 | 4,11,14 | 2,2,7 | 225 | 150 | 1.27E-07 | 4.32E-06 | 1.0,1.0,1.0 | 0.571,0.786,0.57 | 0.357 | ENSG00000138 | [[[[26346143, 263 | 3 |
| ENSG00000065 | YBX1 | chr1 | 42696198 | 42696288 | 42693489 | 42693523 | 42700780 | 42701046 | 625,622,918 | 0,0,0 | 7,461,660,956 | 68,164,94 | 240 | 150 | 0 | 0 | 1.0,1.0,1.0 | 0.873,0.864,0.86 | 0.133 | ENSG00000065 | [[[[42682418, 426 | 3 |
| ENSG00000065 | YBX1 | chr1 | 42697179 | 42697262 | 42693489 | 42693523 | 42700780 | 42701046 | 116,911,871,882 | 0,0,0 | 145,123,741,371 | 68,164,94 | 233 | 150 | 0 | 0 | 1.0,1.0,1.0 | 0.932,0.903,0.90 | 0.087 | ENSG00000065 | [[[[42682418, 426 | 3 |
| ENSG00000065 | YBX1 | chr1 | 42697179 | 42697262 | 42696198 | 42696288 | 42700780 | 42701046 | 116,911,871,882 | 0,0,0 | 144,823,771,372 | 172,281,158 | 233 | 150 | 0 | 0 | 1.0,1.0,1.0 | 0.844,0.845,0.84 | 0.154 | ENSG00000065 | [[[[42682418, 426 | 2 |
| ENSG00000058 | YIPF1 | chr1 | 53866199 | 53866382 | 53860055 | 53860153 | 53871371 | 53871488 | 40,34,45 | 1,0,2 | 26,69,37 | 8,10,1 | 300 | 150 | 0.000160977 | 0.002410326 | 0.952,1.0,0.918 | 0.619,0.775,0.94 | 0.176 | ENSG00000058 | [[[[53889713, 538 | 2 |
| ENSG00000066 | KDM4A | chr1 | 43693993 | 43694102 | 43692255 | 43692311 | 43703616 | 43703736 | 37,26,48 | 0,0,0 | 22,46,42 | 4,9,1 | 259 | 150 | 9.25E-07 | 2.57E-05 | 1.0,1.0,1.0 | 0.761,0.747,0.96 | 0.177 | ENSG00000066 | [[[[43650149, 436 | 3 |
| ENSG00000144 | UBA3 | chr3 | 69077797 | 69077918 | 69067927 | 69068008 | 69080111 | 69080153 | 70,55,152 | 0,0,0 | 49,131,76 | 8,18,12 | 271 | 150 | 2.62E-14 | 2.96E-12 | 1.0,1.0,1.0 | 0.772,0.801,0.77 | 0.216 | ENSG00000144 | [[[[69080334, 690 | 3 |
| ENSG00000169 | THBS3 | chr1 | 155205059 | 155205316 | 155203512 | 155203539 | 155206199 | 155206406 | 4,9,11 | 0,0,0 | 5,6,1 | 1,6,2 | 300 | 150 | 9.59E-09 | 4.28E-07 | 1.0,1.0,1.0 | 0.714,0.333,0.2 | 0.584 | ENSG00000169 | [[[[155207798, 15 | 2 |
| ENSG00000147 | EIF3H | chr8 | 116657214 | 116657314 | 116655855 | 116656005 | 116726015 | 116726160 | 103,106,150 | 1,2,0 | 113,274,136 | 13,29,28 | 250 | 150 | 5.84E-14 | 6.22E-12 | 0.984,0.97,1.0 | 0.839,0.85,0.745 | 0.173 | ENSG00000147 | [[[[116755666, 11 | 2 |
| ENSG00000147 | EIF3H | chr8 | 116657214 | 116657314 | 116655855 | 116656005 | 116755665 | 116755784 | 102,106,149 | 0,0,0 | 113,275,134 | 4,25,6 | 250 | 150 | 1.50E-09 | 8.04E-08 | 1.0,1.0,1.0 | 0.944,0.868,0.93 | 0.086 | ENSG00000147 | [[[[116755666, 11 | 3 |
| ENSG00000147 | EIF3H | chr8 | 116658812 | 116658980 | 116655855 | 116656005 | 116726015 | 116726160 | 248,229,402 | 1,2,0 | 297,633,414 | 13,29,28 | 300 | 150 | 3.53E-13 | 3.39E-11 | 0.992,0.983,1.0 | 0.92,0.916,0.881 | 0.086 | ENSG00000147 | [[[[116755666, 11 | 2 |
| ENSG00000147 | EIF3H | chr8 | 116726015 | 116726172 | 116655855 | 116656005 | 116755665 | 116755797 | 248,353,566 | 0,0,0 | 443,889,607 | 4,25,6 | 300 | 150 | 6.21E-06 | 0.000140601 | 1.0,1.0,1.0 | 0.982,0.947,0.98 | 0.03 | ENSG00000147 | [[[[116755666, 11 | 3 |
| ENSG00000047 | SCML1 | chrX | 17744070 | 17744219 | 17737468 | 17737680 | 17746017 | 17746098 | 0,0,2 | 5,5,11 | 1,5,9 | 2,13,11 | 299 | 150 | 6.89E-05 | 0.001154559 | 0.0,0.0,0.084 | 0.201,0.162,0.29 | -0.19 | ENSG00000047 | [[[[17737493, 177 | 2 |
| ENSG00000047 | SCML1 | chrX | 17746014 | 17746098 | 17737448 | 17737680 | 17749399 | 17749504 | 22,14,33 | 0,0,0 | 14,41,30 | 8,17,24 | 234 | 150 | 7.77E-15 | 9.43E-13 | 1.0,1.0,1.0 | 0.529,0.607,0.44 | 0.473 | ENSG00000047 | [[[[17737493, 177 | 2 |
| ENSG00000047 | SCML1 | chrX | 17746014 | 17746098 | 17744070 | 17744219 | 17749399 | 17749504 | 22,14,19 | 0,0,0 | 7,30,27 | 2,1,6 | 234 | 150 | 2.00E-05 | 0.000393036 | 1.0,1.0,1.0 | 0.692,0.951,0.74 | 0.205 | ENSG00000047 | [[[[17737493, 177 | 2 |
| ENSG00000047 | SCML1 | chrX | 17746017 | 17746098 | 17737467 | 17737680 | 17749399 | 17749504 | 27,19,29 | 0,0,0 | 9,43,33 | 8,17,24 | 231 | 150 | 5.33E-15 | 6.60E-13 | 1.0,1.0,1.0 | 0.422,0.622,0.47 | 0.495 | ENSG00000047 | [[[[17737493, 177 | 3 |
| ENSG00000047 | SCML1 | chrX | 17746017 | 17746098 | 17744070 | 17744219 | 17749399 | 17749504 | 22,14,18 | 0,0,0 | 8,30,23 | 2,1,6 | 231 | 150 | 1.61E-05 | 0.000324275 | 1.0,1.0,1.0 | 0.722,0.951,0.71 | 0.205 | ENSG00000047 | [[[[17737493, 177 | 2 |
| ENSG00000185 | ANKFY1 | chr17 | 4181253 | 4181372 | 4179720 | 4179877 | 4183397 | 4183551 | 33,31,74 | 0,0,0 | 35,93,70 | 0,12,6 | 269 | 150 | 1.37E-06 | 3.64E-05 | 1.0,1.0,1.0 | 1.0,0.812,0.867 | 0.107 | ENSG00000185 | [[[[4263932, 4263 | 2 |
| ENSG00000185 | ANKFY1 | chr17 | 4182180 | 4182349 | 4179720 | 4179877 | 4183397 | 4183551 | 26,18,66 | 0,0,0 | 27,46,41 | 0,12,6 | 300 | 150 | 3.09E-07 | 9.65E-06 | 1.0,1.0,1.0 | 1.0,0.657,0.774 | 0.19 | ENSG00000185 | [[[[4263932, 4263 | 2 |
| ENSG00000185 | ANKFY1 | chr17 | 4182180 | 4182352 | 4179720 | 4179877 | 4183397 | 4183551 | 9,4,11 | 0,0,0 | 12,17,14 | 0,12,6 | 300 | 150 | 4.86E-07 | 1.45E-05 | 1.0,1.0,1.0 | 1.0,0.415,0.538 | 0.349 | ENSG00000185 | [[[[4263932, 4263 | 2 |
| ENSG00000137 | IFI44 | chr1 | 78660553 | 78660654 | 78659311 | 78659483 | 78663764 | 78664078 | 94,91,137 | 12,10,9 | 54,122,65 | 18,31,18 | 251 | 150 | 6.57E-06 | 0.000147692 | 0.824,0.845,0.90 | 0.642,0.702,0.68 | 0.181 | ENSG00000137 | [[[[78649831, 786 | 2 |
| ENSG00000137 | IFI44 | chr1 | 78662703 | 78662878 | 78659311 | 78659483 | 78663764 | 78663897 | 150,125,192 | 12,10,9 | 96,183,78 | 18,31,18 | 300 | 150 | 6.11E-06 | 0.000138628 | 0.862,0.862,0.91 | 0.727,0.747,0.68 | 0.16 | ENSG00000137 | [[[[78649831, 786 | 2 |
| ENSG00000198 | SREBF2 | chr22 | 41873797 | 41874019 | 41870888 | 41871035 | 41880715 | 41880992 | 69,93,170 | 0,0,1 | 61,154,132 | 0,7,6 | 300 | 150 | 0.000216903 | 0.003126665 | 1.0,1.0,0.988 | 1.0,0.917,0.917 | 0.051 | ENSG00000198 | [[[[41833105, 418 | 5 |
| ENSG00000198 | SREBF2 | chr22 | 41877228 | 41877421 | 41875542 | 41875724 | 41880715 | 41880992 | 133,115,200 | 1,0,6 | 119,146,134 | 7,11,10 | 300 | 150 | 1.88E-05 | 0.000372693 | 0.985,1.0,0.943 | 0.895,0.869,0.87 | 0.098 | ENSG00000198 | [[[[41833105, 418 | 2 |
| ENSG00000007 | PAFAH1B1 | chr17 | 2676504 | 2676606 | 2673664 | 2674288 | 2681728 | 2682038 | 186,148,244 | 0,0,0 | 166,319,218 | 3,11,3 | 252 | 150 | 0.000463583 | 0.006031095 | 1.0,1.0,1.0 | 0.971,0.945,0.97 | 0.036 | ENSG00000007 | [[[[2593654, 2594 | 2 |
| ENSG00000007 | PAFAH1B1 | chr17 | 2680163 | 2680320 | 2674059 | 2674096 | 2681728 | 2681910 | 177,146,266 | 0,0,0 | 100,210,124 | 12,21,13 | 300 | 150 | 0 | 0 | 1.0,1.0,1.0 | 0.806,0.833,0.82 | 0.178 | ENSG00000007 | [[[[2593654, 2594 | 2 |
| ENSG00000007 | PAFAH1B1 | chr17 | 2680163 | 2680320 | 2674059 | 2674288 | 2681728 | 2681910 | 177,149,266 | 0,0,0 | 101,200,123 | 3,11,3 | 300 | 150 | 1.50E-06 | 3.96E-05 | 1.0,1.0,1.0 | 0.944,0.901,0.95 | 0.067 | ENSG00000007 | [[[[2593654, 2594 | 2 |
| ENSG00000144 | SPOPL | chr2 | 138550156 | 138550294 | 138501800 | 138502119 | 138550973 | 138551054 | 5,3,16 | 0,0,0 | 5,5,12 | 0,10,9 | 288 | 150 | 2.98E-07 | 9.33E-06 | 1.0,1.0,1.0 | 1.0,0.207,0.41 | 0.461 | ENSG00000144 | [[[[138501770, 13 | 2 |
| ENSG00000144 | SPOPL | chr2 | 138550482 | 138550604 | 138501800 | 138502119 | 138550973 | 138551054 | 2,3,3 | 0,0,0 | 3,10,16 | 0,10,9 | 272 | 150 | 2.49E-07 | 7.93E-06 | 1.0,1.0,1.0 | 1.0,0.355,0.495 | 0.383 | ENSG00000144 | [[[[138501770, 13 | 2 |
| ENSG00000144 | SPOPL | chr2 | 138550902 | 138551054 | 138550482 | 138550604 | 138559021 | 138559199 | 11,7,13 | 0,0,0 | 6,7,2 | 3,2,2 | 300 | 150 | 3.23E-07 | 1.00E-05 | 1.0,1.0,1.0 | 0.5,0.636,0.333 | 0.51 | ENSG00000144 | [[[[138501770, 13 | 2 |
| ENSG00000144 | SPOPL | chr2 | 138552553 | 138552681 | 138550902 | 138551054 | 138559281 | 138559337 | 12,5,18 | 0,0,0 | 9,31,24 | 2,7,6 | 278 | 150 | 1.84E-07 | 6.03E-06 | 1.0,1.0,1.0 | 0.708,0.705,0.68 | 0.301 | ENSG00000144 | [[[[138501770, 13 | 2 |
| ENSG00000144 | SPOPL | chr2 | 138559021 | 138559199 | 138550902 | 138551054 | 138559281 | 138559337 | 9,8,22 | 0,0,0 | 13,36,24 | 2,7,6 | 300 | 150 | 1.77E-07 | 5.83E-06 | 1.0,1.0,1.0 | 0.765,0.72,0.667 | 0.283 | ENSG00000144 | [[[[138501770, 13 | 2 |
| ENSG00000143 | TARS2 | chr1 | 150496809 | 150496908 | 150492184 | 150492489 | 150497529 | 150497587 | 8,15,20 | 7,15,27 | 29,37,25 | 5,10,9 | 249 | 150 | 5.18E-06 | 0.000119664 | 0.408,0.376,0.30 | 0.777,0.69,0.626 | -0.333 | ENSG00000143 | [[[[150487419, 15 | 2 |
| ENSG00000196 | GDAP2 | chr1 | 117886576 | 117886653 | 117882903 | 117883627 | 117896832 | 117896989 | 18,10,16 | 1,0,0 | 9,20,22 | 0,8,6 | 227 | 150 | 1.38E-05 | 0.000283566 | 0.922,1.0,1.0 | 1.0,0.623,0.708 | 0.197 | ENSG00000196 | [[[[117929448, 11 | 2 |
| ENSG00000196 | GDAP2 | chr1 | 117887697 | 117887774 | 117882903 | 117883627 | 117896832 | 117896989 | 7,8,23 | 1,0,0 | 13,28,17 | 0,8,6 | 227 | 150 | 3.88E-05 | 0.000701389 | 0.822,1.0,1.0 | 1.0,0.698,0.652 | 0.157 | ENSG00000196 | [[[[117929448, 11 | 2 |
| ENSG00000164 | SLU7 | chr5 | 160412450 | 160412519 | 160408649 | 160408697 | 160413898 | 160413979 | 12,18,30 | 0,0,0 | 19,78,40 | 14,25,10 | 219 | 150 | 7.17E-11 | 4.81E-09 | 1.0,1.0,1.0 | 0.482,0.681,0.73 | 0.368 | ENSG00000164 | [[[[160419023, 16 | 2 |
| ENSG00000033 | ALG1 | chr16 | 5082558 | 5082673 | 5079747 | 5079807 | 5083681 | 5083757 | 43,31,19 | 0,0,0 | 53,101,36 | 6,5,2 | 265 | 150 | 3.34E-05 | 0.000614289 | 1.0,1.0,1.0 | 0.833,0.92,0.911 | 0.112 | ENSG00000033 | [[[[5071843, 5072 | 2 |
| ENSG00000164 | TENT2 | chr5 | 79619611 | 79619785 | 79612419 | 79613075 | 79640850 | 79640965 | 29,10,36 | 0,0,1 | 18,52,26 | 2,15,8 | 300 | 150 | 2.39E-08 | 9.79E-07 | 1.0,1.0,0.947 | 0.818,0.634,0.61 | 0.292 | ENSG00000164 | [[[[79612441, 796 | 3 |
| ENSG00000164 | TENT2 | chr5 | 79619611 | 79619785 | 79612419 | 79613075 | 79642843 | 79642910 | 29,12,36 | 0,0,0 | 15,38,24 | 6,5,5 | 300 | 150 | 1.42E-08 | 6.14E-07 | 1.0,1.0,1.0 | 0.556,0.792,0.70 | 0.315 | ENSG00000164 | [[[[79612441, 796 | 5 |
| ENSG00000164 | TENT2 | chr5 | 79619993 | 79620083 | 79619611 | 79619785 | 79640850 | 79640965 | 16,11,17 | 0,0,0 | 14,33,28 | 6,20,5 | 240 | 150 | 1.71E-09 | 9.06E-08 | 1.0,1.0,1.0 | 0.593,0.508,0.77 | 0.374 | ENSG00000164 | [[[[79612441, 796 | 2 |
| ENSG00000164 | TENT2 | chr5 | 79623251 | 79623489 | 79612419 | 79613075 | 79640850 | 79640965 | 37,30,45 | 0,0,1 | 28,80,55 | 2,15,8 | 300 | 150 | 6.27E-08 | 2.32E-06 | 1.0,1.0,0.957 | 0.875,0.727,0.77 | 0.193 | ENSG00000164 | [[[[79612441, 796 | 3 |
| ENSG00000164 | TENT2 | chr5 | 79642843 | 79642910 | 79613010 | 79613075 | 79645122 | 79645192 | 13,7,8 | 0,0,0 | 17,40,20 | 2,3,4 | 217 | 150 | 0.000104132 | 0.001652024 | 1.0,1.0,1.0 | 0.855,0.902,0.77 | 0.156 | ENSG00000164 | [[[[79612441, 796 | 6 |
| ENSG00000113 | MRPS27 | chr5 | 72295530 | 72295589 | 72238013 | 72238128 | 72320148 | 72320191 | 16,24,25 | 0,0,0 | 17,35,32 | 3,14,3 | 209 | 150 | 6.67E-08 | 2.45E-06 | 1.0,1.0,1.0 | 0.803,0.642,0.88 | 0.224 | ENSG00000113 | [[[[72320149, 723 | 3 |
| ENSG00000113 | MRPS27 | chr5 | 72297631 | 72297702 | 72295314 | 72295589 | 72320148 | 72320191 | 27,25,44 | 0,0,0 | 23,54,60 | 2,3,7 | 221 | 150 | 3.74E-05 | 0.000679703 | 1.0,1.0,1.0 | 0.886,0.924,0.85 | 0.112 | ENSG00000113 | [[[[72320149, 723 | 2 |
| ENSG00000080 | SRCAP | chr16 | 30712261 | 30712439 | 30711834 | 30712157 | 30713207 | 30713377 | 34,23,48 | 0,0,0 | 23,58,44 | 1,4,6 | 300 | 150 | 9.58E-06 | 0.000205828 | 1.0,1.0,1.0 | 0.92,0.879,0.786 | 0.138 | ENSG00000080 | [[[[30699171, 306 | 2 |
| ENSG00000080 | SRCAP | chr16 | 30712678 | 30712815 | 30711834 | 30712157 | 30713207 | 30713377 | 12,9,37 | 0,0,0 | 8,25,27 | 1,4,6 | 287 | 150 | 5.60E-07 | 1.64E-05 | 1.0,1.0,1.0 | 0.807,0.766,0.70 | 0.242 | ENSG00000080 | [[[[30699171, 306 | 2 |
| ENSG00000180 | RCC1 | chr1 | 28508829 | 28508905 | 28505942 | 28506084 | 28516724 | 28516853 | 6,10,20 | 1,0,0 | 20,45,25 | 1,5,8 | 226 | 150 | 0.000115961 | 0.001814004 | 0.799,1.0,1.0 | 0.93,0.857,0.675 | 0.112 | ENSG00000180 | [[[[28506043, 285 | 2 |
| ENSG00000180 | RCC1 | chr1 | 28530522 | 28530573 | 28529857 | 28529939 | 28535870 | 28536026 | 5,10,34 | 0,0,0 | 7,14,7 | 1,3,2 | 201 | 150 | 0.000553869 | 0.007025756 | 1.0,1.0,1.0 | 0.839,0.777,0.72 | 0.22 | ENSG00000180 | [[[[28506043, 285 | 4 |
| ENSG00000180 | RCC1 | chr1 | 28531802 | 28531990 | 28529857 | 28529939 | 28535049 | 28535146 | 78,155,236 | 0,1,0 | 182,451,294 | 12,15,15 | 300 | 150 | 1.40E-08 | 6.08E-07 | 1.0,0.987,1.0 | 0.883,0.938,0.90 | 0.086 | ENSG00000180 | [[[[28506043, 285 | 2 |
| ENSG00000121 | ECHDC2 | chr1 | 52905033 | 52905090 | 52904645 | 52904833 | 52907867 | 52907954 | 2,8,14 | 2,0,4 | 4,4,9 | 20,21,16 | 207 | 150 | 3.17E-05 | 0.000587189 | 0.42,1.0,0.717 | 0.127,0.121,0.29 | 0.533 | ENSG00000121 | [[[[52921553, 529 | 2 |
| ENSG00000121 | ECHDC2 | chr1 | 52911722 | 52911790 | 52907447 | 52907954 | 52913894 | 52913930 | 1,2,2 | 0,0,0 | 7,18,4 | 6,10,3 | 218 | 150 | 1.10E-07 | 3.80E-06 | 1.0,1.0,1.0 | 0.445,0.553,0.47 | 0.508 | ENSG00000121 | [[[[52921553, 529 | 2 |

| | | | | | | | | | | | | | | | | | | | | | | |
|---|---|---|---|---|---|---|---|---|---|---|---|---|---|---|---|---|---|---|---|---|---|---|
| ENSG00000121 | ECHDC2 | chr1 | 52911722 | 52912021 | 52907867 | 52907954 | 52913894 | 52914095 | 4,1,4 | 0,0,0 | 10,23,7 | 6,10,3 | 300 | 150 | 6.69E-08 | 2.45E-06 | 1.0,1.0,1.0 | 0.455,0.535,0.53 | 0.491 | ENSG00000121 | [[[[52921553, 529 | 2 |
| ENSG00000171 | HDAC3 | chr5 | 141627892 | 141627957 | 141626193 | 141626283 | 141628558 | 141628562 | 65,73,112 | 0,2,0 | 75,131,98 | 6,11,7 | 215 | 150 | 9.99E-07 | 2.75E-05 | 1.0,0.962,1.0 | 0.897,0.893,0.90 | 0.088 | ENSG00000171 | [[[[141636736, 14 | 2 |
| ENSG00000171 | HDAC3 | chr5 | 141628113 | 141628187 | 141626193 | 141626283 | 141628558 | 141628562 | 107,79,133 | 0,2,0 | 87,168,92 | 6,11,7 | 224 | 150 | 5.85E-07 | 1.70E-05 | 1.0,0.964,1.0 | 0.907,0.911,0.89 | 0.083 | ENSG00000171 | [[[[141636736, 14 | 2 |
| ENSG00000151 | MBIP | chr14 | 36311572 | 36311725 | 36300784 | 36300823 | 36314511 | 36314608 | 3,2,8 | 0,0,0 | 3,8,2 | 2,2,2 | 300 | 150 | 6.67E-06 | 0.000149402 | 1.0,1.0,1.0 | 0.429,0.667,0.33 | 0.524 | ENSG00000151 | [[[[36320460, 363 | 3 |
| ENSG00000108 | UBTF | chr17 | 44215653 | 44215809 | 44213217 | 44213282 | 44216528 | 44216704 | 90,57,83 | 0,0,0 | 47,151,76 | 12,31,12 | 300 | 150 | 1.11E-16 | 1.65E-14 | 1.0,1.0,1.0 | 0.662,0.709,0.76 | 0.29 | ENSG00000108 | [[[[44219445, 442 | 2 |
| ENSG00000182 | SKA2 | chr17 | 59119318 | 59119495 | 59109950 | 59112345 | 59155130 | 59155184 | 114,124,166 | 33,22,20 | 107,230,138 | 9,21,9 | 300 | 150 | 0.00067956 | 0.008345066 | 0.633,0.738,0.80 | 0.856,0.846,0.88 | -0.137 | ENSG00000182 | [[[[59155131, 591 | 2 |
| ENSG00000163 | HDAC11 | chr3 | 13500712 | 13500789 | 13483463 | 13483564 | 13501870 | 13501933 | 2,4,9 | 0,0,1 | 2,7,11 | 2,2,2 | 227 | 150 | 0.000518247 | 0.006625735 | 1.0,1.0,0.856 | 0.398,0.698,0.78 | 0.325 | ENSG00000163 | [[[[13480306, 134 | 3 |
| ENSG00000146 | NCAPG2 | chr7 | 158644288 | 158644388 | 158631310 | 158631717 | 158650831 | 158650972 | 122,122,149 | 2,0,0 | 61,121,71 | 16,25,14 | 250 | 150 | 0 | 0 | 0.973,1.0,1.0 | 0.696,0.744,0.75 | 0.26 | ENSG00000146 | [[[[158704724, 15 | 3 |
| ENSG00000146 | NCAPG2 | chr7 | 158644288 | 158644388 | 158631310 | 158631717 | 158652292 | 158652480 | 122,122,149 | 3,0,1 | 59,118,70 | 4,15,4 | 250 | 150 | 1.20E-07 | 4.11E-06 | 0.961,1.0,0.989 | 0.898,0.825,0.91 | 0.105 | ENSG00000146 | [[[[158704724, 15 | 4 |
| ENSG00000146 | NCAPG2 | chr7 | 158650831 | 158650972 | 158631310 | 158631717 | 158652292 | 158652480 | 99,94,153 | 3,0,1 | 86,212,109 | 4,15,4 | 291 | 150 | 5.18E-05 | 0.000900665 | 0.944,1.0,0.987 | 0.917,0.879,0.93 | 0.067 | ENSG00000146 | [[[[158704724, 15 | 4 |
| ENSG00000146 | NCAPG2 | chr7 | 158671513 | 158671666 | 158664527 | 158664750 | 158679959 | 158680085 | 94,83,196 | 0,0,1 | 59,185,134 | 9,17,10 | 300 | 150 | 1.06E-11 | 8.09E-10 | 1.0,1.0,0.99 | 0.766,0.845,0.87 | 0.17 | ENSG00000146 | [[[[158704724, 15 | 2 |
| ENSG00000146 | NCAPG2 | chr7 | 158675476 | 158675616 | 158664527 | 158664750 | 158679959 | 158680085 | 2,2,10 | 0,0,1 | 1,5,1 | 9,17,10 | 290 | 150 | 8.03E-12 | 6.25E-10 | 1.0,1.0,0.838 | 0.054,0.132,0.04 | 0.868 | ENSG00000146 | [[[[158704724, 15 | 2 |
| ENSG00000146 | NCAPG2 | chr7 | 158675476 | 158675656 | 158664527 | 158664750 | 158679959 | 158680085 | 71,54,134 | 0,0,1 | 43,105,57 | 9,17,10 | 300 | 150 | 4.30E-14 | 4.68E-12 | 1.0,1.0,0.985 | 0.705,0.755,0.74 | 0.262 | ENSG00000146 | [[[[158704724, 15 | 2 |
| ENSG00000146 | NCAPG2 | chr7 | 158689818 | 158689953 | 158687347 | 158687442 | 158693308 | 158693339 | 43,25,45 | 0,0,0 | 26,89,51 | 0,9,5 | 285 | 150 | 7.61E-06 | 0.000168356 | 1.0,1.0,1.0 | 1.0,0.839,0.843 | 0.106 | ENSG00000146 | [[[[158704724, 15 | 3 |
| ENSG00000146 | NCAPG2 | chr7 | 158692841 | 158692956 | 158689818 | 158689953 | 158693308 | 158693339 | 52,26,62 | 0,0,0 | 36,77,58 | 1,7,4 | 265 | 150 | 2.34E-05 | 0.000452422 | 1.0,1.0,1.0 | 0.953,0.862,0.89 | 0.098 | ENSG00000146 | [[[[158704724, 15 | 2 |
| ENSG00000105 | NAMPT | chr7 | 106261587 | 106261707 | 106254363 | 106254504 | 106272529 | 106272658 | 41,21,58 | 0,0,0 | 21,43,27 | 3,6,10 | 270 | 150 | 1.52E-09 | 8.16E-08 | 1.0,1.0,1.0 | 0.795,0.799,0.6 | 0.269 | ENSG00000105 | [[[[106284828, 1( | 4 |
| ENSG00000105 | NAMPT | chr7 | 106269153 | 106269312 | 106254363 | 106254504 | 106272529 | 106272658 | 51,51,114 | 0,0,0 | 54,157,114 | 3,6,10 | 300 | 150 | 1.11E-06 | 3.00E-05 | 1.0,1.0,1.0 | 0.9,0.929,0.851 | 0.107 | ENSG00000105 | [[[[106284828, 1( | 4 |
| ENSG00000080 | CRYBG3 | chr3 | 97895958 | 97896085 | 97889354 | 97889390 | 97898882 | 97899025 | 6,4,15 | 0,0,0 | 7,6,12 | 4,2,0 | 277 | 150 | 0.000411729 | 0.005433485 | 1.0,1.0,1.0 | 0.487,0.619,1.0 | 0.298 | ENSG00000080 | [[[[97822011, 978 | 2 |
| ENSG00000009 | REV3L | chr6 | 111367114 | 111368028 | 111365264 | 111365344 | 111381324 | 111381444 | 10,5,12 | 0,0,0 | 14,20,28 | 0,4,4 | 300 | 150 | 4.14E-05 | 0.000742454 | 1.0,1.0,1.0 | 1.0,0.714,0.778 | 0.169 | ENSG00000009 | [[[[111482750, 11 | 4 |
| ENSG00000164 | SPIDR | chr8 | 47407860 | 47407961 | 47396375 | 47396626 | 47595810 | 47595984 | 16,11,28 | 0,1,0 | 15,38,26 | 2,6,9 | 251 | 150 | 2.25E-06 | 5.68E-05 | 1.0,0.868,1.0 | 0.818,0.791,0.63 | 0.209 | ENSG00000164 | [[[[47260938, 472 | 2 |
| ENSG00000164 | SPIDR | chr8 | 47440322 | 47440542 | 47396375 | 47396626 | 47595810 | 47595984 | 24,23,38 | 0,1,0 | 19,73,47 | 2,6,9 | 300 | 150 | 4.47E-06 | 0.000104521 | 1.0,0.92,1.0 | 0.826,0.859,0.72 | 0.171 | ENSG00000164 | [[[[47260938, 472 | 2 |
| ENSG00000156 | EIF4A2 | chr3 | 186785882 | 186786051 | 186784563 | 186784696 | 186788309 | 186788416 | 111,110,194 | 0,0,0 | 377,956,539 | 7,20,6 | 300 | 150 | 0.000366677 | 0.004912099 | 1.0,1.0,1.0 | 0.964,0.96,0.978 | 0.033 | ENSG00000156 | [[[[186783577, 18 | 7 |
| ENSG00000156 | EIF4A2 | chr3 | 186787478 | 186787584 | 186787126 | 186787264 | 186789124 | 186789140 | 9,8,11 | 0,0,0 | 38,53,28 | 5,12,8 | 256 | 150 | 5.17E-07 | 1.53E-05 | 1.0,1.0,1.0 | 0.817,0.721,0.67 | 0.263 | ENSG00000156 | [[[[186783577, 18 | 2 |
| ENSG00000156 | EIF4A2 | chr3 | 186787494 | 186787584 | 186787126 | 186787264 | 186788309 | 186788416 | 235,244,345 | 0,0,1 | 8,301,670,992 | 70,164,107 | 240 | 150 | 0 | 0 | 1.0,1.0,0.995 | 0.881,0.864,0.85 | 0.132 | ENSG00000156 | [[[[186783577, 18 | 2 |
| ENSG00000108 | DRG2 | chr17 | 18098269 | 18098359 | 18093812 | 18093968 | 18099632 | 18099723 | 3,5,5 | 0,0,0 | 6,7,0 | 0,4,9 | 240 | 150 | 2.26E-06 | 5.70E-05 | 1.0,1.0,1.0 | 1.0,0.522,0.0 | 0.493 | ENSG00000108 | [[[[18087948, 18( | 2 |
| ENSG00000108 | DRG2 | chr17 | 18099016 | 18099077 | 18093812 | 18093968 | 18099632 | 18099662 | 15,9,38 | 0,0,0 | 28,57,36 | 0,4,9 | 211 | 150 | 3.71E-05 | 0.000674954 | 1.0,1.0,1.0 | 1.0,0.91,0.74 | 0.117 | ENSG00000108 | [[[[18087948, 18( | 2 |
| ENSG00000205 | PDE7A | chr8 | 65727169 | 65727301 | 65717509 | 65719495 | 65734793 | 65734894 | 7,5,20 | 0,0,0 | 18,22,14 | 3,6,3 | 282 | 150 | 4.59E-07 | 1.37E-05 | 1.0,1.0,1.0 | 0.761,0.661,0.71 | 0.288 | ENSG00000205 | [[[[65841371, 658 | 5 |
| ENSG00000125 | THOC2 | chrX | 123622757 | 123622860 | 123619400 | 123619436 | 123623104 | 123623283 | 29,15,18 | 0,0,0 | 15,15,9 | 1,6,5 | 253 | 150 | 6.61E-08 | 2.43E-06 | 1.0,1.0,1.0 | 0.899,0.597,0.51 | 0.329 | ENSG00000125 | [[[[123732952, 12 | 3 |
| ENSG00000125 | THOC2 | chrX | 123623104 | 123623283 | 123619400 | 123619436 | 123623786 | 123623971 | 25,20,52 | 0,0,0 | 8,26,15 | 3,4,0 | 300 | 150 | 0.000244811 | 0.003457021 | 1.0,1.0,1.0 | 0.571,0.765,1.0 | 0.221 | ENSG00000125 | [[[[123732952, 12 | 4 |
| ENSG00000125 | THOC2 | chrX | 123631687 | 123631852 | 123627692 | 123627968 | 123633952 | 123634070 | 28,18,37 | 0,0,0 | 9,14,8 | 1,4,1 | 300 | 150 | 0.00024115 | 0.003413547 | 1.0,1.0,1.0 | 0.818,0.636,0.8 | 0.249 | ENSG00000125 | [[[[123732952, 12 | 2 |
| ENSG00000107 | MPDZ | chr9 | 13113010 | 13113054 | 13112023 | 13112146 | 13115247 | 13115334 | 4,5,12 | 0,0,0 | 3,10,6 | 1,4,5 | 194 | 150 | 4.70E-08 | 1.79E-06 | 1.0,1.0,1.0 | 0.699,0.659,0.48 | 0.387 | ENSG00000107 | [[[[13279400, 132 | 2 |
| ENSG00000171 | SLC25A33 | chr1 | 9567283 | 9567361 | 9553625 | 9553805 | 9579953 | 9580234 | 29,24,28 | 0,0,0 | 25,81,59 | 2,9,5 | 228 | 150 | 5.20E-06 | 0.000119954 | 1.0,1.0,1.0 | 0.892,0.856,0.88 | 0.122 | ENSG00000171 | [[[[9539465, 953 | 3 |
| ENSG00000171 | SLC25A33 | chr1 | 9570257 | 9570358 | 9567283 | 9567361 | 9579953 | 9580234 | 34,38,42 | 0,0,0 | 44,84,52 | 5,18,10 | 251 | 150 | 1.97E-10 | 1.23E-08 | 1.0,1.0,1.0 | 0.84,0.736,0.757 | 0.222 | ENSG00000171 | [[[[9539465, 953 | 2 |
| ENSG00000171 | SLC25A33 | chr1 | 9573345 | 9573412 | 9553625 | 9553805 | 9579953 | 9580234 | 17,17,15 | 0,0,0 | 35,59,48 | 2,9,5 | 217 | 150 | 1.04E-05 | 0.000221668 | 1.0,1.0,1.0 | 0.924,0.819,0.86 | 0.129 | ENSG00000171 | [[[[9539465, 953 | 3 |
| ENSG00000171 | SLC25A33 | chr1 | 9573345 | 9573412 | 9567283 | 9567361 | 9579953 | 9580234 | 18,18,22 | 0,0,0 | 48,68,64 | 5,18,10 | 217 | 150 | 5.97E-08 | 2.22E-06 | 1.0,1.0,1.0 | 0.869,0.723,0.81 | 0.197 | ENSG00000171 | [[[[9539465, 953 | 2 |
| ENSG00000197 | DNM3 | chr1 | 172081831 | 172081902 | 172068818 | 172068905 | 172131174 | 172131288 | 7,4,3 | 0,0,0 | 1,1,2 | 0,4,2 | 221 | 150 | 2.39E-05 | 0.000459721 | 1.0,1.0,1.0 | 1.0,0.145,0.404 | 0.484 | ENSG00000197 | [[[[171841498, 17 | 2 |
| ENSG00000066 | MTFR1 | chr8 | 65669872 | 65670018 | 65644733 | 65644784 | 65693643 | 65693759 | 20,28,53 | 0,0,0 | 25,47,42 | 1,3,5 | 296 | 150 | 0.000103969 | 0.00164994 | 1.0,1.0,1.0 | 0.927,0.888,0.81 | 0.125 | ENSG00000066 | [[[[65644734, 656 | 2 |
| ENSG00000066 | MTFR1 | chr8 | 65682352 | 65682451 | 65644733 | 65644784 | 65693643 | 65693759 | 10,15,20 | 0,0,0 | 19,51,23 | 1,3,5 | 249 | 150 | 6.91E-05 | 0.001157295 | 1.0,1.0,1.0 | 0.92,0.911,0.735 | 0.145 | ENSG00000066 | [[[[65644734, 656 | 2 |
| ENSG00000125 | HNRNPR | chr1 | 23323555 | 23323732 | 23321527 | 23321663 | 23337753 | 23337861 | 301,258,381 | 0,0,1 | 227,534,329 | 24,49,33 | 300 | 150 | 0 | 0 | 1.0,1.0,0.995 | 0.825,0.845,0.83 | 0.164 | ENSG00000125 | [[[[23344211, 233 | 2 |
| ENSG00000125 | HNRNPR | chr1 | 23333517 | 23333631 | 23321527 | 23321663 | 23337753 | 23337861 | 322,291,425 | 0,0,1 | 269,722,391 | 24,49,33 | 264 | 150 | 0 | 0 | 1.0,1.0,0.996 | 0.864,0.893,0.87 | 0.123 | ENSG00000125 | [[[[23344211, 233 | 2 |
| ENSG00000116 | DOCK7 | chr1 | 62475207 | 62475351 | 62457537 | 62457705 | 62475706 | 62475943 | 40,35,38 | 0,0,0 | 19,75,32 | 0,8,4 | 294 | 150 | 8.01E-06 | 0.000175856 | 1.0,1.0,1.0 | 1.0,0.827,0.803 | 0.123 | ENSG00000116 | [[[[62688227, 626 | 2 |
| ENSG00000151 | FLI1 | chr11 | 128805365 | 128805431 | 128781957 | 128782023 | 128809156 | 128809204 | 3,7,4 | 0,1,0 | 1,1,2 | 0,3,3 | 216 | 150 | 3.97E-05 | 0.00071513 | 1.0,0.829,1.0 | 1.0,0.188,0.316 | 0.442 | ENSG00000151 | [[[[128694072, 12 | 2 |
| ENSG00000151 | FLI1 | chr11 | 128807179 | 128807239 | 128781957 | 128782023 | 128809156 | 128809204 | 1,8,8 | 0,1,0 | 2,4,1 | 0,3,3 | 210 | 150 | 0.000237438 | 0.003367984 | 1.0,0.851,1.0 | 1.0,0.488,0.192 | 0.39 | ENSG00000151 | [[[[128694072, 12 | 2 |
| ENSG00000175 | PSMD2 | chr3 | 184302369 | 184302528 | 184301536 | 184301658 | 184302678 | 184302692 | 529,450,694 | 0,0,0 | 4,431,066,691 | 29,52,29 | 300 | 150 | 0 | 0 | 1.0,1.0,1.0 | 0.884,0.911,0.92 | 0.094 | ENSG00000175 | [[[[184299241, 18 | 2 |
| ENSG00000175 | PSMD2 | chr3 | 184307613 | 184307708 | 184306750 | 184306834 | 184307889 | 184307902 | 341,575,698 | 0,0,0 | 353,772,395 | 8,15,9 | 245 | 150 | 2.72E-06 | 6.73E-05 | 1.0,1.0,1.0 | 0.964,0.969,0.96 | 0.034 | ENSG00000175 | [[[[184299241, 18 | 2 |
| ENSG00000175 | PSMD2 | chr3 | 184307880 | 184308016 | 184307356 | 184307525 | 184308448 | 184308534 | 411,604,662 | 0,0,0 | 357,838,435 | 9,17,8 | 277 | 150 | 3.21E-07 | 9.87E-06 | 1.0,1.0,1.0 | 0.956,0.964,0.96 | 0.038 | ENSG00000175 | [[[[184299241, 18 | 2 |
| ENSG00000092 | HNRNPC | chr14 | 21263310 | 21263336 | 21212959 | 21213117 | 21269297 | 21269403 | 10,11,12 | 0,0,0 | 15,10,10 | 3,3,1 | 176 | 150 | 0.000299085 | 0.00411812 | 1.0,1.0,1.0 | 0.81,0.74,0.895 | 0.185 | ENSG00000092 | [[[[21269298, 212 | 4 |
| ENSG00000073 | PICALM | chr11 | 86022366 | 86022469 | 86014869 | 86014963 | 86068650 | 86068733 | 58,51,60 | 3,0,7 | 34,56,55 | 7,14,12 | 253 | 150 | 9.17E-05 | 0.00147818 | 0.92,1.0,0.836 | 0.742,0.703,0.73 | 0.193 | ENSG00000073 | [[[[86068651, 86( | 3 |
| ENSG00000157 | LRP8 | chr1 | 53276691 | 53277078 | 53275630 | 53275753 | 53289566 | 53289689 | 9,2,24 | 0,0,2 | 12,12,13 | 4,5,6 | 300 | 150 | 9.51E-06 | 0.000204762 | 1.0,1.0,0.857 | 0.6,0.545,0.52 | 0.397 | ENSG00000157 | [[[[53327789, 533 | 2 |
| ENSG00000117 | MESD | chr15 | 80970461 | 80970659 | 80951958 | 80952296 | 80989578 | 80989878 | 1,2,8 | 0,0,1 | 19,15,13 | 1,2,5 | 300 | 150 | 0.000222182 | 0.003192498 | 1.0,1.0,0.8 | 0.905,0.789,0.56 | 0.18 | ENSG00000117 | [[[[80989579, 809 | 2 |
| ENSG00000090 | PABPC4 | chr1 | 39564700 | 39564773 | 39564422 | 39564530 | 39567750 | 39567846 | 6,8,6 | 0,0,0 | 5,3,3 | 2,6,0 | 223 | 150 | 2.21E-05 | 0.000429133 | 1.0,1.0,1.0 | 0.627,0.252,1.0 | 0.374 | ENSG00000090 | [[[[39575759, 395 | 2 |
| ENSG00000159 | GPBP1L1 | chr1 | 45654542 | 45654829 | 45642426 | 45642499 | 45659027 | 45659142 | 62,66,150 | 0,0,0 | 148,297,175 | 8,22,8 | 300 | 150 | 9.05E-09 | 4.05E-07 | 1.0,1.0,1.0 | 0.902,0.871,0.91 | 0.104 | ENSG00000159 | [[[[45686412, 456 | 2 |
| ENSG00000159 | GPBP1L1 | chr1 | 45685575 | 45685620 | 45660183 | 45661225 | 45686992 | 45687133 | 20,7,26 | 18,25,34 | 28,76,56 | 9,26,41 | 195 | 150 | 0.000521767 | 0.006665153 | 0.461,0.177,0.37 | 0.705,0.692,0.51 | -0.3 | ENSG00000159 | [[[[45686412, 456 | 2 |
| ENSG00000162 | ZNF326 | chr1 | 90007344 | 90007750 | 90005002 | 90005092 | 90010087 | 90010286 | 46,41,54 | 0,6,2 | 49,133,72 | 6,25,18 | 300 | 150 | 0.000587094 | 0.007371132 | 1.0,0.74,0.931 | 0.803,0.727,0.66 | 0.169 | ENSG00000162 | [[[[89995110, 899 | 2 |
| ENSG00000112 | RPS12 | chr6 | 132816959 | 132817061 | 132814971 | 132815088 | 132817479 | 132817564 | 293,306,345 | 0,0,0 | 292,738,381 | 122,227,144 | 252 | 150 | 0 | 0 | 1.0,1.0,1.0 | 0.588,0.659,0.61 | 0.38 | ENSG00000112 | [[[[132814569, 13 | 2 |
| ENSG00000197 | RPF2 | chr6 | 110997182 | 110997264 | 110991746 | 110991786 | 111008037 | 111008137 | 22,17,33 | 0,0,0 | 25,57,30 | 12,7,6 | 232 | 150 | 9.44E-09 | 4.22E-07 | 1.0,1.0,1.0 | 0.574,0.84,0.764 | 0.274 | ENSG00000197 | [[[[110982038, 11 | 2 |
| ENSG00000197 | RPF2 | chr6 | 110999710 | 110999787 | 110991746 | 110991786 | 111008037 | 111008137 | 24,28,29 | 0,0,0 | 18,35,26 | 12,7,6 | 227 | 150 | 3.46E-10 | 2.09E-08 | 1.0,1.0,1.0 | 0.498,0.768,0.74 | 0.331 | ENSG00000197 | [[[[110982038, 11 | 2 |
| ENSG00000179 | RCC2 | chr1 | 17429105 | 17429199 | 17416479 | 17416646 | 17438229 | 17438522 | 2,3,6 | 0,0,0 | 4,19,13 | 1,6,0 | 244 | 150 | 0.000349034 | 0.004701377 | 1.0,1.0,1.0 | 0.711,0.661,1.0 | 0.209 | ENSG00000179 | [[[[17439545, 174 | 5 |
| ENSG00000092 | PPP2R3C | chr14 | 35096708 | 35096764 | 35096557 | 35096633 | 35109847 | 35109931 | 12,2,2 | 0,0,0 | 3,12,4 | 1,4,2 | 206 | 150 | 1.83E-05 | 0.000366334 | 1.0,1.0,1.0 | 0.686,0.686,0.59 | 0.345 | ENSG00000092 | [[[[35121902, 351 | 4 |
| ENSG00000092 | PPP2R3C | chr14 | 35099251 | 35099384 | 35096708 | 35096764 | 35109847 | 35109931 | 27,24,28 | 0,0,0 | 34,61,20 | 2,5,2 | 283 | 150 | 0.000181629 | 0.002679586 | 1.0,1.0,1.0 | 0.9,0.866,0.841 | 0.131 | ENSG00000092 | [[[[35121902, 351 | 3 |
| ENSG00000137 | DENND4C | chr9 | 19352489 | 19352665 | 19350701 | 19350879 | 19356971 | 19357154 | 30,9,18 | 0,0,0 | 15,19,17 | 1,7,4 | 300 | 150 | 1.01E-07 | 3.55E-06 | 1.0,1.0,1.0 | 0.882,0.576,0.68 | 0.287 | ENSG00000137 | [[[[19230693, 192 | 2 |
| ENSG00000163 | NEPRO | chr3 | 113013948 | 113014032 | 113011044 | 113011406 | 113017447 | 113017602 | 9,3,17 | 0,0,0 | 16,26,20 | 2,2,4 | 234 | 150 | 0.00014422 | 0.002186925 | 1.0,1.0,1.0 | 0.837,0.893,0.76 | 0.169 | ENSG00000163 | [[[[113019542, 11 | 3 |
| ENSG00000163 | NEPRO | chr3 | 113013952 | 113014032 | 113011044 | 113011406 | 113017447 | 113017602 | 9,3,18 | 0,0,0 | 16,30,22 | 2,2,4 | 230 | 150 | 0.000239994 | 0.003400975 | 1.0,1.0,1.0 | 0.839,0.907,0.78 | 0.157 | ENSG00000163 | [[[[113019542, 11 | 3 |
| ENSG00000142 | PADI3 | chr1 | 17262132 | 17262205 | 17249097 | 17249229 | 17265658 | 17265720 | 316,258,370 | 74,72,104 | 88,179,128 | 56,81,56 | 223 | 150 | 2.68E-05 | 0.000507113 | 0.742,0.707,0.70 | 0.514,0.598,0.60 | 0.145 | ENSG00000142 | [[[[17249098, 172 | 2 |
| ENSG00000142 | PADI3 | chr1 | 17274634 | 17274786 | 17273339 | 17273447 | 17280349 | 17280429 | 429,428,782 | 1,2,1 | 150,225,161 | 17,22,11 | 300 | 150 | 0 | 0 | 0.995,0.991,0.99 | 0.815,0.836,0.88 | 0.151 | ENSG00000142 | [[[[17249098, 172 | 3 |
| ENSG00000142 | PADI3 | chr1 | 17276518 | 17276663 | 17271066 | 17271178 | 17280349 | 17280429 | 4,2,7 | 0,0,0 | 5,6,8 | 4,4,1 | 295 | 150 | 3.43E-08 | 1.35E-06 | 1.0,1.0,1.0 | 0.389,0.433,0.80 | 0.458 | ENSG00000142 | [[[[17249098, 172 | 4 |
| ENSG00000142 | PADI3 | chr1 | 17276518 | 17276663 | 17273339 | 17273447 | 17280349 | 17280429 | 13,15,20 | 1,2,1 | 12,12,12 | 17,22,11 | 295 | 150 | 7.92E-11 | 5.29E-09 | 0.869,0.792,0.91 | 0.264,0.217,0.35 | 0.578 | ENSG00000142 | [[[[17249098, 172 | 3 |
| ENSG00000142 | PADI3 | chr1 | 17276518 | 17276663 | 17274634 | 17274786 | 17280349 | 17280429 | 3,936,401,068 | 0,1,0 | 172,253,217 | 5,11,7 | 295 | 150 | 2.35E-09 | 1.21E-07 | 1.0,0.997,1.0 | 0.946,0.921,0.94 | 0.063 | ENSG00000142 | [[[[17249098, 172 | 2 |
| ENSG00000142 | PADI3 | chr1 | 17276773 | 17276876 | 17273339 | 17273447 | 17280349 | 17280429 | 358,266,405 | 1,2,1 | 105,131,88 | 17,22,11 | 253 | 150 | 0 | 0 | 0.995,0.987,0.99 | 0.785,0.779,0.82 | 0.196 | ENSG00000142 | [[[[17249098, 172 | 2 |
| ENSG00000116 | CRYZ | chr1 | 74706898 | 74706994 | 74705481 | 74706457 | 74710097 | 74710247 | 84,50,111 | 0,0,0 | 65,167,127 | 10,23,9 | 246 | 150 | 8.04E-12 | 6.26E-10 | 1.0,1.0,1.0 | 0.799,0.816,0.89 | 0.163 | ENSG00000116 | [[[[74732956, 747 | 2 |
| ENSG00000116 | CRYZ | chr1 | 74707102 | 74707204 | 74706898 | 74706994 | 74723117 | 74723270 | 66,75,109 | 0,0,0 | 42,90,103 | 1,13,7 | 252 | 150 | 3.31E-08 | 1.31E-06 | 1.0,1.0,1.0 | 0.962,0.805,0.89 | 0.112 | ENSG00000116 | [[[[74732956, 747 | 4 |
| ENSG00000116 | CRYZ | chr1 | 74710097 | 74710247 | 74706898 | 74706994 | 74723117 | 74723270 | 13,9,29 | 0,0,0 | 24,57,38 | 1,13,7 | 300 | 150 | 3.54E-07 | 1.09E-05 | 1.0,1.0,1.0 | 0.923,0.687,0.73 | 0.22 | ENSG00000116 | [[[[74732956, 747 | 4 |
| ENSG00000116 | CRYZ | chr1 | 74710097 | 74710247 | 74707102 | 74707204 | 74723117 | 74723270 | 91,79,133 | 0,0,1 | 73,116,80 | 2,8,9 | 300 | 150 | 2.77E-07 | 8.74E-06 | 1.0,1.0,0.985 | 0.948,0.879,0.81 | 0.114 | ENSG00000116 | [[[[74732956, 747 | 3 |
| ENSG00000116 | CRYZ | chr1 | 74714578 | 74714630 | 74706898 | 74706994 | 74723117 | 74723270 | 1,7,2 | 0,0,0 | 3,5,1 | 1,13,7 | 202 | 150 | 7.18E-11 | 4.81E-09 | 1.0,1.0,1.0 | 0.69,0.222,0.096 | 0.664 | ENSG00000116 | [[[[74732956, 747 | 4 |
| ENSG00000113 | PPWD1 | chr5 | 65567512 | 65567615 | 65563238 | 65563506 | 65576878 | 65577069 | 15,12,44 | 0,0,2 | 18,38,31 | 3,8,0 | 253 | 150 | 0.00065877 | 0.008126431 | 1.0,1.0,0.929 | 0.781,0.738,1.0 | 0.137 | ENSG00000113 | [[[[65563296, 655 | 4 |
| ENSG00000113 | PPWD1 | chr5 | 65569631 | 65569732 | 65567512 | 65567615 | 65576878 | 65577069 | 36,19,26 | 0,0,0 | 7,15,18 | 2,9,6 | 251 | 150 | 4.28E-11 | 2.97E-09 | 1.0,1.0,1.0 | 0.677,0.499,0.64 | 0.394 | ENSG00000113 | [[[[65563296, 655 | 3 |
| ENSG00000113 | PPWD1 | chr5 | 65571838 | 65572286 | 65567512 | 65567615 | 65576878 | 65576926 | 29,24,21 | 0,0,0 | 16,40,39 | 2,9,6 | 300 | 150 | 3.50E-08 | 1.38E-06 | 1.0,1.0,1.0 | 0.8,0.69,0.765 | 0.248 | ENSG00000113 | [[[[65563296, 655 | 3 |
| ENSG00000175 | PHYKPL | chr5 | 178215275 | 178215430 | 178214795 | 178214885 | 178222851 | 178222934 | 21,19,25 | 1,0,0 | 8,18,12 | 1,7,0 | 300 | 150 | 0.000184964 | 0.002726091 | 0.913,1.0,1.0 | 0.8,0.563,1.0 | 0.183 | ENSG00000175 | [[[[178232492, 17 | 2 |
| ENSG00000175 | PHYKPL | chr5 | 178222354 | 178222580 | 178214795 | 178214885 | 178222851 | 178222934 | 14,21,28 | 1,0,0 | 21,38,21 | 1,7,0 | 300 | 150 | 0.000294823 | 0.00406752 | 0.875,1.0,1.0 | 0.913,0.731,1.0 | 0.077 | ENSG00000175 | [[[[178232492, 17 | 2 |
| ENSG00000175 | PHYKPL | chr5 | 178224641 | 178224729 | 178224447 | 178224564 | 178229939 | 178230099 | 6,10,10 | 0,0,0 | 5,8,14 | 3,4,1 | 238 | 150 | 1.72E-06 | 4.48E-05 | 1.0,1.0,1.0 | 0.512,0.558,0.89 | 0.344 | ENSG00000175 | [[[[178232492, 17 | 2 |

| | | | | | | | | | | | | | | | | | | | | | | |
|---|---|---|---|---|---|---|---|---|---|---|---|---|---|---|---|---|---|---|---|---|---|---|
| ENSG000001753 | PHYKPL | chr5 | 178225354 | 178225429 | 178224447 | 178224564 | 178229939 | 178230049 | 9,8,21 | 0,0,0 | 6,15,6 | 3,4,1 | 225 | 150 | 2.07E-06 | 5.27E-05 | 1.0,1.0,1.0 | 0.571,0.714,0.8 | 0.305 | ENSG000001753 | [[[[178232492, 17 | 2 |
| ENSG000001164 | ATP5PB | chr1 | 111454210 | 111454356 | 111449681 | 111449873 | 111456629 | 111456748 | 88,710,221,453 | 1,0,0 | 81,515,761,054 | 11,21,21 | 296 | 150 | 3.57E-08 | 1.40E-06 | 0.998,1.0,1.0 | 0.974,0.974,0.96 | 0.029 | ENSG000001164 | [[[[111449464, 11 | 2 |
| ENSG000001164 | ATP5PB | chr1 | 111456085 | 111456249 | 111449836 | 111449873 | 111456629 | 111456748 | 479,440,665 | 1,0,0 | 270,582,343 | 11,21,21 | 300 | 150 | 2.22E-16 | 3.17E-14 | 0.996,1.0,1.0 | 0.925,0.933,0.89 | 0.082 | ENSG000001164 | [[[[111449464, 11 | 2 |
| ENSG000001164 | ATP5PB | chr1 | 111456226 | 111456249 | 111449836 | 111449873 | 111456629 | 111456755 | 65,53,75 | 1,0,0 | 138,297,159 | 11,21,21 | 173 | 150 | 3.62E-08 | 1.42E-06 | 0.983,1.0,1.0 | 0.916,0.925,0.86 | 0.091 | ENSG000001164 | [[[[111449464, 11 | 2 |
| ENSG000001452 | OCIAD2 | chr4 | 48892771 | 48892889 | 48885018 | 48885565 | 48899828 | 48899925 | 693,644,794 | 0,0,0 | 5,241,039,531 | 12,21,12 | 268 | 150 | 4.82E-09 | 2.30E-07 | 1.0,1.0,1.0 | 0.961,0.965,0.96 | 0.038 | ENSG000001452 | [[[[48906658, 489 | 3 |
| ENSG000001452 | OCIAD2 | chr4 | 48892771 | 48892889 | 48885018 | 48885565 | 48904483 | 48904611 | 693,644,793 | 1,6,1 | 475,950,478 | 13,39,20 | 268 | 150 | 1.38E-14 | 1.62E-12 | 0.997,0.984,0.99 | 0.953,0.932,0.93 | 0.055 | ENSG000001452 | [[[[48906658, 489 | 4 |
| ENSG000001452 | OCIAD2 | chr4 | 48892771 | 48894053 | 48885381 | 48885565 | 48899828 | 48899925 | 696,645,793 | 0,0,0 | 476,939,474 | 12,21,12 | 300 | 150 | 9.20E-11 | 6.08E-09 | 1.0,1.0,1.0 | 0.952,0.957,0.95 | 0.046 | ENSG000001452 | [[[[48906658, 489 | 3 |
| ENSG000001452 | OCIAD2 | chr4 | 48894005 | 48894053 | 48885387 | 48885565 | 48899828 | 48899925 | 26,18,22 | 0,0,0 | 24,51,21 | 12,21,12 | 198 | 150 | 1.58E-13 | 1.61E-11 | 1.0,1.0,1.0 | 0.602,0.648,0.57 | 0.393 | ENSG000001452 | [[[[48906658, 489 | 3 |
| ENSG000001452 | OCIAD2 | chr4 | 48894005 | 48894053 | 48885387 | 48885565 | 48904483 | 48904611 | 65,57,71 | 1,6,1 | 36,90,53 | 13,39,20 | 198 | 150 | 3.82E-13 | 3.66E-11 | 0.98,0.878,0.982 | 0.677,0.636,0.66 | 0.286 | ENSG000001452 | [[[[48906658, 489 | 4 |
| ENSG000001452 | OCIAD2 | chr4 | 48894005 | 48894053 | 48892770 | 48892889 | 48899828 | 48899925 | 445,342,491 | 0,0,2 | 255,531,314 | 55,109,62 | 198 | 150 | 0 | 0 | 1.0,1.0,0.995 | 0.778,0.787,0.79 | 0.212 | ENSG000001452 | [[[[48906658, 489 | 2 |
| ENSG000001452 | OCIAD2 | chr4 | 48894005 | 48894053 | 48892770 | 48892889 | 48904483 | 48904611 | 467,361,511 | 0,0,1 | 253,562,338 | 5,18,9 | 198 | 150 | 1.36E-06 | 3.63E-05 | 1.0,1.0,0.997 | 0.975,0.959,0.96 | 0.032 | ENSG000001452 | [[[[48906658, 489 | 3 |
| ENSG000001452 | OCIAD2 | chr4 | 48897803 | 48897857 | 48885018 | 48885565 | 48899828 | 48899925 | 596,568,683 | 0,0,0 | 398,789,462 | 12,21,12 | 204 | 150 | 5.10E-09 | 2.42E-07 | 1.0,1.0,1.0 | 0.961,0.965,0.96 | 0.036 | ENSG000001452 | [[[[48906658, 489 | 3 |
| ENSG000001452 | OCIAD2 | chr4 | 48897803 | 48897857 | 48885018 | 48885565 | 48904483 | 48904611 | 105,103,152 | 1,6,1 | 45,91,48 | 13,39,20 | 204 | 150 | 0 | 0 | 0.987,0.927,0.99 | 0.718,0.632,0.63 | 0.306 | ENSG000001452 | [[[[48906658, 489 | 4 |
| ENSG000001452 | OCIAD2 | chr4 | 48897803 | 48897857 | 48892771 | 48892889 | 48899828 | 48899925 | 598,571,686 | 0,0,2 | 398,790,470 | 55,109,62 | 204 | 150 | 0 | 0 | 1.0,1.0,0.996 | 0.842,0.842,0.84 | 0.155 | ENSG000001452 | [[[[48906658, 489 | 2 |
| ENSG000001452 | OCIAD2 | chr4 | 48897803 | 48897857 | 48892771 | 48892889 | 48904483 | 48904611 | 105,105,158 | 0,0,1 | 45,91,49 | 5,18,9 | 204 | 150 | 7.99E-15 | 9.68E-13 | 1.0,1.0,0.991 | 0.869,0.788,0.8 | 0.178 | ENSG000001452 | [[[[48906658, 489 | 3 |
| ENSG000001452 | OCIAD2 | chr4 | 48899828 | 48899925 | 48885018 | 48885565 | 48904483 | 48904611 | 107,510,341,453 | 1,6,1 | 93,317,331,071 | 13,39,20 | 247 | 150 | 1.54E-09 | 8.22E-08 | 0.998,0.991,0.99 | 0.978,0.964,0.97 | 0.025 | ENSG000001452 | [[[[48906658, 489 | 4 |
| ENSG000001590 | PAXBP1 | chr21 | 32743247 | 32743314 | 32741392 | 32741622 | 32744791 | 32744913 | 1,3,1 | 0,0,0 | 6,15,7 | 2,2,2 | 217 | 150 | 0.000631312 | 0.007843209 | 1.0,1.0,1.0 | 0.675,0.838,0.70 | 0.26 | ENSG000001590 | [[[[32771326, 327 | 2 |
| ENSG000001138 | TIMMDC1 | chr3 | 119503953 | 119504021 | 119498580 | 119498927 | 119517204 | 119517315 | 1,0,2 | 3,3,3 | 2,8,3 | 0,2,1 | 218 | 150 | 8.31E-06 | 0.000181642 | 0.187,0.0,0.314 | 1.0,0.733,0.674 | -0.635 | ENSG000001138 | [[[[119498547, 11 | 4 |
| ENSG000001737 | AGFG1 | chr2 | 227531090 | 227531210 | 227524761 | 227524915 | 227534844 | 227535025 | 25,22,35 | 0,0,0 | 23,35,17 | 4,3,5 | 270 | 150 | 4.35E-07 | 1.31E-05 | 1.0,1.0,1.0 | 0.762,0.866,0.65 | 0.239 | ENSG000001737 | [[[[227472156, 22 | 2 |
| ENSG000001737 | AGFG1 | chr2 | 227533548 | 227533758 | 227524761 | 227524915 | 227534844 | 227535025 | 35,29,55 | 0,0,0 | 30,57,31 | 4,3,5 | 300 | 150 | 1.95E-06 | 5.00E-05 | 1.0,1.0,1.0 | 0.789,0.905,0.75 | 0.183 | ENSG000001737 | [[[[227472156, 22 | 2 |
| ENSG000001234 | HJURP | chr2 | 233847396 | 233847461 | 233845727 | 233845820 | 233852564 | 233852620 | 37,33,47 | 0,0,0 | 31,52,34 | 7,8,1 | 215 | 150 | 3.87E-07 | 1.18E-05 | 1.0,1.0,1.0 | 0.755,0.819,0.96 | 0.155 | ENSG000001234 | [[[[233854384, 23 | 2 |
| ENSG000001234 | HJURP | chr2 | 233849762 | 233849859 | 233845727 | 233845820 | 233852564 | 233852620 | 29,45,68 | 0,0,0 | 65,170,92 | 7,8,1 | 247 | 150 | 7.69E-05 | 0.00126629 | 1.0,1.0,1.0 | 0.849,0.928,0.98 | 0.08 | ENSG000001234 | [[[[233854384, 23 | 2 |
| ENSG000001382 | DNAJC13 | chr3 | 132514570 | 132514670 | 132507236 | 132507353 | 132516421 | 132516496 | 21,20,16 | 0,0,0 | 24,31,11 | 2,6,0 | 250 | 150 | 0.000145878 | 0.002209867 | 1.0,1.0,1.0 | 0.878,0.756,1.0 | 0.122 | ENSG000001382 | [[[[132417502, 13 | 3 |
| ENSG000002669 | AARSD1 | chr17 | 42955841 | 42955972 | 42955089 | 42955367 | 42956403 | 42956525 | 1,1,3 | 0,0,0 | 29,77,24 | 2,6,2 | 281 | 150 | 0.00035634 | 0.004787096 | 1.0,1.0,1.0 | 0.886,0.873,0.86 | 0.125 | ENSG000002669 | [[[[42964402, 429 | 2 |
| ENSG000002669 | AARSD1 | chr17 | 42955841 | 42955972 | 42955157 | 42955224 | 42956403 | 42956525 | 37,52,59 | 0,0,0 | 70,153,74 | 5,9,12 | 281 | 150 | 9.66E-08 | 3.41E-06 | 1.0,1.0,1.0 | 0.882,0.901,0.76 | 0.15 | ENSG000002669 | [[[[42964402, 429 | 2 |
| ENSG000002669 | AARSD1 | chr17 | 42956203 | 42956320 | 42955157 | 42955224 | 42956403 | 42956504 | 70,53,65 | 0,0,0 | 76,137,67 | 5,9,12 | 267 | 150 | 1.77E-08 | 7.49E-07 | 1.0,1.0,1.0 | 0.895,0.895,0.75 | 0.151 | ENSG000002669 | [[[[42964402, 429 | 2 |
| ENSG000001978 | GPAA1 | chr8 | 144083713 | 144083861 | 144083388 | 144083500 | 144084285 | 144084330 | 23,31,89 | 0,0,0 | 21,36,20 | 4,3,4 | 298 | 150 | 2.49E-07 | 7.93E-06 | 1.0,1.0,1.0 | 0.725,0.858,0.71 | 0.234 | ENSG000001978 | [[[[144082634, 14 | 2 |
| ENSG000001635 | STT3B | chr3 | 31615104 | 31615203 | 31600359 | 31600459 | 31617939 | 31617988 | 59,30,89 | 0,0,0 | 19,66,35 | 1,8,3 | 249 | 150 | 2.42E-06 | 6.04E-05 | 1.0,1.0,1.0 | 0.92,0.832,0.875 | 0.124 | ENSG000001635 | [[[[31532925, 315 | 2 |
| ENSG000001635 | STT3B | chr3 | 31622096 | 31622308 | 31619675 | 31619830 | 31629297 | 31629411 | 97,82,121 | 0,0,0 | 27,79,55 | 5,17,8 | 300 | 150 | 1.33E-15 | 1.77E-13 | 1.0,1.0,1.0 | 0.73,0.699,0.775 | 0.265 | ENSG000001635 | [[[[31532925, 315 | 4 |
| ENSG000001635 | STT3B | chr3 | 31623673 | 31623861 | 31622096 | 31622308 | 31629297 | 31629411 | 112,93,183 | 0,0,0 | 44,93,66 | 3,10,9 | 300 | 150 | 1.63E-11 | 1.20E-09 | 1.0,1.0,1.0 | 0.88,0.823,0.786 | 0.17 | ENSG000001635 | [[[[31532925, 315 | 3 |
| ENSG000001635 | STT3B | chr3 | 31625953 | 31626127 | 31619675 | 31619830 | 31629297 | 31629411 | 155,96,152 | 0,0,0 | 55,72,40 | 5,17,8 | 300 | 150 | 6.66E-16 | 9.20E-14 | 1.0,1.0,1.0 | 0.846,0.679,0.71 | 0.254 | ENSG000001635 | [[[[31532925, 315 | 4 |
| ENSG000001635 | STT3B | chr3 | 31625953 | 31626127 | 31623673 | 31623861 | 31629297 | 31629411 | 157,98,152 | 0,0,0 | 55,72,39 | 6,14,11 | 300 | 150 | 1.11E-16 | 1.65E-14 | 1.0,1.0,1.0 | 0.821,0.72,0.639 | 0.273 | ENSG000001635 | [[[[31532925, 315 | 2 |
| ENSG000001434 | POGZ | chr1 | 151424954 | 151425061 | 151412295 | 151412396 | 151427822 | 151428014 | 18,20,45 | 0,0,0 | 28,59,51 | 2,4,7 | 257 | 150 | 8.20E-06 | 0.000179416 | 1.0,1.0,1.0 | 0.891,0.896,0.81 | 0.134 | ENSG000001434 | [[[[151459152, 15 | 3 |
| ENSG000001526 | SLC30A6 | chr2 | 32197706 | 32197826 | 32197317 | 32197392 | 32220212 | 32221524 | 13,10,14 | 0,0,0 | 7,18,9 | 1,4,3 | 270 | 150 | 2.83E-06 | 6.96E-05 | 1.0,1.0,1.0 | 0.795,0.714,0.62 | 0.289 | ENSG000001526 | [[[[32165864, 321 | 4 |
| ENSG000001526 | SLC30A6 | chr2 | 32204589 | 32204692 | 32197706 | 32197826 | 32220212 | 32221524 | 20,7,17 | 0,0,0 | 7,7,22 | 5,7,3 | 253 | 150 | 5.29E-09 | 2.50E-07 | 1.0,1.0,1.0 | 0.454,0.372,0.81 | 0.454 | ENSG000001526 | [[[[32165864, 321 | 3 |
| ENSG000001351 | DMTF1 | chr7 | 87188091 | 87188301 | 87185828 | 87185980 | 87193724 | 87194102 | 23,15,30 | 0,0,0 | 15,32,18 | 2,3,11 | 300 | 150 | 9.03E-08 | 3.21E-06 | 1.0,1.0,1.0 | 0.789,0.842,0.45 | 0.306 | ENSG000001351 | [[[[87152453, 871 | 3 |
| ENSG000001018 | CSTF2 | chrX | 100831514 | 100831656 | 100828039 | 100828102 | 100837338 | 100837449 | 21,29,43 | 0,0,0 | 25,64,22 | 8,8,1 | 292 | 150 | 1.90E-07 | 6.21E-06 | 1.0,1.0,1.0 | 0.616,0.804,0.91 | 0.22 | ENSG000001018 | [[[[100820391, 10 | 3 |
| ENSG000001342 | AP4B1 | chr1 | 113896257 | 113896465 | 113895756 | 113896038 | 113904604 | 113904793 | 14,14,28 | 0,0,0 | 16,39,13 | 0,3,5 | 300 | 150 | 0.00011224 | 0.001762666 | 1.0,1.0,1.0 | 1.0,0.867,0.565 | 0.189 | ENSG000001342 | [[[[113904605, 11 | 7 |
| ENSG000001342 | AP4B1 | chr1 | 113901235 | 113901383 | 113899903 | 113900400 | 113904604 | 113904793 | 26,15,21 | 0,3,0 | 11,16,11 | 11,14,7 | 298 | 150 | 3.13E-09 | 1.57E-07 | 1.0,0.716,1.0 | 0.335,0.365,0.44 | 0.525 | ENSG000001342 | [[[[113904605, 11 | 3 |
| ENSG000001342 | AP4B1 | chr1 | 113901754 | 113901885 | 113899941 | 113900400 | 113902637 | 113902734 | 15,19,35 | 2,0,4 | 12,7,5 | 6,23,17 | 281 | 150 | 4.45E-09 | 2.15E-07 | 0.8,1.0,0.824 | 0.516,0.14,0.136 | 0.611 | ENSG000001342 | [[[[113904605, 11 | 2 |
| ENSG000001342 | AP4B1 | chr1 | 113902637 | 113902862 | 113899903 | 113900400 | 113904604 | 113904793 | 20,13,33 | 0,3,0 | 14,67,42 | 11,14,7 | 300 | 150 | 0.000113152 | 0.001774431 | 1.0,0.684,1.0 | 0.389,0.705,0.75 | 0.28 | ENSG000001342 | [[[[113904605, 11 | 3 |
| ENSG000001217 | ZCCHC17 | chr1 | 31310043 | 31310164 | 31296981 | 31297075 | 31346639 | 31346740 | 22,23,27 | 0,0,0 | 16,51,42 | 2,3,5 | 271 | 150 | 4.75E-05 | 0.000834627 | 1.0,1.0,1.0 | 0.816,0.904,0.82 | 0.152 | ENSG000001217 | [[[[31297032, 312 | 4 |
| ENSG000000775 | CAPZB | chr1 | 19356634 | 19356751 | 19345186 | 19345252 | 19378539 | 19378653 | 421,367,532 | 0,0,0 | 484,881,529 | 7,30,5 | 267 | 150 | 1.59E-07 | 5.30E-06 | 1.0,1.0,1.0 | 0.975,0.943,0.98 | 0.033 | ENSG000000775 | [[[[19485436, 194 | 2 |
| ENSG000000775 | CAPZB | chr1 | 19357421 | 19357563 | 19345186 | 19345252 | 19378539 | 19378653 | 460,421,643 | 0,0,0 | 432,945,612 | 7,30,5 | 292 | 150 | 5.44E-08 | 2.04E-06 | 1.0,1.0,1.0 | 0.969,0.942,0.98 | 0.035 | ENSG000000775 | [[[[19485436, 194 | 2 |
| ENSG000001533 | RANBP2 | chr2 | 108777129 | 108777231 | 108772867 | 108773046 | 108781268 | 108781429 | 29,20,47 | 0,0,0 | 30,65,37 | 11,11,5 | 252 | 150 | 1.20E-09 | 6.54E-08 | 1.0,1.0,1.0 | 0.619,0.779,0.81 | 0.262 | ENSG000001533 | [[[[108719482, 10 | 2 |
| ENSG000001435 | ADAM15 | chr1 | 155060762 | 155060832 | 155060204 | 155060343 | 155061903 | 155061975 | 1,18,20 | 0,0,0 | 9,26,22 | 5,2,5 | 220 | 150 | 1.19E-06 | 3.20E-05 | 1.0,1.0,1.0 | 0.551,0.899,0.75 | 0.267 | ENSG000001435 | [[[[155051316, 15 | 2 |
| ENSG000001435 | ADAM15 | chr1 | 155061903 | 155061975 | 155060204 | 155060343 | 155062244 | 155062369 | 0,0,1 | 1,10,35 | 5,2,4 | 21,28,24 | 222 | 150 | 0.000704479 | 0.008613592 | 0.0,0.0,0.019 | 0.139,0.046,0.10 | -0.089 | ENSG000001435 | [[[[155051316, 15 | 3 |
| ENSG000001208 | ARL1 | chr12 | 101402864 | 101402946 | 101396142 | 101396577 | 101405843 | 101405981 | 47,46,72 | 0,0,0 | 36,93,47 | 4,6,0 | 232 | 150 | 0.000522518 | 0.006671391 | 1.0,1.0,1.0 | 0.853,0.909,1.0 | 0.079 | ENSG000001208 | [[[[101407642, 10 | 2 |
| ENSG000001625 | KIAA1522 | chr1 | 32769672 | 32769786 | 32768651 | 32768775 | 32772864 | 32774970 | 17,24,43 | 0,0,0 | 18,31,22 | 6,19,17 | 264 | 150 | 2.44E-15 | 3.14E-13 | 1.0,1.0,1.0 | 0.63,0.481,0.424 | 0.488 | ENSG000001625 | [[[[32741830, 327 | 2 |
| ENSG000001346 | PUM1 | chr1 | 31005852 | 31006031 | 30995053 | 30995220 | 31059298 | 31059577 | 37,32,74 | 0,0,0 | 29,125,90 | 1,6,6 | 300 | 150 | 5.35E-05 | 0.000926567 | 1.0,1.0,1.0 | 0.935,0.912,0.88 | 0.09 | ENSG000001346 | [[[[31065616, 310 | 3 |
| ENSG000001346 | PUM1 | chr1 | 31028795 | 31028864 | 30995053 | 30995220 | 31059298 | 31059577 | 4,3,1 | 0,0,0 | 2,13,8 | 1,6,6 | 219 | 150 | 2.42E-08 | 9.90E-07 | 1.0,1.0,1.0 | 0.578,0.597,0.47 | 0.449 | ENSG000001346 | [[[[31065616, 310 | 3 |
| ENSG000002137 | DDX47 | chr12 | 12823869 | 12824016 | 12823202 | 12823319 | 12827245 | 12827375 | 102,66,156 | 0,0,0 | 69,148,93 | 7,6,1 | 297 | 150 | 1.57E-05 | 0.000317296 | 1.0,1.0,1.0 | 0.833,0.926,0.97 | 0.087 | ENSG000002137 | [[[[12813346, 128 | 3 |
| ENSG000002137 | DDX47 | chr12 | 12824539 | 12824677 | 12823202 | 12823319 | 12825999 | 12826070 | 122,91,141 | 0,0,0 | 80,191,86 | 4,5,6 | 288 | 150 | 1.53E-05 | 0.0003095 | 1.0,1.0,1.0 | 0.912,0.952,0.88 | 0.085 | ENSG000002137 | [[[[12813346, 128 | 2 |
| ENSG000002137 | DDX47 | chr12 | 12825999 | 12826070 | 12823202 | 12823319 | 12827245 | 12827375 | 79,71,124 | 0,0,0 | 71,152,78 | 7,6,1 | 221 | 150 | 9.05E-05 | 0.001462608 | 1.0,1.0,1.0 | 0.873,0.945,0.98 | 0.067 | ENSG000002137 | [[[[12813346, 128 | 3 |
| ENSG000001661 | API5 | chr11 | 43320820 | 43320914 | 43318639 | 43318801 | 43326506 | 43326611 | 97,71,152 | 0,0,0 | 65,158,93 | 3,13,13 | 244 | 150 | 2.43E-10 | 1.51E-08 | 1.0,1.0,1.0 | 0.93,0.882,0.815 | 0.124 | ENSG000001661 | [[[[43311996, 433 | 4 |
| ENSG000001661 | API5 | chr11 | 43321410 | 43321476 | 43320820 | 43320914 | 43326506 | 43326611 | 70,33,90 | 0,0,0 | 41,143,72 | 3,11,1 | 216 | 150 | 2.59E-05 | 0.000491366 | 1.0,1.0,1.0 | 0.905,0.9,0.98 | 0.072 | ENSG000001661 | [[[[43311996, 433 | 3 |
| ENSG000001661 | API5 | chr11 | 43330507 | 43330564 | 43329964 | 43330058 | 43342427 | 43342664 | 48,69,91 | 0,0,0 | 37,80,42 | 8,14,12 | 207 | 150 | 2.80E-14 | 3.14E-12 | 1.0,1.0,1.0 | 0.77,0.805,0.717 | 0.236 | ENSG000001661 | [[[[43311996, 433 | 3 |
| ENSG000000711 | SNX13 | chr7 | 17801587 | 17801659 | 17799008 | 17799154 | 17814833 | 17814944 | 20,18,28 | 0,0,0 | 42,68,47 | 4,8,5 | 222 | 150 | 4.35E-06 | 0.000102176 | 1.0,1.0,1.0 | 0.876,0.852,0.86 | 0.136 | ENSG000000711 | [[[[17940284, 179 | 2 |
| ENSG000000711 | SNX13 | chr7 | 17875668 | 17875790 | 17873527 | 17873616 | 17890362 | 17890484 | 22,21,44 | 0,0,0 | 35,60,47 | 0,9,5 | 272 | 150 | 7.70E-06 | 0.000170066 | 1.0,1.0,1.0 | 1.0,0.786,0.838 | 0.125 | ENSG000000711 | [[[[17940284, 179 | 2 |
| ENSG000001518 | GUF1 | chr4 | 44683234 | 44683318 | 44680440 | 44680552 | 44685958 | 44686023 | 9,10,16 | 0,0,0 | 13,41,29 | 1,9,3 | 234 | 150 | 4.40E-06 | 0.000103214 | 1.0,1.0,1.0 | 0.893,0.745,0.86 | 0.167 | ENSG000001518 | [[[[44678420, 446 | 4 |
| ENSG000001518 | GUF1 | chr4 | 44689285 | 44689409 | 44686509 | 44686713 | 44694411 | 44694513 | 1,1,5 | 0,0,0 | 10,7,11 | 2,6,1 | 274 | 150 | 1.46E-06 | 3.85E-05 | 1.0,1.0,1.0 | 0.732,0.39,0.858 | 0.34 | ENSG000001518 | [[[[44678420, 446 | 5 |
| ENSG000001379 | KYAT3 | chr1 | 88961380 | 88961506 | 88949090 | 88949277 | 88962058 | 88962145 | 35,31,38 | 0,0,0 | 19,36,17 | 6,8,0 | 276 | 150 | 3.33E-07 | 1.03E-05 | 1.0,1.0,1.0 | 0.632,0.71,1.0 | 0.219 | ENSG000001379 | [[[[88992585, 889 | 4 |
| ENSG000001136 | TTC33 | chr5 | 40728344 | 40728476 | 40714474 | 40716498 | 40746797 | 40747019 | 10,8,24 | 0,0,2 | 14,45,14 | 6,9,5 | 282 | 150 | 1.63E-05 | 0.000328519 | 1.0,1.0,0.865 | 0.554,0.727,0.59 | 0.329 | ENSG000001136 | [[[[40755824, 407 | 2 |
| ENSG000001136 | TTC33 | chr5 | 40730261 | 40730343 | 40716144 | 40716498 | 40746797 | 40747019 | 5,4,8 | 0,0,2 | 13,16,10 | 6,9,5 | 232 | 150 | 0.000115616 | 0.001809717 | 1.0,1.0,0.721 | 0.583,0.535,0.56 | 0.346 | ENSG000001136 | [[[[40755824, 407 | 2 |
| ENSG000001620 | SLC30A7 | chr1 | 100921705 | 100921841 | 100918076 | 100918127 | 100974809 | 100974902 | 12,13,31 | 0,0,0 | 16,39,48 | 3,7,0 | 286 | 150 | 4.52E-05 | 0.000800077 | 1.0,1.0,1.0 | 0.737,0.745,1.0 | 0.173 | ENSG000001620 | [[[[100896090, 10 | 3 |
| ENSG000001620 | SLC30A7 | chr1 | 100965768 | 100965918 | 100918076 | 100918127 | 100974809 | 100974902 | 9,8,30 | 0,0,0 | 6,40,31 | 3,7,0 | 300 | 150 | 4.21E-05 | 0.00075327 | 1.0,1.0,1.0 | 0.5,0.741,1.0 | 0.253 | ENSG000001620 | [[[[100896090, 10 | 3 |
| ENSG000001620 | SLC30A7 | chr1 | 100965768 | 100965918 | 100921705 | 100921841 | 100974809 | 100974902 | 9,8,31 | 0,0,0 | 6,41,31 | 1,3,8 | 300 | 150 | 1.19E-06 | 3.21E-05 | 1.0,1.0,1.0 | 0.75,0.872,0.66 | 0.239 | ENSG000001620 | [[[[100896090, 10 | 2 |
| ENSG000001620 | SLC30A7 | chr1 | 100974809 | 100974902 | 100918076 | 100918127 | 100975218 | 100981753 | 5,7,15 | 0,0,0 | 8,22,15 | 1,4,2 | 243 | 150 | 0.000207754 | 0.003012447 | 1.0,1.0,1.0 | 0.832,0.772,0.82 | 0.191 | ENSG000001620 | [[[[100896090, 10 | 3 |
| ENSG000001346 | YARS | chr1 | 32810590 | 32810766 | 32806481 | 32806611 | 32817187 | 32817244 | 250,174,371 | 3,2,5 | 313,623,476 | 47,52,54 | 300 | 150 | 0 | 0 | 0.977,0.978,0.97 | 0.769,0.857,0.81 | 0.163 | ENSG000001346 | [[[[32817188, 328 | 2 |
| ENSG000001849 | WWTR1 | chr3 | 149526012 | 149526125 | 149517234 | 149520989 | 149542334 | 149542537 | 156,129,304 | 0,0,1 | 227,411,397 | 3,12,10 | 263 | 150 | 7.93E-05 | 0.001300392 | 1.0,1.0,0.994 | 0.977,0.951,0.95 | 0.036 | ENSG000001849 | [[[[149657765, 14 | 2 |
| ENSG000001169 | GNPAT | chr1 | 231266013 | 231266165 | 231265292 | 231265420 | 231266276 | 231266407 | 31,49,48 | 0,0,2 | 21,63,34 | 7,15,12 | 300 | 150 | 2.28E-11 | 1.64E-09 | 1.0,1.0,0.923 | 0.6,0.677,0.586 | 0.353 | ENSG000001169 | [[[[231241212, 23 | 2 |
| ENSG000000689 | ERLEC1 | chr2 | 53801712 | 53801842 | 53799046 | 53799081 | 53809213 | 53809273 | 20,16,10 | 4,0,0 | 20,32,28 | 24,22,18 | 280 | 150 | 5.32E-07 | 1.55E-05 | 0.728,1.0,1.0 | 0.309,0.438,0.45 | 0.509 | ENSG000000689 | [[[[53787044, 537 | 3 |
| ENSG000002440 | DDOST | chr1 | 20653626 | 20653774 | 20651766 | 20652528 | 20654222 | 20654371 | 438,678,997 | 0,0,0 | 6,411,372,774 | 11,14,15 | 298 | 150 | 6.61E-06 | 0.000148487 | 1.0,1.0,1.0 | 0.967,0.98,0.963 | 0.03 | ENSG000002440 | [[[[20661197, 206 | 3 |
| ENSG000001021 | MAGT1 | chrX | 77855500 | 77855590 | 77853900 | 77853964 | 77857356 | 77857497 | 78,56,129 | 0,0,0 | 107,160,147 | 3,7,4 | 240 | 150 | 0.000529548 | 0.00675435 | 1.0,1.0,1.0 | 0.957,0.935,0.95 | 0.05 | ENSG000001021 | [[[[77895309, 778 | 2 |
| ENSG000001021 | MAGT1 | chrX | 77856732 | 77856873 | 77855500 | 77855590 | 77870807 | 77870925 | 89,68,141 | 0,0,0 | 94,185,128 | 18,31,28 | 291 | 150 | 0 | 0 | 1.0,1.0,1.0 | 0.729,0.755,0.70 | 0.271 | ENSG000001021 | [[[[77895309, 778 | 2 |
| ENSG000001021 | MAGT1 | chrX | 77857356 | 77857497 | 77855500 | 77855590 | 77870807 | 77870925 | 122,117,205 | 0,0,0 | 179,332,210 | 18,31,28 | 291 | 150 | 0 | 0 | 1.0,1.0,1.0 | 0.837,0.847,0.79 | 0.174 | ENSG000001021 | [[[[77895309, 778 | 2 |
| ENSG000001443 | SCRN3 | chr2 | 174404102 | 174404315 | 174399921 | 174400103 | 174422884 | 174423047 | 22,34,30 | 0,0,0 | 13,22,13 | 2,2,2 | 300 | 150 | 0.000748338 | 0.009061153 | 1.0,1.0,1.0 | 0.765,0.846,0.76 | 0.208 | ENSG000001443 | [[[[174395730, 17 | 2 |
| ENSG000001137 | BNIP1 | chr5 | 173158743 | 173158845 | 173154321 | 173154413 | 173163724 | 173164340 | 15,19,33 | 0,0,1 | 39,78,38 | 2,10,3 | 252 | 150 | 6.67E-05 | 0.00112156 | 1.0,1.0,0.952 | 0.921,0.823,0.88 | 0.108 | ENSG000001137 | [[[[173144531, 17 | 2 |
| ENSG000001137 | BNIP1 | chr5 | 173159932 | 173160051 | 173154321 | 173154413 | 173163724 | 173164057 | 34,38,44 | 0,0,1 | 52,103,50 | 2,10,3 | 269 | 150 | 4.58E-05 | 0.000809934 | 1.0,1.0,0.961 | 0.935,0.852,0.90 | 0.09 | ENSG000001137 | [[[[173144531, 17 | 2 |
| ENSG000001219 | GPSM2 | chr1 | 108897958 | 108898101 | 108885274 | 108885578 | 108898641 | 108898724 | 41,40,68 | 0,0,0 | 53,69,42 | 0,10,6 | 293 | 150 | 1.00E-06 | 2.77E-05 | 1.0,1.0,1.0 | 1.0,0.779,0.782 | 0.146 | ENSG000001219 | [[[[108876985, 10 | 3 |

| | | | | | | | | | | | | | | | | | | | | | | |
|---|---|---|---|---|---|---|---|---|---|---|---|---|---|---|---|---|---|---|---|---|---|---|
| ENSG00000121 | GPSM2 | chr1 | 108923999 | 108924214 | 108918612 | 108918789 | 108929700 | 108929975 | 44,59,71 | 0,0,0 | 26,72,32 | 3,9,4 | 300 | 150 | 1.49E-08 | 6.39E-07 | 1.0,1.0,1.0 | 0.813,0.8,0.8 | 0.196 | ENSG00000121 | [[[[108876985, 1 | 2 |
| ENSG00000082 | BZW1 | chr2 | 200815666 | 200815761 | 200815340 | 200815517 | 200817973 | 200818083 | 225,212,359 | 0,0,0 | 162,400,278 | 5,7,6 | 245 | 150 | 0.000771255 | 0.009301691 | 1.0,1.0,1.0 | 0.952,0.972,0.96 | 0.037 | ENSG00000082 | [[[[200811910, 2 | 3 |
| ENSG00000134 | DPH6 | chr15 | 35538273 | 35538467 | 35454746 | 35454820 | 35546118 | 35546165 | 9,8,19 | 0,0,0 | 7,9,4 | 3,2,2 | 300 | 150 | 1.55E-06 | 4.08E-05 | 1.0,1.0,1.0 | 0.538,0.692,0.5 | 0.423 | ENSG00000134 | [[[[35546119, 35 | 2 |
| ENSG00000126 | KDM5C | chrX | 53210413 | 53210576 | 53201853 | 53201973 | 53211496 | 53211655 | 57,32,77 | 0,0,0 | 56,118,57 | 13,13,19 | 300 | 150 | 6.37E-13 | 5.94E-11 | 1.0,1.0,1.0 | 0.683,0.819,0.6 | 0.299 | ENSG00000126 | [[[[53224740, 53 | 2 |
| ENSG00000119 | FCF1 | chr14 | 74714871 | 74714943 | 74713484 | 74713552 | 74732730 | 74732812 | 20,4,33 | 0,0,0 | 13,40,19 | 3,9,8 | 222 | 150 | 7.64E-09 | 3.49E-07 | 1.0,1.0,1.0 | 0.745,0.75,0.616 | 0.296 | ENSG00000119 | [[[[74713144, 74 | 3 |
| ENSG00000135 | MTO1 | chr6 | 73482039 | 73482244 | 73480674 | 73480805 | 73492233 | 73492352 | 25,17,21 | 1,0,0 | 15,45,25 | 5,1,3 | 300 | 150 | 0.000165422 | 0.002467173 | 0.926,1.0,1.0 | 0.6,0.957,0.806 | 0.188 | ENSG00000135 | [[[[73461737, 73 | 2 |
| ENSG00000135 | MTO1 | chr6 | 73482448 | 73482620 | 73480674 | 73480805 | 73492233 | 73492325 | 17,13,15 | 1,0,0 | 7,33,14 | 5,1,3 | 300 | 150 | 7.83E-05 | 0.00128544 | 0.895,1.0,1.0 | 0.412,0.943,0.7 | 0.28 | ENSG00000135 | [[[[73461737, 73 | 2 |
| ENSG00000110 | MLEC | chr12 | 120694823 | 120695000 | 120693993 | 120694269 | 120696315 | 120696578 | 195,152,224 | 0,0,4 | 92,210,145 | 11,26,15 | 300 | 150 | 1.67E-15 | 2.19E-13 | 1.0,1.0,0.966 | 0.807,0.802,0.82 | 0.176 | ENSG00000110 | [[[[120687149, 1 | 2 |
| ENSG00000110 | MLEC | chr12 | 120695094 | 120695152 | 120693993 | 120694269 | 120696315 | 120696353 | 65,36,68 | 0,0,4 | 74,167,82 | 11,26,15 | 208 | 150 | 7.79E-08 | 2.81E-06 | 1.0,1.0,0.925 | 0.829,0.822,0.79 | 0.159 | ENSG00000110 | [[[[120687149, 1 | 2 |
| ENSG00000134 | BTF3L4 | chr1 | 52064824 | 52064938 | 52056271 | 52056379 | 52083339 | 52083541 | 158,155,170 | 0,0,0 | 187,292,179 | 5,11,7 | 264 | 150 | 3.15E-06 | 7.67E-05 | 1.0,1.0,1.0 | 0.955,0.938,0.93 | 0.057 | ENSG00000134 | [[[[52056295, 52 | 2 |
| ENSG00000067 | RRP15 | chr1 | 218305027 | 218305125 | 218302293 | 218302559 | 218330947 | 218337983 | 17,10,26 | 0,0,0 | 22,29,29 | 14,9,12 | 248 | 150 | 2.06E-11 | 1.49E-09 | 1.0,1.0,1.0 | 0.487,0.661,0.59 | 0.419 | ENSG00000067 | [[[[218285293, 2 | 2 |
| ENSG00000067 | RRP15 | chr1 | 218307430 | 218307632 | 218302293 | 218302559 | 218330947 | 218337983 | 44,37,43 | 0,0,0 | 46,69,56 | 14,9,12 | 300 | 150 | 1.44E-11 | 1.08E-09 | 1.0,1.0,1.0 | 0.622,0.793,0.7 | 0.295 | ENSG00000067 | [[[[218285293, 2 | 2 |
| ENSG00000077 | DYNC1I2 | chr2 | 171727820 | 171727967 | 171726557 | 171726916 | 171728716 | 171728850 | 109,119,130 | 0,0,0 | 87,159,127 | 4,6,2 | 297 | 150 | 0.000398324 | 0.005276684 | 1.0,1.0,1.0 | 0.917,0.93,0.97 | 0.061 | ENSG00000077 | [[[[171687469, 1 | 2 |
| ENSG00000196 | ARHGEF12 | chr11 | 120429711 | 120429831 | 120424357 | 120424415 | 120431770 | 120431911 | 13,6,19 | 0,0,0 | 3,20,21 | 0,2,5 | 270 | 150 | 0.00022826 | 0.003260873 | 1.0,1.0,1.0 | 1.0,0.847,0.7 | 0.151 | ENSG00000196 | [[[[120336413, 1 | 3 |
| ENSG00000054 | SDCCAG8 | chr1 | 243293090 | 243293219 | 243256055 | 243256240 | 243304712 | 243304777 | 7,2,10 | 0,0,0 | 3,7,5 | 3,13,4 | 279 | 150 | 6.39E-12 | 5.12E-10 | 1.0,1.0,1.0 | 0.35,0.225,0.402 | 0.674 | ENSG00000054 | [[[[243256041, 2 | 5 |
| ENSG00000174 | IL20RB | chr3 | 136995413 | 136995556 | 136989440 | 136989565 | 137010112 | 137011079 | 3,2,1 | 0,0,0 | 7,6,7 | 0,4,5 | 293 | 150 | 2.55E-06 | 6.34E-05 | 1.0,1.0,1.0 | 1.0,0.434,0.417 | 0.383 | ENSG00000174 | [[[[136957983, 1 | 2 |
| ENSG00000036 | AC118549.1 | chr1 | 77631849 | 77633405 | 77584516 | 77584655 | 77641521 | 77641655 | 31,41,56 | 0,4,0 | 56,105,81 | 13,25,5 | 300 | 150 | 2.62E-05 | 0.000497085 | 1.0,0.837,1.0 | 0.683,0.677,0.89 | 0.196 | ENSG00000036 | [[[[77682585, 77 | 3 |
| ENSG00000036 | AC118549.1 | chr1 | 77639448 | 77639602 | 77584516 | 77584655 | 77641383 | 77641446 | 26,8,26 | 0,0,0 | 30,91,87 | 4,12,15 | 300 | 150 | 7.16E-08 | 2.61E-06 | 1.0,1.0,1.0 | 0.789,0.791,0.74 | 0.225 | ENSG00000036 | [[[[77682585, 77 | 2 |
| ENSG00000036 | AC118549.1 | chr1 | 77639448 | 77639602 | 77631849 | 77633405 | 77641521 | 77641655 | 31,20,53 | 0,0,0 | 29,114,52 | 0,8,2 | 300 | 150 | 0.000335373 | 0.004542763 | 1.0,1.0,1.0 | 1.0,0.877,0.929 | 0.065 | ENSG00000036 | [[[[77682585, 77 | 2 |
| ENSG00000160 | UBQLN4 | chr1 | 156041871 | 156041987 | 156041484 | 156041671 | 156042773 | 156042913 | 5,8,47 | 0,0,0 | 22,68,62 | 0,1,8 | 266 | 150 | 0.000797994 | 0.009580207 | 1.0,1.0,1.0 | 1.0,0.975,0.814 | 0.07 | ENSG00000160 | [[[[156053594, 1 | 2 |
| ENSG00000160 | UBQLN4 | chr1 | 156043997 | 156044223 | 156042773 | 156042913 | 156053593 | 156053794 | 16,37,59 | 0,0,0 | 64,119,79 | 4,15,5 | 300 | 150 | 3.02E-07 | 9.45E-06 | 1.0,1.0,1.0 | 0.889,0.799,0.88 | 0.141 | ENSG00000160 | [[[[156053594, 1 | 5 |
| ENSG00000160 | UBQLN4 | chr1 | 156051109 | 156051327 | 156050290 | 156050553 | 156053593 | 156053650 | 1,13,41 | 0,0,0 | 10,33,20 | 0,5,2 | 300 | 150 | 0.000474128 | 0.006149386 | 1.0,1.0,1.0 | 1.0,0.767,0.833 | 0.133 | ENSG00000160 | [[[[156053594, 1 | 2 |
| ENSG00000162 | MYSM1 | chr1 | 58667037 | 58667226 | 58661405 | 58661511 | 58667846 | 58667921 | 12,8,14 | 1,0,0 | 21,36,27 | 4,3,10 | 300 | 150 | 1.77E-05 | 0.000352567 | 0.857,1.0,1.0 | 0.724,0.857,0.57 | 0.234 | ENSG00000162 | [[[[58699985, 58 | 2 |
| ENSG00000172 | FGGY | chr1 | 59339957 | 59340069 | 59321535 | 59321750 | 59378748 | 59378837 | 2,6,9 | 0,1,0 | 4,2,7 | 3,2,3 | 262 | 150 | 6.78E-06 | 0.000151477 | 1.0,0.775,1.0 | 0.433,0.364,0.57 | 0.469 | ENSG00000172 | [[[[59297094, 59 | 2 |
| ENSG00000172 | FGGY | chr1 | 59346246 | 59346398 | 59321535 | 59321750 | 59378748 | 59378837 | 8,5,11 | 0,1,0 | 7,12,11 | 3,2,3 | 300 | 150 | 8.03E-05 | 0.001313728 | 1.0,0.714,1.0 | 0.538,0.75,0.647 | 0.26 | ENSG00000172 | [[[[59297094, 59 | 2 |
| ENSG00000091 | LAMB1 | chr7 | 107935414 | 107935656 | 107932173 | 107932493 | 107951225 | 107951322 | 3,5,8 | 0,0,0 | 7,11,3 | 1,7,1 | 300 | 150 | 1.06E-07 | 3.68E-06 | 1.0,1.0,1.0 | 0.778,0.44,0.6 | 0.394 | ENSG00000091 | [[[[108003111, 1 | 3 |
| ENSG00000119 | ITGB1BP1 | chr2 | 9414177 | 9414256 | 9412067 | 9412405 | 9423372 | 9423522 | 53,45,61 | 7,2,6 | 73,135,83 | 2,3,1 | 229 | 150 | 0.000704015 | 0.008609298 | 0.832,0.936,0.86 | 0.96,0.967,0.982 | -0.091 | ENSG00000119 | [[[[9423373, 942 | 2 |
| ENSG00000144 | ZC3H8 | chr2 | 112234119 | 112234236 | 112233257 | 112233371 | 112250190 | 112250272 | 11,1,11 | 0,0,0 | 5,10,19 | 7,6,9 | 267 | 150 | 8.70E-10 | 4.89E-08 | 1.0,1.0,1.0 | 0.286,0.484,0.54 | 0.562 | ENSG00000144 | [[[[112254908, 1 | 3 |
| ENSG00000077 | DNAJC10 | chr2 | 182736248 | 182736386 | 182731029 | 182731107 | 182740298 | 182740388 | 58,42,66 | 1,2,0 | 32,81,50 | 3,2,8 | 288 | 150 | 0.000710529 | 0.008669472 | 0.968,0.916,1.0 | 0.847,0.955,0.76 | 0.106 | ENSG00000077 | [[[[182716257, 1 | 2 |
| ENSG00000069 | GNB5 | chr15 | 52141139 | 52141272 | 52135612 | 52135756 | 52149883 | 52149925 | 6,15,44 | 0,0,0 | 18,47,27 | 0,8,0 | 283 | 150 | 0.0008128 | 0.009741937 | 1.0,1.0,1.0 | 1.0,0.757,1.0 | 0.081 | ENSG00000069 | [[[[52191322, 52 | 2 |
| ENSG00000114 | EIF4G1 | chr3 | 184315488 | 184315545 | 184314494 | 184314674 | 184317716 | 184317816 | 2,2,29 | 0,1,0 | 4,15,19 | 1,7,10 | 207 | 150 | 4.44E-06 | 0.000104013 | 1.0,0.592,1.0 | 0.743,0.608,0.57 | 0.221 | ENSG00000114 | [[[[184314606, 1 | 4 |
| ENSG00000114 | EIF4G1 | chr3 | 184315762 | 184315856 | 184314494 | 184314674 | 184316131 | 184316218 | 11,14,55 | 0,0,0 | 5,46,34 | 3,8,7 | 244 | 150 | 3.17E-08 | 1.26E-06 | 1.0,1.0,1.0 | 0.506,0.779,0.74 | 0.322 | ENSG00000114 | [[[[184314606, 1 | 2 |
| ENSG00000124 | DEK | chr6 | 18255730 | 18255851 | 18249650 | 18249839 | 18257952 | 18258031 | 235,136,331 | 0,0,0 | 127,397,293 | 7,11,16 | 271 | 150 | 1.85E-09 | 9.73E-08 | 1.0,1.0,1.0 | 0.909,0.952,0.91 | 0.076 | ENSG00000124 | [[[[18264385, 18 | 2 |
| ENSG00000124 | DEK | chr6 | 18256360 | 18256455 | 18249650 | 18249839 | 18257952 | 18258031 | 82,64,103 | 0,0,0 | 33,166,95 | 7,11,16 | 245 | 150 | 3.97E-11 | 2.77E-09 | 1.0,1.0,1.0 | 0.743,0.902,0.78 | 0.19 | ENSG00000124 | [[[[18264385, 18 | 2 |
| ENSG00000124 | DEK | chr6 | 18257952 | 18258062 | 18256360 | 18256455 | 18263842 | 18263996 | 128,84,165 | 0,0,0 | 57,224,137 | 0,9,10 | 260 | 150 | 2.90E-06 | 7.14E-05 | 1.0,1.0,1.0 | 1.0,0.935,0.888 | 0.059 | ENSG00000124 | [[[[18264385, 18 | 2 |
| ENSG00000124 | DEK | chr6 | 18258303 | 18258405 | 18249650 | 18249839 | 18263842 | 18263996 | 36,31,97 | 0,0,0 | 35,109,63 | 2,5,5 | 252 | 150 | 8.63E-05 | 0.001400449 | 1.0,1.0,1.0 | 0.912,0.928,0.88 | 0.093 | ENSG00000124 | [[[[18264385, 18 | 4 |
| ENSG00000124 | DEK | chr6 | 18258303 | 18258405 | 18256360 | 18256455 | 18263842 | 18263996 | 36,31,97 | 0,0,0 | 35,111,63 | 0,9,10 | 252 | 150 | 6.99E-07 | 2.00E-05 | 1.0,1.0,1.0 | 1.0,0.88,0.789 | 0.11 | ENSG00000124 | [[[[18264385, 18 | 2 |
| ENSG00000123 | ACSL3 | chr2 | 222927016 | 222927189 | 222924455 | 222924595 | 222941496 | 222942222 | 216,147,253 | 0,0,0 | 209,399,281 | 5,10,3 | 300 | 150 | 0.00040255 | 0.00532619 | 1.0,1.0,1.0 | 0.954,0.952,0.97 | 0.038 | ENSG00000123 | [[[[222861036, 2 | 5 |
| ENSG00000123 | ACSL3 | chr2 | 222928861 | 222928936 | 222927016 | 222927189 | 222941496 | 222942222 | 118,112,192 | 0,0,0 | 117,258,124 | 5,22,9 | 225 | 150 | 1.22E-10 | 7.92E-09 | 1.0,1.0,1.0 | 0.94,0.887,0.902 | 0.09 | ENSG00000123 | [[[[222861036, 2 | 4 |
| ENSG00000123 | ACSL3 | chr2 | 222934529 | 222934687 | 222927016 | 222927189 | 222941496 | 222941527 | 234,185,361 | 0,0,0 | 210,449,220 | 5,22,9 | 300 | 150 | 3.78E-10 | 2.26E-08 | 1.0,1.0,1.0 | 0.955,0.911,0.92 | 0.07 | ENSG00000123 | [[[[222861036, 2 | 4 |
| ENSG00000116 | CAPZA1 | chr1 | 112647209 | 112647273 | 112619172 | 112619883 | 112653597 | 112653661 | 2,3,11 | 0,0,0 | 2,21,17 | 7,20,41 | 214 | 150 | 1.96E-11 | 1.42E-09 | 1.0,1.0,1.0 | 0.167,0.424,0.22 | 0.728 | ENSG00000116 | [[[[112619832, 1 | 2 |
| ENSG00000116 | CAPZA1 | chr1 | 112649417 | 112649469 | 112619172 | 112619883 | 112653597 | 112653661 | 101,98,191 | 0,0,0 | 99,230,204 | 7,20,41 | 202 | 150 | 1.22E-15 | 1.64E-13 | 1.0,1.0,1.0 | 0.913,0.895,0.78 | 0.135 | ENSG00000116 | [[[[112619832, 1 | 2 |
| ENSG00000112 | STK38 | chr6 | 36517716 | 36517840 | 36515337 | 36515492 | 36524340 | 36524463 | 23,16,48 | 0,0,0 | 21,58,34 | 5,9,3 | 274 | 150 | 1.17E-07 | 4.02E-06 | 1.0,1.0,1.0 | 0.697,0.779,0.86 | 0.221 | ENSG00000112 | [[[[36547190, 36 | 2 |
| ENSG00000088 | ALG6 | chr1 | 63428732 | 63428801 | 63419369 | 63419440 | 63436822 | 63437126 | 20,14,4 | 0,0,0 | 13,8,7 | 1,2,4 | 219 | 150 | 5.12E-05 | 0.000891159 | 1.0,1.0,1.0 | 0.899,0.733,0.54 | 0.274 | ENSG00000088 | [[[[63367627, 63 | 2 |
| ENSG00000144 | RABL2A | chr2 | 113635050 | 113635130 | 113634152 | 113634232 | 113641352 | 113641450 | 5,9,9 | 0,1,0 | 13,20,21 | 1,6,5 | 230 | 150 | 8.25E-05 | 0.001344735 | 1.0,0.854,1.0 | 0.894,0.685,0.73 | 0.181 | ENSG00000144 | [[[[113627266, 1 | 2 |
| ENSG00000144 | RABL2A | chr2 | 113640893 | 113641005 | 113634152 | 113634232 | 113641352 | 113641450 | 4,2,2 | 0,1,0 | 2,6,4 | 1,6,5 | 262 | 150 | 1.28E-05 | 0.000264734 | 1.0,0.534,1.0 | 0.534,0.364,0.31 | 0.441 | ENSG00000144 | [[[[113627266, 1 | 2 |
| ENSG00000168 | CEP120 | chr5 | 123391109 | 123391337 | 123389923 | 123390140 | 123399135 | 123399284 | 19,13,26 | 0,0,0 | 8,26,18 | 0,10,2 | 300 | 150 | 3.58E-06 | 8.59E-05 | 1.0,1.0,1.0 | 1.0,0.565,0.818 | 0.206 | ENSG00000168 | [[[[123422950, 1 | 2 |
| ENSG00000112 | MRPL2 | chr6 | 43055544 | 43055618 | 43054028 | 43054486 | 43058064 | 43058233 | 19,37,22 | 7,5,14 | 45,102,42 | 4,5,4 | 224 | 150 | 0.000156225 | 0.002345153 | 0.645,0.832,0.51 | 0.883,0.932,0.87 | -0.233 | ENSG00000112 | [[[[43059286, 43 | 4 |
| ENSG00000124 | ABCC10 | chr6 | 43445598 | 43445942 | 43445124 | 43445314 | 43447247 | 43447408 | 6,12,14 | 0,0,0 | 7,21,14 | 1,3,2 | 300 | 150 | 0.000293987 | 0.004059666 | 1.0,1.0,1.0 | 0.778,0.778,0.77 | 0.222 | ENSG00000124 | [[[[43427541, 43 | 2 |
| ENSG00000066 | LRRC40 | chr1 | 70155688 | 70155796 | 70148486 | 70148672 | 70159329 | 70159438 | 19,27,26 | 0,0,0 | 13,29,20 | 6,20,16 | 258 | 150 | 4.44E-16 | 6.24E-14 | 1.0,1.0,1.0 | 0.557,0.457,0.42 | 0.522 | ENSG00000066 | [[[[70205390, 70 | 3 |
| ENSG00000116 | CTH | chr1 | 70415955 | 70416037 | 70411217 | 70411583 | 70421565 | 70421675 | 5,5,21 | 0,0,0 | 20,48,45 | 5,6,0 | 232 | 150 | 0.000100274 | 0.001601821 | 1.0,1.0,1.0 | 0.721,0.838,1.0 | 0.147 | ENSG00000116 | [[[[70411268, 70 | 2 |
| ENSG00000116 | CTH | chr1 | 70417936 | 70418032 | 70411271 | 70411583 | 70421565 | 70421675 | 3,4,8 | 0,0,0 | 31,75,40 | 5,6,0 | 246 | 150 | 0.000326615 | 0.004447305 | 1.0,1.0,1.0 | 0.791,0.884,1.0 | 0.108 | ENSG00000116 | [[[[70411268, 70 | 2 |
| ENSG00000116 | CTH | chr1 | 70421565 | 70421675 | 70411217 | 70411583 | 70424284 | 70424416 | 6,4,13 | 0,0,0 | 49,96,82 | 1,6,5 | 260 | 150 | 0.000540309 | 0.006876639 | 1.0,1.0,1.0 | 0.966,0.902,0.90 | 0.076 | ENSG00000116 | [[[[70411268, 70 | 3 |
| ENSG00000139 | TMEM19 | chr12 | 71697279 | 71697534 | 71696435 | 71696573 | 71700831 | 71705046 | 18,27,52 | 0,0,0 | 9,32,22 | 0,10,6 | 300 | 150 | 8.47E-08 | 3.03E-06 | 1.0,1.0,1.0 | 1.0,0.615,0.647 | 0.246 | ENSG00000139 | [[[[71686082, 71 | 2 |
| ENSG00000143 | LBR | chr1 | 225410290 | 225410416 | 225406561 | 225406832 | 225412453 | 225412645 | 90,67,113 | 0,0,0 | 67,136,93 | 1,11,5 | 276 | 150 | 1.84E-06 | 4.76E-05 | 1.0,1.0,1.0 | 0.973,0.87,0.91 | 0.082 | ENSG00000143 | [[[[225427954, 2 | 2 |
| ENSG00000143 | LBR | chr1 | 225411336 | 225411440 | 225406663 | 225406832 | 225412453 | 225412645 | 88,67,102 | 0,0,0 | 45,64,70 | 1,11,5 | 254 | 150 | 9.26E-08 | 3.28E-06 | 1.0,1.0,1.0 | 0.964,0.775,0.89 | 0.123 | ENSG00000143 | [[[[225427954, 2 | 2 |
| ENSG00000137 | RABGGTB | chr1 | 75789158 | 75789356 | 75786700 | 75787604 | 75790151 | 75790274 | 426,295,478 | 0,0,0 | 97,018,351,152 | 12,29,14 | 300 | 150 | 2.63E-05 | 0.000497756 | 1.0,1.0,1.0 | 0.976,0.969,0.97 | 0.026 | ENSG00000137 | [[[[75786256, 75 | 2 |
| ENSG00000137 | RABGGTB | chr1 | 75789158 | 75789356 | 75786700 | 75787604 | 75792180 | 75792286 | 426,295,478 | 0,0,0 | 97,418,631,168 | 19,21,13 | 300 | 150 | 4.84E-05 | 0.000847862 | 1.0,1.0,1.0 | 0.962,0.978,0.97 | 0.027 | ENSG00000137 | [[[[75786256, 75 | 4 |
| ENSG00000137 | RABGGTB | chr1 | 75789951 | 75790057 | 75789158 | 75789356 | 75792180 | 75792286 | 126,93,155 | 0,0,0 | 289,560,317 | 4,30,17 | 256 | 150 | 4.93E-08 | 1.86E-06 | 1.0,1.0,1.0 | 0.977,0.916,0.91 | 0.064 | ENSG00000137 | [[[[75786256, 75 | 3 |
| ENSG00000137 | RABGGTB | chr1 | 75789951 | 75790274 | 75787496 | 75787604 | 75791284 | 75791337 | 4,2,11 | 0,1,2 | 6,7,5 | 12,28,9 | 300 | 150 | 1.29E-05 | 0.000266805 | 1.0,0.5,0.733 | 0.2,0.111,0.217 | 0.568 | ENSG00000137 | [[[[75786256, 75 | 2 |
| ENSG00000137 | RABGGTB | chr1 | 75789951 | 75790274 | 75787496 | 75787604 | 75792180 | 75792254 | 4,2,11 | 0,0,0 | 7,11,6 | 19,21,13 | 300 | 150 | 5.12E-13 | 4.85E-11 | 1.0,1.0,1.0 | 0.156,0.208,0.18 | 0.816 | ENSG00000137 | [[[[75786256, 75 | 4 |
| ENSG00000137 | RABGGTB | chr1 | 75790151 | 75790274 | 75789951 | 75790057 | 75792180 | 75792306 | 31,26,18 | 0,0,0 | 85,160,69 | 12,18,11 | 273 | 150 | 4.41E-08 | 1.69E-06 | 1.0,1.0,1.0 | 0.796,0.83,0.775 | 0.2 | ENSG00000137 | [[[[75786256, 75 | 2 |
| ENSG00000137 | RABGGTB | chr1 | 75790151 | 75790274 | 75789951 | 75790057 | 75794083 | 75794233 | 31,26,18 | 0,0,0 | 83,157,67 | 5,5,3 | 273 | 150 | 0.000635779 | 0.007892649 | 1.0,1.0,1.0 | 0.901,0.945,0.92 | 0.076 | ENSG00000137 | [[[[75786256, 75 | 3 |
| ENSG00000137 | RABGGTB | chr1 | 75791460 | 75791571 | 75786700 | 75787604 | 75792180 | 75792250 | 141,148,155 | 0,0,0 | 245,468,235 | 19,21,13 | 261 | 150 | 1.34E-10 | 8.66E-09 | 1.0,1.0,1.0 | 0.881,0.928,0.91 | 0.093 | ENSG00000137 | [[[[75786256, 75 | 4 |
| ENSG00000137 | RABGGTB | chr1 | 75791460 | 75791571 | 75789158 | 75789356 | 75792180 | 75792250 | 141,148,155 | 0,0,0 | 245,466,233 | 4,30,17 | 261 | 150 | 5.27E-10 | 3.08E-08 | 1.0,1.0,1.0 | 0.972,0.899,0.88 | 0.081 | ENSG00000137 | [[[[75786256, 75 | 3 |
| ENSG00000137 | RABGGTB | chr1 | 75791460 | 75791571 | 75789951 | 75790057 | 75792180 | 75792250 | 141,148,155 | 0,0,0 | 244,470,230 | 12,18,11 | 261 | 150 | 2.02E-08 | 8.44E-07 | 1.0,1.0,1.0 | 0.921,0.938,0.92 | 0.073 | ENSG00000137 | [[[[75786256, 75 | 2 |
| ENSG00000173 | NABP1 | chr2 | 191684229 | 191684296 | 191678989 | 191679128 | 191685598 | 191685894 | 11,8,9 | 0,0,0 | 9,17,13 | 7,7,2 | 217 | 150 | 3.66E-08 | 1.43E-06 | 1.0,1.0,1.0 | 0.471,0.627,0.81 | 0.361 | ENSG00000173 | [[[[191678136, 1 | 3 |
| ENSG00000104 | DECR1 | chr8 | 90017123 | 90017326 | 90001404 | 90001561 | 90020908 | 90020912 | 65,26,61 | 0,0,0 | 27,52,41 | 1,16,5 | 300 | 150 | 7.98E-09 | 3.63E-07 | 1.0,1.0,1.0 | 0.931,0.619,0.80 | 0.215 | ENSG00000104 | [[[[90001477, 90 | 3 |
| ENSG00000104 | DECR1 | chr8 | 90017123 | 90017326 | 90001404 | 90001561 | 90042727 | 90042800 | 65,26,61 | 0,2,0 | 27,47,40 | 7,22,14 | 300 | 150 | 7.06E-14 | 7.46E-12 | 1.0,0.867,1.0 | 0.659,0.516,0.58 | 0.368 | ENSG00000104 | [[[[90001477, 90 | 5 |
| ENSG00000104 | DECR1 | chr8 | 90020908 | 90021056 | 90001404 | 90001561 | 90036840 | 90036879 | 77,68,102 | 0,1,0 | 61,117,70 | 3,9,5 | 298 | 150 | 1.74E-06 | 4.52E-05 | 1.0,0.972,1.0 | 0.911,0.867,0.87 | 0.106 | ENSG00000104 | [[[[90001477, 90 | 4 |
| ENSG00000104 | DECR1 | chr8 | 90036840 | 90036940 | 90001404 | 90001561 | 90042727 | 90042748 | 81,67,81 | 0,2,0 | 76,117,89 | 7,22,14 | 250 | 150 | 3.08E-11 | 2.18E-09 | 1.0,0.953,1.0 | 0.867,0.761,0.79 | 0.178 | ENSG00000104 | [[[[90001477, 90 | 5 |
| ENSG00000170 | USP47 | chr11 | 11950363 | 11950482 | 11947944 | 11948120 | 11952740 | 11952777 | 25,37,38 | 0,0,0 | 41,72,43 | 3,8,0 | 269 | 150 | 7.16E-05 | 0.001191779 | 1.0,1.0,1.0 | 0.884,0.834,1.0 | 0.094 | ENSG00000170 | [[[[11841972, 11 | 3 |
| ENSG00000170 | USP47 | chr11 | 11954896 | 11954944 | 11950872 | 11952871 | 11956000 | 11956421 | 10,4,19 | 0,0,0 | 22,25,17 | 10,9,3 | 198 | 150 | 3.91E-08 | 1.52E-06 | 1.0,1.0,1.0 | 0.625,0.678,0.81 | 0.295 | ENSG00000170 | [[[[11841972, 11 | 2 |
| ENSG00000170 | USP47 | chr11 | 11955033 | 11955164 | 11950872 | 11952871 | 11956000 | 11956421 | 41,39,33 | 0,0,0 | 54,64,43 | 10,9,3 | 281 | 150 | 2.00E-08 | 8.35E-07 | 1.0,1.0,1.0 | 0.742,0.791,0.88 | 0.194 | ENSG00000170 | [[[[11841972, 11 | 2 |
| ENSG00000077 | USP33 | chr1 | 77715741 | 77715868 | 77713199 | 77713281 | 77717866 | 77718047 | 58,48,69 | 0,3,8 | 17,53,40 | 8,15,7 | 277 | 150 | 0.000101483 | 0.001616382 | 1.0,0.897,0.824 | 0.535,0.657,0.75 | 0.258 | ENSG00000077 | [[[[77759643, 77 | 2 |
| ENSG00000077 | USP33 | chr1 | 77715741 | 77715868 | 77714613 | 77714783 | 77722023 | 77722196 | 49,36,59 | 0,0,0 | 18,51,33 | 2,10,2 | 277 | 150 | 1.18E-07 | 4.06E-06 | 1.0,1.0,1.0 | 0.83,0.734,0.899 | 0.179 | ENSG00000077 | [[[[77759643, 77 | 5 |
| ENSG00000116 | PHTF1 | chr1 | 113705889 | 113706162 | 113704665 | 113704797 | 113710253 | 113710475 | 30,31,37 | 0,0,0 | 16,36,26 | 10,17,15 | 300 | 150 | 0 | 0 | 1.0,1.0,1.0 | 0.444,0.514,0.46 | 0.526 | ENSG00000116 | [[[[113759023, 1 | 2 |
| ENSG00000115 | RAB3GAP1 | chr2 | 135115215 | 135115381 | 135058010 | 135058086 | 135120818 | 135120918 | 57,53,86 | 1,0,0 | 44,99,67 | 3,8,3 | 300 | 150 | 1.80E-05 | 0.000357103 | 0.966,1.0,1.0 | 0.88,0.861,0.918 | 0.102 | ENSG00000115 | [[[[135052292, 1 | 4 |
| ENSG00000204 | MBD5 | chr2 | 148468340 | 148470461 | 148458202 | 148458871 | 148483109 | 148483436 | 6,1,6 | 0,0,0 | 9,23,8 | 0,6,3 | 300 | 150 | 1.02E-05 | 0.000217802 | 1.0,1.0,1.0 | 1.0,0.657,0.571 | 0.257 | ENSG00000204 | [[[[148021091, 1 | 3 |
| ENSG00000150 | PDCD4 | chr10 | 110885252 | 110885366 | 110883002 | 110883097 | 110890555 | 110890670 | 9,11,15 | 0,0,0 | 24,32,37 | 6,0,3 | 264 | 150 | 0.000100044 | 0.001599166 | 1.0,1.0,1.0 | 0.694,1.0,0.875 | 0.144 | ENSG00000150 | [[[[110871928, 1 | 3 |

| | | | | | | | | | | | | | | | | | | | | | | |
|---|---|---|---|---|---|---|---|---|---|---|---|---|---|---|---|---|---|---|---|---|---|---|
| ENSG000001505 | PDCD4 | chr10 | 110885252 | 110885366 | 110883002 | 110883097 | 110894090 | 110894198 | 9,11,15 | 0,0,0 | 24,38,38 | 5,6,10 | 264 | 150 | 2.21E-07 | 7.10E-06 | 1.0,1.0,1.0 | 0.732,0.783,0.68 | 0.267 | ENSG000001505 | [[[[110871928, 11 | 4 |
| ENSG000001505 | PDCD4 | chr10 | 110890555 | 110890670 | 110883002 | 110883097 | 110894090 | 110894179 | 19,12,15 | 0,0,0 | 31,55,35 | 5,6,10 | 265 | 150 | 3.27E-07 | 1.02E-05 | 1.0,1.0,1.0 | 0.778,0.838,0.66 | 0.24 | ENSG000001505 | [[[[110871928, 11 | 4 |
| ENSG000001505 | PDCD4 | chr10 | 110890555 | 110890670 | 110884770 | 110885366 | 110894090 | 110894179 | 19,12,15 | 0,0,0 | 26,57,32 | 1,7,1 | 265 | 150 | 0.000220806 | 0.003175118 | 1.0,1.0,1.0 | 0.936,0.822,0.94 | 0.098 | ENSG000001505 | [[[[110871928, 11 | 3 |
| ENSG000001133 | FAM172A | chr5 | 93958776 | 93958968 | 93881310 | 93881688 | 94050730 | 94050831 | 12,6,12 | 0,0,0 | 4,17,6 | 3,2,3 | 300 | 150 | 2.34E-07 | 7.48E-06 | 1.0,1.0,1.0 | 0.4,0.81,0.5 | 0.43 | ENSG000001133 | [[[[94111511, 941 | 2 |
| ENSG000001133 | FAM172A | chr5 | 93964457 | 93964523 | 93881469 | 93881688 | 94050730 | 94050831 | 5,2,7 | 0,0,0 | 1,2,1 | 3,2,3 | 216 | 150 | 5.30E-11 | 3.62E-09 | 1.0,1.0,1.0 | 0.188,0.41,0.188 | 0.738 | ENSG000001133 | [[[[94111511, 941 | 2 |
| ENSG000000891 | TASP1 | chr20 | 13580896 | 13580981 | 13534021 | 13534141 | 13587249 | 13587370 | 10,7,17 | 0,0,0 | 5,11,10 | 3,2,1 | 235 | 150 | 0.000453414 | 0.00591394 | 1.0,1.0,1.0 | 0.515,0.778,0.86 | 0.281 | ENSG000000891 | [[[[13638894, 136 | 3 |
| ENSG000001858 | SLC52A2 | chr8 | 144359183 | 144359423 | 144358556 | 144358610 | 144360114 | 144360493 | 2,5,15 | 0,0,0 | 11,13,11 | 1,6,1 | 300 | 150 | 4.39E-06 | 0.00010304 | 1.0,1.0,1.0 | 0.846,0.52,0.846 | 0.263 | ENSG000001858 | [[[[144358613, 14 | 2 |
| ENSG000001287 | HAT1 | chr2 | 171965337 | 171965517 | 171952880 | 171953001 | 171966842 | 171966949 | 192,140,185 | 0,0,0 | 125,272,154 | 6,4,4 | 300 | 150 | 0.000401048 | 0.005308162 | 1.0,1.0,1.0 | 0.912,0.971,0.95 | 0.055 | ENSG000001287 | [[[[171922461, 17 | 3 |
| ENSG000001287 | HAT1 | chr2 | 171965786 | 171965908 | 171965337 | 171965517 | 171966842 | 171966949 | 116,105,146 | 0,0,0 | 105,197,125 | 4,10,5 | 272 | 150 | 2.69E-06 | 6.65E-05 | 1.0,1.0,1.0 | 0.935,0.916,0.93 | 0.072 | ENSG000001287 | [[[[171922461, 17 | 2 |
| ENSG000001287 | HAT1 | chr2 | 171966408 | 171966513 | 171952880 | 171953001 | 171966842 | 171966949 | 72,56,119 | 0,0,0 | 59,117,47 | 6,4,4 | 255 | 150 | 6.20E-06 | 0.000140569 | 1.0,1.0,1.0 | 0.853,0.945,0.87 | 0.109 | ENSG000001287 | [[[[171922461, 17 | 3 |
| ENSG000001278 | METTL16 | chr17 | 2467760 | 2467876 | 2464207 | 2464350 | 2477685 | 2477885 | 14,4,20 | 0,0,1 | 15,25,23 | 3,5,2 | 266 | 150 | 7.50E-05 | 0.001241117 | 1.0,1.0,0.919 | 0.738,0.738,0.86 | 0.192 | ENSG000001278 | [[[[2511759, 2511 | 2 |
| ENSG000001654 | DDIAS | chr11 | 82930156 | 82930274 | 82914688 | 82914851 | 82931731 | 82931832 | 2,10,9 | 0,0,0 | 3,9,12 | 1,3,2 | 268 | 150 | 0.000147797 | 0.002235371 | 1.0,1.0,1.0 | 0.627,0.627,0.77 | 0.325 | ENSG000001654 | [[[[82901736, 829 | 2 |
| ENSG000001680 | PBK | chr8 | 27822318 | 27822488 | 27820564 | 27820694 | 27828104 | 27828198 | 177,153,219 | 0,1,0 | 95,171,110 | 3,20,3 | 300 | 150 | 1.68E-09 | 8.90E-08 | 1.0,0.987,1.0 | 0.941,0.81,0.948 | 0.096 | ENSG000001680 | [[[[27837652, 278 | 2 |
| ENSG000001834 | SF3A3 | chr1 | 37976883 | 37976953 | 37969570 | 37969735 | 37979464 | 37979533 | 68,69,74 | 0,1,1 | 52,140,95 | 2,13,6 | 220 | 150 | 1.03E-05 | 0.000219487 | 1.0,0.979,0.981 | 0.947,0.88,0.915 | 0.073 | ENSG000001834 | [[[[37989870, 379 | 3 |
| ENSG000001758 | ZDHHC21 | chr9 | 14662214 | 14662326 | 14639895 | 14640012 | 14672829 | 14672928 | 16,16,15 | 0,0,0 | 6,13,5 | 0,8,7 | 262 | 150 | 3.62E-07 | 1.11E-05 | 1.0,1.0,1.0 | 1.0,0.482,0.29 | 0.409 | ENSG000001758 | [[[[14693229, 146 | 2 |
| ENSG000001568 | MFSD14A | chr1 | 100049908 | 100050004 | 100038096 | 100038316 | 100067976 | 100068195 | 29,28,54 | 0,0,0 | 40,70,56 | 4,9,3 | 246 | 150 | 1.63E-06 | 4.25E-05 | 1.0,1.0,1.0 | 0.859,0.826,0.91 | 0.132 | ENSG000001568 | [[[[100038095, 10 | 4 |
| ENSG000001568 | MFSD14A | chr1 | 100061834 | 100061949 | 100038096 | 100038316 | 100067976 | 100068195 | 43,41,69 | 0,0,0 | 48,84,61 | 4,9,3 | 265 | 150 | 1.06E-06 | 2.89E-05 | 1.0,1.0,1.0 | 0.872,0.841,0.92 | 0.122 | ENSG000001568 | [[[[100038095, 10 | 4 |
| ENSG000000702 | TMEM260 | chr14 | 56605569 | 56605683 | 56585760 | 56585912 | 56609105 | 56609189 | 3,3,12 | 0,0,0 | 6,7,2 | 0,2,5 | 264 | 150 | 8.80E-05 | 0.001425234 | 1.0,1.0,1.0 | 1.0,0.665,0.185 | 0.383 | ENSG000000702 | [[[[56579798, 565 | 2 |
| ENSG000001668 | NEMP1 | chr12 | 57069233 | 57069306 | 57064645 | 57064739 | 57078618 | 57078763 | 9,4,9 | 0,0,0 | 15,33,14 | 2,1,8 | 223 | 150 | 1.76E-05 | 0.000350225 | 1.0,1.0,1.0 | 0.835,0.957,0.54 | 0.222 | ENSG000001668 | [[[[57078619, 570 | 3 |
| ENSG000001668 | NEMP1 | chr12 | 57070673 | 57070893 | 57069233 | 57069306 | 57078618 | 57078769 | 19,8,34 | 0,0,0 | 17,49,27 | 7,9,3 | 300 | 150 | 4.35E-08 | 1.67E-06 | 1.0,1.0,1.0 | 0.548,0.731,0.81 | 0.301 | ENSG000001668 | [[[[57078619, 570 | 2 |
| ENSG000001639 | RFC4 | chr3 | 186790325 | 186790406 | 186790141 | 186790255 | 186797534 | 186797614 | 61,44,62 | 0,0,0 | 41,87,40 | 4,7,2 | 231 | 150 | 1.24E-05 | 0.000258612 | 1.0,1.0,1.0 | 0.869,0.89,0.929 | 0.104 | ENSG000001639 | [[[[186806290, 18 | 5 |
| ENSG000001639 | RFC4 | chr3 | 186791724 | 186791850 | 186790141 | 186790255 | 186792803 | 186792947 | 1,1,1 | 0,0,0 | 3,10,5 | 3,3,0 | 276 | 150 | 0.000237072 | 0.003364583 | 1.0,1.0,1.0 | 0.352,0.644,1.0 | 0.335 | ENSG000001639 | [[[[186806290, 18 | 3 |
| ENSG000001365 | NDUFB5 | chr3 | 179614951 | 179615059 | 179604810 | 179604939 | 179616982 | 179617044 | 5,0,7 | 16,43,72 | 8,34,16 | 13,47,39 | 258 | 150 | 1.98E-08 | 8.30E-07 | 0.154,0.0,0.054 | 0.264,0.296,0.19 | -0.182 | ENSG000001365 | [[[[179604794, 17 | 2 |
| ENSG000001199 | HELLS | chr10 | 94597034 | 94597111 | 94596859 | 94596956 | 94607909 | 94608026 | 11,11,6 | 0,0,0 | 1,6,5 | 0,2,5 | 227 | 150 | 3.08E-05 | 0.000572896 | 1.0,1.0,1.0 | 1.0,0.665,0.398 | 0.312 | ENSG000001199 | [[[[94545788, 945 | 2 |
| ENSG000001241 | VAPB | chr20 | 58434601 | 58434705 | 58418210 | 58418363 | 58440906 | 58441083 | 30,35,51 | 0,1,0 | 43,73,57 | 3,6,6 | 254 | 150 | 2.11E-05 | 0.000411597 | 1.0,0.954,1.0 | 0.894,0.878,0.84 | 0.111 | ENSG000001241 | [[[[58389229, 583 | 2 |
| ENSG000001241 | VAPB | chr20 | 58434601 | 58434705 | 58418210 | 58418363 | 58444076 | 58445552 | 30,35,49 | 5,10,7 | 41,72,59 | 0,0,1 | 254 | 150 | 3.66E-10 | 2.20E-08 | 0.78,0.674,0.805 | 1.0,1.0,0.972 | -0.238 | ENSG000001241 | [[[[58389229, 583 | 3 |
| ENSG000001241 | VAPB | chr20 | 58438944 | 58439025 | 58418210 | 58418363 | 58440906 | 58441083 | 25,31,26 | 0,1,0 | 31,106,48 | 3,6,6 | 231 | 150 | 0.000100118 | 0.001599995 | 1.0,0.953,1.0 | 0.87,0.92,0.839 | 0.108 | ENSG000001241 | [[[[58389229, 583 | 2 |
| ENSG000001241 | VAPB | chr20 | 58440906 | 58441083 | 58418210 | 58418363 | 58444076 | 58444531 | 100,70,111 | 5,10,7 | 153,283,153 | 0,0,1 | 300 | 150 | 2.30E-10 | 1.43E-08 | 0.909,0.778,0.88 | 1.0,1.0,0.987 | -0.137 | ENSG000001241 | [[[[58389229, 583 | 3 |
| ENSG000001434 | SELENBP1 | chr1 | 151369003 | 151369189 | 151368198 | 151368319 | 151369712 | 151369769 | 14,15,29 | 2,0,2 | 6,17,16 | 3,8,10 | 300 | 150 | 7.07E-06 | 0.000157126 | 0.778,1.0,0.879 | 0.5,0.515,0.444 | 0.399 | ENSG000001434 | [[[[151372638, 15 | 2 |
| ENSG000001434 | SELENBP1 | chr1 | 151369003 | 151369554 | 151368198 | 151368319 | 151369712 | 151369769 | 44,39,57 | 2,0,2 | 29,39,29 | 3,8,10 | 300 | 150 | 6.52E-06 | 0.000146865 | 0.917,1.0,0.934 | 0.829,0.709,0.59 | 0.24 | ENSG000001434 | [[[[151372638, 15 | 2 |
| ENSG000001434 | SELENBP1 | chr1 | 151369441 | 151369554 | 151368198 | 151368319 | 151369712 | 151369769 | 28,24,28 | 2,0,2 | 26,23,14 | 3,8,10 | 263 | 150 | 9.23E-05 | 0.001485885 | 0.889,1.0,0.889 | 0.832,0.621,0.44 | 0.294 | ENSG000001434 | [[[[151372638, 15 | 2 |
| ENSG000002055 | NAP1L4 | chr11 | 2959769 | 2959909 | 2951779 | 2951809 | 2964679 | 2964712 | 201,135,220 | 0,0,0 | 142,367,175 | 3,17,7 | 290 | 150 | 8.13E-08 | 2.92E-06 | 1.0,1.0,1.0 | 0.961,0.918,0.92 | 0.064 | ENSG000002055 | [[[[2992254, 2992 | 4 |
| ENSG000001973 | TRIM33 | chr1 | 114463103 | 114463236 | 114433616 | 114433733 | 114464269 | 114464388 | 9,8,21 | 0,0,0 | 12,17,21 | 4,5,6 | 283 | 150 | 2.65E-08 | 1.08E-06 | 1.0,1.0,1.0 | 0.614,0.643,0.65 | 0.364 | ENSG000001973 | [[[[114510551, 11 | 2 |
| ENSG000002042 | DAXX | chr6 | 33320379 | 33320591 | 33320010 | 33320224 | 33322861 | 33322930 | 61,51,102 | 0,0,0 | 87,174,102 | 1,5,6 | 300 | 150 | 0.000669786 | 0.008246296 | 1.0,1.0,1.0 | 0.978,0.946,0.89 | 0.06 | ENSG000002042 | [[[[33322862, 333 | 3 |
| ENSG000002042 | DAXX | chr6 | 33321718 | 33321974 | 33320010 | 33320224 | 33322861 | 33323010 | 15,22,47 | 0,0,0 | 45,77,33 | 1,5,6 | 300 | 150 | 2.56E-05 | 0.000487736 | 1.0,1.0,1.0 | 0.957,0.885,0.73 | 0.142 | ENSG000002042 | [[[[33322862, 333 | 3 |
| ENSG000002042 | DAXX | chr6 | 33321718 | 33321977 | 33320010 | 33320224 | 33322861 | 33323014 | 36,37,84 | 0,0,0 | 92,133,90 | 1,5,6 | 300 | 150 | 0.000387224 | 0.005146636 | 1.0,1.0,1.0 | 0.979,0.93,0.882 | 0.07 | ENSG000002042 | [[[[33322862, 333 | 3 |
| ENSG000001439 | EML4 | chr2 | 42303103 | 42303229 | 42301240 | 42301392 | 42304483 | 42304551 | 26,15,41 | 0,0,0 | 12,51,23 | 1,5,9 | 276 | 150 | 2.35E-07 | 7.52E-06 | 1.0,1.0,1.0 | 0.867,0.847,0.58 | 0.235 | ENSG000001439 | [[[[42169353, 421 | 2 |
| ENSG000001439 | EML4 | chr2 | 42315961 | 42316050 | 42304483 | 42304551 | 42325466 | 42325554 | 22,22,36 | 0,1,0 | 12,33,26 | 1,3,6 | 239 | 150 | 3.33E-05 | 0.000613217 | 1.0,0.932,1.0 | 0.883,0.873,0.73 | 0.148 | ENSG000001439 | [[[[42169353, 421 | 2 |
| ENSG000001113 | NAA25 | chr12 | 112054387 | 112054568 | 112053557 | 112053657 | 112061180 | 112061388 | 25,17,21 | 0,0,0 | 24,50,32 | 2,6,0 | 300 | 150 | 0.00044998 | 0.005871161 | 1.0,1.0,1.0 | 0.857,0.806,1.0 | 0.112 | ENSG000001113 | [[[[112108716, 11 | 2 |
| ENSG000001113 | NAA25 | chr12 | 112060269 | 112060359 | 112053557 | 112053657 | 112061180 | 112061388 | 9,3,4 | 0,0,0 | 4,8,6 | 2,6,0 | 240 | 150 | 1.46E-05 | 0.000297646 | 1.0,1.0,1.0 | 0.556,0.455,1.0 | 0.33 | ENSG000001113 | [[[[112108716, 11 | 2 |
| ENSG000001218 | POLR3GL | chr1 | 145975306 | 145975436 | 145974824 | 145974991 | 145977482 | 145977539 | 22,14,37 | 0,0,1 | 12,36,32 | 5,9,3 | 280 | 150 | 5.22E-07 | 1.54E-05 | 1.0,1.0,0.952 | 0.563,0.682,0.85 | 0.285 | ENSG000001218 | [[[[145964690, 14 | 2 |
| ENSG000001218 | POLR3GL | chr1 | 145977083 | 145977152 | 145974824 | 145974991 | 145977482 | 145977539 | 18,11,19 | 0,0,1 | 13,16,27 | 5,9,3 | 219 | 150 | 7.99E-07 | 2.24E-05 | 1.0,1.0,0.929 | 0.64,0.549,0.86 | 0.293 | ENSG000001218 | [[[[145964690, 14 | 2 |
| ENSG000002658 | SEC22B | chr1 | 120163209 | 120163370 | 120160383 | 120160530 | 120176306 | 120176515 | 238,210,343 | 0,0,0 | 227,368,271 | 6,12,14 | 300 | 150 | 3.34E-09 | 1.66E-07 | 1.0,1.0,1.0 | 0.95,0.939,0.906 | 0.068 | ENSG000002658 | [[[[120176307, 12 | 2 |
| ENSG000001692 | SLC50A1 | chr1 | 155136827 | 155136951 | 155136298 | 155136376 | 155137978 | 155138098 | 51,51,67 | 10,8,13 | 45,132,97 | 5,7,3 | 274 | 150 | 0.000202766 | 0.002946861 | 0.736,0.777,0.73 | 0.831,0.912,0.94 | -0.146 | ENSG000001692 | [[[[155135852, 15 | 2 |
| ENSG000001280 | OSGEPL1 | chr2 | 189754140 | 189754345 | 189753915 | 189754064 | 189761646 | 189761660 | 6,3,8 | 0,0,0 | 1,8,11 | 2,6,4 | 300 | 150 | 4.89E-10 | 2.87E-08 | 1.0,1.0,1.0 | 0.2,0.4,0.579 | 0.607 | ENSG000001280 | [[[[189762685, 18 | 2 |
| ENSG000001154 | FN1 | chr2 | 215367862 | 215368027 | 215365504 | 215365630 | 215373321 | 215373411 | 291,367,589 | 0,0,0 | 3,491,103,694 | 3,20,12 | 300 | 150 | 1.91E-05 | 0.000378307 | 1.0,1.0,1.0 | 0.983,0.965,0.96 | 0.028 | ENSG000001154 | [[[[215435655, 21 | 3 |
| ENSG000001154 | FN1 | chr2 | 215371908 | 215372015 | 215365504 | 215365630 | 215373321 | 215373411 | 16,16,20 | 0,0,0 | 22,65,41 | 3,20,12 | 257 | 150 | 3.30E-09 | 1.65E-07 | 1.0,1.0,1.0 | 0.811,0.655,0.66 | 0.289 | ENSG000001154 | [[[[215435655, 21 | 3 |
| ENSG000001154 | FN1 | chr2 | 215371908 | 215372375 | 215365504 | 215365630 | 215373321 | 215373411 | 238,334,493 | 0,0,0 | 242,856,519 | 3,20,12 | 300 | 150 | 7.13E-07 | 2.03E-05 | 1.0,1.0,1.0 | 0.976,0.955,0.95 | 0.038 | ENSG000001154 | [[[[215435655, 21 | 3 |
| ENSG000001560 | PSD3 | chr8 | 18799294 | 18799353 | 18765448 | 18765538 | 18804521 | 18804602 | 4,5,12 | 0,0,0 | 2,22,13 | 2,2,7 | 209 | 150 | 1.83E-06 | 4.72E-05 | 1.0,1.0,1.0 | 0.418,0.888,0.57 | 0.374 | ENSG000001560 | [[[[19013563, 190 | 2 |
| ENSG000001388 | SEC24B | chr4 | 109524817 | 109524941 | 109521419 | 109521626 | 109527321 | 109527432 | 15,9,20 | 0,0,0 | 12,48,36 | 0,5,5 | 274 | 150 | 4.26E-05 | 0.000760907 | 1.0,1.0,1.0 | 1.0,0.84,0.798 | 0.121 | ENSG000001388 | [[[[109433815, 10 | 3 |
| ENSG000001388 | SEC24B | chr4 | 109525345 | 109525504 | 109521419 | 109521626 | 109526225 | 109526399 | 25,16,32 | 0,0,0 | 24,45,22 | 1,7,0 | 300 | 150 | 0.000314363 | 0.00430128 | 1.0,1.0,1.0 | 0.923,0.763,1.0 | 0.105 | ENSG000001388 | [[[[109433815, 10 | 2 |
| ENSG000001388 | SEC24B | chr4 | 109526225 | 109526399 | 109521419 | 109521626 | 109527321 | 109527432 | 23,14,38 | 0,0,0 | 13,16,17 | 0,5,5 | 300 | 150 | 1.48E-06 | 3.90E-05 | 1.0,1.0,1.0 | 1.0,0.615,0.63 | 0.252 | ENSG000001388 | [[[[109433815, 10 | 3 |
| ENSG000001090 | NSD2 | chr4 | 1939653 | 1939778 | 1935143 | 1935262 | 1951071 | 1951180 | 53,29,62 | 7,1,2 | 46,151,68 | 11,30,17 | 275 | 150 | 0.000214123 | 0.003093032 | 0.805,0.941,0.94 | 0.695,0.733,0.68 | 0.192 | ENSG000001090 | [[[[1871393, 1871 | 2 |
| ENSG000001090 | NSD2 | chr4 | 1955982 | 1956188 | 1955578 | 1955849 | 1959470 | 1959740 | 81,38,88 | 0,0,0 | 55,113,84 | 4,17,7 | 300 | 150 | 5.07E-10 | 2.97E-08 | 1.0,1.0,1.0 | 0.873,0.769,0.85 | 0.167 | ENSG000001090 | [[[[1871393, 1871 | 2 |
| ENSG000000884 | TGDS | chr13 | 94576313 | 94576411 | 94574050 | 94574852 | 94578729 | 94578773 | 17,13,19 | 0,0,0 | 14,59,24 | 3,4,1 | 248 | 150 | 0.000628043 | 0.007810613 | 1.0,1.0,1.0 | 0.738,0.899,0.93 | 0.142 | ENSG000000884 | [[[[94596051, 945 | 3 |
| ENSG000001628 | B3GALNT2 | chr1 | 235458602 | 235458786 | 235455558 | 235455684 | 235494680 | 235494828 | 17,18,22 | 0,0,0 | 18,51,28 | 2,7,0 | 300 | 150 | 5.60E-05 | 0.00096215 | 1.0,1.0,1.0 | 0.818,0.785,1.0 | 0.132 | ENSG000001628 | [[[[235504141, 23 | 6 |
| ENSG000001628 | B3GALNT2 | chr1 | 235470849 | 235470960 | 235458602 | 235458786 | 235480053 | 235480149 | 27,13,37 | 3,0,0 | 24,51,38 | 0,8,13 | 261 | 150 | 0.000822402 | 0.009836039 | 0.838,1.0,1.0 | 1.0,0.786,0.627 | 0.142 | ENSG000001628 | [[[[235504141, 23 | 2 |
| ENSG000000828 | XPO1 | chr2 | 61485767 | 61485962 | 61483936 | 61484105 | 61490641 | 61490776 | 181,156,259 | 0,0,0 | 139,324,155 | 31,33,19 | 300 | 150 | 0 | 0 | 1.0,1.0,1.0 | 0.692,0.831,0.80 | 0.225 | ENSG000000828 | [[[[61537562, 615 | 3 |
| ENSG000000828 | XPO1 | chr2 | 61485767 | 61485962 | 61483936 | 61484105 | 61492034 | 61492198 | 181,156,259 | 0,0,0 | 135,304,157 | 9,12,4 | 300 | 150 | 3.14E-08 | 1.25E-06 | 1.0,1.0,1.0 | 0.882,0.927,0.95 | 0.08 | ENSG000000828 | [[[[61537562, 615 | 4 |
| ENSG000000828 | XPO1 | chr2 | 61488164 | 61488271 | 61483936 | 61484105 | 61488587 | 61488663 | 136,124,153 | 0,0,0 | 97,192,132 | 6,4,4 | 257 | 150 | 0.000195363 | 0.002856701 | 1.0,1.0,1.0 | 0.904,0.966,0.95 | 0.06 | ENSG000000828 | [[[[61537562, 615 | 2 |
| ENSG000000828 | XPO1 | chr2 | 61488587 | 61488771 | 61481184 | 61481281 | 61492034 | 61492198 | 4,1,1 | 0,0,0 | 1,4,1 | 2,4,0 | 300 | 150 | 2.07E-05 | 0.000406021 | 1.0,1.0,1.0 | 0.2,0.333,1.0 | 0.489 | ENSG000000828 | [[[[61537562, 615 | 7 |
| ENSG000000828 | XPO1 | chr2 | 61488587 | 61488771 | 61483936 | 61484105 | 61490641 | 61490776 | 233,220,331 | 0,0,0 | 134,396,222 | 31,33,19 | 300 | 150 | 0 | 0 | 1.0,1.0,1.0 | 0.684,0.857,0.85 | 0.202 | ENSG000000828 | [[[[61537562, 615 | 3 |
| ENSG000000828 | XPO1 | chr2 | 61488587 | 61488771 | 61483936 | 61484105 | 61492034 | 61492198 | 4,1,1 | 0,0,0 | 6,6,5 | 9,12,4 | 300 | 150 | 5.91E-09 | 2.76E-07 | 1.0,1.0,1.0 | 0.25,0.2,0.385 | 0.722 | ENSG000000828 | [[[[61537562, 615 | 4 |
| ENSG000000828 | XPO1 | chr2 | 61490641 | 61490776 | 61483936 | 61484105 | 61492034 | 61492198 | 185,162,258 | 0,0,0 | 174,322,229 | 9,12,4 | 285 | 150 | 4.26E-07 | 1.28E-05 | 1.0,1.0,1.0 | 0.911,0.934,0.96 | 0.062 | ENSG000000828 | [[[[61537562, 615 | 4 |
| ENSG000000828 | XPO1 | chr2 | 61492034 | 61492198 | 61490641 | 61490776 | 61493893 | 61494091 | 204,184,286 | 0,0,0 | 130,276,181 | 7,18,17 | 300 | 150 | 1.47E-14 | 1.72E-12 | 1.0,1.0,1.0 | 0.903,0.885,0.84 | 0.123 | ENSG000000828 | [[[[61537562, 615 | 4 |
| ENSG000000828 | XPO1 | chr2 | 61492324 | 61492481 | 61492034 | 61492198 | 61493893 | 61494091 | 240,152,279 | 0,0,0 | 125,296,185 | 9,12,2 | 300 | 150 | 1.24E-07 | 4.23E-06 | 1.0,1.0,1.0 | 0.874,0.925,0.97 | 0.074 | ENSG000000828 | [[[[61537562, 615 | 3 |
| ENSG000001633 | POGLUT1 | chr3 | 119480050 | 119480172 | 119477312 | 119477448 | 119488928 | 119488987 | 3,12,12 | 0,0,0 | 4,10,3 | 2,7,3 | 272 | 150 | 1.35E-10 | 8.70E-09 | 1.0,1.0,1.0 | 0.524,0.441,0.35 | 0.56 | ENSG000001633 | [[[[119468963, 11 | 3 |
| ENSG000001434 | ANP32E | chr1 | 150229071 | 150229114 | 150226607 | 150226795 | 150235732 | 150235829 | 21,27,34 | 0,0,0 | 22,37,32 | 7,3,1 | 193 | 150 | 2.35E-05 | 0.000453824 | 1.0,1.0,1.0 | 0.71,0.906,0.961 | 0.141 | ENSG000001434 | [[[[150235733, 15 | 3 |
| ENSG000001434 | ANP32E | chr1 | 150229071 | 150229144 | 150226607 | 150226795 | 150235732 | 150235983 | 31,39,58 | 0,0,0 | 42,84,62 | 7,3,1 | 223 | 150 | 0.000219115 | 0.003154376 | 1.0,1.0,1.0 | 0.801,0.95,0.977 | 0.091 | ENSG000001434 | [[[[150235733, 15 | 3 |
| ENSG000001434 | ANP32E | chr1 | 150230570 | 150230693 | 150226607 | 150226795 | 150235732 | 150236064 | 6,1,2 | 0,0,0 | 8,30,26 | 7,3,1 | 273 | 150 | 1.65E-05 | 0.000330678 | 1.0,1.0,1.0 | 0.386,0.846,0.93 | 0.278 | ENSG000001434 | [[[[150235733, 15 | 3 |
| ENSG000001431 | MPC2 | chr1 | 167919978 | 167920090 | 167916728 | 167918359 | 167924496 | 167924537 | 64,62,62 | 0,0,0 | 41,118,63 | 5,8,14 | 262 | 150 | 1.03E-09 | 5.70E-08 | 1.0,1.0,1.0 | 0.824,0.894,0.72 | 0.187 | ENSG000001431 | [[[[167936939, 16 | 2 |
| ENSG000001167 | BCAS2 | chr1 | 114575589 | 114575751 | 114570699 | 114570750 | 114581298 | 114581391 | 106,115,147 | 0,1,0 | 142,294,170 | 7,24,11 | 300 | 150 | 5.30E-10 | 3.09E-08 | 1.0,0.983,1.0 | 0.91,0.86,0.885 | 0.109 | ENSG000001167 | [[[[114581499, 11 | 2 |
| ENSG000001167 | BCAS2 | chr1 | 114576687 | 114576758 | 114570699 | 114570750 | 114581298 | 114581391 | 59,57,87 | 0,1,0 | 71,146,84 | 7,24,11 | 221 | 150 | 5.85E-11 | 3.96E-09 | 1.0,0.975,1.0 | 0.873,0.805,0.83 | 0.153 | ENSG000001167 | [[[[114581499, 11 | 2 |
| ENSG000000873 | GMCL1 | chr2 | 69871744 | 69871832 | 69849651 | 69849742 | 69878908 | 69878990 | 5,6,19 | 0,0,0 | 6,16,23 | 1,7,0 | 238 | 150 | 0.000123808 | 0.001918629 | 1.0,1.0,1.0 | 0.791,0.59,1.0 | 0.206 | ENSG000000873 | [[[[69829660, 698 | 5 |
| ENSG000001791 | TCAIM | chr3 | 44400354 | 44400587 | 44392854 | 44392977 | 44401202 | 44401334 | 20,17,28 | 0,0,0 | 17,52,26 | 0,6,2 | 300 | 150 | 0.000288645 | 0.003993135 | 1.0,1.0,1.0 | 1.0,0.813,0.867 | 0.107 | ENSG000001791 | [[[[44338766, 443 | 3 |
| ENSG000001368 | ECPAS | chr9 | 111410979 | 111411142 | 111397029 | 111397153 | 111412013 | 111412148 | 70,43,106 | 0,0,0 | 33,121,94 | 1,5,6 | 300 | 150 | 7.23E-05 | 0.001201482 | 1.0,1.0,1.0 | 0.943,0.924,0.88 | 0.082 | ENSG000001368 | [[[[111484116, 11 | 3 |
| ENSG000001440 | SFXN5 | chr2 | 72968447 | 72968533 | 72942035 | 72945099 | 72988257 | 72988348 | 7,7,11 | 1,2,0 | 6,6,4 | 2,8,2 | 236 | 150 | 0.000700711 | 0.008573016 | 0.816,0.69,1.0 | 0.656,0.323,0.56 | 0.322 | ENSG000001440 | [[[[73071604, 730 | 3 |
| ENSG000001470 | TATDN1 | chr8 | 124515745 | 124515788 | 124508602 | 124508688 | 124518817 | 124518881 | 16,15,37 | 0,0,0 | 34,85,37 | 2,7,4 | 193 | 150 | 9.02E-05 | 0.001458801 | 1.0,1.0,1.0 | 0.93,0.904,0.878 | 0.096 | ENSG000001470 | [[[[124539025, 12 | 2 |
| ENSG000001470 | TATDN1 | chr8 | 124515886 | 124516030 | 124508602 | 124508688 | 124518817 | 124518881 | 59,41,63 | 0,0,0 | 49,104,64 | 2,7,4 | 294 | 150 | 2.34E-05 | 0.000451166 | 1.0,1.0,1.0 | 0.926,0.883,0.89 | 0.1 | ENSG000001470 | [[[[124539025, 12 | 2 |
| ENSG000001470 | TATDN1 | chr8 | 124515886 | 124516030 | 124508602 | 124508688 | 124539024 | 124539458 | 1,0,0 | 2,2,5 | 5,13,0 | 1,2,2 | 294 | 150 | 2.66E-05 | 0.000502515 | 0.203,0.0,0.0 | 0.718,0.768,0.0 | -0.428 | ENSG000001470 | [[[[124539025, 12 | 5 |

| | | | | | | | | | | | | | | | | | | | | | | |
|---|---|---|---|---|---|---|---|---|---|---|---|---|---|---|---|---|---|---|---|---|---|---|
| ENSG000001476 | TATDN1 | chr8 | 124518741 | 124518881 | 124515886 | 124516030 | 124522936 | 124523002 | 38,34,41 | 4,2,6 | 32,81,32 | 7,32,20 | 290 | 150 | 0.000177964 | 0.002631464 | 0.831,0.898,0.77 | 0.703,0.567,0.45 | 0.262 | ENSG000001476 | [[[[124539025, 12 | 2 |
| ENSG000001476 | TATDN1 | chr8 | 124518817 | 124518881 | 124508602 | 124508688 | 124539024 | 124539076 | 0,0,0 | 2,2,5 | 0,4,3 | 1,2,2 | 214 | 150 | 5.06E-05 | 0.000882505 | 0.0,0.0,0.0 | 0.0,0.584,0.513 | -0.366 | ENSG000001476 | [[[[124539025, 12 | 5 |
| ENSG000001476 | TATDN1 | chr8 | 124518817 | 124518881 | 124515886 | 124516030 | 124522936 | 124523002 | 50,44,63 | 4,2,6 | 45,114,47 | 7,32,20 | 214 | 150 | 4.61E-05 | 0.00081301 | 0.898,0.939,0.88 | 0.818,0.714,0.62 | 0.188 | ENSG000001476 | [[[[124539025, 12 | 2 |
| ENSG000001476 | TATDN1 | chr8 | 124518817 | 124518881 | 124515886 | 124516030 | 124539024 | 124539076 | 30,21,28 | 1,0,0 | 24,54,24 | 3,13,0 | 214 | 150 | 1.64E-05 | 0.000329686 | 0.955,1.0,1.0 | 0.849,0.744,1.0 | 0.121 | ENSG000001476 | [[[[124539025, 12 | 3 |
| ENSG000001476 | TATDN1 | chr8 | 124522936 | 124523002 | 124515886 | 124516030 | 124539024 | 124539070 | 4,2,6 | 1,0,0 | 6,32,17 | 3,13,0 | 216 | 150 | 0.000299793 | 0.004124141 | 0.735,1.0,1.0 | 0.581,0.631,1.0 | 0.174 | ENSG000001476 | [[[[124539025, 12 | 3 |
| ENSG000001138 | CRBN | chr3 | 3153959 | 3154075 | 3150010 | 3151045 | 3154746 | 3154831 | 5,8,16 | 0,0,0 | 9,29,18 | 1,9,0 | 266 | 150 | 2.97E-05 | 0.000554667 | 1.0,1.0,1.0 | 0.835,0.645,1.0 | 0.173 | ENSG000001138 | [[[[3179621, 3179 | 3 |
| ENSG000001210 | COPS5 | chr8 | 67059210 | 67059445 | 67057379 | 67057445 | 67061853 | 67062317 | 126,124,140 | 0,0,0 | 109,265,146 | 9,29,4 | 300 | 150 | 1.71E-11 | 1.26E-09 | 1.0,1.0,1.0 | 0.858,0.82,0.948 | 0.125 | ENSG000001210 | [[[[67061854, 670 | 2 |
| ENSG000001571 | SMG1 | chr16 | 18838356 | 18838466 | 18836359 | 18836532 | 18838550 | 18838689 | 17,5,15 | 0,0,0 | 0,13,11 | 2,5,2 | 260 | 150 | 1.26E-06 | 3.36E-05 | 1.0,1.0,1.0 | 0.0,0.6,0.76 | 0.547 | ENSG000001571 | [[[[18925950, 189 | 3 |
| ENSG000001627 | FLVCR1 | chr1 | 212888488 | 212888594 | 212885292 | 212885396 | 212889145 | 212889257 | 2,3,10 | 0,0,0 | 11,19,13 | 0,12,1 | 256 | 150 | 4.20E-05 | 0.000751715 | 1.0,1.0,1.0 | 1.0,0.481,0.884 | 0.212 | ENSG000001627 | [[[[212858275, 21 | 2 |
| ENSG000002653 | TIMM23 | chr10 | 45975453 | 45975512 | 45972448 | 45972730 | 45988736 | 45988847 | 39,82,144 | 0,0,0 | 74,200,127 | 9,24,10 | 209 | 150 | 1.71E-11 | 1.26E-09 | 1.0,1.0,1.0 | 0.855,0.857,0.90 | 0.129 | ENSG000002653 | [[[[45972489, 459 | 4 |
| ENSG000001146 | MRPL3 | chr3 | 131489980 | 131490080 | 131471170 | 131471279 | 131500429 | 131500521 | 1,1,2 | 0,0,0 | 27,53,26 | 4,4,5 | 250 | 150 | 4.03E-05 | 0.000724032 | 1.0,1.0,1.0 | 0.802,0.888,0.75 | 0.184 | ENSG000001146 | [[[[131502730, 13 | 3 |
| ENSG000001146 | MRPL3 | chr3 | 131498178 | 131498277 | 131471170 | 131471279 | 131500429 | 131500484 | 168,173,233 | 0,0,0 | 95,177,96 | 4,4,5 | 249 | 150 | 0.000190369 | 0.002794668 | 1.0,1.0,1.0 | 0.935,0.964,0.92 | 0.06 | ENSG000001146 | [[[[131502730, 13 | 3 |
| ENSG000001146 | MRPL3 | chr3 | 131501530 | 131501715 | 131489980 | 131490080 | 131502729 | 131502951 | 59,108,240 | 0,0,0 | 79,131,84 | 3,6,2 | 300 | 150 | 0.000256037 | 0.003594899 | 1.0,1.0,1.0 | 0.929,0.916,0.95 | 0.067 | ENSG000001146 | [[[[131502730, 13 | 3 |
| ENSG000001146 | MRPL3 | chr3 | 131501530 | 131501715 | 131489980 | 131490080 | 131502848 | 131502983 | 3,3,5 | 0,0,0 | 3,13,3 | 1,6,6 | 300 | 150 | 5.20E-10 | 3.04E-08 | 1.0,1.0,1.0 | 0.6,0.52,0.2 | 0.56 | ENSG000001146 | [[[[131502730, 13 | 3 |
| ENSG000001430 | MTF2 | chr1 | 93127231 | 93127299 | 93120548 | 93120672 | 93133702 | 93133808 | 18,17,26 | 0,0,1 | 28,68,55 | 5,5,2 | 218 | 150 | 0.000476769 | 0.006181528 | 1.0,1.0,0.947 | 0.794,0.903,0.95 | 0.1 | ENSG000001430 | [[[[93079283, 930 | 2 |
| ENSG000001430 | MTF2 | chr1 | 93129277 | 93129448 | 93120548 | 93120672 | 93133702 | 93133808 | 23,18,26 | 0,0,1 | 33,70,26 | 5,5,2 | 300 | 150 | 8.96E-05 | 0.001449224 | 1.0,1.0,0.929 | 0.767,0.875,0.86 | 0.14 | ENSG000001430 | [[[[93079283, 930 | 2 |
| ENSG000001430 | MTF2 | chr1 | 93133702 | 93133808 | 93127231 | 93127299 | 93133927 | 93133980 | 41,42,53 | 0,0,0 | 41,122,70 | 6,17,11 | 256 | 150 | 3.22E-10 | 1.95E-08 | 1.0,1.0,1.0 | 0.8,0.808,0.789 | 0.201 | ENSG000001430 | [[[[93079283, 930 | 2 |
| ENSG000001430 | MTF2 | chr1 | 93133702 | 93133808 | 93127231 | 93127299 | 93134090 | 93134195 | 4,1,5 | 0,0,0 | 3,15,6 | 1,8,5 | 256 | 150 | 3.19E-09 | 1.60E-07 | 1.0,1.0,1.0 | 0.637,0.523,0.41 | 0.476 | ENSG000001430 | [[[[93079283, 930 | 3 |
| ENSG000001430 | MTF2 | chr1 | 93133927 | 93133980 | 93127231 | 93127299 | 93134090 | 93134195 | 11,13,10 | 0,0,0 | 36,44,42 | 1,8,5 | 203 | 150 | 3.54E-05 | 0.00064762 | 1.0,1.0,1.0 | 0.964,0.803,0.86 | 0.124 | ENSG000001430 | [[[[93079283, 930 | 3 |
| ENSG000000507 | MAPK9 | chr5 | 180269279 | 180269409 | 180248972 | 180249138 | 180279795 | 180280036 | 2,1,13 | 0,0,0 | 2,17,3 | 4,1,5 | 280 | 150 | 1.06E-07 | 3.68E-06 | 1.0,1.0,1.0 | 0.211,0.901,0.24 | 0.548 | ENSG000000507 | [[[[180291848, 18 | 3 |
| ENSG000000507 | MAPK9 | chr5 | 180269279 | 180269409 | 180248972 | 180249138 | 180280439 | 180280581 | 36,29,58 | 0,0,0 | 19,64,25 | 2,9,2 | 280 | 150 | 4.87E-07 | 1.45E-05 | 1.0,1.0,1.0 | 0.836,0.792,0.87 | 0.167 | ENSG000000507 | [[[[180291848, 18 | 3 |
| ENSG000000507 | MAPK9 | chr5 | 180279795 | 180280036 | 180248972 | 180249138 | 180280439 | 180280581 | 1,1,2 | 0,0,0 | 30,25,32 | 2,9,2 | 300 | 150 | 9.61E-06 | 0.000206493 | 1.0,1.0,1.0 | 0.882,0.581,0.88 | 0.216 | ENSG000000507 | [[[[180291848, 18 | 3 |
| ENSG000000359 | RFC1 | chr4 | 39302736 | 39302874 | 39300259 | 39300414 | 39303059 | 39303151 | 28,21,27 | 0,0,0 | 24,49,29 | 1,18,2 | 288 | 150 | 6.49E-07 | 1.87E-05 | 1.0,1.0,1.0 | 0.926,0.586,0.88 | 0.202 | ENSG000000359 | [[[[39366239, 393 | 3 |
| ENSG000001516 | ADAM17 | chr2 | 9521202 | 9521316 | 9509978 | 9510131 | 9523248 | 9523338 | 36,13,41 | 0,0,0 | 39,76,50 | 10,28,19 | 264 | 150 | 4.29E-14 | 4.68E-12 | 1.0,1.0,1.0 | 0.689,0.607,0.59 | 0.368 | ENSG000001516 | [[[[9555509, 9555 | 3 |
| ENSG000001206 | MTRF1 | chr13 | 41223255 | 41223354 | 41216368 | 41217228 | 41233889 | 41234007 | 2,2,8 | 0,0,0 | 3,4,2 | 4,4,1 | 249 | 150 | 4.25E-10 | 2.51E-08 | 1.0,1.0,1.0 | 0.311,0.376,0.54 | 0.589 | ENSG000001206 | [[[[41263485, 412 | 2 |
| ENSG000001206 | MTRF1 | chr13 | 41226431 | 41226568 | 41216368 | 41217228 | 41233889 | 41234007 | 5,5,10 | 0,0,0 | 15,5,10 | 4,4,1 | 287 | 150 | 5.38E-07 | 1.58E-05 | 1.0,1.0,1.0 | 0.662,0.395,0.83 | 0.368 | ENSG000001206 | [[[[41263485, 412 | 2 |
| ENSG000001005 | PSMC6 | chr14 | 52708482 | 52708522 | 52707183 | 52707304 | 52708763 | 52708816 | 69,72,103 | 0,0,0 | 72,196,103 | 27,27,28 | 190 | 150 | 0 | 0 | 1.0,1.0,1.0 | 0.678,0.851,0.74 | 0.242 | ENSG000001005 | [[[[52707200, 527 | 2 |
| ENSG000001005 | PSMC6 | chr14 | 52708763 | 52708816 | 52707218 | 52707304 | 52711100 | 52711168 | 111,102,142 | 0,0,0 | 152,315,193 | 9,15,6 | 203 | 150 | 9.33E-07 | 2.58E-05 | 1.0,1.0,1.0 | 0.926,0.939,0.96 | 0.058 | ENSG000001005 | [[[[52707200, 527 | 3 |
| ENSG000001005 | PSMC6 | chr14 | 52711100 | 52711168 | 52708763 | 52708816 | 52718080 | 52718142 | 94,87,147 | 0,0,0 | 124,255,145 | 1,9,6 | 218 | 150 | 0.000711828 | 0.008683939 | 1.0,1.0,1.0 | 0.988,0.951,0.94 | 0.039 | ENSG000001005 | [[[[52707200, 527 | 3 |
| ENSG000001005 | PSMC6 | chr14 | 52713880 | 52713968 | 52708763 | 52708816 | 52718080 | 52718142 | 112,99,146 | 0,0,0 | 111,255,150 | 1,9,6 | 238 | 150 | 0.000334464 | 0.004532362 | 1.0,1.0,1.0 | 0.986,0.947,0.94 | 0.042 | ENSG000001005 | [[[[52707200, 527 | 3 |
| ENSG000001005 | PSMC6 | chr14 | 52718080 | 52718142 | 52713880 | 52713968 | 52718976 | 52719038 | 166,149,263 | 0,0,0 | 151,424,257 | 7,14,8 | 212 | 150 | 1.57E-06 | 4.12E-05 | 1.0,1.0,1.0 | 0.939,0.955,0.95 | 0.049 | ENSG000001005 | [[[[52707200, 527 | 2 |
| ENSG000001005 | PSMC6 | chr14 | 52718228 | 52718352 | 52713880 | 52713968 | 52718976 | 52719038 | 111,106,132 | 0,0,0 | 138,244,162 | 7,14,8 | 274 | 150 | 3.96E-08 | 1.54E-06 | 1.0,1.0,1.0 | 0.915,0.905,0.91 | 0.088 | ENSG000001005 | [[[[52707200, 527 | 2 |
| ENSG000001108 | PPFIBP1 | chr12 | 27647728 | 27647842 | 27646061 | 27646148 | 27656615 | 27656730 | 26,12,48 | 0,0,0 | 5,17,12 | 4,3,1 | 264 | 150 | 4.21E-06 | 9.92E-05 | 1.0,1.0,1.0 | 0.415,0.763,0.87 | 0.317 | ENSG000001108 | [[[[27524206, 275 | 3 |
| ENSG000001588 | B4GALT3 | chr1 | 161174992 | 161175228 | 161173604 | 161174049 | 161176433 | 161176508 | 39,37,88 | 0,0,0 | 74,175,107 | 14,16,11 | 300 | 150 | 3.53E-10 | 2.13E-08 | 1.0,1.0,1.0 | 0.725,0.845,0.82 | 0.2 | ENSG000001588 | [[[[161177423, 16 | 2 |
| ENSG000001588 | B4GALT3 | chr1 | 161175807 | 161176074 | 161173858 | 161174049 | 161176433 | 161176579 | 32,56,73 | 0,0,0 | 42,83,74 | 14,16,11 | 300 | 150 | 3.22E-13 | 3.12E-11 | 1.0,1.0,1.0 | 0.6,0.722,0.771 | 0.302 | ENSG000001588 | [[[[161177423, 16 | 2 |
| ENSG000001588 | B4GALT3 | chr1 | 161175807 | 161176095 | 161173858 | 161174049 | 161176433 | 161176508 | 15,23,52 | 0,0,0 | 45,72,62 | 14,16,11 | 300 | 150 | 3.24E-11 | 2.28E-09 | 1.0,1.0,1.0 | 0.616,0.692,0.73 | 0.318 | ENSG000001588 | [[[[161177423, 16 | 2 |
| ENSG000001622 | TAF6L | chr11 | 62778868 | 62778963 | 62778284 | 62778335 | 62782692 | 62782825 | 13,20,24 | 0,0,0 | 23,41,18 | 7,22,8 | 245 | 150 | 5.49E-12 | 4.46E-10 | 1.0,1.0,1.0 | 0.668,0.533,0.57 | 0.407 | ENSG000001622 | [[[[62771396, 627 | 3 |
| ENSG000001622 | TAF6L | chr11 | 62782112 | 62782333 | 62778284 | 62778335 | 62782692 | 62782825 | 36,25,39 | 0,0,0 | 22,44,25 | 7,22,8 | 300 | 150 | 9.99E-15 | 1.20E-12 | 1.0,1.0,1.0 | 0.611,0.5,0.61 | 0.426 | ENSG000001622 | [[[[62771396, 627 | 3 |
| ENSG000001622 | TAF6L | chr11 | 62782112 | 62782333 | 62778868 | 62778963 | 62782692 | 62782825 | 36,25,38 | 0,0,0 | 22,46,27 | 2,7,2 | 300 | 150 | 2.00E-06 | 5.12E-05 | 1.0,1.0,1.0 | 0.846,0.767,0.87 | 0.172 | ENSG000001622 | [[[[62771396, 627 | 2 |
| ENSG000000921 | SCFD1 | chr14 | 30630476 | 30630565 | 30622318 | 30622399 | 30633946 | 30634037 | 17,21,46 | 0,3,0 | 14,33,21 | 4,8,4 | 239 | 150 | 3.34E-05 | 0.000614289 | 1.0,0.815,1.0 | 0.687,0.721,0.76 | 0.213 | ENSG000000921 | [[[[30622319, 306 | 2 |
| ENSG000000921 | SCFD1 | chr14 | 30638124 | 30638247 | 30633946 | 30634037 | 30643315 | 30643405 | 14,14,30 | 0,0,0 | 18,23,18 | 4,12,3 | 273 | 150 | 4.35E-09 | 2.10E-07 | 1.0,1.0,1.0 | 0.712,0.513,0.76 | 0.336 | ENSG000000921 | [[[[30622319, 306 | 2 |
| ENSG000000872 | ADD1 | chr4 | 2884514 | 2884666 | 2875895 | 2876110 | 2894012 | 2894093 | 66,37,82 | 0,0,0 | 19,76,29 | 12,34,15 | 300 | 150 | 0 | 0 | 1.0,1.0,1.0 | 0.442,0.528,0.49 | 0.513 | ENSG000000872 | [[[[2843844, 2844 | 2 |
| ENSG000001428 | ITGB3BP | chr1 | 63490082 | 63490218 | 63454889 | 63454968 | 63523128 | 63523255 | 2,6,5 | 0,0,0 | 5,3,5 | 0,3,3 | 286 | 150 | 6.56E-05 | 0.001106861 | 1.0,1.0,1.0 | 1.0,0.344,0.466 | 0.397 | ENSG000001428 | [[[[63523129, 635 | 3 |
| ENSG000001462 | FAXC | chr6 | 99291703 | 99291820 | 99271168 | 99281453 | 99333350 | 99333547 | 55,33,61 | 0,0,0 | 71,147,112 | 7,5,11 | 267 | 150 | 1.05E-06 | 2.88E-05 | 1.0,1.0,1.0 | 0.851,0.943,0.85 | 0.118 | ENSG000001462 | [[[[99349107, 993 | 2 |
| ENSG000001967 | CD47 | chr3 | 108051938 | 108051970 | 108043093 | 108047292 | 108060744 | 108060852 | 18,14,12 | 0,0,0 | 14,38,15 | 9,18,11 | 182 | 150 | 6.85E-12 | 5.43E-10 | 1.0,1.0,1.0 | 0.562,0.635,0.52 | 0.425 | ENSG000001967 | [[[[108090863, 10 | 6 |
| ENSG000001967 | CD47 | chr3 | 108057476 | 108057569 | 108043093 | 108047292 | 108060744 | 108060852 | 9,5,10 | 0,0,0 | 7,11,1 | 9,18,11 | 243 | 150 | 6.77E-14 | 7.19E-12 | 1.0,1.0,1.0 | 0.324,0.274,0.05 | 0.783 | ENSG000001967 | [[[[108090863, 10 | 6 |
| ENSG000001967 | CD47 | chr3 | 108057476 | 108057569 | 108051938 | 108051970 | 108060744 | 108060852 | 57,29,61 | 0,0,0 | 24,50,39 | 8,5,6 | 243 | 150 | 3.26E-09 | 1.63E-07 | 1.0,1.0,1.0 | 0.649,0.861,0.8 | 0.23 | ENSG000001967 | [[[[108090863, 10 | 3 |
| ENSG000001967 | CD47 | chr3 | 108059451 | 108059544 | 108043093 | 108047292 | 108060744 | 108060852 | 57,70,79 | 0,0,0 | 35,94,42 | 9,18,11 | 243 | 150 | 2.22E-15 | 2.87E-13 | 1.0,1.0,1.0 | 0.706,0.763,0.70 | 0.276 | ENSG000001967 | [[[[108090863, 10 | 6 |
| ENSG000001354 | CDK4 | chr12 | 57750655 | 57750671 | 57749453 | 57749504 | 57751296 | 57751342 | 45,75,84 | 0,0,0 | 80,131,74 | 11,27,13 | 166 | 150 | 8.29E-14 | 8.69E-12 | 1.0,1.0,1.0 | 0.868,0.814,0.83 | 0.16 | ENSG000001354 | [[[[57752175, 577 | 2 |
| ENSG000001354 | CDK4 | chr12 | 57750655 | 57750671 | 57749453 | 57749504 | 57751499 | 57751736 | 45,75,84 | 0,0,0 | 80,132,73 | 31,60,34 | 166 | 150 | 0 | 0 | 1.0,1.0,1.0 | 0.7,0.665,0.66 | 0.325 | ENSG000001354 | [[[[57752175, 577 | 3 |
| ENSG000001354 | CDK4 | chr12 | 57750655 | 57750765 | 57749453 | 57749504 | 57751206 | 57751342 | 295,384,543 | 0,0,0 | 440,660,373 | 15,36,19 | 260 | 150 | 2.22E-16 | 3.17E-14 | 1.0,1.0,1.0 | 0.944,0.914,0.91 | 0.074 | ENSG000001354 | [[[[57752175, 577 | 2 |
| ENSG000001354 | CDK4 | chr12 | 57750655 | 57750765 | 57749453 | 57749504 | 57751499 | 57751659 | 295,384,536 | 0,0,0 | 468,693,391 | 31,60,34 | 260 | 150 | 0 | 0 | 1.0,1.0,1.0 | 0.897,0.87,0.869 | 0.121 | ENSG000001354 | [[[[57752175, 577 | 3 |
| ENSG000001354 | CDK4 | chr12 | 57750922 | 57751090 | 57749453 | 57749504 | 57751206 | 57751342 | 538,546,754 | 0,0,0 | 629,980,637 | 15,36,19 | 300 | 150 | 5.88E-15 | 7.27E-13 | 1.0,1.0,1.0 | 0.954,0.932,0.94 | 0.057 | ENSG000001354 | [[[[57752175, 577 | 2 |
| ENSG000001354 | CDK4 | chr12 | 57750922 | 57751090 | 57749453 | 57749504 | 57751499 | 57751736 | 1,4,2 | 0,0,0 | 6,13,8 | 31,60,34 | 300 | 150 | 9.68E-10 | 5.38E-08 | 1.0,1.0,1.0 | 0.088,0.098,0.10 | 0.903 | ENSG000001354 | [[[[57752175, 577 | 3 |
| ENSG000001354 | CDK4 | chr12 | 57750922 | 57751090 | 57750655 | 57750765 | 57751499 | 57751736 | 8,127,281,023 | 0,0,0 | 7,081,485,840 | 31,34,18 | 300 | 150 | 1.11E-16 | 1.65E-14 | 1.0,1.0,1.0 | 0.919,0.956,0.95 | 0.055 | ENSG000001354 | [[[[57752175, 577 | 2 |
| ENSG000001354 | CDK4 | chr12 | 57751206 | 57751342 | 57749453 | 57749504 | 57751499 | 57751659 | 477,466,758 | 0,0,0 | 602,995,578 | 31,60,34 | 286 | 150 | 0 | 0 | 1.0,1.0,1.0 | 0.911,0.897,0.89 | 0.098 | ENSG000001354 | [[[[57752175, 577 | 3 |
| ENSG000001354 | CDK4 | chr12 | 57751206 | 57751342 | 57750655 | 57750765 | 57751499 | 57751659 | 477,466,763 | 0,0,0 | 591,962,559 | 31,34,18 | 286 | 150 | 0 | 0 | 1.0,1.0,1.0 | 0.909,0.937,0.94 | 0.071 | ENSG000001354 | [[[[57752175, 577 | 2 |
| ENSG000001354 | CDK4 | chr12 | 57751296 | 57751342 | 57749453 | 57749504 | 57751499 | 57751736 | 135,165,242 | 0,0,0 | 204,354,190 | 31,60,34 | 196 | 150 | 0 | 0 | 1.0,1.0,1.0 | 0.834,0.819,0.81 | 0.179 | ENSG000001354 | [[[[57752175, 577 | 3 |
| ENSG000001354 | CDK4 | chr12 | 57751296 | 57751342 | 57750678 | 57750765 | 57751499 | 57751736 | 135,165,242 | 0,0,0 | 193,332,183 | 31,34,18 | 196 | 150 | 0 | 0 | 1.0,1.0,1.0 | 0.827,0.882,0.88 | 0.135 | ENSG000001354 | [[[[57752175, 577 | 2 |
| ENSG000001154 | GLS | chr2 | 190931544 | 190931637 | 190927003 | 190927482 | 190953564 | 190953626 | 10,8,12 | 0,0,0 | 8,12,5 | 6,9,7 | 243 | 150 | 1.37E-11 | 1.03E-09 | 1.0,1.0,1.0 | 0.451,0.451,0.30 | 0.597 | ENSG000001154 | [[[[190880821, 19 | 2 |
| ENSG000001427 | MAP3K6 | chr1 | 27362090 | 27362250 | 27361704 | 27361867 | 27363441 | 27363548 | 13,10,36 | 0,0,0 | 8,45,20 | 3,7,2 | 300 | 150 | 6.70E-07 | 1.92E-05 | 1.0,1.0,1.0 | 0.571,0.763,0.83 | 0.278 | ENSG000001427 | [[[[27366258, 273 | 3 |
| ENSG000001427 | MAP3K6 | chr1 | 27362850 | 27363021 | 27361704 | 27361867 | 27363441 | 27363548 | 6,13,15 | 0,0,0 | 0,26,9 | 3,7,2 | 300 | 150 | 8.92E-08 | 3.17E-06 | 1.0,1.0,1.0 | 0.0,0.65,0.692 | 0.553 | ENSG000001427 | [[[[27366258, 273 | 3 |
| ENSG000001261 | HECTD3 | chr1 | 45006950 | 45007015 | 45006691 | 45006795 | 45007412 | 45007595 | 19,20,33 | 0,0,0 | 25,24,25 | 2,5,1 | 215 | 150 | 0.000327715 | 0.004457956 | 1.0,1.0,1.0 | 0.897,0.77,0.946 | 0.129 | ENSG000001261 | [[[[45010889, 450 | 2 |
| ENSG000001762 | ZBTB8OS | chr1 | 32633644 | 32633727 | 32620819 | 32621948 | 32650432 | 32650525 | 6,2,0 | 4,1,3 | 0,0,2 | 5,17,6 | 233 | 150 | 0.000638316 | 0.007915302 | 0.491,0.563,0.0 | 0.0,0.0,0.177 | 0.292 | ENSG000001762 | [[[[32650433, 326 | 5 |
| ENSG000001762 | ZBTB8OS | chr1 | 32633950 | 32634072 | 32627507 | 32627544 | 32650401 | 32650525 | 1,1,3 | 1,0,0 | 4,4,2 | 3,3,2 | 272 | 150 | 0.000104434 | 0.001655428 | 0.355,1.0,1.0 | 0.424,0.424,0.35 | 0.384 | ENSG000001762 | [[[[32650433, 326 | 4 |
| ENSG000001762 | ZBTB8OS | chr1 | 32634767 | 32634792 | 32627507 | 32627544 | 32650401 | 32650525 | 2,2,2 | 1,0,0 | 1,4,0 | 3,3,2 | 175 | 150 | 4.30E-06 | 0.000101179 | 0.632,1.0,1.0 | 0.222,0.533,0.0 | 0.626 | ENSG000001762 | [[[[32650433, 326 | 4 |
| ENSG000001762 | ZBTB8OS | chr1 | 32634767 | 32634792 | 32627507 | 32627544 | 32650432 | 32650535 | 26,14,15 | 9,3,1 | 16,24,17 | 12,20,12 | 175 | 150 | 0.000766737 | 0.009248672 | 0.712,0.8,0.928 | 0.533,0.507,0.54 | 0.284 | ENSG000001762 | [[[[32650433, 326 | 4 |
| ENSG000001072 | RAPGEF1 | chr9 | 131638634 | 131638791 | 131630235 | 131630324 | 131650809 | 131650949 | 18,20,75 | 0,0,0 | 14,46,44 | 4,2,3 | 300 | 150 | 3.25E-05 | 0.000599222 | 1.0,1.0,1.0 | 0.636,0.92,0.88 | 0.188 | ENSG000001072 | [[[[131739770, 13 | 3 |
| ENSG000001072 | RAPGEF1 | chr9 | 131643238 | 131643417 | 131630235 | 131630324 | 131650809 | 131650949 | 19,11,44 | 0,0,0 | 12,27,21 | 4,2,3 | 300 | 150 | 2.94E-06 | 7.22E-05 | 1.0,1.0,1.0 | 0.6,0.871,0.778 | 0.25 | ENSG000001072 | [[[[131739770, 13 | 3 |
| ENSG000001072 | RAPGEF1 | chr9 | 131650128 | 131650242 | 131630235 | 131630324 | 131650809 | 131650949 | 21,13,35 | 0,0,0 | 11,19,31 | 4,2,3 | 264 | 150 | 5.86E-06 | 0.000133441 | 1.0,1.0,1.0 | 0.61,0.844,0.854 | 0.231 | ENSG000001072 | [[[[131739770, 13 | 3 |
| ENSG000001427 | PLK4 | chr4 | 127885707 | 127886626 | 127883261 | 127883357 | 127887395 | 127887496 | 2,2,8 | 0,0,0 | 4,5,5 | 1,6,2 | 300 | 150 | 5.63E-09 | 2.65E-07 | 1.0,1.0,1.0 | 0.667,0.294,0.55 | 0.494 | ENSG000001427 | [[[[127880893, 12 | 2 |
| ENSG000001427 | PLK4 | chr4 | 127885707 | 127886728 | 127883261 | 127883357 | 127887395 | 127887496 | 13,17,27 | 0,0,0 | 4,16,20 | 1,6,2 | 300 | 150 | 7.76E-07 | 2.19E-05 | 1.0,1.0,1.0 | 0.667,0.571,0.83 | 0.31 | ENSG000001427 | [[[[127880893, 12 | 2 |
| ENSG000001683 | ARF4 | chr3 | 57583897 | 57584007 | 57577315 | 57577387 | 57597073 | 57597325 | 209,172,323 | 0,1,0 | 159,352,228 | 15,32,17 | 260 | 150 | 0 | 0 | 1.0,0.99,1.0 | 0.859,0.864,0.88 | 0.127 | ENSG000001683 | [[[[57597074, 575 | 2 |
| ENSG000001683 | ARF4 | chr3 | 57584383 | 57584464 | 57577315 | 57577387 | 57597073 | 57597198 | 272,240,448 | 0,1,0 | 231,439,312 | 15,32,17 | 231 | 150 | 1.11E-16 | 1.65E-14 | 1.0,0.994,1.0 | 0.909,0.899,0.92 | 0.088 | ENSG000001683 | [[[[57597074, 575 | 2 |
| ENSG000000853 | SCAMP1 | chr5 | 78449932 | 78450034 | 78416540 | 78416649 | 78459244 | 78459362 | 15,18,23 | 0,0,0 | 11,22,16 | 0,6,2 | 252 | 150 | 4.99E-05 | 0.000871763 | 1.0,1.0,1.0 | 1.0,0.686,0.826 | 0.163 | ENSG000000853 | [[[[78360617, 783 | 3 |
| ENSG000001383 | FASTKD1 | chr2 | 169538012 | 169538141 | 169537001 | 169537340 | 169544720 | 169544835 | 14,12,13 | 0,0,0 | 18,21,15 | 7,4,2 | 279 | 150 | 9.85E-08 | 3.46E-06 | 1.0,1.0,1.0 | 0.58,0.738,0.801 | 0.294 | ENSG000001383 | [[[[169573669, 16 | 2 |
| ENSG000001383 | FASTKD1 | chr2 | 169540050 | 169540179 | 169537226 | 169537340 | 169544720 | 169544835 | 25,16,26 | 0,0,0 | 15,22,16 | 7,4,2 | 279 | 150 | 3.62E-08 | 1.42E-06 | 1.0,1.0,1.0 | 0.535,0.747,0.81 | 0.302 | ENSG000001383 | [[[[169573669, 16 | 2 |
| ENSG000001383 | FASTKD1 | chr2 | 169560386 | 169560785 | 169557186 | 169557297 | 169569183 | 169569252 | 17,20,28 | 0,0,0 | 6,29,9 | 4,7,11 | 300 | 150 | 2.74E-11 | 1.95E-09 | 1.0,1.0,1.0 | 0.429,0.674,0.29 | 0.536 | ENSG000001383 | [[[[169573669, 16 | 2 |
| ENSG000001383 | FASTKD1 | chr2 | 169560386 | 169560785 | 169557186 | 169557297 | 169571652 | 169572171 | 17,20,26 | 0,2,0 | 6,30,13 | 1,11,4 | 300 | 150 | 1.68E-06 | 4.36E-05 | 1.0,0.833,1.0 | 0.75,0.577,0.619 | 0.296 | ENSG000001383 | [[[[169573669, 16 | 3 |
| ENSG000001383 | FASTKD1 | chr2 | 169563224 | 169563350 | 169557186 | 169557297 | 169569183 | 169569252 | 10,6,17 | 0,0,0 | 4,7,10 | 4,7,11 | 276 | 150 | 6.46E-13 | 6.02E-11 | 1.0,1.0,1.0 | 0.352,0.352,0.33 | 0.655 | ENSG000001383 | [[[[169573669, 16 | 2 |

| | | | | | | | | | | | | | | | | | | | | | |
|---|---|---|---|---|---|---|---|---|---|---|---|---|---|---|---|---|---|---|---|---|---|
| ENSG000001383 | FASTKD1 | chr2 | 169569183 | 169569252 | 169557186 | 169557297 | 169571652 | 169572171 | 14,8,12 | 0,2,0 | 7,23,21 | 1,11,4 | 219 | 150 | 0.000162571 | 0.002430607 | 1.0,0.733,1.0 | 0.827,0.589,0.78 | 0.178 | ENSG000001383 | [[[[169573669, 16 | 3 |
| ENSG000001584 | EIF5B | chr2 | 99394508 | 99394585 | 99394266 | 99394398 | 99398747 | 99398909 | 52,33,30 | 0,0,0 | 57,96,57 | 5,10,7 | 227 | 150 | 2.60E-07 | 8.24E-06 | 1.0,1.0,1.0 | 0.883,0.864,0.84 | 0.137 | ENSG000001584 | [[[[99337389, 993 | 3 |
| ENSG000001584 | EIF5B | chr2 | 99396759 | 99396898 | 99394266 | 99394398 | 99398747 | 99398909 | 86,70,88 | 0,0,0 | 71,139,71 | 5,10,7 | 289 | 150 | 2.18E-08 | 9.03E-07 | 1.0,1.0,1.0 | 0.881,0.878,0.84 | 0.134 | ENSG000001584 | [[[[99337389, 993 | 3 |
| ENSG000001370 | APTX | chr9 | 32984630 | 32984857 | 32974457 | 32974561 | 32988082 | 32988124 | 74,69,82 | 0,0,0 | 64,128,80 | 4,6,4 | 300 | 150 | 1.74E-05 | 0.000347933 | 1.0,1.0,1.0 | 0.889,0.914,0.90 | 0.096 | ENSG000001370 | [[[[33001567, 330 | 3 |
| ENSG000001370 | APTX | chr9 | 32984630 | 32984857 | 32974457 | 32974561 | 32989758 | 32989895 | 74,69,82 | 0,0,0 | 64,127,79 | 12,21,10 | 300 | 150 | 3.45E-14 | 3.84E-12 | 1.0,1.0,1.0 | 0.727,0.751,0.79 | 0.241 | ENSG000001370 | [[[[33001567, 330 | 4 |
| ENSG000001370 | APTX | chr9 | 32988082 | 32988124 | 32974457 | 32974561 | 32989758 | 32989895 | 15,11,21 | 0,0,0 | 14,31,25 | 12,21,10 | 192 | 150 | 2.12E-12 | 1.85E-10 | 1.0,1.0,1.0 | 0.477,0.536,0.66 | 0.442 | ENSG000001370 | [[[[33001567, 330 | 4 |
| ENSG000001370 | APTX | chr9 | 32988082 | 32988129 | 32974457 | 32974561 | 32989758 | 32989895 | 42,23,43 | 0,0,0 | 62,93,41 | 12,21,10 | 197 | 150 | 1.67E-11 | 1.23E-09 | 1.0,1.0,1.0 | 0.797,0.771,0.75 | 0.225 | ENSG000001370 | [[[[33001567, 330 | 4 |
| ENSG000001631 | RNF149 | chr2 | 101286080 | 101286177 | 101281858 | 101282057 | 101294930 | 101295181 | 37,24,39 | 0,0,1 | 61,137,65 | 2,13,12 | 247 | 150 | 4.12E-06 | 9.74E-05 | 1.0,1.0,0.959 | 0.949,0.865,0.76 | 0.126 | ENSG000001631 | [[[[101308129, 10 | 3 |
| ENSG000001615 | FDXR | chr17 | 74863895 | 74864347 | 74863075 | 74863246 | 74864479 | 74864564 | 38,65,88 | 0,0,0 | 38,76,49 | 3,3,5 | 300 | 150 | 2.22E-05 | 0.000431795 | 1.0,1.0,1.0 | 0.864,0.927,0.83 | 0.126 | ENSG000001615 | [[[[74872866, 748 | 2 |
| ENSG000001615 | FDXR | chr17 | 74864147 | 74864347 | 74863075 | 74863246 | 74864479 | 74864564 | 3,20,37 | 0,0,0 | 11,22,7 | 3,3,5 | 300 | 150 | 2.88E-08 | 1.16E-06 | 1.0,1.0,1.0 | 0.647,0.786,0.41 | 0.385 | ENSG000001615 | [[[[74872866, 748 | 2 |
| ENSG000001329 | ZMYM5 | chr13 | 19837655 | 19837821 | 19835476 | 19835689 | 19851354 | 19852190 | 10,7,12 | 0,0,2 | 18,33,25 | 8,15,3 | 300 | 150 | 9.71E-05 | 0.001557703 | 1.0,1.0,0.75 | 0.529,0.524,0.80 | 0.297 | ENSG000001329 | [[[[19863450, 198 | 2 |
| ENSG000001329 | ZMYM5 | chr13 | 19838699 | 19838985 | 19835476 | 19835689 | 19851354 | 19851448 | 4,11,18 | 0,0,2 | 15,46,25 | 8,15,3 | 300 | 150 | 6.61E-05 | 0.00111327 | 1.0,1.0,0.818 | 0.484,0.605,0.80 | 0.308 | ENSG000001329 | [[[[19863450, 198 | 2 |
| ENSG000001267 | DLGAP5 | chr14 | 55170701 | 55170787 | 55154616 | 55154806 | 55175345 | 55175472 | 40,28,60 | 0,0,0 | 7,33,28 | 1,3,3 | 236 | 150 | 0.000658559 | 0.008125146 | 1.0,1.0,1.0 | 0.816,0.875,0.85 | 0.151 | ENSG000001267 | [[[[55191463, 551 | 4 |
| ENSG000001103 | EIF4G2 | chr11 | 10799212 | 10799424 | 10798991 | 10799113 | 10800089 | 10800348 | 11,118,161,309 | 0,0,1 | 5,951,468,841 | 30,75,37 | 300 | 150 | 0 | 0 | 1.0,1.0,0.998 | 0.908,0.907,0.91 | 0.088 | ENSG000001103 | [[[[10808705, 108 | 2 |
| ENSG000001103 | EIF4G2 | chr11 | 10799551 | 10799756 | 10798991 | 10799113 | 10800089 | 10800348 | 647,528,943 | 0,0,1 | 281,691,394 | 30,75,37 | 300 | 150 | 0 | 0 | 1.0,1.0,0.998 | 0.824,0.822,0.84 | 0.17 | ENSG000001103 | [[[[10808705, 108 | 2 |
| ENSG000001156 | PPP1R7 | chr2 | 241159212 | 241159343 | 241158465 | 241158549 | 241163284 | 241163401 | 146,139,200 | 1,0,0 | 63,114,93 | 2,15,5 | 281 | 150 | 1.91E-09 | 1.00E-07 | 0.987,1.0,1.0 | 0.944,0.802,0.90 | 0.111 | ENSG000001156 | [[[[241150467, 24 | 2 |
| ENSG000001156 | PPP1R7 | chr2 | 241160331 | 241160494 | 241158483 | 241158549 | 241163284 | 241163401 | 161,134,184 | 1,0,0 | 84,205,101 | 2,15,5 | 300 | 150 | 4.57E-08 | 1.74E-06 | 0.988,1.0,1.0 | 0.955,0.872,0.91 | 0.084 | ENSG000001156 | [[[[241150467, 24 | 2 |
| ENSG000000866 | ZFAND6 | chr15 | 80121711 | 80121820 | 80120327 | 80120498 | 80131179 | 80131233 | 110,95,161 | 0,0,0 | 110,206,146 | 13,24,9 | 259 | 150 | 2.01E-13 | 2.02E-11 | 1.0,1.0,1.0 | 0.831,0.833,0.90 | 0.144 | ENSG000000866 | [[[[80059676, 800 | 2 |
| ENSG000001064 | MEST | chr7 | 130503932 | 130503996 | 130500461 | 130500532 | 130504938 | 130505298 | 91,80,157 | 0,0,0 | 105,151,82 | 0,13,6 | 214 | 150 | 2.22E-06 | 5.61E-05 | 1.0,1.0,1.0 | 1.0,0.891,0.905 | 0.068 | ENSG000001064 | [[[[130492085, 13 | 3 |
| ENSG000001630 | CCDC138 | chr2 | 108788851 | 108788966 | 108786787 | 108786915 | 108791674 | 108791802 | 7,11,7 | 0,0,0 | 3,5,3 | 5,20,5 | 265 | 150 | 1.44E-15 | 1.91E-13 | 1.0,1.0,1.0 | 0.254,0.124,0.25 | 0.789 | ENSG000001630 | [[[[108786771, 10 | 2 |
| ENSG000001630 | CCDC138 | chr2 | 108788851 | 108788966 | 108786787 | 108786915 | 108791679 | 108791802 | 3,1,3 | 0,0,0 | 1,2,0 | 2,6,4 | 265 | 150 | 3.33E-16 | 4.71E-14 | 1.0,1.0,1.0 | 0.221,0.159,0.0 | 0.873 | ENSG000001630 | [[[[108786771, 10 | 2 |
| ENSG000001630 | CCDC138 | chr2 | 108846737 | 108846930 | 108815940 | 108816105 | 108873450 | 108873589 | 3,1,3 | 0,0,0 | 0,1,6 | 1,2,2 | 300 | 150 | 1.50E-05 | 0.000304457 | 1.0,1.0,1.0 | 0.0,0.2,0.6 | 0.733 | ENSG000001630 | [[[[108786771, 10 | 3 |
| ENSG000000048 | CDC27 | chr17 | 47154671 | 47154786 | 47151805 | 47151918 | 47157229 | 47157384 | 39,33,61 | 0,0,0 | 27,93,54 | 0,4,6 | 265 | 150 | 0.000248286 | 0.003501548 | 1.0,1.0,1.0 | 1.0,0.929,0.836 | 0.078 | ENSG000000048 | [[[[47189146, 471 | 2 |
| ENSG000000048 | CDC27 | chr17 | 47156912 | 47156983 | 47154697 | 47154786 | 47181561 | 47181605 | 36,31,61 | 0,0,0 | 12,40,20 | 4,4,6 | 221 | 150 | 2.53E-08 | 1.03E-06 | 1.0,1.0,1.0 | 0.671,0.872,0.69 | 0.255 | ENSG000000048 | [[[[47189146, 471 | 5 |
| ENSG000000048 | CDC27 | chr17 | 47156912 | 47157124 | 47151805 | 47151918 | 47157229 | 47157267 | 34,28,77 | 0,0,0 | 13,26,31 | 0,4,6 | 300 | 150 | 2.21E-06 | 5.58E-05 | 1.0,1.0,1.0 | 1.0,0.765,0.721 | 0.171 | ENSG000000048 | [[[[47189146, 471 | 2 |
| ENSG000000048 | CDC27 | chr17 | 47156912 | 47157124 | 47154671 | 47154786 | 47169916 | 47170042 | 58,45,99 | 0,0,0 | 28,71,33 | 6,7,6 | 300 | 150 | 4.42E-10 | 2.61E-08 | 1.0,1.0,1.0 | 0.7,0.835,0.733 | 0.244 | ENSG000000048 | [[[[47189146, 471 | 3 |
| ENSG000000048 | CDC27 | chr17 | 47158205 | 47158303 | 47154671 | 47154786 | 47169916 | 47169977 | 36,20,51 | 0,0,0 | 7,25,20 | 6,7,6 | 248 | 150 | 3.08E-11 | 2.17E-09 | 1.0,1.0,1.0 | 0.414,0.684,0.66 | 0.411 | ENSG000000048 | [[[[47189146, 471 | 3 |
| ENSG000000048 | CDC27 | chr17 | 47169916 | 47170042 | 47158205 | 47158303 | 47181561 | 47181605 | 44,27,87 | 0,3,3 | 16,39,32 | 3,17,8 | 276 | 150 | 5.97E-07 | 1.73E-05 | 1.0,0.83,0.94 | 0.743,0.555,0.68 | 0.262 | ENSG000000048 | [[[[47189146, 471 | 2 |
| ENSG000000048 | CDC27 | chr17 | 47171916 | 47172064 | 47154671 | 47154786 | 47181561 | 47181637 | 58,41,96 | 0,0,0 | 48,74,48 | 4,4,6 | 298 | 150 | 6.58E-07 | 1.89E-05 | 1.0,1.0,1.0 | 0.858,0.903,0.80 | 0.146 | ENSG000000048 | [[[[47189146, 471 | 5 |
| ENSG000001256 | POLR1B | chr2 | 112559292 | 112559574 | 112552644 | 112552816 | 112564365 | 112564499 | 13,13,25 | 0,0,0 | 26,28,40 | 3,4,4 | 300 | 150 | 8.48E-06 | 0.000184624 | 1.0,1.0,1.0 | 0.813,0.778,0.83 | 0.192 | ENSG000001256 | [[[[112542465, 11 | 2 |
| ENSG000001353 | MAP3K7 | chr6 | 90552048 | 90552179 | 90550467 | 90550549 | 90556499 | 90556624 | 29,10,24 | 0,0,0 | 8,36,34 | 1,3,3 | 281 | 150 | 0.000550771 | 0.006989943 | 1.0,1.0,1.0 | 0.81,0.865,0.858 | 0.156 | ENSG000001353 | [[[[90586764, 905 | 2 |
| ENSG000001353 | MAP3K7 | chr6 | 90553457 | 90553586 | 90552048 | 90552179 | 90561621 | 90561667 | 15,10,48 | 0,0,0 | 12,27,33 | 3,6,3 | 279 | 150 | 3.86E-07 | 1.17E-05 | 1.0,1.0,1.0 | 0.683,0.708,0.85 | 0.251 | ENSG000001353 | [[[[90586764, 905 | 3 |
| ENSG000002065 | ANKRD28 | chr3 | 15735409 | 15735497 | 15724381 | 15724524 | 15751749 | 15751820 | 9,8,14 | 0,0,0 | 7,18,13 | 1,0,6 | 238 | 150 | 0.000456731 | 0.005954144 | 1.0,1.0,1.0 | 0.815,1.0,0.577 | 0.203 | ENSG000002065 | [[[[15796405, 157 | 2 |
| ENSG000001392 | SCAF11 | chr12 | 45945248 | 45945313 | 45934446 | 45934505 | 45961699 | 45961857 | 14,3,14 | 0,0,0 | 5,23,11 | 0,4,4 | 215 | 150 | 3.53E-05 | 0.000646257 | 1.0,1.0,1.0 | 1.0,0.8,0.657 | 0.181 | ENSG000001392 | [[[[45990353, 459 | 3 |
| ENSG000001450 | RUBCN | chr3 | 197696953 | 197697049 | 197693714 | 197693816 | 197700612 | 197701146 | 18,4,22 | 0,0,0 | 19,37,20 | 5,3,6 | 246 | 150 | 5.85E-07 | 1.70E-05 | 1.0,1.0,1.0 | 0.699,0.883,0.67 | 0.249 | ENSG000001450 | [[[[197736655, 19 | 3 |
| ENSG000000067 | VPS41 | chr7 | 38754701 | 38754752 | 38752175 | 38752313 | 38763454 | 38763547 | 12,23,41 | 0,0,0 | 33,40,24 | 7,7,8 | 201 | 150 | 6.01E-09 | 2.81E-07 | 1.0,1.0,1.0 | 0.779,0.81,0.691 | 0.24 | ENSG000000067 | [[[[38909154, 389 | 4 |
| ENSG000001672 | HDHD2 | chr18 | 47134495 | 47134704 | 47115315 | 47115348 | 47136338 | 47136449 | 8,6,17 | 0,0,0 | 6,20,12 | 3,1,5 | 300 | 150 | 1.17E-06 | 3.16E-05 | 1.0,1.0,1.0 | 0.5,0.909,0.545 | 0.349 | ENSG000001672 | [[[[47150378, 471 | 2 |
| ENSG000001672 | HDHD2 | chr18 | 47136338 | 47136449 | 47115130 | 47115348 | 47150377 | 47150498 | 4,4,19 | 0,0,0 | 6,9,13 | 2,7,5 | 261 | 150 | 1.47E-09 | 7.89E-08 | 1.0,1.0,1.0 | 0.633,0.425,0.59 | 0.448 | ENSG000001672 | [[[[47150378, 471 | 3 |
| ENSG000001478 | UHRF2 | chr9 | 6433913 | 6434173 | 6413150 | 6413643 | 6460572 | 6460581 | 21,14,25 | 0,0,1 | 11,46,27 | 0,4,4 | 300 | 150 | 0.000577761 | 0.007267121 | 1.0,1.0,0.926 | 1.0,0.852,0.771 | 0.101 | ENSG000001478 | [[[[6413199, 6413 | 2 |
| ENSG000001491 | EIF3M | chr11 | 32587011 | 32587144 | 32583797 | 32583929 | 32593865 | 32593949 | 341,343,512 | 3,0,6 | 401,933,588 | 29,49,27 | 283 | 150 | 1.06E-13 | 1.10E-11 | 0.984,1.0,0.978 | 0.88,0.91,0.92 | 0.084 | ENSG000001491 | [[[[32583831, 325 | 4 |
| ENSG000001491 | EIF3M | chr11 | 32588593 | 32588732 | 32583797 | 32583929 | 32589546 | 32589641 | 4,6,8 | 24,25,34 | 21,28,16 | 16,38,35 | 289 | 150 | 0.000108061 | 0.001706534 | 0.08,0.111,0.109 | 0.405,0.277,0.19 | -0.191 | ENSG000001491 | [[[[32583831, 325 | 3 |
| ENSG000001491 | EIF3M | chr11 | 32589546 | 32589641 | 32583831 | 32583929 | 32593865 | 32593949 | 381,332,503 | 3,0,6 | 351,677,424 | 29,49,27 | 245 | 150 | 0 | 0 | 0.987,1.0,0.981 | 0.881,0.894,0.90 | 0.096 | ENSG000001491 | [[[[32583831, 325 | 4 |
| ENSG000001491 | EIF3M | chr11 | 32589546 | 32589641 | 32588593 | 32588732 | 32593865 | 32593949 | 362,309,469 | 0,0,0 | 345,659,393 | 62,103,59 | 245 | 150 | 0 | 0 | 1.0,1.0,1.0 | 0.773,0.797,0.80 | 0.209 | ENSG000001491 | [[[[32583831, 325 | 2 |
| ENSG000001491 | EIF3M | chr11 | 32595965 | 32596047 | 32593768 | 32593949 | 32600688 | 32600793 | 172,167,214 | 1,0,1 | 142,314,184 | 3,12,7 | 232 | 150 | 3.22E-05 | 0.000595755 | 0.991,1.0,0.993 | 0.968,0.944,0.94 | 0.043 | ENSG000001491 | [[[[32583831, 325 | 2 |
| ENSG000001491 | EIF3M | chr11 | 32600688 | 32600832 | 32593768 | 32593949 | 32601761 | 32601822 | 482,433,602 | 0,0,0 | 521,953,585 | 37,47,20 | 294 | 150 | 0 | 0 | 1.0,1.0,1.0 | 0.878,0.912,0.93 | 0.091 | ENSG000001491 | [[[[32583831, 325 | 3 |
| ENSG000001442 | AMMECR1L | chr2 | 127869453 | 127869544 | 127866899 | 127866996 | 127871248 | 127871359 | 8,3,11 | 0,0,0 | 7,21,17 | 4,4,3 | 241 | 150 | 5.69E-07 | 1.66E-05 | 1.0,1.0,1.0 | 0.521,0.766,0.77 | 0.311 | ENSG000001442 | [[[[127885810, 12 | 2 |
| ENSG000001442 | AMMECR1L | chr2 | 127870813 | 127870928 | 127866899 | 127866996 | 127871248 | 127871359 | 11,5,13 | 0,0,0 | 9,43,23 | 4,4,3 | 265 | 150 | 4.98E-06 | 0.000115432 | 1.0,1.0,1.0 | 0.56,0.859,0.813 | 0.256 | ENSG000001442 | [[[[127885810, 12 | 2 |
| ENSG000000745 | BCS1L | chr2 | 218661405 | 218661545 | 218660036 | 218660166 | 218661758 | 218661916 | 36,27,44 | 0,0,0 | 48,75,27 | 4,8,10 | 290 | 150 | 3.30E-08 | 1.31E-06 | 1.0,1.0,1.0 | 0.861,0.829,0.58 | 0.242 | ENSG000000745 | [[[[218659683, 21 | 2 |
| ENSG000001321 | LRRC41 | chr1 | 46279491 | 46279614 | 46279181 | 46279257 | 46280395 | 46280560 | 60,101,129 | 0,0,0 | 75,126,80 | 5,8,2 | 273 | 150 | 6.38E-06 | 0.000144106 | 1.0,1.0,1.0 | 0.892,0.896,0.95 | 0.085 | ENSG000001321 | [[[[46303124, 463 | 2 |
| ENSG000001321 | LRRC41 | chr1 | 46279491 | 46279614 | 46279181 | 46279257 | 46281124 | 46281161 | 60,103,129 | 0,0,0 | 72,128,80 | 15,19,12 | 273 | 150 | 2.22E-16 | 3.17E-14 | 1.0,1.0,1.0 | 0.725,0.787,0.78 | 0.234 | ENSG000001321 | [[[[46303124, 463 | 3 |
| ENSG000001321 | LRRC41 | chr1 | 46280191 | 46280290 | 46279181 | 46279257 | 46280395 | 46280560 | 70,89,104 | 0,0,0 | 110,152,110 | 5,8,2 | 249 | 150 | 0.000101286 | 0.001613589 | 1.0,1.0,1.0 | 0.93,0.92,0.971 | 0.06 | ENSG000001321 | [[[[46303124, 463 | 2 |
| ENSG000001321 | LRRC41 | chr1 | 46280395 | 46280560 | 46279181 | 46279257 | 46281124 | 46281161 | 111,128,203 | 0,0,0 | 163,321,172 | 15,19,12 | 300 | 150 | 3.17E-12 | 2.68E-10 | 1.0,1.0,1.0 | 0.845,0.894,0.87 | 0.128 | ENSG000001321 | [[[[46303124, 463 | 3 |
| ENSG000001564 | SH3RF2 | chr5 | 146047771 | 146047863 | 146013746 | 146014061 | 146055980 | 146056213 | 21,2,10 | 0,0,0 | 28,46,22 | 1,6,5 | 242 | 150 | 1.55E-05 | 0.000312584 | 1.0,1.0,1.0 | 0.946,0.826,0.73 | 0.165 | ENSG000001564 | [[[[145936578, 14 | 2 |
| ENSG000001629 | WDR26 | chr1 | 224404429 | 224404570 | 224400949 | 224401069 | 224419517 | 224419615 | 106,76,178 | 0,0,0 | 77,177,120 | 7,13,8 | 291 | 150 | 4.17E-10 | 2.47E-08 | 1.0,1.0,1.0 | 0.85,0.875,0.885 | 0.13 | ENSG000001629 | [[[[224433684, 22 | 3 |
| ENSG000001629 | WDR26 | chr1 | 224418259 | 224418416 | 224400949 | 224401069 | 224419517 | 224419615 | 69,63,146 | 0,0,0 | 71,125,127 | 7,13,8 | 300 | 150 | 5.07E-10 | 2.97E-08 | 1.0,1.0,1.0 | 0.835,0.828,0.88 | 0.15 | ENSG000001629 | [[[[224433684, 22 | 3 |
| ENSG000001043 | NBN | chr8 | 89980733 | 89980852 | 89978277 | 89978323 | 89982721 | 89982855 | 25,14,18 | 0,0,0 | 22,58,34 | 6,3,0 | 269 | 150 | 0.000202504 | 0.002943608 | 1.0,1.0,1.0 | 0.672,0.915,1.0 | 0.138 | ENSG000001043 | [[[[89984525, 899 | 2 |
| ENSG000000822 | CCNT2 | chr2 | 134942611 | 134942674 | 134939001 | 134939062 | 134947735 | 134947899 | 1,3,8 | 0,0,0 | 6,10,15 | 0,4,4 | 213 | 150 | 1.67E-05 | 0.000335312 | 1.0,1.0,1.0 | 1.0,0.638,0.725 | 0.212 | ENSG000000822 | [[[[134918822, 13 | 2 |
| ENSG000001697 | UGP2 | chr2 | 63856305 | 63856433 | 63840939 | 63841232 | 63882465 | 63882503 | 59,40,96 | 5,3,2 | 32,64,54 | 9,16,11 | 278 | 150 | 3.96E-06 | 9.39E-05 | 0.864,0.878,0.96 | 0.657,0.683,0.72 | 0.213 | ENSG000001697 | [[[[63841896, 638 | 3 |
| ENSG000001697 | UGP2 | chr2 | 63857828 | 63857936 | 63840939 | 63841232 | 63882465 | 63882503 | 84,78,106 | 5,3,2 | 59,116,93 | 9,16,11 | 258 | 150 | 5.97E-05 | 0.001019884 | 0.907,0.938,0.96 | 0.792,0.808,0.83 | 0.128 | ENSG000001697 | [[[[63841896, 638 | 3 |
| ENSG000001396 | ESD | chr13 | 46777455 | 46777623 | 46771255 | 46771496 | 46781495 | 46781615 | 262,179,317 | 0,0,0 | 233,441,241 | 16,45,14 | 300 | 150 | 0 | 0 | 1.0,1.0,1.0 | 0.879,0.831,0.89 | 0.131 | ENSG000001396 | [[[[46797105, 467 | 2 |
| ENSG000000093 | CSDE1 | chr1 | 114726210 | 114726386 | 114725220 | 114725333 | 114730257 | 114730422 | 471,415,646 | 0,0,0 | 187,454,266 | 3,8,8 | 300 | 150 | 5.65E-05 | 0.000970644 | 1.0,1.0,1.0 | 0.969,0.966,0.94 | 0.041 | ENSG000000093 | [[[[114757925, 11 | 2 |
| ENSG000001700 | CNTROB | chr17 | 7948486 | 7948619 | 7947570 | 7947722 | 7949081 | 7949157 | 12,8,15 | 0,0,0 | 17,53,20 | 6,4,8 | 283 | 150 | 2.81E-07 | 8.85E-06 | 1.0,1.0,1.0 | 0.6,0.875,0.57 | 0.318 | ENSG000001700 | [[[[7932206, 7933 | 3 |
| ENSG000001700 | CNTROB | chr17 | 7948486 | 7948619 | 7947570 | 7947722 | 7949084 | 7949137 | 20,38,32 | 0,0,0 | 28,60,20 | 4,5,0 | 283 | 150 | 8.45E-05 | 0.001373972 | 1.0,1.0,1.0 | 0.788,0.864,1.0 | 0.116 | ENSG000001700 | [[[[7932206, 7933 | 3 |
| ENSG000001340 | MRPS36 | chr5 | 69226337 | 69226401 | 69217759 | 69217871 | 69229207 | 69229875 | 14,21,37 | 0,0,0 | 31,18,15 | 8,7,7 | 214 | 150 | 1.42E-10 | 9.08E-09 | 1.0,1.0,1.0 | 0.731,0.643,0.6 | 0.342 | ENSG000001340 | [[[[69217760, 692 | 2 |
| ENSG000001340 | MRPS36 | chr5 | 69228199 | 69228387 | 69217785 | 69217871 | 69229207 | 69229231 | 91,76,90 | 0,0,0 | 55,145,51 | 8,7,7 | 300 | 150 | 4.75E-09 | 2.28E-07 | 1.0,1.0,1.0 | 0.775,0.912,0.78 | 0.176 | ENSG000001340 | [[[[69217760, 692 | 2 |
| ENSG000001384 | IDH1 | chr2 | 208243426 | 208243604 | 208239862 | 208240003 | 208245318 | 208245424 | 174,120,227 | 0,0,0 | 150,328,221 | 6,19,4 | 300 | 150 | 3.64E-08 | 1.43E-06 | 1.0,1.0,1.0 | 0.926,0.896,0.96 | 0.071 | ENSG000001384 | [[[[208254939, 20 | 2 |
| ENSG000001248 | RUNX2 | chr6 | 45491940 | 45492114 | 45437946 | 45438051 | 45546826 | 45547209 | 9,5,24 | 0,0,0 | 7,13,16 | 0,3,3 | 300 | 150 | 0.000836781 | 0.009984499 | 1.0,1.0,1.0 | 1.0,0.684,0.727 | 0.196 | ENSG000001248 | [[[[45328330, 453 | 3 |
| ENSG000001658 | NSMF | chr9 | 137455238 | 137455307 | 137454390 | 137454443 | 137456410 | 137456486 | 31,54,99 | 2,0,15 | 72,124,81 | 9,42,27 | 219 | 150 | 0.000348803 | 0.0046991 | 0.914,1.0,0.819 | 0.846,0.669,0.67 | 0.182 | ENSG000001658 | [[[[137459032, 13 | 2 |
| ENSG000000612 | HDAC7 | chr12 | 47786584 | 47786703 | 47785450 | 47785885 | 47788044 | 47788164 | 33,44,87 | 0,0,0 | 100,203,87 | 4,13,11 | 269 | 150 | 2.97E-07 | 9.33E-06 | 1.0,1.0,1.0 | 0.933,0.897,0.81 | 0.118 | ENSG000000612 | [[[[47819767, 478 | 2 |
| ENSG000000612 | HDAC7 | chr12 | 47787711 | 47787809 | 47785751 | 47785885 | 47788044 | 47788164 | 26,38,47 | 0,0,0 | 21,60,20 | 4,13,11 | 248 | 150 | 2.96E-11 | 2.09E-09 | 1.0,1.0,1.0 | 0.761,0.736,0.52 | 0.326 | ENSG000000612 | [[[[47819767, 478 | 2 |
| ENSG000001578 | FMNL2 | chr2 | 152578887 | 152578964 | 152575135 | 152575244 | 152607338 | 152607413 | 18,11,42 | 0,0,0 | 4,39,34 | 3,1,11 | 227 | 150 | 3.02E-06 | 7.39E-05 | 1.0,1.0,1.0 | 0.468,0.963,0.67 | 0.299 | ENSG000001578 | [[[[152335174, 15 | 2 |
| ENSG000000852 | AK6 | chr5 | 69355894 | 69355953 | 69350983 | 69352253 | 69366502 | 69366595 | 46,60,94 | 0,0,0 | 45,67,43 | 37,49,25 | 209 | 150 | 0 | 0 | 1.0,1.0,1.0 | 0.466,0.495,0.55 | 0.496 | ENSG000000852 | [[[[69369463, 693 | 2 |
| ENSG000001151 | ACVR1 | chr2 | 157760879 | 157761077 | 157738439 | 157738570 | 157774087 | 157774187 | 53,25,81 | 0,0,0 | 48,106,55 | 1,8,4 | 300 | 150 | 1.26E-05 | 0.000261938 | 1.0,1.0,1.0 | 0.96,0.869,0.873 | 0.099 | ENSG000001151 | [[[[157875796, 15 | 3 |
| ENSG000001151 | ACVR1 | chr2 | 157770367 | 157770514 | 157738439 | 157738570 | 157774087 | 157774187 | 18,35,59 | 0,0,0 | 12,26,16 | 1,8,4 | 297 | 150 | 8.51E-09 | 3.83E-07 | 1.0,1.0,1.0 | 0.858,0.621,0.66 | 0.284 | ENSG000001151 | [[[[157875796, 15 | 3 |
| ENSG000002728 | DCP1A | chr3 | 53319406 | 53319473 | 53312240 | 53312379 | 53344901 | 53344936 | 1,1,7 | 0,0,0 | 14,35,26 | 0,5,3 | 217 | 150 | 0.000460761 | 0.005999498 | 1.0,1.0,1.0 | 1.0,0.829,0.857 | 0.105 | ENSG000002728 | [[[[53347383, 533 | 2 |
| ENSG000002728 | DCP1A | chr3 | 53342143 | 53342271 | 53312240 | 53312379 | 53344901 | 53344936 | 8,4,9 | 0,0,0 | 14,40,40 | 0,5,3 | 278 | 150 | 0.000378731 | 0.005049636 | 1.0,1.0,1.0 | 1.0,0.812,0.878 | 0.103 | ENSG000002728 | [[[[53347383, 533 | 2 |
| ENSG000001384 | WDR12 | chr2 | 202883608 | 202883741 | 202882710 | 202882783 | 202892616 | 202892702 | 50,58,74 | 0,0,0 | 40,104,47 | 7,6,6 | 283 | 150 | 3.77E-08 | 1.47E-06 | 1.0,1.0,1.0 | 0.752,0.902,0.80 | 0.18 | ENSG000001384 | [[[[202911436, 20 | 3 |
| ENSG000001384 | WDR12 | chr2 | 202884197 | 202884303 | 202883608 | 202883741 | 202892616 | 202892702 | 56,41,65 | 0,0,0 | 34,71,50 | 4,3,3 | 256 | 150 | 0.000155623 | 0.002338432 | 1.0,1.0,1.0 | 0.833,0.933,0.90 | 0.109 | ENSG000001384 | [[[[202911436, 20 | 2 |
| ENSG000000679 | ROCK1 | chr18 | 21045291 | 21045467 | 21044101 | 21044186 | 21049779 | 21049880 | 25,17,37 | 0,0,0 | 6,30,24 | 4,1,5 | 300 | 150 | 2.60E-06 | 6.45E-05 | 1.0,1.0,1.0 | 0.429,0.938,0.70 | 0.309 | ENSG000000679 | [[[[21110818, 211 | 2 |
| ENSG000001158 | GORASP2 | chr2 | 170956435 | 170956559 | 170951327 | 170951458 | 170961662 | 170961749 | 194,143,260 | 2,3,1 | 225,335,195 | 18,31,24 | 274 | 150 | 2.41E-13 | 2.37E-11 | 0.982,0.963,0.99 | 0.872,0.855,0.81 | 0.132 | ENSG000001158 | [[[[170929253, 17 | 2 |

| | | | | | | | | | | | | | | | | | | | | | | |
|---|---|---|---|---|---|---|---|---|---|---|---|---|---|---|---|---|---|---|---|---|---|---|
| ENSG00000138 | OLA1 | chr2 | 174123594 | 174123675 | 174081923 | 174082064 | 174141824 | 174142000 | 111,102,122 | 3,0,0 | 105,230,113 | 4,7,10 | 231 | 150 | 9.92E-05 | 0.001587229 | 0.96,1.0,1.0 | 0.945,0.955,0.88 | 0.06 | ENSG00000138 | [[[[174248452, 17 | 2 |
| ENSG00000138 | OLA1 | chr2 | 174223032 | 174223160 | 174141824 | 174142000 | 174246714 | 174246815 | 129,121,200 | 0,0,1 | 110,219,141 | 12,19,6 | 278 | 150 | 2.38E-11 | 1.71E-09 | 1.0,1.0,0.991 | 0.832,0.861,0.92 | 0.124 | ENSG00000138 | [[[[174248452, 17 | 2 |
| ENSG00000125 | IRF1 | chr5 | 132485666 | 132485716 | 132484361 | 132484497 | 132486556 | 132486686 | 6,10,8 | 0,0,0 | 15,36,27 | 4,16,7 | 200 | 150 | 9.08E-08 | 3.22E-06 | 1.0,1.0,1.0 | 0.738,0.628,0.74 | 0.297 | ENSG00000125 | [[[[132490545, 13 | 2 |
| ENSG00000125 | IRF1 | chr5 | 132486250 | 132486373 | 132484361 | 132484497 | 132486556 | 132486686 | 6,17,15 | 0,0,0 | 33,58,44 | 4,16,7 | 273 | 150 | 2.07E-07 | 6.71E-06 | 1.0,1.0,1.0 | 0.819,0.666,0.77 | 0.247 | ENSG00000125 | [[[[132490545, 13 | 2 |
| ENSG00000164 | TMEM161B | chr5 | 88220562 | 88220719 | 88207028 | 88207180 | 88240812 | 88240916 | 10,11,20 | 0,0,0 | 15,43,23 | 3,4,4 | 300 | 150 | 4.13E-06 | 9.77E-05 | 1.0,1.0,1.0 | 0.714,0.843,0.74 | 0.234 | ENSG00000164 | [[[[88268721, 882 | 3 |
| ENSG00000065 | WDR3 | chr1 | 117938479 | 117938558 | 117933287 | 117933389 | 117939476 | 117939572 | 23,26,41 | 0,0,0 | 15,55,47 | 3,9,5 | 229 | 150 | 1.89E-07 | 6.18E-06 | 1.0,1.0,1.0 | 0.766,0.8,0.86 | 0.191 | ENSG00000065 | [[[[117929739, 11 | 3 |
| ENSG00000065 | WDR3 | chr1 | 117939476 | 117939572 | 117933287 | 117933389 | 117940826 | 117940940 | 42,21,57 | 0,0,0 | 22,76,48 | 3,5,7 | 246 | 150 | 1.08E-06 | 2.94E-05 | 1.0,1.0,1.0 | 0.817,0.903,0.80 | 0.158 | ENSG00000065 | [[[[117929739, 11 | 4 |
| ENSG00000055 | PUM2 | chr2 | 20278582 | 20278819 | 20263192 | 20263460 | 20290651 | 20290790 | 28,25,79 | 0,0,0 | 36,60,74 | 6,7,10 | 300 | 150 | 3.34E-09 | 1.66E-07 | 1.0,1.0,1.0 | 0.75,0.811,0.787 | 0.217 | ENSG00000055 | [[[[20350597, 203 | 3 |
| ENSG00000055 | PUM2 | chr2 | 20278582 | 20278819 | 20263192 | 20263460 | 20297552 | 20297678 | 28,25,79 | 0,0,0 | 35,54,71 | 2,5,2 | 300 | 150 | 0.000429551 | 0.00563258 | 1.0,1.0,1.0 | 0.897,0.844,0.94 | 0.104 | ENSG00000055 | [[[[20350597, 203 | 5 |
| ENSG00000055 | PUM2 | chr2 | 20283342 | 20283486 | 20263192 | 20263460 | 20290651 | 20290790 | 46,31,61 | 0,0,0 | 25,44,32 | 6,7,10 | 294 | 150 | 1.39E-11 | 1.04E-09 | 1.0,1.0,1.0 | 0.68,0.762,0.62 | 0.313 | ENSG00000055 | [[[[20350597, 203 | 3 |
| ENSG00000023 | SNAPC1 | chr14 | 61767211 | 61767352 | 61762356 | 61762588 | 61768635 | 61768740 | 53,62,58 | 0,0,0 | 42,133,79 | 23,33,33 | 291 | 150 | 0 | 0 | 1.0,1.0,1.0 | 0.485,0.675,0.55 | 0.429 | ENSG00000023 | [[[[61762420, 617 | 2 |
| ENSG00000023 | SNAPC1 | chr14 | 61768635 | 61768740 | 61762356 | 61762588 | 61776094 | 61776253 | 39,40,44 | 0,0,0 | 94,169,131 | 0,9,5 | 255 | 150 | 0.000706381 | 0.008632685 | 1.0,1.0,1.0 | 1.0,0.917,0.939 | 0.048 | ENSG00000023 | [[[[61762420, 617 | 3 |
| ENSG00000188 | PLSCR1 | chr3 | 146517005 | 146517167 | 146515179 | 146516101 | 146525604 | 146525647 | 49,34,66 | 0,0,0 | 51,99,69 | 1,4,8 | 300 | 150 | 2.42E-05 | 0.000464256 | 1.0,1.0,1.0 | 0.962,0.925,0.81 | 0.1 | ENSG00000188 | [[[[146544467, 14 | 3 |
| ENSG00000188 | PLSCR1 | chr3 | 146521832 | 146522053 | 146515179 | 146516101 | 146525604 | 146525647 | 58,37,59 | 0,0,0 | 36,134,72 | 1,4,8 | 300 | 150 | 3.65E-05 | 0.000664127 | 1.0,1.0,1.0 | 0.947,0.944,0.81 | 0.097 | ENSG00000188 | [[[[146544467, 14 | 3 |
| ENSG00000064 | PMS1 | chr2 | 189843963 | 189844080 | 189795768 | 189795951 | 189852654 | 189852762 | 16,12,26 | 0,0,0 | 10,26,10 | 4,1,2 | 267 | 150 | 0.000135901 | 0.002076209 | 1.0,1.0,1.0 | 0.584,0.936,0.73 | 0.248 | ENSG00000064 | [[[[189784450, 18 | 3 |
| ENSG00000064 | PMS1 | chr2 | 189873495 | 189873656 | 189863742 | 189864228 | 189877271 | 189877436 | 32,27,26 | 0,0,0 | 26,21,8 | 4,1,2 | 300 | 150 | 0.000183983 | 0.002713726 | 1.0,1.0,1.0 | 0.765,0.913,0.66 | 0.218 | ENSG00000064 | [[[[189784450, 18 | 2 |
| ENSG00000053 | ANAPC4 | chr4 | 25416424 | 25416598 | 25415465 | 25415540 | 25418154 | 25418261 | 17,15,13 | 0,0,0 | 7,15,8 | 0,7,2 | 300 | 150 | 4.22E-06 | 9.94E-05 | 1.0,1.0,1.0 | 1.0,0.517,0.667 | 0.272 | ENSG00000053 | [[[[25377263, 253 | 2 |
| ENSG00000053 | ANAPC4 | chr4 | 25417615 | 25417739 | 25415465 | 25415540 | 25418154 | 25418261 | 35,27,39 | 0,0,0 | 21,56,18 | 0,7,2 | 274 | 150 | 6.31E-05 | 0.001070732 | 1.0,1.0,1.0 | 1.0,0.814,0.831 | 0.118 | ENSG00000053 | [[[[25377263, 253 | 2 |
| ENSG00000115 | SF3B1 | chr2 | 197398460 | 197398581 | 197389783 | 197392461 | 197398968 | 197399028 | 11,10,6 | 0,0,0 | 9,11,9 | 5,4,0 | 271 | 150 | 3.69E-06 | 8.84E-05 | 1.0,1.0,1.0 | 0.499,0.604,1.0 | 0.299 | ENSG00000115 | [[[[197434972, 19 | 4 |
| ENSG00000115 | SF3B1 | chr2 | 197398460 | 197398581 | 197389783 | 197392461 | 197400054 | 197400166 | 161,129,204 | 0,0,0 | 112,242,125 | 8,13,4 | 271 | 150 | 1.31E-08 | 5.69E-07 | 1.0,1.0,1.0 | 0.886,0.912,0.94 | 0.086 | ENSG00000115 | [[[[197434972, 19 | 4 |
| ENSG00000188 | UROS | chr10 | 125816436 | 125816525 | 125815033 | 125815130 | 125823028 | 125823039 | 2,5,28 | 0,0,0 | 4,3,4 | 8,6,8 | 239 | 150 | 1.33E-15 | 1.77E-13 | 1.0,1.0,1.0 | 0.239,0.239,0.23 | 0.761 | ENSG00000188 | [[[[125823029, 12 | 2 |
| ENSG00000121 | KIF18A | chr11 | 28084631 | 28084808 | 28083168 | 28083243 | 28090616 | 28090727 | 23,16,24 | 0,0,0 | 8,17,17 | 4,6,2 | 300 | 150 | 1.39E-08 | 6.02E-07 | 1.0,1.0,1.0 | 0.5,0.586,0.81 | 0.368 | ENSG00000121 | [[[[28108064, 281 | 2 |
| ENSG00000121 | CLCC1 | chr1 | 108943474 | 108943615 | 108941404 | 108941498 | 108947610 | 108947718 | 39,16,57 | 0,0,0 | 18,32,24 | 9,16,19 | 291 | 150 | 0 | 0 | 1.0,1.0,1.0 | 0.508,0.508,0.39 | 0.53 | ENSG00000121 | [[[[108963361, 10 | 2 |
| ENSG00000121 | CLCC1 | chr1 | 108943835 | 108944057 | 108939635 | 108939782 | 108947610 | 108947718 | 29,17,34 | 0,0,0 | 5,17,16 | 0,6,2 | 300 | 150 | 1.52E-05 | 0.000307332 | 1.0,1.0,1.0 | 1.0,0.586,0.8 | 0.205 | ENSG00000121 | [[[[108963361, 10 | 4 |
| ENSG00000121 | CLCC1 | chr1 | 108943835 | 108944057 | 108941404 | 108941498 | 108947610 | 108947718 | 29,17,34 | 0,0,0 | 5,19,16 | 9,16,19 | 300 | 150 | 0 | 0 | 1.0,1.0,1.0 | 0.217,0.373,0.29 | 0.705 | ENSG00000121 | [[[[108963361, 10 | 2 |
| ENSG00000121 | CLCC1 | chr1 | 108950308 | 108950448 | 108940044 | 108940142 | 108963360 | 108963457 | 7,11,20 | 0,0,0 | 3,35,24 | 2,2,3 | 290 | 150 | 0.000479612 | 0.006209939 | 1.0,1.0,1.0 | 0.437,0.901,0.80 | 0.286 | ENSG00000121 | [[[[108963361, 10 | 7 |
| ENSG00000121 | CLCC1 | chr1 | 108950308 | 108950448 | 108941404 | 108941498 | 108963360 | 108963457 | 7,11,20 | 0,0,0 | 8,35,27 | 0,6,4 | 290 | 150 | 9.58E-06 | 0.000205828 | 1.0,1.0,1.0 | 1.0,0.751,0.777 | 0.157 | ENSG00000121 | [[[[108963361, 10 | 6 |
| ENSG00000055 | NOP58 | chr2 | 202292776 | 202292903 | 202291124 | 202291270 | 202297378 | 202297513 | 145,134,153 | 0,0,0 | 229,451,223 | 16,58,27 | 277 | 150 | 0 | 0 | 1.0,1.0,1.0 | 0.886,0.808,0.81 | 0.163 | ENSG00000055 | [[[[202265763, 20 | 2 |
| ENSG00000055 | NOP58 | chr2 | 202295673 | 202295837 | 202291124 | 202291270 | 202297378 | 202297513 | 110,153,149 | 0,0,0 | 194,421,195 | 16,58,27 | 300 | 150 | 0 | 0 | 1.0,1.0,1.0 | 0.858,0.784,0.78 | 0.192 | ENSG00000055 | [[[[202265763, 20 | 2 |
| ENSG00000136 | NUPL2 | chr7 | 23187051 | 23187146 | 23182065 | 23182206 | 23195838 | 23195915 | 36,32,45 | 0,0,0 | 32,76,33 | 3,23,12 | 245 | 150 | 3.25E-11 | 2.28E-09 | 1.0,1.0,1.0 | 0.867,0.669,0.62 | 0.279 | ENSG00000136 | [[[[23182048, 231 | 2 |
| ENSG00000136 | NUPL2 | chr7 | 23187051 | 23187146 | 23182065 | 23182206 | 23196679 | 23196766 | 1,1,5 | 0,0,0 | 12,36,21 | 3,13,3 | 245 | 150 | 2.34E-06 | 5.88E-05 | 1.0,1.0,1.0 | 0.71,0.629,0.811 | 0.283 | ENSG00000136 | [[[[23182048, 231 | 3 |
| ENSG00000136 | NUPL2 | chr7 | 23195838 | 23195915 | 23181826 | 23182206 | 23196679 | 23196766 | 38,31,42 | 0,0,0 | 42,94,47 | 3,13,3 | 227 | 150 | 6.91E-07 | 1.98E-05 | 1.0,1.0,1.0 | 0.902,0.827,0.91 | 0.12 | ENSG00000136 | [[[[23182048, 231 | 3 |
| ENSG00000136 | NUPL2 | chr7 | 23196679 | 23196766 | 23181826 | 23182206 | 23199457 | 23199542 | 33,47,32 | 0,0,0 | 56,101,46 | 4,8,4 | 237 | 150 | 6.58E-06 | 0.000147947 | 1.0,1.0,1.0 | 0.899,0.889,0.87 | 0.111 | ENSG00000136 | [[[[23182048, 231 | 4 |
| ENSG00000136 | NUPL2 | chr7 | 23196679 | 23196766 | 23187051 | 23187146 | 23199457 | 23199542 | 33,48,35 | 0,0,0 | 59,113,53 | 4,5,5 | 237 | 150 | 5.88E-05 | 0.001004842 | 1.0,1.0,1.0 | 0.903,0.935,0.87 | 0.097 | ENSG00000136 | [[[[23182048, 231 | 2 |
| ENSG00000075 | DLG1 | chr3 | 197051576 | 197051668 | 197042559 | 197044729 | 197065707 | 197065809 | 93,107,161 | 0,0,0 | 86,178,95 | 3,15,10 | 242 | 150 | 8.22E-10 | 4.64E-08 | 1.0,1.0,1.0 | 0.947,0.88,0.855 | 0.106 | ENSG00000075 | [[[[197298536, 19 | 3 |
| ENSG00000075 | DLG1 | chr3 | 197065275 | 197065448 | 197051576 | 197051668 | 197065707 | 197065809 | 161,117,178 | 0,0,0 | 56,141,86 | 26,51,20 | 300 | 150 | 0 | 0 | 1.0,1.0,1.0 | 0.519,0.58,0.683 | 0.406 | ENSG00000075 | [[[[197298536, 19 | 2 |
| ENSG00000075 | DLG1 | chr3 | 197136541 | 197136678 | 197119409 | 197119530 | 197138221 | 197138391 | 103,99,223 | 0,0,0 | 79,254,164 | 6,3,5 | 287 | 150 | 0.00033697 | 0.004557328 | 1.0,1.0,1.0 | 0.873,0.978,0.94 | 0.068 | ENSG00000075 | [[[[197298536, 19 | 2 |
| ENSG00000136 | SLC41A2 | chr12 | 104889032 | 104889177 | 104802552 | 104805337 | 104895273 | 104895345 | 19,6,15 | 0,0,0 | 1,8,3 | 2,1,3 | 295 | 150 | 6.56E-06 | 0.000147601 | 1.0,1.0,1.0 | 0.203,0.803,0.33 | 0.552 | ENSG00000136 | [[[[104958088, 10 | 6 |
| ENSG00000009 | UBE3C | chr7 | 157223253 | 157223351 | 157216866 | 157216971 | 157231079 | 157231327 | 1,1,3 | 0,0,0 | 5,8,4 | 4,2,0 | 248 | 150 | 0.000334275 | 0.004530607 | 1.0,1.0,1.0 | 0.431,0.708,1.0 | 0.287 | ENSG00000009 | [[[[157138926, 15 | 3 |
| ENSG00000009 | UBE3C | chr7 | 157223253 | 157223351 | 157220688 | 157220776 | 157231079 | 157231327 | 36,60,98 | 2,0,0 | 47,83,54 | 0,11,10 | 248 | 150 | 2.14E-06 | 5.43E-05 | 0.916,1.0,1.0 | 1.0,0.82,0.766 | 0.11 | ENSG00000009 | [[[[157138926, 15 | 2 |
| ENSG00000009 | UBE3C | chr7 | 157225406 | 157225539 | 157220688 | 157220776 | 157231079 | 157231327 | 97,85,134 | 2,0,0 | 74,171,110 | 0,11,10 | 283 | 150 | 5.38E-06 | 0.000123787 | 0.963,1.0,1.0 | 1.0,0.892,0.854 | 0.072 | ENSG00000009 | [[[[157138926, 15 | 2 |
| ENSG00000009 | UBE3C | chr7 | 157254243 | 157254310 | 157253953 | 157254142 | 157267584 | 157269116 | 41,69,94 | 0,0,0 | 43,103,44 | 3,7,2 | 217 | 150 | 6.34E-05 | 0.001073736 | 1.0,1.0,1.0 | 0.908,0.91,0.938 | 0.081 | ENSG00000009 | [[[[157138926, 15 | 2 |
| ENSG00000107 | GLIS3 | chr9 | 4125733 | 4125941 | 4117767 | 4118881 | 4299420 | 4299916 | 15,10,36 | 3,0,4 | 27,58,52 | 0,1,0 | 300 | 150 | 0.000574188 | 0.00723053 | 0.714,1.0,0.818 | 1.0,0.967,1.0 | -0.145 | ENSG00000107 | [[[[4299421, 4300 | 2 |
| ENSG00000107 | GLIS3 | chr9 | 4286037 | 4286523 | 4118744 | 4118881 | 4299420 | 4299594 | 11,11,31 | 3,0,4 | 20,46,60 | 0,1,0 | 300 | 150 | 0.000441202 | 0.005771456 | 0.647,1.0,0.795 | 1.0,0.958,1.0 | -0.172 | ENSG00000107 | [[[[4299421, 4300 | 2 |
| ENSG00000198 | ATG9A | chr2 | 219227921 | 219228053 | 219225331 | 219225572 | 219229534 | 219229636 | 0,0,4 | 1,1,2 | 5,7,14 | 1,0,0 | 282 | 150 | 6.24E-05 | 0.001059783 | 0.0,0.0,0.515 | 0.727,1.0,1.0 | -0.737 | ENSG00000198 | [[[[219229535, 21 | 4 |
| ENSG00000123 | DNPEP | chr2 | 219385630 | 219385706 | 219384365 | 219384443 | 219385967 | 219386098 | 21,54,76 | 0,0,0 | 58,122,73 | 4,12,10 | 226 | 150 | 3.47E-08 | 1.37E-06 | 1.0,1.0,1.0 | 0.906,0.871,0.82 | 0.131 | ENSG00000123 | [[[[219387759, 21 | 2 |
| ENSG00000105 | TYK2 | chr19 | 10364613 | 10364771 | 10362548 | 10362657 | 10365516 | 10365863 | 7,24,47 | 0,0,1 | 5,32,20 | 4,1,4 | 300 | 150 | 2.67E-05 | 0.000504463 | 1.0,1.0,0.959 | 0.385,0.941,0.71 | 0.306 | ENSG00000105 | [[[[10380380, 103 | 2 |
| ENSG00000105 | TYK2 | chr19 | 10368054 | 10368202 | 10366416 | 10366580 | 10378213 | 10378426 | 43,63,84 | 0,0,0 | 31,84,51 | 4,5,7 | 298 | 150 | 9.30E-08 | 3.30E-06 | 1.0,1.0,1.0 | 0.796,0.894,0.78 | 0.175 | ENSG00000105 | [[[[10380380, 103 | 2 |
| ENSG00000105 | TYK2 | chr19 | 10368294 | 10368418 | 10366416 | 10366580 | 10378213 | 10378426 | 23,29,52 | 0,0,0 | 12,29,14 | 4,5,7 | 274 | 150 | 3.39E-10 | 2.05E-08 | 1.0,1.0,1.0 | 0.622,0.76,0.523 | 0.365 | ENSG00000105 | [[[[10380380, 103 | 2 |
| ENSG00000077 | UBE2T | chr1 | 202333449 | 202333555 | 202333009 | 202333093 | 202334988 | 202335058 | 113,118,189 | 0,0,0 | 131,237,150 | 9,6,2 | 256 | 150 | 6.53E-05 | 0.00110163 | 1.0,1.0,1.0 | 0.895,0.959,0.97 | 0.056 | ENSG00000077 | [[[[202341895, 20 | 2 |
| ENSG00000197 | HTT | chr4 | 3236148 | 3236254 | 3233162 | 3233353 | 3238446 | 3238609 | 48,39,58 | 0,0,0 | 30,43,22 | 8,8,4 | 256 | 150 | 7.93E-11 | 5.29E-09 | 1.0,1.0,1.0 | 0.687,0.759,0.76 | 0.264 | ENSG00000197 | [[[[3074681, 3075 | 3 |
| ENSG00000157 | RAB28 | chr4 | 13460698 | 13460828 | 13381490 | 13381594 | 13484075 | 13484365 | 17,9,21 | 0,0,0 | 9,22,14 | 3,4,8 | 280 | 150 | 5.10E-09 | 2.42E-07 | 1.0,1.0,1.0 | 0.616,0.747,0.48 | 0.384 | ENSG00000157 | [[[[13484076, 134 | 3 |
| ENSG00000157 | RAB28 | chr4 | 13479429 | 13479526 | 13460698 | 13460828 | 13484075 | 13484331 | 8,2,17 | 0,0,0 | 4,12,4 | 2,1,3 | 247 | 150 | 0.000131206 | 0.002018914 | 1.0,1.0,1.0 | 0.548,0.879,0.44 | 0.375 | ENSG00000157 | [[[[13484076, 134 | 2 |
| ENSG00000105 | TTC26 | chr7 | 139160947 | 139161022 | 139142255 | 139142305 | 139165131 | 139165234 | 8,3,13 | 2,0,0 | 7,17,13 | 4,6,4 | 225 | 150 | 0.000191863 | 0.002813077 | 0.727,1.0,1.0 | 0.538,0.654,0.68 | 0.284 | ENSG00000105 | [[[[139133778, 13 | 3 |
| ENSG00000153 | TRIP12 | chr2 | 229774096 | 229774261 | 229771518 | 229771632 | 229785756 | 229785855 | 114,107,138 | 0,0,0 | 60,148,97 | 10,13,5 | 300 | 150 | 5.73E-11 | 3.88E-09 | 1.0,1.0,1.0 | 0.75,0.851,0.907 | 0.164 | ENSG00000153 | [[[[229921880, 22 | 4 |
| ENSG00000153 | TRIP12 | chr2 | 229777314 | 229777479 | 229771518 | 229771632 | 229785756 | 229785855 | 1,2,1 | 0,0,0 | 2,2,3 | 10,13,5 | 300 | 150 | 9.26E-10 | 5.18E-08 | 1.0,1.0,1.0 | 0.091,0.071,0.23 | 0.869 | ENSG00000153 | [[[[229921880, 22 | 4 |
| ENSG00000153 | TRIP12 | chr2 | 229777314 | 229777479 | 229771518 | 229771632 | 229792152 | 229792226 | 1,2,1 | 0,0,0 | 1,1,3 | 3,7,1 | 300 | 150 | 2.47E-12 | 2.12E-10 | 1.0,1.0,1.0 | 0.143,0.067,0.6 | 0.73 | ENSG00000153 | [[[[229921880, 22 | 10 |
| ENSG00000100 | POLE2 | chr14 | 49654783 | 49654838 | 49654154 | 49654214 | 49663314 | 49663387 | 10,6,4 | 0,0,0 | 8,13,10 | 2,6,3 | 205 | 150 | 2.85E-07 | 8.95E-06 | 1.0,1.0,1.0 | 0.745,0.613,0.70 | 0.311 | ENSG00000100 | [[[[49688136, 496 | 3 |
| ENSG00000115 | NCL | chr2 | 231456009 | 231456136 | 231455400 | 231455624 | 231457000 | 231457124 | 172,016,531,940 | 0,0,0 | 170,532,571,714 | 19,48,27 | 277 | 150 | 5.59E-10 | 3.24E-08 | 1.0,1.0,1.0 | 0.98,0.974,0.972 | 0.025 | ENSG00000115 | [[[[231464336, 23 | 2 |
| ENSG00000115 | NCL | chr2 | 231456630 | 231456764 | 231455400 | 231455624 | 231457000 | 231457124 | 8,908,391,054 | 0,0,0 | 7,671,740,868 | 19,48,27 | 284 | 150 | 0 | 0 | 1.0,1.0,1.0 | 0.955,0.95,0.944 | 0.05 | ENSG00000115 | [[[[231464336, 23 | 2 |
| ENSG00000115 | NCL | chr2 | 231457642 | 231457800 | 231457000 | 231457124 | 231459000 | 231459125 | 9,458,841,082 | 0,0,0 | 7,441,531,875 | 67,140,72 | 300 | 150 | 0 | 0 | 1.0,1.0,1.0 | 0.847,0.845,0.85 | 0.15 | ENSG00000115 | [[[[231464336, 23 | 2 |
| ENSG00000137 | ATAT1 | chr6 | 30640376 | 30640422 | 30628328 | 30628430 | 30642175 | 30642247 | 17,8,14 | 1,0,0 | 33,47,50 | 14,50,12 | 196 | 150 | 1.13E-08 | 4.97E-07 | 0.929,1.0,1.0 | 0.643,0.418,0.76 | 0.369 | ENSG00000137 | [[[[30626874, 306 | 2 |
| ENSG00000137 | ATAT1 | chr6 | 30640534 | 30640603 | 30628328 | 30628430 | 30642175 | 30642247 | 2,3,3 | 1,0,0 | 7,10,9 | 14,50,12 | 219 | 150 | 3.53E-07 | 1.09E-05 | 0.578,1.0,1.0 | 0.255,0.12,0.339 | 0.621 | ENSG00000137 | [[[[30626874, 306 | 2 |
| ENSG00000131 | ACTR10 | chr14 | 58213630 | 58213698 | 58208998 | 58209107 | 58215204 | 58215284 | 39,45,60 | 0,2,1 | 46,69,50 | 4,17,5 | 218 | 150 | 2.07E-06 | 5.27E-05 | 1.0,0.939,0.976 | 0.888,0.736,0.87 | 0.139 | ENSG00000131 | [[[[58200149, 582 | 2 |
| ENSG00000115 | ORC4 | chr2 | 147972738 | 147972829 | 147958591 | 147958866 | 148020632 | 148020723 | 16,10,24 | 0,0,2 | 12,30,31 | 6,20,3 | 241 | 150 | 3.98E-06 | 9.44E-05 | 1.0,1.0,0.882 | 0.555,0.483,0.86 | 0.326 | ENSG00000115 | [[[[148020633, 14 | 3 |
| ENSG00000151 | GXYLT1 | chr12 | 42097441 | 42097614 | 42081844 | 42087947 | 42105817 | 42106069 | 21,14,43 | 0,0,0 | 40,80,40 | 1,8,0 | 300 | 150 | 0.000571114 | 0.00720016 | 1.0,1.0,1.0 | 0.952,0.833,1.0 | 0.072 | ENSG00000151 | [[[[42144426, 421 | 2 |
| ENSG00000105 | GTPBP10 | chr7 | 90354457 | 90354549 | 90352815 | 90353009 | 90372154 | 90372228 | 17,5,25 | 0,0,1 | 19,51,21 | 10,3,5 | 242 | 150 | 6.79E-06 | 0.000151619 | 1.0,1.0,0.939 | 0.541,0.913,0.72 | 0.254 | ENSG00000105 | [[[[90346716, 903 | 2 |
| ENSG00000105 | GTPBP10 | chr7 | 90355085 | 90355230 | 90352815 | 90353009 | 90372154 | 90372228 | 12,7,19 | 0,0,1 | 7,22,22 | 10,3,5 | 295 | 150 | 1.99E-06 | 5.08E-05 | 1.0,1.0,0.906 | 0.263,0.789,0.69 | 0.388 | ENSG00000105 | [[[[90346716, 903 | 2 |
| ENSG00000123 | PFDN5 | chr12 | 53296429 | 53296551 | 53295524 | 53295639 | 53297849 | 53297924 | 0,0,1 | 91,57,95 | 25,30,17 | 91,151,85 | 272 | 150 | 8.88E-16 | 1.21E-13 | 0.0,0.0,0.006 | 0.132,0.099,0.09 | -0.108 | ENSG00000123 | [[[[53295542, 532 | 2 |
| ENSG00000123 | PFDN5 | chr12 | 53296429 | 53296551 | 53295524 | 53295639 | 53298044 | 53298150 | 0,0,0 | 5,1,2 | 4,2,3 | 4,1,1 | 272 | 150 | 2.69E-07 | 8.50E-06 | 0.0,0.0,0.0 | 0.355,0.524,0.62 | -0.501 | ENSG00000123 | [[[[53295542, 532 | 3 |
| ENSG00000123 | PFDN5 | chr12 | 53296453 | 53296551 | 53295541 | 53295639 | 53297849 | 53297924 | 0,1,1 | 91,57,95 | 14,21,16 | 91,151,85 | 248 | 150 | 4.79E-11 | 3.30E-09 | 0.0,0.01,0.006 | 0.085,0.078,0.10 | -0.083 | ENSG00000123 | [[[[53295542, 532 | 2 |
| ENSG00000053 | AKR7A2 | chr1 | 19307313 | 19307410 | 19307001 | 19307101 | 19308454 | 19308642 | 139,161,264 | 1,1,0 | 133,245,155 | 2,13,12 | 247 | 150 | 9.80E-08 | 3.45E-06 | 0.988,0.99,1.0 | 0.976,0.92,0.887 | 0.065 | ENSG00000053 | [[[[19311827, 193 | 2 |
| ENSG00000053 | AKR7A2 | chr1 | 19308157 | 19308262 | 19307001 | 19307101 | 19308454 | 19308505 | 52,77,182 | 1,1,0 | 100,204,147 | 2,13,12 | 255 | 150 | 1.40E-06 | 3.71E-05 | 0.968,0.978,1.0 | 0.967,0.902,0.87 | 0.066 | ENSG00000053 | [[[[19311827, 193 | 2 |
| ENSG00000134 | PRPF38B | chr1 | 108696276 | 108696337 | 108692322 | 108692867 | 108698603 | 108698827 | 8,4,13 | 0,0,0 | 5,16,9 | 0,2,8 | 211 | 150 | 3.27E-05 | 0.000602631 | 1.0,1.0,1.0 | 1.0,0.85,0.444 | 0.235 | ENSG00000134 | [[[[108692310, 10 | 3 |
| ENSG00000134 | PRPF38B | chr1 | 108696276 | 108696337 | 108695701 | 108695770 | 108698603 | 108698827 | 8,4,13 | 0,0,0 | 7,21,9 | 1,10,8 | 211 | 150 | 2.78E-08 | 1.12E-06 | 1.0,1.0,1.0 | 0.833,0.599,0.44 | 0.375 | ENSG00000134 | [[[[108692310, 10 | 2 |
| ENSG00000136 | SMPD4 | chr2 | 130156584 | 130156675 | 130156026 | 130156135 | 130164373 | 130164445 | 4,19,31 | 0,0,0 | 27,47,40 | 0,6,4 | 241 | 150 | 0.000102133 | 0.001625382 | 1.0,1.0,1.0 | 1.0,0.83,0.862 | 0.103 | ENSG00000136 | [[[[130181530, 13 | 3 |
| ENSG00000136 | SMPD4 | chr2 | 130156584 | 130156813 | 130156034 | 130156135 | 130164373 | 130164445 | 4,19,32 | 0,0,0 | 24,58,44 | 0,6,4 | 300 | 150 | 7.51E-05 | 0.001242559 | 1.0,1.0,1.0 | 1.0,0.829,0.846 | 0.108 | ENSG00000136 | [[[[130181530, 13 | 3 |
| ENSG00000136 | SMPD4 | chr2 | 130157250 | 130157396 | 130156584 | 130156626 | 130164373 | 130164445 | 4,3,29 | 0,0,0 | 29,32,41 | 1,5,4 | 296 | 150 | 3.81E-05 | 0.000690527 | 1.0,1.0,1.0 | 0.936,0.764,0.83 | 0.154 | ENSG00000136 | [[[[130181530, 13 | 2 |
| ENSG00000100 | CDKN3 | chr14 | 54401523 | 54401579 | 54396848 | 54397077 | 54408744 | 54408789 | 46,29,61 | 0,0,0 | 27,75,29 | 8,5,8 | 206 | 150 | 6.38E-09 | 2.96E-07 | 1.0,1.0,1.0 | 0.711,0.916,0.72 | 0.216 | ENSG00000100 | [[[[54397007, 543 | 2 |
| ENSG00000160 | ANO10 | chr3 | 43574808 | 43574864 | 43565652 | 43565727 | 43605713 | 43605863 | 5,3,10 | 0,0,0 | 3,8,9 | 4,12,2 | 206 | 150 | 3.08E-09 | 1.54E-07 | 1.0,1.0,1.0 | 0.353,0.327,0.76 | 0.518 | ENSG00000160 | [[[[43621909, 436 | 5 |

| | | | | | | | | | | | | | | | | | | | | | | |
|---|---|---|---|---|---|---|---|---|---|---|---|---|---|---|---|---|---|---|---|---|---|---|
| ENSG000001607 | ANO10 | chr3 | 43598531 | 43598666 | 43565652 | 43565727 | 43605713 | 43605863 | 1,3,3 | 0,0,0 | 3,1,1 | 4,12,2 | 285 | 150 | 3.49E-14 | 3.87E-12 | 1.0,1.0,1.0 | 0.283,0.042,0.20 | 0.822 | ENSG000001607 | [[[[43621909, 436 | 5 |
| ENSG000001607 | ANO10 | chr3 | 43598531 | 43598666 | 43580352 | 43580472 | 43605713 | 43605863 | 34,21,29 | 0,0,0 | 22,23,25 | 2,5,2 | 285 | 150 | 6.74E-06 | 0.000150687 | 1.0,1.0,1.0 | 0.853,0.708,0.86 | 0.19 | ENSG000001607 | [[[[43621909, 436 | 2 |
| ENSG000001013 | SAMHD1 | chr20 | 36916721 | 36916830 | 36911217 | 36912552 | 36919363 | 36919519 | 18,9,17 | 0,0,0 | 2,21,6 | 3,2,8 | 259 | 150 | 2.34E-08 | 9.59E-07 | 1.0,1.0,1.0 | 0.279,0.859,0.30 | 0.52 | ENSG000001013 | [[[[36951436, 369 | 2 |
| ENSG000001093 | UBE2D3 | chr4 | 102809067 | 102809159 | 102797402 | 102797460 | 102809671 | 102809703 | 2,1,3 | 0,0,0 | 7,4,4 | 1,5,1 | 242 | 150 | 8.22E-06 | 0.000179811 | 1.0,1.0,1.0 | 0.813,0.331,0.71 | 0.381 | ENSG000001093 | [[[[102827427, 1( | 3 |
| ENSG000001327 | RAF1 | chr3 | 12585121 | 12585253 | 12583600 | 12584657 | 12585680 | 12585799 | 87,59,79 | 0,0,0 | 71,170,94 | 6,19,4 | 282 | 150 | 4.78E-09 | 2.29E-07 | 1.0,1.0,1.0 | 0.863,0.826,0.92 | 0.128 | ENSG000001327 | [[[[12663813, 126 | 2 |
| ENSG000002737 | CYFIP1 | chr15 | 22873490 | 22873729 | 22867051 | 22870192 | 22874549 | 22874644 | 99,127,140 | 0,0,0 | 46,127,56 | 2,13,7 | 300 | 150 | 2.10E-10 | 1.31E-08 | 1.0,1.0,1.0 | 0.92,0.83,0.8 | 0.15 | ENSG000002737 | [[[[22980287, 229 | 2 |
| ENSG000001224 | RWDD3 | chr1 | 95239776 | 95239909 | 95234220 | 95234315 | 95246755 | 95247217 | 10,3,14 | 2,0,0 | 6,17,11 | 2,7,7 | 283 | 150 | 3.28E-05 | 0.000604889 | 0.726,1.0,1.0 | 0.614,0.563,0.45 | 0.365 | ENSG000001224 | [[[[95234210, 952 | 2 |
| ENSG000001224 | RWDD3 | chr1 | 95244210 | 95244715 | 95234210 | 95234315 | 95246755 | 95247223 | 13,7,26 | 2,0,0 | 9,17,12 | 2,7,7 | 300 | 150 | 3.90E-06 | 9.27E-05 | 0.765,1.0,1.0 | 0.692,0.548,0.46 | 0.354 | ENSG000001224 | [[[[95234210, 952 | 2 |
| ENSG000001224 | RWDD3 | chr1 | 95246541 | 95246657 | 95234154 | 95234315 | 95246755 | 95247217 | 49,41,64 | 2,0,0 | 41,109,60 | 2,7,7 | 266 | 150 | 6.82E-05 | 0.001145154 | 0.933,1.0,1.0 | 0.92,0.898,0.829 | 0.095 | ENSG000001224 | [[[[95234210, 952 | 2 |
| ENSG000001108 | ASIC1 | chr12 | 50080497 | 50080727 | 50078923 | 50078980 | 50081101 | 50081181 | 2,2,2 | 0,0,1 | 0,1,3 | 2,2,0 | 300 | 150 | 0.000116599 | 0.001822485 | 1.0,1.0,0.5 | 0.0,0.2,1.0 | 0.433 | ENSG000001108 | [[[[50057596, 50( | 2 |
| ENSG000001808 | PPA1 | chr10 | 70206263 | 70206333 | 70204872 | 70204915 | 70214499 | 70214586 | 213,154,165 | 0,0,0 | 176,335,224 | 6,16,6 | 220 | 150 | 7.40E-07 | 2.10E-05 | 1.0,1.0,1.0 | 0.952,0.935,0.96 | 0.05 | ENSG000001808 | [[[[70233264, 702 | 4 |
| ENSG000001808 | PPA1 | chr10 | 70209204 | 70209290 | 70203588 | 70206333 | 70214499 | 70214586 | 231,183,232 | 0,0,0 | 236,384,223 | 6,12,8 | 236 | 150 | 3.56E-06 | 8.56E-05 | 1.0,1.0,1.0 | 0.962,0.953,0.94 | 0.046 | ENSG000001808 | [[[[70233264, 702 | 3 |
| ENSG000001808 | PPA1 | chr10 | 70209557 | 70209685 | 70209204 | 70209290 | 70214499 | 70214586 | 251,219,316 | 0,0,0 | 285,500,303 | 23,19,22 | 278 | 150 | 2.11E-15 | 2.75E-13 | 1.0,1.0,1.0 | 0.87,0.934,0.881 | 0.105 | ENSG000001808 | [[[[70233264, 702 | 2 |
| ENSG000001808 | PPA1 | chr10 | 70213462 | 70213589 | 70204872 | 70204915 | 70214499 | 70214586 | 248,221,315 | 0,0,0 | 189,442,250 | 6,16,6 | 277 | 150 | 1.80E-07 | 5.93E-06 | 1.0,1.0,1.0 | 0.945,0.937,0.95 | 0.053 | ENSG000001808 | [[[[70233264, 702 | 4 |
| ENSG000001808 | PPA1 | chr10 | 70213462 | 70213589 | 70206263 | 70206333 | 70214499 | 70214586 | 248,221,315 | 0,0,0 | 186,438,247 | 6,12,8 | 277 | 150 | 6.29E-07 | 1.82E-05 | 1.0,1.0,1.0 | 0.944,0.952,0.94 | 0.053 | ENSG000001808 | [[[[70233264, 702 | 3 |
| ENSG000001701 | UBE2E1 | chr3 | 23887566 | 23887699 | 23811457 | 23811510 | 23890508 | 23890872 | 202,161,246 | 1,0,1 | 141,225,180 | 27,27,20 | 283 | 150 | 0 | 0 | 0.991,1.0,0.992 | 0.735,0.815,0.82 | 0.202 | ENSG000001701 | [[[[23805955, 238 | 2 |
| ENSG000001701 | UBE2E1 | chr3 | 23889111 | 23889259 | 23811457 | 23811510 | 23890508 | 23890872 | 293,254,318 | 1,0,1 | 163,283,199 | 27,27,20 | 298 | 150 | 0 | 0 | 0.993,1.0,0.994 | 0.752,0.841,0.83 | 0.187 | ENSG000001701 | [[[[23805955, 238 | 2 |
| ENSG000001262 | UBA2 | chr19 | 34431860 | 34431931 | 34428372 | 34428570 | 34433347 | 34433412 | 26,38,46 | 0,0,0 | 21,55,52 | 4,12,10 | 221 | 150 | 2.99E-10 | 1.81E-08 | 1.0,1.0,1.0 | 0.781,0.757,0.77 | 0.228 | ENSG000001262 | [[[[34428381, 344 | 2 |
| ENSG000001887 | FAM221A | chr7 | 23689268 | 23689459 | 23680183 | 23680283 | 23691389 | 23691399 | 19,16,44 | 0,0,0 | 29,52,25 | 4,2,3 | 300 | 150 | 5.51E-05 | 0.000949744 | 1.0,1.0,1.0 | 0.784,0.929,0.80 | 0.16 | ENSG000001887 | [[[[23680199, 236 | 2 |
| ENSG000001487 | MKI67 | chr10 | 128122880 | 128122996 | 128119252 | 128119319 | 128125575 | 128125756 | 33,5,74 | 0,0,0 | 6,51,32 | 9,41,15 | 266 | 150 | 0 | 0 | 1.0,1.0,1.0 | 0.273,0.412,0.54 | 0.59 | ENSG000001487 | [[[[128126099, 12 | 2 |
| ENSG000000770 | TOP2B | chr3 | 25618653 | 25618849 | 25615430 | 25615586 | 25619861 | 25620062 | 61,45,65 | 0,0,0 | 33,78,48 | 2,5,9 | 300 | 150 | 1.66E-07 | 5.48E-06 | 1.0,1.0,1.0 | 0.892,0.886,0.72 | 0.165 | ENSG000000770 | [[[[25664229, 256 | 2 |
| ENSG000000770 | TOP2B | chr3 | 25620681 | 25620816 | 25619861 | 25620062 | 25624681 | 25624803 | 48,47,74 | 0,0,0 | 34,76,54 | 9,7,2 | 285 | 150 | 1.02E-07 | 3.58E-06 | 1.0,1.0,1.0 | 0.665,0.851,0.93 | 0.183 | ENSG000000770 | [[[[25664229, 256 | 3 |
| ENSG000001510 | NGLY1 | chr3 | 25736003 | 25736149 | 25733871 | 25733982 | 25751097 | 25751263 | 26,27,39 | 0,0,2 | 15,30,25 | 6,11,4 | 296 | 150 | 4.01E-08 | 1.55E-06 | 1.0,1.0,0.908 | 0.559,0.58,0.76 | 0.336 | ENSG000001510 | [[[[25783260, 257 | 3 |
| ENSG000001510 | NGLY1 | chr3 | 25739576 | 25739799 | 25733871 | 25733982 | 25751097 | 25751263 | 30,21,32 | 0,0,2 | 17,36,22 | 6,11,4 | 300 | 150 | 1.10E-07 | 3.81E-06 | 1.0,1.0,0.889 | 0.586,0.621,0.73 | 0.316 | ENSG000001510 | [[[[25783260, 257 | 3 |
| ENSG000001122 | CCNC | chr6 | 99558496 | 99558548 | 99551839 | 99551895 | 99561596 | 99561658 | 15,13,16 | 0,0,0 | 6,20,10 | 2,7,1 | 202 | 150 | 1.13E-06 | 3.06E-05 | 1.0,1.0,1.0 | 0.69,0.68,0.881 | 0.25 | ENSG000001122 | [[[[99568496, 99( | 2 |
| ENSG000001122 | CCNC | chr6 | 99558496 | 99558548 | 99551839 | 99551895 | 99562841 | 99562948 | 15,13,16 | 0,0,0 | 7,21,12 | 1,2,8 | 202 | 150 | 2.03E-06 | 5.19E-05 | 1.0,1.0,1.0 | 0.839,0.886,0.52 | 0.249 | ENSG000001122 | [[[[99568496, 99( | 3 |
| ENSG000001122 | CCNC | chr6 | 99561596 | 99561681 | 99551839 | 99551895 | 99562841 | 99562948 | 57,46,80 | 0,0,0 | 33,103,50 | 1,2,8 | 235 | 150 | 0.000119455 | 0.001860254 | 1.0,1.0,1.0 | 0.955,0.97,0.8 | 0.092 | ENSG000001122 | [[[[99568496, 99( | 3 |
| ENSG000001043 | PLAT | chr8 | 42182718 | 42182765 | 42181936 | 42182022 | 42187397 | 42187572 | 69,135,200 | 0,0,0 | 153,346,240 | 3,17,3 | 197 | 150 | 0.000129096 | 0.001990454 | 1.0,1.0,1.0 | 0.975,0.939,0.98 | 0.034 | ENSG000001043 | [[[[42207494, 422 | 2 |
| ENSG000001690 | BUB1 | chr2 | 110657545 | 110657645 | 110657035 | 110657117 | 110661581 | 110661841 | 40,24,43 | 0,0,0 | 12,25,20 | 1,5,3 | 250 | 150 | 5.62E-06 | 0.000128678 | 1.0,1.0,1.0 | 0.878,0.75,0.8 | 0.191 | ENSG000001690 | [[[[110677970, 11 | 4 |
| ENSG000001365 | TANK | chr2 | 161204674 | 161204793 | 161179482 | 161179761 | 161223914 | 161223991 | 16,17,25 | 0,0,0 | 7,35,29 | 4,8,2 | 269 | 150 | 4.02E-07 | 1.22E-05 | 1.0,1.0,1.0 | 0.494,0.709,0.89 | 0.302 | ENSG000001365 | [[[[161160428, 16 | 2 |
| ENSG000001365 | TANK | chr2 | 161224630 | 161224746 | 161204674 | 161204793 | 161230970 | 161231017 | 20,16,27 | 0,0,0 | 21,36,12 | 4,11,2 | 266 | 150 | 1.12E-08 | 4.94E-07 | 1.0,1.0,1.0 | 0.748,0.649,0.77 | 0.277 | ENSG000001365 | [[[[161160428, 16 | 2 |
| ENSG000001303 | QRSL1 | chr6 | 106655614 | 106655732 | 106652466 | 106652582 | 106662979 | 106663185 | 17,11,28 | 0,0,0 | 17,24,34 | 12,8,5 | 268 | 150 | 2.52E-09 | 1.29E-07 | 1.0,1.0,1.0 | 0.442,0.627,0.79 | 0.38 | ENSG000001303 | [[[[106629578, 1( | 2 |
| ENSG000001231 | WWP1 | chr8 | 86398571 | 86398638 | 86381504 | 86381629 | 86402018 | 86402203 | 1,8,9 | 0,0,1 | 8,10,7 | 2,2,2 | 217 | 150 | 0.000765494 | 0.009236596 | 1.0,1.0,0.862 | 0.734,0.776,0.70 | 0.215 | ENSG000001231 | [[[[86342547, 86( | 2 |
| ENSG000001154 | USP34 | chr2 | 61190270 | 61190414 | 61188909 | 61189069 | 61192900 | 61192980 | 45,28,59 | 0,0,0 | 26,79,38 | 5,2,2 | 294 | 150 | 0.000189648 | 0.002787039 | 1.0,1.0,1.0 | 0.726,0.953,0.90 | 0.138 | ENSG000001154 | [[[[61470650, 614 | 2 |
| ENSG000001116 | KRR1 | chr12 | 75505197 | 75505254 | 75503903 | 75504074 | 75506483 | 75506609 | 18,20,56 | 0,0,0 | 22,79,42 | 5,16,2 | 207 | 150 | 8.51E-08 | 3.04E-06 | 1.0,1.0,1.0 | 0.761,0.782,0.93 | 0.173 | ENSG000001116 | [[[[75511513, 755 | 2 |
| ENSG000001663 | TPP1 | chr11 | 6616974 | 6617153 | 6616314 | 6616503 | 6617300 | 6617428 | 72,95,134 | 0,0,0 | 58,104,74 | 1,1,8 | 300 | 150 | 0.000513302 | 0.006572442 | 1.0,1.0,1.0 | 0.967,0.981,0.82 | 0.077 | ENSG000001663 | [[[[6619384, 661( | 2 |
| ENSG000001842 | IRAK1 | chrX | 154013042 | 154013433 | 154011730 | 154011917 | 154016031 | 154016097 | 1,4,4 | 0,0,0 | 12,13,9 | 2,7,5 | 300 | 150 | 1.51E-08 | 6.48E-07 | 1.0,1.0,1.0 | 0.75,0.481,0.474 | 0.432 | ENSG000001842 | [[[[154019677, 15 | 3 |
| ENSG000000120 | ERCC1 | chr19 | 45413676 | 45413745 | 45407332 | 45409725 | 45416820 | 45416897 | 20,29,30 | 0,5,0 | 33,57,39 | 5,13,16 | 219 | 150 | 0.000346306 | 0.004670413 | 1.0,0.799,1.0 | 0.819,0.75,0.625 | 0.202 | ENSG000000120 | [[[[45423781, 454 | 3 |
| ENSG000000120 | ERCC1 | chr19 | 45414860 | 45414960 | 45407332 | 45409725 | 45416820 | 45416859 | 70,104,132 | 0,5,0 | 115,215,155 | 5,13,16 | 250 | 150 | 3.53E-05 | 0.000645353 | 1.0,0.926,1.0 | 0.932,0.908,0.85 | 0.078 | ENSG000000120 | [[[[45423781, 454 | 3 |
| ENSG000000478 | MAP4 | chr3 | 47909037 | 47912421 | 47877416 | 47877523 | 47914816 | 47914939 | 13,7,9 | 97,42,137 | 18,55,44 | 118,214,183 | 300 | 150 | 0.000371569 | 0.004970649 | 0.063,0.077,0.03 | 0.071,0.114,0.10 | -0.04 | ENSG000000478 | [[[[48016334, 48( | 2 |
| ENSG000001142 | COL7A1 | chr3 | 48571100 | 48571160 | 48570638 | 48570710 | 48571242 | 48571278 | 8,38,90 | 0,0,3 | 10,32,13 | 4,6,2 | 210 | 150 | 1.24E-05 | 0.000257547 | 1.0,1.0,0.955 | 0.641,0.792,0.82 | 0.233 | ENSG000001142 | [[[[48595268, 48( | 2 |
| ENSG000001155 | DARS | chr2 | 135924386 | 135924498 | 135920452 | 135920600 | 135932782 | 135932842 | 51,53,79 | 0,0,0 | 29,71,43 | 7,2,2 | 262 | 150 | 1.78E-05 | 0.000353911 | 1.0,1.0,1.0 | 0.703,0.953,0.92 | 0.14 | ENSG000001155 | [[[[135985403, 1: | 2 |
| ENSG000001830 | SLC25A10 | chr17 | 81716811 | 81716855 | 81715920 | 81716050 | 81717481 | 81717491 | 1,16,31 | 0,0,0 | 10,32,24 | 5,5,3 | 194 | 150 | 1.41E-06 | 3.74E-05 | 1.0,1.0,1.0 | 0.607,0.832,0.86 | 0.233 | ENSG000001830 | [[[[81712284, 817 | 2 |
| ENSG000001442 | LNPK | chr2 | 175937343 | 175937514 | 175923891 | 175930199 | 175939551 | 175939657 | 58,28,45 | 0,0,0 | 28,73,53 | 2,6,4 | 300 | 150 | 1.14E-05 | 0.000239142 | 1.0,1.0,1.0 | 0.875,0.859,0.86 | 0.132 | ENSG000001442 | [[[[176002160, 17 | 2 |
| ENSG000000045 | RBM6 | chr3 | 49962575 | 49962685 | 49940006 | 49940225 | 50054334 | 50054395 | 12,16,41 | 0,0,0 | 25,40,29 | 5,4,2 | 260 | 150 | 7.85E-06 | 0.000172814 | 1.0,1.0,1.0 | 0.743,0.852,0.89 | 0.171 | ENSG000000045 | [[[[49940150, 49( | 6 |
| ENSG000000045 | RBM6 | chr3 | 49999439 | 49999513 | 49962575 | 49962685 | 50048244 | 50048319 | 14,17,33 | 19,11,14 | 7,14,19 | 37,74,44 | 224 | 150 | 2.32E-05 | 0.000448585 | 0.33,0.509,0.383 | 0.112,0.112,0.22 | 0.258 | ENSG000000045 | [[[[49940150, 49( | 4 |
| ENSG000000045 | RBM6 | chr3 | 50048244 | 50048319 | 49940006 | 49940225 | 50054334 | 50054395 | 28,21,41 | 0,0,0 | 51,63,57 | 5,4,2 | 225 | 150 | 0.000318371 | 0.004347534 | 1.0,1.0,1.0 | 0.872,0.913,0.95 | 0.088 | ENSG000000045 | [[[[49940150, 49( | 6 |
| ENSG000001464 | TMEM181 | chr6 | 158623549 | 158623607 | 158608658 | 158608750 | 158631322 | 158631389 | 5,3,13 | 0,0,0 | 1,12,8 | 1,10,8 | 208 | 150 | 2.58E-10 | 1.59E-08 | 1.0,1.0,1.0 | 0.419,0.464,0.41 | 0.566 | ENSG000001464 | [[[[158560091, 15 | 5 |
| ENSG000001323 | ILKAP | chr2 | 238189852 | 238189972 | 238188130 | 238188257 | 238194804 | 238194870 | 56,63,74 | 0,1,0 | 102,174,90 | 10,16,7 | 270 | 150 | 3.77E-08 | 1.47E-06 | 1.0,0.972,1.0 | 0.85,0.858,0.877 | 0.129 | ENSG000001323 | [[[[238203499, 23 | 2 |
| ENSG000001323 | ILKAP | chr2 | 238194274 | 238194331 | 238188130 | 238188257 | 238194804 | 238194870 | 9,19,19 | 0,1,0 | 20,40,34 | 10,16,7 | 207 | 150 | 2.76E-08 | 1.12E-06 | 1.0,0.932,1.0 | 0.592,0.644,0.77 | 0.306 | ENSG000001323 | [[[[238203499, 23 | 2 |
| ENSG000001639 | GNL3 | chr3 | 52689073 | 52689206 | 52688108 | 52688192 | 52690944 | 52691071 | 97,91,164 | 0,0,0 | 109,311,219 | 4,10,7 | 283 | 150 | 1.94E-05 | 0.000382641 | 1.0,1.0,1.0 | 0.935,0.943,0.94 | 0.06 | ENSG000001639 | [[[[52686032, 52( | 2 |
| ENSG000001639 | GNL3 | chr3 | 52690944 | 52691071 | 52690591 | 52690704 | 52692871 | 52692929 | 118,120,165 | 0,0,0 | 97,293,184 | 7,4,6 | 277 | 150 | 9.87E-05 | 0.00158074 | 1.0,1.0,1.0 | 0.882,0.975,0.94 | 0.067 | ENSG000001639 | [[[[52686032, 52( | 2 |
| ENSG000001014 | RPRD1B | chr20 | 38040434 | 38040564 | 38033545 | 38034098 | 38057531 | 38057644 | 1,3,16 | 0,0,0 | 9,17,21 | 1,14,4 | 280 | 150 | 1.48E-07 | 4.95E-06 | 1.0,1.0,1.0 | 0.828,0.394,0.73 | 0.347 | ENSG000001014 | [[[[38033746, 38( | 2 |
| ENSG000001014 | RPRD1B | chr20 | 38048347 | 38048481 | 38033724 | 38034098 | 38057531 | 38057644 | 21,15,41 | 0,0,0 | 15,34,31 | 1,14,4 | 284 | 150 | 6.24E-08 | 2.31E-06 | 1.0,1.0,1.0 | 0.888,0.562,0.80 | 0.249 | ENSG000001014 | [[[[38033746, 38( | 2 |
| ENSG000001633 | NAXE | chr1 | 156592556 | 156592670 | 156592100 | 156592209 | 156593407 | 156593555 | 201,197,273 | 0,0,0 | 241,390,238 | 26,46,15 | 264 | 150 | 0 | 0 | 1.0,1.0,1.0 | 0.84,0.828,0.9 | 0.144 | ENSG000001633 | [[[[156591776, 15 | 2 |
| ENSG000001040 | PPP2CB | chr8 | 30797580 | 30797754 | 30794191 | 30794281 | 30812319 | 30812872 | 232,150,232 | 0,0,0 | 80,142,110 | 4,7,2 | 300 | 150 | 2.31E-05 | 0.00044678 | 1.0,1.0,1.0 | 0.909,0.91,0.965 | 0.072 | ENSG000001040 | [[[[30812320, 30( | 2 |
| ENSG000001040 | PPP2CB | chr8 | 30799545 | 30799755 | 30794191 | 30794281 | 30812319 | 30812872 | 166,175,287 | 0,0,0 | 84,228,159 | 4,7,2 | 300 | 150 | 0.000328175 | 0.00446222 | 1.0,1.0,1.0 | 0.913,0.942,0.97 | 0.057 | ENSG000001040 | [[[[30812320, 30( | 2 |
| ENSG000001040 | PPP2CB | chr8 | 30799716 | 30799755 | 30794197 | 30794281 | 30812319 | 30812597 | 13,13,16 | 0,0,0 | 47,91,73 | 4,7,2 | 189 | 150 | 0.000683091 | 0.008384376 | 1.0,1.0,1.0 | 0.903,0.912,0.96 | 0.073 | ENSG000001040 | [[[[30812320, 30( | 2 |
| ENSG000001204 | TCP1 | chr6 | 159779626 | 159779790 | 159778497 | 159779261 | 159780442 | 159780566 | 393,422,435 | 0,0,0 | 507,986,464 | 30,40,22 | 300 | 150 | 0 | 0 | 1.0,1.0,1.0 | 0.894,0.925,0.91 | 0.089 | ENSG000001204 | [[[[159789405, 15 | 2 |
| ENSG000001204 | TCP1 | chr6 | 159779626 | 159779790 | 159778497 | 159779261 | 159780934 | 159781110 | 393,422,435 | 0,0,0 | 5,221,007,480 | 25,28,23 | 300 | 150 | 5.11E-15 | 6.35E-13 | 1.0,1.0,1.0 | 0.913,0.947,0.91 | 0.076 | ENSG000001204 | [[[[159789405, 15 | 3 |
| ENSG000001204 | TCP1 | chr6 | 159779894 | 159780087 | 159778984 | 159779261 | 159780442 | 159780566 | 595,519,707 | 0,0,0 | 5,721,252,666 | 30,40,22 | 300 | 150 | 0 | 0 | 1.0,1.0,1.0 | 0.905,0.94,0.938 | 0.072 | ENSG000001204 | [[[[159789405, 15 | 2 |
| ENSG000001204 | TCP1 | chr6 | 159779894 | 159780087 | 159779626 | 159779790 | 159780934 | 159781110 | 451,498,602 | 0,0,0 | 509,924,521 | 16,25,17 | 300 | 150 | 1.19E-12 | 1.07E-10 | 1.0,1.0,1.0 | 0.941,0.949,0.93 | 0.057 | ENSG000001204 | [[[[159789405, 15 | 2 |
| ENSG000001204 | TCP1 | chr6 | 159780442 | 159780566 | 159778497 | 159779261 | 159780934 | 159781110 | 484,482,577 | 0,0,0 | 6,301,312,743 | 25,28,23 | 274 | 150 | 9.55E-13 | 8.68E-11 | 1.0,1.0,1.0 | 0.932,0.962,0.94 | 0.053 | ENSG000001204 | [[[[159789405, 15 | 3 |
| ENSG000001204 | TCP1 | chr6 | 159780442 | 159780566 | 159779626 | 159779790 | 159780934 | 159781110 | 484,482,577 | 0,0,0 | 6,011,276,722 | 16,25,17 | 274 | 150 | 2.08E-09 | 1.08E-07 | 1.0,1.0,1.0 | 0.954,0.965,0.95 | 0.041 | ENSG000001204 | [[[[159789405, 15 | 2 |
| ENSG000001630 | SLMAP | chr3 | 57864806 | 57864857 | 57864547 | 57864716 | 57871635 | 57871698 | 14,7,15 | 0,0,0 | 6,34,10 | 2,5,4 | 201 | 150 | 1.86E-06 | 4.80E-05 | 1.0,1.0,1.0 | 0.691,0.835,0.65 | 0.274 | ENSG000001630 | [[[[57756309, 577 | 2 |
| ENSG000001682 | PDHB | chr3 | 58428472 | 58428614 | 58427629 | 58428179 | 58431876 | 58431984 | 136,117,178 | 0,0,0 | 105,205,124 | 3,7,9 | 292 | 150 | 1.08E-06 | 2.94E-05 | 1.0,1.0,1.0 | 0.947,0.938,0.87 | 0.08 | ENSG000001682 | [[[[58433768, 584 | 6 |
| ENSG000001682 | PDHB | chr3 | 58431730 | 58431793 | 58427629 | 58428179 | 58431876 | 58431984 | 40,60,105 | 0,0,0 | 32,112,55 | 3,7,9 | 213 | 150 | 9.69E-08 | 3.42E-06 | 1.0,1.0,1.0 | 0.883,0.918,0.81 | 0.129 | ENSG000001682 | [[[[58433768, 584 | 6 |
| ENSG000001643 | GFM2 | chr5 | 74725639 | 74725755 | 74722378 | 74722561 | 74733021 | 74733098 | 42,14,31 | 0,0,0 | 47,79,20 | 5,6,0 | 266 | 150 | 6.11E-05 | 0.00104056 | 1.0,1.0,1.0 | 0.841,0.881,1.0 | 0.093 | ENSG000001643 | [[[[74766938, 747 | 3 |
| ENSG000001643 | GFM2 | chr5 | 74725940 | 74726126 | 74722378 | 74722561 | 74733021 | 74733098 | 4,4,7 | 0,0,0 | 16,24,20 | 5,6,0 | 300 | 150 | 4.64E-06 | 0.000108084 | 1.0,1.0,1.0 | 0.615,0.667,1.0 | 0.239 | ENSG000001643 | [[[[74766938, 747 | 3 |
| ENSG000001643 | GFM2 | chr5 | 74725940 | 74726126 | 74725639 | 74725755 | 74733021 | 74733098 | 23,30,54 | 0,0,0 | 56,96,59 | 2,8,1 | 300 | 150 | 0.000139969 | 0.002131364 | 1.0,1.0,1.0 | 0.933,0.857,0.96 | 0.081 | ENSG000001643 | [[[[74766938, 747 | 2 |
| ENSG000001643 | GFM2 | chr5 | 74738317 | 74738417 | 74736668 | 74736985 | 74741528 | 74741609 | 36,22,43 | 0,0,0 | 40,78,56 | 9,6,7 | 250 | 150 | 9.45E-08 | 3.35E-06 | 1.0,1.0,1.0 | 0.727,0.886,0.82 | 0.186 | ENSG000001643 | [[[[74766938, 747 | 3 |
| ENSG000001643 | GFM2 | chr5 | 74759368 | 74759426 | 74751367 | 74751493 | 74760901 | 74760986 | 6,7,17 | 0,0,0 | 12,22,19 | 8,9,5 | 208 | 150 | 1.27E-08 | 5.55E-07 | 1.0,1.0,1.0 | 0.52,0.638,0.733 | 0.37 | ENSG000001643 | [[[[74766938, 747 | 2 |
| ENSG000000671 | TRAM1 | chr8 | 70596262 | 70596321 | 70594500 | 70594590 | 70600018 | 70600082 | 66,39,101 | 0,0,0 | 74,197,112 | 6,10,5 | 209 | 150 | 7.27E-06 | 0.000161003 | 1.0,1.0,1.0 | 0.898,0.934,0.94 | 0.076 | ENSG000000671 | [[[[70608077, 70( | 3 |
| ENSG000000671 | TRAM1 | chr8 | 70597894 | 70598011 | 70596262 | 70596321 | 70600018 | 70600082 | 135,124,222 | 0,0,0 | 188,373,262 | 7,23,10 | 267 | 150 | 2.46E-09 | 1.26E-07 | 1.0,1.0,1.0 | 0.938,0.901,0.93 | 0.075 | ENSG000000671 | [[[[70608077, 70( | 2 |
| ENSG000000671 | TRAM1 | chr8 | 70598133 | 70598255 | 70594500 | 70594590 | 70600018 | 70600082 | 118,101,233 | 0,0,0 | 154,302,260 | 6,10,5 | 272 | 150 | 3.07E-05 | 0.000572025 | 1.0,1.0,1.0 | 0.934,0.943,0.96 | 0.052 | ENSG000000671 | [[[[70608077, 70( | 3 |
| ENSG000001483 | GPR107 | chr9 | 130100628 | 130100702 | 130099456 | 130099532 | 130104419 | 130104550 | 21,11,31 | 0,0,0 | 20,51,35 | 2,5,6 | 224 | 150 | 5.38E-06 | 0.000123787 | 1.0,1.0,1.0 | 0.87,0.872,0.796 | 0.154 | ENSG000001483 | [[[[130053906, 1: | 2 |
| ENSG000001013 | CDK5RAP1 | chr20 | 33374114 | 33374212 | 33372641 | 33372697 | 33401427 | 33401452 | 19,15,33 | 0,0,0 | 16,44,23 | 2,9,0 | 248 | 150 | 1.58E-05 | 0.000319234 | 1.0,1.0,1.0 | 0.829,0.747,1.0 | 0.141 | ENSG000001013 | [[[[33401428, 334 | 8 |
| ENSG000001013 | CDK5RAP1 | chr20 | 33385649 | 33385770 | 33372641 | 33372697 | 33392141 | 33392242 | 3,1,4 | 0,0,0 | 2,9,5 | 3,7,1 | 271 | 150 | 5.73E-09 | 2.69E-07 | 1.0,1.0,1.0 | 0.27,0.416,0.735 | 0.526 | ENSG000001013 | [[[[33401428, 334 | 4 |
| ENSG000001013 | CDK5RAP1 | chr20 | 33387322 | 33387533 | 33372641 | 33372697 | 33392141 | 33392242 | 23,30,22 | 0,0,0 | 16,43,21 | 3,7,1 | 300 | 150 | 1.90E-06 | 4.89E-05 | 1.0,1.0,1.0 | 0.727,0.754,0.91 | 0.202 | ENSG000001013 | [[[[33401428, 334 | 4 |
| ENSG000001013 | CDK5RAP1 | chr20 | 33396760 | 33397084 | 33372641 | 33372697 | 33401427 | 33401561 | 10,13,19 | 0,0,0 | 6,17,27 | 2,9,0 | 300 | 150 | 9.29E-06 | 0.000200626 | 1.0,1.0,1.0 | 0.6,0.486,1.0 | 0.305 | ENSG000001013 | [[[[33401428, 334 | 8 |

| | | | | | | | | | | | | | | | | | | | | | | |
|---|---|---|---|---|---|---|---|---|---|---|---|---|---|---|---|---|---|---|---|---|---|---|
| ENSG000001342 | NOTCH2 | chr1 | 119948413 | 119948566 | 119941525 | 119941754 | 119950723 | 119950837 | 32,9,70 | 0,0,0 | 23,59,54 | 0,8,3 | 300 | 150 | 2.62E-05 | 0.000497085 | 1.0,1.0,1.0 | 1.0,0.787,0.9 | 0.104 | ENSG000001342 | [[[[120069334, 12 | 2 |
| ENSG000001342 | NOTCH2 | chr1 | 119949006 | 119949126 | 119941525 | 119941754 | 119950723 | 119950837 | 32,8,50 | 0,0,0 | 12,58,51 | 0,8,3 | 270 | 150 | 2.83E-05 | 0.000530733 | 1.0,1.0,1.0 | 1.0,0.801,0.904 | 0.098 | ENSG000001342 | [[[[120069334, 12 | 2 |
| ENSG000001342 | NOTCH2 | chr1 | 119986959 | 119987082 | 119969510 | 119969744 | 120005328 | 120005588 | 18,14,46 | 0,0,0 | 11,72,68 | 1,4,4 | 273 | 150 | 0.000639435 | 0.007925124 | 1.0,1.0,1.0 | 0.858,0.908,0.90 | 0.11 | ENSG000001342 | [[[[120069334, 12 | 2 |
| ENSG000001384 | ABI2 | chr2 | 203391045 | 203391143 | 203380207 | 203380384 | 203395655 | 203395780 | 23,23,34 | 0,0,0 | 13,18,23 | 0,10,5 | 248 | 150 | 4.78E-07 | 1.42E-05 | 1.0,1.0,1.0 | 1.0,0.521,0.736 | 0.248 | ENSG000001384 | [[[[203328394, 20 | 3 |
| ENSG000001384 | ABI2 | chr2 | 203394699 | 203394846 | 203380207 | 203380384 | 203395655 | 203395780 | 64,36,79 | 0,0,0 | 13,41,35 | 0,10,5 | 297 | 150 | 6.78E-08 | 2.49E-06 | 1.0,1.0,1.0 | 1.0,0.674,0.78 | 0.182 | ENSG000001384 | [[[[203328394, 20 | 3 |
| ENSG000001384 | ABI2 | chr2 | 203394699 | 203394846 | 203380207 | 203380384 | 203395658 | 203395780 | 5,4,13 | 0,0,0 | 3,7,1 | 2,4,0 | 297 | 150 | 7.42E-05 | 0.001228004 | 1.0,1.0,1.0 | 0.431,0.469,1.0 | 0.367 | ENSG000001384 | [[[[203328394, 20 | 3 |
| ENSG000001384 | ABI2 | chr2 | 203411284 | 203411371 | 203380207 | 203380384 | 203416907 | 203417081 | 10,7,10 | 0,0,0 | 12,18,7 | 2,3,2 | 237 | 150 | 0.000115475 | 0.001808249 | 1.0,1.0,1.0 | 0.792,0.792,0.68 | 0.242 | ENSG000001384 | [[[[203328394, 20 | 7 |
| ENSG000001606 | SHC1 | chr1 | 154965542 | 154965781 | 154962297 | 154963931 | 154967670 | 154967797 | 296,298,549 | 0,0,1 | 453,833,573 | 16,32,18 | 300 | 150 | 1.97E-12 | 1.73E-10 | 1.0,1.0,0.996 | 0.934,0.929,0.94 | 0.064 | ENSG000001606 | [[[[154970032, 15 | 4 |
| ENSG000001606 | SHC1 | chr1 | 154965945 | 154966077 | 154965542 | 154965781 | 154967670 | 154967797 | 74,140,230 | 0,0,0 | 110,258,208 | 7,4,5 | 282 | 150 | 0.000162587 | 0.002430607 | 1.0,1.0,1.0 | 0.893,0.972,0.95 | 0.059 | ENSG000001606 | [[[[154970032, 15 | 3 |
| ENSG000001606 | SHC1 | chr1 | 154965945 | 154966080 | 154965542 | 154965781 | 154967670 | 154967797 | 78,144,236 | 0,0,0 | 111,269,210 | 7,4,5 | 285 | 150 | 0.000171611 | 0.002548509 | 1.0,1.0,1.0 | 0.893,0.973,0.95 | 0.059 | ENSG000001606 | [[[[154970032, 15 | 3 |
| ENSG000001606 | SHC1 | chr1 | 154966161 | 154966231 | 154965945 | 154966077 | 154967670 | 154967797 | 27,56,106 | 0,0,0 | 62,165,108 | 8,15,12 | 220 | 150 | 1.58E-09 | 8.43E-08 | 1.0,1.0,1.0 | 0.841,0.882,0.86 | 0.139 | ENSG000001606 | [[[[154970032, 15 | 2 |
| ENSG000001606 | SHC1 | chr1 | 154966161 | 154966231 | 154965945 | 154966080 | 154967670 | 154967797 | 68,84,168 | 0,0,0 | 95,210,153 | 5,12,5 | 220 | 150 | 2.70E-06 | 6.69E-05 | 1.0,1.0,1.0 | 0.928,0.923,0.95 | 0.065 | ENSG000001606 | [[[[154970032, 15 | 2 |
| ENSG000000909 | TNRC6A | chr16 | 24804176 | 24804319 | 24797914 | 24797966 | 24805604 | 24805733 | 59,40,107 | 0,0,0 | 51,144,86 | 7,6,5 | 293 | 150 | 1.04E-06 | 2.86E-05 | 1.0,1.0,1.0 | 0.789,0.925,0.89 | 0.129 | ENSG000000909 | [[[[24729651, 247 | 3 |
| ENSG000000909 | TNRC6A | chr16 | 24805013 | 24805151 | 24797914 | 24797966 | 24805604 | 24805733 | 45,47,86 | 0,0,0 | 33,74,68 | 7,6,5 | 288 | 150 | 5.27E-08 | 1.98E-06 | 1.0,1.0,1.0 | 0.711,0.865,0.87 | 0.183 | ENSG000000909 | [[[[24729651, 247 | 3 |
| ENSG000000821 | STRADB | chr2 | 201477618 | 201477790 | 201475618 | 201475742 | 201478562 | 201478601 | 63,54,65 | 0,0,0 | 50,64,56 | 7,5,2 | 300 | 150 | 1.04E-06 | 2.87E-05 | 1.0,1.0,1.0 | 0.781,0.865,0.93 | 0.14 | ENSG000000821 | [[[[201451740, 20 | 2 |
| ENSG000000821 | STRADB | chr2 | 201478562 | 201478601 | 201477618 | 201477790 | 201479488 | 201479531 | 10,14,21 | 0,0,0 | 13,13,16 | 2,6,0 | 189 | 150 | 0.00011065 | 0.001742002 | 1.0,1.0,1.0 | 0.838,0.632,1.0 | 0.177 | ENSG000000821 | [[[[201451740, 20 | 2 |
| ENSG000000742 | EED | chr11 | 86264171 | 86264263 | 86257514 | 86257596 | 86276979 | 86277138 | 19,20,37 | 0,0,1 | 15,37,32 | 1,8,1 | 242 | 150 | 0.00011631 | 0.001818346 | 1.0,1.0,0.958 | 0.903,0.741,0.95 | 0.121 | ENSG000000742 | [[[[86244753, 862 | 3 |
| ENSG000001174 | AKR1A1 | chr1 | 45566568 | 45566688 | 45550845 | 45551155 | 45567981 | 45568068 | 13,10,11 | 0,0,0 | 6,18,11 | 2,3,2 | 270 | 150 | 4.46E-05 | 0.000790883 | 1.0,1.0,1.0 | 0.625,0.769,0.75 | 0.284 | ENSG000001174 | [[[[45550826, 455 | 3 |
| ENSG000001174 | AKR1A1 | chr1 | 45566568 | 45566688 | 45561788 | 45561878 | 45567981 | 45568068 | 383,311,581 | 0,2,0 | 380,658,396 | 42,70,55 | 270 | 150 | 0 | 0 | 1.0,0.989,1.0 | 0.834,0.839,0.8 | 0.172 | ENSG000001174 | [[[[45550826, 455 | 2 |
| ENSG000001174 | AKR1A1 | chr1 | 45566868 | 45567020 | 45561788 | 45561878 | 45567981 | 45568068 | 253,296,488 | 0,2,0 | 305,688,442 | 42,70,55 | 300 | 150 | 0 | 0 | 1.0,0.987,1.0 | 0.784,0.831,0.80 | 0.19 | ENSG000001174 | [[[[45550826, 455 | 2 |
| ENSG000000570 | DCBLD2 | chr3 | 98808080 | 98808174 | 98801382 | 98801649 | 98817768 | 98817893 | 384,298,606 | 0,0,0 | 156,416,343 | 19,35,17 | 244 | 150 | 0 | 0 | 1.0,1.0,1.0 | 0.835,0.88,0.925 | 0.12 | ENSG000000570 | [[[[98901122, 989 | 4 |
| ENSG000000570 | DCBLD2 | chr3 | 98811467 | 98811554 | 98801382 | 98801649 | 98812331 | 98812482 | 283,201,559 | 0,0,0 | 137,315,266 | 1,8,9 | 237 | 150 | 0.000224451 | 0.003218157 | 1.0,1.0,1.0 | 0.989,0.961,0.94 | 0.034 | ENSG000000570 | [[[[98901122, 989 | 3 |
| ENSG000000570 | DCBLD2 | chr3 | 98812331 | 98812482 | 98801599 | 98801649 | 98817768 | 98817828 | 481,278,890 | 0,0,0 | 201,513,389 | 19,35,17 | 300 | 150 | 0 | 0 | 1.0,1.0,1.0 | 0.841,0.88,0.92 | 0.12 | ENSG000000570 | [[[[98901122, 989 | 4 |
| ENSG000001842 | CMSS1 | chr3 | 100167737 | 100167840 | 100162302 | 100162432 | 100171838 | 100171899 | 53,52,65 | 0,0,0 | 35,124,48 | 16,17,8 | 253 | 150 | 2.73E-12 | 2.32E-10 | 1.0,1.0,1.0 | 0.565,0.812,0.78 | 0.281 | ENSG000001842 | [[[[99817862, 998 | 2 |
| ENSG000001530 | CENPH | chr5 | 69202505 | 69202569 | 69195716 | 69195750 | 69202918 | 69202970 | 21,17,15 | 0,0,0 | 14,23,6 | 6,10,14 | 214 | 150 | 2.89E-11 | 2.05E-09 | 1.0,1.0,1.0 | 0.621,0.617,0.23 | 0.51 | ENSG000001530 | [[[[69189583, 691 | 2 |
| ENSG000000931 | LRRFIP2 | chr3 | 37065809 | 37065942 | 37063741 | 37063791 | 37075023 | 37075116 | 179,100,188 | 0,1,0 | 104,193,156 | 2,18,11 | 283 | 150 | 8.49E-10 | 4.78E-08 | 1.0,0.981,1.0 | 0.965,0.85,0.883 | 0.094 | ENSG000000931 | [[[[37174539, 371 | 3 |
| ENSG000000931 | LRRFIP2 | chr3 | 37072789 | 37072882 | 37065809 | 37065942 | 37075023 | 37075116 | 1,1,3 | 90,46,85 | 11,5,6 | 49,81,66 | 243 | 150 | 5.96E-05 | 0.001017715 | 0.007,0.013,0.02 | 0.122,0.037,0.05 | -0.057 | ENSG000000931 | [[[[37174539, 371 | 2 |
| ENSG000001627 | NCSTN | chr1 | 160344721 | 160344826 | 160343271 | 160343481 | 160350104 | 160350205 | 55,42,123 | 0,0,0 | 73,138,150 | 1,18,9 | 255 | 150 | 7.54E-08 | 2.73E-06 | 1.0,1.0,1.0 | 0.977,0.819,0.90 | 0.099 | ENSG000001627 | [[[[160343383, 16 | 3 |
| ENSG000001456 | OSMR | chr5 | 38924421 | 38924595 | 38918839 | 38919062 | 38925203 | 38925371 | 35,22,44 | 0,0,0 | 24,43,22 | 0,12,1 | 300 | 150 | 1.15E-05 | 0.000241968 | 1.0,1.0,1.0 | 1.0,0.642,0.917 | 0.147 | ENSG000001456 | [[[[38846012, 388 | 3 |
| ENSG000001624 | NOL9 | chr1 | 6541829 | 6541927 | 6533279 | 6533441 | 6549570 | 6549698 | 6,5,6 | 0,0,1 | 2,18,9 | 0,8,4 | 248 | 150 | 5.84E-05 | 0.000999195 | 1.0,1.0,0.784 | 1.0,0.576,0.576 | 0.211 | ENSG000001624 | [[[[6554107, 6554 | 3 |
| ENSG000001624 | NOL9 | chr1 | 6544825 | 6544922 | 6533279 | 6533441 | 6545044 | 6545180 | 11,9,9 | 0,0,0 | 17,22,10 | 3,3,2 | 247 | 150 | 6.73E-05 | 0.001130258 | 1.0,1.0,1.0 | 0.775,0.817,0.75 | 0.219 | ENSG000001624 | [[[[6554107, 6554 | 2 |
| ENSG000001624 | NOL9 | chr1 | 6545044 | 6545180 | 6533279 | 6533441 | 6549570 | 6549698 | 13,7,19 | 0,0,1 | 22,47,42 | 0,8,4 | 286 | 150 | 0.000316737 | 0.004328328 | 1.0,1.0,0.909 | 1.0,0.755,0.846 | 0.103 | ENSG000001624 | [[[[6554107, 6554 | 3 |
| ENSG000001391 | PEX5 | chr12 | 7202611 | 7202704 | 7201750 | 7201841 | 7203455 | 7203551 | 12,3,11 | 0,0,0 | 13,16,15 | 4,2,1 | 243 | 150 | 0.0001925 | 0.002821876 | 1.0,1.0,1.0 | 0.667,0.832,0.90 | 0.199 | ENSG000001391 | [[[[7189686, 7189 | 2 |
| ENSG000000722 | TFRC | chr3 | 196068031 | 196068130 | 196067517 | 196067657 | 196071395 | 196071498 | 153,125,305 | 0,0,0 | 65,174,167 | 4,19,11 | 249 | 150 | 2.22E-13 | 2.21E-11 | 1.0,1.0,1.0 | 0.907,0.847,0.90 | 0.115 | ENSG000000722 | [[[[196082043, 19 | 2 |
| ENSG000001754 | CCDC14 | chr3 | 123948895 | 123949132 | 123946802 | 123947319 | 123955842 | 123955965 | 20,15,18 | 0,0,0 | 17,41,21 | 2,3,2 | 300 | 150 | 0.000452913 | 0.005908418 | 1.0,1.0,1.0 | 0.81,0.872,0.84 | 0.159 | ENSG000001754 | [[[[123961144, 12 | 2 |
| ENSG000000688 | PSME4 | chr2 | 53901349 | 53901559 | 53899880 | 53900017 | 53906597 | 53906693 | 37,40,56 | 0,0,0 | 19,78,40 | 3,11,4 | 300 | 150 | 1.65E-08 | 7.03E-07 | 1.0,1.0,1.0 | 0.76,0.78,0.833 | 0.209 | ENSG000000688 | [[[[53970543, 539 | 2 |
| ENSG000000748 | ENO1 | chr1 | 8874823 | 8874914 | 8870451 | 8870510 | 8878579 | 8879250 | 34,51,103 | 0,0,0 | 45,92,70 | 8,16,16 | 241 | 150 | 7.25E-14 | 7.65E-12 | 1.0,1.0,1.0 | 0.778,0.782,0.73 | 0.236 | ENSG000000748 | [[[[8878580, 8878 | 2 |
| ENSG000001403 | WDR61 | chr15 | 78286040 | 78286133 | 78278207 | 78278301 | 78289618 | 78289712 | 0,0,1 | 3,2,2 | 7,25,2 | 4,7,1 | 243 | 150 | 1.80E-07 | 5.92E-06 | 0.0,0.0,0.236 | 0.519,0.688,0.55 | -0.508 | ENSG000001403 | [[[[78299562, 782 | 4 |
| ENSG000001403 | WDR61 | chr15 | 78292615 | 78292788 | 78289950 | 78290099 | 78294945 | 78294973 | 324,257,356 | 0,2,0 | 226,345,233 | 37,44,15 | 300 | 150 | 0 | 0 | 1.0,0.985,1.0 | 0.753,0.797,0.88 | 0.183 | ENSG000001403 | [[[[78299562, 782 | 2 |
| ENSG000001403 | WDR61 | chr15 | 78292615 | 78292885 | 78289950 | 78290099 | 78294945 | 78294973 | 324,257,356 | 0,2,0 | 232,334,227 | 37,44,15 | 300 | 150 | 0 | 0 | 1.0,0.985,1.0 | 0.758,0.791,0.88 | 0.184 | ENSG000001403 | [[[[78299562, 782 | 2 |
| ENSG000001403 | WDR61 | chr15 | 78293166 | 78293272 | 78289950 | 78290099 | 78294945 | 78294973 | 270,213,442 | 0,2,0 | 228,418,238 | 37,44,15 | 256 | 150 | 0 | 0 | 1.0,0.984,1.0 | 0.783,0.848,0.90 | 0.15 | ENSG000001403 | [[[[78299562, 782 | 2 |
| ENSG000001696 | ASPSCR1 | chr17 | 82011542 | 82011605 | 82009485 | 82009567 | 82012230 | 82012283 | 2,10,20 | 0,0,0 | 10,27,15 | 6,2,1 | 213 | 150 | 2.83E-05 | 0.000531448 | 1.0,1.0,1.0 | 0.54,0.905,0.914 | 0.214 | ENSG000001696 | [[[[81977629, 819 | 2 |
| ENSG000000852 | MYNN | chr3 | 169782464 | 169782643 | 169778767 | 169779561 | 169783476 | 169783560 | 36,16,29 | 0,0,0 | 29,54,26 | 3,10,3 | 300 | 150 | 1.40E-07 | 4.71E-06 | 1.0,1.0,1.0 | 0.829,0.73,0.813 | 0.209 | ENSG000000852 | [[[[169773396, 16 | 2 |
| ENSG000000852 | MYNN | chr3 | 169782464 | 169782643 | 169778767 | 169779561 | 169784621 | 169784676 | 4,8,5 | 0,0,0 | 12,29,8 | 5,10,5 | 300 | 150 | 3.06E-08 | 1.22E-06 | 1.0,1.0,1.0 | 0.545,0.592,0.44 | 0.473 | ENSG000000852 | [[[[169773396, 16 | 3 |
| ENSG000000852 | MYNN | chr3 | 169783476 | 169783560 | 169778767 | 169779561 | 169784621 | 169784708 | 15,20,22 | 0,0,0 | 16,45,28 | 5,10,5 | 234 | 150 | 2.90E-08 | 1.17E-06 | 1.0,1.0,1.0 | 0.672,0.743,0.78 | 0.268 | ENSG000000852 | [[[[169773396, 16 | 3 |
| ENSG000001631 | ERCC3 | chr2 | 127261227 | 127261346 | 127259295 | 127259448 | 127272864 | 127272961 | 73,70,74 | 0,0,0 | 64,117,68 | 4,4,4 | 269 | 150 | 0.000105981 | 0.001677854 | 1.0,1.0,1.0 | 0.899,0.942,0.90 | 0.085 | ENSG000001631 | [[[[127294054, 12 | 2 |
| ENSG000001631 | ERCC3 | chr2 | 127271335 | 127271453 | 127259295 | 127259448 | 127272864 | 127272961 | 55,55,54 | 0,0,0 | 23,24,16 | 4,4,4 | 268 | 150 | 2.01E-08 | 8.40E-07 | 1.0,1.0,1.0 | 0.763,0.771,0.69 | 0.258 | ENSG000001631 | [[[[127294054, 12 | 2 |
| ENSG000001093 | NFKB1 | chr4 | 102612433 | 102612606 | 102610574 | 102610699 | 102613424 | 102613581 | 26,51,38 | 0,0,0 | 47,69,51 | 0,20,8 | 300 | 150 | 1.33E-07 | 4.50E-06 | 1.0,1.0,1.0 | 1.0,0.633,0.761 | 0.202 | ENSG000001093 | [[[[102501359, 10 | 2 |
| ENSG000001093 | NFKB1 | chr4 | 102613424 | 102613581 | 102610574 | 102610699 | 102616433 | 102616915 | 32,29,66 | 0,0,0 | 28,96,76 | 2,5,3 | 300 | 150 | 0.000368889 | 0.004938269 | 1.0,1.0,1.0 | 0.875,0.906,0.92 | 0.097 | ENSG000001093 | [[[[102501359, 10 | 3 |
| ENSG000000172 | ATP2C1 | chr3 | 130999517 | 130999659 | 130998293 | 130998389 | 131016151 | 131016684 | 73,64,95 | 0,3,0 | 46,96,49 | 12,29,20 | 292 | 150 | 0 | 0 | 1.0,0.916,1.0 | 0.663,0.63,0.557 | 0.355 | ENSG000000172 | [[[[130894084, 13 | 2 |
| ENSG000001637 | TOPBP1 | chr3 | 133649779 | 133649943 | 133649382 | 133649633 | 133653344 | 133653524 | 30,23,72 | 0,0,0 | 14,41,55 | 8,8,4 | 300 | 150 | 7.35E-09 | 3.37E-07 | 1.0,1.0,1.0 | 0.467,0.719,0.87 | 0.314 | ENSG000001637 | [[[[133661769, 13 | 2 |
| ENSG000001445 | GMPPA | chr2 | 219500120 | 219500218 | 219498866 | 219498938 | 219501475 | 219501579 | 78,80,133 | 3,1,1 | 87,157,110 | 6,13,3 | 248 | 150 | 0.000669913 | 0.008246533 | 0.94,0.98,0.988 | 0.898,0.88,0.957 | 0.058 | ENSG000001445 | [[[[219498891, 21 | 2 |
| ENSG000001445 | GMPPA | chr2 | 219501475 | 219501579 | 219498866 | 219498938 | 219501850 | 219502037 | 69,67,105 | 0,0,0 | 70,150,110 | 3,20,11 | 254 | 150 | 2.80E-10 | 1.71E-08 | 1.0,1.0,1.0 | 0.932,0.816,0.85 | 0.132 | ENSG000001445 | [[[[219498891, 21 | 3 |
| ENSG000000803 | RIF1 | chr2 | 151436826 | 151437003 | 151433076 | 151433228 | 151437240 | 151437351 | 11,9,21 | 0,0,0 | 16,26,22 | 5,1,1 | 300 | 150 | 0.000460896 | 0.006000226 | 1.0,1.0,1.0 | 0.615,0.929,0.91 | 0.18 | ENSG000000803 | [[[[151409902, 15 | 2 |
| ENSG000001742 | PRPF8 | chr17 | 1661270 | 1661406 | 1660697 | 1660827 | 1661905 | 1662038 | 1,1,2 | 0,0,0 | 2,4,5 | 1,14,8 | 286 | 150 | 1.04E-10 | 6.84E-09 | 1.0,1.0,1.0 | 0.512,0.13,0.247 | 0.704 | ENSG000001742 | [[[[1684780, 1684 | 3 |
| ENSG000001875 | USP7 | chr16 | 8896998 | 8897099 | 8895641 | 8895741 | 8899120 | 8899188 | 76,63,111 | 0,0,0 | 61,122,67 | 3,12,8 | 251 | 150 | 5.65E-09 | 2.66E-07 | 1.0,1.0,1.0 | 0.924,0.859,0.83 | 0.128 | ENSG000001875 | [[[[8963207, 8963 | 3 |
| ENSG000001245 | WRNIP1 | chr6 | 2770119 | 2770361 | 2765413 | 2766444 | 2779262 | 2779492 | 93,126,211 | 1,2,5 | 91,167,89 | 4,15,9 | 300 | 150 | 3.15E-05 | 0.000584399 | 0.979,0.969,0.95 | 0.919,0.848,0.83 | 0.101 | ENSG000001245 | [[[[2765393, 2766 | 2 |
| ENSG000001245 | WRNIP1 | chr6 | 2770194 | 2770361 | 2765413 | 2766444 | 2779262 | 2779492 | 93,125,211 | 1,2,5 | 91,165,91 | 4,15,9 | 300 | 150 | 3.16E-05 | 0.000586044 | 0.979,0.969,0.95 | 0.919,0.846,0.83 | 0.101 | ENSG000001245 | [[[[2765393, 2766 | 2 |
| ENSG000001399 | SLC38A6 | chr14 | 61037624 | 61037683 | 61030444 | 61030523 | 61043146 | 61043208 | 3,3,5 | 0,0,0 | 6,4,6 | 0,2,4 | 209 | 150 | 0.000410458 | 0.005419556 | 1.0,1.0,1.0 | 1.0,0.589,0.518 | 0.298 | ENSG000001399 | [[[[60981245, 609 | 2 |
| ENSG000001399 | SLC38A6 | chr14 | 61043449 | 61043503 | 61030444 | 61030523 | 61045345 | 61045425 | 6,10,11 | 0,0,0 | 2,7,7 | 0,8,1 | 204 | 150 | 9.55E-05 | 0.001534067 | 1.0,1.0,1.0 | 1.0,0.391,0.837 | 0.257 | ENSG000001399 | [[[[60981245, 609 | 4 |
| ENSG000001393 | POC1B | chr12 | 89467616 | 89467685 | 89466672 | 89466922 | 89491935 | 89492115 | 6,3,14 | 0,0,0 | 10,19,7 | 2,4,1 | 219 | 150 | 0.000142933 | 0.00216913 | 1.0,1.0,1.0 | 0.774,0.765,0.82 | 0.211 | ENSG000001393 | [[[[89525881, 895 | 4 |
| ENSG000001058 | DNAJC2 | chr7 | 103327331 | 103327378 | 103323597 | 103323625 | 103327655 | 103327754 | 1,3,1 | 0,0,0 | 5,17,15 | 0,5,2 | 197 | 150 | 0.000422637 | 0.005554341 | 1.0,1.0,1.0 | 1.0,0.721,0.851 | 0.143 | ENSG000001058 | [[[[103344559, 10 | 2 |
| ENSG000001245 | MRS2 | chr6 | 24418083 | 24418236 | 24412221 | 24412395 | 24418460 | 24418561 | 30,20,36 | 0,0,0 | 59,73,65 | 13,19,21 | 300 | 150 | 2.17E-12 | 1.88E-10 | 1.0,1.0,1.0 | 0.694,0.658,0.60 | 0.347 | ENSG000001245 | [[[[24402936, 244 | 3 |
| ENSG000000073 | ST7L | chr1 | 112550600 | 112550693 | 112523517 | 112526111 | 112555867 | 112556018 | 1,1,2 | 0,0,0 | 5,13,5 | 4,2,0 | 243 | 150 | 0.000817358 | 0.009781847 | 1.0,1.0,1.0 | 0.436,0.8,1.0 | 0.255 | ENSG000000073 | [[[[112618909, 11 | 2 |
| ENSG000001688 | GFM1 | chr3 | 158681994 | 158682157 | 158662627 | 158662684 | 158684523 | 158684668 | 63,49,98 | 0,0,0 | 61,70,68 | 6,9,0 | 300 | 150 | 1.55E-06 | 4.08E-05 | 1.0,1.0,1.0 | 0.836,0.795,1.0 | 0.123 | ENSG000001688 | [[[[158644527, 15 | 3 |
| ENSG000001315 | NDFIP1 | chr5 | 142135729 | 142135817 | 142132211 | 142132342 | 142140562 | 142140629 | 130,83,181 | 2,4,3 | 90,203,130 | 41,121,55 | 238 | 150 | 0 | 0 | 0.976,0.929,0.97 | 0.58,0.514,0.598 | 0.396 | ENSG000001315 | [[[[142108779, 14 | 2 |
| ENSG000001315 | NDFIP1 | chr5 | 142137733 | 142137858 | 142132211 | 142132342 | 142140562 | 142140629 | 240,188,211 | 2,4,3 | 138,263,154 | 41,121,55 | 275 | 150 | 0 | 0 | 0.985,0.962,0.97 | 0.647,0.542,0.60 | 0.376 | ENSG000001315 | [[[[142108779, 14 | 2 |
| ENSG000001315 | NDFIP1 | chr5 | 142140562 | 142140629 | 142132211 | 142132342 | 142144570 | 142144676 | 34,32,65 | 0,0,0 | 87,172,95 | 27,23,17 | 217 | 150 | 3.65E-12 | 3.06E-10 | 1.0,1.0,1.0 | 0.69,0.838,0.794 | 0.226 | ENSG000001315 | [[[[142108779, 14 | 3 |
| ENSG000000908 | AARS | chr16 | 70276965 | 70277154 | 70271780 | 70271972 | 70282619 | 70282784 | 283,203,439 | 0,0,0 | 276,677,466 | 10,15,16 | 300 | 150 | 7.03E-09 | 3.25E-07 | 1.0,1.0,1.0 | 0.932,0.958,0.93 | 0.058 | ENSG000000908 | [[[[70289421, 702 | 2 |
| ENSG000001738 | PHC3 | chr3 | 170171372 | 170171450 | 170147247 | 170149244 | 170178772 | 170178938 | 11,3,12 | 0,0,0 | 4,16,20 | 3,8,7 | 228 | 150 | 8.50E-09 | 3.83E-07 | 1.0,1.0,1.0 | 0.467,0.568,0.65 | 0.437 | ENSG000001738 | [[[[170181702, 17 | 2 |
| ENSG000000435 | TRIT1 | chr1 | 39847219 | 39847297 | 39844530 | 39844640 | 39883317 | 39883480 | 33,21,28 | 0,0,0 | 13,44,16 | 3,4,1 | 228 | 150 | 0.000234444 | 0.003333484 | 1.0,1.0,1.0 | 0.74,0.879,0.913 | 0.156 | ENSG000000435 | [[[[39883318, 398 | 7 |
| ENSG000001365 | MRPL47 | chr3 | 179593764 | 179593895 | 179588284 | 179588995 | 179601729 | 179601790 | 4,5,5 | 0,0,1 | 11,15,9 | 6,5,1 | 281 | 150 | 5.84E-05 | 0.000999607 | 1.0,1.0,0.727 | 0.495,0.616,0.82 | 0.263 | ENSG000001365 | [[[[179604527, 17 | 3 |
| ENSG000001671 | CERCAM | chr9 | 128428929 | 128429036 | 128428301 | 128428421 | 128431170 | 128431303 | 19,26,33 | 0,0,0 | 17,41,31 | 4,2,4 | 257 | 150 | 1.28E-05 | 0.000265425 | 1.0,1.0,1.0 | 0.713,0.923,0.81 | 0.182 | ENSG000001671 | [[[[128420857, 12 | 2 |
| ENSG000001224 | RPL5 | chr1 | 92833388 | 92833458 | 92831982 | 92832117 | 92836189 | 92836354 | 4,4,12 | 0,0,0 | 239,480,257 | 63,98,65 | 220 | 150 | 4.46E-07 | 1.34E-05 | 1.0,1.0,1.0 | 0.721,0.77,0.729 | 0.26 | ENSG000001224 | [[[[92832040, 928 | 3 |
| ENSG000001224 | RPL5 | chr1 | 92833388 | 92833458 | 92831982 | 92832117 | 92837455 | 92837469 | 4,4,12 | 0,0,0 | 5,10,8 | 2,7,1 | 220 | 150 | 2.22E-07 | 7.14E-06 | 1.0,1.0,1.0 | 0.63,0.493,0.845 | 0.344 | ENSG000001224 | [[[[92832040, 928 | 4 |
| ENSG000001224 | RPL5 | chr1 | 92833388 | 92833458 | 92831982 | 92832117 | 92840550 | 92840639 | 4,4,12 | 0,0,0 | 7,22,12 | 13,33,11 | 220 | 150 | 4.31E-12 | 3.58E-10 | 1.0,1.0,1.0 | 0.269,0.313,0.42 | 0.664 | ENSG000001224 | [[[[92832040, 928 | 5 |
| ENSG000001224 | RPL5 | chr1 | 92833544 | 92833660 | 92832012 | 92832117 | 92840550 | 92840639 | 3,4,2 | 0,0,0 | 7,11,5 | 13,33,11 | 266 | 150 | 5.04E-10 | 2.96E-08 | 1.0,1.0,1.0 | 0.233,0.158,0.20 | 0.802 | ENSG000001224 | [[[[92832040, 928 | 5 |
| ENSG000001224 | RPL5 | chr1 | 92833544 | 92833660 | 92833388 | 92833458 | 92836189 | 92836354 | 325,833,154,871 | 0,0,0 | 314,466,264,158 | 234,479,254 | 266 | 150 | 0 | 0 | 1.0,1.0,1.0 | 0.883,0.886,0.90 | 0.11 | ENSG000001224 | [[[[92832040, 928 | 2 |
| ENSG000001224 | RPL5 | chr1 | 92834778 | 92834913 | 92831982 | 92832117 | 92836189 | 92836271 | 254,621,693,186 | 0,0,0 | 212,343,262,554 | 63,98,65 | 285 | 150 | 0 | 0 | 1.0,1.0,1.0 | 0.947,0.959,0.95 | 0.047 | ENSG000001224 | [[[[92832040, 928 | 3 |
| ENSG000001224 | RPL5 | chr1 | 92834778 | 92834913 | 92831982 | 92832117 | 92840550 | 92840639 | 4,3,5 | 0,0,0 | 27,50,32 | 13,33,11 | 285 | 150 | 9.81E-08 | 3.45E-06 | 1.0,1.0,1.0 | 0.522,0.444,0.60 | 0.476 | ENSG000001224 | [[[[92832040, 928 | 5 |

| | | | | | | | | | | | | | | | | | | | | | | |
|---|---|---|---|---|---|---|---|---|---|---|---|---|---|---|---|---|---|---|---|---|---|---|
| ENSG00000122 | RPL5 | chr1 | 92834778 | 92834913 | 92833388 | 92833458 | 92836189 | 92836271 | 254,621,693,186 | 0,0,0 | 212,343,272,554 | 234,479,254 | 285 | 150 | 0 | 0 | 1.0,1.0,1.0 | 0.827,0.826,0.84 | 0.169 | ENSG00000122 | [[[[92832040, 928 | 2 |
| ENSG00000122 | RPL5 | chr1 | 92834778 | 92834913 | 92833388 | 92833458 | 92840550 | 92840639 | 4,3,5 | 0,0,0 | 27,51,32 | 2,19,8 | 285 | 150 | 2.29E-06 | 5.78E-05 | 1.0,1.0,1.0 | 0.877,0.586,0.67 | 0.286 | ENSG00000122 | [[[[92832040, 928 | 4 |
| ENSG00000122 | RPL5 | chr1 | 92837455 | 92837633 | 92834778 | 92834913 | 92840550 | 92840639 | 221,220,992,537 | 5,3,5 | 234,045,652,584 | 31,64,35 | 300 | 150 | 1.35E-11 | 1.01E-09 | 0.995,0.997,0.99 | 0.974,0.973,0.97 | 0.022 | ENSG00000122 | [[[[92832040, 928 | 2 |
| ENSG00000175 | PARL | chr3 | 183842297 | 183842384 | 183840569 | 183840640 | 183862752 | 183862801 | 52,34,45 | 0,0,0 | 66,106,66 | 1,14,6 | 237 | 150 | 1.15E-06 | 3.11E-05 | 1.0,1.0,1.0 | 0.977,0.827,0.87 | 0.107 | ENSG00000175 | [[[[183884722, 18 | 2 |
| ENSG00000073 | PDE8A | chr15 | 85117640 | 85117839 | 85115983 | 85116119 | 85126206 | 85126374 | 49,38,80 | 0,0,0 | 34,73,43 | 4,3,3 | 300 | 150 | 6.03E-05 | 0.001027442 | 1.0,1.0,1.0 | 0.81,0.924,0.878 | 0.129 | ENSG00000073 | [[[[84981856, 84 | 3 |
| ENSG00000161 | DVL3 | chr3 | 184166445 | 184166522 | 184164795 | 184164931 | 184166605 | 184166673 | 18,52,69 | 0,0,0 | 26,55,56 | 2,5,2 | 227 | 150 | 0.000519229 | 0.006636059 | 1.0,1.0,1.0 | 0.896,0.879,0.94 | 0.092 | ENSG00000161 | [[[[184155377, 18 | 4 |
| ENSG00000163 | SENP2 | chr3 | 185614563 | 185614740 | 185613344 | 185613408 | 185621825 | 185621905 | 37,18,53 | 0,0,0 | 39,115,70 | 0,6,4 | 300 | 150 | 0.000710134 | 0.008667433 | 1.0,1.0,1.0 | 1.0,0.906,0.897 | 0.066 | ENSG00000163 | [[[[185586295, 18 | 3 |
| ENSG00000163 | SENP2 | chr3 | 185617479 | 185617611 | 185614502 | 185614740 | 185621825 | 185621905 | 34,25,43 | 0,0,0 | 27,50,27 | 3,8,5 | 282 | 150 | 4.09E-08 | 1.58E-06 | 1.0,1.0,1.0 | 0.827,0.769,0.74 | 0.221 | ENSG00000163 | [[[[185586295, 18 | 2 |
| ENSG00000163 | SENP2 | chr3 | 185619298 | 185619502 | 185613344 | 185613408 | 185621825 | 185621905 | 25,13,26 | 0,0,0 | 12,59,36 | 0,6,4 | 300 | 150 | 3.41E-05 | 0.00062645 | 1.0,1.0,1.0 | 1.0,0.831,0.818 | 0.117 | ENSG00000163 | [[[[185586295, 18 | 3 |
| ENSG00000163 | SENP2 | chr3 | 185619298 | 185619502 | 185614563 | 185614740 | 185621825 | 185621905 | 25,14,27 | 0,0,0 | 14,69,38 | 3,8,5 | 300 | 150 | 5.70E-07 | 1.66E-05 | 1.0,1.0,1.0 | 0.7,0.812,0.792 | 0.232 | ENSG00000163 | [[[[185586295, 18 | 2 |
| ENSG00000136 | TRA2B | chr3 | 185923795 | 185923984 | 185922010 | 185922126 | 185931576 | 185931852 | 466,360,593 | 0,0,0 | 5,531,079,626 | 37,92,50 | 300 | 150 | 0 | 0 | 1.0,1.0,1.0 | 0.882,0.854,0.86 | 0.134 | ENSG00000136 | [[[[185937825, 18 | 3 |
| ENSG00000136 | TRA2B | chr3 | 185923795 | 185923984 | 185922010 | 185922126 | 185937824 | 185938029 | 466,360,594 | 0,0,0 | 5,671,105,638 | 172,332,190 | 300 | 150 | 0 | 0 | 1.0,1.0,1.0 | 0.622,0.625,0.62 | 0.375 | ENSG00000136 | [[[[185937825, 18 | 3 |
| ENSG00000136 | TRA2B | chr3 | 185925463 | 185925626 | 185922010 | 185922126 | 185937824 | 185937998 | 17,14,28 | 0,0,0 | 23,56,27 | 172,332,190 | 300 | 150 | 0 | 0 | 1.0,1.0,1.0 | 0.063,0.078,0.06 | 0.931 | ENSG00000136 | [[[[185937825, 18 | 3 |
| ENSG00000136 | TRA2B | chr3 | 185925463 | 185925626 | 185922010 | 185923809 | 185937824 | 185937998 | 17,14,27 | 0,0,0 | 23,54,30 | 15,44,19 | 300 | 150 | 1.11E-16 | 1.65E-14 | 1.0,1.0,1.0 | 0.434,0.38,0.441 | 0.582 | ENSG00000136 | [[[[185937825, 18 | 2 |
| ENSG00000136 | TRA2B | chr3 | 185925463 | 185925626 | 185923795 | 185923984 | 185937824 | 185937998 | 453,399,651 | 0,0,1 | 5,571,223,648 | 15,26,12 | 300 | 150 | 6.85E-09 | 3.17E-07 | 1.0,1.0,0.997 | 0.949,0.959,0.96 | 0.042 | ENSG00000136 | [[[[185937825, 18 | 2 |
| ENSG00000136 | TRA2B | chr3 | 185931576 | 185931852 | 185922010 | 185922126 | 185937824 | 185938029 | 12,33,31 | 0,0,0 | 164,307,223 | 172,332,190 | 300 | 150 | 0 | 0 | 1.0,1.0,1.0 | 0.323,0.316,0.37 | 0.664 | ENSG00000136 | [[[[185937825, 18 | 3 |
| ENSG00000136 | TRA2B | chr3 | 185931576 | 185931852 | 185922010 | 185923809 | 185937824 | 185938029 | 12,33,31 | 0,0,0 | 143,240,185 | 15,44,19 | 300 | 150 | 1.36E-08 | 5.92E-07 | 1.0,1.0,1.0 | 0.827,0.732,0.83 | 0.204 | ENSG00000136 | [[[[185937825, 18 | 2 |
| ENSG00000136 | TRA2B | chr3 | 185931576 | 185931852 | 185923795 | 185923984 | 185937824 | 185938029 | 12,33,31 | 0,0,1 | 128,215,173 | 15,26,12 | 300 | 150 | 2.80E-06 | 6.89E-05 | 1.0,1.0,0.939 | 0.81,0.805,0.878 | 0.149 | ENSG00000136 | [[[[185937825, 18 | 2 |
| ENSG00000204 | MSH5 | chr6 | 31761865 | 31761955 | 31760089 | 31760216 | 31762111 | 31762185 | 5,9,16 | 0,0,0 | 4,21,7 | 0,6,4 | 240 | 150 | 3.37E-06 | 8.15E-05 | 1.0,1.0,1.0 | 1.0,0.686,0.522 | 0.264 | ENSG00000204 | [[[[31740005, 317 | 4 |
| ENSG00000136 | SUCLA2 | chr13 | 47948939 | 47949028 | 47942655 | 47943445 | 47954139 | 47954282 | 53,36,53 | 0,0,0 | 36,63,32 | 12,12,15 | 239 | 150 | 2.62E-14 | 2.96E-12 | 1.0,1.0,1.0 | 0.653,0.767,0.57 | 0.336 | ENSG00000136 | [[[[48001180, 480 | 2 |
| ENSG00000118 | WDR35 | chr2 | 19960553 | 19960614 | 19953833 | 19953978 | 19973562 | 19973708 | 3,3,2 | 0,0,0 | 5,3,7 | 1,7,0 | 211 | 150 | 0.000661341 | 0.008156821 | 1.0,1.0,1.0 | 0.78,0.234,1.0 | 0.329 | ENSG00000118 | [[[[19989992, 19 | 4 |
| ENSG00000128 | MTX2 | chr2 | 176326824 | 176326901 | 176297848 | 176297895 | 176328292 | 176328322 | 51,66,66 | 0,0,0 | 48,97,67 | 5,22,7 | 227 | 150 | 2.79E-11 | 1.98E-09 | 1.0,1.0,1.0 | 0.864,0.744,0.86 | 0.176 | ENSG00000128 | [[[[176269442, 17 | 2 |
| ENSG00000041 | PSMA4 | chr15 | 78542482 | 78542645 | 78542176 | 78542219 | 78548789 | 78548820 | 110,710,991,504 | 0,0,0 | 136,724,741,567 | 14,37,19 | 300 | 150 | 1.01E-07 | 3.55E-06 | 1.0,1.0,1.0 | 0.98,0.971,0.976 | 0.024 | ENSG00000041 | [[[[78540444, 78 | 5 |
| ENSG00000041 | PSMA4 | chr15 | 78542575 | 78542645 | 78542176 | 78542219 | 78545633 | 78545764 | 9,6,8 | 0,0,0 | 6,19,4 | 13,7,6 | 220 | 150 | 3.04E-10 | 1.85E-08 | 1.0,1.0,1.0 | 0.239,0.649,0.31 | 0.6 | ENSG00000041 | [[[[78540444, 78 | 3 |
| ENSG00000041 | PSMA4 | chr15 | 78542575 | 78542645 | 78542176 | 78542219 | 78548789 | 78549037 | 9,6,8 | 0,0,0 | 10,23,10 | 14,37,19 | 220 | 150 | 4.23E-14 | 4.63E-12 | 1.0,1.0,1.0 | 0.328,0.298,0.26 | 0.703 | ENSG00000041 | [[[[78540444, 78 | 5 |
| ENSG00000041 | PSMA4 | chr15 | 78544189 | 78544267 | 78540443 | 78542645 | 78548789 | 78549176 | 309,231,289 | 0,0,0 | 295,500,267 | 10,15,10 | 228 | 150 | 6.59E-08 | 2.43E-06 | 1.0,1.0,1.0 | 0.951,0.956,0.94 | 0.049 | ENSG00000041 | [[[[78540444, 78 | 4 |
| ENSG00000041 | PSMA4 | chr15 | 78544189 | 78544267 | 78540460 | 78542219 | 78545633 | 78545764 | 11,13,16 | 0,0,0 | 23,50,33 | 13,7,6 | 228 | 150 | 1.18E-07 | 4.06E-06 | 1.0,1.0,1.0 | 0.538,0.825,0.78 | 0.285 | ENSG00000041 | [[[[78540444, 78 | 3 |
| ENSG00000041 | PSMA4 | chr15 | 78544189 | 78544267 | 78540460 | 78542219 | 78548789 | 78549176 | 11,13,16 | 0,0,0 | 24,48,32 | 14,37,19 | 228 | 150 | 2.93E-13 | 2.87E-11 | 1.0,1.0,1.0 | 0.53,0.46,0.526 | 0.495 | ENSG00000041 | [[[[78540444, 78 | 5 |
| ENSG00000041 | PSMA4 | chr15 | 78545633 | 78545764 | 78540443 | 78542645 | 78548789 | 78548970 | 14,9,15 | 0,0,0 | 16,30,20 | 10,15,10 | 281 | 150 | 2.71E-11 | 1.93E-09 | 1.0,1.0,1.0 | 0.461,0.516,0.51 | 0.502 | ENSG00000041 | [[[[78540444, 78 | 4 |
| ENSG00000041 | PSMA4 | chr15 | 78545633 | 78545764 | 78540460 | 78542219 | 78548789 | 78548970 | 14,9,15 | 0,0,0 | 28,35,26 | 14,37,19 | 281 | 150 | 3.19E-13 | 3.10E-11 | 1.0,1.0,1.0 | 0.516,0.336,0.42 | 0.575 | ENSG00000041 | [[[[78540444, 78 | 5 |
| ENSG00000041 | PSMA4 | chr15 | 78546574 | 78546698 | 78540443 | 78542645 | 78548789 | 78548820 | 395,413,573 | 0,0,0 | 468,997,547 | 10,15,10 | 274 | 150 | 1.02E-05 | 0.00021776 | 1.0,1.0,1.0 | 0.962,0.973,0.96 | 0.032 | ENSG00000041 | [[[[78540444, 78 | 4 |
| ENSG00000169 | PLEKHA2 | chr8 | 38943788 | 38943837 | 38935993 | 38936050 | 38950849 | 38950990 | 8,5,21 | 1,0,0 | 7,9,11 | 8,9,11 | 199 | 150 | 5.24E-11 | 3.58E-09 | 0.858,1.0,1.0 | 0.397,0.43,0.43 | 0.534 | ENSG00000169 | [[[38901346, 38 | 2 |
| ENSG00000169 | PLEKHA2 | chr8 | 38946123 | 38946221 | 38935993 | 38936050 | 38950849 | 38950990 | 20,14,68 | 1,0,0 | 27,87,68 | 8,9,11 | 248 | 150 | 9.90E-08 | 3.48E-06 | 0.924,1.0,1.0 | 0.671,0.854,0.78 | 0.203 | ENSG00000169 | [[[38901346, 38 | 2 |
| ENSG00000118 | FBXL5 | chr4 | 15630665 | 15630791 | 15604538 | 15605799 | 15636493 | 15636676 | 40,25,82 | 0,0,0 | 38,112,62 | 1,5,5 | 276 | 150 | 0.000228926 | 0.003269457 | 1.0,1.0,1.0 | 0.954,0.924,0.87 | 0.084 | ENSG00000118 | [[[[15655204, 15 | 5 |
| ENSG00000197 | ENTPD4 | chr8 | 23441568 | 23441723 | 23439748 | 23439915 | 23443849 | 23443953 | 26,31,42 | 0,0,0 | 16,54,37 | 2,6,2 | 300 | 150 | 1.68E-05 | 0.000335648 | 1.0,1.0,1.0 | 0.8,0.818,0.902 | 0.16 | ENSG00000197 | [[[[23457557, 23 | 2 |
| ENSG00000197 | ENTPD4 | chr8 | 23441592 | 23441723 | 23439748 | 23439915 | 23443849 | 23443953 | 10,6,11 | 0,0,0 | 14,22,16 | 2,6,2 | 281 | 150 | 1.84E-06 | 4.74E-05 | 1.0,1.0,1.0 | 0.789,0.662,0.81 | 0.246 | ENSG00000197 | [[[[23457557, 23 | 2 |
| ENSG00000163 | PACRGL | chr4 | 20707802 | 20707870 | 20704659 | 20704814 | 20713431 | 20713521 | 8,4,10 | 0,5,6 | 8,24,15 | 0,1,0 | 218 | 150 | 9.30E-06 | 0.000200765 | 1.0,0.355,0.534 | 1.0,0.943,1.0 | -0.351 | ENSG00000163 | [[[[20700458, 20 | 3 |
| ENSG00000163 | PACRGL | chr4 | 20712787 | 20712922 | 20704659 | 20704814 | 20713431 | 20713516 | 17,19,29 | 0,5,6 | 25,29,27 | 0,1,0 | 285 | 150 | 1.71E-05 | 0.000342868 | 1.0,0.667,0.718 | 1.0,0.939,1.0 | -0.185 | ENSG00000163 | [[[[20700458, 20 | 3 |
| ENSG00000109 | DHX15 | chr4 | 24532863 | 24533054 | 24529600 | 24529770 | 24540107 | 24540299 | 185,192,265 | 0,0,0 | 99,249,136 | 9,10,2 | 300 | 150 | 8.10E-08 | 2.91E-06 | 1.0,1.0,1.0 | 0.846,0.926,0.97 | 0.086 | ENSG00000109 | [[[[24584323, 24 | 2 |
| ENSG00000166 | MARS | chr12 | 57515919 | 57515991 | 57514953 | 57515014 | 57516244 | 57516337 | 114,100,141 | 0,0,0 | 156,346,144 | 21,47,41 | 222 | 150 | 0 | 0 | 1.0,1.0,1.0 | 0.834,0.833,0.70 | 0.21 | ENSG00000166 | [[[[57488068, 57 | 2 |
| ENSG00000136 | SCYL2 | chr12 | 100298030 | 100298175 | 100267139 | 100267396 | 100311043 | 100311193 | 21,17,38 | 0,0,0 | 22,36,39 | 2,7,4 | 295 | 150 | 7.97E-07 | 2.24E-05 | 1.0,1.0,1.0 | 0.848,0.723,0.83 | 0.199 | ENSG00000136 | [[[[100267177, 10 | 3 |
| ENSG00000108 | KPNB1 | chr17 | 47676408 | 47676491 | 47674637 | 47674782 | 47680019 | 47680134 | 274,194,377 | 0,0,0 | 104,329,208 | 8,7,2 | 233 | 150 | 0.000244153 | 0.003449652 | 1.0,1.0,1.0 | 0.893,0.968,0.98 | 0.051 | ENSG00000108 | [[[[47649919, 47 | 4 |
| ENSG00000108 | KPNB1 | chr17 | 47678307 | 47678413 | 47674637 | 47674782 | 47680019 | 47680134 | 343,289,399 | 0,0,0 | 93,278,159 | 8,7,2 | 256 | 150 | 1.38E-05 | 0.000283387 | 1.0,1.0,1.0 | 0.872,0.959,0.97 | 0.063 | ENSG00000108 | [[[[47649919, 47 | 4 |
| ENSG00000065 | TBC1D1 | chr4 | 38020590 | 38020695 | 38014508 | 38014973 | 38021585 | 38021718 | 7,6,38 | 0,0,0 | 11,17,12 | 2,5,7 | 255 | 150 | 6.10E-09 | 2.84E-07 | 1.0,1.0,1.0 | 0.764,0.667,0.50 | 0.356 | ENSG00000065 | [[[[37891084, 37 | 2 |
| ENSG00000148 | SLC30A9 | chr4 | 42078211 | 42078325 | 42070525 | 42070691 | 42086081 | 42086369 | 33,24,50 | 0,1,0 | 9,25,18 | 0,5,2 | 264 | 150 | 0.000597769 | 0.007486666 | 1.0,0.932,1.0 | 1.0,0.74,0.836 | 0.119 | ENSG00000148 | [[[[41990530, 41 | 2 |
| ENSG00000060 | WNK1 | chr12 | 900475 | 900670 | 890452 | 890513 | 907846 | 908034 | 59,58,128 | 0,0,0 | 49,133,77 | 1,12,1 | 300 | 150 | 1.29E-05 | 0.000266075 | 1.0,1.0,1.0 | 0.961,0.847,0.97 | 0.072 | ENSG00000060 | [[[[752579, 75432 | 4 |
| ENSG00000047 | WDR37 | chr10 | 1105125 | 1105267 | 1096169 | 1096246 | 1124217 | 1124352 | 6,2,5 | 0,0,0 | 7,26,20 | 1,4,4 | 292 | 150 | 1.73E-05 | 0.000346499 | 1.0,1.0,1.0 | 0.782,0.77,0.72 | 0.243 | ENSG00000047 | [[[[1056385, 105 | 2 |
| ENSG00000122 | FAM126A | chr7 | 22946951 | 22947247 | 22941268 | 22946163 | 22964414 | 22964531 | 23,9,49 | 0,0,0 | 10,26,22 | 29,50,36 | 300 | 150 | 0 | 0 | 1.0,1.0,1.0 | 0.147,0.206,0.23 | 0.804 | ENSG00000122 | [[[[23013925, 23 | 2 |
| ENSG00000099 | CIRBP | chr19 | 1270866 | 1271036 | 1269265 | 1269410 | 1271328 | 1271467 | 16,13,41 | 0,0,0 | 52,138,75 | 4,12,12 | 300 | 150 | 5.63E-07 | 1.65E-05 | 1.0,1.0,1.0 | 0.867,0.852,0.75 | 0.174 | ENSG00000099 | [[[[1269332, 126 | 2 |
| ENSG00000099 | CIRBP | chr19 | 1270927 | 1271036 | 1269267 | 1269410 | 1271328 | 1271467 | 82,179,308 | 0,0,0 | 147,321,243 | 4,12,12 | 259 | 150 | 1.07E-07 | 3.71E-06 | 1.0,1.0,1.0 | 0.955,0.939,0.92 | 0.062 | ENSG00000099 | [[[[1269332, 126 | 2 |
| ENSG00000099 | CIRBP | chr19 | 1271328 | 1271361 | 1271139 | 1271246 | 1274306 | 1274810 | 66,55,89 | 0,0,0 | 55,111,73 | 2,9,6 | 183 | 150 | 5.59E-06 | 0.000128077 | 1.0,1.0,1.0 | 0.958,0.91,0.909 | 0.074 | ENSG00000099 | [[[[1269332, 126 | 2 |
| ENSG00000099 | CIRBP | chr19 | 1271328 | 1271365 | 1271139 | 1271246 | 1274306 | 1274810 | 70,55,90 | 0,0,0 | 57,120,85 | 2,9,6 | 187 | 150 | 9.83E-06 | 0.000210438 | 1.0,1.0,1.0 | 0.958,0.914,0.91 | 0.07 | ENSG00000099 | [[[[1269332, 126 | 2 |
| ENSG00000099 | CIRBP | chr19 | 1271980 | 1272051 | 1271139 | 1271246 | 1274306 | 1274440 | 2,4,6 | 0,0,0 | 28,43,28 | 2,9,6 | 221 | 150 | 1.13E-05 | 0.000237095 | 1.0,1.0,1.0 | 0.905,0.764,0.76 | 0.19 | ENSG00000099 | [[[[1269332, 126 | 2 |
| ENSG00000122 | SMU1 | chr9 | 33056836 | 33056964 | 33053122 | 33053290 | 33057597 | 33057714 | 143,108,170 | 0,0,0 | 102,195,147 | 7,22,19 | 278 | 150 | 2.33E-15 | 3.01E-13 | 1.0,1.0,1.0 | 0.887,0.827,0.80 | 0.16 | ENSG00000122 | [[[[33076583, 33 | 2 |
| ENSG00000122 | SMU1 | chr9 | 33057597 | 33057714 | 33053122 | 33053290 | 33060464 | 33060584 | 84,73,145 | 0,0,0 | 75,182,116 | 9,18,6 | 267 | 150 | 1.34E-10 | 8.64E-09 | 1.0,1.0,1.0 | 0.824,0.85,0.916 | 0.137 | ENSG00000122 | [[[[33076583, 33 | 3 |
| ENSG00000129 | PSMA1 | chr11 | 14517641 | 14517745 | 14513459 | 14513699 | 14517879 | 14517981 | 581,498,766 | 1,0,0 | 5,741,210,698 | 54,88,46 | 254 | 150 | 0 | 0 | 0.997,1.0,1.0 | 0.863,0.89,0.9 | 0.115 | ENSG00000129 | [[[[14520297, 14 | 3 |
| ENSG00000186 | KIF24 | chr9 | 34286616 | 34286704 | 34271808 | 34271930 | 34306251 | 34306441 | 4,1,6 | 0,1,0 | 1,5,5 | 0,4,3 | 238 | 150 | 0.000243563 | 0.003442237 | 1.0,0.387,1.0 | 1.0,0.441,0.512 | 0.145 | ENSG00000186 | [[[[34329106, 34 | 3 |
| ENSG00000221 | FANCG | chr9 | 35075961 | 35076028 | 35075464 | 35075754 | 35076970 | 35077101 | 8,9,19 | 0,0,0 | 3,22,8 | 2,5,0 | 217 | 150 | 0.000715706 | 0.008725658 | 1.0,1.0,1.0 | 0.509,0.753,1.0 | 0.246 | ENSG00000221 | [[[[35079441, 35 | 3 |
| ENSG00000221 | FANCG | chr9 | 35076431 | 35076583 | 35075464 | 35075754 | 35076970 | 35077101 | 3,1,2 | 0,0,0 | 2,0,2 | 2,5,0 | 300 | 150 | 1.42E-06 | 3.77E-05 | 1.0,1.0,1.0 | 0.333,0.0,1.0 | 0.556 | ENSG00000221 | [[[[35079441, 35 | 3 |
| ENSG00000137 | ARHGEF39 | chr9 | 35660093 | 35660967 | 35658874 | 35659173 | 35663321 | 35663392 | 27,13,17 | 0,0,0 | 31,61,41 | 7,16,10 | 300 | 150 | 1.18E-09 | 6.45E-08 | 1.0,1.0,1.0 | 0.689,0.656,0.67 | 0.328 | ENSG00000137 | [[[[35665032, 35 | 4 |
| ENSG00000137 | ARHGEF39 | chr9 | 35662511 | 35662741 | 35658874 | 35659173 | 35662945 | 35663074 | 2,13,26 | 0,0,0 | 3,18,11 | 0,7,6 | 300 | 150 | 2.19E-07 | 7.04E-06 | 1.0,1.0,1.0 | 1.0,0.563,0.478 | 0.32 | ENSG00000137 | [[[[35665032, 35 | 3 |
| ENSG00000137 | ARHGEF39 | chr9 | 35662945 | 35663074 | 35658874 | 35659173 | 35663321 | 35663392 | 9,16,27 | 0,0,0 | 13,30,26 | 7,16,10 | 279 | 150 | 4.42E-12 | 3.66E-10 | 1.0,1.0,1.0 | 0.5,0.502,0.583 | 0.472 | ENSG00000137 | [[[[35665032, 35 | 4 |
| ENSG00000070 | GBA2 | chr9 | 35741671 | 35741890 | 35740208 | 35740362 | 35744296 | 35744412 | 19,33,30 | 0,2,0 | 18,47,24 | 0,5,10 | 300 | 150 | 0.000193944 | 0.002838679 | 1.0,0.892,1.0 | 1.0,0.825,0.545 | 0.174 | ENSG00000070 | [[[[35748346, 35 | 3 |
| ENSG00000122 | GLIPR2 | chr9 | 36148546 | 36148650 | 36147785 | 36147894 | 36162361 | 36162910 | 150,178,261 | 0,0,4 | 171,271,170 | 10,12,7 | 254 | 150 | 8.58E-07 | 2.39E-05 | 1.0,1.0,0.975 | 0.91,0.93,0.935 | 0.067 | ENSG00000122 | [[[[36136733, 36 | 2 |
| ENSG00000122 | GLIPR2 | chr9 | 36150871 | 36150949 | 36147785 | 36147894 | 36162361 | 36163902 | 131,125,164 | 0,0,4 | 133,171,94 | 10,12,7 | 228 | 150 | 1.89E-07 | 6.17E-06 | 1.0,1.0,0.964 | 0.897,0.904,0.89 | 0.088 | ENSG00000122 | [[[[36136733, 36 | 2 |
| ENSG00000165 | MELK | chr9 | 36594627 | 36594771 | 36589535 | 36589652 | 36599393 | 36599486 | 73,65,104 | 5,0,0 | 47,101,81 | 9,29,6 | 294 | 150 | 6.31E-08 | 2.33E-06 | 0.882,1.0,1.0 | 0.727,0.64,0.873 | 0.214 | ENSG00000165 | [[[[36572895, 36 | 2 |
| ENSG00000165 | MELK | chr9 | 36594627 | 36594771 | 36589535 | 36589652 | 36630298 | 36630367 | 73,65,102 | 0,0,0 | 45,91,73 | 5,16,7 | 294 | 150 | 1.04E-11 | 7.99E-10 | 1.0,1.0,1.0 | 0.821,0.744,0.84 | 0.198 | ENSG00000165 | [[[[36572895, 36 | 4 |
| ENSG00000165 | MELK | chr9 | 36597221 | 36597290 | 36589535 | 36589652 | 36599393 | 36599486 | 77,58,99 | 5,0,0 | 40,102,64 | 9,29,6 | 219 | 150 | 2.87E-09 | 1.45E-07 | 0.913,1.0,1.0 | 0.753,0.707,0.88 | 0.191 | ENSG00000165 | [[[[36572895, 36 | 2 |
| ENSG00000165 | MELK | chr9 | 36665349 | 36665581 | 36651745 | 36651877 | 36674833 | 36674937 | 26,13,36 | 0,0,0 | 10,48,10 | 2,10,1 | 300 | 150 | 9.02E-08 | 3.20E-06 | 1.0,1.0,1.0 | 0.714,0.706,0.83 | 0.249 | ENSG00000165 | [[[[36572895, 36 | 4 |
| ENSG00000165 | MELK | chr9 | 36674833 | 36674937 | 36665349 | 36665581 | 36677159 | 36677417 | 98,104,93 | 0,0,0 | 70,152,81 | 0,10,6 | 254 | 150 | 6.66E-06 | 0.000149348 | 1.0,1.0,1.0 | 1.0,0.9,0.889 | 0.07 | ENSG00000165 | [[[[36572895, 36 | 3 |
| ENSG00000137 | RMDN3 | chr15 | 40740132 | 40740193 | 40738500 | 40738576 | 40751425 | 40751670 | 26,25,28 | 0,0,0 | 13,41,18 | 2,4,2 | 211 | 150 | 0.000222653 | 0.003196858 | 1.0,1.0,1.0 | 0.822,0.879,0.86 | 0.145 | ENSG00000137 | [[[[40755083, 40 | 3 |
| ENSG00000197 | MCMBP | chr10 | 119838534 | 119838700 | 119835539 | 119835704 | 119840842 | 119840960 | 55,62,118 | 0,0,0 | 96,158,109 | 3,7,3 | 300 | 150 | 0.000251847 | 0.00354521 | 1.0,1.0,1.0 | 0.941,0.919,0.94 | 0.064 | ENSG00000197 | [[[[119872527, 11 | 2 |
| ENSG00000197 | MCMBP | chr10 | 119840842 | 119840960 | 119838534 | 119838700 | 119843253 | 119843426 | 45,50,104 | 0,0,0 | 85,157,102 | 1,5,9 | 268 | 150 | 6.68E-05 | 0.0011227 | 1.0,1.0,1.0 | 0.979,0.946,0.86 | 0.07 | ENSG00000197 | [[[[119872527, 11 | 2 |
| ENSG00000003 | RBM5 | chr3 | 50090381 | 50090451 | 50088907 | 50089029 | 50093719 | 50093783 | 11,10,13 | 0,0,0 | 12,26,8 | 1,11,8 | 220 | 150 | 6.01E-08 | 2.23E-06 | 1.0,1.0,1.0 | 0.891,0.617,0.40 | 0.362 | ENSG00000003 | [[[[50088919, 50 | 2 |
| ENSG00000003 | RBM5 | chr3 | 50090381 | 50090451 | 50088907 | 50089029 | 50099981 | 50100010 | 10,10,12 | 0,0,0 | 22,32,20 | 10,11,5 | 220 | 150 | 1.46E-08 | 6.31E-07 | 1.0,1.0,1.0 | 0.6,0.665,0.732 | 0.334 | ENSG00000003 | [[[[50088919, 50 | 3 |
| ENSG00000003 | RBM5 | chr3 | 50093719 | 50093875 | 50088907 | 50089029 | 50099981 | 50100004 | 91,48,110 | 0,0,0 | 64,132,105 | 10,11,5 | 300 | 150 | 3.45E-09 | 1.71E-07 | 1.0,1.0,1.0 | 0.762,0.857,0.91 | 0.156 | ENSG00000003 | [[[[50088919, 50 | 3 |
| ENSG00000003 | RBM5 | chr3 | 50110678 | 50110770 | 50109602 | 50109688 | 50113382 | 50113489 | 52,45,84 | 0,0,0 | 48,104,49 | 17,24,15 | 242 | 150 | 0 | 0 | 1.0,1.0,1.0 | 0.636,0.729,0.66 | 0.322 | ENSG00000003 | [[[[50088919, 50 | 2 |
| ENSG00000197 | COL13A1 | chr10 | 69880502 | 69880553 | 69867797 | 69867805 | 69887455 | 69887491 | 1,3,7 | 2,6,10 | 0,2,0 | 4,4,3 | 201 | 150 | 0.000821222 | 0.009825013 | 0.272,0.272,0.34 | 0.0,0.272,0.0 | 0.205 | ENSG00000197 | [[[[69801906, 69 | 4 |
| ENSG00000116 | MRPS15 | chr1 | 36461263 | 36461312 | 36457922 | 36457981 | 36463805 | 36463850 | 1,0,2 | 10,10,6 | 9,15,8 | 12,27,9 | 199 | 150 | 2.42E-05 | 0.000464256 | 0.07,0.0,0.201 | 0.361,0.295,0.40 | -0.262 | ENSG00000116 | [[[[36464146, 36 | 3 |
| ENSG00000116 | MRPS15 | chr1 | 36461263 | 36461312 | 36460691 | 36460776 | 36463805 | 36463850 | 89,81,109 | 0,0,0 | 123,236,144 | 4,9,6 | 199 | 150 | 0.000140541 | 0.002138367 | 1.0,1.0,1.0 | 0.959,0.952,0.94 | 0.047 | ENSG00000116 | [[[[36464146, 36 | 2 |

| | | | | | | | | | | | | | | | | | | | | | | |
|---|---|---|---|---|---|---|---|---|---|---|---|---|---|---|---|---|---|---|---|---|---|---|
| ENSG00000116 | MRPS15 | chr1 | 36462087 | 36462163 | 36460691 | 36460776 | 36463805 | 36463850 | 32,100,165 | 0,0,0 | 136,282,203 | 4,9,6 | 226 | 150 | 0.000308796 | 0.00423499 | 1.0,1.0,1.0 | 0.958,0.954,0.95 | 0.044 | ENSG00000116 | [[[[36464146, 364 | 2 |
| ENSG00000119 | TJP2 | chr9 | 69252814 | 69252900 | 69248011 | 69248224 | 69254208 | 69254520 | 16,7,24 | 0,0,0 | 4,16,11 | 2,4,5 | 236 | 150 | 2.82E-08 | 1.14E-06 | 1.0,1.0,1.0 | 0.56,0.718,0.583 | 0.38 | ENSG00000119 | [[[[69174277, 69 | 3 |
| ENSG00000135 | TTLL4 | chr2 | 218747596 | 218747725 | 218747289 | 218747372 | 218748835 | 218748934 | 17,19,30 | 0,0,0 | 18,43,20 | 6,13,10 | 279 | 150 | 2.00E-11 | 1.45E-09 | 1.0,1.0,1.0 | 0.617,0.64,0.518 | 0.408 | ENSG00000135 | [[[[218710835, 2 | 2 |
| ENSG00000135 | CEMIP2 | chr9 | 71734805 | 71734994 | 71732350 | 71732520 | 71746200 | 71746341 | 133,71,167 | 0,0,0 | 80,162,136 | 7,5,2 | 300 | 150 | 4.41E-05 | 0.000783018 | 1.0,1.0,1.0 | 0.851,0.942,0.97 | 0.079 | ENSG00000135 | [[[[71768357, 71 | 3 |
| ENSG00000135 | CEMIP2 | chr9 | 71740062 | 71740232 | 71732350 | 71732520 | 71746200 | 71746341 | 12,9,21 | 0,0,0 | 7,17,19 | 7,5,2 | 300 | 150 | 3.03E-08 | 1.21E-06 | 1.0,1.0,1.0 | 0.333,0.63,0.826 | 0.404 | ENSG00000135 | [[[[71768357, 71 | 3 |
| ENSG00000135 | CEMIP2 | chr9 | 71740062 | 71740232 | 71734805 | 71734994 | 71746200 | 71746341 | 72,64,93 | 2,0,0 | 38,85,83 | 3,3,14 | 300 | 150 | 9.00E-07 | 2.50E-05 | 0.947,1.0,1.0 | 0.864,0.934,0.74 | 0.134 | ENSG00000135 | [[[[71768357, 71 | 2 |
| ENSG00000178 | PLEC | chr8 | 143937164 | 143937242 | 143935847 | 143936014 | 143938150 | 143938240 | 11,16,63 | 0,0,0 | 22,65,60 | 6,15,8 | 228 | 150 | 1.64E-09 | 8.70E-08 | 1.0,1.0,1.0 | 0.707,0.74,0.831 | 0.241 | ENSG00000178 | [[[[143939350, 1 | 2 |
| ENSG00000148 | CEP78 | chr9 | 78265371 | 78265543 | 78254835 | 78254964 | 78265858 | 78265906 | 18,22,29 | 0,0,0 | 14,44,30 | 4,4,1 | 300 | 150 | 2.36E-05 | 0.000455491 | 1.0,1.0,1.0 | 0.636,0.846,0.93 | 0.193 | ENSG00000148 | [[[[78236075, 78 | 3 |
| ENSG00000135 | NAA35 | chr9 | 85977362 | 85977446 | 85976684 | 85976735 | 86003584 | 86003644 | 12,5,20 | 0,0,0 | 14,30,32 | 1,11,0 | 234 | 150 | 5.06E-05 | 0.000882505 | 1.0,1.0,1.0 | 0.9,0.636,1.0 | 0.155 | ENSG00000135 | [[[[85941146, 85 | 3 |
| ENSG00000166 | VPS39 | chr15 | 42187270 | 42187363 | 42184191 | 42184700 | 42191124 | 42191167 | 18,23,45 | 0,0,0 | 27,36,35 | 1,5,3 | 243 | 150 | 0.000127814 | 0.001973088 | 1.0,1.0,1.0 | 0.943,0.816,0.87 | 0.121 | ENSG00000166 | [[[[42208081, 42 | 3 |
| ENSG00000166 | VPS39 | chr15 | 42189113 | 42189208 | 42184191 | 42184700 | 42191124 | 42191167 | 5,13,19 | 0,0,0 | 12,55,35 | 1,5,3 | 245 | 150 | 0.000361287 | 0.004846075 | 1.0,1.0,1.0 | 0.88,0.871,0.877 | 0.124 | ENSG00000166 | [[[[42208081, 42 | 3 |
| ENSG00000160 | VPS11 | chr11 | 119073197 | 119073399 | 119070233 | 119070397 | 119073799 | 119073951 | 9,18,17 | 0,0,0 | 11,26,16 | 2,4,2 | 300 | 150 | 3.17E-05 | 0.000587951 | 1.0,1.0,1.0 | 0.733,0.765,0.8 | 0.234 | ENSG00000160 | [[[[119067818, 1 | 2 |
| ENSG00000135 | PHF21A | chr11 | 45965314 | 45965608 | 45953526 | 45953625 | 45971115 | 45971367 | 15,21,29 | 0,0,0 | 21,67,50 | 6,14,16 | 300 | 150 | 2.12E-10 | 1.32E-08 | 1.0,1.0,1.0 | 0.636,0.705,0.61 | 0.35 | ENSG00000135 | [[[[46120935, 46 | 2 |
| ENSG00000136 | XPA | chr9 | 97687095 | 97687261 | 97684922 | 97685040 | 97693648 | 97693759 | 30,10,8 | 0,0,0 | 18,36,14 | 2,4,4 | 300 | 150 | 2.99E-06 | 7.33E-05 | 1.0,1.0,1.0 | 0.818,0.818,0.63 | 0.243 | ENSG00000136 | [[[[97697121, 97 | 2 |
| ENSG00000136 | XPA | chr9 | 97689533 | 97689639 | 97684922 | 97685040 | 97693648 | 97693759 | 5,12,11 | 0,0,0 | 8,19,8 | 2,4,4 | 256 | 150 | 2.30E-07 | 7.36E-06 | 1.0,1.0,1.0 | 0.701,0.736,0.54 | 0.341 | ENSG00000136 | [[[[97697121, 97 | 2 |
| ENSG00000241 | TMEFF1 | chr9 | 100550094 | 100550160 | 100516674 | 100516771 | 100561396 | 100561520 | 2,6,12 | 0,0,0 | 5,15,6 | 4,3,1 | 216 | 150 | 2.72E-06 | 6.73E-05 | 1.0,1.0,1.0 | 0.465,0.776,0.80 | 0.318 | ENSG00000241 | [[[[100473149, 1 | 2 |
| ENSG00000175 | PPP2R2D | chr10 | 131934893 | 131935006 | 131934457 | 131934555 | 131943967 | 131944145 | 3,1,1 | 0,0,0 | 2,9,2 | 2,4,5 | 263 | 150 | 1.63E-10 | 1.03E-08 | 1.0,1.0,1.0 | 0.363,0.562,0.18 | 0.63 | ENSG00000175 | [[[[131901008, 1 | 2 |
| ENSG00000175 | PPP2R2D | chr10 | 131934893 | 131935006 | 131934457 | 131934555 | 131944094 | 131944145 | 3,1,1 | 0,0,0 | 4,7,5 | 1,3,2 | 263 | 150 | 3.23E-05 | 0.000597653 | 1.0,1.0,1.0 | 0.695,0.571,0.58 | 0.382 | ENSG00000175 | [[[[131901008, 1 | 2 |
| ENSG00000175 | PPP2R2D | chr10 | 131940030 | 131940196 | 131934457 | 131934555 | 131943967 | 131944145 | 25,41,61 | 0,0,0 | 37,99,55 | 2,4,5 | 300 | 150 | 0.000112839 | 0.001770976 | 1.0,1.0,1.0 | 0.902,0.925,0.84 | 0.109 | ENSG00000175 | [[[[131901008, 1 | 2 |
| ENSG00000136 | SMC2 | chr9 | 104096147 | 104096297 | 104095298 | 104095552 | 104099643 | 104099682 | 26,17,53 | 0,0,0 | 10,24,21 | 4,11,4 | 300 | 150 | 3.37E-11 | 2.36E-09 | 1.0,1.0,1.0 | 0.556,0.522,0.72 | 0.399 | ENSG00000136 | [[[[104094309, 1 | 2 |
| ENSG00000136 | SMC2 | chr9 | 104098445 | 104098568 | 104095298 | 104095552 | 104099643 | 104099682 | 32,14,38 | 0,0,0 | 5,9,8 | 4,11,4 | 273 | 150 | 8.99E-15 | 1.08E-12 | 1.0,1.0,1.0 | 0.407,0.31,0.524 | 0.586 | ENSG00000136 | [[[[104094309, 1 | 2 |
| ENSG00000106 | FSD1L | chr9 | 105471903 | 105472005 | 105468192 | 105468324 | 105484380 | 105484502 | 3,4,1 | 0,0,0 | 7,37,21 | 6,4,5 | 252 | 150 | 1.07E-06 | 2.93E-05 | 1.0,1.0,1.0 | 0.41,0.846,0.714 | 0.343 | ENSG00000106 | [[[[105448034, 1 | 2 |
| ENSG00000106 | FSD1L | chr9 | 105535066 | 105535318 | 105534492 | 105534593 | 105546357 | 105546575 | 3,2,1 | 0,0,0 | 18,36,19 | 0,5,2 | 300 | 150 | 0.000838039 | 0.009996383 | 1.0,1.0,1.0 | 1.0,0.783,0.826 | 0.13 | ENSG00000106 | [[[[105448034, 1 | 2 |
| ENSG00000136 | NCBP1 | chr9 | 97645116 | 97645224 | 97641561 | 97641662 | 97645610 | 97645732 | 22,17,26 | 0,0,0 | 13,27,16 | 2,12,2 | 258 | 150 | 2.77E-08 | 1.12E-06 | 1.0,1.0,1.0 | 0.791,0.567,0.82 | 0.273 | ENSG00000136 | [[[[97633821, 97 | 2 |
| ENSG00000119 | PTBP3 | chr9 | 112268048 | 112268195 | 112262434 | 112262599 | 112297840 | 112297916 | 21,11,65 | 0,0,0 | 23,74,69 | 0,3,12 | 297 | 150 | 1.18E-05 | 0.000246588 | 1.0,1.0,1.0 | 1.0,0.926,0.744 | 0.11 | ENSG00000119 | [[[[112333470, 1 | 2 |
| ENSG00000157 | TMEM268 | chr9 | 114628100 | 114628250 | 114617117 | 114617301 | 114633767 | 114633878 | 13,25,36 | 0,0,0 | 32,67,48 | 1,3,7 | 300 | 150 | 4.55E-05 | 0.000804439 | 1.0,1.0,1.0 | 0.941,0.918,0.77 | 0.122 | ENSG00000157 | [[[[114611291, 1 | 3 |
| ENSG00000119 | FBXW2 | chr9 | 120771347 | 120771517 | 120756973 | 120764847 | 120776092 | 120776226 | 84,65,85 | 0,0,0 | 84,184,110 | 6,8,0 | 300 | 150 | 0.000118182 | 0.001845071 | 1.0,1.0,1.0 | 0.875,0.92,1.0 | 0.068 | ENSG00000119 | [[[[120793354, 1 | 2 |
| ENSG00000119 | FBXW2 | chr9 | 120776092 | 120776226 | 120756973 | 120764847 | 120778350 | 120778545 | 67,83,148 | 0,0,0 | 110,233,116 | 2,7,8 | 284 | 150 | 3.11E-05 | 0.000578784 | 1.0,1.0,1.0 | 0.967,0.946,0.88 | 0.067 | ENSG00000119 | [[[[120793354, 1 | 3 |
| ENSG00000095 | PSMD5 | chr9 | 120824493 | 120824685 | 120821354 | 120821464 | 120831330 | 120831459 | 58,46,97 | 0,0,0 | 60,127,96 | 5,8,0 | 300 | 150 | 6.31E-05 | 0.001070024 | 1.0,1.0,1.0 | 0.857,0.888,1.0 | 0.085 | ENSG00000095 | [[[[120842737, 1 | 3 |
| ENSG00000095 | PSMD5 | chr9 | 120829098 | 120829208 | 120821354 | 120821464 | 120831330 | 120831459 | 43,26,53 | 0,0,0 | 48,92,73 | 5,8,0 | 260 | 150 | 5.94E-05 | 0.001014216 | 1.0,1.0,1.0 | 0.847,0.869,1.0 | 0.095 | ENSG00000095 | [[[[120842737, 1 | 3 |
| ENSG00000095 | PSMD5 | chr9 | 120829098 | 120829208 | 120826764 | 120826907 | 120831831 | 120831945 | 35,15,61 | 0,0,0 | 18,50,46 | 0,7,4 | 260 | 150 | 1.55E-05 | 0.000312585 | 1.0,1.0,1.0 | 1.0,0.805,0.869 | 0.109 | ENSG00000095 | [[[[120842737, 1 | 2 |
| ENSG00000106 | GRB10 | chr7 | 50603997 | 50604085 | 50595436 | 50595530 | 50605289 | 50605406 | 113,81,194 | 0,0,1 | 144,317,205 | 10,17,5 | 238 | 150 | 3.29E-07 | 1.02E-05 | 1.0,1.0,0.992 | 0.901,0.922,0.96 | 0.069 | ENSG00000106 | [[[[50782424, 50 | 2 |
| ENSG00000106 | GRB10 | chr7 | 50604310 | 50604377 | 50595436 | 50595530 | 50605289 | 50605406 | 42,47,123 | 0,0,1 | 112,141,146 | 10,17,5 | 217 | 150 | 1.71E-07 | 5.65E-06 | 1.0,1.0,0.988 | 0.886,0.851,0.95 | 0.099 | ENSG00000106 | [[[[50782424, 50 | 2 |
| ENSG00000106 | GRB10 | chr7 | 50616209 | 50616347 | 50614769 | 50614880 | 50626821 | 50626978 | 144,82,244 | 0,0,0 | 200,397,291 | 13,15,6 | 288 | 150 | 8.96E-08 | 3.18E-06 | 1.0,1.0,1.0 | 0.889,0.932,0.96 | 0.072 | ENSG00000106 | [[[[50782424, 50 | 3 |
| ENSG00000106 | GRB10 | chr7 | 50618070 | 50618139 | 50616209 | 50616347 | 50626821 | 50626978 | 77,59,138 | 0,0,0 | 79,197,157 | 4,13,13 | 219 | 150 | 2.67E-08 | 1.08E-06 | 1.0,1.0,1.0 | 0.931,0.912,0.89 | 0.088 | ENSG00000106 | [[[[50782424, 50 | 2 |
| ENSG00000135 | SNX14 | chr6 | 85513799 | 85513895 | 85507967 | 85508059 | 85514505 | 85514629 | 59,55,48 | 0,0,0 | 22,45,24 | 12,23,5 | 246 | 150 | 2.22E-16 | 3.17E-14 | 1.0,1.0,1.0 | 0.528,0.544,0.74 | 0.394 | ENSG00000135 | [[[[85593579, 85 | 2 |
| ENSG00000136 | PSMB7 | chr9 | 124384597 | 124384656 | 124356763 | 124356915 | 124412351 | 124412492 | 294,285,383 | 0,0,0 | 536,970,435 | 65,88,32 | 209 | 150 | 0 | 0 | 1.0,1.0,1.0 | 0.855,0.888,0.90 | 0.117 | ENSG00000136 | [[[[124415364, 1 | 2 |
| ENSG00000136 | PSMB7 | chr9 | 124405316 | 124405432 | 124356763 | 124356915 | 124412351 | 124412492 | 462,403,582 | 0,0,0 | 542,971,531 | 65,88,32 | 266 | 150 | 0 | 0 | 1.0,1.0,1.0 | 0.825,0.862,0.90 | 0.137 | ENSG00000136 | [[[[124415364, 1 | 2 |
| ENSG00000160 | MCM3AP | chr21 | 46256788 | 46256986 | 46254775 | 46254844 | 46260792 | 46260906 | 14,23,40 | 0,0,0 | 7,40,25 | 10,2,5 | 300 | 150 | 2.23E-07 | 7.16E-06 | 1.0,1.0,1.0 | 0.259,0.909,0.71 | 0.373 | ENSG00000160 | [[[[46284068, 46 | 2 |
| ENSG00000136 | FPGS | chr9 | 127807585 | 127807775 | 127806972 | 127807087 | 127808233 | 127808311 | 2,4,3 | 0,0,0 | 6,9,5 | 2,3,3 | 300 | 150 | 8.20E-08 | 2.94E-06 | 1.0,1.0,1.0 | 0.6,0.6,0.455 | 0.448 | ENSG00000136 | [[[[127802882, 1 | 3 |
| ENSG00000121 | TBC1D15 | chr12 | 71884810 | 71885021 | 71872928 | 71873003 | 71893221 | 71893324 | 4,15,11 | 1,0,0 | 13,33,17 | 0,4,4 | 300 | 150 | 0.000398428 | 0.005277142 | 0.667,1.0,1.0 | 1.0,0.805,0.68 | 0.061 | ENSG00000121 | [[[[71839759, 71 | 2 |
| ENSG00000135 | ACVR1B | chr12 | 51986817 | 51986942 | 51980968 | 51981199 | 51991862 | 51991993 | 20,14,21 | 0,1,2 | 9,33,11 | 5,6,5 | 275 | 150 | 0.000100738 | 0.001607969 | 1.0,0.884,0.851 | 0.495,0.75,0.545 | 0.315 | ENSG00000135 | [[[[51951699, 51 | 3 |
| ENSG00000140 | BNIP2 | chr15 | 59668102 | 59668138 | 59663378 | 59664120 | 59677910 | 59678016 | 10,10,14 | 0,0,0 | 16,30,18 | 6,7,7 | 186 | 150 | 5.94E-08 | 2.21E-06 | 1.0,1.0,1.0 | 0.683,0.776,0.67 | 0.289 | ENSG00000140 | [[[[59689135, 59 | 4 |
| ENSG00000140 | BNIP2 | chr15 | 59668891 | 59668990 | 59659145 | 59664120 | 59677910 | 59678016 | 31,29,39 | 0,0,0 | 72,135,84 | 6,7,7 | 249 | 150 | 9.55E-06 | 0.00020555 | 1.0,1.0,1.0 | 0.878,0.921,0.87 | 0.108 | ENSG00000140 | [[[[59689135, 59 | 4 |
| ENSG00000140 | BNIP2 | chr15 | 59672636 | 59672739 | 59659145 | 59664120 | 59677910 | 59678016 | 83,72,131 | 0,0,0 | 102,237,173 | 6,7,7 | 253 | 150 | 1.98E-05 | 0.000390393 | 1.0,1.0,1.0 | 0.91,0.953,0.936 | 0.067 | ENSG00000140 | [[[[59689135, 59 | 4 |
| ENSG00000130 | EXOSC2 | chr9 | 130698161 | 130698251 | 130697581 | 130697627 | 130702133 | 130702310 | 37,33,58 | 0,0,0 | 37,77,49 | 20,21,8 | 240 | 150 | 2.82E-13 | 2.76E-11 | 1.0,1.0,1.0 | 0.536,0.696,0.79 | 0.325 | ENSG00000130 | [[[[130693780, 1 | 3 |
| ENSG00000126 | NUP214 | chr9 | 131195232 | 131195294 | 131189052 | 131189131 | 131197215 | 131197330 | 8,8,12 | 0,0,0 | 9,15,20 | 0,4,3 | 212 | 150 | 0.000488715 | 0.006308506 | 1.0,1.0,1.0 | 1.0,0.726,0.825 | 0.15 | ENSG00000126 | [[[[131125586, 1 | 2 |
| ENSG00000148 | SURF1 | chr9 | 133352445 | 133352608 | 133352060 | 133352142 | 133353748 | 133353940 | 147,111,188 | 0,0,0 | 115,278,166 | 3,12,4 | 300 | 150 | 8.80E-06 | 0.000191144 | 1.0,1.0,1.0 | 0.95,0.921,0.954 | 0.058 | ENSG00000148 | [[[[133356400, 1 | 2 |
| ENSG00000148 | SURF1 | chr9 | 133353748 | 133353940 | 133352060 | 133352142 | 133354658 | 133354741 | 85,95,129 | 0,0,0 | 117,204,114 | 4,14,2 | 300 | 150 | 1.85E-06 | 4.76E-05 | 1.0,1.0,1.0 | 0.936,0.879,0.96 | 0.073 | ENSG00000148 | [[[[133356400, 1 | 3 |
| ENSG00000116 | MAP7D1 | chr1 | 36174897 | 36175008 | 36173363 | 36173478 | 36176696 | 36176834 | 47,86,155 | 0,0,0 | 97,194,153 | 5,5,4 | 261 | 150 | 0.000463884 | 0.00603398 | 1.0,1.0,1.0 | 0.918,0.957,0.95 | 0.056 | ENSG00000116 | [[[[36156160, 36 | 2 |
| ENSG00000116 | MAP7D1 | chr1 | 36176198 | 36176581 | 36173363 | 36173478 | 36176696 | 36176834 | 17,25,82 | 0,0,0 | 41,88,71 | 5,5,4 | 300 | 150 | 7.74E-06 | 0.000170595 | 1.0,1.0,1.0 | 0.804,0.898,0.89 | 0.133 | ENSG00000116 | [[[[36156160, 36 | 2 |
| ENSG00000069 | NEDD4 | chr15 | 55869578 | 55869681 | 55860660 | 55860778 | 55872414 | 55872476 | 33,16,45 | 0,0,0 | 27,55,32 | 4,9,9 | 253 | 150 | 3.33E-09 | 1.66E-07 | 1.0,1.0,1.0 | 0.8,0.784,0.678 | 0.246 | ENSG00000069 | [[[[55993511, 55 | 2 |
| ENSG00000137 | KLC4 | chr6 | 43066995 | 43067083 | 43061310 | 43061593 | 43070353 | 43070455 | 13,9,29 | 0,0,0 | 7,28,20 | 2,9,1 | 238 | 150 | 1.03E-06 | 2.83E-05 | 1.0,1.0,1.0 | 0.688,0.662,0.92 | 0.241 | ENSG00000137 | [[[[43059631, 43 | 4 |
| ENSG00000175 | ETV4 | chr17 | 43533858 | 43533985 | 43530106 | 43530181 | 43536425 | 43536479 | 8,7,15 | 0,0,0 | 14,19,21 | 3,11,3 | 277 | 150 | 7.06E-08 | 2.57E-06 | 1.0,1.0,1.0 | 0.716,0.483,0.79 | 0.337 | ENSG00000175 | [[[[43546185, 43 | 3 |
| ENSG00000140 | NIPA2 | chr15 | 22839654 | 22839790 | 22838640 | 22838921 | 22858539 | 22858630 | 3,9,4 | 0,0,0 | 5,3,3 | 1,2,2 | 286 | 150 | 0.000279195 | 0.003879303 | 1.0,1.0,1.0 | 0.724,0.44,0.44 | 0.465 | ENSG00000140 | [[[[22838666, 22 | 4 |
| ENSG00000140 | NIPA2 | chr15 | 22845145 | 22845267 | 22839654 | 22839790 | 22858539 | 22858630 | 6,12,8 | 1,0,0 | 16,17,18 | 5,1,3 | 272 | 150 | 0.000137415 | 0.002097511 | 0.768,1.0,1.0 | 0.638,0.904,0.76 | 0.153 | ENSG00000140 | [[[[22838666, 22 | 3 |
| ENSG00000140 | NIPA2 | chr15 | 22851638 | 22851870 | 22839654 | 22839790 | 22858539 | 22858630 | 15,14,38 | 1,0,0 | 25,38,32 | 5,1,3 | 300 | 150 | 0.000226845 | 0.003246269 | 0.882,1.0,1.0 | 0.714,0.95,0.842 | 0.125 | ENSG00000140 | [[[[22838666, 22 | 3 |
| ENSG00000140 | NIPA2 | chr15 | 22851638 | 22851870 | 22845145 | 22845267 | 22858539 | 22858630 | 27,20,48 | 0,0,0 | 36,72,30 | 8,5,6 | 300 | 150 | 6.63E-08 | 2.44E-06 | 1.0,1.0,1.0 | 0.692,0.878,0.71 | 0.239 | ENSG00000140 | [[[[22838666, 22 | 2 |
| ENSG00000150 | ITGB1 | chr10 | 32908367 | 32908534 | 32900318 | 32901635 | 32911526 | 32911670 | 600,552,749 | 0,0,0 | 425,839,503 | 8,14,8 | 300 | 150 | 6.54E-06 | 0.000147211 | 1.0,1.0,1.0 | 0.964,0.968,0.96 | 0.033 | ENSG00000150 | [[[[32958145, 32 | 2 |
| ENSG00000150 | ITGB1 | chr10 | 32920244 | 32920385 | 32900318 | 32901635 | 32922256 | 32922346 | 333,297,544 | 0,0,0 | 160,445,298 | 2,5,16 | 291 | 150 | 1.68E-06 | 4.38E-05 | 1.0,1.0,1.0 | 0.976,0.979,0.90 | 0.046 | ENSG00000150 | [[[[32958145, 32 | 6 |
| ENSG00000150 | ITGB1 | chr10 | 32920244 | 32920385 | 32911885 | 32912124 | 32922256 | 32922346 | 333,297,544 | 0,0,0 | 159,448,297 | 8,18,11 | 291 | 150 | 5.52E-12 | 4.48E-10 | 1.0,1.0,1.0 | 0.911,0.928,0.93 | 0.076 | ENSG00000150 | [[[[32958145, 32 | 2 |
| ENSG00000150 | ITGB1 | chr10 | 32922256 | 32922346 | 32920244 | 32920385 | 32923584 | 32923740 | 239,209,350 | 0,0,0 | 121,268,203 | 3,4,10 | 240 | 150 | 9.08E-05 | 0.00146719 | 1.0,1.0,1.0 | 0.962,0.977,0.92 | 0.045 | ENSG00000150 | [[[[32958145, 32 | 2 |
| ENSG00000162 | ZNF75A | chr16 | 3313048 | 3313175 | 3312676 | 3312768 | 3317189 | 3317925 | 4,10,10 | 0,0,0 | 8,21,8 | 1,2,3 | 277 | 150 | 0.000743074 | 0.009008865 | 1.0,1.0,1.0 | 0.812,0.85,0.591 | 0.249 | ENSG00000162 | [[[[3305497, 330 | 2 |
| ENSG00000111 | LTBR | chr12 | 6386065 | 6386162 | 6385021 | 6385379 | 6388397 | 6388505 | 30,31,84 | 0,0,0 | 53,82,76 | 17,24,21 | 247 | 150 | 1.11E-16 | 1.65E-14 | 1.0,1.0,1.0 | 0.654,0.675,0.68 | 0.328 | ENSG00000111 | [[[[6384162, 638 | 2 |
| ENSG00000111 | LTBR | chr12 | 6386346 | 6386444 | 6385226 | 6385379 | 6388397 | 6388505 | 47,63,88 | 0,0,0 | 50,106,65 | 17,24,21 | 248 | 150 | 0 | 0 | 1.0,1.0,1.0 | 0.64,0.728,0.652 | 0.327 | ENSG00000111 | [[[[6384162, 638 | 2 |
| ENSG00000107 | EIF3A | chr10 | 119049800 | 119049985 | 119037118 | 119037309 | 119050520 | 119050674 | 85,84,151 | 0,0,0 | 71,158,78 | 14,9,10 | 300 | 150 | 5.05E-12 | 4.13E-10 | 1.0,1.0,1.0 | 0.717,0.898,0.79 | 0.196 | ENSG00000107 | [[[[119080628, 1 | 4 |
| ENSG00000107 | EIF3A | chr10 | 119050520 | 119050674 | 119037118 | 119037309 | 119051198 | 119051321 | 153,124,207 | 0,0,0 | 108,312,178 | 4,8,8 | 300 | 150 | 5.30E-06 | 0.000122176 | 1.0,1.0,1.0 | 0.931,0.951,0.91 | 0.067 | ENSG00000107 | [[[[119080628, 1 | 5 |
| ENSG00000266 | NCOA4 | chr10 | 46010222 | 46011206 | 46005087 | 46006597 | 46015125 | 46015266 | 1,1,1 | 0,0,0 | 2,2,1 | 0,2,6 | 300 | 150 | 2.20E-06 | 5.57E-05 | 1.0,1.0,1.0 | 1.0,0.333,0.077 | 0.53 | ENSG00000266 | [[[[46030526, 46 | 6 |
| ENSG00000204 | TIMM23B | chr10 | 49943234 | 49943358 | 49942048 | 49942300 | 49958369 | 49958480 | 1,1,8 | 0,0,0 | 13,9,10 | 0,6,1 | 274 | 150 | 0.00049261 | 0.006351252 | 1.0,1.0,1.0 | 1.0,0.451,0.846 | 0.234 | ENSG00000204 | [[[[49942053, 49 | 4 |
| ENSG00000196 | NF1 | chr17 | 31230841 | 31230925 | 31229024 | 31229974 | 31232072 | 31232189 | 2,2,1 | 0,0,0 | 1,2,8 | 2,1,2 | 234 | 150 | 0.00069811 | 0.008548063 | 1.0,1.0,1.0 | 0.243,0.562,0.71 | 0.492 | ENSG00000196 | [[[[31094977, 31 | 2 |
| ENSG00000170 | CDK1 | chr10 | 60785663 | 60785787 | 60780140 | 60780202 | 60788059 | 60788230 | 166,132,191 | 0,0,0 | 82,193,139 | 5,7,9 | 274 | 150 | 8.52E-08 | 3.05E-06 | 1.0,1.0,1.0 | 0.9,0.938,0.894 | 0.089 | ENSG00000170 | [[[[60778478, 60 | 2 |
| ENSG00000182 | RTKN2 | chr10 | 62236065 | 62236263 | 62204856 | 62205022 | 62239647 | 62239765 | 8,3,17 | 0,0,0 | 1,10,6 | 2,2,2 | 300 | 150 | 2.70E-05 | 0.000509118 | 1.0,1.0,1.0 | 0.2,0.714,0.6 | 0.495 | ENSG00000182 | [[[[62268551, 62 | 4 |
| ENSG00000171 | JMJD1C | chr10 | 63198512 | 63198727 | 63197410 | 63197563 | 63206594 | 63208801 | 11,10,20 | 0,0,0 | 20,49,40 | 2,13,5 | 300 | 150 | 3.65E-07 | 1.12E-05 | 1.0,1.0,1.0 | 0.833,0.653,0.8 | 0.238 | ENSG00000171 | [[[[63465495, 63 | 2 |
| ENSG00000171 | JMJD1C | chr10 | 63198512 | 63198727 | 63197410 | 63197563 | 63217206 | 63217331 | 11,10,20 | 0,0,0 | 20,49,41 | 0,3,7 | 300 | 150 | 7.00E-05 | 0.001168909 | 1.0,1.0,1.0 | 1.0,0.891,0.745 | 0.121 | ENSG00000171 | [[[[63465495, 63 | 7 |
| ENSG00000096 | HNRNPH3 | chr10 | 68337857 | 68337996 | 68337198 | 68337333 | 68339139 | 68339201 | 187,168,195 | 0,0,2 | 121,296,182 | 78,145,90 | 289 | 150 | 0 | 0 | 1.0,1.0,0.981 | 0.446,0.514,0.51 | 0.503 | ENSG00000096 | [[[[68332064, 68 | 2 |
| ENSG00000096 | HNRNPH3 | chr10 | 68338502 | 68338642 | 68337198 | 68337333 | 68339139 | 68339226 | 95,72,140 | 0,0,2 | 154,313,197 | 78,145,90 | 290 | 150 | 0 | 0 | 1.0,1.0,0.973 | 0.505,0.528,0.53 | 0.47 | ENSG00000096 | [[[[68332064, 68 | 2 |
| ENSG00000096 | HNRNPH3 | chr10 | 68338502 | 68338687 | 68337198 | 68337333 | 68339139 | 68339201 | 205,155,281 | 0,0,2 | 149,315,174 | 78,145,90 | 300 | 150 | 0 | 0 | 1.0,1.0,0.986 | 0.489,0.521,0.49 | 0.495 | ENSG00000096 | [[[[68332064, 68 | 2 |
| ENSG00000023 | ZDHHC6 | chr10 | 112438335 | 112438389 | 112434296 | 112434464 | 112442191 | 112442351 | 28,24,16 | 0,0,0 | 30,62,50 | 5,8,2 | 204 | 150 | 1.12E-05 | 0.000236613 | 1.0,1.0,1.0 | 0.815,0.851,0.94 | 0.129 | ENSG00000023 | [[[[112446705, 1 | 2 |
| ENSG00000060 | CCAR1 | chr10 | 68753851 | 68754077 | 68749523 | 68749685 | 68756272 | 68756483 | 42,37,69 | 0,0,0 | 41,102,62 | 7,38,20 | 300 | 150 | 5.55E-16 | 7.76E-14 | 1.0,1.0,1.0 | 0.745,0.573,0.60 | 0.358 | ENSG00000060 | [[[[68721239, 68 | 3 |
| ENSG00000122 | VPS26A | chr10 | 69155811 | 69155887 | 69132897 | 69133047 | 69171155 | 69171359 | 36,27,50 | 0,0,0 | 44,86,71 | 4,9,7 | 226 | 150 | 6.29E-07 | 1.82E-05 | 1.0,1.0,1.0 | 0.88,0.864,0.871 | 0.128 | ENSG00000122 | [[[[69124178, 69 | 6 |

| | | | | | | | | | | | | | | | | | | | | | | |
|---|---|---|---|---|---|---|---|---|---|---|---|---|---|---|---|---|---|---|---|---|---|---|
| ENSG00000122 | VPS26A | chr10 | 69157006 | 69157163 | 69132897 | 69133047 | 69171155 | 69171260 | 4,3,4 | 0,0,0 | 7,14,14 | 4,9,7 | 300 | 150 | 2.27E-08 | 9.34E-07 | 1.0,1.0,1.0 | 0.467,0.438,0.5 | 0.532 | ENSG00000122 | [[[[69124178, 691 | 6 |
| ENSG00000079 | SAR1A | chr10 | 70161618 | 70161738 | 70161003 | 70161069 | 70170412 | 70170523 | 114,70,173 | 0,1,0 | 121,250,158 | 33,64,39 | 270 | 150 | 0 | 0 | 1.0,0.975,1.0 | 0.671,0.685,0.69 | 0.309 | ENSG00000079 | [[[[70170413, 701 | 2 |
| ENSG00000196 | CD55 | chr1 | 207336692 | 207336818 | 207326751 | 207326837 | 207337328 | 207337409 | 17,19,23 | 0,0,0 | 32,57,46 | 2,4,4 | 276 | 150 | 0.000193538 | 0.002833826 | 1.0,1.0,1.0 | 0.897,0.886,0.86 | 0.118 | ENSG00000196 | [[[[207321678, 20 | 2 |
| ENSG00000138 | ANXA7 | chr10 | 73396518 | 73396583 | 73388311 | 73388414 | 73400802 | 73400857 | 76,65,119 | 0,0,0 | 93,177,132 | 8,4,3 | 215 | 150 | 0.000363571 | 0.00487478 | 1.0,1.0,1.0 | 0.89,0.969,0.968 | 0.058 | ENSG00000138 | [[[[73414012, 734 | 3 |
| ENSG00000138 | ANXA7 | chr10 | 73396518 | 73396583 | 73388311 | 73388414 | 73414011 | 73414067 | 76,65,119 | 0,1,0 | 89,176,125 | 5,6,6 | 215 | 150 | 0.000155904 | 0.002340798 | 1.0,0.978,1.0 | 0.925,0.953,0.93 | 0.055 | ENSG00000138 | [[[[73414012, 734 | 4 |
| ENSG00000088 | ANKRD10 | chr13 | 110883697 | 110883793 | 110878539 | 110880112 | 110906032 | 110906124 | 45,24,35 | 0,0,0 | 45,99,61 | 9,24,12 | 246 | 150 | 1.05E-11 | 8.04E-10 | 1.0,1.0,1.0 | 0.753,0.716,0.75 | 0.258 | ENSG00000088 | [[[[110914725, 11 | 2 |
| ENSG00000138 | RPS24 | chr10 | 78048980 | 78049225 | 78037193 | 78037304 | 78054530 | 78054676 | 0,4,0 | 1,0,3 | 7,10,7 | 0,0,0 | 300 | 150 | 3.98E-05 | 0.000717173 | 0.0,1.0,0.0 | 1.0,1.0,1.0 | -0.667 | ENSG00000138 | [[[[78033863, 780 | 2 |
| ENSG00000111 | TSPAN9 | chr12 | 3206279 | 3206418 | 3201176 | 3201256 | 3280381 | 3280483 | 4,2,3 | 0,0,1 | 2,11,4 | 2,6,0 | 289 | 150 | 0.000164468 | 0.002455361 | 1.0,1.0,0.609 | 0.342,0.488,1.0 | 0.26 | ENSG00000111 | [[[[3077379, 3077 | 2 |
| ENSG00000111 | TSPAN9 | chr12 | 3278991 | 3279066 | 3201176 | 3201256 | 3280381 | 3280471 | 41,32,51 | 0,0,1 | 10,34,19 | 2,6,0 | 225 | 150 | 0.000325102 | 0.004428335 | 1.0,1.0,0.971 | 0.769,0.791,1.0 | 0.137 | ENSG00000111 | [[[[3077379, 3077 | 2 |
| ENSG00000122 | ANXA11 | chr10 | 80164052 | 80164143 | 80163348 | 80163405 | 80166083 | 80166197 | 98,61,73 | 0,0,0 | 70,133,91 | 2,17,2 | 241 | 150 | 6.56E-07 | 1.89E-05 | 1.0,1.0,1.0 | 0.956,0.83,0.966 | 0.083 | ENSG00000122 | [[[[80205343, 802 | 2 |
| ENSG00000078 | NRDC | chr1 | 51803813 | 51803964 | 51798248 | 51798411 | 51805509 | 51805561 | 268,236,330 | 0,0,0 | 104,234,130 | 5,11,4 | 300 | 150 | 3.71E-08 | 1.45E-06 | 1.0,1.0,1.0 | 0.912,0.914,0.94 | 0.077 | ENSG00000078 | [[[[51878275, 518 | 2 |
| ENSG00000078 | NRDC | chr1 | 51819799 | 51819873 | 51814692 | 51814813 | 51821497 | 51821555 | 76,58,133 | 0,0,0 | 36,101,59 | 2,4,4 | 224 | 150 | 0.000590926 | 0.007414358 | 1.0,1.0,1.0 | 0.923,0.944,0.90 | 0.075 | ENSG00000078 | [[[[51878275, 518 | 3 |
| ENSG00000078 | NRDC | chr1 | 51823663 | 51823786 | 51821497 | 51821555 | 51827795 | 51827869 | 105,77,192 | 0,0,0 | 42,134,86 | 8,15,7 | 273 | 150 | 9.49E-13 | 8.63E-11 | 1.0,1.0,1.0 | 0.743,0.831,0.87 | 0.185 | ENSG00000078 | [[[[51878275, 518 | 2 |
| ENSG00000078 | NRDC | chr1 | 51825286 | 51825382 | 51821497 | 51821555 | 51827795 | 51827869 | 148,88,179 | 0,0,0 | 28,106,71 | 8,15,7 | 246 | 150 | 3.38E-14 | 3.76E-12 | 1.0,1.0,1.0 | 0.681,0.812,0.86 | 0.215 | ENSG00000078 | [[[[51878275, 518 | 2 |
| ENSG00000148 | GLUD1 | chr10 | 87076575 | 87076656 | 87068062 | 87068157 | 87094324 | 87094866 | 3,13,21 | 0,0,0 | 8,12,22 | 3,2,7 | 231 | 150 | 1.31E-07 | 4.45E-06 | 1.0,1.0,1.0 | 0.634,0.796,0.67 | 0.3 | ENSG00000148 | [[[[87094325, 870 | 3 |
| ENSG00000049 | RFC2 | chr7 | 74246661 | 74246763 | 74243145 | 74243246 | 74249738 | 74249780 | 64,54,83 | 0,0,1 | 60,104,79 | 6,7,2 | 252 | 150 | 2.69E-05 | 0.000507838 | 1.0,1.0,0.98 | 0.856,0.898,0.95 | 0.089 | ENSG00000049 | [[[[74254271, 742 | 2 |
| ENSG00000049 | RFC2 | chr7 | 74249011 | 74249118 | 74243145 | 74243246 | 74249738 | 74249780 | 44,52,74 | 0,0,1 | 33,57,35 | 6,7,2 | 257 | 150 | 5.56E-07 | 1.63E-05 | 1.0,1.0,0.977 | 0.762,0.826,0.91 | 0.159 | ENSG00000049 | [[[[74254271, 742 | 2 |
| ENSG00000049 | RFC2 | chr7 | 74249011 | 74249220 | 74243145 | 74243246 | 74249738 | 74249780 | 8,11,17 | 0,0,1 | 12,36,18 | 6,7,2 | 300 | 150 | 9.45E-06 | 0.000203761 | 1.0,1.0,0.895 | 0.5,0.72,0.818 | 0.286 | ENSG00000049 | [[[[74254271, 742 | 2 |
| ENSG00000107 | TNKS2 | chr10 | 91848382 | 91848635 | 91842171 | 91842391 | 91849511 | 91849594 | 28,21,51 | 0,0,1 | 47,83,57 | 3,7,5 | 300 | 150 | 2.97E-05 | 0.000554667 | 1.0,1.0,0.962 | 0.887,0.856,0.85 | 0.123 | ENSG00000107 | [[[[91798426, 917 | 3 |
| ENSG00000138 | KIF11 | chr10 | 92632485 | 92632693 | 92630175 | 92630364 | 92637183 | 92637309 | 82,52,100 | 0,1,1 | 20,58,37 | 5,6,10 | 300 | 150 | 5.91E-10 | 3.41E-08 | 1.0,0.963,0.98 | 0.667,0.829,0.64 | 0.266 | ENSG00000138 | [[[[92593130, 925 | 2 |
| ENSG00000148 | FRA10AC1 | chr10 | 93692008 | 93692093 | 93681479 | 93681598 | 93692645 | 93692729 | 12,7,13 | 0,0,0 | 14,30,28 | 1,5,2 | 235 | 150 | 0.000286222 | 0.003963207 | 1.0,1.0,1.0 | 0.899,0.793,0.89 | 0.136 | ENSG00000148 | [[[[93702375, 937 | 4 |
| ENSG00000149 | TMEM25 | chr11 | 118532916 | 118533207 | 118531201 | 118532461 | 118533856 | 118533887 | 6,9,19 | 0,0,0 | 6,13,5 | 4,1,2 | 300 | 150 | 5.71E-06 | 0.000130442 | 1.0,1.0,1.0 | 0.429,0.867,0.55 | 0.383 | ENSG00000149 | [[[[118531192, 11 | 2 |
| ENSG00000149 | RPS3 | chr11 | 75401639 | 75401733 | 75400693 | 75400777 | 75404019 | 75404207 | 6,15,17 | 0,0,0 | 32,108,51 | 3,7,2 | 244 | 150 | 0.000347271 | 0.004681764 | 1.0,1.0,1.0 | 0.868,0.905,0.94 | 0.096 | ENSG00000149 | [[[[75399518, 753 | 2 |
| ENSG00000149 | RPS3 | chr11 | 75401639 | 75401733 | 75400693 | 75400777 | 75404671 | 75404730 | 4,15,16 | 0,0,0 | 16,53,22 | 14,39,28 | 244 | 150 | 6.00E-14 | 6.38E-12 | 1.0,1.0,1.0 | 0.413,0.455,0.32 | 0.602 | ENSG00000149 | [[[[75399518, 753 | 3 |
| ENSG00000149 | RPS3 | chr11 | 75404148 | 75404207 | 75400691 | 75400777 | 75404671 | 75404868 | 96,510,841,544 | 0,0,0 | 131,723,261,361 | 14,39,28 | 209 | 150 | 4.94E-07 | 1.47E-05 | 1.0,1.0,1.0 | 0.985,0.977,0.97 | 0.022 | ENSG00000149 | [[[[75399518, 753 | 3 |
| ENSG00000139 | C12orf10 | chr12 | 53305907 | 53306060 | 53300092 | 53300262 | 53306679 | 53306861 | 2,4,11 | 0,2,0 | 7,20,12 | 4,16,11 | 300 | 150 | 0.000111361 | 0.001752121 | 1.0,0.5,1.0 | 0.467,0.385,0.35 | 0.432 | ENSG00000139 | [[[[53299695, 532 | 3 |
| ENSG00000139 | C12orf10 | chr12 | 53305907 | 53306060 | 53303033 | 53303193 | 53306679 | 53306861 | 91,84,117 | 0,0,0 | 142,199,140 | 22,36,13 | 300 | 150 | 2.22E-16 | 3.17E-14 | 1.0,1.0,1.0 | 0.763,0.734,0.84 | 0.22 | ENSG00000139 | [[[[53299695, 532 | 2 |
| ENSG00000139 | C12orf10 | chr12 | 53306197 | 53306320 | 53299717 | 53300262 | 53306679 | 53306861 | 99,99,133 | 0,2,0 | 151,226,143 | 4,16,11 | 273 | 150 | 3.62E-07 | 1.11E-05 | 1.0,0.965,1.0 | 0.954,0.886,0.87 | 0.083 | ENSG00000139 | [[[[53299695, 532 | 3 |
| ENSG00000139 | C12orf10 | chr12 | 53306197 | 53306320 | 53303033 | 53303193 | 53306679 | 53306861 | 97,99,131 | 0,0,0 | 149,223,138 | 22,36,13 | 273 | 150 | 0 | 0 | 1.0,1.0,1.0 | 0.788,0.773,0.85 | 0.195 | ENSG00000139 | [[[[53299695, 532 | 2 |
| ENSG00000171 | ZDHHC16 | chr10 | 97453798 | 97453846 | 97452414 | 97452503 | 97454713 | 97454799 | 12,11,9 | 3,1,2 | 9,14,12 | 15,20,12 | 198 | 150 | 3.47E-06 | 8.36E-05 | 0.752,0.893,0.77 | 0.313,0.347,0.43 | 0.442 | ENSG00000171 | [[[[97446175, 974 | 3 |
| ENSG00000155 | MMS19 | chr10 | 97460694 | 97460751 | 97460045 | 97460232 | 97461827 | 97461896 | 35,20,34 | 0,0,0 | 16,33,27 | 6,11,1 | 207 | 150 | 5.29E-08 | 1.98E-06 | 1.0,1.0,1.0 | 0.659,0.685,0.95 | 0.235 | ENSG00000155 | [[[[97498273, 974 | 3 |
| ENSG00000075 | SEC31B | chr10 | 100490007 | 100490322 | 100489702 | 100489761 | 100496257 | 100496431 | 7,4,10 | 0,0,0 | 4,11,2 | 1,5,0 | 300 | 150 | 0.000374798 | 0.005008568 | 1.0,1.0,1.0 | 0.667,0.524,1.0 | 0.27 | ENSG00000075 | [[[[100519782, 10 | 3 |
| ENSG00000198 | OGA | chr10 | 101807729 | 101807901 | 101806044 | 101806143 | 101813029 | 101813127 | 27,22,45 | 0,0,0 | 14,46,30 | 4,12,5 | 300 | 150 | 3.87E-10 | 2.31E-08 | 1.0,1.0,1.0 | 0.636,0.657,0.75 | 0.319 | ENSG00000198 | [[[[101817824, 10 | 2 |
| ENSG00000120 | ARMH3 | chr10 | 101939862 | 101939938 | 101889411 | 101889490 | 101975211 | 101975300 | 27,15,17 | 0,0,0 | 7,28,15 | 5,4,7 | 226 | 150 | 9.17E-09 | 4.10E-07 | 1.0,1.0,1.0 | 0.482,0.823,0.58 | 0.369 | ENSG00000120 | [[[[102056085, 10 | 3 |
| ENSG00000120 | ARMH3 | chr10 | 102023674 | 102023749 | 102013967 | 102014024 | 102025155 | 102025248 | 18,5,14 | 0,0,0 | 7,16,6 | 2,2,3 | 225 | 150 | 4.07E-05 | 0.000731053 | 1.0,1.0,1.0 | 0.7,0.842,0.571 | 0.296 | ENSG00000120 | [[[[102056085, 10 | 2 |
| ENSG00000166 | NOLC1 | chr10 | 102161555 | 102161662 | 102159895 | 102160024 | 102161832 | 102161925 | 90,67,127 | 0,0,0 | 116,210,118 | 1,15,7 | 257 | 150 | 1.06E-06 | 2.89E-05 | 1.0,1.0,1.0 | 0.985,0.891,0.90 | 0.072 | ENSG00000166 | [[[[102152389, 10 | 3 |
| ENSG00000077 | NFKB2 | chr10 | 102396903 | 102397055 | 102396448 | 102396489 | 102397301 | 102397408 | 21,11,19 | 0,0,0 | 13,31,32 | 1,4,2 | 300 | 150 | 0.000579479 | 0.007285088 | 1.0,1.0,1.0 | 0.867,0.795,0.88 | 0.15 | ENSG00000077 | [[[[102395705, 10 | 2 |
| ENSG00000156 | SFXN2 | chr10 | 102732151 | 102732218 | 102729722 | 102729808 | 102732858 | 102732908 | 7,10,20 | 0,0,0 | 4,6,4 | 4,1,1 | 217 | 150 | 0.000109012 | 0.001720126 | 1.0,1.0,1.0 | 0.409,0.806,0.73 | 0.35 | ENSG00000156 | [[[[102714636, 10 | 2 |
| ENSG00000123 | ADGRE5 | chr19 | 14396341 | 14396473 | 14390923 | 14391079 | 14397657 | 14397804 | 20,14,26 | 30,40,39 | 31,97,70 | 23,61,39 | 282 | 150 | 8.09E-07 | 2.27E-05 | 0.262,0.157,0.26 | 0.418,0.458,0.48 | -0.228 | ENSG00000123 | [[[[14381444, 14 | 2 |
| ENSG00000161 | MFSD12 | chr19 | 3542775 | 3542977 | 3538274 | 3538756 | 3547271 | 3547364 | 3,14,14 | 0,0,0 | 30,77,46 | 9,18,9 | 300 | 150 | 7.27E-08 | 2.64E-06 | 1.0,1.0,1.0 | 0.625,0.681,0.71 | 0.325 | ENSG00000161 | [[[[3557106, 3557 | 4 |
| ENSG00000161 | MFSD12 | chr19 | 3544808 | 3544939 | 3544198 | 3544732 | 3547271 | 3547364 | 2,5,17 | 0,0,0 | 4,13,15 | 9,14,12 | 281 | 150 | 8.56E-12 | 6.59E-10 | 1.0,1.0,1.0 | 0.192,0.331,0.4 | 0.692 | ENSG00000161 | [[[[3557106, 3557 | 3 |
| ENSG00000161 | MFSD12 | chr19 | 3546073 | 3546168 | 3544675 | 3544939 | 3547271 | 3547364 | 7,14,35 | 0,0,0 | 20,48,34 | 0,6,3 | 245 | 150 | 0.000148994 | 0.00225125 | 1.0,1.0,1.0 | 1.0,0.83,0.874 | 0.099 | ENSG00000161 | [[[[3557106, 3557 | 2 |
| ENSG00000165 | PDZD8 | chr10 | 117318871 | 117318974 | 117290185 | 117290348 | 117374355 | 117375467 | 20,18,36 | 0,0,0 | 10,63,35 | 2,5,2 | 253 | 150 | 0.000163889 | 0.002448149 | 1.0,1.0,1.0 | 0.748,0.882,0.91 | 0.153 | ENSG00000165 | [[[[117374356, 11 | 2 |
| ENSG00000165 | PDZD8 | chr10 | 117340979 | 117341102 | 117290185 | 117290348 | 117374355 | 117375467 | 12,10,21 | 0,0,0 | 26,73,42 | 2,5,2 | 273 | 150 | 0.000817213 | 0.009781847 | 1.0,1.0,1.0 | 0.877,0.889,0.92 | 0.105 | ENSG00000165 | [[[[117374356, 11 | 2 |
| ENSG00000159 | CYB5R1 | chr1 | 202965886 | 202965993 | 202964611 | 202964695 | 202966527 | 202966560 | 41,52,65 | 0,0,0 | 56,78,74 | 25,38,18 | 257 | 150 | 0 | 0 | 1.0,1.0,1.0 | 0.567,0.545,0.70 | 0.394 | ENSG00000159 | [[[[202967191, 20 | 2 |
| ENSG00000159 | CYB5R1 | chr1 | 202966748 | 202966898 | 202964611 | 202964695 | 202967190 | 202967225 | 3,10,46 | 0,0,0 | 19,33,42 | 2,7,0 | 300 | 150 | 5.20E-05 | 0.000903585 | 1.0,1.0,1.0 | 0.826,0.702,1.0 | 0.157 | ENSG00000159 | [[[[202967191, 20 | 4 |
| ENSG00000154 | BUB3 | chr10 | 123162235 | 123162413 | 123157728 | 123157880 | 123162611 | 123162693 | 237,287,335 | 0,0,0 | 130,309,165 | 46,121,55 | 300 | 150 | 0 | 0 | 1.0,1.0,1.0 | 0.586,0.561,0.6 | 0.418 | ENSG00000154 | [[[[123154402, 12 | 2 |
| ENSG00000176 | SEC24C | chr10 | 73746804 | 73747004 | 73746629 | 73746716 | 73759621 | 73759794 | 19,10,29 | 0,0,0 | 2,15,11 | 4,5,4 | 300 | 150 | 4.17E-10 | 2.47E-08 | 1.0,1.0,1.0 | 0.2,0.6,0.579 | 0.54 | ENSG00000176 | [[[[73744372, 737 | 2 |
| ENSG00000176 | SEC24C | chr10 | 73751107 | 73751243 | 73746629 | 73746716 | 73759621 | 73759794 | 32,35,135 | 0,0,0 | 23,74,40 | 4,5,4 | 286 | 150 | 4.75E-07 | 1.42E-05 | 1.0,1.0,1.0 | 0.751,0.886,0.84 | 0.174 | ENSG00000176 | [[[[73744372, 737 | 2 |
| ENSG00000176 | SEC24C | chr10 | 73767836 | 73768007 | 73763855 | 73763983 | 73768809 | 73768906 | 87,68,135 | 0,0,0 | 23,60,57 | 0,5,5 | 300 | 150 | 2.53E-05 | 0.000482611 | 1.0,1.0,1.0 | 1.0,0.857,0.851 | 0.097 | ENSG00000176 | [[[[73744372, 737 | 7 |
| ENSG00000148 | MTG1 | chr10 | 133399528 | 133399619 | 133399169 | 133399226 | 133402148 | 133402245 | 86,81,148 | 2,0,1 | 88,143,104 | 9,18,10 | 241 | 150 | 7.39E-10 | 4.19E-08 | 0.964,1.0,0.989 | 0.859,0.832,0.86 | 0.132 | ENSG00000148 | [[[[133394157, 13 | 2 |
| ENSG00000148 | MTG1 | chr10 | 133399528 | 133399653 | 133399169 | 133399226 | 133402148 | 133402773 | 89,90,154 | 2,0,1 | 66,132,92 | 9,18,10 | 275 | 150 | 2.56E-11 | 1.84E-09 | 0.96,1.0,0.988 | 0.8,0.8,0.834 | 0.171 | ENSG00000148 | [[[[133394157, 13 | 2 |
| ENSG00000148 | MTG1 | chr10 | 133401528 | 133401590 | 133399169 | 133399226 | 133402148 | 133402245 | 47,48,78 | 2,0,1 | 38,79,46 | 9,18,10 | 212 | 150 | 1.02E-10 | 6.71E-09 | 0.943,1.0,0.982 | 0.749,0.756,0.76 | 0.218 | ENSG00000148 | [[[[133394157, 13 | 2 |
| ENSG00000177 | ARIH2 | chr3 | 48927461 | 48927813 | 48918820 | 48918998 | 48964918 | 48964982 | 1,0,0 | 0,1,1 | 7,12,5 | 0,0,0 | 300 | 150 | 0.000271374 | 0.003785121 | 1.0,0.0,0.0 | 1.0,1.0,1.0 | -0.667 | ENSG00000177 | [[[[48918842, 489 | 3 |
| ENSG00000177 | ARIH2 | chr3 | 48945135 | 48945181 | 48926717 | 48927813 | 48968533 | 48968655 | 2,1,1 | 0,0,0 | 9,17,4 | 0,7,2 | 196 | 150 | 2.71E-05 | 0.000511596 | 1.0,1.0,1.0 | 1.0,0.65,0.605 | 0.248 | ENSG00000177 | [[[[48918842, 489 | 3 |
| ENSG00000177 | ARIH2 | chr3 | 48964918 | 48964982 | 48926717 | 48927813 | 48967124 | 48967151 | 33,26,47 | 0,0,1 | 28,67,43 | 3,8,8 | 214 | 150 | 7.67E-07 | 2.17E-05 | 1.0,1.0,0.971 | 0.867,0.854,0.79 | 0.153 | ENSG00000177 | [[[[48918842, 489 | 2 |
| ENSG00000142 | SIRT3 | chr11 | 230451 | 230552 | 218831 | 219041 | 232982 | 233215 | 8,23,36 | 0,0,0 | 9,12,13 | 0,9,2 | 251 | 150 | 7.63E-06 | 0.000168615 | 1.0,1.0,1.0 | 1.0,0.443,0.795 | 0.254 | ENSG00000142 | [[[[236048, 23634 | 2 |
| ENSG00000177 | DEAF1 | chr11 | 679687 | 679816 | 678693 | 678822 | 684897 | 684963 | 2,12,30 | 0,0,0 | 9,9,16 | 2,6,2 | 279 | 150 | 1.52E-07 | 5.08E-06 | 1.0,1.0,1.0 | 0.708,0.446,0.81 | 0.345 | ENSG00000177 | [[[[694759, 69522 | 2 |
| ENSG00000145 | RNF145 | chr5 | 159168872 | 159169055 | 159162931 | 159163079 | 159194715 | 159194824 | 228,172,285 | 0,0,0 | 104,204,173 | 6,21,16 | 300 | 150 | 1.11E-16 | 1.65E-14 | 1.0,1.0,1.0 | 0.897,0.829,0.84 | 0.143 | ENSG00000145 | [[[[159209231, 15 | 5 |
| ENSG00000145 | RNF145 | chr5 | 159173982 | 159174158 | 159169678 | 159169819 | 159194715 | 159194824 | 188,128,251 | 0,1,2 | 93,185,151 | 8,26,10 | 300 | 150 | 6.81E-14 | 7.21E-12 | 1.0,0.985,0.984 | 0.853,0.781,0.88 | 0.151 | ENSG00000145 | [[[[159209231, 15 | 3 |
| ENSG00000145 | RNF145 | chr5 | 159176631 | 159176867 | 159162931 | 159163079 | 159194715 | 159194824 | 2,1,3 | 0,0,0 | 1,14,7 | 6,21,16 | 300 | 150 | 3.01E-08 | 1.21E-06 | 1.0,1.0,1.0 | 0.077,0.25,0.179 | 0.831 | ENSG00000145 | [[[[159209231, 15 | 5 |
| ENSG00000145 | RNF145 | chr5 | 159176631 | 159176867 | 159173982 | 159174158 | 159194715 | 159194824 | 165,131,250 | 1,0,0 | 46,148,95 | 23,65,42 | 300 | 150 | 0 | 0 | 0.988,1.0,1.0 | 0.5,0.532,0.531 | 0.475 | ENSG00000145 | [[[[159209231, 15 | 2 |
| ENSG00000145 | RNF145 | chr5 | 159181959 | 159182051 | 159162931 | 159163079 | 159194715 | 159194824 | 177,105,251 | 0,0,0 | 151,256,172 | 6,21,16 | 242 | 150 | 6.82E-13 | 6.34E-11 | 1.0,1.0,1.0 | 0.94,0.883,0.87 | 0.102 | ENSG00000145 | [[[[159209231, 15 | 5 |
| ENSG00000145 | RNF145 | chr5 | 159181959 | 159182051 | 159169678 | 159169819 | 159194715 | 159194824 | 177,105,251 | 0,1,2 | 151,257,169 | 8,26,10 | 242 | 150 | 5.69E-11 | 3.86E-09 | 1.0,0.985,0.987 | 0.921,0.86,0.913 | 0.093 | ENSG00000145 | [[[[159209231, 15 | 3 |
| ENSG00000145 | RNF145 | chr5 | 159181959 | 159182051 | 159173982 | 159174158 | 159194715 | 159194824 | 177,105,251 | 1,0,0 | 161,281,191 | 23,65,42 | 242 | 150 | 0 | 0 | 0.991,1.0,1.0 | 0.813,0.728,0.73 | 0.237 | ENSG00000145 | [[[[159209231, 15 | 2 |
| ENSG00000104 | GYS1 | chr19 | 48981529 | 48981636 | 48978097 | 48978157 | 48985460 | 48985605 | 47,36,81 | 0,0,0 | 46,88,53 | 1,7,5 | 257 | 150 | 1.12E-05 | 0.000236865 | 1.0,1.0,1.0 | 0.964,0.88,0.861 | 0.098 | ENSG00000104 | [[[[48992995, 489 | 3 |
| ENSG00000119 | PPP6C | chr9 | 125153532 | 125153742 | 125146572 | 125149921 | 125171084 | 125171180 | 109,156,167 | 0,0,0 | 159,314,203 | 3,13,6 | 300 | 150 | 5.33E-06 | 0.00012296 | 1.0,1.0,1.0 | 0.964,0.924,0.94 | 0.056 | ENSG00000119 | [[[[125189644, 12 | 4 |
| ENSG00000119 | PPP6C | chr9 | 125158240 | 125158382 | 125153905 | 125153985 | 125171084 | 125171180 | 126,113,165 | 0,4,5 | 149,270,141 | 30,61,40 | 292 | 150 | 0 | 0 | 1.0,0.936,0.944 | 0.718,0.695,0.64 | 0.274 | ENSG00000119 | [[[[125189644, 12 | 2 |
| ENSG00000119 | PPP6C | chr9 | 125160840 | 125160906 | 125146572 | 125149921 | 125171084 | 125171180 | 59,64,91 | 0,0,0 | 61,139,75 | 3,13,6 | 216 | 150 | 1.48E-07 | 4.95E-06 | 1.0,1.0,1.0 | 0.934,0.881,0.89 | 0.096 | ENSG00000119 | [[[[125189644, 12 | 4 |
| ENSG00000119 | PPP6C | chr9 | 125160840 | 125160906 | 125153905 | 125153985 | 125171084 | 125171180 | 59,64,92 | 0,4,5 | 62,137,74 | 30,61,40 | 216 | 150 | 0 | 0 | 1.0,0.917,0.927 | 0.589,0.609,0.56 | 0.361 | ENSG00000119 | [[[[125189644, 12 | 2 |
| ENSG00000005 | ZNF195 | chr11 | 3369356 | 3369502 | 3360477 | 3360565 | 3370974 | 3371070 | 1,2,5 | 16,13,15 | 12,17,19 | 15,41,27 | 296 | 150 | 0.000692979 | 0.008496153 | 0.031,0.072,0.14 | 0.288,0.174,0.26 | -0.159 | ENSG00000005 | [[[[3379038, 3379 | 2 |
| ENSG00000130 | HABP4 | chr9 | 96471010 | 96471094 | 96458378 | 96458541 | 96484461 | 96484633 | 31,17,33 | 0,0,0 | 36,37,26 | 1,5,3 | 234 | 150 | 0.000150478 | 0.002270067 | 1.0,1.0,1.0 | 0.958,0.826,0.84 | 0.123 | ENSG00000130 | [[[[96450169, 964 | 3 |
| ENSG00000132 | ARFIP2 | chr11 | 6478737 | 6478959 | 6478040 | 6478198 | 6479971 | 6480068 | 71,98,137 | 0,0,0 | 115,204,107 | 13,19,14 | 300 | 150 | 2.27E-13 | 2.25E-11 | 1.0,1.0,1.0 | 0.816,0.843,0.79 | 0.183 | ENSG00000132 | [[[[6481231, 6481 | 2 |
| ENSG00000166 | ILK | chr11 | 6608045 | 6608211 | 6603815 | 6604360 | 6608883 | 6608967 | 48,93,198 | 0,0,0 | 70,167,99 | 2,11,14 | 300 | 150 | 1.18E-09 | 6.42E-08 | 1.0,1.0,1.0 | 0.946,0.884,0.78 | 0.13 | ENSG00000166 | [[[[6603774, 6603 | 3 |
| ENSG00000166 | ILK | chr11 | 6608393 | 6608489 | 6604225 | 6604360 | 6608883 | 6608967 | 1,2,1 | 0,0,0 | 11,19,8 | 2,11,14 | 246 | 150 | 6.69E-08 | 2.45E-06 | 1.0,1.0,1.0 | 0.77,0.513,0.258 | 0.486 | ENSG00000166 | [[[[6603774, 6603 | 3 |
| ENSG00000166 | ILK | chr11 | 6608693 | 6608790 | 6603815 | 6604360 | 6608883 | 6608967 | 110,102,171 | 0,0,0 | 94,206,125 | 2,11,14 | 247 | 150 | 1.11E-08 | 4.90E-07 | 1.0,1.0,1.0 | 0.966,0.919,0.84 | 0.09 | ENSG00000166 | [[[[6603774, 6603 | 3 |
| ENSG00000125 | CNOT1 | chr16 | 58530245 | 58530347 | 58525178 | 58525359 | 58531957 | 58532075 | 48,41,55 | 0,0,0 | 50,88,74 | 3,8,8 | 252 | 150 | 3.92E-07 | 1.19E-05 | 1.0,1.0,1.0 | 0.908,0.868,0.84 | 0.126 | ENSG00000125 | [[[[58629728, 586 | 3 |
| ENSG00000125 | CNOT1 | chr16 | 58542230 | 58542335 | 58541500 | 58541620 | 58543606 | 58543903 | 92,57,100 | 0,2,0 | 61,139,59 | 13,33,19 | 255 | 150 | 0 | 0 | 1.0,0.944,1.0 | 0.734,0.712,0.64 | 0.284 | ENSG00000125 | [[[[58629728, 586 | 2 |
| ENSG00000125 | CNOT1 | chr16 | 58586544 | 58586748 | 58585337 | 58585506 | 58587344 | 58587413 | 69,30,102 | 0,0,0 | 32,114,92 | 6,5,6 | 300 | 150 | 7.07E-07 | 2.02E-05 | 1.0,1.0,1.0 | 0.727,0.919,0.88 | 0.156 | ENSG00000125 | [[[[58629728, 586 | 2 |
| ENSG00000166 | TMEM41B | chr11 | 9283370 | 9283593 | 9280653 | 9281041 | 9288441 | 9288535 | 8,7,12 | 0,0,0 | 47,86,42 | 6,16,5 | 300 | 150 | 1.63E-06 | 4.25E-05 | 1.0,1.0,1.0 | 0.797,0.729,0.80 | 0.222 | ENSG00000166 | [[[[9314321, 931 | 2 |

| | | | | | | | | | | | | | | | | | | | | | | |
|---|---|---|---|---|---|---|---|---|---|---|---|---|---|---|---|---|---|---|---|---|---|---|
| ENSG00000166 | TMEM41B | chr11 | 9287701 | 9287806 | 9283546 | 9283593 | 9288441 | 9288535 | 10,15,19 | 0,0,1 | 15,21,20 | 10,15,5 | 255 | 150 | 3.66E-09 | 1.80E-07 | 1.0,1.0,0.918 | 0.469,0.452,0.70 | 0.432 | ENSG00000166 | [[[9314321, 9314 | 2 |
| ENSG00000137 | PPP2R1B | chr11 | 111743375 | 111743530 | 111742052 | 111742144 | 111747953 | 111747994 | 43,26,70 | 0,0,0 | 24,93,47 | 4,6,8 | 300 | 150 | 7.25E-08 | 2.64E-06 | 1.0,1.0,1.0 | 0.75,0.886,0.746 | 0.206 | ENSG00000137 | [[[111766248, 11 | 2 |
| ENSG00000137 | PPP2R1B | chr11 | 111764804 | 111764905 | 111760818 | 111761051 | 111766247 | 111766368 | 22,28,59 | 0,0,0 | 23,75,40 | 3,3,6 | 251 | 150 | 1.19E-05 | 0.000248782 | 1.0,1.0,1.0 | 0.821,0.937,0.79 | 0.148 | ENSG00000137 | [[[111766248, 11 | 2 |
| ENSG00000137 | PPP2R1B | chr11 | 111765293 | 111765384 | 111760818 | 111761051 | 111766247 | 111766367 | 25,49,55 | 0,0,0 | 44,87,66 | 3,3,6 | 241 | 150 | 0.000121893 | 0.001893577 | 1.0,1.0,1.0 | 0.901,0.948,0.87 | 0.093 | ENSG00000137 | [[[111766248, 11 | 2 |
| ENSG00000148 | BTBD10 | chr11 | 13445023 | 13445181 | 13405658 | 13405856 | 13463091 | 13463226 | 16,13,41 | 1,0,0 | 34,49,28 | 2,6,2 | 300 | 150 | 0.000253648 | 0.003566611 | 0.889,1.0,1.0 | 0.895,0.803,0.87 | 0.105 | ENSG00000148 | [[[13463092, 134 | 5 |
| ENSG00000113 | LNPEP | chr5 | 96998013 | 96998145 | 96993816 | 96993971 | 97003414 | 97003546 | 7,4,6 | 0,0,0 | 16,20,11 | 0,7,4 | 282 | 150 | 3.31E-06 | 8.01E-05 | 1.0,1.0,1.0 | 1.0,0.603,0.594 | 0.268 | ENSG00000113 | [[[96936080, 969 | 2 |
| ENSG00000166 | SAAL1 | chr11 | 18083534 | 18083731 | 18081410 | 18081503 | 18087142 | 18087225 | 44,30,46 | 0,0,0 | 37,113,48 | 8,21,1 | 300 | 150 | 2.37E-08 | 9.71E-07 | 1.0,1.0,1.0 | 0.698,0.729,0.96 | 0.204 | ENSG00000166 | [[[18105907, 181 | 2 |
| ENSG00000166 | SAAL1 | chr11 | 18086865 | 18087054 | 18081410 | 18081503 | 18087142 | 18087206 | 58,49,56 | 0,0,0 | 102,174,73 | 8,21,1 | 300 | 150 | 1.03E-07 | 3.62E-06 | 1.0,1.0,1.0 | 0.864,0.806,0.97 | 0.119 | ENSG00000166 | [[[18105907, 181 | 2 |
| ENSG00000166 | SAAL1 | chr11 | 18087142 | 18087225 | 18086856 | 18087054 | 18090433 | 18090493 | 38,23,38 | 0,0,0 | 81,113,66 | 4,9,3 | 233 | 150 | 4.42E-05 | 0.000784417 | 1.0,1.0,1.0 | 0.929,0.89,0.934 | 0.082 | ENSG00000166 | [[[18105907, 181 | 3 |
| ENSG00000110 | GTF2H1 | chr11 | 18341527 | 18341607 | 18338108 | 18338274 | 18347587 | 18347715 | 26,27,36 | 1,0,0 | 34,68,53 | 7,16,3 | 230 | 150 | 3.35E-07 | 1.04E-05 | 0.944,1.0,1.0 | 0.76,0.735,0.92 | 0.176 | ENSG00000110 | [[[18322567, 183 | 3 |
| ENSG00000110 | GTF2H1 | chr11 | 18351880 | 18351969 | 18347831 | 18347919 | 18358524 | 18358640 | 24,15,19 | 0,0,0 | 11,57,25 | 4,4,2 | 239 | 150 | 3.49E-05 | 0.000639962 | 1.0,1.0,1.0 | 0.633,0.899,0.88 | 0.194 | ENSG00000110 | [[[18322567, 183 | 3 |
| ENSG00000110 | GTF2H1 | chr11 | 18351880 | 18351969 | 18347831 | 18347919 | 18365782 | 18366261 | 24,15,19 | 0,0,0 | 13,47,28 | 2,4,5 | 239 | 150 | 8.56E-06 | 0.00018635 | 1.0,1.0,1.0 | 0.803,0.881,0.77 | 0.179 | ENSG00000110 | [[[18322567, 183 | 5 |
| ENSG00000110 | GTF2H1 | chr11 | 18352328 | 18352446 | 18347831 | 18347919 | 18357951 | 18358042 | 69,40,96 | 0,0,0 | 30,66,23 | 20,40,21 | 268 | 150 | 0 | 0 | 1.0,1.0,1.0 | 0.456,0.48,0.38 | 0.561 | ENSG00000110 | [[[18322567, 183 | 2 |
| ENSG00000110 | GTF2H1 | chr11 | 18360614 | 18360707 | 18347831 | 18347919 | 18365782 | 18365898 | 42,29,41 | 0,0,0 | 37,76,43 | 2,4,5 | 243 | 150 | 0.00010205 | 0.0016244 | 1.0,1.0,1.0 | 0.919,0.921,0.84 | 0.106 | ENSG00000110 | [[[18322567, 183 | 5 |
| ENSG00000074 | TSG101 | chr11 | 18519518 | 18519603 | 18509541 | 18509665 | 18526774 | 18526913 | 3,12,43 | 0,0,0 | 4,17,15 | 17,30,12 | 235 | 150 | 0 | 0 | 1.0,1.0,1.0 | 0.131,0.266,0.44 | 0.72 | ENSG00000074 | [[[18526775, 185 | 3 |
| ENSG00000164 | GRIK2 | chr6 | 101682552 | 101682606 | 101676622 | 101676804 | 101799647 | 101799791 | 3,1,3 | 0,0,0 | 0,1,2 | 3,3,3 | 204 | 150 | 3.57E-13 | 3.42E-11 | 1.0,1.0,1.0 | 0.0,0.197,0.329 | 0.825 | ENSG00000164 | [[[101393708, 10 | 2 |
| ENSG00000130 | ZNF317 | chr19 | 9156611 | 9156748 | 9140428 | 9140592 | 9157267 | 9157394 | 8,8,26 | 0,0,0 | 15,48,34 | 6,43,29 | 287 | 150 | 2.00E-13 | 2.01E-11 | 1.0,1.0,1.0 | 0.566,0.368,0.38 | 0.562 | ENSG00000130 | [[[9140397, 9140 | 2 |
| ENSG00000130 | ZNF317 | chr19 | 9158825 | 9158908 | 9157267 | 9157394 | 9163054 | 9163424 | 2,2,2 | 0,0,0 | 8,22,20 | 16,34,16 | 233 | 150 | 8.93E-08 | 3.18E-06 | 1.0,1.0,1.0 | 0.244,0.294,0.44 | 0.672 | ENSG00000130 | [[[9140397, 9140 | 2 |
| ENSG00000148 | IMMP1L | chr11 | 31460625 | 31460714 | 31456259 | 31456386 | 31509518 | 31509574 | 9,3,10 | 7,6,7 | 11,12,23 | 1,0,2 | 239 | 150 | 9.40E-08 | 3.33E-06 | 0.447,0.239,0.47 | 0.873,1.0,0.878 | -0.531 | ENSG00000148 | [[[31509519, 315 | 2 |
| ENSG00000148 | IMMP1L | chr11 | 31463171 | 31463305 | 31433459 | 31433570 | 31509518 | 31509574 | 23,25,34 | 54,63,69 | 17,53,20 | 20,25,16 | 284 | 150 | 3.10E-05 | 0.000576512 | 0.184,0.173,0.20 | 0.31,0.528,0.398 | -0.224 | ENSG00000148 | [[[31509519, 315 | 3 |
| ENSG00000148 | IMMP1L | chr11 | 31463171 | 31463305 | 31456312 | 31456386 | 31509518 | 31509574 | 16,18,25 | 7,6,7 | 15,43,16 | 1,0,2 | 284 | 150 | 2.51E-06 | 6.24E-05 | 0.547,0.613,0.65 | 0.888,1.0,0.809 | -0.294 | ENSG00000148 | [[[31509519, 315 | 2 |
| ENSG00000007 | PAX6 | chr11 | 31793437 | 31793553 | 31790709 | 31790860 | 31794031 | 31794114 | 9,18,26 | 0,0,0 | 17,38,25 | 3,8,5 | 266 | 150 | 1.17E-07 | 4.04E-06 | 1.0,1.0,1.0 | 0.762,0.728,0.73 | 0.257 | ENSG00000007 | [[[31811115, 318 | 2 |
| ENSG00000007 | PAX6 | chr11 | 31793651 | 31793802 | 31790709 | 31790860 | 31794031 | 31794114 | 24,19,34 | 0,0,0 | 32,53,33 | 3,8,5 | 300 | 150 | 2.66E-07 | 8.42E-06 | 1.0,1.0,1.0 | 0.842,0.768,0.76 | 0.208 | ENSG00000007 | [[[31811115, 318 | 2 |
| ENSG00000275 | MLLT6 | chr17 | 38717431 | 38717613 | 38715611 | 38715828 | 38717844 | 38717953 | 7,6,27 | 0,0,0 | 17,33,34 | 2,4,4 | 300 | 150 | 2.05E-05 | 0.000402475 | 1.0,1.0,1.0 | 0.81,0.805,0.81 | 0.192 | ENSG00000275 | [[[38705273, 387 | 2 |
| ENSG00000110 | HIPK3 | chr11 | 33348165 | 33348228 | 33347851 | 33348013 | 33351607 | 33351843 | 1,3,5 | 0,0,0 | 7,15,7 | 5,3,3 | 213 | 150 | 1.42E-07 | 4.78E-06 | 1.0,1.0,1.0 | 0.496,0.779,0.62 | 0.368 | ENSG00000110 | [[[33257381, 332 | 3 |
| ENSG00000110 | HIPK3 | chr11 | 33348521 | 33348818 | 33347851 | 33348013 | 33351607 | 33351843 | 15,5,24 | 0,0,0 | 9,33,45 | 5,3,3 | 300 | 150 | 5.29E-06 | 0.000121982 | 1.0,1.0,1.0 | 0.474,0.846,0.88 | 0.266 | ENSG00000110 | [[[33257381, 332 | 3 |
| ENSG00000110 | HIPK3 | chr11 | 33348521 | 33348818 | 33348165 | 33348228 | 33351607 | 33351843 | 6,6,9 | 0,0,0 | 7,24,29 | 2,4,1 | 300 | 150 | 0.000291241 | 0.004025397 | 1.0,1.0,1.0 | 0.636,0.75,0.935 | 0.226 | ENSG00000110 | [[[33257381, 332 | 2 |
| ENSG00000135 | CAPRIN1 | chr11 | 34082954 | 34083041 | 34082824 | 34082877 | 34086304 | 34086413 | 270,201,413 | 0,0,0 | 260,590,418 | 6,15,5 | 237 | 150 | 0.000134347 | 0.002056454 | 1.0,1.0,1.0 | 0.965,0.961,0.98 | 0.031 | ENSG00000135 | [[[34051731, 340 | 2 |
| ENSG00000135 | CAPRIN1 | chr11 | 34086063 | 34086219 | 34082824 | 34082877 | 34086304 | 34086413 | 235,210,421 | 0,0,0 | 229,524,359 | 6,15,5 | 300 | 150 | 4.48E-06 | 0.000104831 | 1.0,1.0,1.0 | 0.95,0.946,0.973 | 0.044 | ENSG00000135 | [[[34051731, 340 | 2 |
| ENSG00000135 | CAPRIN1 | chr11 | 34090528 | 34090678 | 34090025 | 34090289 | 34096478 | 34096673 | 411,352,569 | 0,0,0 | 352,851,537 | 12,12,5 | 300 | 150 | 2.58E-05 | 0.000489965 | 1.0,1.0,1.0 | 0.936,0.973,0.98 | 0.036 | ENSG00000135 | [[[34051731, 340 | 2 |
| ENSG00000110 | PDHX | chr11 | 34966639 | 34966814 | 34947505 | 34947606 | 34970138 | 34970282 | 147,131,181 | 0,0,0 | 157,347,217 | 10,35,17 | 300 | 150 | 6.44E-15 | 7.90E-13 | 1.0,1.0,1.0 | 0.887,0.832,0.86 | 0.139 | ENSG00000110 | [[[34916618, 349 | 3 |
| ENSG00000026 | CD44 | chr11 | 35189834 | 35189972 | 35176574 | 35176740 | 35214851 | 35214914 | 7,11,12 | 0,0,0 | 9,22,20 | 1,2,5 | 288 | 150 | 1.53E-05 | 0.00030966 | 1.0,1.0,1.0 | 0.824,0.851,0.67 | 0.216 | ENSG00000026 | [[[35139171, 351 | 12 |
| ENSG00000026 | CD44 | chr11 | 35198120 | 35198246 | 35189834 | 35190065 | 35201081 | 35201195 | 57,35,68 | 4,1,2 | 77,145,117 | 11,29,25 | 276 | 150 | 7.76E-05 | 0.886,0.95,0.949 | 0.792,0.731,0.71 | | 0.181 | ENSG00000026 | [[[35139171, 351 | 2 |
| ENSG00000026 | CD44 | chr11 | 35198120 | 35198246 | 35189834 | 35190065 | 35214851 | 35214914 | 87,56,102 | 175,612,592,357 | 108,188,176 | 162,630,331,951 | 276 | 150 | 7.39E-05 | 0.001223468 | 0.026,0.024,0.02 | 0.035,0.033,0.04 | -0.014 | ENSG00000026 | [[[35139171, 351 | 9 |
| ENSG00000026 | CD44 | chr11 | 35201081 | 35201195 | 35189834 | 35190065 | 35206111 | 35206243 | 4,1,2 | 1,0,0 | 8,23,20 | 0,6,6 | 264 | 150 | 0.000444402 | 0.005807333 | 0.694,1.0,1.0 | 1.0,0.685,0.654 | 0.118 | ENSG00000026 | [[[35139171, 351 | 5 |
| ENSG00000026 | CD44 | chr11 | 35201081 | 35201195 | 35197856 | 35198246 | 35214851 | 35214914 | 2,0,3 | 34,22,39 | 4,30,5 | 35,79,66 | 264 | 150 | 0.000181559 | 0.002679586 | 0.032,0.0,0.042 | 0.061,0.177,0.04 | -0.068 | ENSG00000026 | [[[35139171, 351 | 7 |
| ENSG00000026 | CD44 | chr11 | 35201667 | 35201787 | 35189834 | 35190065 | 35204511 | 35204640 | 8,3,8 | 55,47,88 | 33,49,45 | 53,98,105 | 270 | 150 | 1.48E-11 | 1.10E-09 | 0.075,0.034,0.04 | 0.257,0.217,0.19 | -0.17 | ENSG00000026 | [[[35139171, 351 | 4 |
| ENSG00000026 | CD44 | chr11 | 35201670 | 35201787 | 35198120 | 35198246 | 35214851 | 35214914 | 1,1,3 | 34,22,39 | 20,21,14 | 35,79,66 | 267 | 150 | 6.47E-07 | 1.86E-05 | 0.016,0.025,0.04 | 0.243,0.13,0.106 | -0.132 | ENSG00000026 | [[[35139171, 351 | 7 |
| ENSG00000026 | CD44 | chr11 | 35204511 | 35204640 | 35201667 | 35201787 | 35214851 | 35214914 | 31,16,36 | 1,3,3 | 57,92,76 | 24,27,17 | 279 | 150 | 0.000374487 | 0.005005286 | 0.943,0.741,0.86 | 0.561,0.647,0.70 | 0.212 | ENSG00000026 | [[[35139171, 351 | 5 |
| ENSG00000026 | CD44 | chr11 | 35208104 | 35208206 | 35189834 | 35189972 | 35214851 | 35214914 | 0,0,2 | 9,11,12 | 4,2,5 | 4,14,15 | 252 | 150 | 0.000467366 | 0.00607409 | 0.0,0.0,0.09 | 0.373,0.078,0.16 | -0.176 | ENSG00000026 | [[[35139171, 351 | 9 |
| ENSG00000026 | CD44 | chr11 | 35208104 | 35208206 | 35198120 | 35198246 | 35211245 | 35211253 | 4,7,10 | 0,0,1 | 0,10,5 | 3,3,3 | 252 | 150 | 2.54E-06 | 6.32E-05 | 1.0,1.0,0.856 | 0.0,0.665,0.498 | 0.564 | ENSG00000026 | [[[35139171, 351 | 6 |
| ENSG00000026 | CD44 | chr11 | 35211245 | 35211449 | 35201620 | 35201787 | 35214851 | 35214861 | 159,135,176 | 1,3,3 | 142,290,179 | 24,27,17 | 300 | 150 | 6.50E-11 | 4.37E-09 | 0.988,0.957,0.96 | 0.747,0.843,0.84 | 0.161 | ENSG00000026 | [[[35139171, 351 | 5 |
| ENSG00000103 | BFAR | chr16 | 14649803 | 14649973 | 14644287 | 14644609 | 14661891 | 14662065 | 3,4,10 | 2,4,5 | 11,23,16 | 1,2,0 | 300 | 150 | 7.23E-06 | 0.000160178 | 0.429,0.333,0.5 | 0.846,0.852,1.0 | -0.479 | ENSG00000103 | [[[14632951, 146 | 3 |
| ENSG00000103 | BFAR | chr16 | 14655065 | 14655210 | 14644273 | 14644609 | 14661891 | 14662065 | 66,45,120 | 2,4,5 | 130,200,144 | 1,2,0 | 295 | 150 | 0.000319231 | 0.004356932 | 0.944,0.851,0.92 | 0.985,0.981,1.0 | -0.082 | ENSG00000103 | [[[14632951, 146 | 3 |
| ENSG00000110 | PRDM4 | chr12 | 107751414 | 107752209 | 107746274 | 107746424 | 107756831 | 107756965 | 21,21,55 | 0,0,0 | 75,135,85 | 19,28,17 | 300 | 150 | 5.10E-12 | 4.17E-10 | 1.0,1.0,1.0 | 0.664,0.707,0.71 | 0.305 | ENSG00000110 | [[[107760957, 10 | 2 |
| ENSG00000110 | PRDM4 | chr12 | 107753923 | 107754109 | 107746274 | 107746424 | 107756831 | 107756965 | 43,37,63 | 0,0,0 | 71,224,147 | 19,28,17 | 300 | 150 | 2.88E-11 | 2.05E-09 | 1.0,1.0,1.0 | 0.651,0.8,0.812 | 0.246 | ENSG00000110 | [[[107760957, 10 | 2 |
| ENSG00000052 | TTC17 | chr11 | 43396708 | 43396818 | 43389651 | 43389821 | 43397346 | 43397491 | 32,15,57 | 0,0,0 | 28,44,39 | 1,6,2 | 260 | 150 | 0.000149319 | 0.002255264 | 1.0,1.0,1.0 | 0.942,0.809,0.91 | 0.11 | ENSG00000052 | [[[43358920, 433 | 3 |
| ENSG00000052 | TTC17 | chr11 | 43397973 | 43398113 | 43397346 | 43397491 | 43403997 | 43404144 | 36,25,56 | 0,0,0 | 41,94,74 | 4,4,4 | 290 | 150 | 8.11E-05 | 0.001324353 | 1.0,1.0,1.0 | 0.841,0.924,0.90 | 0.11 | ENSG00000052 | [[[43358920, 433 | 3 |
| ENSG00000052 | TTC17 | chr11 | 43401445 | 43401558 | 43397346 | 43397491 | 43403997 | 43404144 | 24,15,36 | 0,0,0 | 9,33,39 | 4,4,4 | 263 | 150 | 1.21E-06 | 3.24E-05 | 1.0,1.0,1.0 | 0.562,0.825,0.84 | 0.255 | ENSG00000052 | [[[43358920, 433 | 3 |
| ENSG00000052 | TTC17 | chr11 | 43414589 | 43414776 | 43405785 | 43405951 | 43436163 | 43436334 | 29,28,45 | 0,0,0 | 30,81,67 | 0,8,2 | 300 | 150 | 0.000193193 | 0.002829853 | 1.0,1.0,1.0 | 1.0,0.835,0.944 | 0.074 | ENSG00000052 | [[[43358920, 433 | 3 |
| ENSG00000151 | EXT2 | chr11 | 44124788 | 44124984 | 44114184 | 44114301 | 44130044 | 44130138 | 126,79,145 | 1,0,1 | 118,217,127 | 3,8,4 | 300 | 150 | 0.000631604 | 0.007844653 | 0.984,1.0,0.986 | 0.952,0.931,0.94 | 0.049 | ENSG00000151 | [[[44095678, 441 | 2 |
| ENSG00000138 | RAP1GDS1 | chr4 | 98433935 | 98434062 | 98417366 | 98417498 | 98436939 | 98437068 | 33,22,43 | 0,0,0 | 18,43,27 | 4,9,2 | 277 | 150 | 6.04E-08 | 2.24E-06 | 1.0,1.0,1.0 | 0.709,0.721,0.88 | 0.23 | ENSG00000138 | [[[98261384, 982 | 4 |
| ENSG00000138 | DBR1 | chr3 | 138171632 | 138171713 | 138170106 | 138170192 | 138174598 | 138174866 | 9,9,14 | 0,0,0 | 12,37,23 | 7,3,2 | 231 | 150 | 6.36E-06 | 0.000143782 | 1.0,1.0,1.0 | 0.527,0.889,0.88 | 0.234 | ENSG00000138 | [[[138174599, 13 | 2 |
| ENSG00000138 | DBR1 | chr3 | 138173501 | 138173626 | 138170106 | 138170192 | 138174598 | 138174859 | 8,8,12 | 0,0,0 | 10,18,8 | 7,3,2 | 275 | 150 | 3.83E-08 | 1.49E-06 | 1.0,1.0,1.0 | 0.438,0.766,0.68 | 0.37 | ENSG00000138 | [[[138174599, 13 | 2 |
| ENSG00000205 | SNX2 | chr5 | 122816914 | 122817028 | 122808276 | 122808355 | 122817279 | 122817340 | 64,63,78 | 0,0,0 | 71,143,95 | 2,11,3 | 264 | 150 | 1.32E-05 | 0.000271567 | 1.0,1.0,1.0 | 0.953,0.881,0.94 | 0.073 | ENSG00000205 | [[[122775080, 12 | 2 |
| ENSG00000175 | CKAP5 | chr11 | 46784487 | 46784673 | 46780427 | 46780485 | 46788680 | 46788773 | 105,59,182 | 0,0,0 | 41,109,72 | 0,8,3 | 300 | 150 | 7.70E-05 | 0.001267075 | 1.0,1.0,1.0 | 1.0,0.872,0.923 | 0.068 | ENSG00000175 | [[[46846220, 468 | 2 |
| ENSG00000125 | GPCPD1 | chr20 | 5561464 | 5561530 | 5544403 | 5547850 | 5565016 | 5565078 | 9,7,11 | 0,0,0 | 10,16,9 | 0,5,3 | 216 | 150 | 2.45E-05 | 0.000469445 | 1.0,1.0,1.0 | 1.0,0.69,0.676 | 0.211 | ENSG00000125 | [[[5610842, 5611 | 4 |
| ENSG00000149 | ARFGAP2 | chr11 | 47173758 | 47173840 | 47172025 | 47172333 | 47175014 | 47175098 | 17,31,54 | 1,0,0 | 20,49,34 | 4,6,3 | 232 | 150 | 9.95E-06 | 0.000212648 | 0.917,1.0,1.0 | 0.764,0.841,0.88 | 0.144 | ENSG00000149 | [[[47176782, 471 | 2 |
| ENSG00000149 | ARFGAP2 | chr11 | 47173758 | 47173882 | 47172280 | 47172333 | 47175014 | 47175098 | 4,4,8 | 1,0,0 | 3,5,2 | 4,6,3 | 274 | 150 | 8.30E-08 | 2.97E-06 | 0.686,1.0,1.0 | 0.291,0.313,0.26 | 0.605 | ENSG00000149 | [[[47176782, 471 | 2 |
| ENSG00000149 | ARFGAP2 | chr11 | 47175181 | 47175313 | 47172280 | 47172333 | 47175850 | 47175923 | 44,23,58 | 0,0,0 | 35,82,47 | 1,5,7 | 282 | 150 | 8.45E-06 | 0.000184032 | 1.0,1.0,1.0 | 0.949,0.897,0.78 | 0.124 | ENSG00000149 | [[[47176782, 471 | 4 |
| ENSG00000134 | DDB2 | chr11 | 47216335 | 47216472 | 47214941 | 47215263 | 47232813 | 47232959 | 15,38,56 | 0,0,0 | 9,27,19 | 5,9,2 | 287 | 150 | 1.52E-09 | 8.13E-08 | 1.0,1.0,1.0 | 0.485,0.611,0.83 | 0.357 | ENSG00000134 | [[[47214974, 472 | 2 |
| ENSG00000080 | CLDND1 | chr3 | 98518884 | 98518995 | 98515472 | 98516879 | 98521132 | 98521283 | 92,59,136 | 1,0,0 | 96,198,129 | 1,16,4 | 261 | 150 | 9.24E-06 | 0.000199665 | 0.981,1.0,1.0 | 0.982,0.877,0.94 | 0.058 | ENSG00000080 | [[[98522849, 985 | 2 |
| ENSG00000109 | MTCH2 | chr11 | 47635544 | 47635571 | 47634671 | 47634734 | 47642378 | 47642452 | 25,34,36 | 0,0,0 | 55,85,51 | 3,10,7 | 177 | 150 | 2.22E-06 | 5.60E-05 | 1.0,1.0,1.0 | 0.94,0.878,0.861 | 0.107 | ENSG00000109 | [[[47642379, 476 | 3 |
| ENSG00000109 | MTCH2 | chr11 | 47638698 | 47638805 | 47634671 | 47634734 | 47642378 | 47642452 | 20,16,20 | 0,0,0 | 20,30,13 | 3,10,7 | 257 | 150 | 4.39E-09 | 2.12E-07 | 1.0,1.0,1.0 | 0.796,0.636,0.52 | 0.349 | ENSG00000109 | [[[47642379, 476 | 3 |
| ENSG00000109 | MTCH2 | chr11 | 47638698 | 47638805 | 47635544 | 47635571 | 47642378 | 47642452 | 358,310,511 | 0,0,0 | 298,567,348 | 16,33,18 | 257 | 150 | 0 | 0 | 1.0,1.0,1.0 | 0.916,0.909,0.91 | 0.085 | ENSG00000109 | [[[47642379, 476 | 2 |
| ENSG00000153 | RBMS1 | chr2 | 160282256 | 160282304 | 160281313 | 160281364 | 160286977 | 160287084 | 7,7,12 | 0,0,0 | 5,4,8 | 4,7,0 | 198 | 150 | 2.54E-06 | 6.32E-05 | 1.0,1.0,1.0 | 0.486,0.302,1.0 | 0.404 | ENSG00000153 | [[[160493289, 16 | 2 |
| ENSG00000153 | RBMS1 | chr2 | 160284994 | 160285044 | 160281313 | 160281364 | 160286977 | 160287084 | 11,11,16 | 0,0,0 | 6,27,27 | 4,7,0 | 200 | 150 | 2.50E-05 | 0.000477928 | 1.0,1.0,1.0 | 0.529,0.743,1.0 | 0.243 | ENSG00000153 | [[[160493289, 16 | 2 |
| ENSG00000110 | SLC35F2 | chr11 | 107804717 | 107804770 | 107803000 | 107803155 | 107806716 | 107806876 | 57,47,78 | 0,6,2 | 32,93,70 | 9,16,5 | 203 | 150 | 0.000707909 | 0.008647206 | 1.0,0.853,0.966 | 0.724,0.811,0.91 | 0.124 | ENSG00000110 | [[[107858658, 10 | 2 |
| ENSG00000110 | SLC35F2 | chr11 | 107805358 | 107805515 | 107803000 | 107803155 | 107806716 | 107806876 | 115,117,187 | 0,6,2 | 73,166,99 | 9,16,5 | 300 | 150 | 1.23E-05 | 0.000255573 | 1.0,0.907,0.979 | 0.802,0.838,0.90 | 0.113 | ENSG00000110 | [[[107858658, 10 | 2 |
| ENSG00000143 | HNRNPLL | chr2 | 38569803 | 38569925 | 38561977 | 38564237 | 38573209 | 38573427 | 48,60,69 | 0,0,0 | 37,55,27 | 2,20,10 | 272 | 150 | 2.55E-11 | 1.83E-09 | 1.0,1.0,1.0 | 0.911,0.603,0.59 | 0.296 | ENSG00000143 | [[[38602438, 386 | 4 |
| ENSG00000213 | HEXA | chr15 | 72351132 | 72351234 | 72348047 | 72348134 | 72353067 | 72353178 | 123,96,122 | 0,0,0 | 134,184,84 | 10,3,4 | 252 | 150 | 1.55E-05 | 0.00031337 | 1.0,1.0,1.0 | 0.889,0.973,0.92 | 0.071 | ENSG00000213 | [[[72375720, 723 | 3 |
| ENSG00000213 | HEXA | chr15 | 72353067 | 72353178 | 72348047 | 72348134 | 72355558 | 72355624 | 4,10,14 | 0,0,0 | 31,24,17 | 1,4,6 | 261 | 150 | 6.19E-06 | 0.000140413 | 1.0,1.0,1.0 | 0.947,0.775,0.62 | 0.219 | ENSG00000213 | [[[72375720, 723 | 5 |
| ENSG00000213 | HEXA | chr15 | 72353690 | 72353737 | 72353067 | 72353178 | 72375719 | 72375844 | 42,33,49 | 0,0,0 | 36,79,38 | 1,14,4 | 197 | 150 | 3.66E-07 | 1.12E-05 | 1.0,1.0,1.0 | 0.965,0.811,0.87 | 0.115 | ENSG00000213 | [[[72375720, 723 | 3 |
| ENSG00000213 | HEXA | chr15 | 72356524 | 72356617 | 72353690 | 72353737 | 72375719 | 72375831 | 46,70,155 | 0,0,0 | 75,185,118 | 3,10,5 | 243 | 150 | 1.09E-05 | 0.00023084 | 1.0,1.0,1.0 | 0.939,0.919,0.93 | 0.069 | ENSG00000213 | [[[72375720, 723 | 2 |
| ENSG00000213 | TMX2 | chr11 | 57737835 | 57738026 | 57737607 | 57737668 | 57738663 | 57738770 | 5,9,2 | 0,0,0 | 10,12,14 | 2,3,5 | 300 | 150 | 2.83E-07 | 8.90E-06 | 1.0,1.0,1.0 | 0.714,0.667,0.58 | 0.345 | ENSG00000213 | [[[57712593, 577 | 2 |
| ENSG00000213 | TMX2 | chr11 | 57737835 | 57738026 | 57737607 | 57737668 | 57738973 | 57739039 | 5,9,2 | 0,0,0 | 10,11,14 | 6,5,7 | 300 | 150 | 1.00E-08 | 4.45E-07 | 1.0,1.0,1.0 | 0.455,0.524,0.5 | 0.507 | ENSG00000213 | [[[57712593, 577 | 3 |
| ENSG00000213 | TMX2 | chr11 | 57737912 | 57738026 | 57737607 | 57737668 | 57738973 | 57739039 | 82,70,126 | 0,0,0 | 76,185,131 | 6,5,7 | 264 | 150 | 8.40E-06 | 0.000183289 | 1.0,1.0,1.0 | 0.878,0.955,0.91 | 0.084 | ENSG00000213 | [[[57712593, 577 | 3 |
| ENSG00000213 | TMX2 | chr11 | 57738353 | 57738430 | 57737607 | 57737668 | 57738973 | 57739039 | 10,6,11 | 0,0,0 | 17,22,20 | 6,5,7 | 227 | 150 | 9.64E-08 | 3.41E-06 | 1.0,1.0,1.0 | 0.652,0.744,0.65 | 0.317 | ENSG00000213 | [[[57712593, 577 | 3 |
| ENSG00000213 | TMX2 | chr11 | 57738663 | 57738770 | 57737607 | 57737668 | 57738973 | 57739039 | 79,89,123 | 0,0,0 | 94,148,139 | 6,5,7 | 257 | 150 | 6.73E-06 | 0.000150664 | 1.0,1.0,1.0 | 0.901,0.945,0.92 | 0.078 | ENSG00000213 | [[[57712593, 577 | 3 |
| ENSG00000213 | TMX2 | chr11 | 57738663 | 57739039 | 57737607 | 57737668 | 57739130 | 57739181 | 45,38,64 | 0,0,0 | 46,102,67 | 2,9,2 | 300 | 150 | 1.73E-05 | 0.000345451 | 1.0,1.0,1.0 | 0.92,0.85,0.944 | 0.095 | ENSG00000213 | [[[57712593, 577 | 4 |

| | | | | | | | | | | | | | | | | | | | | | | |
|---|---|---|---|---|---|---|---|---|---|---|---|---|---|---|---|---|---|---|---|---|---|---|
| ENSG000002135 | TMX2 | chr11 | 57738973 | 57739039 | 57737607 | 57737668 | 57739130 | 57739212 | 45,38,61 | 0,0,0 | 49,102,69 | 2,9,2 | 216 | 150 | 0.00011831 | 0.00184657 | 1.0,1.0,1.0 | 0.944,0.887,0.96 | 0.07 | ENSG000002135 | [[[[57712593, 577 | 4 |
| ENSG000001366 | KCTD3 | chr1 | 215578999 | 215579137 | 215577669 | 215577728 | 215579908 | 215579999 | 45,35,74 | 0,0,0 | 37,90,62 | 3,9,9 | 288 | 150 | 1.62E-08 | 6.89E-07 | 1.0,1.0,1.0 | 0.865,0.839,0.78 | 0.171 | ENSG000001366 | [[[[215567304, 2 | 2 |
| ENSG000001830 | GAS6 | chr13 | 113832633 | 113832752 | 113828546 | 113828711 | 113834550 | 113834672 | 5,15,25 | 0,0,0 | 7,9,15 | 1,6,3 | 269 | 150 | 1.28E-07 | 4.35E-06 | 1.0,1.0,1.0 | 0.796,0.455,0.73 | 0.338 | ENSG000001830 | [[[[113863833, 11 | 2 |
| ENSG000001101 | PRPF19 | chr11 | 60899148 | 60899304 | 60898861 | 60898931 | 60901294 | 60901369 | 244,207,304 | 0,0,0 | 335,651,394 | 9,18,22 | 300 | 150 | 5.46E-10 | 3.18E-08 | 1.0,1.0,1.0 | 0.949,0.948,0.9 | 0.068 | ENSG000001101 | [[[[60906364, 609 | 3 |
| ENSG000001101 | PRPF19 | chr11 | 60900853 | 60900929 | 60898861 | 60898931 | 60901294 | 60901369 | 55,91,111 | 0,0,0 | 94,207,146 | 9,18,22 | 226 | 150 | 4.82E-12 | 3.97E-10 | 1.0,1.0,1.0 | 0.874,0.884,0.81 | 0.142 | ENSG000001101 | [[[[60906364, 609 | 3 |
| ENSG000001607 | NBEAL2 | chr3 | 47007240 | 47007350 | 47006332 | 47006449 | 47007524 | 47007697 | 12,15,28 | 0,0,0 | 5,14,17 | 2,9,3 | 260 | 150 | 3.48E-09 | 1.72E-07 | 1.0,1.0,1.0 | 0.591,0.473,0.76 | 0.39 | ENSG000001607 | [[[[46979666, 469 | 2 |
| ENSG000000890 | ESF1 | chr20 | 13771330 | 13771483 | 13766776 | 13766924 | 13772514 | 13772615 | 19,19,22 | 0,0,0 | 21,49,20 | 8,27,6 | 300 | 150 | 1.87E-12 | 1.65E-10 | 1.0,1.0,1.0 | 0.568,0.476,0.62 | 0.444 | ENSG000000890 | [[[[13784880, 137 | 2 |
| ENSG000001837 | TMEM50A | chr1 | 25340473 | 25340579 | 25337916 | 25338456 | 25351625 | 25351693 | 62,90,206 | 0,1,0 | 103,180,158 | 9,19,15 | 256 | 150 | 4.98E-12 | 4.08E-10 | 1.0,0.981,1.0 | 0.87,0.847,0.861 | 0.134 | ENSG000001837 | [[[[25338334, 253 | 2 |
| ENSG000000725 | MARK2 | chr11 | 63906087 | 63906114 | 63904785 | 63905043 | 63908876 | 63909579 | 3,4,4 | 0,0,1 | 1,1,5 | 0,7,10 | 177 | 150 | 6.43E-06 | 0.000145069 | 1.0,1.0,0.772 | 1.0,0.108,0.298 | 0.455 | ENSG000000725 | [[[[63839110, 638 | 2 |
| ENSG000000725 | MARK2 | chr11 | 63908259 | 63908304 | 63904785 | 63905043 | 63908876 | 63909237 | 5,28,51 | 0,0,1 | 20,89,40 | 0,7,10 | 195 | 150 | 3.85E-05 | 0.000697403 | 1.0,1.0,0.975 | 1.0,0.907,0.755 | 0.104 | ENSG000000725 | [[[[63839110, 638 | 2 |
| ENSG000001109 | SLC11A2 | chr12 | 50988381 | 50988435 | 50979534 | 50981803 | 50991598 | 50991672 | 14,16,24 | 0,0,0 | 14,34,20 | 3,4,2 | 204 | 150 | 7.15E-05 | 0.00119017 | 1.0,1.0,1.0 | 0.774,0.862,0.88 | 0.161 | ENSG000001109 | [[[[51026310, 510 | 2 |
| ENSG000001109 | SLC11A2 | chr12 | 50988381 | 50988435 | 50979534 | 50981803 | 50994543 | 50994630 | 14,16,24 | 0,0,0 | 14,33,20 | 11,4,4 | 204 | 150 | 8.64E-08 | 3.08E-06 | 1.0,1.0,1.0 | 0.483,0.858,0.78 | 0.291 | ENSG000001109 | [[[[51026310, 510 | 5 |
| ENSG000000873 | SF3B2 | chr11 | 66061890 | 66061998 | 66061685 | 66061775 | 66063399 | 66063542 | 269,225,377 | 0,0,0 | 266,495,298 | 9,29,11 | 258 | 150 | 3.13E-12 | 2.65E-10 | 1.0,1.0,1.0 | 0.945,0.908,0.94 | 0.069 | ENSG000000873 | [[[[66052364, 660 | 2 |
| ENSG000001871 | NAP1L1 | chr12 | 76053769 | 76053909 | 76053090 | 76053110 | 76055018 | 76055090 | 761,641,890 | 0,0,0 | 7,621,587,868 | 27,64,34 | 290 | 150 | 0 | 0 | 1.0,1.0,1.0 | 0.936,0.928,0.93 | 0.069 | ENSG000001871 | [[[[76084567, 760 | 2 |
| ENSG000001871 | NAP1L1 | chr12 | 76053832 | 76053909 | 76053090 | 76053110 | 76055018 | 76055090 | 395,331,492 | 0,0,0 | 460,963,520 | 27,64,34 | 227 | 150 | 0 | 0 | 1.0,1.0,1.0 | 0.918,0.909,0.91 | 0.088 | ENSG000001871 | [[[[76084567, 760 | 2 |
| ENSG000001871 | NAP1L1 | chr12 | 76055018 | 76055090 | 76053090 | 76053110 | 76056032 | 76056158 | 301,236,319 | 0,0,0 | 347,585,377 | 9,14,14 | 222 | 150 | 4.67E-07 | 1.40E-05 | 1.0,1.0,1.0 | 0.963,0.966,0.94 | 0.041 | ENSG000001871 | [[[[76084567, 760 | 3 |
| ENSG000001871 | NAP1L1 | chr12 | 76056032 | 76056158 | 76053090 | 76053110 | 76059797 | 76059878 | 6,6,5 | 0,0,0 | 27,33,24 | 8,36,15 | 276 | 150 | 9.58E-09 | 4.28E-07 | 1.0,1.0,1.0 | 0.647,0.333,0.46 | 0.518 | ENSG000001871 | [[[[76084567, 760 | 4 |
| ENSG000001871 | NAP1L1 | chr12 | 76056032 | 76056161 | 76053090 | 76053110 | 76059797 | 76059878 | 425,325,475 | 0,0,0 | 444,922,617 | 8,36,15 | 279 | 150 | 6.78E-11 | 4.56E-09 | 1.0,1.0,1.0 | 0.968,0.932,0.95 | 0.048 | ENSG000001871 | [[[[76084567, 760 | 4 |
| ENSG000001871 | NAP1L1 | chr12 | 76060137 | 76060279 | 76053090 | 76053110 | 76074202 | 76074239 | 36,50,75 | 0,0,0 | 43,92,56 | 10,9,13 | 292 | 150 | 2.00E-11 | 1.45E-09 | 1.0,1.0,1.0 | 0.688,0.84,0.689 | 0.261 | ENSG000001871 | [[[[76084567, 760 | 8 |
| ENSG000001871 | NAP1L1 | chr12 | 76068908 | 76068994 | 76053090 | 76053110 | 76074202 | 76074239 | 179,199,376 | 0,0,0 | 207,512,399 | 10,9,13 | 236 | 150 | 1.08E-06 | 2.94E-05 | 1.0,1.0,1.0 | 0.929,0.973,0.95 | 0.049 | ENSG000001871 | [[[[76084567, 760 | 8 |
| ENSG000001337 | IMPA1 | chr8 | 81673840 | 81673949 | 81660515 | 81660667 | 81679125 | 81679230 | 10,8,5 | 1,0,0 | 17,37,18 | 2,5,2 | 259 | 150 | 0.000699185 | 0.008555719 | 0.853,1.0,1.0 | 0.831,0.811,0.83 | 0.124 | ENSG000001337 | [[[[81686252, 816 | 3 |
| ENSG000000857 | CTTN | chr11 | 70420399 | 70420510 | 70419745 | 70419856 | 70422939 | 70422995 | 198,148,233 | 1,4,2 | 138,335,200 | 17,31,26 | 261 | 150 | 1.30E-13 | 1.34E-11 | 0.991,0.955,0.98 | 0.823,0.861,0.81 | 0.144 | ENSG000000857 | [[[[70398529, 703 | 2 |
| ENSG000000857 | CTTN | chr11 | 70421469 | 70421580 | 70419745 | 70419856 | 70422939 | 70422995 | 88,80,103 | 1,4,2 | 61,93,78 | 17,31,26 | 261 | 150 | 2.22E-16 | 3.17E-14 | 0.981,0.92,0.967 | 0.673,0.633,0.63 | 0.31 | ENSG000000857 | [[[[70398529, 703 | 2 |
| ENSG000001375 | RNF121 | chr11 | 71982806 | 71982915 | 71960749 | 71960891 | 71990596 | 71990663 | 0,1,1 | 4,6,8 | 22,27,13 | 2,13,2 | 259 | 150 | 1.80E-11 | 1.32E-09 | 0.0,0.088,0.068 | 0.864,0.546,0.79 | -0.681 | ENSG000001375 | [[[[71929046, 719 | 2 |
| ENSG000001654 | INPPL1 | chr11 | 72228726 | 72228847 | 72228347 | 72228498 | 72229464 | 72229558 | 2,25,56 | 0,0,0 | 16,31,39 | 0,5,3 | 271 | 150 | 0.000261174 | 0.003657225 | 1.0,1.0,1.0 | 1.0,0.774,0.878 | 0.116 | ENSG000001654 | [[[[72224767, 722 | 2 |
| ENSG000001654 | INPPL1 | chr11 | 72229089 | 72229230 | 72228726 | 72228847 | 72229923 | 72230019 | 7,40,73 | 0,0,0 | 41,83,59 | 2,8,1 | 291 | 150 | 6.69E-05 | 0.001124961 | 1.0,1.0,1.0 | 0.914,0.842,0.96 | 0.092 | ENSG000001654 | [[[[72224767, 722 | 3 |
| ENSG000001402 | TCF12 | chr15 | 57091788 | 57091891 | 56921025 | 56921098 | 57166401 | 57166428 | 32,25,40 | 0,0,0 | 23,60,53 | 3,7,10 | 253 | 150 | 3.67E-08 | 1.43E-06 | 1.0,1.0,1.0 | 0.82,0.836,0.759 | 0.195 | ENSG000001402 | [[[[56918644, 569 | 2 |
| ENSG000001402 | TCF12 | chr15 | 57166401 | 57166466 | 56921025 | 56921098 | 57192157 | 57192239 | 30,18,37 | 0,0,0 | 19,46,49 | 2,6,5 | 215 | 150 | 6.21E-06 | 0.000140601 | 1.0,1.0,1.0 | 0.869,0.842,0.87 | 0.139 | ENSG000001402 | [[[[56918644, 569 | 3 |
| ENSG000001402 | TCF12 | chr15 | 57231151 | 57231257 | 57197772 | 57197825 | 57232711 | 57232856 | 42,24,69 | 0,0,0 | 32,85,57 | 6,17,8 | 256 | 150 | 4.94E-11 | 3.39E-09 | 1.0,1.0,1.0 | 0.758,0.746,0.80 | 0.23 | ENSG000001402 | [[[[56918644, 569 | 2 |
| ENSG000001402 | TCF12 | chr15 | 57231151 | 57231257 | 57219036 | 57219590 | 57232711 | 57232856 | 38,41,64 | 0,0,0 | 33,64,43 | 3,9,3 | 256 | 150 | 3.46E-07 | 1.07E-05 | 1.0,1.0,1.0 | 0.866,0.806,0.89 | 0.145 | ENSG000001402 | [[[[56918644, 569 | 2 |
| ENSG000000419 | TNC | chr9 | 115048259 | 115048532 | 115046409 | 115046682 | 115073602 | 115073866 | 128,84,165 | 4,3,2 | 88,222,154 | 6,39,19 | 300 | 150 | 3.54E-08 | 1.39E-06 | 0.941,0.933,0.97 | 0.88,0.74,0.802 | 0.143 | ENSG000000419 | [[[[115117982, 11 | 6 |
| ENSG000000419 | TNC | chr9 | 115057152 | 115057425 | 115046409 | 115046682 | 115073602 | 115073866 | 254,224,337 | 4,3,2 | 137,378,279 | 6,39,19 | 300 | 150 | 1.92E-11 | 1.40E-09 | 0.969,0.974,0.98 | 0.919,0.829,0.88 | 0.101 | ENSG000000419 | [[[[115117982, 11 | 6 |
| ENSG000000419 | TNC | chr9 | 115059729 | 115060002 | 115042218 | 115042341 | 115073602 | 115073866 | 0,0,1 | 11,3,6 | 19,61,48 | 5,14,12 | 300 | 150 | 2.22E-16 | 3.17E-14 | 0.0,0.0,0.077 | 0.655,0.685,0.66 | -0.643 | ENSG000000419 | [[[[115117982, 11 | 7 |
| ENSG000000419 | TNC | chr9 | 115062916 | 115063189 | 115059729 | 115060002 | 115073602 | 115073866 | 80,135,302 | 0,0,1 | 109,226,173 | 19,61,48 | 300 | 150 | 0 | 0 | 1.0,1.0,0.993 | 0.741,0.649,0.64 | 0.32 | ENSG000000419 | [[[[115117982, 11 | 3 |
| ENSG000000419 | TNC | chr9 | 115063795 | 115064068 | 115059729 | 115060002 | 115073602 | 115073866 | 54,40,78 | 0,0,1 | 40,112,88 | 19,61,48 | 300 | 150 | 0 | 0 | 1.0,1.0,0.975 | 0.513,0.479,0.47 | 0.502 | ENSG000000419 | [[[[115117982, 11 | 3 |
| ENSG000000419 | TNC | chr9 | 115063795 | 115064068 | 115062916 | 115063189 | 115073602 | 115073866 | 279,221,437 | 0,0,0 | 144,394,267 | 19,42,23 | 300 | 150 | 0 | 0 | 1.0,1.0,1.0 | 0.791,0.824,0.85 | 0.177 | ENSG000000419 | [[[[115117982, 11 | 2 |
| ENSG000001755 | RAB6A | chr11 | 73716250 | 73716362 | 73679653 | 73679720 | 73718784 | 73718890 | 33,39,44 | 0,0,0 | 13,56,35 | 4,13,8 | 262 | 150 | 1.85E-11 | 1.35E-09 | 1.0,1.0,1.0 | 0.65,0.712,0.715 | 0.308 | ENSG000001755 | [[[[73760566, 737 | 3 |
| ENSG000001755 | RAB6A | chr11 | 73718612 | 73718718 | 73679653 | 73679720 | 73720845 | 73720899 | 20,21,34 | 2,2,0 | 27,50,34 | 9,15,12 | 256 | 150 | 5.47E-06 | 0.000125537 | 0.854,0.86,1.0 | 0.637,0.661,0.62 | 0.264 | ENSG000001755 | [[[[73760566, 737 | 3 |
| ENSG000001755 | RAB6A | chr11 | 73718784 | 73718890 | 73679653 | 73679720 | 73720845 | 73720899 | 23,25,38 | 2,2,0 | 46,77,43 | 9,15,12 | 256 | 150 | 4.67E-05 | 0.000823205 | 0.871,0.88,1.0 | 0.75,0.75,0.677 | 0.191 | ENSG000001755 | [[[[73760566, 737 | 3 |
| ENSG000001755 | MRPL48 | chr11 | 73808312 | 73808350 | 73787315 | 73787992 | 73825707 | 73825796 | 12,6,6 | 0,1,2 | 5,19,24 | 7,8,10 | 188 | 150 | 0.000707768 | 0.008646863 | 1.0,0.827,0.705 | 0.363,0.655,0.65 | 0.286 | ENSG000001755 | [[[[73787874, 737 | 2 |
| ENSG000001755 | MRPL48 | chr11 | 73825707 | 73825796 | 73787929 | 73787992 | 73844806 | 73844941 | 35,22,25 | 0,0,0 | 25,48,42 | 11,8,15 | 239 | 150 | 4.31E-11 | 2.99E-09 | 1.0,1.0,1.0 | 0.588,0.79,0.637 | 0.328 | ENSG000001755 | [[[[73787874, 737 | 3 |
| ENSG000001755 | PAAF1 | chr11 | 73887353 | 73887457 | 73878776 | 73878819 | 73909398 | 73909512 | 16,17,23 | 0,0,0 | 18,12,13 | 4,3,5 | 254 | 150 | 4.06E-08 | 1.57E-06 | 1.0,1.0,1.0 | 0.727,0.703,0.60 | 0.321 | ENSG000001755 | [[[[73876999, 738 | 4 |
| ENSG000001680 | C2CD3 | chr11 | 74103130 | 74103625 | 74098008 | 74098255 | 74106370 | 74106493 | 5,3,12 | 0,0,0 | 1,5,1 | 0,2,3 | 300 | 150 | 0.000230899 | 0.003290456 | 1.0,1.0,1.0 | 1.0,0.556,0.143 | 0.434 | ENSG000001680 | [[[[74170738, 741 | 2 |
| ENSG000001329 | USPL1 | chr13 | 30637743 | 30637857 | 30630834 | 30631474 | 30653147 | 30653305 | 10,3,6 | 0,0,0 | 1,4,7 | 6,0,2 | 264 | 150 | 5.19E-05 | 0.000901678 | 1.0,1.0,1.0 | 0.087,1.0,0.665 | 0.416 | ENSG000001329 | [[[[30617920, 306 | 3 |
| ENSG000001590 | ALG8 | chr11 | 78112649 | 78112770 | 78109441 | 78109581 | 78114265 | 78114376 | 49,40,55 | 0,0,0 | 21,39,20 | 5,3,2 | 271 | 150 | 2.03E-06 | 5.19E-05 | 1.0,1.0,1.0 | 0.699,0.878,0.84 | 0.192 | ENSG000001590 | [[[[78139494, 781 | 2 |
| ENSG000001590 | ALG8 | chr11 | 78124020 | 78124214 | 78121064 | 78121174 | 78139493 | 78139624 | 41,23,61 | 0,0,0 | 33,57,40 | 3,18,7 | 300 | 150 | 1.88E-10 | 1.19E-08 | 1.0,1.0,1.0 | 0.846,0.613,0.74 | 0.267 | ENSG000001590 | [[[[78139494, 781 | 2 |
| ENSG000001481 | C9orf3 | chr9 | 94923985 | 94924175 | 94800756 | 94801002 | 94955176 | 94955279 | 2,4,6 | 1,0,0 | 0,0,7 | 1,7,3 | 300 | 150 | 3.16E-07 | 9.84E-06 | 0.5,1.0,1.0 | 0.0,0.0,0.538 | 0.654 | ENSG000001481 | [[[[94726699, 947 | 2 |
| ENSG000001375 | PRCP | chr11 | 82838386 | 82838574 | 82823501 | 82825122 | 82849048 | 82849218 | 72,46,76 | 0,0,0 | 45,131,63 | 12,22,11 | 300 | 150 | 1.82E-14 | 2.11E-12 | 1.0,1.0,1.0 | 0.652,0.749,0.74 | 0.286 | ENSG000001375 | [[[[82900235, 829 | 2 |
| ENSG000001712 | TMEM126A | chr11 | 85650248 | 85650341 | 85647966 | 85648089 | 85655593 | 85655708 | 27,22,25 | 0,0,0 | 9,38,18 | 2,9,4 | 243 | 150 | 1.86E-08 | 7.84E-07 | 1.0,1.0,1.0 | 0.735,0.723,0.73 | 0.269 | ENSG000001712 | [[[[85647967, 856 | 2 |
| ENSG000001712 | TMEM126A | chr11 | 85654062 | 85654256 | 85647976 | 85648089 | 85655593 | 85655708 | 53,55,67 | 0,0,0 | 48,91,44 | 2,9,4 | 300 | 150 | 6.66E-07 | 1.92E-05 | 1.0,1.0,1.0 | 0.923,0.835,0.84 | 0.132 | ENSG000001712 | [[[[85647967, 856 | 2 |
| ENSG000000930 | CDC45 | chr22 | 19507765 | 19507864 | 19507385 | 19507517 | 19514748 | 19514784 | 45,38,49 | 0,0,0 | 32,58,31 | 3,7,8 | 249 | 150 | 1.98E-08 | 8.30E-07 | 1.0,1.0,1.0 | 0.865,0.833,0.7 | 0.201 | ENSG000000930 | [[[[19479920, 194 | 2 |
| ENSG000000930 | CDC45 | chr22 | 19514748 | 19514887 | 19508529 | 19508691 | 19516526 | 19516645 | 54,71,154 | 0,0,0 | 48,86,39 | 0,7,7 | 289 | 150 | 9.87E-07 | 2.73E-05 | 1.0,1.0,1.0 | 1.0,0.864,0.743 | 0.131 | ENSG000000930 | [[[[19479920, 194 | 2 |
| ENSG000000930 | CDC45 | chr22 | 19514964 | 19515048 | 19508529 | 19508691 | 19516526 | 19516645 | 14,41,53 | 0,0,0 | 36,64,38 | 0,7,7 | 234 | 150 | 6.39E-06 | 0.00014426 | 1.0,1.0,1.0 | 1.0,0.854,0.777 | 0.123 | ENSG000000930 | [[[[19479920, 194 | 2 |
| ENSG000001101 | CHORDC1 | chr11 | 90210535 | 90210594 | 90206201 | 90206272 | 90214017 | 90214175 | 18,4,14 | 0,0,0 | 21,54,36 | 2,7,1 | 209 | 150 | 0.000167414 | 0.002492982 | 1.0,1.0,1.0 | 0.883,0.847,0.96 | 0.102 | ENSG000001101 | [[[[90222891, 902 | 2 |
| ENSG000001660 | TAF1D | chr11 | 93739236 | 93739331 | 93734624 | 93734660 | 93741321 | 93741476 | 3,8,25 | 0,0,0 | 41,76,62 | 2,8,3 | 245 | 150 | 0.000120684 | 0.0018771 | 1.0,1.0,1.0 | 0.926,0.853,0.92 | 0.098 | ENSG000001660 | [[[[93741322, 937 | 5 |
| ENSG000000870 | MTMR2 | chr11 | 95858530 | 95858632 | 95857551 | 95857635 | 95865600 | 95865676 | 34,18,45 | 0,0,0 | 6,36,18 | 2,9,3 | 252 | 150 | 1.06E-08 | 4.71E-07 | 1.0,1.0,1.0 | 0.641,0.704,0.78 | 0.291 | ENSG000000870 | [[[[95923875, 959 | 3 |
| ENSG000000870 | MTMR2 | chr11 | 95861991 | 95862102 | 95858530 | 95858632 | 95865600 | 95865676 | 18,3,21 | 0,0,0 | 5,14,8 | 0,6,2 | 261 | 150 | 1.08E-05 | 0.000227566 | 1.0,1.0,1.0 | 1.0,0.573,0.697 | 0.243 | ENSG000000870 | [[[[95923875, 959 | 2 |
| ENSG000000870 | MTMR2 | chr11 | 95862271 | 95862366 | 95857551 | 95857635 | 95865600 | 95865676 | 18,11,32 | 0,0,0 | 10,23,12 | 2,9,3 | 245 | 150 | 7.27E-09 | 3.34E-07 | 1.0,1.0,1.0 | 0.754,0.61,0.71 | 0.309 | ENSG000000870 | [[[[95923875, 959 | 3 |
| ENSG000001010 | CEP192 | chr18 | 13037236 | 13037301 | 13030464 | 13030608 | 13040829 | 13040956 | 5,1,11 | 0,0,0 | 2,7,0 | 0,3,3 | 215 | 150 | 0.00024574 | 0.003468401 | 1.0,1.0,1.0 | 1.0,0.619,0.0 | 0.46 | ENSG000001010 | [[[[12991362, 129 | 2 |
| ENSG000001377 | RDX | chr11 | 110263959 | 110264234 | 110258105 | 110258189 | 110272535 | 110272619 | 63,28,110 | 1,0,1 | 28,120,75 | 4,6,3 | 300 | 150 | 0.000208398 | 0.003020062 | 0.969,1.0,0.982 | 0.778,0.909,0.92 | 0.113 | ENSG000001377 | [[[[110296467, 11 | 2 |
| ENSG000002579 | TEN1 | chr17 | 75986186 | 75986284 | 75979219 | 75979511 | 76000140 | 76000521 | 36,33,86 | 0,0,0 | 42,128,78 | 5,3,6 | 248 | 150 | 3.62E-05 | 0.000660918 | 1.0,1.0,1.0 | 0.836,0.963,0.88 | 0.105 | ENSG000002579 | [[[[75979240, 759 | 2 |
| ENSG000002579 | TEN1 | chr17 | 75991465 | 75991623 | 75979219 | 75979511 | 76000140 | 76000586 | 15,64,101 | 0,0,0 | 60,156,92 | 5,3,6 | 300 | 150 | 4.40E-05 | 0.000781507 | 1.0,1.0,1.0 | 0.857,0.963,0.88 | 0.098 | ENSG000002579 | [[[[75979240, 759 | 2 |
| ENSG000001507 | DLAT | chr11 | 112033403 | 112033530 | 112028791 | 112028945 | 112043465 | 112043533 | 42,25,54 | 0,0,0 | 22,36,24 | 6,7,4 | 277 | 150 | 9.71E-10 | 5.39E-08 | 1.0,1.0,1.0 | 0.665,0.736,0.76 | 0.278 | ENSG000001507 | [[[[112025408, 11 | 3 |
| ENSG000001507 | DLAT | chr11 | 112037272 | 112037460 | 112028791 | 112028945 | 112039243 | 112039397 | 65,49,87 | 0,0,0 | 35,59,65 | 4,1,5 | 300 | 150 | 5.92E-05 | 0.001011543 | 1.0,1.0,1.0 | 0.814,0.967,0.86 | 0.117 | ENSG000001507 | [[[[112025408, 11 | 2 |
| ENSG000001507 | DLAT | chr11 | 112039243 | 112039397 | 112028791 | 112028945 | 112043465 | 112043533 | 42,39,54 | 0,0,0 | 39,72,33 | 6,7,4 | 300 | 150 | 4.86E-08 | 1.84E-06 | 1.0,1.0,1.0 | 0.765,0.837,0.80 | 0.198 | ENSG000001507 | [[[[112025408, 11 | 3 |
| ENSG000001354 | HNRNPA1 | chr12 | 54282799 | 54282874 | 54282393 | 54282486 | 54283078 | 54283234 | 93,82,128 | 0,0,0 | 223,370,222 | 6,14,3 | 225 | 150 | 0.000225669 | 0.003233453 | 1.0,1.0,1.0 | 0.961,0.946,0.98 | 0.038 | ENSG000001354 | [[[[54280726, 542 | 2 |
| ENSG000001354 | HNRNPA1 | chr12 | 54283078 | 54283234 | 54281385 | 54281502 | 54283811 | 54283967 | 135,101,181 | 0,0,0 | 290,474,283 | 18,18,12 | 300 | 150 | 4.11E-09 | 2.00E-07 | 1.0,1.0,1.0 | 0.89,0.929,0.922 | 0.086 | ENSG000001354 | [[[[54280726, 542 | 6 |
| ENSG000001354 | HNRNPA1 | chr12 | 54283078 | 54283234 | 54282799 | 54282874 | 54286661 | 54286732 | 186,164,252 | 0,0,0 | 421,741,472 | 8,15,9 | 300 | 150 | 4.32E-05 | 0.000769957 | 1.0,1.0,1.0 | 0.963,0.961,0.96 | 0.038 | ENSG000001354 | [[[[54280726, 542 | 3 |
| ENSG000001354 | HNRNPA1 | chr12 | 54284257 | 54284317 | 54282799 | 54282874 | 54286661 | 54286732 | 16,25,18 | 0,0,0 | 55,131,41 | 8,15,9 | 210 | 150 | 3.07E-07 | 9.60E-06 | 1.0,1.0,1.0 | 0.831,0.862,0.76 | 0.181 | ENSG000001354 | [[[[54280726, 542 | 3 |
| ENSG000000044 | AK2 | chr1 | 33021366 | 33021461 | 33014364 | 33014594 | 33024441 | 33024567 | 223,238,369 | 0,2,2 | 252,547,317 | 13,26,13 | 245 | 150 | 7.33E-10 | 4.16E-08 | 1.0,0.986,0.991 | 0.922,0.928,0.93 | 0.063 | ENSG000000044 | [[[[33036736, 330 | 2 |
| ENSG000000044 | AK2 | chr1 | 33021592 | 33021703 | 33014482 | 33014594 | 33024441 | 33024567 | 285,253,443 | 0,2,2 | 274,551,358 | 13,26,13 | 261 | 150 | 1.39E-10 | 8.93E-09 | 1.0,0.986,0.992 | 0.924,0.924,0.94 | 0.063 | ENSG000000044 | [[[[33036736, 330 | 2 |
| ENSG000001378 | MAPKBP1 | chr15 | 41822938 | 41823006 | 41822592 | 41822677 | 41824483 | 41824569 | 6,3,21 | 0,1,1 | 2,15,10 | 4,2,2 | 218 | 150 | 0.00082636 | 0.009878723 | 1.0,0.674,0.935 | 0.256,0.838,0.77 | 0.247 | ENSG000001378 | [[[[41774484, 417 | 2 |
| ENSG000001103 | UBE4A | chr11 | 118374895 | 118375229 | 118372506 | 118372666 | 118376573 | 118376694 | 15,13,41 | 0,0,0 | 12,38,31 | 2,9,1 | 300 | 150 | 1.63E-06 | 4.26E-05 | 1.0,1.0,1.0 | 0.75,0.679,0.939 | 0.211 | ENSG000001103 | [[[[118359600, 11 | 3 |
| ENSG000000900 | PAPOLA | chr14 | 96525309 | 96525391 | 96520963 | 96521072 | 96527952 | 96528006 | 55,29,83 | 0,0,0 | 70,165,118 | 5,12,8 | 232 | 150 | 6.89E-07 | 1.97E-05 | 1.0,1.0,1.0 | 0.901,0.899,0.90 | 0.098 | ENSG000000900 | [[[[96502382, 965 | 2 |
| ENSG000000900 | PAPOLA | chr14 | 96527429 | 96527539 | 96521005 | 96521072 | 96527952 | 96528002 | 84,68,117 | 0,0,0 | 60,161,106 | 5,12,8 | 260 | 150 | 1.14E-08 | 5.00E-07 | 1.0,1.0,1.0 | 0.874,0.886,0.88 | 0.119 | ENSG000000900 | [[[[96502382, 965 | 2 |
| ENSG000000900 | PAPOLA | chr14 | 96532330 | 96532338 | 96531474 | 96531586 | 96534490 | 96534889 | 11,8,12 | 0,0,0 | 12,25,19 | 0,6,2 | 158 | 150 | 0.000572227 | 0.007211805 | 1.0,1.0,1.0 | 1.0,0.798,0.9 | 0.101 | ENSG000000900 | [[[[96502382, 965 | 2 |
| ENSG000000900 | PAPOLA | chr14 | 96536975 | 96537060 | 96534490 | 96534563 | 96542242 | 96542296 | 32,39,77 | 0,0,0 | 29,60,46 | 2,7,1 | 235 | 150 | 0.000122109 | 0.001896552 | 1.0,1.0,1.0 | 0.902,0.845,0.96 | 0.095 | ENSG000000900 | [[[[96502382, 965 | 2 |
| ENSG000000191 | PHLDB1 | chr11 | 118643799 | 118643940 | 118639161 | 118639251 | 118644071 | 118644174 | 26,43,102 | 0,0,0 | 54,93,72 | 0,10,5 | 291 | 150 | 5.89E-06 | 0.000134118 | 1.0,1.0,1.0 | 1.0,0.827,0.881 | 0.097 | ENSG000000191 | [[[[118607615, 11 | 2 |
| ENSG000001434 | MRPL9 | chr1 | 151760815 | 151760899 | 151759653 | 151760031 | 151762104 | 151762155 | 8,11,14 | 0,0,0 | 17,50,15 | 20,23,14 | 234 | 150 | 8.05E-12 | 6.26E-10 | 1.0,1.0,1.0 | 0.353,0.582,0.40 | 0.553 | ENSG000001434 | [[[[151763327, 15 | 2 |
| ENSG000001722 | DPAGT1 | chr11 | 119097854 | 119098043 | 119097139 | 119097297 | 119098402 | 119098435 | 35,31,37 | 0,0,0 | 23,40,16 | 4,4,1 | 300 | 150 | 1.15E-05 | 0.000241185 | 1.0,1.0,1.0 | 0.742,0.833,0.88 | 0.179 | ENSG000001722 | [[[[119101495, 11 | 2 |

| | | | | | | | | | | | | | | | | | | | | | | |
|---|---|---|---|---|---|---|---|---|---|---|---|---|---|---|---|---|---|---|---|---|---|---|
| ENSG000000627 | VMP1 | chr17 | 59731420 | 59731522 | 59707464 | 59707748 | 59737452 | 59737543 | 50,49,106 | 0,0,0 | 36,86,68 | 0,6,4 | 252 | 150 | 0.000484926 | 0.006268035 | 1.0,1.0,1.0 | 1.0,0.895,0.91 | 0.065 | ENSG000000627 | [[[[59707654, 597 | 2 |
| ENSG000000627 | VMP1 | chr17 | 59731420 | 59731522 | 59707464 | 59707748 | 59764970 | 59765068 | 50,49,100 | 0,0,8 | 37,86,65 | 6,13,13 | 252 | 150 | 3.08E-05 | 0.000573668 | 1.0,1.0,0.882 | 0.786,0.797,0.74 | 0.183 | ENSG000000627 | [[[[59707654, 597 | 4 |
| ENSG000000627 | VMP1 | chr17 | 59735337 | 59735473 | 59707464 | 59707748 | 59737452 | 59737474 | 63,57,76 | 0,0,0 | 42,89,51 | 0,6,4 | 286 | 150 | 0.000198267 | 0.002891401 | 1.0,1.0,1.0 | 1.0,0.886,0.87 | 0.081 | ENSG000000627 | [[[[59707654, 597 | 2 |
| ENSG000000627 | VMP1 | chr17 | 59737452 | 59737543 | 59735337 | 59735473 | 59764970 | 59765111 | 51,52,63 | 3,3,1 | 32,72,39 | 7,21,12 | 241 | 150 | 4.29E-08 | 1.64E-06 | 0.914,0.915,0.97 | 0.74,0.681,0.669 | 0.238 | ENSG000000627 | [[[[59707654, 597 | 2 |
| ENSG000000627 | VMP1 | chr17 | 59738836 | 59738947 | 59707464 | 59707748 | 59764970 | 59765068 | 76,59,102 | 0,0,8 | 52,94,69 | 6,13,13 | 261 | 150 | 1.95E-05 | 0.00038447 | 1.0,1.0,0.88 | 0.833,0.806,0.75 | 0.163 | ENSG000000627 | [[[[59707654, 597 | 4 |
| ENSG000000627 | VMP1 | chr17 | 59738836 | 59738947 | 59735337 | 59735473 | 59764970 | 59765068 | 79,63,102 | 3,3,1 | 53,97,72 | 7,21,12 | 261 | 150 | 7.28E-08 | 2.65E-06 | 0.938,0.923,0.98 | 0.813,0.726,0.77 | 0.177 | ENSG000000627 | [[[[59707654, 597 | 2 |
| ENSG000000627 | VMP1 | chr17 | 59738836 | 59738947 | 59735337 | 59735473 | 59765051 | 59765138 | 2,4,2 | 0,0,0 | 1,3,4 | 3,3,1 | 261 | 150 | 5.20E-08 | 1.96E-06 | 1.0,1.0,1.0 | 0.161,0.365,0.69 | 0.592 | ENSG000000627 | [[[[59707654, 597 | 2 |
| ENSG000000627 | VMP1 | chr17 | 59738836 | 59738947 | 59735337 | 59735473 | 59773753 | 59773881 | 2,4,3 | 0,0,0 | 2,3,4 | 3,3,2 | 261 | 150 | 2.92E-09 | 1.47E-07 | 1.0,1.0,1.0 | 0.277,0.365,0.53 | 0.608 | ENSG000000627 | [[[[59707654, 597 | 3 |
| ENSG000000627 | VMP1 | chr17 | 59764970 | 59765138 | 59738836 | 59738947 | 59808795 | 59808876 | 105,74,132 | 0,0,0 | 65,120,88 | 5,4,2 | 300 | 150 | 0.000211204 | 0.00305608 | 1.0,1.0,1.0 | 0.867,0.938,0.95 | 0.079 | ENSG000000627 | [[[[59707654, 597 | 2 |
| ENSG000000627 | VMP1 | chr17 | 59773753 | 59773885 | 59735337 | 59735473 | 59808795 | 59808876 | 130,131,181 | 2,0,0 | 82,190,111 | 8,6,3 | 282 | 150 | 1.61E-05 | 0.000324979 | 0.972,1.0,1.0 | 0.845,0.944,0.95 | 0.077 | ENSG000000627 | [[[[59707654, 597 | 4 |
| ENSG000001568 | FBRS | chr16 | 30661303 | 30661333 | 30660262 | 30660442 | 30662419 | 30662468 | 10,5,13 | 0,0,0 | 13,22,27 | 3,5,4 | 180 | 150 | 6.55E-06 | 0.000147312 | 1.0,1.0,1.0 | 0.783,0.786,0.84 | 0.194 | ENSG000001568 | [[[[30658431, 306 | 2 |
| ENSG000001829 | SRPRA | chr11 | 126264190 | 126264289 | 126262918 | 126264044 | 126264958 | 126265172 | 116,109,193 | 0,0,0 | 135,285,174 | 27,27,26 | 249 | 150 | 0 | 0 | 1.0,1.0,1.0 | 0.751,0.864,0.80 | 0.195 | ENSG000001829 | [[[[126268688, 12 | 2 |
| ENSG000001829 | SRPRA | chr11 | 126267548 | 126267712 | 126267174 | 126267335 | 126268687 | 126268982 | 131,127,192 | 0,0,0 | 141,339,232 | 2,6,10 | 300 | 150 | 0.000132819 | 0.002039131 | 1.0,1.0,1.0 | 0.972,0.966,0.92 | 0.047 | ENSG000001829 | [[[[126268688, 12 | 2 |
| ENSG000001403 | UBE2Q2 | chr15 | 75876186 | 75876271 | 75873427 | 75873568 | 75879097 | 75879188 | 25,19,37 | 0,0,0 | 22,31,36 | 3,0,5 | 235 | 150 | 0.000544636 | 0.006918999 | 1.0,1.0,1.0 | 0.824,1.0,0.821 | 0.118 | ENSG000001403 | [[[[75843447, 758 | 2 |
| ENSG000001403 | UBE2Q2 | chr15 | 75877960 | 75878021 | 75873427 | 75873568 | 75879097 | 75879188 | 21,18,11 | 0,0,0 | 18,26,9 | 3,0,5 | 211 | 150 | 0.00012296 | 0.001907037 | 1.0,1.0,1.0 | 0.81,1.0,0.561 | 0.21 | ENSG000001403 | [[[[75843447, 758 | 2 |
| ENSG000000068 | ADIPOR2 | chr12 | 1777853 | 1778025 | 1772713 | 1772961 | 1780888 | 1781076 | 63,38,124 | 0,0,0 | 71,157,109 | 5,7,3 | 300 | 150 | 3.08E-05 | 0.000572896 | 1.0,1.0,1.0 | 0.877,0.918,0.94 | 0.086 | ENSG000000068 | [[[[1691070, 1691 | 2 |
| ENSG000000068 | ADIPOR2 | chr12 | 1777853 | 1778025 | 1772713 | 1772961 | 1783879 | 1784073 | 63,38,124 | 0,0,0 | 70,162,108 | 1,7,7 | 300 | 150 | 2.39E-05 | 0.000460025 | 1.0,1.0,1.0 | 0.972,0.92,0.885 | 0.074 | ENSG000000068 | [[[[1691070, 1691 | 3 |
| ENSG000000068 | ADIPOR2 | chr12 | 1780450 | 1780637 | 1772841 | 1772961 | 1780888 | 1781076 | 59,50,80 | 0,0,0 | 57,144,91 | 5,7,3 | 300 | 150 | 1.34E-05 | 0.000275565 | 1.0,1.0,1.0 | 0.851,0.911,0.93 | 0.1 | ENSG000000068 | [[[[1691070, 1691 | 2 |
| ENSG000000068 | ADIPOR2 | chr12 | 1780888 | 1781076 | 1772841 | 1772961 | 1783879 | 1784073 | 44,38,113 | 0,0,0 | 56,116,92 | 1,7,7 | 300 | 150 | 6.98E-06 | 0.000155276 | 1.0,1.0,1.0 | 0.966,0.892,0.86 | 0.091 | ENSG000000068 | [[[[1691070, 1691 | 3 |
| ENSG000000671 | TNFRSF1A | chr12 | 6330852 | 6330926 | 6329867 | 6329899 | 6333068 | 6333147 | 114,88,151 | 0,0,0 | 133,236,117 | 7,13,9 | 224 | 150 | 2.69E-08 | 1.09E-06 | 1.0,1.0,1.0 | 0.927,0.924,0.89 | 0.084 | ENSG000000671 | [[[[6341776, 6342 | 3 |
| ENSG000001116 | TPI1 | chr12 | 6868863 | 6868987 | 6867118 | 6867681 | 6869257 | 6869295 | 106,722,323,862 | 0,0,0 | 158,531,032,248 | 11,46,13 | 274 | 150 | 0.000689462 | 0.008458483 | 1.0,1.0,1.0 | 0.987,0.974,0.99 | 0.016 | ENSG000001116 | [[[[6867531, 6867 | 2 |
| ENSG000001116 | TPI1 | chr12 | 6869257 | 6869390 | 6867118 | 6867681 | 6869687 | 6869770 | 186,317,062,699 | 0,0,0 | 162,128,891,791 | 14,31,22 | 283 | 150 | 7.00E-05 | 0.001167853 | 1.0,1.0,1.0 | 0.984,0.98,0.977 | 0.02 | ENSG000001116 | [[[[6867531, 6867 | 3 |
| ENSG000000237 | STRAP | chr12 | 15894046 | 15894143 | 15890596 | 15890669 | 15897881 | 15898018 | 127,77,130 | 0,0,0 | 71,157,99 | 6,24,18 | 247 | 150 | 8.88E-16 | 1.21E-13 | 1.0,1.0,1.0 | 0.878,0.799,0.77 | 0.184 | ENSG000000237 | [[[[15882387, 158 | 2 |
| ENSG000001117 | GOLT1B | chr12 | 21506884 | 21506976 | 21501814 | 21501948 | 21512294 | 21512376 | 49,46,78 | 0,1,0 | 94,235,140 | 4,8,8 | 242 | 150 | 0.000205184 | 0.002979726 | 1.0,0.966,1.0 | 0.936,0.948,0.91 | 0.055 | ENSG000001117 | [[[[21501789, 215 | 2 |
| ENSG000001117 | GOLT1B | chr12 | 21508382 | 21508561 | 21501792 | 21501948 | 21512294 | 21512376 | 54,40,90 | 0,1,0 | 87,89,68 | 4,8,8 | 300 | 150 | 8.09E-07 | 2.27E-05 | 1.0,0.952,1.0 | 0.916,0.848,0.81 | 0.126 | ENSG000001117 | [[[[21501789, 215 | 2 |
| ENSG000001117 | CMAS | chr12 | 22060831 | 22060926 | 22058566 | 22058700 | 22065120 | 22065663 | 70,51,82 | 0,0,0 | 76,165,117 | 3,10,9 | 245 | 150 | 1.19E-06 | 3.20E-05 | 1.0,1.0,1.0 | 0.939,0.91,0.888 | 0.088 | ENSG000001117 | [[[[22046218, 220 | 3 |
| ENSG000001117 | CMAS | chr12 | 22061280 | 22061452 | 22060641 | 22060926 | 22065120 | 22065663 | 113,92,92 | 0,0,0 | 66,130,83 | 11,21,13 | 300 | 150 | 2.22E-16 | 3.17E-14 | 1.0,1.0,1.0 | 0.75,0.756,0.761 | 0.244 | ENSG000001117 | [[[[22046218, 220 | 2 |
| ENSG000001117 | CMAS | chr12 | 22062280 | 22062434 | 22058566 | 22058700 | 22065120 | 22065663 | 87,66,132 | 0,0,0 | 100,185,90 | 3,10,9 | 300 | 150 | 1.64E-07 | 5.43E-06 | 1.0,1.0,1.0 | 0.943,0.902,0.83 | 0.107 | ENSG000001117 | [[[[22046218, 220 | 3 |
| ENSG000001117 | CMAS | chr12 | 22062280 | 22062434 | 22060831 | 22060926 | 22065120 | 22065663 | 89,66,133 | 0,0,0 | 108,224,108 | 11,21,13 | 300 | 150 | 1.60E-12 | 1.41E-10 | 1.0,1.0,1.0 | 0.831,0.842,0.80 | 0.174 | ENSG000001117 | [[[[22046218, 220 | 2 |
| ENSG000001561 | DRAM2 | chr1 | 111124741 | 111124881 | 111120515 | 111120693 | 111137522 | 111137586 | 28,33,31 | 0,0,0 | 50,89,70 | 5,13,3 | 290 | 150 | 7.17E-07 | 2.04E-05 | 1.0,1.0,1.0 | 0.838,0.78,0.923 | 0.153 | ENSG000001561 | [[[[111140038, 11 | 3 |
| ENSG000001561 | DRAM2 | chr1 | 111124741 | 111124881 | 111120515 | 111120693 | 111139500 | 111139666 | 28,33,31 | 0,0,0 | 49,88,69 | 2,6,2 | 290 | 150 | 0.000731262 | 0.00888971 | 1.0,1.0,1.0 | 0.927,0.884,0.94 | 0.081 | ENSG000001561 | [[[[111140038, 11 | 4 |
| ENSG000001561 | DRAM2 | chr1 | 111131423 | 111131568 | 111120515 | 111120693 | 111137522 | 111137586 | 54,22,84 | 0,0,0 | 70,106,78 | 5,13,3 | 295 | 150 | 2.37E-07 | 7.59E-06 | 1.0,1.0,1.0 | 0.877,0.806,0.93 | 0.129 | ENSG000001561 | [[[[111140038, 11 | 3 |
| ENSG000001561 | DRAM2 | chr1 | 111137522 | 111137586 | 111120515 | 111120693 | 111139500 | 111139666 | 11,11,22 | 0,0,0 | 23,50,28 | 2,6,2 | 214 | 150 | 0.00017848 | 0.002637668 | 1.0,1.0,1.0 | 0.89,0.854,0.908 | 0.116 | ENSG000001561 | [[[[111140038, 11 | 4 |
| ENSG000000609 | BCAT1 | chr12 | 24878529 | 24878649 | 24849785 | 24849949 | 24894274 | 24894475 | 95,76,163 | 0,0,0 | 60,154,111 | 11,26,18 | 270 | 150 | 0 | 0 | 1.0,1.0,1.0 | 0.752,0.767,0.77 | 0.236 | ENSG000000609 | [[[[24948927, 249 | 2 |
| ENSG000000874 | DNM1L | chr12 | 32720663 | 32720795 | 32718642 | 32718763 | 32731013 | 32731061 | 22,29,62 | 0,0,0 | 19,42,52 | 9,17,0 | 282 | 150 | 1.08E-07 | 3.75E-06 | 1.0,1.0,1.0 | 0.529,0.568,1.0 | 0.301 | ENSG000000874 | [[[[32679301, 326 | 2 |
| ENSG000001764 | CLK2 | chr1 | 155265859 | 155265954 | 155264644 | 155264774 | 155268009 | 155268126 | 25,24,35 | 3,0,1 | 22,76,26 | 10,6,7 | 245 | 150 | 0.000581066 | 0.007302662 | 0.836,1.0,0.955 | 0.574,0.886,0.69 | 0.212 | ENSG000001764 | [[[[155273201, 15 | 2 |
| ENSG000001764 | CLK2 | chr1 | 155266728 | 155266895 | 155264644 | 155264774 | 155268009 | 155268126 | 52,48,95 | 3,0,1 | 56,114,82 | 10,6,7 | 300 | 150 | 0.000138965 | 0.002117341 | 0.897,1.0,0.979 | 0.737,0.905,0.85 | 0.127 | ENSG000001764 | [[[[155273201, 15 | 2 |
| ENSG000001342 | PPHLN1 | chr12 | 42384939 | 42384996 | 42355160 | 42355222 | 42387455 | 42387535 | 34,24,28 | 5,0,4 | 21,44,31 | 10,15,11 | 207 | 150 | 0.000351507 | 0.004728831 | 0.831,1.0,0.835 | 0.603,0.68,0.671 | 0.237 | ENSG000001342 | [[[[42326167, 423 | 2 |
| ENSG000001890 | ARID2 | chr12 | 45846855 | 45846937 | 45839328 | 45839496 | 45850035 | 45852896 | 7,11,10 | 0,0,0 | 1,8,4 | 3,3,6 | 232 | 150 | 1.47E-10 | 9.37E-09 | 1.0,1.0,1.0 | 0.177,0.633,0.30 | 0.63 | ENSG000001890 | [[[[45729706, 457 | 3 |
| ENSG000001341 | MEIS2 | chr15 | 37096288 | 37096407 | 37095563 | 37095614 | 37100891 | 37101279 | 13,23,18 | 1,0,0 | 13,45,48 | 4,4,2 | 269 | 150 | 0.000336839 | 0.004556442 | 0.879,1.0,1.0 | 0.644,0.863,0.93 | 0.147 | ENSG000001341 | [[[[37099455, 371 | 3 |
| ENSG000000661 | SMARCD1 | chr12 | 50087362 | 50087485 | 50086755 | 50086878 | 50089883 | 50089985 | 48,29,65 | 1,0,2 | 79,139,87 | 10,13,13 | 273 | 150 | 5.05E-06 | 0.000116869 | 0.963,1.0,0.947 | 0.813,0.855,0.78 | 0.152 | ENSG000000661 | [[[[50085342, 500 | 2 |
| ENSG000000661 | SMARCD1 | chr12 | 50088520 | 50088637 | 50086755 | 50086878 | 50089883 | 50089985 | 55,53,77 | 1,0,2 | 85,172,111 | 10,13,13 | 267 | 150 | 1.88E-06 | 4.85E-05 | 0.969,1.0,0.956 | 0.827,0.881,0.82 | 0.13 | ENSG000000661 | [[[[50085342, 500 | 2 |
| ENSG000000504 | LETMD1 | chr12 | 51052091 | 51052207 | 51048342 | 51049185 | 51055986 | 51056021 | 16,6,9 | 0,0,0 | 12,37,35 | 11,22,23 | 266 | 150 | 1.22E-11 | 9.28E-10 | 1.0,1.0,1.0 | 0.381,0.487,0.46 | 0.557 | ENSG000000504 | [[[[51048327, 510 | 2 |
| ENSG000000504 | LETMD1 | chr12 | 51053777 | 51053860 | 51052087 | 51052207 | 51056143 | 51056245 | 2,5,9 | 0,0,0 | 2,12,11 | 2,3,2 | 233 | 150 | 2.56E-05 | 0.000486969 | 1.0,1.0,1.0 | 0.392,0.72,0.78 | 0.369 | ENSG000000504 | [[[[51048327, 510 | 2 |
| ENSG000000504 | LETMD1 | chr12 | 51055834 | 51056021 | 51052091 | 51052207 | 51056143 | 51056221 | 26,19,29 | 0,0,0 | 20,41,31 | 2,3,2 | 300 | 150 | 0.000789749 | 0.009488692 | 1.0,1.0,1.0 | 0.833,0.872,0.88 | 0.136 | ENSG000000504 | [[[[51048327, 510 | 2 |
| ENSG000000504 | LETMD1 | chr12 | 51055986 | 51056021 | 51052091 | 51052207 | 51056143 | 51056245 | 12,5,7 | 0,0,0 | 7,12,18 | 2,3,2 | 185 | 150 | 0.000314458 | 0.004301809 | 1.0,1.0,1.0 | 0.739,0.764,0.87 | 0.206 | ENSG000000504 | [[[[51048327, 510 | 2 |
| ENSG000000504 | LETMD1 | chr12 | 51056143 | 51056245 | 51049033 | 51049185 | 51056349 | 51056428 | 19,19,15 | 4,3,13 | 26,64,40 | 65,92,38 | 252 | 150 | 0.000651868 | 0.008051687 | 0.739,0.79,0.407 | 0.192,0.293,0.38 | 0.355 | ENSG000000504 | [[[[51048327, 510 | 4 |
| ENSG000001004 | FKBP3 | chr14 | 45120886 | 45120954 | 45118027 | 45118125 | 45134348 | 45134529 | 124,125,155 | 0,0,0 | 112,219,141 | 11,7,7 | 218 | 150 | 1.29E-07 | 4.39E-06 | 1.0,1.0,1.0 | 0.875,0.956,0.93 | 0.079 | ENSG000001004 | [[[[45134349, 451 | 4 |
| ENSG000001004 | FKBP3 | chr14 | 45129793 | 45129901 | 45121484 | 45121620 | 45134348 | 45134483 | 170,156,254 | 0,0,0 | 124,237,135 | 65,123,79 | 258 | 150 | 0 | 0 | 1.0,1.0,1.0 | 0.526,0.528,0.49 | 0.483 | ENSG000001004 | [[[[45134349, 451 | 2 |
| ENSG000000992 | RAB18 | chr10 | 27533734 | 27533853 | 27509874 | 27509930 | 27533927 | 27533994 | 32,26,64 | 0,0,0 | 32,41,39 | 3,3,2 | 269 | 150 | 0.000460096 | 0.005991863 | 1.0,1.0,1.0 | 0.856,0.884,0.91 | 0.115 | ENSG000000992 | [[[[27504304, 275 | 3 |
| ENSG000000727 | ACADVL | chr17 | 7221537 | 7221682 | 7220923 | 7221058 | 7222176 | 7222302 | 143,213,405 | 0,0,0 | 169,381,258 | 30,51,29 | 295 | 150 | 0 | 0 | 1.0,1.0,1.0 | 0.741,0.792,0.81 | 0.216 | ENSG000000727 | [[[[7219938, 7220 | 2 |
| ENSG000001735 | GMPPB | chr3 | 49722230 | 49722358 | 49719923 | 49722147 | 49722971 | 49723114 | 5,24,38 | 0,0,0 | 12,25,22 | 0,6,6 | 278 | 150 | 8.51E-07 | 2.38E-05 | 1.0,1.0,1.0 | 1.0,0.692,0.664 | 0.215 | ENSG000001735 | [[[[49723598, 497 | 3 |
| ENSG000001735 | GMPPB | chr3 | 49722595 | 49722754 | 49719923 | 49722147 | 49722971 | 49723114 | 26,54,83 | 0,0,0 | 49,98,65 | 0,6,6 | 300 | 150 | 4.30E-05 | 0.000766791 | 1.0,1.0,1.0 | 1.0,0.891,0.844 | 0.088 | ENSG000001735 | [[[[49723598, 497 | 3 |
| ENSG000001414 | OSBPL1A | chr18 | 24196124 | 24196200 | 24179737 | 24179835 | 24225041 | 24225198 | 5,3,6 | 0,0,0 | 5,14,12 | 2,0,6 | 226 | 150 | 3.68E-05 | 0.000668911 | 1.0,1.0,1.0 | 0.624,1.0,0.57 | 0.269 | ENSG000001414 | [[[[24397655, 243 | 2 |
| ENSG000000624 | CS | chr12 | 56282419 | 56282608 | 56275995 | 56276195 | 56285915 | 56286023 | 207,160,289 | 0,0,0 | 136,276,202 | 9,16,13 | 300 | 150 | 9.61E-13 | 8.72E-11 | 1.0,1.0,1.0 | 0.883,0.896,0.88 | 0.112 | ENSG000000624 | [[[[56300160, 563 | 3 |
| ENSG000000624 | CS | chr12 | 56282859 | 56282991 | 56276147 | 56276195 | 56285915 | 56286023 | 1,2,3 | 0,0,0 | 66,176,123 | 9,16,13 | 282 | 150 | 4.90E-05 | 0.000856317 | 1.0,1.0,1.0 | 0.796,0.854,0.83 | 0.172 | ENSG000000624 | [[[[56300160, 563 | 3 |
| ENSG000000624 | CS | chr12 | 56283791 | 56283857 | 56275995 | 56276195 | 56285915 | 56285961 | 77,68,112 | 0,0,0 | 78,222,133 | 9,16,13 | 216 | 150 | 6.38E-10 | 3.65E-08 | 1.0,1.0,1.0 | 0.858,0.906,0.87 | 0.12 | ENSG000000624 | [[[[56300160, 563 | 3 |
| ENSG000001980 | PRIM1 | chr12 | 56741434 | 56741576 | 56739293 | 56739363 | 56744064 | 56744123 | 48,36,47 | 0,0,0 | 22,68,26 | 2,9,0 | 292 | 150 | 1.18E-05 | 0.000247111 | 1.0,1.0,1.0 | 0.85,0.795,1.0 | 0.118 | ENSG000001980 | [[[[56752196, 567 | 3 |
| ENSG000001669 | MBD6 | chr12 | 57524952 | 57525115 | 57524719 | 57524822 | 57527847 | 57528017 | 6,11,16 | 0,0,0 | 29,73,44 | 3,8,1 | 300 | 150 | 5.20E-05 | 0.000903446 | 1.0,1.0,1.0 | 0.829,0.82,0.957 | 0.131 | ENSG000001669 | [[[[57522824, 575 | 4 |
| ENSG000001669 | MBD6 | chr12 | 57527506 | 57527660 | 57524719 | 57524822 | 57527847 | 57528017 | 9,10,25 | 0,0,0 | 36,77,43 | 3,8,1 | 300 | 150 | 5.67E-05 | 0.000973769 | 1.0,1.0,1.0 | 0.857,0.828,0.95 | 0.12 | ENSG000001669 | [[[[57522824, 575 | 4 |
| ENSG000001415 | SAT2 | chr17 | 7627142 | 7627226 | 7626706 | 7626793 | 7627362 | 7627414 | 99,114,151 | 0,0,0 | 229,382,185 | 48,56,32 | 234 | 150 | 0 | 0 | 1.0,1.0,1.0 | 0.754,0.814,0.78 | 0.215 | ENSG000001415 | [[[[7627570, 7627 | 2 |
| ENSG000001415 | SAT2 | chr17 | 7627142 | 7627226 | 7626706 | 7626793 | 7627552 | 7627802 | 13,12,34 | 0,0,0 | 32,53,20 | 5,9,3 | 234 | 150 | 4.13E-07 | 1.24E-05 | 1.0,1.0,1.0 | 0.804,0.791,0.81 | 0.198 | ENSG000001415 | [[[[7627570, 7627 | 3 |
| ENSG000001415 | SAT2 | chr17 | 7627142 | 7627226 | 7626706 | 7626793 | 7627569 | 7627805 | 13,12,33 | 0,0,0 | 30,53,18 | 1,8,3 | 234 | 150 | 1.29E-05 | 0.000266582 | 1.0,1.0,1.0 | 0.951,0.809,0.79 | 0.149 | ENSG000001415 | [[[[7627570, 7627 | 3 |
| ENSG000000058 | ZFX | chrX | 24152719 | 24152830 | 24149728 | 24149794 | 24172714 | 24172800 | 3,2,4 | 0,0,0 | 1,13,7 | 3,4,2 | 261 | 150 | 1.56E-07 | 5.21E-06 | 1.0,1.0,1.0 | 0.161,0.651,0.66 | 0.507 | ENSG000000058 | [[[[24149729, 241 | 2 |
| ENSG000000058 | ZFX | chrX | 24161714 | 24161842 | 24149744 | 24149794 | 24172714 | 24172800 | 3,1,2 | 0,0,0 | 0,8,12 | 3,4,2 | 278 | 150 | 1.20E-07 | 4.11E-06 | 1.0,1.0,1.0 | 0.0,0.519,0.764 | 0.572 | ENSG000000058 | [[[[24149729, 241 | 2 |
| ENSG000001233 | SPATS2 | chr12 | 49494740 | 49495002 | 49490681 | 49490731 | 49512861 | 49512905 | 25,20,30 | 0,0,0 | 18,58,44 | 1,5,5 | 300 | 150 | 1.50E-05 | 0.000304429 | 1.0,1.0,1.0 | 0.9,0.853,0.815 | 0.144 | ENSG000001233 | [[[[49367462, 493 | 3 |
| ENSG000001845 | XPOT | chr12 | 64420069 | 64420254 | 64418875 | 64419094 | 64423004 | 64423043 | 88,72,166 | 0,0,0 | 102,255,176 | 2,19,10 | 300 | 150 | 1.58E-08 | 6.73E-07 | 1.0,1.0,1.0 | 0.962,0.87,0.898 | 0.09 | ENSG000001845 | [[[[64404392, 644 | 3 |
| ENSG000001845 | XPOT | chr12 | 64420352 | 64420521 | 64420069 | 64420254 | 64423004 | 64423043 | 85,62,135 | 0,0,0 | 86,183,125 | 4,10,11 | 300 | 150 | 3.28E-08 | 1.30E-06 | 1.0,1.0,1.0 | 0.915,0.901,0.85 | 0.111 | ENSG000001845 | [[[[64404392, 644 | 2 |
| ENSG000001884 | CHM | chrX | 85978766 | 85978891 | 85963664 | 85964052 | 86027490 | 86027557 | 34,22,69 | 0,1,0 | 21,59,45 | 3,5,5 | 275 | 150 | 9.34E-06 | 0.000201527 | 1.0,0.923,1.0 | 0.792,0.866,0.83 | 0.145 | ENSG000001884 | [[[[86047484, 860 | 2 |
| ENSG000001167 | WLS | chr1 | 68137779 | 68137933 | 68098472 | 68098753 | 68144568 | 68144652 | 181,183,285 | 0,0,0 | 195,367,220 | 13,17,18 | 300 | 150 | 1.39E-13 | 1.43E-11 | 1.0,1.0,1.0 | 0.882,0.915,0.85 | 0.115 | ENSG000001167 | [[[[68232194, 682 | 2 |
| ENSG000001115 | CAND1 | chr12 | 67307396 | 67307492 | 67305103 | 67306597 | 67311692 | 67311800 | 68,49,95 | 0,0,0 | 49,79,39 | 2,7,3 | 246 | 150 | 1.43E-05 | 0.000292629 | 1.0,1.0,1.0 | 0.937,0.873,0.88 | 0.101 | ENSG000001115 | [[[[67269358, 672 | 3 |
| ENSG000001527 | HNRNPDL | chr4 | 82426036 | 82426129 | 82422563 | 82424883 | 82427189 | 82427304 | 257,191,266 | 0,0,0 | 125,279,155 | 4,14,4 | 243 | 150 | 3.89E-07 | 1.18E-05 | 1.0,1.0,1.0 | 0.951,0.925,0.96 | 0.055 | ENSG000001527 | [[[[82429248, 824 | 2 |
| ENSG000001527 | HNRNPDL | chr4 | 82426036 | 82426129 | 82422563 | 82424883 | 82429247 | 82429412 | 245,182,260 | 0,0,0 | 87,197,106 | 1,11,3 | 243 | 150 | 2.74E-05 | 0.000515445 | 1.0,1.0,1.0 | 0.982,0.917,0.95 | 0.048 | ENSG000001527 | [[[[82429248, 824 | 6 |
| ENSG000001527 | HNRNPDL | chr4 | 82426036 | 82426129 | 82425562 | 82425667 | 82427189 | 82427304 | 44,61,53 | 0,0,0 | 4,881,069,633 | 39,66,45 | 243 | 150 | 4.19E-08 | 1.61E-06 | 1.0,1.0,1.0 | 0.885,0.909,0.89 | 0.103 | ENSG000001527 | [[[[82429248, 824 | 2 |
| ENSG000001527 | HNRNPDL | chr4 | 82426462 | 82426633 | 82425562 | 82425667 | 82427189 | 82427304 | 402,389,571 | 0,0,0 | 463,921,540 | 39,66,45 | 300 | 150 | 0 | 0 | 1.0,1.0,1.0 | 0.856,0.875,0.85 | 0.137 | ENSG000001527 | [[[[82429248, 824 | 2 |
| ENSG000001527 | HNRNPDL | chr4 | 82427189 | 82427304 | 82426036 | 82426129 | 82428017 | 82428179 | 10,12,6 | 0,0,0 | 64,129,77 | 6,15,0 | 265 | 150 | 8.63E-05 | 0.001401181 | 1.0,1.0,1.0 | 0.858,0.83,1.0 | 0.104 | ENSG000001527 | [[[[82429248, 824 | 3 |
| ENSG000001527 | HNRNPDL | chr4 | 82427189 | 82427304 | 82426036 | 82426129 | 82429247 | 82429397 | 10,7,6 | 0,0,0 | 71,135,84 | 41,76,51 | 265 | 150 | 4.06E-10 | 2.41E-08 | 1.0,1.0,1.0 | 0.495,0.501,0.48 | 0.507 | ENSG000001527 | [[[[82429248, 824 | 5 |
| ENSG000001527 | HNRNPDL | chr4 | 82427432 | 82427564 | 82427189 | 82427304 | 82429247 | 82429425 | 466,406,593 | 0,0,0 | 6,511,314,728 | 11,15,9 | 282 | 150 | 0.000379712 | 0.005059156 | 1.0,1.0,1.0 | 0.969,0.979,0.97 | 0.025 | ENSG000001527 | [[[[82429248, 824 | 3 |
| ENSG000001527 | HNRNPDL | chr4 | 82428017 | 82428179 | 82427432 | 82427564 | 82429247 | 82429397 | 458,414,583 | 0,0,0 | 5,571,149,689 | 14,39,18 | 300 | 150 | 8.27E-13 | 7.62E-11 | 1.0,1.0,1.0 | 0.952,0.936,0.95 | 0.054 | ENSG000001527 | [[[[82429248, 824 | 2 |
| ENSG000001527 | HNRNPDL | chr4 | 82428277 | 82428446 | 82424624 | 82426129 | 82429247 | 82429397 | 383,405,736 | 0,0,0 | 5,781,159,727 | 41,76,51 | 300 | 150 | 0 | 0 | 1.0,1.0,1.0 | 0.876,0.884,0.87 | 0.121 | ENSG000001527 | [[[[82429248, 824 | 5 |

| | | | | | | | | | | | | | | | | | | | | | | |
|---|---|---|---|---|---|---|---|---|---|---|---|---|---|---|---|---|---|---|---|---|---|---|
| ENSG000001527 | HNRNPDL | chr4 | 82428277 | 82428446 | 82427189 | 82427304 | 82429247 | 82429397 | 383,405,736 | 0,0,0 | 5,791,159,726 | 11,15,9 | 300 | 150 | 5.07E-05 | 0.0008832 | 1.0,1.0,1.0 | 0.963,0.975,0.97 | 0.029 | ENSG000001527 | [[[[82429248, 824 | 3 |
| ENSG000001527 | HNRNPDL | chr4 | 82428277 | 82428446 | 82427432 | 82427564 | 82429247 | 82429397 | 383,405,736 | 0,0,0 | 5,781,160,723 | 14,39,18 | 300 | 150 | 8.89E-13 | 8.12E-11 | 1.0,1.0,1.0 | 0.954,0.937,0.95 | 0.052 | ENSG000001527 | [[[[82429248, 824 | 2 |
| ENSG000001356 | MDM2 | chr12 | 68813553 | 68813628 | 68809207 | 68809292 | 68820324 | 68820374 | 11,5,19 | 0,0,0 | 27,60,47 | 2,5,4 | 225 | 150 | 0.0002354 | 0.003344944 | 1.0,1.0,1.0 | 0.9,0.889,0.887 | 0.108 | ENSG000001356 | [[[[68808177, 688 | 2 |
| ENSG000001356 | MDM2 | chr12 | 68816811 | 68816945 | 68809211 | 68809292 | 68820324 | 68820374 | 14,4,18 | 0,0,0 | 24,70,70 | 2,5,4 | 284 | 150 | 0.000229599 | 0.0032756 | 1.0,1.0,1.0 | 0.864,0.881,0.90 | 0.118 | ENSG000001356 | [[[[68808177, 688 | 2 |
| ENSG000001658 | ZFYVE1 | chr14 | 72978860 | 72978969 | 72977926 | 72978002 | 72981788 | 72981895 | 9,12,18 | 0,0,0 | 13,14,21 | 4,2,1 | 259 | 150 | 0.000198373 | 0.002892391 | 1.0,1.0,1.0 | 0.653,0.802,0.92 | 0.207 | ENSG000001658 | [[[[73026926, 730 | 2 |
| ENSG000001662 | CCT2 | chr12 | 69597637 | 69597766 | 69597155 | 69597275 | 69599862 | 69600004 | 528,498,565 | 0,0,0 | 358,812,495 | 18,33,9 | 279 | 150 | 1.27E-14 | 1.50E-12 | 1.0,1.0,1.0 | 0.914,0.93,0.967 | 0.063 | ENSG000001662 | [[[[69585459, 695 | 3 |
| ENSG000001122 | CRYBG1 | chr6 | 106539402 | 106539529 | 106530175 | 106530315 | 106551851 | 106551978 | 86,61,146 | 0,0,0 | 28,107,70 | 1,3,8 | 277 | 150 | 1.62E-05 | 0.000326285 | 1.0,1.0,1.0 | 0.938,0.951,0.82 | 0.095 | ENSG000001122 | [[[[106360717, 10 | 5 |
| ENSG000001122 | CRYBG1 | chr6 | 106544570 | 106544697 | 106541585 | 106541621 | 106544787 | 106544933 | 74,59,134 | 0,0,0 | 21,51,35 | 6,8,7 | 277 | 150 | 5.77E-13 | 5.43E-11 | 1.0,1.0,1.0 | 0.655,0.775,0.73 | 0.28 | ENSG000001122 | [[[[106360717, 10 | 2 |
| ENSG000001122 | CRYBG1 | chr6 | 106544787 | 106544933 | 106541585 | 106541621 | 106551851 | 106551978 | 110,75,145 | 0,0,0 | 49,98,82 | 3,11,7 | 296 | 150 | 6.61E-10 | 3.78E-08 | 1.0,1.0,1.0 | 0.892,0.819,0.85 | 0.144 | ENSG000001122 | [[[[106360717, 10 | 3 |
| ENSG000001122 | CRYBG1 | chr6 | 106558485 | 106558625 | 106553454 | 106553567 | 106560802 | 106560926 | 112,106,153 | 0,0,0 | 62,86,53 | 7,10,7 | 290 | 150 | 2.17E-12 | 1.88E-10 | 1.0,1.0,1.0 | 0.821,0.816,0.79 | 0.189 | ENSG000001122 | [[[[106360717, 10 | 2 |
| ENSG000001122 | CRYBG1 | chr6 | 106560802 | 106560926 | 106558485 | 106558625 | 106563763 | 106563926 | 109,101,143 | 2,2,0 | 60,87,49 | 7,6,7 | 274 | 150 | 3.07E-07 | 9.58E-06 | 0.968,0.965,1.0 | 0.824,0.888,0.79 | 0.143 | ENSG000001122 | [[[[106360717, 10 | 2 |
| ENSG00000070 | ATP2B1 | chr12 | 89601325 | 89601433 | 89599116 | 89599299 | 89603711 | 89603925 | 23,18,33 | 0,0,0 | 11,53,40 | 0,2,7 | 258 | 150 | 0.000138259 | 0.002108272 | 1.0,1.0,1.0 | 1.0,0.939,0.769 | 0.097 | ENSG00000070 | [[[[89708596, 897 | 2 |
| ENSG00000070 | ATP2B1 | chr12 | 89604154 | 89604346 | 89603711 | 89603925 | 89611192 | 89611372 | 42,15,45 | 0,0,0 | 12,43,20 | 3,12,2 | 300 | 150 | 3.32E-09 | 1.65E-07 | 1.0,1.0,1.0 | 0.667,0.642,0.83 | 0.286 | ENSG00000070 | [[[[89708596, 897 | 3 |
| ENSG00000070 | ATP2B1 | chr12 | 89610420 | 89610508 | 89603711 | 89603925 | 89611192 | 89611372 | 10,11,25 | 0,0,0 | 8,34,11 | 3,12,2 | 238 | 150 | 1.21E-08 | 5.31E-07 | 1.0,1.0,1.0 | 0.627,0.641,0.77 | 0.319 | ENSG00000070 | [[[[89708596, 897 | 3 |
| ENSG00000185 | PARPBP | chr12 | 102148229 | 102148463 | 102120184 | 102120286 | 102153868 | 102153976 | 13,8,19 | 0,0,1 | 10,21,22 | 1,6,3 | 300 | 150 | 3.49E-05 | 0.000639312 | 1.0,1.0,0.905 | 0.833,0.636,0.78 | 0.217 | ENSG00000185 | [[[[102120183, 10 | 2 |
| ENSG00000185 | PARPBP | chr12 | 102148234 | 102148463 | 102120246 | 102120286 | 102153868 | 102153976 | 11,8,19 | 0,0,1 | 10,23,22 | 1,6,3 | 300 | 150 | 4.24E-05 | 0.000757172 | 1.0,1.0,0.905 | 0.833,0.657,0.78 | 0.21 | ENSG00000185 | [[[[102120183, 10 | 2 |
| ENSG00000185 | PARPBP | chr12 | 102178591 | 102178770 | 102165728 | 102165883 | 102182548 | 102182627 | 11,14,17 | 9,8,5 | 4,4,1 | 9,14,7 | 300 | 150 | 2.02E-05 | 0.000396998 | 0.379,0.467,0.63 | 0.182,0.125,0.06 | 0.367 | ENSG00000185 | [[[[102120183, 10 | 2 |
| ENSG00000075 | SART3 | chr12 | 108543027 | 108543152 | 108537487 | 108537595 | 108544426 | 108544478 | 42,24,48 | 0,0,0 | 24,89,60 | 4,13,9 | 275 | 150 | 2.54E-09 | 1.30E-07 | 1.0,1.0,1.0 | 0.766,0.789,0.78 | 0.22 | ENSG00000075 | [[[[108560843, 10 | 3 |
| ENSG00000075 | SART3 | chr12 | 108544426 | 108544478 | 108538933 | 108539089 | 108545138 | 108545323 | 10,13,18 | 0,0,0 | 11,23,9 | 4,3,0 | 202 | 150 | 0.000604779 | 0.007562023 | 1.0,1.0,1.0 | 0.671,0.851,1.0 | 0.159 | ENSG00000075 | [[[[108560843, 10 | 2 |
| ENSG00000075 | SART3 | chr12 | 108544426 | 108544532 | 108538933 | 108539089 | 108545138 | 108545323 | 7,3,6 | 0,0,0 | 4,14,9 | 4,3,0 | 256 | 150 | 9.92E-05 | 0.001587229 | 1.0,1.0,1.0 | 0.369,0.732,1.0 | 0.3 | ENSG00000075 | [[[[108560843, 10 | 2 |
| ENSG00000023 | ERP44 | chr9 | 100060099 | 100060172 | 100022041 | 100022226 | 100098783 | 100099040 | 12,27,43 | 0,0,0 | 29,61,77 | 6,4,4 | 223 | 150 | 2.36E-05 | 0.000455491 | 1.0,1.0,1.0 | 0.765,0.911,0.92 | 0.132 | ENSG00000023 | [[[[100098784, 10 | 3 |
| ENSG00000139 | TCHP | chr12 | 109911062 | 109911235 | 109908814 | 109908937 | 109914441 | 109914528 | 8,7,8 | 0,0,0 | 14,24,18 | 3,2,2 | 300 | 150 | 0.000206156 | 0.002992129 | 1.0,1.0,1.0 | 0.7,0.857,0.818 | 0.208 | ENSG00000139 | [[[[109900274, 10 | 2 |
| ENSG00000139 | TCHP | chr12 | 109912990 | 109913072 | 109908814 | 109908937 | 109914441 | 109914528 | 3,5,9 | 0,0,0 | 7,28,12 | 3,2,2 | 232 | 150 | 0.000275171 | 0.003831073 | 1.0,1.0,1.0 | 0.601,0.901,0.79 | 0.234 | ENSG00000139 | [[[[109900274, 10 | 2 |
| ENSG00000196 | ANAPC7 | chr12 | 110386326 | 110386469 | 110377392 | 110377617 | 110387738 | 110387892 | 46,56,47 | 0,0,0 | 20,26,30 | 2,8,1 | 293 | 150 | 1.09E-06 | 2.98E-05 | 1.0,1.0,1.0 | 0.837,0.625,0.93 | 0.2 | ENSG00000196 | [[[[110403527, 11 | 3 |
| ENSG00000089 | MAPKAPK5 | chr12 | 111867571 | 111867669 | 111866155 | 111866231 | 111870270 | 111870307 | 20,10,29 | 0,2,0 | 17,28,15 | 7,26,17 | 248 | 150 | 3.57E-10 | 2.15E-08 | 1.0,0.752,1.0 | 0.595,0.394,0.34 | 0.472 | ENSG00000089 | [[[[111842228, 11 | 2 |
| ENSG00000089 | MAPKAPK5 | chr12 | 111867571 | 111867669 | 111866155 | 111866231 | 111871084 | 111871180 | 20,8,36 | 0,0,5 | 16,30,15 | 3,13,9 | 248 | 150 | 0.000226203 | 0.003239279 | 1.0,1.0,0.813 | 0.763,0.583,0.50 | 0.322 | ENSG00000089 | [[[[111842228, 11 | 3 |
| ENSG00000089 | MAPKAPK5 | chr12 | 111868752 | 111868861 | 111866155 | 111866231 | 111870270 | 111870360 | 21,27,43 | 0,2,0 | 39,60,70 | 7,26,17 | 259 | 150 | 3.92E-09 | 1.91E-07 | 1.0,0.887,1.0 | 0.763,0.572,0.70 | 0.282 | ENSG00000089 | [[[[111842228, 11 | 2 |
| ENSG00000089 | MAPKAPK5 | chr12 | 111868752 | 111868861 | 111867571 | 111867669 | 111871084 | 111871180 | 28,31,37 | 2,0,8 | 38,72,57 | 0,2,0 | 259 | 150 | 0.000200817 | 0.002922994 | 0.89,1.0,0.728 | 1.0,0.954,1.0 | -0.112 | ENSG00000089 | [[[[111842228, 11 | 2 |
| ENSG00000137 | ADAM10 | chr15 | 58611807 | 58611991 | 58609157 | 58610517 | 58621470 | 58621510 | 120,84,141 | 0,0,0 | 88,190,130 | 5,11,6 | 300 | 150 | 1.12E-07 | 3.88E-06 | 1.0,1.0,1.0 | 0.898,0.896,0.91 | 0.097 | ENSG00000137 | [[[[58749480, 587 | 2 |
| ENSG00000137 | ADAM10 | chr15 | 58679123 | 58679282 | 58665096 | 58665197 | 58749479 | 58749770 | 50,34,78 | 0,0,0 | 33,96,70 | 3,6,5 | 300 | 150 | 4.58E-06 | 0.000106798 | 1.0,1.0,1.0 | 0.846,0.889,0.87 | 0.13 | ENSG00000137 | [[[[58749480, 587 | 3 |
| ENSG00000137 | ADAM10 | chr15 | 58717576 | 58717727 | 58665096 | 58665197 | 58749479 | 58749720 | 25,25,80 | 0,0,0 | 17,53,38 | 3,6,5 | 300 | 150 | 9.66E-08 | 3.41E-06 | 1.0,1.0,1.0 | 0.739,0.815,0.79 | 0.218 | ENSG00000137 | [[[[58749480, 587 | 3 |
| ENSG00000134 | TMEM106C | chr12 | 47966088 | 47966172 | 47965281 | 47965345 | 47966682 | 47966732 | 108,98,134 | 0,0,2 | 79,177,76 | 6,45,29 | 234 | 150 | 1.78E-15 | 2.33E-13 | 1.0,1.0,0.977 | 0.894,0.716,0.62 | 0.247 | ENSG00000134 | [[[[47963596, 479 | 2 |
| ENSG00000176 | WSB2 | chr12 | 118036337 | 118036510 | 118034674 | 118035324 | 118042840 | 118042972 | 217,166,294 | 0,0,0 | 207,340,239 | 23,47,25 | 300 | 150 | 0 | 0 | 1.0,1.0,1.0 | 0.818,0.783,0.82 | 0.191 | ENSG00000176 | [[[[118061036, 11 | 2 |
| ENSG0000013 | CLK1 | chr2 | 200856681 | 200856811 | 200854615 | 200854695 | 200856890 | 200856985 | 73,57,72 | 0,0,0 | 48,80,75 | 8,10,6 | 280 | 150 | 7.39E-10 | 4.19E-08 | 1.0,1.0,1.0 | 0.763,0.811,0.87 | 0.185 | ENSG0000013 | [[[[200864564, 20 | 2 |
| ENSG0000013 | CLK1 | chr2 | 200856890 | 200856985 | 200854615 | 200854695 | 200857717 | 200857884 | 24,13,39 | 0,0,0 | 19,49,36 | 5,1,3 | 245 | 150 | 0.000103863 | 0.001649807 | 1.0,1.0,1.0 | 0.699,0.968,0.88 | 0.151 | ENSG0000013 | [[[[200864564, 20 | 3 |
| ENSG00000184 | KNTC1 | chr12 | 122597730 | 122597938 | 122593415 | 122594385 | 122602568 | 122602740 | 30,18,26 | 2,0,0 | 5,15,14 | 2,12,6 | 300 | 150 | 7.76E-10 | 4.39E-08 | 0.882,1.0,1.0 | 0.556,0.385,0.53 | 0.468 | ENSG00000184 | [[[[122527249, 12 | 2 |
| ENSG00000184 | KNTC1 | chr12 | 122601535 | 122601625 | 122594275 | 122594385 | 122602568 | 122602740 | 21,13,21 | 2,0,0 | 18,23,12 | 2,12,6 | 240 | 150 | 2.74E-06 | 6.78E-05 | 0.868,1.0,1.0 | 0.849,0.545,0.55 | 0.306 | ENSG00000184 | [[[[122527249, 12 | 2 |
| ENSG00000139 | DENR | chr12 | 122762844 | 122762929 | 122753692 | 122753807 | 122765303 | 122765387 | 24,29,39 | 0,0,0 | 14,41,37 | 1,5,3 | 235 | 150 | 0.000103909 | 0.00164984 | 1.0,1.0,1.0 | 0.899,0.84,0.887 | 0.125 | ENSG00000139 | [[[[122752824, 12 | 2 |
| ENSG00000165 | MICU2 | chr13 | 21495160 | 21495318 | 21492699 | 21493353 | 21502925 | 21503098 | 74,92,91 | 0,0,0 | 50,119,47 | 5,13,3 | 300 | 150 | 1.94E-09 | 1.02E-07 | 1.0,1.0,1.0 | 0.833,0.821,0.88 | 0.153 | ENSG00000165 | [[[[21603939, 216 | 2 |
| ENSG00000165 | MICU2 | chr13 | 21495160 | 21496160 | 21492690 | 21493353 | 21502925 | 21503073 | 112,138,150 | 0,0,0 | 63,156,79 | 5,13,3 | 300 | 150 | 3.75E-09 | 1.84E-07 | 1.0,1.0,1.0 | 0.863,0.857,0.92 | 0.117 | ENSG00000165 | [[[[21603939, 216 | 2 |
| ENSG00000127 | TNFRSF19 | chr13 | 23590149 | 23590252 | 23570369 | 23570848 | 23615866 | 23616045 | 32,11,29 | 2,1,3 | 29,61,54 | 10,16,14 | 253 | 150 | 0.000638331 | 0.007915302 | 0.905,0.867,0.85 | 0.632,0.693,0.69 | 0.201 | ENSG00000127 | [[[[23570412, 235 | 2 |
| ENSG00000127 | TNFRSF19 | chr13 | 23593344 | 23593455 | 23570369 | 23570848 | 23615866 | 23616045 | 81,53,143 | 2,1,3 | 106,214,181 | 10,16,14 | 261 | 150 | 8.21E-05 | 0.001340711 | 0.959,0.968,0.96 | 0.859,0.885,0.88 | 0.089 | ENSG00000127 | [[[[23570412, 235 | 2 |
| ENSG00000127 | TNFRSF19 | chr13 | 23659049 | 23659214 | 23626706 | 23626792 | 23667979 | 23668082 | 26,33,110 | 0,0,3 | 66,114,135 | 6,19,6 | 300 | 150 | 1.46E-05 | 0.000297212 | 1.0,1.0,0.948 | 0.846,0.75,0.918 | 0.145 | ENSG00000127 | [[[[23570412, 235 | 2 |
| ENSG00000127 | TNFRSF19 | chr13 | 23659049 | 23659214 | 23626706 | 23626792 | 23673371 | 23674889 | 27,34,106 | 0,0,0 | 64,112,132 | 3,9,0 | 300 | 150 | 0.000276227 | 0.003844377 | 1.0,1.0,1.0 | 0.914,0.862,1.0 | 0.075 | ENSG00000127 | [[[[23570412, 235 | 4 |
| ENSG00000122 | RPL21 | chr13 | 27253764 | 27253843 | 27251308 | 27251585 | 27255241 | 27255354 | 5,6,14 | 0,0,0 | 24,32,25 | 2,5,2 | 229 | 150 | 0.000170414 | 0.00253419 | 1.0,1.0,1.0 | 0.887,0.807,0.89 | 0.138 | ENSG00000122 | [[[[27251558, 272 | 2 |
| ENSG00000278 | MYO19 | chr17 | 36509061 | 36509135 | 36507802 | 36507924 | 36511364 | 36511455 | 22,28,40 | 0,0,0 | 13,21,9 | 6,26,15 | 224 | 150 | 0 | 0 | 1.0,1.0,1.0 | 0.592,0.351,0.28 | 0.59 | ENSG00000278 | [[[[36534761, 365 | 2 |
| ENSG00000120 | HSPH1 | chr13 | 31152851 | 31152951 | 31151608 | 31151742 | 31155513 | 31155654 | 112,71,126 | 0,0,0 | 125,274,179 | 1,12,3 | 250 | 150 | 0.000817284 | 0.009781847 | 1.0,1.0,1.0 | 0.987,0.932,0.97 | 0.036 | ENSG00000120 | [[[[31161476, 311 | 2 |
| ENSG00000157 | DYRK1A | chr21 | 37486466 | 37486614 | 37472683 | 37472880 | 37490174 | 37490461 | 16,13,21 | 0,0,0 | 15,47,29 | 0,4,6 | 298 | 150 | 2.72E-05 | 0.000511903 | 1.0,1.0,1.0 | 1.0,0.855,0.709 | 0.145 | ENSG00000157 | [[[[37366773, 373 | 3 |
| ENSG00000102 | SUPT20H | chr13 | 37019341 | 37019397 | 37017244 | 37017328 | 37022010 | 37022080 | 1,5,6 | 0,0,0 | 4,6,3 | 5,7,3 | 206 | 150 | 1.07E-09 | 5.91E-08 | 1.0,1.0,1.0 | 0.368,0.384,0.42 | 0.609 | ENSG00000102 | [[[[37059559, 370 | 2 |
| ENSG00000102 | SUPT20H | chr13 | 37019341 | 37019397 | 37017244 | 37017328 | 37024034 | 37024193 | 1,5,6 | 0,0,0 | 4,6,2 | 1,3,4 | 206 | 150 | 1.05E-07 | 3.67E-06 | 1.0,1.0,1.0 | 0.744,0.593,0.26 | 0.465 | ENSG00000102 | [[[[37059559, 370 | 3 |
| ENSG00000102 | SUPT20H | chr13 | 37022010 | 37022080 | 37017244 | 37017328 | 37024034 | 37024193 | 5,6,19 | 0,0,0 | 10,16,10 | 1,3,4 | 220 | 150 | 9.98E-06 | 0.00021328 | 1.0,1.0,1.0 | 0.872,0.784,0.63 | 0.238 | ENSG00000102 | [[[[37059559, 370 | 3 |
| ENSG00000120 | PROSER1 | chr13 | 39022325 | 39022412 | 39017499 | 39017544 | 39024472 | 39024556 | 15,10,41 | 0,0,1 | 20,54,22 | 10,25,9 | 237 | 150 | 5.56E-12 | 4.51E-10 | 1.0,1.0,0.963 | 0.559,0.578,0.60 | 0.406 | ENSG00000120 | [[[[39037198, 390 | 2 |
| ENSG00000120 | PROSER1 | chr13 | 39023051 | 39023130 | 39017499 | 39017544 | 39024472 | 39024556 | 8,15,17 | 0,0,1 | 10,44,28 | 10,25,9 | 229 | 150 | 3.05E-09 | 1.54E-07 | 1.0,1.0,0.918 | 0.396,0.535,0.67 | 0.439 | ENSG00000120 | [[[[39037198, 390 | 2 |
| ENSG00000120 | PROSER1 | chr13 | 39024472 | 39024556 | 39017499 | 39017544 | 39026276 | 39026387 | 9,4,21 | 0,0,0 | 29,77,42 | 0,12,1 | 234 | 150 | 0.000161153 | 0.002412012 | 1.0,1.0,1.0 | 1.0,0.804,0.964 | 0.077 | ENSG00000120 | [[[[39037198, 390 | 3 |
| ENSG00000102 | VWA8 | chr13 | 41727193 | 41727313 | 41721369 | 41721575 | 41732079 | 41732155 | 11,14,14 | 0,0,0 | 2,9,6 | 6,4,1 | 270 | 150 | 1.04E-08 | 4.62E-07 | 1.0,1.0,1.0 | 0.156,0.556,0.76 | 0.506 | ENSG00000102 | [[[[41960853, 419 | 2 |
| ENSG00000168 | MFF | chr2 | 227328677 | 227330846 | 227325264 | 227325427 | 227332418 | 227332588 | 103,90,155 | 8,5,13 | 141,258,166 | 3,8,1 | 300 | 150 | 0.000184799 | 0.002724183 | 0.866,0.9,0.856 | 0.959,0.942,0.98 | -0.089 | ENSG00000168 | [[[[227325251, 22 | 2 |
| ENSG00000168 | MFF | chr2 | 227330625 | 227330846 | 227325258 | 227325427 | 227332418 | 227332588 | 96,84,130 | 8,5,13 | 129,240,151 | 3,8,1 | 300 | 150 | 0.000129936 | 0.002001788 | 0.857,0.894,0.83 | 0.956,0.938,0.98 | -0.099 | ENSG00000168 | [[[[227325251, 22 | 2 |
| ENSG00000168 | MFF | chr2 | 227347225 | 227347384 | 227340291 | 227340380 | 227355676 | 227355761 | 42,24,53 | 26,15,29 | 28,46,44 | 44,80,48 | 300 | 150 | 1.96E-05 | 0.000386317 | 0.447,0.444,0.47 | 0.241,0.223,0.31 | 0.197 | ENSG00000168 | [[[[227325251, 22 | 2 |
| ENSG00000168 | MFF | chr2 | 227352513 | 227352573 | 227340291 | 227340380 | 227355676 | 227355761 | 42,37,47 | 26,15,29 | 30,55,37 | 44,80,48 | 210 | 150 | 1.11E-05 | 0.000234534 | 0.536,0.638,0.53 | 0.328,0.329,0.35 | 0.233 | ENSG00000168 | [[[[227325251, 22 | 2 |
| ENSG00000168 | MFF | chr2 | 227352513 | 227352573 | 227342744 | 227342819 | 227355676 | 227355761 | 31,30,35 | 1,4,2 | 25,45,32 | 5,14,15 | 210 | 150 | 6.65E-05 | 0.001118875 | 0.957,0.843,0.92 | 0.781,0.697,0.60 | 0.215 | ENSG00000168 | [[[[227325251, 22 | 2 |
| ENSG00000136 | RCBTB2 | chr13 | 48522307 | 48522403 | 48515585 | 48515741 | 48524658 | 48524757 | 7,3,5 | 1,2,4 | 4,15,13 | 0,0,1 | 246 | 150 | 1.37E-05 | 0.00028168 | 0.81,0.478,0.433 | 1.0,1.0,0.888 | -0.389 | ENSG00000136 | [[[[48533028, 485 | 2 |
| ENSG00000136 | RCBTB1 | chr13 | 49551325 | 49551468 | 49549457 | 49549648 | 49555514 | 49555673 | 3,20,31 | 0,0,0 | 14,25,17 | 4,5,7 | 293 | 150 | 4.58E-09 | 2.21E-07 | 1.0,1.0,1.0 | 0.642,0.719,0.55 | 0.362 | ENSG00000136 | [[[[49585444, 495 | 2 |
| ENSG00000136 | RCBTB1 | chr13 | 49552177 | 49552285 | 49549457 | 49549648 | 49555514 | 49555673 | 16,12,26 | 0,0,0 | 11,18,26 | 4,5,7 | 258 | 150 | 8.26E-09 | 3.74E-07 | 1.0,1.0,1.0 | 0.615,0.677,0.68 | 0.342 | ENSG00000136 | [[[[49585444, 495 | 2 |
| ENSG00000005 | ITGA3 | chr17 | 50071985 | 50072182 | 50070843 | 50070930 | 50073915 | 50074004 | 342,359,628 | 13,9,16 | 184,465,288 | 7,39,16 | 300 | 150 | 0.000221437 | 0.00318348 | 0.929,0.952,0.95 | 0.929,0.856,0.9 | 0.049 | ENSG00000005 | [[[[50056110, 500 | 2 |
| ENSG00000088 | TMEM40 | chr3 | 12748654 | 12748792 | 12742453 | 12742507 | 12749759 | 12749840 | 3,2,2 | 0,1,1 | 0,1,2 | 3,2,1 | 288 | 150 | 8.28E-05 | 0.001349061 | 1.0,0.51,0.51 | 0.0,0.207,0.51 | 0.434 | ENSG00000088 | [[[[12759191, 127 | 2 |
| ENSG00000141 | UTP4 | chr16 | 69154392 | 69154457 | 69153583 | 69153680 | 69160355 | 69160462 | 28,21,28 | 2,0,1 | 32,57,38 | 7,10,7 | 215 | 150 | 5.70E-05 | 0.000978283 | 0.907,1.0,0.951 | 0.761,0.799,0.79 | 0.169 | ENSG00000141 | [[[[69132649, 691 | 3 |
| ENSG00000141 | UTP4 | chr16 | 69155870 | 69155993 | 69154392 | 69154457 | 69160355 | 69160462 | 51,67,80 | 0,0,0 | 52,104,64 | 6,4,4 | 273 | 150 | 5.94E-06 | 0.000135162 | 1.0,1.0,1.0 | 0.826,0.935,0.89 | 0.114 | ENSG00000141 | [[[[69132649, 691 | 2 |
| ENSG00000024 | DEPDC1 | chr1 | 68482045 | 68482897 | 68481439 | 68481612 | 68488915 | 68489034 | 9,12,30 | 0,0,0 | 19,40,42 | 8,6,15 | 300 | 150 | 3.05E-09 | 1.54E-07 | 1.0,1.0,1.0 | 0.543,0.769,0.58 | 0.368 | ENSG00000024 | [[[[68496952, 684 | 4 |
| ENSG00000157 | RER1 | chr1 | 2397115 | 2397220 | 2394004 | 2395871 | 2400856 | 2400935 | 156,150,210 | 0,0,0 | 198,342,256 | 2,16,2 | 255 | 150 | 0.000175206 | 0.002596843 | 1.0,1.0,1.0 | 0.983,0.926,0.98 | 0.035 | ENSG00000157 | [[[[2391841, 239 | 2 |
| ENSG00000068 | FTSJ1 | chrX | 48479037 | 48479116 | 48476020 | 48476396 | 48481150 | 48481203 | 28,14,33 | 0,0,0 | 23,62,40 | 2,5,2 | 229 | 150 | 0.000573244 | 0.007222236 | 1.0,1.0,1.0 | 0.883,0.89,0.929 | 0.099 | ENSG00000068 | [[[[48476199, 484 | 4 |
| ENSG00000169 | UPF3A | chr13 | 114282836 | 114282943 | 114281924 | 114282127 | 114286518 | 114286629 | 16,7,29 | 1,0,1 | 18,47,38 | 3,9,8 | 257 | 150 | 3.04E-05 | 0.000567178 | 0.903,1.0,0.944 | 0.778,0.753,0.73 | 0.194 | ENSG00000169 | [[[[114281601, 11 | 2 |
| ENSG00000258 | CEP95 | chr17 | 64516722 | 64516828 | 64514228 | 64514358 | 64522701 | 64522895 | 4,4,16 | 0,0,0 | 9,33,12 | 3,17,4 | 256 | 150 | 1.12E-08 | 4.95E-07 | 1.0,1.0,1.0 | 0.637,0.532,0.63 | 0.398 | ENSG00000258 | [[[[64506977, 645 | 3 |
| ENSG00000258 | CEP95 | chr17 | 64516722 | 64516828 | 64514228 | 64514358 | 64526070 | 64526200 | 4,4,16 | 0,0,0 | 10,33,20 | 6,5,5 | 256 | 150 | 2.29E-07 | 7.35E-06 | 1.0,1.0,1.0 | 0.494,0.795,0.70 | 0.337 | ENSG00000258 | [[[[64506977, 645 | 5 |
| ENSG00000258 | CEP95 | chr17 | 64519320 | 64519436 | 64514247 | 64514358 | 64522701 | 64522754 | 2,2,6 | 0,0,0 | 7,20,16 | 3,17,4 | 266 | 150 | 9.75E-08 | 3.44E-06 | 1.0,1.0,1.0 | 0.568,0.399,0.69 | 0.447 | ENSG00000258 | [[[[64506977, 645 | 3 |
| ENSG00000258 | CEP95 | chr17 | 64519320 | 64519436 | 64516722 | 64516828 | 64522701 | 64522754 | 4,5,10 | 0,0,0 | 3,2,5 | 0,5,3 | 266 | 150 | 3.92E-06 | 9.32E-05 | 1.0,1.0,1.0 | 1.0,0.184,0.484 | 0.444 | ENSG00000258 | [[[[64506977, 645 | 2 |
| ENSG00000258 | CEP95 | chr17 | 64525769 | 64525882 | 64514228 | 64514358 | 64526070 | 64526200 | 12,19,11 | 0,0,0 | 18,55,15 | 6,5,5 | 263 | 150 | 3.54E-07 | 1.09E-05 | 1.0,1.0,1.0 | 0.631,0.863,0.63 | 0.292 | ENSG00000258 | [[[[64506977, 645 | 5 |
| ENSG00000258 | CEP95 | chr17 | 64529287 | 64529427 | 64526070 | 64526200 | 64530925 | 64531018 | 21,5,22 | 0,0,0 | 32,84,33 | 9,32,18 | 290 | 150 | 3.39E-11 | 2.37E-09 | 1.0,1.0,1.0 | 0.648,0.576,0.48 | 0.43 | ENSG00000258 | [[[[64506977, 645 | 2 |
| ENSG00000258 | CEP95 | chr17 | 64530925 | 64531018 | 64526070 | 64526200 | 64531889 | 64532022 | 8,7,11 | 0,0,0 | 33,80,50 | 1,8,3 | 243 | 150 | 0.000179107 | 0.002646418 | 1.0,1.0,1.0 | 0.953,0.861,0.91 | 0.092 | ENSG00000258 | [[[[64506977, 645 | 3 |
| ENSG00000258 | CEP95 | chr17 | 64530925 | 64531022 | 64526070 | 64526200 | 64531889 | 64531912 | 3,2,2 | 0,0,0 | 16,49,28 | 1,8,3 | 247 | 150 | 3.46E-05 | 0.00063424 | 1.0,1.0,1.0 | 0.907,0.788,0.85 | 0.152 | ENSG00000258 | [[[[64506977, 645 | 3 |
| ENSG00000136 | TTC5 | chr14 | 20295311 | 20295526 | 20291982 | 20292127 | 20296389 | 20296446 | 22,22,38 | 0,0,0 | 17,15,10 | 0,7,1 | 300 | 150 | 9.20E-05 | 0.001482418 | 1.0,1.0,1.0 | 1.0,0.517,0.833 | 0.217 | ENSG00000136 | [[[[20305887, 203 | 2 |

| | | | | | | | | | | | | | | | | | | | | | | |
|---|---|---|---|---|---|---|---|---|---|---|---|---|---|---|---|---|---|---|---|---|---|---|
| ENSG000001383 | ZNF365 | chr10 | 62459740 | 62459797 | 62388395 | 62388576 | 62479875 | 62480288 | 6,4,8 | 0,0,2 | 0,7,3 | 2,3,4 | 207 | 150 | 4.04E-05 | 0.00072658 | 1.0,1.0,0.743 | 0.0,0.628,0.352 | 0.588 | ENSG000001383 | [[[[62374369, 623 | 2 |
| ENSG000001657 | METTL17 | chr14 | 20994793 | 20994901 | 20993117 | 20993191 | 20995164 | 20995233 | 62,44,64 | 0,0,0 | 64,108,50 | 3,4,8 | 258 | 150 | 6.42E-06 | 0.000144836 | 1.0,1.0,1.0 | 0.925,0.94,0.784 | 0.117 | ENSG000001657 | [[[[20989980, 209 | 3 |
| ENSG000001658 | METTL3 | chr14 | 21499305 | 21499371 | 21499024 | 21499137 | 21503172 | 21503577 | 23,20,18 | 0,0,0 | 19,39,15 | 0,7,2 | 216 | 150 | 6.13E-05 | 0.001043806 | 1.0,1.0,1.0 | 1.0,0.795,0.839 | 0.122 | ENSG000001658 | [[[[21511124, 215 | 6 |
| ENSG000001658 | METTL3 | chr14 | 21500912 | 21501129 | 21499763 | 21499802 | 21503172 | 21503577 | 0,0,4 | 2,2,2 | 2,24,13 | 1,1,2 | 300 | 150 | 2.76E-05 | 0.000519569 | 0.0,0.0,0.5 | 0.5,0.923,0.765 | -0.563 | ENSG000001658 | [[[[21511124, 215 | 3 |
| ENSG000001394 | NUP58 | chr13 | 25312882 | 25313032 | 25309246 | 25309282 | 25319325 | 25319350 | 42,28,97 | 0,0,0 | 34,88,73 | 5,8,11 | 300 | 150 | 1.39E-09 | 7.47E-08 | 1.0,1.0,1.0 | 0.773,0.846,0.76 | 0.204 | ENSG000001394 | [[[[25301625, 253 | 3 |
| ENSG000001394 | NUP58 | chr13 | 25312882 | 25313032 | 25309246 | 25309282 | 25320529 | 25320595 | 42,28,97 | 0,0,0 | 34,89,71 | 5,15,4 | 300 | 150 | 3.21E-09 | 1.61E-07 | 1.0,1.0,1.0 | 0.773,0.748,0.89 | 0.193 | ENSG000001394 | [[[[25301625, 253 | 4 |
| ENSG000001394 | NUP58 | chr13 | 25315356 | 25315467 | 25309246 | 25309282 | 25319325 | 25319350 | 17,12,39 | 0,0,0 | 14,58,23 | 5,8,11 | 261 | 150 | 2.70E-09 | 1.37E-07 | 1.0,1.0,1.0 | 0.617,0.806,0.54 | 0.344 | ENSG000001394 | [[[[25301625, 253 | 3 |
| ENSG000001394 | NUP58 | chr13 | 25319325 | 25319350 | 25309246 | 25309282 | 25320529 | 25320595 | 9,9,17 | 0,0,0 | 15,33,27 | 5,15,4 | 175 | 150 | 7.54E-08 | 2.73E-06 | 1.0,1.0,1.0 | 0.72,0.653,0.853 | 0.258 | ENSG000001394 | [[[[25301625, 253 | 4 |
| ENSG000001569 | GALK2 | chr15 | 49225204 | 49225349 | 49170223 | 49170375 | 49235850 | 49235941 | 3,1,4 | 0,0,0 | 16,31,20 | 1,4,2 | 295 | 150 | 0.000622311 | 0.007748933 | 1.0,1.0,1.0 | 0.891,0.798,0.83 | 0.158 | ENSG000001569 | [[[[49170247, 491 | 2 |
| ENSG000001554 | OXA1L | chr14 | 22767957 | 22768171 | 22766521 | 22766764 | 22769790 | 22769934 | 119,128,198 | 0,0,0 | 180,301,233 | 17,45,41 | 300 | 150 | 0 | 0 | 1.0,1.0,1.0 | 0.841,0.77,0.74 | 0.216 | ENSG000001554 | [[[[22766688, 227 | 2 |
| ENSG000001174 | SLC19A2 | chr1 | 169468643 | 169468737 | 169468110 | 169468252 | 169477154 | 169477757 | 5,9,15 | 0,0,0 | 8,9,6 | 2,4,2 | 244 | 150 | 9.88E-07 | 2.73E-05 | 1.0,1.0,1.0 | 0.711,0.58,0.648 | 0.354 | ENSG000001174 | [[[[169485563, 16 | 2 |
| ENSG000001649 | ALDH7A1 | chr5 | 126550195 | 126550293 | 126549503 | 126550002 | 126559239 | 126559325 | 112,84,92 | 0,0,0 | 39,119,58 | 1,15,2 | 248 | 150 | 1.36E-07 | 4.61E-06 | 1.0,1.0,1.0 | 0.959,0.828,0.94 | 0.089 | ENSG000001649 | [[[[126595007, 12 | 4 |
| ENSG000001008 | PCK2 | chr14 | 24099048 | 24099236 | 24098202 | 24098678 | 24099994 | 24100213 | 9,12,30 | 0,0,0 | 32,57,57 | 13,8,2 | 300 | 150 | 9.29E-07 | 2.58E-05 | 1.0,1.0,1.0 | 0.552,0.781,0.93 | 0.244 | ENSG000001008 | [[[[24094311, 240 | 2 |
| ENSG000001008 | PCK2 | chr14 | 24099048 | 24099236 | 24098202 | 24098678 | 24102752 | 24102890 | 9,12,30 | 0,0,0 | 32,58,55 | 3,3,3 | 300 | 150 | 0.000577235 | 0.007262907 | 1.0,1.0,1.0 | 0.842,0.906,0.90 | 0.117 | ENSG000001008 | [[[[24094311, 240 | 3 |
| ENSG000001009 | GMPR2 | chr14 | 24233218 | 24233340 | 24232421 | 24232788 | 24235736 | 24235810 | 11,25,57 | 0,0,0 | 24,55,47 | 6,15,24 | 272 | 150 | 2.98E-13 | 2.91E-11 | 1.0,1.0,1.0 | 0.688,0.669,0.51 | 0.375 | ENSG000001009 | [[[[24232929, 242 | 3 |
| ENSG000001009 | GMPR2 | chr14 | 24233218 | 24233340 | 24232421 | 24232847 | 24235736 | 24235810 | 10,7,26 | 0,0,0 | 9,6,10 | 3,6,4 | 272 | 150 | 3.20E-10 | 1.94E-08 | 1.0,1.0,1.0 | 0.623,0.355,0.58 | 0.481 | ENSG000001009 | [[[[24232929, 242 | 3 |
| ENSG000001009 | GMPR2 | chr14 | 24233218 | 24233340 | 24232871 | 24232977 | 24235736 | 24235810 | 15,13,32 | 0,0,0 | 1,19,19 | 1,11,1 | 272 | 150 | 3.76E-06 | 8.97E-05 | 1.0,1.0,1.0 | 0.355,0.488,0.91 | 0.415 | ENSG000001009 | [[[[24232929, 242 | 2 |
| ENSG000001009 | GMPR2 | chr14 | 24233478 | 24233598 | 24232421 | 24232788 | 24235736 | 24235810 | 40,48,72 | 0,0,0 | 32,77,49 | 6,15,24 | 270 | 150 | 4.25E-14 | 4.65E-12 | 1.0,1.0,1.0 | 0.748,0.74,0.531 | 0.327 | ENSG000001009 | [[[[24232929, 242 | 3 |
| ENSG000001009 | GMPR2 | chr14 | 24233478 | 24233598 | 24232749 | 24232847 | 24235736 | 24235810 | 40,48,72 | 0,0,0 | 32,76,49 | 3,6,4 | 270 | 150 | 4.36E-06 | 0.000102437 | 1.0,1.0,1.0 | 0.856,0.876,0.87 | 0.132 | ENSG000001009 | [[[[24232929, 242 | 3 |
| ENSG000001009 | GMPR2 | chr14 | 24233478 | 24233598 | 24232871 | 24232977 | 24235736 | 24235810 | 40,49,72 | 0,0,0 | 32,76,49 | 1,11,1 | 270 | 150 | 7.79E-06 | 0.000171542 | 1.0,1.0,1.0 | 0.947,0.793,0.96 | 0.098 | ENSG000001009 | [[[[24232929, 242 | 2 |
| ENSG000001009 | GMPR2 | chr14 | 24234106 | 24234241 | 24232489 | 24232788 | 24235736 | 24235820 | 1,4,1 | 0,0,0 | 17,15,6 | 6,15,24 | 285 | 150 | 1.66E-08 | 7.06E-07 | 1.0,1.0,1.0 | 0.599,0.345,0.11 | 0.647 | ENSG000001009 | [[[[24232929, 242 | 3 |
| ENSG000001009 | GMPR2 | chr14 | 24234106 | 24234241 | 24232749 | 24232847 | 24235736 | 24235820 | 1,4,1 | 0,0,0 | 12,11,6 | 3,6,4 | 285 | 150 | 7.47E-09 | 3.42E-07 | 1.0,1.0,1.0 | 0.678,0.491,0.44 | 0.463 | ENSG000001009 | [[[[24232929, 242 | 3 |
| ENSG000001009 | GMPR2 | chr14 | 24234106 | 24234241 | 24232897 | 24232977 | 24235736 | 24235820 | 1,4,1 | 0,0,0 | 13,11,7 | 1,11,1 | 285 | 150 | 1.82E-06 | 4.71E-05 | 1.0,1.0,1.0 | 0.872,0.345,0.78 | 0.332 | ENSG000001009 | [[[[24232929, 242 | 2 |
| ENSG000001421 | APP | chr21 | 25997359 | 25997416 | 25982343 | 25982477 | 26021839 | 26022042 | 121,97,177 | 22,13,44 | 123,254,177 | 51,104,69 | 207 | 150 | 1.98E-05 | 0.000389877 | 0.799,0.844,0.74 | 0.636,0.639,0.65 | 0.154 | ENSG000001421 | [[[[26170564, 261 | 2 |
| ENSG000001295 | EAPP | chr14 | 34524696 | 34524807 | 34515928 | 34516586 | 34533443 | 34533539 | 20,19,31 | 0,0,0 | 46,96,64 | 1,7,3 | 261 | 150 | 0.000500921 | 0.006436624 | 1.0,1.0,1.0 | 0.964,0.887,0.92 | 0.075 | ENSG000001295 | [[[[34539555, 345 | 2 |
| ENSG000001008 | SRP54 | chr14 | 34999557 | 34999649 | 34996676 | 34996787 | 35007282 | 35007387 | 45,20,44 | 1,0,0 | 7,40,22 | 5,11,4 | 242 | 150 | 4.61E-09 | 2.22E-07 | 0.965,1.0,1.0 | 0.465,0.693,0.77 | 0.345 | ENSG000001008 | [[[[34982992, 349 | 2 |
| ENSG000001008 | SRP54 | chr14 | 35000935 | 35001020 | 34996676 | 34996787 | 35007282 | 35007387 | 43,25,43 | 1,0,0 | 41,137,86 | 5,11,4 | 235 | 150 | 2.87E-05 | 0.000537951 | 0.965,1.0,1.0 | 0.84,0.888,0.932 | 0.102 | ENSG000001008 | [[[[34982992, 349 | 2 |
| ENSG000001008 | SRP54 | chr14 | 35014743 | 35014830 | 35007282 | 35007387 | 35018691 | 35018765 | 85,70,95 | 0,0,0 | 57,134,81 | 5,3,3 | 237 | 150 | 0.000826036 | 0.009876395 | 1.0,1.0,1.0 | 0.878,0.966,0.94 | 0.07 | ENSG000001008 | [[[[34982992, 349 | 6 |
| ENSG000001008 | SRP54 | chr14 | 35018965 | 35019074 | 35014743 | 35014830 | 35022909 | 35023080 | 71,52,112 | 0,0,0 | 70,138,96 | 9,14,14 | 259 | 150 | 7.70E-12 | 6.02E-10 | 1.0,1.0,1.0 | 0.818,0.851,0.79 | 0.177 | ENSG000001008 | [[[[34982992, 349 | 2 |
| ENSG000001349 | TPP2 | chr13 | 102664794 | 102664925 | 102657055 | 102657207 | 102674282 | 102674490 | 34,23,32 | 0,0,0 | 6,25,20 | 5,2,4 | 281 | 150 | 2.46E-07 | 7.85E-06 | 1.0,1.0,1.0 | 0.39,0.87,0.727 | 0.338 | ENSG000001349 | [[[[102596986, 10 | 2 |
| ENSG000001852 | PRPF39 | chr14 | 45110602 | 45110817 | 45109615 | 45109780 | 45112317 | 45112502 | 50,22,34 | 0,0,0 | 67,140,62 | 4,5,2 | 300 | 150 | 0.00078771 | 0.009467302 | 1.0,1.0,1.0 | 0.893,0.933,0.93 | 0.078 | ENSG000001852 | [[[[45084116, 450 | 2 |
| ENSG000001399 | TMX1 | chr14 | 51249692 | 51249765 | 51245312 | 51245358 | 51254340 | 51255654 | 48,31,60 | 0,0,0 | 52,85,62 | 4,12,14 | 223 | 150 | 1.17E-09 | 6.39E-08 | 1.0,1.0,1.0 | 0.897,0.827,0.74 | 0.176 | ENSG000001399 | [[[[51240247, 512 | 4 |
| ENSG000001005 | GNPNAT1 | chr14 | 52780678 | 52780740 | 52775193 | 52778458 | 52791427 | 52791483 | 23,10,23 | 0,0,0 | 9,36,17 | 4,0,4 | 212 | 150 | 0.000135195 | 0.002068178 | 1.0,1.0,1.0 | 0.614,1.0,0.75 | 0.212 | ENSG000001005 | [[[[52791428, 527 | 4 |
| ENSG000001005 | AHSA1 | chr14 | 77464597 | 77464686 | 77462641 | 77462759 | 77469076 | 77469247 | 193,172,232 | 0,0,2 | 364,613,336 | 16,41,16 | 239 | 150 | 4.07E-11 | 2.84E-09 | 1.0,1.0,0.986 | 0.935,0.904,0.92 | 0.073 | ENSG000001005 | [[[[77458093, 774 | 4 |
| ENSG000001005 | AHSA1 | chr14 | 77468456 | 77468508 | 77462641 | 77462759 | 77469076 | 77469247 | 144,150,186 | 0,0,2 | 230,483,268 | 16,41,16 | 202 | 150 | 7.94E-12 | 6.19E-10 | 1.0,1.0,0.986 | 0.914,0.897,0.92 | 0.083 | ENSG000001005 | [[[[77458093, 774 | 4 |
| ENSG000001267 | KTN1 | chr14 | 55663954 | 55664041 | 55661273 | 55661612 | 55671784 | 55671877 | 88,43,85 | 0,0,0 | 72,163,84 | 3,9,7 | 237 | 150 | 2.68E-06 | 6.65E-05 | 1.0,1.0,1.0 | 0.938,0.92,0.884 | 0.086 | ENSG000001267 | [[[[55580207, 555 | 4 |
| ENSG000001267 | KTN1 | chr14 | 55667240 | 55667330 | 55661521 | 55661612 | 55671784 | 55671877 | 66,63,58 | 0,0,0 | 48,129,86 | 3,9,7 | 240 | 150 | 9.30E-07 | 2.58E-05 | 1.0,1.0,1.0 | 0.909,0.9,0.885 | 0.102 | ENSG000001267 | [[[[55580207, 555 | 4 |
| ENSG000001267 | KTN1 | chr14 | 55667240 | 55667330 | 55663342 | 55664041 | 55671784 | 55671877 | 60,46,69 | 0,0,0 | 31,119,54 | 6,18,14 | 240 | 150 | 5.88E-13 | 5.51E-11 | 1.0,1.0,1.0 | 0.764,0.805,0.70 | 0.241 | ENSG000001267 | [[[[55580207, 555 | 3 |
| ENSG000001267 | KTN1 | chr14 | 55671565 | 55671655 | 55661273 | 55661612 | 55671784 | 55671877 | 126,107,156 | 0,0,0 | 110,225,109 | 3,9,7 | 240 | 150 | 6.24E-06 | 0.000141391 | 1.0,1.0,1.0 | 0.958,0.94,0.907 | 0.065 | ENSG000001267 | [[[[55580207, 555 | 4 |
| ENSG000001267 | KTN1 | chr14 | 55671565 | 55671655 | 55663342 | 55664041 | 55671784 | 55671877 | 126,107,156 | 0,0,0 | 111,227,109 | 6,18,14 | 240 | 150 | 9.63E-12 | 7.39E-10 | 1.0,1.0,1.0 | 0.92,0.887,0.83 | 0.121 | ENSG000001267 | [[[[55580207, 555 | 3 |
| ENSG000000720 | RDH11 | chr14 | 67690211 | 67690421 | 67685014 | 67685204 | 67692437 | 67692483 | 275,226,401 | 0,0,0 | 264,513,397 | 16,40,21 | 300 | 150 | 0 | 0 | 1.0,1.0,1.0 | 0.892,0.865,0.90 | 0.113 | ENSG000000720 | [[[[67695630, 676 | 2 |
| ENSG000000720 | RDH11 | chr14 | 67691139 | 67691244 | 67684509 | 67685204 | 67692437 | 67692483 | 140,133,305 | 0,0,0 | 186,319,210 | 16,40,21 | 255 | 150 | 0 | 0 | 1.0,1.0,1.0 | 0.872,0.824,0.85 | 0.15 | ENSG000000720 | [[[[67695630, 676 | 2 |
| ENSG000000720 | RDH11 | chr14 | 67691139 | 67691244 | 67684509 | 67685204 | 67695629 | 67695717 | 4,0,4 | 0,1,0 | 0,1,0 | 2,0,1 | 255 | 150 | 1.40E-06 | 3.72E-05 | 1.0,0.0,1.0 | 0.0,1.0,0.0 | 0.333 | ENSG000000720 | [[[[67695630, 676 | 4 |
| ENSG000001838 | IQGAP3 | chr1 | 156548072 | 156548243 | 156544388 | 156544472 | 156550260 | 156550351 | 43,56,67 | 0,2,0 | 17,46,19 | 1,5,8 | 300 | 150 | 2.10E-06 | 5.34E-05 | 1.0,0.933,1.0 | 0.895,0.821,0.54 | 0.225 | ENSG000001838 | [[[[156572493, 15 | 3 |
| ENSG000000501 | JKAMP | chr14 | 59495017 | 59495224 | 59484442 | 59484593 | 59498726 | 59498777 | 150,100,160 | 0,0,0 | 90,149,86 | 5,11,5 | 300 | 150 | 8.42E-09 | 3.80E-07 | 1.0,1.0,1.0 | 0.9,0.871,0.896 | 0.111 | ENSG000000501 | [[[[59484517, 594 | 3 |
| ENSG000001267 | PCNX4 | chr14 | 60107585 | 60108327 | 60091918 | 60092419 | 60115718 | 60115819 | 7,8,15 | 0,0,0 | 7,21,21 | 4,8,13 | 300 | 150 | 6.35E-10 | 3.64E-08 | 1.0,1.0,1.0 | 0.467,0.568,0.44 | 0.506 | ENSG000001267 | [[[[60091923, 600 | 3 |
| ENSG000001267 | PCNX4 | chr14 | 60114973 | 60115461 | 60091910 | 60092419 | 60115718 | 60115819 | 34,33,51 | 0,0,0 | 19,66,44 | 4,8,13 | 300 | 150 | 2.63E-10 | 1.62E-08 | 1.0,1.0,1.0 | 0.704,0.805,0.62 | 0.287 | ENSG000001267 | [[[[60091923, 600 | 3 |
| ENSG000001267 | PCNX4 | chr14 | 60124217 | 60125251 | 60115940 | 60116060 | 60125636 | 60125823 | 42,27,40 | 0,0,0 | 51,93,58 | 7,4,9 | 300 | 150 | 3.42E-07 | 1.05E-05 | 1.0,1.0,1.0 | 0.785,0.921,0.76 | 0.177 | ENSG000001267 | [[[[60091923, 600 | 3 |
| ENSG000001267 | PCNX4 | chr14 | 60125636 | 60125823 | 60115940 | 60116060 | 60133969 | 60134085 | 52,28,42 | 0,0,0 | 57,93,92 | 1,9,2 | 300 | 150 | 0.000132733 | 0.0020387 | 1.0,1.0,1.0 | 0.966,0.838,0.95 | 0.079 | ENSG000001267 | [[[[60091923, 600 | 4 |
| ENSG000001006 | HIF1A | chr14 | 61740504 | 61740627 | 61738086 | 61738373 | 61744704 | 61744813 | 396,283,621 | 0,1,1 | 266,521,364 | 6,21,11 | 273 | 150 | 1.04E-09 | 5.75E-08 | 1.0,0.994,0.997 | 0.961,0.932,0.94 | 0.05 | ENSG000001006 | [[[[61695513, 616 | 2 |
| ENSG000001006 | HIF1A | chr14 | 61740754 | 61741188 | 61738086 | 61738373 | 61744704 | 61744813 | 573,445,736 | 0,1,1 | 399,849,470 | 6,21,11 | 300 | 150 | 4.15E-08 | 1.59E-06 | 1.0,0.996,0.997 | 0.971,0.953,0.95 | 0.038 | ENSG000001006 | [[[[61695513, 616 | 2 |
| ENSG000001007 | MTHFD1 | chr14 | 64424803 | 64424931 | 64417887 | 64418024 | 64425729 | 64425827 | 149,137,184 | 1,0,0 | 71,154,93 | 3,4,4 | 278 | 150 | 0.000773386 | 0.009322961 | 0.988,1.0,1.0 | 0.927,0.954,0.92 | 0.06 | ENSG000001007 | [[[[64388353, 643 | 2 |
| ENSG000001637 | IL17RC | chr3 | 9930027 | 9930149 | 9929027 | 9929897 | 9930894 | 9930943 | 20,13,33 | 0,0,0 | 25,22,24 | 0,7,1 | 272 | 150 | 0.000337042 | 0.004557328 | 1.0,1.0,1.0 | 1.0,0.634,0.93 | 0.145 | ENSG000001637 | [[[[9917098, 9917 | 2 |
| ENSG000001637 | IL17RC | chr3 | 9930399 | 9930459 | 9929851 | 9929897 | 9930894 | 9930943 | 5,16,21 | 0,0,0 | 15,13,7 | 0,7,1 | 210 | 150 | 0.000120096 | 0.001868704 | 1.0,1.0,1.0 | 1.0,0.57,0.833 | 0.199 | ENSG000001637 | [[[[9917098, 9917 | 2 |
| ENSG000001006 | SRSF5 | chr14 | 69768137 | 69768282 | 69767156 | 69767251 | 69769181 | 69769251 | 120,124,247 | 0,0,0 | 143,230,170 | 1,11,5 | 295 | 150 | 3.33E-05 | 0.000613542 | 1.0,1.0,1.0 | 0.986,0.914,0.94 | 0.052 | ENSG000001006 | [[[[69767142, 697 | 3 |
| ENSG000001006 | SRSF5 | chr14 | 69769181 | 69769251 | 69766113 | 69767255 | 69770466 | 69770540 | 57,50,54 | 0,0,0 | 31,55,39 | 6,26,14 | 220 | 150 | 2.22E-16 | 3.17E-14 | 1.0,1.0,1.0 | 0.779,0.591,0.65 | 0.325 | ENSG000001006 | [[[[69767142, 697 | 4 |
| ENSG000001006 | SRSF5 | chr14 | 69769181 | 69769251 | 69767156 | 69767251 | 69770466 | 69770540 | 57,50,54 | 0,0,0 | 27,59,42 | 6,7,0 | 220 | 150 | 4.50E-06 | 0.00010521 | 1.0,1.0,1.0 | 0.754,0.852,1.0 | 0.131 | ENSG000001006 | [[[[69767142, 697 | 4 |
| ENSG000001007 | PCNX1 | chr14 | 70988566 | 70988699 | 70976941 | 70978648 | 71009633 | 71009724 | 2,5,14 | 3,4,8 | 11,8,9 | 0,1,2 | 283 | 150 | 0.000130327 | 0.002006595 | 0.261,0.399,0.48 | 1.0,0.809,0.705 | -0.458 | ENSG000001007 | [[[[70907459, 709 | 2 |
| ENSG000001007 | PCNX1 | chr14 | 70988566 | 70988699 | 70976941 | 70978648 | 71012999 | 71013202 | 2,5,13 | 0,0,0 | 11,9,9 | 1,9,3 | 283 | 150 | 7.22E-08 | 2.63E-06 | 1.0,1.0,1.0 | 0.854,0.346,0.61 | 0.395 | ENSG000001007 | [[[[70907459, 709 | 4 |
| ENSG000001007 | PCNX1 | chr14 | 71009633 | 71009724 | 70976941 | 70978648 | 71012999 | 71013202 | 1,1,4 | 0,0,0 | 0,3,2 | 1,9,3 | 241 | 150 | 1.49E-13 | 1.53E-11 | 1.0,1.0,1.0 | 0.0,0.172,0.293 | 0.845 | ENSG000001007 | [[[[70907459, 709 | 4 |
| ENSG000001007 | PCNX1 | chr14 | 71009633 | 71009724 | 70995740 | 70995925 | 71012999 | 71013202 | 6,3,3 | 0,0,0 | 0,5,5 | 1,6,3 | 241 | 150 | 2.36E-09 | 1.21E-07 | 1.0,1.0,1.0 | 0.0,0.342,0.509 | 0.716 | ENSG000001007 | [[[[70907459, 709 | 2 |
| ENSG000001339 | NUMB | chr14 | 73316389 | 73316422 | 73292786 | 73292825 | 73323129 | 73323204 | 2,3,7 | 0,0,0 | 5,12,0 | 0,4,6 | 183 | 150 | 6.23E-05 | 0.001059332 | 1.0,1.0,1.0 | 1.0,0.711,0.0 | 0.43 | ENSG000001339 | [[[[73458493, 734 | 2 |
| ENSG000001196 | ABCD4 | chr14 | 74299547 | 74299675 | 74296332 | 74296449 | 74300149 | 74300268 | 12,21,21 | 1,2,0 | 17,22,14 | 6,4,7 | 278 | 150 | 8.23E-05 | 0.001341915 | 0.866,0.85,1.0 | 0.605,0.748,0.51 | 0.281 | ENSG000001196 | [[[[74302875, 743 | 2 |
| ENSG000001196 | AREL1 | chr14 | 74667464 | 74667594 | 74667318 | 74667377 | 74670761 | 74670871 | 26,29,37 | 0,0,0 | 23,50,40 | 1,5,5 | 280 | 150 | 8.30E-06 | 0.000181481 | 1.0,1.0,1.0 | 0.925,0.843,0.81 | 0.14 | ENSG000001196 | [[[[74712933, 747 | 3 |
| ENSG000001196 | AREL1 | chr14 | 74669648 | 74669774 | 74667464 | 74667594 | 74670761 | 74670871 | 31,26,31 | 0,0,0 | 13,18,19 | 3,14,16 | 276 | 150 | 1.34E-13 | 1.38E-11 | 1.0,1.0,1.0 | 0.702,0.411,0.39 | 0.498 | ENSG000001196 | [[[[74712933, 747 | 2 |
| ENSG000001196 | AREL1 | chr14 | 74676140 | 74676321 | 74675698 | 74675946 | 74683295 | 74683395 | 26,21,52 | 0,0,0 | 35,49,41 | 6,9,11 | 300 | 150 | 2.16E-10 | 1.35E-08 | 1.0,1.0,1.0 | 0.745,0.731,0.65 | 0.291 | ENSG000001196 | [[[[74712933, 747 | 2 |
| ENSG000001197 | EIF2B2 | chr14 | 75005865 | 75005961 | 75004736 | 75004900 | 75007721 | 75007788 | 68,51,75 | 0,0,0 | 67,100,78 | 4,6,3 | 246 | 150 | 7.54E-05 | 0.001245923 | 1.0,1.0,1.0 | 0.911,0.91,0.941 | 0.079 | ENSG000001197 | [[[[75002921, 750 | 2 |
| ENSG000001448 | EIF2A | chr3 | 150562541 | 150562660 | 150558387 | 150558462 | 150567692 | 150567766 | 49,52,94 | 0,0,0 | 70,136,103 | 7,16,8 | 269 | 150 | 1.58E-09 | 8.43E-08 | 1.0,1.0,1.0 | 0.848,0.826,0.87 | 0.149 | ENSG000001448 | [[[[150546787, 15 | 3 |
| ENSG000001448 | EIF2A | chr3 | 150564298 | 150564381 | 150558387 | 150558462 | 150567692 | 150567766 | 33,38,59 | 0,0,0 | 28,60,52 | 7,16,8 | 233 | 150 | 8.46E-12 | 6.53E-10 | 1.0,1.0,1.0 | 0.72,0.707,0.807 | 0.255 | ENSG000001448 | [[[[150546787, 15 | 3 |
| ENSG000001448 | EIF2A | chr3 | 150568012 | 150568046 | 150564298 | 150564381 | 150568175 | 150568292 | 28,19,47 | 0,0,0 | 21,50,43 | 3,5,3 | 184 | 150 | 7.87E-05 | 0.00129255 | 1.0,1.0,1.0 | 0.851,0.891,0.92 | 0.112 | ENSG000001448 | [[[[150546787, 15 | 2 |
| ENSG000001448 | EIF2A | chr3 | 150568016 | 150568046 | 150558387 | 150558462 | 150568175 | 150568292 | 28,19,46 | 0,0,0 | 19,48,42 | 1,2,9 | 180 | 150 | 2.40E-05 | 0.000460848 | 1.0,1.0,1.0 | 0.941,0.952,0.79 | 0.104 | ENSG000001448 | [[[[150546787, 15 | 5 |
| ENSG000001448 | EIF2A | chr3 | 150568016 | 150568046 | 150564298 | 150564381 | 150568175 | 150568292 | 28,19,46 | 0,0,0 | 20,48,41 | 3,5,3 | 180 | 150 | 6.85E-05 | 0.001148514 | 1.0,1.0,1.0 | 0.847,0.889,0.91 | 0.115 | ENSG000001448 | [[[[150546787, 15 | 2 |
| ENSG000001279 | AKAP9 | chr7 | 92065230 | 92065463 | 92062273 | 92062486 | 92070904 | 92071009 | 13,1,12 | 0,0,0 | 3,6,6 | 1,4,2 | 300 | 150 | 3.87E-07 | 1.18E-05 | 1.0,1.0,1.0 | 0.6,0.429,0.6 | 0.457 | ENSG000001279 | [[[[91940862, 919 | 3 |
| ENSG000001279 | AKAP9 | chr7 | 92089384 | 92089529 | 92085494 | 92085686 | 92093096 | 92093316 | 24,13,23 | 0,0,0 | 7,20,17 | 5,2,2 | 295 | 150 | 1.79E-06 | 4.64E-05 | 1.0,1.0,1.0 | 0.416,0.836,0.81 | 0.312 | ENSG000001279 | [[[[91940862, 919 | 2 |
| ENSG000001279 | AKAP9 | chr7 | 92107292 | 92107422 | 92105677 | 92105763 | 92110121 | 92110668 | 31,29,40 | 0,0,0 | 29,60,20 | 3,7,5 | 280 | 150 | 1.95E-07 | 6.34E-06 | 1.0,1.0,1.0 | 0.838,0.821,0.68 | 0.22 | ENSG000001279 | [[[[91940862, 919 | 2 |
| ENSG000001593 | ADPGK | chr15 | 72755555 | 72755654 | 72751368 | 72752892 | 72760406 | 72760527 | 43,53,68 | 2,2,0 | 43,87,62 | 22,41,28 | 249 | 150 | 0 | 0 | 0.928,0.941,1.0 | 0.541,0.561,0.57 | 0.398 | ENSG000001593 | [[[[72783459, 727 | 2 |
| ENSG000001593 | ADPGK | chr15 | 72755555 | 72755654 | 72751368 | 72752895 | 72760406 | 72760527 | 45,52,84 | 1,2,4 | 56,100,57 | 23,69,31 | 249 | 150 | 0 | 0 | 0.964,0.94,0.927 | 0.595,0.466,0.52 | 0.415 | ENSG000001593 | [[[[72783459, 727 | 2 |
| ENSG000001054 | CARD8 | chr19 | 48249522 | 48249633 | 48240961 | 48241063 | 48255791 | 48255871 | 7,2,4 | 0,1,0 | 5,9,8 | 2,2,2 | 261 | 150 | 0.000817847 | 0.00978617 | 1.0,0.535,1.0 | 0.59,0.721,0.697 | 0.176 | ENSG000001054 | [[[[48255792, 482 | 2 |
| ENSG000001659 | IFI27L1 | chr14 | 94101813 | 94102059 | 94096886 | 94096965 | 94102476 | 94102641 | 17,8,27 | 0,0,1 | 1,5,9 | 3,4,0 | 300 | 150 | 3.77E-05 | 0.000685053 | 1.0,1.0,0.931 | 0.143,0.385,1.0 | 0.468 | ENSG000001659 | [[[[94081310, 940 | 2 |
| ENSG000001835 | SETD3 | chr14 | 99413875 | 99413934 | 99406515 | 99406590 | 99458278 | 99458535 | 23,15,15 | 1,0,0 | 17,26,21 | 3,7,4 | 209 | 150 | 2.78E-06 | 6.86E-05 | 0.943,1.0,1.0 | 0.803,0.727,0.79 | 0.208 | ENSG000001835 | [[[[99480728, 994 | 2 |

| | | | | | | | | | | | | | | | | | | | | | | |
|---|---|---|---|---|---|---|---|---|---|---|---|---|---|---|---|---|---|---|---|---|---|---|
| ENSG00000183 | SETD3 | chr14 | 99459112 | 99459185 | 99458278 | 99458535 | 99463485 | 99463578 | 11,19,16 | 0,0,0 | 15,19,10 | 4,5,1 | 223 | 150 | 1.41E-06 | 3.75E-05 | 1.0,1.0,1.0 | 0.716,0.719,0.87 | 0.231 | ENSG00000183 | [[[[99480728, 99 | 2 |
| ENSG00000140 | WARS | chr14 | 100375282 | 100375350 | 100361707 | 100361921 | 100376259 | 100376307 | 1,1,5 | 8,6,17 | 24,31,16 | 14,14,12 | 218 | 150 | 1.37E-07 | 4.64E-06 | 0.079,0.103,0.16 | 0.541,0.604,0.47 | -0.424 | ENSG00000140 | [[[[100375283, 1 | 2 |
| ENSG00000140 | WARS | chr14 | 100375282 | 100375406 | 100361707 | 100361921 | 100376259 | 100376285 | 1,6,14 | 8,6,17 | 32,44,37 | 14,14,12 | 274 | 150 | 0.000138241 | 0.002108272 | 0.064,0.354,0.31 | 0.556,0.632,0.62 | -0.362 | ENSG00000140 | [[[[100375283, 1 | 2 |
| ENSG00000065 | PKN2 | chr1 | 88805890 | 88806017 | 88805496 | 88805671 | 88821940 | 88822003 | 36,19,71 | 0,0,0 | 27,58,44 | 3,4,6 | 277 | 150 | 1.30E-06 | 3.46E-05 | 1.0,1.0,1.0 | 0.83,0.887,0.799 | 0.161 | ENSG00000065 | [[[[88684273, 88 | 5 |
| ENSG00000122 | POLM | chr7 | 44074130 | 44074233 | 44073782 | 44074025 | 44076508 | 44076629 | 4,20,31 | 1,0,0 | 15,36,18 | 2,4,4 | 253 | 150 | 0.000163816 | 0.002447545 | 0.703,1.0,1.0 | 0.816,0.842,0.72 | 0.106 | ENSG00000122 | [[[[44082251, 44 | 2 |
| ENSG00000122 | POLM | chr7 | 44074130 | 44074530 | 44073782 | 44074025 | 44076508 | 44076629 | 6,21,56 | 1,0,0 | 24,49,24 | 2,4,4 | 300 | 150 | 0.000175421 | 0.002599519 | 0.75,1.0,1.0 | 0.857,0.86,0.75 | 0.094 | ENSG00000122 | [[[[44082251, 44 | 2 |
| ENSG00000145 | SLIT2 | chr4 | 20596414 | 20596655 | 20568864 | 20569004 | 20598264 | 20598395 | 20,18,21 | 0,0,0 | 3,13,10 | 2,7,3 | 300 | 150 | 2.89E-10 | 1.76E-08 | 1.0,1.0,1.0 | 0.429,0.481,0.62 | 0.488 | ENSG00000145 | [[[[20251905, 20 | 3 |
| ENSG00000278 | GGNBP2 | chr17 | 36585299 | 36585450 | 36581343 | 36581538 | 36586049 | 36586153 | 62,43,52 | 0,0,0 | 51,104,55 | 5,6,2 | 300 | 150 | 1.25E-05 | 0.000259605 | 1.0,1.0,1.0 | 0.836,0.897,0.93 | 0.112 | ENSG00000278 | [[[[36544919, 36 | 2 |
| ENSG00000164 | PTTG1 | chr5 | 160422708 | 160422893 | 160421909 | 160422403 | 160427714 | 160427873 | 667,658,776 | 0,3,0 | 6,181,217,668 | 36,44,25 | 300 | 150 | 0 | 0 | 1.0,0.991,1.0 | 0.896,0.933,0.93 | 0.077 | ENSG00000164 | [[[[160421855, 1 | 2 |
| ENSG00000164 | PTTG1 | chr5 | 160424236 | 160424330 | 160421909 | 160422403 | 160427714 | 160427873 | 166,145,184 | 0,3,0 | 145,251,154 | 36,44,25 | 244 | 150 | 0 | 0 | 1.0,0.967,1.0 | 0.712,0.778,0.79 | 0.229 | ENSG00000164 | [[[[160421855, 1 | 2 |
| ENSG00000137 | RTCA | chr1 | 100275598 | 100275723 | 100273393 | 100273452 | 100277257 | 100277316 | 76,85,114 | 0,0,0 | 120,215,139 | 9,25,17 | 275 | 150 | 9.76E-13 | 8.84E-11 | 1.0,1.0,1.0 | 0.879,0.824,0.81 | 0.16 | ENSG00000137 | [[[[100266216, 1 | 2 |
| ENSG00000114 | UBE3A | chr15 | 25354527 | 25354683 | 25354032 | 25354426 | 25360382 | 25360527 | 36,20,33 | 0,0,0 | 18,55,39 | 3,4,1 | 300 | 150 | 0.000442997 | 0.00579294 | 1.0,1.0,1.0 | 0.75,0.873,0.951 | 0.142 | ENSG00000114 | [[[[25438489, 25 | 3 |
| ENSG00000114 | UBE3A | chr15 | 25411907 | 25411971 | 25405460 | 25405502 | 25438488 | 25438513 | 0,0,0 | 3,4,5 | 1,7,0 | 2,3,4 | 214 | 150 | 0.000513481 | 0.006573632 | 0.0,0.0,0.0 | 0.26,0.621,0.0 | -0.294 | ENSG00000114 | [[[[25438489, 25 | 2 |
| ENSG00000105 | AVL9 | chr7 | 32553726 | 32553767 | 32552228 | 32552295 | 32558928 | 32559464 | 7,5,20 | 0,0,0 | 23,57,31 | 3,6,10 | 191 | 150 | 3.73E-06 | 8.92E-05 | 1.0,1.0,1.0 | 0.858,0.882,0.70 | 0.184 | ENSG00000105 | [[[[32495489, 32 | 3 |
| ENSG00000105 | AVL9 | chr7 | 32554557 | 32554596 | 32553726 | 32553767 | 32558928 | 32559464 | 8,1,8 | 0,0,0 | 19,50,19 | 5,6,5 | 189 | 150 | 9.02E-06 | 0.00019563 | 1.0,1.0,1.0 | 0.751,0.869,0.75 | 0.21 | ENSG00000105 | [[[[32495489, 32 | 2 |
| ENSG00000105 | AVL9 | chr7 | 32558558 | 32558628 | 32552228 | 32552295 | 32558928 | 32559464 | 17,9,19 | 0,0,0 | 7,30,13 | 3,6,10 | 220 | 150 | 5.75E-09 | 2.70E-07 | 1.0,1.0,1.0 | 0.614,0.773,0.47 | 0.381 | ENSG00000105 | [[[[32495489, 32 | 3 |
| ENSG00000105 | AVL9 | chr7 | 32558558 | 32558628 | 32553726 | 32553767 | 32558928 | 32559464 | 17,9,19 | 0,0,0 | 6,28,11 | 5,6,5 | 220 | 150 | 8.56E-09 | 3.85E-07 | 1.0,1.0,1.0 | 0.45,0.761,0.6 | 0.396 | ENSG00000105 | [[[[32495489, 32 | 2 |
| ENSG00000214 | CPNE1 | chr20 | 35627279 | 35627413 | 35626562 | 35626803 | 35630740 | 35630795 | 143,177,265 | 0,0,0 | 180,433,286 | 12,22,20 | 284 | 150 | 6.08E-13 | 5.68E-11 | 1.0,1.0,1.0 | 0.888,0.912,0.88 | 0.106 | ENSG00000214 | [[[[35664760, 35 | 2 |
| ENSG00000214 | CPNE1 | chr20 | 35630438 | 35630490 | 35626566 | 35626803 | 35630740 | 35630795 | 76,55,91 | 0,0,0 | 102,196,122 | 12,22,20 | 202 | 150 | 2.20E-12 | 1.90E-10 | 1.0,1.0,1.0 | 0.863,0.869,0.81 | 0.15 | ENSG00000214 | [[[[35664760, 35 | 2 |
| ENSG00000070 | CFAP20 | chr16 | 58116040 | 58116152 | 58114809 | 58115457 | 58129031 | 58129361 | 93,90,117 | 0,0,0 | 82,190,97 | 15,15,11 | 262 | 150 | 1.14E-12 | 1.03E-10 | 1.0,1.0,1.0 | 0.758,0.879,0.83 | 0.176 | ENSG00000070 | [[[[58129032, 58 | 2 |
| ENSG00000070 | CFAP20 | chr16 | 58116871 | 58116951 | 58114809 | 58115457 | 58129031 | 58129300 | 15,24,42 | 0,0,0 | 25,53,44 | 15,15,11 | 230 | 150 | 6.11E-12 | 4.91E-10 | 1.0,1.0,1.0 | 0.521,0.697,0.72 | 0.353 | ENSG00000070 | [[[[58129032, 58 | 2 |
| ENSG00000198 | ARHGAP11A | chr15 | 32632978 | 32633108 | 32629594 | 32629762 | 32636256 | 32636342 | 109,74,158 | 0,0,0 | 49,125,71 | 6,8,5 | 280 | 150 | 1.13E-08 | 4.97E-07 | 1.0,1.0,1.0 | 0.814,0.893,0.88 | 0.136 | ENSG00000198 | [[[[32615504, 32 | 3 |
| ENSG00000137 | KNL1 | chr15 | 40628610 | 40628678 | 40628069 | 40628208 | 40640911 | 40641027 | 9,3,12 | 0,0,0 | 9,13,12 | 3,9,2 | 218 | 150 | 5.18E-08 | 1.95E-06 | 1.0,1.0,1.0 | 0.674,0.498,0.80 | 0.341 | ENSG00000137 | [[[[40594249, 40 | 2 |
| ENSG00000137 | KNL1 | chr15 | 40629323 | 40629371 | 40628073 | 40628208 | 40640911 | 40641027 | 15,4,5 | 0,0,0 | 1,12,7 | 3,9,2 | 198 | 150 | 1.20E-08 | 5.24E-07 | 1.0,1.0,1.0 | 0.202,0.503,0.72 | 0.523 | ENSG00000137 | [[[[40594249, 40 | 2 |
| ENSG00000103 | RPAP1 | chr15 | 41521737 | 41521880 | 41518005 | 41518182 | 41522097 | 41522250 | 4,12,24 | 0,0,0 | 6,25,14 | 0,1,6 | 293 | 150 | 0.000695139 | 0.008517152 | 1.0,1.0,1.0 | 1.0,0.928,0.544 | 0.176 | ENSG00000103 | [[[[41544219, 41 | 2 |
| ENSG00000103 | RPAP1 | chr15 | 41523244 | 41523354 | 41522097 | 41522250 | 41523770 | 41523844 | 4,20,43 | 0,0,0 | 10,47,13 | 5,7,0 | 260 | 150 | 6.45E-06 | 0.000145425 | 1.0,1.0,1.0 | 0.536,0.795,1.0 | 0.223 | ENSG00000103 | [[[[41544219, 41 | 2 |
| ENSG00000137 | TMEM62 | chr15 | 43181180 | 43181299 | 43169592 | 43169677 | 43184259 | 43184385 | 24,9,19 | 0,0,0 | 26,48,23 | 1,6,4 | 269 | 150 | 1.35E-05 | 0.000278725 | 1.0,1.0,1.0 | 0.935,0.817,0.76 | 0.162 | ENSG00000137 | [[[[43133586, 43 | 2 |
| ENSG00000137 | TUBGCP4 | chr15 | 43399111 | 43399174 | 43398037 | 43398179 | 43403682 | 43403799 | 3,2,2 | 0,0,0 | 8,18,25 | 3,4,1 | 213 | 150 | 6.20E-05 | 0.001054759 | 1.0,1.0,1.0 | 0.653,0.76,0.946 | 0.214 | ENSG00000137 | [[[[43371101, 43 | 2 |
| ENSG00000137 | TUBGCP4 | chr15 | 43400043 | 43400221 | 43398037 | 43398179 | 43403682 | 43403799 | 13,20,34 | 0,0,0 | 24,44,14 | 3,4,1 | 300 | 150 | 0.00013497 | 0.002065142 | 1.0,1.0,1.0 | 0.8,0.846,0.875 | 0.16 | ENSG00000137 | [[[[43371101, 43 | 2 |
| ENSG00000137 | TUBGCP4 | chr15 | 43401715 | 43401850 | 43399111 | 43399174 | 43403682 | 43403799 | 17,36,25 | 0,0,0 | 34,90,63 | 3,1,10 | 285 | 150 | 2.71E-05 | 0.000510849 | 1.0,1.0,1.0 | 0.856,0.979,0.76 | 0.132 | ENSG00000137 | [[[[43371101, 43 | 2 |
| ENSG00000005 | KMT2E | chr7 | 105063350 | 105063580 | 105040838 | 105041023 | 105066726 | 105066807 | 14,7,21 | 0,0,0 | 11,47,15 | 7,46,16 | 300 | 150 | 3.00E-15 | 3.83E-13 | 1.0,1.0,1.0 | 0.44,0.338,0.319 | 0.634 | ENSG00000005 | [[[[105014205, 1 | 2 |
| ENSG00000069 | PIGB | chr15 | 55339266 | 55339318 | 55333866 | 55334007 | 55350698 | 55350912 | 9,5,16 | 0,0,0 | 17,37,18 | 4,0,4 | 202 | 150 | 0.000359511 | 0.004825442 | 1.0,1.0,1.0 | 0.759,1.0,0.77 | 0.157 | ENSG00000069 | [[[[55319222, 55 | 3 |
| ENSG00000069 | PIGB | chr15 | 55350698 | 55350912 | 55321269 | 55321390 | 55354797 | 55354978 | 19,24,22 | 0,0,0 | 15,22,16 | 0,4,3 | 300 | 150 | 0.000229215 | 0.003272569 | 1.0,1.0,1.0 | 1.0,0.733,0.727 | 0.18 | ENSG00000069 | [[[[55319222, 55 | 7 |
| ENSG00000069 | PIGB | chr15 | 55350698 | 55350912 | 55339266 | 55339318 | 55354797 | 55354978 | 19,24,22 | 0,0,0 | 22,32,20 | 2,14,1 | 300 | 150 | 5.42E-07 | 1.59E-05 | 1.0,1.0,1.0 | 0.846,0.533,0.90 | 0.237 | ENSG00000069 | [[[[55319222, 55 | 3 |
| ENSG00000126 | RHOT1 | chr17 | 32211115 | 32211238 | 32209349 | 32209445 | 32253084 | 32253374 | 2,6,3 | 0,0,0 | 9,9,6 | 6,0,1 | 273 | 150 | 0.000223836 | 0.003211422 | 1.0,1.0,1.0 | 0.452,1.0,0.767 | 0.26 | ENSG00000126 | [[[[32142502, 32 | 2 |
| ENSG00000117 | ABCD3 | chr1 | 94475142 | 94475240 | 94473765 | 94473835 | 94478523 | 94478723 | 32,32,61 | 0,0,0 | 30,63,54 | 4,4,10 | 248 | 150 | 1.13E-07 | 3.90E-06 | 1.0,1.0,1.0 | 0.819,0.905,0.76 | 0.17 | ENSG00000117 | [[[[94418389, 94 | 3 |
| ENSG00000112 | PHF1 | chr6 | 33414965 | 33415144 | 33414724 | 33414829 | 33415589 | 33415670 | 3,4,19 | 0,0,0 | 11,17,12 | 8,9,1 | 300 | 150 | 2.40E-08 | 9.82E-07 | 1.0,1.0,1.0 | 0.407,0.486,0.85 | 0.417 | ENSG00000112 | [[[[33411014, 33 | 2 |
| ENSG00000112 | PHF1 | chr6 | 33415234 | 33415329 | 33414724 | 33414829 | 33415589 | 33415670 | 19,33,38 | 0,0,0 | 39,61,29 | 8,9,1 | 245 | 150 | 3.50E-07 | 1.08E-05 | 1.0,1.0,1.0 | 0.749,0.806,0.94 | 0.166 | ENSG00000112 | [[[[33411014, 33 | 2 |
| ENSG00000112 | PHF1 | chr6 | 33415260 | 33415329 | 33414724 | 33414829 | 33415589 | 33415670 | 19,23,37 | 0,0,0 | 40,62,29 | 8,9,1 | 219 | 150 | 8.28E-07 | 2.32E-05 | 1.0,1.0,1.0 | 0.774,0.825,0.95 | 0.15 | ENSG00000112 | [[[[33411014, 33 | 2 |
| ENSG00000182 | ANXA2 | chr15 | 60386027 | 60386086 | 60364428 | 60364523 | 60397942 | 60397986 | 23,33,54 | 23,25,40 | 80,140,59 | 25,38,28 | 209 | 150 | 5.59E-05 | 0.000960377 | 0.418,0.486,0.49 | 0.697,0.726,0.60 | -0.21 | ENSG00000182 | [[[[60397943, 60 | 2 |
| ENSG00000128 | ICE2 | chr15 | 60453602 | 60453784 | 60448847 | 60449841 | 60456656 | 60456794 | 21,7,24 | 0,0,0 | 18,35,26 | 2,4,3 | 300 | 150 | 4.24E-05 | 0.000756486 | 1.0,1.0,1.0 | 0.818,0.814,0.81 | 0.185 | ENSG00000128 | [[[[60479003, 60 | 3 |
| ENSG00000128 | ICE2 | chr15 | 60455325 | 60455442 | 60448847 | 60449841 | 60456656 | 60456794 | 19,18,36 | 0,0,0 | 17,26,22 | 2,4,3 | 267 | 150 | 1.69E-05 | 0.000337947 | 1.0,1.0,1.0 | 0.827,0.785,0.80 | 0.194 | ENSG00000128 | [[[[60479003, 60 | 3 |
| ENSG00000128 | ICE2 | chr15 | 60476062 | 60476167 | 60468060 | 60468162 | 60479002 | 60479110 | 2,3,5 | 0,0,0 | 1,9,5 | 2,5,4 | 255 | 150 | 4.67E-10 | 2.75E-08 | 1.0,1.0,1.0 | 0.227,0.514,0.42 | 0.612 | ENSG00000128 | [[[[60479003, 60 | 2 |
| ENSG00000128 | ICE2 | chr15 | 60477936 | 60478069 | 60468060 | 60468162 | 60479002 | 60479073 | 12,17,43 | 0,0,0 | 19,43,25 | 2,5,4 | 283 | 150 | 4.73E-06 | 0.00011001 | 1.0,1.0,1.0 | 0.834,0.82,0.768 | 0.193 | ENSG00000128 | [[[[60479003, 60 | 2 |
| ENSG00000115 | RTN4 | chr2 | 54973820 | 54973867 | 54973512 | 54973621 | 54982514 | 54982653 | 303,251,284 | 3,3,5 | 259,489,256 | 13,19,13 | 197 | 150 | 2.46E-05 | 0.00047001 | 0.987,0.985,0.97 | 0.938,0.951,0.93 | 0.041 | ENSG00000115 | [[[[55049745, 55 | 2 |
| ENSG00000115 | RTN4 | chr2 | 54974694 | 54974764 | 54973512 | 54973621 | 54982514 | 54982653 | 465,354,476 | 3,3,5 | 414,769,441 | 13,19,13 | 220 | 150 | 4.41E-05 | 0.000782058 | 0.991,0.988,0.98 | 0.956,0.965,0.95 | 0.028 | ENSG00000115 | [[[[55049745, 55 | 2 |
| ENSG00000129 | RPAIN | chr17 | 5425970 | 5426082 | 5422768 | 5422829 | 5428070 | 5428211 | 50,32,83 | 1,0,0 | 68,141,85 | 19,35,15 | 262 | 150 | 1.19E-14 | 1.41E-12 | 0.966,1.0,1.0 | 0.672,0.698,0.76 | 0.277 | ENSG00000129 | [[[[5420182, 542 | 2 |
| ENSG00000129 | RPAIN | chr17 | 5425970 | 5426299 | 5422768 | 5422829 | 5428070 | 5428211 | 110,73,138 | 1,0,0 | 123,232,156 | 19,35,15 | 300 | 150 | 1.33E-15 | 1.77E-13 | 0.982,1.0,1.0 | 0.764,0.768,0.83 | 0.204 | ENSG00000129 | [[[[5420182, 542 | 2 |
| ENSG00000129 | RPAIN | chr17 | 5425970 | 5428211 | 5422768 | 5422829 | 5432541 | 5432704 | 136,111,204 | 80,56,117 | 209,426,257 | 82,145,79 | 300 | 150 | 0.000237333 | 0.003367655 | 0.459,0.498,0.46 | 0.56,0.595,0.619 | -0.117 | ENSG00000129 | [[[[5420182, 542 | 3 |
| ENSG00000129 | RPAIN | chr17 | 5426235 | 5426299 | 5422768 | 5422829 | 5428070 | 5428211 | 78,55,80 | 1,0,0 | 107,179,109 | 19,35,15 | 214 | 150 | 4.04E-14 | 4.44E-12 | 0.982,1.0,1.0 | 0.798,0.782,0.83 | 0.189 | ENSG00000129 | [[[[5420182, 542 | 2 |
| ENSG00000129 | RPAIN | chr17 | 5428070 | 5428211 | 5422768 | 5422829 | 5432541 | 5432876 | 87,78,120 | 80,56,117 | 172,348,198 | 82,145,79 | 291 | 150 | 1.41E-07 | 4.75E-06 | 0.359,0.418,0.34 | 0.52,0.553,0.564 | -0.171 | ENSG00000129 | [[[[5420182, 542 | 3 |
| ENSG00000166 | CLPX | chr15 | 65156777 | 65156932 | 65155691 | 65155856 | 65158686 | 65158751 | 4,3,3 | 0,0,0 | 5,10,14 | 3,0,3 | 300 | 150 | 0.000511593 | 0.006553859 | 1.0,1.0,1.0 | 0.455,1.0,0.7 | 0.282 | ENSG00000166 | [[[[65185075, 65 | 2 |
| ENSG00000106 | BRAT1 | chr7 | 2543589 | 2543962 | 2543203 | 2543323 | 2547323 | 2547478 | 34,53,126 | 0,0,0 | 94,216,177 | 8,2,7 | 300 | 150 | 7.90E-05 | 0.001295064 | 1.0,1.0,1.0 | 0.855,0.982,0.92 | 0.079 | ENSG00000106 | [[[[2555487, 255 | 2 |
| ENSG00000156 | NPTN | chr15 | 73591965 | 73592137 | 73573661 | 73573795 | 73633124 | 73633243 | 102,119,287 | 0,2,5 | 149,234,243 | 2,17,9 | 300 | 150 | 0.000718025 | 0.008746932 | 1.0,0.967,0.966 | 0.974,0.873,0.93 | 0.052 | ENSG00000156 | [[[[73633125, 73 | 3 |
| ENSG00000140 | SCAMP2 | chr15 | 74848599 | 74848645 | 74845472 | 74845593 | 74851352 | 74851481 | 31,38,67 | 0,0,0 | 85,130,76 | 6,12,8 | 196 | 150 | 4.27E-07 | 1.29E-05 | 1.0,1.0,1.0 | 0.916,0.892,0.87 | 0.104 | ENSG00000140 | [[[[74873199, 74 | 2 |
| ENSG00000140 | SCAMP2 | chr15 | 74850513 | 74850673 | 74843731 | 74845593 | 74851352 | 74851481 | 115,105,158 | 0,0,0 | 126,260,167 | 6,12,8 | 300 | 150 | 1.08E-07 | 3.73E-06 | 1.0,1.0,1.0 | 0.913,0.915,0.91 | 0.086 | ENSG00000140 | [[[[74873199, 74 | 2 |
| ENSG00000140 | COMMD4 | chr15 | 75339263 | 75339344 | 75338984 | 75339104 | 75339964 | 75340271 | 68,187,246 | 0,1,0 | 157,339,232 | 7,13,2 | 231 | 150 | 7.26E-05 | 0.001205717 | 1.0,0.992,1.0 | 0.936,0.944,0.98 | 0.042 | ENSG00000140 | [[[[75336063, 75 | 2 |
| ENSG00000140 | COMMD4 | chr15 | 75339701 | 75339878 | 75338984 | 75339104 | 75339964 | 75340073 | 95,239,399 | 0,1,0 | 179,347,262 | 7,13,2 | 300 | 150 | 7.97E-06 | 0.000175152 | 1.0,0.992,1.0 | 0.927,0.93,0.985 | 0.05 | ENSG00000140 | [[[[75336063, 75 | 2 |
| ENSG00000140 | MAN2C1 | chr15 | 75362641 | 75362748 | 75362342 | 75362453 | 75364487 | 75364665 | 5,11,18 | 0,0,0 | 5,10,10 | 0,2,5 | 257 | 150 | 6.85E-05 | 0.001148867 | 1.0,1.0,1.0 | 1.0,0.745,0.539 | 0.239 | ENSG00000140 | [[[[75368483, 75 | 2 |
| ENSG00000167 | FBXO22 | chr15 | 75929883 | 75930049 | 75914109 | 75914205 | 75932684 | 75932798 | 50,57,60 | 0,0,0 | 28,83,33 | 7,5,7 | 300 | 150 | 5.15E-09 | 2.44E-07 | 1.0,1.0,1.0 | 0.667,0.892,0.70 | 0.246 | ENSG00000167 | [[[[75903878, 75 | 2 |
| ENSG00000136 | IREB2 | chr15 | 78483317 | 78483434 | 78478296 | 78478397 | 78485704 | 78485840 | 32,41,75 | 0,0,0 | 40,100,52 | 2,1,7 | 267 | 150 | 0.000324633 | 0.004422736 | 1.0,1.0,1.0 | 0.918,0.983,0.80 | 0.097 | ENSG00000136 | [[[[78438216, 78 | 2 |
| ENSG00000064 | BTBD1 | chr15 | 83041727 | 83041925 | 83030135 | 83030328 | 83056388 | 83056479 | 145,103,234 | 0,0,0 | 145,260,228 | 6,19,8 | 300 | 150 | 1.33E-09 | 7.18E-08 | 1.0,1.0,1.0 | 0.924,0.872,0.93 | 0.09 | ENSG00000064 | [[[[83066751, 83 | 2 |
| ENSG00000064 | BTBD1 | chr15 | 83050072 | 83050178 | 83030135 | 83030328 | 83056388 | 83056545 | 75,57,117 | 0,0,0 | 45,114,97 | 6,19,8 | 256 | 150 | 6.09E-12 | 4.90E-10 | 1.0,1.0,1.0 | 0.815,0.779,0.87 | 0.176 | ENSG00000064 | [[[[83066751, 83 | 2 |
| ENSG00000163 | HIPK1 | chr1 | 113956626 | 113956811 | 113955562 | 113955649 | 113958065 | 113958291 | 23,11,40 | 0,0,0 | 22,47,47 | 1,5,4 | 300 | 150 | 3.90E-05 | 0.000703997 | 1.0,1.0,1.0 | 0.917,0.825,0.85 | 0.134 | ENSG00000163 | [[[[113929324, 1 | 2 |
| ENSG00000140 | FANCI | chr15 | 89292941 | 89293063 | 89292687 | 89292864 | 89294914 | 89295094 | 27,30,74 | 0,0,0 | 17,25,18 | 2,8,1 | 272 | 150 | 3.95E-07 | 1.20E-05 | 1.0,1.0,1.0 | 0.824,0.633,0.90 | 0.212 | ENSG00000140 | [[[[89243979, 89 | 2 |
| ENSG00000140 | FANCI | chr15 | 89307475 | 89307529 | 89306006 | 89306194 | 89312903 | 89312972 | 25,14,31 | 0,0,0 | 6,15,20 | 2,2,13 | 204 | 150 | 6.13E-09 | 2.85E-07 | 1.0,1.0,1.0 | 0.688,0.847,0.53 | 0.311 | ENSG00000140 | [[[[89243979, 89 | 2 |
| ENSG00000143 | PPOX | chr1 | 161167370 | 161167486 | 161167099 | 161167234 | 161169659 | 161169720 | 1,9,12 | 0,0,3 | 1,12,8 | 6,11,6 | 266 | 150 | 4.95E-05 | 0.000865503 | 1.0,1.0,0.693 | 0.086,0.381,0.42 | 0.599 | ENSG00000143 | [[[[161166426, 1 | 4 |
| ENSG00000143 | PPOX | chr1 | 161167994 | 161168127 | 161167099 | 161167486 | 161169659 | 161169720 | 7,14,30 | 0,0,0 | 7,19,10 | 1,7,2 | 283 | 150 | 1.23E-07 | 4.20E-06 | 1.0,1.0,1.0 | 0.788,0.59,0.726 | 0.299 | ENSG00000143 | [[[[161166426, 1 | 3 |
| ENSG00000143 | PPOX | chr1 | 161168992 | 161169183 | 161167099 | 161167234 | 161169659 | 161169720 | 17,25,47 | 0,0,3 | 11,30,16 | 6,11,6 | 300 | 150 | 1.86E-08 | 7.85E-07 | 1.0,1.0,0.887 | 0.478,0.577,0.57 | 0.42 | ENSG00000143 | [[[[161166426, 1 | 4 |
| ENSG00000143 | PPOX | chr1 | 161168992 | 161169183 | 161167338 | 161167486 | 161169659 | 161169720 | 17,25,47 | 0,0,0 | 11,29,18 | 1,7,2 | 300 | 150 | 7.05E-07 | 2.01E-05 | 1.0,1.0,1.0 | 0.846,0.674,0.81 | 0.221 | ENSG00000143 | [[[[161166426, 1 | 3 |
| ENSG00000140 | POLG | chr15 | 89320765 | 89321260 | 89319227 | 89319262 | 89321735 | 89321853 | 10,19,43 | 0,0,0 | 29,57,70 | 4,10,2 | 300 | 150 | 2.98E-06 | 7.30E-05 | 1.0,1.0,1.0 | 0.784,0.74,0.946 | 0.177 | ENSG00000140 | [[[[89334673, 89 | 2 |
| ENSG00000086 | EIF2AK1 | chr7 | 6037423 | 6037524 | 6028917 | 6029032 | 6040891 | 6041219 | 196,176,311 | 1,0,0 | 255,417,226 | 18,40,16 | 251 | 150 | 0 | 0 | 0.992,1.0,1.0 | 0.894,0.862,0.89 | 0.114 | ENSG00000086 | [[[[6058966, 605 | 2 |
| ENSG00000086 | EIF2AK1 | chr7 | 6038559 | 6038671 | 6028917 | 6029032 | 6040891 | 6041219 | 312,217,430 | 1,0,0 | 263,583,379 | 18,40,16 | 262 | 150 | 1.11E-16 | 1.65E-14 | 0.994,1.0,1.0 | 0.893,0.893,0.93 | 0.092 | ENSG00000086 | [[[[6058966, 605 | 2 |
| ENSG00000173 | CHD2 | chr15 | 92998148 | 92998261 | 92997252 | 92997403 | 93002176 | 93002317 | 4,1,6 | 0,0,0 | 4,3,7 | 1,4,3 | 263 | 150 | 7.53E-08 | 2.73E-06 | 1.0,1.0,1.0 | 0.695,0.3,0.571 | 0.478 | ENSG00000173 | [[[[92900324, 92 | 2 |
| ENSG00000100 | GSS | chr20 | 34941712 | 34941829 | 34936942 | 34937023 | 34942930 | 34943006 | 69,68,123 | 0,1,0 | 50,87,63 | 6,8,14 | 267 | 150 | 5.15E-11 | 3.53E-09 | 1.0,0.974,1.0 | 0.824,0.859,0.71 | 0.191 | ENSG00000100 | [[[[34955727, 34 | 2 |
| ENSG00000100 | GSS | chr20 | 34942487 | 34942585 | 34936942 | 34937023 | 34942930 | 34942980 | 5,1,2 | 0,1,0 | 2,2,1 | 6,8,14 | 248 | 150 | 1.41E-07 | 4.74E-06 | 1.0,0.377,1.0 | 0.168,0.131,0.04 | 0.679 | ENSG00000100 | [[[[34955727, 34 | 2 |
| ENSG00000100 | GSS | chr20 | 34942487 | 34942627 | 34936942 | 34937023 | 34942930 | 34943006 | 61,68,114 | 0,1,0 | 57,102,64 | 6,8,14 | 290 | 150 | 4.12E-10 | 2.45E-08 | 1.0,0.972,1.0 | 0.831,0.868,0.70 | 0.19 | ENSG00000100 | [[[[34955727, 34 | 2 |
| ENSG00000103 | TTC23 | chr15 | 99234987 | 99235080 | 99228532 | 99228732 | 99249416 | 99249610 | 0,0,0 | 2,1,4 | 2,1,4 | 3,9,6 | 243 | 150 | 0.000479051 | 0.006204779 | 0.0,0.0,0.0 | 0.292,0.064,0.29 | -0.216 | ENSG00000103 | [[[[99249171, 99 | 3 |
| ENSG00000163 | GTPBP8 | chr3 | 112996891 | 112996991 | 112995134 | 112995265 | 113014376 | 113015060 | 16,16,18 | 0,0,0 | 9,6,9 | 3,5,1 | 250 | 150 | 2.43E-07 | 7.74E-06 | 1.0,1.0,1.0 | 0.643,0.419,0.84 | 0.365 | ENSG00000163 | [[[[112990984, 1 | 3 |
| ENSG00000176 | CSTF3 | chr11 | 33086987 | 33087141 | 33085712 | 33085773 | 33090531 | 33090727 | 86,54,75 | 0,0,0 | 58,135,72 | 9,20,10 | 300 | 150 | 4.95E-13 | 4.69E-11 | 1.0,1.0,1.0 | 0.763,0.771,0.78 | 0.228 | ENSG00000176 | [[[[33161299, 33 | 2 |

| | | | | | | | | | | | | | | | | | | | | | | |
|---|---|---|---|---|---|---|---|---|---|---|---|---|---|---|---|---|---|---|---|---|---|---|
| ENSG00000176 | CSTF3 | chr11 | 33090531 | 33090727 | 33085712 | 33085773 | 33092270 | 33092340 | 41,53,67 | 0,0,0 | 57,86,59 | 3,4,5 | 300 | 150 | 3.82E-05 | 0.000692443 | 1.0,1.0,1.0 | 0.905,0.915,0.85 | 0.108 | ENSG00000176 | [[[[33161299, 33 | 3 |
| ENSG00000176 | CSTF3 | chr11 | 33092270 | 33092340 | 33085712 | 33085773 | 33096305 | 33096408 | 30,45,35 | 0,0,0 | 41,67,44 | 6,6,2 | 220 | 150 | 6.74E-06 | 0.000150751 | 1.0,1.0,1.0 | 0.823,0.884,0.93 | 0.118 | ENSG00000176 | [[[[33161299, 33 | 4 |
| ENSG00000131 | SNRPA1 | chr15 | 101289813 | 101289922 | 101287655 | 101287702 | 101295096 | 101295222 | 0,0,0 | 3,1,3 | 8,16,3 | 3,9,6 | 259 | 150 | 5.19E-08 | 1.96E-06 | 0.0,0.0,0.0 | 0.607,0.507,0.22 | -0.446 | ENSG00000131 | [[[[101295097, 1 | 2 |
| ENSG00000198 | TSEN15 | chr1 | 184054353 | 184054435 | 184051650 | 184051890 | 184072156 | 184072298 | 1,10,20 | 0,0,0 | 6,18,8 | 2,2,2 | 232 | 150 | 0.000725433 | 0.00883153 | 1.0,1.0,1.0 | 0.66,0.853,0.721 | 0.255 | ENSG00000198 | [[[[184051730, 1 | 2 |
| ENSG00000169 | NMD3 | chr3 | 161233398 | 161233479 | 161224929 | 161225018 | 161234726 | 161234836 | 34,15,32 | 0,0,0 | 16,44,35 | 7,19,17 | 231 | 150 | 1.20E-14 | 1.42E-12 | 1.0,1.0,1.0 | 0.597,0.601,0.57 | 0.41 | ENSG00000169 | [[[[161221311, 1 | 2 |
| ENSG00000169 | NMD3 | chr3 | 161234726 | 161234855 | 161227246 | 161227343 | 161235121 | 161235212 | 35,26,64 | 0,0,0 | 49,104,98 | 2,4,5 | 279 | 150 | 0.000621929 | 0.007745956 | 1.0,1.0,1.0 | 0.929,0.933,0.91 | 0.075 | ENSG00000169 | [[[[161221311, 1 | 2 |
| ENSG00000135 | UBQLN1 | chr9 | 83665029 | 83665145 | 83663874 | 83664043 | 83669184 | 83669327 | 262,222,405 | 0,0,0 | 219,427,313 | 17,54,21 | 266 | 150 | 0 | 0 | 1.0,1.0,1.0 | 0.879,0.817,0.89 | 0.137 | ENSG00000135 | [[[[83707500, 83 | 2 |
| ENSG00000205 | RNPS1 | chr16 | 2264175 | 2264331 | 2263095 | 2263287 | 2268054 | 2268071 | 111,261,397 | 0,0,0 | 251,614,357 | 19,35,19 | 300 | 150 | 6.66E-16 | 9.20E-14 | 1.0,1.0,1.0 | 0.869,0.898,0.90 | 0.11 | ENSG00000205 | [[[[2268055, 226 | 2 |
| ENSG00000119 | ATAD2B | chr2 | 23857414 | 23857503 | 23833918 | 23834078 | 23864808 | 23864924 | 5,2,5 | 0,0,0 | 1,8,12 | 0,6,7 | 239 | 150 | 2.80E-07 | 8.82E-06 | 1.0,1.0,1.0 | 1.0,0.456,0.518 | 0.342 | ENSG00000119 | [[[[23926555, 23 | 2 |
| ENSG00000156 | CCT8 | chr21 | 29064967 | 29065105 | 29063351 | 29063530 | 29067555 | 29067705 | 372,364,432 | 0,0,0 | 355,713,364 | 20,50,22 | 288 | 150 | 0 | 0 | 1.0,1.0,1.0 | 0.902,0.881,0.89 | 0.107 | ENSG00000156 | [[[[29073531, 29 | 3 |
| ENSG00000167 | DUS2 | chr16 | 68038005 | 68038149 | 68023252 | 68023351 | 68049504 | 68049533 | 35,27,46 | 0,0,0 | 16,33,37 | 4,13,4 | 294 | 150 | 6.13E-10 | 3.52E-08 | 1.0,1.0,1.0 | 0.671,0.564,0.82 | 0.313 | ENSG00000167 | [[[[68023284, 68 | 2 |
| ENSG00000168 | ANKS3 | chr16 | 4700969 | 4701134 | 4699051 | 4699176 | 4702101 | 4702242 | 12,35,28 | 0,0,0 | 45,47,31 | 1,3,6 | 300 | 150 | 4.38E-05 | 0.000778382 | 1.0,1.0,1.0 | 0.957,0.887,0.72 | 0.145 | ENSG00000168 | [[[[4733938, 473 | 2 |
| ENSG00000170 | RALGAPB | chr20 | 38535085 | 38535207 | 38532729 | 38532859 | 38546242 | 38546430 | 3,1,9 | 0,0,0 | 2,4,3 | 2,4,0 | 272 | 150 | 7.50E-05 | 0.001241117 | 1.0,1.0,1.0 | 0.355,0.355,1.0 | 0.43 | ENSG00000170 | [[[[38472843, 38 | 3 |
| ENSG00000140 | PMM2 | chr16 | 8806315 | 8806407 | 8801798 | 8801910 | 8811078 | 8811111 | 49,31,50 | 0,0,2 | 46,113,65 | 21,20,6 | 242 | 150 | 1.34E-08 | 5.81E-07 | 1.0,1.0,0.939 | 0.576,0.778,0.87 | 0.238 | ENSG00000140 | [[[[8797839, 879 | 2 |
| ENSG00000171 | RSL1D1 | chr16 | 11847667 | 11847806 | 11846694 | 11846843 | 11851407 | 11851580 | 151,151,197 | 0,0,0 | 308,602,362 | 12,38,19 | 289 | 150 | 1.34E-11 | 1.00E-09 | 1.0,1.0,1.0 | 0.93,0.892,0.908 | 0.09 | ENSG00000171 | [[[[11851408, 11 | 2 |
| ENSG00000140 | PARN | chr16 | 14584748 | 14584791 | 14584346 | 14584422 | 14593300 | 14593378 | 12,14,19 | 0,0,2 | 8,24,13 | 6,3,1 | 193 | 150 | 0.000628602 | 0.007815736 | 1.0,1.0,0.881 | 0.509,0.861,0.91 | 0.2 | ENSG00000140 | [[[[14630107, 14 | 2 |
| ENSG00000179 | PDXDC1 | chr16 | 15001775 | 15001788 | 14998339 | 14998405 | 15006393 | 15006583 | 1,6,1 | 0,0,0 | 3,4,3 | 3,2,2 | 163 | 150 | 2.70E-06 | 6.69E-05 | 1.0,1.0,1.0 | 0.479,0.648,0.58 | 0.431 | ENSG00000179 | [[[[14975026, 14 | 2 |
| ENSG00000179 | PDXDC1 | chr16 | 15006393 | 15006583 | 15001775 | 15001788 | 15008778 | 15008840 | 90,44,120 | 0,0,0 | 38,118,87 | 4,13,5 | 300 | 150 | 3.53E-09 | 1.74E-07 | 1.0,1.0,1.0 | 0.826,0.819,0.89 | 0.153 | ENSG00000179 | [[[[14975026, 14 | 2 |
| ENSG00000103 | ABCC1 | chr16 | 16083365 | 16083542 | 16079351 | 16079478 | 16090404 | 16090588 | 20,28,55 | 0,0,0 | 17,58,44 | 1,4,3 | 300 | 150 | 0.000420001 | 0.005526108 | 1.0,1.0,1.0 | 0.895,0.879,0.88 | 0.115 | ENSG00000103 | [[[[15949616, 15 | 2 |
| ENSG00000196 | ZNF140 | chr12 | 133083486 | 133083561 | 133081272 | 133081329 | 133105509 | 133107526 | 3,1,7 | 0,0,0 | 7,9,3 | 1,3,3 | 225 | 150 | 9.74E-06 | 0.000208972 | 1.0,1.0,1.0 | 0.824,0.667,0.4 | 0.37 | ENSG00000196 | [[[[133080923, 1 | 2 |
| ENSG00000196 | ZNF140 | chr12 | 133100986 | 133101053 | 133081272 | 133081329 | 133105509 | 133107428 | 2,2,6 | 0,0,0 | 4,9,5 | 1,3,3 | 217 | 150 | 7.20E-06 | 0.000159661 | 1.0,1.0,1.0 | 0.734,0.675,0.53 | 0.352 | ENSG00000196 | [[[[133080923, 1 | 2 |
| ENSG00000146 | ZMYM4 | chr1 | 35389947 | 35390098 | 35387453 | 35387604 | 35392211 | 35392352 | 43,44,54 | 0,0,0 | 10,41,31 | 3,5,6 | 300 | 150 | 1.13E-08 | 4.96E-07 | 1.0,1.0,1.0 | 0.625,0.804,0.72 | 0.283 | ENSG00000146 | [[[[35268709, 35 | 2 |
| ENSG00000186 | KIF18B | chr17 | 44934506 | 44934617 | 44934232 | 44934430 | 44935258 | 44935416 | 25,35,54 | 0,0,1 | 41,71,43 | 2,7,5 | 261 | 150 | 1.94E-05 | 0.000383359 | 1.0,1.0,0.969 | 0.922,0.854,0.83 | 0.12 | ENSG00000186 | [[[[44947628, 44 | 2 |
| ENSG00000186 | KIF18B | chr17 | 44934830 | 44934935 | 44934232 | 44934430 | 44935258 | 44935416 | 16,18,24 | 0,0,1 | 23,58,40 | 2,7,5 | 255 | 150 | 3.93E-05 | 0.000709031 | 1.0,1.0,0.934 | 0.871,0.83,0.825 | 0.136 | ENSG00000186 | [[[[44947628, 44 | 2 |
| ENSG00000112 | MRPS30 | chr5 | 44813105 | 44813282 | 44811008 | 44811154 | 44814912 | 44815066 | 93,76,103 | 0,0,0 | 75,155,68 | 9,12,2 | 300 | 150 | 1.52E-08 | 6.51E-07 | 1.0,1.0,1.0 | 0.806,0.866,0.94 | 0.128 | ENSG00000112 | [[[[44808947, 44 | 2 |
| ENSG00000169 | XPO6 | chr16 | 28117317 | 28117462 | 28112903 | 28113050 | 28125688 | 28125848 | 64,50,96 | 0,0,0 | 64,135,89 | 1,7,10 | 295 | 150 | 1.64E-06 | 4.29E-05 | 1.0,1.0,1.0 | 0.97,0.907,0.819 | 0.101 | ENSG00000169 | [[[[28211366, 28 | 2 |
| ENSG00000169 | XPO6 | chr16 | 28121669 | 28121762 | 28112903 | 28113050 | 28125688 | 28125848 | 15,26,47 | 0,0,0 | 10,47,19 | 1,7,10 | 243 | 150 | 2.69E-08 | 1.09E-06 | 1.0,1.0,1.0 | 0.861,0.806,0.54 | 0.264 | ENSG00000169 | [[[[28211366, 28 | 2 |
| ENSG00000169 | XPO6 | chr16 | 28175897 | 28176095 | 28169749 | 28169909 | 28180940 | 28181031 | 74,34,88 | 0,0,0 | 25,49,50 | 6,16,8 | 300 | 150 | 2.28E-14 | 2.59E-12 | 1.0,1.0,1.0 | 0.676,0.605,0.75 | 0.32 | ENSG00000169 | [[[[28211366, 28 | 2 |
| ENSG00000169 | XPO6 | chr16 | 28175897 | 28176095 | 28169749 | 28169909 | 28211365 | 28211550 | 73,31,87 | 1,0,2 | 25,51,50 | 10,3,14 | 300 | 150 | 5.40E-08 | 2.03E-06 | 0.973,1.0,0.956 | 0.556,0.895,0.64 | 0.279 | ENSG00000169 | [[[[28211366, 28 | 3 |
| ENSG00000067 | RHOA | chr3 | 49362495 | 49362626 | 49359144 | 49360382 | 49375433 | 49375591 | 7,286,531,021 | 6,1,4 | 5,371,291,746 | 86,132,111 | 281 | 150 | 0 | 0 | 0.985,0.997,0.99 | 0.769,0.839,0.78 | 0.195 | ENSG00000067 | [[[[49411820, 49 | 2 |
| ENSG00000067 | RHOA | chr3 | 49368427 | 49368548 | 49359144 | 49360382 | 49375433 | 49375591 | 495,483,759 | 6,1,4 | 490,862,549 | 86,132,111 | 271 | 150 | 0 | 0 | 0.979,0.996,0.99 | 0.759,0.783,0.73 | 0.231 | ENSG00000067 | [[[[49411820, 49 | 2 |
| ENSG00000188 | CLN3 | chr16 | 28486346 | 28486490 | 28484005 | 28484118 | 28487455 | 28487541 | 11,35,44 | 0,0,1 | 66,95,93 | 9,16,7 | 294 | 150 | 5.06E-07 | 1.50E-05 | 1.0,1.0,0.957 | 0.789,0.752,0.87 | 0.182 | ENSG00000188 | [[[[28492020, 28 | 2 |
| ENSG00000169 | SPNS1 | chr16 | 28981469 | 28981615 | 28979404 | 28979471 | 28982856 | 28982922 | 13,45,84 | 0,0,1 | 71,121,71 | 5,3,4 | 296 | 150 | 0.000714128 | 0.008709208 | 1.0,1.0,0.977 | 0.878,0.953,0.9 | 0.082 | ENSG00000169 | [[[[28974778, 28 | 3 |
| ENSG00000169 | SPNS1 | chr16 | 28981900 | 28982056 | 28981454 | 28981615 | 28982856 | 28982922 | 17,56,98 | 0,0,0 | 103,149,125 | 6,6,7 | 300 | 150 | 8.08E-06 | 0.000177233 | 1.0,1.0,1.0 | 0.896,0.925,0.89 | 0.093 | ENSG00000169 | [[[[28974778, 28 | 2 |
| ENSG00000169 | SPNS1 | chr16 | 28981900 | 28982056 | 28981454 | 28981615 | 28983191 | 28983290 | 17,56,98 | 0,0,0 | 102,145,123 | 2,10,0 | 300 | 150 | 0.000765876 | 0.009239744 | 1.0,1.0,1.0 | 0.962,0.879,1.0 | 0.053 | ENSG00000169 | [[[[28974778, 28 | 3 |
| ENSG00000169 | SPNS1 | chr16 | 28982355 | 28982545 | 28977907 | 28978044 | 28982856 | 28982922 | 28,42,76 | 0,0,0 | 72,119,78 | 7,13,4 | 300 | 150 | 8.29E-08 | 2.97E-06 | 1.0,1.0,1.0 | 0.837,0.821,0.90 | 0.145 | ENSG00000169 | [[[[28974778, 28 | 5 |
| ENSG00000169 | SPNS1 | chr16 | 28982355 | 28982545 | 28981469 | 28981615 | 28982856 | 28982922 | 29,48,88 | 0,0,0 | 76,135,84 | 6,6,7 | 300 | 150 | 1.48E-06 | 3.89E-05 | 1.0,1.0,1.0 | 0.864,0.918,0.85 | 0.12 | ENSG00000169 | [[[[28974778, 28 | 2 |
| ENSG00000169 | SPNS1 | chr16 | 28982856 | 28982922 | 28981454 | 28981615 | 28983191 | 28983290 | 12,26,62 | 0,0,0 | 70,78,63 | 2,10,0 | 216 | 150 | 0.000350437 | 0.004717777 | 1.0,1.0,1.0 | 0.96,0.844,1.0 | 0.065 | ENSG00000169 | [[[[28974778, 28 | 3 |
| ENSG00000117 | MTFR1L | chr1 | 25823237 | 25823323 | 25823018 | 25823128 | 25826614 | 25826806 | 2,4,4 | 0,0,0 | 4,9,4 | 0,10,6 | 236 | 150 | 4.56E-07 | 1.36E-05 | 1.0,1.0,1.0 | 1.0,0.364,0.298 | 0.446 | ENSG00000117 | [[[[25819954, 25 | 2 |
| ENSG00000156 | PHKG2 | chr16 | 30756181 | 30756272 | 30753393 | 30753557 | 30756589 | 30756715 | 54,49,96 | 0,0,0 | 58,127,80 | 4,9,1 | 241 | 150 | 4.21E-05 | 0.00075327 | 1.0,1.0,1.0 | 0.9,0.898,0.98 | 0.074 | ENSG00000156 | [[[[30748425, 30 | 2 |
| ENSG00000156 | PHKG2 | chr16 | 30756366 | 30756520 | 30753393 | 30753557 | 30756589 | 30756715 | 28,58,50 | 0,0,0 | 55,85,66 | 4,9,1 | 300 | 150 | 1.01E-05 | 0.000215429 | 1.0,1.0,1.0 | 0.873,0.825,0.97 | 0.11 | ENSG00000156 | [[[[30748425, 30 | 2 |
| ENSG00000089 | FUS | chr16 | 31183860 | 31184002 | 31182512 | 31182664 | 31186801 | 31186836 | 175,198,328 | 0,0,0 | 191,431,247 | 2,12,8 | 292 | 150 | 1.49E-05 | 0.000302473 | 1.0,1.0,1.0 | 0.98,0.949,0.941 | 0.043 | ENSG00000089 | [[[[31180139, 31 | 3 |
| ENSG00000089 | FUS | chr16 | 31184938 | 31185179 | 31182512 | 31182664 | 31188324 | 31188357 | 3,10,12 | 0,0,0 | 21,50,29 | 137,241,147 | 300 | 150 | 0 | 0 | 1.0,1.0,1.0 | 0.071,0.094,0.09 | 0.915 | ENSG00000089 | [[[[31180139, 31 | 4 |
| ENSG00000089 | FUS | chr16 | 31186801 | 31186836 | 31182512 | 31182664 | 31188324 | 31188357 | 199,215,295 | 0,0,0 | 230,502,257 | 137,241,147 | 185 | 150 | 0 | 0 | 1.0,1.0,1.0 | 0.576,0.628,0.58 | 0.403 | ENSG00000089 | [[[[31180139, 31 | 4 |
| ENSG00000185 | ZNF267 | chr16 | 31890206 | 31890250 | 31873806 | 31873969 | 31914475 | 31914558 | 2,1,1 | 0,0,0 | 0,2,1 | 1,3,1 | 194 | 150 | 3.29E-06 | 7.96E-05 | 1.0,1.0,1.0 | 0.0,0.34,0.436 | 0.741 | ENSG00000185 | [[[[31873807, 31 | 2 |
| ENSG00000100 | CINP | chr14 | 102349918 | 102350048 | 102341101 | 102344389 | 102355767 | 102355897 | 100,59,81 | 0,0,0 | 50,90,47 | 3,18,5 | 280 | 150 | 8.27E-11 | 5.50E-09 | 1.0,1.0,1.0 | 0.899,0.728,0.83 | 0.18 | ENSG00000100 | [[[[102362845, 1 | 2 |
| ENSG00000135 | DYNC1LI2 | chr16 | 66725827 | 66725944 | 66720892 | 66723822 | 66728200 | 66728219 | 59,36,95 | 0,0,0 | 49,90,53 | 1,6,3 | 267 | 150 | 0.000279505 | 0.003882908 | 1.0,1.0,1.0 | 0.965,0.894,0.90 | 0.078 | ENSG00000135 | [[[[66751485, 66 | 2 |
| ENSG00000135 | DYNC1LI2 | chr16 | 66729039 | 66729099 | 66725827 | 66725944 | 66730111 | 66730219 | 21,14,38 | 0,0,0 | 16,32,18 | 0,7,3 | 210 | 150 | 1.45E-05 | 0.000295958 | 1.0,1.0,1.0 | 1.0,0.766,0.811 | 0.141 | ENSG00000135 | [[[[66751485, 66 | 3 |
| ENSG00000159 | NAE1 | chr16 | 66805911 | 66806026 | 66805776 | 66805826 | 66810373 | 66810413 | 133,119,117 | 0,0,0 | 82,170,86 | 8,18,21 | 265 | 150 | 2.00E-15 | 2.61E-13 | 1.0,1.0,1.0 | 0.853,0.842,0.69 | 0.202 | ENSG00000159 | [[[[66830847, 66 | 3 |
| ENSG00000159 | NAE1 | chr16 | 66808988 | 66809075 | 66805776 | 66805826 | 66810373 | 66810413 | 84,63,77 | 0,0,0 | 80,136,87 | 8,18,21 | 237 | 150 | 3.25E-13 | 3.14E-11 | 1.0,1.0,1.0 | 0.864,0.827,0.72 | 0.195 | ENSG00000159 | [[[[66830847, 66 | 3 |
| ENSG00000159 | NAE1 | chr16 | 66824854 | 66824885 | 66823341 | 66823600 | 66830846 | 66830917 | 17,29,37 | 0,0,0 | 21,29,28 | 1,6,2 | 181 | 150 | 0.000158425 | 0.00237632 | 1.0,1.0,1.0 | 0.946,0.8,0.921 | 0.111 | ENSG00000159 | [[[[66830847, 66 | 3 |
| ENSG00000172 | CES2 | chr16 | 66938155 | 66938241 | 66934455 | 66934913 | 66939216 | 66939358 | 8,8,14 | 0,0,0 | 7,14,20 | 1,8,2 | 236 | 150 | 1.22E-06 | 3.27E-05 | 1.0,1.0,1.0 | 0.816,0.527,0.86 | 0.264 | ENSG00000172 | [[[[66935516, 66 | 2 |
| ENSG00000103 | SLC7A6 | chr16 | 68291208 | 68291332 | 68290395 | 68290540 | 68294704 | 68294801 | 10,5,14 | 0,0,0 | 6,48,23 | 3,6,4 | 274 | 150 | 1.44E-06 | 3.82E-05 | 1.0,1.0,1.0 | 0.523,0.814,0.75 | 0.301 | ENSG00000103 | [[[[68264526, 68 | 2 |
| ENSG00000103 | SLC7A6 | chr16 | 68291557 | 68291661 | 68290395 | 68290540 | 68294704 | 68294801 | 3,7,22 | 0,0,0 | 13,33,19 | 3,6,4 | 254 | 150 | 1.20E-06 | 3.22E-05 | 1.0,1.0,1.0 | 0.719,0.765,0.73 | 0.26 | ENSG00000103 | [[[[68264526, 68 | 2 |
| ENSG00000242 | EIF6 | chr20 | 35280653 | 35280829 | 35279941 | 35280118 | 35284380 | 35284492 | 250,333,365 | 3,1,4 | 333,710,356 | 8,21,9 | 300 | 150 | 0.000209655 | 0.003036544 | 0.977,0.994,0.97 | 0.954,0.944,0.95 | 0.033 | ENSG00000242 | [[[[35284726, 35 | 2 |
| ENSG00000166 | R3HCC1L | chr10 | 98162882 | 98162975 | 98134670 | 98134706 | 98163292 | 98163397 | 11,8,7 | 1,0,1 | 3,13,1 | 1,5,1 | 243 | 150 | 0.000692015 | 0.008485701 | 0.872,1.0,0.812 | 0.649,0.616,0.38 | 0.346 | ENSG00000166 | [[[[98134657, 98 | 2 |
| ENSG00000119 | ATL2 | chr2 | 38309378 | 38309506 | 38300271 | 38300328 | 38313150 | 38313243 | 48,40,56 | 0,0,0 | 46,85,68 | 15,28,18 | 278 | 150 | 0 | 0 | 1.0,1.0,1.0 | 0.623,0.621,0.67 | 0.362 | ENSG00000119 | [[[[38377143, 38 | 2 |
| ENSG00000119 | ATL2 | chr2 | 38318534 | 38318639 | 38314607 | 38314654 | 38318884 | 38319019 | 30,26,65 | 0,0,0 | 25,89,64 | 2,4,4 | 255 | 150 | 0.000384551 | 0.005115573 | 1.0,1.0,1.0 | 0.88,0.929,0.904 | 0.096 | ENSG00000119 | [[[[38377143, 38 | 2 |
| ENSG00000119 | ATL2 | chr2 | 38318884 | 38319019 | 38318325 | 38318639 | 38377142 | 38377273 | 37,32,75 | 0,0,0 | 43,112,95 | 0,7,7 | 285 | 150 | 3.49E-05 | 0.000639312 | 1.0,1.0,1.0 | 1.0,0.894,0.877 | 0.076 | ENSG00000119 | [[[[38377143, 38 | 2 |
| ENSG00000119 | ATL2 | chr2 | 38343267 | 38343512 | 38318534 | 38318639 | 38377142 | 38377256 | 6,14,39 | 0,0,0 | 9,16,6 | 0,7,7 | 300 | 150 | 5.10E-07 | 1.51E-05 | 1.0,1.0,1.0 | 1.0,0.533,0.3 | 0.389 | ENSG00000119 | [[[[38377143, 38 | 2 |
| ENSG00000182 | IST1 | chr16 | 71920738 | 71920822 | 71916461 | 71916642 | 71921342 | 71921400 | 96,68,151 | 0,0,0 | 163,355,237 | 0,19,4 | 234 | 150 | 0.000109176 | 0.001721641 | 1.0,1.0,1.0 | 1.0,0.923,0.974 | 0.034 | ENSG00000182 | [[[[71895531, 71 | 2 |
| ENSG00000168 | RFWD3 | chr16 | 74636345 | 74636577 | 74632522 | 74632673 | 74644361 | 74644453 | 26,25,70 | 0,1,3 | 32,56,48 | 6,6,11 | 300 | 150 | 6.13E-05 | 0.001044304 | 1.0,0.926,0.921 | 0.727,0.824,0.68 | 0.203 | ENSG00000168 | [[[[74666786, 74 | 2 |
| ENSG00000168 | RFWD3 | chr16 | 74637855 | 74637970 | 74632522 | 74632673 | 74644361 | 74644453 | 23,21,50 | 0,1,3 | 22,57,28 | 6,6,11 | 265 | 150 | 0.000109148 | 0.001721554 | 1.0,0.922,0.904 | 0.675,0.843,0.59 | 0.239 | ENSG00000168 | [[[[74666786, 74 | 2 |
| ENSG00000166 | IDH3A | chr15 | 78164991 | 78165076 | 78162208 | 78162367 | 78166149 | 78166220 | 43,69,88 | 0,0,0 | 45,94,98 | 3,4,6 | 235 | 150 | 5.15E-05 | 0.000895823 | 1.0,1.0,1.0 | 0.905,0.938,0.90 | 0.085 | ENSG00000166 | [[[[78149362, 78 | 3 |
| ENSG00000072 | DERL2 | chr17 | 5480053 | 5480144 | 5471250 | 5474789 | 5481100 | 5481142 | 59,65,75 | 0,0,0 | 94,214,103 | 5,10,6 | 241 | 150 | 7.64E-06 | 0.00016885 | 1.0,1.0,1.0 | 0.921,0.93,0.914 | 0.078 | ENSG00000072 | [[[[5486069, 548 | 2 |
| ENSG00000122 | HNRNPA2B1 | chr7 | 26190992 | 26191128 | 26190124 | 26190460 | 26195846 | 26195909 | 244,184,240 | 0,0,0 | 209,544,289 | 12,32,10 | 286 | 150 | 2.30E-12 | 1.99E-10 | 1.0,1.0,1.0 | 0.901,0.899,0.93 | 0.087 | ENSG00000122 | [[[[26200572, 26 | 3 |
| ENSG00000122 | HNRNPA2B1 | chr7 | 26192494 | 26192577 | 26189926 | 26192338 | 26195846 | 26195855 | 136,911,011,634 | 0,0,0 | 117,520,231,140 | 74,124,69 | 233 | 150 | 0 | 0 | 1.0,1.0,1.0 | 0.911,0.913,0.91 | 0.087 | ENSG00000122 | [[[[26200572, 26 | 3 |
| ENSG00000122 | HNRNPA2B1 | chr7 | 26192494 | 26192577 | 26190124 | 26190460 | 26195846 | 26195855 | 90,82,122 | 0,0,0 | 77,165,64 | 12,32,10 | 233 | 150 | 0 | 0 | 1.0,1.0,1.0 | 0.805,0.768,0.80 | 0.207 | ENSG00000122 | [[[[26200572, 26 | 3 |
| ENSG00000122 | HNRNPA2B1 | chr7 | 26192494 | 26192577 | 26190124 | 26190460 | 26197314 | 26197461 | 90,82,122 | 0,0,0 | 70,157,56 | 2,9,1 | 233 | 150 | 0.000319015 | 0.004355486 | 1.0,1.0,1.0 | 0.958,0.918,0.97 | 0.05 | ENSG00000122 | [[[[26200572, 26 | 7 |
| ENSG00000122 | HNRNPA2B1 | chr7 | 26192494 | 26192577 | 26190992 | 26191128 | 26195846 | 26195855 | 106,85,102 | 0,0,0 | 119,239,145 | 8,12,3 | 233 | 150 | 3.41E-06 | 8.23E-05 | 1.0,1.0,1.0 | 0.905,0.928,0.96 | 0.066 | ENSG00000122 | [[[[26200572, 26 | 3 |
| ENSG00000122 | HNRNPA2B1 | chr7 | 26193250 | 26193373 | 26190124 | 26190460 | 26195846 | 26195855 | 15,15,25 | 0,0,0 | 11,25,21 | 12,32,10 | 273 | 150 | 2.22E-15 | 2.87E-13 | 1.0,1.0,1.0 | 0.335,0.3,0.536 | 0.61 | ENSG00000122 | [[[[26200572, 26 | 3 |
| ENSG00000275 | SYNRG | chr17 | 37535978 | 37536127 | 37520537 | 37520648 | 37539191 | 37539245 | 24,31,21 | 0,0,0 | 28,67,50 | 11,10,9 | 299 | 150 | 2.02E-09 | 1.06E-07 | 1.0,1.0,1.0 | 0.561,0.771,0.73 | 0.311 | ENSG00000275 | [[[[37609279, 37 | 2 |
| ENSG00000275 | SYNRG | chr17 | 37538323 | 37538420 | 37520537 | 37520648 | 37539191 | 37539245 | 4,10,9 | 0,0,0 | 17,19,14 | 11,10,9 | 247 | 150 | 1.11E-09 | 6.12E-08 | 1.0,1.0,1.0 | 0.484,0.536,0.48 | 0.498 | ENSG00000275 | [[[[37609279, 37 | 2 |
| ENSG00000275 | SYNRG | chr17 | 37568790 | 37568924 | 37561470 | 37561589 | 37576340 | 37576418 | 10,6,19 | 0,0,0 | 15,33,21 | 0,4,5 | 284 | 150 | 2.83E-05 | 0.000530625 | 1.0,1.0,1.0 | 1.0,0.813,0.689 | 0.166 | ENSG00000275 | [[[[37609279, 37 | 3 |
| ENSG00000197 | SPG7 | chr16 | 89532462 | 89532636 | 89530682 | 89530808 | 89536764 | 89536948 | 9,19,17 | 0,0,0 | 3,13,12 | 18,15,10 | 300 | 150 | 2.22E-16 | 3.17E-14 | 1.0,1.0,1.0 | 0.077,0.302,0.37 | 0.749 | ENSG00000197 | [[[[89508403, 89 | 2 |
| ENSG00000197 | SPG7 | chr16 | 89532462 | 89532636 | 89530682 | 89530808 | 89540943 | 89540977 | 5,1,10 | 0,0,0 | 1,14,7 | 1,5,2 | 300 | 150 | 1.83E-07 | 6.01E-06 | 1.0,1.0,1.0 | 0.333,0.583,0.63 | 0.483 | ENSG00000197 | [[[[89508403, 89 | 2 |
| ENSG00000197 | SPG7 | chr16 | 89532462 | 89532636 | 89530682 | 89530808 | 89544647 | 89544772 | 11,22,26 | 0,0,0 | 8,26,10 | 1,5,2 | 300 | 150 | 9.44E-06 | 0.000203621 | 1.0,1.0,1.0 | 0.8,0.722,0.714 | 0.255 | ENSG00000197 | [[[[89508403, 89 | 2 |
| ENSG00000197 | SPG7 | chr16 | 89540943 | 89542279 | 89530682 | 89530808 | 89544647 | 89544772 | 2,2,2 | 0,0,0 | 6,18,5 | 1,5,2 | 300 | 150 | 7.05E-07 | 2.01E-05 | 1.0,1.0,1.0 | 0.75,0.643,0.556 | 0.35 | ENSG00000197 | [[[[89508403, 89 | 2 |
| ENSG00000141 | TCF25 | chr16 | 89893788 | 89893858 | 89887651 | 89887681 | 89895082 | 89895137 | 2,4,7 | 0,0,0 | 4,10,10 | 1,1,7 | 220 | 150 | 2.42E-06 | 6.05E-05 | 1.0,1.0,1.0 | 0.732,0.872,0.49 | 0.301 | ENSG00000141 | [[[[89873592, 89 | 2 |
| ENSG00000164 | CAMLG | chr5 | 134741062 | 134741523 | 134738500 | 134738792 | 134750758 | 134752160 | 36,27,47 | 7,6,14 | 108,168,93 | 5,4,4 | 300 | 150 | 2.90E-08 | 1.17E-06 | 0.72,0.692,0.627 | 0.915,0.955,0.92 | -0.251 | ENSG00000164 | [[[[134738548, 1 | 2 |

| | | | | | | | | | | | | | | | | | | | | | | |
|---|---|---|---|---|---|---|---|---|---|---|---|---|---|---|---|---|---|---|---|---|---|---|
| ENSG00000164 | CAMLG | chr5 | 134743986 | 134744052 | 134738582 | 134738792 | 134750758 | 134751377 | 33,22,26 | 7,6,14 | 36,70,42 | 5,4,4 | 216 | 150 | 0.000436319 | 0.005716407 | 0.766,0.718,0.56 | 0.833,0.924,0.87 | -0.196 | ENSG00000164 | [[[[134738548, 13 | 2 |
| ENSG00000108 | YWHAE | chr17 | 1361901 | 1362008 | 1354210 | 1354347 | 1364858 | 1365058 | 590,481,923 | 0,0,0 | 518,868,619 | 11,17,10 | 257 | 150 | 8.85E-07 | 2.46E-05 | 1.0,1.0,1.0 | 0.965,0.968,0.97 | 0.031 | ENSG00000108 | [[[[1400047, 140 | 2 |
| ENSG00000108 | YWHAE | chr17 | 1361929 | 1362008 | 1354210 | 1354347 | 1364858 | 1365058 | 450,357,709 | 0,0,0 | 437,693,480 | 11,17,10 | 229 | 150 | 2.54E-07 | 8.06E-06 | 1.0,1.0,1.0 | 0.963,0.964,0.96 | 0.035 | ENSG00000108 | [[[[1400047, 140 | 2 |
| ENSG00000196 | TRAPPC4 | chr11 | 119020149 | 119020253 | 119018765 | 119019317 | 119023320 | 119023394 | 70,53,111 | 0,0,0 | 64,122,49 | 5,8,1 | 254 | 150 | 1.10E-05 | 0.00023272 | 1.0,1.0,1.0 | 0.883,0.9,0.967 | 0.083 | ENSG00000196 | [[[[119018766, 11 | 2 |
| ENSG00000153 | ANAPC1 | chr2 | 111873612 | 111873664 | 111872629 | 111872712 | 111878809 | 111878971 | 7,4,11 | 0,0,1 | 3,6,7 | 0,2,4 | 202 | 150 | 0.000802324 | 0.009626118 | 1.0,1.0,0.891 | 1.0,0.69,0.565 | 0.212 | ENSG00000153 | [[[[111883942, 11 | 2 |
| ENSG00000111 | SLC38A1 | chr12 | 46197719 | 46197817 | 46183062 | 46189071 | 46198624 | 46198743 | 23,22,47 | 0,0,0 | 31,88,60 | 12,30,24 | 248 | 150 | 4.00E-15 | 5.05E-13 | 1.0,1.0,1.0 | 0.61,0.64,0.602 | 0.383 | ENSG00000111 | [[[[46268526, 462 | 2 |
| ENSG00000111 | SLC38A1 | chr12 | 46201097 | 46201198 | 46183062 | 46189071 | 46203009 | 46203089 | 35,28,77 | 0,0,0 | 75,149,98 | 6,7,9 | 251 | 150 | 2.17E-06 | 5.50E-05 | 1.0,1.0,1.0 | 0.882,0.927,0.86 | 0.108 | ENSG00000111 | [[[[46268526, 462 | 4 |
| ENSG00000111 | SLC38A1 | chr12 | 46239678 | 46239893 | 46229563 | 46229639 | 46268525 | 46268995 | 30,26,73 | 0,0,0 | 63,163,156 | 2,4,7 | 300 | 150 | 0.000780268 | 0.009394034 | 1.0,1.0,1.0 | 0.94,0.953,0.918 | 0.063 | ENSG00000111 | [[[[46268526, 462 | 2 |
| ENSG00000111 | SLC38A1 | chr12 | 46239678 | 46239893 | 46229563 | 46229639 | 46268864 | 46269130 | 30,26,69 | 1,0,0 | 63,162,156 | 2,16,9 | 300 | 150 | 8.83E-06 | 0.00019162 | 0.938,1.0,1.0 | 0.94,0.835,0.897 | 0.089 | ENSG00000111 | [[[[46268526, 462 | 2 |
| ENSG00000111 | SLC38A1 | chr12 | 46243199 | 46243314 | 46229563 | 46229639 | 46268525 | 46269012 | 15,3,22 | 0,0,0 | 33,71,78 | 2,4,7 | 265 | 150 | 8.80E-05 | 0.001425254 | 1.0,1.0,1.0 | 0.903,0.909,0.86 | 0.108 | ENSG00000111 | [[[[46268526, 462 | 2 |
| ENSG00000111 | SLC38A1 | chr12 | 46243199 | 46243314 | 46229563 | 46229639 | 46268864 | 46269119 | 10,7,46 | 1,0,0 | 9,52,45 | 2,16,9 | 265 | 150 | 6.71E-08 | 2.46E-06 | 0.85,1.0,1.0 | 0.718,0.648,0.73 | 0.248 | ENSG00000111 | [[[[46268526, 462 | 2 |
| ENSG00000108 | RNF167 | chr17 | 4943425 | 4943519 | 4942850 | 4942941 | 4944557 | 4944638 | 123,169,230 | 1,5,14 | 233,414,212 | 23,42,24 | 244 | 150 | 1.65E-05 | 0.000330896 | 0.987,0.954,0.91 | 0.862,0.858,0.84 | 0.096 | ENSG00000108 | [[[[4940268, 494 | 2 |
| ENSG00000108 | CAMTA2 | chr17 | 4981677 | 4981742 | 4981224 | 4981359 | 4982756 | 4982892 | 6,7,12 | 0,0,1 | 12,42,18 | 3,5,2 | 215 | 150 | 0.000467015 | 0.006070555 | 1.0,1.0,0.893 | 0.736,0.854,0.86 | 0.147 | ENSG00000108 | [[[[4987593, 4987 | 2 |
| ENSG00000108 | CAMTA2 | chr17 | 4981677 | 4981831 | 4981224 | 4981359 | 4982756 | 4982892 | 11,10,38 | 0,0,1 | 14,45,19 | 3,5,2 | 300 | 150 | 4.64E-05 | 0.000817465 | 1.0,1.0,0.95 | 0.7,0.818,0.826 | 0.202 | ENSG00000108 | [[[[4987593, 4987 | 2 |
| ENSG00000108 | CAMTA2 | chr17 | 4982088 | 4982160 | 4981224 | 4981359 | 4982756 | 4982892 | 3,2,5 | 0,0,1 | 3,8,3 | 3,5,2 | 222 | 150 | 1.23E-05 | 0.000255339 | 1.0,1.0,0.772 | 0.403,0.519,0.50 | 0.449 | ENSG00000108 | [[[[4987593, 4987 | 2 |
| ENSG00000144 | CNOT9 | chr2 | 218580560 | 218580740 | 218568579 | 218568978 | 218592303 | 218592402 | 21,27,64 | 0,0,0 | 37,68,60 | 3,2,5 | 300 | 150 | 0.000165597 | 0.002469314 | 1.0,1.0,1.0 | 0.86,0.944,0.857 | 0.113 | ENSG00000144 | [[[[218568839, 2 | 4 |
| ENSG00000144 | CNOT9 | chr2 | 218584611 | 218584721 | 218582970 | 218583086 | 218592303 | 218592402 | 72,48,118 | 0,0,0 | 53,124,86 | 2,12,14 | 260 | 150 | 9.33E-10 | 5.21E-08 | 1.0,1.0,1.0 | 0.939,0.856,0.78 | 0.142 | ENSG00000144 | [[[[218568839, 2 | 2 |
| ENSG00000144 | CNOT9 | chr2 | 218587585 | 218587695 | 218568579 | 218568978 | 218592303 | 218592402 | 66,72,76 | 0,0,0 | 54,94,69 | 3,2,5 | 260 | 150 | 0.000567399 | 0.007163993 | 1.0,1.0,1.0 | 0.912,0.964,0.88 | 0.079 | ENSG00000144 | [[[[218568839, 2 | 4 |
| ENSG00000144 | CNOT9 | chr2 | 218587585 | 218587695 | 218582970 | 218583086 | 218592303 | 218592402 | 66,72,76 | 0,0,0 | 57,98,69 | 2,12,14 | 260 | 150 | 7.12E-10 | 4.05E-08 | 1.0,1.0,1.0 | 0.943,0.825,0.74 | 0.164 | ENSG00000144 | [[[[218568839, 2 | 2 |
| ENSG00000004 | DVL2 | chr17 | 7229377 | 7229447 | 7228968 | 7229045 | 7229587 | 7229678 | 6,24,57 | 0,0,0 | 24,40,37 | 9,11,13 | 220 | 150 | 2.64E-12 | 2.26E-10 | 1.0,1.0,1.0 | 0.645,0.713,0.66 | 0.327 | ENSG00000004 | [[[[7234069, 7234 | 2 |
| ENSG00000129 | MPDU1 | chr17 | 7585731 | 7585797 | 7583646 | 7583965 | 7586655 | 7586777 | 96,138,160 | 0,0,2 | 118,200,124 | 14,21,10 | 216 | 150 | 5.06E-12 | 4.13E-10 | 1.0,1.0,0.982 | 0.854,0.869,0.89 | 0.121 | ENSG00000129 | [[[[7583852, 758 | 2 |
| ENSG00000129 | MPDU1 | chr17 | 7585731 | 7585797 | 7583646 | 7583965 | 7586691 | 7586777 | 96,139,162 | 0,1,3 | 123,220,137 | 19,38,24 | 216 | 150 | 0 | 0 | 1.0,0.99,0.974 | 0.818,0.801,0.79 | 0.182 | ENSG00000129 | [[[[7583852, 758 | 2 |
| ENSG00000129 | MPDU1 | chr17 | 7585945 | 7586078 | 7583646 | 7583965 | 7586655 | 7586777 | 15,9,8 | 0,0,2 | 20,48,15 | 14,21,10 | 283 | 150 | 2.09E-06 | 5.32E-05 | 1.0,1.0,0.68 | 0.431,0.548,0.44 | 0.419 | ENSG00000129 | [[[[7583852, 758 | 2 |
| ENSG00000129 | MPDU1 | chr17 | 7585945 | 7586078 | 7583646 | 7583965 | 7586691 | 7586746 | 218,206,332 | 0,1,3 | 220,449,229 | 19,38,24 | 283 | 150 | 0 | 0 | 1.0,0.991,0.983 | 0.86,0.862,0.835 | 0.139 | ENSG00000129 | [[[[7583852, 758 | 2 |
| ENSG00000129 | MPDU1 | chr17 | 7586655 | 7586777 | 7583665 | 7583965 | 7586898 | 7587017 | 82,93,127 | 0,0,0 | 105,184,131 | 7,9,0 | 272 | 150 | 4.53E-05 | 0.000801519 | 1.0,1.0,1.0 | 0.892,0.919,1.0 | 0.063 | ENSG00000129 | [[[[7583852, 758 | 3 |
| ENSG00000129 | MPDU1 | chr17 | 7586691 | 7586777 | 7583894 | 7583965 | 7586898 | 7586961 | 79,89,120 | 0,0,0 | 105,186,135 | 7,9,0 | 236 | 150 | 0.000113467 | 0.001778639 | 1.0,1.0,1.0 | 0.905,0.929,1.0 | 0.055 | ENSG00000129 | [[[[7583852, 758 | 3 |
| ENSG00000133 | MEN1 | chr11 | 64806231 | 64806368 | 64805634 | 64805770 | 64807178 | 64807219 | 27,49,53 | 0,0,0 | 38,62,45 | 13,10,1 | 287 | 150 | 2.64E-08 | 1.07E-06 | 1.0,1.0,1.0 | 0.604,0.764,0.95 | 0.224 | ENSG00000133 | [[[[64810514, 64 | 2 |
| ENSG00000178 | AURKB | chr17 | 8206749 | 8206888 | 8206490 | 8206639 | 8210529 | 8210568 | 144,134,177 | 0,0,0 | 110,288,130 | 5,10,6 | 289 | 150 | 1.17E-06 | 3.15E-05 | 1.0,1.0,1.0 | 0.919,0.937,0.91 | 0.075 | ENSG00000178 | [[[[8210530, 821 | 5 |
| ENSG00000178 | AURKB | chr17 | 8206749 | 8206934 | 8206490 | 8206639 | 8210529 | 8210600 | 148,139,179 | 0,0,0 | 119,303,142 | 5,10,6 | 300 | 150 | 1.62E-06 | 4.24E-05 | 1.0,1.0,1.0 | 0.922,0.938,0.92 | 0.073 | ENSG00000178 | [[[[8210530, 821 | 5 |
| ENSG00000178 | AURKB | chr17 | 8207737 | 8207840 | 8207570 | 8207625 | 8210529 | 8210581 | 133,177,229 | 0,0,3 | 171,320,211 | 1,18,1 | 253 | 150 | 0.000499457 | 0.006422537 | 1.0,1.0,0.978 | 0.99,0.913,0.992 | 0.028 | ENSG00000178 | [[[[8210530, 821 | 2 |
| ENSG00000178 | AURKB | chr17 | 8207737 | 8207901 | 8207570 | 8207625 | 8210529 | 8210568 | 143,175,215 | 0,0,3 | 149,290,179 | 1,18,1 | 300 | 150 | 0.000143047 | 0.002169561 | 1.0,1.0,0.973 | 0.987,0.89,0.989 | 0.036 | ENSG00000178 | [[[[8210530, 821 | 2 |
| ENSG00000178 | AURKB | chr17 | 8209953 | 8210055 | 8206749 | 8206888 | 8210176 | 8210254 | 1,2,5 | 0,0,0 | 2,3,3 | 0,4,2 | 252 | 150 | 6.97E-05 | 0.001164316 | 1.0,1.0,1.0 | 1.0,0.309,0.472 | 0.406 | ENSG00000178 | [[[[8210530, 821 | 3 |
| ENSG00000198 | DDX42 | chr17 | 63809559 | 63809659 | 63805043 | 63805175 | 63810512 | 63810560 | 85,35,87 | 0,0,0 | 43,68,54 | 4,9,3 | 250 | 150 | 1.28E-07 | 4.35E-06 | 1.0,1.0,1.0 | 0.866,0.819,0.91 | 0.133 | ENSG00000198 | [[[[63774170, 63 | 4 |
| ENSG00000141 | NCOR1 | chr17 | 16039432 | 16039654 | 16034764 | 16034944 | 16046950 | 16047035 | 74,43,83 | 0,0,0 | 42,86,52 | 13,10,2 | 300 | 150 | 2.33E-09 | 1.20E-07 | 1.0,1.0,1.0 | 0.618,0.811,0.92 | 0.214 | ENSG00000141 | [[[[16215362, 16 | 2 |
| ENSG00000141 | COPS3 | chr17 | 17261599 | 17261692 | 17260373 | 17260474 | 17270757 | 17270807 | 1,3,5 | 0,0,0 | 5,16,13 | 2,7,2 | 243 | 150 | 2.34E-07 | 7.48E-06 | 1.0,1.0,1.0 | 0.607,0.585,0.8 | 0.336 | ENSG00000141 | [[[[17281132, 172 | 3 |
| ENSG00000141 | COPS3 | chr17 | 17261965 | 17262106 | 17260300 | 17260474 | 17267884 | 17267977 | 180,161,220 | 1,0,1 | 133,280,154 | 3,15,13 | 291 | 150 | 7.71E-09 | 3.52E-07 | 0.989,1.0,0.991 | 0.958,0.906,0.85 | 0.086 | ENSG00000141 | [[[[17281132, 172 | 2 |
| ENSG00000141 | COPS3 | chr17 | 17264801 | 17264981 | 17260300 | 17260474 | 17267884 | 17267977 | 174,183,222 | 1,0,1 | 135,301,178 | 3,15,13 | 300 | 150 | 1.26E-08 | 5.49E-07 | 0.989,1.0,0.991 | 0.957,0.909,0.87 | 0.08 | ENSG00000141 | [[[[17281132, 172 | 2 |
| ENSG00000141 | COPS3 | chr17 | 17267884 | 17268045 | 17260300 | 17260474 | 17270757 | 17270807 | 11,4,11 | 0,0,0 | 18,36,25 | 2,7,2 | 300 | 150 | 5.44E-06 | 0.000124948 | 1.0,1.0,1.0 | 0.818,0.72,0.862 | 0.2 | ENSG00000141 | [[[[17281132, 172 | 3 |
| ENSG00000119 | PHF19 | chr9 | 120865709 | 120865830 | 120862587 | 120862749 | 120866027 | 120866096 | 19,35,61 | 0,0,0 | 17,51,25 | 5,16,8 | 271 | 150 | 2.26E-13 | 2.25E-11 | 1.0,1.0,1.0 | 0.653,0.638,0.63 | 0.358 | ENSG00000119 | [[[[120877091, 12 | 2 |
| ENSG00000108 | BLMH | chr17 | 30272560 | 30272628 | 30271268 | 30271388 | 30274041 | 30274197 | 48,49,61 | 0,0,0 | 15,47,13 | 10,17,8 | 218 | 150 | 0 | 0 | 1.0,1.0,1.0 | 0.508,0.655,0.52 | 0.436 | ENSG00000108 | [[[[30291807, 302 | 2 |
| ENSG00000108 | BLMH | chr17 | 30287805 | 30287947 | 30286813 | 30286902 | 30291310 | 30291508 | 82,77,93 | 0,0,0 | 24,63,33 | 1,6,5 | 292 | 150 | 3.73E-07 | 1.14E-05 | 1.0,1.0,1.0 | 0.925,0.844,0.77 | 0.153 | ENSG00000108 | [[[[30291807, 302 | 2 |
| ENSG00000108 | BLMH | chr17 | 30289372 | 30289482 | 30286813 | 30286902 | 30291310 | 30291508 | 16,33,79 | 0,0,0 | 22,27,30 | 1,6,5 | 260 | 150 | 3.54E-07 | 1.09E-05 | 1.0,1.0,1.0 | 0.927,0.722,0.77 | 0.192 | ENSG00000108 | [[[[30291807, 302 | 2 |
| ENSG00000108 | CPD | chr17 | 30462369 | 30462469 | 30456461 | 30456526 | 30464587 | 30465017 | 36,17,30 | 0,0,0 | 32,57,38 | 6,17,4 | 250 | 150 | 3.27E-09 | 1.63E-07 | 1.0,1.0,1.0 | 0.762,0.668,0.85 | 0.24 | ENSG00000108 | [[[[30378927, 303 | 3 |
| ENSG00000108 | GOSR1 | chr17 | 30490125 | 30490217 | 30481142 | 30481257 | 30492678 | 30492753 | 17,11,26 | 0,0,0 | 38,76,57 | 0,6,6 | 242 | 150 | 0.000159411 | 0.00239016 | 1.0,1.0,1.0 | 1.0,0.887,0.855 | 0.086 | ENSG00000108 | [[[[30477408, 304 | 3 |
| ENSG00000176 | CRLF3 | chr17 | 30796159 | 30796337 | 30792439 | 30792572 | 30797310 | 30797398 | 47,44,45 | 0,0,0 | 46,90,49 | 0,7,5 | 300 | 150 | 3.08E-05 | 0.000573652 | 1.0,1.0,1.0 | 1.0,0.865,0.831 | 0.101 | ENSG00000176 | [[[[30824523, 308 | 2 |
| ENSG00000104 | EIF3E | chr8 | 108214606 | 108214716 | 108203400 | 108203503 | 108228266 | 108228391 | 202,240,246 | 4,4,8 | 243,486,275 | 33,68,35 | 260 | 150 | 1.91E-14 | 2.20E-12 | 0.967,0.972,0.94 | 0.809,0.805,0.81 | 0.151 | ENSG00000104 | [[[[108248613, 10 | 3 |
| ENSG00000104 | EIF3E | chr8 | 108217333 | 108217460 | 108203400 | 108203503 | 108228266 | 108228391 | 214,206,311 | 4,4,8 | 229,509,366 | 33,68,35 | 277 | 150 | 5.57E-14 | 5.96E-12 | 0.967,0.965,0.95 | 0.79,0.802,0.85 | 0.148 | ENSG00000104 | [[[[108248613, 10 | 3 |
| ENSG00000104 | EIF3E | chr8 | 108236160 | 108236203 | 108217333 | 108217460 | 108239957 | 108240075 | 53,59,76 | 0,0,0 | 63,159,85 | 4,9,1 | 193 | 150 | 0.00042839 | 0.005620261 | 1.0,1.0,1.0 | 0.924,0.932,0.98 | 0.053 | ENSG00000104 | [[[[108248613, 10 | 4 |
| ENSG00000104 | EIF3E | chr8 | 108241798 | 108241913 | 108229069 | 108229195 | 108248612 | 108248668 | 162,165,187 | 4,2,2 | 153,323,218 | 36,77,28 | 265 | 150 | 0 | 0 | 0.958,0.979,0.98 | 0.706,0.704,0.81 | 0.231 | ENSG00000104 | [[[[108248613, 10 | 4 |
| ENSG00000099 | HNRNPM | chr19 | 8471325 | 8471427 | 8468773 | 8468834 | 8483157 | 8483211 | 288,228,348 | 0,1,0 | 403,646,391 | 11,12,11 | 252 | 150 | 5.85E-06 | 0.000133336 | 1.0,0.993,1.0 | 0.956,0.97,0.955 | 0.037 | ENSG00000099 | [[[[8444975, 844 | 3 |
| ENSG00000099 | HNRNPM | chr19 | 8473663 | 8473708 | 8471325 | 8471427 | 8483157 | 8483211 | 174,191,187 | 5,2,3 | 181,336,182 | 36,65,33 | 195 | 150 | 0 | 0 | 0.964,0.987,0.98 | 0.795,0.799,0.80 | 0.176 | ENSG00000099 | [[[[8444975, 844 | 2 |
| ENSG00000099 | HNRNPM | chr19 | 8474166 | 8474244 | 8468773 | 8468834 | 8483157 | 8483211 | 94,116,208 | 0,1,0 | 175,250,174 | 11,12,11 | 228 | 150 | 2.05E-08 | 8.53E-07 | 1.0,0.987,1.0 | 0.913,0.932,0.91 | 0.077 | ENSG00000099 | [[[[8444975, 844 | 3 |
| ENSG00000099 | HNRNPM | chr19 | 8474166 | 8474244 | 8471325 | 8471427 | 8483157 | 8483211 | 104,125,218 | 5,2,3 | 189,264,192 | 36,65,33 | 228 | 150 | 0 | 0 | 0.932,0.976,0.98 | 0.775,0.728,0.79 | 0.197 | ENSG00000099 | [[[[8444975, 844 | 2 |
| ENSG00000099 | HNRNPM | chr19 | 8474166 | 8474244 | 8471325 | 8471427 | 8485602 | 8486273 | 11,29,56 | 0,0,1 | 28,32,29 | 2,11,15 | 228 | 150 | 8.46E-09 | 3.82E-07 | 1.0,1.0,0.974 | 0.902,0.657,0.56 | 0.285 | ENSG00000099 | [[[[8444975, 844 | 3 |
| ENSG00000099 | HNRNPM | chr19 | 8483157 | 8483211 | 8471325 | 8471427 | 8485602 | 8485648 | 12,20,54 | 0,0,1 | 57,91,75 | 2,11,15 | 204 | 150 | 3.15E-06 | 7.68E-05 | 1.0,1.0,0.975 | 0.954,0.859,0.78 | 0.125 | ENSG00000099 | [[[[8444975, 844 | 3 |
| ENSG00000108 | PSMD3 | chr17 | 39995175 | 39995295 | 39990093 | 39990197 | 39996182 | 39996338 | 0,0,0 | 2,4,1 | 11,7,5 | 17,23,23 | 270 | 150 | 2.08E-05 | 0.000407247 | 0.0,0.0,0.0 | 0.264,0.145,0.10 | -0.172 | ENSG00000108 | [[[[39980807, 39 | 3 |
| ENSG00000108 | PSMD3 | chr17 | 39995423 | 39995527 | 39990093 | 39990197 | 39996182 | 39996338 | 422,467,677 | 2,4,1 | 4,781,002,581 | 17,23,23 | 254 | 150 | 1.20E-09 | 6.55E-08 | 0.992,0.986,0.99 | 0.943,0.963,0.93 | 0.044 | ENSG00000108 | [[[[39980807, 39 | 3 |
| ENSG00000125 | GGA3 | chr17 | 75242830 | 75242911 | 75241614 | 75242473 | 75243446 | 75243570 | 21,26,41 | 0,0,0 | 28,50,45 | 1,6,2 | 231 | 150 | 0.000504359 | 0.006476438 | 1.0,1.0,1.0 | 0.948,0.844,0.93 | 0.091 | ENSG00000125 | [[[[75261548, 75 | 2 |
| ENSG00000151 | SLC7A11 | chr4 | 138179216 | 138179394 | 138164096 | 138172017 | 138214584 | 138214629 | 36,21,70 | 0,0,0 | 23,73,60 | 7,3,13 | 300 | 150 | 1.51E-08 | 6.49E-07 | 1.0,1.0,1.0 | 0.622,0.924,0.69 | 0.252 | ENSG00000151 | [[[[138241793, 1 | 5 |
| ENSG00000151 | SLC7A11 | chr4 | 138180640 | 138180790 | 138164096 | 138172017 | 138182296 | 138182393 | 58,20,75 | 0,0,0 | 17,76,56 | 7,34,24 | 300 | 150 | 0 | 0 | 1.0,1.0,1.0 | 0.548,0.528,0.53 | 0.462 | ENSG00000151 | [[[[138241793, 1 | 2 |
| ENSG00000151 | SLC7A11 | chr4 | 138182296 | 138182393 | 138164096 | 138172017 | 138183201 | 138183305 | 22,15,69 | 0,0,0 | 37,127,119 | 2,7,9 | 247 | 150 | 1.82E-05 | 0.000362014 | 1.0,1.0,1.0 | 0.918,0.917,0.88 | 0.092 | ENSG00000151 | [[[[138241793, 1 | 3 |
| ENSG00000141 | FKBP10 | chr17 | 41819209 | 41819399 | 41818381 | 41818527 | 41820268 | 41820461 | 97,122,207 | 0,0,2 | 182,285,241 | 17,19,18 | 300 | 150 | 2.70E-11 | 1.92E-09 | 1.0,1.0,0.981 | 0.843,0.882,0.87 | 0.129 | ENSG00000141 | [[[[41813004, 41 | 2 |
| ENSG00000068 | IFI35 | chr17 | 43013266 | 43013373 | 43012178 | 43012277 | 43013475 | 43013648 | 11,12,17 | 0,0,0 | 18,29,19 | 3,12,14 | 257 | 150 | 2.89E-09 | 1.46E-07 | 1.0,1.0,1.0 | 0.778,0.585,0.44 | 0.398 | ENSG00000068 | [[[[43006784, 43 | 2 |
| ENSG00000068 | IFI35 | chr17 | 43013266 | 43013379 | 43012178 | 43012277 | 43013475 | 43013662 | 5,17,28 | 0,0,0 | 23,38,30 | 3,12,14 | 263 | 150 | 4.14E-09 | 2.01E-07 | 1.0,1.0,1.0 | 0.814,0.644,0.55 | 0.331 | ENSG00000068 | [[[[43006784, 43 | 2 |
| ENSG00000120 | EXOSC8 | chr13 | 37005919 | 37006025 | 37004217 | 37004561 | 37009183 | 37009401 | 119,70,124 | 0,0,0 | 109,256,120 | 9,4,6 | 256 | 150 | 1.42E-05 | 0.000291347 | 1.0,1.0,1.0 | 0.876,0.974,0.92 | 0.076 | ENSG00000120 | [[[[37000786, 37 | 5 |
| ENSG00000120 | EXOSC8 | chr13 | 37005956 | 37006025 | 37002933 | 37004561 | 37009183 | 37009401 | 1,2,3 | 0,0,0 | 28,35,11 | 9,4,6 | 219 | 150 | 3.65E-06 | 8.76E-05 | 1.0,1.0,1.0 | 0.681,0.857,0.55 | 0.302 | ENSG00000120 | [[[[37000786, 37 | 5 |
| ENSG00000120 | EXOSC8 | chr13 | 37006114 | 37006160 | 37005919 | 37006025 | 37009183 | 37009401 | 31,28,38 | 0,0,0 | 48,78,41 | 0,13,0 | 196 | 150 | 0.000177967 | 0.002631464 | 1.0,1.0,1.0 | 1.0,0.821,1.0 | 0.06 | ENSG00000120 | [[[[37000786, 37 | 4 |
| ENSG00000115 | MPV17 | chr2 | 27311898 | 27311951 | 27309491 | 27309981 | 27312493 | 27312589 | 39,33,56 | 0,0,0 | 16,30,21 | 6,4,0 | 203 | 150 | 9.88E-06 | 0.000211576 | 1.0,1.0,1.0 | 0.663,0.847,1.0 | 0.163 | ENSG00000115 | [[[[27323052, 27 | 2 |
| ENSG00000115 | MPV17 | chr2 | 27312679 | 27312772 | 27312213 | 27312246 | 27312993 | 27313074 | 41,55,124 | 0,0,0 | 32,77,65 | 24,28,20 | 243 | 150 | 0 | 0 | 1.0,1.0,1.0 | 0.451,0.629,0.66 | 0.418 | ENSG00000115 | [[[[27323052, 27 | 2 |
| ENSG00000115 | MPV17 | chr2 | 27312679 | 27312772 | 27312213 | 27312246 | 27322447 | 27322522 | 4,3,12 | 0,0,0 | 0,12,6 | 15,7,5 | 243 | 150 | 2.13E-11 | 1.54E-09 | 1.0,1.0,1.0 | 0.0,0.514,0.426 | 0.687 | ENSG00000115 | [[[[27323052, 27 | 3 |
| ENSG00000115 | MPV17 | chr2 | 27312993 | 27313074 | 27309529 | 27309981 | 27322447 | 27322522 | 3,3,4 | 0,0,0 | 4,11,14 | 3,2,3 | 231 | 150 | 2.35E-06 | 5.90E-05 | 1.0,1.0,1.0 | 0.464,0.781,0.75 | 0.334 | ENSG00000115 | [[[[27323052, 27 | 5 |
| ENSG00000115 | MPV17 | chr2 | 27312993 | 27313109 | 27312213 | 27312246 | 27322447 | 27322522 | 36,76,144 | 0,0,0 | 85,93,86 | 15,7,5 | 266 | 150 | 7.17E-10 | 4.08E-08 | 1.0,1.0,1.0 | 0.762,0.882,0.90 | 0.15 | ENSG00000115 | [[[[27323052, 27 | 3 |
| ENSG00000006 | REX1BD | chr19 | 18590853 | 18590933 | 18588994 | 18589077 | 18592107 | 18592327 | 25,39,64 | 0,0,0 | 24,33,20 | 9,8,13 | 230 | 150 | 4.62E-14 | 5.01E-12 | 1.0,1.0,1.0 | 0.635,0.729,0.50 | 0.378 | ENSG00000006 | [[[[18588798, 18 | 2 |
| ENSG00000141 | NPEPPS | chr17 | 47579389 | 47579511 | 47569416 | 47569494 | 47586364 | 47586398 | 22,11,45 | 0,0,0 | 11,23,9 | 1,8,4 | 272 | 150 | 1.52E-08 | 6.49E-07 | 1.0,1.0,1.0 | 0.858,0.613,0.55 | 0.325 | ENSG00000141 | [[[[47531108, 47 | 4 |
| ENSG00000141 | NPEPPS | chr17 | 47585499 | 47585700 | 47579389 | 47579511 | 47586147 | 47586244 | 77,40,97 | 0,0,0 | 21,22,27 | 2,5,0 | 300 | 150 | 0.000616786 | 0.00769322 | 1.0,1.0,1.0 | 0.84,0.688,1.0 | 0.157 | ENSG00000141 | [[[[47531108, 47 | 2 |
| ENSG00000141 | NPEPPS | chr17 | 47586147 | 47586244 | 47569416 | 47569494 | 47586364 | 47586398 | 40,21,60 | 0,0,0 | 17,29,18 | 1,8,4 | 247 | 150 | 5.44E-08 | 2.04E-06 | 1.0,1.0,1.0 | 0.912,0.688,0.73 | 0.223 | ENSG00000141 | [[[[47531108, 47 | 4 |
| ENSG00000108 | RAD51C | chr17 | 58694930 | 58695189 | 58692572 | 58692685 | 58703195 | 58703329 | 15,6,13 | 0,0,0 | 6,9,7 | 4,15,11 | 300 | 150 | 5.44E-14 | 5.85E-12 | 1.0,1.0,1.0 | 0.429,0.231,0.24 | 0.7 | ENSG00000108 | [[[[58692602, 58 | 2 |
| ENSG00000108 | RAD51C | chr17 | 58694930 | 58695189 | 58692572 | 58692685 | 58709858 | 58709911 | 14,6,9 | 0,0,0 | 5,5,11 | 10,17,5 | 300 | 150 | 1.37E-12 | 1.22E-10 | 1.0,1.0,1.0 | 0.2,0.128,0.524 | 0.716 | ENSG00000108 | [[[[58692602, 58 | 3 |
| ENSG00000108 | RAD51C | chr17 | 58696692 | 58696859 | 58692572 | 58692685 | 58703195 | 58703329 | 26,24,32 | 0,0,0 | 34,62,28 | 4,15,11 | 300 | 150 | 6.39E-10 | 3.66E-08 | 1.0,1.0,1.0 | 0.81,0.674,0.56 | 0.319 | ENSG00000108 | [[[[58692602, 58 | 2 |
| ENSG00000108 | RAD51C | chr17 | 58703195 | 58703329 | 58692572 | 58692685 | 58709858 | 58709911 | 57,30,48 | 0,0,0 | 59,108,80 | 10,17,5 | 284 | 150 | 2.49E-09 | 1.27E-07 | 1.0,1.0,1.0 | 0.757,0.77,0.894 | 0.193 | ENSG00000108 | [[[[58692602, 58 | 3 |

| | | | | | | | | | | | | | | | | | | | | | | |
|---|---|---|---|---|---|---|---|---|---|---|---|---|---|---|---|---|---|---|---|---|---|---|
| ENSG000001411 | TOM1L1 | chr17 | 54903707 | 54903792 | 54900393 | 54900923 | 54912665 | 54912815 | 12,9,36 | 1,2,2 | 10,21,15 | 5,10,6 | 235 | 150 | 0.000186258 | 0.002743042 | 0.885,0.742,0.92 | 0.561,0.573,0.61 | 0.266 | ENSG000001411 | [[[[54900852, 549 | 2 |
| ENSG000001411 | TOM1L1 | chr17 | 54905488 | 54905567 | 54900805 | 54900923 | 54912665 | 54912815 | 20,12,18 | 1,2,2 | 5,24,17 | 5,10,6 | 229 | 150 | 0.00067387 | 0.008287203 | 0.929,0.797,0.85 | 0.396,0.611,0.65 | 0.308 | ENSG000001411 | [[[[54900852, 549 | 2 |
| ENSG000001662 | STXBP4 | chr17 | 54999344 | 54999451 | 54990824 | 54990957 | 55031167 | 55031264 | 22,2,11 | 0,0,0 | 3,10,10 | 4,3,0 | 257 | 150 | 5.88E-05 | 0.001005513 | 1.0,1.0,1.0 | 0.304,0.661,1.0 | 0.345 | ENSG000001662 | [[[[54968765, 549 | 4 |
| ENSG000001539 | MSI2 | chr17 | 57257465 | 57257547 | 57257012 | 57257138 | 57262150 | 57262192 | 24,18,80 | 0,0,2 | 40,54,77 | 7,30,6 | 232 | 150 | 8.31E-08 | 2.97E-06 | 1.0,1.0,0.963 | 0.787,0.538,0.89 | 0.249 | ENSG000001539 | [[[[57256523, 572 | 2 |
| ENSG000001539 | MSI2 | chr17 | 57258269 | 57258354 | 57257012 | 57257138 | 57262150 | 57262192 | 100,53,108 | 0,0,2 | 58,188,123 | 7,30,6 | 235 | 150 | 9.85E-10 | 5.47E-08 | 1.0,1.0,0.972 | 0.841,0.8,0.929 | 0.134 | ENSG000001539 | [[[[57256523, 572 | 2 |
| ENSG000001539 | MSI2 | chr17 | 57674917 | 57675126 | 57652098 | 57652161 | 57679548 | 57680121 | 17,15,54 | 0,2,0 | 25,35,51 | 6,11,0 | 300 | 150 | 0.000152563 | 0.002296746 | 1.0,0.789,1.0 | 0.676,0.614,1.0 | 0.166 | ENSG000001539 | [[[[57256523, 572 | 2 |
| ENSG000001539 | MSI2 | chr17 | 57676986 | 57677059 | 57652098 | 57652161 | 57679548 | 57680121 | 44,35,76 | 0,2,0 | 39,120,93 | 6,11,0 | 223 | 150 | 0.000264728 | 0.003701234 | 1.0,0.922,1.0 | 0.814,0.88,1.0 | 0.076 | ENSG000001539 | [[[[57256523, 572 | 2 |
| ENSG000001808 | CUEDC1 | chr17 | 57869121 | 57869193 | 57866473 | 57866544 | 57871285 | 57871365 | 60,50,107 | 1,0,0 | 33,63,26 | 5,4,4 | 222 | 150 | 2.05E-06 | 5.24E-05 | 0.976,1.0,1.0 | 0.817,0.914,0.81 | 0.143 | ENSG000001808 | [[[[57955225, 579 | 3 |
| ENSG000001887 | ZDHHC9 | chrX | 129841778 | 129841956 | 129828980 | 129829141 | 129843695 | 129843909 | 23,26,40 | 0,1,0 | 25,54,40 | 6,8,5 | 300 | 150 | 3.42E-07 | 1.05E-05 | 1.0,0.929,1.0 | 0.676,0.771,0.8 | 0.227 | ENSG000001887 | [[[[129843696, 12 | 2 |
| ENSG000001887 | ZDHHC9 | chrX | 129841778 | 129842080 | 129828980 | 129829141 | 129843695 | 129843884 | 23,26,40 | 0,1,0 | 25,53,43 | 6,8,5 | 300 | 150 | 3.98E-07 | 1.20E-05 | 1.0,0.929,1.0 | 0.676,0.768,0.81 | 0.225 | ENSG000001887 | [[[[129843696, 12 | 2 |
| ENSG000001083 | MTMR4 | chr17 | 58505471 | 58505583 | 58504302 | 58504488 | 58506742 | 58506871 | 5,3,19 | 0,0,0 | 10,36,29 | 3,7,5 | 262 | 150 | 8.05E-07 | 2.26E-05 | 1.0,1.0,1.0 | 0.656,0.746,0.76 | 0.276 | ENSG000001083 | [[[[58514363, 585 | 2 |
| ENSG000000926 | PHGDH | chr1 | 119740385 | 119740518 | 119737113 | 119737266 | 119742806 | 119743044 | 476,497,822 | 6,3,2 | 7,151,310,747 | 46,72,50 | 283 | 150 | 0 | 0 | 0.977,0.989,0.99 | 0.892,0.906,0.88 | 0.092 | ENSG000000926 | [[[[119711934, 11 | 2 |
| ENSG000000926 | PHGDH | chr1 | 119741766 | 119741897 | 119737113 | 119737266 | 119742806 | 119743044 | 81,174,335 | 6,3,2 | 122,279,214 | 46,72,50 | 281 | 150 | 0 | 0 | 0.878,0.969,0.98 | 0.586,0.674,0.69 | 0.293 | ENSG000000926 | [[[[119711934, 11 | 2 |
| ENSG000001084 | RPS6KB1 | chr17 | 59931621 | 59931722 | 59930116 | 59930174 | 59935192 | 59935300 | 28,15,46 | 0,0,0 | 22,73,42 | 2,4,3 | 251 | 150 | 0.000480963 | 0.006224251 | 1.0,1.0,1.0 | 0.868,0.916,0.89 | 0.108 | ENSG000001084 | [[[[59893121, 598 | 3 |
| ENSG000001988 | OSTC | chr4 | 108655563 | 108655657 | 108650583 | 108650794 | 108667246 | 108667818 | 467,325,572 | 2,0,0 | 354,819,546 | 48,118,67 | 244 | 150 | 0 | 0 | 0.993,1.0,1.0 | 0.819,0.81,0.834 | 0.177 | ENSG000001988 | [[[[108650599, 10 | 2 |
| ENSG000001468 | TLK2 | chr17 | 62606129 | 62606241 | 62600650 | 62600820 | 62608040 | 62608148 | 20,33,43 | 0,0,0 | 35,81,54 | 7,17,11 | 262 | 150 | 1.16E-10 | 7.51E-09 | 1.0,1.0,1.0 | 0.741,0.732,0.73 | 0.263 | ENSG000001468 | [[[[62478825, 624 | 2 |
| ENSG000000087 | PKD1 | chr16 | 2093810 | 2094013 | 2093543 | 2093738 | 2097147 | 2097241 | 7,33,55 | 0,0,0 | 13,36,30 | 2,6,3 | 300 | 150 | 9.49E-07 | 2.62E-05 | 1.0,1.0,1.0 | 0.765,0.75,0.833 | 0.217 | ENSG000000087 | [[[[2135475, 2135 | 2 |
| ENSG000001200 | GNA13 | chr17 | 65018252 | 65018303 | 65010714 | 65014829 | 65056310 | 65056839 | 3,3,10 | 0,0,0 | 17,43,46 | 0,6,6 | 201 | 150 | 0.000100045 | 0.001599166 | 1.0,1.0,1.0 | 1.0,0.842,0.851 | 0.102 | ENSG000001200 | [[[[65056311, 650 | 2 |
| ENSG000001200 | GNA13 | chr17 | 65053501 | 65053728 | 65010714 | 65014829 | 65056310 | 65056839 | 26,10,26 | 0,0,0 | 24,48,16 | 0,6,6 | 300 | 150 | 7.08E-06 | 0.000157146 | 1.0,1.0,1.0 | 1.0,0.8,0.571 | 0.21 | ENSG000001200 | [[[[65056311, 650 | 2 |
| ENSG000001716 | BPTF | chr17 | 67928958 | 67929120 | 67928354 | 67928601 | 67940438 | 67940545 | 4,2,8 | 0,0,0 | 1,9,9 | 1,8,5 | 300 | 150 | 1.90E-10 | 1.19E-08 | 1.0,1.0,1.0 | 0.333,0.36,0.474 | 0.611 | ENSG000001716 | [[[[67825503, 678 | 2 |
| ENSG000001716 | BPTF | chr17 | 67929335 | 67929487 | 67928354 | 67928601 | 67940438 | 67940545 | 51,26,52 | 0,0,0 | 37,62,73 | 1,8,5 | 300 | 150 | 3.73E-06 | 8.91E-05 | 1.0,1.0,1.0 | 0.949,0.795,0.88 | 0.125 | ENSG000001716 | [[[[67825503, 678 | 2 |
| ENSG000001716 | BPTF | chr17 | 67931910 | 67932019 | 67928354 | 67928601 | 67940438 | 67940545 | 35,30,57 | 0,0,0 | 27,38,29 | 1,8,5 | 259 | 150 | 2.01E-07 | 6.53E-06 | 1.0,1.0,1.0 | 0.94,0.733,0.771 | 0.185 | ENSG000001716 | [[[[67825503, 678 | 2 |
| ENSG000000705 | WIPI1 | chr17 | 68427134 | 68427253 | 68421281 | 68421820 | 68428828 | 68428936 | 11,18,38 | 0,0,0 | 25,50,29 | 6,17,6 | 269 | 150 | 2.95E-10 | 1.79E-08 | 1.0,1.0,1.0 | 0.699,0.621,0.72 | 0.317 | ENSG000000705 | [[[[68457342, 684 | 2 |
| ENSG000001415 | NARF | chr17 | 82468763 | 82468896 | 82464286 | 82464430 | 82478799 | 82478918 | 147,126,205 | 0,0,0 | 206,368,252 | 4,6,9 | 283 | 150 | 0.000256391 | 0.003599207 | 1.0,1.0,1.0 | 0.965,0.97,0.937 | 0.043 | ENSG000001415 | [[[[82458741, 824 | 2 |
| ENSG000001415 | NARF | chr17 | 82478570 | 82478667 | 82464286 | 82464430 | 82478799 | 82478918 | 2,4,7 | 0,0,0 | 1,4,9 | 4,6,9 | 247 | 150 | 2.90E-11 | 2.05E-09 | 1.0,1.0,1.0 | 0.132,0.288,0.37 | 0.734 | ENSG000001415 | [[[[82458741, 824 | 2 |
| ENSG000001415 | NARF | chr17 | 82478799 | 82478918 | 82464286 | 82464430 | 82481081 | 82481093 | 48,91,153 | 0,0,0 | 179,282,227 | 7,14,4 | 269 | 150 | 9.67E-06 | 0.000207609 | 1.0,1.0,1.0 | 0.934,0.918,0.96 | 0.06 | ENSG000001415 | [[[[82458741, 824 | 3 |
| ENSG000001615 | SAP30BP | chr17 | 75703310 | 75703371 | 75702479 | 75702571 | 75704755 | 75704768 | 71,59,67 | 1,1,2 | 55,103,67 | 19,36,11 | 211 | 150 | 6.58E-13 | 6.12E-11 | 0.981,0.977,0.96 | 0.673,0.67,0.812 | 0.254 | ENSG000001615 | [[[[75667338, 756 | 2 |
| ENSG000001615 | SAP30BP | chr17 | 75703807 | 75703859 | 75702479 | 75702571 | 75704755 | 75704768 | 61,54,116 | 1,1,2 | 104,183,146 | 19,36,11 | 202 | 150 | 1.31E-09 | 7.08E-08 | 0.978,0.976,0.97 | 0.803,0.791,0.90 | 0.143 | ENSG000001615 | [[[[75667338, 756 | 2 |
| ENSG000001197 | COQ6 | chr14 | 73955804 | 73955928 | 73955450 | 73955509 | 73958970 | 73959078 | 16,18,36 | 0,0,0 | 9,14,14 | 4,7,7 | 274 | 150 | 6.28E-12 | 5.04E-10 | 1.0,1.0,1.0 | 0.552,0.523,0.52 | 0.467 | ENSG000001197 | [[[[73950302, 739 | 2 |
| ENSG000001197 | COQ6 | chr14 | 73958146 | 73958277 | 73955450 | 73955509 | 73958970 | 73959078 | 20,17,34 | 0,0,0 | 11,15,18 | 4,7,7 | 281 | 150 | 2.79E-11 | 1.98E-09 | 1.0,1.0,1.0 | 0.595,0.534,0.57 | 0.431 | ENSG000001197 | [[[[73950302, 739 | 2 |
| ENSG000001678 | MGAT5B | chr17 | 76903302 | 76903376 | 76902554 | 76902670 | 76906017 | 76906187 | 11,19,31 | 0,1,1 | 44,48,42 | 6,6,8 | 224 | 150 | 6.43E-05 | 0.00108758 | 1.0,0.927,0.954 | 0.831,0.843,0.77 | 0.143 | ENSG000001678 | [[[[76868404, 768 | 3 |
| ENSG000001678 | MGAT5B | chr17 | 76904251 | 76904422 | 76903302 | 76903376 | 76906017 | 76906187 | 10,34,58 | 2,0,1 | 27,85,67 | 11,8,9 | 300 | 150 | 0.000115422 | 0.001807792 | 0.714,1.0,0.967 | 0.551,0.842,0.78 | 0.167 | ENSG000001678 | [[[[76868404, 768 | 2 |
| ENSG000001678 | MGAT5B | chr17 | 76924965 | 76925097 | 76906017 | 76906187 | 76937987 | 76938143 | 10,27,84 | 0,3,0 | 51,79,76 | 6,16,5 | 282 | 150 | 5.58E-05 | 0.000959813 | 1.0,0.827,1.0 | 0.819,0.724,0.89 | 0.131 | ENSG000001678 | [[[[76868404, 768 | 4 |
| ENSG000001678 | MGAT5B | chr17 | 76932644 | 76932775 | 76926596 | 76926730 | 76937987 | 76938143 | 24,56,98 | 8,18,26 | 29,99,67 | 37,64,56 | 281 | 150 | 3.45E-06 | 8.33E-05 | 0.616,0.624,0.66 | 0.295,0.452,0.39 | 0.257 | ENSG000001678 | [[[[76868404, 768 | 2 |
| ENSG000001824 | NPLOC4 | chr17 | 81567416 | 81567533 | 81561765 | 81565607 | 81572016 | 81572073 | 58,79,125 | 1,4,0 | 64,114,78 | 13,28,19 | 267 | 150 | 1.20E-13 | 1.24E-11 | 0.97,0.917,1.0 | 0.734,0.696,0.69 | 0.253 | ENSG000001824 | [[[[81636916, 816 | 2 |
| ENSG000001824 | NPLOC4 | chr17 | 81569015 | 81569111 | 81561765 | 81565607 | 81572016 | 81572073 | 89,95,174 | 1,4,0 | 122,214,174 | 13,28,19 | 246 | 150 | 1.84E-10 | 1.16E-08 | 0.982,0.935,1.0 | 0.851,0.823,0.84 | 0.132 | ENSG000001824 | [[[[81636916, 816 | 2 |
| ENSG000001824 | NPLOC4 | chr17 | 81600340 | 81600427 | 81597244 | 81597316 | 81610209 | 81610258 | 142,121,233 | 0,0,0 | 125,292,212 | 3,12,7 | 237 | 150 | 7.73E-06 | 0.000170512 | 1.0,1.0,1.0 | 0.963,0.939,0.95 | 0.049 | ENSG000001824 | [[[[81636916, 816 | 4 |
| ENSG000001824 | NPLOC4 | chr17 | 81604547 | 81604727 | 81572016 | 81572088 | 81610209 | 81610258 | 2,4,5 | 0,0,0 | 7,5,0 | 5,2,2 | 300 | 150 | 3.45E-09 | 1.71E-07 | 1.0,1.0,1.0 | 0.412,0.556,0.0 | 0.677 | ENSG000001824 | [[[[81636916, 816 | 7 |
| ENSG000001824 | NPLOC4 | chr17 | 81604547 | 81604727 | 81597244 | 81597316 | 81610209 | 81610258 | 2,5,6 | 0,0,0 | 7,9,1 | 3,12,7 | 300 | 150 | 5.58E-11 | 3.79E-09 | 1.0,1.0,1.0 | 0.538,0.273,0.06 | 0.707 | ENSG000001824 | [[[[81636916, 816 | 4 |
| ENSG000001856 | P4HB | chr17 | 81845560 | 81845742 | 81845143 | 81845230 | 81859180 | 81859387 | 218,322,363,254 | 0,0,0 | 280,456,413,468 | 35,97,74 | 300 | 150 | 0 | 0 | 1.0,1.0,1.0 | 0.976,0.967,0.95 | 0.033 | ENSG000001856 | [[[[81860327, 818 | 7 |
| ENSG000001856 | P4HB | chr17 | 81855141 | 81855279 | 81845143 | 81845230 | 81859178 | 81859387 | 3,1,5 | 0,0,0 | 5,12,6 | 4,5,1 | 288 | 150 | 2.33E-08 | 9.56E-07 | 1.0,1.0,1.0 | 0.394,0.556,0.75 | 0.431 | ENSG000001856 | [[[[81860327, 818 | 7 |
| ENSG000001370 | RNF38 | chr9 | 36390466 | 36390616 | 36375933 | 36376127 | 36400935 | 36401198 | 2,4,7 | 2,4,1 | 15,27,17 | 0,1,0 | 300 | 150 | 1.06E-06 | 2.91E-05 | 0.333,0.333,0.77 | 1.0,0.931,1.0 | -0.496 | ENSG000001370 | [[[[36400097, 364 | 2 |
| ENSG000000809 | NDC80 | chr18 | 2577745 | 2577869 | 2571510 | 2571683 | 2577968 | 2578141 | 130,105,111 | 0,0,0 | 75,188,106 | 4,25,11 | 274 | 150 | 5.32E-13 | 5.03E-11 | 1.0,1.0,1.0 | 0.911,0.805,0.84 | 0.148 | ENSG000000809 | [[[[2571557, 2571 | 3 |
| ENSG000000809 | NDC80 | chr18 | 2578926 | 2579029 | 2571510 | 2571683 | 2585112 | 2585202 | 41,42,65 | 0,0,0 | 28,89,49 | 3,17,3 | 253 | 150 | 7.51E-09 | 3.44E-07 | 1.0,1.0,1.0 | 0.847,0.756,0.90 | 0.164 | ENSG000000809 | [[[[2571557, 2571 | 5 |
| ENSG000000954 | CWF19L1 | chr10 | 100256261 | 100256476 | 100253420 | 100253539 | 100260965 | 100261044 | 20,23,19 | 0,0,0 | 13,22,25 | 2,7,4 | 300 | 150 | 4.50E-08 | 1.72E-06 | 1.0,1.0,1.0 | 0.765,0.611,0.75 | 0.289 | ENSG000000954 | [[[[100267571, 10 | 2 |
| ENSG000000954 | CWF19L1 | chr10 | 100260217 | 100260319 | 100253420 | 100253539 | 100260965 | 100261028 | 15,17,23 | 0,0,0 | 28,29,16 | 2,7,4 | 252 | 150 | 5.85E-07 | 1.70E-05 | 1.0,1.0,1.0 | 0.893,0.711,0.70 | 0.231 | ENSG000000954 | [[[[100267571, 10 | 2 |
| ENSG000000954 | CWF19L1 | chr10 | 100260965 | 100261044 | 100253420 | 100253539 | 100261978 | 100262063 | 13,21,21 | 0,0,0 | 15,32,16 | 6,7,0 | 229 | 150 | 2.02E-06 | 5.17E-05 | 1.0,1.0,1.0 | 0.621,0.75,1.0 | 0.21 | ENSG000000954 | [[[[100267571, 10 | 3 |
| ENSG000001548 | PPP4R1 | chr18 | 9588710 | 9588853 | 9584720 | 9584828 | 9593767 | 9593874 | 69,38,118 | 0,0,0 | 32,80,50 | 5,4,5 | 293 | 150 | 3.39E-07 | 1.05E-05 | 1.0,1.0,1.0 | 0.766,0.911,0.83 | 0.162 | ENSG000001548 | [[[[9614478, 9614 | 2 |
| ENSG000001413 | AFG3L2 | chr18 | 12351084 | 12351210 | 12344131 | 12344247 | 12351305 | 12351413 | 86,55,122 | 0,0,0 | 30,113,91 | 6,26,15 | 276 | 150 | 0 | 0 | 1.0,1.0,1.0 | 0.731,0.703,0.76 | 0.266 | ENSG000001413 | [[[[12376969, 123 | 2 |
| ENSG000001753 | PTPN2 | chr18 | 12801969 | 12802151 | 12792152 | 12794485 | 12825809 | 12825944 | 40,45,64 | 0,0,0 | 48,104,59 | 0,10,6 | 300 | 150 | 3.35E-06 | 8.11E-05 | 1.0,1.0,1.0 | 1.0,0.839,0.831 | 0.11 | ENSG000001753 | [[[[12884073, 128 | 3 |
| ENSG000001753 | PTPN2 | chr18 | 12814202 | 12814355 | 12801969 | 12802151 | 12825809 | 12825944 | 43,42,55 | 0,0,0 | 48,81,47 | 0,4,11 | 300 | 150 | 7.77E-06 | 0.00017134 | 1.0,1.0,1.0 | 1.0,0.91,0.681 | 0.136 | ENSG000001753 | [[[[12884073, 128 | 2 |
| ENSG000001193 | GLE1 | chr9 | 128508875 | 128509097 | 128504699 | 128504904 | 128522667 | 128522816 | 24,17,40 | 0,0,0 | 8,13,19 | 2,5,5 | 300 | 150 | 3.78E-09 | 1.85E-07 | 1.0,1.0,1.0 | 0.667,0.565,0.65 | 0.371 | ENSG000001193 | [[[[128504719, 12 | 2 |
| ENSG000001193 | GLE1 | chr9 | 128515528 | 128515639 | 128504699 | 128504904 | 128522667 | 128522816 | 35,37,65 | 0,0,0 | 24,44,31 | 2,5,5 | 261 | 150 | 8.60E-07 | 2.40E-05 | 1.0,1.0,1.0 | 0.873,0.835,0.78 | 0.17 | ENSG000001193 | [[[[128504719, 12 | 2 |
| ENSG000001386 | SPPL2A | chr15 | 50722123 | 50722201 | 50719939 | 50720100 | 50726320 | 50726377 | 68,60,97 | 0,0,0 | 56,96,52 | 1,7,4 | 228 | 150 | 7.10E-05 | 0.001183556 | 1.0,1.0,1.0 | 0.974,0.9,0.895 | 0.077 | ENSG000001386 | [[[[50765468, 507 | 2 |
| ENSG000001386 | SPPL2A | chr15 | 50730964 | 50731039 | 50725915 | 50726377 | 50736100 | 50736202 | 30,35,47 | 0,0,0 | 26,48,16 | 6,8,12 | 225 | 150 | 1.22E-10 | 7.92E-09 | 1.0,1.0,1.0 | 0.743,0.8,0.471 | 0.329 | ENSG000001386 | [[[[50765468, 507 | 2 |
| ENSG000001386 | SPPL2A | chr15 | 50732602 | 50732684 | 50726320 | 50726377 | 50736100 | 50736202 | 80,85,105 | 0,0,0 | 61,137,72 | 6,8,12 | 232 | 150 | 1.19E-09 | 6.52E-08 | 1.0,1.0,1.0 | 0.868,0.917,0.79 | 0.14 | ENSG000001386 | [[[[50765468, 507 | 2 |
| ENSG000001386 | SPPL2A | chr15 | 50736100 | 50736202 | 50732602 | 50732684 | 50747494 | 50747508 | 83,76,102 | 0,0,0 | 75,120,81 | 1,4,6 | 252 | 150 | 0.000634048 | 0.007873732 | 1.0,1.0,1.0 | 0.978,0.947,0.88 | 0.062 | ENSG000001386 | [[[[50765468, 507 | 3 |
| ENSG000001347 | ELP2 | chr18 | 36142280 | 36142347 | 36141136 | 36141201 | 36154849 | 36154999 | 10,1,10 | 0,0,1 | 2,14,8 | 1,3,2 | 217 | 150 | 0.000828068 | 0.009893949 | 1.0,1.0,0.874 | 0.58,0.763,0.734 | 0.266 | ENSG000001347 | [[[[36129899, 361 | 5 |
| ENSG000002745 | RCC1L | chr7 | 75055900 | 75056074 | 75052710 | 75052796 | 75058587 | 75058769 | 153,158,212 | 0,2,0 | 136,281,163 | 6,17,11 | 300 | 150 | 2.23E-09 | 1.15E-07 | 1.0,0.975,1.0 | 0.919,0.892,0.88 | 0.094 | ENSG000002745 | [[[[75073414, 750 | 2 |
| ENSG000002745 | RCC1L | chr7 | 75057528 | 75057616 | 75052710 | 75052796 | 75058587 | 75058769 | 56,77,102 | 0,2,0 | 88,101,68 | 6,17,11 | 238 | 150 | 9.12E-10 | 5.11E-08 | 1.0,0.96,1.0 | 0.902,0.789,0.79 | 0.158 | ENSG000002745 | [[[[75073414, 750 | 2 |
| ENSG000001753 | SMAD2 | chr18 | 47845339 | 47845484 | 47808956 | 47841950 | 47851273 | 47851327 | 35,29,53 | 0,0,0 | 28,41,24 | 2,17,1 | 295 | 150 | 2.52E-07 | 8.00E-06 | 1.0,1.0,1.0 | 0.877,0.551,0.92 | 0.216 | ENSG000001753 | [[[[47930361, 479 | 3 |
| ENSG000001753 | SMAD2 | chr18 | 47848474 | 47848687 | 47808956 | 47841950 | 47851273 | 47851327 | 30,27,34 | 0,0,0 | 37,108,61 | 2,17,1 | 300 | 150 | 3.30E-06 | 7.99E-05 | 1.0,1.0,1.0 | 0.902,0.761,0.96 | 0.123 | ENSG000001753 | [[[[47930361, 479 | 3 |
| ENSG000001527 | GPR180 | chr13 | 94619467 | 94619517 | 94612189 | 94612390 | 94625965 | 94626043 | 5,2,4 | 0,0,0 | 0,0,3 | 6,1,5 | 200 | 150 | 6.33E-15 | 7.80E-13 | 1.0,1.0,1.0 | 0.0,0.0,0.31 | 0.897 | ENSG000001527 | [[[[94601857, 946 | 4 |
| ENSG000001664 | SERPINB8 | chr18 | 63985092 | 63985245 | 63981720 | 63981838 | 63986873 | 63989371 | 10,7,19 | 0,0,0 | 23,43,32 | 2,6,2 | 300 | 150 | 2.35E-05 | 0.000453426 | 1.0,1.0,1.0 | 0.852,0.782,0.88 | 0.159 | ENSG000001664 | [[[[63970081, 639 | 2 |
| ENSG000001664 | SERPINB8 | chr18 | 63985092 | 63985245 | 63981720 | 63981838 | 63986962 | 63987396 | 5,3,1 | 0,0,0 | 2,21,14 | 2,7,0 | 300 | 150 | 9.02E-05 | 0.00145833 | 1.0,1.0,1.0 | 0.333,0.6,1.0 | 0.356 | ENSG000001664 | [[[[63970081, 639 | 2 |
| ENSG000001569 | VPS8 | chr3 | 184886109 | 184886156 | 184870715 | 184870805 | 184900920 | 184900931 | 5,1,11 | 0,0,0 | 2,4,7 | 2,3,1 | 197 | 150 | 8.69E-05 | 0.001409431 | 1.0,1.0,1.0 | 0.432,0.504,0.84 | 0.407 | ENSG000001569 | [[[[184812166, 18 | 3 |
| ENSG000001569 | VPS8 | chr3 | 184898564 | 184898654 | 184870715 | 184870805 | 184900920 | 184900931 | 11,7,16 | 0,0,0 | 2,8,3 | 2,3,1 | 240 | 150 | 2.71E-05 | 0.00051059 | 1.0,1.0,1.0 | 0.385,0.625,0.65 | 0.446 | ENSG000001569 | [[[[184812166, 18 | 3 |
| ENSG000000089 | SEC62 | chr3 | 169988239 | 169988359 | 169983160 | 169983253 | 169992593 | 169992721 | 94,66,122 | 0,0,0 | 119,236,156 | 13,31,19 | 270 | 150 | 1.78E-14 | 2.07E-12 | 1.0,1.0,1.0 | 0.836,0.809,0.82 | 0.178 | ENSG000000089 | [[[[169966807, 16 | 2 |
| ENSG000000089 | SEC62 | chr3 | 169992593 | 169993101 | 169983160 | 169983253 | 169993654 | 169993831 | 2,8,3 | 0,0,0 | 40,79,37 | 5,16,3 | 300 | 150 | 9.23E-06 | 0.00019955 | 1.0,1.0,1.0 | 0.8,0.712,0.86 | 0.209 | ENSG000000089 | [[[[169966807, 16 | 2 |
| ENSG000001290 | VPS13C | chr15 | 61882595 | 61882736 | 61881746 | 61881828 | 61890164 | 61890400 | 12,18,19 | 0,0,0 | 3,15,8 | 1,2,4 | 291 | 150 | 7.59E-06 | 0.000167903 | 1.0,1.0,1.0 | 0.607,0.794,0.50 | 0.364 | ENSG000001290 | [[[[62060275, 620 | 2 |
| ENSG000001963 | LONP1 | chr19 | 5707059 | 5707143 | 5705771 | 5705992 | 5711770 | 5712002 | 70,72,109 | 0,0,0 | 103,195,134 | 12,10,3 | 234 | 150 | 6.40E-07 | 1.85E-05 | 1.0,1.0,1.0 | 0.846,0.926,0.96 | 0.087 | ENSG000001963 | [[[[5719704, 5720 | 3 |
| ENSG000001963 | LONP1 | chr19 | 5708341 | 5708403 | 5705771 | 5705992 | 5711770 | 5711950 | 10,15,45 | 0,0,0 | 33,44,71 | 12,10,3 | 212 | 150 | 1.64E-07 | 5.42E-06 | 1.0,1.0,1.0 | 0.661,0.757,0.94 | 0.213 | ENSG000001963 | [[[[5719704, 5720 | 3 |
| ENSG000000857 | DDHD2 | chr8 | 38245741 | 38245950 | 38240274 | 38240364 | 38246232 | 38246300 | 21,9,10 | 0,0,0 | 8,21,16 | 1,8,4 | 300 | 150 | 2.38E-08 | 9.77E-07 | 1.0,1.0,1.0 | 0.8,0.568,0.667 | 0.322 | ENSG000000857 | [[[[38231585, 382 | 2 |
| ENSG000001077 | MINPP1 | chr10 | 87513123 | 87513221 | 87508335 | 87508533 | 87552081 | 87553388 | 34,32,54 | 0,0,0 | 19,39,37 | 5,10,1 | 248 | 150 | 1.08E-07 | 3.74E-06 | 1.0,1.0,1.0 | 0.697,0.702,0.95 | 0.215 | ENSG000001077 | [[[[87504893, 875 | 2 |
| ENSG000001275 | EPS15L1 | chr19 | 16377121 | 16377254 | 16355238 | 16355851 | 16385128 | 16385211 | 27,19,20 | 0,8,2 | 8,9,7 | 6,13,7 | 283 | 150 | 3.63E-05 | 0.000662475 | 1.0,0.557,0.841 | 0.414,0.268,0.34 | 0.457 | ENSG000001275 | [[[[16471913, 164 | 2 |
| ENSG000001053 | BABAM1 | chr19 | 17273903 | 17274024 | 17268793 | 17269091 | 17276494 | 17276555 | 3,8,4 | 0,0,0 | 48,50,18 | 3,7,0 | 271 | 150 | 0.000174727 | 0.002591251 | 1.0,1.0,1.0 | 0.899,0.798,1.0 | 0.101 | ENSG000001053 | [[[[17267443, 172 | 4 |
| ENSG000001053 | BABAM1 | chr19 | 17274106 | 17274185 | 17273903 | 17274024 | 17276494 | 17276525 | 81,127,181 | 2,6,4 | 160,282,214 | 40,51,15 | 229 | 150 | 1.47E-08 | 6.31E-07 | 0.964,0.933,0.96 | 0.724,0.784,0.90 | 0.151 | ENSG000001053 | [[[[17267443, 172 | 2 |
| ENSG000001153 | AUP1 | chr2 | 74527247 | 74527363 | 74526940 | 74527059 | 74527939 | 74528006 | 151,194,246 | 0,0,0 | 249,519,285 | 9,25,13 | 266 | 150 | 5.09E-10 | 2.98E-08 | 1.0,1.0,1.0 | 0.94,0.921,0.925 | 0.071 | ENSG000001153 | [[[[74529580, 745 | 3 |
| ENSG000001153 | AUP1 | chr2 | 74527735 | 74527838 | 74526940 | 74527059 | 74527939 | 74528006 | 113,101,155 | 0,0,0 | 172,248,145 | 9,25,13 | 253 | 150 | 4.15E-12 | 3.45E-10 | 1.0,1.0,1.0 | 0.919,0.855,0.86 | 0.119 | ENSG000001153 | [[[[74529580, 745 | 3 |
| ENSG000001153 | AUP1 | chr2 | 74527735 | 74527838 | 74527247 | 74527363 | 74527939 | 74528006 | 114,101,158 | 0,0,0 | 179,266,154 | 16,23,6 | 253 | 150 | 2.80E-11 | 1.99E-09 | 1.0,1.0,1.0 | 0.869,0.873,0.93 | 0.107 | ENSG000001153 | [[[[74529580, 745 | 2 |

| | | | | | | | | | | | | | | | | | | | | | | |
|---|---|---|---|---|---|---|---|---|---|---|---|---|---|---|---|---|---|---|---|---|---|---|
| ENSG00000115 | AUP1 | chr2 | 74528750 | 74528935 | 74527939 | 74528006 | 74529131 | 74529282 | 134,118,188 | 0,0,0 | 239,399,264 | 21,27,15 | 300 | 150 | 6.23E-13 | 5.81E-11 | 1.0,1.0,1.0 | 0.851,0.881,0.89 | 0.123 | ENSG00000115 | [[[[74529580, 74 | 3 |
| ENSG00000257 | LSM14A | chr19 | 34219390 | 34219573 | 34215595 | 34215661 | 34221506 | 34221738 | 45,41,63 | 0,0,0 | 43,94,73 | 16,47,23 | 300 | 150 | 0 | 0 | 1.0,1.0,1.0 | 0.573,0.5,0.613 | 0.438 | ENSG00000257 | [[[[34172504, 34 | 2 |
| ENSG00000257 | LSM14A | chr19 | 34219705 | 34219877 | 34215595 | 34215661 | 34221506 | 34221738 | 154,142,187 | 0,0,0 | 184,285,193 | 16,47,23 | 300 | 150 | 0 | 0 | 1.0,1.0,1.0 | 0.852,0.752,0.80 | 0.196 | ENSG00000257 | [[[[34172504, 34 | 2 |
| ENSG00000075 | WDR62 | chr19 | 36071555 | 36071716 | 36067827 | 36068010 | 36081432 | 36081570 | 9,10,27 | 0,0,2 | 10,12,11 | 2,4,3 | 300 | 150 | 0.000108021 | 0.001706251 | 1.0,1.0,0.871 | 0.714,0.6,0.647 | 0.303 | ENSG00000075 | [[[[36054897, 36 | 2 |
| ENSG00000075 | WDR62 | chr19 | 36073341 | 36073531 | 36067827 | 36068010 | 36081432 | 36081570 | 20,24,24 | 0,0,2 | 13,19,6 | 2,4,3 | 300 | 150 | 5.38E-05 | 0.000931521 | 1.0,1.0,0.857 | 0.765,0.704,0.5 | 0.296 | ENSG00000075 | [[[[36054897, 36 | 2 |
| ENSG00000178 | EIF3K | chr19 | 38632429 | 38632496 | 38626027 | 38626102 | 38634992 | 38635118 | 255,195,370 | 0,2,1 | 312,613,352 | 8,37,10 | 217 | 150 | 2.24E-09 | 1.15E-07 | 1.0,0.985,0.996 | 0.964,0.92,0.961 | 0.045 | ENSG00000178 | [[[[38619188, 38 | 2 |
| ENSG00000167 | AXL | chr19 | 41239163 | 41239314 | 41237943 | 41238154 | 41242882 | 41243015 | 16,26,32 | 0,0,0 | 21,38,19 | 2,9,6 | 300 | 150 | 1.54E-08 | 6.57E-07 | 1.0,1.0,1.0 | 0.84,0.679,0.613 | 0.289 | ENSG00000167 | [[[[41219223, 41 | 3 |
| ENSG00000167 | AXL | chr19 | 41239693 | 41239720 | 41237943 | 41238154 | 41242882 | 41243015 | 4,3,3 | 0,0,0 | 1,9,8 | 2,9,6 | 177 | 150 | 3.23E-09 | 1.61E-07 | 1.0,1.0,1.0 | 0.298,0.459,0.53 | 0.571 | ENSG00000167 | [[[[41219223, 41 | 3 |
| ENSG00000104 | ERCC2 | chr19 | 45364423 | 45364547 | 45364234 | 45364331 | 45368629 | 45368743 | 13,27,30 | 0,1,0 | 22,53,37 | 2,4,5 | 274 | 150 | 0.000140106 | 0.002133023 | 1.0,0.937,1.0 | 0.858,0.879,0.80 | 0.133 | ENSG00000104 | [[[[45370536, 45 | 3 |
| ENSG00000063 | RPL18 | chr19 | 48615876 | 48615946 | 48615327 | 48615447 | 48616725 | 48616824 | 20,71,164 | 0,0,0 | 93,179,156 | 12,26,22 | 220 | 150 | 2.82E-14 | 3.16E-12 | 1.0,1.0,1.0 | 0.841,0.824,0.82 | 0.169 | ENSG00000063 | [[[[48619141, 48 | 2 |
| ENSG00000135 | SYNCRIP | chr6 | 85614980 | 85615347 | 85607784 | 85612923 | 85622481 | 85622687 | 51,98,183 | 0,0,0 | 101,219,193 | 9,14,7 | 300 | 150 | 1.04E-08 | 4.62E-07 | 1.0,1.0,1.0 | 0.849,0.887,0.93 | 0.111 | ENSG00000135 | [[[[85642797, 85 | 3 |
| ENSG00000135 | SYNCRIP | chr6 | 85619267 | 85619417 | 85607784 | 85612923 | 85622481 | 85622687 | 349,251,429 | 0,0,0 | 177,421,289 | 9,14,7 | 300 | 150 | 3.35E-09 | 1.66E-07 | 1.0,1.0,1.0 | 0.908,0.938,0.95 | 0.067 | ENSG00000135 | [[[[85642797, 85 | 3 |
| ENSG00000135 | SYNCRIP | chr6 | 85622481 | 85622687 | 85619267 | 85619417 | 85637242 | 85637356 | 349,251,429 | 0,0,0 | 174,418,283 | 1,10,8 | 300 | 150 | 6.53E-05 | 0.001101505 | 1.0,1.0,1.0 | 0.989,0.954,0.94 | 0.037 | ENSG00000135 | [[[[85642797, 85 | 3 |
| ENSG00000135 | SYNCRIP | chr6 | 85636966 | 85637143 | 85619267 | 85619417 | 85637242 | 85637356 | 306,281,415 | 0,0,0 | 188,410,278 | 1,10,8 | 300 | 150 | 6.86E-05 | 0.001149763 | 1.0,1.0,1.0 | 0.989,0.953,0.94 | 0.037 | ENSG00000135 | [[[[85642797, 85 | 3 |
| ENSG00000039 | PNKP | chr19 | 49865126 | 49865426 | 49864178 | 49864236 | 49866398 | 49866445 | 40,49,35 | 0,0,0 | 34,65,51 | 5,13,4 | 300 | 150 | 5.44E-09 | 2.57E-07 | 1.0,1.0,1.0 | 0.773,0.714,0.86 | 0.216 | ENSG00000039 | [[[[49867469, 49 | 2 |
| ENSG00000105 | VRK3 | chr19 | 49989638 | 49989771 | 49987700 | 49988492 | 49994813 | 49994919 | 13,28,57 | 0,0,0 | 38,92,73 | 1,8,1 | 283 | 150 | 0.000559377 | 0.007085007 | 1.0,1.0,1.0 | 0.953,0.859,0.97 | 0.071 | ENSG00000105 | [[[[50025267, 50 | 2 |
| ENSG00000105 | VRK3 | chr19 | 49992859 | 49992952 | 49987700 | 49988492 | 49994813 | 49994919 | 12,14,29 | 0,0,0 | 36,61,39 | 1,8,1 | 243 | 150 | 0.000197131 | 0.002878313 | 1.0,1.0,1.0 | 0.957,0.825,0.96 | 0.086 | ENSG00000105 | [[[[50025267, 50 | 2 |
| ENSG00000019 | ZRANB1 | chr10 | 124983174 | 124983304 | 124981410 | 124981829 | 124984773 | 124988189 | 69,54,78 | 0,0,0 | 91,219,130 | 3,14,4 | 280 | 150 | 6.93E-06 | 0.000154342 | 1.0,1.0,1.0 | 0.942,0.893,0.94 | 0.073 | ENSG00000019 | [[[[124942123, 1 | 2 |
| ENSG00000164 | SERINC5 | chr5 | 80166382 | 80166478 | 80158835 | 80158962 | 80174953 | 80175047 | 46,39,108 | 0,0,0 | 75,201,166 | 6,2,8 | 246 | 150 | 0.000218624 | 0.003147898 | 1.0,1.0,1.0 | 0.884,0.984,0.92 | 0.068 | ENSG00000164 | [[[[80255896, 80 | 2 |
| ENSG00000057 | MTA3 | chr2 | 42707902 | 42708054 | 42697775 | 42697834 | 42708873 | 42709014 | 43,36,43 | 0,0,0 | 28,51,37 | 4,5,3 | 300 | 150 | 1.85E-06 | 4.78E-05 | 1.0,1.0,1.0 | 0.778,0.836,0.86 | 0.175 | ENSG00000057 | [[[[42568632, 42 | 2 |
| ENSG00000198 | SIRPA | chr20 | 1924763 | 1924877 | 1915098 | 1915455 | 1927874 | 1927899 | 65,43,70 | 1,0,0 | 61,104,48 | 0,1,10 | 264 | 150 | 0.000764128 | 0.009224496 | 0.974,1.0,1.0 | 1.0,0.983,0.732 | 0.086 | ENSG00000198 | [[[[1895407, 189 | 3 |
| ENSG00000101 | IDH3B | chr20 | 2660262 | 2660365 | 2659698 | 2659793 | 2660456 | 2660590 | 67,101,101 | 0,0,0 | 95,196,128 | 6,6,3 | 253 | 150 | 0.000196657 | 0.002873317 | 1.0,1.0,1.0 | 0.904,0.951,0.96 | 0.061 | ENSG00000101 | [[[[2664153, 266 | 2 |
| ENSG00000089 | SNX5 | chr20 | 17950290 | 17950396 | 17950131 | 17950207 | 17955364 | 17955475 | 130,94,151 | 0,0,0 | 132,349,218 | 2,14,4 | 256 | 150 | 0.000119911 | 0.001866588 | 1.0,1.0,1.0 | 0.975,0.936,0.97 | 0.04 | ENSG00000089 | [[[[17968375, 17 | 4 |
| ENSG00000089 | SNX5 | chr20 | 17952586 | 17952710 | 17943109 | 17943195 | 17955364 | 17955475 | 1,3,1 | 0,0,0 | 4,7,1 | 5,17,13 | 274 | 150 | 5.38E-09 | 2.54E-07 | 1.0,1.0,1.0 | 0.305,0.184,0.04 | 0.824 | ENSG00000089 | [[[[17968375, 17 | 8 |
| ENSG00000089 | SNX5 | chr20 | 17952586 | 17952710 | 17949063 | 17949103 | 17955364 | 17955475 | 1,3,1 | 0,0,0 | 4,6,1 | 1,3,2 | 274 | 150 | 9.08E-06 | 0.000196678 | 1.0,1.0,1.0 | 0.686,0.523,0.21 | 0.525 | ENSG00000089 | [[[[17968375, 17 | 5 |
| ENSG00000089 | SNX5 | chr20 | 17952586 | 17952710 | 17950131 | 17950207 | 17955364 | 17955475 | 1,3,1 | 0,0,0 | 4,13,3 | 2,14,4 | 274 | 150 | 3.24E-09 | 1.62E-07 | 1.0,1.0,1.0 | 0.523,0.337,0.29 | 0.616 | ENSG00000089 | [[[[17968375, 17 | 4 |
| ENSG00000089 | SNX5 | chr20 | 17952586 | 17952710 | 17951499 | 17951595 | 17955364 | 17955475 | 189,149,249 | 1,0,0 | 158,442,192 | 9,25,19 | 274 | 150 | 5.20E-13 | 4.92E-11 | 0.99,1.0,1.0 | 0.906,0.906,0.84 | 0.11 | ENSG00000089 | [[[[17968375, 17 | 2 |
| ENSG00000089 | SNX5 | chr20 | 17956932 | 17957037 | 17943109 | 17943195 | 17968374 | 17968457 | 3,10,35 | 0,0,0 | 7,16,11 | 2,7,0 | 255 | 150 | 6.80E-06 | 0.000151775 | 1.0,1.0,1.0 | 0.673,0.573,1.0 | 0.251 | ENSG00000089 | [[[[17968375, 17 | 10 |
| ENSG00000089 | SNX5 | chr20 | 17956932 | 17957037 | 17953995 | 17954117 | 17961139 | 17961326 | 30,16,22 | 0,0,0 | 24,38,21 | 6,6,3 | 255 | 150 | 1.44E-07 | 4.83E-06 | 1.0,1.0,1.0 | 0.702,0.788,0.80 | 0.235 | ENSG00000089 | [[[[17968375, 17 | 2 |
| ENSG00000089 | SNX5 | chr20 | 17956932 | 17957037 | 17953995 | 17954117 | 17968374 | 17968457 | 6,10,37 | 0,0,0 | 8,18,12 | 2,5,5 | 255 | 150 | 2.07E-08 | 8.61E-07 | 1.0,1.0,1.0 | 0.702,0.679,0.58 | 0.345 | ENSG00000089 | [[[[17968375, 17 | 2 |
| ENSG00000088 | KIZ | chr20 | 21214534 | 21214700 | 21205490 | 21205530 | 21229010 | 21229115 | 2,2,4 | 0,0,0 | 14,19,18 | 4,1,2 | 300 | 150 | 0.000140659 | 0.002139307 | 1.0,1.0,1.0 | 0.636,0.905,0.81 | 0.214 | ENSG00000088 | [[[[21126082, 21 | 2 |
| ENSG00000197 | ENTPD6 | chr20 | 25218549 | 25218614 | 25217501 | 25217581 | 25222837 | 25222978 | 7,20,60 | 0,0,1 | 39,68,65 | 5,12,9 | 215 | 150 | 3.85E-07 | 1.17E-05 | 1.0,1.0,0.977 | 0.845,0.798,0.83 | 0.167 | ENSG00000197 | [[[[25195712, 25 | 2 |
| ENSG00000197 | ENTPD6 | chr20 | 25221231 | 25221333 | 25217501 | 25217581 | 25222837 | 25222970 | 37,33,78 | 0,0,1 | 75,142,94 | 5,12,9 | 252 | 150 | 1.36E-06 | 3.61E-05 | 1.0,1.0,0.979 | 0.899,0.876,0.86 | 0.114 | ENSG00000197 | [[[[25195712, 25 | 2 |
| ENSG00000100 | ABHD12 | chr20 | 25317047 | 25317078 | 25309445 | 25309575 | 25323324 | 25323430 | 1,0,0 | 2,3,1 | 2,3,2 | 0,3,0 | 181 | 150 | 1.06E-07 | 3.70E-06 | 0.293,0.0,0.0 | 1.0,0.453,1.0 | -0.72 | ENSG00000100 | [[[[25390513, 25 | 3 |
| ENSG00000101 | HM13 | chr20 | 31549028 | 31549114 | 31544946 | 31545035 | 31550063 | 31550121 | 129,81,135 | 0,0,0 | 180,337,173 | 2,10,8 | 236 | 150 | 0.000194929 | 0.002851836 | 1.0,1.0,1.0 | 0.983,0.955,0.93 | 0.043 | ENSG00000101 | [[[[31514442, 31 | 2 |
| ENSG00000101 | HM13 | chr20 | 31566209 | 31566295 | 31561633 | 31561736 | 31569119 | 31569174 | 452,370,465 | 15,9,11 | 6,041,226,616 | 61,125,54 | 236 | 150 | 9.08E-13 | 8.28E-11 | 0.95,0.963,0.964 | 0.863,0.862,0.87 | 0.091 | ENSG00000101 | [[[[31514442, 31 | 2 |
| ENSG00000101 | TM9SF4 | chr20 | 32145090 | 32145209 | 32142981 | 32143105 | 32146784 | 32146855 | 41,33,95 | 0,0,0 | 65,110,59 | 8,11,10 | 269 | 150 | 5.88E-10 | 3.39E-08 | 1.0,1.0,1.0 | 0.819,0.848,0.76 | 0.189 | ENSG00000101 | [[[[32109714, 32 | 2 |
| ENSG00000198 | NCOA6 | chr20 | 34746806 | 34746928 | 34736689 | 34736758 | 34757219 | 34758104 | 2,5,1 | 0,0,0 | 3,19,5 | 0,5,4 | 272 | 150 | 6.26E-06 | 0.000141628 | 1.0,1.0,1.0 | 1.0,0.677,0.408 | 0.305 | ENSG00000198 | [[[[34825472, 34 | 4 |
| ENSG00000085 | ATRX | chrX | 77595711 | 77595785 | 77593695 | 77593849 | 77616612 | 77616730 | 2,2,3 | 0,0,0 | 1,23,12 | 4,15,3 | 224 | 150 | 2.17E-08 | 8.98E-07 | 1.0,1.0,1.0 | 0.143,0.507,0.72 | 0.541 | ENSG00000085 | [[[[77785982, 77 | 3 |
| ENSG00000085 | ATRX | chrX | 77599410 | 77599580 | 77593695 | 77593849 | 77616612 | 77616730 | 45,36,61 | 0,0,0 | 21,45,42 | 4,15,3 | 300 | 150 | 8.93E-10 | 5.01E-08 | 1.0,1.0,1.0 | 0.724,0.6,0.875 | 0.267 | ENSG00000085 | [[[[77785982, 77 | 3 |
| ENSG00000085 | ATRX | chrX | 77616612 | 77616730 | 77600433 | 77600564 | 77620394 | 77620532 | 19,7,29 | 0,0,0 | 4,16,9 | 2,3,5 | 268 | 150 | 2.68E-08 | 1.08E-06 | 1.0,1.0,1.0 | 0.528,0.749,0.50 | 0.407 | ENSG00000085 | [[[[77785982, 77 | 2 |
| ENSG00000101 | IFT52 | chr20 | 43604182 | 43604258 | 43596434 | 43596522 | 43613849 | 43613976 | 1,1,1 | 0,0,0 | 1,3,2 | 2,5,4 | 226 | 150 | 2.27E-09 | 1.17E-07 | 1.0,1.0,1.0 | 0.249,0.285,0.24 | 0.739 | ENSG00000101 | [[[[43590937, 43 | 3 |
| ENSG00000101 | IFT52 | chr20 | 43605001 | 43605073 | 43596434 | 43596522 | 43613849 | 43613976 | 27,26,41 | 0,0,0 | 23,46,38 | 2,5,4 | 222 | 150 | 2.73E-05 | 0.000514732 | 1.0,1.0,1.0 | 0.886,0.861,0.86 | 0.129 | ENSG00000101 | [[[[43590937, 43 | 3 |
| ENSG00000115 | ITGA4 | chr2 | 181478756 | 181478824 | 181475158 | 181475288 | 181481597 | 181481683 | 11,10,15 | 0,0,0 | 6,14,8 | 3,4,4 | 218 | 150 | 4.11E-08 | 1.58E-06 | 1.0,1.0,1.0 | 0.579,0.707,0.57 | 0.378 | ENSG00000115 | [[[[181457386, 1 | 2 |
| ENSG00000064 | CTSA | chr20 | 45896964 | 45897040 | 45894993 | 45895133 | 45898004 | 45898109 | 137,105,172 | 0,0,2 | 210,382,227 | 7,12,5 | 226 | 150 | 0.000336635 | 0.004554498 | 1.0,1.0,0.983 | 0.952,0.955,0.96 | 0.036 | ENSG00000064 | [[[[45891335, 45 | 2 |
| ENSG00000064 | CTSA | chr20 | 45897716 | 45897806 | 45894993 | 45895133 | 45898004 | 45898109 | 107,156,248 | 0,0,2 | 239,389,252 | 7,12,5 | 240 | 150 | 0.000201793 | 0.002935599 | 1.0,1.0,0.987 | 0.955,0.953,0.96 | 0.037 | ENSG00000064 | [[[[45891335, 45 | 2 |
